# Inhibition of the transcription factor PU.1 suppresses tumor growth in mice by promoting the recruitment of cytotoxic lymphocytes through the CXCL9-CXCR3 axis

**DOI:** 10.1101/2024.10.10.617590

**Authors:** Nichita Sleapnicov, Soon-Duck Ha, Shanshan Jenny Zhong, Jackie Duchscher, Sally Ezra, Shawn Shun-Cheng Li, Sung O. Kim

## Abstract

Tumor-associated macrophages (TAMs) are among the most abundant immune cells associated with tumors, which often exhibit immune regulatory phenotypes that promote tumor growth and confer resistance to anti-tumor immune therapies. Despite extensive efforts in developing immunotherapeutic strategies aimed at controlling the recruitment or reprogramming of TAMs, success has been limited due to strategic caveats, underscoring the need for a novel approach targeting the TAMs. PU.1, a lineage-dependent transcription factor, is highly expressed throughout the lifespan of macrophages. We have found that inhibition of PU.1 by the small molecule DB2313 suppresses melanoma tumor growth in mice through enhanced tumor recruitment of CD4+ T helper cells and cytotoxic T/NK cells mediated by TAMs. Whole transcriptome and targeted gene expression analyses revealed that DB2313 upregulates CXCL9 expression in bone marrow-derived macrophages (BMDMs) and TAMs. The anti-tumor effects of DB2313 were abolished by depleting macrophages with clodronate or inhibiting the CXCL9-CXCR3 chemokine axis using neutralizing antibodies against CXCL9 or CXCR3. Collectively, these results suggest that pharmacological inhibition of PU.1 suppresses tumor growth by promoting tumor infiltrating lymphocytes through the CXCL9-CXCR3 chemokine axis. Our study establishes a framework for developing TAM-modulating immunotherapies by targeting the transcriptional factor PU.1.

## Introduction

Tumor establishment, growth, and responses to treatment are critically influenced by the tumor microenvironment (TME) which is composed of a variety of cell types. Tumor-associated macrophages (TAMs) make up the largest portion of the immune cell populations within the TME and play a key role in determining the immunological characteristics of TME [1]. TAMs are heterogeneous exhibiting a spectrum of phenotypes from pro-inflammatory M1-like to anti-inflammatory M2-like [2, 3]. These heterogeneous TAMs largely promote tumor growth/metastasis and render resistance to anti-tumor immune therapies. Hence, TAMs have been a potential target for anti-tumor therapies, and various strategies have been developed to prevent the generation of TAMs or re-programming them from tumor-supportive to tumor-suppressive phenotypes [4]. These TAM-targeted therapeutic strategies include depleting TAMs by targeting macrophage colony-stimulating factor receptor (M-CSFR), inhibiting M2-signaling proteins such as phosphoinositide 3 kinase γ, Janus kinase 2, receptor-interacting protein kinase 1, or Bruton’s tyrosine kinase, and reprogramming TAMs by toll-like receptor agonists or CD40 ligands. However, these approaches have been met with limited success, and other potential challenges [5], underscoring the need for novel strategies for targeting the TAM for immunotherapy.

PU.1, a member of the E26 transformation-specific (ETS) family transcription factor (TF), plays a key role in the early development of myeloid and lymphoid cells as genetic PU.1 knockout mice lack myeloid cells and B cells and are impaired in T cells and NK cells [6, 7]. In myeloid cells, a high level of PU.1 expression is required for their proper development, function, and differentiation, as well as for optimal expression of inflammatory cytokines [8] and M2-like macrophage polarization [9]. While genetic studies employing complete or partial depletion of PU.1 underscores its crucial role in the development of immune cells, small molecule inhibitors of PU.1 such as DB2313 (DB) have shown anti-tumor effects by selectively inducing cell death of leukemia cells that require PU.1 for survival [10]. Notably, the PU.1 inhibitors showed minimal and reversible impacts on the development of lymphocytes and myeloid cells, suggesting that pharmacological inhibition of PU.1 is a viable strategy for regulating immune responses. However, how the pharmacological inhibition of PU.1 affects the TAMs and the growth of solid tumors is unknown.

To evaluate PU.1 as a potential therapeutic target for anti-cancer immunotherapy by reprogramming TAMs, we have investigated the effects and mechanisms of action of the PU.1 inhibitor DB using an established mouse melanoma model [11]. DB is a heterocyclic diamine derived from the clinically tested compound furamidine [12]. It has been shown to selectively inhibit PU.1, among ETS TFs, by binding to the minor DNA groove adjacent to the major groove that comprises the ETS-binding core motif (5’-GGAA/T-3’)[10]. We found that PU.1 inhibition by DB suppressed tumor growth by enhancing tumor-infiltration of lymphocytes, including CD4^+^ type 1 helper (Th1), CD8^+^ cytotoxic T lymphocytes (CTLs), and NK cells, through the CXCL9-CXCR3 axis.

## Methods and Materials

### Materials and reagents

DB2313 (DB) was purchased from Glixx Laboratories (MA, USA). For *in vitro* use, it was prepared in 500 mM stock solution in DMSO. For *in vivo* experiments, DB was prepared in 30% Polyethylene glycol 300 (PEG300; MediChemExpress; NJ, USA) and 70% saline. Matrigel Basement Membrane Matrix was purchased from BD Biosciences (CA, USA). InVivoMAb anti-mouse PD-1(CD279), rat IgG2a isotype control, anti-mouse CXCR3 (CD183), anti-mouse CXCL9 (MIG), and polyclonal Armenian hamster IgG isotype control antibodies were purchased from Bio X Cell (NH, USA). Antibodies used for flow cytometry and immunohistochemistry are listed in Supplementary Table S1. For immunoblotting analysis, PU.1 antibody (C-3) and anti-β-actin were purchased from Santa Cruz Biotechnology (Dallas, Texas, USA) and Rockland (PA, USA), respectively. RNA protect reagent, RNeasy Mini Kit and tissue shredder were purchased from Qiagen (ON, Canada). PureLink^TM^ RNA Mini kit and Lipofectamine RNAi MAX reagent were obtained from Invitrogen through Thermofisher Scientific (MA, USA).

### Mice

C57BL/6 female mice aged 6-7 weeks were purchased from Charles River Laboratories Canada and housed in the Animal Care and Veterinary Service facility at Western University. Mice were housed under specific-pathogen-free conditions in groups of six animals; they were provided a 12-hour light-dark cycle, bedding and nesting materials, and food and water ad libitum. All efforts were done to minimize mice suffering and all mice experiments were conducted under the guidelines of approved animal care and use protocol (AUP2022-138), at Western University.

### Cell culture

B16-OVA melanoma cells (Sigma-Aldrich) were cultured in complete DMEM (Sigma) supplemented with 10 % fetal bovine serum (WISENT INC, Quebec, Canada) and 100 U/mL penicillin-streptomycin (Gibco). G418 (200 µg/mL, WISENT INC) was also added to cell culture to maintain OVA-expressing clones but removed 24 h before inoculation. Bone marrow-derived macrophages (BMDMs) were prepared after isolating bone marrow cells from the femurs of 7-9 weeks old C57BL/6 mice (Charles River Laboratories Canada) and cultured in the presence of cell culture media obtained from L929-feeder cells producing macrophage colony-stimulating factor (M-CSF) as previously described [13].

### Mouse Tumor Model

B16-OVA cells were harvested when they reached around 80 % confluence after three ∼ four passages and washed twice with serum free DMEM, and 5 X 10^4^ cells were suspended with 50 µL of serum-free DMEM. The cell suspensions were mixed with 50 uL of Matrixgel Basement Membrane Matrix and subcutaneously injected into the right hind flank of C57BL/6 mice (body weight; ∼20g). Mice were monitored daily, and drug injection was initiated when tumor volume reached around 50∼60 mm^3^, mice were randomly divided into four or six groups (n=5∼6 per group) prior to treatment. DB2313 (DB; Glixx Laboratories, MA, USA) were prepared in 30% PEG300 in saline, and 100 µL of vehicle (30% PEG300 and 70% saline) or DB (100 µL, 17 mg/kg) were injected every 2 days via intraperitoneal (IP) injection for 12 days (six doses).

For macrophage depletion experiments, mice were treated with i.p injection of 1 mg per mouse clodronate or control liposomes (www.clodronateliposomes.com) every 4 days as described previously [14] with or without the vehicle or DB administration every 2 days for 12 days. All antibodies for mice injection were diluted in phosphate-buffered saline (PBS) to make experimental dosages, and MAb rat IgG2a isotype control (100 µL, 1 mg/kg, Clone;2A3, Bio X Cell, Lebanon, NH, USA) or anti-PD1 (100 µL, 1 mg/kg, Clone; RMP 1-14, Bio X Cell) were administered by i.p injection every 3 days. Armenian hamster IgG isotype control (100 µL,140 µg/mouse, every 4 days, Clone; Polyclonal, Bio X Cell), anti-CXCR3 (100 µL, 140 µg/mouse, every 4 days, Clone; CXCR3-173, Bio X Cell), and anti-mouse CXCL9 (100 µL, 200 µg/mouse, every 3 days, Clone; MIG-2F5.5, Bio X Cell) were administered by i.p. injection. Tumor size was measured blindly using a caliper every 1-2 days in the morning as indicated in the figure legend. Tumor volumes were calculated using the formula V(mm^3^) = (W^2^ × L)/2 (V, volume; W, width; L, length) as previously reported [15] and represented graphically. On the 13^th^ day post initiation of treatment, all mice were euthanized with Isoflurane (5 %), following the institutional guidelines and tumor tissues were harvested from randomly selected three or four mice per group to study further. No apparent adverse events were observed in each experimental group.

### Single-cell suspension preparation from tumors

Dissociation of tumors into single-cell suspension was prepared following the STEMCELL Technologies protocol. Briefly, tumor tissues were excised into millimeter-sized pieces using razor blades and digested in RPMI 1640 media containing collagenase/hyaluronidase (STEMCELL Technologies) and DNase I (Sigma-Aldrich) for 30 min at 37°C on a shaking platform. Tumor cell suspensions were passed with 70 μm filters and treated with RBC lysis buffer (BioLegend, CA, USA) by following the manufacturer’s instructions. Single-cell suspensions were then washed with 1X Phosphate Buffered Saline (PBS) containing 2% FBS for further processing.

### Tumor-associated macrophage (TAM) isolation

F4/80 positive macrophages from tumors were isolated from single-cell suspensions using the PE positive selection kit (STEMCELL Technologies) and F4/80-PE antibody (Bio-Rad) according to manufacturer instructions. Purified F4/80 positive cells were used for the RNA isolation, followed by RNA sequencing and quantitative real-time PCR (RT-qPCR).

### Flow Cytometry

Single cells prepared from tumor tissues were counted and 2-5 million cells were plated on a 96-well plate (VWR, round bottom) and washed with PBS. Cells were then incubated with PBS containing 1:8000 dilution of fixable viability dye (FVD) for 25 minutes in the dark at 4°C. After washing twice with PBS, cells were incubated with FACS buffer containing 2% fetal bovine serum, monocyte block (Biolegend), Fc block (Biolegend), and surface staining antibodies at room temperature for 25 min. Fluorescence minus one (FMO) control used the same amounts of monocyte block, Fc block, FVD, and surface stain antibodies as samples except for the antibody being controlled for. FMO controls for FVD were incubated in only PBS for the same timeframe.

For the intracellular staining, surface-stained cells were washed with first PBS and then incubated in fixation buffer (Invitrogen) for 20 min at room temperature. Cells were then washed twice with PBS and permeabilized using the permeabilization buffer (Invitrogen). Antibodies with intracellular targets were diluted in the permeabilization buffer and exposed to cells for 30 min in the dark at room temperature. Fluorescence minus one (FMO) controls were also incubated in the permeabilization buffer. Cells were washed twice with PBS and suspended in 200 μL PBS or FACS buffer for FACS analysis.

The OVA peptide (SIINFEKL)-conjugate MHC-I tetramers were generated following the vendor’s protocol. Briefly, Flex-T™ H-2Kb monomers that harbor the peptide (Biolegend) were combined with PE-conjugated streptavidin (Biolegend) in the presence of D-biotin (50 mM, Sigma-Aldrich), and NaN3 (10%, Sigma-Aldrich) on ice overnight in 1X PBS. After centrifuging down tetramers, they were resuspended and administered to samples in the same manner as surface stain antibodies.

### Immunohistochemical (IHC) staining

Tumor tissues were fixed with 10% formalin in PBS for 24 hours and then paraffin-imbedded for IHC staining. Briefly, tumor tissues were dehydrated by sequential immersion once in 50%, 70%, 90%, and twice in 100% ethanol. The tissues were then immersed twice in xylene and dehydrated. Tissues were then paraffin-embedded, sectioned, and mounted on slides. For the IHC staining, tissue slides were deparaffinized in xylene, dehydrated by running through ethanol washes in reverse order as described above, and stained with antibodies for CD8 and granzyme B. Secondary antibodies conjugated with HRP were used and the signal was detected by the 3,3’-diaminobenzidine tetrahydrochloride substrate kit from ThermoFisher Scientific. Counterstaining for nuclei was done via hematoxylin staining. Images were taken at 100× magnification using a Qimaging cooled charged-coupled device camera on an Axioscope 2 (Carl Zeiss) microscope. The optical density of secreted Granzyme B was performed by using the color deconvolution tool on the Fiji-ImageJ software (converting DAB color into optical density using the formula: OD = log_10_ (255/mean value) [16]. CD8+ cells were quantified by cell counting in the field of view (FOV), counting cells with clear DAB color around the cell margins.

### mRNA sequencing and transcriptomic analysis

Total RNAs were prepared from BMDMs and TAMs using the Qiagen RNeasy kit (Qiagen, RNeasy Mini Kit) or PureLink^TM^ RNA Mini Kit (Invitrogen by Thermo Fisher Scientific) according to the manufacturer’s protocol. For bulk tumor RNA preparations, tumor tissues were excised into ∼30 mg pieces using razor blades and submerged into RNA protect tissue reagent (Qiagen) overnight at 4 °C. Tissues were then transferred to -80 °C and pulverized using a liquid nitrogen cold hammer. Total RNAs were then isolated using the Qiagen RNeasy Mini kit according to the manufacturer’s protocol. mRNA enriched sequencing was performed by Genome Quebec, using an Illumina NovaSeq PE100 sequencer with 50 million reads per sample, following the RNA-seq User Guides provided by Genome Quebec. BAM files aligned with NCBI37/mm9 mouse genome were counted for matched genes using the FeatureCount tool (with built-in gene annotation file, using default settings) [17], and fold of changes and dispersions were estimated using the DESeq2 tool (c7 < 0.05) [18] in the Galaxy platform. The differential gene expression and gene counts were then visualized in volcano plots and bubble heat maps using GraphPad Prism 10 and R Studio. Gene ontology and hallmark annotation analysis for the genes that significantly changed were performed using the Metascape online tool (https://metascape.org/gp/index.html#/main/) [19]. For Gene Set Enrichment Analysis (GSEA) for bulk tumors, the Limma-voom tool [20] was used to filter out counts lower than 10 in the Galaxy, and the resulting differential expression tables were used for GSEA [21], followed by Cytoscape visualization [22] with a node cut-off of 0.01 q-value and an edge cut-off of 0.5.

### Transfection of small interfering (si)RNA and LPS stimulation in BMDMs

BMDMs were transfected with mouse PU.1 specific siRNAs (Invitrogen by life technologies, Cat#:10620318 and 10620319, Spi1 MSS277025 and Spi I MSS247676) for 18∼20 h using Lipofectamine RNAiMAX (Invitrogen, Life Technologies), according to the manufacturer’s instructions. Cells were re-plated to 6 well plates and incubated for 18 h. Cells were then activated by LPS (100ng/mL) for 5 h and harvested. Total mRNAs and cell lysates were prepared for RT-qPCR and immunoblotting, respectively.

### RT-qPCR

RT-qPCR was carried out as previously described [23]. Briefly, isolation of total cellular RNAs and reverse transcription were performed using the Qiagen RNeasy kit or Trizol (Ambion) and M-MuLV reverse transcriptase (New England Biotechnology) by following the manufacturer’s protocol, respectively. qPCR was conducted using the Rotor-Gene RG3000 instrument (Montreal Biotech Inc) with the 2 x Universal Sybr Green Fast qPCR Mix (Ab Clonal). Data were calculated relative to the levels of the glyceraldehyde 3-phosphate dehydrogenase (GAPDH) housekeeping gene. The primer list and the sequences used in this study are in Supplementary Table S2.

### Murine cytokine/chemokine multiplex analysis

BMDMs (60,000 cells) in 96 well plates were treated with DB (500 nM) overnight and stimulated with LPS (100 ng /mL) for 18 hours. Cell cultured media were collected after centrifugation at 3000 x g for 10 min at 4°C. The supernatants were then analyzed for the 32 cytokines using the Mouse Cytokine/Chemokine 32-Plex Discovery Assay® performed by Eve Technologies.

### Immunoblotting

Immunoblotting was performed as previously described [23]. Briefly, BMDMs were lysed in ice-cold lysis buffer containing 20 mM MOPS (pH7.2), 2 mM EGTA, 5 mM EDTA, 1 mM Na3VO4, 40 mM β-glycerophosphate, 30 mM sodium fluoride, 20 mM sodium pyrophosphate, 0.1% SDS, 1% Triton X-100, and protease inhibitor tablets (Pierce, ThermoScientific) and incubated on ice for 10 min. Whole lysates were centrifuged at 14,000 x g for 15 min at 4 °C, and supernatants were electrophoretically resolved in 10% SDS-polyacrylamide gels, followed by transfer onto 0.2 μm nitrocellulose membranes. The membranes were blocked at room temperature for 1 h with 5% (w/v) skim milk and then immunoblotted with primary antibodies overnight at room temperature, followed by the corresponding secondary antibody for 1 hour at room temperature. The bands were developed with chemiluminescence (BioRad Clarity Max Western ECL system) and images were obtained using the BioRad Chemidoc XR+ System.

### Statistics

Data were analyzed using GraphPad Prism Version 4.0 software, and the results were presented as scatter plots or bar graphs. Comparisons of means between groups were conducted with either Student’s T-tests or one-way ANOVA followed by Tukey’s multiple comparisons test.

## Results

### The PU.1 inhibitor DB2313 (DB) changes the tumor-infiltrated immune cell repertoire and suppresses tumor growth in a melanoma mouse model

To examine the effect of PU.1 inhibition on solid tumors, we tested the PU.1 inhibitor DB in the B16-OVA melanoma and 4T1 breast tumor models. Six days post tumor cell inoculation, DB (17 mg/kg, every 2 days), αPD-1 (1 mg/kg, every 3 days), or both DB and αPD-1 were administered via i.p. injection. As expected, αPD-1 significantly suppressed B16-OVA (Fig. 1A) and 4T1 (Supplementary Fig. S1) tumor growth. αPD-1 suppressed tumor growth by ∼65% when compared to vehicle control, and DB more substantially suppressed by ∼80% than αPD-1 DB. Co-treatment with αPD-1 and DB resulted in the most pronounced tumor suppression among treatments but failed to reach statistical significance when compared with DB alone. Since tumor-associated immune cells play a key role in determining tumor growth, we focused on the B16-OVA melanoma, and enumerated immune cells infiltrating the tumors in DB-treated mice in comparison to vehicle treatment. Overall, DB recruited more hematogenous immune cells (CD45^+^; ∼50% of live cells isolated) into the tumors, compared to ∼35% in vehicle-treated tumors (Fig. 1B). Using FACS with appropriate gating strategy (Supplementary Fig. S2), we profiled tumor-associated immune cells, including macrophages, T cells, B cells, and NK cells. Within the CD45^+^ cell population, no significant changes were detected in the ratios of F4/80^+^/CD11b^+^ macrophages relative to the total number of cells or CD45^+^ cells (Fig. 1C, upper panels). Additionally, within the macrophage population, the ratio between M1 (CD86^+^/CD163^-^) and M2 (CD86^-^/CD163^+^) macrophage subtypes remained unchanged (Fig. 1C, lower panels). These findings suggest that DB treatment does not significantly affect the recruitment or polarization of macrophages. In contrast, significant changes in both CD4+ and CD8+ T cells were observed in tumors treated with DB compared to vehicle control. Specifically, a significantly higher ratio of CD4^+^ T cells, relative to the total tumor cells (but not CD45^+^ cells), was detected in DB-treated tumors (Fig. 1D, upper panels). Within the CD4^+^ T cell population, the ratios of Th1 (T-bet^+^) and Th1/Th2 hybrid cells [24] were significantly increased, whereas the ratio of Th2 (Gata^+^) cells remained unchanged following DB treatment (Fig. 2D, upper and lower panels). In contrast, the Treg (FoxP3^+^) cell population was significantly decreased by DB treatment (Fig. 2D, lower far right panel). The ratios of CD8^+^ T cells, relative to the total live cells, were substantially increased in DB-treated tumors over, and the ratios of CD8^+^/granzyme B^+^ (GnzB^+^) cytotoxic T cells tended to be higher in DB-treated tumors, though this increase did not reach statistical significance (Fig. 1E, right panel). Moreover, NK cell and GnzB^+^-NK cell populations tended to be higher in DB-treated tumors without reaching statistical significance (Fig. 1F). In comparison, the ratios of plasma B cells and memory B cells remained unchanged by DB treatment (Fig. 1G). Collectively, these data suggest that DB treatment enhances tumor infiltration of anti-tumor Th1 (CD4+) and CD8^+^ cytotoxic T cells, while decreasing the infiltration of Treg cells.

**Figure 1.**
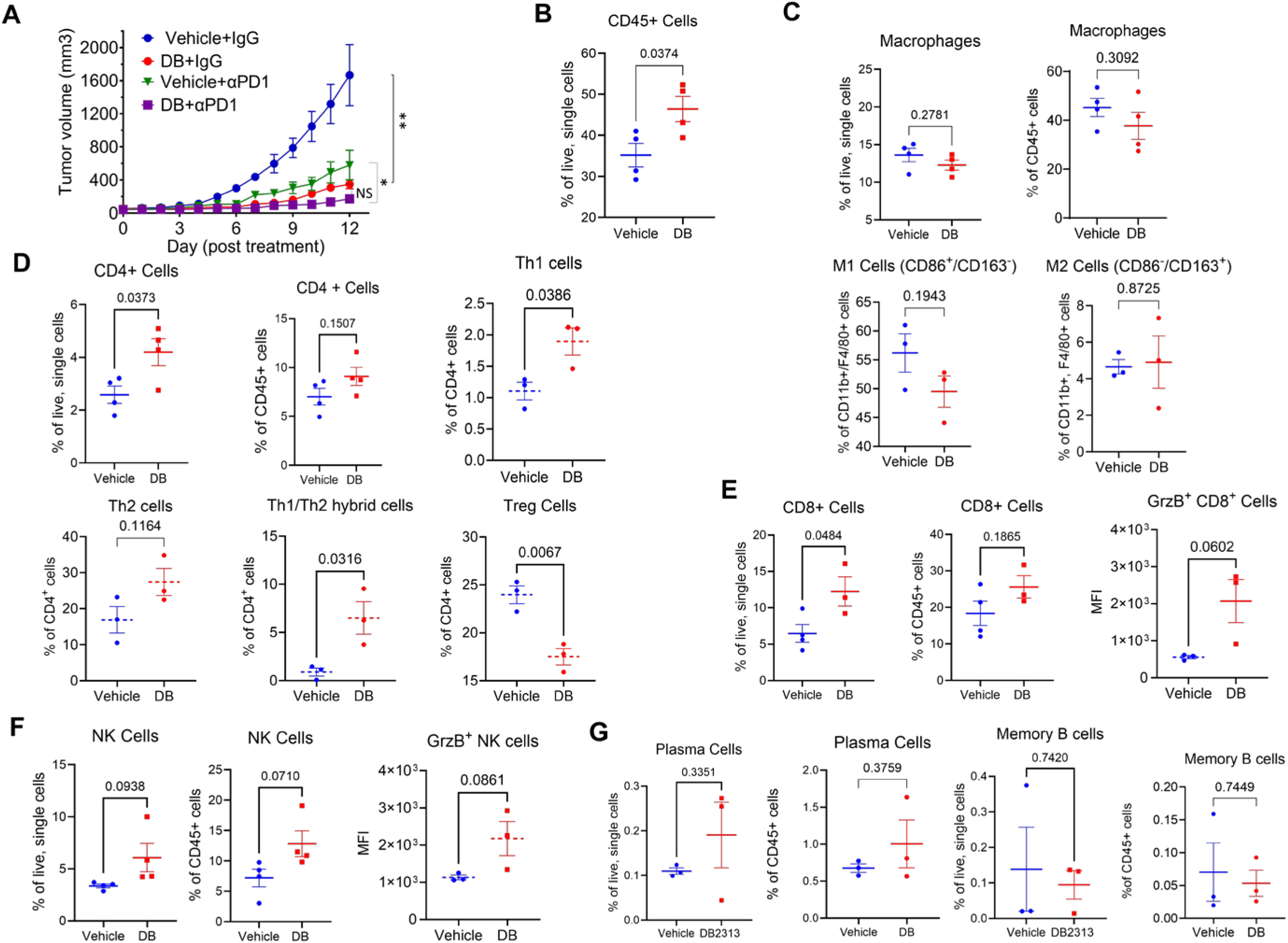
DB recruits cytotoxic immune lymphocytes and suppresses tumor growth in a mouse melanoma model. **A)** Growth curves of B16-OVA tumors treated with the PU.1 inhibitor DB (DB; 17 mg/kg), αPD-1 (1 mg/kg), or both. Error bars are expressed as mean ± s.d (n=5-6). Statistical significance was determined using one-way ANOVA with Tukey’s multiple comparisons test (NS, not significant; *, p<0.05; **, p<0.001). **B)** Flow cytometry measurement of frequencies of CD45+ cells as a percentage of live single cells between DB- and vehicle-treated tumors. **C)** Relative frequencies of macrophages (CD45^+^, CD11b^+^, F4/80^+^) as a % of live single cells or CD45^+^ immune cells, and subtypes M-1 (CD86^+^, CD163^-^) and M-2(CD86^-^, CD163^+^) as a % of macrophages between vehicle and DB-treated tumors. **D)** Relative frequencies of CD4+ T cells (CD45^+^, CD3^+^, CD4^+^) and subtypes Th1 (Foxp3^-^,Tbet^+^, GATA3^-^), Th2 (Foxp3^-^, Tbet^-^, GATA3^+^), Tregs (Foxp3^+^ CD25^+^), and Th1/Th2 (Foxp3^-^, Tbet^+^, GATA3^+^) between DB- and vehicle-treated tumors. **E)** Relative frequencies of CD8^+^ T cells (CD45^+^, CD3^+^, CD8^+^, NK1.1^-^). Cytotoxic CD8+ T cells were identified via expression of granzyme B as measured by median fluorescence intensity (MFI). **F**) Relative frequencies of total and cytotoxic NK cells (CD3^-^, NK1.1^+^, GrzB^+^). **G**) Relative frequencies of plasma cells and B cells between DB- and vehicle-treated tumors. Statistical analysis in B-G by student’s unpaired t-tests, significance defined as p<0.05 (n=3-4).

**Figure 2.**
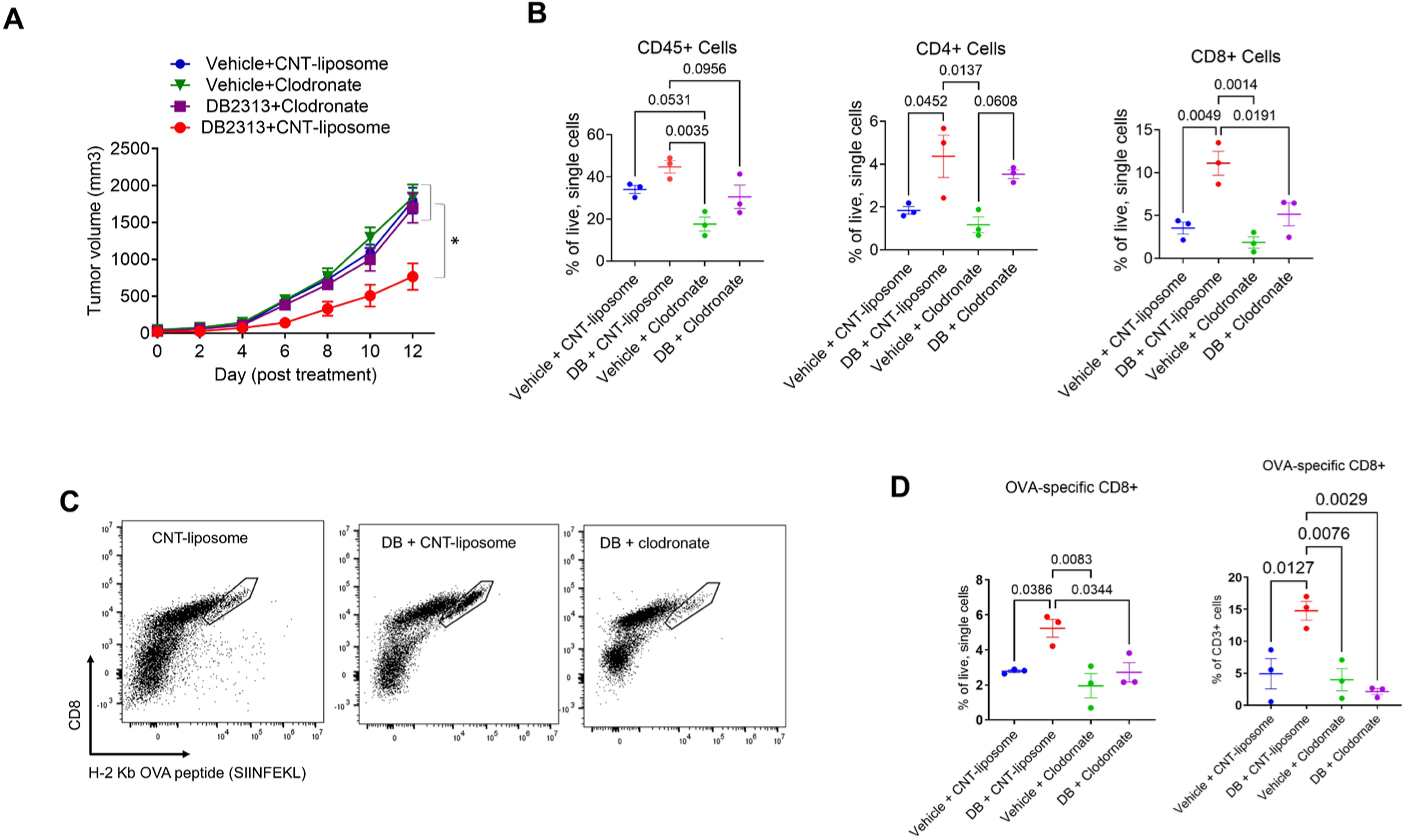
Macrophage depletion inhibits DB-induced T cell recruitment to the tumor. **A)** Macrophage depletion abolished the anti-tumor effect of DB. Shown are growth curves of B16-OVA tumors) treated with vehicle or DB (DB, 17mg/kg) together with either control liposome (1 mg /mouse) or clodronate (1 mg/mouse). Statistical significance was determined using one-way ANOVA with Tukey’s multiple comparisons test (n=5-6; *, p<0.05). **B)** Relative frequencies of immune cells (CD45^+^), CD4 T cells (CD45^+^/CD3^+^/CD4^+^), and CD8 T cells (CD45^+^/CD3^+^/CD8^+^) as a percentage of live, single cells, measured by flow cytometry in the various treatment groups. **C-D)** Representative FACS plots (8000 events in CD3+/NK1.1-gate) and relative frequencies of CD8^+^/OVA-tetramer+ cells corresponding to the different treatment groups. Statistical significance was tested by one-way ANOVA with Tukey’s multiple comparisons test, with significance defined as p < 0.05 (n=3).

### Depletion of tumor-associated macrophages (TAMs) abolished the anti-tumor effects of DB

Given that macrophages play a crucial role in the recruitment of T cells to tumors and that PU.1 expression is typically low (or absent) in T cells but high in macrophages, the observed effect of DB in enhancing T cell recruitment to tumors may be mediated by macrophages. To investigate this possibility, we examine the anti-tumor effects of DB in B16-OVA tumor-bearing mice, with or without macrophage depletion using clodronate. Treatment with clodronate liposomes led to ∼50% reduction in TAMs, compared to the control liposome (Supplementary Fig. S3). Notably, clodronate significantly reversed the tumor growth-suppressing effect of DB, whereas clodronate or the control liposome alone had no effect (Fig. 2A). In line with these results, flow cytometry analysis showed that the enhanced recruitment of CD45^+^ immune cells, as well as CD4^+^ and CD8^+^ T cells into tumors by DB, was substantially diminished when macrophages were depleted (Fig. 2B). To further examine if DB enhanced recruitment of tumor-specific cytotoxic T cells in a macrophage-dependent manner, we used tetramers of H-2 Kb (C57BL6 mouse MHC-I molecule) linked to the OVA peptide (SIINFEKL). DB significantly increased the recruitment of OVA-specific CD8^+^ T cells into tumors by more than 2-fold per given number of live tumor cells, as well as that of CD3^+^ lymphocytes (Fig. 2D). Importantly, this increase in OVA-specific CD8^+^ T cell recruitment was no longer observed when TAMs were depleted using clodronate (Fig. 2C, D). These results suggest that TAMs play a key role in DB-induced recruitment of cytotoxic T cells and the suppression of melanoma growth.

### DB affects the expression of specific cytokines and chemokines in activated bone marrow-derived macrophages (BMDMs)

While this study identified a crucial role for TAMs in mediating the anti-tumor effect of DB through promoting T cell tumor infiltration, the underlying mechanism remains unclear. Because TAMs are partially derived from monocytes recruited to the tumor [2], we next examined the effects of DB on gene transcription in activated BMDMs (e.g, stimulated with 100 ng/ml lipopolysaccharide (LPS) for 5 h), through RNA-sequencing analysis, particularly focusing on cytokine/chemokine transcript expressions. LPS induced the expression of 3,134 genes and suppressed the expression of 2,658 genes (FC > 2, p < 0.05; Fig. 3A; Supplementary Table S3). Among the LPS-induced genes, DB further enhanced the expression of 689 genes and suppressed the expression of 544 genes (Fig 3B). The Metascape gene ontology (GO) and hallmark signature gene analysis program [19] identified the top 20 gene categories among the LPS-induced genes. As expected, the top categories of the LPS-induced genes (L-Up) were related to inflammatory and innate immune responses, cytokine production pathways, and anti-viral responses. Interestingly, the genes enhanced by DB (DB-Up) overlapped with the LPS-induced GO/hallmark terms (19 out of 20). Similarly, the genes inhibited by DB (DB-Down) coincided with the majority of the GO/hallmark terms found to be enhanced by DB (14 out of 19). These results suggest that DB has both positive and negative effects on LPS-induced genes related to inflammatory responses.

**Figure 3.**
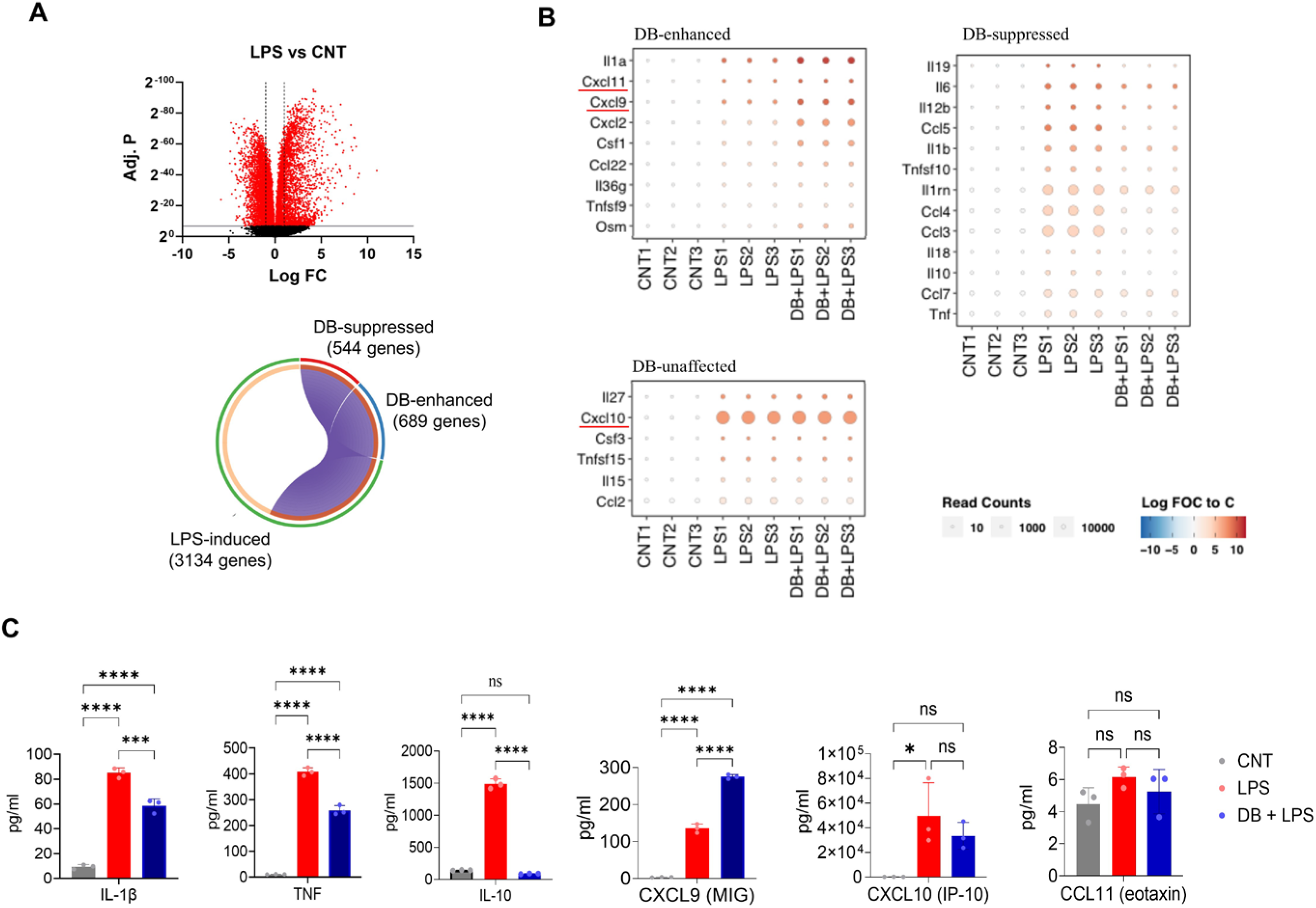
DB regulates cytokines expression in LPS-activated BMDMs. M-CSF-derived BMDMs were exposed to the drug vehicle or DB (500 nM) for 18-20 hours and then activated by LPS (100 ng/mL, 5 h). Total mRNAs were sequenced by Illumina sequencing, and differential gene expression analysis was performed using the FeatureCount and Limma-voom tools in the Galaxy platform. **A)** Volcano plot showing differential gene expression between LPS-activated and control BMDMs. Red dots indicate genes with an adjusted P value smaller than 0.01. **B**) A circos plot highlighting DB- and LPS-regulated genes. Among the LPS-induced gene (3134 genes, FC >2, adj. p <0.01), 544 were DB-suppressed (FC <0.5, adj. p <0.01), while 689 were DB-enhanced (FC >2, adj. p <0.01). **C)** Bubble heat map plots highlighting 28 cytokines/chemokines of which DB enhanced the expression of 9 (top-left panel) suppressed 13 (top-right panel), and did not affect 6 (bottom-left panel). **D)** Bar graphs showing the effect of DB and LPS on the expression of selected cytokines, measured by cytokine array (see Methods for details). Statistical analysis was by one-way ANOVA test followed by Tukey’s Multiple comparisons test (n=3; * adj. p≤ 0.05, ** adj. p ≤ 0.01, *** adj. p ≤ 0.001, **** adj. p ≤ 0.0001).

Since macrophages regulate the TME primarily through cytokines/chemokines, we focused on the effects of DB on 112 annotated murine cytokines/chemokines (Supplementary Table S4) in LPS-activated BMDMs. Among the 3,134 LPS-induced genes, 28 cytokines/chemokines with over 1000 reads per million counts were included in the analysis (Fig. 3B; Supplementary Table S5). Of the 28 cytokine/chemokine genes, DB inhibited the expression of 13, further enhanced the expression of 9 (including Cxcl9 and Cxcl11), and had no effects on 6 genes (such as Cxcl10). The expression patterns of 6 representative cytokine/chemokine genes, including DB-inhibited IL-1β, TNF & IL-10; DB-enhanced Cxcl9 & Cxcl11; DB-unaffected Cxcl10, were confirmed by qPCR (Supplementary Fig. S4). A multiplexed cytokine array analysis further confirmed the significant effect of DB treatment on suppressingIL-1β, TNF & IL-10 and enhancing CXCL9 gene expression 24 hrs post-LPS stimulation (Fig. 3D). Our findings that DB inhibited the expression of pro-inflammatory cytokines such as IL-1β, IL-6, IL-8 (CCL8), and TNF are consistent with previous reports [8]. However, it was unexpected that DB significantly increased the expression of CXCL9, a chemokine that plays a crucial role in the recruitment of tumor-infiltrating lymphocytes (TILs) and anti-tumor immunity [25]. To confirm the results of PU.1 inhibition in cytokine/chemokine gene transcription, we depleted PU.1 from BMDMs using siRNAs. The two specific siRNA oligos were able to knock down PU.1 by ∼80% on both the mRNA and protein levels in 48 h post-transfection (Fig. 4A). Like DB, the PU.1 siRNAs significantly enhanced the expression of CXCL9 while inhibiting the expression IL-1β, TNF-α, and IL-10 (Fig. 4B) in LPS-activated BMDMs. Intriguingly, PU.1 depletion also significantly increased the expression of CXCL11 over the levels induced by LPS while having no effects on the expression of LPS-induced CXCL10. Therefore, the depletion of PU.1 from BMDMs by siRNAs corroborated the results obtained from pharmacological inhibition on cytokine/chemokine gene transcription.

**Figure 4.**
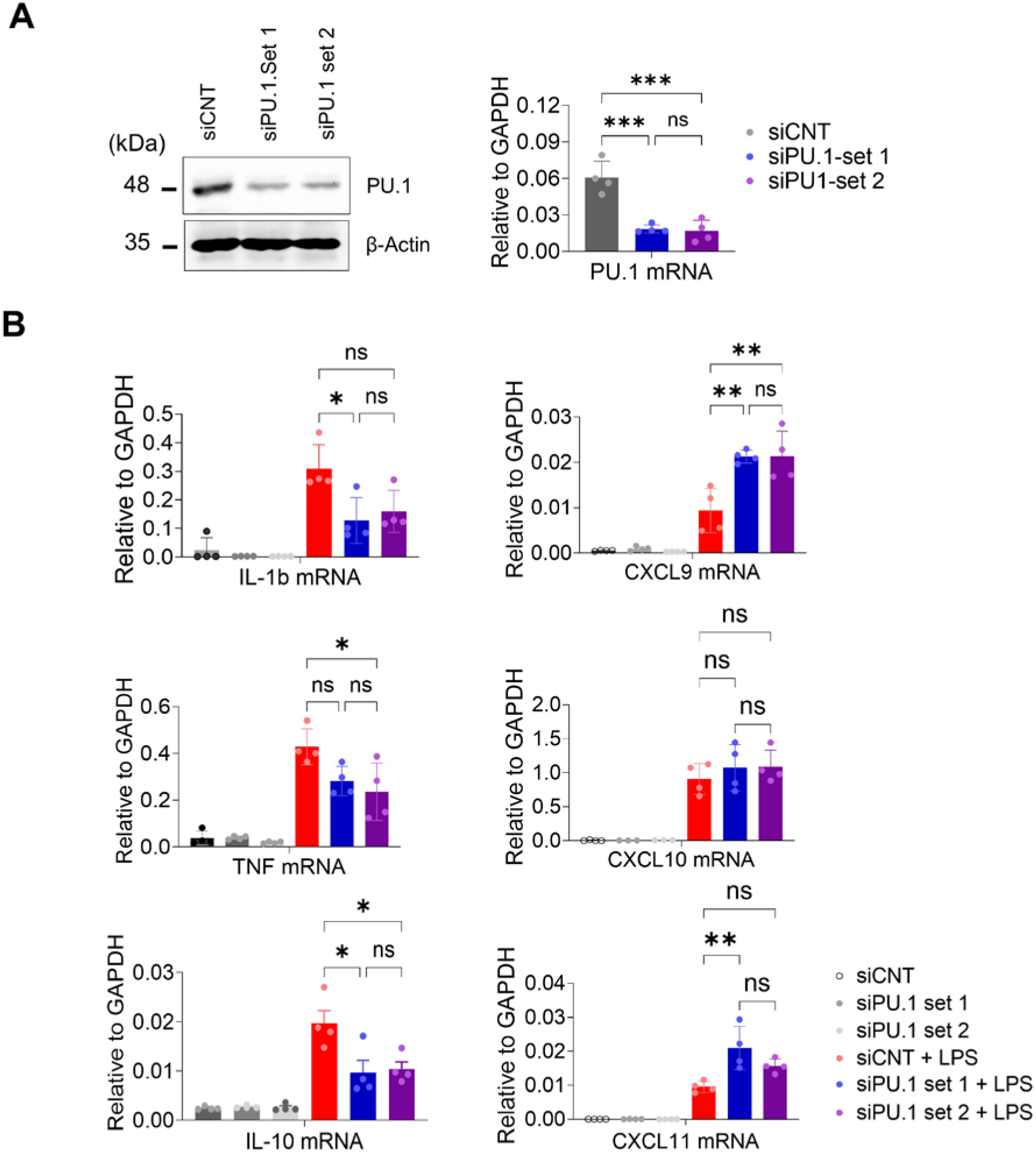
PU.1 knockdown by siRNAs produces similar effects on cytokine transcription as DB in BMDMs. **A)** Left panel: Westen blot of total cell lysates of BMDMs transfected with PU.1-specific siRNA or control oligos (siCNT). β-actin was included as a loading control. Right panel: PU.1 mRNA levels (relative to GAPDH) from the same cells, determined by RT-qPCR. **B)** Expression levels of 6 selected cytokines in BMDMs transfected with PU.1-specific or control siRNA, with or without LPS treatment.* adj. p ≤ 0.05, ** p ≤ 0.01, *** p ≤ 0.001, **** p ≤ 0.0001, one-way ANOVA test followed by Tukey’s Multiple comparisons test (n=3)

### DB selectively enhances the expression of Cxcl9 in melanoma TAMs in vivo

To confirm the effect of DB on TAMs, F4/80^+^ macrophages were isolated from tumors of mice 13 days after treatment with vehicle or DB (3 mice in each group). Transcriptomic analysis of the isolated TAMs identified 259 genes that were significantly altered (>2-fold, p<0.05) by DB treatment, including 255 genes up-regulated and 4 down-regulated genes (Fig. 5A). Metascape analysis of the 255 up-regulated genes identified vasculature development, positive regulation of cell motility, leukocyte migration as the top-ranked GO terms (Fig. 5B). Among the 5 cytokine/chemokine genes induced by DB (Fig. 5C), the transcript for CXCL9 showed the largest increase in expression in DB-treated TAMs compared to vehicle-treated TAMs, with a 2.1-fold increase (p < 0.01; Supplementary Table S6). This increase in CXCL9 mRNA expression in DB-treated TAMs was subsequently confirmed by qPCR (Fig. 5D). Importantly, the increase in CXCL9 mRNA levels in DB-treated tumors was abolished when macrophages were depleted using clodronate (Fig. 5E). Collectively, these results suggest that DB elevates CXCL9 production within the TME via TAMs.

**Figure 5.**
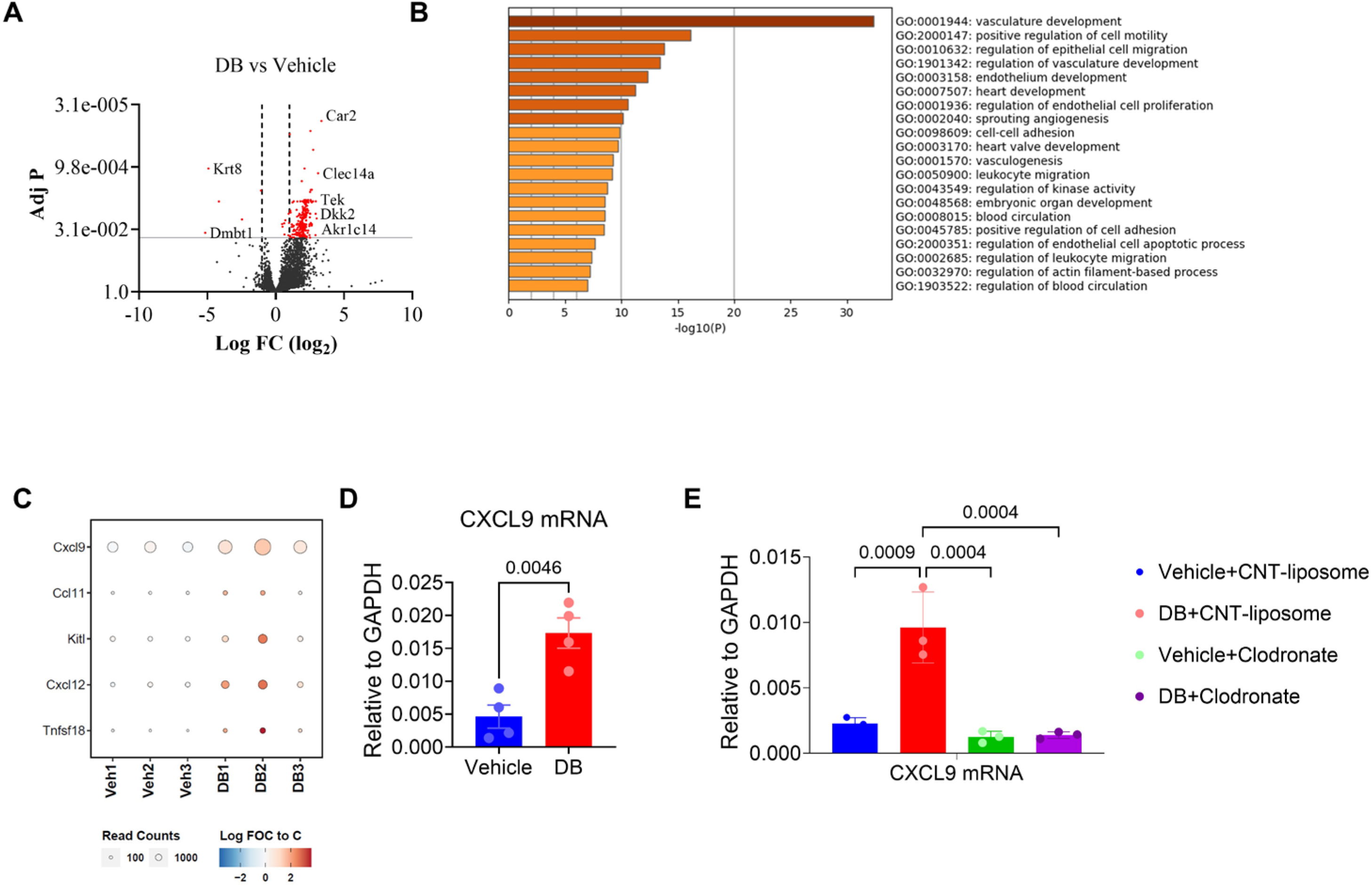
DB enhances the expression of Cxcl9 in tumor-associated macrophages (TAMs). **A)** A volcano plot showing differentially expressed genes between TAMs isolated from DB- or vehicle-treated melanoma tumors. Red dots indicate genes with an adj. p< 0.01. **B)** DB-affected genes (FC > 1.5, adj. p <0.05) were analyzed for GO terms using the Metascape online program. The top 20 GO and hallmark terms are presented. **C)** A bubble heatmap showing 5 cytokines displaying increased expression in DB-treated TAMs, compared to vehicle-treated TAMs. **D)** CXCL9 mRNA levels in TAMs isolated from DB- or vehicle-treated tumors were determined by real-time PCR. **E)** TAM depletion by clodronate abolished the effect of DB on CXCL9 mRNA expression. Shown are mRNA levels of CXCL9 (relative to GAPDH) in TAMs isolated from tumors under the specified treatments. Statistical significance was determined by a one-way ANOVA test followed by Tukey’s Multiple comparisons test (n=3-4).

### The CXCL9-CXCR3 chemokine axis plays a key role in the recruitment of cytotoxic lymphocytes into tumors and the tumor growth suppression induced by DB

While our study demonstrated that DB increases CXCL9 expression in BMDMs and TAMs, effective T cell recruitment into tumors requires the CXCL9 receptor, CXCR3 expressed on T cells. Indeed, the CXCR3-CXCL9 axis plays a critical role in tumor-infiltration and reinvigoration of CD8^+^ T cells in response to PD-1 blockade [26]. Therefore, we next examined the role of CXCL9 and CXCR3 in mediating the anti-tumor effects of DB using the B16-OVA tumor model. Specifically, tumor-bearing mice were treated with DB together with control IgG, or neutralizing antibodies for CXCR3 (αCXCR3, 140 µg/mouse) every 4 days or CXCL9 (αCXCL9, 200 µg/mouse) every 3 days, starting 6 days after B16-OVA cell inoculation. While αCXCR3 and αCXCL9 alone did not affect tumor growth in vehicle-treated mice, the tumor-suppressive effect of DB, observed in DB+control IgG-treated mice, was abolished in mice treated with DB in combination with either αCXCR3 or αCXCL9. Mechanistically, αCXCR3 or αCXCL9 treatment significantly blocked the recruitment of CD45^+^ immune cells, CD4^+^/CD8^+^ T cells, and NK cells, but not macrophages, into tumors induced by DB (Fig. 6B). In line with these data, immunohistochemistry analysis of tumor tissues confirmed that DB enhanced recruitments of CD8^+^ cells into tumors (Fig. 6C), and production of GrzB in tumors (in both intracellular and extracellular forms; Fig. 6D), both of which were inhibited by αCXCR3.

**Figure 6.**
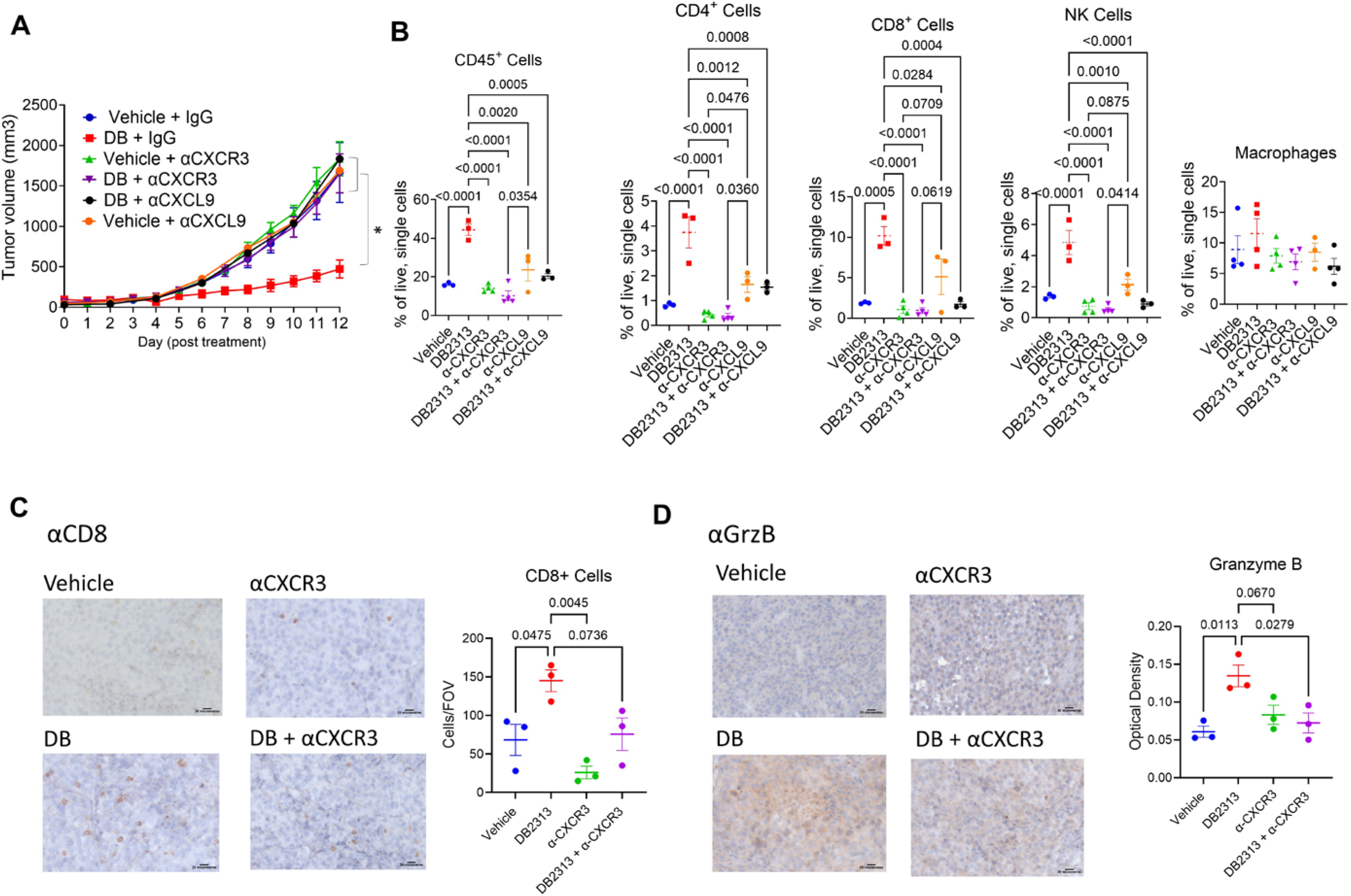
CXCL9- and CXCR3-neutralizing antibodies abolish the anti-tumor effect of DB by reducing TILs. **A**) Growth curves of B16-OVA melanoma treated with DB or vehicle (every 2 days) in combination with neutralizing antibodies for CXCL9 (αCXCL9, every 3 days), CXCR3 (αCXCR3, every 4 days), control IgG (every 4 days). **B)** Relative frequencies of immune cells (CD45+), CD4 or CD8 T cells (CD45^+^/CD3^+^, CD4^+^ or CD8+), and NK cells (CD45^+^/CD3^-^/NK1.1^+^). Macrophages (CD45^+^/CD11b^+^/F4/80^+^) were included for comparison (n = 3-4 per group). **C-D)** Immunohistochemical staining of CD8 and Granzyme B (GrzB) in tumors under the specified treatments. Tumor tissues were stained with CD8- and GrzB-specific antibodies, followed by incubation with HRP-conjugated secondary antibodies. HRP signals were detected by the DAB substrate kit as described in Methods. Representative images of the tumor tissue slides are presented in C & D (left panels). The right panels in C & D are quantification of CD8 IHC, expressed as cells/FOV, and GrzB IHC, expressed as the optical density of DAB. Statistical significance was determined by a one-way ANOVA test followed by Tukey’s Multiple comparisons test. *, P<0.05. n=5-6 for panel A, n=3-4 for other panels.

### The CXCL9-CXCR3 chemokine axis is responsible for the DB-induced global transcript changes that promote anti-tumor immune responses

To examine the role of the CXCL9-CXCR3 chemokine axis in global transcript changes of tumors, total mRNAs extracted from the whole tumor tissues were sequenced, and differential expression of transcripts was analyzed using Gene Set Enrichment Analysis (<u>https://</u>www.gsea-msigdb.org/gsea/index.jsp). The data were then visualized using the Cytoscape platform (Gene Ontology Biological Process dataset; FDR <0.5, https://cytoscape.org/). Global transcripts of DB-treated tumors were highly enriched in genes related to both innate and adaptive immune responses, while genes associated with nonimmune functions such as cell development and differentiation were suppressed (Fig. 7A; node list and NES in Supplementary Table S7). Compared to control IgG-treated tumors, αCXCR3 treatment largely down-regulated GO and hallmark terms related to lymphocyte immune responses, such as T, B, and NK cell activation and differentiation. In contrast, genes associated with nonimmune cell function, such as keratinocyte and vascular cell growth and migration, were up-regulated (Fig. 7B; Supplementary Table S7). When comparing the transcriptomics of DB+αCXCR3 with those of DB-treated tumors, we observed substantial enrichments of various GO/hallmark terms related to nonimmune cell proliferation/differentiation, as well as leukocyte chemotaxis (Fig. 7C; Supplementary Table S7). Also, when comparing the transcripts of DB+αCXCR3-treated tumors with those of IgG-treated tumors, most of the DB-enriched GO/hallmark terms were absent (Fig. 7D; Supplementary Table S7). Altogether, these data suggest that CXCR3 plays a key role in the global enrichment of immune response gene transcription in DB-treated tumors.

**Figure 7.**
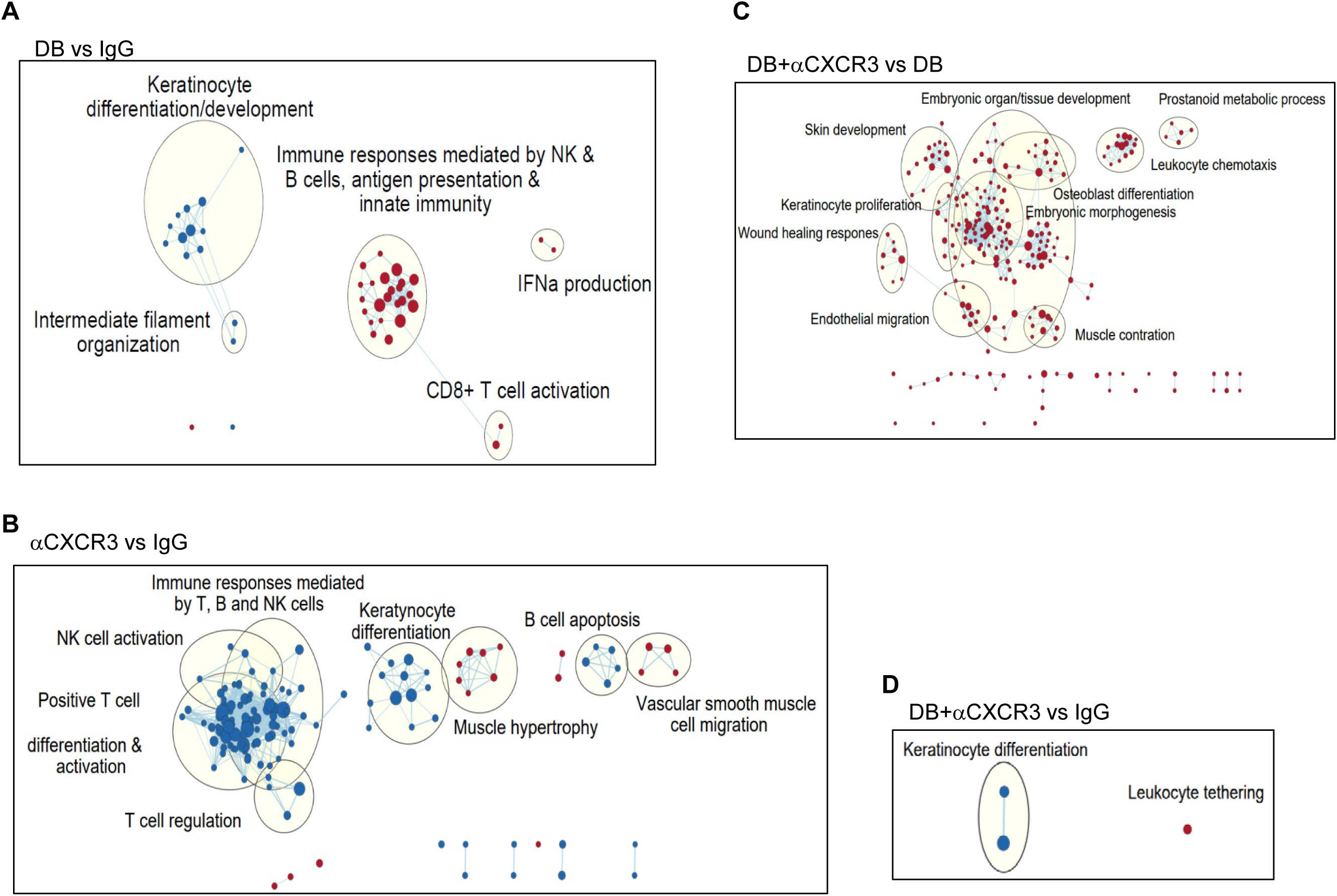
The CXCL9-CXCR3 axis is responsible for DB-induced transcriptional changes in tumors. B16-OVA tumors under the same treatments as described in Figure 6A were harvested on day 13 for mRNA sequencing. Transcript counts were analyzed by GSEA and visualized using the Cytoscape program. Enriched GO nodes (dots) and leading edge (grouped in circles with key functional annotation) are presented in pair-wise comparisons (upregulation in red, downregulation in blue), including DB+IgG vs Vehicle + IgG (**A**), Vehicle + αCXCR3 vs Vehicle + IgG (**B**), DB + αCXCR3 vs DB + IgG (**C**), DB + αCXCR3 vs Vehicle + IgG (**D**).

## Discussion

PU.1 is a member of the ETS-family transcription factors that are considered oncogenic. Previous studies on leukemia have shown that PU.1 can be either oncogenic [27] or tumor-suppressive [28] depending on the type of leukemia. In solid tumors, however, high expression levels of PU.1 have been associated with shorter survival rates across different cancers (Supplementary Fig. S6), including melanoma and breast cancer, suggesting a pro-tumour role for PU.1 [29]. In this study, we showed that pharmacological inhibition of PU.1 by DB significantly suppressed tumor growth in the B16-OVA and 4T1 mouse tumor models. Our in-depth investigation using the B16-OVA melanoma model revealed that the anti-tumor effect of DB is mediated by enhanced tumor infiltration of cytotoxic lymphocytes, particularly cytotoxic CD8^+^ T cells, which depends on the increased expression of CXCL9 by tumor-associated macrophages (TAMs) and the CXCL9-CXCR3 chemokine axis.

TAMs are the most abundant immune cells in the tumor microenvironment and play crucial roles in tumor development [1]. We also found that TAMs comprise approximately 40% of the hematogenous immune cells in our melanoma mouse model (Fig. 1B). The crucial involvement of TAMs in mediating the antitumor effect of DB was substantiated by macrophage depletion experiments, which showed a nearly complete abatement of the DB’s tumor-suppressive effect (Fig. 2A). Moreover, the enhanced recruitment of CD45^+^ immune cells, CD4^+^ and CD8^+^ T lymphocytes, and OVA-specific CD8^+^ cells induced by DB were significantly abrogated by macrophage depletion (Fig. 2B). Since TAMs modulate TME mainly by releasing cytokines and chemokines, we examine how PU.1 inhibition (by DB and siRNAs) affects the expression of cytokines/chemokines in LPS-activated BMDMs. LPS potently activates the toll-like receptor 4 (TLR4), which is activated in TAMs and other cells within tumors and plays a key role in modulating TME, including in melanoma [30]. In BMDMs, LPS activated the transcription of almost 50% of all cytokine/chemokine genes (49 out of 112 genes; FC > 2, p < 0.05; Supplementary Table S6). As expected for the transactivating role of PU.1, DB suppressed the expression of cytokine genes such as *Il1b*, *Il6*, *Tnf*, and *Il10*, as well as chemokines *Ccl3/4/5/7* (Fig. 3C & Supplementary Fig. S4), all of which can promote tumor growth. At the same time, PU.1 exhibited a repressive role in the transcription of various cytokines and chemokines, as DB enhanced the expression of *Il1a*, *Cxcl2*, and *Ccl22*, all of which are inherently tumor-supportive [31, 32]. Interestingly, DB had opposing effects on the expression of *Il1a* (encoding IL-1α) and *Il1b* (encoding IL-1β), both of which bind to the same receptor, IL-1R, but exert either complementary or contrasting effects in different types of cancers [31]. Therefore, how each of these cytokines and chemokines contributes to the anti-tumor effects of DB is yet to be delineated.

Notably, PU.1 inhibition (through either DB or PU.1-targeting siRNAs) further enhanced the expression of CXCL9 and 11 induced by LPS, while not affecting CXCL10 (Fig. 3 & 4). Given that these chemokines are involved in the recruitment of cytotoxic lymphocytes into tumors through binding to the common receptor CXCR3 [33], thereby suppressing tumor growth, our finding that DB specifically enhanced the expression of CXCL9 and CXCL11 underscores the importance of the CXCL9/11-CXCR3 axis in mediating the anti-tumor effect of DB. Our data also demonstrated that the source of *Cxcl9* transcription is TAMs as the increase in CXCL9 transcript expression by DB was completely abrogated when macrophages were depleted (Fig. 5E). Nevertheless, the exact mechanism underlying how DB enhances *Cxcl9* expression in BMDMs and TAMs is unknown. Our findings are also at odds with a report showing that PU.1 depletion inhibited the expression of CXCL9/10/11 in interferon γ-activated microglia [34]. PU.1 has been shown to have both transactivation and repression effects on different genes and cell types, potentially by interacting with different epigenetic histone modifiers and transcription factors [35]. Currently, we are examining the mechanisms by which PU.1 and DB2313 regulate Cxcl9/10/11 expression in BMDMs, noting that these mechanisms may differ in microglia.

A crucial role for the CXCL9/10/11-CXCR3 chemokine axis in anti-tumor immunity has been demonstrated in various tumors [36], and high levels of CXCL9/10 are associated with increased infiltration of immune cells and better prognosis in melanoma patients [37]. Particularly, CXCL9/10/11 produced by TAMs are required for antitumor immune responses following immune checkpoint inhibitor treatments as they play an essential role in recruiting CTLs and NK cells into tumors [38]. In line with these studies, we demonstrated that enhanced expression of *Cxcl9*, and possibly *Cxcl11*, by DB treatment was associated with increased infiltration of Th1 (CD4^+^/T-bet^+^), CTLs (CD8^+^/GrzB^+^), and NK (NK1.1^+^/GrzB^+^) into tumors (Fig. 1), which was abrogated by CXCL9- and CXCR3-neutralizing antibodies (Fig. 6B-D). Importantly, blocking the CXCR9-CXCR3 axis abolished both the overall tumor growth-suppressing effects (Fig. 6A) and the overall immune-responsive transcriptomic changes in the tumors (Fig. 7D) induced by DB, suggesting that the CXCR9-CXCR3 chemokine axis is the primary driver of anti- tumor immune responses triggered by DB.

To date, enhancing CXCL9/10/11-CXCR3 chemokine axis by administering recombinant CXCL9/10/11 proteins or using expression vector systems has demonstrated positive outcomes in various pre-clinical tumor models, including skin, lung, kidney, and colon tumors. However, unlike the paracrine effects that recruit anti-tumor immune cells by chemokines released by macrophages, the autocrine effects of the CXCL9/10/11-CXCR3 axis within tumor cells are associated with increased tumor growth and metastatic potential [33]. Thus, it has been suggested that selectively activating the CXCL9/10/11-CXCR3 paracrine axis is a more effective anti-tumor strategy. This study, for the first time, suggests that pharmacological inhibition of PU.1 enhances the CXCL9-CXCR3 paracrine axis, likely through targeting TAMs. In addition, the effectiveness of immune checkpoint inhibitors (such as PD-1/PD-L1 inhibitors) is limited in part due to TAMs, thus targeting TAMs has been suggested as a combinatory therapy. The anti-tumor effects in the melanoma (Fig. 1A) and breast tumor (Supplementary Fig. S1) models in our study are in line with the indispensable role of the CXCL9/10/11-CXCR3 axis in effective αPD-1 treatment [26]. However, further detailed studies are required to elucidate the mechanism by which PU.1 inhibition enhances the anti-tumor effects of αPD-1.

In addition to macrophages, PU.1 is involved in early T cell development, where its expression is turned off at the thymic stage [39]. This dynamics in PU.1 expression is crucial for avoiding T cell malignancy [40]. Among mature T cells, PU.1 is uniquely expressed in helper T cell 9 (Th9) and Th2 cells [41]. IL-9-producing Th9 cells are a new subgroup of CD4^+^ T cells that are differentiated in response to TGF-β and IL-4 [42]. These cells have dual roles in tumorigenesis: on one hand, they enhance the function of immunosuppressive regulatory T cells and promote tumor growth [43]; on the other hand, they suppress tumor growth by indirectly recruiting cytotoxic lymphocytes [44]. The effects of DB on Th9 cell function and its role in the anti-tumor response remain to be investigated. In contrast, Th2 cells also express PU.1, albeit at low levels, and knocking down PU.1 by siRNAs increases the production of Th2 cytokines [45]. We observed that DB-treated tumors showed no differences in the Th2 cell population from controls (Fig.1D), ruling out the involvement of Th2 cells. Of interest, DB substantially increased the population of Th1/Th2 hybrid cells while decreasing Treg cells (Fig. 1D). Th1/Th2 hybrid T cells are known to naturally develop with intermediate phenotypes characteristic of both Th1 and Th2 cells [24]. While the role of Th1/Th2 hybrid T cells in tumor growth is unclear, the involvement of Treg cells in tumor progression is well established. Intriguingly, Treg cells also express CXCR3 and can be recruited into tumors by CXCL9-producing BATF3^+^ dendritic cells [46]. We observed a substantial decrease of Treg cells in DB-treated tumors, likely due to the overriding inhibitory effects of DB on other cytokines/chemokines, such as IL-10 and CCLs, which are also required for the generation and recruitment of Treg cells [47].

In addition, PU.1 plays a key role in early B cell development, likely up to the common lymphoid progenitor stage. Although PU.1 continues to be expressed at low levels, it is dispensable for B cell differentiation [48]. However, PU.1 promotes the generation of B1 B cells [48] and plasma cells [49]. Plasma cells can infiltrate tumors, where they may confer anti-tumor effects by producing antibodies and presenting tumor antigens to T cells. However, they can also support tumor growth, likely by releasing immune suppressive cytokines and ligands after being programmed to become immunosuppressive B cells [50]. We observed no differences in the populations of plasma and memory B cells (Fig. 1G), ruling out their involvement in the anti- tumor effects of DB.

In summary, this study demonstrated that pharmacological inhibition of PU.1 suppressed tumor growth and enhanced the anti-tumor effects of the immune checkpoint inhibitor αPD-1 in melanoma and likely breast cancer mouse models, at least in part by selectively recruiting CTLs and NK cells through the CXCL9-CXCR3 chemokine axis. These results suggest a novel immunotherapy strategy that reprograms TAMs by targeting the transcription factor PU.1. However, the precise role of PU.1 in TAMs in supporting tumor growth still needs to be addressed in more PU.1-specific or genetic approaches. Also, further studies on the anti- tumour effects of PU.1 inhibition in different tumor models will substantiate PU.1 as a potential therapeutic target.

## Declarations

This work was supported by grants from the Cancer Research Society (to SOK and SSCL; 863914) the Canadian Institute of Health Research (to SSCL; MOP 201809), and the Natural Science Engineering Research Council (to SOK; RGPIN 05514). SSCL is a Wolfe Medical Research Professor and Canada Research Chair (Tier I) in the Molecular and Epigenetic Basis of Cancer. SJZ held a Translational Breast Cancer Research Scholarship from the Canadian Cancer Society.

## Notes

### Competing Interest Statement

The authors have declared no competing interest.

