## Supplemental figures for "Inhibition of the transcription factor PU.1 suppresses tumor growth in mice by promoting the recruitment of cytotoxic lymphocytes through the CXCL9-CXCR3 axis"

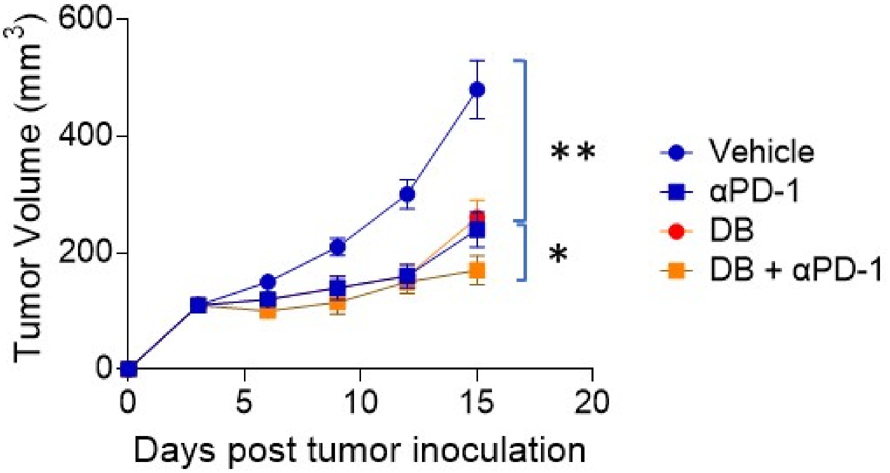


**Supplementary Figure S1. DB2313 suppressed 4T1 breast tumor growth in mice. A)** DB2313 significantly slowed down tumor growth in the 4T1 mouse tumor models, while co-administration of a PD-1 antibody (aPD-1, 1 mg/kg) substantiated the anti-tumor effect of DB2313. 4T1 tumor cells (5 X $10^4$) cells were subcutaneously injected into the mammary fat pad of BALB/c mice. Peritoneal injections of the PU.1 inhibitor DB2313 (DB; 17 mg/kg; every 2 days), αPD-1 (1 mg/kg; every 3 days) were started on day 3 after human breast tumor 4T1 cell injection. Tumor sizes were measured every 3 days for 15 days . *, $p<0.05$, **, $p<0.002$; One-way ANOVA (n= 5). All error bars are expressed as mean ± s.d.

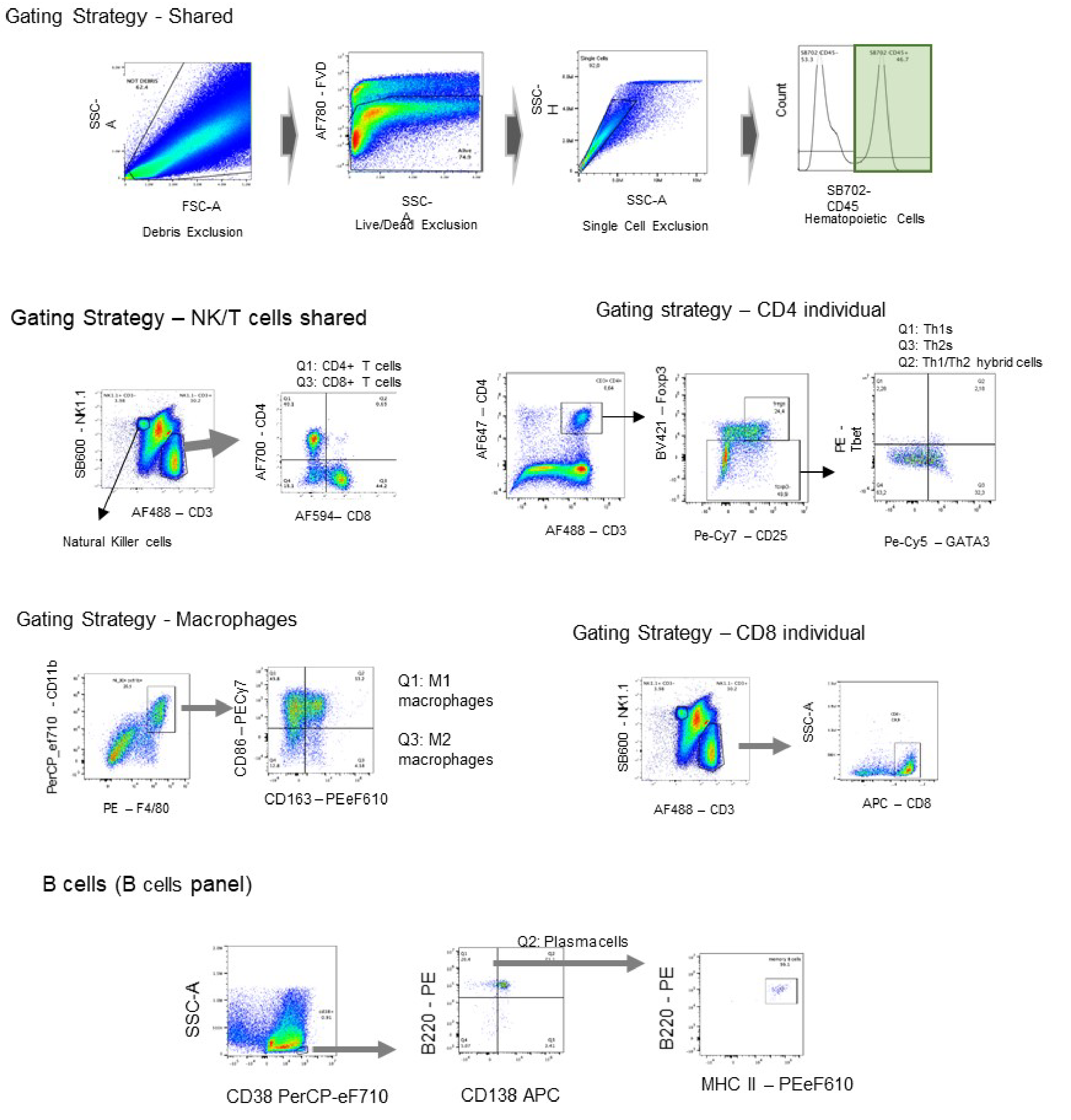


**Supplementary Figure S2. Gating strategy for specific immune cell subtypes and representative flow cytometric plots.** Flow cytometry gating strategy for all cell types. All samples went through shared gates before cell-lineage sorting. T cells were either sorted with CD4/CD8 in one panel or with CD4 and CD8 in separate panels with subsets. NK cells were always phenotyped in the same panel with CD8+ T cells. B cells and macrophages were phenotyped in their own panels.

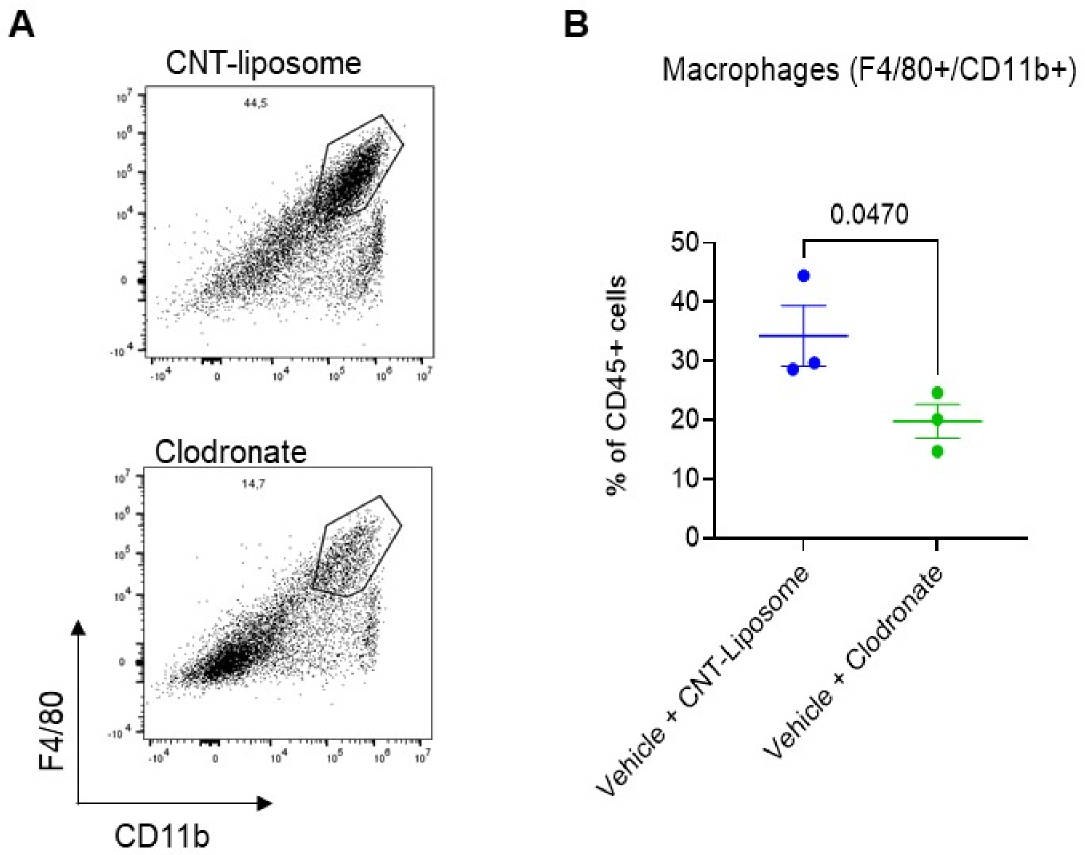


**Supplementary Figure S3. Depletion of macrophages by clodronate liposome.** (**A**) FACS plots showing depletion of F4/80$^+$/CD11b$^+$ macrophages by clodronate gated on live, single CD45+ cells. B16-OVA bearing mice were injected with clodronate (1mg/kg) or control (CNT) liposome every 4 days for 2 weeks once tumors reached ~100 mm3. Tumors were homogenized and macrophage markers (F4/80, CD11b) were analyzed by flow cytometry (**B**) Relative frequencies of macrophages (CD45+/CD11b+/F4/80+) in control-liposome and clodronate-treated tumors. N = 3, statistical significance by unpaired student t-test.

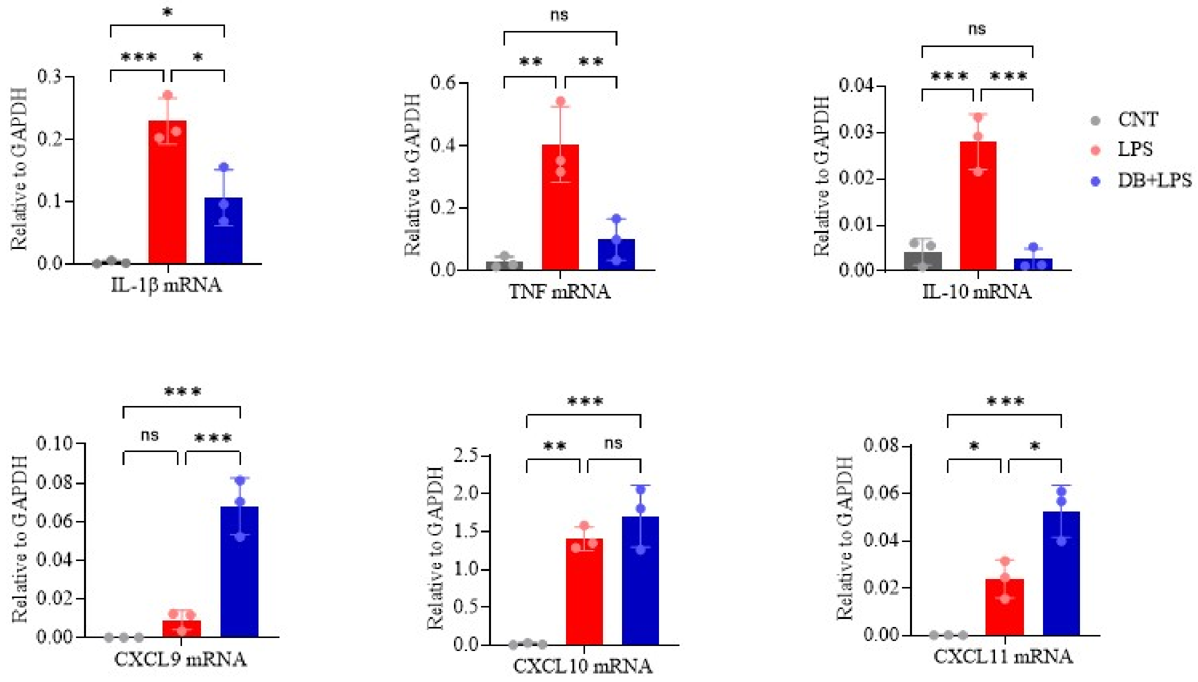


**Supplementary Figure S4. Confirmation of RNA-seq analysis by qPCR.** BMDMs were exposed to the drug vehicle or DB (500 nM) for 18~20 hrs and then activated by LPS (100 ng/mL, 5 h). RNAs from BMDMs were isolated and mRNA expression of representative cytokines/chemokines was analyzed by RT-qPCR. one-way ANOVA test followed by Tukey's Multiple comparisons test (n=3; * $p \leq 0.05$, ** $p \leq 0.01$, *** $p \leq 0.001$)

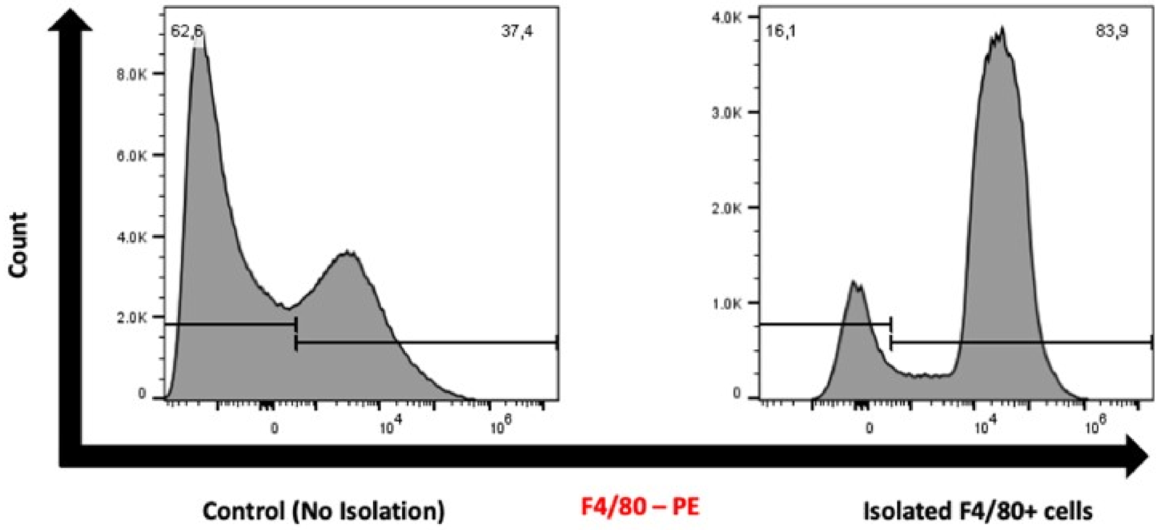


**Supplementary Figure S5. F4/80+ cell isolation efficiency from tumor tissue**. F4/80+ macrophages were isolated from murine vehicle tumor cell suspensions by magnetic-activated cell sorting of cells positive for F4/80 - PE antibody. Representative FACS plots shown of both tumor samples with no isolation and isolated F4/80+ cells, gated on single cells. PE-positive cells increased from 37.4% in control to 83.9% in isolated samples, with a more intense peak seen in the isolated sample.

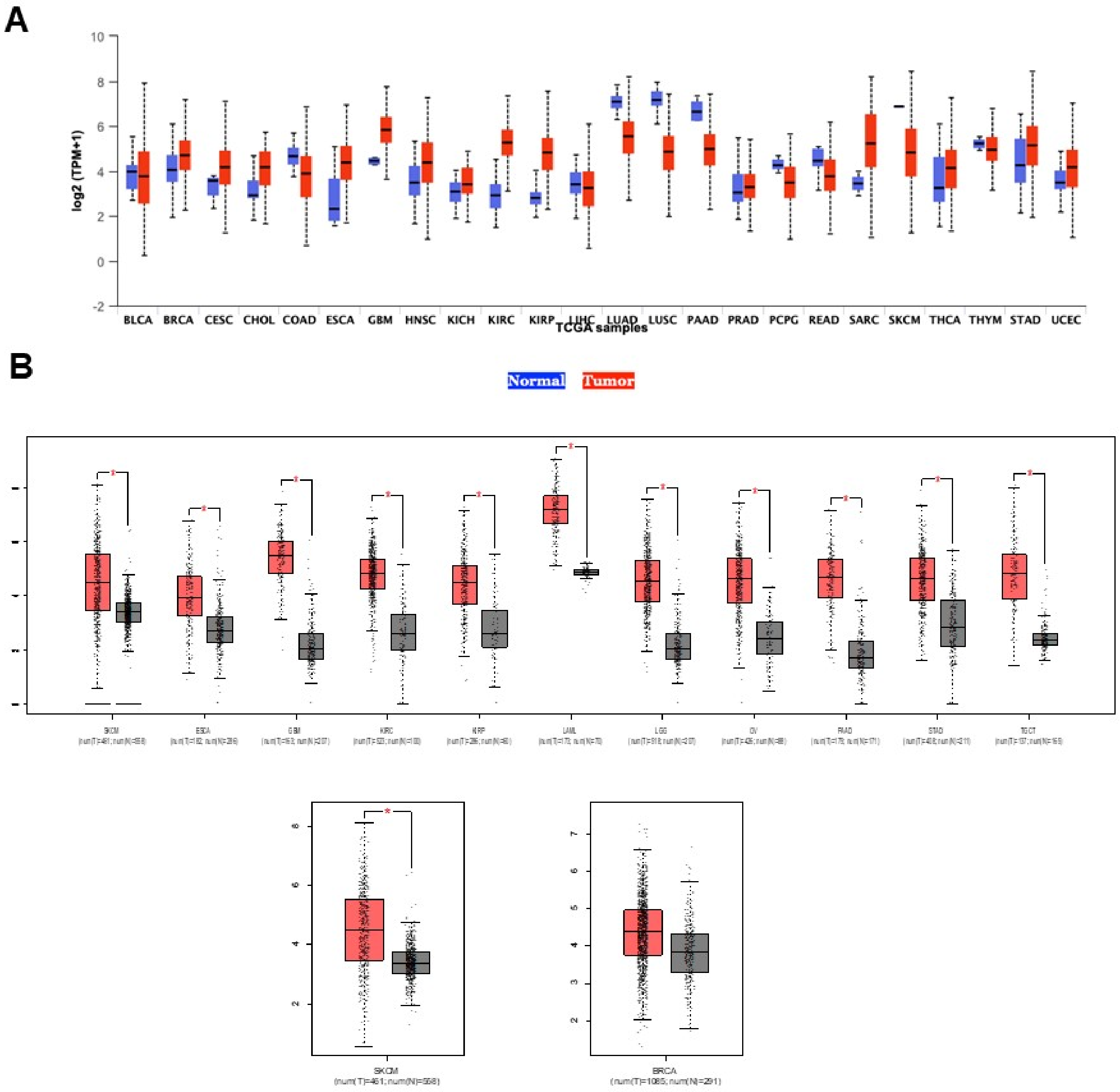


**Supplementary Figure S6. PU.1/SPI1 is over-expressed in multiple cancers.** (**A**) Box plots showing the relative mRNA levels of SPI1 in tumors vs. adjacent normal tissues based on data from the TCGA. (**B**) SPI1 is significantly over-expressed in multiple cancers including melanoma (SKCM) and breast cancer (BRCA). *, p< 0.01, unpaired student t-test.
