## Supplemental tables for "Inhibition of the transcription factor PU.1 suppresses tumor growth in mice by promoting the recruitment of cytotoxic lymphocytes through the CXCL9-CXCR3 axis"

**Supplementary Table S1**

| Target | Catalog # | Source | Fluorophore |
|---|---|---|---|
| CD3 | 53-0032-82 | ebioscience | AF-488 |
| F4/80 | MCA497PE | Bio-rad | PE |
| Tbet | 12-5825-82 | ebioscience | PE |
| GATA3 | 15-9966-42 | ebioscience | PE-Cy5 |
| CD8a | 100758 | BioLegend | AF-594 |
| CD80 | 25-0801-80 | ebioscience | PE-Cyanine7 |
| CD25 | 25-0251-82 | Invitrogen | PE-Cyanine7 |
| CD45 | 67-0451-82 | Invitrogen | Super Bright™ 702 |
| CD206 | 56-2061-82 | Invitrogen | AF700 |
| CD4 | 100536 | BioLegend | AF700 |
| FOXP3 | 126419 | BioLegend | BV421 |
| CD11b | 46-0112-82 | Invitrogen | Percp-eF710 |
| NK1.1 | 63-5941-82 | Invitrogen | SB600 |
| Granzyme B | 46-8898-82 | Invitrogen | Percp-eF710 |
| Fixable Viability Dye | 65-0865-14 | Invitrogen | AF780 |
| MHC Class II (I-A/I-E) | 12-5321-81 | ebioscience | PE |
| CD8a | 100712 | BioLegend | APC |
| CD163 | 61-1631-82 | Invitrogen | PE-eF610 |
| CD86 | 25-0862-80 | Invitrogen | PE-Cyanine7 |
| CD38 | 46-0381-80 | Invitrogen | PerCP-ef710 |
| CD138 | 142505 | Biolegend | APC |
| B220 | 12-0452-82 | Invitrogen | PE |
| CD4 (mAb) | Ab183685 | ABcam | N/A |
| CD8 (mAb) | Ab217344 | ABcam | N/A |
| Granzyme B (mAb) | Ab4059 | ABcam | N/A |
| Goat anti-Mouse IgG (HRP conjugated pAb) | Ab6721 | Abcam | N/A |

| | | | |
|---|---|---|---|
| CD163 (mAb) | Ab182422 | ABcam | N/A |
| CD68 (mAb) | Ab125212 | ABcam | N/A |

**Supplementary Table S2**

| Target ger | Primer`s name and It`s sequence | |
|---|---|---|
| GAPDH | Forward | GCATTGTGGAAGGGCTCATG |
| | Reverse | TTGCTGTTGAAGTCGCAGGAG |
| PU.1 | Forward | CTGGAACAGATGCACGTCC |
| | Reverse | CTGGTACAGGCGAATCTTTTTC |
| IL-1β | Forward | GTGGACCTTCCAGGATGAGG |
| | Reverse | GCTTGGGATCCACACTCTCC |
| TNF-α | Forward | ATGAGAAGTTCCCAAATGGCC |
| | Reverse | TCCACTTGGTGGTTTGCTACG |
| IL-10 | Forward | TGCACTACCAAAGCCACAAAGCAG |
| | Reverse | AGTAAGAGCAGGCAGCATAGCAGT |
| CXCL9 | Forward | AGGCACGGTCCACTACAAAT |
| | Reverse | TCCGGATCTAGGCAGGTTTG |
| CXCL10 | Forward | TGATGGTCAAGCCATGGTCC |
| | Reverse | GTCGCACCTCCACATAGCTT |
| CXCL11 | Forward | CCACGCTACCTTCTGTGGTT |
| | Reverse | ATGTTCGTGTGCCTCGTGAT |

**Supplementary Table S3**

| SYMBOL | ENTREZID | LPS vs CNT logFC | AveExpr | t | P.Value | adj.P.Val | DB+LPS vs LPS logFC | AveExpr | t | P.Value | adj.P.Val | CNT1 | CNT2 | CNT3 | LPS1 | LPS2 | LPS3 | DB+LPS1 | DB+LPS2 | DB+LPS3 | IL4-1 | IL4-2 | IL4-3 | DB+IL4-1 | DB+IL4-2 | DB+IL4-3 |
|---|---|---|---|---|---|---|---|---|---|---|---|---|---|---|---|---|---|---|---|---|---|---|---|---|---|---|
| Trip12 | 14897 | 1.000139 | 9.348034 | 53.24393 | 3.47E-24 | 4.54E-22 | -0.49954 | 9.348034 | -26.626 | 7.34E-18 | 2.60E-16 | 31672.19 | 32256.09 | 32392.63 | 68421.19 | 70282.96 | 66931.35 | 51141.23 | 50675.98 | 49321.76 | 35966.16 | 35604.71 | 36281.14 | 22454.24 | 22248.5 | 22830.24 |
| Slx4ip | 74243 | 1.00137 | 3.899671 | 13.80175 | 4.14E-12 | 3.63E-11 | -0.55532 | 3.899671 | -7.59261 | 1.69E-07 | 8.03E-07 | 714.5275 | 792.1272 | 764.6812 | 1609.101 | 1647.356 | 1582.621 | 1196.702 | 1125.954 | 1103.189 | 645.3709 | 698.4755 | 599.312 | 634.9228 | 587.9898 | 657.6147 |
| Cdkn2d | 12581 | 1.001508 | 1.579692 | 3.879694 | 0.000845 | 0.002458 | 0.074888 | 1.579692 | 0.320734 | 0.751531 | 0.827353 | 135.7825 | 196.2557 | 87.66493 | 289.6121 | 386.2087 | 204.2528 | 385.1617 | 316.3395 | 255.1125 | 165.3811 | 139.5426 | 105.9805 | 130.155 | 91.97167 | 119.0509 |
| Anxa7 | 11750 | 1.002178 | 7.269622 | 23.04249 | 1.45E-16 | 2.90E-15 | 0.509886 | 7.269622 | 13.41516 | 7.14E-12 | 6.89E-11 | 6087.952 | 6004.893 | 6383.916 | 13126.05 | 13381.08 | 12995.08 | 20487.48 | 19143.9 | 18897.86 | 6852.932 | 6241.29 | 7068.598 | 6478.549 | 6178.609 | 6218.994 |
| Stard7 | 99138 | 1.002182 | 5.759015 | 24.27664 | 4.96E-17 | 1.08E-15 | -3.11435 | 5.759015 | -47.3286 | 4.20E-23 | 8.30E-21 | 4500.113 | 4214.614 | 4620.202 | 9623.435 | 9523.204 | 9376.693 | 1264.922 | 1072.337 | 1099.742 | 4300.678 | 4196.191 | 4474.316 | 1795.472 | 1610.683 | 1706.397 |
| H2-K1 | 14972 | 1.003422 | 8.891709 | 15.01066 | 6.31E-09 | 3.49E-08 | 0.574522 | 8.891709 | 10.086 | 4.56E-07 | 2.03E-06 | 14651.15 | 15800.81 | 14971.61 | 32587.21 | 32211.21 | 32200.26 | 53696.66 | 50213.54 | 46985.52 | 26720.97 | 26715.16 | 28835.64 | 27307.52 | 26017.76 | 28913.99 |
| Rnf34 | 80751 | 1.004024 | 5.53892 | 19.6105 | 3.91E-15 | 5.77E-14 | -0.46075 | 5.53892 | -9.05553 | 9.28E-09 | 5.28E-08 | 2376.564 | 2511.363 | 2398.2 | 5157.174 | 5449.054 | 4992.697 | 4054.857 | 3782.67 | 3920.919 | 2563.023 | 2433.226 | 2613.687 | 1209.774 | 1321.405 | 1443.189 |
| Ccdc6 | 76551 | 1.004517 | 6.820302 | 25.99096 | 1.21E-17 | 2.95E-16 | -1.02887 | 6.820302 | -24.615 | 3.73E-17 | 1.13E-15 | 5735.511 | 5950.722 | 6267.609 | 12866.31 | 13261.7 | 12270.05 | 6796.469 | 6498.365 | 6273.24 | 5809.108 | 6147.5 | 6111.042 | 4583.792 | 4711.779 | 4752.319 |
| Gprc5c | 70355 | 1.00477 | 1.651582 | 3.177056 | 0.004476 | 0.011511 | 2.269448 | 1.651582 | 9.818947 | 2.27E-09 | 1.42E-08 | 37.84102 | 39.07354 | 60.75787 | 72.72771 | 112.3516 | 108.2134 | 447.6972 | 514.722 | 480.347 | 244.6102 | 240.1963 | 175.3902 | 489.7499 | 464.5748 | 581.4869 |
| Gm20257 | 100504501 | 1.005795 | 0.248063 | 2.966485 | 0.00728 | 0.017981 | 0.555812 | 0.248063 | 1.929339 | 0.06709 | 0.11118 | 29.67923 | 36.40943 | 45.13442 | 80.51997 | 81.45493 | 71.69139 | 177.6576 | 128.6805 | 80.44089 | 57.69108 | 67.86498 | 46.27316 | 88.43865 | 105.3351 | 73.69819 |
| Ttc9c | 70387 | 1.007032 | 5.908809 | 22.94123 | 1.59E-16 | 3.16E-15 | -1.25587 | 5.908809 | -25.6529 | 1.59E-17 | 5.24E-16 | 3285.491 | 3187.158 | 3247.942 | 7103.939 | 7116.071 | 6660.536 | 3096.927 | 3129.885 | 2871.74 | 3302.238 | 3458.064 | 3293.604 | 2717.403 | 2550.052 | 2560 |
| Sat1 | 20229 | 1.007448 | 7.207743 | 18.99127 | 7.48E-15 | 1.05E-13 | -0.04695 | 7.207743 | -0.94352 | 0.355974 | 0.453455 | 5275.483 | 5704.737 | 4654.053 | 11424.74 | 11487.95 | 10552.16 | 11608.86 | 11184.48 | 10876.76 | 12763.58 | 12354.48 | 12907.97 | 6706.32 | 6585.014 | 6341.284 |
| 4930404I0 | 67394 | 1.008029 | -0.15145 | 3.992061 | 0.000645 | 0.001911 | -0.13615 | -0.15145 | -0.57161 | 0.573558 | 0.667514 | 28.93725 | 48.84192 | 50.34224 | 96.10448 | 85.66811 | 81.16006 | 88.11817 | 77.74446 | 82.7392 | 36.92229 | 37.36386 | 41.04877 | 40.88202 | 33.01547 | 38.87377 |
| Kpna4 | 16649 | 1.008048 | 7.680404 | 43.99082 | 1.97E-22 | 1.43E-20 | -1.09211 | 7.680404 | -43.3308 | 2.71E-22 | 3.97E-20 | 11080.74 | 11063.14 | 11188.13 | 23745.6 | 24376.09 | 23475.55 | 11598.91 | 11333.27 | 11875.37 | 11438.22 | 11495.11 | 11626.5 | 7800.122 | 7937.076 | 7727.782 |
| Sumo1 | 22218 | 1.009045 | 5.39177 | 8.13796 | 1.71E-06 | 6.90E-06 | -0.52619 | 5.39177 | -4.24171 | 0.000938 | 0.002583 | 2230.394 | 2277.81 | 1439.962 | 4137.687 | 4541.814 | 3920.031 | 3096.927 | 3116.481 | 2871.74 | 2579.176 | 2523.205 | 2537.56 | 1619.429 | 1586.315 | 1551.711 |
| Ptpn2 | 19255 | 1.009399 | 5.386495 | 21.11859 | 8.66E-16 | 1.46E-14 | 0.008427 | 5.386495 | 0.188552 | 0.852225 | 0.914574 | 1963.281 | 1945.685 | 1880.022 | 4294.831 | 4149.988 | 4005.249 | 4350.479 | 4452.881 | 4211.655 | 2200.723 | 2194.555 | 2254.697 | 1235.638 | 1306.469 | 1265.827 |
| Inpp5k | 19062 | 1.009535 | 5.947716 | 15.32552 | 5.38E-13 | 5.46E-12 | 0.630899 | 5.947716 | 10.99047 | 3.00E-10 | 2.17E-09 | 2164.358 | 2087.77 | 2083.995 | 4449.378 | 4752.474 | 4444.866 | 7536.946 | 7696.702 | 6790.36 | 2959.168 | 3191.942 | 2724.892 | 3313.947 | 3262.243 | 3285.644 |
| Ptpn1 | 19246 | 1.009721 | 7.440535 | 31.23177 | 2.64E-19 | 9.46E-18 | -1.67162 | 7.440535 | -42.5498 | 3.98E-22 | 5.26E-20 | 10827.73 | 11111.09 | 10533.68 | 23165.08 | 23478.68 | 23153.61 | 7905.053 | 7842.808 | 7092.588 | 10482.85 | 10823.32 | 11136.16 | 5637.547 | 5946.715 | 5910.433 |
| Chst7 | 60322 | 1.009972 | 3.778326 | 11.51449 | 1.27E-10 | 8.95E-10 | -0.89642 | 3.778326 | -9.53875 | 3.78E-09 | 2.29E-08 | 497.1271 | 576.3347 | 486.063 | 1154.552 | 1077.171 | 1102.424 | 680.7839 | 609.8919 | 581.4727 | 928.4418 | 996.6239 | 1050.102 | 1041.24 | 1006.972 | 1077.937 |
| Plekha5 | 109135 | 1.010338 | 3.739964 | 11.65851 | 1.01E-10 | 7.22E-10 | 0.808296 | 3.739964 | 11.09848 | 2.51E-10 | 1.83E-09 | 480.0615 | 485.7551 | 525.1216 | 1062.344 | 1174.074 | 975.2734 | 2045.194 | 1948.973 | 1845.544 | 510.7584 | 478.8675 | 478.4047 | 644.1004 | 620.2192 | 663.2837 |
| Arid4a | 238247 | 1.010777 | 6.45402 | 24.54349 | 3.96E-17 | 8.78E-16 | -1.24016 | 6.45402 | -27.0075 | 5.46E-18 | 2.01E-16 | 4790.97 | 5057.359 | 4682.696 | 10942.92 | 10153.78 | 10127.42 | 4660.314 | 4672.71 | 4412.757 | 5149.891 | 5367.434 | 5186.325 | 3260.55 | 3343.209 | 3215.185 |
| AW04620 | 100502619 | 1.011682 | 1.56388 | 6.259255 | 3.07E-06 | 1.20E-05 | 0.826339 | 1.56388 | 6.240388 | 3.20E-06 | 1.27E-05 | 132.0726 | 145.6377 | 111.1001 | 283.1186 | 321.6065 | 235.3642 | 528.709 | 534.8283 | 479.1979 | 68.46009 | 86.92817 | 86.57559 | 110.1312 | 109.2655 | 127.1496 |
| Mab21l2 | 23937 | 1.011773 | -0.52339 | 2.679907 | 0.0139 | 0.032252 | 0.837027 | -0.52339 | 2.679738 | 0.013905 | 0.028763 | 23.74338 | 15.98463 | 16.49142 | 28.5716 | 39.32307 | 59.51738 | 61.11422 | 72.38277 | 93.0816 | 24.61486 | 26.68847 | 23.88292 | 97.61625 | 76.25001 | 77.74755 |
| Ktn1 | 16709 | 1.012292 | 5.261514 | 20.23438 | 2.07E-15 | 3.23E-14 | -1.41282 | 5.261514 | -24.6482 | 3.62E-17 | 1.10E-15 | 2192.553 | 2197.887 | 2196.831 | 4597.43 | 4961.728 | 4645.061 | 1809.265 | 1959.697 | 1773.147 | 2480.717 | 2401.963 | 2320.375 | 1389.154 | 1440.103 | 1300.652 |
| Psmb8 | 16913 | 1.014283 | 6.102817 | 15.00338 | 8.17E-13 | 8.05E-12 | 0.004047 | 6.102817 | 0.064133 | 0.949461 | 0.957984 | 2886.305 | 2966.925 | 3083.896 | 6635.105 | 6514.99 | 6103.237 | 7337.97 | 6917.917 | 5973.311 | 3834.534 | 3753.162 | 3895.155 | 2596.425 | 2177.449 | 2388.307 |
| Lgals3bp | 19039 | 1.014322 | 9.242145 | 33.76294 | 5.17E-20 | 2.17E-18 | 0.298326 | 9.242145 | 11.33542 | 1.70E-10 | 1.28E-09 | 22220.84 | 21792.38 | 22122.81 | 48202.89 | 46977.02 | 47496.22 | 62873.74 | 59605.87 | 59962.94 | 33230.83 | 32566.8 | 33651.03 | 29127.19 | 28127.61 | 31056.9 |
| Fndc7 | 320181 | 1.014759 | 0.888609 | 3.748379 | 0.001158 | 0.003298 | 0.430473 | 0.888609 | 1.850202 | 0.078178 | 0.126627 | 63.06836 | 51.50603 | 40.79457 | 90.90964 | 146.0571 | 105.5081 | 154.9174 | 183.6378 | 136.7495 | 159.2274 | 136.4925 | 179.1219 | 124.3147 | 130.4897 | 93.13508 |
| Gm10584 | 100043682 | 1.014887 | -0.6656 | 2.471239 | 0.021958 | 0.048465 | 1.797948 | -0.6656 | 5.792467 | 8.90E-06 | 3.33E-05 | 23.74338 | 9.768385 | 13.01954 | 29.87031 | 30.8967 | 36.52203 | 116.5434 | 99.19121 | 139.0478 | 15.38429 | 19.82572 | 17.16585 | 73.42077 | 85.68301 | 63.16988 |
| Bst2 | 69550 | 1.01543 | 7.463971 | 19.559 | 1.95E-14 | 2.55E-13 | -0.19239 | 7.463971 | -3.89217 | 0.000916 | 0.002528 | 8109.85 | 8377.722 | 8230.956 | 18044.27 | 17727.68 | 17530.57 | 17337.96 | 16739.19 | 14728.73 | 13952.78 | 13609.6 | 14996.23 | 3886.295 | 3674.15 | 4112.521 |
| Prkcg | 18752 | 1.015911 | -0.15273 | 3.555374 | 0.001836 | 0.005058 | -0.05162 | -0.15273 | -0.19465 | 0.847509 | 0.910779 | 31.16319 | 37.29747 | 48.6063 | 96.10448 | 109.5428 | 52.75404 | 73.90556 | 92.4891 | 81.59005 | 30.76858 | 34.31375 | 34.3317 | 53.39692 | 48.73712 | 47.78235 |
| Itprip | 414801 | 1.016224 | 4.284215 | 11.602 | 1.11E-10 | 7.86E-10 | -0.26399 | 4.284215 | -3.13001 | 0.004993 | 0.011668 | 987.5764 | 1006.144 | 953.8986 | 2249.364 | 1987.219 | 2130.452 | 1954.234 | 2014.654 | 1599.625 | 835.3669 | 972.9855 | 850.829 | 689.154 | 606.8558 | 594.4448 |
| Heca | 380629 | 1.016842 | 5.327224 | 18.01136 | 2.18E-14 | 2.81E-13 | -0.81662 | 5.327224 | -13.8802 | 3.71E-12 | 3.76E-11 | 2018.93 | 1864.873 | 1852.247 | 4271.454 | 4193.524 | 3951.142 | 2460.203 | 2491.844 | 2371.857 | 2170.723 | 2103.052 | 2125.58 | 1819.667 | 1866.16 | 2125.1 |
| Keap1 | 50868 | 1.01688 | 6.20572 | 19.77659 | 3.29E-15 | 4.94E-14 | -0.17354 | 6.20572 | -3.53826 | 0.001912 | 0.004922 | 3370.077 | 3473.993 | 3486.634 | 7551.994 | 7634.293 | 7123.148 | 7326.6 | 7055.98 | 6277.837 | 4201.449 | 4064.273 | 4125.028 | 2253.517 | 2385.761 | 2262.778 |
| Grina | 66168 | 1.017253 | 8.140109 | 27.12551 | 4.99E-18 | 1.35E-16 | 0.83847 | 8.140109 | 26.84427 | 6.19E-18 | 2.23E-16 | 9244.338 | 8956.721 | 9112.813 | 20101.42 | 19501.43 | 19458.13 | 37674.78 | 36893.77 | 35358.37 | 11882.06 | 11958.72 | 12303.44 | 17192.97 | 16606 | 17946.73 |
| Grhl1 | 195733 | 1.019277 | 1.288619 | 5.790135 | 8.95E-06 | 3.33E-05 | 1.023807 | 1.288619 | 7.24709 | 3.51E-07 | 1.59E-06 | 108.3292 | 138.5335 | 98.94853 | 262.3393 | 224.7032 | 251.5962 | 487.4925 | 548.2325 | 524.0149 | 59.22951 | 50.32684 | 31.34633 | 101.7879 | 99.04641 | 94.75482 |
| Olfr99 | 258508 | 1.019492 | 0.227555 | 3.844577 | 0.00092 | 0.002658 | 1.315293 | 0.227555 | 6.336201 | 2.58E-06 | 1.04E-05 | 43.77686 | 48.84192 | 52.07818 | 100.0006 | 94.09448 | 119.0348 | 258.6695 | 231.893 | 316.0178 | 59.99873 | 64.05234 | 49.25852 | 24.19548 | 31.4433 | 12.14805 |
| Lrmda | 76633 | 1.019691 | 1.331928 | 4.802629 | 9.18E-05 | 0.000302 | -0.63039 | 1.331928 | -2.91935 | 0.008107 | 0.017949 | 128.3627 | 140.3095 | 132.7993 | 223.378 | 349.6944 | 313.8189 | 170.5513 | 296.2332 | 149.3902 | 122.3051 | 130.3923 | 107.4731 | 63.40885 | 98.26033 | 92.32521 |
| Prdm9 | 213389 | 1.020258 | 2.033691 | 6.888926 | 7.59E-07 | 3.20E-06 | 0.626474 | 2.033691 | 5.045975 | 5.14E-05 | 0.000173 | 143.9443 | 134.9813 | 144.083 | 372.7295 | 273.8571 | 277.2969 | 496.02 | 489.2539 | 487.242 | 144.6123 | 152.5056 | 158.2243 | 249.4637 | 281.4176 | 285.8842 |
| Notch3 | 18131 | 1.021252 | 0.3779 | 2.916108 | 0.008167 | 0.019964 | 1.203537 | 0.3779 | 4.346545 | 0.000275 | 0.000829 | 37.84102 | 20.4248 | 15.62345 | 58.44191 | 42.13186 | 54.10671 | 106.5946 | 116.6167 | 145.9428 | 87.69045 | 89.21576 | 74.63413 | 220.2623 | 238.183 | 221.0946 |
| Ly96 | 17087 | 1.021565 | 2.981861 | 11.68989 | 9.62E-11 | 6.90E-10 | -2.09069 | 2.981861 | -17.2462 | 5.19E-14 | 7.69E-13 | 592.8426 | 656.2579 | 567.6521 | 1401.307 | 1217.611 | 1308.03 | 353.894 | 339.1267 | 276.9465 | 622.2945 | 643.5735 | 574.6828 | 171.8713 | 142.281 | 137.6779 |
| Mycbp2 | 105689 | 1.02174 | 8.135478 | 32.61728 | 1.07E-19 | 4.18E-18 | -0.91445 | 8.135478 | -27.3459 | 4.22E-18 | 1.59E-16 | 15363.45 | 15128.56 | 16217.14 | 34143.06 | 33539.77 | 33482.58 | 19242.45 | 19209.58 | 17507.39 | 14545.85 | 14301.21 | 13984.94 | 9620.624 | 9560.337 | 9648.794 |
| Stx3 | 20908 | 1.021939 | 5.520416 | 13.93531 | 3.43E-12 | 3.05E-11 | 2.035967 | 5.520416 | 35.92918 | 1.40E-20 | 1.09E-18 | 949.7354 | 875.6025 | 1082.358 | 2161.052 | 2071.483 | 2047.939 | 9054.853 | 9102.804 | 8633.606 | 2326.105 | 2145.753 | 2306.194 | 3868.774 | 3788.918 | 3779.665 |
| Daam2 | 76441 | 1.021993 | 2.007262 | 5.868787 | 7.47E-06 | 2.80E-05 | 0.57444 | 2.007262 | 3.920083 | 0.000767 | 0.002148 | 159.5259 | 124.3249 | 139.7431 | 319.4825 | 258.4087 | 348.9883 | 540.0791 | 434.2966 | 461.9605 | 231.5336 | 187.5819 | 180.6146 | 226.9369 | 173.7243 | 186.2702 |
| Phlpp2 | 244650 | 1.022704 | 4.605288 | 19.89291 | 2.93E-15 | 4.43E-14 | -0.98937 | 4.605288 | -17.9522 | 2.32E-14 | 3.68E-13 | 1420.893 | 1342.709 | 1345.353 | 2958.459 | 3091.074 | 2882.535 | 1615.974 | 1545.506 | 1513.438 | 1474.584 | 1480.067 | 1497.161 | 706.6749 | 689.3945 | 677.0515 |
| Ccdc74a | 72315 | 1.023183 | 1.290344 | 4.694584 | 0.000119 | 0.000386 | -0.85997 | 1.290344 | -3.72332 | 0.00123 | 0.003304 | 105.3613 | 97.68385 | 98.94853 | 210.3909 | 231.7252 | 216.4268 | 119.3859 | 176.9357 | 96.52907 | 163.0735 | 261.5471 | 226.1414 | 100.9536 | 106.1212 | 124.72 |
| Sdr39u1 | 654795 | 1.023311 | 3.890826 | 9.875975 | 2.05E-09 | 1.22E-08 | 0.120212 | 3.890826 | 1.282496 | 0.213423 | 0.296214 | 396.9597 | 428.0329 | 496.4786 | 848.0571 | 1022.4 | 987.4475 | 1186.753 | 1073.678 | 976.7823 | 751.5225 | 729.7392 | 782.1656 | 1656.973 | 1615.4 | 1580.867 |
| Atp2b4 | 381290 | 1.024344 | 3.436142 | 11.28937 | 1.83E-10 | 1.26E-09 | 0.621406 | 3.436142 | 7.996424 | 7.39E-08 | 3.69E-07 | 533.4842 | 574.5586 | 504.2903 | 1123.383 | 1203.567 | 1171.41 | 2019.612 | 1761.314 | 1820.262 | 103.0747 | 108.279 | 67.17071 | 1043.743 | 1065.928 | 1081.177 |
| Pik3r5 | 320207 | 1.024589 | 8.997601 | 48.13956 | 2.93E-23 | 2.83E-21 | -1.65511 | 8.997601 | -61.4469 | 1.66E-25 | 6.72E-23 | 40007.6 | 40832.74 | 38980.51 | 87636.89 | 87455.91 | 85103.09 | 29584.97 | 29363.28 | 27144.2 | 31414.72 | 31136.3 | 32092.67 | 8294.044 | 8631.973 | 8573.287 |
| Arl5b | 75869 | 1.024885 | 4.842007 | 13.32194 | 8.16E-12 | 6.85E-11 | 2.374917 | 4.842007 | 40.8632 | 9.34E-22 | 1.04E-19 | 819.8887 | 849.8495 | 814.1555 | 1915.596 | 1738.641 | 1739.531 | 9535.239 | 9522.356 | 9944.792 | 1016.132 | 1136.166 | 1027.712 | 950.2984 | 1080.864 | 893.2869 |
| Abca13 | 268379 | 1.026025 | 4.292655 | 12.73605 | 1.92E-11 | 1.53E-10 | 0.435153 | 4.292655 | 6.109792 | 4.30E-06 | 1.68E-05 | 912.6363 | 869.3862 | 945.2189 | 1803.907 | 1994.241 | 2154.8 | 2674.813 | 2934.183 | 2722.35 | 531.5272 | 657.299 | 601.5511 | 707.5092 | 798.6599 | 723.2141 |
| Krit1 | 79264 | 1.026692 | 5.795157 | 18.57883 | 1.17E-14 | 1.59E-13 | 0.083156 | 5.795157 | 1.628215 | 0.118152 | 0.179719 | 2281.591 | 2022.944 | 2136.941 | 4500.027 | 4874.656 | 4677.525 | 5342.52 | 5006.475 | 5098.803 | 3192.24 | 3037.911 | 2915.209 | 2478.785 | 2689.975 | 2507.358 |
| Gm15663 | 327812 | 1.027194 | -0.97833 | 2.572132 | 0.017631 | 0.039844 | 0.708872 | -0.97833 | 2.122247 | 0.045678 | 0.080134 | 23.0014 | 15.09659 | 15.62345 | 36.36386 | 44.94065 | 36.52203 | 51.16539 | 85.78699 | 66.65102 | 16.1535 | 10.67539 | 24.62926 | 22.52683 | 38.51805 | 37.25403 |
| Foxp1 | 108655 | 1.028799 | 6.506755 | 35.53233 | 1.77E-20 | 8.45E-19 | -2.57952 | 6.506755 | -62.7342 | 1.07E-25 | 5.12E-23 | 7353.029 | 7208.18 | 7661.568 | 16110.49 | 16382.27 | 15911.43 | 2768.616 | 2849.737 | 2783.255 | 5118.353 | 5138.675 | 5116.916 | 2653.16 | 2800.026 | 2848.314 |
| Gmppb | 331026 | 1.029757 | 4.406746 | 16.18324 | 1.84E-13 | 2.02E-12 | -0.85419 | 4.406746 | -12.8099 | 1.72E-11 | 1.55E-10 | 1268.045 | 1286.763 | 1239.461 | 2736.38 | 2907.098 | 2637.702 | 1755.257 | 1601.804 | 1435.295 | 994.5943 | 1008.062 | 1011.292 | 597.3781 | 609.214 | 553.1414 |
| Gm17359 | 100233207 | 1.030492 | -0.67443 | 3.232717 | 0.00393 | 0.010224 | -1.32667 | -0.67443 | -3.61433 | 0.001595 | 0.00418 | 22.25942 | 23.97694 | 37.32269 | 59.74062 | 63.19779 | 54.10671 | 35.53152 | 24.12759 | 17.23733 | 47.6913 | 41.1765 | 39.55609 | 46.72231 | 47.95104 | 30.77507 |
| E230008O | 319862 | 1.031626 | 2.125957 | 7.175801 | 4.09E-07 | 1.78E-06 | -0.73789 | 2.125957 | -4.94728 | 6.50E-05 | 0.000215 | 250.0475 | 210.4643 | 194.4252 | 492.2108 | 440.9801 | 500.4871 | 252.9844 | 349.8501 | 294.1838 | 149.2276 | 179.9566 | 129.117 | 221.931 | 275.915 | 279.4052 |
| Tfec | 21426 | 1.032384 | 4.614299 | 11.64721 | 1.03E-10 | 7.35E-10 | 0.562205 | 4.614299 | 7.305238 | 3.10E-07 | 1.42E-06 | 672.9765 | 673.1305 | 852.3462 | 1489.619 | 1683.87 | 1596.148 | 2363.557 | 2470.397 | 2453.447 | 3649.153 | 4001.746 | 3898.887 | 668.2959 | 622.5774 | 576.6276 |
| Myadm | 50918 | 1.032604 | 8.476731 | 39.81271 | 1.62E-21 | 9.67E-20 | -1.07659 | 8.476731 | -37.616 | 5.34E-21 | 4.70E-19 | 22234.94 | 21787.94 | 23243.36 | 50100.3 | 49739.47 | 47039.02 | 24503.96 | 24191.93 | 23661.11 | 18574.22 | 18129.1 | 18308.5 | 8862.221 | 8710.582 | 9523.264 |
| Alox12 | 11684 | 1.033193 | 1.041657 | 3.567051 | 0.001785 | 0.004928 | 2.665977 | 1.041657 | 12.91997 | 1.46E-11 | 1.33E-10 | 46.74479 | 58.61031 | 42.53051 | 119.4812 | 94.09448 | 106.8608 | 838.5439 | 705.0618 | 613.6491 | 68.46009 | 46.5142 | 50.00486 | 115.1371 | 155.6444 | 143.347 |
| Fam169b | 434197 | 1.033194 | -0.67947 | 2.583162 | 0.017209 | 0.038983 | 0.911017 | -0.67947 | 2.792102 | 0.010816 | 0.023129 | 29.67923 | 14.20856 | 18.22736 | 63.63675 | 30.8967 | 43.28537 | 73.90556 | 80.4253 | 106.8715 | 13.07665 | 19.82572 | 26.86829 | 41.71635 | 42.44846 | 59.12053 |
| Map4k3 | 225028 | 1.033676 | 5.802163 | 24.38521 | 4.52E-17 | 9.96E-16 | -2.46242 | 5.802163 | -42.1092 | 4.96E-22 | 6.24E-20 | 3881.301 | 3898.474 | 4279.958 | 8811.742 | 9103.29 | 8409.535 | 1716.883 | 1663.463 | 1585.835 | 3116.857 | 3186.604 | 3062.238 | 2355.305 | 2187.668 | 2261.968 |
| Zfp385c | 278304 | 1.033953 | 0.35537 | 4.269662 | 0.000331 | 0.001017 | -0.06681 | 0.35537 | -0.29907 | 0.767777 | 0.841598 | 69.74619 | 47.95389 | 45.13442 | 132.4683 | 106.734 | 116.3294 | 125.071 | 91.14868 | 137.8987 | 49.22973 | 50.32684 | 36.57072 | 69.24913 | 76.25001 | 86.65612 |
| Slc25a12 | 78830 | 1.033968 | 5.449299 | 26.77998 | 6.51E-18 | 1.72E-16 | -2.08357 | 5.449299 | -42.0497 | 5.11E-22 | 6.36E-20 | 3349.301 | 3274.185 | 3313.04 | 7124.718 | 7360.436 | 7259.768 | 1706.934 | 1885.973 | 1735.225 | 2567.638 | 2549.893 | 2631.599 | 1141.359 | 1064.356 | 1105.473 |
| Slc11a2 | 18174 | 1.03459 | 7.122365 | 26.64728 | 7.21E-18 | 1.87E-16 | -1.09865 | 7.122365 | -25.8953 | 1.31E-17 | 4.40E-16 | 8986.871 | 8926.528 | 8714.415 | 19563.75 | 19733.16 | 18980.63 | 9918.979 | 9334.697 | 9069.136 | 14257.39 | 13987.05 | 14462.6 | 1704.53 | 1525 | 1992.281 |
| Dync1h1 | 13424 | 1.035759 | 10.5074 | 22.30652 | 5.06E-10 | 3.28E-09 | -0.4425 | 10.5074 | -9.76049 | 1.64E-06 | 6.80E-06 | 72651.04 | 74017.72 | 72738.46 | 163232.2 | 162386 | 154427.3 | 127171.6 | 129378.8 | 113124 | 72183.85 | 72786.33 | 72396.59 | 47650.91 | 47519.48 | 48299.85 |
| Orai2 | 269717 | 1.036 | 5.272677 | 12.55297 | 2.53E-11 | 1.97E-10 | -0.13712 | 5.272677 | -1.75423 | 0.09374 | 0.147614 | 1919.504 | 1748.541 | 1443.433 | 3981.842 | 3522.223 | 3665.73 | 3629.9 | 3718.33 | 3247.514 | 1900.729 | 1907.082 | 2133.043 | 1454.232 | 1471.547 | 1577.627 |
| Ube2f | 67921 | 1.037292 | 5.998366 | 22.67546 | 2.02E-16 | 3.92E-15 | -0.27435 | 5.998366 | -6.21976 | 3.35E-06 | 1.33E-05 | 3229.1 | 3425.151 | 3097.784 | 7203.94 | 7087.983 | 7047.399 | 6316.083 | 6283.897 | 5779.103 | 3413.005 | 3340.635 | 3463.023 | 1685.34 | 1661.779 | 1725.834 |
| Col20a1 | 73368 | 1.037549 | 2.240151 | 5.763462 | 9.52E-06 | 3.53E-05 | 0.950738 | 2.240151 | 6.564054 | 1.55E-06 | 6.44E-06 | 108.3292 | 104.7881 | 135.4033 | 264.9367 | 262.6219 | 231.3062 | 598.3508 | 465.1263 | 478.0487 | 328.4546 | 329.412 | 291.0731 | 349.583 | 421.3403 | 368.491 |
| Sgms2 | 74442 | 1.039244 | 4.249053 | 13.60713 | 5.44E-12 | 4.68E-11 | 0.584677 | 4.249053 | 8.846471 | 1.38E-08 | 7.65E-08 | 666.2987 | 729.9648 | 763.8133 | 1493.516 | 1752.685 | 1490.64 | 2445.99 | 2343.057 | 2544.23 | 532.2964 | 517.7564 | 559.7559 | 1238.975 | 1260.877 | 1330.617 |
| Tent5c | 74645 | 1.039609 | 3.812097 | 7.93033 | 8.45E-08 | 4.03E-07 | 1.171234 | 3.812097 | 11.228 | 2.03E-10 | 1.50E-09 | 254.4994 | 239.7694 | 340.2441 | 549.354 | 619.3383 | 643.8698 | 1452.529 | 1511.996 | 1288.203 | 1399.201 | 1332.136 | 1345.653 | 1278.189 | 1363.067 | 1277.165 |
| Cd44 | 12505 | 1.03994 | 10.18338 | 48.61245 | 2.38E-23 | 2.37E-21 | 0.273554 | 10.18338 | 14.88337 | 9.56E-13 | 1.07E-11 | 44975.9 | 44666.38 | 46665.52 | 100691.5 | 101527.9 | 96913.23 | 125402.1 | 123159.3 | 126455.4 | 55356.52 | 53977.82 | 55241.19 | 51868.44 | 50351.74 | 54577.97 |
| Cited2 | 17684 | 1.040083 | 7.33444 | 25.57525 | 1.69E-17 | 3.98E-16 | 0.340299 | 7.33444 | 9.435728 | 4.56E-09 | 2.73E-08 | 6139.891 | 6067.055 | 5753.771 | 13336.44 | 13170.42 | 12949.09 | 18177.93 | 17169.46 | 16662.76 | 6106.024 | 6184.101 | 5943.862 | 10056.14 | 9807.953 | 10388.21 |
| 4930453N | 67609 | 1.041811 | 4.477147 | 14.46687 | 1.66E-12 | 1.55E-11 | -0.45885 | 4.477147 | -6.44539 | 2.02E-06 | 8.24E-06 | 957.8971 | 1191.743 | 1111.869 | 2331.183 | 2499.824 | 2279.245 | 1893.119 | 1698.314 | 1780.042 | 1197.667 | 1145.317 | 1095.629 | 818.4747 | 768.7888 | 796.9123 |
| Mkx | 210719 | 1.041911 | 0.457227 | 3.206745 | 0.004176 | 0.010806 | 1.619618 | 0.457227 | 6.547668 | 1.61E-06 | 6.66E-06 | 28.19527 | 40.84961 | 26.03909 | 46.75353 | 92.69009 | 74.39673 | 181.9214 | 230.5525 | 242.4718 | 29.23015 | 32.02617 | 20.15121 | 310.3696 | 282.9897 | 278.5954 |
| Cnp | 12799 | 1.041922 | 6.078469 | 24.43623 | 4.33E-17 | 9.58E-16 | -0.67625 | 6.078469 | -15.5821 | 3.88E-13 | 4.69E-12 | 3298.846 | 3320.363 | 3192.392 | 7242.901 | 7363.244 | 6974.355 | 4792.492 | 4841.603 | 4424.249 | 4380.676 | 4325.82 | 4269.818 | 2245.174 | 2209.678 | 2316.229 |
| Cd47 | 16423 | 1.042844 | 8.214725 | 38.92009 | 2.61E-21 | 1.49E-19 | -0.3471 | 8.214725 | -13.3716 | 7.60E-12 | 7.27E-11 | 13438.76 | 14140.18 | 14193.04 | 30185.9 | 31423.34 | 30150.96 | 25586.96 | 23890.34 | 25344.63 | 18629.6 | 18503.5 | 18236.85 | 8942.316 | 8976.277 | 9128.048 |
| Fbxl3 | 50789 | 1.042925 | 6.168074 | 23.72549 | 7.96E-17 | 1.66E-15 | 0.048423 | 6.168074 | 1.189858 | 0.24717 | 0.334835 | 2749.781 | 2825.727 | 2923.322 | 5980.556 | 6474.262 | 6239.856 | 6797.891 | 6625.705 | 6627.18 | 3741.459 | 3596.844 | 3662.297 | 3304.769 | 3325.129 | 3552.901 |
| Snx20 | 71607 | 1.043023 | 5.999702 | 19.56434 | 4.10E-15 | 6.04E-14 | -0.46207 | 5.999702 | -8.77226 | 1.59E-08 | 8.75E-08 | 3105.931 | 3414.495 | 3146.39 | 7075.368 | 7187.695 | 6954.065 | 5503.122 | 5617.708 | 4943.667 | 2795.325 | 2758.826 | 2952.526 | 2322.766 | 2250.555 | 2549.472 |
| Gm13570 | 667103 | 1.045392 | 1.14977 | 4.284248 | 0.000319 | 0.000984 | -0.46979 | 1.14977 | -1.94954 | 0.064492 | 0.107525 | 108.3292 | 157.1822 | 81.58914 | 300.0018 | 233.1296 | 208.3108 | 152.0749 | 215.8079 | 186.1632 | 99.22866 | 112.8541 | 92.54632 | 52.5626 | 131.2758 | 73.69819 |
| Bcl6 | 12053 | 1.045477 | 7.207231 | 29.61972 | 7.99E-19 | 2.54E-17 | -1.04396 | 7.207231 | -27.3732 | 4.13E-18 | 1.56E-16 | 8330.96 | 8350.193 | 8550.369 | 18904.01 | 18969.17 | 17733.47 | 9961.617 | 9341.399 | 8807.128 | 4775.283 | 4899.241 | 4843.755 | 7548.156 | 7496.87 | 7813.628 |
| Cyp26c1 | 546726 | 1.045558 | 0.311762 | 3.793691 | 0.001039 | 0.002983 | 1.807712 | 0.311762 | 8.763214 | 1.62E-08 | 8.89E-08 | 30.42121 | 32.85729 | 46.87036 | 79.22126 | 92.69009 | 67.63339 | 298.4648 | 302.9353 | 271.2007 | 41.53758 | 32.7887 | 36.57072 | 86.77 | 58.9562 | 69.64884 |
| Mcmbp | 210711 | 1.046656 | 7.098525 | 30.9879 | 3.11E-19 | 1.09E-17 | -1.69294 | 7.098525 | -41.4304 | 6.98E-22 | 8.05E-20 | 9447.641 | 9227.571 | 9388.827 | 21035.19 | 21346.81 | 19590.69 | 6691.296 | 6731.598 | 6478.939 | 8035.214 | 7574.189 | 7342.505 | 3631.825 | 3836.083 | 3833.926 |
| Cplx2 | 12890 | 1.047326 | -1.28584 | 2.69982 | 0.013298 | 0.031008 | 0.464481 | -1.28584 | 1.385361 | 0.180251 | 0.256981 | 17.06556 | 17.7607 | 9.547666 | 31.16902 | 30.8967 | 33.81669 | 34.11026 | 49.5956 | 55.15947 | 9.230573 | 12.20045 | 7.463413 | 40.88202 | 38.51805 | 42.11325 |
| Slc2a1 | 20525 | 1.050531 | 6.733942 | 18.20244 | 1.76E-14 | 2.33E-13 | 0.641668 | 6.733942 | 12.95835 | 1.38E-11 | 1.27E-10 | 4480.822 | 4616.894 | 4869.309 | 10459.8 | 10379.89 | 9990.804 | 17450.24 | 17367.84 | 15406.73 | 3530.694 | 3326.146 | 3458.545 | 3929.68 | 3483.918 | 3991.041 |
| Pdk3 | 236900 | 1.051435 | 4.114639 | 15.9653 | 2.41E-13 | 2.60E-12 | -0.597 | 4.114639 | -9.01128 | 1.01E-08 | 5.69E-08 | 928.2179 | 915.5641 | 856.686 | 1914.297 | 2068.674 | 2004.654 | 1429.788 | 1327.018 | 1352.556 | 1171.514 | 1172.768 | 1267.287 | 390.465 | 373.3892 | 444.6188 |
| Fbxw11 | 103583 | 1.052938 | 7.433754 | 31.43518 | 2.31E-19 | 8.36E-18 | -0.72947 | 7.433754 | -21.1969 | 8.03E-16 | 1.77E-14 | 9806.018 | 10284.33 | 9676.993 | 22709.23 | 22279.33 | 20902.77 | 14161.44 | 13721.9 | 13420.99 | 8274.44 | 8003.492 | 7899.276 | 4991.778 | 4753.442 | 4900.525 |
| Psmd10 | 53380 | 1.053548 | 4.47322 | 16.95867 | 7.26E-14 | 8.61E-13 | 0.027457 | 4.47322 | 0.478097 | 0.637442 | 0.726339 | 959.3811 | 1058.538 | 988.6174 | 2292.222 | 2143.107 | 2207.554 | 2409.037 | 2332.334 | 2296.013 | 1253.05 | 1312.31 | 1378.492 | 589.0348 | 622.5774 | 597.6843 |
| Arhgap17 | 70497 | 1.053692 | 7.155026 | 35.42 | 1.89E-20 | 8.96E-19 | -2.63382 | 7.155026 | -61.3795 | 1.70E-25 | 6.77E-23 | 9613.103 | 9991.282 | 9804.585 | 21930 | 21622.07 | 21554.76 | 3862.987 | 3612.437 | 3453.213 | 10754.39 | 11153.49 | 11187.66 | 5050.181 | 5352.437 | 5363.771 |
| Ccnl2 | 56036 | 1.054254 | 5.868334 | 19.29082 | 5.45E-15 | 7.88E-14 | -0.24698 | 5.868334 | -4.71308 | 0.000114 | 0.000365 | 2633.29 | 2697.85 | 2681.158 | 5692.242 | 6142.825 | 5940.917 | 5139.279 | 5660.601 | 4810.365 | 2776.864 | 2954.795 | 2956.258 | 2135.877 | 2246.624 | 2166.403 |
| 4931440P | 71004 | 1.054404 | 1.741474 | 6.812019 | 8.97E-07 | 3.74E-06 | -0.48951 | 1.741474 | -3.21239 | 0.004122 | 0.00983 | 166.2037 | 166.0625 | 153.6306 | 342.8592 | 320.2021 | 423.385 | 270.0396 | 317.68 | 225.2345 | 139.997 | 105.9914 | 112.6975 | 159.3564 | 169.7938 | 181.4109 |
| Nfix | 18032 | 1.054521 | 3.450709 | 12.5836 | 2.42E-11 | 1.89E-10 | -2.09487 | 3.450709 | -18.044 | 2.10E-14 | 3.34E-13 | 596.5525 | 562.1261 | 583.2756 | 1467.541 | 1214.802 | 1195.758 | 306.9923 | 324.3821 | 307.9737 | 1001.517 | 947.8221 | 975.468 | 525.626 | 485.7991 | 553.1414 |
| Tbrg3 | 21378 | 1.055045 | 4.634815 | 12.84566 | 1.63E-11 | 1.31E-10 | -0.80294 | 4.634815 | -9.43107 | 4.60E-09 | 2.75E-08 | 1206.461 | 1228.152 | 1405.243 | 2749.367 | 2800.364 | 2939.347 | 1770.891 | 1722.442 | 1576.641 | 1101.515 | 1131.591 | 883.668 | 1076.282 | 1253.802 | 1112.762 |
| Nup210l | 77595 | 1.05584 | 1.653323 | 7.817344 | 1.06E-07 | 5.01E-07 | -1.47841 | 1.653323 | -9.05712 | 9.25E-09 | 5.27E-08 | 213.6905 | 193.5916 | 200.501 | 515.5875 | 415.701 | 424.7377 | 170.5513 | 172.9144 | 160.8818 | 173.8425 | 167.7561 | 126.878 | 101.7879 | 111.6237 | 82.60677 |
| Itga4 | 16401 | 1.056064 | 8.246356 | 53.37311 | 3.30E-24 | 4.36E-22 | -0.9311 | 8.246356 | -43.9447 | 2.01E-22 | 3.17E-20 | 23150.54 | 24046.21 | 24104.38 | 53060.06 | 53404.94 | 51605.63 | 28514.76 | 29042.92 | 28495.61 | 14741.23 | 14991.3 | 14685.76 | 3740.288 | 3722.887 | 3654.135 |
| Csta2 | 76770 | 1.05836 | -0.56203 | 4.753635 | 0.000103 | 0.000337 | -1.82952 | -0.56203 | -6.328 | 2.63E-06 | 1.06E-05 | 86.06977 | 73.7069 | 85.92899 | 200.0012 | 144.6527 | 208.3108 | 45.48035 | 69.70193 | 47.11538 | 15.38429 | 18.30067 | 23.13658 | 6.674615 | 17.29382 | 2.429611 |
| Sppl2a | 66552 | 1.059457 | 8.409926 | 47.27798 | 4.29E-23 | 3.95E-21 | -0.40022 | 8.409926 | -18.3531 | 1.49E-14 | 2.45E-13 | 17447.68 | 17945.41 | 17101.61 | 39001.53 | 40032.29 | 37777.3 | 30444.83 | 30293.53 | 31012.26 | 19672.66 | 19182.15 | 19469.8 | 8540.17 | 8432.308 | 8765.226 |
| Acpp | 56318 | 1.06006 | 2.84199 | 8.50457 | 2.68E-08 | 1.35E-07 | -1.41002 | 2.84199 | -9.55184 | 3.69E-09 | 2.24E-08 | 328.6975 | 295.7156 | 316.8089 | 741.5629 | 610.9119 | 753.4359 | 316.9412 | 269.4248 | 244.7701 | 289.2246 | 323.3118 | 241.0682 | 892.7298 | 865.477 | 775.0458 |
| Klhdc8b | 78267 | 1.060317 | 1.450976 | 5.383184 | 2.31E-05 | 8.18E-05 | 0.667046 | 1.450976 | 4.086048 | 0.000515 | 0.001485 | 88.29571 | 90.57957 | 66.83366 | 197.4038 | 190.9978 | 158.2621 | 261.512 | 313.6587 | 321.7636 | 124.6127 | 140.3051 | 144.0439 | 206.9131 | 205.1676 | 167.6431 |
| Brd2 | 14312 | 1.060611 | 8.78345 | 34.56141 | 3.17E-20 | 1.40E-18 | -0.38661 | 8.78345 | -13.0047 | 1.29E-11 | 1.19E-10 | 20217.49 | 19571.4 | 20279.24 | 44244.42 | 45555.77 | 44101.03 | 36648.63 | 36702.09 | 33418.59 | 25569.46 | 24902.63 | 24729.27 | 15854.71 | 15851.36 | 16048.39 |
| Fzd5 | 14367 | 1.062267 | 5.085239 | 20.10511 | 2.36E-15 | 3.63E-14 | 0.519464 | 5.085239 | 11.22999 | 2.02E-10 | 1.50E-09 | 1282.143 | 1216.608 | 1286.331 | 2735.081 | 2842.496 | 2873.066 | 4185.613 | 4182.116 | 4194.418 | 1329.972 | 1395.426 | 1315.8 | 1707.033 | 1687.719 | 1752.559 |
| Rptoros | 319454 | 1.062733 | 0.346472 | 4.993992 | 5.81E-05 | 0.000196 | -0.37633 | 0.346472 | -1.83025 | 0.081214 | 0.130716 | 59.35846 | 55.05817 | 45.13442 | 110.3903 | 112.3516 | 135.2668 | 83.85439 | 105.8933 | 95.37991 | 33.07622 | 38.88892 | 32.09267 | 119.3087 | 120.2706 | 118.2411 |
| Caml | 12328 | 1.062965 | 4.357814 | 14.64982 | 1.30E-12 | 1.24E-11 | -1.35268 | 4.357814 | -16.2935 | 1.61E-13 | 2.13E-12 | 1020.966 | 1272.554 | 1033.752 | 2375.339 | 2412.751 | 2572.774 | 950.8235 | 999.9546 | 1030.793 | 1153.052 | 1109.478 | 1224 | 846.8418 | 918.1445 | 889.2376 |
| Tm6sf2 | 107770 | 1.063592 | -0.5626 | 2.589016 | 0.01699 | 0.038538 | 0.372807 | -0.5626 | 1.041471 | 0.309312 | 0.403993 | 13.35565 | 28.41712 | 30.37894 | 51.94837 | 68.81537 | 33.81669 | 71.06304 | 69.70193 | 60.90525 | 24.61486 | 19.82572 | 34.3317 | 54.23125 | 75.46393 | 36.44416 |
| Cpne3 | 70568 | 1.063974 | 8.067474 | 38.38103 | 3.50E-21 | 1.94E-19 | 0.143922 | 8.067474 | 5.766692 | 9.45E-06 | 3.52E-05 | 10051.61 | 10074.76 | 10423.45 | 22793.64 | 22675.37 | 22685.59 | 26055.98 | 25854.05 | 26219.13 | 15246.6 | 15464.83 | 14687.25 | 11572.11 | 11810.11 | 11508.26 |
| Slc10a1 | 20493 | 1.064524 | 0.84079 | 5.120336 | 4.30E-05 | 0.000147 | -0.63618 | 0.84079 | -3.04081 | 0.006137 | 0.014012 | 100.1674 | 91.4676 | 61.62584 | 216.8844 | 167.123 | 177.1995 | 143.5473 | 138.0634 | 101.1257 | 89.99809 | 102.1787 | 105.9805 | 67.58048 | 82.53868 | 65.59949 |
| Usp32 | 237898 | 1.065339 | 6.505853 | 25.824 | 1.38E-17 | 3.31E-16 | -0.37284 | 6.505853 | -9.28986 | 5.98E-09 | 3.51E-08 | 3680.225 | 3479.321 | 3582.979 | 8145.504 | 8145.493 | 7742.67 | 6624.497 | 6310.706 | 6343.339 | 5201.428 | 5466.562 | 5476.652 | 4908.345 | 4582.076 | 4827.637 |
| Ap3b1 | 11774 | 1.065451 | 7.678875 | 32.59808 | 1.08E-19 | 4.22E-18 | -0.43914 | 7.678875 | -13.7108 | 4.70E-12 | 4.66E-11 | 9042.519 | 9440.7 | 9580.648 | 21361.17 | 21037.84 | 20203.45 | 16173.95 | 16530.08 | 15341.23 | 10425.16 | 10315.48 | 10303.24 | 8446.726 | 8515.633 | 8627.548 |
| Pkp4 | 227937 | 1.065535 | 6.847191 | 27.32975 | 4.27E-18 | 1.17E-16 | -2.45117 | 6.847191 | -45.5985 | 9.23E-23 | 1.63E-20 | 7465.068 | 7907.063 | 7073.084 | 16661.14 | 17326.02 | 16133.27 | 3129.616 | 3336.31 | 3056.754 | 6643.705 | 6749.134 | 7274.588 | 5763.53 | 5537.166 | 5689.339 |
| Marcks | 17118 | 1.065804 | 9.583669 | 42.94035 | 2.15E-20 | 1.01E-18 | -1.12312 | 9.583669 | -40.2953 | 7.11E-20 | 4.59E-18 | 43823.61 | 47080.06 | 42944.53 | 100348.7 | 100658.6 | 97513.82 | 49069.03 | 47654.67 | 45915.66 | 60607.95 | 60089.48 | 61089.52 | 15725.39 | 15944.9 | 15993.32 |
| Dnah1 | 110084 | 1.066658 | 0.519035 | 3.725513 | 0.001223 | 0.003472 | 0.735231 | 0.519035 | 3.110987 | 0.005218 | 0.012134 | 43.03488 | 52.39406 | 41.66254 | 87.01351 | 98.30767 | 123.0928 | 147.8111 | 182.2974 | 196.5056 | 59.99873 | 30.50111 | 32.83902 | 142.6699 | 169.7938 | 116.6213 |
| Csmd2 | 329942 | 1.067033 | 0.148327 | 3.209139 | 0.004153 | 0.010751 | 0.879887 | 0.148327 | 3.259687 | 0.00369 | 0.008913 | 40.80894 | 37.29747 | 22.56721 | 59.74062 | 75.83734 | 89.27607 | 130.756 | 107.2337 | 190.7598 | 33.07622 | 22.11331 | 31.34633 | 146.0072 | 128.1315 | 114.1917 |
| Tlr1 | 21897 | 1.067294 | 7.27752 | 33.69408 | 5.40E-20 | 2.25E-18 | -0.87769 | 7.27752 | -26.4008 | 8.75E-18 | 3.05E-16 | 9803.792 | 9579.233 | 8934.011 | 21253.38 | 21436.69 | 20726.93 | 12185.89 | 11726.01 | 11916.74 | 11059 | 10549.57 | 10906.28 | 2618.118 | 2574.421 | 2787.573 |
| Akna | 100182 | 1.06755 | 7.281431 | 25.76775 | 1.45E-17 | 3.45E-16 | -1.11835 | 7.281431 | -24.7413 | 3.35E-17 | 1.03E-15 | 8919.351 | 9272.861 | 9116.285 | 20800.13 | 20561.75 | 19688.08 | 10085.27 | 10479.42 | 8838.156 | 8274.44 | 8762.208 | 8503.812 | 4424.436 | 4549.846 | 4564.429 |
| Slc25a37 | 67712 | 1.069098 | 6.111398 | 25.68616 | 1.54E-17 | 3.68E-16 | -1.31251 | 6.111398 | -28.2137 | 2.20E-18 | 9.06E-17 | 4700.448 | 4403.765 | 4541.217 | 10692.27 | 9896.773 | 10013.8 | 4522.452 | 4233.052 | 4077.204 | 2354.565 | 2405.775 | 2308.433 | 2664.84 | 2532.758 | 2547.852 |
| Cdk3 | 69681 | 1.069203 | -1.19273 | 2.632068 | 0.015453 | 0.035493 | -0.31353 | -1.19273 | -0.79961 | 0.432745 | 0.531751 | 19.2915 | 13.32052 | 16.49142 | 35.06515 | 36.51428 | 39.22736 | 15.63387 | 30.8297 | 52.86116 | 17.69193 | 22.11331 | 14.18048 | 36.71038 | 38.51805 | 31.58494 |
| Mapk1ip1l | 218975 | 1.069809 | 7.795657 | 33.05459 | 8.06E-20 | 3.22E-18 | -0.40312 | 7.795657 | -12.7974 | 1.76E-11 | 1.57E-10 | 10565.06 | 10274.56 | 9807.189 | 23132.61 | 23725.85 | 21942.98 | 17878.04 | 18450.91 | 17683.21 | 10972.07 | 10962.86 | 10760 | 9090.826 | 8756.174 | 9153.964 |
| Slpi | 20568 | 1.070728 | 1.879883 | 6.331362 | 2.61E-06 | 1.03E-05 | 1.219913 | 1.879883 | 9.200043 | 7.07E-09 | 4.10E-08 | 180.3013 | 175.8309 | 256.919 | 490.9121 | 407.2746 | 451.791 | 1227.969 | 1018.721 | 1041.135 | 102.3055 | 67.10245 | 127.6244 | 58.40288 | 43.23454 | 61.55014 |
| Sesn3 | 75747 | 1.072283 | 3.116615 | 8.486992 | 2.77E-08 | 1.40E-07 | 0.638353 | 3.116615 | 6.082388 | 4.58E-06 | 1.78E-05 | 245.5956 | 200.6959 | 237.8237 | 515.5875 | 491.5383 | 531.5984 | 810.1187 | 940.9761 | 757.2935 | 433.837 | 404.9023 | 387.3511 | 926.1029 | 954.3043 | 943.4989 |
| Ss18l1 | 269397 | 1.073282 | 3.240466 | 12.60521 | 2.34E-11 | 1.83E-10 | -1.36391 | 3.240466 | -13.7214 | 4.63E-12 | 4.60E-11 | 496.3851 | 428.0329 | 460.0239 | 1045.461 | 1081.384 | 994.2108 | 397.953 | 461.1051 | 402.2045 | 550.7575 | 620.6977 | 581.3998 | 494.7559 | 499.9485 | 471.3445 |
| Il18rap | 16174 | 1.073734 | 2.376186 | 8.389188 | 3.36E-08 | 1.69E-07 | 0.581015 | 2.376186 | 5.424908 | 2.10E-05 | 7.43E-05 | 191.431 | 190.0395 | 200.501 | 480.5224 | 411.4878 | 416.6217 | 622.5123 | 751.9766 | 663.0628 | 159.2274 | 146.4053 | 164.1951 | 347.08 | 308.9305 | 303.7013 |
| Colgalt2 | 269132 | 1.074303 | 0.56497 | 6.307767 | 2.75E-06 | 1.08E-05 | -2.47296 | 0.56497 | -9.53831 | 3.78E-09 | 2.29E-08 | 107.5872 | 104.7881 | 105.8923 | 211.6896 | 247.1736 | 262.4175 | 48.32287 | 37.53181 | 47.11538 | 72.30616 | 80.06542 | 61.94632 | 62.57452 | 86.46909 | 52.64157 |
| Plscr1 | 22038 | 1.074962 | 4.27029 | 14.11115 | 2.69E-12 | 2.43E-11 | 0.796305 | 4.27029 | 12.36082 | 3.39E-11 | 2.88E-10 | 828.7925 | 883.5948 | 856.686 | 1994.817 | 1800.435 | 1974.895 | 3733.652 | 3318.884 | 3379.667 | 763.0607 | 741.9396 | 782.1656 | 480.5723 | 514.8841 | 413.8437 |
| Cyp4f37 | 677156 | 1.075347 | -0.30428 | 2.674502 | 0.014068 | 0.032598 | 0.961854 | -0.30428 | 2.958687 | 0.007411 | 0.016575 | 17.80754 | 13.32052 | 18.22736 | 54.54578 | 29.4923 | 31.11136 | 61.11422 | 101.8721 | 70.09849 | 34.61465 | 44.22662 | 44.78048 | 117.6401 | 146.2114 | 138.4878 |
| Morc3 | 338467 | 1.076541 | 7.012495 | 29.44801 | 9.02E-19 | 2.83E-17 | -0.66873 | 7.012495 | -18.0637 | 2.05E-14 | 3.28E-13 | 7468.778 | 7458.606 | 7297.888 | 16665.04 | 17143.45 | 16273.95 | 11459.63 | 11042.39 | 10315.97 | 6410.633 | 6368.633 | 6099.847 | 3332.302 | 3130.181 | 3198.988 |
| Snord3a | 19850 | 1.07756 | 4.59263 | 15.04041 | 7.78E-13 | 7.69E-12 | -0.71982 | 4.59263 | -9.86017 | 2.11E-09 | 1.33E-08 | 1282.143 | 1188.191 | 1185.646 | 2849.368 | 2655.711 | 2745.916 | 1922.966 | 1762.655 | 1564.001 | 1319.203 | 1400.001 | 1248.629 | 770.9181 | 867.0491 | 788.0038 |
| Heg1 | 77446 | 1.077933 | 5.305409 | 24.65982 | 3.59E-17 | 8.03E-16 | -3.95611 | 5.305409 | -47.2723 | 4.31E-23 | 8.45E-21 | 6255.64 | 6167.403 | 5581.045 | 13637.74 | 13972.33 | 13005.9 | 994.8826 | 961.0824 | 792.9174 | 2511.485 | 2633.771 | 2801.019 | 359.5949 | 371.8171 | 398.4562 |
| Gtf2f1 | 98053 | 1.07859 | 5.944792 | 22.77466 | 1.85E-16 | 3.62E-15 | -0.36992 | 5.944792 | -8.04196 | 6.74E-08 | 3.38E-07 | 2786.88 | 2929.627 | 2927.661 | 6663.677 | 6350.675 | 6444.109 | 5484.646 | 5239.709 | 4960.905 | 3080.704 | 3100.438 | 3299.575 | 2223.481 | 2104.343 | 2256.299 |
| Lilrb4b | 14727 | 1.078737 | 8.527331 | 35.9849 | 1.36E-20 | 6.68E-19 | 0.553742 | 8.527331 | 21.85877 | 4.28E-16 | 1.01E-14 | 13838.68 | 12907.59 | 13737.35 | 30288.5 | 30738 | 30384.98 | 47688.99 | 45629.3 | 46069.65 | 18984.98 | 18078.77 | 18286.11 | 15024.56 | 14315.35 | 15117.04 |
| Ppp6r1 | 243819 | 1.080668 | 8.158444 | 29.91284 | 6.51E-19 | 2.12E-17 | 0.00263 | 8.158444 | 0.079835 | 0.937112 | 0.948071 | 13201.32 | 13269.91 | 12872.86 | 29536.54 | 30959.89 | 28449.31 | 31715.44 | 31919.46 | 29158.67 | 10623.62 | 10999.46 | 10730.15 | 11326.82 | 11105.78 | 11224.8 |
| Prkg1 | 19091 | 1.082089 | 0.54113 | 3.488689 | 0.002151 | 0.005859 | 1.210059 | 0.54113 | 4.979126 | 6.02E-05 | 0.0002 | 25.96933 | 23.08891 | 34.71878 | 54.54578 | 77.24174 | 58.16471 | 132.1773 | 174.2548 | 148.2411 | 76.92145 | 84.64059 | 93.29266 | 225.2683 | 193.3763 | 239.7216 |
| Zfp319 | 79233 | 1.082575 | 5.981808 | 20.44718 | 1.67E-15 | 2.66E-14 | -0.11068 | 5.981808 | -2.22712 | 0.036826 | 0.066755 | 2542.768 | 2464.297 | 2604.777 | 5783.152 | 5817.005 | 5616.276 | 5889.705 | 5479.644 | 5244.746 | 3096.088 | 3451.201 | 3474.965 | 2916.807 | 2869.988 | 2800.531 |
| Gm9903 | 433873 | 1.082843 | -0.35637 | 3.502735 | 0.00208 | 0.005679 | -0.70474 | -0.35637 | -2.23182 | 0.036468 | 0.066199 | 43.77686 | 27.52908 | 44.26645 | 120.78 | 68.81537 | 75.74939 | 51.16539 | 48.25518 | 64.35271 | 29.99936 | 34.31375 | 25.3756 | 60.90587 | 38.51805 | 28.34546 |
| Ly6a2 | 546643 | 1.083396 | 3.56182 | 12.46792 | 2.88E-11 | 2.22E-10 | -0.53352 | 3.56182 | -6.2015 | 3.49E-06 | 1.38E-05 | 887.409 | 755.7178 | 915.7079 | 1968.843 | 1726.002 | 2106.104 | 1471.005 | 1427.549 | 1276.712 | 491.528 | 482.6801 | 467.2096 | 183.5519 | 135.9923 | 162.7839 |
| Spint2 | 20733 | 1.085046 | 0.983965 | 3.039691 | 0.006153 | 0.015439 | 2.924372 | 0.983965 | 11.71492 | 9.24E-11 | 7.27E-10 | 31.16319 | 29.30515 | 28.643 | 89.61093 | 49.15383 | 68.98605 | 513.0752 | 611.2323 | 501.0318 | 46.15287 | 61.00223 | 87.32193 | 371.2755 | 321.5078 | 304.5112 |
| Hipk2 | 15258 | 1.085463 | 7.009109 | 24.69878 | 3.47E-17 | 7.80E-16 | -0.37392 | 7.009109 | -8.77693 | 1.58E-08 | 8.67E-08 | 4371.751 | 4329.171 | 4148.895 | 9692.266 | 9725.437 | 9714.86 | 8065.655 | 8093.467 | 7278.752 | 7688.298 | 7734.32 | 7765.681 | 10429.09 | 10920.26 | 11016.67 |
| Utrn | 22288 | 1.08597 | 7.095697 | 21.33608 | 7.03E-16 | 1.22E-14 | 0.080684 | 7.095697 | 1.739627 | 0.096328 | 0.151038 | 5227.254 | 5104.425 | 4919.652 | 11326.04 | 11753.38 | 11534.2 | 13082.71 | 13447.11 | 11676.57 | 5773.724 | 5859.264 | 5958.042 | 8752.924 | 8231.071 | 8101.132 |
| Vps37c | 107305 | 1.088071 | 5.604649 | 22.51292 | 2.34E-16 | 4.48E-15 | -0.31511 | 5.604649 | -6.76839 | 9.87E-07 | 4.20E-06 | 2189.585 | 2258.273 | 2273.212 | 4924.705 | 5308.614 | 5025.161 | 4320.633 | 4396.583 | 4034.685 | 2676.097 | 2684.861 | 2624.882 | 1657.808 | 1560.374 | 1629.459 |
| 1700109H | 77036 | 1.088481 | 0.200841 | 4.666855 | 0.000127 | 0.000411 | 0.092724 | 0.200841 | 0.447629 | 0.658931 | 0.745144 | 63.81034 | 42.62568 | 49.47427 | 109.0916 | 132.0132 | 114.9768 | 135.0198 | 156.8293 | 109.1698 | 39.99915 | 49.56431 | 53.73657 | 39.21337 | 40.8763 | 36.44416 |
| Gm5150 | 381484 | 1.090344 | 4.162546 | 20.48236 | 1.62E-15 | 2.58E-14 | -6.17406 | 4.162546 | -29.2218 | 1.06E-18 | 4.74E-17 | 2859.594 | 2682.754 | 2532.735 | 6185.752 | 6186.361 | 6009.903 | 85.27565 | 92.4891 | 85.03751 | 1009.979 | 1001.199 | 934.4192 | 991.1804 | 867.8352 | 1111.142 |
| Ptprj | 19271 | 1.090984 | 9.909959 | 33.302 | 6.90E-13 | 6.88E-12 | -1.34778 | 9.909959 | -34.6129 | 4.42E-13 | 5.30E-12 | 63515.04 | 65228.83 | 58293.71 | 142729.4 | 144659.7 | 137932.9 | 59485.45 | 58244.01 | 56121.31 | 59907.96 | 59669.33 | 61440.3 | 20232.43 | 19972.79 | 20948.1 |
| Rn18s | 19791 | 1.091086 | 7.410896 | 3.563869 | 0.005036 | 0.012834 | -0.68139 | 7.410896 | -2.20347 | 0.051771 | 0.089258 | 9543.356 | 10398 | 3438.896 | 15903.99 | 16107.01 | 16167.08 | 10545.76 | 11180.46 | 9623.029 | 9827.484 | 11289.99 | 10826.43 | 8773.782 | 8829.28 | 8651.844 |
| Sphk1 | 20698 | 1.091177 | -0.15097 | 2.678109 | 0.013955 | 0.032375 | 2.139985 | -0.15097 | 7.195581 | 3.92E-07 | 1.77E-06 | 17.80754 | 14.20856 | 17.35939 | 36.36386 | 64.60218 | 22.99535 | 169.13 | 175.5952 | 179.2683 | 72.30616 | 51.85189 | 61.94632 | 68.41481 | 52.66754 | 72.88832 |
| Irak3 | 73914 | 1.091493 | 7.313281 | 34.05306 | 4.32E-20 | 1.86E-18 | -0.70181 | 7.313281 | -21.5332 | 5.82E-16 | 1.34E-14 | 11728.49 | 12113.69 | 11774.88 | 27401.46 | 27308.47 | 26272.87 | 17633.58 | 17969.69 | 16275.49 | 8076.752 | 7992.054 | 8259.759 | 1713.707 | 1799.343 | 1677.241 |
| Txlnb | 378431 | 1.093065 | 1.924042 | 11.41943 | 1.48E-10 | 1.03E-09 | -2.43748 | 1.924042 | -16.6734 | 1.02E-13 | 1.41E-12 | 316.0838 | 317.9165 | 339.3761 | 714.29 | 712.0284 | 791.3106 | 136.441 | 119.2975 | 166.6276 | 325.3777 | 340.0874 | 356.0048 | 52.5626 | 47.95104 | 55.88105 |
| Gm9975 | 327988 | 1.094254 | 3.31058 | 13.1392 | 1.06E-11 | 8.80E-11 | -0.34173 | 3.31058 | -4.28692 | 0.000317 | 0.000946 | 441.4785 | 420.0405 | 443.5325 | 941.5641 | 959.202 | 1084.84 | 797.3273 | 814.9764 | 823.9446 | 566.9111 | 548.2575 | 559.0096 | 350.4173 | 388.3248 | 382.2588 |
| Polr2g | 67710 | 1.09464 | 4.997547 | 19.86527 | 3.01E-15 | 4.55E-14 | -1.02167 | 4.997547 | -17.4188 | 4.25E-14 | 6.44E-13 | 1619.744 | 1681.05 | 1630.915 | 3807.815 | 3610.7 | 3818.581 | 2036.667 | 1919.484 | 1812.218 | 1735.348 | 1759.914 | 1751.663 | 1168.892 | 1192.487 | 1323.328 |
| Armcx3 | 71703 | 1.095872 | 4.503073 | 9.124847 | 8.14E-09 | 4.44E-08 | 0.645891 | 4.503073 | 6.342638 | 2.54E-06 | 1.02E-05 | 787.2416 | 703.3237 | 479.9872 | 1571.438 | 1505.512 | 1367.547 | 2465.888 | 2503.908 | 2268.433 | 1400.74 | 1250.546 | 1359.087 | 1451.729 | 1450.322 | 1576.817 |
| Sult6b1 | 73671 | 1.096898 | 2.71914 | 9.607078 | 3.33E-09 | 1.92E-08 | -0.25 | 2.71914 | -2.33213 | 0.029557 | 0.055228 | 347.989 | 319.6926 | 289.9018 | 742.8616 | 822.9756 | 638.4592 | 679.3627 | 634.0195 | 607.9033 | 291.5323 | 304.2486 | 270.1755 | 185.2206 | 238.183 | 237.292 |
| Mitf | 17342 | 1.097597 | 7.639121 | 34.35744 | 3.59E-20 | 1.57E-18 | -0.20909 | 7.639121 | -6.93399 | 6.88E-07 | 2.99E-06 | 7000.588 | 6939.993 | 7121.691 | 16178.02 | 16460.92 | 15502.92 | 14587.82 | 14624 | 14057.62 | 10828.23 | 10972.78 | 10630.14 | 13803.94 | 13666.83 | 13570.19 |
| Stau1 | 20853 | 1.098423 | 6.66512 | 30.47738 | 4.41E-19 | 1.49E-17 | -0.73069 | 6.66512 | -19.9028 | 2.90E-15 | 5.63E-14 | 4304.23 | 4466.816 | 4321.621 | 10218.24 | 9732.459 | 9952.929 | 6291.922 | 6345.557 | 6090.525 | 4849.897 | 4850.44 | 4887.789 | 6185.7 | 6135.375 | 6343.714 |
| Lrp10 | 65107 | 1.09866 | 7.597472 | 31.42817 | 2.32E-19 | 8.39E-18 | -0.41297 | 7.597472 | -12.166 | 4.56E-11 | 3.78E-10 | 8377.705 | 8526.024 | 8409.757 | 19849.47 | 18953.72 | 19044.21 | 15541.49 | 15268.74 | 14425.35 | 9643.642 | 9418.744 | 9918.875 | 8654.473 | 8540.002 | 9157.203 |
| Morf4l1b | 627352 | 1.099304 | 0.563305 | 5.368206 | 2.40E-05 | 8.46E-05 | -1.24701 | 0.563305 | -5.36109 | 2.44E-05 | 8.57E-05 | 69.00421 | 84.36332 | 121.5157 | 205.196 | 202.2329 | 197.4895 | 109.4371 | 83.10615 | 76.99342 | 57.69108 | 68.62751 | 46.27316 | 65.91183 | 62.88661 | 46.16261 |
| Macir | 52392 | 1.099657 | 6.382016 | 29.59955 | 8.11E-19 | 2.57E-17 | -1.24825 | 6.382016 | -30.4184 | 4.59E-19 | 2.28E-17 | 5339.293 | 5745.586 | 5313.71 | 12329.94 | 13125.48 | 12040.1 | 5616.823 | 5316.113 | 5438.953 | 4052.222 | 3976.583 | 4177.272 | 1980.692 | 2089.408 | 2028.725 |
| Cd80 | 12519 | 1.100091 | 4.529945 | 13.62803 | 5.28E-12 | 4.56E-11 | 2.528929 | 4.529945 | 42.73662 | 3.63E-22 | 4.89E-20 | 464.4799 | 507.068 | 480.8552 | 1101.305 | 1137.56 | 1078.076 | 6631.603 | 6703.449 | 6543.292 | 925.365 | 828.8678 | 837.3949 | 1636.115 | 1601.25 | 1679.671 |
| Itgal | 16408 | 1.100637 | 8.45508 | 31.47029 | 8.75E-14 | 1.02E-12 | -1.66242 | 8.45508 | -37.9628 | 7.59E-15 | 1.33E-13 | 41521.24 | 41888.61 | 38645.48 | 93708.36 | 96236.19 | 89748.15 | 32235.62 | 31209.04 | 28703.61 | 14819.69 | 13987.81 | 15376.12 | 2308.583 | 2403.055 | 2410.984 |

| | | | | | | | | | | | | | | | | | | | | | | | | | | |
|---|---|---|---|---|---|---|---|---|---|---|---|---|---|---|---|---|---|---|---|---|---|---|---|---|---|---|
| Map2k4 | 26398 | 1.102986 | 6.704588 | 27.14564 | 4.91E-18 | 1.33E-16 | 0.164429 | 6.704588 | 4.490036 | 0.000194 | 0.000603 | 4405.882 | 4425.078 | 4668.809 | 10158.5 | 10860.19 | 9948.871 | 12386.29 | 11892.22 | 11754.71 | 4026.068 | 4126.801 | 3938.443 | 4054.829 | 4188.248 | 4340.905 |
| Rtp4 | 67775 | 1.10341 | 7.411431 | 37.1292 | 7.03E-21 | 3.73E-19 | -0.61296 | 7.411431 | -20.6152 | 1.42E-15 | 2.95E-14 | 9523.323 | 9503.75 | 8929.671 | 21420.91 | 21963.34 | 20804.03 | 14607.72 | 14561 | 14390.88 | 10215.94 | 9954.039 | 10291.3 | 3855.425 | 3592.398 | 3698.678 |
| B3galt2 | 26878 | 1.103895 | 2.694886 | 11.26513 | 1.91E-10 | 1.31E-09 | -0.48991 | 2.694886 | -5.10426 | 4.47E-05 | 0.000152 | 281.9527 | 316.1405 | 327.2245 | 667.5365 | 751.3515 | 696.6239 | 538.6579 | 513.3815 | 510.2251 | 250.7639 | 236.3836 | 225.3951 | 320.3815 | 302.6418 | 345.0047 |
| Rab9 | 56382 | 1.104638 | 4.816621 | 18.7937 | 9.24E-15 | 1.28E-13 | -0.95527 | 4.816621 | -15.3952 | 4.93E-13 | 5.86E-12 | 1187.169 | 1209.504 | 1253.348 | 2755.861 | 2956.252 | 2671.519 | 1428.367 | 1538.804 | 1509.99 | 2326.105 | 2128.978 | 2259.921 | 1163.886 | 1254.588 | 1168.643 |
| Wnt4 | 22417 | 1.104883 | -0.09795 | 3.893682 | 0.000817 | 0.00238 | -1.07072 | -0.09795 | -3.46835 | 0.002257 | 0.005722 | 51.93865 | 42.62568 | 41.66254 | 93.50706 | 103.9252 | 119.0348 | 73.90556 | 36.19139 | 51.712 | 32.30701 | 48.80178 | 42.54145 | 33.37308 | 65.24486 | 64.78962 |
| Col1a1 | 12842 | 1.10494 | -0.25094 | 2.999121 | 0.006755 | 0.016803 | 1.108324 | -0.25094 | 3.813605 | 0.000991 | 0.002714 | 22.25942 | 19.53677 | 22.56721 | 49.35095 | 63.19779 | 39.22736 | 123.6497 | 111.255 | 103.424 | 36.92229 | 19.0632 | 37.31706 | 90.10731 | 101.4047 | 67.21923 |
| Atp2a1 | 11937 | 1.105348 | 0.284547 | 4.955182 | 6.38E-05 | 0.000213 | -0.29626 | 0.284547 | -1.40243 | 0.175167 | 0.250779 | 55.64856 | 56.83424 | 49.47427 | 142.858 | 126.3956 | 105.5081 | 109.4371 | 93.82952 | 111.4681 | 36.92229 | 48.80178 | 39.55609 | 82.59836 | 55.81187 | 68.83897 |
| Zfp691 | 195522 | 1.105484 | 2.48779 | 7.867781 | 9.60E-08 | 4.55E-07 | 0.279592 | 2.48779 | 2.301209 | 0.031547 | 0.058377 | 278.2428 | 261.9703 | 237.8237 | 576.6269 | 589.846 | 626.2852 | 756.1108 | 872.6146 | 659.6153 | 126.1512 | 90.74081 | 153.7463 | 266.1503 | 324.6521 | 303.7013 |
| Prickle1 | 106042 | 1.105855 | 2.384041 | 14.82509 | 1.03E-12 | 9.94E-12 | -3.46145 | 2.384041 | -23.3857 | 1.07E-16 | 2.95E-15 | 526.8063 | 567.4544 | 540.7451 | 1283.125 | 1241.485 | 1224.164 | 140.7048 | 113.9358 | 103.424 | 316.1471 | 298.1484 | 295.5511 | 110.9655 | 117.9124 | 131.199 |
| Srgap2 | 14270 | 1.107237 | 8.368711 | 38.40849 | 3.44E-21 | 1.92E-19 | -1.29088 | 8.368711 | -39.609 | 1.80E-21 | 1.83E-19 | 18664.53 | 18493.33 | 18193.51 | 42469.09 | 43873.31 | 41115.69 | 18435.17 | 18869.12 | 16963.83 | 24001.8 | 24651.76 | 24120.26 | 8611.922 | 8749.886 | 8939.348 |
| Serpind1 | 15160 | 1.108494 | 0.625815 | 4.427926 | 0.000226 | 0.000708 | -0.57226 | 0.625815 | -2.31675 | 0.030532 | 0.056747 | 77.90798 | 46.17782 | 65.96569 | 102.598 | 160.1011 | 188.0208 | 122.2284 | 95.16994 | 90.78329 | 98.45945 | 85.40312 | 85.82924 | 67.58048 | 89.61342 | 80.98703 |
| Ppp4r2 | 232314 | 1.109331 | 5.966771 | 5.904519 | 0.000133 | 0.000429 | 0.369865 | 5.966771 | 2.230684 | 0.049024 | 0.08517 | 3012.442 | 3218.239 | 1614.423 | 5762.373 | 6127.377 | 5514.826 | 7876.628 | 7801.255 | 7653.376 | 3357.621 | 3382.574 | 3421.975 | 1876.401 | 1897.603 | 1959.076 |
| Ppp1r15a | 17872 | 1.109331 | 5.255424 | 10.16611 | 1.23E-09 | 7.52E-09 | 2.699545 | 5.255424 | 33.8032 | 5.04E-20 | 3.35E-18 | 767.9501 | 831.2007 | 716.9429 | 1837.673 | 1692.296 | 1792.285 | 12978.95 | 12197.84 | 10988.23 | 1379.971 | 1520.481 | 1646.429 | 2613.112 | 2432.926 | 2441.759 |
| Sdk1 | 330222 | 1.111475 | -0.09919 | 3.057362 | 0.005907 | 0.014869 | 1.114162 | -0.09919 | 3.925929 | 0.000756 | 0.00212 | 51.19667 | 15.09659 | 30.37894 | 64.93546 | 57.58021 | 86.57074 | 127.9135 | 170.2336 | 167.7767 | 25.38408 | 12.96297 | 19.40487 | 102.6222 | 87.25517 | 84.22651 |
| AU040972 | 104522 | 1.11363 | 0.995945 | 5.041045 | 5.20E-05 | 0.000176 | -0.85288 | 0.995945 | -3.69548 | 0.001314 | 0.003511 | 105.3613 | 79.03511 | 126.7236 | 251.9496 | 223.2988 | 234.0115 | 129.3347 | 144.7655 | 133.302 | 93.07495 | 87.6907 | 40.30243 | 118.4744 | 107.6933 | 89.8956 |
| Mob1b | 68473 | 1.113837 | 5.989149 | 23.88233 | 6.95E-17 | 1.47E-15 | -0.10426 | 5.989149 | -2.39295 | 0.025976 | 0.04945 | 2606.578 | 2666.769 | 2560.51 | 5840.295 | 6451.792 | 5839.467 | 5884.02 | 5782.579 | 5808.982 | 3093.011 | 3240.743 | 3299.575 | 2667.343 | 2568.918 | 2795.672 |
| Gm5475 | 432982 | 1.113861 | -0.9075 | 2.672792 | 0.014121 | 0.032714 | -0.15152 | -0.9075 | -0.38996 | 0.70043 | 0.781913 | 20.03348 | 23.97694 | 23.43518 | 64.93546 | 30.8967 | 67.63339 | 34.11026 | 48.25518 | 64.35271 | 33.07622 | 41.1765 | 16.41951 | 13.34923 | 28.29897 | 20.24676 |
| Scarna3a | 100217414 | 1.114595 | -0.54605 | 3.774662 | 0.001087 | 0.003113 | -0.8727 | -0.54605 | -2.82754 | 0.009985 | 0.021541 | 40.80894 | 32.85729 | 30.37894 | 81.81868 | 77.24174 | 82.51273 | 65.378 | 28.14886 | 50.56285 | 39.22994 | 37.36386 | 32.09267 | 34.2074 | 24.36856 | 16.19741 |
| Dlgap4 | 228836 | 1.11561 | 6.895751 | 23.61242 | 8.78E-17 | 1.82E-15 | -0.16384 | 6.895751 | -3.70006 | 0.0013 | 0.003478 | 4966.077 | 4896.625 | 4798.136 | 11228.64 | 11678.95 | 11051.3 | 10956.5 | 10915.05 | 9742.541 | 5199.89 | 4967.869 | 5488.594 | 6038.858 | 5831.161 | 6192.268 |
| Washc4 | 319277 | 1.115979 | 7.849341 | 46.61258 | 5.79E-23 | 4.98E-21 | -0.28902 | 7.849341 | -12.7124 | 1.99E-11 | 1.77E-10 | 10418.15 | 10264.8 | 10314.08 | 23818.33 | 24439.29 | 23531.01 | 20264.34 | 20563.41 | 20153.89 | 12995.11 | 12759.38 | 13079.63 | 7649.109 | 7672.952 | 7631.408 |
| Gm19967 | 100503922 | 1.116139 | 1.185071 | 6.14267 | 3.99E-06 | 1.54E-05 | -0.95935 | 1.185071 | -4.9366 | 6.66E-05 | 0.000221 | 97.19948 | 126.989 | 134.5353 | 250.6509 | 282.2834 | 284.0602 | 140.7048 | 170.2336 | 128.7054 | 83.84438 | 138.7801 | 102.2488 | 93.44461 | 80.18043 | 102.8535 |
| Cds1 | 74596 | 1.116215 | 5.079025 | 15.42348 | 4.75E-13 | 4.87E-12 | 0.850076 | 5.079025 | 13.97105 | 3.27E-12 | 3.35E-11 | 1423.119 | 1394.215 | 1443.433 | 3359.761 | 3477.283 | 3043.502 | 6287.658 | 6493.003 | 5757.27 | 1719.964 | 1701.962 | 1693.448 | 793.4449 | 616.2888 | 655.185 |
| Adgb | 215772 | 1.117127 | 3.208655 | 16.89079 | 7.86E-14 | 9.28E-13 | -2.63417 | 3.208655 | -25.841 | 1.36E-17 | 4.55E-16 | 888.8929 | 933.3248 | 980.8057 | 2113 | 2096.762 | 2269.776 | 345.3664 | 328.4033 | 403.3536 | 136.9202 | 153.2681 | 154.4926 | 459.7141 | 522.7449 | 460.8162 |
| Prdm1 | 12142 | 1.117551 | 5.549872 | 21.29299 | 7.32E-16 | 1.26E-14 | 0.24834 | 5.549872 | 5.286507 | 2.90E-05 | 0.000101 | 2332.045 | 2375.494 | 2404.276 | 5371.461 | 5640.051 | 5464.778 | 6800.733 | 7070.725 | 6480.088 | 1446.892 | 1551.744 | 1564.331 | 1293.207 | 1176.766 | 1385.688 |
| Vaultrc5 | 378472 | 1.119801 | 0.417795 | 3.636223 | 0.001514 | 0.004228 | -0.43595 | 0.417795 | -1.45762 | 0.159502 | 0.231939 | 41.55092 | 47.95389 | 58.15396 | 145.4554 | 68.81537 | 139.3248 | 90.96069 | 67.02109 | 102.2748 | 68.46009 | 73.9652 | 119.4146 | 85.10135 | 77.82218 | 76.93768 |
| Clint1 | 216705 | 1.120065 | 7.690505 | 33.06948 | 7.99E-20 | 3.21E-18 | -0.11949 | 7.690505 | -3.80757 | 0.001005 | 0.002749 | 9093.716 | 9505.526 | 9129.304 | 21258.57 | 22702.05 | 20415.81 | 20851.32 | 20630.43 | 20025.19 | 9629.796 | 9573.537 | 9527.792 | 7211.922 | 7317.643 | 7611.971 |
| Chmp4b | 75608 | 1.12276 | 7.982454 | 18.91089 | 2.09E-11 | 1.65E-10 | -0.04762 | 7.982454 | -0.87783 | 0.394769 | 0.493644 | 11059.97 | 11297.58 | 10695.99 | 26259.9 | 26051.53 | 24542.8 | 27251.26 | 27126.11 | 23420.94 | 12793.57 | 12495.54 | 13153.52 | 8541.839 | 7931.574 | 8887.516 |
| Rnf225 | 381845 | 1.122822 | 1.359038 | 5.896803 | 7.00E-06 | 2.64E-05 | 2.392771 | 1.359038 | 17.70382 | 3.07E-14 | 4.80E-13 | 97.19948 | 87.91546 | 89.40087 | 225.9754 | 235.9384 | 182.6101 | 1186.753 | 1205.039 | 1119.278 | 59.99873 | 39.65145 | 53.73657 | 55.8999 | 91.97167 | 72.88832 |
| Cd46 | 17221 | 1.123334 | -0.41141 | 2.960383 | 0.007382 | 0.018206 | 0.184851 | -0.41141 | 0.548515 | 0.589031 | 0.682532 | 13.35565 | 35.5214 | 33.85081 | 67.53288 | 51.96262 | 52.75404 | 52.58665 | 77.74446 | 73.54596 | 48.46051 | 43.46409 | 25.3756 | 41.71635 | 53.45362 | 48.59222 |
| Jarid2 | 16468 | 1.126133 | 7.2473 | 28.43354 | 1.87E-18 | 5.55E-17 | -0.24421 | 7.2473 | -6.5431 | 1.62E-06 | 6.73E-06 | 8517.197 | 7937.257 | 8454.024 | 19823.5 | 19231.79 | 19041.5 | 18041.49 | 17197.61 | 15943.38 | 4692.208 | 4790.962 | 4648.213 | 4733.137 | 4869.782 | 4766.087 |
| Gm11696 | 100036768 | 1.12618 | 2.352441 | 10.39865 | 8.18E-10 | 5.13E-09 | -0.75068 | 2.352441 | -6.79455 | 9.32E-07 | 3.98E-06 | 329.4395 | 249.5378 | 295.1097 | 727.2771 | 664.279 | 649.2805 | 453.3822 | 399.4457 | 410.2485 | 229.9951 | 237.1462 | 204.4975 | 147.6759 | 135.2062 | 144.1569 |
| Rgcc | 66214 | 1.12664 | 1.822863 | 4.924512 | 6.86E-05 | 0.000229 | 3.983386 | 1.822863 | 25.82657 | 1.38E-17 | 4.59E-16 | 54.16459 | 67.49066 | 52.07818 | 133.767 | 154.4835 | 116.3294 | 2322.34 | 2135.292 | 2176.501 | 63.8448 | 55.66453 | 41.04877 | 465.5544 | 433.1315 | 489.1616 |
| Polm | 54125 | 1.12698 | 4.264775 | 12.68635 | 2.07E-11 | 1.64E-10 | -0.18251 | 4.264775 | -2.19354 | 0.039474 | 0.070771 | 812.4689 | 913.788 | 861.8938 | 1975.337 | 1971.771 | 2066.876 | 1883.171 | 2111.164 | 1581.238 | 688.4469 | 635.9482 | 663.4974 | 905.2447 | 929.9357 | 1008.288 |
| Slc25a18 | 71803 | 1.127349 | 2.379624 | 7.617842 | 1.61E-07 | 7.38E-07 | -0.88742 | 2.379624 | -5.72263 | 1.05E-05 | 3.87E-05 | 220.3683 | 189.1515 | 236.9557 | 564.9385 | 457.8329 | 485.6077 | 275.7246 | 341.8075 | 242.4718 | 309.9934 | 313.3989 | 263.4585 | 317.0442 | 371.8171 | 281.8349 |
| Abat | 268860 | 1.127627 | -0.57043 | 2.999491 | 0.006749 | 0.01679 | -0.09713 | -0.57043 | -0.28204 | 0.78063 | 0.852582 | 24.48536 | 29.30515 | 15.62345 | 54.54578 | 51.96262 | 51.40137 | 46.90161 | 57.63814 | 49.41369 | 35.38386 | 29.73859 | 24.62926 | 45.88798 | 88.82734 | 42.11325 |
| Mocs1 | 56738 | 1.128505 | 5.228201 | 14.37357 | 1.88E-12 | 1.74E-11 | 0.247072 | 5.228201 | 3.521309 | 0.00199 | 0.005102 | 1418.667 | 1463.482 | 1471.208 | 3554.567 | 3300.329 | 3297.804 | 4432.913 | 4556.094 | 3656.613 | 2189.954 | 2146.516 | 2262.16 | 1271.514 | 1154.755 | 1316.849 |
| Mylk | 107589 | 1.128586 | 1.733818 | 4.229847 | 0.000364 | 0.001114 | 0.409091 | 1.733818 | 1.815018 | 0.083601 | 0.133893 | 60.10044 | 82.58725 | 48.6063 | 166.2348 | 140.4395 | 129.8561 | 208.9253 | 281.4886 | 144.7936 | 403.0684 | 407.1899 | 423.1755 | 390.465 | 396.9717 | 425.1819 |
| Cask | 12361 | 1.128707 | 4.951658 | 20.9721 | 9.98E-16 | 1.67E-14 | -2.44873 | 4.951658 | -33.0527 | 8.07E-20 | 5.12E-18 | 2473.022 | 2587.734 | 2415.559 | 5889.646 | 5835.262 | 5715.021 | 963.6149 | 1121.933 | 1220.403 | 2363.027 | 2163.292 | 2065.873 | 630.7512 | 649.3042 | 583.1066 |
| Dclre1c | 227525 | 1.128785 | 6.43387 | 26.60156 | 7.48E-18 | 1.93E-16 | -0.45473 | 6.43387 | -10.9945 | 2.98E-10 | 2.15E-09 | 3799.683 | 3889.593 | 3568.223 | 8775.378 | 8870.16 | 8634.078 | 6907.328 | 6747.683 | 6310.013 | 5696.033 | 5567.978 | 5564.72 | 3053.637 | 3013.841 | 2952.787 |
| Cerkl | 228094 | 1.131897 | 0.682954 | 4.198814 | 0.000392 | 0.001195 | -0.5173 | 0.682954 | -1.96473 | 0.062598 | 0.104932 | 63.81034 | 69.26673 | 36.45472 | 128.5722 | 169.9318 | 97.39208 | 75.32683 | 125.9996 | 85.03751 | 103.0747 | 102.1787 | 122.4 | 105.9595 | 121.8428 | 107.7127 |
| Zfp456 | 408065 | 1.132136 | 3.129983 | 9.979574 | 1.70E-09 | 1.03E-08 | 0.55404 | 3.129983 | 5.84045 | 7.97E-06 | 3.00E-05 | 332.4074 | 372.9747 | 261.2589 | 742.8616 | 755.5647 | 742.6146 | 1175.383 | 1111.21 | 1131.918 | 460.7595 | 472.0047 | 532.8877 | 312.0383 | 311.2887 | 264.8276 |
| Fnbp1l | 214459 | 1.13238 | 7.374014 | 44.01982 | 1.94E-22 | 1.42E-20 | -4.14852 | 7.374014 | -79.0111 | 7.94E-28 | 1.85E-24 | 18651.17 | 19284.57 | 18459.11 | 44615.85 | 45377.42 | 41988.16 | 2453.096 | 2710.333 | 2556.871 | 10900.54 | 10814.17 | 10750.3 | 4255.902 | 3963.429 | 4287.453 |
| Myo10 | 17909 | 1.133379 | 6.942596 | 26.8617 | 6.11E-18 | 1.63E-16 | -0.37297 | 6.942596 | -9.19822 | 7.09E-09 | 4.11E-08 | 6624.404 | 6902.696 | 7381.214 | 16014.38 | 16955.26 | 15903.31 | 13801.86 | 13192.43 | 12317.8 | 5048.354 | 4904.579 | 4940.779 | 3004.411 | 2824.395 | 2921.202 |
| Slc26a9 | 320718 | 1.133859 | -0.92383 | 3.004914 | 0.006665 | 0.016598 | 0.963582 | -0.92383 | 3.178327 | 0.004462 | 0.010556 | 13.35565 | 20.4248 | 25.17112 | 37.66257 | 39.32307 | 58.16471 | 90.96069 | 73.7232 | 105.7223 | 9.999788 | 11.43792 | 7.463413 | 38.37904 | 30.65722 | 53.45144 |
| Sirpb1c | 100038947 | 1.135981 | 3.004551 | 10.61881 | 5.61E-10 | 3.61E-09 | -3.17963 | 3.004551 | -16.7683 | 9.09E-14 | 1.27E-12 | 986.0924 | 1165.102 | 716.0749 | 2144.169 | 2412.751 | 2064.171 | 270.0396 | 245.2972 | 242.4718 | 382.2996 | 330.9371 | 396.3072 | 130.155 | 115.5541 | 150.6359 |
| Myo15 | 17910 | 1.136015 | 0.322013 | 3.280124 | 0.003517 | 0.009228 | 0.975971 | 0.322013 | 3.567052 | 0.001785 | 0.004625 | 54.90658 | 24.86498 | 34.71878 | 96.10448 | 92.69009 | 74.39673 | 117.9647 | 206.4249 | 221.787 | 20.76879 | 26.68847 | 34.3317 | 146.8415 | 175.2964 | 134.4385 |
| Cav1 | 12389 | 1.136143 | 4.393188 | 19.91799 | 2.85E-15 | 4.33E-14 | 0.35004 | 4.393188 | 6.946197 | 6.70E-07 | 2.92E-06 | 1874.985 | 1727.228 | 2103.09 | 4479.248 | 4405.588 | 4463.804 | 5808.693 | 5982.302 | 5867.588 | 206.9187 | 227.9958 | 251.517 | 246.9608 | 293.9949 | 286.6941 |
| Mapk10 | 26414 | 1.136596 | -1.64183 | 2.845942 | 0.009578 | 0.023085 | 0.243552 | -1.64183 | 0.695027 | 0.494537 | 0.592511 | 13.35565 | 9.768385 | 12.15157 | 25.97418 | 29.4923 | 28.40602 | 27.00396 | 40.21265 | 36.77298 | 6.153716 | 3.812639 | 4.478048 | 49.22529 | 40.8763 | 35.63429 |
| D330050G | 320975 | 1.137585 | 0.703144 | 4.198893 | 0.000392 | 0.001195 | 2.052577 | 0.703144 | 10.48382 | 7.06E-10 | 4.81E-09 | 40.80894 | 40.84961 | 46.00239 | 120.78 | 71.62416 | 113.6241 | 469.0161 | 443.6796 | 396.4587 | 47.6913 | 54.13948 | 61.19998 | 88.43865 | 70.74744 | 98.80417 |
| Kif17 | 16559 | 1.138128 | -0.54542 | 2.779605 | 0.011124 | 0.026406 | 0.652936 | -0.54542 | 1.927628 | 0.067314 | 0.111494 | 20.03348 | 12.43249 | 26.03909 | 49.35095 | 54.77142 | 32.46403 | 54.00791 | 89.80826 | 78.14258 | 30.76858 | 22.11331 | 33.58536 | 65.91183 | 81.75259 | 46.97248 |
| Apol7c | 108956 | 1.138287 | 2.452846 | 6.930208 | 6.94E-07 | 2.94E-06 | 1.312244 | 2.452846 | 10.45776 | 7.39E-10 | 5.01E-09 | 103.8773 | 118.9967 | 115.44 | 271.4302 | 257.0043 | 262.4175 | 645.2524 | 656.8067 | 733.1613 | 566.9111 | 579.5212 | 505.273 | 342.9084 | 305.0001 | 365.2515 |
| Tmtc1 | 387314 | 1.139018 | -0.18966 | 3.221088 | 0.004039 | 0.010484 | 0.586752 | -0.18966 | 1.98689 | 0.059921 | 0.101102 | 21.51744 | 21.31284 | 27.77503 | 57.1432 | 47.74944 | 60.87005 | 62.53548 | 89.80826 | 108.0206 | 39.22994 | 57.18959 | 29.85365 | 77.5924 | 95.90208 | 82.60677 |
| Gna14 | 14675 | 1.13932 | -0.20774 | 3.473158 | 0.002231 | 0.006059 | -0.0257 | -0.20774 | -0.08647 | 0.9319 | 0.944266 | 21.51744 | 34.63336 | 32.11488 | 57.1432 | 94.09448 | 56.81205 | 75.32683 | 64.34024 | 66.65102 | 24.61486 | 41.1765 | 28.36097 | 109.2968 | 73.10568 | 80.98703 |
| Celf3 | 78784 | 1.140642 | 1.44927 | 6.489999 | 1.83E-06 | 7.33E-06 | -0.4258 | 1.44927 | -2.51124 | 0.020135 | 0.039711 | 99.42542 | 141.1976 | 96.34463 | 281.8199 | 244.3648 | 250.2435 | 166.2875 | 207.7654 | 224.0853 | 130.7665 | 129.6297 | 119.4146 | 166.0311 | 176.8686 | 138.4878 |
| Tes | 21753 | 1.140748 | 6.987863 | 29.05186 | 1.20E-18 | 3.69E-17 | -0.02692 | 6.987863 | -0.74821 | 0.462497 | 0.561179 | 5396.426 | 5457.863 | 5282.463 | 12610.47 | 13125.48 | 12279.52 | 13443.71 | 13058.39 | 12326.99 | 5572.959 | 5449.787 | 5672.194 | 5212.875 | 4938.957 | 5229.332 |
| Gm10933 | 100169866 | 1.140896 | 2.334804 | 7.640116 | 1.54E-07 | 7.07E-07 | -0.98438 | 2.334804 | -6.17871 | 3.68E-06 | 1.45E-05 | 257.4673 | 224.6729 | 281.2222 | 620.783 | 622.1471 | 553.2411 | 318.3624 | 340.4671 | 288.4381 | 221.5338 | 308.0613 | 175.3902 | 196.0668 | 251.5464 | 235.6722 |
| Myh3 | 17883 | 1.141522 | -0.69949 | 2.458974 | 0.022546 | 0.049557 | 0.980469 | -0.69949 | 2.65763 | 0.014604 | 0.029994 | 24.48536 | 9.768385 | 19.09533 | 59.74062 | 22.47032 | 47.34337 | 93.80322 | 68.36151 | 89.63414 | 13.84586 | 14.48803 | 16.41951 | 84.26702 | 96.68816 | 53.45144 |
| Robo1 | 19876 | 1.14193 | -0.07094 | 5.133042 | 4.18E-05 | 0.000143 | -0.68263 | -0.07094 | -3.04467 | 0.006083 | 0.013902 | 51.93865 | 44.40175 | 69.43757 | 122.0787 | 129.2044 | 133.9141 | 79.59061 | 80.4253 | 87.33582 | 22.30722 | 16.01308 | 13.43414 | 72.58644 | 51.09537 | 54.26131 |
| Crebl2 | 232430 | 1.142038 | 3.28888 | 10.20914 | 1.14E-09 | 7.00E-09 | 0.051115 | 3.28888 | 0.51239 | 0.61364 | 0.705299 | 314.5998 | 387.1833 | 307.2612 | 751.9526 | 849.6591 | 760.1993 | 881.1817 | 863.2316 | 799.8123 | 488.4512 | 500.9808 | 556.0242 | 538.9752 | 658.7372 | 582.2967 |
| Ostn | 239790 | 1.142039 | -1.14804 | 2.950914 | 0.007544 | 0.018572 | 0.809117 | -1.14804 | 2.557093 | 0.018221 | 0.036384 | 11.87169 | 25.75301 | 21.69924 | 41.55869 | 42.13186 | 47.34337 | 72.4843 | 76.40404 | 89.63414 | 26.92251 | 30.50111 | 32.09267 | 10.84625 | 10.21907 | 2.429611 |
| Gtf2a1 | 83602 | 1.143299 | 7.706517 | 27.50285 | 3.74E-18 | 1.05E-16 | -1.46975 | 7.706517 | -30.6894 | 3.81E-19 | 1.96E-17 | 11807.14 | 12431.6 | 10498.09 | 27710.56 | 28370.19 | 25750.74 | 10490.33 | 9932.525 | 10225.19 | 11756.67 | 11694.89 | 12088.49 | 7673.305 | 8081.715 | 8148.915 |
| Ppp4r1 | 70351 | 1.144087 | 6.919363 | 31.31275 | 2.50E-19 | 9.03E-18 | -1.57183 | 6.919363 | -37.0552 | 7.33E-21 | 6.16E-19 | 7082.948 | 7048.334 | 6862.168 | 16263.73 | 16956.67 | 16365.93 | 6003.406 | 5975.6 | 5399.882 | 7132.156 | 6868.088 | 7270.856 | 4294.281 | 3957.14 | 4294.742 |
| Brwd3 | 382236 | 1.144654 | 5.164058 | 19.84866 | 3.06E-15 | 4.62E-14 | -0.55215 | 5.164058 | -9.70248 | 2.80E-09 | 1.73E-08 | 1693.942 | 1716.572 | 1844.435 | 4024.7 | 4350.817 | 4024.187 | 2990.333 | 2856.439 | 2921.154 | 1741.502 | 1884.969 | 1687.478 | 1138.022 | 1172.835 | 1320.898 |
| Lrch3 | 70144 | 1.144888 | 6.296666 | 33.60179 | 5.72E-20 | 2.37E-18 | -1.99478 | 6.296666 | -47.0865 | 4.68E-23 | 9.05E-21 | 4713.062 | 4805.157 | 4803.344 | 11387.08 | 11448.63 | 10966.08 | 3061.396 | 2868.503 | 2883.231 | 5952.181 | 5940.854 | 5966.252 | 2509.655 | 2320.516 | 2480.633 |
| Fabp3 | 14077 | 1.146231 | 1.76781 | 6.826885 | 8.69E-07 | 3.63E-06 | -1.68095 | 1.76781 | -8.10385 | 5.95E-08 | 3.02E-07 | 230.756 | 131.4292 | 144.9509 | 385.7166 | 457.8329 | 350.3409 | 143.5473 | 138.0634 | 108.0206 | 366.9153 | 343.1375 | 335.8536 | 120.1431 | 112.4098 | 123.1003 |
| Tm4sf20 | 66261 | 1.146307 | 0.172039 | 3.706885 | 0.001279 | 0.003617 | -0.00988 | 0.172039 | -0.03553 | 0.971991 | 0.976847 | 29.67923 | 53.2821 | 52.07818 | 81.81868 | 115.1604 | 110.9188 | 112.2796 | 127.3401 | 82.7392 | 45.38365 | 59.47717 | 41.79511 | 39.21337 | 70.74744 | 74.50806 |
| Angptl7 | 654812 | 1.146544 | 1.418692 | 6.885579 | 7.64E-07 | 3.22E-06 | -1.44844 | 1.418692 | -7.42982 | 2.38E-07 | 1.11E-06 | 193.657 | 151.854 | 132.7993 | 416.8856 | 360.9296 | 353.0463 | 150.6537 | 195.7016 | 104.5732 | 131.5357 | 120.4794 | 106.7268 | 101.7879 | 96.68816 | 101.2338 |
| Ttn | 22138 | 1.146623 | 3.778573 | 12.37059 | 3.34E-11 | 2.54E-10 | 0.520214 | 3.778573 | 6.659348 | 1.26E-06 | 5.27E-06 | 408.0894 | 389.8474 | 415.7574 | 937.668 | 985.8855 | 949.5728 | 1347.355 | 1451.677 | 1468.621 | 769.9837 | 838.7806 | 807.5412 | 867.7 | 892.9899 | 753.1793 |
| Acot9 | 56360 | 1.146776 | 5.25004 | 22.62939 | 2.11E-16 | 4.08E-15 | -0.36602 | 5.25004 | -7.51398 | 2.00E-07 | 9.38E-07 | 2081.256 | 1995.415 | 2030.181 | 4801.328 | 4850.781 | 4800.618 | 4131.605 | 3927.436 | 3634.779 | 2663.79 | 2630.721 | 2735.341 | 608.2243 | 558.9047 | 621.1705 |
| Lmbr1l | 74775 | 1.148508 | 3.76549 | 14.33175 | 1.99E-12 | 1.83E-11 | -0.27951 | 3.76549 | -3.70226 | 0.001293 | 0.003461 | 484.5134 | 575.4467 | 566.7842 | 1346.761 | 1221.824 | 1249.865 | 1135.587 | 1087.082 | 1051.477 | 639.2172 | 664.9243 | 661.2584 | 723.3614 | 712.1909 | 725.6438 |
| Pitpnm1 | 18739 | 1.148602 | 7.01518 | 26.0036 | 1.20E-17 | 2.93E-16 | -0.91188 | 7.01518 | -19.8469 | 3.07E-15 | 5.90E-14 | 7286.251 | 7456.83 | 7288.341 | 17137.77 | 17347.09 | 17657.72 | 10096.64 | 10248.86 | 8648.545 | 6004.488 | 6338.894 | 6146.867 | 3641.003 | 3667.862 | 3627.409 |
| Dlgap3 | 242667 | 1.15029 | -0.86641 | 2.837448 | 0.009764 | 0.023502 | 0.851939 | -0.86641 | 2.598002 | 0.016658 | 0.033678 | 16.32358 | 10.65642 | 23.43518 | 25.97418 | 37.91867 | 58.16471 | 63.95674 | 65.68067 | 87.33582 | 12.30743 | 9.150334 | 6.717071 | 77.5924 | 85.68301 | 93.13508 |
| Usp50 | 75083 | 1.153983 | -0.67905 | 2.881426 | 0.008838 | 0.021453 | -0.44638 | -0.67905 | -1.15882 | 0.259331 | 0.348687 | 28.93725 | 15.98463 | 38.19066 | 66.23417 | 67.41097 | 55.45938 | 52.58665 | 42.8935 | 48.26453 | 13.07665 | 37.36386 | 43.28779 | 25.86413 | 47.16496 | 25.10598 |
| Heatr6 | 217026 | 1.154339 | 5.790156 | 28.58525 | 1.68E-18 | 5.01E-17 | -2.05446 | 5.790156 | -40.6073 | 1.07E-21 | 1.15E-19 | 3574.121 | 3424.263 | 3618.565 | 8592.26 | 8475.525 | 8175.524 | 2117.679 | 2210.355 | 1997.232 | 3510.695 | 3554.142 | 3504.072 | 1879.739 | 1721.521 | 1775.236 |
| Etv6 | 14011 | 1.15482 | 7.472108 | 34.17308 | 4.01E-20 | 1.75E-18 | 0.178883 | 7.472108 | 5.987639 | 5.68E-06 | 2.19E-05 | 8645.56 | 8835.948 | 8554.708 | 20311.81 | 21429.67 | 20185.86 | 24998.56 | 24567.25 | 23338.2 | 6646.013 | 6630.18 | 6499.886 | 4800.717 | 4615.877 | 4670.522 |
| Prr18 | 320111 | 1.155854 | 0.160554 | 3.972644 | 0.000676 | 0.001995 | -1.17887 | 0.160554 | -3.61821 | 0.001581 | 0.004146 | 31.90517 | 56.83424 | 60.75787 | 137.6632 | 95.49888 | 105.5081 | 34.11026 | 49.5956 | 72.3968 | 73.07537 | 69.39003 | 66.42437 | 50.89394 | 73.10568 | 89.08573 |
| 2310001H | 76432 | 1.157499 | 1.986073 | 8.834179 | 1.41E-08 | 7.44E-08 | -0.65323 | 1.986073 | -4.98685 | 5.91E-05 | 0.000197 | 186.9791 | 200.6959 | 177.9338 | 470.1327 | 508.3911 | 377.3943 | 258.6695 | 323.0416 | 305.6754 | 404.6068 | 394.2269 | 376.9023 | 74.2551 | 78.60826 | 73.69819 |
| Cttn | 13043 | 1.157533 | 3.308629 | 13.49219 | 6.40E-12 | 5.47E-11 | -1.49486 | 3.308629 | -14.7583 | 1.13E-12 | 1.25E-11 | 552.0337 | 516.8364 | 451.3442 | 1162.345 | 1228.846 | 1236.338 | 434.9058 | 491.9348 | 414.8452 | 907.6731 | 813.6172 | 867.9949 | 297.0204 | 313.647 | 316.6593 |
| Ifitm1 | 68713 | 1.157977 | 2.712271 | 9.462469 | 4.34E-09 | 2.46E-08 | 1.742567 | 2.712271 | 19.01485 | 7.30E-15 | 1.28E-13 | 254.4994 | 314.3644 | 284.694 | 576.6269 | 740.1163 | 715.5612 | 2347.923 | 2400.695 | 2263.837 | 499.2202 | 444.5537 | 492.5852 | 38.37904 | 58.17011 | 34.82442 |
| Tenm3 | 23965 | 1.158142 | -0.69099 | 2.78386 | 0.011018 | 0.02618 | 1.239654 | -0.69099 | 3.862316 | 0.000881 | 0.002441 | 25.22735 | 8.88035 | 16.49142 | 38.96127 | 39.32307 | 36.52203 | 96.64574 | 81.76573 | 105.7223 | 6.92293 | 16.77561 | 14.18048 | 86.77 | 113.982 | 90.70547 |
| Gm5065 | 272350 | 1.158449 | -1.21756 | 2.528603 | 0.019389 | 0.043371 | 1.275509 | -1.21756 | 3.571694 | 0.001766 | 0.00458 | 11.87169 | 15.98463 | 9.547666 | 31.16902 | 21.06593 | 37.8747 | 52.58665 | 80.4253 | 93.0816 | 10.769 | 7.625279 | 7.463413 | 55.8999 | 48.73712 | 33.20468 |
| Clcn6 | 26372 | 1.15853 | 4.066329 | 15.10202 | 7.18E-13 | 7.15E-12 | -1.37106 | 4.066329 | -15.7554 | 3.12E-13 | 3.86E-12 | 875.5373 | 854.2897 | 959.1064 | 2213 | 2186.643 | 2003.301 | 795.9061 | 966.4441 | 817.0496 | 1016.132 | 1005.774 | 958.3022 | 702.5033 | 576.9846 | 583.9165 |
| Dtnb | 13528 | 1.159352 | 2.73024 | 10.4277 | 7.78E-10 | 4.90E-09 | -1.15917 | 2.73024 | -9.44672 | 4.47E-09 | 2.68E-08 | 318.3097 | 255.7541 | 277.7503 | 737.6668 | 647.4262 | 653.3385 | 306.9923 | 305.6162 | 330.9568 | 429.2217 | 407.9524 | 448.5511 | 367.9382 | 342.732 | 427.6115 |
| Aicda | 11628 | 1.161971 | -0.66277 | 2.614213 | 0.016074 | 0.036733 | 1.259547 | -0.66277 | 3.647908 | 0.001472 | 0.003887 | 12.61367 | 15.98463 | 19.09533 | 22.07806 | 37.91867 | 63.57538 | 72.4843 | 93.82952 | 116.0647 | 23.84565 | 35.07628 | 36.57072 | 33.37308 | 51.88145 | 42.11325 |
| ND4L | 17720 | 1.162312 | -1.97048 | 3.012523 | 0.00655 | 0.016333 | -2.40169 | -1.97048 | -4.56635 | 0.000162 | 0.000507 | 17.80754 | 25.75301 | 15.62345 | 49.35095 | 30.8967 | 63.57538 | 7.106304 | 9.382952 | 9.193245 | 19.23036 | 20.58825 | 35.07804 | 5.840288 | 1.572165 | 4.859222 |
| Fsip2 | 241516 | 1.163103 | -0.45628 | 2.830728 | 0.009914 | 0.023827 | 1.062154 | -0.45628 | 3.260641 | 0.003682 | 0.008898 | 12.61367 | 21.31284 | 29.51097 | 58.44191 | 32.30109 | 54.10671 | 76.74809 | 108.5742 | 124.1088 | 18.46115 | 17.53814 | 15.67317 | 73.42077 | 117.1263 | 85.84625 |
| Ccdc39 | 51938 | 1.163417 | -0.27127 | 3.155302 | 0.004708 | 0.012062 | 0.0037 | -0.27127 | 0.011267 | 0.991115 | 0.992587 | 42.2929 | 19.53677 | 19.9633 | 51.94837 | 60.389 | 79.8074 | 90.96069 | 40.21265 | 75.84427 | 76.15223 | 61.00223 | 54.48291 | 42.55067 | 42.44846 | 42.92312 |
| 1700001K | 66323 | 1.165647 | 1.059604 | 6.119286 | 4.21E-06 | 1.62E-05 | -0.82708 | 1.059604 | -4.22643 | 0.000367 | 0.001085 | 131.3306 | 129.6531 | 94.60869 | 236.3651 | 297.7318 | 323.2876 | 156.3387 | 206.4249 | 142.4953 | 83.07516 | 101.4162 | 84.33656 | 55.8999 | 46.37887 | 81.7969 |
| Slc24a1 | 214111 | 1.166181 | -0.34616 | 3.781278 | 0.00107 | 0.003067 | -0.53708 | -0.34616 | -1.78128 | 0.089105 | 0.141499 | 39.32498 | 23.08891 | 53.81412 | 98.7019 | 77.24174 | 90.62874 | 61.11422 | 50.93603 | 78.14258 | 14.61507 | 26.68847 | 10.44878 | 62.57452 | 79.39434 | 60.74027 |
| Iglv1 | 16142 | 1.167008 | -1.03343 | 2.519278 | 0.019787 | 0.044147 | -0.02958 | -1.03343 | -0.07051 | 0.944446 | 0.954059 | 14.83961 | 8.88035 | 19.09533 | 33.76644 | 36.51428 | 29.75869 | 18.47639 | 56.29771 | 35.62382 | 30.76858 | 31.26364 | 42.54145 | 30.03577 | 52.66754 | 32.39481 |
| Oas1c | 114643 | 1.168033 | 3.10746 | 11.46145 | 1.39E-10 | 9.69E-10 | 0.443894 | 3.10746 | 5.178461 | 3.75E-05 | 0.000129 | 326.4715 | 322.3567 | 302.0534 | 751.9526 | 745.7339 | 787.2526 | 1121.375 | 1111.21 | 1008.959 | 459.9902 | 417.8653 | 476.9121 | 319.5472 | 274.3428 | 332.0468 |
| Arf4 | 11843 | 1.169225 | 7.319925 | 37.02238 | 7.46E-21 | 3.94E-19 | -0.67445 | 7.319925 | -21.3522 | 6.92E-16 | 1.55E-14 | 7058.463 | 7225.053 | 6874.319 | 17072.83 | 17043.74 | 16674.34 | 11065.94 | 10925.78 | 11031.89 | 8868.273 | 8516.674 | 8819.515 | 6740.527 | 6760.311 | 7279.924 |
| Rnf145 | 74315 | 1.169519 | 6.536672 | 26.90592 | 5.90E-18 | 1.58E-16 | -0.99708 | 6.536672 | -21.8501 | 4.32E-16 | 1.02E-14 | 4706.384 | 4820.254 | 4884.065 | 11436.43 | 11785.69 | 11370.53 | 6166.851 | 6005.089 | 5837.71 | 4252.987 | 4619.394 | 4086.218 | 3666.867 | 3465.052 | 3888.187 |
| Cacnb3 | 12297 | 1.169654 | 3.210479 | 10.17925 | 1.20E-09 | 7.35E-09 | 0.035299 | 3.210479 | 0.346328 | 0.732487 | 0.811062 | 308.664 | 344.5576 | 269.0706 | 814.2906 | 714.8372 | 677.6865 | 864.1266 | 794.8701 | 709.029 | 506.1431 | 549.0201 | 524.6779 | 568.1766 | 529.8197 | 570.1487 |
| Inpp4a | 269180 | 1.17004 | 6.189136 | 26.67192 | 7.08E-18 | 1.84E-16 | -1.27053 | 6.189136 | -26.4966 | 8.12E-18 | 2.85E-16 | 3196.453 | 3093.026 | 3143.786 | 7639.007 | 7336.561 | 7704.795 | 3384.022 | 3222.374 | 3168.222 | 4877.589 | 5115.799 | 4744.491 | 4101.551 | 4082.913 | 4268.016 |
| Sbds | 66711 | 1.17006 | 5.14129 | 20.60439 | 1.43E-15 | 2.30E-14 | -0.86519 | 5.14129 | -14.8159 | 1.04E-12 | 1.17E-11 | 1584.871 | 1784.062 | 1596.196 | 4035.089 | 4103.643 | 3757.711 | 2292.494 | 2349.759 | 2150.07 | 1955.343 | 2013.836 | 2024.077 | 1447.557 | 1345.773 | 1439.139 |
| Pcsk7 | 18554 | 1.170537 | 5.984499 | 23.45986 | 1.00E-16 | 2.05E-15 | -0.65218 | 5.984499 | -13.1273 | 1.08E-11 | 1.01E-10 | 3080.704 | 2887.002 | 3096.916 | 7180.563 | 7406.781 | 7220.54 | 5162.019 | 4893.88 | 4431.144 | 3946.839 | 3876.692 | 3778.726 | 1859.715 | 1848.866 | 1984.992 |
| Zfp513 | 101023 | 1.17085 | 5.376086 | 20.15114 | 2.25E-15 | 3.49E-14 | -0.71691 | 5.376086 | -12.2772 | 3.85E-11 | 3.24E-10 | 2150.26 | 2192.558 | 2091.807 | 5411.721 | 5016.5 | 5037.335 | 3435.187 | 3448.905 | 2953.33 | 1897.652 | 2029.087 | 2021.092 | 1355.781 | 1256.946 | 1334.666 |
| Spag1 | 26942 | 1.17162 | 2.021902 | 7.744458 | 1.24E-07 | 5.77E-07 | -0.56739 | 2.021902 | -3.81846 | 0.000979 | 0.002683 | 181.0433 | 201.5839 | 154.4986 | 380.5218 | 490.1339 | 424.7377 | 291.3585 | 257.361 | 349.3433 | 230.7643 | 226.4708 | 175.3902 | 165.1967 | 174.5103 | 187.8899 |
| Asxl3 | 211961 | 1.171676 | -1.25643 | 2.531307 | 0.019275 | 0.043142 | 1.484332 | -1.25643 | 4.225333 | 0.000368 | 0.001087 | 16.32358 | 11.54445 | 10.41564 | 20.77935 | 37.91867 | 37.8747 | 68.22052 | 88.46784 | 117.2139 | 6.92293 | 3.812639 | 4.478048 | 49.22529 | 37.73197 | 66.40936 |
| Igfn1 | 226438 | 1.171993 | -0.9426 | 3.014252 | 0.006524 | 0.016278 | 0.798448 | -0.9426 | 2.531142 | 0.019282 | 0.038234 | 14.83961 | 11.54445 | 27.77503 | 36.36386 | 47.74944 | 39.22736 | 73.90556 | 60.31898 | 88.48498 | 9.999788 | 9.150334 | 8.956095 | 49.22529 | 73.10568 | 48.59222 |
| Tnip2 | 231130 | 1.172308 | 4.592747 | 22.13892 | 3.30E-16 | 6.12E-15 | -1.63602 | 4.592747 | -26.2319 | 9.99E-18 | 3.45E-16 | 1366.729 | 1265.45 | 1289.803 | 3279.241 | 3151.463 | 3023.212 | 1084.422 | 1061.614 | 1017.003 | 1312.28 | 1275.709 | 1329.234 | 1101.312 | 1081.65 | 1118.431 |
| Dach1 | 13134 | 1.172505 | 3.495821 | 9.701435 | 2.81E-09 | 1.64E-08 | 0.827712 | 3.495821 | 8.506416 | 2.67E-08 | 1.42E-07 | 266.3711 | 274.4028 | 326.3566 | 629.8739 | 692.3669 | 758.8466 | 1247.867 | 1347.124 | 1231.895 | 401.5299 | 439.9786 | 520.9462 | 1191.419 | 1194.059 | 1250.44 |
| Plek2 | 27260 | 1.172735 | -0.2788 | 3.764809 | 0.001113 | 0.003182 | 1.376492 | -0.2788 | 5.842596 | 7.93E-06 | 2.99E-05 | 34.8731 | 26.64105 | 26.90706 | 76.62384 | 64.60218 | 73.04406 | 193.2915 | 199.7228 | 188.4615 | 26.15329 | 12.20045 | 23.13658 | 52.5626 | 27.51289 | 29.15533 |
| Trim13 | 66597 | 1.17294 | 3.499947 | 10.57595 | 6.03E-10 | 3.86E-09 | 1.436305 | 3.499947 | 16.92352 | 7.56E-14 | 1.07E-12 | 311.6319 | 317.9165 | 322.8847 | 838.9661 | 672.7053 | 784.5473 | 2315.234 | 2065.59 | 2082.27 | 375.3767 | 428.5407 | 397.0535 | 716.6868 | 647.7321 | 668.143 |
| Psd4 | 215632 | 1.174593 | 7.54631 | 25.35362 | 2.02E-17 | 4.72E-16 | -0.30424 | 7.54631 | -6.96683 | 6.41E-07 | 2.80E-06 | 9023.228 | 8790.658 | 9043.375 | 21218.31 | 21849.58 | 21717.08 | 19098.9 | 19255.16 | 16511.07 | 7112.926 | 7153.274 | 7639.549 | 6826.463 | 6545.71 | 6928.44 |
| Rnaset2b | 68195 | 1.175155 | 0.704479 | 4.915189 | 7.01E-05 | 0.000234 | -0.0096 | 0.704479 | -0.04484 | 0.96465 | 0.970997 | 51.19667 | 79.03511 | 118.9118 | 192.209 | 179.7626 | 183.9628 | 206.0828 | 183.6378 | 186.1632 | 36.15308 | 50.32684 | 67.17071 | 69.24913 | 66.81702 | 51.02183 |
| 4932441J | 319216 | 1.17586 | -0.17156 | 4.61413 | 0.000144 | 0.000463 | -0.65072 | -0.17156 | -2.58136 | 0.017278 | 0.034756 | 61.5844 | 65.71459 | 72.04148 | 157.1438 | 186.7846 | 139.3248 | 136.441 | 123.3188 | 74.69511 | 28.46093 | 39.65145 | 56.72194 | 5.005962 | 18.86598 | 6.478962 |
| Cldnd1 | 224250 | 1.176316 | 5.037124 | 19.64742 | 3.76E-15 | 5.59E-14 | -0.6404 | 5.037124 | -10.7616 | 4.40E-10 | 3.09E-09 | 1643.487 | 1561.165 | 1617.895 | 3833.789 | 4164.032 | 3672.493 | 2744.455 | 2478.44 | 2548.827 | 1879.96 | 1750.764 | 1831.521 | 889.3925 | 1058.853 | 1026.915 |
| Wfdc17 | 100034251 | 1.176411 | 4.952383 | 23.78561 | 7.56E-17 | 1.59E-15 | -2.72127 | 4.952383 | -38.095 | 4.09E-21 | 3.76E-19 | 2673.357 | 2693.41 | 2767.955 | 6601.339 | 6588.018 | 6431.935 | 1071.631 | 916.8485 | 1098.593 | 2566.869 | 2467.54 | 2836.097 | 453.8738 | 414.2655 | 500.4998 |
| Naip5 | 17951 | 1.176857 | 4.932329 | 23.21632 | 1.24E-16 | 2.51E-15 | -2.19048 | 4.932329 | -33.0188 | 8.25E-20 | 5.20E-18 | 1761.462 | 1659.737 | 1642.198 | 4176.649 | 4027.806 | 4033.655 | 952.2448 | 932.9335 | 902.0871 | 2736.865 | 2947.17 | 2845.799 | 1117.998 | 1153.183 | 1069.839 |
| D130012P | 319377 | 1.178201 | 1.236304 | 6.811534 | 8.98E-07 | 3.74E-06 | -1.52755 | 1.236304 | -7.42216 | 2.42E-07 | 1.12E-06 | 137.2664 | 111.0044 | 123.2517 | 340.2618 | 238.7472 | 330.0509 | 110.8583 | 109.9146 | 103.424 | 104.6132 | 127.3422 | 91.05363 | 111.7998 | 128.1315 | 145.7766 |
| Etv3 | 27049 | 1.178477 | 7.63322 | 24.18285 | 5.37E-17 | 1.16E-15 | 0.940442 | 7.63322 | 23.82349 | 7.31E-17 | 2.10E-15 | 5737.737 | 6049.294 | 5881.362 | 14759.83 | 14542.51 | 13363 | 29579.28 | 28750.71 | 26945.4 | 16335.04 | 16575.83 | 16493.4 | 5993.805 | 5771.419 | 6284.593 |
| Sh2d5 | 230863 | 1.178978 | 0.42035 | 4.551477 | 0.000168 | 0.000534 | 0.574059 | 0.42035 | 2.686931 | 0.013685 | 0.028353 | 37.84102 | 44.40175 | 51.21021 | 138.9619 | 88.4769 | 96.03941 | 170.5513 | 155.4889 | 167.7767 | 38.46072 | 47.27673 | 50.00486 | 85.93567 | 114.7681 | 78.55742 |
| Tmem132 | 98170 | 1.180684 | 7.728636 | 25.66334 | 1.57E-17 | 3.73E-16 | -0.26124 | 7.728636 | -6.05922 | 4.83E-06 | 1.87E-05 | 7864.996 | 8156.601 | 7638.133 | 19142.97 | 19758.44 | 18362.46 | 17539.78 | 16932.21 | 15378 | 10929.77 | 10863.73 | 11483.21 | 11465.32 | 11612.01 | 12452.57 |
| Fbxl22 | 74165 | 1.18124 | 1.602188 | 6.554081 | 1.58E-06 | 6.41E-06 | -0.26453 | 1.602188 | -1.58855 | 0.126868 | 0.190867 | 149.1381 | 116.3326 | 138.0072 | 323.3786 | 286.4966 | 374.689 | 243.0356 | 357.8926 | 259.7092 | 149.9968 | 99.89115 | 126.878 | 113.4685 | 128.9175 | 140.1076 |
| Mm2pr | 433966 | 1.181899 | -1.12923 | 2.883498 | 0.008797 | 0.021369 | -0.50062 | -1.12923 | -1.25221 | 0.224041 | 0.308509 | 15.5816 | 18.64873 | 19.09533 | 76.62384 | 29.4923 | 33.81669 | 24.16143 | 28.14886 | 42.51876 | 11.53822 | 18.30067 | 21.6439 | 40.04769 | 29.08506 | 40.49351 |
| Trim75 | 333307 | 1.182227 | -1.78218 | 2.529478 | 0.019352 | 0.043296 | 0.337213 | -1.78218 | 0.834528 | 0.413227 | 0.512293 | 10.38773 | 7.10428 | 10.41564 | 22.07806 | 28.08791 | 18.93735 | 22.74017 | 37.53181 | 31.0272 | 13.84586 | 7.625279 | 9.702436 | 15.85221 | 30.65722 | 21.05663 |
| Gm20022 | 100504031 | 1.183977 | 3.09503 | 11.91688 | 6.72E-11 | 4.92E-10 | -1.11894 | 3.09503 | -10.3249 | 9.29E-10 | 6.21E-09 | 442.9625 | 499.0757 | 404.4738 | 1114.292 | 1065.936 | 1069.96 | 466.1736 | 490.5944 | 589.5168 | 362.3 | 414.0526 | 312.717 | 383.7904 | 389.897 | 384.6884 |
| Omg | 18377 | 1.184715 | 1.64623 | 8.618463 | 2.15E-08 | 1.10E-07 | -1.00373 | 1.64623 | -6.85196 | 8.22E-07 | 3.53E-06 | 174.3655 | 171.3908 | 165.7822 | 410.3921 | 457.8329 | 380.0996 | 190.449 | 235.9142 | 217.1904 | 136.9202 | 118.9543 | 97.7707 | 134.3266 | 142.281 | 110.1424 |
| Hk3 | 212032 | 1.185902 | 8.788772 | 16.78113 | 3.53E-09 | 2.03E-08 | -0.11125 | 8.788772 | -1.73468 | 0.110719 | 0.170274 | 15394.62 | 15504.2 | 15618.25 | 37715.81 | 37639.2 | 37605.52 | 39636.12 | 37975.49 | 32225.77 | 24021.03 | 24412.33 | 25059.15 | 25372.72 | 24021.11 | 24892.17 |
| Mtm1 | 17772 | 1.186666 | 5.292573 | 17.43517 | 4.17E-14 | 5.15E-13 | 1.049573 | 5.292573 | 18.88136 | 8.41E-15 | 1.46E-13 | 1042.483 | 1212.168 | 1110.133 | 2722.094 | 2737.166 | 2676.929 | 5834.276 | 5944.77 | 5715.9 | 2446.871 | 2320.372 | 2294.999 | 1562.694 | 1818.995 | 1705.587 |
| Rnf139 | 75841 | 1.186667 | 5.977231 | 26.4205 | 8.61E-18 | 2.18E-16 | 0.059506 | 5.977231 | 1.462108 | 0.158281 | 0.230583 | 2555.382 | 2515.803 | 2604.777 | 6184.453 | 6543.078 | 5950.385 | 6878.903 | 6478.258 | 6819.089 | 2624.56 | 2838.129 | 2706.98 | 2644.816 | 2594.073 | 2555.951 |
| Psme1 | 19186 | 1.188522 | 7.309757 | 26.51635 | 7.99E-18 | 2.03E-16 | 0.771299 | 7.309757 | 20.85048 | 1.12E-15 | 2.39E-14 | 5233.19 | 5568.867 | 5304.162 | 12952.03 | 13160.59 | 13028.9 | 24235.34 | 23543.17 | 21860.39 | 7572.147 | 7696.194 | 7728.364 | 6099.764 | 6109.434 | 6275.685 |
| Gm8234 | 666680 | 1.188773 | -2.02441 | 2.678743 | 0.013936 | 0.032333 | -0.80192 | -2.02441 | -1.77223 | 0.090632 | 0.143529 | 16.32358 | 8.88035 | 9.547666 | 40.25998 | 19.66153 | 27.05335 | 11.37009 | 13.40422 | 26.43058 | 9.230573 | 9.150334 | 9.702436 | 12.5149 | 15.72165 | 9.718443 |
| Mark4 | 232944 | 1.189251 | 5.045842 | 18.06221 | 2.06E-14 | 2.67E-13 | -0.45267 | 5.045842 | -7.11531 | 4.65E-07 | 2.07E-06 | 1333.339 | 1399.543 | 1451.245 | 3497.424 | 3298.924 | 3366.79 | 2733.085 | 2696.929 | 2335.084 | 1583.813 | 1691.287 | 1576.273 | 1606.079 | 1480.194 | 1482.063 |
| Spata21 | 329972 | 1.189711 | 0.688052 | 4.061246 | 0.000546 | 0.001633 | 1.640088 | 0.688052 | 7.648798 | 1.51E-07 | 7.20E-07 | 69.00421 | 44.40175 | 41.66254 | 161.0399 | 124.9912 | 96.03941 | 488.9137 | 415.5307 | 342.4484 | 47.6913 | 49.56431 | 35.82438 | 65.91183 | 90.3995 | 59.9304 |
| Hspb1 | 15507 | 1.190289 | 0.735313 | 3.362573 | 0.002898 | 0.007712 | 2.076719 | 0.735313 | 8.183013 | 5.07E-08 | 2.59E-07 | 25.22735 | 17.7607 | 20.83127 | 54.54578 | 49.15383 | 52.75404 | 264.3545 | 201.0633 | 232.1294 | 24.61486 | 27.451 | 19.40487 | 1445.889 | 1541.508 | 1669.143 |
| Lipm | 78753 | 1.191458 | -1.41197 | 2.895877 | 0.008552 | 0.020829 | 0.315847 | -1.41197 | 0.888603 | 0.384122 | 0.482764 | 7.419807 | 15.09659 | 17.35939 | 32.46773 | 28.08791 | 29.75869 | 54.00791 | 30.8297 | 36.77298 | 10.769 | 12.20045 | 11.19512 | 29.20144 | 31.4433 | 29.9652 |
| Kcnrg | 328424 | 1.192218 | 1.779956 | 7.192166 | 3.95E-07 | 1.72E-06 | 0.618601 | 1.779956 | 4.605486 | 0.000147 | 0.000465 | 122.4268 | 95.90778 | 107.6282 | 262.3393 | 251.3868 | 285.4129 | 412.1656 | 457.0838 | 407.9502 | 110.7669 | 115.9042 | 88.06827 | 284.5055 | 258.6212 | 306.131 |
| Dync1i2 | 13427 | 1.192741 | 7.206715 | 35.34269 | 1.98E-20 | 9.37E-19 | -1.08864 | 7.206715 | -30.308 | 4.95E-19 | 2.44E-17 | 7341.158 | 7423.972 | 7695.419 | 17970.24 | 19268.3 | 17602.27 | 9280.833 | 8904.422 | 8608.325 | 7834.449 | 7887.588 | 7900.769 | 5678.429 | 5480.568 | 5591.344 |
| Myoz3 | 170947 | 1.192801 | -0.80026 | 3.103677 | 0.005307 | 0.013475 | -0.4631 | -0.80026 | -1.25773 | 0.222077 | 0.306241 | 33.38913 | 15.09659 | 30.37894 | 88.31222 | 49.15383 | 54.10671 | 35.53152 | 54.95729 | 50.56285 | 13.07665 | 27.451 | 7.463413 | 52.5626 | 38.51805 | 43.73299 |
| Gm6341 | 622673 | 1.19358 | -0.4337 | 3.680914 | 0.001361 | 0.003832 | -1.36239 | -0.4337 | -3.72896 | 0.001213 | 0.003264 | 23.0014 | 23.08891 | 25.17112 | 66.23417 | 43.53625 | 67.63339 | 41.21656 | 16.08506 | 19.53565 | 77.69066 | 76.25279 | 67.17071 | 64.24317 | 66.81702 | 68.83897 |
| Ilrun | 224647 | 1.194439 | 7.982272 | 29.57226 | 8.27E-19 | 2.61E-17 | -0.34733 | 7.982272 | -9.10548 | 8.44E-09 | 4.84E-08 | 11127.49 | 11687.43 | 11443.31 | 28331.34 | 28568.21 | 26721.95 | 24034.94 | 23367.57 | 21192.73 | 12758.19 | 13008.73 | 13049.03 | 8464.247 | 8029.048 | 8706.915 |
| Lrp1b | 94217 | 1.195571 | -0.77072 | 2.61724 | 0.015967 | 0.036517 | 1.239584 | -0.77072 | 3.512257 | 0.002034 | 0.005205 | 15.5816 | 10.65642 | 25.17112 | 45.45482 | 36.51428 | 37.8747 | 79.59061 | 81.76573 | 137.8987 | 12.30743 | 9.150334 | 5.224389 | 94.27894 | 118.6985 | 69.64884 |
| Stk38l | 232533 | 1.196561 | 4.697921 | 15.03346 | 7.85E-13 | 7.75E-12 | 0.759047 | 4.697921 | 11.40525 | 1.52E-10 | 1.15E-09 | 1052.871 | 1090.507 | 882.7251 | 2549.366 | 2688.013 | 2177.795 | 4384.59 | 4372.456 | 4231.191 | 783.0603 | 791.5039 | 873.9656 | 1052.921 | 1017.977 | 1013.958 |
| Eml2 | 72205 | 1.197995 | 5.187703 | 23.42977 | 1.03E-16 | 2.10E-15 | -1.71815 | 5.187703 | -28.6569 | 1.59E-18 | 6.82E-17 | 2586.545 | 2592.174 | 2662.931 | 6272.765 | 6639.981 | 6304.784 | 2244.171 | 2045.484 | 1823.71 | 1878.422 | 1987.91 | 1889.736 | 719.1898 | 801.8043 | 736.9819 |
| Rnf14 | 56736 | 1.19855 | 6.782253 | 29.94833 | 6.35E-19 | 2.08E-17 | 0.432224 | 6.782253 | 12.56946 | 2.47E-11 | 2.15E-10 | 4613.636 | 4320.29 | 4536.009 | 10788.38 | 11273.08 | 10999.89 | 15491.74 | 15850.49 | 15005.67 | 5126.814 | 5063.185 | 4981.082 | 3050.299 | 3178.132 | 3349.623 |
| Pcsk5 | 18552 | 1.198578 | -0.91757 | 3.132944 | 0.004959 | 0.012653 | 0.831305 | -0.91757 | 2.691633 | 0.013542 | 0.028098 | 13.35565 | 29.30515 | 18.22736 | 37.66257 | 47.74944 | 56.81205 | 73.90556 | 75.06362 | 111.4681 | 11.53822 | 4.575167 | 3.731706 | 63.40885 | 58.17011 | 68.0291 |
| Ier2 | 15936 | 1.200837 | 4.26224 | 17.67614 | 3.17E-14 | 3.98E-13 | -1.32848 | 4.26224 | -17.6859 | 3.14E-14 | 4.88E-13 | 1293.272 | 1390.663 | 1431.282 | 3388.332 | 3393.019 | 3288.335 | 1445.422 | 1426.209 | 1306.59 | 632.2943 | 672.5496 | 819.4827 | 497.2588 | 415.8377 | 411.4141 |
| Dennd1b | 329260 | 1.201085 | 6.974885 | 30.59885 | 4.05E-19 | 1.39E-17 | 0.347652 | 6.974885 | 10.21355 | 1.13E-09 | 7.44E-09 | 4704.9 | 4555.619 | 4760.813 | 11339.03 | 11805.35 | 11305.6 | 15470.42 | 15460.42 | 14644.84 | 7320.614 | 7516.237 | 7110.393 | 4083.196 | 3979.15 | 4033.154 |
| Megf11 | 214058 | 1.202045 | 3.987699 | 13.64123 | 5.18E-12 | 4.48E-11 | -0.76489 | 3.987699 | -8.60337 | 2.21E-08 | 1.19E-07 | 843.6321 | 914.676 | 795.9281 | 2181.831 | 1999.859 | 2077.698 | 1220.863 | 1454.358 | 1176.735 | 693.0622 | 761.0028 | 579.1608 | 500.5962 | 504.665 | 498.0702 |
| Piezo2 | 667742 | 1.203179 | -0.73887 | 2.946986 | 0.007612 | 0.018734 | 1.13412 | -0.73887 | 3.566104 | 0.001789 | 0.004635 | 14.83961 | 12.43249 | 28.643 | 37.66257 | 44.94065 | 47.34337 | 79.59061 | 91.14868 | 126.4071 | 7.692145 | 12.96297 | 7.463413 | 103.4565 | 78.60826 | 65.59949 |
| Scn5a | 20271 | 1.20413 | -0.44749 | 2.888124 | 0.008705 | 0.021161 | 1.146412 | -0.44749 | 3.555167 | 0.001837 | 0.004748 | 23.74338 | 12.43249 | 19.9633 | 63.63675 | 44.94065 | 32.46403 | 82.43313 | 113.9358 | 120.6613 | 9.999788 | 25.16342 | 15.67317 | 134.3266 | 106.9072 | 96.37456 |
| Eppk1 | 223650 | 1.204331 | 0.364949 | 3.736847 | 0.00119 | 0.003385 | 2.806104 | 0.364949 | 12.62199 | 2.28E-11 | 2.00E-10 | 24.48536 | 39.96157 | 25.17112 | 81.81868 | 57.58021 | 75.74939 | 480.3862 | 575.0409 | 511.3742 | 13.84586 | 20.58825 | 18.65853 | 138.4983 | 162.7191 | 114.1917 |
| Plcb3 | 18797 | 1.205245 | 5.931555 | 19.60953 | 3.91E-15 | 5.78E-14 | -0.48199 | 5.931555 | -8.10592 | 5.92E-08 | 3.00E-07 | 2863.304 | 2805.303 | 2713.273 | 7107.835 | 6835.192 | 6706.527 | 5351.047 | 5643.176 | 4512.734 | 3023.782 | 3154.578 | 3133.887 | 2124.196 | 2049.317 | 2139.677 |
| Pkd1l1 | 171395 | 1.206282 | -0.03282 | 3.446123 | 0.002379 | 0.006433 | 1.077427 | -0.03282 | 3.986775 | 0.000653 | 0.001852 | 40.06696 | 15.09659 | 27.77503 | 62.33804 | 71.62416 | 62.22272 | 150.6537 | 117.9571 | 163.1801 | 13.84586 | 25.16342 | 17.91219 | 171.8713 | 139.9227 | 131.199 |
| Gpsm2 | 76123 | 1.206335 | 4.894511 | 19.55745 | 4.13E-15 | 6.08E-14 | -1.16078 | 4.894511 | -17.5563 | 3.63E-14 | 5.57E-13 | 1664.263 | 1908.387 | 1874.814 | 4485.741 | 4628.887 | 4246.024 | 2130.47 | 1884.633 | 2179.948 | 1682.272 | 1773.64 | 1568.809 | 632.4198 | 680.7475 | 602.5435 |
| Ehd1 | 13660 | 1.20753 | 8.706643 | 23.00822 | 6.43E-11 | 4.72E-10 | -0.4264 | 8.706643 | -8.57902 | 2.60E-06 | 1.05E-05 | 25202.12 | 25588.73 | 25258.78 | 63954.93 | 62477.33 | 60993.14 | 51692.68 | 50229.62 | 44131.02 | 17334.25 | 17634.98 | 18616.74 | 7157.691 | 6790.968 | 7280.734 |
| Gm17821 | 100316870 | 1.208398 | -1.01116 | 2.660321 | 0.014517 | 0.033542 | 0.728086 | -1.01116 | 1.972604 | 0.061634 | 0.103581 | 13.35565 | 9.768385 | 24.30315 | 31.16902 | 56.17581 | 28.40602 | 51.16539 | 81.76573 | 60.90525 | 7.692145 | 16.01308 | 17.91219 | 35.04173 | 56.59795 | 51.02183 |
| Ep400 | 75560 | 1.208434 | 8.385271 | 41.80244 | 5.78E-22 | 3.86E-20 | -0.86843 | 8.385271 | -29.2505 | 1.04E-18 | 4.68E-17 | 16252.35 | 16536.99 | 15894.26 | 40396.35 | 40445.18 | 39263.89 | 23302.99 | 23714.74 | 21535.18 | 16496.57 | 16633.02 | 16624.01 | 12816.1 | 12822.58 | 13261.63 |
| E330033B | 319722 | 1.208435 | 3.004666 | 9.629714 | 3.20E-09 | 1.85E-08 | -1.03424 | 3.004666 | -7.7492 | 1.23E-07 | 5.93E-07 | 343.5371 | 383.6311 | 400.134 | 925.9796 | 931.1141 | 907.6401 | 469.0161 | 554.9346 | 401.0553 | 445.3752 | 517.7564 | 356.7511 | 318.7129 | 371.8171 | 384.6884 |
| Tifa | 211550 | 1.208808 | 5.864082 | 23.778 | 7.61E-17 | 1.60E-15 | -1.27126 | 5.864082 | -23.0036 | 1.50E-16 | 4.00E-15 | 3525.893 | 3710.21 | 3286.133 | 8522.13 | 8908.079 | 8520.454 | 3850.196 | 4010.542 | 3371.623 | 2916.092 | 3009.697 | 2996.56 | 1480.93 | 1507.706 | 1571.148 |
| Nfxl1 | 100978 | 1.208974 | 6.407845 | 44.4075 | 1.61E-22 | 1.21E-20 | -3.18912 | 6.407845 | -74.2816 | 2.95E-27 | 3.70E-24 | 9035.842 | 9108.575 | 9078.094 | 22545.59 | 22577.06 | 22062.01 | 2451.675 | 2679.503 | 2516.651 | 6384.48 | 6515.8 | 6258.071 | 1057.092 | 962.1651 | 1037.444 |
| Tmod3 | 50875 | 1.210019 | 7.094794 | 39.66149 | 1.75E-21 | 1.04E-19 | -1.4873 | 7.094794 | -42.9131 | 3.33E-22 | 4.58E-20 | 7434.647 | 7413.316 | 7614.697 | 18211.8 | 19268.3 | 18040.53 | 7038.084 | 6467.535 | 7023.639 | 8235.21 | 8300.116 | 8128.403 | 4678.071 | 4593.867 | 4757.178 |
| Hmcn1 | 545370 | 1.210607 | 0.064638 | 3.384067 | 0.002755 | 0.007359 | 0.995234 | 0.064638 | 3.55262 | 0.001848 | 0.004774 | 33.38913 | 46.17782 | 16.49142 | 77.92255 | 73.02855 | 68.98605 | 144.9686 | 115.2763 | 201.1022 | 26.15329 | 17.53814 | 19.40487 | 126.8177 | 160.3609 | 123.1003 |
| Btla | 208154 | 1.212027 | 1.121196 | 7.433866 | 2.36E-07 | 1.06E-06 | -1.87359 | 1.121196 | -8.88368 | 1.29E-08 | 7.16E-08 | 280.4687 | 174.9429 | 184.0096 | 520.7824 | 525.2438 | 526.1878 | 147.8111 | 99.19121 | 207.9972 | 69.2293 | 68.62751 | 59.7073 | 25.02981 | 40.8763 | 41.30338 |
| Xrn2 | 24128 | 1.212202 | 7.38526 | 29.57403 | 8.25E-19 | 2.61E-17 | -0.32469 | 7.38526 | -8.41248 | 3.21E-08 | 1.69E-07 | 7611.98 | 8293.359 | 7747.497 | 19788.43 | 19941.01 | 18662.76 | 16984.07 | 16227.15 | 15351.57 | 8387.514 | 8518.961 | 8482.168 | 5015.973 | 4567.14 | 4673.761 |

| | | | | | | | | | | | | | | | | | | | | | | | | | | |
|---|---|---|---|---|---|---|---|---|---|---|---|---|---|---|---|---|---|---|---|---|---|---|---|---|---|---|
| Sco1 | 52892 | 1.212797 | 3.763908 | 17.8595 | 2.58E-14 | 3.29E-13 | -2.38008 | 3.763908 | -24.855 | 3.05E-17 | 9.44E-16 | 1016.514 | 893.3632 | 869.7055 | 2261.053 | 2367.81 | 2268.424 | 480.3862 | 443.6796 | 451.6181 | 966.1334 | 916.5585 | 865.7559 | 357.9262 | 405.6186 | 423.5621 |
| Rbfox1 | 268859 | 1.214234 | 0.332012 | 4.185768 | 0.000405 | 0.001232 | 0.488616 | 0.332012 | 2.022689 | 0.055812 | 0.095186 | 33.38913 | 78.14708 | 72.04148 | 153.2477 | 102.5209 | 166.3781 | 186.1852 | 195.7016 | 220.6379 | 9.999788 | 22.11331 | 24.62926 | 85.93567 | 91.18558 | 72.88832 |
| Gna13 | 14674 | 1.218583 | 8.810858 | 54.33123 | 2.26E-24 | 3.15E-22 | -1.0391 | 8.810858 | -43.6071 | 2.37E-22 | 3.62E-20 | 24466.07 | 24999.96 | 25485.32 | 62443.24 | 62658.5 | 60933.62 | 31881.72 | 32135.27 | 30187.17 | 23572.58 | 23170.93 | 24265.05 | 12132.78 | 11955.53 | 12253.34 |
| Smox | 228608 | 1.21934 | 6.016151 | 16.88147 | 7.95E-14 | 9.35E-13 | 2.192282 | 6.016151 | 40.94417 | 8.96E-22 | 9.98E-20 | 1843.822 | 1885.298 | 2040.597 | 4684.444 | 4807.245 | 4824.966 | 24061.95 | 22541.87 | 21582.29 | 1356.125 | 1461.003 | 1641.204 | 2753.279 | 2708.055 | 2967.365 |
| Uaca | 72565 | 1.219662 | 3.538976 | 12.90401 | 1.50E-11 | 1.21E-10 | 0.303692 | 3.538976 | 3.753803 | 0.001143 | 0.003093 | 437.7686 | 420.9286 | 498.2146 | 1090.916 | 1103.855 | 1171.41 | 1594.655 | 1442.294 | 1311.187 | 487.682 | 484.2052 | 443.3267 | 468.8917 | 463.7887 | 472.1544 |
| Ppp1r32 | 67752 | 1.220372 | -0.39421 | 3.066538 | 0.005783 | 0.014585 | 1.284681 | -0.39421 | 4.233265 | 0.000361 | 0.001068 | 21.51744 | 33.74533 | 15.62345 | 59.74062 | 49.15383 | 59.51738 | 109.4371 | 159.5102 | 160.8818 | 28.46093 | 22.11331 | 19.40487 | 45.88798 | 69.17527 | 28.34546 |
| Zdhhc23 | 332175 | 1.2215 | -1.46521 | 2.924862 | 0.008006 | 0.019615 | -1.96346 | -1.46521 | -3.78458 | 0.001062 | 0.00289 | 16.32358 | 18.64873 | 12.15157 | 58.44191 | 36.51428 | 27.05335 | 4.263783 | 14.74464 | 13.78987 | 41.53758 | 26.68847 | 32.09267 | 17.52087 | 26.72681 | 21.8665 |
| Chst1 | 76969 | 1.222375 | -0.83572 | 3.771998 | 0.001094 | 0.003131 | -0.02332 | -0.83572 | -0.08153 | 0.93578 | 0.947087 | 34.13111 | 27.52908 | 23.43518 | 75.32513 | 75.83734 | 62.22272 | 98.067 | 80.4253 | 51.712 | 8.461359 | 7.625279 | 9.702436 | 42.55067 | 21.22423 | 31.58494 |
| Strc | 140476 | 1.222434 | -1.17092 | 2.660536 | 0.01451 | 0.033535 | 1.336607 | -1.17092 | 3.798825 | 0.001026 | 0.0028 | 11.87169 | 10.65642 | 8.679696 | 22.07806 | 40.72746 | 20.29002 | 82.43313 | 52.27645 | 78.14258 | 13.07665 | 15.25056 | 9.702436 | 52.5626 | 57.38403 | 41.30338 |
| Zbtb32 | 58206 | 1.222647 | 1.798052 | 5.093183 | 4.59E-05 | 0.000157 | 3.688955 | 1.798052 | 22.97029 | 1.55E-16 | 4.10E-15 | 66.77827 | 79.92315 | 102.4204 | 207.7935 | 174.145 | 227.2482 | 2758.667 | 2971.715 | 2469.535 | 62.30637 | 54.13948 | 47.0195 | 156.8535 | 183.9433 | 132.8187 |
| Ppp1r9a | 243725 | 1.222814 | 2.08166 | 5.994629 | 5.59E-06 | 2.13E-05 | 1.495172 | 2.08166 | 9.942566 | 1.82E-09 | 1.16E-08 | 100.1674 | 107.4522 | 59.8899 | 212.9883 | 230.3208 | 208.3108 | 646.6737 | 662.1683 | 605.605 | 177.6885 | 153.2681 | 155.9853 | 541.4782 | 566.7656 | 482.6827 |
| Pdgfra | 18595 | 1.223173 | -1.21863 | 2.522621 | 0.019643 | 0.043866 | 1.147641 | -1.21863 | 3.025503 | 0.006357 | 0.014457 | 8.903769 | 13.32052 | 17.35939 | 25.97418 | 32.30109 | 37.8747 | 42.63783 | 88.46784 | 97.67823 | 4.615287 | 12.20045 | 4.478048 | 46.72231 | 61.31444 | 53.45144 |
| Adamts17 | 233332 | 1.225378 | -1.13281 | 2.923081 | 0.008039 | 0.01968 | 0.810773 | -1.13281 | 2.415617 | 0.024748 | 0.047386 | 17.06556 | 13.32052 | 11.2836 | 38.96127 | 36.51428 | 29.75869 | 78.16935 | 57.63814 | 59.75609 | 9.230573 | 6.862751 | 2.239024 | 68.41481 | 95.116 | 80.98703 |
| Ttll9 | 74711 | 1.225622 | -0.52472 | 3.715286 | 0.001254 | 0.00355 | -0.25558 | -0.52472 | -0.83462 | 0.413175 | 0.512252 | 20.03348 | 21.31284 | 32.11488 | 55.84449 | 67.41097 | 56.81205 | 35.53152 | 57.63814 | 64.35271 | 23.07643 | 34.31375 | 17.91219 | 67.58048 | 66.03094 | 63.97975 |
| 4930432E1 | 243900 | 1.226011 | -0.62421 | 2.781341 | 0.011081 | 0.026315 | 1.035793 | -0.62421 | 2.998359 | 0.006767 | 0.015283 | 16.32358 | 15.98463 | 26.90706 | 41.55869 | 50.55823 | 52.75404 | 90.96069 | 72.38277 | 152.8377 | 10.769 | 12.20045 | 9.702436 | 105.9595 | 90.3995 | 55.07118 |
| H2-Q1 | 15006 | 1.226659 | -1.21739 | 2.718841 | 0.012746 | 0.029859 | 0.964549 | -1.21739 | 2.698283 | 0.013343 | 0.027747 | 11.87169 | 15.09659 | 6.075787 | 29.87031 | 21.06593 | 28.40602 | 66.79926 | 61.6594 | 40.22045 | 13.07665 | 14.48803 | 14.18048 | 44.21933 | 50.30929 | 51.8317 |
| Il2rb | 16185 | 1.227113 | -0.56064 | 3.038495 | 0.00617 | 0.015478 | 1.276101 | -0.56064 | 4.174567 | 0.000416 | 0.001218 | 24.48536 | 14.20856 | 13.01954 | 25.97418 | 54.77142 | 55.45938 | 126.4922 | 119.2975 | 87.33582 | 28.46093 | 25.16342 | 27.61463 | 49.22529 | 52.66754 | 43.73299 |
| Gng3 | 14704 | 1.22733 | -0.13696 | 4.901615 | 7.24E-05 | 0.000241 | -1.15021 | -0.13696 | -4.24125 | 0.000354 | 0.001049 | 51.93865 | 40.84961 | 46.00239 | 109.0916 | 112.3516 | 128.5034 | 45.48035 | 67.02109 | 51.712 | 66.15244 | 50.32684 | 50.75121 | 17.52087 | 33.80155 | 38.0639 |
| Cdo1 | 12583 | 1.227734 | -0.47769 | 3.360121 | 0.002914 | 0.007751 | -0.11496 | -0.47769 | -0.34749 | 0.731625 | 0.810278 | 16.32358 | 21.31284 | 28.643 | 44.15611 | 42.13186 | 82.51273 | 52.58665 | 52.27645 | 49.41369 | 39.22994 | 48.03925 | 26.12194 | 69.24913 | 60.52836 | 59.9304 |
| Ubald2 | 319370 | 1.22853 | 4.629842 | 5.11772 | 0.000354 | 0.001085 | 0.473234 | 4.629842 | 2.322617 | 0.04078 | 0.072775 | 830.2765 | 1023.016 | 406.2098 | 1676.634 | 1867.846 | 1755.763 | 2824.045 | 2591.035 | 2282.223 | 1083.054 | 1048.476 | 1101.6 | 2041.598 | 2147.578 | 2047.352 |
| Npepps | 19155 | 1.231527 | 7.09279 | 28.88936 | 1.35E-18 | 4.11E-17 | -1.13138 | 7.09279 | -24.9453 | 2.83E-17 | 8.82E-16 | 5944.008 | 6418.717 | 6389.992 | 15758.54 | 16262.9 | 14902.34 | 7963.325 | 7513.064 | 6870.801 | 8137.52 | 8356.543 | 8370.964 | 6037.19 | 5967.153 | 6020.576 |
| Nox1 | 237038 | 1.233897 | 1.536831 | 7.158549 | 4.24E-07 | 1.84E-06 | 0.376632 | 1.536831 | 2.63684 | 0.015291 | 0.031217 | 116.491 | 98.57188 | 108.4962 | 274.0276 | 268.2395 | 273.2389 | 469.0161 | 323.0416 | 334.4043 | 46.92208 | 48.80178 | 50.75121 | 302.8607 | 277.4872 | 296.4125 |
| Kcna1 | 16485 | 1.23592 | -1.07138 | 2.577023 | 0.017443 | 0.039453 | 1.337451 | -1.07138 | 3.630479 | 0.001535 | 0.004038 | 6.677827 | 18.64873 | 19.9633 | 50.64966 | 25.27911 | 28.40602 | 90.96069 | 69.70193 | 106.8715 | 9.230573 | 14.48803 | 4.478048 | 51.72827 | 53.45362 | 44.54286 |
| Tgfbr3 | 21814 | 1.237419 | 1.942317 | 6.326842 | 2.63E-06 | 1.04E-05 | 0.155482 | 1.942317 | 0.92835 | 0.363607 | 0.461412 | 106.1032 | 85.25136 | 74.64539 | 193.5077 | 233.1296 | 248.8909 | 224.5592 | 278.8077 | 271.2007 | 401.5299 | 371.3511 | 321.6731 | 342.9084 | 356.0954 | 334.4764 |
| Atp11b | 76295 | 1.23742 | 7.01235 | 36.89881 | 8.01E-21 | 4.21E-19 | 0.636718 | 7.01235 | 22.74771 | 1.89E-16 | 4.88E-15 | 4408.108 | 4451.719 | 4618.466 | 11402.67 | 11242.18 | 11255.55 | 18013.06 | 18225.71 | 18430.16 | 5046.047 | 4939.655 | 5160.203 | 5902.029 | 5974.228 | 5705.536 |
| Tbk1 | 56480 | 1.238083 | 6.621024 | 31.98945 | 1.60E-19 | 6.01E-18 | -0.4669 | 6.621024 | -12.5571 | 2.51E-11 | 2.19E-10 | 4431.851 | 4819.366 | 4425.777 | 11574.1 | 11583.45 | 11216.32 | 8824.609 | 8687.273 | 8346.317 | 4650.671 | 4436.387 | 4651.945 | 3350.657 | 3432.823 | 3485.682 |
| Armc3 | 70882 | 1.239142 | -0.01129 | 2.774426 | 0.011254 | 0.026676 | 3.393415 | -0.01129 | 11.37373 | 1.60E-10 | 1.21E-09 | 26.71131 | 8.88035 | 9.547666 | 28.5716 | 40.72746 | 36.52203 | 351.0514 | 450.3817 | 365.4315 | 29.99936 | 25.16342 | 35.07804 | 126.8177 | 128.9175 | 132.0089 |
| Insm1 | 53626 | 1.23957 | -0.10457 | 2.620324 | 0.015859 | 0.036301 | 0.70083 | -0.10457 | 1.828442 | 0.081495 | 0.131062 | 11.87169 | 17.7607 | 14.75548 | 20.77935 | 47.74944 | 51.40137 | 72.4843 | 69.70193 | 50.56285 | 83.84438 | 97.60356 | 127.6244 | 148.5102 | 117.1263 | 167.6431 |
| Liph | 239759 | 1.239819 | -0.11816 | 4.858041 | 8.04E-05 | 0.000266 | -0.94355 | -0.11816 | -3.55741 | 0.001827 | 0.004725 | 62.32638 | 55.05817 | 29.51097 | 127.2735 | 132.0132 | 101.4501 | 58.27169 | 54.95729 | 79.29174 | 18.46115 | 19.0632 | 20.89756 | 68.41481 | 76.25001 | 66.40936 |
| Gadd45a | 13197 | 1.239849 | 4.411728 | 9.270263 | 6.20E-09 | 3.44E-08 | 1.153728 | 4.411728 | 11.18537 | 2.17E-10 | 1.60E-09 | 354.6668 | 277.0669 | 301.1855 | 818.1868 | 838.424 | 708.7979 | 1982.659 | 1872.569 | 1643.292 | 397.6839 | 528.4318 | 431.3852 | 16653.17 | 16395.33 | 16867.98 |
| Znrf1 | 170737 | 1.240078 | 6.690968 | 29.76222 | 7.23E-19 | 2.33E-17 | -0.32869 | 6.690968 | -8.38659 | 3.38E-08 | 1.77E-07 | 5013.564 | 4871.76 | 4974.334 | 12322.15 | 12809.49 | 12386.38 | 10891.12 | 10546.44 | 9700.022 | 5363.732 | 5002.183 | 5347.535 | 2622.29 | 2696.263 | 2697.678 |
| Gm973 | 381260 | 1.240218 | -1.41123 | 2.856615 | 0.00935 | 0.022575 | -0.49424 | -1.41123 | -1.1906 | 0.246884 | 0.334568 | 22.25942 | 15.98463 | 13.88751 | 40.25998 | 33.70549 | 62.22272 | 34.11026 | 28.14886 | 34.47467 | 9.999788 | 13.7255 | 1.492683 | 35.04173 | 35.37372 | 32.39481 |
| Galnt3 | 14425 | 1.240568 | 1.973588 | 7.220541 | 3.71E-07 | 1.63E-06 | 1.331109 | 1.973588 | 10.3734 | 8.54E-10 | 5.74E-09 | 153.59 | 144.7497 | 112.836 | 305.1967 | 355.312 | 381.4523 | 808.6974 | 971.8058 | 926.2194 | 72.30616 | 48.03925 | 55.97559 | 278.6652 | 257.8351 | 329.6172 |
| Sertad1 | 55942 | 1.241963 | 4.566376 | 17.24595 | 5.19E-14 | 6.29E-13 | -0.03157 | 4.566376 | -0.48498 | 0.632629 | 0.722138 | 959.3811 | 909.3478 | 895.7446 | 2235.078 | 2363.597 | 2400.985 | 2484.364 | 2465.036 | 2190.291 | 1182.283 | 1125.491 | 1238.926 | 938.6178 | 879.6265 | 1016.387 |
| Tcirg1 | 27060 | 1.243218 | 9.408169 | 23.35552 | 8.03E-11 | 5.83E-10 | 0.121148 | 9.408169 | 2.660533 | 0.021939 | 0.042727 | 26131.82 | 25785.87 | 27389.65 | 66952.35 | 67047.23 | 66325.36 | 77886.52 | 79830.16 | 70012.3 | 34173.12 | 33805.91 | 33893.6 | 23891.79 | 24238.07 | 24856.54 |
| Trim15 | 69097 | 1.243361 | 0.369785 | 3.911967 | 0.000782 | 0.002285 | 0.847965 | 0.369785 | 3.381166 | 0.002773 | 0.006895 | 36.35706 | 49.72996 | 33.85081 | 132.4683 | 75.83734 | 96.03941 | 166.2875 | 206.4249 | 187.3124 | 45.38365 | 60.2397 | 32.09267 | 73.42077 | 66.03094 | 115.8114 |
| D5Ertd683 | 100504556 | 1.243554 | -0.50603 | 4.009258 | 0.000619 | 0.001838 | -0.37173 | -0.50603 | -1.27716 | 0.215266 | 0.298253 | 23.0014 | 30.19319 | 32.98284 | 84.4161 | 53.36702 | 79.8074 | 63.95674 | 53.61687 | 55.15947 | 28.46093 | 21.35078 | 28.36097 | 44.21933 | 29.87114 | 54.26131 |
| Minpp1 | 17330 | 1.244124 | 5.060415 | 21.97635 | 3.84E-16 | 7.00E-15 | -1.99869 | 5.060415 | -28.6041 | 1.65E-18 | 7.00E-17 | 1983.315 | 2006.071 | 1932.1 | 4768.86 | 5199.071 | 5030.571 | 1370.095 | 1333.72 | 1206.613 | 1669.965 | 1569.282 | 1666.58 | 1631.109 | 1360.709 | 1529.035 |
| Syk | 20963 | 1.244772 | 9.252062 | 29.08722 | 4.23E-13 | 4.37E-12 | -0.26755 | 9.252062 | -6.84731 | 1.26E-05 | 4.59E-05 | 27182.46 | 27798.16 | 27605.77 | 70080.94 | 71322.21 | 67483.24 | 62124.73 | 63007.86 | 56035.12 | 32157.01 | 31735.65 | 32973.36 | 17288.92 | 16680.67 | 17759.65 |
| Riox2 | 67014 | 1.245283 | 5.161123 | 22.49029 | 2.39E-16 | 4.56E-15 | -0.18503 | 5.161123 | -3.60634 | 0.001626 | 0.004252 | 1552.966 | 1618 | 1678.653 | 3990.933 | 4147.179 | 4120.226 | 3862.987 | 3788.032 | 3565.83 | 3301.468 | 3476.364 | 3482.428 | 486.4126 | 580.129 | 527.2255 |
| Gm10785 | 100038392 | 1.246471 | 1.191769 | 7.124804 | 4.56E-07 | 1.97E-06 | -1.07072 | 1.191769 | -5.72828 | 1.03E-05 | 3.82E-05 | 100.9094 | 111.8924 | 84.19305 | 251.9496 | 266.8351 | 231.3062 | 135.0198 | 91.14868 | 147.0919 | 127.6896 | 144.8803 | 119.4146 | 125.149 | 116.3402 | 117.4312 |
| Ankrd22 | 52024 | 1.247076 | 0.241108 | 5.490976 | 1.80E-05 | 6.44E-05 | -0.15033 | 0.241108 | -0.73663 | 0.469367 | 0.567904 | 61.5844 | 42.62568 | 65.96569 | 138.9619 | 132.0132 | 158.2621 | 115.1221 | 156.8293 | 131.0037 | 24.61486 | 28.21353 | 28.36097 | 46.72231 | 73.10568 | 60.74027 |
| Idnk | 75731 | 1.247668 | 4.025871 | 15.7922 | 2.98E-13 | 3.16E-12 | 0.611885 | 4.025871 | 9.338328 | 5.46E-09 | 3.23E-08 | 544.6139 | 581.6629 | 511.2341 | 1359.748 | 1401.586 | 1385.132 | 2131.891 | 2238.504 | 2201.782 | 786.9064 | 806.7545 | 740.3705 | 709.1779 | 622.5774 | 713.4957 |
| Cmtr1 | 74157 | 1.247966 | 7.267484 | 30.72078 | 3.73E-19 | 1.29E-17 | -0.13069 | 7.267484 | -3.53027 | 0.001949 | 0.005005 | 6711.216 | 6791.692 | 6659.063 | 17409.2 | 16861.17 | 16843.42 | 17121.93 | 16436.25 | 15158.51 | 7209.078 | 7192.163 | 7267.871 | 5053.518 | 4993.983 | 4773.375 |
| Ttc16 | 338348 | 1.248654 | -1.44283 | 2.624996 | 0.015696 | 0.035983 | 0.28835 | -1.44283 | 0.704902 | 0.488493 | 0.586794 | 14.09763 | 11.54445 | 5.207818 | 25.97418 | 33.70549 | 17.58468 | 27.00396 | 25.46801 | 43.66791 | 16.1535 | 16.01308 | 17.91219 | 30.03577 | 43.23454 | 38.87377 |
| Cubn | 65969 | 1.249141 | 1.018377 | 4.916038 | 7.00E-05 | 0.000233 | 2.814358 | 1.018377 | 16.33051 | 1.54E-13 | 2.04E-12 | 46.00281 | 38.1855 | 33.85081 | 87.01351 | 123.5868 | 94.68674 | 739.0556 | 723.8277 | 744.6528 | 43.84522 | 31.26364 | 31.34633 | 268.6533 | 256.2629 | 225.1439 |
| Marf1 | 223989 | 1.250559 | 8.106876 | 40.80694 | 9.61E-22 | 6.05E-20 | -0.35543 | 8.106876 | -12.3868 | 3.25E-11 | 2.78E-10 | 12015.64 | 11718.51 | 11814.8 | 29926.16 | 29839.19 | 30618.99 | 24714.3 | 24880.91 | 23836.93 | 13209.72 | 13793.37 | 13027.39 | 10057.81 | 10438.39 | 10794.76 |
| Dnajc13 | 235567 | 1.25101 | 8.399745 | 45.22575 | 1.10E-22 | 8.63E-21 | -0.85159 | 8.399745 | -30.2917 | 5.00E-19 | 2.47E-17 | 15434.68 | 14853.27 | 15055.8 | 38319.71 | 39931.17 | 37180.78 | 22573.89 | 22348.85 | 21539.77 | 23876.42 | 23528.56 | 23205.24 | 11368.54 | 10715.09 | 11121.14 |
| Rnf114 | 81018 | 1.251455 | 7.651848 | 34.89496 | 2.59E-20 | 1.19E-18 | -0.53012 | 7.651848 | -15.3247 | 5.39E-13 | 6.34E-12 | 8390.318 | 8727.608 | 8375.907 | 21748.18 | 21855.2 | 21154.37 | 15625.34 | 16302.21 | 14764.35 | 10310.55 | 10308.61 | 10182.33 | 8271.517 | 8529.783 | 8928.01 |
| Mapk8ip3 | 30957 | 1.252149 | 6.799569 | 28.31017 | 2.05E-18 | 6.03E-17 | -1.50781 | 6.799569 | -30.2215 | 5.25E-19 | 2.58E-17 | 6044.917 | 6111.457 | 6034.993 | 15174.12 | 15458.18 | 15642.25 | 5884.02 | 6057.366 | 5075.82 | 5780.647 | 6179.526 | 5810.267 | 4330.991 | 4075.838 | 4131.958 |
| Rab22a | 19334 | 1.255569 | 5.786558 | 23.90173 | 6.84E-17 | 1.45E-15 | -0.61651 | 5.786558 | -11.9786 | 6.10E-11 | 4.95E-10 | 2212.587 | 2401.247 | 2393.86 | 5729.905 | 6132.994 | 5969.323 | 4155.767 | 4203.563 | 3754.291 | 3113.78 | 3140.09 | 3145.082 | 2309.417 | 2120.065 | 2264.397 |
| Ambp | 11699 | 1.256107 | -2.22559 | 2.606853 | 0.016336 | 0.037242 | -0.81324 | -2.22559 | -1.66559 | 0.110407 | 0.169868 | 11.87169 | 12.43249 | 20.83127 | 28.5716 | 43.53625 | 41.9327 | 12.79135 | 33.51054 | 22.98311 | 0 | 6.100223 | 2.985365 | 14.18356 | 13.3634 | 11.33818 |
| Cmya5 | 76469 | 1.256123 | -0.09343 | 2.937733 | 0.007774 | 0.019092 | 1.02142 | -0.09343 | 3.079617 | 0.005611 | 0.012939 | 25.96933 | 13.32052 | 32.11488 | 75.32513 | 73.02855 | 35.16936 | 92.38196 | 111.255 | 168.9259 | 20.76879 | 33.55123 | 44.78048 | 118.4744 | 99.83249 | 103.6634 |
| Plk2 | 20620 | 1.256769 | 7.760988 | 36.17111 | 1.22E-20 | 6.09E-19 | 0.50002 | 7.760988 | 17.03123 | 6.67E-14 | 9.60E-13 | 4476.37 | 4636.431 | 4726.094 | 11950.72 | 11716.87 | 11588.3 | 17205.78 | 16863.85 | 17630.35 | 13744.32 | 13917.66 | 13985.69 | 28604.9 | 27725.92 | 28466.94 |
| 1110038F1 | 117171 | 1.257458 | 3.904271 | 14.80517 | 1.06E-12 | 1.02E-11 | 1.100172 | 3.904271 | 16.34885 | 1.50E-13 | 2.01E-12 | 533.4842 | 601.1997 | 577.1998 | 1498.71 | 1391.756 | 1459.528 | 3325.75 | 3395.288 | 3007.34 | 572.2956 | 565.7957 | 613.4925 | 366.2695 | 392.2552 | 343.385 |
| A3galt2 | 215493 | 1.258677 | -0.80942 | 2.69913 | 0.013318 | 0.031042 | 0.993753 | -0.80942 | 2.750948 | 0.011863 | 0.025073 | 27.45329 | 13.32052 | 11.2836 | 32.46773 | 42.13186 | 56.81205 | 76.74809 | 113.9358 | 80.44089 | 15.38429 | 23.63836 | 9.702436 | 30.8701 | 44.80671 | 68.83897 |
| Lap3 | 66988 | 1.25952 | 5.181703 | 24.04709 | 6.03E-17 | 1.30E-15 | -1.24058 | 5.181703 | -22.0421 | 3.61E-16 | 8.70E-15 | 2145.808 | 2174.798 | 2172.528 | 5280.551 | 5849.306 | 5493.184 | 2373.506 | 2625.886 | 2312.101 | 2051.495 | 1997.823 | 2160.658 | 837.6642 | 740.4898 | 812.2999 |
| Fam171a2 | 217219 | 1.259868 | 1.235947 | 7.142228 | 4.39E-07 | 1.90E-06 | -1.15768 | 1.235947 | -6.04744 | 4.96E-06 | 1.92E-05 | 111.2971 | 104.7881 | 105.0243 | 228.5728 | 300.5406 | 304.3502 | 98.067 | 147.4464 | 139.0478 | 129.9972 | 106.7539 | 94.78534 | 143.5042 | 123.415 | 133.6286 |
| Lcor | 212391 | 1.260685 | 4.656598 | 19.89446 | 2.92E-15 | 4.43E-14 | -1.17365 | 4.656598 | -17.3543 | 4.58E-14 | 6.89E-13 | 1271.013 | 1293.867 | 1268.104 | 3207.812 | 3518.01 | 3098.962 | 1564.808 | 1481.166 | 1472.068 | 1137.668 | 1102.615 | 1163.546 | 998.6893 | 1221.572 | 1060.93 |
| Tagln | 21345 | 1.260705 | 1.328372 | 7.618258 | 1.61E-07 | 7.37E-07 | -0.44407 | 1.328372 | -2.85159 | 0.009457 | 0.020554 | 94.97354 | 94.13171 | 124.9876 | 277.9238 | 254.1955 | 262.4175 | 196.134 | 237.2546 | 180.4174 | 153.0737 | 149.4555 | 132.8487 | 90.94163 | 86.46909 | 101.2338 |
| Nrg1 | 211323 | 1.26099 | 0.408124 | 2.600048 | 0.016583 | 0.037716 | 1.941856 | 0.408124 | 5.677736 | 1.16E-05 | 4.27E-05 | 52.68063 | 8.88035 | 25.17112 | 58.44191 | 43.53625 | 91.98141 | 299.886 | 269.4248 | 198.8039 | 244.6102 | 286.7105 | 220.1707 | 40.04769 | 31.4433 | 30.77507 |
| Chd7 | 320790 | 1.26128 | 6.270736 | 23.6112 | 8.79E-17 | 1.82E-15 | -0.08633 | 6.270736 | -1.77581 | 0.090025 | 0.142738 | 3507.343 | 3509.514 | 3532.636 | 9185.77 | 8660.906 | 9152.15 | 9113.125 | 9201.995 | 8209.568 | 2581.484 | 2677.235 | 2338.287 | 2863.41 | 3006.766 | 3022.436 |
| Ldb3 | 24131 | 1.261932 | -1.22732 | 2.819815 | 0.010161 | 0.024354 | 0.621192 | -1.22732 | 1.688905 | 0.105796 | 0.163852 | 8.903769 | 21.31284 | 10.41564 | 35.06515 | 25.27911 | 36.52203 | 62.53548 | 37.53181 | 57.45778 | 5.384501 | 9.150334 | 15.67317 | 53.39692 | 55.02578 | 45.35273 |
| Casp8 | 12370 | 1.262871 | 6.88647 | 35.5609 | 1.74E-20 | 8.32E-19 | -0.30146 | 6.88647 | -9.08862 | 8.71E-09 | 4.98E-08 | 5237.642 | 5551.107 | 5295.483 | 13906.58 | 13730.77 | 13513.15 | 11839.1 | 11524.95 | 11337.57 | 5792.185 | 5966.78 | 5992.374 | 3595.115 | 3340.851 | 3617.69 |
| B3gnt5 | 108105 | 1.2636 | 2.999768 | 6.196592 | 3.53E-06 | 1.37E-05 | 5.096573 | 2.999768 | 38.7533 | 2.85E-21 | 2.74E-19 | 71.97213 | 51.50603 | 63.36178 | 180.5206 | 155.8879 | 144.7354 | 5623.929 | 5668.644 | 5806.683 | 225.3798 | 231.8085 | 252.2633 | 1803.815 | 1821.353 | 1955.837 |
| Tmem86b | 68255 | 1.26364 | 2.537743 | 10.51593 | 6.68E-10 | 4.26E-09 | -0.23648 | 2.537743 | -2.16732 | 0.041661 | 0.074116 | 301.2442 | 244.2096 | 295.1097 | 792.2126 | 724.668 | 645.2225 | 622.5123 | 672.8917 | 610.2016 | 167.6888 | 165.4685 | 234.3512 | 201.0728 | 220.1031 | 211.3761 |
| Naip6 | 17952 | 1.264285 | 5.362744 | 22.20112 | 3.12E-16 | 5.83E-15 | -2.43066 | 5.362744 | -31.9689 | 1.62E-19 | 9.24E-18 | 2743.103 | 2422.559 | 2392.124 | 6410.428 | 6411.064 | 6599.666 | 1169.698 | 1371.251 | 1206.613 | 3573.77 | 3674.622 | 3282.409 | 1314.899 | 1216.07 | 1263.398 |
| Rcan1 | 54720 | 1.264934 | 4.753398 | 21.63112 | 5.31E-16 | 9.32E-15 | -2.55786 | 4.753398 | -31.4036 | 2.36E-19 | 1.28E-17 | 1839.37 | 1804.487 | 1848.775 | 4807.821 | 4814.267 | 4482.741 | 869.8116 | 914.1676 | 729.7138 | 1384.586 | 1236.82 | 1400.883 | 1069.607 | 1106.804 | 1225.334 |
| Sash1 | 70097 | 1.265224 | 8.655735 | 54.93465 | 1.79E-24 | 2.58E-22 | -1.79607 | 8.655735 | -63.5051 | 8.25E-26 | 4.35E-23 | 23528.95 | 24010.69 | 24112.2 | 61589.98 | 62666.93 | 59541.73 | 19248.14 | 18466.99 | 17417.75 | 21586.47 | 21839.56 | 22223.06 | 13628.73 | 13992.27 | 13936.25 |
| Rab29 | 226422 | 1.266591 | 3.937298 | 13.45417 | 6.75E-12 | 5.75E-11 | -1.45909 | 3.937298 | -13.564 | 5.78E-12 | 5.66E-11 | 634.3935 | 824.0965 | 710.8671 | 1636.374 | 2040.586 | 1862.623 | 763.2171 | 592.4664 | 730.863 | 967.6718 | 940.9594 | 1030.697 | 696.663 | 703.5439 | 753.1793 |
| Chic2 | 74277 | 1.266653 | 4.933908 | 20.23786 | 2.06E-15 | 3.22E-14 | -0.17836 | 4.933908 | -3.09241 | 0.005447 | 0.012603 | 1230.204 | 1218.384 | 1275.047 | 3250.669 | 3314.373 | 3006.98 | 3109.719 | 2832.311 | 2852.204 | 1424.585 | 1393.901 | 1547.912 | 1286.532 | 1214.498 | 1125.72 |
| Efna2 | 13637 | 1.266921 | -0.08242 | 3.657086 | 0.001441 | 0.004037 | 1.304446 | -0.08242 | 4.980519 | 6.00E-05 | 0.0002 | 18.54952 | 28.41712 | 24.30315 | 75.32513 | 57.58021 | 48.69604 | 132.1773 | 143.4251 | 186.1632 | 29.23015 | 35.07628 | 19.40487 | 90.94163 | 110.8376 | 84.22651 |
| Tox4 | 268741 | 1.266924 | 7.388028 | 37.9709 | 4.38E-21 | 2.40E-19 | -0.50917 | 7.388028 | -15.9071 | 2.59E-13 | 3.25E-12 | 7707.696 | 7524.32 | 7286.605 | 19336.48 | 19882.02 | 18720.92 | 14353.31 | 14447.07 | 13537.05 | 8732.122 | 8650.878 | 8576.954 | 5707.63 | 5572.54 | 5784.903 |
| Ssc5d | 269855 | 1.269939 | 1.312956 | 6.298993 | 2.80E-06 | 1.10E-05 | -0.89067 | 1.312956 | -4.28252 | 0.000321 | 0.000955 | 61.5844 | 95.01974 | 110.2321 | 232.4689 | 190.9978 | 235.3642 | 103.752 | 115.2763 | 147.0919 | 89.22888 | 105.9914 | 108.2195 | 315.3756 | 301.0696 | 309.3704 |
| P2ry10 | 78826 | 1.271617 | 2.339273 | 8.795253 | 1.52E-08 | 7.99E-08 | 0.545453 | 2.339273 | 4.676762 | 0.000124 | 0.000396 | 131.3306 | 126.989 | 113.704 | 322.0799 | 335.6505 | 304.3502 | 538.6579 | 427.5945 | 497.5844 | 169.9964 | 173.0938 | 176.8829 | 728.3674 | 768.7888 | 757.2287 |
| Trem1 | 58217 | 1.271627 | 3.837444 | 9.322757 | 5.62E-09 | 3.14E-08 | 4.630694 | 3.837444 | 51.93699 | 5.87E-24 | 1.50E-21 | 326.4715 | 249.5378 | 284.694 | 794.81 | 693.7713 | 735.8513 | 19304.99 | 19332.9 | 18672.63 | 76.15223 | 74.72773 | 100.7561 | 1241.478 | 1283.673 | 1542.803 |
| Arrdc4 | 66412 | 1.271806 | 5.270217 | 20.63293 | 1.39E-15 | 2.24E-14 | 1.05431 | 5.270217 | 21.26196 | 7.54E-16 | 1.68E-14 | 1050.645 | 1006.144 | 971.258 | 2545.47 | 2568.639 | 2708.041 | 5871.229 | 5491.708 | 5517.096 | 2544.561 | 2412.638 | 2465.165 | 1771.276 | 1834.717 | 1746.08 |
| Slc6a4 | 15567 | 1.2721 | 0.736184 | 5.314546 | 2.72E-05 | 9.54E-05 | 2.589127 | 0.736184 | 15.73552 | 3.20E-13 | 3.95E-12 | 50.45469 | 79.92315 | 92.87275 | 188.3128 | 214.8725 | 151.4988 | 1199.544 | 1196.997 | 1061.82 | 15.38429 | 23.63836 | 13.43414 | 50.89394 | 29.87114 | 31.58494 |
| Kirrel3 | 67703 | 1.272437 | 0.02146 | 3.484175 | 0.002174 | 0.005917 | -0.04745 | 0.02146 | -0.14633 | 0.885033 | 0.918721 | 24.48536 | 41.73764 | 33.85081 | 89.61093 | 61.79339 | 102.8027 | 78.16935 | 84.44657 | 87.33582 | 25.38408 | 41.93903 | 21.6439 | 81.76404 | 144.6392 | 156.305 |
| Tnik | 665113 | 1.273114 | 1.731367 | 7.814236 | 1.07E-07 | 5.04E-07 | -0.56688 | 1.731367 | -3.62241 | 0.001565 | 0.004109 | 117.9749 | 117.2206 | 145.8189 | 340.2618 | 348.29 | 290.8236 | 264.3545 | 197.042 | 228.682 | 179.9962 | 246.2965 | 214.9463 | 157.6878 | 144.6392 | 156.305 |
| Mthfsl | 100039707 | 1.273804 | 3.820098 | 9.664091 | 3.00E-09 | 1.75E-08 | 0.576598 | 3.820098 | 5.341855 | 2.55E-05 | 8.94E-05 | 391.0239 | 458.226 | 328.9605 | 1068.838 | 1124.921 | 841.3593 | 1688.458 | 1548.187 | 1454.831 | 686.9085 | 844.8809 | 867.2485 | 858.5224 | 905.5672 | 881.1388 |
| Adam21 | 56622 | 1.274739 | -1.32148 | 2.875323 | 0.008961 | 0.021727 | -0.56451 | -1.32148 | -1.32522 | 0.199111 | 0.279535 | 28.93725 | 16.87266 | 12.15157 | 48.05224 | 35.10988 | 66.28072 | 31.26774 | 29.48928 | 40.22045 | 12.30743 | 12.20045 | 13.43414 | 28.36712 | 7.860826 | 30.77507 |
| Gm20199 | 100504377 | 1.276028 | 3.767652 | 10.60347 | 5.75E-10 | 3.69E-09 | -0.07463 | 3.767652 | -0.69253 | 0.496073 | 0.594082 | 515.6766 | 589.6552 | 510.3661 | 1320.787 | 1344.006 | 1502.814 | 1340.249 | 1411.464 | 1349.109 | 616.1408 | 655.0114 | 426.1609 | 688.3197 | 638.2991 | 567.7191 |
| Hck | 15162 | 1.276606 | 8.000863 | 41.61007 | 6.37E-22 | 4.20E-20 | -0.81154 | 8.000863 | -26.3243 | 9.29E-18 | 3.22E-16 | 19439.9 | 20115.77 | 18409.64 | 49744.46 | 51437.38 | 48742.03 | 29960.18 | 30719.79 | 28224.41 | 9162.883 | 9377.568 | 9698.705 | 3079.501 | 3078.3 | 3232.192 |
| Pdzd2 | 68070 | 1.277951 | 2.468491 | 10.06733 | 1.46E-09 | 8.87E-09 | -0.0149 | 2.468491 | -0.13423 | 0.894479 | 0.920122 | 218.8843 | 190.9275 | 230.0119 | 524.6785 | 537.8834 | 597.8791 | 594.087 | 521.4241 | 587.2185 | 193.0728 | 247.059 | 232.8585 | 240.2862 | 257.8351 | 299.652 |
| Eng | 13805 | 1.278174 | 7.054424 | 25.64109 | 1.60E-17 | 3.78E-16 | 0.940041 | 7.054424 | 23.47663 | 9.89E-17 | 2.76E-15 | 4050.473 | 4057.432 | 4184.481 | 10410.45 | 11303.98 | 10157.18 | 21886 | 21688.02 | 20038.98 | 5789.877 | 5722.772 | 5813.998 | 5836.117 | 5843.738 | 6172.831 |
| Cmtm6 | 67213 | 1.278194 | 7.69608 | 35.28194 | 2.05E-20 | 9.70E-19 | 0.122837 | 7.69608 | 3.886026 | 0.000833 | 0.002317 | 8444.483 | 8546.449 | 8228.352 | 21624.81 | 22123.44 | 21577.76 | 25309.81 | 25202.61 | 23535.86 | 8509.05 | 8759.92 | 8971.022 | 7659.955 | 7118.764 | 7462.955 |
| Dusp5 | 240672 | 1.279344 | 3.088874 | 8.948392 | 1.14E-08 | 6.08E-08 | 1.756196 | 3.088874 | 16.94593 | 7.37E-14 | 1.05E-12 | 290.1145 | 204.248 | 205.7088 | 535.0682 | 626.3603 | 662.8072 | 2265.49 | 2156.739 | 1975.398 | 299.9936 | 272.985 | 299.2828 | 373.7785 | 432.3454 | 374.1601 |
| Il17ra | 16172 | 1.279682 | 5.87183 | 29.90153 | 6.56E-19 | 2.14E-17 | -0.26241 | 5.87183 | -6.59901 | 1.43E-06 | 5.98E-06 | 3469.502 | 3145.42 | 3258.358 | 8445.506 | 8837.859 | 8368.955 | 7623.643 | 7269.107 | 7310.928 | 1385.355 | 1315.361 | 1403.868 | 1888.916 | 1737.243 | 1916.153 |
| Pdcd10 | 56426 | 1.280702 | 6.297404 | 21.87015 | 4.24E-16 | 7.61E-15 | -0.13042 | 6.297404 | -2.43818 | 0.023579 | 0.045422 | 3488.051 | 3587.661 | 2782.711 | 8362.388 | 8801.345 | 8318.907 | 8050.021 | 7900.446 | 8175.093 | 3967.608 | 3941.506 | 4031.735 | 2612.278 | 2604.292 | 2543.803 |
| Fcgr1 | 14129 | 1.28096 | 7.517728 | 39.2239 | 2.21E-21 | 1.28E-19 | -0.8383 | 7.517728 | -25.4642 | 1.85E-17 | 6.05E-16 | 11917.69 | 11835.73 | 11983.19 | 30878.11 | 31264.65 | 30601.4 | 19179.92 | 17999.18 | 16925.91 | 7218.308 | 7570.376 | 7311.905 | 3363.172 | 3141.972 | 3181.98 |
| Bcl3 | 12051 | 1.281431 | 6.159619 | 33.36022 | 6.65E-20 | 2.70E-18 | -1.6445 | 6.159619 | -37.2599 | 6.53E-21 | 5.65E-19 | 5690.25 | 5766.011 | 5298.086 | 14597.49 | 14602.9 | 14282.82 | 5011.366 | 4892.539 | 4577.087 | 5224.505 | 5179.852 | 5159.457 | 747.5569 | 631.2243 | 736.9819 |
| Eps8l1 | 67425 | 1.282527 | 1.71651 | 7.191591 | 3.95E-07 | 1.72E-06 | -0.13236 | 1.71651 | -0.83611 | 0.412358 | 0.511383 | 115.749 | 109.2283 | 110.2321 | 290.9109 | 306.1582 | 277.2969 | 267.197 | 290.8715 | 270.0516 | 143.0739 | 188.3444 | 108.9658 | 210.2504 | 219.3171 | 246.2006 |
| Epha1 | 13835 | 1.283564 | 3.022212 | 10.70473 | 4.84E-10 | 3.15E-09 | 0.595714 | 3.022212 | 6.160071 | 3.84E-06 | 1.51E-05 | 244.8536 | 269.9626 | 247.3713 | 636.3675 | 702.1976 | 641.1645 | 1068.788 | 1117.912 | 938.8601 | 374.6074 | 398.0395 | 328.3902 | 448.0336 | 396.9717 | 423.5621 |
| Syn2 | 20965 | 1.28389 | -1.19739 | 2.491607 | 0.021011 | 0.046556 | 1.159685 | -1.19739 | 2.896557 | 0.008539 | 0.018789 | 7.419807 | 7.10428 | 13.88751 | 27.27289 | 22.47032 | 21.64268 | 42.63783 | 67.02109 | 59.75609 | 8.461359 | 15.25056 | 5.97073 | 113.4685 | 107.6933 | 74.50806 |
| Stard3 | 59045 | 1.284972 | 6.862921 | 31.50373 | 2.20E-19 | 8.04E-18 | -0.39618 | 6.862921 | -10.3037 | 9.64E-10 | 6.43E-09 | 5313.324 | 5084.888 | 5147.928 | 13810.47 | 13312.26 | 13358.95 | 11242.17 | 10799.78 | 10034.43 | 5863.722 | 5850.114 | 6079.696 | 3585.937 | 3643.493 | 3817.728 |
| Igsf6 | 80719 | 1.287633 | 6.511419 | 29.3845 | 9.44E-19 | 2.95E-17 | 0.406796 | 6.511419 | 10.92961 | 3.32E-10 | 2.38E-09 | 4582.473 | 4731.45 | 4762.549 | 12284.49 | 12685.9 | 11708.69 | 17131.88 | 17042.12 | 16350.19 | 5664.495 | 5393.359 | 5819.223 | 983.6714 | 900.8507 | 789.6235 |
| Yju2 | 72886 | 1.287664 | 2.767689 | 12.45321 | 2.94E-11 | 2.26E-10 | -0.30694 | 2.767689 | -3.24213 | 0.003845 | 0.00924 | 313.8579 | 317.0285 | 335.9042 | 829.8752 | 862.2987 | 826.48 | 773.1659 | 734.5511 | 628.5881 | 299.9936 | 292.0482 | 366.4536 | 182.7176 | 180.799 | 183.0307 |
| Fbxw17 | 109082 | 1.287689 | 5.283797 | 21.86056 | 4.28E-16 | 7.67E-15 | -0.19663 | 5.283797 | -3.61822 | 0.00158 | 0.004146 | 1737.719 | 1871.978 | 1849.643 | 4787.042 | 4887.296 | 4531.437 | 4297.893 | 4558.774 | 4047.326 | 2513.024 | 2745.1 | 2564.429 | 671.6332 | 765.6445 | 694.8687 |
| Sav1 | 64010 | 1.288172 | 6.404022 | 33.8886 | 4.78E-20 | 2.03E-18 | -0.91906 | 6.404022 | -23.725 | 7.96E-17 | 2.27E-15 | 4590.635 | 4441.951 | 4273.882 | 11283.19 | 12267.39 | 11233.91 | 6374.355 | 6322.769 | 6362.875 | 5096.815 | 5051.747 | 5335.594 | 1948.153 | 1934.549 | 2036.824 |
| Plekha2 | 83436 | 1.28876 | 7.80365 | 36.12022 | 1.25E-20 | 6.26E-19 | -0.51012 | 7.80365 | -14.9901 | 8.31E-13 | 9.43E-12 | 11200.2 | 11585.3 | 11029.29 | 29622.26 | 30537.17 | 28061.09 | 22109.13 | 21844.85 | 20496.34 | 10525.93 | 10723.43 | 10579.39 | 5811.087 | 5280.903 | 5815.678 |
| Obsl1 | 98733 | 1.289177 | -0.66759 | 3.69249 | 0.001324 | 0.003735 | 0.591878 | -0.66759 | 2.082473 | 0.049509 | 0.085882 | 16.32358 | 15.98463 | 21.69924 | 46.75353 | 39.32307 | 55.45938 | 65.378 | 64.34024 | 90.78329 | 13.07665 | 11.43792 | 14.18048 | 77.5924 | 105.3351 | 88.27586 |
| A630001G | 319997 | 1.289254 | 5.875031 | 26.11698 | 1.09E-17 | 2.70E-16 | -0.7245 | 5.875031 | -14.8331 | 1.02E-12 | 1.14E-11 | 3660.933 | 3968.628 | 3598.602 | 10031.23 | 9988.059 | 9236.015 | 6426.942 | 6305.344 | 5721.646 | 2563.023 | 2430.939 | 2439.79 | 976.1625 | 900.8507 | 997.7602 |
| Pabpc1l | 381404 | 1.289696 | -0.77274 | 3.162296 | 0.004632 | 0.011882 | -0.68819 | -0.77274 | -1.73874 | 0.096486 | 0.151223 | 25.96933 | 21.31284 | 29.51097 | 79.22126 | 56.17581 | 66.28072 | 66.79926 | 57.63814 | 22.98311 | 20.76879 | 25.16342 | 7.463413 | 45.05365 | 51.88145 | 29.15533 |
| Pirb | 18733 | 1.290107 | 8.792562 | 19.53235 | 7.10E-10 | 4.51E-09 | 0.412441 | 8.792562 | 7.611838 | 1.06E-05 | 3.91E-05 | 19135.68 | 19038.58 | 15224.19 | 46647.04 | 46471.44 | 45962.3 | 65449.06 | 64577.5 | 62510.62 | 24026.41 | 24419.95 | 25118.11 | 9857.573 | 10366.86 | 10180.07 |
| Lgals9 | 16859 | 1.290251 | 8.49775 | 31.10442 | 9.09E-18 | 2.29E-16 | 0.305278 | 8.49775 | 8.77391 | 4.12E-08 | 2.14E-07 | 14238.61 | 14238.75 | 13447.45 | 36623.6 | 37425.73 | 35512.94 | 48851.58 | 48410.67 | 43977.04 | 18269.61 | 18060.47 | 18319.69 | 10607.63 | 10799.99 | 10923.53 |
| Cth | 107869 | 1.292389 | -0.81192 | 2.572325 | 0.017623 | 0.039831 | 0.092752 | -0.81192 | 0.211375 | 0.8346 | 0.899771 | 20.77546 | 42.62568 | 12.15157 | 81.81868 | 51.96262 | 44.63804 | 46.90161 | 45.57434 | 110.3189 | 22.30722 | 28.97606 | 33.58536 | 27.53279 | 35.37372 | 7.288832 |
| AI182371 | 98870 | 1.292533 | -1.02299 | 2.955729 | 0.007461 | 0.018386 | -0.8889 | -1.02299 | -1.99576 | 0.058878 | 0.099624 | 18.54952 | 11.54445 | 26.03909 | 70.13029 | 49.15383 | 29.75869 | 14.21261 | 41.55307 | 28.72889 | 29.99936 | 33.55123 | 21.6439 | 33.37308 | 25.15464 | 45.35273 |
| Hunk | 26559 | 1.293061 | -1.74163 | 2.600811 | 0.016555 | 0.037662 | 1.715604 | -1.74163 | 4.621604 | 0.000142 | 0.000448 | 3.709904 | 12.43249 | 11.2836 | 23.37676 | 16.85274 | 22.99535 | 62.53548 | 68.36151 | 87.33582 | 3.076858 | 3.812639 | 2.239024 | 37.54471 | 40.8763 | 42.92312 |
| Rnd3 | 74194 | 1.294543 | 5.610488 | 28.85031 | 1.38E-18 | 4.21E-17 | 0.044282 | 5.610488 | 1.105093 | 0.281422 | 0.373133 | 2644.419 | 2442.096 | 2540.547 | 6757.184 | 6790.251 | 6460.341 | 7410.454 | 6972.874 | 7051.219 | 1347.664 | 1400.764 | 1308.336 | 1287.366 | 1249.085 | 1358.152 |
| Prkcd | 18753 | 1.294857 | 8.403059 | 28.31657 | 3.85E-16 | 7.02E-15 | -0.95228 | 8.403059 | -20.3063 | 1.14E-13 | 1.56E-12 | 12333.2 | 12478.67 | 13023.02 | 32863.84 | 33678.8 | 32492.43 | 18673.95 | 18960.27 | 15949.13 | 33984.66 | 33497.09 | 35166.85 | 13710.49 | 12997.88 | 13432.51 |
| Gm6345 | 622708 | 1.295477 | -0.52019 | 3.030867 | 0.006279 | 0.015719 | -0.05655 | -0.52019 | -0.1482 | 0.883579 | 0.918721 | 11.12971 | 24.86498 | 46.87036 | 51.94837 | 67.41097 | 62.22272 | 42.63783 | 77.74446 | 63.20356 | 42.30679 | 26.68847 | 33.58536 | 42.55067 | 28.29897 | 60.74027 |
| Tlr7 | 170743 | 1.297214 | 7.705344 | 41.25158 | 7.65E-22 | 4.94E-20 | -0.65161 | 7.705344 | -21.2181 | 7.87E-16 | 1.74E-14 | 8361.381 | 8348.417 | 9003.449 | 21972.86 | 23020.85 | 22425.88 | 14987.2 | 14296.94 | 15192.99 | 16348.88 | 15560.14 | 15897.07 | 6657.094 | 6757.952 | 6530.794 |
| Max | 17187 | 1.297604 | 7.075529 | 34.85763 | 2.65E-20 | 1.21E-18 | -0.58146 | 7.075529 | -16.149 | 1.92E-13 | 2.50E-12 | 5665.765 | 5904.545 | 5958.611 | 15444.25 | 16018.53 | 14522.24 | 10636.72 | 10679.14 | 10553.84 | 6567.553 | 6627.13 | 6708.862 | 5519.073 | 5104.821 | 5591.344 |
| Stoml1 | 69106 | 1.297777 | 4.227515 | 13.92041 | 3.51E-12 | 3.11E-11 | 0.349289 | 4.227515 | 4.400534 | 0.000241 | 0.000735 | 647.0072 | 743.2853 | 611.9186 | 1798.712 | 1747.068 | 1682.719 | 2572.482 | 2335.015 | 2074.226 | 849.982 | 809.8046 | 739.6242 | 785.1016 | 848.9692 | 823.6381 |
| Otud5 | 54644 | 1.298084 | 6.936616 | 33.80735 | 5.03E-20 | 2.12E-18 | -0.03193 | 6.936616 | -0.92878 | 0.363391 | 0.461204 | 4974.981 | 4923.266 | 4997.769 | 12955.92 | 12879.71 | 13287.26 | 13715.17 | 13437.73 | 12678.63 | 5907.567 | 6167.325 | 6228.218 | 4058.166 | 4048.325 | 3940.019 |
| Inpp1 | 16329 | 1.298198 | 3.143023 | 11.53452 | 1.23E-10 | 8.68E-10 | 1.749048 | 3.143023 | 21.3174 | 7.15E-16 | 1.59E-14 | 245.5956 | 255.7541 | 284.694 | 700.0042 | 653.0438 | 702.0346 | 2339.395 | 2435.546 | 2397.139 | 255.3792 | 263.8346 | 232.1121 | 424.6724 | 394.6135 | 331.2369 |
| Crim1 | 50766 | 1.298523 | 5.954833 | 22.91572 | 1.63E-16 | 3.22E-15 | 0.292778 | 5.954833 | 5.960511 | 6.05E-06 | 2.32E-05 | 2407.728 | 2411.015 | 2302.723 | 6132.505 | 6766.376 | 5857.051 | 8261.789 | 8045.211 | 7559.145 | 2103.802 | 1913.182 | 2165.882 | 2932.659 | 2914.794 | 2821.588 |
| Unc79 | 217843 | 1.299171 | -0.83482 | 2.939077 | 0.00775 | 0.019038 | 1.144829 | -0.83482 | 3.390927 | 0.00271 | 0.006759 | 21.51744 | 13.32052 | 8.679696 | 31.16902 | 35.10988 | 45.9907 | 71.06304 | 64.34024 | 126.4071 | 18.46115 | 14.48803 | 11.94146 | 61.74019 | 64.45877 | 62.36001 |
| Shank2 | 210274 | 1.299193 | -0.61796 | 2.862107 | 0.009234 | 0.022326 | 0.970116 | -0.61796 | 2.756296 | 0.011722 | 0.024811 | 22.25942 | 12.43249 | 18.22736 | 59.74062 | 32.30109 | 50.04871 | 98.067 | 64.34024 | 127.5563 | 16.92272 | 9.912862 | 5.97073 | 88.43865 | 112.4098 | 140.1076 |
| Col6a5 | 665033 | 1.299984 | -1.17054 | 2.605761 | 0.016376 | 0.037316 | 1.176534 | -1.17054 | 3.066359 | 0.005786 | 0.013292 | 9.64575 | 14.20856 | 11.2836 | 29.87031 | 58.9846 | 16.23201 | 61.11422 | 60.31898 | 97.67823 | 10.769 | 10.67539 | 5.97073 | 63.40885 | 43.23454 | 53.45144 |
| Nfkbiz | 80859 | 1.300708 | 8.052465 | 36.45499 | 1.27E-15 | 2.07E-14 | -1.25347 | 8.052465 | -32.0533 | 7.94E-15 | 1.38E-13 | 30784.04 | 32602.43 | 28693.34 | 81294 | 81995.62 | 78550.77 | 35891.1 | 36180.66 | 33550.75 | 11712.06 | 11871.03 | 12029.53 | 936.9491 | 904.7811 | 1092.515 |
| Trim69 | 70928 | 1.301799 | -1.25686 | 3.594234 | 0.001673 | 0.004636 | -0.50557 | -1.25686 | -1.4628 | 0.158094 | 0.230375 | 17.06556 | 19.53677 | 22.56721 | 57.1432 | 42.13186 | 56.81205 | 28.42522 | 38.87223 | 45.96622 | 9.230573 | 10.67539 | 17.16585 | 11.68058 | 29.08506 | 22.67637 |
| Lpar2 | 53978 | 1.302562 | 2.345158 | 9.569045 | 3.57E-09 | 2.05E-08 | -0.15128 | 2.345158 | -1.24966 | 0.224955 | 0.309564 | 201.0768 | 183.8232 | 215.2565 | 461.0418 | 596.868 | 535.6564 | 532.9728 | 415.5307 | 533.2082 | 249.9947 | 210.4577 | 231.3658 | 226.1026 | 194.1624 | 227.5735 |
| Lysmd2 | 70082 | 1.302693 | 1.090194 | 5.215367 | 3.44E-05 | 0.000119 | 2.473491 | 1.090194 | 14.54625 | 1.49E-12 | 1.63E-11 | 95.71552 | 76.37101 | 48.6063 | 163.6374 | 217.6813 | 193.4315 | 1026.15 | 1188.954 | 1080.206 | 36.15308 | 17.53814 | 35.07804 | 60.90587 | 95.116 | 81.7969 |
| Cxcr5 | 12145 | 1.302702 | 0.362023 | 4.881985 | 7.59E-05 | 0.000252 | -0.51273 | 0.362023 | -2.03316 | 0.05466 | 0.093545 | 52.68063 | 71.0428 | 38.19066 | 119.4812 | 157.2923 | 140.6774 | 95.22448 | 131.3613 | 82.7392 | 60.76794 | 39.65145 | 58.96096 | 82.59836 | 51.88145 | 72.88832 |
| Pcdhgb2 | 93700 | 1.304043 | 0.929198 | 6.5894 | 1.47E-06 | 5.95E-06 | -1.17738 | 0.929198 | -5.55295 | 1.55E-05 | 5.60E-05 | 84.58581 | 95.90778 | 92.87275 | 250.6509 | 235.9384 | 231.3062 | 142.1261 | 124.6592 | 79.29174 | 116.1514 | 102.9413 | 97.02436 | 98.45058 | 67.60311 | 58.31066 |
| Rbfox3 | 52897 | 1.304617 | -1.19731 | 2.548333 | 0.018573 | 0.041758 | 1.091274 | -1.19731 | 2.741929 | 0.012105 | 0.02552 | 8.161788 | 10.65642 | 11.2836 | 15.58451 | 23.87472 | 50.04871 | 56.85043 | 46.91476 | 74.69511 | 9.230573 | 6.862751 | 8.956095 | 105.1252 | 81.75259 | 66.40936 |
| Gm6807 | 627889 | 1.305114 | -0.30873 | 3.274737 | 0.003562 | 0.009338 | -1.01621 | -0.30873 | -2.45304 | 0.022836 | 0.04422 | 45.26083 | 55.9462 | 12.15157 | 79.22126 | 73.02855 | 104.1554 | 41.21656 | 36.19139 | 51.712 | 59.22951 | 59.47717 | 52.24389 | 23.36115 | 47.95104 | 51.02183 |
| Msantd3 | 66665 | 1.305307 | 2.428139 | 9.792689 | 2.38E-09 | 1.41E-08 | 1.496643 | 2.428139 | 15.22894 | 6.10E-13 | 7.10E-12 | 208.4966 | 217.5686 | 240.4276 | 603.8998 | 558.9493 | 588.4105 | 1829.163 | 1671.506 | 1651.337 | 106.9208 | 156.3182 | 173.1512 | 119.3087 | 92.75775 | 113.3818 |
| Mss51 | 74843 | 1.305795 | 0.949772 | 7.393875 | 2.57E-07 | 1.15E-06 | -1.15404 | 0.949772 | -6.10799 | 4.32E-06 | 1.69E-05 | 107.5872 | 112.7804 | 84.19305 | 305.1967 | 286.4966 | 216.4268 | 117.9647 | 146.106 | 112.6172 | 119.9975 | 100.6537 | 117.1756 | 59.23721 | 51.88145 | 49.40209 |
| Col4a3 | 12828 | 1.306471 | -0.10587 | 2.897851 | 0.008514 | 0.020746 | 1.317942 | -0.10587 | 3.908564 | 0.000789 | 0.002205 | 18.54952 | 16.87266 | 34.71878 | 41.55869 | 74.43295 | 66.28072 | 136.441 | 130.0209 | 191.909 | 33.07622 | 25.92595 | 11.19512 | 114.3028 | 162.7191 | 91.51534 |
| Wnt10b | 22410 | 1.307352 | -0.56015 | 3.905912 | 0.000794 | 0.002316 | -0.45692 | -0.56015 | -1.44861 | 0.16198 | 0.234983 | 28.19527 | 33.74533 | 28.643 | 67.53288 | 77.24174 | 97.39208 | 48.32287 | 61.6594 | 70.09849 | 16.92272 | 12.20045 | 8.956095 | 47.55663 | 95.116 | 43.73299 |
| Nrcam | 319504 | 1.307608 | 0.44303 | 3.572514 | 0.001762 | 0.00487 | 0.184826 | 0.44303 | 0.592633 | 0.559654 | 0.654257 | 22.25942 | 23.97694 | 17.35939 | 67.53288 | 66.00658 | 39.22736 | 78.16935 | 48.25518 | 75.84427 | 55.38344 | 51.85189 | 37.31706 | 979.4998 | 889.0594 | 882.7586 |
| Tram2 | 170829 | 1.308128 | 3.026881 | 14.85933 | 9.86E-13 | 9.56E-12 | -2.62908 | 3.026881 | -19.7938 | 3.24E-15 | 6.18E-14 | 534.9681 | 502.6278 | 579.8037 | 1450.658 | 1461.975 | 1360.784 | 304.1498 | 243.9568 | 191.909 | 446.9136 | 475.8174 | 500.0486 | 305.3637 | 286.9202 | 340.1455 |
| Ldhd | 52815 | 1.30981 | 1.342734 | 5.402087 | 2.21E-05 | 7.85E-05 | -0.45889 | 1.342734 | -2.02814 | 0.055209 | 0.094351 | 114.265 | 155.4061 | 80.72117 | 244.1573 | 405.8702 | 271.8862 | 261.512 | 218.4887 | 204.5497 | 69.99852 | 148.6929 | 83.59022 | 117.6401 | 96.68816 | 109.3325 |
| Tmem243 | 652925 | 1.310406 | 4.729248 | 21.70732 | 4.94E-16 | 8.73E-15 | 0.473109 | 4.729248 | 9.242444 | 6.53E-09 | 3.81E-08 | 1133.747 | 1051.433 | 1061.527 | 2854.563 | 2797.555 | 2962.342 | 4002.271 | 4125.818 | 4261.069 | 951.5183 | 982.8984 | 900.0876 | 775.924 | 823.8146 | 882.7586 |
| Cd164 | 53599 | 1.312828 | 8.676227 | 35.24881 | 2.09E-20 | 9.85E-19 | 2.056604 | 8.676227 | 77.42084 | 1.22E-27 | 1.91E-24 | 6664.471 | 6836.093 | 6922.058 | 18150.76 | 17966.43 | 17998.6 | 77696.07 | 76583.66 | 79173.37 | 17400.4 | 17088.25 | 17268.84 | 52350.68 | 51797.34 | 54996.67 |
| Tmem170l | 621976 | 1.314079 | 6.95299 | 36.47252 | 1.02E-20 | 5.22E-19 | -0.50968 | 6.95299 | -14.837 | 1.02E-12 | 1.14E-11 | 6195.539 | 5910.761 | 6002.01 | 15522.17 | 16967.9 | 15711.24 | 12094.93 | 11590.63 | 11453.63 | 5411.424 | 5378.871 | 5748.32 | 3517.522 | 3600.258 | 3569.098 |
| Rbm20 | 73713 | 1.314405 | -0.54451 | 2.588753 | 0.017 | 0.038554 | 1.532687 | -0.54451 | 4.075525 | 0.000528 | 0.001519 | 13.35565 | 5.32821 | 15.62345 | 31.16902 | 16.85274 | 43.28537 | 68.22052 | 79.08488 | 114.9156 | 16.92272 | 19.82572 | 14.92683 | 262.813 | 248.4021 | 221.9045 |
| Zfp985 | 433804 | 1.314944 | -1.4773 | 2.602401 | 0.016497 | 0.037547 | 0.929501 | -1.4773 | 2.325512 | 0.029973 | 0.055874 | 9.64575 | 4.440175 | 13.01954 | 19.48064 | 21.06593 | 28.40602 | 46.90161 | 33.51054 | 57.45778 | 13.84586 | 10.67539 | 10.44878 | 50.89394 | 30.65722 | 43.73299 |
| Map2k1 | 26395 | 1.315392 | 6.961237 | 30.1331 | 5.58E-19 | 1.85E-17 | -0.10291 | 6.961237 | -2.6172 | 0.015969 | 0.032452 | 5167.896 | 5340.642 | 5088.906 | 13803.98 | 13840.32 | 13780.98 | 14201.24 | 13408.24 | 12610.83 | 5001.432 | 4656.758 | 4916.15 | 5085.223 | 4721.998 | 4928.87 |
| Abracl | 73112 | 1.315993 | 6.310275 | 30.64662 | 3.92E-19 | 1.35E-17 | -0.54732 | 6.310275 | -13.2674 | 8.83E-12 | 8.36E-11 | 4094.25 | 4448.167 | 4038.663 | 11540.33 | 11339.09 | 10510.23 | 8223.415 | 7782.489 | 7736.115 | 3585.309 | 3414.6 | 3445.858 | 1884.745 | 1855.155 | 1871.61 |
| Pkhd1l1 | 192190 | 1.316101 | -0.61201 | 3.259691 | 0.00369 | 0.009653 | 1.219199 | -0.61201 | 4.011888 | 0.000615 | 0.001752 | 22.25942 | 10.65642 | 19.09533 | 61.03933 | 32.30109 | 45.9907 | 98.067 | 108.5742 | 122.9596 | 9.999788 | 20.58825 | 7.463413 | 90.10731 | 79.39434 | 99.61404 |
| Dyrk2 | 69181 | 1.316646 | 5.391359 | 21.9855 | 3.81E-16 | 6.95E-15 | -0.05992 | 5.391359 | -1.1079 | 0.280236 | 0.372002 | 1575.225 | 1738.772 | 1657.822 | 4459.767 | 4460.359 | 4266.314 | 4415.857 | 4674.051 | 4094.441 | 1745.348 | 1748.476 | 1658.37 | 1737.903 | 1793.054 | 1906.435 |
| Astn1 | 11899 | 1.317969 | -1.41269 | 2.839968 | 0.009709 | 0.023384 | 0.312904 | -1.41269 | 0.796546 | 0.434486 | 0.533493 | 9.64575 | 16.87266 | 14.75548 | 31.16902 | 32.30109 | 44.63804 | 42.63783 | 34.85097 | 62.0544 | 7.692145 | 2.287584 | 14.92683 | 36.71038 | 43.23454 | 28.34546 |
| Atxn7l1 | 380753 | 1.31857 | 6.572509 | 30.42337 | 4.57E-19 | 1.54E-17 | -0.13322 | 6.572509 | -3.39286 | 0.002698 | 0.00673 | 4328.716 | 4144.459 | 4324.225 | 11710.46 | 11357.34 | 11033.71 | 10973.56 | 11196.54 | 10212.55 | 3553.771 | 3450.439 | 3357.043 | 3424.078 | 3245.735 | 3233.812 |
| Cdkn1a | 12575 | 1.318916 | 9.218038 | 29.23856 | 2.04E-13 | 2.23E-12 | 0.404968 | 9.218038 | 11.08504 | 4.50E-08 | 2.32E-07 | 22749.87 | 22252.38 | 22016.92 | 60565.3 | 60283.67 | 57833.31 | 85316.87 | 84088.68 | 77456.53 | 29164.77 | 29579.22 | 30055.16 | 17651.02 | 17065.07 | 18282.01 |
| Fmr1 | 14265 | 1.319238 | 6.283594 | 35.79388 | 1.52E-20 | 7.36E-19 | -0.60427 | 6.283594 | -16.9188 | 7.61E-14 | 1.08E-12 | 3733.647 | 3745.732 | 3702.758 | 9811.748 | 10180.46 | 9817.662 | 6456.788 | 6745.002 | 7097.185 | 3769.151 | 3715.036 | 3810.072 | 2387.009 | 2234.047 | 2346.194 |
| Togaram2 | 320159 | 1.319327 | -1.62453 | 2.88122 | 0.008842 | 0.021462 | 1.351364 | -1.62453 | 3.915968 | 0.000775 | 0.002168 | 13.35565 | 7.992315 | 6.075787 | 23.37676 | 33.70549 | 17.58468 | 66.79926 | 62.99982 | 64.35271 | 3.846072 | 4.575167 | 3.731706 | 50.05962 | 34.58764 | 42.11325 |
| Leng9 | 243813 | 1.319625 | 1.935528 | 9.071695 | 9.00E-09 | 4.88E-08 | -0.85758 | 1.935528 | -5.85949 | 7.63E-06 | 2.88E-05 | 172.1395 | 199.8079 | 177.9338 | 501.3017 | 483.112 | 473.4337 | 309.8349 | 272.1056 | 259.7092 | 128.4588 | 154.0306 | 128.3707 | 227.7712 | 191.8042 | 138.4878 |
| Bnipl | 171388 | 1.319647 | -1.09581 | 3.174705 | 0.0045 | 0.011563 | -0.18826 | -1.09581 | -0.50285 | 0.620218 | 0.710964 | 17.80754 | 7.992315 | 25.17112 | 31.16902 | 46.34504 | 51.40137 | 35.53152 | 30.8297 | 48.26453 | 13.07665 | 25.16342 | 13.43414 | 39.21337 | 37.73197 | 40.49351 |
| Ublcp1 | 79560 | 1.319914 | 3.880611 | 21.69166 | 5.01E-16 | 8.84E-15 | -1.29913 | 3.880611 | -19.6029 | 3.94E-15 | 7.37E-14 | 811.7269 | 821.4324 | 859.2899 | 2150.662 | 2244.224 | 2246.781 | 994.8826 | 860.5508 | 948.0534 | 562.2958 | 533.007 | 556.0242 | 546.4841 | 560.4769 | 570.1487 |
| Gm6813 | 627939 | 1.319945 | -0.75938 | 3.968983 | 0.000682 | 0.002011 | -0.44025 | -0.75938 | -1.4127 | 0.172161 | 0.247108 | 22.25942 | 23.97694 | 24.30315 | 89.61093 | 57.58021 | 47.34337 | 35.53152 | 61.6594 | 49.41369 | 31.53779 | 35.07628 | 21.6439 | 25.86413 | 26.72681 | 23.48624 |
| Rufy3 | 52822 | 1.320693 | 5.851554 | 22.94691 | 1.58E-16 | 3.14E-15 | -0.34979 | 5.851554 | -6.49699 | 1.80E-06 | 7.41E-06 | 2277.139 | 2352.405 | 2255.853 | 6084.452 | 6165.295 | 6107.295 | 5309.831 | 5241.049 | 4512.734 | 3493.003 | 3498.478 | 3334.653 | 2094.161 | 2024.949 | 1951.787 |
| Ngef | 53972 | 1.321774 | -1.98118 | 2.503967 | 0.020456 | 0.045459 | 0.994492 | -1.98118 | 2.369391 | 0.027313 | 0.051636 | 8.161788 | 5.32821 | 6.075787 | 20.77935 | 14.04395 | 18.93735 | 32.689 | 30.8297 | 49.41369 | 5.384501 | 3.050111 | 2.239024 | 36.71038 | 59.74228 | 41.30338 |
| Cdhr4 | 69398 | 1.32193 | -0.14279 | 2.820648 | 0.010142 | 0.02432 | 2.091543 | -0.14279 | 6.402352 | 2.22E-06 | 9.03E-06 | 28.93725 | 15.98463 | 12.15157 | 55.84449 | 51.96262 | 40.58003 | 235.9293 | 288.1907 | 164.3292 | 12.30743 | 24.40089 | 23.88292 | 108.4625 | 99.04641 | 144.9668 |
| 9930111J2 | 245240 | 1.324261 | 2.663904 | 9.285922 | 6.02E-09 | 3.35E-08 | 0.223823 | 2.663904 | 1.860826 | 0.076602 | 0.124488 | 204.0447 | 269.0746 | 373.2269 | 715.5887 | 686.7493 | 764.2573 | 847.0715 | 855.1891 | 915.877 | 227.6875 | 218.083 | 208.9756 | 239.4518 | 193.3763 | 189.5096 |

| | | | | | | | | | | | | | | | | | | | | | | | | | | |
|---|---|---|---|---|---|---|---|---|---|---|---|---|---|---|---|---|---|---|---|---|---|---|---|---|---|---|
| C78197 | 100504352 | 1.32567 | 1.348541 | 6.972574 | 6.33E-07 | 2.69E-06 | -0.83634 | 1.348541 | -4.40638 | 0.000238 | 0.000726 | 112.7811 | 150.9659 | 84.19305 | 279.2225 | 271.0483 | 362.515 | 196.134 | 194.3612 | 145.9428 | 126.9204 | 115.1417 | 132.8487 | 116.8058 | 107.6933 | 97.18443 |
| Gas7 | 14457 | 1.326018 | 7.820141 | 37.32879 | 6.28E-21 | 3.36E-19 | 0.69252 | 7.820141 | 24.37673 | 4.56E-17 | 1.35E-15 | 10023.42 | 10179.54 | 10321.03 | 28084.59 | 26995.29 | 26559.63 | 46971.25 | 46288.78 | 44096.55 | 19769.58 | 20357.21 | 20139.27 | 1668.654 | 1819.781 | 1817.349 |
| Hif1a | 15251 | 1.326112 | 7.945707 | 50.68894 | 9.83E-24 | 1.09E-21 | -0.39922 | 7.945707 | -16.3858 | 1.44E-13 | 1.93E-12 | 11459.15 | 11957.39 | 11392.1 | 31044.34 | 32157.84 | 29954.83 | 24739.89 | 23897.04 | 24614.91 | 11820.52 | 11718.53 | 11708.6 | 6601.195 | 6786.251 | 6801.29 |
| Frmpd1 | 666060 | 1.326186 | -1.25115 | 3.023147 | 0.006392 | 0.015977 | 1.026387 | -1.25115 | 3.003606 | 0.006685 | 0.015123 | 9.64575 | 9.768385 | 8.679696 | 29.87031 | 15.44835 | 35.16936 | 51.16539 | 56.29771 | 56.30862 | 13.84586 | 16.77561 | 18.65853 | 38.37904 | 44.80671 | 43.73299 |
| Atl1 | 73991 | 1.326645 | 1.490899 | 6.33655 | 2.58E-06 | 1.02E-05 | -0.45436 | 1.490899 | -2.32839 | 0.029791 | 0.055586 | 83.10184 | 93.24367 | 94.60869 | 298.7031 | 216.2769 | 213.7215 | 220.2954 | 154.1485 | 180.4174 | 91.53652 | 104.4663 | 115.6829 | 369.6068 | 268.8403 | 335.2863 |
| Tmem140 | 68487 | 1.327028 | 2.755208 | 10.238 | 1.08E-09 | 6.68E-09 | 0.900418 | 2.755208 | 9.063536 | 9.14E-09 | 5.21E-08 | 179.5593 | 143.8617 | 154.4986 | 390.9115 | 471.8768 | 428.7957 | 818.6463 | 845.8061 | 827.392 | 262.3021 | 285.9479 | 282.8633 | 664.9586 | 709.8326 | 663.2837 |
| Samd15 | 238333 | 1.327757 | -0.90445 | 3.658939 | 0.001434 | 0.004021 | -0.65715 | -0.90445 | -1.86716 | 0.075675 | 0.12321 | 24.48536 | 16.87266 | 24.30315 | 37.66257 | 81.45493 | 67.63339 | 32.689 | 37.53181 | 44.81707 | 29.23015 | 20.58825 | 14.18048 | 26.69846 | 28.29897 | 38.87377 |
| Fam177a2 | 100101807 | 1.32804 | -1.60176 | 2.920001 | 0.008095 | 0.019806 | 0.951537 | -1.60176 | 2.659381 | 0.014547 | 0.029891 | 8.161788 | 8.88035 | 9.547666 | 35.06515 | 16.85274 | 22.99535 | 59.69296 | 40.21265 | 49.41369 | 16.1535 | 18.30067 | 20.89756 | 15.01788 | 10.21907 | 16.19741 |
| Tlr6 | 21899 | 1.329095 | 4.706255 | 22.43465 | 2.51E-16 | 4.78E-15 | -0.49579 | 4.706255 | -8.78353 | 1.56E-08 | 8.57E-08 | 1289.563 | 1393.327 | 1212.554 | 3588.333 | 3641.597 | 3212.586 | 2535.529 | 2578.971 | 2558.02 | 1775.347 | 1743.901 | 1692.702 | 453.0395 | 459.8583 | 471.3445 |
| C1ra | 50909 | 1.330316 | 1.557791 | 8.890038 | 1.27E-08 | 6.74E-08 | -0.92375 | 1.557791 | -6.03878 | 5.06E-06 | 1.96E-05 | 141.7183 | 168.7266 | 153.6306 | 441.5611 | 376.3779 | 420.6797 | 193.2915 | 230.5525 | 250.5159 | 231.5336 | 254.6843 | 322.4194 | 45.88798 | 40.8763 | 44.54286 |
| Furin | 18550 | 1.33091 | 8.251582 | 25.42175 | 1.14E-14 | 1.56E-13 | 0.295545 | 8.251582 | 6.717065 | 4.22E-06 | 1.65E-05 | 10090.94 | 10380.24 | 10099.69 | 27528.74 | 27496.65 | 27038.48 | 37320.89 | 35739.67 | 32240.71 | 12221.28 | 12892.82 | 12384.04 | 16241.01 | 16115.48 | 16980.55 |
| Daam1 | 208846 | 1.331188 | 5.893665 | 34.18436 | 3.99E-20 | 1.74E-18 | -1.75544 | 5.893665 | -38.8898 | 2.65E-21 | 2.58E-19 | 3986.663 | 3787.469 | 3821.67 | 10648.12 | 10329.33 | 10207.23 | 3071.345 | 3278.672 | 3229.127 | 3320.699 | 3146.952 | 3374.209 | 1284.863 | 1315.116 | 1447.238 |
| Gm9984 | 791265 | 1.332235 | 1.716083 | 9.129436 | 8.07E-09 | 4.41E-08 | -3.61291 | 1.716083 | -12.3668 | 3.35E-11 | 2.85E-10 | 373.9583 | 276.1789 | 278.6182 | 798.7061 | 806.1229 | 899.524 | 110.8583 | 34.85097 | 87.33582 | 234.6104 | 295.8608 | 221.6634 | 86.77 | 55.02578 | 56.69092 |
| Serpinc1 | 11905 | 1.332499 | -0.64469 | 4.006124 | 0.000624 | 0.00185 | -0.9304 | -0.64469 | -2.74381 | 0.012054 | 0.025426 | 29.67923 | 25.75301 | 28.643 | 72.72771 | 73.02855 | 82.51273 | 48.32287 | 26.80843 | 51.712 | 14.61507 | 37.36386 | 36.57072 | 35.04173 | 36.94588 | 26.72572 |
| Zbtb5 | 230119 | 1.333095 | 4.750909 | 18.81731 | 9.01E-15 | 1.26E-13 | -1.48195 | 4.750909 | -18.7807 | 9.37E-15 | 1.61E-13 | 1233.914 | 1268.114 | 1334.937 | 3327.293 | 3529.245 | 3456.066 | 1321.773 | 1349.805 | 1179.034 | 2686.866 | 2725.275 | 2554.726 | 701.6689 | 847.3971 | 631.6988 |
| Ptprb | 19263 | 1.333132 | -0.77965 | 3.644356 | 0.001485 | 0.004151 | 0.882263 | -0.77965 | 3.098648 | 0.005369 | 0.012447 | 17.80754 | 16.87266 | 17.35939 | 40.25998 | 58.9846 | 44.63804 | 100.9095 | 69.70193 | 104.5732 | 6.153716 | 10.67539 | 8.209754 | 80.92971 | 90.3995 | 62.36001 |
| Ndufa11b | 239760 | 1.333705 | -2.55034 | 2.52271 | 0.019639 | 0.043864 | -0.76399 | -2.55034 | -1.45558 | 0.16006 | 0.232557 | 6.677827 | 19.53677 | 3.471878 | 19.48064 | 22.47032 | 21.64268 | 14.21261 | 9.382952 | 14.93902 | 3.076858 | 3.050111 | 7.463413 | 11.68058 | 10.21907 | 8.098703 |
| Arl4a | 11861 | 1.333725 | 3.470365 | 10.03762 | 1.54E-09 | 9.32E-09 | 2.636537 | 3.470365 | 28.74825 | 1.49E-18 | 6.41E-17 | 254.4994 | 293.0515 | 255.1831 | 776.6281 | 780.8438 | 605.9951 | 4936.039 | 4651.263 | 4363.344 | 279.9941 | 343.9001 | 331.3755 | 467.2231 | 423.6985 | 463.2458 |
| Gm6548 | 625054 | 1.334112 | 6.445913 | 30.53871 | 4.22E-19 | 1.44E-17 | -0.2227 | 6.445913 | -5.56116 | 1.52E-05 | 5.50E-05 | 3344.849 | 3394.958 | 3406.781 | 8894.859 | 9243.73 | 9179.203 | 8580.152 | 8121.615 | 7680.956 | 4866.82 | 5082.248 | 4893.013 | 3132.898 | 3090.877 | 3093.704 |
| Mcmdc2 | 240697 | 1.334888 | 0.855112 | 6.844547 | 8.36E-07 | 3.50E-06 | -0.93011 | 0.855112 | -4.66556 | 0.000128 | 0.000406 | 90.52165 | 93.24367 | 85.92899 | 270.1315 | 255.5999 | 205.6055 | 117.9647 | 113.9358 | 163.1801 | 98.45945 | 100.6537 | 62.69267 | 53.39692 | 66.03094 | 60.74027 |
| Zc3hav1 | 78781 | 1.335142 | 7.836727 | 36.46022 | 1.03E-20 | 5.25E-19 | -0.28366 | 7.836727 | -8.47066 | 2.86E-08 | 1.52E-07 | 9459.513 | 10220.39 | 9462.605 | 26246.91 | 26206.02 | 25887.36 | 23455.07 | 22500.32 | 21130.67 | 10309.78 | 10484 | 10765.97 | 8051.255 | 7936.29 | 8003.138 |
| Dnaaf9 | 228602 | 1.335471 | 6.066172 | 23.38132 | 1.08E-16 | 2.18E-15 | -0.47724 | 6.066172 | -8.79742 | 1.52E-08 | 8.36E-08 | 2453.73 | 2608.159 | 2767.087 | 7110.433 | 6919.455 | 6990.587 | 5399.37 | 5590.899 | 4783.935 | 2841.478 | 2883.118 | 2912.224 | 3917.165 | 3590.825 | 3701.107 |
| Fstl4 | 320027 | 1.336995 | -1.79157 | 2.489411 | 0.021111 | 0.046762 | 0.949149 | -1.79157 | 2.222598 | 0.037173 | 0.067268 | 6.677827 | 7.10428 | 6.943757 | 32.46773 | 12.63956 | 16.23201 | 39.7953 | 32.17012 | 44.81707 | 7.692145 | 6.100223 | 6.717071 | 26.69846 | 53.45362 | 37.25403 |
| Gm6833 | 628058 | 1.337107 | -1.56357 | 2.722271 | 0.012648 | 0.029657 | 0.697209 | -1.56357 | 1.752573 | 0.094031 | 0.147992 | 6.677827 | 12.43249 | 8.679696 | 19.48064 | 23.87472 | 31.11136 | 44.05909 | 33.51054 | 47.11538 | 20.76879 | 6.100223 | 8.956095 | 27.53279 | 29.87114 | 37.25403 |
| 4833419F2 | 73915 | 1.337115 | 1.585959 | 7.922605 | 8.58E-08 | 4.09E-07 | -0.70301 | 1.585959 | -4.26057 | 0.000338 | 0.001004 | 120.2009 | 174.9429 | 135.4033 | 427.2753 | 405.8702 | 312.4662 | 206.0828 | 260.0418 | 257.4109 | 154.6121 | 188.3444 | 169.4195 | 80.92971 | 102.9768 | 70.45871 |
| Ogfrl1 | 70155 | 1.33782 | 6.352538 | 35.8821 | 1.44E-20 | 7.02E-19 | -2.08679 | 6.352538 | -45.3931 | 1.01E-22 | 1.74E-20 | 5075.148 | 5568.867 | 5182.647 | 14057.23 | 14726.49 | 13848.61 | 3365.546 | 3502.522 | 3527.908 | 3974.531 | 3782.138 | 3921.277 | 2785.818 | 2855.838 | 2619.93 |
| Sall3 | 20689 | 1.338267 | -0.81327 | 2.531248 | 0.019278 | 0.043144 | 1.352409 | -0.81327 | 3.4292 | 0.002476 | 0.00623 | 14.09763 | 6.216245 | 18.22736 | 37.66257 | 44.94065 | 20.29002 | 76.74809 | 84.44657 | 99.97654 | 28.46093 | 9.912862 | 5.97073 | 105.1252 | 88.82734 | 121.4805 |
| Pla2g4a | 18783 | 1.338431 | 6.184547 | 33.4366 | 6.34E-20 | 2.58E-18 | -0.03093 | 6.184547 | -0.86404 | 0.397174 | 0.496228 | 3232.81 | 3136.54 | 3320.852 | 8554.597 | 8839.264 | 8782.872 | 8926.939 | 8644.38 | 8993.292 | 3307.622 | 3256.756 | 3361.521 | 1761.264 | 1873.235 | 2017.387 |
| Gm16364 | 100504626 | 1.339155 | 0.307816 | 4.680805 | 0.000123 | 0.000398 | -1.07686 | 0.307816 | -3.58011 | 0.001731 | 0.004497 | 33.38913 | 39.96157 | 54.68209 | 118.1825 | 147.4615 | 81.16006 | 46.90161 | 58.97856 | 58.60694 | 106.1516 | 118.1918 | 104.4878 | 85.10135 | 93.54383 | 55.88105 |
| Lyn | 17096 | 1.339184 | 8.947647 | 35.36798 | 7.59E-14 | 8.99E-13 | -1.23808 | 8.947647 | -30.0458 | 5.63E-13 | 6.60E-12 | 38477.64 | 39332.85 | 34119.02 | 101634.4 | 103606.5 | 97057.97 | 44921.79 | 44275.47 | 43867.87 | 23797.19 | 23239.56 | 23836.65 | 5740.169 | 5496.29 | 5861.841 |
| ATP8 | 17706 | 1.339234 | -1.18626 | 3.415846 | 0.002555 | 0.006865 | -2.78165 | -1.18626 | -4.755 | 0.000103 | 0.000333 | 59.35846 | 43.51371 | 11.2836 | 94.80577 | 74.43295 | 90.62874 | 8.527565 | 10.72337 | 19.53565 | 46.92208 | 54.90201 | 70.90242 | 5.005962 | 5.502578 | 7.288832 |
| Nr3c1 | 14815 | 1.340993 | 7.910652 | 43.28722 | 2.77E-22 | 1.96E-20 | -0.02893 | 7.910652 | -1.06403 | 0.299207 | 0.393044 | 10712.72 | 11004.53 | 11055.33 | 29733.95 | 30521.72 | 28355.97 | 30909.58 | 30024.11 | 29328.75 | 9401.339 | 9887.699 | 9684.524 | 6738.859 | 6374.344 | 6604.492 |
| Slit3 | 20564 | 1.34122 | -1.11814 | 2.733197 | 0.012343 | 0.02901 | 1.172843 | -1.11814 | 3.123505 | 0.005069 | 0.011822 | 10.38773 | 10.65642 | 8.679696 | 29.87031 | 19.66153 | 33.81669 | 52.58665 | 61.6594 | 79.29174 | 22.30722 | 12.96297 | 17.16585 | 32.53875 | 62.88661 | 57.50079 |
| Hs3st3a1 | 15478 | 1.341466 | -0.79501 | 4.088012 | 0.000512 | 0.001538 | -1.56978 | -0.79501 | -4.14234 | 0.000449 | 0.001308 | 34.13111 | 31.96926 | 21.69924 | 79.22126 | 68.81537 | 89.27607 | 18.47639 | 34.85097 | 29.87805 | 30.76858 | 12.20045 | 10.44878 | 51.72827 | 54.2397 | 36.44416 |
| Sgk3 | 170755 | 1.341514 | 6.890908 | 44.0596 | 1.91E-22 | 1.39E-20 | -1.46576 | 6.890908 | -43.3243 | 2.72E-22 | 3.97E-20 | 8919.351 | 8523.36 | 8314.281 | 23184.56 | 24162.62 | 22488.1 | 8592.943 | 8948.656 | 8681.87 | 8423.667 | 8263.514 | 8156.764 | 1257.331 | 1203.492 | 1267.447 |
| Gpr34 | 23890 | 1.342677 | 0.915714 | 6.086076 | 4.54E-06 | 1.75E-05 | -1.7612 | 0.915714 | -6.49938 | 1.79E-06 | 7.37E-06 | 51.93865 | 71.93083 | 97.2126 | 229.8715 | 157.2923 | 189.3735 | 45.48035 | 52.27645 | 76.99342 | 62.30637 | 60.2397 | 62.69267 | 364.6009 | 370.2449 | 384.6884 |
| Csmd1 | 94109 | 1.343589 | -0.63802 | 2.831302 | 0.009901 | 0.023803 | 1.20651 | -0.63802 | 3.371418 | 0.002838 | 0.007039 | 12.61367 | 15.09659 | 26.90706 | 58.44191 | 42.13186 | 40.58003 | 147.8111 | 79.08488 | 121.8105 | 7.692145 | 20.58825 | 14.18048 | 80.09538 | 82.53868 | 42.11325 |
| Pcdhga5 | 93713 | 1.343698 | 0.352846 | 5.747035 | 9.89E-06 | 3.66E-05 | -1.51998 | 0.352846 | -5.64746 | 1.25E-05 | 4.56E-05 | 66.03629 | 56.83424 | 84.19305 | 163.6374 | 210.6593 | 185.3155 | 81.01187 | 50.93603 | 71.24765 | 69.99852 | 44.22662 | 54.48291 | 39.21337 | 80.96651 | 49.40209 |
| Kdelr3 | 105785 | 1.343969 | 0.659192 | 5.89177 | 7.08E-06 | 2.66E-05 | -0.0774 | 0.659192 | -0.38833 | 0.701622 | 0.782917 | 58.61648 | 52.39406 | 54.68209 | 146.7541 | 174.145 | 133.9141 | 129.3347 | 151.4677 | 162.0309 | 91.53652 | 55.66453 | 62.69267 | 81.76404 | 103.7629 | 78.55742 |
| Fbxo40 | 207215 | 1.344145 | 0.67338 | 5.463264 | 1.92E-05 | 6.85E-05 | -0.42241 | 0.67338 | -1.85763 | 0.077073 | 0.125122 | 83.10184 | 95.01974 | 59.8899 | 166.2348 | 223.2988 | 259.7122 | 171.9726 | 143.4251 | 176.97 | 41.53758 | 89.21576 | 62.69267 | 60.90587 | 47.95104 | 33.20468 |
| Gbp8 | 76074 | 1.345484 | 1.896905 | 6.385952 | 2.31E-06 | 9.15E-06 | 0.057199 | 1.896905 | 0.318659 | 0.753083 | 0.828807 | 76.42402 | 101.236 | 66.83366 | 227.2741 | 210.6593 | 213.7215 | 288.516 | 223.8504 | 206.848 | 432.2985 | 411.765 | 394.8145 | 362.0979 | 272.7707 | 319.8988 |
| Mug2 | 17837 | 1.347317 | -2.10786 | 2.815044 | 0.010271 | 0.024576 | 0.44785 | -2.10786 | 1.122229 | 0.274232 | 0.365564 | 9.64575 | 9.768385 | 6.075787 | 23.37676 | 28.08791 | 18.93735 | 27.00396 | 28.14886 | 44.81707 | 4.615287 | 3.812639 | 1.492683 | 27.53279 | 21.22423 | 17.00728 |
| Fbln2 | 14115 | 1.347337 | -2.04209 | 2.612599 | 0.016131 | 0.036851 | 0.817073 | -2.04209 | 1.97956 | 0.060795 | 0.102432 | 7.419807 | 7.992315 | 8.679696 | 27.27289 | 22.47032 | 17.58468 | 25.5827 | 54.95729 | 47.11538 | 2.307643 | 0.762528 | 3.731706 | 36.71038 | 26.72681 | 42.11325 |
| Ifi207 | 226691 | 1.34841 | 9.208136 | 78.41254 | 9.33E-28 | 4.02E-25 | -0.52343 | 9.208136 | -32.4437 | 1.19E-19 | 7.14E-18 | 26421.93 | 25773.44 | 25653.71 | 71262.77 | 71357.32 | 69229.53 | 50942.25 | 51171.94 | 50871.97 | 35820.78 | 35476.61 | 35331.05 | 16596.43 | 16572.19 | 16726.25 |
| Mir22hg | 100042498 | 1.349095 | 5.174591 | 24.18634 | 5.36E-17 | 1.16E-15 | -0.50233 | 5.174591 | -9.47158 | 4.27E-09 | 2.57E-08 | 1667.231 | 1629.544 | 1647.406 | 4468.858 | 4501.087 | 4488.152 | 3396.813 | 3471.692 | 3042.964 | 1571.505 | 1704.25 | 1602.395 | 1128.844 | 1209.781 | 1209.136 |
| 6030445D1 | 320133 | 1.3491 | -0.70867 | 3.292847 | 0.003414 | 0.008979 | -0.55363 | -0.70867 | -1.41502 | 0.171489 | 0.246332 | 20.03348 | 31.08122 | 16.49142 | 36.36386 | 78.64613 | 74.39673 | 51.16539 | 20.10633 | 65.50187 | 27.69172 | 34.31375 | 31.34633 | 40.88202 | 33.01547 | 38.0639 |
| Ly6c2 | 100041546 | 1.349698 | 5.811438 | 24.3965 | 4.48E-17 | 9.89E-16 | 0.461028 | 5.811438 | 9.903924 | 1.95E-09 | 1.24E-08 | 3169.742 | 2945.612 | 2954.569 | 8215.634 | 7710.13 | 8823.452 | 11758.09 | 11705.9 | 11837.45 | 1527.66 | 1566.995 | 1319.531 | 941.1208 | 995.9667 | 1038.254 |
| Depp1 | 213393 | 1.350649 | 2.890545 | 9.985935 | 1.69E-09 | 1.02E-08 | -1.58211 | 2.890545 | -10.1143 | 1.34E-09 | 8.75E-09 | 458.5441 | 500.8517 | 479.9872 | 1301.307 | 1200.758 | 1413.538 | 544.3429 | 532.1474 | 336.7026 | 322.3009 | 455.2291 | 444.8194 | 196.9012 | 115.5541 | 139.2977 |
| Rnaset2a | 100037283 | 1.351442 | 1.328162 | 7.031037 | 5.58E-07 | 2.39E-06 | -0.01862 | 1.328162 | -0.11292 | 0.911147 | 0.928876 | 101.6514 | 105.6762 | 89.40087 | 315.5863 | 242.9604 | 252.9489 | 336.8388 | 285.5098 | 228.682 | 77.69066 | 112.0916 | 101.5024 | 95.11327 | 113.982 | 106.9029 |
| Doc2b | 13447 | 1.351549 | -1.30711 | 2.828953 | 0.009954 | 0.023916 | 0.248183 | -1.30711 | 0.614231 | 0.545555 | 0.640675 | 11.87169 | 8.88035 | 21.69924 | 31.16902 | 33.70549 | 45.9907 | 52.58665 | 25.46801 | 64.35271 | 9.230573 | 3.050111 | 8.956095 | 41.71635 | 60.52836 | 59.9304 |
| Sla2 | 77799 | 1.353489 | -0.13593 | 3.636172 | 0.001514 | 0.004228 | -0.21655 | -0.13593 | -0.65526 | 0.519301 | 0.615897 | 54.90658 | 15.09659 | 42.53051 | 68.83159 | 98.30767 | 120.3874 | 65.378 | 88.46784 | 95.37991 | 25.38408 | 22.87584 | 23.13658 | 90.94163 | 88.04125 | 54.26131 |
| Ppp1r26 | 241289 | 1.353715 | 0.387598 | 4.689614 | 0.00012 | 0.00039 | 0.314505 | 0.387598 | 1.317792 | 0.201544 | 0.282393 | 26.71131 | 55.05817 | 37.32269 | 75.32513 | 119.3736 | 120.3874 | 133.5985 | 155.4889 | 113.7664 | 60.76794 | 61.76476 | 50.75121 | 89.27298 | 99.04641 | 95.56469 |
| Flvcr2 | 217721 | 1.353902 | 0.776683 | 5.174916 | 3.78E-05 | 0.00013 | -0.39649 | 0.776683 | -1.63867 | 0.11594 | 0.176896 | 37.09904 | 59.49834 | 87.66493 | 168.8322 | 165.7186 | 132.5614 | 146.3899 | 92.4891 | 133.302 | 93.84416 | 107.5164 | 94.78534 | 91.77596 | 95.90208 | 121.4805 |
| Asah2 | 54447 | 1.354169 | 4.478408 | 18.29588 | 1.59E-14 | 2.12E-13 | -1.87127 | 4.478408 | -20.9255 | 1.04E-15 | 2.25E-14 | 1170.846 | 1031.009 | 1146.588 | 2927.29 | 3248.366 | 2992.101 | 825.7526 | 792.1893 | 968.7382 | 2490.716 | 2601.745 | 2337.541 | 525.626 | 560.4769 | 660.8541 |
| Aoah | 27052 | 1.354233 | 6.754471 | 37.0883 | 7.19E-21 | 3.81E-19 | -1.51541 | 6.754471 | -37.1335 | 7.01E-21 | 5.97E-19 | 8309.442 | 8548.225 | 8068.645 | 22802.74 | 22876.19 | 22339.31 | 8352.75 | 8840.081 | 7631.542 | 9582.874 | 9236.5 | 9590.485 | 715.8525 | 753.0671 | 813.1097 |
| Pi4k2a | 84095 | 1.354357 | 7.594309 | 29.92344 | 6.46E-19 | 2.11E-17 | -0.49804 | 7.594309 | -11.6623 | 1.01E-10 | 7.87E-10 | 7614.206 | 7831.58 | 7982.716 | 21820.91 | 21398.77 | 20662 | 17207.21 | 15740.57 | 14381.68 | 10052.09 | 9988.352 | 9865.885 | 7906.916 | 7930.788 | 7822.537 |
| Gng4 | 14706 | 1.355953 | -0.53934 | 2.888769 | 0.008692 | 0.02114 | 1.655418 | -0.53934 | 4.925751 | 6.84E-05 | 0.000226 | 25.96933 | 11.54445 | 8.679696 | 25.97418 | 58.9846 | 40.58003 | 144.9686 | 140.7443 | 111.4681 | 44.61444 | 29.73859 | 24.62926 | 42.55067 | 41.66238 | 60.74027 |
| Naalad2 | 72560 | 1.35668 | -0.3686 | 4.166856 | 0.000424 | 0.001286 | 0.620335 | -0.3686 | 2.396656 | 0.025772 | 0.049114 | 25.22735 | 36.40943 | 17.35939 | 59.74062 | 67.41097 | 82.51273 | 108.0158 | 115.2763 | 109.1698 | 37.69151 | 44.98914 | 29.10731 | 23.36115 | 40.09021 | 24.29611 |
| Igf2bp1 | 140486 | 1.356834 | 0.648085 | 4.597535 | 0.00015 | 0.000481 | 2.237832 | 0.648085 | 11.00615 | 2.92E-10 | 2.11E-09 | 24.48536 | 33.74533 | 33.85081 | 90.90964 | 82.85932 | 74.39673 | 396.5318 | 414.1903 | 409.0994 | 16.92272 | 19.82572 | 8.956095 | 486.4126 | 418.196 | 498.8801 |
| Txlng | 353170 | 1.358071 | 4.071935 | 18.23585 | 1.70E-14 | 2.25E-13 | 0.106625 | 4.071935 | 1.647793 | 0.11404 | 0.174493 | 653.685 | 758.3819 | 622.3342 | 1809.102 | 1922.617 | 1809.869 | 2023.875 | 2128.59 | 2037.453 | 644.6017 | 605.4471 | 649.3169 | 594.0408 | 624.1496 | 670.5726 |
| Ush1c | 72088 | 1.358273 | -1.64001 | 2.538802 | 0.018963 | 0.04253 | 1.354064 | -1.64001 | 3.340753 | 0.00305 | 0.007513 | 8.903769 | 4.440175 | 9.547666 | 19.48064 | 18.25714 | 24.34802 | 52.58665 | 41.55307 | 74.69511 | 9.999788 | 4.575167 | 3.731706 | 60.90587 | 38.51805 | 46.97248 |
| Gm15337 | 100503620 | 1.358677 | 1.304395 | 6.376118 | 2.36E-06 | 9.34E-06 | 0.302847 | 1.304395 | 1.730226 | 0.098026 | 0.153262 | 92.74759 | 155.4061 | 72.04148 | 315.5863 | 285.0922 | 231.3062 | 318.3624 | 402.1265 | 343.5975 | 99.22866 | 93.79093 | 73.14144 | 83.43269 | 76.25001 | 79.36729 |
| Tmem50b | 77975 | 1.358991 | 6.023833 | 33.59039 | 5.76E-20 | 2.38E-18 | -2.16602 | 6.023833 | -43.088 | 3.05E-22 | 4.29E-20 | 4291.617 | 4249.247 | 4614.126 | 11880.59 | 12181.72 | 11933.23 | 2927.797 | 2810.864 | 2616.627 | 3559.924 | 3639.545 | 3478.697 | 1616.926 | 1686.147 | 1828.687 |
| Spen | 56381 | 1.360653 | 7.797188 | 43.10228 | 3.03E-22 | 2.11E-20 | -2.34369 | 7.797188 | -56.3864 | 1.03E-24 | 3.07E-22 | 11859.08 | 12516.85 | 11672.46 | 33036.56 | 33117.04 | 32565.48 | 7079.3 | 7030.512 | 6196.247 | 14456.62 | 14844.89 | 14876.82 | 10610.97 | 10350.35 | 10734.02 |
| Stxbp6 | 217517 | 1.361542 | 1.737674 | 7.518587 | 1.98E-07 | 8.97E-07 | 1.91885 | 1.737674 | 15.21628 | 6.20E-13 | 7.20E-12 | 82.35986 | 70.15476 | 85.06102 | 212.9883 | 234.534 | 206.9582 | 828.5951 | 944.9973 | 807.8564 | 125.382 | 114.3792 | 127.6244 | 176.043 | 160.3609 | 179.7912 |
| Rab43 | 69834 | 1.361563 | 6.754215 | 36.28457 | 1.14E-20 | 5.74E-19 | -1.6058 | 6.754215 | -37.9786 | 4.37E-21 | 3.96E-19 | 4808.777 | 4801.605 | 4751.266 | 13172.81 | 13890.87 | 12402.61 | 4526.716 | 4533.306 | 4390.924 | 6900.623 | 6901.64 | 7250.705 | 5591.659 | 5573.326 | 5790.572 |
| Abcc9 | 20928 | 1.362 | -1.19438 | 2.643632 | 0.015063 | 0.034683 | 1.242521 | -1.19438 | 3.223521 | 0.004016 | 0.009609 | 21.51744 | 6.216245 | 9.547666 | 29.87031 | 23.87472 | 43.28537 | 65.378 | 77.74446 | 90.78329 | 11.53822 | 8.387806 | 5.97073 | 65.91183 | 55.81187 | 31.58494 |
| Tbx18 | 76365 | 1.363573 | -1.84343 | 2.589396 | 0.016976 | 0.038509 | 1.348681 | -1.84343 | 3.359663 | 0.002918 | 0.007213 | 5.193865 | 9.768385 | 6.943757 | 16.88322 | 23.87472 | 18.93735 | 39.7953 | 52.27645 | 67.80018 | 2.307643 | 6.100223 | 2.985365 | 30.03577 | 51.09537 | 43.73299 |
| Vasn | 246154 | 1.364167 | 4.889262 | 22.37186 | 2.66E-16 | 5.04E-15 | -2.02024 | 4.889262 | -27.2057 | 4.69E-18 | 1.75E-16 | 1802.271 | 1798.271 | 1868.739 | 4983.147 | 4926.619 | 5119.847 | 1334.564 | 1377.954 | 1161.796 | 1506.891 | 1571.57 | 1294.156 | 1040.406 | 1021.121 | 997.7602 |
| Gsdmd | 69146 | 1.365069 | 6.120289 | 20.94241 | 1.03E-15 | 1.71E-14 | 0.265698 | 6.120289 | 4.741075 | 0.000106 | 0.000343 | 2579.867 | 2605.495 | 2517.98 | 7033.809 | 7219.996 | 6941.891 | 9486.916 | 9349.442 | 7861.373 | 3181.471 | 3141.615 | 3191.355 | 2551.372 | 2381.044 | 2378.589 |
| Adgrg2 | 237175 | 1.365137 | -1.73534 | 2.528533 | 0.019392 | 0.043374 | -0.07364 | -1.73534 | -0.15476 | 0.878464 | 0.918721 | 20.77546 | 3.55214 | 13.01954 | 20.77935 | 29.4923 | 40.58003 | 19.89765 | 24.12759 | 44.81707 | 4.615287 | 6.100223 | 3.731706 | 51.72827 | 40.09021 | 32.39481 |
| Dtx2 | 74198 | 1.367815 | 5.797211 | 29.38819 | 9.41E-19 | 2.94E-17 | -1.96849 | 5.797211 | -35.3661 | 1.95E-20 | 1.48E-18 | 3433.145 | 3703.994 | 3436.292 | 9950.71 | 9728.246 | 9402.393 | 2531.266 | 2821.588 | 2397.139 | 3656.846 | 3550.33 | 3431.677 | 1339.095 | 1253.016 | 1308.75 |
| Casp7 | 12369 | 1.368662 | 5.506095 | 30.01468 | 6.06E-19 | 2.00E-17 | 0.484206 | 5.506095 | 12.61293 | 2.31E-11 | 2.03E-10 | 1618.26 | 1611.783 | 1574.497 | 4435.092 | 4304.471 | 4513.852 | 6382.882 | 6368.344 | 6477.79 | 1984.573 | 1967.322 | 2028.556 | 1620.263 | 1661.779 | 1624.6 |
| Gm11702 | 100503586 | 1.368722 | -0.10737 | 5.488898 | 1.80E-05 | 6.47E-05 | -1.61924 | -0.10737 | -5.59259 | 1.42E-05 | 5.14E-05 | 68.26223 | 53.2821 | 41.66254 | 145.4554 | 176.9538 | 129.8561 | 52.58665 | 57.63814 | 42.51876 | 57.69108 | 35.83881 | 29.85365 | 32.53875 | 40.8763 | 23.48624 |
| Dpysl5 | 65254 | 1.370557 | -1.21922 | 3.28367 | 0.003488 | 0.009159 | 0.829605 | -1.21922 | 2.532944 | 0.019207 | 0.038102 | 13.35565 | 8.88035 | 13.01954 | 32.46773 | 23.87472 | 44.63804 | 49.74413 | 52.27645 | 81.59005 | 10.769 | 8.387806 | 10.44878 | 47.55663 | 43.23454 | 44.54286 |
| Tas1r1 | 110326 | 1.371806 | -1.1333 | 2.738912 | 0.012187 | 0.028677 | 0.841767 | -1.1333 | 2.1235 | 0.045561 | 0.079967 | 5.193865 | 12.43249 | 14.75548 | 22.07806 | 35.10988 | 25.70069 | 51.16539 | 48.25518 | 52.86116 | 19.23036 | 12.20045 | 20.89756 | 57.56856 | 43.23454 | 80.17716 |
| Thsd7a | 330267 | 1.371902 | -1.09015 | 3.099768 | 0.005355 | 0.013591 | 0.831184 | -1.09015 | 2.393331 | 0.025955 | 0.049417 | 11.12971 | 11.54445 | 24.30315 | 42.8574 | 47.74944 | 32.46403 | 54.00791 | 67.02109 | 109.1698 | 7.692145 | 6.100223 | 2.239024 | 70.08346 | 66.81702 | 84.22651 |
| Xdh | 22436 | 1.372371 | 9.221535 | 37.37572 | 6.11E-21 | 3.28E-19 | 0.129412 | 9.221535 | 4.163467 | 0.000427 | 0.001248 | 14249.74 | 14280.49 | 14354.48 | 39508.03 | 40227.5 | 38826.97 | 46439.7 | 46350.44 | 42418.78 | 46149.79 | 46360.17 | 45532.79 | 49988.7 | 48448.63 | 49867.76 |
| Mrgpra2a | 668727 | 1.372577 | -2.12023 | 2.608868 | 0.016264 | 0.037097 | 0.11613 | -2.12023 | 0.260327 | 0.797107 | 0.866962 | 19.2915 | 19.53677 | 6.943757 | 66.23417 | 16.85274 | 51.40137 | 63.95674 | 46.91476 | 31.0272 | 1.538429 | 1.525056 | 2.985365 | 6.674615 | 5.502578 | 12.95792 |
| Mustn1 | 66175 | 1.372694 | -1.34444 | 3.645911 | 0.001479 | 0.004136 | -0.73633 | -1.34444 | -1.99592 | 0.05886 | 0.099599 | 14.09763 | 15.98463 | 10.41564 | 37.66257 | 40.72746 | 33.81669 | 21.31891 | 17.42548 | 31.0272 | 18.46115 | 18.30067 | 14.18048 | 35.87606 | 39.30413 | 28.34546 |
| Il4ra | 16190 | 1.373293 | 8.171983 | 53.38392 | 3.28E-24 | 4.36E-22 | -1.98907 | 8.171983 | -62.145 | 1.31E-25 | 5.54E-23 | 18555.45 | 18965.76 | 17610.24 | 49944.46 | 51357.33 | 51225.53 | 13456.5 | 13641.47 | 12842.96 | 14655.84 | 14500.23 | 14749.94 | 8280.695 | 8122.592 | 8719.873 |
| Trpm1 | 17364 | 1.37381 | -1.22775 | 2.656964 | 0.014625 | 0.033766 | 1.388895 | -1.22775 | 3.636364 | 0.001514 | 0.003988 | 19.2915 | 11.54445 | 6.075787 | 31.16902 | 32.30109 | 32.46403 | 73.90556 | 71.04235 | 121.8105 | 3.076858 | 6.100223 | 6.717071 | 80.09538 | 67.60311 | 41.30338 |
| Hydin | 244653 | 1.373911 | 0.385806 | 5.409791 | 2.17E-05 | 7.72E-05 | 0.768086 | 0.385806 | 3.891111 | 0.000822 | 0.002292 | 35.61508 | 47.95389 | 37.32269 | 96.10448 | 117.9692 | 119.0348 | 174.8151 | 174.2548 | 235.5769 | 30.76858 | 25.92595 | 23.13658 | 130.9893 | 109.2655 | 134.4385 |
| Tnk1 | 83813 | 1.374921 | -0.57004 | 3.067985 | 0.005764 | 0.014539 | 1.740522 | -0.57004 | 5.40916 | 2.18E-05 | 7.69E-05 | 9.64575 | 17.7607 | 17.35939 | 41.55869 | 29.4923 | 50.04871 | 170.5513 | 151.4677 | 109.1698 | 32.30701 | 19.0632 | 17.16585 | 70.91779 | 50.30929 | 48.59222 |
| Acaa1a | 113868 | 1.376074 | 5.386623 | 18.18848 | 1.79E-14 | 2.36E-13 | -0.49496 | 5.386623 | -6.91724 | 7.13E-07 | 3.10E-06 | 1956.603 | 1970.55 | 1613.555 | 5181.85 | 5252.438 | 4895.305 | 3786.239 | 4255.839 | 3352.087 | 2281.49 | 2267.758 | 2371.126 | 1215.614 | 1140.606 | 1223.714 |
| Eva1b | 230752 | 1.377334 | 3.592744 | 14.84641 | 1.00E-12 | 9.70E-12 | -0.30798 | 3.592744 | -3.66379 | 0.001418 | 0.003756 | 474.1257 | 521.2765 | 411.4176 | 1316.891 | 1307.492 | 1260.686 | 1192.438 | 1070.997 | 1015.854 | 1176.898 | 1193.356 | 1332.965 | 247.7951 | 229.5361 | 240.5315 |
| Arhgap20 | 244867 | 1.377462 | -1.30934 | 2.805923 | 0.010484 | 0.02502 | 0.978669 | -1.30934 | 2.560112 | 0.018101 | 0.036173 | 5.935846 | 10.65642 | 15.62345 | 19.48064 | 32.30109 | 33.81669 | 49.74413 | 52.27645 | 68.94934 | 11.53822 | 9.912862 | 4.478048 | 51.72827 | 73.10568 | 49.40209 |
| Reln | 19699 | 1.378792 | -0.42314 | 3.088128 | 0.005502 | 0.013934 | 1.115442 | -0.42314 | 3.309861 | 0.00328 | 0.008014 | 17.06556 | 15.09659 | 26.03909 | 46.75353 | 68.81537 | 45.9907 | 102.3308 | 87.12741 | 176.97 | 17.69193 | 16.01308 | 14.92683 | 75.92375 | 128.1315 | 68.83897 |
| Mtdh | 67154 | 1.378955 | 8.093358 | 48.4638 | 2.54E-23 | 2.52E-21 | -0.38016 | 8.093358 | -14.5487 | 1.49E-12 | 1.62E-11 | 12454.15 | 12546.16 | 12112.52 | 34654.76 | 35390.76 | 33076.78 | 27457.34 | 27959.86 | 26874.15 | 11169.76 | 11193.15 | 11804.13 | 8655.307 | 8559.654 | 8997.659 |
| Mgam2-ps | 638904 | 1.378977 | -1.17072 | 2.683086 | 0.013802 | 0.032061 | 1.644105 | -1.17072 | 4.387743 | 0.000249 | 0.000757 | 8.161788 | 9.768385 | 11.2836 | 22.07806 | 33.70549 | 27.05335 | 83.85439 | 85.78699 | 97.67823 | 9.230573 | 1.525056 | 11.94146 | 75.08942 | 103.7629 | 80.98703 |
| Rundc3b | 242819 | 1.380723 | -0.314 | 3.682845 | 0.001355 | 0.003817 | 1.620603 | -0.314 | 5.965621 | 5.98E-06 | 2.29E-05 | 11.12971 | 12.43249 | 14.75548 | 35.06515 | 35.10988 | 36.52203 | 108.0158 | 116.6167 | 119.5122 | 26.15329 | 30.50111 | 20.15121 | 171.8713 | 198.8789 | 178.9813 |
| Ms4a6d | 68774 | 1.38253 | 8.728605 | 47.75441 | 3.47E-23 | 3.27E-21 | -0.49135 | 8.728605 | -18.1951 | 1.77E-14 | 2.87E-13 | 15194.28 | 15226.25 | 15458.54 | 43167.79 | 43621.92 | 40942.55 | 31570.47 | 31191.61 | 31500.65 | 20016.5 | 18633.13 | 19677.29 | 25920.87 | 25725.34 | 27504 |
| Wnk3 | 279561 | 1.383839 | -1.41802 | 2.462555 | 0.022373 | 0.049245 | 1.512663 | -1.41802 | 3.648411 | 0.001471 | 0.003884 | 11.87169 | 8.88035 | 5.207818 | 18.18193 | 32.30109 | 22.99535 | 62.53548 | 64.34024 | 87.33582 | 14.61507 | 3.812639 | 2.985365 | 56.73423 | 84.89692 | 49.40209 |
| Xirp1 | 22437 | 1.384229 | -0.59913 | 4.092713 | 0.000506 | 0.001521 | 1.209922 | -0.59913 | 4.835108 | 8.49E-05 | 0.000277 | 26.71131 | 23.97694 | 14.75548 | 53.24708 | 68.81537 | 59.51738 | 136.441 | 180.9569 | 126.4071 | 13.84586 | 12.20045 | 7.463413 | 41.71635 | 43.23454 | 46.97248 |
| Lnpep | 240028 | 1.384586 | 8.883025 | 68.42919 | 1.69E-26 | 5.21E-24 | -0.81421 | 8.883025 | -40.8167 | 9.57E-22 | 1.06E-19 | 23480.72 | 23196.36 | 23172.18 | 66053.65 | 66547.27 | 62280.88 | 37558.24 | 38007.66 | 39249.41 | 21502.62 | 21096.1 | 21439.4 | 14458.89 | 14458.42 | 14567.95 |
| Hdac1 | 433759 | 1.390011 | 6.511637 | 29.69031 | 7.61E-19 | 2.43E-17 | -0.19081 | 6.511637 | -4.51146 | 0.000185 | 0.000574 | 4201.837 | 4187.973 | 3595.998 | 11559.81 | 11063.83 | 10886.27 | 10270.03 | 10314.55 | 9927.555 | 4036.837 | 3957.52 | 4047.409 | 2589.751 | 2689.189 | 2689.579 |
| Myh10 | 77579 | 1.390132 | 2.689626 | 10.04189 | 1.53E-09 | 9.26E-09 | -0.71771 | 2.689626 | -5.36686 | 2.40E-05 | 8.46E-05 | 229.2721 | 161.6224 | 216.1244 | 553.2501 | 528.0526 | 618.1692 | 295.6223 | 368.616 | 403.3536 | 564.6034 | 561.983 | 555.2779 | 350.4173 | 371.8171 | 416.2733 |
| Ap3b2 | 11775 | 1.390488 | -1.41992 | 3.025752 | 0.006354 | 0.01589 | 0.342373 | -1.41992 | 0.895242 | 0.380643 | 0.478918 | 10.38773 | 10.65642 | 11.2836 | 45.45482 | 19.66153 | 31.11136 | 39.7953 | 33.51054 | 48.26453 | 14.61507 | 10.67539 | 4.478048 | 41.71635 | 46.37887 | 38.0639 |
| Cecr2 | 330409 | 1.3905 | 0.517758 | 5.664089 | 1.20E-05 | 4.38E-05 | 0.210286 | 0.517758 | 1.028582 | 0.315193 | 0.410329 | 37.84102 | 46.17782 | 41.66254 | 135.0658 | 103.9252 | 112.2714 | 143.5473 | 111.255 | 167.7767 | 72.30616 | 62.52728 | 55.22925 | 103.4565 | 111.6237 | 99.61404 |
| Meg3 | 17263 | 1.390566 | -0.39122 | 3.938825 | 0.000733 | 0.002152 | 0.690961 | -0.39122 | 2.490339 | 0.021069 | 0.041267 | 19.2915 | 25.75301 | 19.09533 | 46.75353 | 54.77142 | 82.51273 | 88.11817 | 109.9146 | 102.2748 | 19.99958 | 19.82572 | 15.67317 | 69.24913 | 106.9072 | 68.83897 |
| Khnyn | 219094 | 1.395009 | 5.355938 | 18.19115 | 1.78E-14 | 2.35E-13 | -0.11529 | 5.355938 | -1.67832 | 0.107869 | 0.166572 | 1167.136 | 1169.542 | 1121.417 | 3227.292 | 3318.586 | 3172.006 | 3374.073 | 3254.544 | 2768.316 | 1664.58 | 1925.383 | 1824.058 | 4280.097 | 4026.315 | 4252.629 |
| Larp1 | 73158 | 1.39569 | 8.951667 | 51.31199 | 7.59E-24 | 8.83E-22 | -0.93328 | 8.951667 | -34.1217 | 4.14E-20 | 2.82E-18 | 28555.13 | 29293.61 | 27441.73 | 80591.4 | 80814.52 | 78132.79 | 45422.08 | 44158.85 | 41200.68 | 22024.92 | 21735.09 | 22426.06 | 10516.69 | 10147.54 | 10584.19 |
| Phc2 | 54383 | 1.395813 | 6.989105 | 34.73171 | 2.86E-20 | 1.29E-18 | 0.526632 | 6.989105 | 15.9297 | 2.51E-13 | 3.17E-12 | 4495.661 | 4747.435 | 4625.41 | 12949.43 | 13146.54 | 12819.23 | 20268.6 | 19413.33 | 18670.33 | 4542.981 | 4569.067 | 4484.018 | 4824.913 | 4648.107 | 4866.51 |
| Zfp821 | 75871 | 1.39597 | 3.55294 | 18.19756 | 1.77E-14 | 2.34E-13 | -3.28714 | 3.55294 | -24.8312 | 3.11E-17 | 9.60E-16 | 1004.642 | 1068.306 | 907.8962 | 2875.342 | 2772.276 | 2710.746 | 328.3113 | 250.6589 | 312.5703 | 699.9852 | 703.8132 | 710.5169 | 336.2337 | 227.964 | 332.8567 |
| Tcf4 | 21413 | 1.397676 | 7.466298 | 41.03309 | 8.56E-22 | 5.49E-20 | -0.76197 | 7.466298 | -22.9054 | 1.64E-16 | 4.31E-15 | 8834.765 | 8781.778 | 8462.704 | 24476.77 | 25089.52 | 23863.76 | 15525.85 | 14826.4 | 14651.73 | 8489.82 | 8510.573 | 7934.354 | 4837.427 | 4858.777 | 5252.009 |
| Cybb | 13058 | 1.397801 | 9.5472 | 93.45998 | 2.23E-29 | 3.25E-26 | -1.24293 | 9.5472 | -76.2419 | 1.70E-27 | 2.23E-24 | 70621.73 | 71047.24 | 71893.05 | 200088.2 | 204558.6 | 196111.1 | 90527.21 | 87253.41 | 86137.26 | 55193.44 | 53915.29 | 55170.29 | 2555.543 | 2460.439 | 2812.679 |
| Vcpip1 | 70675 | 1.397966 | 7.53931 | 45.34763 | 1.04E-22 | 8.26E-21 | -0.59293 | 7.53931 | -20.2261 | 2.09E-15 | 4.16E-14 | 8168.466 | 8316.448 | 8250.919 | 23557.29 | 23763.77 | 22275.73 | 16085.83 | 16248.59 | 15599.79 | 10755.93 | 11160.36 | 10807.77 | 5179.502 | 4968.042 | 5157.254 |
| Morc1 | 17450 | 1.398039 | -2.29945 | 2.65664 | 0.014636 | 0.033781 | 0.726658 | -2.29945 | 1.700591 | 0.103547 | 0.160795 | 5.935846 | 4.440175 | 7.811726 | 19.48064 | 12.63956 | 20.29002 | 22.74017 | 37.53181 | 31.0272 | 2.307643 | 1.525056 | 4.478048 | 25.02981 | 33.01547 | 21.05663 |
| Gm10555 | 100038695 | 1.398607 | -1.67422 | 2.974582 | 0.007146 | 0.017689 | -0.24326 | -1.67422 | -0.57648 | 0.570321 | 0.664217 | 11.12971 | 10.65642 | 17.35939 | 42.8574 | 47.74944 | 22.99535 | 38.37404 | 28.14886 | 29.87805 | 2.307643 | 6.100223 | 14.92683 | 17.52087 | 31.4433 | 20.24676 |
| Irf2 | 16363 | 1.399602 | 6.576992 | 32.24674 | 1.35E-19 | 5.19E-18 | -0.86281 | 6.576992 | -20.0139 | 2.59E-15 | 5.07E-14 | 4228.548 | 4298.089 | 4325.093 | 11923.45 | 12133.98 | 12136.13 | 7181.631 | 6999.682 | 6540.994 | 5055.277 | 5270.593 | 5636.369 | 3324.793 | 3019.343 | 3236.242 |
| Eqtn | 67753 | 1.399741 | -0.96808 | 3.427257 | 0.002487 | 0.006698 | 0.184923 | -0.96808 | 0.541309 | 0.5939 | 0.686812 | 17.80754 | 10.65642 | 15.62345 | 27.27289 | 51.96262 | 51.40137 | 51.16539 | 57.63814 | 41.3696 | 13.07665 | 16.77561 | 23.88292 | 54.23125 | 37.73197 | 51.02183 |
| Ccbe1 | 320924 | 1.39985 | -0.85163 | 2.777972 | 0.011165 | 0.026491 | 0.339853 | -0.85163 | 0.813722 | 0.42479 | 0.52369 | 14.83961 | 12.43249 | 10.41564 | 45.45482 | 43.53625 | 22.99535 | 32.689 | 41.55307 | 71.24765 | 36.15308 | 32.02617 | 55.22925 | 63.40885 | 48.73712 | 29.15533 |
| Sema3b | 20347 | 1.40011 | -0.47434 | 3.358383 | 0.002926 | 0.007779 | 1.46496 | -0.47434 | 4.872454 | 7.77E-05 | 0.000255 | 25.96933 | 11.54445 | 23.43518 | 53.24708 | 85.66811 | 37.8747 | 146.3899 | 160.8506 | 172.3733 | 17.69193 | 22.87584 | 8.209754 | 54.23125 | 68.38919 | 42.11325 |
| Parp3 | 235587 | 1.400349 | 5.61166 | 18.51345 | 1.25E-14 | 1.70E-13 | 0.857006 | 5.61166 | 14.05411 | 2.91E-12 | 3.01E-11 | 1284.369 | 1325.836 | 1446.037 | 4035.089 | 3662.663 | 3702.252 | 7603.746 | 7341.49 | 6612.241 | 2841.478 | 3040.199 | 3092.092 | 2064.125 | 1830 | 1928.301 |
| Frmd4a | 209630 | 1.404139 | 7.27209 | 40.06545 | 1.42E-21 | 8.52E-20 | -1.70301 | 7.27209 | -42.1972 | 4.74E-22 | 6.00E-20 | 10480.48 | 10445.96 | 10013.77 | 29188.49 | 29899.58 | 28415.49 | 9738.479 | 9529.058 | 8739.328 | 8982.886 | 8682.142 | 8135.866 | 2840.049 | 2811.818 | 2728.453 |
| Nsmaf | 18201 | 1.404287 | 6.040488 | 27.35739 | 4.18E-18 | 1.15E-16 | -1.20108 | 6.040488 | -22.3902 | 2.62E-16 | 6.57E-15 | 3239.488 | 2964.261 | 3216.695 | 9018.236 | 8776.066 | 8849.152 | 3847.353 | 4436.796 | 3795.661 | 3066.089 | 3161.44 | 2894.311 | 2798.332 | 2826.753 | 2688.769 |
| Dck | 13178 | 1.405199 | 7.045805 | 44.1711 | 1.81E-22 | 1.33E-20 | -2.82976 | 7.045805 | -62.3677 | 1.21E-25 | 5.40E-23 | 8928.254 | 8829.732 | 9135.38 | 25639.12 | 26213.04 | 24238.45 | 3872.936 | 3381.884 | 3850.82 | 15078.91 | 14944.02 | 14608.14 | 2467.105 | 2521.753 | 2391.547 |
| Armc8 | 74125 | 1.40528 | 5.338257 | 28.48688 | 1.80E-18 | 5.35E-17 | -1.27589 | 5.338257 | -24.4518 | 4.28E-17 | 1.28E-15 | 2004.832 | 1959.893 | 2123.922 | 5745.489 | 5864.755 | 5602.75 | 2474.415 | 2553.503 | 2362.664 | 2009.957 | 2138.128 | 2056.17 | 1390.823 | 1311.972 | 1486.112 |
| Lgals8 | 56048 | 1.405576 | 7.402301 | 44.40592 | 1.62E-22 | 1.21E-20 | 0.106637 | 7.402301 | 3.916032 | 0.000775 | 0.002168 | 6337.258 | 6351.226 | 6153.904 | 17563.74 | 18251.52 | 17523.81 | 20284.23 | 19673.37 | 19667.8 | 10053.63 | 9577.35 | 9681.539 | 4968.417 | 4882.359 | 4780.664 |
| Birc2 | 11797 | 1.406541 | 4.560167 | 22.75046 | 1.89E-16 | 3.69E-15 | -0.79611 | 4.560167 | -13.1087 | 1.11E-11 | 1.04E-10 | 951.2193 | 1052.321 | 1005.977 | 2849.368 | 2838.283 | 2804.08 | 1820.635 | 1601.804 | 1667.425 | 1412.278 | 1350.437 | 1400.136 | 884.3865 | 839.5362 | 783.1445 |
| Scn3a | 20269 | 1.406655 | 0.453942 | 3.118932 | 0.005123 | 0.013032 | 1.347869 | 0.453942 | 4.042909 | 0.000571 | 0.001636 | 9.64575 | 11.54445 | 17.35939 | 24.67547 | 61.79339 | 29.75869 | 88.11817 | 85.78699 | 109.1698 | 1259.204 | 1231.482 | 1129.214 | 64.24317 | 75.46393 | 66.40936 |
| St3gal3 | 20441 | 1.406693 | 3.549979 | 15.95972 | 2.42E-13 | 2.61E-12 | -1.69534 | 3.549979 | -16.4106 | 1.39E-13 | 1.88E-12 | 586.1648 | 642.0493 | 541.613 | 1745.465 | 1508.321 | 1747.647 | 604.0359 | 553.5942 | 466.5572 | 604.6026 | 591.7216 | 593.3413 | 500.5962 | 543.9692 | 556.3809 |
| Iqsec2 | 245666 | 1.408143 | 6.517118 | 26.61451 | 7.40E-18 | 1.91E-16 | 0.549642 | 6.517118 | 12.59809 | 2.36E-11 | 2.07E-10 | 3178.646 | 3063.721 | 2957.172 | 8719.533 | 8787.301 | 8592.145 | 13837.4 | 13707.15 | 12314.35 | 4223.757 | 4341.834 | 4174.287 | 3335.639 | 3215.864 | 3267.017 |
| Dnaja1 | 15502 | 1.408223 | 7.396852 | 25.53634 | 1.32E-15 | 2.14E-14 | -0.51352 | 7.396852 | -9.9338 | 9.75E-09 | 5.51E-08 | 8014.876 | 7942.585 | 6409.956 | 20970.26 | 21550.45 | 20735.04 | 15837.11 | 15479.19 | 14775.84 | 9079.038 | 9014.604 | 9264.334 | 4624.674 | 4625.31 | 4966.124 |
| Col6a4 | 68553 | 1.408963 | -1.04103 | 3.208194 | 0.004162 | 0.010771 | 0.906309 | -1.04103 | 2.687597 | 0.013664 | 0.028316 | 18.54952 | 7.10428 | 17.35939 | 35.06515 | 35.10988 | 47.34337 | 58.27169 | 65.68067 | 106.8715 | 9.999788 | 4.575167 | 8.956095 | 59.23721 | 66.81702 | 69.64884 |
| Frem2 | 242022 | 1.408995 | -0.79232 | 3.210273 | 0.004142 | 0.010726 | 1.227883 | -0.79232 | 3.75652 | 0.001136 | 0.003075 | 14.83961 | 9.768385 | 13.88751 | 46.75353 | 29.4923 | 35.16936 | 69.64178 | 99.19121 | 102.2748 | 6.153716 | 15.25056 | 16.41951 | 79.26106 | 96.68816 | 106.9029 |
| Sirpb1b | 668101 | 1.409137 | 2.116288 | 12.88342 | 1.55E-11 | 1.24E-10 | -3.50747 | 2.116288 | -16.7008 | 9.85E-14 | 1.36E-12 | 489.7073 | 516.8364 | 393.1902 | 1292.216 | 1402.991 | 1262.039 | 105.1733 | 97.85079 | 157.4343 | 189.996 | 250.1091 | 240.3219 | 65.0775 | 54.2397 | 102.8535 |
| Dusp27 | 240892 | 1.409949 | -1.55968 | 2.757424 | 0.011692 | 0.027604 | 0.981346 | -1.55968 | 2.457001 | 0.022642 | 0.043891 | 5.935846 | 11.54445 | 6.075787 | 24.67547 | 19.66153 | 20.29002 | 46.90161 | 33.51054 | 55.15947 | 11.53822 | 10.67539 | 4.478048 | 50.05962 | 42.44846 | 59.9304 |
| Fpr1 | 14293 | 1.410954 | 6.951475 | 52.7047 | 4.30E-24 | 5.44E-22 | -2.65461 | 6.951475 | -70.037 | 1.03E-26 | 9.44E-24 | 17882.48 | 16948.15 | 17037.38 | 49309.39 | 50170.62 | 48041.35 | 8627.053 | 8003.658 | 7764.844 | 4861.435 | 5082.248 | 5307.233 | 649.1063 | 560.4769 | 691.6292 |
| Dhx40 | 67487 | 1.411604 | 5.577159 | 27.8057 | 2.98E-18 | 8.50E-17 | 0.633408 | 5.577159 | 15.166 | 6.61E-13 | 7.63E-12 | 1268.045 | 1222.824 | 1253.348 | 3631.191 | 3508.179 | 3503.409 | 5898.233 | 5568.112 | 5684.873 | 2705.327 | 2659.697 | 2751.014 | 2805.841 | 2730.065 | 2752.749 |
| Dbnl | 13169 | 1.411918 | 7.587666 | 33.87081 | 4.84E-20 | 2.05E-18 | -0.59042 | 7.587666 | -14.928 | 9.01E-13 | 1.02E-11 | 7065.883 | 7265.014 | 7053.121 | 20142.98 | 20389.01 | 20199.39 | 14293.62 | 14678.96 | 13077.39 | 9169.036 | 9032.905 | 8807.573 | 10727.78 | 10774.05 | 11675.09 |
| Fer1l4 | 74562 | 1.412107 | -0.80675 | 3.002908 | 0.006696 | 0.016668 | 0.959399 | -0.80675 | 2.661451 | 0.014481 | 0.029772 | 14.83961 | 17.7607 | 10.41564 | 45.45482 | 37.91867 | 37.8747 | 61.11422 | 67.02109 | 124.1088 | 7.692145 | 16.77561 | 11.19512 | 90.94163 | 104.549 | 60.74027 |
| Resf1 | 67246 | 1.413083 | 8.460078 | 58.22294 | 5.21E-25 | 9.02E-23 | -0.93153 | 8.460078 | -38.272 | 3.71E-21 | 3.47E-19 | 13304.46 | 13802.73 | 13929.18 | 38648.29 | 39424.18 | 38468.52 | 20940.86 | 21328.79 | 21121.48 | 19458.82 | 19191.3 | 18928.71 | 18809.07 | 19004.33 | 19746.26 |
| Gm20069 | 100504113 | 1.414057 | -1.48519 | 3.178977 | 0.004456 | 0.011469 | 0.232804 | -1.48519 | 0.627621 | 0.53691 | 0.632164 | 11.12971 | 21.31284 | 12.15157 | 53.24708 | 32.30109 | 37.8747 | 49.74413 | 72.38277 | 35.62382 | 13.07665 | 9.150334 | 10.44878 | 18.35519 | 6.288661 | 17.00728 |
| Tnip1 | 57783 | 1.415638 | 6.97807 | 38.93707 | 2.58E-21 | 1.48E-19 | -1.23633 | 6.97807 | -32.3009 | 1.31E-19 | 7.71E-18 | 7004.298 | 6945.322 | 6956.776 | 19915.71 | 20537.88 | 19149.72 | 9172.818 | 9063.932 | 8141.767 | 6924.469 | 6906.215 | 7200.7 | 2889.274 | 2821.251 | 2845.884 |
| Tmtc2 | 278279 | 1.415983 | 2.241655 | 9.61602 | 3.28E-09 | 1.89E-08 | 0.417773 | 2.241655 | 3.553976 | 0.001842 | 0.004761 | 168.4296 | 149.1899 | 172.726 | 451.9508 | 539.2878 | 417.9743 | 591.2445 | 754.6574 | 610.2016 | 254.61 | 210.4577 | 229.1268 | 165.1967 | 155.6444 | 150.6359 |
| Orm2 | 18406 | 1.416567 | 0.735768 | 8.146685 | 5.45E-08 | 2.67E-07 | -1.4411 | 0.735768 | -7.45246 | 2.27E-07 | 1.06E-06 | 108.3292 | 114.5565 | 78.11726 | 279.2225 | 280.8791 | 290.8236 | 113.7009 | 120.638 | 94.23076 | 82.30595 | 66.33992 | 51.49755 | 50.89394 | 45.59279 | 46.16261 |
| Odad4 | 74407 | 1.417463 | -1.31357 | 2.807545 | 0.010446 | 0.02494 | 0.304037 | -1.31357 | 0.72016 | 0.47924 | 0.577684 | 11.87169 | 14.20856 | 6.943757 | 29.87031 | 37.91867 | 25.70069 | 19.89765 | 38.87223 | 70.09849 | 10.769 | 14.48803 | 11.94146 | 55.8999 | 50.30929 | 37.25403 |
| Gfra1 | 14585 | 1.417626 | -0.64911 | 2.92026 | 0.008091 | 0.019798 | 1.257642 | -0.64911 | 3.424049 | 0.002506 | 0.0063 | 7.419807 | 7.992315 | 6.943757 | 19.48064 | 15.44835 | 33.81669 | 51.16539 | 50.93603 | 62.0544 | 22.30722 | 14.48803 | 17.91219 | 453.8738 | 422.9124 | 481.0629 |
| Cbln3 | 56410 | 1.418006 | -1.03561 | 2.584344 | 0.017165 | 0.038895 | 0.808688 | -1.03561 | 1.875015 | 0.074541 | 0.121626 | 9.64575 | 10.65642 | 10.41564 | 29.87031 | 37.91867 | 22.99535 | 28.42522 | 61.6594 | 82.7392 | 7.692145 | 16.77561 | 22.39024 | 80.09538 | 69.17527 | 103.6634 |
| Cdh24 | 239096 | 1.418853 | 1.825732 | 7.250622 | 3.48E-07 | 1.53E-06 | -0.12021 | 1.825732 | -0.71277 | 0.483706 | 0.5824 | 109.0712 | 95.01974 | 141.479 | 272.7289 | 386.2087 | 328.6983 | 365.264 | 272.1056 | 303.3771 | 129.9972 | 185.2943 | 120.9073 | 270.3219 | 260.9794 | 238.9117 |
| Pfpl | 56093 | 1.41896 | -1.24019 | 3.186742 | 0.004376 | 0.011281 | 0.119597 | -1.24019 | 0.31418 | 0.756435 | 0.831835 | 10.38773 | 22.20087 | 12.15157 | 48.05224 | 35.10988 | 37.8747 | 41.21656 | 28.14886 | 73.54596 | 8.461359 | 12.20045 | 17.16585 | 35.04173 | 36.1598 | 23.48624 |
| Psme2 | 19188 | 1.419543 | 7.149206 | 26.65011 | 7.20E-18 | 1.87E-16 | 0.052464 | 7.149206 | 1.13813 | 0.267681 | 0.358029 | 5974.429 | 6213.581 | 5173.099 | 16348.15 | 16583.1 | 16560.71 | 18455.07 | 18059.5 | 16932.81 | 7136.772 | 6647.718 | 7094.72 | 3968.893 | 3468.983 | 3767.516 |
| Itgad | 381924 | 1.420051 | -1.13114 | 2.77662 | 0.011199 | 0.026555 | 0.783323 | -1.13114 | 1.94167 | 0.065493 | 0.108884 | 9.64575 | 13.32052 | 6.943757 | 25.97418 | 44.94065 | 18.93735 | 36.95278 | 40.21265 | 79.29174 | 25.38408 | 18.30067 | 11.94146 | 56.73423 | 59.74228 | 56.69092 |
| Tapbpl | 213233 | 1.42016 | 4.668237 | 16.76268 | 9.15E-14 | 1.07E-12 | 2.015599 | 4.668237 | 32.95315 | 8.60E-20 | 5.39E-18 | 637.3615 | 584.327 | 616.2584 | 1793.517 | 1775.156 | 1693.54 | 7733.08 | 7307.979 | 7106.378 | 979.9792 | 969.9354 | 1097.122 | 1021.216 | 1088.724 | 920.0126 |
| Elavl4 | 15572 | 1.420247 | -1.65087 | 2.789741 | 0.010873 | 0.025862 | -0.16183 | -1.65087 | -0.35508 | 0.726011 | 0.805145 | 9.64575 | 21.31284 | 6.943757 | 33.76644 | 23.87472 | 41.9327 | 15.63387 | 28.14886 | 54.01031 | 6.92293 | 10.67539 | 5.97073 | 29.20144 | 19.65207 | 34.01455 |
| Ythdf1 | 228994 | 1.42029 | 6.635937 | 44.17808 | 1.80E-22 | 1.33E-20 | -1.86541 | 6.635937 | -49.5721 | 1.58E-23 | 3.67E-21 | 5354.875 | 5290.024 | 5582.78 | 15054.64 | 15945.5 | 15390.65 | 4410.172 | 4451.541 | 4346.106 | 5673.726 | 5805.887 | 5823.701 | 3386.533 | 3509.073 | 3595.824 |
| A930007I1 | 77779 | 1.421243 | 1.094298 | 5.70886 | 1.08E-05 | 3.97E-05 | 1.273356 | 1.094298 | 7.068185 | 5.15E-07 | 2.28E-06 | 62.32638 | 58.61031 | 36.45472 | 168.8322 | 120.778 | 156.9095 | 410.7444 | 391.4031 | 328.6585 | 73.84459 | 67.10245 | 91.79997 | 157.6878 | 153.2861 | 146.5865 |
| Icosl | 50723 | 1.423074 | 5.149487 | 26.62285 | 7.35E-18 | 1.90E-16 | 0.061969 | 5.149487 | 1.324123 | 0.199469 | 0.279948 | 1489.897 | 1481.242 | 1445.169 | 4375.351 | 4270.766 | 4009.307 | 4590.673 | 4707.561 | 4429.995 | 1569.197 | 1557.082 | 1668.073 | 921.9312 | 918.9306 | 1001.81 |
| Gucy1b2 | 239134 | 1.424456 | -2.37099 | 2.45733 | 0.022626 | 0.049724 | 0.320027 | -2.37099 | 0.650081 | 0.522576 | 0.619028 | 5.193865 | 3.55214 | 11.2836 | 27.27289 | 23.87472 | 8.116006 | 18.47639 | 25.46801 | 25.28142 | 5.384501 | 4.575167 | 2.239024 | 16.68654 | 22.01031 | 14.57766 |
| Gm9883 | 791409 | 1.426251 | -1.53818 | 2.487302 | 0.021208 | 0.046944 | -0.28819 | -1.53818 | -0.55727 | 0.583139 | 0.676807 | 6.677827 | 27.52908 | 12.15157 | 32.46773 | 30.8967 | 50.04871 | 46.90161 | 32.17012 | 21.83396 | 3.076858 | 15.25056 | 23.88292 | 22.52683 | 12.57732 | 27.53559 |
| Tmco3 | 234076 | 1.426582 | 5.785675 | 27.65399 | 3.34E-18 | 9.46E-17 | 0.409999 | 5.785675 | 9.462619 | 4.34E-09 | 2.61E-08 | 2061.964 | 1906.611 | 2204.643 | 5997.439 | 5845.093 | 5855.699 | 8293.057 | 8376.295 | 7802.766 | 2687.635 | 2618.521 | 2718.921 | 1500.12 | 1461.328 | 1461.006 |
| Pla2g4d | 78390 | 1.426803 | -1.73122 | 2.542138 | 0.018826 | 0.04225 | 0.217906 | -1.73122 | 0.460164 | 0.650052 | 0.737481 | 8.161788 | 8.88035 | 10.41564 | 20.77935 | 33.70549 | 27.05335 | 28.42522 | 33.51054 | 34.47467 | 12.30743 | 8.387806 | 7.463413 | 9.177596 | 44.02063 | 29.15533 |
| Cept1 | 99712 | 1.427406 | 7.013611 | 38.46486 | 3.34E-21 | 1.87E-19 | -0.90819 | 7.013611 | -24.545 | 3.95E-17 | 1.19E-15 | 5435.009 | 5724.273 | 5840.567 | 16388.41 | 16511.48 | 15824.86 | 9246.723 | 8289.168 | 9395.496 | 7236.77 | 7147.174 | 7138.008 | 4599.644 | 4561.637 | 4779.044 |
| Phex | 18675 | 1.429448 | -1.39523 | 2.97706 | 0.007105 | 0.017595 | 0.149374 | -1.39523 | 0.370884 | 0.71438 | 0.794613 | 19.2915 | 13.32052 | 9.547666 | 37.66257 | 39.32307 | 43.28537 | 65.378 | 26.80843 | 55.15947 | 13.07665 | 2.287584 | 9.702436 | 25.02981 | 41.66238 | 30.77507 |
| Irf8 | 15900 | 1.42999 | 6.656179 | 32.35113 | 1.27E-19 | 4.87E-18 | -0.76937 | 6.656179 | -17.8714 | 2.54E-14 | 4.01E-13 | 5075.148 | 5269.6 | 4792.06 | 14581.91 | 14699.81 | 14239.53 | 9431.487 | 8952.677 | 8254.385 | 5096.815 | 4904.579 | 5081.091 | 2391.181 | 2351.959 | 2589.965 |
| Lmo4 | 16911 | 1.431654 | 7.06181 | 46.17334 | 7.08E-23 | 5.89E-21 | 0.093364 | 7.06181 | 3.508518 | 0.002052 | 0.005247 | 6312.03 | 6262.423 | 6133.941 | 17876.73 | 18632.11 | 17437.24 | 19522.44 | 20158.6 | 19982.67 | 4912.204 | 5053.272 | 5021.384 | 2910.132 | 2825.967 | 3015.957 |
| Ddhd1 | 114874 | 1.431688 | 5.364176 | 21.19227 | 8.07E-16 | 1.37E-14 | 0.559992 | 5.364176 | 10.06453 | 1.47E-09 | 9.50E-09 | 1004.642 | 888.035 | 1005.109 | 2746.77 | 2895.863 | 2717.509 | 4320.633 | 4040.031 | 4411.608 | 4658.363 | 4622.444 | 4283.252 | 1728.725 | 1767.9 | 1753.369 |

| | | | | | | | | | | | | | | | | | | | | | | | | | | |
|---|---|---|---|---|---|---|---|---|---|---|---|---|---|---|---|---|---|---|---|---|---|---|---|---|---|---|
| Qrich2 | 217341 | 1.431872 | -0.76366 | 3.436221 | 0.002435 | 0.006572 | 0.39303 | -0.76366 | 1.168683 | 0.25542 | 0.344185 | 26.71131 | 10.65642 | 18.22736 | 84.4161 | 49.15383 | 32.46403 | 62.53548 | 68.36151 | 80.44089 | 14.61507 | 10.67539 | 12.6878 | 80.09538 | 62.88661 | 51.02183 |
| Maml2 | 270118 | 1.432666 | 5.44894 | 28.07146 | 2.45E-18 | 7.09E-17 | 0.043697 | 5.44894 | 0.97741 | 0.339315 | 0.436291 | 1445.378 | 1339.157 | 1360.108 | 4009.115 | 4013.762 | 3943.026 | 4178.507 | 4314.818 | 4298.991 | 1588.428 | 1691.287 | 1671.804 | 3105.365 | 3042.926 | 3242.721 |
| Zc3h12a | 230738 | 1.432749 | 7.038816 | 34.82973 | 2.69E-20 | 1.23E-18 | -0.56665 | 7.038816 | -14.5964 | 1.40E-12 | 1.53E-11 | 5888.359 | 6195.82 | 5638.331 | 17123.48 | 17627.97 | 16309.11 | 11927.22 | 12428.39 | 11481.21 | 10516.7 | 10165.26 | 10802.54 | 2341.121 | 2294.575 | 2521.126 |
| Sh3d21 | 66938 | 1.434241 | 1.9654 | 8.703801 | 1.82E-08 | 9.44E-08 | 0.999172 | 1.9654 | 8.169889 | 5.21E-08 | 2.66E-07 | 115.749 | 134.9813 | 103.2884 | 337.6644 | 356.7164 | 321.9349 | 722.0005 | 723.8277 | 669.9577 | 209.2263 | 208.9326 | 212.7073 | 151.8475 | 115.5541 | 89.8956 |
| Adhfe1 | 76187 | 1.436071 | 1.472135 | 7.501669 | 2.05E-07 | 9.27E-07 | -0.26768 | 1.472135 | -1.58905 | 0.126754 | 0.190718 | 88.29571 | 101.236 | 101.5524 | 307.7941 | 249.9824 | 280.0022 | 291.3585 | 245.2972 | 202.2514 | 199.2265 | 214.2703 | 253.0097 | 97.61625 | 66.03094 | 95.56469 |
| Abcb11 | 27413 | 1.437054 | -1.34855 | 2.475615 | 0.021751 | 0.048048 | 0.762859 | -1.34855 | 1.669098 | 0.109702 | 0.168977 | 16.32358 | 3.55214 | 7.811726 | 15.58451 | 23.87472 | 36.52203 | 22.74017 | 53.61687 | 57.45778 | 13.84586 | 9.912862 | 8.956095 | 72.58644 | 69.96135 | 85.03638 |
| Rictor | 78757 | 1.438057 | 6.805648 | 36.89365 | 8.03E-21 | 4.22E-19 | -0.58832 | 6.805648 | -15.9269 | 2.52E-13 | 3.18E-12 | 4510.501 | 4737.667 | 4534.273 | 13313.07 | 13423.21 | 13097.88 | 9211.192 | 9474.101 | 8861.139 | 4958.356 | 5073.098 | 5228.867 | 4470.324 | 4684.266 | 4941.018 |
| Trpm4 | 68667 | 1.438545 | 3.699449 | 16.78474 | 8.91E-14 | 1.04E-12 | -1.78204 | 3.699449 | -17.4869 | 3.93E-14 | 5.99E-13 | 601.7464 | 703.3237 | 598.0311 | 1777.933 | 1806.052 | 1896.44 | 586.9807 | 532.1474 | 537.8048 | 947.6722 | 822.005 | 1032.19 | 506.4364 | 478.7243 | 441.3793 |
| Kat2b | 18519 | 1.439905 | 6.368177 | 33.55601 | 5.88E-20 | 2.42E-18 | -0.26026 | 6.368177 | -6.69153 | 1.17E-06 | 4.93E-06 | 3539.99 | 3726.195 | 3272.245 | 10079.28 | 10493.64 | 9929.934 | 8943.995 | 8663.146 | 8812.874 | 3369.159 | 3584.643 | 3377.941 | 2764.959 | 2710.413 | 2667.713 |
| Iqcg | 69707 | 1.440708 | 0.272202 | 6.233514 | 3.25E-06 | 1.27E-05 | -0.99014 | 0.272202 | -4.2487 | 0.000348 | 0.001031 | 55.64856 | 50.61799 | 49.47427 | 150.6503 | 140.4395 | 163.6728 | 73.90556 | 83.10615 | 79.29174 | 71.53694 | 73.9652 | 42.54145 | 50.89394 | 59.74228 | 42.11325 |
| Fam110c | 104943 | 1.441314 | 1.1469 | 3.262973 | 0.003662 | 0.009586 | 4.66069 | 1.1469 | 16.73811 | 9.42E-14 | 1.31E-12 | 14.83961 | 11.54445 | 30.37894 | 44.15611 | 68.81537 | 43.28537 | 1331.721 | 1282.784 | 1419.207 | 30.76858 | 58.71464 | 58.21462 | 638.2601 | 674.4589 | 771.8064 |
| Chsy3 | 78923 | 1.44141 | -0.28435 | 3.614703 | 0.001594 | 0.004436 | 1.811738 | -0.28435 | 6.502768 | 1.78E-06 | 7.32E-06 | 21.51744 | 13.32052 | 15.62345 | 44.15611 | 44.94065 | 59.51738 | 171.9726 | 166.2123 | 201.1022 | 9.999788 | 22.11331 | 14.92683 | 107.6282 | 107.6933 | 159.5444 |
| Il1rap | 16180 | 1.442394 | 5.199572 | 21.49309 | 6.05E-16 | 1.06E-14 | -0.87801 | 5.199572 | -13.2192 | 9.47E-12 | 8.93E-11 | 1026.159 | 1004.368 | 1183.043 | 3016.901 | 3093.883 | 3185.533 | 1679.93 | 1820.293 | 1745.567 | 2249.183 | 2421.026 | 2385.307 | 3558.404 | 3811.715 | 3906.814 |
| Adcy1 | 432530 | 1.444037 | -0.94841 | 3.448346 | 0.002366 | 0.006402 | 1.106832 | -0.94841 | 3.528463 | 0.001957 | 0.005026 | 15.5816 | 12.43249 | 13.01954 | 50.64966 | 37.91867 | 33.81669 | 69.64178 | 79.08488 | 127.5563 | 8.461359 | 12.96297 | 5.97073 | 69.24913 | 61.31444 | 52.64157 |
| Ms4a6b | 69774 | 1.444348 | 7.552509 | 44.83507 | 1.32E-22 | 1.01E-20 | -0.05701 | 7.552509 | -2.03248 | 0.054734 | 0.09366 | 8054.943 | 7794.283 | 7426.348 | 22624.81 | 22541.95 | 22535.44 | 21623.06 | 23060.62 | 22799.25 | 6968.314 | 6968.742 | 6945.452 | 6547.798 | 6302.024 | 6638.507 |
| Gm19474 | 100502960 | 1.44479 | -0.50734 | 3.559358 | 0.001818 | 0.005014 | 0.715506 | -0.50734 | 2.270951 | 0.033613 | 0.061723 | 20.77546 | 26.64105 | 7.811726 | 54.54578 | 46.34504 | 44.63804 | 102.3308 | 60.31898 | 91.93245 | 37.69151 | 44.22662 | 49.25852 | 40.88202 | 47.16496 | 42.11325 |
| Slc29a4 | 243328 | 1.445577 | -1.45902 | 2.665314 | 0.014357 | 0.033211 | 0.874969 | -1.45902 | 2.063799 | 0.051406 | 0.088715 | 14.09763 | 4.440175 | 5.207818 | 24.67547 | 16.85274 | 22.99535 | 39.7953 | 28.14886 | 58.60694 | 11.53822 | 9.912862 | 10.44878 | 57.56856 | 82.53868 | 61.55014 |
| Nlgn1 | 192167 | 1.445996 | -1.99657 | 2.573836 | 0.017565 | 0.039713 | 1.451842 | -1.99657 | 3.416373 | 0.002552 | 0.006406 | 6.677827 | 4.440175 | 4.339848 | 18.18193 | 11.23516 | 17.58468 | 29.84648 | 52.27645 | 55.15947 | 4.615287 | 2.287584 | 5.224389 | 37.54471 | 47.95104 | 51.02183 |
| Gvin3 | 434223 | 1.447548 | 5.737787 | 28.53509 | 1.74E-18 | 5.18E-17 | -0.57109 | 5.737787 | -11.898 | 6.92E-11 | 5.56E-10 | 2881.853 | 3067.273 | 2899.886 | 8581.87 | 8288.741 | 8868.09 | 5949.398 | 6187.387 | 5858.395 | 4478.367 | 4555.341 | 4248.921 | 499.7618 | 426.8429 | 556.3809 |
| Ppp1r1a | 58200 | 1.448052 | 0.704658 | 7.315842 | 3.03E-07 | 1.34E-06 | -0.54109 | 0.704658 | -2.95942 | 0.007398 | 0.016553 | 63.06836 | 75.48297 | 57.28599 | 179.2219 | 213.4681 | 177.1995 | 136.441 | 152.8081 | 119.5122 | 63.07559 | 82.35301 | 82.09754 | 65.0775 | 62.88661 | 77.74755 |
| Muc1 | 17829 | 1.449025 | -0.69926 | 3.37138 | 0.002838 | 0.007565 | 1.601857 | -0.69926 | 5.261392 | 3.08E-05 | 0.000107 | 21.51744 | 13.32052 | 9.547666 | 27.27289 | 63.19779 | 44.63804 | 153.4962 | 128.6805 | 125.258 | 32.30701 | 21.35078 | 23.88292 | 22.52683 | 32.22939 | 38.87377 |
| Rnf144a | 108089 | 1.449455 | 3.398855 | 17.41721 | 4.26E-14 | 5.25E-13 | -0.22532 | 3.398855 | -3.06082 | 0.00586 | 0.013442 | 451.8663 | 458.226 | 479.9872 | 1357.151 | 1259.743 | 1436.533 | 1196.702 | 1234.528 | 1168.691 | 373.8382 | 330.9371 | 350.034 | 354.5889 | 363.9563 | 414.6536 |
| Lrrc15 | 74488 | 1.449632 | -2.08799 | 2.45545 | 0.022718 | 0.049909 | 0.788111 | -2.08799 | 1.686745 | 0.106216 | 0.164377 | 7.419807 | 10.65642 | 6.943757 | 25.97418 | 15.44835 | 35.16936 | 21.31891 | 48.25518 | 73.54596 | 3.846072 | 1.525056 | 0.746341 | 26.69846 | 22.01031 | 38.0639 |
| Pcdhga6 | 93714 | 1.45039 | -0.65349 | 3.853965 | 0.000899 | 0.002604 | -0.90371 | -0.65349 | -2.4467 | 0.02315 | 0.04474 | 59.35846 | 22.20087 | 22.56721 | 120.78 | 68.81537 | 98.74475 | 39.7953 | 71.04235 | 49.41369 | 13.07665 | 28.21353 | 35.82438 | 24.19548 | 25.15464 | 14.57766 |
| Tor1aip1 | 208263 | 1.450657 | 8.653236 | 52.74294 | 4.24E-24 | 5.39E-22 | -0.48994 | 8.653236 | -19.4168 | 4.78E-15 | 8.75E-14 | 18976.16 | 19772.1 | 18054.64 | 55515.92 | 57326.01 | 52909.6 | 41486.6 | 40538.37 | 40475.56 | 20608.79 | 20361.02 | 20940.1 | 9266.869 | 8988.069 | 9252.768 |
| Gvin-ps1 | 77700 | 1.451545 | 0.302969 | 5.283954 | 2.92E-05 | 0.000102 | -0.35409 | 0.302969 | -1.44629 | 0.162622 | 0.235719 | 55.64856 | 42.62568 | 53.81412 | 118.1825 | 144.6527 | 193.4315 | 116.5434 | 148.7868 | 103.424 | 60.76794 | 90.74081 | 70.15608 | 30.8701 | 20.43815 | 50.21196 |
| Gm19955 | 100503907 | 1.451907 | 3.655998 | 12.42895 | 3.05E-11 | 2.34E-10 | 0.42987 | 3.655998 | 4.571397 | 0.00016 | 0.000501 | 351.6989 | 341.8935 | 374.0949 | 1046.76 | 1015.378 | 1055.081 | 1414.155 | 1621.91 | 1350.258 | 759.2147 | 918.0835 | 702.3071 | 582.3602 | 572.2681 | 556.3809 |
| Fbln1 | 14114 | 1.452076 | -1.67856 | 2.810464 | 0.010378 | 0.024799 | 0.753079 | -1.67856 | 1.835054 | 0.080474 | 0.129699 | 11.12971 | 5.32821 | 4.339848 | 25.97418 | 14.04395 | 20.29002 | 32.689 | 37.53181 | 34.47467 | 11.53822 | 14.48803 | 13.43414 | 40.04769 | 39.30413 | 25.10598 |
| Oas1a | 246730 | 1.452371 | 7.166838 | 27.49297 | 3.77E-18 | 1.06E-16 | -0.24345 | 7.166838 | -5.13894 | 4.12E-05 | 0.000141 | 5958.105 | 5940.954 | 4892.745 | 16276.72 | 16809.21 | 15911.43 | 14678.78 | 14817.02 | 13592.21 | 8669.816 | 8707.306 | 8768.017 | 3993.089 | 4078.983 | 4246.96 |
| BC034090 | 207792 | 1.452441 | -1.09244 | 3.177256 | 0.004473 | 0.011508 | 0.867805 | -1.09244 | 2.462854 | 0.022359 | 0.043415 | 9.64575 | 22.20087 | 11.2836 | 37.66257 | 56.17581 | 28.40602 | 75.32683 | 56.29771 | 94.23076 | 5.384501 | 13.7255 | 5.97073 | 64.24317 | 40.09021 | 55.07118 |
| Oxtr | 18430 | 1.453198 | -1.86994 | 2.745749 | 0.012002 | 0.028282 | 1.108072 | -1.86994 | 2.714215 | 0.012878 | 0.0269 | 5.193865 | 9.768385 | 5.207818 | 19.48064 | 26.68351 | 13.52668 | 31.26774 | 46.91476 | 54.01031 | 3.076858 | 3.812639 | 5.97073 | 35.04173 | 43.23454 | 53.45144 |
| P2ry2 | 18442 | 1.453349 | 4.568751 | 21.94811 | 3.94E-16 | 7.16E-15 | -1.43895 | 4.568751 | -20.0309 | 2.54E-15 | 4.99E-14 | 1159.716 | 1018.576 | 1105.793 | 3237.682 | 3171.125 | 3208.528 | 1226.548 | 1361.868 | 1120.427 | 1108.438 | 1188.781 | 1214.297 | 1103.815 | 1100.516 | 1204.277 |
| Stat5a | 20850 | 1.455337 | 6.084029 | 29.06954 | 1.18E-18 | 3.65E-17 | -0.6799 | 6.084029 | -14.1575 | 2.53E-12 | 2.64E-11 | 2769.814 | 2831.944 | 2915.51 | 8436.415 | 8384.24 | 8093.011 | 5588.398 | 5665.963 | 4973.545 | 3379.928 | 3579.306 | 3294.35 | 2531.348 | 2506.031 | 2523.556 |
| Smim3 | 106878 | 1.455621 | 3.99018 | 20.83518 | 1.14E-15 | 1.88E-14 | -0.50038 | 3.99018 | -7.70531 | 1.34E-07 | 6.45E-07 | 770.176 | 843.6332 | 693.5077 | 2331.183 | 2199.283 | 2202.143 | 1691.3 | 1650.059 | 1606.52 | 286.917 | 301.1985 | 297.7902 | 886.0552 | 904.7811 | 974.2739 |
| Pclo | 26875 | 1.45573 | -0.22331 | 2.542565 | 0.018808 | 0.042225 | 1.419156 | -0.22331 | 3.430811 | 0.002467 | 0.006207 | 16.32358 | 7.992315 | 41.66254 | 64.93546 | 56.17581 | 39.22736 | 129.3347 | 108.5742 | 210.2955 | 35.38386 | 28.97606 | 7.463413 | 152.6818 | 127.3454 | 102.0437 |
| Map2 | 17756 | 1.456517 | 0.875748 | 4.80055 | 9.23E-05 | 0.000303 | -1.16168 | 0.875748 | -3.6579 | 0.001438 | 0.003806 | 44.51884 | 31.96926 | 57.28599 | 125.9748 | 139.0351 | 120.3874 | 48.32287 | 48.25518 | 82.7392 | 639.9864 | 634.4232 | 518.7072 | 92.61029 | 58.17011 | 96.37456 |
| L3mbtl1 | 241764 | 1.456772 | -1.62119 | 2.545064 | 0.018706 | 0.042031 | 0.892001 | -1.62119 | 1.988488 | 0.059732 | 0.100835 | 5.935846 | 6.216245 | 10.41564 | 46.75353 | 15.44835 | 13.52668 | 28.42522 | 49.5956 | 51.712 | 14.61507 | 7.625279 | 12.6878 | 27.53279 | 41.66238 | 29.9652 |
| Ankrd17 | 81702 | 1.457004 | 8.554215 | 59.75132 | 3.01E-25 | 5.69E-23 | -0.95106 | 8.554215 | -39.1028 | 2.36E-21 | 2.35E-19 | 17661.37 | 17915.22 | 17677.07 | 52600.32 | 53007.49 | 50480.21 | 28635.56 | 28458.49 | 26912.07 | 19544.2 | 18990.76 | 19404.87 | 11405.25 | 11578.21 | 11436.99 |
| A230098N | 97209 | 1.457309 | -1.24186 | 3.151677 | 0.004748 | 0.012155 | 1.530921 | -1.24186 | 4.624302 | 0.000141 | 0.000446 | 16.32358 | 6.216245 | 16.49142 | 27.27289 | 39.32307 | 44.63804 | 95.22448 | 113.9358 | 119.5122 | 9.999788 | 4.575167 | 14.92683 | 15.85221 | 29.08506 | 22.67637 |
| Wfdc21 | 66107 | 1.45806 | 2.294284 | 11.49271 | 1.32E-10 | 9.24E-10 | -0.85728 | 2.294284 | -6.92176 | 7.07E-07 | 3.07E-06 | 270.081 | 275.2908 | 272.5425 | 776.6281 | 815.9536 | 808.8953 | 534.3941 | 473.1689 | 393.0112 | 165.3811 | 173.8563 | 226.1414 | 154.3505 | 99.04641 | 96.37456 |
| Pcdh15 | 11994 | 1.460102 | -0.47928 | 3.718463 | 0.001244 | 0.003526 | 1.002033 | -0.47928 | 3.397411 | 0.002669 | 0.006668 | 14.09763 | 15.98463 | 24.30315 | 36.36386 | 68.81537 | 56.81205 | 108.0158 | 103.2125 | 113.7664 | 15.38429 | 16.77561 | 8.956095 | 107.6282 | 124.2011 | 79.36729 |
| Cd200r2 | 271375 | 1.460182 | -1.76771 | 2.610654 | 0.0162 | 0.036983 | -0.74774 | -1.76771 | -1.38223 | 0.181196 | 0.258202 | 8.161788 | 15.09659 | 6.943757 | 25.97418 | 37.91867 | 22.99535 | 17.05513 | 18.7659 | 16.08818 | 18.46115 | 4.575167 | 4.478048 | 50.89394 | 24.36856 | 58.31066 |
| Naaa | 67111 | 1.460677 | 5.062967 | 20.65244 | 1.37E-15 | 2.21E-14 | -0.3262 | 5.062967 | -5.05225 | 5.06E-05 | 0.00017 | 1064.742 | 1301.859 | 1130.096 | 3324.695 | 3689.346 | 3208.528 | 3000.282 | 2700.95 | 2761.421 | 1863.037 | 1908.607 | 1996.463 | 1485.102 | 1428.312 | 1549.282 |
| Cd14 | 12475 | 1.460741 | 8.194366 | 48.31982 | 2.71E-23 | 2.67E-21 | -1.31386 | 8.194366 | -40.5738 | 1.08E-21 | 1.17E-19 | 21236.97 | 20736.5 | 22877.94 | 63967.92 | 63946.33 | 62425.62 | 27218.57 | 27516.18 | 25061.93 | 10195.17 | 9682.579 | 10645.81 | 4422.767 | 4401.277 | 4508.548 |
| Urgcp | 72046 | 1.461184 | 5.430681 | 31.73708 | 1.89E-19 | 7.00E-18 | -1.33366 | 5.430681 | -27.3364 | 4.25E-18 | 1.60E-16 | 1894.277 | 1900.395 | 1880.89 | 5575.358 | 5693.418 | 5424.198 | 2198.691 | 2367.185 | 2301.759 | 2279.952 | 2358.499 | 2242.755 | 2150.06 | 2037.526 | 1997.14 |
| Sipa1l1 | 217692 | 1.461466 | 7.054736 | 31.69732 | 1.94E-19 | 7.15E-18 | -0.50048 | 7.054736 | -11.7179 | 9.20E-11 | 7.23E-10 | 8163.272 | 8444.325 | 7760.516 | 23744.3 | 24614.84 | 23288.88 | 17930.63 | 18775.29 | 16160.58 | 3688.383 | 3446.626 | 3421.228 | 2558.046 | 2579.923 | 2797.292 |
| Ankrd33b | 67434 | 1.461809 | 6.392405 | 33.56219 | 5.86E-20 | 2.41E-18 | -0.6731 | 6.392405 | -16.1459 | 1.93E-13 | 2.51E-12 | 3245.424 | 3331.907 | 3371.194 | 9706.552 | 10205.74 | 9349.639 | 6661.45 | 6381.748 | 6049.155 | 3529.925 | 3564.055 | 3763.799 | 4218.357 | 4248.777 | 4408.124 |
| Srrm4 | 68955 | 1.463244 | -1.19012 | 3.120284 | 0.005107 | 0.012996 | 0.900213 | -1.19012 | 2.518688 | 0.019812 | 0.039167 | 15.5816 | 7.10428 | 12.15157 | 23.37676 | 58.9846 | 28.40602 | 65.378 | 53.61687 | 79.29174 | 11.53822 | 6.862751 | 4.478048 | 54.23125 | 69.96135 | 63.97975 |
| Hc | 15139 | 1.463428 | -0.8364 | 3.055429 | 0.005934 | 0.014931 | 0.722669 | -0.8364 | 1.930884 | 0.066888 | 0.110922 | 13.35565 | 15.09659 | 7.811726 | 31.16902 | 36.51428 | 37.8747 | 48.32287 | 50.93603 | 83.88836 | 9.230573 | 16.01308 | 5.97073 | 194.3982 | 167.4356 | 132.0089 |
| Tmprss6 | 71753 | 1.464216 | -0.8552 | 2.810756 | 0.010371 | 0.02479 | 2.080754 | -0.8552 | 5.693256 | 1.12E-05 | 4.13E-05 | 4.451884 | 22.20087 | 8.679696 | 23.37676 | 30.8967 | 29.75869 | 110.8583 | 99.19121 | 164.3292 | 17.69193 | 18.30067 | 11.94146 | 71.75211 | 68.38919 | 69.64884 |
| Itpkb | 320404 | 1.464352 | 7.912453 | 43.03034 | 3.14E-22 | 2.17E-20 | -0.26057 | 7.912453 | -8.61536 | 2.16E-08 | 1.17E-07 | 10530.19 | 10793.18 | 10465.11 | 31711.88 | 30920.57 | 30965.27 | 27886.56 | 27919.64 | 25633.06 | 9625.95 | 9726.043 | 9557.646 | 7093.447 | 7075.53 | 7339.044 |
| Crb2 | 241324 | 1.465163 | 0.834565 | 5.723804 | 1.04E-05 | 3.85E-05 | 0.097474 | 0.834565 | 0.459025 | 0.650857 | 0.738067 | 68.26223 | 55.9462 | 39.9266 | 174.027 | 151.6747 | 154.2041 | 181.9214 | 131.3613 | 225.2345 | 74.6138 | 73.20267 | 53.73657 | 148.5102 | 141.4949 | 117.4312 |
| Adcy4 | 104110 | 1.465179 | 0.651751 | 2.978651 | 0.007079 | 0.017539 | 4.477756 | 0.651751 | 14.31129 | 2.05E-12 | 2.17E-11 | 6.677827 | 23.08891 | 19.9633 | 35.06515 | 44.94065 | 47.34337 | 1085.843 | 1001.295 | 897.4905 | 25.38408 | 22.87584 | 29.85365 | 434.6843 | 378.8918 | 502.9294 |
| Trpa1 | 277328 | 1.465259 | -2.12583 | 2.6872 | 0.013676 | 0.031807 | 0.801736 | -2.12583 | 1.84381 | 0.07914 | 0.128011 | 5.193865 | 4.440175 | 9.547666 | 15.58451 | 19.66153 | 20.29002 | 21.31891 | 37.53181 | 44.81707 | 7.692145 | 3.812639 | 2.239024 | 22.52683 | 30.65722 | 24.29611 |
| Agrp | 11604 | 1.46532 | 0.138337 | 6.313155 | 2.71E-06 | 1.07E-05 | 0.052617 | 0.138337 | 0.270863 | 0.789099 | 0.859845 | 39.32498 | 41.73764 | 40.79457 | 119.4812 | 129.2044 | 112.2714 | 132.1773 | 125.9996 | 129.8546 | 53.0758 | 70.91509 | 44.03413 | 31.70442 | 35.37372 | 35.63429 |
| Plag1 | 56711 | 1.465336 | 2.204795 | 6.838728 | 8.47E-07 | 3.54E-06 | 1.8222 | 2.204795 | 12.2929 | 3.76E-11 | 3.17E-10 | 81.61788 | 114.5565 | 148.4228 | 301.3005 | 283.6878 | 399.037 | 1218.021 | 1156.784 | 1200.868 | 156.1505 | 162.4184 | 96.27802 | 279.4995 | 275.915 | 257.5387 |
| Atp8b5 | 320571 | 1.466457 | -1.2451 | 3.134242 | 0.004944 | 0.012618 | 1.130894 | -1.2451 | 3.186224 | 0.004381 | 0.010386 | 5.193865 | 12.43249 | 9.547666 | 29.87031 | 16.85274 | 31.11136 | 54.00791 | 57.63814 | 62.0544 | 12.30743 | 17.53814 | 15.67317 | 56.73423 | 48.73712 | 42.11325 |
| Ncoa7 | 211329 | 1.466605 | 6.149748 | 43.66315 | 2.31E-22 | 1.65E-20 | -1.17648 | 6.149748 | -33.9095 | 4.72E-20 | 3.17E-18 | 3965.887 | 4046.775 | 4024.775 | 12092.28 | 11843.27 | 11566.66 | 5464.748 | 5365.708 | 5466.533 | 4465.29 | 4405.123 | 4440.73 | 1138.022 | 1223.931 | 1248.01 |
| Insl6 | 27356 | 1.467437 | 2.838053 | 9.526286 | 3.86E-09 | 2.21E-08 | 1.254476 | 2.838053 | 11.28785 | 1.84E-10 | 1.38E-09 | 126.8787 | 143.8617 | 138.8751 | 355.8463 | 425.5318 | 430.1483 | 1067.367 | 932.9335 | 992.8704 | 462.2979 | 458.2792 | 567.2194 | 533.1349 | 507.8094 | 535.3242 |
| Vasp | 22323 | 1.468926 | 8.364885 | 37.776 | 1.99E-16 | 3.87E-15 | -1.50189 | 8.364885 | -34.7371 | 7.01E-16 | 1.57E-14 | 22507.99 | 23065.82 | 21776.49 | 68361.45 | 67774.71 | 62932.87 | 25584.12 | 25131.57 | 22640.66 | 16024.28 | 15358.07 | 16790.44 | 5126.105 | 4949.962 | 5212.325 |
| Gm19337 | 100502721 | 1.469364 | 2.370261 | 12.57019 | 2.47E-11 | 1.92E-10 | -0.80986 | 2.370261 | -7.14759 | 4.34E-07 | 1.94E-06 | 231.498 | 245.9857 | 249.1073 | 681.8223 | 738.7119 | 726.3826 | 385.1617 | 449.0413 | 433.2317 | 177.6885 | 197.4947 | 145.5365 | 233.6115 | 247.616 | 190.3195 |
| Cyp39a1 | 56050 | 1.469524 | -0.37439 | 4.285776 | 0.000318 | 0.000981 | -0.17572 | -0.37439 | -0.58743 | 0.56308 | 0.657531 | 27.45329 | 22.20087 | 34.71878 | 71.429 | 106.734 | 74.39673 | 58.27169 | 68.36151 | 102.2748 | 32.30701 | 32.7887 | 56.72194 | 33.37308 | 40.09021 | 19.43689 |
| Igfbp5 | 16011 | 1.469801 | -1.81252 | 2.535466 | 0.019101 | 0.042789 | 1.041144 | -1.81252 | 2.331872 | 0.029573 | 0.055246 | 11.12971 | 5.32821 | 3.471878 | 14.2858 | 19.66153 | 22.99535 | 48.32287 | 32.17012 | 42.51876 | 11.53822 | 3.812639 | 4.478048 | 33.37308 | 45.59279 | 59.12053 |
| Myh13 | 544791 | 1.472584 | -1.39712 | 2.980483 | 0.00705 | 0.017476 | 0.556798 | -1.39712 | 1.414796 | 0.171553 | 0.246396 | 13.35565 | 16.87266 | 6.075787 | 45.45482 | 26.68351 | 31.11136 | 56.85043 | 32.17012 | 72.3968 | 10.769 | 3.050111 | 4.478048 | 51.72827 | 58.17011 | 44.54286 |
| Galnt17 | 212996 | 1.473627 | -2.18001 | 2.656679 | 0.014635 | 0.033781 | 0.794729 | -2.18001 | 1.804804 | 0.085235 | 0.136178 | 8.161788 | 5.32821 | 4.339848 | 15.58451 | 16.85274 | 21.64268 | 27.00396 | 41.55307 | 31.0272 | 2.307643 | 7.625279 | 2.239024 | 20.85817 | 38.51805 | 24.29611 |
| Sag | 20215 | 1.474438 | 0.38105 | 4.611169 | 0.000145 | 0.000466 | 0.41082 | 0.38105 | 1.627699 | 0.118262 | 0.179876 | 51.93865 | 47.95389 | 26.90706 | 107.7929 | 153.0791 | 112.2714 | 233.0868 | 138.0634 | 155.136 | 30.76858 | 69.39003 | 52.24389 | 63.40885 | 72.3196 | 66.40936 |
| Bmerb1 | 67254 | 1.47471 | -1.69749 | 4.052832 | 0.000557 | 0.001664 | -1.84842 | -1.69749 | -4.22542 | 0.000368 | 0.001087 | 20.77546 | 30.19319 | 26.90706 | 98.7019 | 71.62416 | 60.87005 | 31.26774 | 8.04253 | 35.62382 | 7.692145 | 6.100223 | 4.478048 | 5.840288 | 4.716496 | 14.57766 |
| Col2a1 | 12824 | 1.475093 | -0.85333 | 2.888493 | 0.008697 | 0.021147 | 1.279533 | -0.85333 | 3.419371 | 0.002534 | 0.006365 | 16.32358 | 11.54445 | 8.679696 | 29.87031 | 58.9846 | 27.05335 | 69.64178 | 73.7232 | 139.0478 | 21.538 | 11.43792 | 8.956095 | 70.08346 | 87.25517 | 61.55014 |
| Ptpru | 19273 | 1.475668 | 0.42435 | 5.685694 | 1.14E-05 | 4.18E-05 | -0.05957 | 0.42435 | -0.2704 | 0.789449 | 0.86012 | 45.26083 | 72.81887 | 48.6063 | 127.2735 | 212.0637 | 154.2041 | 152.0749 | 207.7654 | 132.1529 | 16.92272 | 25.92595 | 15.67317 | 128.4863 | 121.8428 | 108.5226 |
| Fn3k | 63828 | 1.476016 | -1.62659 | 3.136824 | 0.004915 | 0.012549 | 0.795212 | -1.62659 | 2.154341 | 0.042783 | 0.07583 | 6.677827 | 9.768385 | 7.811726 | 28.5716 | 21.06593 | 22.99535 | 35.53152 | 48.25518 | 48.26453 | 4.615287 | 8.387806 | 6.717071 | 35.87606 | 58.17011 | 38.87377 |
| Noxred1 | 71275 | 1.476093 | 0.665763 | 4.963093 | 6.26E-05 | 0.000209 | 1.401167 | 0.665763 | 6.609219 | 1.40E-06 | 5.85E-06 | 29.67923 | 46.17782 | 40.79457 | 97.40319 | 146.0571 | 101.4501 | 287.0947 | 382.0202 | 275.7973 | 69.99852 | 94.55345 | 60.45364 | 68.41481 | 75.46393 | 81.7969 |
| Mir744 | 791070 | 1.476162 | -2.14206 | 3.250512 | 0.00377 | 0.009844 | 0.027227 | -2.14206 | 0.069629 | 0.945137 | 0.954647 | 6.677827 | 14.20856 | 7.811726 | 37.66257 | 22.47032 | 22.99535 | 31.26774 | 24.12759 | 31.0272 | 7.692145 | 4.575167 | 4.478048 | 6.674615 | 11.79124 | 12.14805 |
| Trpc5 | 22067 | 1.476399 | -1.40238 | 2.707394 | 0.013075 | 0.030544 | -0.50116 | -1.40238 | -0.99205 | 0.332288 | 0.428736 | 14.09763 | 15.98463 | 6.943757 | 53.24708 | 15.44835 | 51.40137 | 24.16143 | 17.42548 | 37.92213 | 11.53822 | 18.30067 | 15.67317 | 26.69846 | 55.02578 | 27.53559 |
| Ak4 | 11639 | 1.480572 | 3.570723 | 18.82002 | 8.99E-15 | 1.25E-13 | 0.123226 | 3.570723 | 1.84622 | 0.078776 | 0.127501 | 638.8454 | 651.8177 | 621.4662 | 2012.999 | 1872.059 | 1813.927 | 2130.47 | 2280.057 | 2048.944 | 252.3023 | 270.6974 | 309.7316 | 217.7593 | 260.1933 | 276.1658 |
| Mybpc3 | 17868 | 1.482415 | 2.224317 | 13.78506 | 4.23E-12 | 3.71E-11 | -1.37976 | 2.224317 | -11.817 | 7.87E-11 | 6.27E-10 | 231.498 | 230.8891 | 237.8237 | 729.8745 | 664.279 | 695.2712 | 279.9884 | 313.6587 | 248.2176 | 247.6871 | 216.5579 | 256.7414 | 146.0072 | 161.1469 | 157.9247 |
| Ccnl1 | 56706 | 1.484132 | 6.934834 | 32.81052 | 9.42E-20 | 3.72E-18 | -0.05329 | 6.934834 | -1.35277 | 0.190286 | 0.268883 | 4831.037 | 4863.768 | 4826.779 | 14244.24 | 14437.18 | 14704.85 | 14873.49 | 15165.53 | 13549.69 | 5479.115 | 5458.174 | 5190.803 | 3691.062 | 3880.104 | 3923.821 |
| Enah | 13800 | 1.484243 | -0.14426 | 3.767613 | 0.001106 | 0.003163 | 2.304884 | -0.14426 | 8.679137 | 1.91E-08 | 1.04E-07 | 14.09763 | 14.20856 | 16.49142 | 38.96127 | 56.17581 | 41.9327 | 245.8781 | 217.1483 | 237.8752 | 21.538 | 36.60134 | 27.61463 | 78.42673 | 124.2011 | 100.4239 |
| Mug-ps1 | 17835 | 1.484912 | -1.98304 | 2.641729 | 0.015127 | 0.034814 | 0.880807 | -1.98304 | 2.004579 | 0.057858 | 0.09812 | 6.677827 | 17.7607 | 2.603909 | 19.48064 | 25.27911 | 18.93735 | 39.7953 | 30.8297 | 52.86116 | 4.615287 | 5.337695 | 7.463413 | 14.18356 | 23.58248 | 22.67637 |
| Coch | 12810 | 1.485013 | 0.593631 | 5.714223 | 1.07E-05 | 3.93E-05 | 2.422812 | 0.593631 | 14.10226 | 2.73E-12 | 2.83E-11 | 41.55092 | 31.08122 | 43.39848 | 129.8709 | 92.69009 | 127.1508 | 615.4059 | 647.4237 | 672.256 | 36.15308 | 25.16342 | 46.27316 | 53.39692 | 65.24486 | 56.69092 |
| Mill2 | 243864 | 1.486325 | -0.1394 | 4.544704 | 0.00017 | 0.000543 | 0.847516 | -0.1394 | 3.435213 | 0.002441 | 0.006148 | 22.25942 | 32.85729 | 17.35939 | 63.63675 | 71.62416 | 75.74939 | 149.2324 | 144.7655 | 109.1698 | 69.99852 | 80.82795 | 54.48291 | 27.53279 | 32.22939 | 31.58494 |
| Pde6b | 18587 | 1.486719 | 1.458639 | 6.804021 | 9.13E-07 | 3.80E-06 | 1.18604 | 1.458639 | 7.52464 | 1.95E-07 | 9.18E-07 | 64.55232 | 57.72227 | 58.15396 | 180.5206 | 147.4615 | 221.8375 | 375.2129 | 482.5518 | 430.9333 | 106.9208 | 102.9413 | 88.06827 | 246.1264 | 262.5516 | 259.9684 |
| Cnnm1 | 83674 | 1.486981 | -1.22073 | 2.860631 | 0.009265 | 0.022391 | 0.591342 | -1.22073 | 1.445924 | 0.162724 | 0.235828 | 17.80754 | 4.440175 | 9.547666 | 16.88322 | 51.96262 | 28.40602 | 46.90161 | 44.23392 | 45.96622 | 17.69193 | 9.912862 | 13.43414 | 62.57452 | 51.09537 | 68.83897 |
| Tmem72 | 319776 | 1.487889 | -1.91851 | 2.476723 | 0.021699 | 0.047946 | 0.8843 | -1.91851 | 1.872203 | 0.074945 | 0.122188 | 7.419807 | 1.77607 | 11.2836 | 25.97418 | 14.04395 | 13.52668 | 28.42522 | 46.91476 | 28.72889 | 8.461359 | 7.625279 | 5.97073 | 35.87606 | 43.23454 | 29.15533 |
| Atoh8 | 71093 | 1.488299 | -1.36224 | 3.402223 | 0.002639 | 0.007073 | -0.53141 | -1.36224 | -1.30686 | 0.205167 | 0.2866 | 11.12971 | 21.31284 | 15.62345 | 50.64966 | 40.72746 | 47.34337 | 35.53152 | 22.78717 | 42.51876 | 7.692145 | 1.525056 | 8.956095 | 58.40288 | 65.24486 | 37.25403 |
| Ap3m2 | 64933 | 1.488737 | 4.619838 | 27.44295 | 3.92E-18 | 1.09E-16 | -1.35057 | 4.619838 | -23.4274 | 1.03E-16 | 2.87E-15 | 1176.039 | 1158.886 | 1270.708 | 3625.996 | 3652.832 | 3514.231 | 1371.517 | 1499.932 | 1512.289 | 1303.818 | 1322.986 | 1245.644 | 815.9717 | 856.044 | 851.9835 |
| Gm14964 | 100008567 | 1.48961 | -2.21827 | 3.43735 | 0.002429 | 0.006556 | -2.29043 | -2.21827 | -4.18955 | 0.000401 | 0.001178 | 8.903769 | 32.85729 | 10.41564 | 42.8574 | 30.8967 | 59.51738 | 8.527565 | 9.382952 | 8.044089 | 7.692145 | 3.812639 | 2.985365 | 10.84625 | 13.3634 | 10.52831 |
| Slc15a3 | 65221 | 1.490977 | 9.9192 | 35.9157 | 8.87E-13 | 8.67E-12 | -0.49095 | 9.9192 | -13.2409 | 4.09E-08 | 2.12E-07 | 43694.5 | 42971.12 | 41695.52 | 129361.8 | 131122.8 | 125193.5 | 100751.8 | 96187.32 | 89620.35 | 45583.65 | 45052.43 | 46298.53 | 27078.08 | 26140.39 | 28167.29 |
| Ccng2 | 12452 | 1.491783 | 5.02377 | 20.18136 | 2.18E-15 | 3.39E-14 | -0.33034 | 5.02377 | -4.92252 | 6.89E-05 | 0.000228 | 979.4146 | 1044.329 | 1108.397 | 3140.279 | 3436.555 | 2828.428 | 2650.651 | 2511.95 | 2576.407 | 2231.491 | 2524.73 | 2538.307 | 1342.432 | 1402.371 | 1339.525 |
| Rnf24 | 51902 | 1.491897 | 5.843877 | 32.55882 | 1.11E-19 | 4.31E-18 | 0.264548 | 5.843877 | 6.837488 | 8.49E-07 | 3.64E-06 | 2434.439 | 2593.95 | 2456.354 | 7441.604 | 7836.526 | 7188.076 | 9293.625 | 9462.037 | 9236.913 | 1380.74 | 1459.478 | 1349.385 | 1938.976 | 1888.17 | 2058.69 |
| Icam5 | 15898 | 1.493379 | 0.085379 | 5.670405 | 1.18E-05 | 4.32E-05 | 0.040025 | 0.085379 | 0.182028 | 0.857279 | 0.918721 | 32.64715 | 38.1855 | 27.77503 | 112.9877 | 96.90327 | 86.57074 | 103.752 | 109.9146 | 102.2748 | 30.76858 | 27.451 | 22.39024 | 125.9834 | 112.4098 | 92.32521 |
| Slc15a2 | 57738 | 1.493465 | 1.505645 | 7.274208 | 3.31E-07 | 1.46E-06 | 2.414547 | 1.505645 | 17.78798 | 2.79E-14 | 4.39E-13 | 53.42261 | 52.39406 | 74.64539 | 175.3257 | 178.3582 | 181.2575 | 956.5086 | 1075.018 | 946.9042 | 126.9204 | 121.2419 | 120.1609 | 100.9536 | 119.4846 | 102.8535 |
| Bst1 | 12182 | 1.495042 | 5.034189 | 33.10866 | 7.79E-20 | 3.14E-18 | -0.86786 | 5.034189 | -19.6 | 3.95E-15 | 7.38E-14 | 1626.422 | 1628.656 | 1610.084 | 4864.965 | 4932.236 | 4850.667 | 2860.998 | 2670.12 | 2799.343 | 1099.977 | 1063.726 | 1080.702 | 1193.087 | 1179.124 | 1263.398 |
| Tlnrd1 | 80889 | 1.495982 | 6.529132 | 34.93819 | 2.52E-20 | 1.17E-18 | -2.13904 | 6.529132 | -41.162 | 8.01E-22 | 9.08E-20 | 5075.89 | 5432.11 | 5018.6 | 15488.41 | 15991.85 | 15224.28 | 3945.42 | 3856.393 | 3292.331 | 4709.131 | 4634.644 | 4845.994 | 3671.038 | 3554.666 | 3731.072 |
| Cemip | 80982 | 1.496788 | -0.82662 | 3.130832 | 0.004983 | 0.01271 | 2.477949 | -0.82662 | 7.669848 | 1.44E-07 | 6.91E-07 | 7.419807 | 12.43249 | 14.75548 | 62.33804 | 16.85274 | 33.81669 | 186.1852 | 197.042 | 204.5497 | 9.999788 | 10.67539 | 6.717071 | 65.0775 | 63.67269 | 54.26131 |
| Slfn10-ps | 237887 | 1.497634 | 6.109189 | 34.06513 | 4.29E-20 | 1.85E-18 | -0.07793 | 6.109189 | -2.01786 | 0.056352 | 0.095972 | 2740.135 | 2801.75 | 2738.444 | 8151.997 | 8358.961 | 8456.879 | 8563.097 | 8240.913 | 7816.556 | 3688.383 | 3579.306 | 3549.599 | 1747.915 | 1787.552 | 1746.08 |
| Plod3 | 26433 | 1.497908 | 7.626411 | 31.04397 | 3.00E-19 | 1.06E-17 | 1.035731 | 7.626411 | 28.39605 | 1.93E-18 | 8.03E-17 | 6267.511 | 6363.659 | 6484.601 | 19722.2 | 19351.16 | 18535.61 | 42271.14 | 42240.71 | 38636.91 | 5646.803 | 5497.826 | 5446.052 | 8077.119 | 8512.489 | 8271.205 |
| Rcbtb1 | 71330 | 1.497912 | 5.401665 | 27.9769 | 2.62E-18 | 7.58E-17 | -0.80211 | 5.401665 | -15.4312 | 4.70E-13 | 5.61E-12 | 1611.582 | 1560.277 | 1735.071 | 5076.654 | 4960.324 | 4746.511 | 2851.049 | 3038.736 | 2909.662 | 2414.564 | 2317.322 | 2500.243 | 1731.228 | 1920.4 | 1782.524 |
| Garem2 | 242915 | 1.49887 | 0.013071 | 4.552072 | 0.000167 | 0.000534 | 1.743066 | 0.013071 | 7.655343 | 1.49E-07 | 7.10E-07 | 25.96933 | 31.08122 | 26.03909 | 70.13029 | 95.49888 | 87.9234 | 240.1931 | 327.0629 | 310.272 | 22.30722 | 19.0632 | 6.717071 | 80.92971 | 82.53868 | 73.69819 |
| Atp10b | 319767 | 1.498949 | -1.58547 | 2.619433 | 0.01589 | 0.036363 | 1.685465 | -1.58547 | 4.073221 | 0.000531 | 0.001527 | 12.61367 | 6.216245 | 2.603909 | 27.27289 | 16.85274 | 14.87935 | 59.69296 | 67.02109 | 68.94934 | 11.53822 | 7.625279 | 5.97073 | 49.22529 | 33.01547 | 53.45144 |
| Fabp4 | 11770 | 1.500351 | 2.012127 | 12.75109 | 1.88E-11 | 1.50E-10 | -2.11547 | 2.012127 | -13.8793 | 3.71E-12 | 3.76E-11 | 233.7239 | 272.6267 | 262.1268 | 759.7449 | 783.6526 | 764.2573 | 171.9726 | 199.7228 | 180.4174 | 896.9041 | 863.1815 | 1026.219 | 30.8701 | 14.14949 | 30.77507 |
| Dgkh | 380921 | 1.500844 | 6.38524 | 41.66911 | 6.19E-22 | 4.10E-20 | -2.13569 | 6.38524 | -48.6604 | 2.33E-23 | 5.04E-21 | 6759.445 | 6946.21 | 6348.33 | 20137.78 | 20269.64 | 20169.63 | 4765.488 | 5037.305 | 4542.612 | 2856.862 | 2976.909 | 3071.194 | 1575.209 | 1738.815 | 1574.388 |
| Cebpd | 12609 | 1.501205 | 5.898181 | 30.08956 | 5.76E-19 | 1.91E-17 | -3.53108 | 5.898181 | -43.9243 | 2.03E-22 | 3.18E-20 | 6125.793 | 7000.38 | 5859.663 | 19720.9 | 19526.71 | 17901.2 | 1854.745 | 1768.016 | 1546.763 | 3838.38 | 4098.587 | 4149.657 | 703.3376 | 723.9821 | 651.1357 |
| Nub1 | 53312 | 1.501229 | 7.52877 | 32.92177 | 8.77E-20 | 3.48E-18 | 0.691704 | 7.52877 | 19.23544 | 5.78E-15 | 1.04E-13 | 5348.939 | 5574.196 | 5693.881 | 17281.92 | 17168.73 | 15723.41 | 29385.99 | 28118.03 | 26858.06 | 8384.438 | 8419.07 | 8828.471 | 7414.663 | 7320.787 | 7395.735 |
| Tra2a | 101214 | 1.502285 | 6.455211 | 33.81672 | 5.00E-20 | 2.11E-18 | -0.41392 | 6.455211 | -10.1712 | 1.22E-09 | 7.97E-09 | 3628.286 | 3779.477 | 3939.714 | 11528.64 | 11544.13 | 11182.5 | 9117.388 | 9109.506 | 8526.734 | 3525.31 | 3596.081 | 3246.584 | 2907.629 | 2989.472 | 3063.739 |
| Adar | 56417 | 1.503111 | 8.494477 | 44.91252 | 1.27E-22 | 9.83E-21 | -0.08901 | 8.494477 | -3.11161 | 0.005211 | 0.012121 | 13245.1 | 12898.71 | 12907.58 | 39749.59 | 39372.22 | 39152.97 | 39970.12 | 39364.17 | 36575.32 | 17506.55 | 17302.52 | 16853.88 | 13490.23 | 13582.72 | 13873.08 |
| Ms4a6c | 73656 | 1.503407 | 8.937493 | 36.45369 | 2.38E-16 | 4.55E-15 | -0.06218 | 8.937493 | -1.78974 | 0.093207 | 0.146925 | 19701.81 | 20460.33 | 17285.61 | 58399.05 | 58851.18 | 56395.42 | 58486.31 | 57904.88 | 56471.8 | 15795.05 | 15199.47 | 15442.55 | 21951.14 | 21399.53 | 22457.7 |
| Enam | 13801 | 1.503841 | -1.02831 | 4.300362 | 0.000307 | 0.000949 | -0.19105 | -1.02831 | -0.62944 | 0.53574 | 0.631125 | 20.77546 | 15.98463 | 25.17112 | 66.23417 | 84.26372 | 43.28537 | 48.32287 | 62.99982 | 58.60694 | 5.384501 | 12.20045 | 12.6878 | 30.03577 | 24.36856 | 34.01455 |
| Plac8 | 231507 | 1.504163 | 3.776151 | 21.24702 | 7.65E-16 | 1.31E-14 | -0.3381 | 3.776151 | -5.28029 | 2.95E-05 | 0.000103 | 899.2807 | 880.9307 | 959.9744 | 2687.029 | 2605.153 | 3017.802 | 2328.025 | 2289.44 | 2202.931 | 658.4476 | 744.2272 | 683.6486 | 104.2909 | 78.60826 | 94.75482 |
| E230029C | 319711 | 1.506956 | -0.74679 | 3.52424 | 0.001977 | 0.005415 | 0.492426 | -0.74679 | 1.458184 | 0.159349 | 0.23176 | 19.2915 | 9.768385 | 22.56721 | 50.64966 | 33.70549 | 74.39673 | 65.378 | 57.63814 | 101.1257 | 46.92208 | 51.85189 | 33.58536 | 17.52087 | 18.86598 | 26.72572 |
| B3gat2 | 280645 | 1.507059 | 1.378901 | 7.50367 | 2.04E-07 | 9.24E-07 | -0.02755 | 1.378901 | -0.16338 | 0.871759 | 0.918721 | 85.32779 | 113.6685 | 97.2126 | 367.5347 | 276.6659 | 251.5962 | 247.2994 | 301.5949 | 352.7908 | 124.6127 | 83.87806 | 82.84388 | 110.9655 | 102.9768 | 113.3818 |
| Tasor2 | 105203 | 1.507797 | 6.901675 | 40.7053 | 1.01E-21 | 6.32E-20 | -0.26376 | 6.901675 | -7.98342 | 7.59E-08 | 3.78E-07 | 5454.3 | 5401.917 | 5661.766 | 16613.09 | 17028.29 | 16509.31 | 14474.12 | 14861.26 | 14069.11 | 4474.52 | 4159.589 | 4125.774 | 3254.709 | 3457.977 | 3178.741 |
| Cfap69 | 207686 | 1.508178 | -1.60429 | 2.549832 | 0.018512 | 0.041642 | 2.100546 | -1.60429 | 5.045446 | 5.14E-05 | 0.000173 | 7.419807 | 4.440175 | 6.075787 | 12.98709 | 11.23516 | 43.28537 | 66.79926 | 87.12741 | 101.1257 | 10.769 | 5.337695 | 5.224389 | 43.385 | 34.58764 | 42.92312 |
| Slamf1 | 27218 | 1.509045 | -0.36488 | 4.444718 | 0.000217 | 0.000682 | 1.159054 | -0.36488 | 4.665964 | 0.000127 | 0.000406 | 17.06556 | 23.08891 | 20.83127 | 79.22126 | 68.81537 | 41.9327 | 117.9647 | 163.5315 | 149.3902 | 44.61444 | 47.27673 | 42.54145 | 30.03577 | 22.7964 | 25.10598 |
| Mir99b | 387230 | 1.5097 | -0.84891 | 5.086789 | 4.66E-05 | 0.000159 | -0.05356 | -0.84891 | -0.2137 | 0.83281 | 0.89817 | 26.71131 | 28.41712 | 32.98284 | 76.62384 | 119.3736 | 78.45473 | 88.11817 | 101.8721 | 80.44089 | 23.07643 | 12.20045 | 8.209754 | 11.68058 | 13.3634 | 10.52831 |
| Lztfl1 | 93730 | 1.511086 | 4.37819 | 19.49802 | 4.39E-15 | 6.44E-14 | -0.90074 | 4.37819 | -11.831 | 7.69E-11 | 6.14E-10 | 930.4439 | 976.8385 | 935.6712 | 2772.744 | 3150.059 | 2745.916 | 1657.19 | 1431.57 | 1715.689 | 1068.439 | 947.0596 | 1097.868 | 629.0825 | 602.1393 | 743.4609 |
| Gm5546 | 433634 | 1.511444 | -2.83547 | 2.667511 | 0.014288 | 0.033061 | -0.36475 | -2.83547 | -0.6906 | 0.497258 | 0.595338 | 3.709904 | 5.32821 | 4.339848 | 12.98709 | 14.04395 | 14.87935 | 8.527565 | 8.04253 | 18.38649 | 3.076858 | 6.100223 | 1.492683 | 17.52087 | 12.57732 | 12.95792 |
| Ctsc | 13032 | 1.51151 | 9.327689 | 78.09871 | 1.02E-27 | 4.31E-25 | -1.18152 | 9.327689 | -58.7508 | 4.30E-25 | 1.49E-22 | 51204.83 | 50601.12 | 48956.96 | 154591.8 | 154983.4 | 149764.7 | 71055.94 | 71515.52 | 68104.71 | 28754.77 | 27394.58 | 29069.99 | 5440.646 | 5276.973 | 5637.507 |
| Smg7 | 226517 | 1.511528 | 7.415658 | 46.4084 | 6.36E-23 | 5.45E-21 | -0.93389 | 7.415658 | -29.1027 | 1.15E-18 | 5.12E-17 | 8112.817 | 7718.8 | 7789.159 | 23936.51 | 24432.26 | 23644.63 | 13727.96 | 13162.94 | 12378.7 | 7734.451 | 7875.388 | 7815.686 | 5665.08 | 5782.424 | 5765.466 |
| Plxdc2 | 67448 | 1.512615 | 2.963075 | 7.625278 | 1.58E-07 | 7.28E-07 | -0.03941 | 2.963075 | -0.23637 | 0.8154 | 0.883252 | 60.10044 | 101.236 | 81.58914 | 249.3522 | 251.3868 | 215.0742 | 238.7718 | 246.6376 | 236.7261 | 1784.578 | 1743.901 | 1612.843 | 2845.055 | 2716.702 | 2845.074 |
| Tspan10 | 208634 | 1.513331 | -2.22165 | 2.472881 | 0.02188 | 0.048305 | -0.19672 | -2.22165 | -0.35365 | 0.72707 | 0.806047 | 6.677827 | 4.440175 | 2.603909 | 10.38967 | 12.63956 | 20.29002 | 18.47639 | 5.361687 | 18.38649 | 8.461359 | 4.575167 | 7.463413 | 53.39692 | 62.88661 | 49.40209 |
| Ntrk2 | 18212 | 1.514276 | -0.79342 | 3.251648 | 0.00376 | 0.009821 | 0.951821 | -0.79342 | 2.758437 | 0.011665 | 0.024703 | 31.90517 | 13.32052 | 7.811726 | 29.87031 | 50.55823 | 74.39673 | 88.11817 | 92.4891 | 109.1698 | 5.384501 | 6.100223 | 10.44878 | 109.2968 | 88.04125 | 75.31793 |
| Parp4 | 328417 | 1.515129 | 6.958778 | 39.86303 | 1.57E-21 | 9.45E-20 | -0.54363 | 6.958778 | -15.3855 | 4.99E-13 | 5.92E-12 | 4551.31 | 4667.512 | 4720.019 | 14516.97 | 14010.25 | 13966.29 | 10282.82 | 10323.93 | 9723.005 | 6646.782 | 6564.602 | 7120.842 | 5061.861 | 5018.351 | 5055.21 |
| Fndc3c1 | 333564 | 1.515487 | -1.09415 | 3.551421 | 0.001853 | 0.005101 | 0.023564 | -1.09415 | 0.065734 | 0.948201 | 0.957044 | 20.03348 | 11.54445 | 13.01954 | 32.46773 | 40.72746 | 71.69139 | 58.27169 | 29.48928 | 62.0544 | 15.38429 | 10.67539 | 15.67317 | 39.21337 | 40.09021 | 30.77507 |
| Snph | 241727 | 1.51565 | -1.54441 | 2.561436 | 0.018048 | 0.040683 | 1.421983 | -1.54441 | 3.241557 | 0.00385 | 0.009251 | 6.677827 | 2.664105 | 12.15157 | 18.18193 | 21.06593 | 18.93735 | 32.689 | 54.95729 | 83.88836 | 6.92293 | 6.100223 | 9.702436 | 68.41481 | 69.96135 | 58.31066 |
| Fsd1 | 240121 | 1.515689 | -2.10836 | 2.461528 | 0.022423 | 0.04933 | 1.507049 | -2.10836 | 3.280721 | 0.003513 | 0.008528 | 5.935846 | 3.55214 | 4.339848 | 14.2858 | 11.23516 | 18.93735 | 52.58665 | 61.6594 | 28.72889 | 8.461359 | 5.337695 | 2.239024 | 25.86413 | 29.08506 | 31.58494 |
| Hmcn2 | 665700 | 1.515701 | 2.237995 | 8.17146 | 5.19E-08 | 2.55E-07 | 0.289291 | 2.237995 | 1.957912 | 0.063441 | 0.106042 | 94.23155 | 108.3403 | 98.08057 | 240.2612 | 383.3999 | 312.4662 | 380.8979 | 347.1692 | 429.7842 | 303.8397 | 268.4098 | 251.517 | 554.8274 | 540.8248 | 545.0427 |
| Sstr4 | 20608 | 1.515891 | -1.07717 | 3.024992 | 0.006365 | 0.015912 | 2.060755 | -1.07717 | 6.028411 | 5.18E-06 | 2.00E-05 | 14.83961 | 5.32821 | 19.9633 | 46.75353 | 25.27911 | 41.9327 | 184.7639 | 128.6805 | 178.1191 | 3.846072 | 6.100223 | 10.44878 | 40.88202 | 46.37887 | 25.91585 |
| Ly75 | 17076 | 1.516148 | 2.105022 | 8.669128 | 1.94E-08 | 1.00E-07 | 1.002021 | 2.105022 | 7.8358 | 1.03E-07 | 5.02E-07 | 98.68344 | 129.6531 | 101.5524 | 389.6127 | 337.0549 | 275.9442 | 730.5281 | 629.9982 | 712.4765 | 288.4554 | 317.2116 | 353.7658 | 128.4863 | 146.9974 | 145.7766 |
| Ripor3 | 69553 | 1.517578 | 0.05499 | 5.606987 | 1.37E-05 | 4.98E-05 | -0.10806 | 0.05499 | -0.46873 | 0.644015 | 0.73199 | 32.64715 | 37.29747 | 24.30315 | 97.40319 | 88.4769 | 100.0974 | 73.90556 | 100.5316 | 101.1257 | 26.15329 | 32.7887 | 43.28779 | 95.9476 | 99.04641 | 102.0437 |
| Ogfr | 72075 | 1.518584 | 6.970899 | 34.87701 | 2.62E-20 | 1.20E-18 | -0.32918 | 6.970899 | -8.3995 | 3.30E-08 | 1.73E-07 | 4874.072 | 5036.934 | 4598.503 | 15479.31 | 14754.58 | 14132.67 | 12617.95 | 12282.28 | 11829.41 | 6184.484 | 6528.763 | 6661.842 | 4097.379 | 4148.158 | 4356.292 |
| Kcna2 | 16490 | 1.518953 | -0.83156 | 4.524042 | 0.000179 | 0.000569 | -0.72052 | -0.83156 | -2.25327 | 0.034877 | 0.063736 | 23.0014 | 34.63336 | 18.22736 | 89.61093 | 64.60218 | 71.69139 | 29.84648 | 50.93603 | 63.20356 | 15.38429 | 25.16342 | 11.94146 | 27.53279 | 32.22939 | 25.91585 |
| Mir706 | 735290 | 1.519603 | -0.71413 | 5.137922 | 4.13E-05 | 0.000142 | -2.99851 | -0.71413 | -6.52421 | 1.69E-06 | 6.99E-06 | 29.67923 | 45.28978 | 42.53051 | 155.8451 | 77.24174 | 129.8561 | 9.948826 | 17.42548 | 17.23733 | 66.15244 | 69.39003 | 62.69267 | 20.02385 | 15.72165 | 8.098703 |
| Gm14057 | 100043766 | 1.520636 | 1.707191 | 9.176942 | 7.38E-09 | 4.05E-08 | -0.1684 | 1.707191 | -1.19039 | 0.246965 | 0.334643 | 102.3933 | 135.8694 | 121.5157 | 300.0018 | 407.2746 | 393.6263 | 316.9412 | 351.1905 | 335.5534 | 139.2278 | 112.8541 | 97.7707 | 168.534 | 168.2217 | 165.2135 |
| Stxbp3 | 20912 | 1.522329 | 6.641349 | 37.47927 | 5.77E-21 | 3.10E-19 | -0.58639 | 6.641349 | -15.4244 | 4.75E-13 | 5.66E-12 | 3562.992 | 3586.773 | 3303.492 | 10788.38 | 10545.6 | 10726.66 | 7673.387 | 7357.575 | 7177.626 | 8029.83 | 7930.29 | 7890.32 | 3525.866 | 3360.503 | 3478.393 |
| Trim34a | 94094 | 1.523446 | 5.899845 | 21.72148 | 4.87E-16 | 8.63E-15 | -0.27681 | 5.899845 | -4.38778 | 0.000249 | 0.000757 | 2291.237 | 2556.653 | 1906.929 | 6697.443 | 7305.664 | 6618.603 | 5999.142 | 5747.728 | 5892.87 | 3567.617 | 3563.293 | 3254.794 | 1873.064 | 1929.833 | 1762.278 |
| Synj1 | 104015 | 1.524108 | 8.38798 | 59.92591 | 2.83E-25 | 5.43E-23 | -0.68259 | 8.38798 | -28.5946 | 1.67E-18 | 7.04E-17 | 14487.92 | 14474.08 | 14635.7 | 44135.33 | 45748.18 | 44055.04 | 29569.33 | 29203.77 | 27989.98 | 18105 | 18697.18 | 18163.71 | 9163.413 | 9440.066 | 9501.398 |
| Gm10419 | 100503802 | 1.524178 | 0.419882 | 3.601959 | 0.001643 | 0.00456 | 2.000067 | 0.419882 | 6.978919 | 6.24E-07 | 2.73E-06 | 20.77546 | 11.54445 | 20.83127 | 62.33804 | 37.91867 | 64.92805 | 281.4096 | 230.5525 | 185.0141 | 36.15308 | 61.00223 | 32.83902 | 348.7487 | 364.7423 | 353.1034 |
| Apoa2 | 11807 | 1.524469 | 0.310602 | 7.043813 | 5.42E-07 | 2.33E-06 | -1.32052 | 0.310602 | -5.713 | 1.07E-05 | 3.95E-05 | 69.00421 | 62.16245 | 104.1564 | 246.7547 | 259.8131 | 201.5475 | 73.90556 | 108.5742 | 110.3189 | 57.69108 | 43.46409 | 34.3317 | 29.20144 | 20.43815 | 38.0639 |
| Cenpj | 219103 | 1.525209 | 4.445976 | 22.74329 | 1.90E-16 | 3.71E-15 | -1.11039 | 4.445976 | -16.3477 | 1.51E-13 | 2.01E-12 | 1007.61 | 963.518 | 1029.412 | 3114.305 | 3050.347 | 3059.734 | 1588.97 | 1439.613 | 1422.655 | 1255.358 | 1374.838 | 1165.039 | 701.6689 | 632.7965 | 622.7902 |
| Shank3 | 58234 | 1.525638 | -0.41324 | 3.794677 | 0.001037 | 0.002976 | 1.160135 | -0.41324 | 3.932369 | 0.000745 | 0.00209 | 11.87169 | 21.31284 | 18.22736 | 50.64966 | 60.389 | 43.28537 | 98.067 | 112.5954 | 145.9428 | 15.38429 | 10.67539 | 20.15121 | 87.60433 | 132.0619 | 128.7694 |
| Clcn1 | 12723 | 1.525943 | 0.480054 | 5.513206 | 1.70E-05 | 6.12E-05 | 2.187673 | 0.480054 | 11.89546 | 6.95E-11 | 5.58E-10 | 37.84102 | 36.40943 | 20.83127 | 100.0006 | 102.5209 | 85.21807 | 427.7995 | 450.3817 | 481.4962 | 61.53716 | 51.08937 | 59.7073 | 48.39096 | 62.10053 | 44.54286 |
| Has1 | 15116 | 1.526224 | 1.833735 | 11.03475 | 2.79E-10 | 1.87E-09 | 0.784645 | 1.833735 | 7.573339 | 1.76E-07 | 8.33E-07 | 198.8508 | 194.4797 | 162.3103 | 603.8998 | 556.1405 | 549.1831 | 1007.674 | 1129.976 | 941.1584 | 74.6138 | 64.81487 | 53.73657 | 80.92971 | 60.52836 | 51.02183 |
| Pcdhga7 | 93715 | 1.526626 | -0.5596 | 4.071201 | 0.000533 | 0.001598 | -1.05613 | -0.5596 | -2.74867 | 0.011923 | 0.025186 | 15.5816 | 40.84961 | 44.26645 | 67.53288 | 108.1384 | 105.5081 | 27.00396 | 40.21265 | 75.84427 | 22.30722 | 10.67539 | 20.15121 | 49.22529 | 55.81187 | 45.35273 |
| H3f3b | 15081 | 1.526643 | 8.603254 | 26.56317 | 2.19E-11 | 1.72E-10 | -0.90648 | 8.603254 | -16.1965 | 4.61E-09 | 2.75E-08 | 22450.11 | 22683.97 | 19044.12 | 66890.02 | 65885.8 | 64400.51 | 38486.32 | 37316 | 34042.59 | 18601.91 | 18174.09 | 18627.19 | 7324.556 | 6952.115 | 7245.909 |
| Aknaos | 66783 | 1.527415 | 1.646189 | 8.375341 | 3.46E-08 | 1.73E-07 | -0.73639 | 1.646189 | -4.31 | 0.0003 | 0.0009 | 155.816 | 102.124 | 166.6502 | 459.743 | 349.6944 | 492.3711 | 265.7758 | 331.0842 | 225.2345 | 126.1512 | 137.255 | 134.3414 | 118.4744 | 103.7629 | 97.18443 |
| Tasl | 71398 | 1.527882 | 3.342653 | 22.80576 | 1.80E-16 | 3.53E-15 | -3.59675 | 3.342653 | -29.4713 | 8.88E-19 | 4.10E-17 | 963.091 | 977.7265 | 1054.583 | 3023.395 | 3202.021 | 2988.043 | 291.3585 | 286.8503 | 224.0853 | 936.134 | 986.711 | 956.8095 | 100.9536 | 82.53868 | 136.0582 |
| Etohd2 | 13996 | 1.528303 | 2.212076 | 12.23437 | 4.11E-11 | 3.09E-10 | -0.29452 | 2.212076 | -2.70949 | 0.013014 | 0.027146 | 178.0754 | 192.7036 | 175.3299 | 574.0295 | 564.5669 | 541.0671 | 510.2326 | 432.9562 | 481.4962 | 162.3042 | 143.3552 | 135.8341 | 256.1384 | 237.397 | 195.1787 |
| Dusp15 | 252864 | 1.528603 | -1.49889 | 2.481318 | 0.021484 | 0.0475 | 0.826291 | -1.49889 | 1.709901 | 0.101784 | 0.158376 | 2.967923 | 7.992315 | 18.22736 | 31.16902 | 14.04395 | 28.40602 | 22.74017 | 48.25518 | 67.80018 | 7.692145 | 7.625279 | 5.97073 | 76.75808 | 69.17527 | 60.74027 |
| Ankib1 | 70797 | 1.529172 | 7.20873 | 47.02733 | 4.81E-23 | 4.30E-21 | -0.2637 | 7.20873 | -9.12313 | 8.17E-09 | 4.69E-08 | 5357.843 | 5680.76 | 5617.499 | 17175.43 | 17323.22 | 16743.32 | 14778.27 | 15089.13 | 14471.32 | 7534.456 | 7654.255 | 7532.076 | 5024.317 | 5127.617 | 5128.908 |
| Sec24b | 99683 | 1.529228 | 7.883804 | 50.22022 | 1.20E-23 | 1.27E-21 | -0.164 | 7.883804 | -6.22128 | 3.34E-06 | 1.33E-05 | 10226.72 | 10061.44 | 10366.16 | 31805.39 | 32369.91 | 30347.1 | 29634.71 | 29768.09 | 28306 | 10088.25 | 9963.189 | 9761.397 | 6153.161 | 5945.929 | 5903.954 |
| Ubr4 | 69116 | 1.53113 | 10.22909 | 27.30743 | 6.74E-11 | 4.93E-10 | -0.17727 | 10.22909 | -3.83147 | 0.003175 | 0.007782 | 47112.81 | 47940.57 | 47277.44 | 146828.2 | 147651.1 | 144535.3 | 139715.6 | 142565.9 | 124166.3 | 49286.65 | 49709.19 | 49336.14 | 40783.57 | 40237.21 | 41821.7 |
| Rab3ip | 216363 | 1.53165 | 2.696308 | 12.69731 | 2.04E-11 | 1.61E-10 | 0.189399 | 2.696308 | 1.94322 | 0.065295 | 0.108595 | 253.0154 | 207.8002 | 269.9385 | 722.0823 | 702.1976 | 834.596 | 898.2369 | 959.742 | 823.9446 | 248.4563 | 242.4839 | 290.3267 | 208.5817 | 206.7397 | 202.4676 |
| Lpcat2 | 270084 | 1.531694 | 6.19549 | 39.28187 | 2.15E-21 | 1.25E-19 | -1.08449 | 6.19549 | -27.6114 | 3.45E-18 | 1.33E-16 | 4378.428 | 4415.31 | 4443.136 | 13997.49 | 13930.2 | 12943.68 | 6863.269 | 6783.874 | 6397.349 | 2387.642 | 2433.226 | 2639.809 | 1606.079 | 1613.042 | 1633.508 |
| Ikzf1 | 22778 | 1.532398 | 7.986011 | 64.56401 | 5.81E-26 | 1.40E-23 | -1.18931 | 7.986011 | -48.7005 | 2.29E-23 | 5.03E-21 | 14550.98 | 14947.4 | 13991.67 | 44913.26 | 45064.24 | 44298.52 | 20696.4 | 20080.86 | 20348.1 | 11651.29 | 11399.79 | 11470.52 | 5028.488 | 5313.919 | 5433.42 |
| Rem2 | 140743 | 1.532637 | -2.20399 | 2.460955 | 0.02245 | 0.049378 | 1.073585 | -2.20399 | 2.227601 | 0.036789 | 0.066698 | 6.677827 | 1.77607 | 8.679696 | 22.07806 | 12.63956 | 13.52668 | 29.84648 | 34.85097 | 40.22045 | 1.538429 | 9.912862 | 2.985365 | 25.86413 | 31.4433 | 32.39481 |
| Cyth1 | 19157 | 1.533367 | 6.344162 | 29.95235 | 6.33E-19 | 2.08E-17 | -0.67467 | 6.344162 | -13.9206 | 3.50E-12 | 3.56E-11 | 2995.376 | 3141.868 | 3161.145 | 9648.11 | 9486.69 | 9555.245 | 6863.269 | 6178.004 | 5750.375 | 3757.613 | 3942.269 | 3766.784 | 3744.459 | 3782.63 | 3923.012 |
| Dnah10 | 56087 | 1.533806 | 0.274348 | 4.762221 | 0.000101 | 0.000331 | 0.950427 | 0.274348 | 3.981537 | 0.000662 | 0.001873 | 25.96933 | 39.96157 | 17.35939 | 90.90964 | 70.21976 | 83.8654 | 130.756 | 156.8293 | 204.5497 | 29.99936 | 22.87584 | 29.10731 | 211.919 | 202.0232 | 194.3689 |
| Ccdc148 | 227933 | 1.533955 | -1.66828 | 2.76936 | 0.011383 | 0.02695 | 0.168238 | -1.66828 | 0.360473 | 0.722037 | 0.801664 | 4.451884 | 10.65642 | 11.2836 | 23.37676 | 12.63956 | 52.75404 | 28.42522 | 22.78717 | 37.92213 | 7.692145 | 9.912862 | 14.92683 | 28.36712 | 33.01547 | 37.25403 |

| | | | | | | | | | | | | | | | | | | | | | | | | | | |
|---|---|---|---|---|---|---|---|---|---|---|---|---|---|---|---|---|---|---|---|---|---|---|---|---|---|---|
| Gm20310 | 100504605 | 1.534273 | -0.67805 | 4.134069 | 0.000458 | 0.001385 | -0.34579 | -0.67805 | -1.05244 | 0.304367 | 0.398703 | 17.80754 | 15.98463 | 24.30315 | 37.66257 | 94.09448 | 60.87005 | 54.00791 | 41.55307 | 50.56285 | 18.46115 | 28.97606 | 25.3756 | 45.88798 | 59.74228 | 49.40209 |
| Adamts16 | 271127 | 1.535462 | -1.9476 | 2.688788 | 0.013628 | 0.031703 | 0.674486 | -1.9476 | 1.490228 | 0.150795 | 0.221296 | 7.419807 | 3.55214 | 13.01954 | 19.48064 | 18.25714 | 32.46403 | 27.00396 | 25.46801 | 70.09849 | 3.846072 | 3.812639 | 3.731706 | 34.2074 | 36.94588 | 27.53559 |
| Adora3 | 11542 | 1.536231 | 1.173439 | 7.50262 | 2.05E-07 | 9.26E-07 | -0.30904 | 1.173439 | -1.74326 | 0.095679 | 0.150164 | 101.6514 | 86.13939 | 90.26884 | 303.8979 | 300.5406 | 262.4175 | 262.9333 | 274.7865 | 202.2514 | 79.9983 | 100.6537 | 53.73657 | 114.3028 | 81.75259 | 74.50806 |
| Ccr9 | 12769 | 1.536307 | 1.034723 | 7.647011 | 1.51E-07 | 6.98E-07 | -1.21699 | 1.034723 | -5.78706 | 9.02E-06 | 3.36E-05 | 89.03769 | 100.348 | 111.1001 | 333.7683 | 294.923 | 294.8816 | 79.59061 | 171.574 | 175.8208 | 72.30616 | 81.59048 | 84.33656 | 94.27894 | 85.68301 | 70.45871 |
| Alpk2 | 225638 | 1.536429 | -0.77833 | 3.757993 | 0.001132 | 0.00323 | 0.961447 | -0.77833 | 3.172868 | 0.004519 | 0.010677 | 19.2915 | 15.09659 | 13.01954 | 40.25998 | 58.9846 | 51.40137 | 132.1773 | 67.02109 | 113.7664 | 9.999788 | 11.43792 | 12.6878 | 65.91183 | 56.59795 | 41.30338 |
| Dnai3 | 242253 | 1.536617 | -1.5906 | 2.502034 | 0.020542 | 0.045631 | 2.005345 | -1.5906 | 4.559901 | 0.000164 | 0.000514 | 5.193865 | 8.88035 | 2.603909 | 25.97418 | 12.63956 | 12.17401 | 45.48035 | 76.40404 | 88.48498 | 10.769 | 13.7255 | 6.717071 | 46.72231 | 51.88145 | 46.16261 |
| Gpr18 | 110168 | 1.536703 | 3.160274 | 20.95072 | 1.02E-15 | 1.70E-14 | -2.54082 | 3.160274 | -24.9622 | 2.79E-17 | 8.71E-16 | 736.0449 | 883.5948 | 828.911 | 2531.184 | 2540.551 | 2457.797 | 451.961 | 467.8072 | 426.3367 | 450.7597 | 423.203 | 520.1999 | 91.77596 | 126.5593 | 91.51534 |
| Junb | 16477 | 1.537592 | 7.688663 | 37.67634 | 5.16E-21 | 2.81E-19 | -0.36154 | 7.688663 | -9.94311 | 1.82E-09 | 1.16E-08 | 10281.63 | 10606.69 | 10031.12 | 32118.38 | 32316.54 | 31365.66 | 27184.46 | 26329.9 | 24265.57 | 10772.08 | 10959.81 | 11689.2 | 3258.881 | 2833.042 | 2941.449 |
| Ankmy1 | 241158 | 1.537838 | -0.91067 | 3.1569 | 0.00469 | 0.012021 | 1.509649 | -0.91067 | 4.35186 | 0.000271 | 0.000819 | 9.64575 | 21.31284 | 12.15157 | 70.13029 | 50.55823 | 20.29002 | 109.4371 | 128.6805 | 136.7495 | 13.84586 | 24.40089 | 8.209754 | 29.20144 | 27.51289 | 33.20468 |
| Mgat5b | 268510 | 1.537865 | -1.55162 | 3.137318 | 0.004909 | 0.012537 | 0.972009 | -1.55162 | 2.618504 | 0.015923 | 0.032371 | 8.903769 | 4.440175 | 14.75548 | 24.67547 | 25.27911 | 31.11136 | 52.58665 | 38.87223 | 78.14258 | 3.076858 | 5.337695 | 2.985365 | 63.40885 | 67.60311 | 63.16988 |
| Tlr2 | 24088 | 1.538018 | 7.034904 | 44.06949 | 1.90E-22 | 1.39E-20 | 1.680226 | 7.034904 | 67.92446 | 1.98E-26 | 1.53E-23 | 5654.635 | 5510.257 | 5869.21 | 17587.12 | 17851.27 | 17376.37 | 59938.83 | 57741.35 | 58138.08 | 3094.55 | 3076.037 | 3254.794 | 1621.097 | 1521.856 | 1686.15 |
| Vrtn | 432677 | 1.538676 | -1.86753 | 2.982279 | 0.007021 | 0.017407 | 0.717432 | -1.86753 | 1.757485 | 0.093171 | 0.146878 | 3.709904 | 8.88035 | 10.41564 | 20.77935 | 19.66153 | 25.70069 | 35.53152 | 29.48928 | 49.41369 | 3.076858 | 6.100223 | 2.985365 | 48.39096 | 49.52321 | 34.01455 |
| Nr1h4 | 20186 | 1.538752 | -2.78077 | 2.585121 | 0.017136 | 0.038842 | 0.567367 | -2.78077 | 1.134601 | 0.269125 | 0.359686 | 3.709904 | 4.440175 | 2.603909 | 15.58451 | 7.021976 | 13.52668 | 18.47639 | 13.40422 | 22.98311 | 5.384501 | 1.525056 | 2.239024 | 17.52087 | 19.65207 | 17.00728 |
| Cfap44 | 212517 | 1.538991 | -0.82936 | 3.579692 | 0.001732 | 0.004791 | 0.827732 | -0.82936 | 2.562233 | 0.018017 | 0.036024 | 23.74338 | 10.65642 | 8.679696 | 51.94837 | 36.51428 | 39.22736 | 82.43313 | 73.7232 | 79.29174 | 18.46115 | 16.77561 | 20.89756 | 63.40885 | 55.81187 | 34.01455 |
| Hivep3 | 16656 | 1.540415 | 6.63154 | 38.24726 | 3.76E-21 | 2.08E-19 | -1.30013 | 6.63154 | -30.9669 | 3.16E-19 | 1.67E-17 | 6200.733 | 6066.167 | 5811.057 | 18294.92 | 19730.35 | 18240.72 | 8102.608 | 8191.317 | 7472.959 | 3306.853 | 3435.95 | 3410.78 | 2290.227 | 2373.183 | 2596.444 |
| Omp | 18378 | 1.540801 | -0.02445 | 5.691297 | 1.13E-05 | 4.13E-05 | -1.7967 | -0.02445 | -5.60157 | 1.39E-05 | 5.04E-05 | 44.51884 | 69.26673 | 72.04148 | 183.118 | 151.6747 | 229.9535 | 61.11422 | 44.23392 | 60.90525 | 24.61486 | 40.41398 | 29.85365 | 19.18952 | 62.88661 | 35.63429 |
| Igsf10 | 242050 | 1.54377 | -0.17523 | 3.646174 | 0.001479 | 0.004135 | 1.652614 | -0.17523 | 5.585922 | 1.44E-05 | 5.22E-05 | 13.35565 | 12.43249 | 12.15157 | 25.97418 | 60.389 | 41.9327 | 123.6497 | 117.9571 | 153.9868 | 10.769 | 7.625279 | 14.92683 | 570.6796 | 506.2372 | 516.6972 |
| Ist1 | 71955 | 1.544375 | 7.12201 | 45.51182 | 9.60E-23 | 7.70E-21 | -1.5292 | 7.12201 | -41.5274 | 6.65E-22 | 7.80E-20 | 7106.692 | 6971.963 | 7132.974 | 22218.32 | 22265.28 | 21602.1 | 8165.144 | 8290.508 | 7413.203 | 6763.703 | 6831.487 | 6714.086 | 4906.677 | 4615.091 | 4809.01 |
| Mirlet7e | 387248 | 1.544663 | -1.26397 | 4.527866 | 0.000178 | 0.000564 | -0.37383 | -1.26397 | -1.23093 | 0.231742 | 0.317202 | 19.2915 | 38.1855 | 24.30315 | 62.33804 | 103.9252 | 82.51273 | 66.79926 | 53.61687 | 73.54596 | 4.615287 | 7.625279 | 17.91219 | 6.674615 | 4.716496 | 11.33818 |
| BC051226 | 407803 | 1.545632 | 2.367918 | 10.84617 | 3.82E-10 | 2.52E-09 | 0.016933 | 2.367918 | 0.144133 | 0.886748 | 0.918721 | 149.1381 | 204.248 | 151.8947 | 546.7566 | 495.7515 | 505.8977 | 623.9335 | 537.5091 | 487.242 | 285.3786 | 303.4861 | 315.7024 | 205.2444 | 206.7397 | 177.3616 |
| Aftph | 216549 | 1.545865 | 7.41552 | 49.11473 | 1.92E-23 | 1.95E-21 | -1.3255 | 7.41552 | -40.1544 | 1.35E-21 | 1.42E-19 | 8299.055 | 8579.306 | 7959.281 | 25836.52 | 26666.66 | 24931.02 | 10943.71 | 10617.48 | 10520.52 | 7768.297 | 8006.542 | 7628.354 | 6098.095 | 6324.821 | 6220.614 |
| Ifitm5 | 73835 | 1.548085 | 0.834263 | 7.535001 | 1.91E-07 | 8.69E-07 | -0.6373 | 0.834263 | -3.38464 | 0.002751 | 0.006844 | 93.48957 | 63.93852 | 85.06102 | 263.638 | 285.0922 | 213.7215 | 173.3938 | 198.3824 | 142.4953 | 109.9977 | 155.5557 | 114.1902 | 28.36712 | 28.29897 | 38.0639 |
| Tspo | 12257 | 1.549392 | 6.765854 | 25.92331 | 1.28E-17 | 3.10E-16 | -0.24474 | 6.765854 | -4.6204 | 0.000142 | 0.000449 | 3979.985 | 4357.588 | 4472.647 | 12980.6 | 13562.25 | 13354.89 | 13138.14 | 11394.93 | 10715.88 | 4959.895 | 4895.429 | 5233.345 | 3485.818 | 3342.423 | 3642.796 |
| Cdc42ep4 | 56699 | 1.549777 | 5.17672 | 21.91804 | 4.05E-16 | 7.33E-15 | 0.075715 | 5.17672 | 1.251282 | 0.224374 | 0.308934 | 1294.756 | 1148.229 | 1229.913 | 4010.414 | 4016.57 | 3495.293 | 4378.905 | 4414.009 | 3861.163 | 1943.036 | 1862.093 | 1985.268 | 1229.798 | 1264.807 | 1223.714 |
| Kpna3 | 16648 | 1.551079 | 7.262588 | 47.09699 | 4.66E-23 | 4.19E-21 | -0.92846 | 7.262588 | -28.7987 | 1.44E-18 | 6.22E-17 | 7347.093 | 7369.802 | 7279.661 | 23344.3 | 23706.19 | 21822.59 | 12502.83 | 12578.52 | 12455.7 | 7212.155 | 7192.163 | 7538.793 | 4240.884 | 4006.663 | 4457.526 |
| Tunar | 69952 | 1.552527 | -2.63774 | 2.495771 | 0.020823 | 0.046184 | 0.720759 | -2.63774 | 1.459987 | 0.158858 | 0.231218 | 5.193865 | 4.440175 | 6.075787 | 14.2858 | 21.06593 | 16.23201 | 44.05909 | 32.17012 | 19.53565 | 0 | 3.050111 | 0 | 40.04769 | 36.94588 | 25.10598 |
| Grm1 | 14816 | 1.552536 | -1.38787 | 3.134044 | 0.004946 | 0.012623 | 0.434331 | -1.38787 | 1.103019 | 0.282301 | 0.374045 | 11.12971 | 19.53677 | 7.811726 | 36.36386 | 32.30109 | 45.9907 | 49.74413 | 57.63814 | 52.86116 | 5.384501 | 14.48803 | 1.492683 | 59.23721 | 44.80671 | 32.39481 |
| Dusp10 | 63953 | 1.552617 | 3.829925 | 11.792 | 8.18E-11 | 5.93E-10 | 2.313699 | 3.829925 | 26.21 | 1.02E-17 | 3.50E-16 | 253.7574 | 240.6575 | 218.7283 | 774.0307 | 766.7998 | 699.3292 | 4151.503 | 3716.99 | 3731.308 | 487.682 | 410.24 | 391.8292 | 1481.765 | 1630.335 | 1428.611 |
| Trmt61b | 68789 | 1.553565 | 2.298422 | 11.49723 | 1.31E-10 | 9.18E-10 | 0.648273 | 2.298422 | 6.343523 | 2.54E-06 | 1.02E-05 | 171.3976 | 170.5027 | 181.4056 | 527.2759 | 605.2944 | 514.0137 | 956.5086 | 853.8487 | 868.7616 | 155.3813 | 125.0546 | 186.5853 | 160.1908 | 186.3016 | 154.6852 |
| Tut7 | 214290 | 1.554577 | 7.520175 | 38.97126 | 2.54E-21 | 1.46E-19 | 0.019286 | 7.520175 | 0.57112 | 0.573884 | 0.667775 | 6680.053 | 6997.716 | 6451.618 | 20950.78 | 21206.37 | 20925.77 | 22858.14 | 22781.81 | 20969.79 | 6395.249 | 6535.626 | 6764.837 | 7611.564 | 7724.834 | 7900.284 |
| Pde7b | 29863 | 1.554665 | 5.159065 | 26.56756 | 7.68E-18 | 1.97E-16 | -2.49217 | 5.159065 | -32.6663 | 1.03E-19 | 6.33E-18 | 1831.208 | 2083.33 | 1721.184 | 5862.373 | 5863.35 | 5870.578 | 1110.005 | 1003.976 | 1128.471 | 3409.928 | 3321.571 | 3107.019 | 1091.3 | 1195.632 | 1141.917 |
| Tbx21 | 57765 | 1.556176 | -1.86773 | 2.681798 | 0.013842 | 0.032128 | 0.88284 | -1.86773 | 1.968742 | 0.062105 | 0.104259 | 11.12971 | 1.77607 | 7.811726 | 25.97418 | 19.66153 | 12.17401 | 46.90161 | 34.85097 | 29.87805 | 6.92293 | 6.862751 | 8.209754 | 30.8701 | 40.09021 | 37.25403 |
| Fnbp4 | 55935 | 1.557984 | 7.258976 | 34.70836 | 2.90E-20 | 1.30E-18 | -0.02257 | 7.258976 | -0.58909 | 0.561987 | 0.656634 | 5818.613 | 5588.404 | 5758.11 | 18196.21 | 17935.53 | 17868.74 | 19067.64 | 19249.8 | 17159.19 | 6309.866 | 6431.922 | 6089.398 | 5228.727 | 5478.21 | 5464.195 |
| Ifi202b | 26388 | 1.55803 | -0.57867 | 2.842986 | 0.009643 | 0.023233 | 4.278907 | -0.57867 | 12.40894 | 3.15E-11 | 2.69E-10 | 9.64575 | 6.216245 | 8.679696 | 22.07806 | 33.70549 | 24.34802 | 493.1775 | 621.9557 | 515.9709 | 27.69172 | 7.625279 | 8.209754 | 59.23721 | 71.53352 | 68.83897 |
| Pde9a | 18585 | 1.558282 | -2.10232 | 2.66796 | 0.014273 | 0.033037 | 0.470927 | -2.10232 | 0.985816 | 0.335268 | 0.432064 | 5.193865 | 3.55214 | 8.679696 | 25.97418 | 19.66153 | 10.82134 | 21.31891 | 32.17012 | 25.28142 | 6.153716 | 3.050111 | 7.463413 | 25.02981 | 47.95104 | 27.53559 |
| Tbc1d13 | 70296 | 1.558448 | 7.551311 | 53.8476 | 2.73E-24 | 3.67E-22 | -3.04106 | 7.551311 | -71.2713 | 7.11E-27 | 7.98E-24 | 12178.87 | 12199.82 | 12333.85 | 38511.92 | 39164.37 | 37786.77 | 4982.941 | 5257.134 | 4416.205 | 13538.94 | 13019.4 | 13364.73 | 5769.371 | 5678.661 | 5959.835 |
| Baiap2l1 | 66898 | 1.559755 | 2.877342 | 10.17708 | 1.20E-09 | 7.38E-09 | -1.2673 | 2.877342 | -7.92489 | 8.55E-08 | 4.23E-07 | 248.5635 | 245.9857 | 260.3909 | 802.6023 | 703.602 | 877.8814 | 349.6302 | 349.8501 | 327.5093 | 533.0656 | 487.2553 | 331.3755 | 564.005 | 519.6006 | 537.7539 |
| Kif26b | 269152 | 1.560043 | -0.68747 | 3.9433 | 0.000725 | 0.002131 | 0.578794 | -0.68747 | 1.884935 | 0.073128 | 0.119626 | 14.09763 | 17.7607 | 13.88751 | 55.84449 | 50.55823 | 39.22736 | 73.90556 | 61.6594 | 89.63414 | 9.230573 | 21.35078 | 6.717071 | 93.44461 | 111.6237 | 122.2904 |
| Samd3 | 268288 | 1.560079 | -2.01959 | 2.732216 | 0.012371 | 0.029058 | 0.770593 | -2.01959 | 1.720363 | 0.099835 | 0.155729 | 3.709904 | 15.98463 | 4.339848 | 12.98709 | 35.10988 | 18.93735 | 38.37404 | 36.19139 | 35.62382 | 3.846072 | 6.100223 | 2.239024 | 35.87606 | 24.36856 | 32.39481 |
| Fgf16 | 80903 | 1.560722 | -2.7221 | 2.551597 | 0.018441 | 0.0415 | 0.917928 | -2.7221 | 1.881999 | 0.073544 | 0.120216 | 3.709904 | 3.55214 | 5.207818 | 18.18193 | 9.830767 | 13.52668 | 24.16143 | 17.42548 | 43.66791 | 0.769214 | 3.050111 | 0.746341 | 25.02981 | 16.50774 | 22.67637 |
| Kcnc1 | 16502 | 1.56099 | -0.73089 | 3.433752 | 0.002449 | 0.006604 | 0.497936 | -0.73089 | 1.401443 | 0.175456 | 0.251112 | 21.51744 | 10.65642 | 16.49142 | 70.13029 | 40.72746 | 44.63804 | 52.58665 | 65.68067 | 110.3189 | 9.999788 | 12.96297 | 14.18048 | 75.92375 | 59.74228 | 111.7621 |
| Dennd1a | 227801 | 1.563699 | 7.596211 | 36.73136 | 8.81E-21 | 4.55E-19 | -0.48772 | 7.596211 | -12.5755 | 2.45E-11 | 2.13E-10 | 6243.026 | 6150.53 | 6522.792 | 19942.98 | 19860.96 | 19873.39 | 15028.41 | 15605.19 | 13756.54 | 11563.6 | 11556.87 | 11836.97 | 9414.545 | 9316.651 | 10046.44 |
| Slc49a4 | 224132 | 1.566177 | 4.732132 | 18.42993 | 1.37E-14 | 1.85E-13 | 0.073913 | 4.732132 | 1.032001 | 0.313625 | 0.408612 | 762.7562 | 611.8561 | 696.9796 | 2263.65 | 2200.687 | 2097.988 | 2542.636 | 2420.802 | 2239.704 | 1375.355 | 1556.319 | 1377.746 | 1948.988 | 1886.598 | 2028.725 |
| Dcp2 | 70640 | 1.566596 | 7.666883 | 50.91337 | 8.95E-24 | 1.01E-21 | -0.70561 | 7.666883 | -24.4098 | 4.43E-17 | 1.32E-15 | 9061.811 | 9202.707 | 8964.39 | 29172.9 | 29221.25 | 27724.28 | 18921.25 | 17958.97 | 17994.63 | 11252.07 | 11124.52 | 11301.1 | 4820.741 | 4754.228 | 5277.925 |
| Adam32 | 353188 | 1.568176 | -0.67661 | 4.808759 | 9.05E-05 | 0.000297 | -0.39783 | -0.67661 | -1.37862 | 0.182291 | 0.259397 | 25.22735 | 15.98463 | 18.22736 | 55.84449 | 74.43295 | 60.87005 | 49.74413 | 41.55307 | 57.45778 | 17.69193 | 23.63836 | 19.40487 | 67.58048 | 57.38403 | 45.35273 |
| Katna1 | 23924 | 1.569287 | 5.426252 | 22.15513 | 3.25E-16 | 6.05E-15 | 1.229375 | 5.426252 | 22.89085 | 1.66E-16 | 4.35E-15 | 1422.377 | 1511.436 | 1354.901 | 4349.377 | 4624.674 | 4609.892 | 11566.22 | 11409.67 | 10233.23 | 1481.507 | 1542.594 | 1606.126 | 1082.122 | 963.7373 | 1007.479 |
| Serpini1 | 20713 | 1.569407 | 0.577629 | 3.550676 | 0.001856 | 0.005109 | 1.720473 | 0.577629 | 5.605346 | 1.37E-05 | 5.00E-05 | 8.903769 | 34.63336 | 19.9633 | 55.84449 | 75.83734 | 43.28537 | 163.445 | 214.4675 | 209.1463 | 26.92251 | 36.60134 | 55.22925 | 650.775 | 709.8326 | 742.651 |
| Gm13889 | 620695 | 1.569628 | 0.251 | 3.624171 | 0.001558 | 0.004343 | 4.550265 | 0.251 | 16.97662 | 7.11E-14 | 1.01E-12 | 11.12971 | 13.32052 | 13.88751 | 28.5716 | 42.13186 | 56.81205 | 1101.477 | 931.5931 | 996.3179 | 47.6913 | 49.56431 | 41.79511 | 65.0775 | 91.18558 | 59.12053 |
| Casp3 | 12367 | 1.569871 | 5.095756 | 22.14439 | 3.28E-16 | 6.10E-15 | -0.03299 | 5.095756 | -0.53905 | 0.595434 | 0.688343 | 973.4787 | 951.0855 | 1169.155 | 3314.306 | 3109.331 | 3327.563 | 3301.589 | 3454.267 | 3159.029 | 1506.891 | 1652.398 | 1615.082 | 2004.053 | 2065.825 | 2032.774 |
| Dlgap1 | 224997 | 1.570245 | -0.4781 | 3.655643 | 0.001446 | 0.00405 | 1.020112 | -0.4781 | 3.232933 | 0.003928 | 0.009425 | 20.03348 | 15.98463 | 7.811726 | 37.66257 | 43.53625 | 54.10671 | 85.27565 | 72.38277 | 128.7054 | 23.07643 | 25.16342 | 29.85365 | 107.6282 | 94.32991 | 79.36729 |
| Clstn2 | 64085 | 1.570735 | -1.86378 | 2.507275 | 0.020309 | 0.045199 | 0.816262 | -1.86378 | 1.676134 | 0.108301 | 0.167151 | 10.38773 | 1.77607 | 10.41564 | 33.76644 | 11.23516 | 20.29002 | 34.11026 | 28.14886 | 50.56285 | 9.999788 | 5.337695 | 11.19512 | 34.2074 | 26.72681 | 22.67637 |
| 1700022I1 | 67317 | 1.571413 | -0.77998 | 3.781273 | 0.00107 | 0.003067 | -0.62766 | -0.77998 | -1.63454 | 0.11681 | 0.178038 | 19.2915 | 46.17782 | 13.88751 | 68.83159 | 82.85932 | 68.98605 | 34.11026 | 76.40404 | 44.81707 | 19.99958 | 16.77561 | 16.41951 | 20.85817 | 60.52836 | 26.72572 |
| Dcdc2a | 195208 | 1.571569 | -1.78474 | 2.551704 | 0.018436 | 0.041494 | 1.093533 | -1.78474 | 2.325025 | 0.030004 | 0.05591 | 3.709904 | 6.216245 | 6.943757 | 7.792255 | 26.68351 | 27.05335 | 46.90161 | 26.80843 | 49.41369 | 6.92293 | 9.912862 | 3.731706 | 50.05962 | 62.10053 | 46.97248 |
| Gm10723 | 100038361 | 1.572036 | -2.2174 | 2.733568 | 0.012333 | 0.028991 | 0.618665 | -2.2174 | 1.329306 | 0.197782 | 0.278042 | 4.451884 | 1.77607 | 12.15157 | 18.18193 | 15.44835 | 13.52668 | 28.42522 | 22.78717 | 25.28142 | 3.076858 | 4.575167 | 5.97073 | 36.71038 | 33.80155 | 29.9652 |
| Scn10a | 20264 | 1.572336 | -1.53408 | 3.158212 | 0.004676 | 0.01199 | 1.809234 | -1.53408 | 5.138218 | 4.12E-05 | 0.000141 | 4.451884 | 7.992315 | 10.41564 | 25.97418 | 18.25714 | 25.70069 | 62.53548 | 88.46784 | 108.0206 | 6.92293 | 4.575167 | 5.224389 | 40.04769 | 44.02063 | 39.68364 |
| Fam217a | 71864 | 1.573483 | -2.26647 | 2.518854 | 0.019805 | 0.044184 | 0.262134 | -2.26647 | 0.503324 | 0.619892 | 0.710683 | 5.193865 | 11.54445 | 3.471878 | 15.58451 | 29.4923 | 16.23201 | 21.31891 | 28.14886 | 24.13227 | 5.384501 | 3.812639 | 7.463413 | 22.52683 | 6.288661 | 25.91585 |
| Abcc5 | 27416 | 1.5749 | 6.60565 | 43.21767 | 2.86E-22 | 2.01E-20 | -2.32639 | 6.60565 | -51.505 | 7.01E-24 | 1.76E-21 | 4355.427 | 4193.301 | 4541.217 | 13809.17 | 14000.42 | 13825.62 | 2978.963 | 2936.864 | 2725.797 | 8625.202 | 8588.351 | 8691.144 | 4612.994 | 4408.351 | 4308.51 |
| Bcl2l15 | 229672 | 1.576505 | -1.8492 | 2.789599 | 0.010877 | 0.025868 | 2.182184 | -1.8492 | 5.589458 | 1.43E-05 | 5.18E-05 | 4.451884 | 7.10428 | 7.811726 | 12.98709 | 37.91867 | 17.58468 | 96.64574 | 93.82952 | 104.5732 | 7.692145 | 4.575167 | 2.985365 | 10.01192 | 18.86598 | 25.91585 |
| D630041G | 320749 | 1.576718 | -1.6883 | 2.596101 | 0.016727 | 0.037995 | -0.4126 | -1.6883 | -0.76025 | 0.455423 | 0.554233 | 19.2915 | 3.55214 | 13.01954 | 45.45482 | 15.44835 | 48.69604 | 11.37009 | 41.55307 | 32.17636 | 9.999788 | 6.100223 | 8.209754 | 25.86413 | 41.66238 | 25.10598 |
| Klra2 | 16633 | 1.577693 | 3.614014 | 31.85337 | 1.75E-19 | 6.56E-18 | -3.03772 | 3.614014 | -41.4465 | 6.93E-22 | 8.02E-20 | 1742.171 | 1970.55 | 1854.851 | 5919.516 | 5961.658 | 5796.181 | 713.4729 | 761.3596 | 754.9952 | 270.7635 | 320.2617 | 329.8828 | 75.08942 | 91.97167 | 85.84625 |
| Mthfs | 107885 | 1.578816 | 3.890901 | 14.75703 | 1.13E-12 | 1.08E-11 | -0.15389 | 3.890901 | -1.6875 | 0.106069 | 0.164189 | 515.6766 | 467.9944 | 339.3761 | 1422.087 | 1467.593 | 1305.324 | 1330.3 | 1328.358 | 1258.325 | 706.1389 | 690.8502 | 675.4388 | 1046.246 | 1015.619 | 940.2594 |
| Homer1 | 26556 | 1.581301 | 5.002091 | 24.85171 | 3.06E-17 | 6.92E-16 | -0.48021 | 5.002091 | -8.25378 | 4.40E-08 | 2.27E-07 | 1029.869 | 1136.685 | 1130.964 | 3441.579 | 3440.768 | 3619.739 | 2750.14 | 2406.057 | 2662.594 | 1658.426 | 1700.437 | 1696.434 | 1636.115 | 1452.681 | 1503.119 |
| Lrp4 | 228357 | 1.581909 | 3.811794 | 16.88142 | 7.95E-14 | 9.35E-13 | 0.057623 | 3.811794 | 0.742436 | 0.465917 | 0.564574 | 413.2833 | 426.2568 | 414.8895 | 1332.476 | 1341.197 | 1335.083 | 1471.005 | 1554.889 | 1330.722 | 728.4461 | 830.3928 | 747.8339 | 639.0944 | 733.4151 | 637.3679 |
| A530083I2 | 331045 | 1.583648 | -1.7937 | 2.592061 | 0.016877 | 0.038304 | 1.003312 | -1.7937 | 2.129619 | 0.044997 | 0.079157 | 4.451884 | 2.664105 | 10.41564 | 28.5716 | 19.66153 | 8.116006 | 31.26774 | 45.57434 | 32.17636 | 7.692145 | 9.912862 | 10.44878 | 35.04173 | 49.52321 | 53.45144 |
| Col9a3 | 12841 | 1.5844 | 1.146675 | 6.417126 | 2.15E-06 | 8.56E-06 | 1.2981 | 1.146675 | 7.48839 | 2.11E-07 | 9.85E-07 | 39.32498 | 59.49834 | 41.66254 | 155.8451 | 144.6527 | 139.3248 | 422.1145 | 396.7648 | 325.211 | 86.92123 | 114.3792 | 85.0829 | 172.7057 | 156.4304 | 164.4037 |
| Tert | 21752 | 1.586941 | -1.10755 | 2.686162 | 0.013708 | 0.031872 | 1.124501 | -1.10755 | 2.564257 | 0.017937 | 0.035887 | 7.419807 | 1.77607 | 24.30315 | 31.16902 | 18.25714 | 22.99535 | 62.53548 | 49.5956 | 52.86116 | 27.69172 | 20.58825 | 24.62926 | 70.91779 | 65.24486 | 61.55014 |
| Pgs1 | 74451 | 1.587959 | 6.265277 | 32.22997 | 1.37E-19 | 5.22E-18 | -0.98329 | 6.265277 | -20.3208 | 1.90E-15 | 3.81E-14 | 4098.702 | 4293.649 | 3940.582 | 13249.43 | 13183.06 | 13116.82 | 7119.096 | 7448.724 | 6331.847 | 3245.316 | 3268.957 | 3241.36 | 1763.767 | 1547.797 | 1729.073 |
| Golga3 | 269682 | 1.588485 | 6.280509 | 31.41245 | 2.34E-19 | 8.46E-18 | -2.56322 | 6.280509 | -39.1875 | 2.26E-21 | 2.26E-19 | 4903.009 | 4853.111 | 4288.638 | 15026.07 | 15416.05 | 14668.33 | 2824.045 | 2832.311 | 2339.681 | 5728.34 | 5978.218 | 5769.964 | 1992.373 | 1861.444 | 2133.198 |
| Ano1 | 101772 | 1.589041 | -1.56194 | 2.526374 | 0.019483 | 0.04356 | 0.62208 | -1.56194 | 1.264288 | 0.21976 | 0.303534 | 14.09763 | 6.216245 | 5.207818 | 28.5716 | 18.25714 | 33.81669 | 36.95278 | 22.78717 | 79.29174 | 7.692145 | 8.387806 | 7.463413 | 65.91183 | 50.30929 | 28.34546 |
| Scube3 | 268935 | 1.589115 | -0.27699 | 3.237914 | 0.003883 | 0.010116 | 2.580148 | -0.27699 | 8.072112 | 6.34E-08 | 3.20E-07 | 14.83961 | 19.53677 | 12.15157 | 46.75353 | 35.10988 | 73.04406 | 328.3113 | 281.4886 | 321.7636 | 17.69193 | 25.16342 | 3.731706 | 101.7879 | 78.60826 | 113.3818 |
| Met | 17295 | 1.589198 | 5.526925 | 29.6169 | 8.01E-19 | 2.54E-17 | -0.36343 | 5.526925 | -7.52023 | 1.97E-07 | 9.26E-07 | 2356.531 | 2400.359 | 2698.517 | 7935.113 | 8190.433 | 7754.844 | 6598.914 | 6124.387 | 6531.8 | 2445.333 | 2346.298 | 2553.98 | 493.9215 | 452.7836 | 601.7336 |
| Fbxo39 | 628100 | 1.590289 | -1.53182 | 2.783955 | 0.011016 | 0.026177 | 0.638533 | -1.53182 | 1.447431 | 0.162306 | 0.235352 | 14.83961 | 19.53677 | 2.603909 | 32.46773 | 30.8967 | 29.75869 | 44.05909 | 50.93603 | 56.30862 | 9.230573 | 22.87584 | 5.224389 | 18.35519 | 28.29897 | 20.24676 |
| Tent4a | 210106 | 1.590768 | 6.778318 | 38.02691 | 4.25E-21 | 2.33E-19 | -0.20847 | 6.778318 | -5.70669 | 1.09E-05 | 4.01E-05 | 5015.79 | 5161.259 | 5160.079 | 16504 | 16584.5 | 16199.55 | 15312.66 | 15015.4 | 14088.65 | 4370.677 | 4687.259 | 4144.433 | 2230.156 | 2246.624 | 2200.418 |
| Cstdc5 | 100034684 | 1.591114 | -1.15425 | 3.704763 | 0.001285 | 0.003632 | -0.04055 | -1.15425 | -0.11285 | 0.911202 | 0.928896 | 17.06556 | 29.30515 | 7.811726 | 40.25998 | 60.389 | 56.81205 | 46.90161 | 57.63814 | 51.712 | 23.07643 | 18.30067 | 8.956095 | 27.53279 | 15.72165 | 16.19741 |
| Igfbp7 | 29817 | 1.591265 | -1.06723 | 3.38668 | 0.002738 | 0.007316 | 1.334804 | -1.06723 | 3.984416 | 0.000657 | 0.001862 | 12.61367 | 12.43249 | 11.2836 | 35.06515 | 33.70549 | 52.75404 | 88.11817 | 115.2763 | 111.4681 | 3.846072 | 8.387806 | 24.62926 | 30.03577 | 35.37372 | 44.54286 |
| Usp42 | 76800 | 1.592176 | 5.306303 | 28.3447 | 2.00E-18 | 5.90E-17 | -0.49814 | 5.306303 | -9.67085 | 2.97E-09 | 1.83E-08 | 1691.716 | 1649.969 | 1841.832 | 5718.216 | 5621.794 | 5326.806 | 4216.881 | 4044.052 | 4000.211 | 1563.813 | 1626.472 | 1647.921 | 1141.359 | 1077.719 | 1305.511 |
| Cpeb3 | 208922 | 1.59258 | 4.911472 | 22.25343 | 2.97E-16 | 5.57E-15 | 0.919004 | 4.911472 | 16.65916 | 1.03E-13 | 1.43E-12 | 743.4647 | 781.4708 | 715.207 | 2276.637 | 2561.617 | 2384.753 | 4904.771 | 4782.625 | 4501.242 | 839.9822 | 822.005 | 799.3315 | 2791.658 | 2623.158 | 2698.488 |
| Mfsd4b1 | 215929 | 1.592866 | -2.12302 | 3.115405 | 0.005165 | 0.013131 | 0.534832 | -2.12302 | 1.31194 | 0.203477 | 0.284707 | 8.161788 | 4.440175 | 6.943757 | 16.88322 | 26.68351 | 21.64268 | 25.5827 | 36.19139 | 35.62382 | 2.307643 | 6.862751 | 2.239024 | 25.86413 | 21.22423 | 34.01455 |
| Rnf165 | 225743 | 1.593063 | -1.35646 | 3.236204 | 0.003898 | 0.010152 | 1.361026 | -1.35646 | 3.823878 | 0.000966 | 0.002653 | 8.903769 | 5.32821 | 9.547666 | 36.36386 | 33.70549 | 13.52668 | 63.95674 | 65.68067 | 78.14258 | 9.230573 | 6.100223 | 5.97073 | 64.24317 | 56.59795 | 69.64884 |
| Pkd1l3 | 244646 | 1.593563 | 1.352231 | 7.742091 | 1.24E-07 | 5.79E-07 | -1.20369 | 1.352231 | -5.67568 | 1.17E-05 | 4.29E-05 | 74.94006 | 118.9967 | 138.8751 | 328.5734 | 348.29 | 344.9303 | 177.6576 | 112.5954 | 175.8208 | 147.6892 | 128.8672 | 121.6536 | 84.26702 | 140.7088 | 114.1917 |
| Zc2hc1a | 67306 | 1.593836 | 3.240654 | 20.13489 | 2.29E-15 | 3.54E-14 | -2.3342 | 3.240654 | -22.2037 | 3.11E-16 | 7.64E-15 | 484.5134 | 525.7167 | 559.8404 | 1728.582 | 1685.274 | 1621.849 | 328.3113 | 339.1267 | 364.2823 | 688.4469 | 590.9591 | 638.1218 | 312.0383 | 316.7913 | 268.0671 |
| Neb | 17996 | 1.595839 | 1.333566 | 6.828842 | 8.65E-07 | 3.61E-06 | -0.44403 | 1.333566 | -2.14967 | 0.043194 | 0.076418 | 80.8759 | 97.68385 | 65.96569 | 276.6251 | 203.6373 | 309.7609 | 180.5001 | 172.9144 | 240.1735 | 103.0747 | 73.20267 | 67.91705 | 257.807 | 259.4073 | 189.5096 |
| Pla2g4c | 232889 | 1.596977 | 1.416227 | 9.123033 | 8.17E-09 | 4.45E-08 | -3.37788 | 1.416227 | -10.9347 | 3.29E-10 | 2.36E-09 | 132.8146 | 126.101 | 132.7993 | 446.756 | 351.0988 | 478.8444 | 21.31891 | 76.40404 | 42.51876 | 156.1505 | 125.8171 | 128.3707 | 335.3994 | 314.433 | 330.4271 |
| Prkag3 | 241113 | 1.597923 | -1.52162 | 2.571922 | 0.017639 | 0.039853 | 0.119784 | -1.52162 | 0.229541 | 0.820634 | 0.888157 | 4.451884 | 6.216245 | 18.22736 | 19.48064 | 30.8967 | 29.75869 | 31.26774 | 14.74464 | 50.56285 | 11.53822 | 18.30067 | 17.91219 | 29.20144 | 62.10053 | 26.72572 |
| Cntnap5c | 620292 | 1.598227 | -2.07644 | 2.704639 | 0.013156 | 0.03071 | 1.095416 | -2.07644 | 2.450799 | 0.022947 | 0.0444 | 9.64575 | 4.440175 | 3.471878 | 19.48064 | 21.06593 | 14.87935 | 27.00396 | 54.95729 | 45.96622 | 6.153716 | 5.337695 | 2.239024 | 18.35519 | 34.58764 | 30.77507 |
| Avpr1b | 26361 | 1.599716 | -1.99974 | 2.970962 | 0.007205 | 0.017819 | 0.569938 | -1.99974 | 1.335938 | 0.195641 | 0.275428 | 8.903769 | 8.88035 | 3.471878 | 14.2858 | 22.47032 | 33.81669 | 36.95278 | 22.78717 | 45.96622 | 3.076858 | 6.100223 | 2.985365 | 30.03577 | 29.87114 | 35.63429 |
| Zyx | 22793 | 1.602077 | 8.4601 | 41.72877 | 6.00E-22 | 4.00E-20 | 0.517124 | 8.4601 | 17.4108 | 4.29E-14 | 6.50E-13 | 10447.09 | 10253.25 | 10399.14 | 33584.62 | 33857.16 | 33417.66 | 50586.94 | 51654.49 | 48011.72 | 17096.56 | 17407.75 | 16586.69 | 13761.39 | 13424.72 | 14575.24 |
| Cxcr6 | 80901 | 1.602367 | 1.51084 | 11.31645 | 1.75E-10 | 1.21E-09 | -1.82818 | 1.51084 | -10.9503 | 3.21E-10 | 2.30E-09 | 149.8801 | 170.5027 | 131.9314 | 466.2366 | 519.6262 | 476.139 | 162.0237 | 119.2975 | 147.0919 | 102.3055 | 90.74081 | 104.4878 | 120.9774 | 125.7732 | 172.5024 |
| Kcnc4 | 99738 | 1.602454 | -2.11485 | 2.600536 | 0.016565 | 0.037682 | 0.692058 | -2.11485 | 1.407832 | 0.17358 | 0.248927 | 3.709904 | 7.992315 | 4.339848 | 10.38967 | 18.25714 | 25.70069 | 18.47639 | 40.21265 | 31.0272 | 2.307643 | 4.575167 | 5.224389 | 38.37904 | 59.74228 | 31.58494 |
| Ppp2r5a | 226849 | 1.60399 | 7.20547 | 45.76646 | 8.54E-23 | 6.91E-21 | 0.079593 | 7.20547 | 2.713232 | 0.012906 | 0.026946 | 5280.677 | 5213.653 | 4962.182 | 17044.26 | 16560.63 | 16585.06 | 18979.52 | 18256.54 | 17923.38 | 5639.88 | 5738.022 | 5746.828 | 5953.757 | 5947.501 | 5830.256 |
| Pcdhga4 | 93712 | 1.604621 | -2.27791 | 2.609742 | 0.016233 | 0.037042 | -0.09626 | -2.27791 | -0.18031 | 0.85861 | 0.918721 | 5.935846 | 7.992315 | 5.207818 | 38.96127 | 14.04395 | 16.23201 | 27.00396 | 12.0638 | 25.28142 | 10.769 | 6.100223 | 9.702436 | 16.68654 | 3.930413 | 15.38754 |
| Slc22a13b | 109280 | 1.604907 | -1.42988 | 2.988175 | 0.006927 | 0.017199 | -0.29526 | -1.42988 | -0.63611 | 0.531467 | 0.627327 | 23.74338 | 15.98463 | 3.471878 | 42.8574 | 32.30109 | 39.22736 | 27.00396 | 49.5956 | 25.28142 | 12.30743 | 21.35078 | 22.39024 | 12.5149 | 20.43815 | 28.34546 |
| Dcn | 13179 | 1.605296 | -0.05976 | 5.851279 | 7.77E-06 | 2.91E-05 | -0.57407 | -0.05976 | -2.3062 | 0.031218 | 0.057841 | 27.45329 | 29.30515 | 42.53051 | 84.4161 | 116.5648 | 120.3874 | 72.4843 | 60.31898 | 87.33582 | 16.92272 | 18.30067 | 28.36097 | 110.9655 | 123.415 | 110.1424 |
| Tet1 | 52463 | 1.605994 | 0.612171 | 6.249984 | 3.13E-06 | 1.22E-05 | 0.645631 | 0.612171 | 3.366 | 0.002874 | 0.007121 | 53.42261 | 31.08122 | 59.02193 | 158.4425 | 176.9538 | 125.7981 | 248.7206 | 229.2121 | 263.1566 | 24.61486 | 30.50111 | 34.3317 | 108.4625 | 150.9279 | 106.9029 |
| Clec9a | 232414 | 1.606432 | -1.16625 | 3.612618 | 0.001602 | 0.004457 | 1.074763 | -1.16625 | 3.36108 | 0.002908 | 0.007194 | 25.96933 | 10.65642 | 10.41564 | 44.15611 | 36.51428 | 70.33872 | 125.071 | 101.8721 | 97.67823 | 5.384501 | 19.0632 | 14.18048 | 14.18356 | 11.79124 | 17.00728 |
| Sh3bp4 | 98402 | 1.606499 | 4.527809 | 27.85047 | 2.88E-18 | 8.24E-17 | -1.93156 | 4.527809 | -28.9523 | 1.29E-18 | 5.63E-17 | 1503.253 | 1433.288 | 1378.336 | 4819.51 | 4721.577 | 4508.442 | 1191.017 | 1379.294 | 1256.027 | 1202.282 | 1128.541 | 1092.644 | 474.732 | 491.3016 | 566.0993 |
| Pik3r6 | 104709 | 1.607363 | 5.25654 | 38.60851 | 3.09E-21 | 1.74E-19 | -2.59864 | 5.25654 | -47.8652 | 3.31E-23 | 6.85E-21 | 2700.068 | 2933.18 | 2708.933 | 8687.066 | 9269.009 | 9154.855 | 1590.391 | 1509.315 | 1542.167 | 2793.018 | 2667.322 | 2759.224 | 594.8751 | 587.9898 | 571.7684 |
| Gm10791 | 100038614 | 1.607921 | 1.901081 | 11.691 | 9.60E-11 | 6.89E-10 | -0.59557 | 1.901081 | -4.81315 | 8.95E-05 | 0.000291 | 165.4617 | 155.4061 | 154.4986 | 489.6134 | 539.2878 | 526.1878 | 361.0003 | 392.7436 | 321.7636 | 142.3047 | 186.8193 | 126.1317 | 128.4863 | 124.2011 | 122.2904 |
| 4930523C( | 67647 | 1.608179 | 2.337841 | 11.10762 | 2.47E-10 | 1.67E-09 | 1.024515 | 2.337841 | 9.889655 | 2.00E-09 | 1.27E-08 | 194.399 | 141.1976 | 148.4228 | 509.094 | 561.7581 | 511.3084 | 1200.965 | 1127.295 | 1030.793 | 239.2257 | 184.5317 | 213.4536 | 136.8296 | 121.8428 | 119.0509 |
| Tax1bp1 | 52440 | 1.60843 | 7.866795 | 56.72223 | 9.07E-25 | 1.41E-22 | 0.140442 | 7.866795 | 6.031651 | 5.14E-06 | 1.99E-05 | 7704.728 | 7886.639 | 8116.384 | 25448.21 | 25971.48 | 25692.57 | 29972.97 | 29302.96 | 29011.58 | 10735.93 | 10645.65 | 11050.33 | 8310.73 | 8007.824 | 8238.81 |
| Slc31a2 | 20530 | 1.609354 | 6.666434 | 40.66532 | 1.03E-21 | 6.43E-20 | -1.38461 | 6.666434 | -33.4943 | 6.11E-20 | 4.01E-18 | 4630.702 | 4715.466 | 4492.611 | 15144.25 | 15274.2 | 14668.33 | 6257.812 | 6155.217 | 5588.344 | 12057.44 | 11551.53 | 12109.39 | 1700.358 | 1701.869 | 1604.353 |
| Sik3 | 70661 | 1.611477 | 6.69765 | 46.03671 | 7.54E-23 | 6.14E-21 | -0.82455 | 6.69765 | -24.6525 | 3.61E-17 | 1.10E-15 | 4142.479 | 4115.154 | 4140.215 | 13646.84 | 13680.21 | 13135.76 | 7876.628 | 7901.786 | 7923.428 | 4759.899 | 5090.636 | 4798.228 | 4211.682 | 4207.114 | 4415.413 |
| Rap2c | 72065 | 1.611586 | 7.339222 | 51.00505 | 8.62E-24 | 9.77E-22 | -0.49731 | 7.339222 | -17.4393 | 4.15E-14 | 6.31E-13 | 7870.932 | 8489.614 | 8045.21 | 26274.19 | 27495.25 | 25752.09 | 19572.18 | 19487.05 | 19380.51 | 6685.243 | 6601.204 | 6742.447 | 3022.766 | 2871.56 | 3208.706 |
| Gm15558 | 100503866 | 1.61285 | -0.69514 | 4.033779 | 0.000584 | 0.001738 | 0.232218 | -0.69514 | 0.724781 | 0.476458 | 0.574831 | 17.06556 | 11.54445 | 20.83127 | 62.33804 | 57.58021 | 41.9327 | 52.58665 | 65.68067 | 76.99342 | 19.23036 | 34.31375 | 23.13658 | 31.70442 | 58.9562 | 55.07118 |
| Slc6a15 | 103098 | 1.612903 | -1.91267 | 2.608684 | 0.016271 | 0.037109 | 1.114155 | -1.91267 | 2.383581 | 0.0265 | 0.050291 | 4.451884 | 3.55214 | 9.547666 | 23.37676 | 16.85274 | 14.87935 | 56.85043 | 28.14886 | 44.81707 | 6.153716 | 15.25056 | 2.985365 | 25.86413 | 40.09021 | 29.15533 |
| Hinfp | 102423 | 1.6137 | 4.363859 | 22.96258 | 1.56E-16 | 3.11E-15 | -2.52437 | 4.363859 | -26.9244 | 5.82E-18 | 2.11E-16 | 1087.744 | 1110.932 | 1145.72 | 3676.646 | 3933.711 | 3335.679 | 618.2485 | 726.5086 | 630.8864 | 1391.509 | 1486.929 | 1246.39 | 775.924 | 764.0723 | 772.6162 |
| Trip6 | 22051 | 1.614021 | 2.495772 | 11.10883 | 2.46E-10 | 1.67E-09 | -0.77885 | 2.495772 | -5.72336 | 1.04E-05 | 3.87E-05 | 149.1381 | 221.1207 | 214.3885 | 640.2636 | 542.0966 | 681.7445 | 375.2129 | 388.7223 | 361.984 | 300.7629 | 270.6974 | 338.8389 | 302.8607 | 314.433 | 348.2442 |
| Notch1 | 18128 | 1.614281 | 8.174954 | 36.20546 | 1.14E-17 | 2.80E-16 | 0.049352 | 8.174954 | 1.3528 | 0.193649 | 0.272929 | 12884.5 | 12869.4 | 12628.09 | 41730.12 | 42598.12 | 41207.67 | 46887.4 | 46904.04 | 41849.95 | 9988.25 | 10492.38 | 10033.81 | 6214.067 | 6316.96 | 6246.529 |
| Cdh16 | 12556 | 1.615362 | -2.52143 | 2.720357 | 0.012703 | 0.029768 | -0.0173 | -2.52143 | -0.03401 | 0.973188 | 0.977723 | 11.87169 | 3.55214 | 4.339848 | 24.67547 | 29.4923 | 10.82134 | 18.47639 | 16.08506 | 27.57973 | 1.538429 | 3.812639 | 3.731706 | 9.177596 | 22.7964 | 12.95792 |
| Pax7 | 18509 | 1.617276 | -2.01827 | 2.536248 | 0.019069 | 0.042734 | 1.087151 | -2.01827 | 2.201396 | 0.038839 | 0.069809 | 2.967923 | 5.32821 | 5.207818 | 12.98709 | 16.85274 | 14.87935 | 15.63387 | 40.21265 | 52.86116 | 11.53822 | 6.100223 | 7.463413 | 35.87606 | 39.30413 | 26.72572 |
| Socs2 | 216233 | 1.617358 | 2.506311 | 6.504432 | 1.77E-06 | 7.11E-06 | 1.624321 | 2.506311 | 9.665076 | 3.00E-09 | 1.85E-08 | 57.13252 | 53.2821 | 62.49381 | 170.1309 | 216.2769 | 185.3155 | 633.8823 | 595.1473 | 595.2626 | 299.9936 | 354.5755 | 257.4877 | 2054.947 | 2256.057 | 2395.596 |
| B430306N( | 320148 | 1.617485 | 6.907426 | 39.63977 | 1.77E-21 | 1.05E-19 | 0.806549 | 6.907426 | 25.635 | 1.61E-17 | 5.30E-16 | 3692.096 | 3782.141 | 3662.832 | 12150.72 | 12454.18 | 11883.19 | 22451.66 | 22366.28 | 21509.89 | 3629.923 | 3585.406 | 3584.677 | 5429.8 | 5031.715 | 5430.18 |
| Trim25 | 217069 | 1.618275 | 8.857252 | 44.15793 | 1.82E-22 | 1.34E-20 | 0.329198 | 8.857252 | 11.4955 | 1.31E-10 | 1.01E-09 | 14084.28 | 13985.66 | 14675.63 | 46823.66 | 47071.12 | 46155.73 | 62418.93 | 63249.14 | 57763.45 | 20728.02 | 20921.48 | 20951.29 | 19292.98 | 19518.43 | 19115.37 |
| Pdzrn3 | 55983 | 1.619671 | -1.88685 | 2.803858 | 0.010533 | 0.025128 | 1.002314 | -1.88685 | 2.277533 | 0.033153 | 0.060977 | 4.451884 | 9.768385 | 5.207818 | 14.2858 | 22.47032 | 27.05335 | 24.16143 | 42.8935 | 70.09849 | 5.384501 | 5.337695 | 2.985365 | 38.37904 | 44.02063 | 34.01455 |
| Cd200r4 | 239849 | 1.620605 | 4.125058 | 18.55246 | 1.20E-14 | 1.63E-13 | -2.25728 | 4.125058 | -20.0685 | 2.45E-15 | 4.81E-14 | 583.9388 | 490.1953 | 582.4076 | 1753.257 | 1921.213 | 1777.405 | 386.583 | 408.8286 | 386.1163 | 1828.423 | 1555.557 | 1788.98 | 1880.573 | 1733.312 | 1843.265 |
| P2ry13 | 74191 | 1.620762 | 1.843207 | 9.466041 | 4.31E-09 | 2.45E-08 | 0.889589 | 1.843207 | 7.199555 | 3.88E-07 | 1.75E-06 | 103.8773 | 94.13171 | 108.4962 | 276.6251 | 335.6505 | 412.5637 | 613.9847 | 668.8705 | 656.1678 | 78.45987 | 93.79093 | 80.60486 | 221.0966 | 253.1186 | 247.0104 |
| Tspan3 | 56434 | 1.621289 | 6.780085 | 47.10899 | 4.63E-23 | 4.19E-21 | -1.03827 | 6.780085 | -30.6027 | 4.04E-19 | 2.07E-17 | 5273.257 | 5188.788 | 5378.808 | 17462.44 | 17921.49 | 16682.45 | 9059.117 | 8429.912 | 8818.62 | 6445.248 | 6324.406 | 6684.232 | 2227.653 | 2408.557 | 2418.273 |
| Sgcb | 24051 | 1.621543 | 4.584433 | 28.83514 | 1.40E-18 | 4.24E-17 | -2.63237 | 4.584433 | -34.3171 | 3.67E-20 | 2.56E-18 | 1208.687 | 1122.476 | 1216.893 | 3857.166 | 3967.417 | 3845.634 | 659.465 | 612.5727 | 676.8526 | 1983.035 | 2058.063 | 2233.799 | 1018.713 | 940.1548 | 955.6469 |
| Adamts6 | 108154 | 1.621788 | 2.93452 | 11.80597 | 8.00E-11 | 5.81E-10 | 1.220587 | 2.93452 | 12.54915 | 2.55E-11 | 2.21E-10 | 263.4032 | 237.9934 | 159.7064 | 729.8745 | 754.1603 | 665.5125 | 1704.092 | 1818.952 | 1680.065 | 299.2244 | 286.7105 | 282.8633 | 251.1324 | 282.2037 | 236.4821 |
| Tdrd6 | 210510 | 1.623817 | -0.81407 | 3.497841 | 0.002104 | 0.005741 | 0.903268 | -0.81407 | 2.665416 | 0.014354 | 0.029552 | 34.8731 | 15.09659 | 5.207818 | 45.45482 | 44.94065 | 58.16471 | 78.16935 | 92.4891 | 117.2139 | 11.53822 | 12.20045 | 12.6878 | 58.40288 | 43.23454 | 55.88105 |
| Chrna2 | 110902 | 1.624833 | 1.329678 | 10.08649 | 1.41E-09 | 8.59E-09 | -1.53351 | 1.329678 | -8.74951 | 1.66E-08 | 9.13E-08 | 140.9763 | 131.4292 | 134.5353 | 424.6779 | 424.1274 | 500.4871 | 170.5513 | 199.7228 | 126.4071 | 69.2293 | 100.6537 | 86.57559 | 81.76404 | 78.60826 | 113.3818 |
| Lrrtm4 | 243499 | 1.625688 | -2.78622 | 2.62662 | 0.01564 | 0.035863 | 0.847811 | -2.78622 | 1.71797 | 0.100278 | 0.156346 | 3.709904 | 6.216245 | 2.603909 | 18.18193 | 11.23516 | 12.17401 | 24.16143 | 17.42548 | 39.07129 | 1.538429 | 1.525056 | 0.746341 | 28.36712 | 18.0799 | 12.14805 |
| Gm11992 | 626870 | 1.626653 | -1.2964 | 3.390022 | 0.002716 | 0.007261 | -0.43729 | -1.2964 | -1.0234 | 0.31758 | 0.412885 | 15.5816 | 11.54445 | 6.075787 | 25.97418 | 53.36702 | 32.46403 | 27.00396 | 17.42548 | 39.07129 | 28.46093 | 19.0632 | 25.3756 | 44.21933 | 25.15464 | 31.58494 |
| Chgb | 12653 | 1.627371 | -2.28808 | 2.563289 | 0.017975 | 0.040543 | 1.752534 | -2.28808 | 3.654413 | 0.00145 | 0.003835 | 1.483961 | 5.32821 | 2.603909 | 15.58451 | 7.021976 | 8.116006 | 38.37404 | 25.46801 | 44.81707 | 6.153716 | 5.337695 | 5.97073 | 36.71038 | 41.66238 | 33.20468 |
| Chrna5 | 110835 | 1.627541 | -1.70424 | 2.663867 | 0.014403 | 0.033306 | 1.547159 | -1.70424 | 3.573503 | 0.001758 | 0.004562 | 8.161788 | 9.768385 | 6.075787 | 62.33804 | 14.04395 | 20.29002 | 62.53548 | 99.19121 | 87.33582 | 2.307643 | 9.912862 | 3.731706 | 23.36115 | 33.80155 | 15.38754 |
| Pigr | 18703 | 1.627796 | -2.21043 | 2.948935 | 0.007578 | 0.018654 | 0.716566 | -2.21043 | 1.667214 | 0.11008 | 0.16944 | 8.903769 | 4.440175 | 5.207818 | 31.16902 | 15.44835 | 17.58468 | 27.00396 | 42.8935 | 37.92213 | 1.538429 | 7.625279 | 2.239024 | 25.86413 | 17.29382 | 21.8665 |
| Slc9c1 | 208169 | 1.628366 | -2.07906 | 3.126175 | 0.005038 | 0.012837 | 0.720267 | -2.07906 | 1.788495 | 0.087902 | 0.139802 | 11.87169 | 3.55214 | 6.075787 | 24.67547 | 19.66153 | 22.99535 | 32.689 | 33.51054 | 50.56285 | 6.92293 | 7.625279 | 4.478048 | 17.52087 | 16.50774 | 8.908573 |
| Gm8378 | 666955 | 1.628874 | -1.47796 | 3.616757 | 0.001586 | 0.004416 | -0.69522 | -1.47796 | -1.67781 | 0.10797 | 0.166698 | 17.80754 | 7.10428 | 21.69924 | 31.16902 | 61.79339 | 58.16471 | 28.42522 | 34.85097 | 28.72889 | 3.076858 | 10.67539 | 5.97073 | 37.54471 | 22.7964 | 40.49351 |
| Mucl3 | 268949 | 1.62983 | -2.45124 | 2.636175 | 0.015314 | 0.035216 | 0.493278 | -2.45124 | 0.988449 | 0.334007 | 0.430624 | 7.419807 | 5.32821 | 3.471878 | 12.98709 | 21.06593 | 21.64268 | 19.89765 | 30.8297 | 29.87805 | 2.307643 | 0 | 4.478048 | 25.86413 | 34.58764 | 35.63429 |
| Slc12a9 | 83704 | 1.630423 | 5.111355 | 29.76899 | 7.20E-19 | 2.32E-17 | -3.84597 | 5.111355 | -39.6078 | 1.80E-21 | 1.83E-19 | 2329.078 | 2415.455 | 2310.535 | 8059.789 | 7731.196 | 7516.775 | 658.0438 | 573.7005 | 481.4962 | 2579.176 | 2703.161 | 2386.053 | 1475.924 | 1348.918 | 1449.668 |
| Sp8 | 320145 | 1.630668 | -1.94041 | 2.739636 | 0.012167 | 0.028636 | 1.353113 | -1.94041 | 3.034534 | 0.006226 | 0.014197 | 2.967923 | 7.10428 | 4.339848 | 20.77935 | 8.426372 | 20.29002 | 38.37404 | 40.21265 | 47.11538 | 3.076858 | 7.625279 | 1.492683 | 71.75211 | 77.82218 | 69.64884 |
| Lcat | 16816 | 1.63118 | 0.330791 | 7.396704 | 2.56E-07 | 1.14E-06 | -0.43679 | 0.330791 | -2.2669 | 0.033899 | 0.062178 | 55.64856 | 50.61799 | 40.79457 | 140.2606 | 161.5055 | 190.7262 | 139.2836 | 103.2125 | 133.302 | 60.76794 | 53.37695 | 52.24389 | 40.04769 | 48.73712 | 39.68364 |
| Wnt5b | 22419 | 1.631469 | -2.17084 | 2.606226 | 0.016359 | 0.037287 | 0.536424 | -2.17084 | 1.062395 | 0.29993 | 0.393675 | 11.12971 | 2.664105 | 3.471878 | 15.58451 | 21.06593 | 14.87935 | 17.05513 | 25.46801 | 36.77298 | 9.230573 | 6.862751 | 2.985365 | 19.18952 | 31.4433 | 35.63429 |
| Cstdc4 | 433016 | 1.633732 | 0.401566 | 5.944232 | 6.28E-06 | 2.37E-05 | 0.122678 | 0.401566 | 0.560803 | 0.580772 | 0.674447 | 53.42261 | 44.40175 | 48.6063 | 129.8709 | 207.8505 | 162.3201 | 187.6064 | 205.0845 | 165.4784 | 32.30701 | 51.85189 | 76.87315 | 52.5626 | 31.4433 | 38.0639 |
| Gm20169 | 100504320 | 1.635634 | 1.650374 | 8.793935 | 1.53E-08 | 8.01E-08 | -0.4673 | 1.650374 | -2.87037 | 0.009063 | 0.019805 | 96.4575 | 115.4445 | 122.3837 | 396.1063 | 296.3274 | 413.9163 | 262.9333 | 340.4671 | 236.7261 | 128.4588 | 165.4685 | 135.8341 | 150.1788 | 130.4897 | 152.2556 |
| Tnfsf9 | 21950 | 1.636028 | 3.887836 | 23.17624 | 1.29E-16 | 2.59E-15 | 1.021555 | 3.887836 | 19.19007 | 6.06E-15 | 1.08E-13 | 667.7827 | 613.6322 | 637.0897 | 2132.48 | 2089.74 | 2157.505 | 4603.464 | 4556.094 | 4315.079 | 347.6849 | 335.5123 | 305.2536 | 290.3458 | 262.5516 | 255.919 |
| Frmpd4 | 333605 | 1.636334 | -1.66276 | 3.210896 | 0.004136 | 0.010714 | 0.786165 | -1.66276 | 2.037079 | 0.054233 | 0.092918 | 10.38773 | 5.32821 | 8.679696 | 20.77935 | 37.91867 | 25.70069 | 48.32287 | 46.91476 | 51.712 | 13.07665 | 3.050111 | 2.985365 | 42.55067 | 44.80671 | 29.9652 |
| Diaph2 | 54004 | 1.637388 | 6.937689 | 40.30292 | 1.25E-21 | 7.57E-20 | -1.09508 | 6.937689 | -27.179 | 4.79E-18 | 1.78E-16 | 5101.86 | 5289.136 | 4877.121 | 16662.44 | 17087.28 | 16951.63 | 8288.793 | 8758.316 | 7686.702 | 6099.101 | 6158.937 | 6368.53 | 4757.332 | 4771.521 | 4894.046 |
| Kdm6b | 216850 | 1.638483 | 6.261503 | 19.72944 | 1.66E-14 | 2.20E-13 | 1.592145 | 6.261503 | 26.48541 | 5.90E-17 | 1.72E-15 | 1962.539 | 1895.955 | 2024.973 | 6828.613 | 6232.706 | 6496.863 | 21600.32 | 21378.39 | 18687.57 | 2419.949 | 2448.477 | 2612.194 | 2945.174 | 3031.135 | 3351.243 |
| Gm10224 | 100043805 | 1.639768 | -1.42953 | 3.035444 | 0.006213 | 0.015574 | -0.28018 | -1.42953 | -0.60622 | 0.550763 | 0.645383 | 22.25942 | 15.09659 | 3.471878 | 36.36386 | 36.51428 | 39.22736 | 38.37404 | 38.87223 | 22.98311 | 29.99936 | 18.30067 | 20.89756 | 9.177596 | 15.72165 | 29.9652 |
| Mitd1 | 69028 | 1.640263 | 4.942379 | 24.37972 | 4.54E-17 | 1.00E-15 | 0.229675 | 4.942379 | 4.121183 | 0.000473 | 0.001372 | 1117.423 | 1093.171 | 1020.732 | 3662.36 | 3502.562 | 3598.096 | 4503.976 | 4352.349 | 4264.516 | 1469.969 | 1436.602 | 1282.214 | 976.9968 | 874.1239 | 825.2578 |
| Map1b | 17755 | 1.640486 | 2.124923 | 7.256412 | 3.44E-07 | 1.51E-06 | 3.864042 | 2.124923 | 27.99169 | 2.60E-18 | 1.04E-16 | 69.74619 | 60.38638 | 59.8899 | 205.196 | 179.7626 | 258.3595 | 3234.79 | 3305.48 | 3121.107 | 39.22994 | 58.71464 | 54.48291 | 488.9156 | 552.6161 | 608.2126 |
| Adamtsl3 | 269959 | 1.641399 | -1.41435 | 2.966791 | 0.007274 | 0.01797 | 0.829015 | -1.41435 | 1.991872 | 0.059334 | 0.100304 | 8.903769 | 12.43249 | 7.811726 | 35.06515 | 14.04395 | 66.28072 | 55.42917 | 49.5956 | 75.84427 | 3.076858 | 7.625279 | 10.44878 | 35.04173 | 47.16496 | 47.78235 |
| Selenow | 20364 | 1.641958 | 5.841438 | 31.71526 | 1.92E-19 | 7.09E-18 | 0.526834 | 5.841438 | 12.6725 | 2.11E-11 | 1.87E-10 | 1585.613 | 1595.799 | 1515.475 | 5219.512 | 5272.1 | 5165.838 | 7997.435 | 7996.956 | 7478.705 | 2269.183 | 2141.178 | 2192.751 | 3287.248 | 3353.428 | 3446.808 |
| Osm | 18413 | 1.64244 | 4.66686 | 17.78033 | 2.82E-14 | 3.57E-13 | 3.494029 | 4.66686 | 59.37689 | 3.44E-25 | 1.27E-22 | 348.731 | 413.8243 | 414.0215 | 1207.8 | 1379.116 | 1312.088 | 15397.94 | 15075.72 | 15041.3 | 883.0582 | 792.2664 | 791.1217 | 1285.698 | 1244.369 | 1292.553 |
| Mob3c | 100465 | 1.642535 | 5.042704 | 25.83527 | 1.37E-17 | 3.29E-16 | -0.32855 | 5.042704 | -5.83077 | 8.15E-06 | 3.06E-05 | 1150.07 | 1109.156 | 1033.752 | 3823.4 | 3502.562 | 3663.024 | 3099.77 | 3084.31 | 2917.706 | 1885.345 | 2084.751 | 2093.487 | 1180.573 | 1248.299 | 1191.319 |
| Tenm4 | 23966 | 1.642929 | 0.515236 | 5.213757 | 3.45E-05 | 0.000119 | 0.986814 | 0.515236 | 4.36291 | 0.000264 | 0.0008 | 32.64715 | 32.85729 | 29.51097 | 115.5851 | 85.66811 | 120.3874 | 251.5632 | 155.4889 | 263.1566 | 41.53758 | 40.41398 | 32.09267 | 137.6639 | 169.7938 | 173.3122 |
| Foxp4 | 74123 | 1.64313 | 5.164123 | 18.75271 | 9.66E-15 | 1.33E-13 | -0.12269 | 5.164123 | -1.62001 | 0.119913 | 0.181976 | 1109.261 | 1186.415 | 1246.404 | 3905.218 | 3891.579 | 3976.843 | 3982.373 | 4061.478 | 3296.927 | 1612.273 | 1305.448 | 1535.224 | 1618.594 | 1863.016 | 1789.003 |
| Fmn2 | 54418 | 1.644292 | -1.46941 | 2.787033 | 0.01094 | 0.026006 | 1.448766 | -1.46941 | 3.387691 | 0.002731 | 0.006801 | 4.451884 | 15.09659 | 3.471878 | 22.07806 | 21.06593 | 20.29002 | 44.05909 | 61.6594 | 79.29174 | 5.384501 | 17.53814 | 7.463413 | 49.22529 | 66.81702 | 55.88105 |
| A830018L1 | 320492 | 1.644592 | -1.6064 | 2.809309 | 0.010405 | 0.024855 | 1.623447 | -1.6064 | 3.886369 | 0.000832 | 0.002317 | 4.451884 | 5.32821 | 9.547666 | 16.88322 | 25.27911 | 21.64268 | 62.53548 | 48.25518 | 98.82738 | 9.999788 | 5.337695 | 10.44878 | 27.53279 | 51.88145 | 29.9652 |
| Pi4k2b | 67073 | 1.644826 | 4.76095 | 27.3704 | 4.14E-18 | 1.14E-16 | 0.940624 | 4.76095 | 20.49046 | 1.60E-15 | 3.28E-14 | 769.434 | 770.8144 | 768.1531 | 2593.522 | 2588.3 | 2528.136 | 5198.972 | 5111.028 | 5059.732 | 1106.9 | 1209.369 | 1218.775 | 935.2805 | 944.0852 | 928.9212 |
| Kcnab1 | 16497 | 1.645071 | 2.402911 | 15.49976 | 4.31E-13 | 4.46E-12 | -0.01733 | 2.402911 | -0.201 | 0.842602 | 0.906762 | 208.4966 | 200.6959 | 190.0853 | 706.4978 | 665.6834 | 634.4012 | 693.5753 | 679.5938 | 684.8967 | 194.6113 | 169.2812 | 174.6439 | 180.2146 | 199.665 | 193.559 |
| Siah2 | 20439 | 1.645145 | 4.73599 | 28.9031 | 1.33E-18 | 4.07E-17 | -2.61001 | 4.73599 | -34.5943 | 3.10E-20 | 2.22E-18 | 1561.127 | 1615.336 | 1721.184 | 5670.164 | 5353.555 | 5294.342 | 952.2448 | 845.8061 | 973.3348 | 2101.494 | 1938.346 | 2173.346 | 673.3018 | 576.9846 | 570.1487 |
| Dhx58 | 80861 | 1.645904 | 7.238832 | 40.61527 | 1.06E-21 | 6.55E-20 | -0.27312 | 7.238832 | -7.76062 | 1.20E-07 | 5.81E-07 | 6701.57 | 6901.808 | 6472.449 | 22485.85 | 22232.98 | 22328.49 | 20093.79 | 19501.8 | 18227.91 | 8472.897 | 8383.231 | 8116.461 | 2569.727 | 2609.794 | 2337.286 |
| Trim12c | 319236 | 1.648513 | 6.233507 | 32.353 | 1.26E-19 | 4.87E-18 | -0.89979 | 6.233507 | -18.3644 | 1.47E-14 | 2.43E-13 | 3321.106 | 3285.729 | 2920.718 | 10690.97 | 10959.9 | 10257.28 | 5812.957 | 5917.962 | 5992.846 | 4300.678 | 4190.091 | 3857.092 | 2412.039 | 2156.225 | 2180.171 |
| Spdye4a | 74673 | 1.648863 | -2.90438 | 2.455737 | 0.022704 | 0.049887 | 1.105557 | -2.90438 | 2.090829 | 0.048681 | 0.084635 | 4.451884 | 1.77607 | 3.471878 | 12.98709 | 8.426372 | 12.17401 | 29.84648 | 16.08506 | 33.32551 | 0.769214 | 3.812639 | 5.97073 | 9.177596 | 12.57732 | 7.288832 |
| Slc5a1 | 20537 | 1.648935 | -1.75457 | 3.441033 | 0.002408 | 0.006503 | 0.337122 | -1.75457 | 0.881819 | 0.387698 | 0.486421 | 5.935846 | 11.54445 | 13.01954 | 16.88322 | 42.13186 | 48.69604 | 39.7953 | 44.23392 | 43.66791 | 4.615287 | 2.287584 | 4.478048 | 33.37308 | 25.94073 | 29.9652 |
| Lama2 | 16773 | 1.649122 | -0.83148 | 3.314226 | 0.003247 | 0.008572 | 1.282629 | -0.83148 | 3.597228 | 0.001662 | 0.004337 | 6.677827 | 15.98463 | 10.41564 | 24.67547 | 49.15383 | 35.16936 | 66.79926 | 91.14868 | 109.1698 | 9.999788 | 13.7255 | 10.44878 | 143.5042 | 94.32991 | 93.13508 |

| | | | | | | | | | | | | | | | | | | | | | | | | | | |
|---|---|---|---|---|---|---|---|---|---|---|---|---|---|---|---|---|---|---|---|---|---|---|---|---|---|---|
| Gm15708 | 100504231 | 1.6497 | 1.72541 | 9.705418 | 2.79E-09 | 1.63E-08 | -1.74257 | 1.72541 | -8.9954 | 1.04E-08 | 5.85E-08 | 126.1367 | 150.9659 | 133.6673 | 476.6263 | 483.112 | 411.211 | 150.6537 | 166.2123 | 116.0647 | 243.841 | 328.6495 | 248.5316 | 124.3147 | 152.5 | 99.61404 |
| Mllt6 | 246198 | 1.649952 | 7.341779 | 44.48815 | 1.55E-22 | 1.18E-20 | -1.21341 | 7.341779 | -32.4071 | 1.22E-19 | 7.28E-18 | 7717.342 | 8094.439 | 7705.834 | 26613.15 | 26273.43 | 25823.78 | 12126.2 | 12369.41 | 10922.72 | 12849.73 | 13143.69 | 13270.69 | 2755.782 | 2587.784 | 2764.087 |
| Casp1 | 12362 | 1.650145 | 6.46173 | 40.95313 | 8.92E-22 | 5.68E-20 | -0.19989 | 6.46173 | -5.71773 | 1.06E-05 | 3.91E-05 | 3778.908 | 3757.276 | 3747.025 | 12767.61 | 12942.91 | 12128.02 | 11608.86 | 11460.61 | 11151.41 | 5675.264 | 5117.324 | 5327.384 | 1364.125 | 1410.232 | 1418.083 |
| Fam221a | 231946 | 1.650874 | 1.857333 | 5.92702 | 6.53E-06 | 2.47E-05 | 1.048918 | 1.857333 | 5.333773 | 2.60E-05 | 9.10E-05 | 69.00421 | 68.37869 | 38.19066 | 201.2999 | 162.9099 | 213.7215 | 386.583 | 415.5307 | 432.0825 | 171.5348 | 98.36609 | 155.9853 | 689.154 | 775.0775 | 836.596 |
| Trim43c | 666731 | 1.651079 | 1.111604 | 6.482381 | 1.86E-06 | 7.45E-06 | 1.733133 | 1.111604 | 10.20673 | 1.14E-09 | 7.52E-09 | 58.61648 | 87.91546 | 39.9266 | 193.5077 | 207.8505 | 193.4315 | 628.1973 | 794.8701 | 646.9746 | 89.22888 | 83.87806 | 67.91705 | 40.88202 | 61.31444 | 65.59949 |
| Kcng3 | 225030 | 1.652399 | -0.13678 | 4.342786 | 0.000277 | 0.00086 | 2.252554 | -0.13678 | 9.084329 | 8.78E-09 | 5.02E-08 | 19.2915 | 14.20856 | 14.75548 | 54.54578 | 74.43295 | 40.58003 | 245.8781 | 256.0206 | 314.8686 | 29.99936 | 35.07628 | 48.51218 | 58.40288 | 43.23454 | 52.64157 |
| Htr4 | 15562 | 1.652961 | -2.30292 | 2.505219 | 0.0204 | 0.045359 | 1.736377 | -2.30292 | 3.570839 | 0.001769 | 0.004587 | 3.709904 | 4.440175 | 2.603909 | 15.58451 | 18.25714 | 6.763339 | 29.84648 | 40.21265 | 68.94934 | 4.615287 | 6.100223 | 4.478048 | 15.85221 | 18.0799 | 30.77507 |
| Wt1 | 22431 | 1.653497 | -2.45268 | 2.521736 | 0.019681 | 0.043934 | 0.980839 | -2.45268 | 1.940174 | 0.065685 | 0.109155 | 5.935846 | 4.440175 | 3.471878 | 19.48064 | 14.04395 | 14.87935 | 31.26774 | 24.12759 | 47.11538 | 0.769214 | 0 | 6.717071 | 33.37308 | 40.09021 | 41.30338 |
| Nrk | 27206 | 1.653587 | -2.14887 | 2.519604 | 0.019773 | 0.04412 | 1.554388 | -2.14887 | 3.235677 | 0.003903 | 0.009369 | 6.677827 | 1.77607 | 6.075787 | 20.77935 | 11.23516 | 14.87935 | 54.00791 | 30.8297 | 63.20356 | 2.307643 | 1.525056 | 8.209754 | 42.55067 | 30.65722 | 33.20468 |
| Ranbp2 | 19386 | 1.654725 | 9.337861 | 80.44762 | 5.41E-28 | 3.03E-25 | -0.68398 | 9.337861 | -36.7766 | 8.59E-21 | 7.08E-19 | 28404.51 | 29953.42 | 28808.78 | 97607.08 | 99210.69 | 95816.22 | 63780.5 | 63442.16 | 62030.27 | 30205.51 | 30489.68 | 29508.84 | 16083.32 | 15985.78 | 16272.72 |
| Axdnd1 | 77352 | 1.654933 | -2.30412 | 3.06161 | 0.005849 | 0.014736 | 0.086063 | -2.30412 | 0.187254 | 0.85323 | 0.91554 | 5.935846 | 5.32821 | 4.339848 | 15.58451 | 23.87472 | 16.23201 | 15.63387 | 16.08506 | 29.87805 | 16.1535 | 8.387806 | 7.463413 | 8.343269 | 11.00516 | 15.38754 |
| Arg2 | 11847 | 1.655091 | 2.527835 | 10.03236 | 1.55E-09 | 9.40E-09 | 1.812129 | 2.527835 | 16.55681 | 1.17E-13 | 1.59E-12 | 97.19948 | 111.0044 | 103.2884 | 319.4825 | 405.8702 | 327.3456 | 1293.347 | 1201.018 | 1313.485 | 286.917 | 256.9719 | 244.7999 | 400.4769 | 364.7423 | 441.3793 |
| Phldb2 | 208177 | 1.655343 | -0.25823 | 3.572413 | 0.001763 | 0.004871 | 3.507231 | -0.25823 | 12.10355 | 5.03E-11 | 4.13E-10 | 8.903769 | 7.10428 | 19.09533 | 37.66257 | 47.74944 | 27.05335 | 451.961 | 440.9988 | 424.0384 | 16.92272 | 5.337695 | 14.92683 | 155.1848 | 139.9227 | 153.0655 |
| Plekha7 | 233765 | 1.655866 | -0.09454 | 3.776604 | 0.001082 | 0.0031 | 1.06118 | -0.09454 | 3.364472 | 0.002885 | 0.007143 | 15.5816 | 19.53677 | 18.22736 | 67.53288 | 87.07251 | 36.52203 | 105.1733 | 123.3188 | 163.1801 | 17.69193 | 46.5142 | 20.89756 | 146.0072 | 169.7938 | 129.5792 |
| Dmp1 | 13406 | 1.655908 | -2.67184 | 2.567141 | 0.017824 | 0.040233 | 1.08028 | -2.67184 | 2.168041 | 0.041599 | 0.07402 | 2.967923 | 5.32821 | 4.339848 | 15.58451 | 14.04395 | 13.52668 | 18.47639 | 38.87223 | 43.66791 | 1.538429 | 3.050111 | 0 | 21.6925 | 26.72681 | 24.29611 |
| Klrg2 | 74253 | 1.657295 | 1.059997 | 7.579711 | 1.74E-07 | 7.95E-07 | -0.37677 | 1.059997 | -2.00766 | 0.057506 | 0.097623 | 85.32779 | 49.72996 | 72.90945 | 189.6115 | 259.8131 | 255.6542 | 170.5513 | 164.8719 | 218.3396 | 98.45945 | 73.9652 | 88.81461 | 123.4804 | 103.7629 | 128.7694 |
| Cdyl2 | 75796 | 1.658366 | 3.974833 | 21.72782 | 4.84E-16 | 8.59E-15 | 0.091475 | 3.974833 | 1.447288 | 0.162346 | 0.235397 | 641.0714 | 690.8912 | 595.4271 | 2254.559 | 2020.925 | 2208.906 | 2373.506 | 2524.014 | 2289.118 | 606.141 | 569.6083 | 604.5364 | 393.8023 | 356.0954 | 366.8712 |
| Spata25 | 75642 | 1.658482 | -0.50963 | 4.993803 | 5.81E-05 | 0.000196 | -3.19411 | -0.50963 | -6.13502 | 4.06E-06 | 1.59E-05 | 35.61508 | 44.40175 | 21.69924 | 128.5722 | 115.1604 | 90.62874 | 8.527565 | 9.382952 | 19.53565 | 96.92102 | 136.4925 | 87.32193 | 27.53279 | 42.44846 | 17.00728 |
| Cacnb2 | 12296 | 1.659714 | 2.214933 | 8.152327 | 5.39E-08 | 2.64E-07 | 1.440676 | 2.214933 | 10.40571 | 8.08E-10 | 5.45E-09 | 73.45609 | 70.15476 | 110.2321 | 264.9367 | 268.2395 | 305.7029 | 825.7526 | 817.6573 | 732.0121 | 122.3051 | 163.181 | 129.117 | 705.0062 | 593.4924 | 654.3752 |
| 4930503L1 | 269033 | 1.660411 | 5.152835 | 35.92697 | 1.40E-20 | 6.87E-19 | -2.52983 | 5.152835 | -42.9011 | 3.34E-22 | 4.58E-20 | 2410.695 | 2669.433 | 2491.073 | 8374.077 | 8891.226 | 8240.452 | 1574.757 | 1562.932 | 1454.831 | 2394.565 | 2280.721 | 2218.126 | 586.5318 | 587.9898 | 556.3809 |
| 4932438A1 | 229227 | 1.660807 | 8.583193 | 76.33492 | 1.65E-27 | 6.30E-25 | -1.48691 | 8.583193 | -63.9383 | 7.14E-26 | 4.00E-23 | 21848.37 | 22285.24 | 20834.74 | 73825.12 | 73844.51 | 71638.64 | 27403.33 | 27482.67 | 26444.37 | 17353.48 | 17418.42 | 17442.74 | 8539.336 | 8507.772 | 8398.355 |
| Ctnnal1 | 54366 | 1.660911 | 0.662108 | 5.953437 | 6.15E-06 | 2.33E-05 | 1.194613 | 0.662108 | 6.126947 | 4.14E-06 | 1.62E-05 | 35.61508 | 35.5214 | 48.6063 | 140.2606 | 176.9538 | 96.03941 | 314.0986 | 272.1056 | 366.5806 | 50.76815 | 55.66453 | 40.30243 | 87.60433 | 84.11084 | 102.8535 |
| Serpinh1 | 12406 | 1.661487 | -1.24219 | 3.003674 | 0.006684 | 0.016644 | 2.406589 | -1.24219 | 6.452514 | 1.99E-06 | 8.13E-06 | 3.709904 | 11.54445 | 5.207818 | 14.2858 | 26.68351 | 24.34802 | 129.3347 | 91.14868 | 136.7495 | 8.461359 | 6.100223 | 13.43414 | 62.57452 | 73.10568 | 63.97975 |
| Astn2 | 56079 | 1.66547 | -2.17355 | 3.356258 | 0.002941 | 0.007812 | 0.908873 | -2.17355 | 2.411569 | 0.024963 | 0.047746 | 4.451884 | 5.32821 | 9.547666 | 19.48064 | 28.08791 | 17.58468 | 36.95278 | 41.55307 | 47.11538 | 2.307643 | 0.762528 | 2.239024 | 30.8701 | 29.87114 | 39.68364 |
| Atp13a4 | 224079 | 1.665726 | 0.569308 | 3.216749 | 0.00408 | 0.010577 | 6.073054 | 0.569308 | 19.28801 | 5.47E-15 | 9.87E-14 | 5.935846 | 14.20856 | 6.943757 | 24.67547 | 30.8967 | 31.11136 | 2129.049 | 2069.611 | 1966.205 | 13.07665 | 13.7255 | 4.478048 | 1129.679 | 1042.346 | 1012.338 |
| Slc26a5 | 80979 | 1.666291 | -2.03863 | 2.71415 | 0.01288 | 0.030138 | 0.886959 | -2.03863 | 1.872062 | 0.074966 | 0.122206 | 2.225942 | 14.20856 | 5.207818 | 14.2858 | 15.44835 | 31.11136 | 25.5827 | 38.87223 | 48.26453 | 6.92293 | 6.862751 | 3.731706 | 23.36115 | 19.65207 | 31.58494 |
| Sap30 | 60406 | 1.666472 | 4.150246 | 30.40071 | 4.64E-19 | 1.56E-17 | -2.07346 | 4.150246 | -32.0707 | 1.52E-19 | 8.70E-18 | 1406.054 | 1325.836 | 1270.708 | 4413.014 | 4609.225 | 4572.017 | 1198.123 | 1080.38 | 1082.505 | 1096.9 | 1073.639 | 1193.4 | 196.0668 | 174.5103 | 162.7839 |
| Magohb | 66441 | 1.666961 | 2.444746 | 17.35077 | 4.60E-14 | 5.62E-13 | -2.79591 | 2.444746 | -19.7076 | 3.54E-15 | 6.70E-14 | 595.8105 | 528.3808 | 511.2341 | 2044.168 | 1717.575 | 1804.459 | 196.134 | 320.3608 | 321.7636 | 193.0728 | 190.632 | 217.9316 | 64.24317 | 61.31444 | 72.07845 |
| Plekhf2 | 71801 | 1.667396 | 6.457485 | 46.13044 | 7.22E-23 | 5.97E-21 | -1.41223 | 6.457485 | -37.5449 | 5.56E-21 | 4.84E-19 | 3595.639 | 3825.655 | 3797.367 | 12513.06 | 12847.41 | 12625.8 | 4936.039 | 4755.816 | 5097.654 | 4634.517 | 4441.725 | 4494.467 | 3770.323 | 3685.941 | 3917.342 |
| Fat4 | 329628 | 1.668391 | 0.755771 | 5.789678 | 8.96E-06 | 3.33E-05 | 2.712008 | 0.755771 | 14.91544 | 9.16E-13 | 1.03E-11 | 32.64715 | 56.83424 | 32.11488 | 135.0658 | 112.3516 | 150.1461 | 875.4967 | 1010.678 | 833.1378 | 10.769 | 24.40089 | 19.40487 | 116.8058 | 137.5645 | 123.1003 |
| Wnk2 | 75607 | 1.669346 | 0.821751 | 4.798276 | 9.28E-05 | 0.000305 | 1.441315 | 0.821751 | 6.065834 | 4.75E-06 | 1.85E-05 | 26.71131 | 23.08891 | 30.37894 | 96.10448 | 81.45493 | 96.03941 | 292.7797 | 257.361 | 233.2786 | 206.9187 | 144.8803 | 151.5073 | 102.6222 | 165.8634 | 108.5226 |
| Atxn7l1os1 | 100504517 | 1.669902 | 1.567871 | 9.605156 | 3.34E-09 | 1.93E-08 | -0.71866 | 1.567871 | -4.54619 | 0.00017 | 0.000531 | 139.4924 | 153.6301 | 113.704 | 506.4966 | 428.3406 | 440.9697 | 287.0947 | 319.0204 | 266.6041 | 88.45966 | 140.3051 | 67.17071 | 98.45058 | 101.4047 | 107.7127 |
| Pde10a | 23984 | 1.670334 | 2.070633 | 14.36657 | 1.90E-12 | 1.75E-11 | -1.38621 | 2.070633 | -11.3697 | 1.61E-10 | 1.22E-09 | 211.4645 | 190.9275 | 185.7455 | 685.7184 | 620.7427 | 700.6819 | 228.823 | 278.8077 | 286.1397 | 219.9953 | 195.9697 | 202.2585 | 130.155 | 155.6444 | 133.6286 |
| Nyap2 | 241134 | 1.67124 | -1.23085 | 2.747109 | 0.011965 | 0.028214 | 0.443856 | -1.23085 | 0.909563 | 0.37321 | 0.471325 | 5.935846 | 7.10428 | 3.471878 | 18.18193 | 14.04395 | 25.70069 | 29.84648 | 40.21265 | 17.23733 | 129.9972 | 89.21576 | 85.82924 | 42.55067 | 32.22939 | 28.34546 |
| Rhoq | 104215 | 1.671459 | 6.933146 | 46.04293 | 7.52E-23 | 6.14E-21 | -1.88282 | 6.933146 | -45.9795 | 7.74E-23 | 1.40E-20 | 4930.462 | 4912.61 | 4844.138 | 16857.24 | 16960.88 | 16160.32 | 4681.633 | 4990.39 | 4436.89 | 9114.422 | 8919.288 | 8972.515 | 5851.969 | 6189.615 | 6051.351 |
| Tek | 21687 | 1.673837 | -1.35435 | 3.524872 | 0.001974 | 0.005407 | 0.582556 | -1.35435 | 1.589233 | 0.126712 | 0.190667 | 8.903769 | 8.88035 | 7.811726 | 22.07806 | 30.8967 | 39.22736 | 34.11026 | 44.23392 | 63.20356 | 10.769 | 8.387806 | 18.65853 | 45.88798 | 51.09537 | 37.25403 |
| Rps15a-ps | 100042335 | 1.675394 | -0.41053 | 5.737615 | 1.01E-05 | 3.73E-05 | -0.12764 | -0.41053 | -0.52813 | 0.60286 | 0.69534 | 23.74338 | 27.52908 | 34.71878 | 85.7148 | 80.05053 | 131.2088 | 99.48826 | 89.80826 | 87.33582 | 23.07643 | 25.92595 | 26.86829 | 31.70442 | 33.01547 | 19.43689 |
| Slc4a10 | 94229 | 1.675821 | -2.05751 | 2.6304 | 0.01551 | 0.0356 | 0.630378 | -2.05751 | 1.245549 | 0.226431 | 0.311277 | 5.193865 | 6.216245 | 4.339848 | 23.37676 | 19.66153 | 13.52668 | 14.21261 | 30.8297 | 54.01031 | 1.538429 | 9.912862 | 4.478048 | 40.04769 | 41.66238 | 43.73299 |
| Rims2 | 116838 | 1.676034 | -0.25892 | 5.706973 | 1.09E-05 | 3.99E-05 | -0.31168 | -0.25892 | -1.24675 | 0.225998 | 0.310748 | 26.71131 | 29.30515 | 22.56721 | 81.81868 | 126.3956 | 70.33872 | 82.43313 | 72.38277 | 71.24765 | 39.22994 | 38.12639 | 29.85365 | 48.39096 | 51.88145 | 42.11325 |
| Gsx2 | 14843 | 1.676496 | -3.21592 | 2.461987 | 0.0224 | 0.049293 | 1.079807 | -3.21592 | 1.925755 | 0.067561 | 0.111812 | 2.967923 | 1.77607 | 0.86797 | 5.194837 | 8.426372 | 6.763339 | 15.63387 | 13.40422 | 17.23733 | 3.076858 | 2.287584 | 0.746341 | 22.52683 | 23.58248 | 14.57766 |
| Tstd1 | 226654 | 1.677493 | 0.530786 | 6.73918 | 1.05E-06 | 4.35E-06 | 0.65123 | 0.530786 | 3.546962 | 0.001873 | 0.004833 | 39.32498 | 42.62568 | 50.34224 | 148.0528 | 144.6527 | 156.9095 | 251.5632 | 253.3397 | 232.1294 | 68.46009 | 30.50111 | 48.51218 | 70.91779 | 58.9562 | 56.69092 |
| Stat1 | 20846 | 1.677809 | 9.471465 | 33.49857 | 1.04E-12 | 1.00E-11 | -0.10574 | 9.471465 | -2.61522 | 0.023486 | 0.045263 | 25121.98 | 25959.04 | 24018.45 | 84983.63 | 87944.64 | 83605.69 | 85864.05 | 84961.29 | 77857.59 | 34921.57 | 34803.3 | 33486.84 | 22080.46 | 23471.64 | 22203.4 |
| Disp2 | 214240 | 1.678872 | -1.24271 | 3.12221 | 0.005084 | 0.012948 | 0.746575 | -1.24271 | 1.853066 | 0.077751 | 0.126074 | 14.09763 | 4.440175 | 14.75548 | 31.16902 | 30.8967 | 44.63804 | 45.48035 | 48.25518 | 96.52907 | 3.076858 | 12.96297 | 2.239024 | 97.61625 | 91.97167 | 92.32521 |
| Gm8221 | 666661 | 1.679615 | 0.696221 | 4.458741 | 0.00021 | 0.00066 | 2.260351 | 0.696221 | 9.233213 | 6.64E-09 | 3.87E-08 | 14.83961 | 28.41712 | 11.2836 | 48.05224 | 53.36702 | 75.74939 | 279.9884 | 321.7012 | 273.499 | 187.6883 | 227.2333 | 203.0048 | 130.155 | 145.4253 | 144.9668 |
| Ascc3 | 77987 | 1.681443 | 8.027425 | 54.79982 | 1.88E-24 | 2.67E-22 | -0.62352 | 8.027425 | -22.4763 | 2.42E-16 | 6.11E-15 | 11084.45 | 11267.39 | 11237.6 | 38246.98 | 38622.27 | 38124.94 | 26353.02 | 26628.82 | 24719.49 | 12558.2 | 12622.12 | 12060.13 | 6671.278 | 6867.218 | 6538.083 |
| Slc45a1 | 242773 | 1.6836 | -2.07588 | 2.896175 | 0.008547 | 0.020817 | -0.07933 | -2.07588 | -0.15983 | 0.874516 | 0.918721 | 8.161788 | 3.55214 | 7.811726 | 22.07806 | 22.47032 | 21.64268 | 17.05513 | 18.7659 | 29.87805 | 3.076858 | 14.48803 | 5.97073 | 33.37308 | 30.65722 | 15.38754 |
| Emx2os | 329078 | 1.68363 | -2.23971 | 2.682438 | 0.013822 | 0.032095 | 1.697764 | -2.23971 | 3.739093 | 0.001184 | 0.003193 | 2.967923 | 5.32821 | 4.339848 | 10.38967 | 25.27911 | 12.17401 | 52.58665 | 57.63814 | 44.81707 | 0.769214 | 4.575167 | 2.239024 | 50.05962 | 34.58764 | 26.72572 |
| Ttc39b | 69863 | 1.684511 | 5.738186 | 30.99358 | 3.10E-19 | 1.09E-17 | 0.17739 | 5.738186 | 3.933337 | 0.000743 | 0.002086 | 1566.321 | 1464.37 | 1516.343 | 5314.318 | 5246.821 | 5065.741 | 6051.729 | 6094.898 | 6180.159 | 2100.725 | 1942.921 | 1971.087 | 3439.93 | 3205.645 | 3630.648 |
| Dnah5 | 110082 | 1.685445 | -0.20334 | 4.646163 | 0.000134 | 0.00043 | 0.356388 | -0.20334 | 1.268174 | 0.218395 | 0.301888 | 22.25942 | 23.97694 | 31.24691 | 97.40319 | 84.26372 | 82.51273 | 110.8583 | 73.7232 | 181.5666 | 13.84586 | 16.01308 | 8.956095 | 124.3147 | 114.7681 | 93.13508 |
| Pnpt1 | 71701 | 1.685451 | 6.532456 | 44.72315 | 1.39E-22 | 1.07E-20 | -0.48053 | 6.532456 | -14.2295 | 2.29E-12 | 2.41E-11 | 4198.869 | 4396.661 | 4243.503 | 14762.43 | 14998.94 | 14297.7 | 10920.97 | 11283.67 | 10611.3 | 4101.451 | 4215.254 | 4003.374 | 1802.146 | 1790.696 | 1923.442 |
| Maff | 17133 | 1.686095 | 3.324386 | 12.41779 | 3.10E-11 | 2.38E-10 | 2.244155 | 3.324386 | 25.27501 | 2.16E-17 | 6.93E-16 | 188.4631 | 230.8891 | 191.8213 | 628.5752 | 745.7339 | 718.2666 | 3649.798 | 3519.948 | 3145.239 | 223.8414 | 215.7954 | 228.3804 | 702.5033 | 683.1058 | 745.8905 |
| Nr4a1 | 15370 | 1.687212 | 2.649428 | 13.52506 | 6.11E-12 | 5.23E-11 | 0.573787 | 2.649428 | 6.162862 | 3.81E-06 | 1.50E-05 | 234.4659 | 226.4489 | 210.0486 | 798.7061 | 800.5053 | 715.5612 | 1286.241 | 1218.443 | 1095.145 | 143.8431 | 194.4446 | 207.4829 | 186.8892 | 203.5954 | 206.5169 |
| Mreg | 381269 | 1.68813 | 1.875383 | 9.788978 | 2.40E-09 | 1.42E-08 | 0.289625 | 1.875383 | 2.193575 | 0.039471 | 0.07077 | 120.9429 | 103.9001 | 111.1001 | 398.7037 | 362.334 | 400.3897 | 451.961 | 552.2538 | 475.7504 | 152.3045 | 120.4794 | 174.6439 | 130.155 | 170.5799 | 184.6504 |
| Pxdn | 69675 | 1.688388 | -1.78896 | 3.309716 | 0.003281 | 0.008657 | 1.142552 | -1.78896 | 3.080937 | 0.005594 | 0.012906 | 11.87169 | 7.992315 | 3.471878 | 29.87031 | 28.08791 | 18.93735 | 45.48035 | 68.36151 | 63.20356 | 2.307643 | 5.337695 | 2.239024 | 39.21337 | 46.37887 | 34.82442 |
| Degs2 | 70059 | 1.688441 | 3.959186 | 18.26263 | 1.65E-14 | 2.19E-13 | -0.73691 | 3.959186 | -8.64401 | 2.04E-08 | 1.11E-07 | 657.3949 | 687.3391 | 583.2756 | 2305.209 | 2113.615 | 2206.201 | 1435.473 | 1584.379 | 1172.139 | 773.8297 | 758.7152 | 729.1754 | 472.229 | 474.0078 | 459.1964 |
| Myo1h | 231646 | 1.688615 | 0.014637 | 5.704696 | 1.09E-05 | 4.01E-05 | -0.0173 | 0.014637 | -0.07228 | 0.943054 | 0.952909 | 36.35706 | 36.40943 | 23.43518 | 115.5851 | 99.71206 | 113.6241 | 106.5946 | 111.255 | 118.363 | 27.69172 | 28.97606 | 36.57072 | 60.07154 | 95.90208 | 46.97248 |
| Acvr1c | 269275 | 1.688653 | -1.14162 | 2.464879 | 0.022261 | 0.049023 | 2.30432 | -1.14162 | 5.108543 | 4.43E-05 | 0.000151 | 16.32358 | 0.888035 | 19.9633 | 40.25998 | 26.68351 | 17.58468 | 130.756 | 139.4039 | 149.3902 | 7.692145 | 8.387806 | 5.97073 | 48.39096 | 78.60826 | 69.64884 |
| Psmb9 | 16912 | 1.689999 | 5.822815 | 28.1672 | 2.28E-18 | 6.64E-17 | 0.436863 | 5.822815 | 9.055942 | 9.27E-09 | 5.28E-08 | 1769.624 | 1800.935 | 1724.656 | 6054.582 | 6059.966 | 6128.938 | 8769.179 | 9110.847 | 7895.848 | 2818.402 | 2821.353 | 2961.482 | 1654.47 | 1655.49 | 1729.073 |
| Pfkp | 56421 | 1.692337 | 6.98115 | 31.79302 | 1.82E-19 | 6.77E-18 | -0.31587 | 6.98115 | -6.81083 | 9.00E-07 | 3.84E-06 | 4780.582 | 5030.718 | 5045.507 | 17432.57 | 17538.09 | 16233.37 | 15069.63 | 15031.49 | 12914.21 | 6029.872 | 6223.752 | 6096.862 | 3245.532 | 3362.075 | 3365.011 |
| Tbc1d1 | 57915 | 1.693201 | 6.798223 | 42.77094 | 3.57E-22 | 2.44E-20 | -1.62354 | 6.798223 | -38.1518 | 3.97E-21 | 3.69E-19 | 5472.108 | 5574.196 | 5620.103 | 19467.65 | 19990.16 | 18105.46 | 6789.363 | 6661.896 | 6016.979 | 5652.957 | 5668.632 | 5810.267 | 3231.348 | 3351.856 | 3332.616 |
| Cpeb4 | 67579 | 1.694441 | 7.485879 | 45.33786 | 1.04E-22 | 8.27E-21 | -0.14063 | 7.485879 | -4.44407 | 0.000217 | 0.000668 | 5856.454 | 5821.957 | 5728.599 | 19841.68 | 20782.24 | 19594.74 | 19506.81 | 19253.82 | 18075.07 | 8324.439 | 8138.46 | 8125.417 | 7662.458 | 7708.326 | 8081.695 |
| Bcl9 | 77578 | 1.694576 | 5.567435 | 31.53664 | 2.16E-19 | 7.89E-18 | -1.07704 | 5.567435 | -20.43 | 1.70E-15 | 3.46E-14 | 1964.765 | 2073.562 | 1964.215 | 7042.9 | 7219.996 | 6481.984 | 3456.506 | 3513.245 | 3249.812 | 2782.249 | 2888.455 | 2655.482 | 1331.586 | 1389.008 | 1486.922 |
| Rnf135 | 71956 | 1.696854 | 4.971028 | 26.08841 | 1.12E-17 | 2.76E-16 | -0.33378 | 4.971028 | -5.82414 | 8.28E-06 | 3.10E-05 | 1125.585 | 1288.539 | 1167.419 | 4133.791 | 4069.937 | 4140.516 | 3641.27 | 3360.437 | 3209.592 | 1579.966 | 1489.217 | 1572.541 | 862.694 | 846.611 | 988.0417 |
| Plagl2 | 54711 | 1.697222 | 8.019354 | 48.26812 | 2.77E-23 | 2.72E-21 | -0.88279 | 8.019354 | -26.6677 | 7.10E-18 | 2.53E-16 | 13081.12 | 12838.32 | 11978.85 | 43952.21 | 44678.03 | 42725.36 | 24910.44 | 25965.31 | 23333.6 | 13625.1 | 13302.3 | 13420.71 | 5092.732 | 4703.918 | 4863.271 |
| Gm20038 | 100504058 | 1.698652 | -0.40582 | 3.364274 | 0.002886 | 0.007684 | 1.139403 | -0.40582 | 3.141455 | 0.004862 | 0.011395 | 10.38773 | 3.55214 | 15.62345 | 25.97418 | 36.51428 | 29.75869 | 52.58665 | 91.14868 | 71.24765 | 56.92187 | 57.18959 | 49.25852 | 185.2206 | 187.0877 | 156.305 |
| Cebpb | 12608 | 1.698661 | 8.115687 | 28.83002 | 1.02E-13 | 1.18E-12 | -0.08254 | 8.115687 | -1.6938 | 0.112771 | 0.172873 | 9785.242 | 10298.54 | 9880.966 | 34253.45 | 35222.23 | 34279.31 | 36388.54 | 34883.14 | 31188.08 | 14865.84 | 15540.32 | 15032.06 | 6674.615 | 6882.939 | 7556.899 |
| Stim2 | 116873 | 1.698818 | 6.070157 | 33.31824 | 6.83E-20 | 2.77E-18 | -0.46079 | 6.070157 | -10.0879 | 1.41E-09 | 9.15E-09 | 2550.93 | 2749.356 | 2666.403 | 9485.772 | 9101.886 | 8965.482 | 7281.119 | 6946.065 | 6621.435 | 2421.487 | 2373.749 | 2577.116 | 2476.282 | 2362.964 | 2709.826 |
| Il31ra | 218624 | 1.699052 | -0.56513 | 2.79652 | 0.010709 | 0.025504 | 0.799498 | -0.56513 | 1.707586 | 0.10222 | 0.158979 | 8.903769 | 3.55214 | 4.339848 | 15.58451 | 19.66153 | 22.99535 | 25.5827 | 25.46801 | 59.75609 | 256.1484 | 244.7714 | 205.9902 | 136.8296 | 112.4098 | 107.7127 |
| Adam33 | 110751 | 1.699319 | 3.145044 | 11.57523 | 1.15E-10 | 8.16E-10 | 1.115096 | 3.145044 | 10.71827 | 4.73E-10 | 3.31E-09 | 232.982 | 243.3216 | 259.5229 | 818.1868 | 859.4899 | 869.7654 | 2153.21 | 2054.867 | 1613.414 | 478.4514 | 518.5189 | 546.3218 | 201.9071 | 224.8196 | 175.7418 |
| Atf3 | 11910 | 1.699439 | 7.915585 | 56.93196 | 8.39E-25 | 1.31E-22 | -2.25795 | 7.915585 | -61.8244 | 1.46E-25 | 5.99E-23 | 10533.16 | 10919.28 | 10423.45 | 37128.8 | 37684.14 | 35677.96 | 8530.408 | 7960.765 | 7578.681 | 21321.09 | 20810.91 | 21716.29 | 9371.16 | 9223.107 | 9566.997 |
| Mbd3l2 | 234988 | 1.699476 | -2.68943 | 2.459123 | 0.022539 | 0.049545 | 2.865162 | -2.68943 | 6.046249 | 4.97E-06 | 1.92E-05 | 2.967923 | 0.888035 | 9.547666 | 10.38967 | 11.23516 | 13.52668 | 89.53943 | 101.8721 | 87.33582 | 1.538429 | 0 | 0.746341 | 23.36115 | 18.86598 | 12.14805 |
| 9930022D1 | 320509 | 1.699526 | 1.431927 | 13.76809 | 4.33E-12 | 3.79E-11 | -1.81152 | 1.431927 | -12.7434 | 1.90E-11 | 1.69E-10 | 187.7211 | 215.7925 | 173.5939 | 680.5236 | 672.7053 | 638.4592 | 207.5041 | 179.6165 | 201.1022 | 129.9972 | 168.5187 | 137.3268 | 34.2074 | 37.73197 | 20.24676 |
| Adgrg4 | 236798 | 1.700911 | -1.23414 | 2.942565 | 0.007689 | 0.018902 | 1.217616 | -1.23414 | 2.918217 | 0.008128 | 0.017981 | 5.193865 | 18.64873 | 6.943757 | 24.67547 | 39.32307 | 29.75869 | 49.74413 | 56.29771 | 132.1529 | 7.692145 | 9.912862 | 6.717071 | 53.39692 | 73.89177 | 54.26131 |
| Clec2d | 93694 | 1.7011 | 7.006463 | 43.54996 | 2.44E-22 | 1.74E-20 | -0.29325 | 7.006463 | -8.67093 | 1.94E-08 | 1.05E-07 | 5529.241 | 5750.915 | 5193.062 | 19192.32 | 19175.61 | 18776.38 | 16364.4 | 16436.25 | 15682.53 | 8820.582 | 8365.693 | 8364.246 | 1980.692 | 1815.065 | 1802.771 |
| Nfasc | 269116 | 1.701448 | -0.60556 | 4.036721 | 0.000579 | 0.001727 | 0.71707 | -0.60556 | 2.287733 | 0.032452 | 0.059838 | 11.87169 | 12.43249 | 16.49142 | 66.23417 | 46.34504 | 33.81669 | 66.79926 | 93.82952 | 83.88836 | 20.76879 | 28.21353 | 16.41951 | 78.42673 | 101.4047 | 63.97975 |
| Sds | 231691 | 1.702675 | -0.19603 | 4.919119 | 6.95E-05 | 0.000232 | -0.47412 | -0.19603 | -1.56527 | 0.132227 | 0.197732 | 16.32358 | 20.4248 | 29.51097 | 57.1432 | 88.4769 | 82.51273 | 51.16539 | 68.36151 | 49.41369 | 85.3828 | 57.18959 | 67.17071 | 46.72231 | 65.24486 | 69.64884 |
| Rnf152 | 320311 | 1.70316 | -0.4665 | 3.670912 | 0.001394 | 0.003917 | 0.612402 | -0.4665 | 1.783469 | 0.088738 | 0.141004 | 35.61508 | 5.32821 | 24.30315 | 61.03933 | 56.17581 | 68.98605 | 89.53943 | 75.06362 | 134.4512 | 13.84586 | 17.53814 | 10.44878 | 92.61029 | 107.6933 | 107.7127 |
| Gm4076 | 100042862 | 1.703359 | -0.83452 | 4.566929 | 0.000162 | 0.000516 | -1.54236 | -0.83452 | -3.92645 | 0.000755 | 0.002118 | 33.38913 | 19.53677 | 19.09533 | 67.53288 | 78.64613 | 108.2134 | 56.85043 | 28.14886 | 17.23733 | 19.99958 | 33.55123 | 43.28779 | 20.02385 | 22.7964 | 22.67637 |
| Khdc3 | 66991 | 1.704676 | -1.43727 | 2.897696 | 0.008517 | 0.020751 | -0.02789 | -1.43727 | -0.05826 | 0.954085 | 0.961985 | 25.22735 | 14.20856 | 4.339848 | 50.64966 | 22.47032 | 67.63339 | 55.42917 | 34.85097 | 43.66791 | 3.846072 | 19.0632 | 5.97073 | 22.52683 | 20.43815 | 44.54286 |
| Sp100 | 20684 | 1.705436 | 7.908217 | 47.83602 | 3.35E-23 | 3.18E-21 | -0.36715 | 7.908217 | -11.8925 | 6.98E-11 | 5.60E-10 | 9148.623 | 9257.765 | 8467.911 | 30704.08 | 31987.91 | 30936.86 | 26023.29 | 25551.12 | 23988.62 | 15098.91 | 15155.24 | 15015.64 | 5638.381 | 5853.171 | 5767.086 |
| Lbx1 | 16814 | 1.707557 | -2.75196 | 2.739787 | 0.012163 | 0.028629 | 0.572392 | -2.75196 | 1.105866 | 0.281094 | 0.372907 | 5.193865 | 2.664105 | 1.735939 | 16.88322 | 9.830767 | 8.116006 | 11.37009 | 17.42548 | 25.28142 | 2.307643 | 5.337695 | 4.478048 | 15.85221 | 18.86598 | 20.24676 |
| Cabp4 | 73660 | 1.707586 | 0.660581 | 5.94015 | 6.34E-06 | 2.40E-05 | 1.209592 | 0.660581 | 6.135108 | 4.06E-06 | 1.59E-05 | 66.03629 | 68.37869 | 29.51097 | 215.5857 | 130.6088 | 206.9582 | 454.8035 | 486.5731 | 381.5197 | 36.15308 | 37.36386 | 49.25852 | 42.55067 | 49.52321 | 42.11325 |
| Plaur | 18793 | 1.707632 | 6.125859 | 26.544 | 7.82E-18 | 2.00E-16 | 0.938011 | 6.125859 | 19.35148 | 5.12E-15 | 9.31E-14 | 2821.753 | 2649.896 | 3114.275 | 10366.3 | 9770.378 | 9740.56 | 21206.63 | 19554.07 | 18922 | 1798.423 | 1554.794 | 1744.946 | 1270.68 | 1245.941 | 1458.576 |
| Wnt1 | 22408 | 1.708213 | -1.15487 | 4.117274 | 0.000477 | 0.001438 | -0.13768 | -1.15487 | -0.40168 | 0.691916 | 0.774222 | 27.45329 | 11.54445 | 12.15157 | 71.429 | 56.17581 | 45.9907 | 39.7953 | 54.95729 | 67.80018 | 4.615287 | 2.287584 | 5.224389 | 49.22529 | 66.81702 | 87.46599 |
| Prelp | 116847 | 1.708383 | -2.27562 | 2.511228 | 0.020136 | 0.044839 | 1.097503 | -2.27562 | 2.134199 | 0.04458 | 0.07854 | 7.419807 | 8.88035 | 0.86797 | 18.18193 | 21.06593 | 9.468674 | 36.95278 | 40.21265 | 29.87805 | 7.692145 | 3.050111 | 2.985365 | 21.6925 | 18.0799 | 33.20468 |
| Unc5a | 107448 | 1.708476 | 3.949871 | 18.19595 | 1.77E-14 | 2.34E-13 | 0.019901 | 3.949871 | 0.259974 | 0.797375 | 0.867211 | 494.1592 | 418.2645 | 450.4762 | 1623.386 | 1678.252 | 1475.761 | 1746.73 | 1821.633 | 1497.35 | 821.521 | 812.0922 | 820.229 | 672.4675 | 698.0414 | 658.4245 |
| Rbm43 | 71684 | 1.709144 | 5.116221 | 30.42003 | 4.58E-19 | 1.55E-17 | -0.5005 | 5.116221 | -9.89222 | 1.99E-09 | 1.27E-08 | 1350.405 | 1435.065 | 1348.825 | 4624.703 | 4998.243 | 4808.734 | 3466.455 | 3545.416 | 3555.487 | 1547.659 | 1448.04 | 1521.043 | 1000.358 | 1075.361 | 1202.657 |
| Lhx3 | 16871 | 1.709906 | -2.06144 | 2.780138 | 0.011111 | 0.026381 | 1.198125 | -2.06144 | 2.574937 | 0.017523 | 0.035186 | 7.419807 | 2.664105 | 2.603909 | 15.58451 | 11.23516 | 16.23201 | 24.16143 | 40.21265 | 41.3696 | 3.076858 | 8.387806 | 9.702436 | 38.37904 | 47.16496 | 37.25403 |
| Ccdc122 | 108811 | 1.710097 | 2.196961 | 12.46897 | 2.87E-11 | 2.21E-10 | -1.30715 | 2.196961 | -9.33353 | 5.51E-09 | 3.25E-08 | 195.1409 | 198.9198 | 183.1416 | 755.8487 | 678.3229 | 591.1158 | 299.886 | 247.978 | 298.7805 | 194.6113 | 256.2094 | 267.1902 | 195.2325 | 184.7294 | 156.305 |
| Orm3 | 18407 | 1.710624 | -0.58776 | 7.225861 | 3.67E-07 | 1.61E-06 | -1.53928 | -0.58776 | -6.08558 | 4.55E-06 | 1.77E-05 | 45.26083 | 39.07354 | 41.66254 | 153.2477 | 154.4835 | 136.6194 | 59.69296 | 49.5956 | 49.41369 | 10.769 | 17.53814 | 29.10731 | 19.18952 | 12.57732 | 17.00728 |
| Znfx1 | 98999 | 1.711205 | 9.889736 | 48.27579 | 7.39E-16 | 1.27E-14 | -0.17381 | 9.889736 | -6.15155 | 3.74E-05 | 0.000128 | 36040.97 | 36892.53 | 35253.45 | 126474.8 | 127412.4 | 124231.7 | 119388.8 | 119502.6 | 110631.5 | 49858.94 | 50618.89 | 50211.6 | 22073.79 | 22037.83 | 22067.35 |
| Trim26 | 22670 | 1.711207 | 6.638624 | 32.5844 | 1.09E-19 | 4.25E-18 | 0.291519 | 6.638624 | 6.876489 | 7.80E-07 | 3.36E-06 | 3243.198 | 3185.381 | 3291.341 | 11236.43 | 11332.07 | 11424.63 | 15228.81 | 14833.11 | 13368.13 | 3909.148 | 3838.565 | 4165.331 | 3595.115 | 3459.55 | 3619.31 |
| Mllt3 | 70122 | 1.711229 | 4.083415 | 18.66585 | 1.06E-14 | 1.45E-13 | 0.017879 | 4.083415 | 0.240502 | 0.812236 | 0.880534 | 358.3767 | 361.4302 | 401.002 | 1376.632 | 1310.301 | 1224.164 | 1394.257 | 1364.549 | 1352.556 | 1249.973 | 1122.441 | 1271.019 | 1238.975 | 1355.206 | 1368.681 |
| Syne2 | 319565 | 1.712127 | 4.992261 | 16.50887 | 1.24E-13 | 1.41E-12 | 2.083736 | 4.992261 | 29.30339 | 1.00E-18 | 4.55E-17 | 662.5888 | 624.2886 | 613.6545 | 2206.507 | 2422.582 | 2051.997 | 10653.77 | 10006.25 | 8897.912 | 1186.129 | 1254.358 | 1221.014 | 1516.806 | 1467.616 | 1443.189 |
| Usp25 | 30940 | 1.713054 | 8.683101 | 58.38982 | 4.91E-25 | 8.60E-23 | -0.61736 | 8.683101 | -23.7026 | 8.12E-17 | 2.31E-15 | 19264.05 | 19326.31 | 18722.1 | 66682.22 | 68476.91 | 65592.21 | 46772.27 | 46469.74 | 43085.29 | 15746.59 | 15628.77 | 15575.4 | 10202.15 | 9831.535 | 9542.701 |
| H2-T23 | 15040 | 1.713251 | 7.059476 | 36.80275 | 8.46E-21 | 4.39E-19 | -0.13048 | 7.059476 | -3.29813 | 0.003372 | 0.008217 | 3816.007 | 3803.454 | 3754.837 | 13048.13 | 13170.42 | 13630.83 | 13182.19 | 12680.39 | 12033.96 | 8628.279 | 8590.639 | 8695.622 | 5969.609 | 5476.638 | 6176.071 |
| D930020B | 216393 | 1.715117 | -1.98932 | 2.54606 | 0.018665 | 0.041951 | 1.650228 | -1.98932 | 3.327292 | 0.003149 | 0.007724 | 1.483961 | 6.216245 | 4.339848 | 9.090964 | 14.04395 | 16.23201 | 34.11026 | 50.93603 | 44.81707 | 9.230573 | 9.912862 | 3.731706 | 43.385 | 34.58764 | 63.16988 |
| Spaca6 | 75202 | 1.716664 | 2.612524 | 16.21275 | 1.77E-13 | 1.96E-12 | -0.40894 | 2.612524 | -4.49673 | 0.000191 | 0.000594 | 345.763 | 321.4687 | 386.2465 | 1196.111 | 1275.191 | 1221.459 | 1023.308 | 994.5929 | 885.999 | 137.6894 | 182.2442 | 145.5365 | 74.2551 | 113.982 | 121.4805 |
| Svbp | 69216 | 1.718365 | 3.606701 | 16.715 | 9.68E-14 | 1.12E-12 | 0.124605 | 3.606701 | 1.50822 | 0.146159 | 0.215309 | 432.5748 | 411.1602 | 496.4786 | 1446.762 | 1658.591 | 1601.559 | 1988.344 | 1806.889 | 1575.492 | 469.2208 | 476.5799 | 495.5706 | 376.2814 | 316.7913 | 334.4764 |
| Cdh17 | 12557 | 1.719308 | -1.11716 | 2.633576 | 0.015402 | 0.035391 | 1.01887 | -1.11716 | 2.122418 | 0.045662 | 0.080111 | 9.64575 | 6.216245 | 5.207818 | 14.2858 | 29.4923 | 37.8747 | 28.42522 | 64.34024 | 75.84427 | 42.30679 | 22.11331 | 42.54145 | 46.72231 | 64.45877 | 33.20468 |
| Rbm11 | 224344 | 1.720268 | -1.07404 | 2.812124 | 0.010339 | 0.024725 | 1.838187 | -1.07404 | 4.426859 | 0.000226 | 0.000694 | 10.38773 | 2.664105 | 11.2836 | 16.88322 | 53.36702 | 18.93735 | 105.1733 | 80.4253 | 105.7223 | 15.38429 | 16.77561 | 5.97073 | 77.5924 | 73.10568 | 95.56469 |
| F730043M | 320046 | 1.721521 | 0.833601 | 9.285886 | 6.02E-09 | 3.35E-08 | -0.15187 | 0.833601 | -1.00474 | 0.326276 | 0.422458 | 71.97213 | 87.02743 | 77.24929 | 258.4431 | 315.9889 | 258.3595 | 241.6143 | 304.2757 | 235.5769 | 69.2293 | 61.76476 | 54.48291 | 40.88202 | 47.16496 | 29.9652 |
| Sertad3 | 170742 | 1.722315 | 4.120066 | 22.5711 | 2.22E-16 | 4.28E-15 | 0.172228 | 4.120066 | 2.790494 | 0.010855 | 0.023187 | 525.3224 | 558.574 | 633.6178 | 2022.09 | 2117.828 | 1881.561 | 2342.238 | 2369.866 | 2322.443 | 684.6009 | 664.9243 | 641.8535 | 639.9287 | 716.1213 | 685.1502 |
| Phlpp1 | 98432 | 1.72251 | 5.537419 | 25.62995 | 1.62E-17 | 3.81E-16 | 0.074728 | 5.537419 | 1.333183 | 0.196528 | 0.276529 | 1256.173 | 1134.909 | 1339.277 | 4581.846 | 4169.65 | 4379.938 | 4974.413 | 4724.987 | 4678.212 | 3736.844 | 3843.903 | 4072.038 | 1661.979 | 1507.706 | 1736.362 |
| Cnksr3 | 215748 | 1.724486 | 4.619868 | 29.79864 | 7.05E-19 | 2.27E-17 | -2.68783 | 4.619868 | -34.8765 | 2.62E-20 | 1.90E-18 | 1578.935 | 1501.667 | 1339.277 | 5320.811 | 5189.241 | 5091.441 | 922.3983 | 832.4019 | 776.8292 | 1348.433 | 1404.576 | 1470.292 | 790.1076 | 750.7089 | 728.8832 |
| Dact2 | 240025 | 1.724639 | -1.78444 | 2.532436 | 0.019228 | 0.043058 | 0.469893 | -1.78444 | 0.866108 | 0.396063 | 0.495027 | 3.709904 | 7.10428 | 8.679696 | 14.2858 | 22.47032 | 33.81669 | 17.05513 | 26.80843 | 62.0544 | 6.92293 | 2.287584 | 19.40487 | 49.22529 | 40.09021 | 48.59222 |
| Oas3 | 246727 | 1.724763 | 9.26023 | 72.1781 | 5.43E-27 | 1.89E-24 | -0.28909 | 9.26023 | -14.5 | 1.59E-12 | 1.72E-11 | 19109.71 | 18868.08 | 19333.15 | 68216 | 67382.88 | 66651.35 | 58260.32 | 57103.31 | 56638.43 | 35730.78 | 36126.28 | 34425.74 | 24987.26 | 25610.57 | 25302.78 |
| Gtpbp2 | 56055 | 1.725058 | 6.552236 | 44.21083 | 1.77E-22 | 1.32E-20 | -1.18921 | 6.552236 | -30.6918 | 3.81E-19 | 1.96E-17 | 3747.003 | 3853.184 | 3902.391 | 13571.51 | 13584.72 | 13407.64 | 6384.304 | 6381.748 | 5774.507 | 5166.813 | 5251.529 | 5236.33 | 3314.781 | 3396.663 | 3265.397 |
| Grin2b | 14812 | 1.726211 | -1.59553 | 3.045487 | 0.006071 | 0.015251 | 0.906953 | -1.59553 | 2.159419 | 0.042341 | 0.075167 | 4.451884 | 8.88035 | 12.15157 | 18.18193 | 28.08791 | 43.28537 | 72.4843 | 37.53181 | 60.90525 | 7.692145 | 3.812639 | 2.239024 | 57.56856 | 62.88661 | 36.44416 |
| Fer1l6 | 631797 | 1.726799 | -1.55558 | 2.884986 | 0.008767 | 0.0213 | 0.895737 | -1.55558 | 1.990177 | 0.059533 | 0.100526 | 3.709904 | 6.216245 | 12.15157 | 28.5716 | 28.08791 | 16.23201 | 44.05909 | 30.8297 | 67.80018 | 21.538 | 6.100223 | 6.717071 | 52.5626 | 43.23454 | 38.0639 |
| Aig1 | 66253 | 1.727153 | 0.82318 | 6.880946 | 7.72E-07 | 3.25E-06 | -0.73079 | 0.82318 | -3.22214 | 0.004029 | 0.009632 | 67.52025 | 83.47529 | 59.8899 | 223.378 | 186.7846 | 357.1043 | 190.449 | 127.3401 | 151.6885 | 136.9202 | 126.5796 | 102.9951 | 52.5626 | 34.58764 | 35.63429 |
| Lrrc4 | 192198 | 1.728528 | 4.301565 | 26.6706 | 7.08E-18 | 1.84E-16 | -0.72959 | 4.301565 | -12.2159 | 4.23E-11 | 3.53E-10 | 859.2137 | 904.0196 | 854.9501 | 3201.318 | 3074.221 | 2979.927 | 2006.82 | 1912.782 | 1883.466 | 669.2166 | 755.6651 | 642.5998 | 600.7154 | 669.7424 | 635.7482 |
| Cadps | 27062 | 1.729553 | -1.50255 | 3.804187 | 0.001013 | 0.002914 | 0.42171 | -1.50255 | 1.196912 | 0.244467 | 0.331734 | 8.903769 | 15.98463 | 6.943757 | 32.46773 | 35.10988 | 40.58003 | 49.74413 | 40.21265 | 60.90525 | 7.692145 | 3.812639 | 7.463413 | 46.72231 | 34.58764 | 23.48624 |
| Clec1a | 243653 | 1.729686 | -0.28002 | 4.500669 | 0.00019 | 0.000601 | 0.711848 | -0.28002 | 2.508019 | 0.020277 | 0.039941 | 11.12971 | 28.41712 | 19.9633 | 74.02642 | 40.72746 | 87.9234 | 125.071 | 97.85079 | 110.3189 | 62.30637 | 54.13948 | 67.17071 | 40.04769 | 29.08506 | 34.01455 |
| Shf | 435684 | 1.732226 | 1.283897 | 7.821844 | 1.06E-07 | 4.97E-07 | 0.724256 | 1.283897 | 4.573655 | 0.000159 | 0.000499 | 81.61788 | 57.72227 | 63.36178 | 212.9883 | 242.9604 | 270.5335 | 493.1775 | 439.6583 | 339.0009 | 102.3055 | 118.9543 | 100.7561 | 95.9476 | 83.32476 | 87.46599 |
| Ndrg2 | 29811 | 1.732302 | -0.73668 | 3.331191 | 0.00312 | 0.008258 | 4.051772 | -0.73668 | 12.49674 | 2.75E-11 | 2.38E-10 | 5.193865 | 6.216245 | 6.943757 | 18.18193 | 32.30109 | 18.93735 | 375.2129 | 414.1903 | 391.8621 | 9.999788 | 3.812639 | 10.44878 | 146.0072 | 115.5541 | 127.9595 |
| Il20rb | 213208 | 1.733279 | 1.835791 | 8.979949 | 1.07E-08 | 5.75E-08 | -0.32546 | 1.835791 | -2.01905 | 0.056218 | 0.095783 | 97.94146 | 73.7069 | 78.98523 | 285.716 | 307.5626 | 301.6449 | 241.6143 | 231.893 | 264.3058 | 436.1446 | 412.5276 | 536.6194 | 151.8475 | 183.9433 | 137.6779 |
| Slc1a2 | 20511 | 1.733617 | 0.634895 | 6.815134 | 8.91E-07 | 3.72E-06 | 0.509467 | 0.634895 | 2.692129 | 0.013527 | 0.028072 | 34.8731 | 40.84961 | 40.79457 | 120.78 | 143.2483 | 151.4988 | 198.9765 | 191.6803 | 218.3396 | 83.07516 | 89.21576 | 53.73657 | 80.92971 | 94.32991 | 77.74755 |
| Prdx5 | 54683 | 1.734287 | 6.384083 | 24.66223 | 1.84E-14 | 2.42E-13 | -0.49571 | 6.384083 | -7.92367 | 5.07E-07 | 2.25E-06 | 4257.486 | 4622.222 | 4036.927 | 15772.82 | 15542.44 | 14481.66 | 12458.77 | 11361.41 | 10190.71 | 2223.03 | 2419.501 | 2611.448 | 1710.37 | 1610.683 | 1891.857 |
| Adamts19 | 240322 | 1.735002 | -1.87657 | 2.501021 | 0.020587 | 0.045723 | 1.288487 | -1.87657 | 2.544302 | 0.018737 | 0.037291 | 8.161788 | 6.216245 | 2.603909 | 19.48064 | 19.66153 | 18.93735 | 32.689 | 37.53181 | 90.78329 | 5.384501 | 1.525056 | 10.44878 | 35.04173 | 58.9562 | 36.44416 |
| Pou4f1 | 18996 | 1.73588 | -0.71827 | 3.497238 | 0.002107 | 0.005749 | 3.65353 | -0.71827 | 11.92646 | 6.62E-11 | 5.33E-10 | 7.419807 | 13.32052 | 6.943757 | 20.77935 | 42.13186 | 37.8747 | 396.5318 | 431.6158 | 443.5741 | 1.538429 | 4.575167 | 10.44878 | 110.9655 | 97.47424 | 105.2831 |
| Ccr7 | 12775 | 1.736299 | 3.745171 | 19.12011 | 6.53E-15 | 9.31E-14 | -0.08256 | 3.745171 | -1.10467 | 0.281599 | 0.373266 | 430.3488 | 429.8089 | 451.3442 | 1493.516 | 1542.026 | 1634.023 | 1505.115 | 1556.23 | 1511.14 | 843.059 | 878.4321 | 704.5461 | 404.6486 | 396.9717 | 384.6884 |
| Wt1os | 329480 | 1.736551 | -2.64997 | 2.755304 | 0.011748 | 0.027728 | 0.40658 | -2.64997 | 0.785711 | 0.440673 | 0.539603 | 4.451884 | 5.32821 | 2.603909 | 16.88322 | 18.25714 | 10.82134 | 25.5827 | 10.72337 | 31.0272 | 1.538429 | 1.525056 | 5.97073 | 23.36115 | 19.65207 | 13.76779 |
| 5430421F1 | 74489 | 1.737423 | -3.15781 | 2.934659 | 0.007829 | 0.019217 | 0.883412 | -3.15781 | 1.943777 | 0.065224 | 0.108504 | 7.419807 | 5.32821 | 1.735939 | 12.98709 | 15.44835 | 20.29002 | 48.32287 | 21.44675 | 29.87805 | 0 | 0 | 0 | 10.84625 | 16.50774 | 15.38754 |
| Chil1 | 12654 | 1.737979 | 3.744313 | 21.69408 | 5.00E-16 | 8.83E-15 | -0.31774 | 3.744313 | -4.58451 | 0.000155 | 0.000487 | 789.4675 | 781.4708 | 728.2265 | 2533.782 | 2755.424 | 2928.526 | 2329.447 | 2379.249 | 2142.026 | 282.3017 | 337.7998 | 308.2389 | 214.422 | 191.8042 | 268.8769 |
| Itga2 | 16398 | 1.739875 | 0.285484 | 4.180021 | 0.00041 | 0.001248 | 1.696831 | 0.285484 | 6.141537 | 4.00E-06 | 1.57E-05 | 24.48536 | 21.31284 | 7.811726 | 76.62384 | 63.19779 | 41.9327 | 207.5041 | 170.2336 | 222.9362 | 16.1535 | 21.35078 | 23.88292 | 507.2708 | 473.2217 | 552.3315 |
| Dock4 | 238130 | 1.739991 | 7.413313 | 41.0768 | 8.37E-22 | 5.38E-20 | -0.37564 | 7.413313 | -10.1944 | 1.17E-09 | 7.67E-09 | 5564.856 | 5245.623 | 5437.83 | 19127.39 | 20247.17 | 18697.93 | 16011.92 | 15940.3 | 14641.39 | 7370.613 | 7331.705 | 7442.515 | 9385.344 | 8772.682 | 9307.029 |
| Slc24a2 | 76376 | 1.741341 | -1.49952 | 2.979587 | 0.007064 | 0.017505 | 1.104813 | -1.49952 | 2.60762 | 0.016309 | 0.033064 | 4.451884 | 14.20856 | 10.41564 | 44.15611 | 30.8967 | 21.64268 | 51.16539 | 60.31898 | 103.424 | 6.153716 | 3.050111 | 1.492683 | 83.43269 | 62.88661 | 42.11325 |
| Itga5 | 16402 | 1.741425 | 8.404024 | 67.36567 | 2.36E-26 | 6.96E-24 | -0.68492 | 8.404024 | -29.5569 | 8.36E-19 | 3.90E-17 | 17015.1 | 16101.85 | 17316.86 | 59604.26 | 61143.16 | 59452.45 | 39681.6 | 38450 | 38301.36 | 12437.43 | 11378.44 | 12292.24 | 7055.903 | 7173.004 | 7449.187 |
| A430093F1 | 403202 | 1.742312 | 0.393498 | 9.802589 | 2.34E-09 | 1.38E-08 | -1.96629 | 0.393498 | -9.38853 | 4.98E-09 | 2.96E-08 | 75.68204 | 64.82655 | 75.51336 | 263.638 | 287.901 | 227.2482 | 72.4843 | 57.63814 | 74.69511 | 80.76752 | 70.15256 | 81.3512 | 34.2074 | 22.7964 | 33.20468 |
| Agtrap | 11610 | 1.74234 | 5.782449 | 32.66782 | 1.03E-19 | 4.06E-18 | -0.83369 | 5.782449 | -16.6558 | 1.04E-13 | 1.43E-12 | 1964.765 | 1970.55 | 1927.76 | 7083.16 | 7127.306 | 6745.754 | 4289.365 | 4207.584 | 3769.23 | 2763.788 | 2992.922 | 3041.341 | 2323.6 | 2421.134 | 2335.666 |
| Traf2 | 22030 | 1.74253 | 5.203634 | 30.73359 | 3.70E-19 | 1.28E-17 | -1.05044 | 5.203634 | -19.1322 | 6.44E-15 | 1.14E-13 | 1495.091 | 1430.624 | 1427.81 | 5332.5 | 5245.416 | 4998.107 | 2777.144 | 2686.205 | 2390.244 | 2106.878 | 2049.675 | 2215.141 | 1243.147 | 1202.706 | 1276.356 |
| Gbgt1 | 227671 | 1.742876 | 3.553617 | 13.84619 | 3.89E-12 | 3.42E-11 | 0.257314 | 3.553617 | 2.615011 | 0.016046 | 0.032588 | 382.1201 | 314.3644 | 467.8356 | 1398.71 | 1310.301 | 1406.774 | 1830.584 | 1900.718 | 1452.533 | 786.1372 | 748.8023 | 756.79 | 238.6175 | 268.8403 | 233.2426 |
| Cdh10 | 320873 | 1.743295 | -3.06074 | 2.581165 | 0.017285 | 0.039133 | 0.338583 | -3.06074 | 0.606482 | 0.550592 | 0.645269 | 5.935846 | 5.32821 | 1.735939 | 12.98709 | 15.44835 | 16.23201 | 22.74017 | 12.0638 | 26.43058 | 0 | 2.287584 | 0.746341 | 7.508942 | 24.36856 | 12.14805 |
| 4632415L0 | 70808 | 1.746768 | 3.810545 | 21.11685 | 8.68E-16 | 1.46E-14 | -0.25967 | 3.810545 | -3.70999 | 0.00127 | 0.003402 | 459.2861 | 520.3885 | 515.5739 | 1655.854 | 1856.611 | 1830.159 | 1510.8 | 1678.208 | 1451.384 | 849.982 | 841.8307 | 882.9217 | 362.0979 | 349.8068 | 346.6245 |
| Trex1 | 22040 | 1.7489 | 7.107433 | 40.57528 | 1.08E-21 | 6.66E-20 | -0.46437 | 7.107433 | -12.2727 | 3.87E-11 | 3.27E-10 | 6329.838 | 6525.281 | 6409.088 | 22555.98 | 23829.78 | 22753.22 | 18152.34 | 17920.1 | 16174.36 | 8283.67 | 8721.794 | 8231.398 | 1675.328 | 1829.214 | 1771.996 |
| Tapbp | 21356 | 1.749091 | 9.022618 | 57.73244 | 6.24E-25 | 1.04E-22 | -0.43844 | 9.022618 | -17.0295 | 6.68E-14 | 9.61E-13 | 20795.49 | 20879.48 | 20406.83 | 75248.51 | 74706.81 | 72881.74 | 58567.32 | 58845.86 | 54028.7 | 33182.37 | 33304.17 | 33121.88 | 10065.32 | 9653.881 | 10180.88 |
| Adamtsl4 | 229595 | 1.751693 | 3.2774 | 16.50241 | 1.25E-13 | 1.42E-12 | -1.12771 | 3.2774 | -10.9246 | 3.35E-10 | 2.40E-09 | 359.1187 | 353.4379 | 326.3566 | 1138.968 | 1256.934 | 1359.431 | 581.2957 | 634.0195 | 566.5337 | 573.834 | 666.4493 | 601.5511 | 393.8023 | 479.5104 | 377.3995 |
| Htra1 | 56213 | 1.751893 | 0.421625 | 3.975489 | 0.000671 | 0.001983 | 4.246118 | 0.421625 | 16.22151 | 1.76E-13 | 2.31E-12 | 27.45329 | 26.64105 | 10.41564 | 90.90964 | 60.389 | 68.98605 | 1476.69 | 1518.698 | 1356.004 | 39.99915 | 22.87584 | 19.40487 | 95.11327 | 44.80671 | 45.35273 |
| Kcnh1 | 16510 | 1.752113 | -0.87314 | 4.305028 | 0.000304 | 0.000938 | 0.351407 | -0.87314 | 1.1186 | 0.275743 | 0.367188 | 11.12971 | 19.53677 | 15.62345 | 54.54578 | 82.85932 | 35.16936 | 66.79926 | 57.63814 | 91.93245 | 12.30743 | 14.48803 | 11.19512 | 56.73423 | 36.94588 | 48.59222 |
| Cacna1b | 12287 | 1.752391 | 2.564471 | 15.31902 | 5.43E-13 | 5.50E-12 | -1.18556 | 2.564471 | -10.5367 | 6.45E-10 | 4.42E-09 | 243.3697 | 214.0164 | 250.8432 | 916.8887 | 810.3361 | 823.7747 | 358.1577 | 439.6583 | 371.1773 | 341.5312 | 313.3989 | 272.4146 | 235.2802 | 220.8892 | 233.2426 |
| Nr2e1 | 21907 | 1.752817 | -2.08803 | 2.62401 | 0.015731 | 0.036048 | 0.666771 | -2.08803 | 1.275329 | 0.2159 | 0.298926 | 2.225942 | 5.32821 | 10.41564 | 16.88322 | 15.44835 | 24.34802 | 45.48035 | 13.40422 | 45.96622 | 3.076858 | 6.862751 | 3.731706 | 40.04769 | 41.66238 | 26.72572 |
| Zfp423 | 94187 | 1.753984 | -1.26588 | 3.937623 | 0.000735 | 0.002158 | 0.780858 | -1.26588 | 2.371391 | 0.027197 | 0.051424 | 9.64575 | 9.768385 | 8.679696 | 31.16902 | 39.32307 | 33.81669 | 38.37404 | 75.06362 | 78.14258 | 7.692145 | 8.387806 | 7.463413 | 55.06558 | 47.16496 | 60.74027 |
| Spred1 | 114715 | 1.754934 | 7.782027 | 60.99926 | 1.94E-25 | 3.92E-23 | -0.96066 | 7.782027 | -35.242 | 2.10E-20 | 1.57E-18 | 10116.17 | 10388.23 | 10687.31 | 37292.43 | 38966.35 | 36085.12 | 20112.26 | 19492.41 | 20223.99 | 9549.028 | 9093.907 | 9503.163 | 5554.114 | 5175.568 | 5525.745 |
| Myrfl | 237558 | 1.756263 | -1.92052 | 3.14263 | 0.004849 | 0.012399 | 0.87964 | -1.92052 | 2.098866 | 0.047896 | 0.083464 | 5.193865 | 7.10428 | 5.207818 | 12.98709 | 32.30109 | 24.34802 | 41.21656 | 40.21265 | 43.66791 | 6.92293 | 6.862751 | 2.239024 | 30.8701 | 44.80671 | 23.48624 |
| Hk1os | 70162 | 1.756435 | 0.047185 | 4.994633 | 5.80E-05 | 0.000195 | -0.07426 | 0.047185 | -0.26213 | 0.795734 | 0.865784 | 31.16319 | 35.5214 | 26.03909 | 119.4812 | 153.0791 | 75.74939 | 122.2284 | 89.80826 | 119.5122 | 29.99936 | 66.33992 | 52.24389 | 62.57452 | 55.02578 | 34.01455 |
| Gm19272 | 100502597 | 1.756455 | -3.71805 | 2.661311 | 0.014485 | 0.033481 | 0.464537 | -3.71805 | 0.838203 | 0.411206 | 0.51022 | 5.193865 | 1.77607 | 1.735939 | 12.98709 | 19.66153 | 4.058003 | 17.05513 | 13.40422 | 14.93902 | 0 | 0 | 0 | 8.343269 | 12.57732 | 8.098703 |
| Slfn9 | 237886 | 1.758915 | 7.841834 | 57.17891 | 7.65E-25 | 1.23E-22 | -0.32408 | 7.841834 | -12.3785 | 3.30E-11 | 2.81E-10 | 9426.123 | 9287.07 | 8644.109 | 33274.23 | 33552.41 | 32115.04 | 27249.83 | 27545.67 | 27229.24 | 11225.15 | 11511.12 | 11008.53 | 5398.095 | 5251.032 | 5098.133 |
| Pnpla1 | 433091 | 1.759529 | -2.1727 | 2.579592 | 0.017345 | 0.039255 | 1.377223 | -2.1727 | 2.723788 | 0.012606 | 0.026428 | 6.677827 | 1.77607 | 3.471878 | 7.792255 | 25.27911 | 13.52668 | 31.26774 | 38.87223 | 44.81707 | 6.153716 | 9.912862 | 1.492683 | 33.37308 | 41.66238 | 34.82442 |
| Tha1 | 71776 | 1.759539 | 0.970035 | 9.069446 | 9.03E-09 | 4.90E-08 | -2.77154 | 0.970035 | -9.85578 | 2.13E-09 | 1.34E-08 | 109.8132 | 99.45992 | 87.66493 | 409.0934 | 304.7538 | 367.9256 | 34.11026 | 61.6594 | 70.09849 | 81.53673 | 121.2419 | 113.4439 | 113.4685 | 130.4897 | 91.51534 |
| Chrna1 | 11435 | 1.760913 | -2.04912 | 3.191197 | 0.00433 | 0.011173 | 0.739245 | -2.04912 | 1.740134 | 0.096237 | 0.150905 | 8.161788 | 2.664105 | 4.339848 | 18.18193 | 12.63956 | 24.34802 | 29.84648 | 40.21265 | 26.43058 | 6.153716 | 9.150334 | 7.463413 | 22.52683 | 33.01547 | 24.29611 |
| Scarf1 | 380713 | 1.760976 | 5.894338 | 32.57014 | 1.10E-19 | 4.28E-18 | -1.01415 | 5.894338 | -19.5363 | 4.22E-15 | 7.84E-14 | 2826.205 | 2731.596 | 2537.075 | 9796.163 | 10003.51 | 9528.192 | 5397.949 | 5214.241 | 4566.744 | 2812.248 | 2906.756 | 2863.711 | 1400.001 | 1538.364 | 1543.613 |
| Xrn1 | 24127 | 1.761298 | 6.568117 | 41.36594 | 7.22E-22 | 4.70E-20 | -0.21363 | 6.568117 | -5.87832 | 7.31E-06 | 2.77E-05 | 3594.897 | 3616.966 | 3334.739 | 12600.08 | 12867.07 | 12721.84 | 11468.15 | 11835.92 | 10949.15 | 4208.372 | 4351.746 | 4138.462 | 2678.189 | 2680.542 | 2791.623 |
| Rcvrn | 19674 | 1.761424 | -2.17072 | 2.993352 | 0.006845 | 0.017015 | 1.038879 | -2.17072 | 2.423646 | 0.024326 | 0.046679 | 8.161788 | 9.768385 | 3.471878 | 25.97418 | 14.04395 | 40.58003 | 51.16539 | 62.99982 | 48.26453 | 3.076858 | 12.96297 | 2.985365 | 12.5149 | 4.716496 | 7.288832 |
| Adm | 11535 | 1.76213 | 2.048253 | 8.931298 | 1.17E-08 | 6.26E-08 | -0.39174 | 2.048253 | -2.36666 | 0.027472 | 0.051902 | 92.00561 | 79.92315 | 58.15396 | 257.1444 | 304.7538 | 270.5335 | 228.823 | 210.4462 | 217.1904 | 89.22888 | 138.7801 | 108.9658 | 1797.975 | 1752.964 | 1869.181 |
| Ofcc1 | 218165 | 1.762199 | -2.72505 | 2.536112 | 0.019075 | 0.042743 | 0.396496 | -2.72505 | 0.682184 | 0.50246 | 0.600118 | 2.225942 | 2.664105 | 6.075787 | 22.07806 | 8.426372 | 10.82134 | 7.106304 | 22.78717 | 29.87805 | 3.846072 | 3.050111 | 1.492683 | 30.03577 | 19.65207 | 18.62702 |
| Lyrm1 | 73919 | 1.762418 | 2.052077 | 14.03025 | 3.01E-12 | 2.70E-11 | -1.00552 | 2.052077 | -8.48384 | 2.79E-08 | 1.48E-07 | 187.7211 | 173.1668 | 139.7431 | 632.4714 | 558.9493 | 622.2272 | 302.7286 | 333.765 | 302.2279 | 208.4571 | 228.7584 | 235.8438 | 125.9834 | 138.3505 | 132.8187 |
| Rhpn1 | 14787 | 1.762481 | -1.48007 | 3.020272 | 0.006434 | 0.016071 | 1.552389 | -1.48007 | 3.862355 | 0.000881 | 0.002441 | 11.87169 | 6.216245 | 4.339848 | 18.18193 | 26.68351 | 37.8747 | 86.69691 | 79.08488 | 79.29174 | 7.692145 | 6.100223 | 0.746341 | 65.0775 | 61.31444 | 89.8956 |

| | | | | | | | | | | | | | | | | | | | | | | | | | |
|---|---|---|---|---|---|---|---|---|---|---|---|---|---|---|---|---|---|---|---|---|---|---|---|---|---|
| Prox1 | 19130 | 1.76262 | -0.40545 | 4.161583 | 0.000429 | 0.001301 | 1.066562 | -0.40545 | 3.573045 | 0.00176 | 0.004567 | 11.12971 | 22.20087 | 19.9633 | 63.63675 | 78.64613 | 45.9907 | 143.5473 | 131.3613 | 128.7054 | 7.692145 | 23.63836 | 18.65853 | 65.91183 | 74.67785 | 114.1917 |
| Prrt4 | 101359 | 1.763408 | -1.41837 | 2.619322 | 0.015894 | 0.036369 | 1.600611 | -1.41837 | 3.380032 | 0.002781 | 0.006911 | 1.483961 | 16.87266 | 11.2836 | 32.46773 | 19.66153 | 21.64268 | 66.79926 | 60.31898 | 108.0206 | 9.999788 | 4.575167 | 3.731706 | 60.07154 | 58.9562 | 90.70547 |
| Eif2ak2 | 19106 | 1.763993 | 8.027363 | 55.02914 | 1.72E-24 | 2.51E-22 | -0.29285 | 8.027363 | -10.8051 | 4.09E-10 | 2.89E-09 | 9245.822 | 9594.33 | 9256.028 | 34189.82 | 34229.33 | 33389.25 | 29222.54 | 29589.81 | 27671.67 | 13488.18 | 13575.28 | 13419.96 | 7584.866 | 7625.001 | 7191.648 |
| Msrb3 | 320183 | 1.76432 | -0.69733 | 3.763955 | 0.001116 | 0.003188 | 0.784333 | -0.69733 | 2.292629 | 0.032121 | 0.059289 | 11.87169 | 12.43249 | 11.2836 | 54.54578 | 35.10988 | 43.28537 | 85.27565 | 67.02109 | 85.03751 | 23.07643 | 22.11331 | 38.80975 | 65.91183 | 79.39434 | 35.63429 |
| Map3k8 | 26410 | 1.765562 | 6.286117 | 39.40277 | 2.01E-21 | 1.18E-19 | -0.75474 | 6.286117 | -18.2414 | 1.69E-14 | 2.73E-13 | 3048.799 | 3141.868 | 2822.637 | 10909.16 | 11355.94 | 10462.88 | 6979.812 | 6530.535 | 6629.479 | 4413.753 | 4554.579 | 4451.179 | 2130.037 | 2314.227 | 2302.461 |
| Rmdn3 | 67809 | 1.766455 | 4.797706 | 27.95256 | 2.67E-18 | 7.70E-17 | 0.199238 | 4.797706 | 3.878045 | 0.000849 | 0.002358 | 1028.385 | 975.9504 | 920.9158 | 3446.774 | 3645.81 | 3557.516 | 4427.228 | 4263.882 | 4031.238 | 1211.513 | 1105.665 | 1120.258 | 735.042 | 745.9924 | 788.0038 |
| Mir350 | 723921 | 1.767831 | -1.6939 | 3.700115 | 0.0013 | 0.00367 | -1.55087 | -1.6939 | -3.07187 | 0.005713 | 0.013148 | 8.903769 | 11.54445 | 10.41564 | 48.05224 | 23.87472 | 45.9907 | 8.527565 | 12.0638 | 19.53565 | 29.23015 | 22.11331 | 11.94146 | 14.18356 | 18.86598 | 26.72572 |
| Wdr37 | 207615 | 1.769221 | 6.052893 | 47.99588 | 3.12E-23 | 2.99E-21 | -2.17001 | 6.052893 | -50.4598 | 1.08E-23 | 2.56E-21 | 3473.212 | 3564.572 | 3444.971 | 12827.35 | 12913.41 | 12403.96 | 3010.23 | 2969.034 | 2832.669 | 3844.534 | 3945.319 | 3723.497 | 1978.189 | 1910.967 | 2008.478 |
| C1qtnf9 | 239126 | 1.769551 | -2.33968 | 3.057464 | 0.005906 | 0.014867 | 0.231894 | -2.33968 | 0.494559 | 0.625964 | 0.716439 | 4.451884 | 7.10428 | 6.075787 | 14.2858 | 26.68351 | 27.05335 | 36.95278 | 16.08506 | 31.0272 | 5.384501 | 6.862751 | 2.239024 | 6.674615 | 18.0799 | 15.38754 |
| Epha6 | 13840 | 1.770253 | -2.43644 | 2.814796 | 0.010277 | 0.024585 | 1.419931 | -2.43644 | 3.056242 | 0.005922 | 0.013575 | 6.677827 | 1.77607 | 3.471878 | 10.38967 | 11.23516 | 22.99535 | 28.42522 | 37.53181 | 55.15947 | 1.538429 | 1.525056 | 3.731706 | 25.02981 | 34.58764 | 27.53559 |
| Igsf9 | 93842 | 1.770572 | 5.460936 | 39.01766 | 2.47E-21 | 1.42E-19 | -2.65105 | 5.460936 | -45.6753 | 8.90E-23 | 1.58E-20 | 4135.801 | 4331.835 | 3994.396 | 15211.78 | 15236.28 | 14911.81 | 2521.317 | 2706.312 | 2307.504 | 1199.205 | 1279.522 | 1059.058 | 629.0825 | 706.6883 | 716.7352 |
| Zscan10 | 332221 | 1.771424 | -1.99214 | 2.743914 | 0.012051 | 0.028386 | 1.426737 | -1.99214 | 3.049138 | 0.00602 | 0.013775 | 8.161788 | 1.77607 | 4.339848 | 11.68838 | 11.23516 | 31.11136 | 52.58665 | 41.55307 | 44.81707 | 4.615287 | 6.100223 | 2.985365 | 35.04173 | 62.88661 | 42.11325 |
| Gm4117 | 100042940 | 1.77167 | 1.065472 | 10.89899 | 3.49E-10 | 2.32E-09 | -0.79048 | 1.065472 | -5.4013 | 2.22E-05 | 7.83E-05 | 111.2971 | 98.57188 | 86.79696 | 409.0934 | 365.1428 | 315.1716 | 218.8742 | 241.2759 | 196.5056 | 104.6132 | 100.6537 | 100.0097 | 32.53875 | 47.16496 | 35.63429 |
| Tnfrsf19 | 29820 | 1.77254 | -1.8087 | 2.524615 | 0.019558 | 0.043704 | 2.534457 | -1.8087 | 5.273504 | 2.99E-05 | 0.000104 | 2.967923 | 1.77607 | 7.811726 | 18.18193 | 8.426372 | 16.23201 | 63.95674 | 87.12741 | 106.8715 | 4.615287 | 6.862751 | 3.731706 | 62.57452 | 75.46393 | 38.87377 |
| Ankrd35 | 213121 | 1.77349 | -1.46761 | 2.601683 | 0.016523 | 0.037597 | 0.818718 | -1.46761 | 1.591335 | 0.126238 | 0.190083 | 1.483961 | 7.992315 | 24.30315 | 24.67547 | 28.08791 | 21.64268 | 34.11026 | 50.93603 | 52.86116 | 16.92272 | 3.812639 | 7.463413 | 70.91779 | 80.18043 | 52.64157 |
| Dnah3 | 381917 | 1.774148 | 0.004832 | 5.052901 | 5.05E-05 | 0.000171 | 0.546961 | 0.004832 | 2.117731 | 0.046099 | 0.08076 | 29.67923 | 23.08891 | 35.58675 | 119.4812 | 126.3956 | 81.16006 | 157.76 | 142.0847 | 187.3124 | 11.53822 | 22.11331 | 27.61463 | 66.74615 | 117.9124 | 57.50079 |
| Tyk2 | 54721 | 1.774189 | 7.262108 | 41.2839 | 7.52E-22 | 4.87E-20 | -0.39249 | 7.262108 | -10.5426 | 6.38E-10 | 4.38E-09 | 5594.535 | 5697.632 | 5825.812 | 20932.59 | 20679.72 | 20847.32 | 16667.13 | 17560.87 | 15402.13 | 7940.601 | 8036.281 | 7775.383 | 4287.606 | 4086.057 | 4184.6 |
| 4833438C( | 100503392 | 1.774469 | 3.22524 | 15.666 | 3.49E-13 | 3.66E-12 | 0.727981 | 3.22524 | 8.860667 | 1.34E-08 | 7.45E-08 | 278.2428 | 258.4182 | 238.6916 | 937.668 | 896.0042 | 1009.09 | 1749.572 | 1624.591 | 1534.123 | 582.2953 | 548.2575 | 535.873 | 285.3398 | 222.4614 | 232.4328 |
| Cdsn | 386463 | 1.77518 | -2.6445 | 2.872368 | 0.009022 | 0.021863 | 0.965583 | -2.6445 | 1.986991 | 0.059909 | 0.101095 | 2.967923 | 5.32821 | 1.735939 | 11.68838 | 9.830767 | 14.87935 | 15.63387 | 26.80843 | 34.47467 | 3.846072 | 1.525056 | 1.492683 | 30.8701 | 31.4433 | 25.10598 |
| Yap1 | 22601 | 1.775356 | -1.16627 | 2.879102 | 0.008885 | 0.021559 | 3.872512 | -1.16627 | 10.19646 | 1.16E-09 | 7.64E-09 | 20.77546 | 3.55214 | 2.603909 | 35.06515 | 14.04395 | 25.70069 | 369.5278 | 388.7223 | 349.3433 | 4.615287 | 3.050111 | 2.239024 | 55.06558 | 74.67785 | 71.26858 |
| Mrgpra2b | 235712 | 1.775951 | 0.047855 | 8.531083 | 2.54E-08 | 1.29E-07 | -0.12047 | 0.047855 | -0.72071 | 0.478906 | 0.577361 | 59.35846 | 53.2821 | 52.07818 | 200.0012 | 202.2329 | 204.2528 | 197.5553 | 180.9569 | 199.9531 | 13.84586 | 13.7255 | 25.3756 | 16.68654 | 36.1598 | 21.8665 |
| Cd55b | 13137 | 1.776189 | -0.92038 | 5.393513 | 2.26E-05 | 8.00E-05 | -2.78883 | -0.92038 | -5.97841 | 5.81E-06 | 2.23E-05 | 20.03348 | 35.5214 | 47.73833 | 142.858 | 106.734 | 102.8027 | 12.79135 | 14.74464 | 24.13227 | 41.53758 | 48.80178 | 29.85365 | 5.005962 | 14.93557 | 20.24676 |
| Slc17a8 | 216227 | 1.776737 | -0.46558 | 4.98879 | 5.88E-05 | 0.000198 | 1.372815 | -0.46558 | 5.668008 | 1.19E-05 | 4.36E-05 | 14.83961 | 12.43249 | 23.43518 | 51.94837 | 61.79339 | 68.98605 | 153.4962 | 148.7868 | 186.1632 | 24.61486 | 24.40089 | 41.04877 | 27.53279 | 34.58764 | 29.9652 |
| Otud1 | 71198 | 1.776934 | 5.264185 | 31.26261 | 2.59E-19 | 9.30E-18 | 0.450036 | 5.264185 | 9.990033 | 1.67E-09 | 1.07E-08 | 1291.046 | 1302.747 | 1346.221 | 4810.419 | 5024.926 | 4586.896 | 7234.218 | 6612.3 | 6637.523 | 851.5204 | 941.7219 | 872.4729 | 1689.512 | 1669.639 | 1657.804 |
| Gpc3 | 14734 | 1.778683 | -2.34681 | 2.846582 | 0.009565 | 0.023057 | 1.090791 | -2.34681 | 2.341251 | 0.028993 | 0.054335 | 5.935846 | 2.664105 | 4.339848 | 11.68838 | 12.63956 | 28.40602 | 55.42917 | 37.53181 | 25.28142 | 3.846072 | 2.287584 | 0.746341 | 30.8701 | 30.65722 | 36.44416 |
| Hivep2 | 15273 | 1.779857 | 6.658791 | 46.68002 | 5.62E-23 | 4.85E-21 | -1.25175 | 6.658791 | -33.0406 | 8.13E-20 | 5.14E-18 | 6162.892 | 6302.384 | 5686.937 | 21942.99 | 22395.89 | 22206.75 | 9995.728 | 9937.887 | 9165.665 | 2649.944 | 2669.61 | 2530.097 | 2434.566 | 2280.426 | 2376.159 |
| Rhou | 69581 | 1.781067 | 2.389954 | 13.52353 | 6.12E-12 | 5.24E-11 | -1.66645 | 2.389954 | -11.5879 | 1.13E-10 | 8.79E-10 | 211.4645 | 246.8737 | 266.4667 | 812.9919 | 844.0416 | 987.4475 | 314.0986 | 265.4035 | 283.8414 | 293.0707 | 309.5863 | 349.2877 | 208.5817 | 133.634 | 136.0582 |
| Alpk1 | 71481 | 1.781275 | 5.499013 | 33.05441 | 8.06E-20 | 3.22E-18 | -1.82136 | 5.499013 | -30.9563 | 3.18E-19 | 1.67E-17 | 2691.164 | 2683.642 | 2509.3 | 10000.06 | 9649.6 | 9298.238 | 2971.856 | 3064.204 | 2537.336 | 2581.484 | 2645.209 | 2461.433 | 770.9181 | 824.6007 | 746.7004 |
| Naa25 | 231713 | 1.781279 | 6.595838 | 42.58109 | 3.92E-22 | 2.66E-20 | -0.7104 | 6.595838 | -18.6433 | 1.09E-14 | 1.84E-13 | 4664.091 | 4582.26 | 4642.769 | 17432.57 | 16994.59 | 16564.77 | 11296.18 | 10960.63 | 10217.14 | 4330.677 | 4484.426 | 4248.921 | 1704.53 | 1649.987 | 1883.758 |
| Gm16754 | 100503569 | 1.781733 | 1.287199 | 9.838643 | 2.19E-09 | 1.30E-08 | 0.09653 | 1.287199 | 0.6885 | 0.498554 | 0.596419 | 70.48817 | 103.9001 | 98.94853 | 333.7683 | 318.7977 | 325.9929 | 392.268 | 406.1478 | 307.9737 | 85.3828 | 106.7539 | 94.78534 | 60.07154 | 69.17527 | 61.55014 |
| Apob | 238055 | 1.783144 | -0.96263 | 3.34391 | 0.003028 | 0.008032 | 0.622046 | -0.96263 | 1.576851 | 0.129537 | 0.194284 | 16.32358 | 7.10428 | 19.09533 | 76.62384 | 30.8967 | 50.04871 | 55.42917 | 68.36151 | 121.8105 | 3.076858 | 6.100223 | 15.67317 | 69.24913 | 93.54383 | 58.31066 |
| Slc6a20a | 102680 | 1.786018 | -1.82197 | 3.323354 | 0.003178 | 0.008404 | 1.268547 | -1.82197 | 3.270516 | 0.003598 | 0.008719 | 4.451884 | 5.32821 | 6.075787 | 18.18193 | 18.25714 | 24.34802 | 51.16539 | 65.68067 | 41.3696 | 3.846072 | 9.150334 | 9.702436 | 24.19548 | 37.73197 | 29.15533 |
| Mgat4a | 269181 | 1.786661 | 6.990355 | 52.5347 | 4.61E-24 | 5.75E-22 | -0.25421 | 6.990355 | -8.78481 | 1.55E-08 | 8.55E-08 | 4609.926 | 4552.955 | 4431.853 | 16771.53 | 17164.52 | 16195.49 | 14899.08 | 14334.47 | 14408.11 | 3756.074 | 3612.857 | 3782.457 | 5998.811 | 6250.143 | 6342.094 |
| Mafk | 17135 | 1.786787 | 5.411186 | 27.72911 | 3.16E-18 | 8.98E-17 | 0.354653 | 5.411186 | 6.889278 | 7.58E-07 | 3.27E-06 | 1351.147 | 1251.241 | 1183.043 | 4822.107 | 4576.924 | 4566.606 | 6534.957 | 6136.451 | 5926.195 | 1599.966 | 1655.448 | 1747.185 | 1631.943 | 1800.915 | 1846.504 |
| Pcdh10 | 18526 | 1.789766 | -1.06327 | 3.285877 | 0.00347 | 0.009115 | 1.026739 | -1.06327 | 2.637377 | 0.015273 | 0.031188 | 6.677827 | 7.992315 | 20.83127 | 27.27289 | 58.9846 | 36.52203 | 65.378 | 65.68067 | 120.6613 | 10.769 | 11.43792 | 4.478048 | 73.42077 | 76.25001 | 46.16261 |
| Zfp536 | 243937 | 1.791377 | -1.07248 | 3.664905 | 0.001414 | 0.003968 | 1.22124 | -1.07248 | 3.586824 | 0.001703 | 0.004435 | 9.64575 | 15.09659 | 9.547666 | 40.25998 | 46.34504 | 39.22736 | 81.01187 | 104.5529 | 120.6613 | 8.461359 | 1.525056 | 14.18048 | 73.42077 | 57.38403 | 48.59222 |
| Col9a1 | 12839 | 1.791871 | -2.07636 | 2.629598 | 0.015538 | 0.035641 | 1.001677 | -2.07636 | 1.991048 | 0.05943 | 0.100403 | 5.193865 | 3.55214 | 9.547666 | 23.37676 | 16.85274 | 25.70069 | 62.53548 | 21.44675 | 67.80018 | 6.153716 | 0.762528 | 7.463413 | 19.18952 | 27.51289 | 24.29611 |
| Wnt9b | 22412 | 1.792322 | -1.82111 | 2.500586 | 0.020606 | 0.045759 | 1.088544 | -1.82111 | 2.038073 | 0.054125 | 0.092758 | 1.483961 | 10.65642 | 6.943757 | 15.58451 | 42.13186 | 9.468674 | 36.95278 | 37.53181 | 49.41369 | 5.384501 | 16.77561 | 4.478048 | 35.04173 | 55.81187 | 35.63429 |
| B4galt5 | 56336 | 1.793611 | 8.689381 | 55.47953 | 1.45E-24 | 2.13E-22 | -0.81605 | 8.689381 | -27.7104 | 3.20E-18 | 1.25E-16 | 14136.96 | 13753 | 14213 | 52353.56 | 51864.32 | 51682.73 | 32114.81 | 31341.74 | 28907.01 | 25464.08 | 26544.36 | 26026.41 | 15765.44 | 15363.2 | 15926.1 |
| Apaf1 | 11783 | 1.795284 | 7.496937 | 46.91967 | 5.04E-23 | 4.44E-21 | -0.89039 | 7.496937 | -24.9768 | 2.76E-17 | 8.63E-16 | 6703.796 | 6613.196 | 6518.452 | 24667.68 | 25096.54 | 23809.66 | 14111.7 | 14236.62 | 12990.05 | 10331.32 | 10593.8 | 9903.202 | 6555.307 | 6283.158 | 6363.961 |
| Vps54 | 245944 | 1.795643 | 6.736929 | 40.36654 | 1.21E-21 | 7.36E-20 | -0.24148 | 6.736929 | -6.39153 | 2.28E-06 | 9.24E-06 | 4138.027 | 4362.916 | 4253.051 | 15568.93 | 16761.46 | 14959.15 | 13635.58 | 14436.34 | 13440.52 | 3909.148 | 3891.18 | 3791.414 | 2793.327 | 2976.109 | 3071.838 |
| Adamts3 | 330119 | 1.796745 | -0.87555 | 3.74121 | 0.001178 | 0.003353 | 1.466486 | -0.87555 | 4.473733 | 0.000202 | 0.000625 | 5.935846 | 10.65642 | 26.90706 | 58.44191 | 29.4923 | 48.69604 | 119.3859 | 138.0634 | 126.4071 | 9.230573 | 12.20045 | 4.478048 | 65.91183 | 69.96135 | 50.21196 |
| Lrrn2 | 16980 | 1.796969 | -1.30648 | 3.072908 | 0.005699 | 0.01439 | 0.586134 | -1.30648 | 1.336952 | 0.195315 | 0.275058 | 8.903769 | 28.41712 | 5.207818 | 48.05224 | 32.30109 | 44.63804 | 46.90161 | 61.6594 | 87.33582 | 4.615287 | 4.575167 | 10.44878 | 62.57452 | 53.45362 | 21.05663 |
| Cpne6 | 12891 | 1.79946 | -2.09777 | 2.772016 | 0.011315 | 0.026804 | 1.087563 | -2.09777 | 2.234563 | 0.036261 | 0.065872 | 2.225942 | 8.88035 | 4.339848 | 18.18193 | 15.44835 | 17.58468 | 27.00396 | 36.19139 | 54.01031 | 3.846072 | 5.337695 | 5.97073 | 34.2074 | 44.80671 | 17.00728 |
| Gm20186 | 100504344 | 1.799564 | 1.129393 | 7.845411 | 1.01E-07 | 4.75E-07 | 0.923215 | 1.129393 | 5.814596 | 8.46E-06 | 3.17E-05 | 59.35846 | 37.29747 | 64.22975 | 164.9361 | 233.1296 | 200.1948 | 385.1617 | 375.3181 | 401.0553 | 81.53673 | 93.0284 | 88.06827 | 98.45058 | 95.116 | 132.0089 |
| Iqca1l | 231045 | 1.800083 | -2.29784 | 2.536041 | 0.019077 | 0.042746 | 0.968051 | -2.29784 | 1.790862 | 0.08751 | 0.139255 | 11.12971 | 4.440175 | 0.86797 | 12.98709 | 12.63956 | 21.64268 | 28.42522 | 22.78717 | 45.96622 | 7.692145 | 2.287584 | 2.985365 | 30.03577 | 22.01031 | 39.68364 |
| Ccm2l | 228788 | 1.801346 | 0.087197 | 6.791341 | 9.39E-07 | 3.90E-06 | -1.59405 | 0.087197 | -5.58425 | 1.44E-05 | 5.23E-05 | 29.67923 | 65.71459 | 59.8899 | 163.6374 | 203.6373 | 170.4361 | 76.74809 | 38.87223 | 73.54596 | 36.92229 | 25.92595 | 23.88292 | 60.90587 | 67.60311 | 70.45871 |
| Ccdc63 | 330188 | 1.801458 | 0.159205 | 4.419617 | 0.00023 | 0.000722 | 3.380097 | 0.159205 | 13.75478 | 4.42E-12 | 4.41E-11 | 18.54952 | 18.64873 | 10.41564 | 74.02642 | 56.17581 | 47.34337 | 658.0438 | 676.913 | 590.666 | 31.53779 | 12.96297 | 12.6878 | 116.8058 | 143.067 | 117.4312 |
| Pgam2 | 56012 | 1.803148 | 2.854579 | 14.42902 | 1.75E-12 | 1.62E-11 | 0.209209 | 2.854579 | 2.211255 | 0.038056 | 0.068626 | 160.2678 | 152.742 | 138.0072 | 518.185 | 605.2944 | 568.1205 | 652.3587 | 722.4873 | 652.7204 | 399.2223 | 393.4644 | 424.6682 | 659.9526 | 630.4383 | 657.6147 |
| Anks1b | 77531 | 1.80476 | -0.06395 | 5.718286 | 1.06E-05 | 3.89E-05 | 0.294599 | -0.06395 | 1.228911 | 0.232483 | 0.318085 | 20.77546 | 29.30515 | 27.77503 | 79.22126 | 95.49888 | 116.3294 | 122.2284 | 107.2337 | 135.6004 | 18.46115 | 16.01308 | 20.89756 | 112.6341 | 132.0619 | 80.17716 |
| Cntnap5b | 241175 | 1.805193 | -2.35892 | 2.501348 | 0.020572 | 0.045695 | 0.93138 | -2.35892 | 1.696331 | 0.104362 | 0.161965 | 5.193865 | 7.10428 | 1.735939 | 27.27289 | 8.426372 | 17.58468 | 39.7953 | 22.78717 | 37.92213 | 7.692145 | 0.762528 | 0.746341 | 38.37904 | 53.45362 | 29.15533 |
| Fasl | 14103 | 1.805283 | -2.87492 | 2.605031 | 0.016402 | 0.037353 | 0.207364 | -2.87492 | 0.354052 | 0.726774 | 0.805787 | 2.225942 | 0.888035 | 13.88751 | 9.090964 | 22.47032 | 10.82134 | 18.47639 | 12.0638 | 17.23733 | 2.307643 | 2.287584 | 4.478048 | 14.18356 | 11.79124 | 12.95792 |
| Cd2ap | 12488 | 1.805564 | 7.084512 | 50.70167 | 9.78E-24 | 1.09E-21 | -3.04954 | 7.084512 | -61.2389 | 1.78E-25 | 7.01E-23 | 8185.532 | 8065.134 | 7433.292 | 29137.84 | 31232.35 | 28205.83 | 3720.861 | 3493.139 | 3858.864 | 8242.902 | 8085.845 | 8047.798 | 5007.63 | 4990.839 | 5118.38 |
| Mt2 | 17750 | 1.805847 | 7.373215 | 26.21734 | 1.01E-17 | 2.51E-16 | 2.909373 | 7.373215 | 65.84184 | 3.83E-26 | 2.52E-23 | 2433.697 | 2610.823 | 2277.552 | 9089.665 | 9447.367 | 8763.934 | 74006.47 | 71211.25 | 68319.6 | 1263.05 | 1264.271 | 1388.195 | 46497.87 | 46325.42 | 46556.2 |
| C1s1 | 50908 | 1.80705 | -0.7661 | 2.91169 | 0.00825 | 0.020149 | 4.26326 | -0.7661 | 11.38886 | 1.56E-10 | 1.18E-09 | 7.419807 | 4.440175 | 15.62345 | 35.06515 | 12.63956 | 64.92805 | 648.0949 | 617.9344 | 629.7373 | 3.076858 | 3.050111 | 14.92683 | 54.23125 | 65.24486 | 48.59222 |
| Dgkb | 217480 | 1.807668 | -2.09754 | 2.7205 | 0.012699 | 0.029764 | 0.248796 | -2.09754 | 0.466581 | 0.645527 | 0.733264 | 10.38773 | 4.440175 | 4.339848 | 22.07806 | 22.47032 | 25.70069 | 12.79135 | 28.14886 | 57.45778 | 5.384501 | 4.575167 | 1.492683 | 23.36115 | 55.02578 | 26.72572 |
| Rps6ka6 | 67071 | 1.807767 | -2.27212 | 2.466965 | 0.022161 | 0.048832 | 0.962462 | -2.27212 | 1.707273 | 0.102279 | 0.159052 | 3.709904 | 1.77607 | 6.943757 | 27.27289 | 19.66153 | 5.410671 | 22.74017 | 21.44675 | 50.56285 | 1.538429 | 8.387806 | 4.478048 | 38.37904 | 36.94588 | 33.20468 |
| Cyp26a1 | 13082 | 1.809367 | -1.6784 | 2.565707 | 0.01788 | 0.040346 | 2.119518 | -1.6784 | 4.411061 | 0.000235 | 0.000719 | 5.193865 | 16.87266 | 0.86797 | 22.07806 | 11.23516 | 22.99535 | 56.85043 | 101.8721 | 96.52907 | 6.153716 | 6.100223 | 5.97073 | 40.04769 | 53.45362 | 42.92312 |
| Gm15628 | 100043176 | 1.809861 | 0.126963 | 8.606359 | 2.20E-08 | 1.12E-07 | 0.51755 | 0.126963 | 3.375602 | 0.00281 | 0.006978 | 45.26083 | 54.17013 | 40.79457 | 194.8064 | 178.3582 | 151.4988 | 255.827 | 243.9568 | 275.7973 | 30.76858 | 22.11331 | 26.12194 | 22.52683 | 18.0799 | 21.8665 |
| Tcp11 | 21463 | 1.810242 | 0.138387 | 5.876862 | 7.33E-06 | 2.75E-05 | 1.263587 | 0.138387 | 6.07059 | 4.70E-06 | 1.83E-05 | 25.22735 | 23.08891 | 18.22736 | 97.40319 | 88.4769 | 67.63339 | 187.6064 | 241.2759 | 203.4005 | 37.69151 | 33.55123 | 32.09267 | 84.26702 | 80.18043 | 104.4733 |
| Gramd2 | 546134 | 1.810363 | -2.65083 | 2.872234 | 0.009024 | 0.021866 | 0.384855 | -2.65083 | 0.750682 | 0.461041 | 0.559774 | 5.193865 | 2.664105 | 3.471878 | 14.2858 | 12.63956 | 17.58468 | 17.05513 | 33.51054 | 14.93902 | 5.384501 | 6.100223 | 2.985365 | 14.18356 | 18.0799 | 5.669092 |
| Ifi35 | 70110 | 1.81093 | 4.318386 | 23.93416 | 6.65E-17 | 1.42E-15 | -0.1186 | 4.318386 | -1.87704 | 0.074251 | 0.121206 | 716.7534 | 782.3588 | 859.2899 | 3051.967 | 2831.261 | 2896.062 | 3058.553 | 2629.907 | 2738.438 | 911.5191 | 1021.787 | 962.0339 | 368.7725 | 402.4743 | 394.4068 |
| Mir125a | 387235 | 1.810964 | -3.20508 | 2.608949 | 0.016261 | 0.037097 | -0.17726 | -3.20508 | -0.29933 | 0.767585 | 0.841422 | 4.451884 | 2.664105 | 10.41564 | 15.58451 | 30.8967 | 16.23201 | 24.16143 | 18.7659 | 13.78987 | 3.846072 | 6.100223 | 0.746341 | 5.840288 | 0 | 1.619741 |
| Gpr55 | 227326 | 1.811855 | -0.66627 | 4.895886 | 7.34E-05 | 0.000244 | -1.35864 | -0.66627 | -3.63329 | 0.001525 | 0.004014 | 11.87169 | 15.98463 | 19.9633 | 68.83159 | 43.53625 | 66.28072 | 29.84648 | 18.7659 | 22.98311 | 42.30679 | 61.76476 | 50.75121 | 77.5924 | 58.17011 | 59.12053 |
| Batf | 53314 | 1.811998 | 3.844233 | 22.40981 | 2.57E-16 | 4.88E-15 | -1.78157 | 3.844233 | -20.1357 | 2.29E-15 | 4.53E-14 | 850.3099 | 749.5015 | 732.5663 | 2670.146 | 3091.074 | 3034.034 | 852.7565 | 990.5717 | 819.3479 | 1436.893 | 1438.128 | 1619.561 | 168.534 | 155.6444 | 140.9174 |
| Art2a | 11871 | 1.81263 | 2.104219 | 10.78367 | 4.24E-10 | 2.78E-09 | -0.37943 | 2.104219 | -2.72403 | 0.012599 | 0.026416 | 145.4282 | 157.1822 | 105.8923 | 546.7566 | 505.5823 | 466.6704 | 368.1066 | 455.7434 | 391.8621 | 115.3822 | 163.9435 | 137.3268 | 261.1443 | 284.5619 | 345.0047 |
| Gstt1 | 14871 | 1.81348 | 1.042 | 8.855001 | 1.36E-08 | 7.18E-08 | -0.91124 | 1.042 | -4.82207 | 8.76E-05 | 0.000286 | 51.19667 | 81.69922 | 59.02193 | 220.7806 | 242.9604 | 238.0695 | 113.7009 | 147.4464 | 126.4071 | 234.6104 | 194.4446 | 217.1853 | 58.40288 | 84.11084 | 64.78962 |
| Ric1 | 226089 | 1.813531 | 7.883851 | 40.47592 | 1.14E-21 | 6.98E-20 | 1.004721 | 7.883851 | 30.94409 | 3.21E-19 | 1.67E-17 | 4178.836 | 4577.82 | 4278.222 | 16294.9 | 16497.43 | 16012.88 | 34195.54 | 34590.92 | 33017.54 | 10088.25 | 10077.57 | 9382.256 | 23650.67 | 23874.9 | 23807.76 |
| Bpifa6 | 545477 | 1.813814 | -3.23246 | 2.700781 | 0.013269 | 0.03095 | 0.969142 | -3.23246 | 1.781021 | 0.089148 | 0.141536 | 2.225942 | 3.55214 | 0.86797 | 14.2858 | 5.617581 | 6.763339 | 11.37009 | 26.80843 | 17.23733 | 2.307643 | 1.525056 | 1.492683 | 13.34923 | 11.00516 | 12.95792 |
| Tmem184l | 223693 | 1.81437 | 7.678197 | 57.08575 | 7.92E-25 | 1.26E-22 | -0.314 | 7.678197 | -11.6913 | 9.60E-11 | 7.53E-10 | 7393.096 | 7045.67 | 7316.116 | 27156.01 | 27967.13 | 26677.31 | 22983.21 | 23217.45 | 22175.26 | 8350.592 | 8688.242 | 8362.007 | 7077.595 | 7125.053 | 7439.468 |
| Gstt2 | 14872 | 1.816977 | 2.37032 | 9.765319 | 2.50E-09 | 1.47E-08 | 0.019625 | 2.37032 | 0.136421 | 0.892767 | 0.918721 | 95.71552 | 71.0428 | 94.60869 | 314.2876 | 352.5032 | 319.2296 | 362.4215 | 361.9139 | 318.3161 | 346.1465 | 397.277 | 372.4243 | 769.2494 | 687.0362 | 845.5046 |
| Cadm3 | 94332 | 1.817028 | -0.90992 | 3.179056 | 0.004455 | 0.011468 | 1.828111 | -0.90992 | 4.833744 | 8.52E-05 | 0.000278 | 10.38773 | 4.440175 | 13.88751 | 33.76644 | 21.06593 | 54.10671 | 83.85439 | 160.8506 | 144.7936 | 10.769 | 9.150334 | 17.91219 | 76.75808 | 52.66754 | 70.45871 |
| Stfa1 | 20861 | 1.817059 | -2.48959 | 2.901722 | 0.008439 | 0.020581 | -0.54366 | -2.48959 | -0.95272 | 0.351398 | 0.448476 | 3.709904 | 5.32821 | 6.943757 | 16.88322 | 22.47032 | 21.64268 | 9.948826 | 5.361687 | 42.51876 | 13.07665 | 11.43792 | 16.41951 | 9.177596 | 3.930413 | 4.859222 |
| Itpka | 228550 | 1.817146 | -1.46534 | 2.951892 | 0.007527 | 0.018537 | 1.183922 | -1.46534 | 2.635947 | 0.015321 | 0.03127 | 2.967923 | 12.43249 | 3.471878 | 32.46773 | 21.06593 | 10.82134 | 38.37404 | 61.6594 | 44.81707 | 22.30722 | 21.35078 | 14.18048 | 50.05962 | 51.88145 | 36.44416 |
| Gata3 | 14462 | 1.820896 | -0.01252 | 3.616976 | 0.001585 | 0.004414 | 1.616945 | -0.01252 | 4.894061 | 7.38E-05 | 0.000243 | 25.96933 | 4.440175 | 19.9633 | 75.32513 | 56.17581 | 35.16936 | 154.9174 | 160.8506 | 195.3565 | 55.38344 | 44.22662 | 49.25852 | 146.0072 | 94.32991 | 113.3818 |
| Marchf1 | 72925 | 1.823323 | 5.582647 | 46.89306 | 5.10E-23 | 4.48E-21 | -4.43413 | 5.582647 | -59.5156 | 3.27E-25 | 1.23E-22 | 4634.412 | 5017.398 | 4396.266 | 17720.89 | 18139.17 | 17126.13 | 901.0794 | 833.7423 | 815.9005 | 5141.429 | 4803.163 | 5063.179 | 559.8334 | 533.7501 | 553.9513 |
| Gm4952 | 240549 | 1.823783 | -2.60843 | 3.118512 | 0.005128 | 0.013043 | 0.150264 | -2.60843 | 0.309189 | 0.760176 | 0.834971 | 2.967923 | 4.440175 | 5.207818 | 15.58451 | 12.63956 | 21.64268 | 27.00396 | 14.74464 | 17.23733 | 8.461359 | 1.525056 | 2.985365 | 16.68654 | 16.50774 | 12.95792 |
| Slc32a1 | 22348 | 1.824401 | -1.41555 | 2.942276 | 0.007694 | 0.018912 | -0.00448 | -1.41555 | -0.00889 | 0.992989 | 0.99416 | 8.161788 | 10.65642 | 8.679696 | 38.96127 | 23.87472 | 45.9907 | 15.63387 | 53.61687 | 51.712 | 16.1535 | 1.525056 | 10.44878 | 58.40288 | 82.53868 | 59.12053 |
| Cst7 | 13011 | 1.825667 | 1.393482 | 9.089741 | 8.70E-09 | 4.73E-08 | -1.84546 | 1.393482 | -8.15006 | 5.42E-08 | 2.76E-07 | 112.7811 | 86.13939 | 109.3642 | 454.5482 | 393.2307 | 328.6983 | 81.01187 | 135.3826 | 124.1088 | 607.6794 | 613.8349 | 548.5608 | 60.90587 | 23.58248 | 32.39481 |
| Gm10862 | 100038349 | 1.8272 | -2.92199 | 2.573249 | 0.017588 | 0.039757 | 1.187749 | -2.92199 | 2.154269 | 0.042789 | 0.075836 | 2.967923 | 1.77607 | 2.603909 | 12.98709 | 5.617581 | 13.52668 | 31.26774 | 17.42548 | 27.57973 | 3.846072 | 0.762528 | 2.985365 | 8.343269 | 22.01031 | 19.43689 |
| Gadl1 | 73748 | 1.827516 | -2.8386 | 2.473387 | 0.021856 | 0.048257 | 0.55408 | -2.8386 | 0.930803 | 0.362365 | 0.460184 | 6.677827 | 1.77607 | 2.603909 | 10.38967 | 16.85274 | 13.52668 | 18.47639 | 21.44675 | 21.83396 | 5.384501 | 0 | 4.478048 | 18.35519 | 21.22423 | 12.14805 |
| Tmigd1 | 66601 | 1.828226 | -1.66994 | 5.238322 | 3.25E-05 | 0.000113 | -1.261 | -1.66994 | -3.67073 | 0.001394 | 0.003702 | 19.2915 | 24.86498 | 13.01954 | 62.33804 | 81.45493 | 70.33872 | 44.05909 | 21.44675 | 29.87805 | 6.92293 | 5.337695 | 5.97073 | 4.171635 | 12.57732 | 10.52831 |
| Enthd1 | 383075 | 1.828542 | -0.94604 | 3.677569 | 0.001372 | 0.00386 | 3.229126 | -0.94604 | 10.40636 | 8.07E-10 | 5.44E-09 | 9.64575 | 5.32821 | 4.339848 | 25.97418 | 29.4923 | 18.93735 | 228.823 | 243.9568 | 255.1125 | 18.46115 | 19.0632 | 30.59999 | 38.37904 | 31.4433 | 26.72572 |
| Dnaja2 | 56445 | 1.829794 | 7.756456 | 57.89789 | 5.87E-25 | 9.83E-23 | 0.194852 | 7.756456 | 7.897525 | 9.04E-08 | 4.46E-07 | 7554.106 | 7986.987 | 7705.834 | 29044.33 | 30759.07 | 28393.85 | 35309.8 | 33998.46 | 35319.3 | 7396.766 | 7102.184 | 7435.798 | 6362.577 | 6263.506 | 6027.055 |
| Cckbr | 12426 | 1.83076 | -2.36506 | 2.913713 | 0.008212 | 0.020064 | 0.745381 | -2.36506 | 1.568758 | 0.131412 | 0.196703 | 8.161788 | 3.55214 | 4.339848 | 19.48064 | 16.85274 | 25.70069 | 32.689 | 33.51054 | 41.3696 | 3.076858 | 3.050111 | 0 | 25.86413 | 45.59279 | 23.48624 |
| Aadacl2fm | 433597 | 1.832147 | -4.27104 | 2.656222 | 0.01465 | 0.03381 | -0.76626 | -4.27104 | -1.15802 | 0.259649 | 0.349061 | 1.483961 | 2.664105 | 0.86797 | 10.38967 | 4.213186 | 6.763339 | 4.263783 | 4.021265 | 3.447467 | 0 | 0.762528 | 1.492683 | 3.337308 | 5.502578 | 4.049351 |
| Ros1 | 19886 | 1.832458 | -1.38876 | 3.103364 | 0.005311 | 0.013483 | 1.038811 | -1.38876 | 2.458498 | 0.02257 | 0.043763 | 8.161788 | 4.440175 | 10.41564 | 19.48064 | 26.68351 | 45.9907 | 46.90161 | 54.95729 | 86.18667 | 13.07665 | 12.20045 | 3.731706 | 52.5626 | 65.24486 | 36.44416 |
| Gm14204 | 100045469 | 1.834304 | -2.05263 | 2.86748 | 0.009122 | 0.02208 | 0.315094 | -2.05263 | 0.621304 | 0.540979 | 0.636101 | 8.903769 | 1.77607 | 6.943757 | 25.97418 | 32.30109 | 9.468674 | 42.63783 | 20.10633 | 21.83396 | 6.153716 | 6.100223 | 9.702436 | 30.03577 | 32.22939 | 20.24676 |
| Hook2 | 170833 | 1.834466 | 3.809535 | 16.98279 | 7.06E-14 | 8.39E-13 | -0.29118 | 3.809535 | -3.26159 | 0.003674 | 0.00888 | 372.4743 | 381.855 | 371.491 | 1511.697 | 1397.373 | 1378.368 | 1355.883 | 1230.507 | 1089.4 | 772.2913 | 792.2664 | 926.9558 | 839.3329 | 745.2063 | 706.2069 |
| H2-T24 | 15042 | 1.83561 | 6.538356 | 37.23604 | 6.61E-21 | 3.53E-19 | -0.93978 | 6.538356 | -20.3608 | 1.82E-15 | 3.68E-14 | 4093.508 | 4132.027 | 3793.895 | 15724.77 | 15067.76 | 15005.14 | 8621.368 | 8592.103 | 7695.895 | 5480.653 | 5400.985 | 5231.852 | 2118.356 | 1896.031 | 1831.117 |
| Fgf1 | 14164 | 1.835984 | -2.24832 | 2.503397 | 0.020481 | 0.045511 | 1.721373 | -2.24832 | 3.235263 | 0.003907 | 0.009378 | 7.419807 | 1.77607 | 1.735939 | 9.090964 | 15.44835 | 13.52668 | 31.26774 | 30.8297 | 78.14258 | 6.92293 | 4.575167 | 4.478048 | 43.385 | 24.36856 | 24.29611 |
| Sbno2 | 216161 | 1.839515 | 7.680549 | 48.22158 | 2.83E-23 | 2.74E-21 | -1.23654 | 7.680549 | -33.1093 | 7.79E-20 | 4.96E-18 | 9974.447 | 10382.9 | 9367.128 | 37889.84 | 37960.8 | 37639.33 | 17488.61 | 17370.53 | 15432.01 | 7708.298 | 7948.59 | 7918.681 | 5564.961 | 5545.813 | 5780.044 |
| Olfr658 | 259051 | 1.840671 | -1.94634 | 3.245912 | 0.003811 | 0.009944 | 0.67167 | -1.94634 | 1.584079 | 0.127882 | 0.192185 | 8.903769 | 6.216245 | 4.339848 | 28.5716 | 22.47032 | 24.34802 | 41.21656 | 49.5956 | 36.77298 | 3.846072 | 5.337695 | 18.65853 | 15.01788 | 25.15464 | 9.718443 |
| Lman1l | 235416 | 1.840913 | -1.64953 | 3.677307 | 0.001373 | 0.003862 | -0.14392 | -1.64953 | -0.34532 | 0.733237 | 0.811618 | 6.677827 | 5.32821 | 7.811726 | 22.07806 | 29.4923 | 27.05335 | 28.42522 | 20.10633 | 25.28142 | 9.230573 | 25.92595 | 18.65853 | 27.53279 | 33.01547 | 34.82442 |
| Gm19816 | 100503654 | 1.841308 | 4.038038 | 27.31303 | 4.32E-18 | 1.18E-16 | -3.21337 | 4.038038 | -31.4119 | 2.34E-19 | 1.27E-17 | 1087.002 | 1080.739 | 1128.36 | 4170.155 | 4491.256 | 3968.727 | 567.0831 | 426.2541 | 437.8283 | 685.3701 | 790.7414 | 706.0388 | 528.9633 | 628.08 | 568.5289 |
| Myom2 | 17930 | 1.841931 | -1.60885 | 3.552007 | 0.00185 | 0.005096 | 0.555705 | -1.60885 | 1.432306 | 0.16654 | 0.240326 | 12.61367 | 11.54445 | 3.471878 | 48.05224 | 23.87472 | 28.40602 | 36.95278 | 56.29771 | 56.30862 | 7.692145 | 8.387806 | 4.478048 | 21.6925 | 35.37372 | 34.01455 |
| Dnah7a | 627872 | 1.84336 | -0.6154 | 4.478265 | 0.0002 | 0.000632 | 0.850178 | -0.6154 | 2.926432 | 0.007977 | 0.017694 | 18.54952 | 18.64873 | 10.41564 | 49.35095 | 57.58021 | 77.10206 | 133.5985 | 80.4253 | 132.1529 | 10.769 | 8.387806 | 4.478048 | 123.4804 | 110.0516 | 88.27586 |
| Mir30d | 387228 | 1.843362 | -4.50921 | 2.521561 | 0.019689 | 0.043943 | -3.54094 | -4.50921 | -4.11995 | 0.000474 | 0.001375 | 3.709904 | 1.77607 | 1.735939 | 9.090964 | 12.63956 | 8.116006 | 0 | 1.340422 | 0 | 0 | 3.812639 | 1.492683 | 5.840288 | 1.572165 | 2.429611 |
| Arhgef28 | 110596 | 1.843822 | -0.7552 | 4.960829 | 6.29E-05 | 0.000211 | 0.191514 | -0.7552 | 0.673243 | 0.508018 | 0.605299 | 23.74338 | 19.53677 | 9.547666 | 58.44191 | 58.9846 | 79.8074 | 62.53548 | 79.08488 | 89.63414 | 4.615287 | 10.67539 | 9.702436 | 87.60433 | 70.74744 | 79.36729 |
| Tshr | 22095 | 1.845067 | -1.31677 | 3.797314 | 0.00103 | 0.002959 | 0.714741 | -1.31677 | 2.001853 | 0.058172 | 0.098582 | 7.419807 | 12.43249 | 5.207818 | 27.27289 | 26.68351 | 40.58003 | 59.69296 | 61.6594 | 42.51876 | 13.07665 | 19.0632 | 7.463413 | 39.21337 | 47.16496 | 44.54286 |
| Pde1b | 18574 | 1.84535 | 6.44903 | 34.12267 | 4.14E-20 | 1.79E-18 | -0.98568 | 6.44903 | -19.3763 | 4.99E-15 | 9.09E-14 | 4378.428 | 4346.043 | 4354.604 | 16866.34 | 16199.7 | 17130.18 | 9339.105 | 9389.654 | 7829.197 | 3686.845 | 3952.944 | 3650.355 | 1621.932 | 1620.116 | 1571.958 |
| Epcam | 17075 | 1.845441 | -1.64689 | 2.95515 | 0.007471 | 0.018408 | 2.012758 | -1.64689 | 4.719473 | 0.000112 | 0.00036 | 3.709904 | 2.664105 | 8.679696 | 18.18193 | 19.66153 | 16.23201 | 68.22052 | 68.36151 | 95.37991 | 12.30743 | 5.337695 | 3.731706 | 76.75808 | 45.59279 | 44.54286 |
| Prpf38a | 230596 | 1.845655 | 4.765363 | 28.58435 | 1.68E-18 | 5.01E-17 | -1.83501 | 4.765363 | -26.0927 | 1.12E-17 | 3.82E-16 | 1457.992 | 1373.79 | 1306.294 | 5589.644 | 5166.77 | 5146.901 | 1547.753 | 1528.081 | 1547.913 | 1389.201 | 1473.204 | 1547.165 | 796.7822 | 596.6367 | 679.4812 |
| Aff1 | 17355 | 1.845668 | 8.945729 | 49.78244 | 2.10E-20 | 9.85E-19 | -0.85025 | 8.945729 | -25.3919 | 2.52E-15 | 4.95E-14 | 17570.1 | 17919.66 | 17297.77 | 68999.12 | 68330.85 | 65202.64 | 40562.78 | 40429.8 | 36290.33 | 30339.36 | 30627.69 | 30797.77 | 15445.06 | 15468.53 | 16153.67 |
| A630014C | 399633 | 1.8479 | -2.22863 | 3.361019 | 0.002908 | 0.007737 | 0.996392 | -2.22863 | 2.400753 | 0.025547 | 0.048742 | 2.967923 | 4.440175 | 4.339848 | 10.38967 | 19.66153 | 18.93735 | 29.84648 | 34.85097 | 34.47467 | 6.153716 | 3.050111 | 5.97073 | 33.37308 | 23.58248 | 25.10598 |
| F11r | 16456 | 1.848023 | 4.739049 | 27.98963 | 2.60E-18 | 7.52E-17 | 1.635467 | 4.739049 | 35.50556 | 1.80E-20 | 1.38E-18 | 801.3392 | 805.4477 | 828.043 | 3123.396 | 3266.623 | 2975.869 | 10164.86 | 10094.72 | 9935.599 | 289.2246 | 216.5579 | 281.3707 | 1513.469 | 1580.812 | 1584.106 |
| Lrp11 | 237253 | 1.84911 | 4.230166 | 22.15457 | 3.25E-16 | 6.05E-15 | -0.85571 | 4.230166 | -11.1708 | 2.23E-10 | 1.64E-09 | 658.8789 | 745.9494 | 810.6836 | 2827.29 | 2900.076 | 2729.684 | 1749.572 | 1668.825 | 1468.621 | 991.5174 | 863.1815 | 1041.146 | 574.8512 | 511.7398 | 555.571 |
| Grin2a | 14811 | 1.849919 | -2.74037 | 2.765924 | 0.011471 | 0.027144 | 0.304333 | -2.74037 | 0.548819 | 0.588826 | 0.682324 | 9.64575 | 0.888035 | 2.603909 | 15.58451 | 12.63956 | 10.82134 | 15.63387 | 18.7659 | 16.08818 | 3.846072 | 3.812639 | 7.463413 | 8.343269 | 18.86598 | 14.57766 |
| Adora2b | 11541 | 1.850176 | 3.568775 | 16.29716 | 1.60E-13 | 1.78E-12 | 1.401775 | 3.568775 | 18.08722 | 2.00E-14 | 3.20E-13 | 439.2526 | 318.8046 | 300.3175 | 1402.606 | 1322.94 | 1341.846 | 4042.066 | 3550.777 | 3610.647 | 572.2956 | 550.5451 | 603.0437 | 181.8833 | 188.6598 | 161.1642 |
| Crlf3 | 54394 | 1.85148 | 7.278804 | 58.89961 | 4.08E-25 | 7.40E-23 | -0.42256 | 7.278804 | -15.6279 | 3.67E-13 | 4.47E-12 | 4999.466 | 5198.557 | 5318.918 | 19823.5 | 20467.66 | 19428.37 | 15558.54 | 15527.45 | 15175.75 | 6377.557 | 6309.918 | 6355.842 | 6874.854 | 6860.929 | 6782.663 |
| Acnat1 | 230161 | 1.852769 | -2.28896 | 2.507714 | 0.02029 | 0.045163 | 0.684078 | -2.28896 | 1.18659 | 0.24843 | 0.33623 | 2.967923 | 12.43249 | 1.735939 | 16.88322 | 39.32307 | 6.763339 | 14.21261 | 32.17012 | 43.66791 | 3.846072 | 3.812639 | 4.478048 | 27.53279 | 34.58764 | 22.67637 |
| Treml2 | 328833 | 1.852827 | 5.62573 | 26.16001 | 1.06E-17 | 2.62E-16 | 1.737964 | 5.62573 | 35.53129 | 1.77E-20 | 1.37E-18 | 778.3378 | 786.799 | 734.3023 | 2925.992 | 2761.041 | 3199.059 | 10430.63 | 10116.16 | 10184.97 | 1783.039 | 1746.189 | 1733.004 | 5446.486 | 5348.506 | 5701.487 |
| Trim12a | 76681 | 1.853567 | 5.116664 | 27.49096 | 3.78E-18 | 1.06E-16 | -0.47059 | 5.116664 | -7.98671 | 7.54E-08 | 3.76E-07 | 1186.427 | 1261.898 | 998.165 | 4477.949 | 4409.801 | 4359.648 | 3395.392 | 3460.969 | 3106.168 | 1411.509 | 1361.112 | 1352.37 | 1680.334 | 1667.281 | 1645.656 |
| Ctrl | 109660 | 1.853676 | 0.727174 | 6.617338 | 1.38E-06 | 5.62E-06 | 1.284182 | 0.727174 | 6.904687 | 7.33E-07 | 3.17E-06 | 36.35706 | 36.40943 | 59.02193 | 216.8844 | 144.6527 | 140.6774 | 478.9649 | 427.5945 | 366.5806 | 54.61423 | 58.71464 | 40.30243 | 64.24317 | 72.3196 | 50.21196 |
| Gm16630 | 100503622 | 1.854008 | -1.979 | 3.100361 | 0.005348 | 0.013573 | 1.088891 | -1.979 | 2.479587 | 0.021565 | 0.04212 | 5.193865 | 1.77607 | 8.679696 | 23.37676 | 16.85274 | 14.87935 | 42.63783 | 44.23392 | 36.77298 | 3.076858 | 9.150334 | 6.717071 | 41.71635 | 37.73197 | 25.10598 |
| Snord123 | 66835 | 1.856519 | -2.15182 | 2.607534 | 0.016312 | 0.037193 | 0.995239 | -2.15182 | 1.868062 | 0.075545 | 0.123028 | 5.193865 | 7.992315 | 1.735939 | 15.58451 | 9.830767 | 33.81669 | 22.74017 | 36.19139 | 55.15947 | 4.615287 | 1.525056 | 7.463413 | 27.53279 | 48.73712 | 28.34546 |
| Galntl6 | 270049 | 1.856844 | -3.27073 | 2.81104 | 0.010364 | 0.024776 | 0.243329 | -3.27073 | 0.420042 | 0.678653 | 0.762528 | 2.967923 | 1.77607 | 0.86797 | 7.792255 | 8.426372 | 6.763339 | 8.527565 | 5.361687 | 17.23733 | 3.076858 | 2.287584 | 2.985365 | 15.85221 | 18.86598 | 13.76779 |
| Spryd7 | 66674 | 1.85742 | 5.055109 | 35.87588 | 1.45E-20 | 7.03E-19 | -0.97007 | 5.055109 | -19.9309 | 2.81E-15 | 5.48E-14 | 1596.743 | 1440.393 | 1604.876 | 5997.439 | 5954.636 | 6020.724 | 3270.321 | 3111.119 | 3147.537 | 1345.356 | 1407.626 | 1513.58 | 790.9419 | 709.0465 | 725.6438 |
| Mylk4 | 238564 | 1.860254 | -1.78497 | 2.469065 | 0.022061 | 0.048664 | 0.998703 | -1.78497 | 1.750867 | 0.094331 | 0.148387 | 2.225942 | 13.32052 | 1.735939 | 27.27289 | 9.830767 | 13.52668 | 34.11026 | 25.46801 | 40.22045 | 3.076858 | 2.287584 | 0 | 493.9215 | 411.1212 | 458.3866 |
| Trdn | 76757 | 1.861173 | -2.39234 | 2.831791 | 0.00989 | 0.023783 | 0.970484 | -2.39234 | 1.972466 | 0.061651 | 0.103596 | 1.483961 | 3.55214 | 13.88751 | 12.98709 | 21.06593 | 18.93735 | 36.95278 | 46.91476 | 27.57973 | 3.076858 | 2.287584 | 1.492683 | 22.52683 | 33.80155 | 16.19741 |
| Synpo2 | 118449 | 1.861445 | 1.00544 | 8.084204 | 6.19E-08 | 3.00E-07 | -1.7603 | 1.00544 | -6.99632 | 6.01E-07 | 2.64E-06 | 94.97354 | 67.49066 | 72.90945 | 227.2741 | 308.967 | 411.211 | 103.752 | 69.70193 | 109.1698 | 79.9983 | 89.21576 | 82.84388 | 109.2968 | 150.1418 | 139.2977 |
| Sytl3 | 83672 | 1.861474 | -0.47119 | 5.331391 | 2.61E-05 | 9.19E-05 | 0.061375 | -0.47119 | 0.226912 | 0.822652 | 0.889698 | 26.71131 | 15.09659 | 34.71878 | 109.0916 | 78.64613 | 100.0974 | 89.53943 | 84.44657 | 137.8987 | 20.76879 | 6.100223 | 6.717071 | 60.90587 | 61.31444 | 64.78962 |
| 2410004P( | 73667 | 1.862319 | -2.24271 | 2.506086 | 0.020362 | 0.045293 | 1.603629 | -2.24271 | 2.841799 | 0.009669 | 0.020948 | 0.741981 | 1.77607 | 5.207818 | 10.38967 | 4.213186 | 13.52668 | 31.26774 | 21.44675 | 33.32551 | 13.84586 | 16.01308 | 20.15121 | 25.86413 | 22.7964 | 43.73299 |
| Ankrd45 | 73844 | 1.862483 | -2.28164 | 2.605035 | 0.016402 | 0.037353 | 0.658167 | -2.28164 | 1.204951 | 0.241413 | 0.328322 | 6.677827 | 3.55214 | 4.339848 | 24.67547 | 15.44835 | 18.93735 | 34.11026 | 29.48928 | 33.32551 | 9.230573 | 6.862751 | 0 | 30.03577 | 29.08506 | 24.29611 |
| Qrfprl | 243407 | 1.864374 | -2.67205 | 2.60981 | 0.016231 | 0.037039 | 1.059842 | -2.67205 | 1.866486 | 0.075774 | 0.123348 | 2.967923 | 1.77607 | 1.735939 | 9.090964 | 12.63956 | 6.763339 | 11.37009 | 17.42548 | 39.07129 | 5.384501 | 7.625279 | 2.985365 | 22.52683 | 35.37372 | 21.8665 |
| Prnp | 19122 | 1.864411 | 4.948838 | 26.15879 | 1.06E-17 | 2.62E-16 | 1.310973 | 4.948838 | 26.00731 | 1.19E-17 | 4.07E-16 | 580.9709 | 589.6552 | 616.2584 | 2351.962 | 2307.421 | 2280.598 | 5957.925 | 6029.217 | 5895.168 | 1323.049 | 1369.5 | 1256.839 | 2006.556 | 1870.091 | 2075.697 |
| D630029K( | 103175 | 1.864541 | -2.3716 | 2.824272 | 0.010059 | 0.02414 | 0.671334 | -2.3716 | 1.304601 | 0.205922 | 0.287425 | 3.709904 | 8.88035 | 1.735939 | 15.58451 | 15.44835 | 17.58468 | 22.74017 | 25.46801 | 33.32551 | 10.769 | 6.862751 | 1.492683 | 25.86413 | 14.93557 | 17.00728 |
| Spdya | 70891 | 1.865138 | 0.297517 | 6.096128 | 4.44E-06 | 1.71E-05 | 0.901743 | 0.297517 | 4.308365 | 0.000301 | 0.000903 | 52.68063 | 51.50603 | 19.9633 | 124.6761 | 175.5494 | 156.9095 | 281.4096 | 345.8288 | 260.8583 | 35.38386 | 26.68847 | 22.39024 | 53.39692 | 41.66238 | 38.87377 |
| Adgrb1 | 107831 | 1.865719 | 0.121124 | 5.773028 | 9.31E-06 | 3.45E-05 | 0.348977 | 0.121124 | 1.46375 | 0.157836 | 0.230075 | 33.38913 | 36.40943 | 26.03909 | 138.9619 | 115.1604 | 119.0348 | 200.3978 | 143.4251 | 157.4343 | 36.92229 | 15.25056 | 8.209754 | 85.10135 | 121.8428 | 102.0437 |
| Socs7 | 192157 | 1.865933 | 5.782065 | 30.59441 | 4.07E-19 | 1.39E-17 | 0.270411 | 5.782065 | 5.573423 | 1.48E-05 | 5.36E-05 | 1911.342 | 1842.673 | 1719.448 | 7381.863 | 7121.688 | 6821.503 | 9083.278 | 9459.356 | 8284.263 | 1978.42 | 1946.734 | 2022.585 | 1654.47 | 1626.405 | 1756.609 |
| Dnajb6 | 23950 | 1.868761 | 6.130362 | 14.17367 | 1.61E-08 | 8.41E-08 | 0.018906 | 6.130362 | 0.175589 | 0.86374 | 0.918721 | 2727.521 | 2695.186 | 1696.013 | 9375.381 | 9266.2 | 8640.842 | 10163.44 | 9773.015 | 8901.359 | 3031.474 | 3119.501 | 3016.711 | 2165.913 | 2046.959 | 2034.394 |
| Dgat2 | 67800 | 1.868822 | 2.380773 | 12.92084 | 1.46E-11 | 1.18E-10 | -0.50144 | 2.380773 | -4.14503 | 0.000446 | 0.0013 | 233.7239 | 190.0395 | 196.1611 | 815.5894 | 724.668 | 894.1134 | 635.3036 | 616.594 | 542.4014 | 209.9955 | 151.743 | 121.6536 | 200.2385 | 175.2964 | 183.0307 |
| Gvin-ps6 | 209380 | 1.869611 | 0.734894 | 6.182457 | 3.65E-06 | 1.42E-05 | 0.128965 | 0.734894 | 0.560629 | 0.580889 | 0.674553 | 45.26083 | 31.96926 | 24.30315 | 138.9619 | 127.8 | 125.7981 | 115.1221 | 139.4039 | 193.0581 | 58.4603 | 72.44015 | 81.3512 | 248.6294 | 219.3171 | 184.6504 |
| Gabrg1 | 14405 | 1.87059 | -2.68223 | 2.636296 | 0.01531 | 0.03521 | 0.906967 | -2.68223 | 1.599843 | 0.124333 | 0.187655 | 2.225942 | 1.77607 | 3.471878 | 14.2858 | 11.23516 | 6.763339 | 12.79135 | 14.74464 | 41.3696 | 7.692145 | 2.287584 | 4.478048 | 30.03577 | 19.65207 | 20.24676 |
| Tnr | 21960 | 1.873115 | -1.47359 | 3.551043 | 0.001855 | 0.005105 | 1.38016 | -1.47359 | 3.732534 | 0.001203 | 0.00324 | 6.677827 | 3.55214 | 6.075787 | 23.37676 | 29.4923 | 14.87935 | 58.27169 | 61.6594 | 59.75609 | 10.769 | 10.67539 | 9.702436 | 56.73423 | 70.74744 | 42.11325 |
| Dnai2 | 432611 | 1.874153 | -2.02041 | 2.508462 | 0.020257 | 0.045094 | 2.014433 | -2.02041 | 3.875603 | 0.000854 | 0.002371 | 11.12971 | 0.888035 | 2.603909 | 20.77935 | 18.25714 | 6.763339 | 65.378 | 53.61687 | 62.0544 | 14.61507 | 2.287584 | 4.478048 | 37.54471 | 36.94588 | 38.0639 |
| Tgif2 | 228839 | 1.874306 | 1.932325 | 10.85851 | 3.74E-10 | 2.47E-09 | -0.99538 | 1.932325 | -6.28715 | 2.88E-06 | 1.15E-05 | 123.1688 | 130.5411 | 140.6111 | 466.2366 | 601.0812 | 485.6077 | 282.8309 | 325.7225 | 214.8921 | 179.9962 | 213.5078 | 233.6048 | 176.8773 | 177.6547 | 132.8187 |
| Drc1 | 381738 | 1.874333 | -2.4619 | 3.203428 | 0.004209 | 0.010884 | 0.479245 | -2.4619 | 1.036037 | 0.311782 | 0.406633 | 2.967923 | 4.440175 | 6.075787 | 28.5716 | 15.44835 | 12.17401 | 22.74017 | 22.78717 | 32.17636 | 3.076858 | 3.812639 | 2.239024 | 30.8701 | 18.0799 | 12.95792 |
| Trim56 | 384309 | 1.877676 | 5.555603 | 34.76049 | 2.81E-20 | 1.28E-18 | -1.37049 | 5.555603 | -25.449 | 1.87E-17 | 6.09E-16 | 2150.26 | 2118.851 | 2018.897 | 8211.738 | 8177.794 | 8301.322 | 3479.247 | 3474.373 | 3021.13 | 2219.953 | 2439.327 | 2379.336 | 1230.632 | 1377.217 | 1320.898 |
| Vmn1r51 | 22296 | 1.87809 | -3.14766 | 2.549407 | 0.018529 | 0.041674 | -0.29049 | -3.14766 | -0.43457 | 0.668235 | 0.753504 | 1.483961 | 4.440175 | 2.603909 | 6.493546 | 15.44835 | 13.52668 | 2.842522 | 13.40422 | 17.23733 | 3.076858 | 4.575167 | 1.492683 | 17.52087 | 14.93557 | 7.288832 |
| Boc | 117606 | 1.878117 | 0.319666 | 5.116263 | 4.35E-05 | 0.000149 | 2.845028 | 0.319666 | 12.7659 | 1.84E-11 | 1.64E-10 | 14.09763 | 31.96926 | 19.9633 | 123.3774 | 56.17581 | 74.39673 | 602.6146 | 604.5302 | 614.7982 | 19.23036 | 32.02617 | 23.88292 | 95.9476 | 76.25001 | 94.75482 |
| Mir202 | 387198 | 1.879582 | -1.35669 | 3.73879 | 0.001185 | 0.003371 | 0.515569 | -1.35669 | 1.393544 | 0.177798 | 0.253938 | 12.61367 | 13.32052 | 5.207818 | 32.46773 | 37.91867 | 48.69604 | 95.22448 | 49.5956 | 44.81707 | 16.1535 | 17.53814 | 8.209754 | 15.85221 | 19.65207 | 28.34546 |
| Creb5 | 231991 | 1.880022 | 6.137476 | 41.54323 | 6.59E-22 | 4.34E-20 | -0.66471 | 6.137476 | -16.3974 | 1.42E-13 | 1.90E-12 | 2382.5 | 2383.486 | 2438.995 | 9724.734 | 9545.675 | 9046.642 | 6250.705 | 6159.238 | 6131.894 | 4529.135 | 4683.446 | 4341.467 | 1981.526 | 2095.696 | 2183.41 |
| Sfmbt2 | 353282 | 1.880262 | 2.28875 | 14.88508 | 9.53E-13 | 9.26E-12 | -1.14844 | 2.28875 | -9.55404 | 3.67E-09 | 2.23E-08 | 163.2358 | 191.8156 | 139.7431 | 618.1856 | 682.5361 | 631.6958 | 284.2522 | 286.8503 | 327.5093 | 269.9943 | 247.059 | 231.3658 | 252.8011 | 284.5619 | 270.4967 |
| Rgs20 | 58175 | 1.88057 | -1.80709 | 3.448044 | 0.002368 | 0.006406 | 0.317197 | -1.80709 | 0.753197 | 0.45956 | 0.558415 | 5.193865 | 11.54445 | 8.679696 | 23.37676 | 35.10988 | 40.58003 | 29.84648 | 45.57434 | 50.56285 | 13.84586 | 1.525056 | 4.478048 | 18.35519 | 33.01547 | 23.48624 |
| Gm17308 | 100503882 | 1.882222 | -3.80639 | 2.593905 | 0.016808 | 0.038163 | -0.13917 | -3.80639 | -0.21486 | 0.831915 | 0.897428 | 2.225942 | 2.664105 | 1.735939 | 18.18193 | 8.426372 | 5.410671 | 14.21261 | 9.382952 | 5.745778 | 0 | 1.525056 | 0 | 15.01788 | 7.074744 | 4.859222 |
| Atp2b3 | 320707 | 1.882568 | -1.71183 | 3.148177 | 0.004787 | 0.012249 | 1.388101 | -1.71183 | 3.301768 | 0.003343 | 0.008155 | 11.12971 | 2.664105 | 3.471878 | 28.5716 | 14.04395 | 20.29002 | 55.42917 | 50.93603 | 62.0544 | 12.30743 | 10.67539 | 7.463413 | 31.70442 | 45.59279 | 21.8665 |
| Tex29 | 75528 | 1.88269 | -1.87397 | 2.730879 | 0.012408 | 0.029134 | 1.06491 | -1.87397 | 2.102861 | 0.04751 | 0.082879 | 0.741981 | 11.54445 | 13.01954 | 16.88322 | 16.85274 | 29.75869 | 35.53152 | 53.61687 | 47.11538 | 11.53822 | 8.387806 | 5.97073 | 30.03577 | 21.22423 | 21.8665 |
| Gzmf | 14943 | 1.884223 | -3.75258 | 2.599271 | 0.016611 | 0.037767 | -0.03066 | -3.75258 | -0.04759 | 0.962484 | 0.969159 | 3.709904 | 0.888035 | 1.735939 | 7.792255 | 14.04395 | 5.410671 | 8.527565 | 13.40422 | 5.745778 | 0 | 1.525056 | 0 | 9.177596 | 18.0799 | 13.76779 |
| Vit | 74199 | 1.884227 | -2.74166 | 2.791859 | 0.010822 | 0.025748 | 0.99726 | -2.74166 | 1.904886 | 0.070359 | 0.115758 | 2.225942 | 2.664105 | 3.471878 | 7.792255 | 14.04395 | 14.87935 | 19.89765 | 30.8297 | 25.28142 | 6.153716 | 0.762528 | 0.746341 | 37.54471 | 25.15464 | 21.8665 |
| Clvs1 | 74438 | 1.884397 | -3.58315 | 2.630352 | 0.015512 | 0.0356 | 0.241477 | -3.58315 | 0.392897 | 0.698292 | 0.779903 | 4.451884 | 0.888035 | 2.603909 | 6.493546 | 16.85274 | 9.468674 | 7.106304 | 10.72337 | 22.98311 | 0.769214 | 1.525056 | 0 | 8.343269 | 11.00516 | 6.478962 |
| Gm3831 | 100042411 | 1.884985 | -1.57288 | 3.42316 | 0.002512 | 0.006755 | 0.34868 | -1.57288 | 0.813146 | 0.425113 | 0.523941 | 2.967923 | 5.32821 | 15.62345 | 25.97418 | 21.06593 | 29.75869 | 38.37404 | 28.14886 | 35.62382 | 8.461359 | 6.862751 | 18.65853 | 51.72827 | 55.81187 | 43.73299 |
| Tmem229l | 268567 | 1.885135 | 7.304257 | 41.66185 | 6.21E-22 | 4.10E-20 | 0.290216 | 7.304257 | 8.289365 | 4.10E-08 | 2.13E-07 | 5128.571 | 4792.725 | 4456.156 | 19200.12 | 18844.18 | 18720.92 | 24650.35 | 24213.38 | 23340.5 | 5542.959 | 5594.667 | 5378.135 | 6574.496 | 5945.143 | 6321.847 |
| Tdrd7 | 100121 | 1.885991 | 6.906765 | 47.26305 | 4.32E-23 | 3.96E-21 | -0.25868 | 6.906765 | -7.73252 | 1.27E-07 | 6.11E-07 | 3945.112 | 3965.964 | 4091.609 | 15980.62 | 16215.15 | 15167.46 | 13845.92 | 14187.02 | 13142.89 | 5462.192 | 5698.371 | 5518.447 | 3951.372 | 3942.99 | 3928.681 |
| Apba1 | 319924 | 1.886267 | 4.075359 | 24.79352 | 3.21E-17 | 7.24E-16 | -1.72068 | 4.075359 | -21.2351 | 7.74E-16 | 1.72E-14 | 779.8218 | 789.4631 | 726.4906 | 3102.616 | 2822.834 | 3143.6 | 1046.048 | 962.4228 | 869.9108 | 1156.899 | 993.5738 | 1046.37 | 430.5127 | 468.5052 | 425.1819 |
| Sod2 | 20656 | 1.88758 | 8.43392 | 85.99058 | 1.31E-28 | 1.16E-25 | -0.2306 | 8.43392 | -13.2755 | 8.73E-12 | 8.29E-11 | 20982.47 | 21642.3 | 20432 | 82629.07 | 84738.4 | 81702.48 | 74447.06 | 73839.81 | 72265.8 | 10309.01 | 10295.65 | 10762.99 | 2807.51 | 2802.385 | 2836.166 |
| Tmem184a | 231832 | 1.888734 | -1.44366 | 2.920336 | 0.008089 | 0.019798 | 0.132905 | -1.44366 | 0.261632 | 0.796113 | 0.866089 | 7.419807 | 7.992315 | 12.15157 | 28.5716 | 26.68351 | 62.22272 | 32.689 | 33.51054 | 59.75609 | 17.69193 | 1.525056 | 18.65853 | 55.06558 | 43.23454 | 29.15533 |
| Gm19279 | 100502606 | 1.88997 | 2.790137 | 13.65753 | 5.06E-12 | 4.38E-11 | -0.55608 | 2.790137 | -4.77715 | 9.76E-05 | 0.000316 | 262.6612 | 212.2404 | 216.1244 | 851.9532 | 917.0701 | 983.3894 | 675.0989 | 714.4448 | 571.1303 | 330.7622 | 283.6604 | 229.1268 | 282.0025 | 322.2939 | 347.4343 |

| | | | | | | | | | | | | | | | | | | | | | | | | | | |
|---|---|---|---|---|---|---|---|---|---|---|---|---|---|---|---|---|---|---|---|---|---|---|---|---|---|---|
| Nfatc4 | 73181 | 1.890221 | -1.7863 | 2.735746 | 0.012273 | 0.028858 | 0.525327 | -1.7863 | 1.004577 | 0.326355 | 0.422474 | 11.87169 | 0.888035 | 15.62345 | 31.16902 | 18.25714 | 28.40602 | 25.5827 | 33.51054 | 59.75609 | 3.846072 | 7.625279 | 2.985365 | 43.385 | 66.81702 | 45.35273 |
| Gm10257 | 544973 | 1.890808 | -2.41338 | 2.84903 | 0.009512 | 0.022943 | -0.25404 | -2.41338 | -0.44549 | 0.660453 | 0.746383 | 6.677827 | 6.216245 | 1.735939 | 12.98709 | 28.08791 | 16.23201 | 14.21261 | 24.12759 | 11.49156 | 10.769 | 24.40089 | 15.67317 | 15.01788 | 3.14433 | 7.288832 |
| Bckdhb | 12040 | 1.891976 | 2.81511 | 17.35381 | 4.58E-14 | 5.61E-13 | -0.30066 | 2.81511 | -3.38982 | 0.002717 | 0.006773 | 301.2442 | 359.6542 | 265.5987 | 1200.007 | 1321.536 | 1126.772 | 1055.997 | 1097.805 | 934.2635 | 311.5319 | 301.1985 | 273.9072 | 122.6461 | 113.1959 | 112.572 |
| Sim1 | 20464 | 1.892069 | -1.99311 | 2.528904 | 0.019376 | 0.043346 | 0.998833 | -1.99311 | 1.833121 | 0.080771 | 0.130114 | 4.451884 | 3.55214 | 9.547666 | 36.36386 | 7.021976 | 39.22736 | 45.48035 | 40.21265 | 54.01031 | 7.692145 | 0.762528 | 1.492683 | 32.53875 | 73.89177 | 46.16261 |
| Pcdhb17 | 93888 | 1.892322 | -1.23393 | 3.95527 | 0.000705 | 0.002075 | -0.23687 | -1.23393 | -0.60915 | 0.548857 | 0.64365 | 14.83961 | 15.98463 | 5.207818 | 51.94837 | 39.32307 | 41.9327 | 59.69296 | 32.17012 | 32.17636 | 23.07643 | 15.25056 | 28.36097 | 28.36712 | 14.93557 | 30.77507 |
| Hpse | 15442 | 1.892384 | 4.438157 | 25.25356 | 2.20E-17 | 5.07E-16 | -2.24256 | 4.438157 | -25.138 | 2.41E-17 | 7.65E-16 | 930.4439 | 801.8956 | 803.7399 | 3415.605 | 3492.731 | 3177.417 | 729.1068 | 788.168 | 699.8358 | 4145.297 | 3872.116 | 3932.472 | 470.5604 | 477.1521 | 373.3502 |
| Mab21l3 | 242125 | 1.895503 | 0.390906 | 6.938579 | 6.81E-07 | 2.89E-06 | 0.5862 | 0.390906 | 3.030792 | 0.00628 | 0.014302 | 46.00281 | 37.29747 | 28.643 | 162.3386 | 148.8659 | 133.9141 | 248.7206 | 193.0207 | 253.9634 | 87.69045 | 79.3029 | 61.19998 | 20.85817 | 42.44846 | 31.58494 |
| 2310014F( | 69625 | 1.896074 | -3.02671 | 2.815264 | 0.010266 | 0.024566 | 0.826325 | -3.02671 | 1.562042 | 0.132985 | 0.198713 | 2.967923 | 7.992315 | 0.86797 | 11.68838 | 15.44835 | 10.82134 | 19.89765 | 20.10633 | 31.0272 | 0.769214 | 1.525056 | 0 | 23.36115 | 24.36856 | 17.00728 |
| Gm9199 | 668489 | 1.897062 | -0.15086 | 6.284899 | 2.89E-06 | 1.13E-05 | -2.9443 | -0.15086 | -6.77134 | 9.81E-07 | 4.17E-06 | 52.68063 | 43.51371 | 26.90706 | 161.0399 | 160.1011 | 159.6148 | 22.74017 | 9.382952 | 39.07129 | 139.2278 | 122.767 | 111.9512 | 40.04769 | 31.4433 | 14.57766 |
| Gm5069 | 277333 | 1.897555 | -0.90842 | 3.591688 | 0.001684 | 0.004661 | 2.171867 | -0.90842 | 6.447053 | 2.01E-06 | 8.22E-06 | 8.903769 | 6.216245 | 11.2836 | 36.36386 | 42.13186 | 28.40602 | 140.7048 | 138.0634 | 225.2345 | 4.615287 | 12.20045 | 15.67317 | 75.08942 | 55.81187 | 55.07118 |
| Rasgrf1 | 19417 | 1.897575 | 2.155804 | 15.2961 | 5.59E-13 | 5.66E-12 | -0.16359 | 2.155804 | -1.67239 | 0.109044 | 0.16808 | 249.3055 | 198.0318 | 223.9362 | 940.2654 | 855.2767 | 883.292 | 770.3234 | 900.7634 | 815.9005 | 95.38259 | 111.3291 | 85.0829 | 60.07154 | 99.04641 | 97.18443 |
| Tor3a | 30935 | 1.898297 | 8.1639 | 54.15567 | 2.42E-24 | 3.33E-22 | -0.79733 | 8.1639 | -25.5116 | 1.78E-17 | 5.83E-16 | 11508.86 | 11375.73 | 11195.07 | 46404.18 | 45850.7 | 43480.15 | 28180.76 | 27872.73 | 25403.23 | 21024.17 | 20971.04 | 21174.45 | 5134.448 | 5019.924 | 4983.942 |
| Samhd1 | 56045 | 1.900203 | 9.674157 | 49.71927 | 8.03E-15 | 1.13E-13 | -0.76146 | 9.674157 | -23.2753 | 4.67E-11 | 3.86E-10 | 38823.4 | 39633.89 | 38342.56 | 155920.4 | 157351.3 | 152106.1 | 99084.62 | 97537.13 | 89865.12 | 43138.32 | 45694.48 | 43706.49 | 10799.53 | 10929.69 | 11015.05 |
| Sp6 | 83395 | 1.900493 | -0.10213 | 5.741595 | 1.00E-05 | 3.70E-05 | -0.86103 | -0.10213 | -2.88724 | 0.008722 | 0.01914 | 19.2915 | 31.96926 | 33.85081 | 150.6503 | 92.69009 | 90.62874 | 59.69296 | 68.36151 | 58.60694 | 17.69193 | 37.36386 | 14.18048 | 146.0072 | 118.6985 | 127.1496 |
| P4ha3 | 320452 | 1.900669 | -2.49823 | 2.796504 | 0.010709 | 0.025504 | 0.167227 | -2.49823 | 0.303305 | 0.764594 | 0.838634 | 6.677827 | 1.77607 | 6.943757 | 18.18193 | 28.08791 | 13.52668 | 17.05513 | 21.44675 | 28.72889 | 0.769214 | 6.100223 | 0 | 42.55067 | 53.45362 | 34.82442 |
| Ikbke | 56489 | 1.902817 | 8.501219 | 32.41705 | 5.64E-13 | 5.70E-12 | 0.000638 | 8.501219 | 0.014111 | 0.988975 | 0.990746 | 14140.67 | 14290.26 | 14303.27 | 58178.27 | 56056.44 | 56283.15 | 61934.28 | 61973.06 | 54541.22 | 19498.05 | 19851.65 | 19470.55 | 5045.175 | 4960.181 | 5166.972 |
| Crocc2 | 381284 | 1.902863 | -1.0952 | 4.136695 | 0.000455 | 0.001376 | 1.045658 | -1.0952 | 3.234697 | 0.003912 | 0.009389 | 7.419807 | 13.32052 | 5.207818 | 25.97418 | 30.8967 | 44.63804 | 61.11422 | 62.99982 | 89.63414 | 12.30743 | 9.912862 | 12.6878 | 66.74615 | 67.60311 | 78.55742 |
| Best1 | 24115 | 1.902976 | 2.33633 | 12.2088 | 4.27E-11 | 3.20E-10 | -1.10383 | 2.33633 | -7.56831 | 1.78E-07 | 8.42E-07 | 210.7225 | 150.9659 | 211.7846 | 732.472 | 655.8526 | 915.7561 | 338.2601 | 355.2118 | 403.3536 | 239.2257 | 298.1484 | 324.6584 | 175.2087 | 154.0722 | 170.8826 |
| Phip | 83946 | 1.904287 | 7.503486 | 84.70173 | 1.81E-28 | 1.34E-25 | -2.77743 | 7.503486 | -94.0545 | 1.94E-29 | 1.28E-25 | 10060.52 | 9982.401 | 9907.005 | 39427.51 | 40414.28 | 39944.28 | 5947.977 | 5891.154 | 6257.152 | 10257.47 | 10140.1 | 10126.36 | 6114.782 | 6022.965 | 6276.495 |
| Thrb | 21834 | 1.904327 | -0.10578 | 4.270054 | 0.00033 | 0.001016 | -0.40676 | -0.10578 | -1.10847 | 0.279994 | 0.371737 | 28.93725 | 5.32821 | 16.49142 | 79.22126 | 50.55823 | 48.69604 | 51.16539 | 45.57434 | 41.3696 | 140.7662 | 118.9543 | 105.2341 | 116.8058 | 113.982 | 106.093 |
| Dmbt1 | 12945 | 1.904624 | -1.52098 | 3.069118 | 0.005749 | 0.014502 | 1.498834 | -1.52098 | 3.510957 | 0.00204 | 0.00522 | 7.419807 | 8.88035 | 2.603909 | 19.48064 | 28.08791 | 24.34802 | 59.69296 | 52.27645 | 104.5732 | 16.1535 | 3.050111 | 8.956095 | 52.5626 | 44.80671 | 43.73299 |
| Nipal4 | 214112 | 1.905068 | -2.61253 | 2.802462 | 0.010566 | 0.025193 | -0.557 | -2.61253 | -0.89901 | 0.378677 | 0.476945 | 6.677827 | 2.664105 | 2.603909 | 14.2858 | 12.63956 | 21.64268 | 8.527565 | 4.021265 | 31.0272 | 1.538429 | 5.337695 | 5.224389 | 27.53279 | 32.22939 | 30.77507 |
| Zfp521 | 225207 | 1.905982 | -1.22479 | 3.594581 | 0.001672 | 0.004633 | 0.26381 | -1.22479 | 0.654298 | 0.519909 | 0.616479 | 14.83961 | 14.20856 | 4.339848 | 29.87031 | 63.19779 | 36.52203 | 48.32287 | 34.85097 | 73.54596 | 9.999788 | 16.01308 | 16.41951 | 24.19548 | 50.30929 | 36.44416 |
| Rasa4 | 54153 | 1.906529 | 7.48962 | 58.29591 | 5.08E-25 | 8.84E-23 | -2.61193 | 7.48962 | -62.679 | 1.09E-25 | 5.12E-23 | 11775.98 | 11925.42 | 11687.21 | 47680.81 | 48278.9 | 45720.17 | 8409.6 | 8467.444 | 7331.613 | 15878.89 | 16547.62 | 15939.61 | 2054.947 | 1870.091 | 2015.767 |
| Lrrc37a | 237954 | 1.906921 | -2.50062 | 2.968313 | 0.007249 | 0.017914 | 1.346324 | -2.50062 | 2.792774 | 0.0108 | 0.0231 | 1.483961 | 3.55214 | 3.471878 | 10.38967 | 15.44835 | 9.468674 | 29.84648 | 33.51054 | 31.0272 | 1.538429 | 5.337695 | 7.463413 | 20.85817 | 22.01031 | 31.58494 |
| Cadps2 | 320405 | 1.907979 | 0.19401 | 7.003579 | 5.92E-07 | 2.53E-06 | -1.02682 | 0.19401 | -4.10476 | 0.000492 | 0.001424 | 46.00281 | 39.07354 | 26.03909 | 110.3903 | 186.7846 | 156.9095 | 72.4843 | 85.78699 | 67.80018 | 76.15223 | 57.95212 | 65.67803 | 61.74019 | 48.73712 | 53.45144 |
| Oas1g | 23960 | 1.908018 | 6.290204 | 27.84497 | 2.89E-18 | 8.27E-17 | -0.02693 | 6.290204 | -0.48199 | 0.634718 | 0.7238 | 2510.121 | 2488.274 | 3070.008 | 10784.48 | 10740.81 | 10591.39 | 11405.62 | 11567.84 | 9972.372 | 3637.615 | 3631.92 | 3402.57 | 2037.426 | 2073.686 | 1908.864 |
| 2900079G | 620760 | 1.908565 | -1.92473 | 3.176858 | 0.004478 | 0.011513 | 0.817414 | -1.92473 | 1.823995 | 0.082187 | 0.132015 | 2.967923 | 12.43249 | 3.471878 | 33.76644 | 14.04395 | 18.93735 | 48.32287 | 25.46801 | 47.11538 | 3.846072 | 4.575167 | 3.731706 | 45.88798 | 57.38403 | 38.0639 |
| Gm19937 | 100503873 | 1.910346 | -2.91343 | 2.860375 | 0.00927 | 0.0224 | -0.26185 | -2.91343 | -0.44479 | 0.660948 | 0.746847 | 2.967923 | 6.216245 | 1.735939 | 12.98709 | 25.27911 | 8.116006 | 7.106304 | 22.78717 | 10.3424 | 2.307643 | 5.337695 | 2.239024 | 16.68654 | 15.72165 | 8.908573 |
| Etv3l | 546801 | 1.911128 | -3.32804 | 2.662443 | 0.014449 | 0.033403 | 1.716228 | -3.32804 | 3.119288 | 0.005119 | 0.011929 | 1.483961 | 0.888035 | 1.735939 | 2.597418 | 7.021976 | 13.52668 | 18.47639 | 28.14886 | 24.13227 | 1.538429 | 2.287584 | 2.985365 | 7.508942 | 13.3634 | 8.098703 |
| Perm1 | 74183 | 1.911523 | 4.379091 | 21.20735 | 7.95E-16 | 1.35E-14 | 1.891144 | 4.379091 | 31.71339 | 1.92E-19 | 1.07E-17 | 403.6375 | 431.585 | 463.4958 | 1716.894 | 1755.494 | 1723.299 | 6799.312 | 6963.491 | 6310.013 | 546.9115 | 546.7325 | 490.3462 | 1087.128 | 1131.959 | 1158.114 |
| Sema7a | 20361 | 1.911571 | 2.161111 | 12.41192 | 3.13E-11 | 2.40E-10 | -0.14821 | 2.161111 | -1.23507 | 0.230229 | 0.315542 | 186.9791 | 162.5104 | 127.5915 | 658.4455 | 690.9625 | 569.4731 | 592.6658 | 628.6578 | 574.5778 | 239.9949 | 169.2812 | 156.7317 | 123.4804 | 120.2706 | 121.4805 |
| Fam53c | 66306 | 1.911898 | 6.719796 | 40.64342 | 1.05E-21 | 6.49E-20 | 0.327182 | 6.719796 | 8.966861 | 1.10E-08 | 6.16E-08 | 3172.71 | 3409.166 | 3310.436 | 13127.35 | 13483.6 | 13068.12 | 18125.34 | 17022.02 | 16654.71 | 4326.831 | 4720.047 | 4557.906 | 2903.458 | 2785.877 | 3049.971 |
| Oaf | 102644 | 1.912355 | 2.510276 | 19.28003 | 5.52E-15 | 7.96E-14 | -2.45531 | 2.510276 | -19.233 | 5.79E-15 | 1.04E-13 | 313.8579 | 264.6344 | 316.8089 | 1225.981 | 1242.89 | 1136.241 | 224.5592 | 227.8717 | 227.5328 | 313.8395 | 303.4861 | 247.7853 | 181.8833 | 192.5902 | 213.8058 |
| Zic2 | 22772 | 1.913634 | -1.69005 | 3.166917 | 0.004582 | 0.011761 | 0.844513 | -1.69005 | 1.8955 | 0.07165 | 0.117522 | 7.419807 | 8.88035 | 1.735939 | 15.58451 | 16.85274 | 36.52203 | 34.11026 | 46.91476 | 40.22045 | 7.692145 | 12.20045 | 3.731706 | 57.56856 | 65.24486 | 57.50079 |
| Ace | 11421 | 1.91364 | -0.66722 | 4.949557 | 6.46E-05 | 0.000216 | 1.759388 | -0.66722 | 7.031059 | 5.58E-07 | 2.46E-06 | 10.38773 | 10.65642 | 12.15157 | 46.75353 | 36.51428 | 54.10671 | 146.3899 | 164.8719 | 170.075 | 9.999788 | 13.7255 | 14.92683 | 75.08942 | 56.59795 | 73.69819 |
| Ntf5 | 78405 | 1.914159 | -2.54982 | 2.621249 | 0.015827 | 0.03625 | 1.518248 | -2.54982 | 2.793049 | 0.010793 | 0.023088 | 2.967923 | 1.77607 | 2.603909 | 9.090964 | 16.85274 | 8.116006 | 21.31891 | 34.85097 | 45.96622 | 4.615287 | 3.050111 | 0.746341 | 38.37904 | 54.2397 | 26.72572 |
| Mfsd9 | 211798 | 1.914929 | 2.693961 | 17.2378 | 5.24E-14 | 6.34E-13 | -2.57162 | 2.693961 | -17.4779 | 3.97E-14 | 6.05E-13 | 267.1131 | 328.5729 | 306.3933 | 1298.709 | 1179.692 | 1122.714 | 214.6104 | 257.361 | 171.2242 | 326.1469 | 343.9001 | 402.2779 | 329.5591 | 323.866 | 301.2717 |
| Col5a3 | 53867 | 1.91517 | -0.01855 | 5.701607 | 1.10E-05 | 4.04E-05 | 2.338764 | -0.01855 | 11.41439 | 1.50E-10 | 1.14E-09 | 26.71131 | 15.09659 | 16.49142 | 93.50706 | 63.19779 | 79.8074 | 426.3783 | 415.5307 | 401.0553 | 14.61507 | 17.53814 | 12.6878 | 80.09538 | 88.04125 | 59.12053 |
| Catsper1 | 225865 | 1.915378 | -2.77192 | 2.667921 | 0.014275 | 0.033037 | 1.160326 | -2.77192 | 2.11784 | 0.046089 | 0.08076 | 0.741981 | 3.55214 | 6.943757 | 9.090964 | 9.830767 | 18.93735 | 21.31891 | 28.14886 | 36.77298 | 3.076858 | 0.762528 | 5.224389 | 11.68058 | 17.29382 | 17.81715 |
| Igdcc3 | 19289 | 1.917046 | -1.74233 | 3.390332 | 0.002714 | 0.007257 | 0.726237 | -1.74233 | 1.73425 | 0.097296 | 0.152329 | 8.161788 | 5.32821 | 3.471878 | 25.97418 | 30.8967 | 14.87935 | 38.37404 | 46.91476 | 35.62382 | 9.999788 | 3.812639 | 5.224389 | 71.75211 | 40.8763 | 48.59222 |
| Rgs14 | 51791 | 1.91769 | 4.875875 | 35.17784 | 2.18E-20 | 1.02E-18 | -3.33714 | 4.875875 | -41.4225 | 7.01E-22 | 8.05E-20 | 1920.988 | 1903.059 | 2089.203 | 7796.151 | 8107.574 | 7927.986 | 881.1817 | 856.5295 | 732.0121 | 1433.816 | 1483.879 | 1269.526 | 893.5641 | 908.7115 | 990.4713 |
| Oas2 | 246728 | 1.917911 | 8.437026 | 32.4873 | 5.82E-14 | 7.00E-13 | 0.16059 | 8.437026 | 3.58262 | 0.003278 | 0.008011 | 10605.13 | 10291.44 | 10686.44 | 42898.96 | 42231.57 | 42331.74 | 50827.13 | 52161.17 | 45849.01 | 14200.47 | 14577.24 | 13776.71 | 12148.63 | 12652 | 11650.79 |
| Amph | 218038 | 1.919373 | -2.45226 | 2.736804 | 0.012244 | 0.028794 | 1.123973 | -2.45226 | 2.109571 | 0.046869 | 0.081897 | 3.709904 | 1.77607 | 3.471878 | 11.68838 | 9.830767 | 17.58468 | 21.31891 | 21.44675 | 50.56285 | 7.692145 | 3.812639 | 1.492683 | 31.70442 | 39.30413 | 21.8665 |
| Mcc | 328949 | 1.919679 | 1.988506 | 14.83317 | 1.02E-12 | 9.85E-12 | -1.17903 | 1.988506 | -9.62526 | 3.22E-09 | 1.98E-08 | 195.8829 | 173.1668 | 177.9338 | 722.0823 | 790.6745 | 710.1506 | 302.7286 | 379.3394 | 336.7026 | 183.073 | 213.5078 | 153 | 106.7938 | 88.82734 | 76.12781 |
| N4bp1 | 80750 | 1.919878 | 8.311466 | 52.17788 | 5.32E-24 | 6.51E-22 | -0.0156 | 8.311466 | -0.54068 | 0.594329 | 0.687248 | 9228.015 | 8926.528 | 9438.301 | 37096.33 | 38105.46 | 36340.77 | 39336.24 | 38899.04 | 36621.29 | 15918.12 | 15541.08 | 16329.95 | 12030.99 | 11288.93 | 12256.58 |
| Rras2 | 66922 | 1.92223 | 1.810928 | 10.89093 | 3.54E-10 | 2.35E-09 | -0.43328 | 1.810928 | -3.00316 | 0.006692 | 0.015136 | 86.06977 | 103.0121 | 77.24929 | 351.9502 | 349.6944 | 369.2783 | 258.6695 | 305.6162 | 262.0075 | 206.9187 | 253.1592 | 247.7853 | 170.2027 | 194.9485 | 231.6229 |
| Vmn1r58 | 81014 | 1.923546 | -3.93846 | 2.75795 | 0.011678 | 0.027579 | 1.240362 | -3.93846 | 2.230146 | 0.036595 | 0.066402 | 1.483961 | 0.888035 | 1.735939 | 9.090964 | 7.021976 | 4.058003 | 17.05513 | 22.78717 | 12.64071 | 0 | 0 | 0 | 6.674615 | 17.29382 | 9.718443 |
| Cdh6 | 12563 | 1.924108 | -1.961 | 3.914551 | 0.000777 | 0.002271 | -0.2843 | -1.961 | -0.68669 | 0.49967 | 0.597383 | 4.451884 | 6.216245 | 9.547666 | 36.36386 | 23.87472 | 21.64268 | 14.21261 | 34.85097 | 22.98311 | 9.230573 | 8.387806 | 8.956095 | 20.85817 | 23.58248 | 18.62702 |
| L3mbtl4 | 320858 | 1.924991 | -2.3995 | 3.006404 | 0.006643 | 0.016546 | 0.17563 | -2.3995 | 0.349082 | 0.730448 | 0.809281 | 3.709904 | 13.32052 | 6.075787 | 32.46773 | 28.08791 | 22.99535 | 44.05909 | 28.14886 | 28.72889 | 0 | 3.812639 | 0.746341 | 40.88202 | 25.15464 | 10.52831 |
| Abi3bp | 320712 | 1.925189 | 1.016094 | 10.88248 | 3.59E-10 | 2.38E-09 | -1.47326 | 1.016094 | -8.19295 | 4.97E-08 | 2.55E-07 | 74.19807 | 108.3403 | 80.72117 | 375.3269 | 344.0768 | 327.3456 | 125.071 | 108.5742 | 156.2852 | 81.53673 | 71.67762 | 80.60486 | 95.9476 | 91.18558 | 67.21923 |
| Pwwp3b | 245631 | 1.925302 | -1.63037 | 3.50568 | 0.002066 | 0.005642 | 0.440131 | -1.63037 | 1.063166 | 0.299588 | 0.393383 | 8.903769 | 2.664105 | 12.15157 | 45.45482 | 26.68351 | 18.93735 | 34.11026 | 44.23392 | 43.66791 | 9.230573 | 11.43792 | 3.731706 | 37.54471 | 40.8763 | 42.11325 |
| Gm9519 | 671132 | 1.925939 | -3.82219 | 2.51495 | 0.019974 | 0.044508 | 0.57917 | -3.82219 | 0.876848 | 0.390333 | 0.48919 | 2.225942 | 0 | 1.735939 | 5.194837 | 5.617581 | 4.058003 | 9.948826 | 9.382952 | 5.745778 | 1.538429 | 3.050111 | 4.478048 | 5.005962 | 8.646909 | 4.859222 |
| Apobr | 171504 | 1.926396 | 6.783348 | 29.1096 | 5.66E-17 | 1.22E-15 | 0.396633 | 6.783348 | 7.855635 | 2.54E-07 | 1.17E-06 | 3782.618 | 3707.546 | 3996.132 | 15749.45 | 15417.45 | 15413.65 | 21844.78 | 22981.53 | 19327.65 | 2856.093 | 2581.919 | 2886.102 | 3497.498 | 3525.581 | 3543.182 |
| Ankrd66 | 100043332 | 1.926804 | 0.961684 | 8.500021 | 2.70E-08 | 1.37E-07 | 0.671164 | 0.961684 | 4.29774 | 0.000309 | 0.000924 | 89.77967 | 60.38638 | 53.81412 | 275.3263 | 282.2834 | 266.4755 | 454.8035 | 541.5304 | 390.7129 | 62.30637 | 39.65145 | 68.6634 | 43.385 | 51.09537 | 41.30338 |
| Faah | 14073 | 1.926875 | -0.19283 | 6.779109 | 9.64E-07 | 4.00E-06 | -0.6651 | -0.19283 | -2.71399 | 0.012884 | 0.02691 | 25.96933 | 25.75301 | 22.56721 | 118.1825 | 106.734 | 82.51273 | 55.42917 | 67.02109 | 75.84427 | 31.53779 | 28.21353 | 39.55609 | 78.42673 | 58.9562 | 71.26858 |
| Col11a1 | 12814 | 1.926982 | -1.33111 | 4.348056 | 0.000274 | 0.00085 | 0.306736 | -1.33111 | 0.918597 | 0.368572 | 0.46634 | 11.87169 | 7.992315 | 9.547666 | 32.46773 | 35.10988 | 59.51738 | 39.7953 | 54.95729 | 63.20356 | 9.999788 | 6.100223 | 6.717071 | 35.04173 | 56.59795 | 42.11325 |
| Csn1s1 | 12990 | 1.927291 | -0.9934 | 4.19835 | 0.000393 | 0.001197 | -1.78279 | -0.9934 | -3.59132 | 0.001685 | 0.00439 | 13.35565 | 24.86498 | 14.75548 | 62.33804 | 85.66811 | 62.22272 | 25.5827 | 12.0638 | 26.43058 | 85.3828 | 53.37695 | 25.3756 | 20.02385 | 18.0799 | 19.43689 |
| Plpp1 | 19012 | 1.92917 | 2.5391 | 19.36687 | 5.04E-15 | 7.33E-14 | -1.87626 | 2.5391 | -16.9876 | 7.02E-14 | 1.00E-12 | 334.6333 | 380.079 | 280.3542 | 1320.787 | 1318.727 | 1378.368 | 342.5239 | 398.1053 | 391.8621 | 247.6871 | 273.7475 | 301.5219 | 112.6341 | 97.47424 | 123.1003 |
| Slc1a1 | 20510 | 1.929778 | -1.46396 | 3.361257 | 0.002907 | 0.007734 | 1.10954 | -1.46396 | 2.750987 | 0.011862 | 0.025072 | 14.09763 | 5.32821 | 2.603909 | 23.37676 | 36.51428 | 20.29002 | 46.90161 | 50.93603 | 79.29174 | 18.46115 | 8.387806 | 12.6878 | 36.71038 | 44.02063 | 32.39481 |
| Mybpc2 | 233199 | 1.931575 | -1.29882 | 3.79497 | 0.001036 | 0.002975 | 0.684662 | -1.29882 | 1.842729 | 0.079304 | 0.128228 | 5.935846 | 7.992315 | 9.547666 | 23.37676 | 42.13186 | 33.81669 | 45.48035 | 45.57434 | 71.24765 | 19.99958 | 21.35078 | 7.463413 | 42.55067 | 36.1598 | 38.0639 |
| Gm16712 | 100504333 | 1.933367 | 1.997364 | 14.65222 | 1.30E-12 | 1.23E-11 | -0.21655 | 1.997364 | -2.0909 | 0.048674 | 0.084633 | 186.2372 | 167.8386 | 163.1783 | 684.4197 | 700.7932 | 734.4986 | 631.0398 | 733.2107 | 553.893 | 140.7662 | 124.292 | 141.8048 | 66.74615 | 67.60311 | 86.65612 |
| Nlrp5 | 23968 | 1.933803 | -2.92063 | 2.470617 | 0.021987 | 0.048518 | 1.418398 | -2.92063 | 2.380372 | 0.026682 | 0.050585 | 2.967923 | 0 | 7.811726 | 9.090964 | 11.23516 | 6.763339 | 34.11026 | 20.10633 | 25.28142 | 3.076858 | 3.050111 | 1.492683 | 19.18952 | 18.86598 | 17.00728 |
| Csf3r | 12986 | 1.933824 | 6.785026 | 50.90572 | 8.98E-24 | 1.01E-21 | -3.63581 | 6.785026 | -62.2943 | 1.24E-25 | 5.44E-23 | 6381.034 | 6051.958 | 5745.959 | 24684.57 | 25267.88 | 24326.38 | 2191.584 | 2120.547 | 1920.239 | 30065.52 | 29893.38 | 29826.04 | 1238.975 | 1293.892 | 1490.161 |
| Nr1i2 | 18171 | 1.93524 | -2.31468 | 2.676538 | 0.014004 | 0.032479 | -1.07691 | -2.31468 | -1.57769 | 0.129343 | 0.194026 | 9.64575 | 13.32052 | 2.603909 | 24.67547 | 39.32307 | 28.40602 | 14.21261 | 12.0638 | 17.23733 | 6.92293 | 16.01308 | 0 | 24.19548 | 14.93557 | 21.8665 |
| Abcg4 | 192663 | 1.935558 | -1.02416 | 4.748484 | 0.000105 | 0.000341 | -0.16101 | -1.02416 | -0.49329 | 0.626842 | 0.717089 | 14.83961 | 8.88035 | 14.75548 | 50.64966 | 50.55823 | 56.81205 | 31.26774 | 46.91476 | 72.3968 | 12.30743 | 12.96297 | 8.209754 | 70.91779 | 53.45362 | 51.8317 |
| Dsg1a | 13510 | 1.937125 | -1.9123 | 2.963496 | 0.00733 | 0.018093 | 0.958575 | -1.9123 | 1.991798 | 0.059342 | 0.100312 | 8.161788 | 3.55214 | 2.603909 | 16.88322 | 15.44835 | 25.70069 | 41.21656 | 20.10633 | 64.35271 | 6.153716 | 5.337695 | 11.19512 | 28.36712 | 43.23454 | 42.92312 |
| Zfand5 | 22682 | 1.937674 | 7.260637 | 48.05174 | 3.05E-23 | 2.93E-21 | -1.64037 | 7.260637 | -39.2585 | 2.17E-21 | 2.18E-19 | 7421.291 | 7502.12 | 6536.679 | 29985.9 | 29176.31 | 28550.76 | 10321.2 | 9448.633 | 9509.263 | 6606.783 | 6787.26 | 6820.813 | 4770.681 | 4888.648 | 4500.449 |
| Olfr477 | 258928 | 1.940165 | -4.77377 | 2.470584 | 0.021989 | 0.048518 | -0.36735 | -4.77377 | -0.50195 | 0.620844 | 0.711557 | 1.483961 | 1.77607 | 0 | 5.194837 | 5.617581 | 2.705335 | 2.842522 | 1.340422 | 8.044089 | 0 | 0 | 0 | 5.840288 | 3.930413 | 5.669092 |
| Fam186b | 545136 | 1.942437 | -1.93536 | 3.374366 | 0.002818 | 0.007519 | 0.53324 | -1.93536 | 1.215377 | 0.237496 | 0.323899 | 4.451884 | 6.216245 | 5.207818 | 28.5716 | 16.85274 | 22.99535 | 41.21656 | 18.7659 | 48.26453 | 4.615287 | 6.862751 | 7.463413 | 22.52683 | 44.02063 | 31.58494 |
| Rs1 | 20147 | 1.942527 | -1.22059 | 4.117927 | 0.000477 | 0.001436 | 0.411665 | -1.22059 | 1.173999 | 0.25333 | 0.341858 | 6.677827 | 9.768385 | 13.01954 | 38.96127 | 50.55823 | 31.11136 | 54.00791 | 65.68067 | 47.11538 | 9.230573 | 7.625279 | 16.41951 | 32.53875 | 64.45877 | 45.35273 |
| Itk | 16428 | 1.942889 | -0.59132 | 5.289166 | 2.89E-05 | 0.000101 | 0.042097 | -0.59132 | 0.14856 | 0.883296 | 0.918721 | 13.35565 | 21.31284 | 23.43518 | 80.51997 | 110.9472 | 51.40137 | 58.27169 | 96.51037 | 95.37991 | 20.76879 | 17.53814 | 20.89756 | 35.87606 | 41.66238 | 41.30338 |
| 4932435O | 319819 | 1.943911 | -2.89671 | 2.605332 | 0.016391 | 0.037344 | 0.731813 | -2.89671 | 1.229567 | 0.232242 | 0.317821 | 0.741981 | 3.55214 | 6.075787 | 6.493546 | 18.25714 | 13.52668 | 15.63387 | 20.10633 | 26.43058 | 1.538429 | 4.575167 | 1.492683 | 16.68654 | 8.646909 | 25.91585 |
| Krt76 | 77055 | 1.944061 | -2.79688 | 2.712258 | 0.012934 | 0.030255 | 0.65137 | -2.79688 | 1.129445 | 0.271245 | 0.362096 | 2.225942 | 4.440175 | 1.735939 | 11.68838 | 7.021976 | 18.93735 | 17.05513 | 13.40422 | 29.87805 | 3.846072 | 3.812639 | 1.492683 | 27.53279 | 25.94073 | 10.52831 |
| Gja1 | 14609 | 1.94653 | 5.313026 | 36.88021 | 8.09E-21 | 4.23E-19 | -2.07218 | 5.313026 | -35.2492 | 2.09E-20 | 1.56E-18 | 2266.009 | 2236.072 | 2172.528 | 9232.523 | 9452.985 | 8816.688 | 2187.32 | 2347.078 | 2248.897 | 1193.821 | 1250.546 | 1413.57 | 1250.656 | 1172.049 | 1432.661 |
| Gpr26 | 233919 | 1.948471 | -2.5878 | 3.0738 | 0.005687 | 0.014363 | 0.096569 | -2.5878 | 0.184802 | 0.855129 | 0.917027 | 4.451884 | 1.77607 | 6.075787 | 12.98709 | 16.85274 | 20.29002 | 22.74017 | 14.74464 | 18.38649 | 3.076858 | 3.812639 | 2.985365 | 38.37904 | 15.72165 | 12.14805 |
| Jakmip3 | 74004 | 1.948776 | -1.51219 | 3.89749 | 0.00081 | 0.00236 | 0.576274 | -1.51219 | 1.564607 | 0.132382 | 0.197907 | 7.419807 | 4.440175 | 12.15157 | 27.27289 | 30.8967 | 37.8747 | 38.37404 | 44.23392 | 66.65102 | 13.07665 | 6.100223 | 4.478048 | 35.04173 | 48.73712 | 46.16261 |
| Calcr | 12311 | 1.94939 | -2.13385 | 3.382871 | 0.002762 | 0.007377 | 0.626293 | -2.13385 | 1.443625 | 0.163363 | 0.236546 | 2.967923 | 7.10428 | 7.811726 | 19.48064 | 32.30109 | 20.29002 | 24.16143 | 38.87223 | 52.86116 | 6.153716 | 3.812639 | 1.492683 | 24.19548 | 17.29382 | 29.9652 |
| Pom121l2 | 195236 | 1.95548 | -1.69999 | 2.836169 | 0.009793 | 0.023566 | 1.366195 | -1.69999 | 2.876882 | 0.00893 | 0.01955 | 4.451884 | 8.88035 | 5.207818 | 27.27289 | 19.66153 | 29.75869 | 76.74809 | 52.27645 | 79.29174 | 0 | 16.77561 | 1.492683 | 79.26106 | 91.18558 | 72.07845 |
| Dtx3l | 209200 | 1.955606 | 8.801829 | 73.45119 | 3.75E-27 | 1.35E-24 | 0.330663 | 8.801829 | 17.06338 | 6.42E-14 | 9.27E-13 | 12788.04 | 13130.49 | 12309.54 | 52400.32 | 54507.39 | 51455.48 | 69265.15 | 69019.66 | 68408.08 | 21731.08 | 21901.32 | 21298.34 | 13567.82 | 13548.13 | 13315.89 |
| Shfl | 319278 | 1.956232 | 3.722238 | 16.00343 | 2.29E-13 | 2.48E-12 | 1.327731 | 3.722238 | 16.19263 | 1.82E-13 | 2.38E-12 | 326.4715 | 311.7003 | 295.9776 | 1255.852 | 1336.984 | 1290.445 | 3600.054 | 3667.394 | 2955.628 | 574.6032 | 597.0593 | 632.151 | 357.0919 | 370.2449 | 386.3081 |
| Cyp3a16 | 13114 | 1.956475 | -3.18607 | 2.610729 | 0.016198 | 0.036981 | 0.553578 | -3.18607 | 0.896152 | 0.380168 | 0.478385 | 2.225942 | 1.77607 | 2.603909 | 5.194837 | 11.23516 | 17.58468 | 7.106304 | 21.44675 | 22.98311 | 2.307643 | 6.862751 | 0.746341 | 10.84625 | 10.21907 | 7.288832 |
| Nhlh2 | 18072 | 1.957319 | -2.81505 | 2.974378 | 0.007149 | 0.017693 | 0.757577 | -2.81505 | 1.497187 | 0.148988 | 0.218962 | 3.709904 | 1.77607 | 5.207818 | 20.77935 | 11.23516 | 13.52668 | 32.689 | 25.46801 | 22.98311 | 0 | 0 | 2.985365 | 33.37308 | 45.59279 | 29.15533 |
| Gm4925 | 237433 | 1.957374 | -0.79269 | 5.045192 | 5.14E-05 | 0.000174 | 0.587202 | -0.79269 | 2.112016 | 0.046637 | 0.081577 | 10.38773 | 19.53677 | 23.43518 | 90.90964 | 70.21976 | 51.40137 | 116.5434 | 77.74446 | 135.6004 | 16.1535 | 15.25056 | 15.67317 | 25.86413 | 26.72681 | 17.81715 |
| Gm5460 | 432838 | 1.957489 | -3.09898 | 2.912008 | 0.008244 | 0.020136 | 0.350317 | -3.09898 | 0.623899 | 0.539306 | 0.634501 | 5.935846 | 0.888035 | 1.735939 | 9.090964 | 8.426372 | 14.87935 | 8.527565 | 25.46801 | 12.64071 | 3.846072 | 3.050111 | 3.731706 | 10.01192 | 10.21907 | 8.098703 |
| Fgd5 | 232237 | 1.959657 | -1.24613 | 2.944338 | 0.007658 | 0.018836 | 0.543737 | -1.24613 | 1.090353 | 0.287715 | 0.380319 | 2.225942 | 14.20856 | 14.75548 | 33.76644 | 25.27911 | 39.22736 | 34.11026 | 32.17012 | 95.37991 | 12.30743 | 15.25056 | 7.463413 | 79.26106 | 78.60826 | 46.97248 |
| Gm5862 | 545739 | 1.960073 | -3.84923 | 2.464634 | 0.022273 | 0.049037 | 1.163348 | -3.84923 | 1.818413 | 0.083064 | 0.133195 | 1.483961 | 1.77607 | 0.86797 | 11.68838 | 8.426372 | 2.705335 | 9.948826 | 14.74464 | 26.43058 | 0.769214 | 0.762528 | 0 | 3.337308 | 12.57732 | 8.098703 |
| Klf6 | 23849 | 1.960325 | 8.976035 | 88.59521 | 6.94E-29 | 7.61E-26 | -1.2057 | 8.976035 | -57.7401 | 6.22E-25 | 2.01E-22 | 23713.7 | 23924.55 | 22945.64 | 97526.56 | 100292.1 | 95622.79 | 45221.68 | 43715.17 | 43259.96 | 24764.09 | 24577.04 | 25266.64 | 9796.667 | 9366.174 | 10264.3 |
| Ube2l6 | 56791 | 1.961307 | 7.392993 | 46.77113 | 5.39E-23 | 4.67E-21 | -0.06608 | 7.392993 | -1.97445 | 0.06141 | 0.103281 | 5522.563 | 5117.746 | 5259.896 | 22322.21 | 21421.24 | 22461.05 | 21763.77 | 22553.94 | 21405.32 | 7969.062 | 8342.817 | 7779.115 | 4713.113 | 5107.965 | 5130.528 |
| Spry4 | 24066 | 1.961497 | -0.3814 | 4.986823 | 5.91E-05 | 0.000199 | 2.432304 | -0.3814 | 10.12775 | 1.31E-09 | 8.57E-09 | 10.38773 | 23.08891 | 16.49142 | 74.02642 | 61.79339 | 60.87005 | 343.9451 | 369.9564 | 391.8621 | 11.53822 | 6.862751 | 8.209754 | 75.92375 | 63.67269 | 37.25403 |
| 2310034G( | 75579 | 1.961525 | -0.33274 | 6.400916 | 2.23E-06 | 8.86E-06 | -0.3651 | -0.33274 | -1.47111 | 0.155853 | 0.227588 | 27.45329 | 20.4248 | 15.62345 | 109.0916 | 78.64613 | 78.45473 | 72.4843 | 61.6594 | 78.14258 | 42.30679 | 29.73859 | 36.57072 | 50.89394 | 47.16496 | 51.8317 |
| Mir29a | 387222 | 1.96241 | -1.26777 | 6.768603 | 9.87E-07 | 4.09E-06 | -1.88089 | -1.26777 | -5.91061 | 6.78E-06 | 2.58E-05 | 21.51744 | 21.31284 | 28.643 | 105.1954 | 99.71206 | 91.98141 | 19.89765 | 32.17012 | 31.0272 | 21.538 | 22.11331 | 11.94146 | 7.508942 | 5.502578 | 9.718443 |
| Il13 | 16163 | 1.965449 | -3.02529 | 2.911599 | 0.008252 | 0.020151 | 0.03998 | -3.02529 | 0.069352 | 0.945355 | 0.954831 | 5.193865 | 0.888035 | 3.471878 | 11.68838 | 9.830767 | 16.23201 | 8.527565 | 17.42548 | 14.93902 | 3.846072 | 2.287584 | 0.746341 | 10.84625 | 18.86598 | 21.05663 |
| Meis2 | 17536 | 1.967776 | -1.48012 | 2.63121 | 0.015483 | 0.035545 | 1.52219 | -1.48012 | 2.926981 | 0.007967 | 0.017675 | 1.483961 | 6.216245 | 11.2836 | 29.87031 | 5.617581 | 47.34337 | 46.90161 | 61.6594 | 80.44089 | 8.461359 | 5.337695 | 5.97073 | 102.6222 | 91.18558 | 93.94495 |
| Scn2b | 72821 | 1.969505 | -1.96161 | 2.715821 | 0.012832 | 0.03005 | 1.092173 | -1.96161 | 2.098108 | 0.04797 | 0.083575 | 6.677827 | 1.77607 | 6.075787 | 9.090964 | 25.27911 | 31.11136 | 46.90161 | 48.25518 | 36.77298 | 7.692145 | 7.625279 | 5.97073 | 31.70442 | 48.73712 | 12.95792 |
| Myod1 | 17927 | 1.969612 | -2.93768 | 2.543237 | 0.01878 | 0.042171 | -0.27629 | -2.93768 | -0.41176 | 0.684623 | 0.767791 | 5.935846 | 1.77607 | 4.339848 | 16.88322 | 14.04395 | 18.93735 | 24.16143 | 4.021265 | 25.28142 | 0.769214 | 4.575167 | 0 | 32.53875 | 22.01031 | 14.57766 |
| Zfp366 | 238803 | 1.969901 | -0.25585 | 5.200553 | 3.56E-05 | 0.000123 | -1.02349 | -0.25585 | -2.91779 | 0.008136 | 0.017996 | 11.87169 | 45.28978 | 41.66254 | 137.6632 | 105.3296 | 101.4501 | 69.64178 | 38.87223 | 70.09849 | 31.53779 | 26.68847 | 37.31706 | 36.71038 | 64.45877 | 56.69092 |
| Plekhn1 | 231002 | 1.971712 | 6.006062 | 24.46662 | 4.22E-17 | 9.35E-16 | 1.713588 | 6.006062 | 31.27464 | 2.57E-19 | 1.38E-17 | 1506.221 | 1377.342 | 1514.607 | 6111.725 | 5863.35 | 6471.162 | 21321.76 | 22205.43 | 19549.43 | 1349.971 | 1517.43 | 1401.629 | 3075.329 | 3274.034 | 2997.33 |
| Myd88 | 17874 | 1.972845 | 7.202699 | 45.63446 | 9.07E-23 | 7.30E-21 | -0.41081 | 7.202699 | -11.3907 | 1.55E-10 | 1.18E-09 | 6282.351 | 6827.213 | 6336.178 | 27715.75 | 27267.74 | 26333.74 | 21843.36 | 21850.21 | 20047.02 | 5150.66 | 5018.958 | 5044.521 | 2776.64 | 2608.222 | 3031.344 |
| Azi2 | 27215 | 1.973641 | 6.844044 | 52.52251 | 4.63E-24 | 5.75E-22 | -1.05401 | 6.844044 | -30.1496 | 5.52E-19 | 2.69E-17 | 4603.249 | 4763.42 | 4660.129 | 20057.26 | 19775.29 | 18949.52 | 10373.78 | 9736.824 | 9374.811 | 5901.413 | 5776.148 | 5775.189 | 2844.22 | 2892.784 | 3020.006 |
| Nmi | 64685 | 1.973927 | 4.633072 | 29.52126 | 8.57E-19 | 2.70E-17 | 0.230539 | 4.633072 | 4.431886 | 0.000224 | 0.000686 | 755.3364 | 767.2622 | 709.9991 | 3277.942 | 3054.56 | 3034.034 | 3899.94 | 3810.819 | 3711.773 | 1101.515 | 1047.713 | 1076.224 | 687.4854 | 798.6599 | 782.3347 |
| Atcay | 16467 | 1.975461 | -2.13839 | 4.045112 | 0.000568 | 0.001694 | -0.02335 | -2.13839 | -0.05944 | 0.953151 | 0.961173 | 6.677827 | 6.216245 | 5.207818 | 29.87031 | 32.30109 | 18.93735 | 19.89765 | 28.14886 | 33.32551 | 3.846072 | 4.575167 | 2.239024 | 19.18952 | 30.65722 | 27.53559 |
| Abca12 | 74591 | 1.975558 | -1.27337 | 3.968337 | 0.000683 | 0.002014 | 0.494769 | -1.27337 | 1.366731 | 0.185935 | 0.263727 | 16.32358 | 4.440175 | 8.679696 | 31.16902 | 50.55823 | 36.52203 | 54.00791 | 44.23392 | 72.3968 | 9.230573 | 13.7255 | 4.478048 | 43.385 | 56.59795 | 59.12053 |
| Jam2 | 67374 | 1.975662 | -0.85817 | 3.901513 | 0.000802 | 0.002339 | 1.884552 | -0.85817 | 5.883839 | 7.21E-06 | 2.73E-05 | 8.161788 | 20.4248 | 13.01954 | 53.24708 | 37.91867 | 78.45473 | 210.3466 | 251.9993 | 179.2683 | 1.538429 | 15.25056 | 1.492683 | 61.74019 | 79.39434 | 55.88105 |
| Gm20236 | 100504453 | 1.97581 | 3.119737 | 14.47315 | 1.65E-12 | 1.54E-11 | -1.02939 | 3.119737 | -8.28547 | 4.13E-08 | 2.14E-07 | 477.0936 | 366.7584 | 433.1168 | 1833.777 | 1723.193 | 1815.28 | 874.0754 | 1083.061 | 810.1547 | 201.5342 | 298.9109 | 135.8341 | 266.1503 | 294.781 | 249.44 |
| Sorcs2 | 81840 | 1.976387 | -1.26168 | 3.029 | 0.006306 | 0.01578 | 1.598756 | -1.26168 | 3.662942 | 0.001421 | 0.003763 | 9.64575 | 1.77607 | 10.41564 | 28.5716 | 25.27911 | 24.34802 | 56.85043 | 68.36151 | 134.4512 | 7.692145 | 20.58825 | 8.209754 | 58.40288 | 69.96135 | 59.9304 |
| 3425401B | 100504518 | 1.97644 | -1.65027 | 3.315517 | 0.003237 | 0.008549 | 1.382804 | -1.65027 | 3.340518 | 0.003052 | 0.007516 | 6.677827 | 1.77607 | 7.811726 | 22.07806 | 25.27911 | 16.23201 | 54.00791 | 54.95729 | 63.20356 | 10.769 | 17.53814 | 4.478048 | 41.71635 | 43.23454 | 34.82442 |
| Rfx4 | 71137 | 1.976591 | -1.6176 | 3.476439 | 0.002214 | 0.006017 | 0.039774 | -1.6176 | 0.089107 | 0.929828 | 0.942768 | 14.09763 | 4.440175 | 5.207818 | 35.06515 | 18.25714 | 45.9907 | 24.16143 | 30.8297 | 45.96622 | 13.07665 | 16.01308 | 5.224389 | 40.88202 | 28.29897 | 32.39481 |
| Nsd3 | 234135 | 1.976767 | 7.981391 | 49.19463 | 1.85E-23 | 1.89E-21 | -0.20991 | 7.981391 | -6.5253 | 1.69E-06 | 6.98E-06 | 8353.961 | 8561.545 | 8865.442 | 35956.06 | 36640.67 | 35626.56 | 33295.88 | 33737.07 | 30512.38 | 10365.93 | 10543.47 | 9911.412 | 7557.333 | 7558.184 | 7707.535 |
| Larp6 | 67557 | 1.976886 | -2.88717 | 2.47555 | 0.021754 | 0.048048 | 1.824252 | -2.88717 | 3.131309 | 0.004978 | 0.011636 | 2.225942 | 3.55214 | 0.86797 | 7.792255 | 15.44835 | 6.763339 | 54.00791 | 28.14886 | 32.17636 | 0 | 1.525056 | 5.224389 | 23.36115 | 30.65722 | 18.62702 |
| Mapk15 | 332110 | 1.97709 | -1.30539 | 2.967892 | 0.007256 | 0.01793 | 0.985449 | -1.30539 | 2.064282 | 0.051356 | 0.088635 | 3.709904 | 12.43249 | 3.471878 | 16.88322 | 21.06593 | 37.8747 | 52.58665 | 40.21265 | 56.30862 | 23.84565 | 6.862751 | 11.19512 | 85.93567 | 125.7732 | 68.0291 |
| Fpr2 | 14289 | 1.978156 | 5.986361 | 55.7834 | 1.29E-24 | 1.93E-22 | -1.39743 | 5.986361 | -40.0053 | 1.46E-21 | 1.51E-19 | 5610.858 | 5474.736 | 5414.394 | 22542.99 | 24002.52 | 22987.24 | 8807.553 | 9419.144 | 9133.489 | 1921.498 | 1933.008 | 1705.39 | 370.4412 | 352.165 | 366.8712 |
| Rcn1 | 19672 | 1.978286 | 3.192389 | 18.77977 | 9.38E-15 | 1.30E-13 | -0.21509 | 3.192389 | -2.57786 | 0.017411 | 0.034996 | 311.6319 | 335.6772 | 318.5448 | 1411.697 | 1353.837 | 1291.798 | 1334.564 | 1060.274 | 1247.983 | 383.838 | 468.1921 | 457.5072 | 223.5996 | 224.8196 | 189.5096 |
| Abcg5 | 27409 | 1.980014 | -2.64599 | 3.010834 | 0.006575 | 0.016391 | 0.184198 | -2.64599 | 0.346891 | 0.73207 | 0.810702 | 5.193865 | 4.440175 | 2.603909 | 9.090964 | 21.06593 | 29.75869 | 21.31891 | 16.08506 | 26.43058 | 2.307643 | 0.762528 | 5.224389 | 10.84625 | 16.50774 | 28.34546 |
| Gm17435 | 100503374 | 1.980263 | 2.479542 | 14.94856 | 8.77E-13 | 8.58E-12 | -1.90764 | 2.479542 | -13.0682 | 1.18E-11 | 1.09E-10 | 240.4018 | 264.6344 | 303.7894 | 1228.579 | 1151.604 | 1005.032 | 402.2168 | 277.4673 | 278.0957 | 252.3023 | 243.2464 | 230.6194 | 221.931 | 184.7294 | 137.6779 |
| Ms4a19 | 69416 | 1.98085 | -3.06064 | 2.829293 | 0.009946 | 0.0239 | 1.924952 | -3.06064 | 3.945885 | 0.000721 | 0.00203 | 5.193865 | 2.664105 | 1.735939 | 18.18193 | 7.021976 | 18.93735 | 54.00791 | 56.29771 | 56.30862 | 3.076858 | 2.287584 | 0 | 5.005962 | 6.288661 | 2.429611 |
| Nxph3 | 104079 | 1.981633 | -2.37132 | 3.347393 | 0.003003 | 0.007968 | 0.417439 | -2.37132 | 0.897534 | 0.379447 | 0.477756 | 4.451884 | 5.32821 | 2.603909 | 20.77935 | 15.44835 | 17.58468 | 21.31891 | 20.10633 | 34.47467 | 2.307643 | 3.812639 | 5.97073 | 31.70442 | 29.08506 | 14.57766 |
| Gm16677 | 100504477 | 1.983079 | -3.87852 | 2.839447 | 0.00972 | 0.023409 | 0.502336 | -3.87852 | 0.858513 | 0.400148 | 0.499327 | 2.967923 | 1.77607 | 0.86797 | 6.493546 | 8.426372 | 10.82134 | 14.21261 | 9.382952 | 14.93902 | 0 | 0 | 0 | 5.840288 | 21.22423 | 10.52831 |
| Aknad1 | 329738 | 1.983627 | -2.71934 | 2.820214 | 0.010152 | 0.02434 | 1.015368 | -2.71934 | 1.902118 | 0.070737 | 0.116272 | 3.709904 | 0.888035 | 5.207818 | 12.98709 | 7.021976 | 21.64268 | 32.689 | 28.14886 | 22.98311 | 3.076858 | 0.762528 | 3.731706 | 15.85221 | 30.65722 | 17.00728 |
| Hoxd9 | 15438 | 1.984824 | -3.12412 | 2.65279 | 0.014761 | 0.034053 | 0.774369 | -3.12412 | 1.297145 | 0.208429 | 0.290431 | 1.483961 | 1.77607 | 4.339848 | 6.493546 | 15.44835 | 12.17401 | 19.89765 | 12.0638 | 28.72889 | 0 | 0 | 2.239024 | 23.36115 | 36.1598 | 46.16261 |
| Pde5a | 242202 | 1.985219 | 3.097636 | 11.61336 | 1.09E-10 | 7.73E-10 | 1.4024 | 3.097636 | 12.70613 | 2.01E-11 | 1.78E-10 | 115.007 | 183.8232 | 167.5181 | 550.6527 | 734.4987 | 639.8118 | 1884.592 | 1817.612 | 1575.492 | 273.0711 | 256.2094 | 262.7121 | 730.0361 | 803.3764 | 794.4827 |
| Fam13c | 71721 | 1.985283 | -2.60726 | 2.488897 | 0.021135 | 0.046802 | 0.7511 | -2.60726 | 1.211092 | 0.2391 | 0.325681 | 8.903769 | 0.888035 | 2.603909 | 19.48064 | 9.830767 | 13.52668 | 15.63387 | 17.42548 | 48.26453 | 7.692145 | 6.100223 | 0.746341 | 15.85221 | 21.22423 | 21.8665 |
| Chrna10 | 504186 | 1.986221 | -2.0211 | 3.475051 | 0.002221 | 0.006035 | 0.883299 | -2.0211 | 2.1178 | 0.046092 | 0.08076 | 2.967923 | 4.440175 | 9.547666 | 16.88322 | 23.87472 | 27.05335 | 29.84648 | 44.23392 | 56.30862 | 1.538429 | 6.100223 | 4.478048 | 40.04769 | 29.87114 | 29.15533 |
| Foxp2 | 114142 | 1.987008 | -1.76521 | 3.260737 | 0.003681 | 0.009632 | 0.336834 | -1.76521 | 0.72955 | 0.473596 | 0.572061 | 15.5816 | 7.992315 | 1.735939 | 31.16902 | 21.06593 | 33.81669 | 25.5827 | 36.19139 | 51.712 | 4.615287 | 7.625279 | 4.478048 | 58.40288 | 33.01547 | 34.82442 |
| Tor1aip2 | 240832 | 1.987243 | 8.853536 | 80.02001 | 6.06E-28 | 3.14E-25 | -0.03593 | 8.853536 | -1.91089 | 0.069543 | 0.114581 | 15549.69 | 15776.83 | 15838.71 | 65769.23 | 69393.98 | 64588.53 | 68027.23 | 66458.11 | 67554.26 | 20911.09 | 20469.3 | 20120.61 | 11441.96 | 10921.83 | 11207.79 |
| Oprd1 | 18386 | 1.989164 | -1.22696 | 3.807647 | 0.001005 | 0.00289 | 1.1324 | -1.22696 | 3.164368 | 0.00461 | 0.010859 | 5.193865 | 12.43249 | 7.811726 | 36.36386 | 25.27911 | 43.28537 | 108.0158 | 65.68067 | 71.24765 | 4.615287 | 16.77561 | 10.44878 | 53.39692 | 51.09537 | 38.0639 |
| Dio3 | 107585 | 1.992148 | -3.30715 | 2.732552 | 0.012361 | 0.029041 | 0.46286 | -3.30715 | 0.763617 | 0.453455 | 0.552323 | 0.741981 | 6.216245 | 1.735939 | 10.38967 | 8.426372 | 10.82134 | 17.05513 | 10.72337 | 16.08818 | 0 | 3.050111 | 0.746341 | 20.02385 | 18.86598 | 13.76779 |
| Sptssb | 66183 | 1.992185 | -2.99214 | 2.539436 | 0.018937 | 0.042482 | 1.946684 | -2.99214 | 3.390758 | 0.002711 | 0.006761 | 0.741981 | 0.888035 | 6.075787 | 11.68838 | 7.021976 | 6.763339 | 35.53152 | 40.21265 | 31.0272 | 9.230573 | 7.625279 | 1.492683 | 5.840288 | 8.646909 | 4.859222 |
| Gm9999 | 629141 | 1.992841 | -4.16004 | 2.694912 | 0.013444 | 0.03131 | 1.227921 | -4.16004 | 2.081894 | 0.049567 | 0.085971 | 0.741981 | 2.664105 | 0.86797 | 6.493546 | 5.617581 | 6.763339 | 19.89765 | 14.74464 | 14.93902 | 0 | 0 | 2.239024 | 2.502981 | 3.930413 | 1.619741 |
| Dnah9 | 237806 | 1.993906 | 0.731275 | 9.31309 | 5.72E-09 | 3.19E-08 | 0.340504 | 0.731275 | 2.218808 | 0.037466 | 0.06771 | 43.03488 | 45.28978 | 45.13442 | 209.0922 | 178.3582 | 182.6101 | 224.5592 | 256.0206 | 266.6041 | 43.84522 | 44.98914 | 33.58536 | 98.45058 | 125.7732 | 108.5226 |
| Stfa3 | 20863 | 1.99489 | -3.17309 | 2.875396 | 0.00896 | 0.021725 | -0.28051 | -3.17309 | -0.45996 | 0.650193 | 0.737609 | 3.709904 | 7.10428 | 0.86797 | 15.58451 | 21.06593 | 8.116006 | 7.106304 | 13.40422 | 16.08818 | 3.076858 | 8.387806 | 4.478048 | 4.171635 | 6.288661 | 1.619741 |
| Lgi1 | 56839 | 1.994947 | -2.59427 | 3.03088 | 0.006279 | 0.015719 | -0.09353 | -2.59427 | -0.17379 | 0.863667 | 0.918721 | 7.419807 | 4.440175 | 3.471878 | 31.16902 | 21.06593 | 16.23201 | 38.37404 | 10.72337 | 24.13227 | 1.538429 | 5.337695 | 0 | 20.02385 | 21.22423 | 24.29611 |
| Adamdec1 | 58860 | 1.995194 | -3.42925 | 2.596988 | 0.016695 | 0.037931 | 0.694767 | -3.42925 | 1.105801 | 0.281122 | 0.372915 | 2.225942 | 1.77607 | 1.735939 | 11.68838 | 9.830767 | 6.763339 | 8.527565 | 13.40422 | 31.0272 | 3.076858 | 1.525056 | 0 | 7.508942 | 8.646909 | 16.19741 |
| Psmb10 | 19171 | 1.995401 | 6.614211 | 33.67822 | 5.45E-20 | 2.27E-18 | 0.355387 | 6.614211 | 7.85667 | 9.82E-08 | 4.82E-07 | 2851.432 | 2859.473 | 2804.41 | 11954.62 | 12334.8 | 11984.64 | 16991.17 | 16820.95 | 14688.51 | 5406.039 | 5264.492 | 5567.706 | 2244.339 | 2270.993 | 2379.399 |
| Cyp4f17 | 208285 | 1.996397 | 1.668246 | 10.60128 | 5.78E-10 | 3.71E-09 | -0.81138 | 1.668246 | -4.97667 | 6.06E-05 | 0.000202 | 62.32638 | 87.02743 | 78.11726 | 329.8721 | 271.0483 | 357.1043 | 177.6576 | 223.8504 | 168.9259 | 193.0728 | 195.2071 | 217.1853 | 266.9846 | 256.2629 | 285.0743 |
| Lrit3 | 242235 | 1.996687 | -3.88802 | 2.488935 | 0.021133 | 0.046802 | 0.762468 | -3.88802 | 1.140051 | 0.266898 | 0.357108 | 2.967923 | 2.664105 | 0 | 9.090964 | 4.213186 | 6.763339 | 11.37009 | 8.04253 | 17.23733 | 0 | 0 | 1.492683 | 12.5149 | 10.21907 | 10.52831 |
| Calcrl | 54598 | 1.996696 | 7.269551 | 60.11253 | 2.65E-25 | 5.15E-23 | -2.27935 | 7.269551 | -58.8696 | 4.12E-25 | 1.45E-22 | 9057.359 | 8684.094 | 9637.934 | 38526.21 | 40084.25 | 38034.31 | 8133.876 | 8003.658 | 8733.582 | 4576.826 | 4448.587 | 4558.652 | 4976.76 | 4707.849 | 5392.926 |
| Pigv | 230801 | 1.997134 | 4.298822 | 30.1807 | 5.40E-19 | 1.80E-17 | -0.87316 | 4.298822 | -14.6785 | 1.25E-12 | 1.38E-11 | 870.3434 | 858.7298 | 828.043 | 3613.009 | 3769.397 | 3534.521 | 2126.206 | 2151.377 | 1930.581 | 602.2949 | 704.5757 | 622.4486 | 533.1349 | 522.7449 | 587.1559 |
| Col28a1 | 213945 | 1.997392 | -2.17407 | 3.358974 | 0.002922 | 0.007769 | 0.972742 | -2.17407 | 2.246605 | 0.035364 | 0.064511 | 7.419807 | 5.32821 | 1.735939 | 20.77935 | 11.23516 | 29.75869 | 31.26774 | 40.21265 | 48.26453 | 1.538429 | 3.812639 | 2.985365 | 43.385 | 33.80155 | 34.01455 |
| Svop | 68666 | 1.998856 | -1.71907 | 3.191655 | 0.004326 | 0.011162 | 0.98764 | -1.71907 | 2.206764 | 0.038411 | 0.069133 | 4.451884 | 11.54445 | 2.603909 | 37.66257 | 15.44835 | 20.29002 | 45.48035 | 67.02109 | 37.92213 | 16.92272 | 6.862751 | 3.731706 | 38.37904 | 38.51805 | 35.63429 |
| Fam83c | 71405 | 1.999697 | -1.94707 | 2.545754 | 0.018678 | 0.041972 | 0.525579 | -1.94707 | 0.890494 | 0.383129 | 0.481608 | 2.967923 | 7.10428 | 10.41564 | 23.37676 | 30.8967 | 25.70069 | 21.31891 | 40.21265 | 65.50187 | 8.461359 | 15.25056 | 0 | 36.71038 | 32.22939 | 34.82442 |
| Crybg1 | 11630 | 2.001596 | 7.167177 | 57.24916 | 7.45E-25 | 1.20E-22 | 0.385003 | 7.167177 | 14.7217 | 1.18E-12 | 1.31E-11 | 4718.998 | 4790.949 | 4715.679 | 20898.83 | 20244.36 | 19663.73 | 28187.87 | 27321.82 | 27024.69 | 5887.567 | 5959.918 | 5948.34 | 3089.513 | 2917.939 | 2947.928 |
| Islr | 26968 | 2.002289 | -3.03736 | 2.827164 | 0.009994 | 0.023993 | 0.963023 | -3.03736 | 1.696134 | 0.1044 | 0.162014 | 1.483961 | 3.55214 | 0.86797 | 6.493546 | 8.426372 | 10.82134 | 11.37009 | 13.40422 | 32.17636 | 4.615287 | 1.525056 | 2.985365 | 15.01788 | 20.43815 | 20.24676 |
| Trim30a | 20128 | 2.002826 | 8.707784 | 78.788 | 8.43E-28 | 3.96E-25 | -0.38805 | 8.707784 | -19.1483 | 6.34E-15 | 1.13E-13 | 18460.48 | 18894.72 | 17089.45 | 77721.25 | 79033.75 | 76197.13 | 61918.65 | 61301.51 | 61526.94 | 25502.54 | 24942.29 | 24820.32 | 4300.121 | 4487.746 | 4669.712 |
| Prkx | 19108 | 2.003066 | 6.876067 | 54.09504 | 2.48E-24 | 3.38E-22 | -1.11347 | 6.876067 | -32.1561 | 1.44E-19 | 8.28E-18 | 4519.405 | 4658.632 | 4486.535 | 19615.7 | 20198.01 | 18680.34 | 9391.692 | 9516.994 | 9158.77 | 8399.822 | 7956.978 | 8212.739 | 2502.146 | 2392.835 | 2638.557 |
| Apon | 28194 | 2.004776 | -0.12778 | 6.693904 | 1.16E-06 | 4.78E-06 | -1.13081 | -0.12778 | -4.11199 | 0.000483 | 0.0014 | 33.38913 | 31.96926 | 38.19066 | 203.8973 | 139.0351 | 113.6241 | 72.4843 | 99.19121 | 50.56285 | 49.99894 | 32.02617 | 53.73657 | 30.8701 | 27.51289 | 29.15533 |
| Trim55 | 381485 | 2.004855 | -3.94144 | 2.509448 | 0.020214 | 0.045001 | 1.251674 | -3.94144 | 1.92482 | 0.067684 | 0.111967 | 0.741981 | 0.888035 | 0.86797 | 2.597418 | 5.617581 | 6.763339 | 8.527565 | 16.08506 | 13.78987 | 3.076858 | 1.525056 | 0 | 5.840288 | 7.860826 | 8.098703 |
| Rbm46 | 633285 | 2.005227 | -3.5415 | 2.56623 | 0.01786 | 0.040307 | 0.834306 | -3.5415 | 1.319069 | 0.201124 | 0.281895 | 2.967923 | 1.77607 | 0.86797 | 7.792255 | 8.426372 | 9.468674 | 8.527565 | 17.42548 | 26.43058 | 2.307643 | 0 | 0 | 8.343269 | 20.43815 | 18.62702 |
| Xaf1 | 327959 | 2.005318 | 6.675728 | 46.18276 | 7.05E-23 | 5.89E-21 | -0.53696 | 6.675728 | -14.4774 | 1.64E-12 | 1.77E-11 | 4146.93 | 4129.363 | 3637.661 | 16946.86 | 17147.67 | 16971.92 | 12168.84 | 12618.73 | 11800.68 | 5467.576 | 5529.089 | 5502.774 | 1927.295 | 1948.699 | 1776.855 |
| Ppp1r15b | 108954 | 2.006549 | 8.288122 | 59.6355 | 3.13E-25 | 5.89E-23 | -0.57452 | 8.288122 | -20.4404 | 1.68E-15 | 3.43E-14 | 12460.08 | 12470.68 | 11983.19 | 53023.7 | 53779.91 | 51657.03 | 38318.61 | 37707.4 | 34884.92 | 12796.65 | 12581.71 | 12838.56 | 7328.728 | 7246.896 | 7778.804 |
| Zcchc2 | 227449 | 2.006879 | 7.849179 | 76.54272 | 1.56E-27 | 6.03E-25 | -0.50037 | 7.849179 | -22.891 | 1.66E-16 | 4.35E-15 | 8721.984 | 8857.261 | 8690.112 | 37865.16 | 37810.53 | 37029.28 | 28392.53 | 27425.03 | 27014.35 | 11886.67 | 11816.89 | 11920.56 | 4530.395 | 4575.787 | 4734.502 |
| Ampd1 | 229665 | 2.007039 | -1.7062 | 3.952266 | 0.00071 | 0.002089 | 0.837122 | -1.7062 | 2.302773 | 0.031444 | 0.058219 | 7.419807 | 2.664105 | 8.679696 | 35.06515 | 26.68351 | 17.58468 | 48.32287 | 54.95729 | 42.51876 | 6.92293 | 9.150334 | 6.717071 | 30.03577 | 35.37372 | 34.82442 |
| Kcnq2 | 16536 | 2.010517 | -1.56192 | 4.07396 | 0.00053 | 0.001588 | 0.343026 | -1.56192 | 0.937073 | 0.359205 | 0.45695 | 11.12971 | 10.65642 | 6.075787 | 33.76644 | 54.77142 | 35.16936 | 36.95278 | 44.23392 | 82.7392 | 3.076858 | 6.100223 | 2.239024 | 35.04173 | 57.38403 | 40.49351 |
| Ppp1r1b | 19049 | 2.010524 | -1.48332 | 4.105092 | 0.000491 | 0.001479 | -0.52235 | -1.48332 | -1.27607 | 0.215644 | 0.29868 | 17.06556 | 5.32821 | 13.88751 | 44.15611 | 60.389 | 44.63804 | 19.89765 | 38.87223 | 51.712 | 3.076858 | 5.337695 | 15.67317 | 27.53279 | 34.58764 | 37.25403 |
| Hspbap1 | 66667 | 2.011514 | 3.673289 | 18.09962 | 1.97E-14 | 2.58E-13 | -0.34275 | 3.673289 | -3.82928 | 0.000954 | 0.002623 | 428.1229 | 356.9901 | 379.3027 | 1758.452 | 1497.085 | 1784.169 | 1434.052 | 1436.932 | 1269.817 | 514.6045 | 569.6083 | 538.8584 | 489.7499 | 626.5078 | 491.5913 |
| Gpr82 | 319200 | 2.013921 | 0.443976 | 10.18531 | 1.19E-09 | 7.28E-09 | -2.46081 | 0.443976 | -9.93341 | 1.85E-09 | 1.18E-08 | 71.97213 | 92.35564 | 81.58914 | 368.8334 | 356.7164 | 325.9929 | 75.32683 | 73.7232 | 52.86116 | 77.69066 | 49.56431 | 32.83902 | 36.71038 | 29.08506 | 43.73299 |
| Rhd | 19746 | 2.014203 | 0.03407 | 5.36719 | 2.40E-05 | 8.48E-05 | 0.970213 | 0.03407 | 3.876017 | 0.000853 | 0.002369 | 20.77546 | 10.65642 | 23.43518 | 102.598 | 78.64613 | 55.45938 | 181.9214 | 163.5315 | 134.4512 | 38.46072 | 40.41398 | 35.07804 | 120.9774 | 99.04641 | 89.08573 |
| Tdgf1 | 21667 | 2.014647 | -2.52916 | 2.808347 | 0.010427 | 0.024904 | -0.68203 | -2.52916 | -1.03524 | 0.312144 | 0.407025 | 5.193865 | 1.77607 | 2.603909 | 14.2858 | 14.04395 | 13.52668 | 11.37009 | 4.021265 | 13.78987 | 12.30743 | 25.92595 | 7.463413 | 15.85221 | 12.57732 | 31.58494 |

| | | | | | | | | | | | | | | | | | | | | | | | | | | |
|---|---|---|---|---|---|---|---|---|---|---|---|---|---|---|---|---|---|---|---|---|---|---|---|---|---|---|
| Fam241b | 69894 | 2.015612 | -3.01933 | 2.631805 | 0.015462 | 0.035511 | 1.911931 | -3.01933 | 3.299406 | 0.003362 | 0.008198 | 1.483961 | 0.888035 | 0.86797 | 6.493546 | 4.213186 | 6.763339 | 14.21261 | 28.14886 | 33.32551 | 6.153716 | 6.100223 | 1.492683 | 19.18952 | 19.65207 | 26.72572 |
| Trim58 | 216781 | 2.016936 | -2.3499 | 3.666939 | 0.001407 | 0.003952 | 0.055522 | -2.3499 | 0.126132 | 0.900808 | 0.923042 | 2.967923 | 7.992315 | 6.943757 | 29.87031 | 16.85274 | 28.40602 | 14.21261 | 36.19139 | 33.32551 | 1.538429 | 1.525056 | 2.985365 | 23.36115 | 28.29897 | 21.05663 |
| Gpr85 | 64450 | 2.018198 | 4.424847 | 31.16562 | 2.76E-19 | 9.85E-18 | -1.03434 | 4.424847 | -17.2904 | 4.93E-14 | 7.36E-13 | 1123.359 | 1195.295 | 1174.363 | 5054.576 | 5159.748 | 4873.662 | 2569.64 | 2498.546 | 2570.661 | 1132.284 | 1131.591 | 1062.044 | 153.5162 | 232.6805 | 261.5881 |
| Il13ra1 | 16164 | 2.018468 | 4.427528 | 27.78481 | 3.03E-18 | 8.63E-17 | 0.859798 | 4.427528 | 16.26858 | 1.66E-13 | 2.19E-12 | 1004.642 | 861.3939 | 999.901 | 3989.635 | 4356.434 | 4078.293 | 7508.521 | 7439.341 | 8364.704 | 369.9922 | 416.3402 | 331.3755 | 300.3577 | 334.8712 | 293.9829 |
| Gm4583 | 100043676 | 2.018832 | 4.648329 | 26.55146 | 7.77E-18 | 1.99E-16 | -1.13704 | 4.648329 | -15.9714 | 2.39E-13 | 3.03E-12 | 1196.815 | 1118.924 | 1117.945 | 4993.537 | 4944.876 | 4944.001 | 2363.557 | 2667.439 | 2068.48 | 750.7533 | 761.7653 | 625.434 | 832.6583 | 807.3069 | 752.3695 |
| Parp9 | 80285 | 2.020157 | 7.838155 | 56.66172 | 9.28E-25 | 1.44E-22 | -0.01052 | 7.838155 | -0.38073 | 0.707165 | 0.787866 | 7074.044 | 7246.365 | 6922.058 | 31315.77 | 30833.5 | 29820.91 | 32517.03 | 32136.61 | 30392.87 | 11138.23 | 10929.31 | 11056.3 | 6347.559 | 6457.669 | 6128.288 |
| Pcsk2 | 18549 | 2.020329 | -2.04822 | 2.86058 | 0.009266 | 0.022391 | 0.54687 | -2.04822 | 1.021994 | 0.318229 | 0.413565 | 0.741981 | 9.768385 | 17.35939 | 22.07806 | 23.87472 | 24.34802 | 32.689 | 25.46801 | 51.712 | 3.846072 | 6.100223 | 4.478048 | 17.52087 | 21.22423 | 37.25403 |
| Tcfl5 | 277353 | 2.020992 | -2.34231 | 3.179616 | 0.004449 | 0.011456 | 0.154404 | -2.34231 | 0.299013 | 0.767823 | 0.841613 | 3.709904 | 5.32821 | 2.603909 | 10.38967 | 16.85274 | 29.75869 | 15.63387 | 16.08506 | 29.87805 | 2.307643 | 5.337695 | 10.44878 | 23.36115 | 23.58248 | 34.82442 |
| Gm10335 | 100039571 | 2.021446 | -1.56135 | 4.792336 | 9.41E-05 | 0.000309 | -0.15669 | -1.56135 | -0.47196 | 0.641747 | 0.730011 | 12.61367 | 6.216245 | 18.22736 | 46.75353 | 47.74944 | 58.16471 | 28.42522 | 56.29771 | 60.90525 | 13.07665 | 16.77561 | 14.92683 | 7.508942 | 7.860826 | 7.288832 |
| Gm4956 | 241041 | 2.021597 | -2.79883 | 2.761906 | 0.011575 | 0.027357 | -0.69992 | -2.79883 | -1.03449 | 0.312485 | 0.407349 | 2.225942 | 0.888035 | 6.943757 | 5.194837 | 28.08791 | 12.17401 | 11.37009 | 6.702109 | 5.745778 | 16.92272 | 16.77561 | 16.41951 | 6.674615 | 14.93557 | 5.669092 |
| Ctla4 | 12477 | 2.021797 | -2.94224 | 2.525872 | 0.019505 | 0.043603 | 1.345128 | -2.94224 | 2.214875 | 0.037772 | 0.0682 | 4.451884 | 0 | 4.339848 | 12.98709 | 5.617581 | 9.468674 | 34.11026 | 25.46801 | 18.38649 | 3.076858 | 1.525056 | 5.224389 | 19.18952 | 14.14949 | 13.76779 |
| Gpr149 | 229357 | 2.021976 | -2.82394 | 2.889099 | 0.008685 | 0.021126 | 0.519951 | -2.82394 | 0.923316 | 0.366164 | 0.46394 | 2.225942 | 7.992315 | 0.86797 | 10.38967 | 12.63956 | 14.87935 | 14.21261 | 18.7659 | 24.13227 | 4.615287 | 0.762528 | 1.492683 | 35.04173 | 22.01031 | 19.43689 |
| Gm10851 | 100038528 | 2.022488 | 1.01122 | 10.88406 | 3.58E-10 | 2.37E-09 | -1.25613 | 1.01122 | -7.17623 | 4.08E-07 | 1.83E-06 | 78.64996 | 97.68385 | 117.1759 | 458.4443 | 396.0395 | 393.6263 | 177.6576 | 214.4675 | 157.4343 | 61.53716 | 99.12862 | 55.22925 | 45.88798 | 57.38403 | 44.54286 |
| Gm1141 | 382221 | 2.0232 | -2.53305 | 3.216253 | 0.004085 | 0.010588 | 0.153989 | -2.53305 | 0.303966 | 0.764098 | 0.838264 | 2.225942 | 2.664105 | 9.547666 | 12.98709 | 25.27911 | 17.58468 | 34.11026 | 14.74464 | 18.38649 | 5.384501 | 9.150334 | 5.97073 | 5.005962 | 11.00516 | 13.76779 |
| Col27a1 | 373864 | 2.025912 | 1.982238 | 9.704092 | 2.79E-09 | 1.63E-08 | 2.411917 | 1.982238 | 19.60514 | 3.93E-15 | 7.35E-14 | 84.58581 | 67.49066 | 66.83366 | 364.9373 | 275.2615 | 320.5823 | 1826.32 | 1875.25 | 1627.204 | 71.53694 | 50.32684 | 57.46828 | 335.3994 | 282.2037 | 281.8349 |
| Tmem171 | 380863 | 2.027222 | 3.857564 | 25.30992 | 2.10E-17 | 4.87E-16 | -1.7774 | 3.857564 | -21.1148 | 8.69E-16 | 1.92E-14 | 834.7283 | 870.2743 | 784.6445 | 3680.542 | 3487.113 | 3660.319 | 1071.631 | 979.8483 | 1219.254 | 997.6711 | 878.4321 | 861.2778 | 187.7236 | 121.8428 | 194.3689 |
| Gm19825 | 100503674 | 2.027265 | -2.44937 | 3.42413 | 0.002506 | 0.006742 | -0.20986 | -2.44937 | -0.41844 | 0.679808 | 0.76363 | 4.451884 | 1.77607 | 6.943757 | 15.58451 | 14.04395 | 27.05335 | 11.37009 | 16.08506 | 21.83396 | 1.538429 | 3.050111 | 5.97073 | 30.03577 | 38.51805 | 32.39481 |
| 9030425P( | 100504699 | 2.029431 | 2.286007 | 15.69244 | 3.38E-13 | 3.55E-12 | -1.34757 | 2.286007 | -10.8547 | 3.76E-10 | 2.67E-09 | 190.6891 | 248.6498 | 216.1244 | 949.3564 | 924.0921 | 948.2201 | 383.7404 | 423.5733 | 351.6416 | 257.6868 | 205.8825 | 172.4048 | 127.652 | 146.9974 | 108.5226 |
| Gm19705 | 100503460 | 2.030572 | -0.37429 | 6.801008 | 9.19E-07 | 3.83E-06 | 0.451532 | -0.37429 | 2.15491 | 0.042733 | 0.075762 | 30.42121 | 15.98463 | 32.11488 | 93.50706 | 137.6307 | 109.5661 | 149.2324 | 168.8931 | 158.5835 | 14.61507 | 9.912862 | 14.92683 | 40.04769 | 36.1598 | 26.72572 |
| Gm19660 | 100503370 | 2.030843 | -2.98531 | 2.790464 | 0.010856 | 0.025825 | 0.968948 | -2.98531 | 1.781377 | 0.089088 | 0.141499 | 2.967923 | 1.77607 | 6.075787 | 24.67547 | 8.426372 | 16.23201 | 31.26774 | 24.12759 | 40.22045 | 2.307643 | 0 | 4.478048 | 5.005962 | 6.288661 | 12.95792 |
| Stxbp1 | 20910 | 2.035023 | 5.247473 | 33.4596 | 6.25E-20 | 2.56E-18 | -0.75925 | 5.247473 | -14.1521 | 2.55E-12 | 2.66E-11 | 964.575 | 981.2787 | 996.4291 | 4532.495 | 4340.986 | 4005.249 | 2551.163 | 2588.354 | 2736.139 | 2684.558 | 2602.508 | 2480.092 | 2079.143 | 2087.835 | 2170.452 |
| Cyp3a25 | 56388 | 2.035847 | -4.03883 | 2.888662 | 0.008694 | 0.021141 | 0.251598 | -4.03883 | 0.406472 | 0.688443 | 0.770964 | 1.483961 | 0.888035 | 0.86797 | 5.194837 | 7.021976 | 5.410671 | 4.263783 | 6.702109 | 12.64071 | 0.769214 | 0.762528 | 0 | 8.343269 | 13.3634 | 9.718443 |
| Epp13 | 627821 | 2.035923 | -2.88341 | 3.051671 | 0.005985 | 0.015048 | 0.841865 | -2.88341 | 1.607743 | 0.122585 | 0.185454 | 1.483961 | 2.664105 | 2.603909 | 12.98709 | 11.23516 | 8.116006 | 29.84648 | 20.10633 | 14.93902 | 4.615287 | 6.862751 | 1.492683 | 17.52087 | 10.21907 | 12.95792 |
| Adam19 | 11492 | 2.037847 | 3.337587 | 26.85012 | 6.16E-18 | 1.64E-16 | -3.78119 | 3.337587 | -28.3217 | 2.03E-18 | 8.41E-17 | 609.1662 | 626.9527 | 704.7913 | 2811.705 | 2995.575 | 2680.987 | 261.512 | 223.8504 | 172.3733 | 461.5287 | 427.7781 | 476.9121 | 453.8738 | 462.2166 | 424.372 |
| Slc8a3 | 110893 | 2.038202 | -1.79608 | 2.869207 | 0.009087 | 0.021998 | 0.504943 | -1.79608 | 0.953813 | 0.350857 | 0.44803 | 7.419807 | 0.888035 | 18.22736 | 36.36386 | 21.06593 | 20.29002 | 29.84648 | 30.8297 | 52.86116 | 11.53822 | 9.912862 | 4.478048 | 44.21933 | 36.94588 | 20.24676 |
| Psg21 | 72242 | 2.038345 | -3.515 | 2.86516 | 0.00917 | 0.022184 | 0.062066 | -3.515 | 0.101755 | 0.919901 | 0.93519 | 2.967923 | 0.888035 | 3.471878 | 11.68838 | 9.830767 | 10.82134 | 15.63387 | 6.702109 | 14.93902 | 0 | 1.525056 | 0.746341 | 17.52087 | 13.3634 | 5.669092 |
| Adamts8 | 30806 | 2.039793 | -2.08403 | 3.039727 | 0.006152 | 0.015439 | 0.419968 | -2.08403 | 0.838519 | 0.411032 | 0.510067 | 10.38773 | 7.10428 | 2.603909 | 31.16902 | 25.27911 | 25.70069 | 27.00396 | 42.8935 | 45.96622 | 7.692145 | 6.100223 | 0 | 29.20144 | 28.29897 | 34.01455 |
| Srgn | 19073 | 2.039869 | 7.583135 | 58.47945 | 4.75E-25 | 8.44E-23 | 0.451871 | 7.583135 | 17.49345 | 3.90E-14 | 5.96E-13 | 3951.789 | 4150.675 | 4213.992 | 17628.68 | 18209.39 | 18164.98 | 25918.11 | 24598.08 | 26066.3 | 4736.053 | 4588.893 | 4663.886 | 22415.86 | 22366.41 | 22733.87 |
| Gm9758 | 381714 | 2.040306 | -2.46737 | 2.929854 | 0.007915 | 0.019415 | 0.813465 | -2.46737 | 1.563475 | 0.132648 | 0.198248 | 8.161788 | 0.888035 | 5.207818 | 14.2858 | 16.85274 | 21.64268 | 28.42522 | 17.42548 | 59.75609 | 1.538429 | 3.050111 | 3.731706 | 23.36115 | 23.58248 | 17.00728 |
| Gm10742 | 100038486 | 2.041572 | -2.92082 | 2.545939 | 0.01867 | 0.041958 | 0.889428 | -2.92082 | 1.461142 | 0.158543 | 0.230889 | 3.709904 | 1.77607 | 3.471878 | 10.38967 | 12.63956 | 20.29002 | 42.63783 | 13.40422 | 34.47467 | 0 | 1.525056 | 1.492683 | 29.20144 | 32.22939 | 8.908573 |
| Slc4a11 | 269356 | 2.042718 | 1.364844 | 6.766359 | 9.92E-07 | 4.11E-06 | 2.44643 | 1.364844 | 13.72841 | 4.58E-12 | 4.56E-11 | 25.96933 | 37.29747 | 63.36178 | 166.2348 | 183.9758 | 165.0255 | 1053.154 | 1003.976 | 885.999 | 90.76731 | 72.44015 | 69.40974 | 144.3386 | 213.0284 | 148.2063 |
| Ccl2 | 20296 | 2.042757 | 8.976411 | 37.75698 | 4.63E-14 | 5.66E-13 | -0.42548 | 8.976411 | -9.82095 | 3.64E-07 | 1.65E-06 | 15253.64 | 15069.07 | 15373.48 | 66995.21 | 67417.99 | 66667.58 | 54131.56 | 50383.77 | 51169.6 | 24231.79 | 23565.16 | 24913.62 | 18597.15 | 17718.3 | 21749.88 |
| Myoz2 | 59006 | 2.043771 | -3.47191 | 2.892958 | 0.008609 | 0.020956 | -0.73692 | -3.47191 | -1.13231 | 0.270064 | 0.360775 | 5.935846 | 1.77607 | 1.735939 | 10.38967 | 16.85274 | 13.52668 | 7.106304 | 10.72337 | 6.894934 | 3.076858 | 2.287584 | 0 | 9.177596 | 11.00516 | 4.049351 |
| Shroom3 | 27428 | 2.044615 | -0.80992 | 3.453322 | 0.002339 | 0.006332 | 1.978411 | -0.80992 | 5.278679 | 2.96E-05 | 0.000103 | 9.64575 | 12.43249 | 2.603909 | 44.15611 | 37.91867 | 18.93735 | 140.7048 | 124.6592 | 132.1529 | 9.230573 | 12.96297 | 23.88292 | 117.6401 | 110.8376 | 76.12781 |
| A230028O | 319487 | 2.044664 | 2.752422 | 15.90961 | 2.58E-13 | 2.77E-12 | 1.096624 | 2.752422 | 12.90985 | 1.49E-11 | 1.35E-10 | 216.6584 | 212.2404 | 228.276 | 933.7719 | 1001.334 | 964.4521 | 2231.38 | 2318.93 | 1932.88 | 149.9968 | 109.804 | 114.9366 | 186.8892 | 209.098 | 166.0234 |
| Shox2 | 20429 | 2.045472 | -1.72605 | 2.778641 | 0.011148 | 0.026458 | 1.069142 | -1.72605 | 2.057813 | 0.052027 | 0.089612 | 1.483961 | 8.88035 | 11.2836 | 19.48064 | 36.51428 | 20.29002 | 51.16539 | 48.25518 | 60.90525 | 0 | 3.812639 | 8.209754 | 117.6401 | 107.6933 | 80.98703 |
| Relb | 19698 | 2.045618 | 5.244871 | 28.3064 | 2.06E-18 | 6.04E-17 | 0.05451 | 5.244871 | 0.952252 | 0.35163 | 0.448703 | 1222.784 | 1125.14 | 1198.666 | 5306.526 | 5624.603 | 4764.096 | 5942.292 | 5863.005 | 5173.498 | 2607.637 | 2632.246 | 2755.492 | 614.8989 | 651.6625 | 665.7134 |
| Tdrd9 | 74691 | 2.045788 | -1.5368 | 3.635919 | 0.001515 | 0.00423 | 1.100925 | -1.5368 | 2.812942 | 0.01032 | 0.022195 | 3.709904 | 5.32821 | 10.41564 | 22.07806 | 21.06593 | 41.9327 | 39.7953 | 64.34024 | 81.59005 | 8.461359 | 9.150334 | 3.731706 | 48.39096 | 46.37887 | 55.07118 |
| 4930590J( | 381798 | 2.045838 | -1.82084 | 2.876169 | 0.008944 | 0.021691 | 0.83206 | -1.82084 | 1.619469 | 0.120029 | 0.182122 | 10.38773 | 8.88035 | 0.86797 | 18.18193 | 21.06593 | 28.40602 | 48.32287 | 30.8297 | 47.11538 | 4.615287 | 6.862751 | 14.92683 | 21.6925 | 53.45362 | 34.82442 |
| Nr4a2 | 18227 | 2.045847 | 2.044628 | 13.44567 | 6.84E-12 | 5.81E-11 | -1.1496 | 2.044628 | -8.25353 | 4.40E-08 | 2.27E-07 | 124.6528 | 188.2634 | 139.7431 | 670.1339 | 705.0064 | 578.9418 | 332.575 | 312.3183 | 274.6482 | 185.3807 | 203.5949 | 150.7609 | 156.0191 | 179.2268 | 180.6011 |
| Prss43 | 272643 | 2.046075 | -3.62545 | 2.676183 | 0.014015 | 0.032497 | 0.799628 | -3.62545 | 1.293177 | 0.209773 | 0.29201 | 2.967923 | 0 | 5.207818 | 7.792255 | 7.021976 | 10.82134 | 17.05513 | 9.382952 | 22.98311 | 0 | 0 | 0 | 22.52683 | 24.36856 | 22.67637 |
| Ccnb1ip1 | 239083 | 2.047449 | -3.78808 | 2.777818 | 0.011169 | 0.026498 | 0.023787 | -3.78808 | 0.036467 | 0.971249 | 0.976297 | 1.483961 | 1.77607 | 0.86797 | 9.090964 | 5.617581 | 6.763339 | 12.79135 | 4.021265 | 8.044089 | 2.307643 | 0.762528 | 0 | 15.85221 | 9.432991 | 13.76779 |
| Tex21 | 80384 | 2.047921 | -3.49059 | 2.634824 | 0.015359 | 0.035309 | 1.264259 | -3.49059 | 2.086461 | 0.049112 | 0.085295 | 0.741981 | 1.77607 | 2.603909 | 9.090964 | 9.830767 | 5.410671 | 15.63387 | 16.08506 | 32.17636 | 0 | 0.762528 | 2.239024 | 5.840288 | 15.72165 | 15.38754 |
| Zc3h7a | 106205 | 2.049414 | 7.502499 | 46.31278 | 6.64E-23 | 5.64E-21 | -0.28754 | 7.502499 | -8.05232 | 6.60E-08 | 3.32E-07 | 5972.203 | 6171.843 | 6291.044 | 27126.14 | 27315.49 | 26934.32 | 23568.77 | 24382.27 | 21590.34 | 6919.084 | 7034.319 | 6723.042 | 5724.317 | 5758.055 | 5403.454 |
| Chrnb3 | 108043 | 2.049429 | -2.29951 | 3.122861 | 0.005076 | 0.012934 | 0.710001 | -2.29951 | 1.462579 | 0.158153 | 0.230448 | 5.193865 | 3.55214 | 5.207818 | 29.87031 | 12.63956 | 25.70069 | 29.84648 | 25.46801 | 60.90525 | 4.615287 | 0.762528 | 1.492683 | 33.37308 | 18.0799 | 41.30338 |
| Mmp28 | 118453 | 2.050045 | 1.262041 | 9.017492 | 9.97E-09 | 5.38E-08 | 1.328579 | 1.262041 | 9.319173 | 5.66E-09 | 3.34E-08 | 67.52025 | 45.28978 | 49.47427 | 184.4167 | 279.4747 | 271.8862 | 667.9926 | 581.743 | 633.1847 | 52.30658 | 41.93903 | 38.80975 | 166.8654 | 175.2964 | 159.5444 |
| Lcp2 | 16822 | 2.051401 | 8.474111 | 70.17339 | 9.89E-27 | 3.25E-24 | -0.85586 | 8.474111 | -33.8152 | 5.01E-20 | 3.33E-18 | 16034.95 | 16531.66 | 15080.97 | 70131.59 | 71358.73 | 69322.87 | 41583.25 | 40672.42 | 38950.63 | 14078.16 | 13455.57 | 14016.29 | 6790.587 | 6818.481 | 7269.396 |
| Prdm6 | 225518 | 2.052916 | -2.19858 | 3.517183 | 0.00201 | 0.005501 | 0.971375 | -2.19858 | 2.26436 | 0.034079 | 0.062465 | 5.193865 | 4.440175 | 1.735939 | 16.88322 | 12.63956 | 21.64268 | 29.84648 | 38.87223 | 35.62382 | 1.538429 | 4.575167 | 1.492683 | 67.58048 | 50.30929 | 63.16988 |
| Arxes1 | 76219 | 2.055454 | -4.64542 | 2.494035 | 0.020901 | 0.046339 | 0.623521 | -4.64542 | 0.869565 | 0.394212 | 0.493128 | 0 | 0.888035 | 0.86797 | 2.597418 | 2.808791 | 4.058003 | 5.685043 | 12.0638 | 2.298311 | 0 | 0 | 0 | 12.5149 | 7.074744 | 10.52831 |
| BB123696 | 105404 | 2.056796 | -2.56145 | 2.65841 | 0.014579 | 0.033667 | 0.866949 | -2.56145 | 1.416977 | 0.170922 | 0.245638 | 2.225942 | 1.77607 | 1.735939 | 12.98709 | 8.426372 | 8.116006 | 27.00396 | 12.0638 | 20.6848 | 10.769 | 12.20045 | 2.239024 | 22.52683 | 45.59279 | 29.9652 |
| Fbll1 | 237730 | 2.060318 | -3.13171 | 2.487739 | 0.021188 | 0.046904 | 1.594395 | -3.13171 | 2.494645 | 0.020873 | 0.040933 | 1.483961 | 3.55214 | 0 | 2.597418 | 14.04395 | 5.410671 | 19.89765 | 16.08506 | 24.13227 | 1.538429 | 3.050111 | 1.492683 | 34.2074 | 37.73197 | 24.29611 |
| Gm5114 | 330513 | 2.060354 | -3.15028 | 2.499706 | 0.020646 | 0.045835 | 0.650401 | -3.15028 | 0.973103 | 0.341402 | 0.438524 | 1.483961 | 0.888035 | 4.339848 | 20.77935 | 5.617581 | 6.763339 | 15.63387 | 18.7659 | 13.78987 | 1.538429 | 6.862751 | 0 | 20.85817 | 22.7964 | 18.62702 |
| Slc4a9 | 240215 | 2.06088 | -2.71937 | 2.632368 | 0.015443 | 0.035476 | 1.075154 | -2.71937 | 1.838114 | 0.080006 | 0.129156 | 4.451884 | 1.77607 | 2.603909 | 23.37676 | 7.021976 | 14.87935 | 18.47639 | 32.17012 | 44.81707 | 4.615287 | 3.812639 | 0 | 20.85817 | 18.86598 | 26.72572 |
| Crem | 12916 | 2.062347 | 3.712622 | 18.07912 | 2.02E-14 | 2.63E-13 | 1.051425 | 3.712622 | 13.79042 | 4.20E-12 | 4.22E-11 | 298.2763 | 281.5071 | 237.8237 | 1279.229 | 1228.846 | 1143.004 | 2649.23 | 2436.887 | 2757.973 | 922.2881 | 751.8525 | 817.2437 | 438.0216 | 406.4047 | 396.0266 |
| Morrbid | 100043424 | 2.063325 | 3.199568 | 21.6318 | 5.30E-16 | 9.32E-15 | 0.079931 | 3.199568 | 1.103438 | 0.282123 | 0.373866 | 435.5427 | 453.7859 | 396.6621 | 1775.335 | 2001.263 | 1973.542 | 2221.431 | 1969.08 | 2110.999 | 142.3047 | 209.6952 | 181.3609 | 144.3386 | 190.232 | 161.9741 |
| A530013C | 329562 | 2.063801 | 1.587513 | 10.92034 | 3.37E-10 | 2.24E-09 | 0.538884 | 1.587513 | 4.184214 | 0.000406 | 0.001192 | 86.06977 | 78.14708 | 68.5696 | 350.6515 | 359.5252 | 334.1089 | 493.1775 | 566.9984 | 517.12 | 95.38259 | 89.21576 | 102.9951 | 156.8535 | 101.4047 | 159.5444 |
| Ccdc25 | 67179 | 2.063862 | 4.945112 | 35.62522 | 1.68E-20 | 8.06E-19 | -0.51248 | 4.945112 | -10.4165 | 7.93E-10 | 5.35E-09 | 1150.812 | 1201.511 | 1022.468 | 5001.329 | 5189.241 | 4861.488 | 3624.215 | 3755.862 | 3573.874 | 1337.664 | 1416.014 | 1450.141 | 744.2196 | 743.6342 | 715.9253 |
| Epn3 | 71889 | 2.064203 | -1.5736 | 3.757393 | 0.001133 | 0.003233 | 1.585978 | -1.5736 | 4.231026 | 0.000363 | 0.001073 | 3.709904 | 7.10428 | 2.603909 | 24.67547 | 16.85274 | 17.58468 | 65.378 | 53.61687 | 67.80018 | 13.07665 | 9.912862 | 8.209754 | 48.39096 | 67.60311 | 46.97248 |
| Gm10864 | 100038573 | 2.064336 | -2.73284 | 2.632251 | 0.015447 | 0.035482 | 0.404447 | -2.73284 | 0.652252 | 0.521202 | 0.617762 | 8.903769 | 1.77607 | 1.735939 | 31.16902 | 8.426372 | 13.52668 | 12.79135 | 28.14886 | 26.43058 | 3.076858 | 4.575167 | 0 | 28.36712 | 25.15464 | 21.8665 |
| Igfbp3 | 16009 | 2.064562 | -2.88455 | 2.687322 | 0.013673 | 0.031801 | -0.02413 | -2.88455 | -0.03711 | 0.970745 | 0.975865 | 2.225942 | 4.440175 | 2.603909 | 12.98709 | 23.87472 | 9.468674 | 17.05513 | 5.361687 | 29.87805 | 3.846072 | 0.762528 | 2.985365 | 24.19548 | 25.94073 | 8.908573 |
| Gjc1 | 14615 | 2.064868 | -1.75792 | 2.988517 | 0.006921 | 0.017187 | 0.819213 | -1.75792 | 1.600709 | 0.12414 | 0.187437 | 2.225942 | 9.768385 | 2.603909 | 15.58451 | 26.68351 | 14.87935 | 25.5827 | 44.23392 | 34.47467 | 15.38429 | 13.7255 | 11.19512 | 40.04769 | 73.89177 | 28.34546 |
| H2-Q6 | 110557 | 2.065562 | 2.915126 | 8.160316 | 5.31E-08 | 2.60E-07 | 2.701189 | 2.915126 | 18.39634 | 1.42E-14 | 2.35E-13 | 90.52165 | 98.57188 | 91.13681 | 424.6779 | 369.356 | 462.6124 | 3227.683 | 2794.779 | 2538.485 | 176.9193 | 199.0198 | 159.717 | 957.8073 | 786.8687 | 1099.804 |
| Gm5431 | 432555 | 2.06619 | 7.005494 | 42.3958 | 4.30E-22 | 2.90E-20 | 0.098665 | 7.005494 | 2.623141 | 0.015761 | 0.032074 | 4129.123 | 4171.1 | 3516.145 | 18066.34 | 18054.91 | 16671.63 | 19634.72 | 18846.33 | 20094.13 | 7421.381 | 7210.463 | 7094.72 | 2641.479 | 2765.439 | 2807.01 |
| Gimap1 | 16205 | 2.067249 | -1.54629 | 3.529684 | 0.001951 | 0.005351 | 2.04702 | -1.54629 | 5.195809 | 3.60E-05 | 0.000124 | 6.677827 | 1.77607 | 2.603909 | 18.18193 | 9.830767 | 21.64268 | 65.378 | 77.74446 | 67.80018 | 23.84565 | 22.11331 | 27.61463 | 29.20144 | 34.58764 | 38.87377 |
| Tent5a | 212943 | 2.068112 | 7.669336 | 58.68542 | 4.41E-25 | 7.94E-23 | -0.19283 | 7.669336 | -6.87856 | 7.76E-07 | 3.35E-06 | 5541.854 | 5217.206 | 5682.597 | 24594.95 | 25022.11 | 24035.55 | 22312.37 | 22241.62 | 22301.66 | 12920.5 | 12972.12 | 12419.86 | 6783.912 | 6805.117 | 7336.615 |
| 4930518I1! | 74704 | 2.068621 | 1.885558 | 13.03658 | 1.23E-11 | 1.01E-10 | -1.35984 | 1.885558 | -9.02869 | 9.76E-09 | 5.52E-08 | 160.2678 | 135.8694 | 132.7993 | 658.4455 | 587.0372 | 687.1552 | 315.5199 | 217.1483 | 258.56 | 167.6888 | 227.2333 | 179.1219 | 89.27298 | 133.634 | 126.3398 |
| Notch2 | 18129 | 2.068831 | 9.384907 | 75.02125 | 2.39E-27 | 8.85E-25 | -1.13823 | 9.384907 | -45.5844 | 9.29E-23 | 1.63E-20 | 23557.89 | 23429.91 | 23738.97 | 106720.1 | 106103.5 | 103989 | 51810.64 | 51319.39 | 47022.3 | 27899.41 | 28505.58 | 28257.23 | 28568.19 | 28901.9 | 29574.03 |
| Irgm2 | 54396 | 2.069022 | 8.029076 | 58.44777 | 4.80E-25 | 8.48E-23 | 0.876158 | 8.029076 | 36.26809 | 1.15E-20 | 9.12E-19 | 7033.977 | 6972.851 | 6563.586 | 31466.42 | 30513.3 | 30211.83 | 58862.94 | 59270.77 | 57718.64 | 10584.39 | 10058.5 | 10261.45 | 7575.688 | 7217.025 | 7110.661 |
| Gm11626 | 100502949 | 2.069634 | 2.131119 | 15.3494 | 5.22E-13 | 5.31E-12 | -0.19576 | 2.131119 | -1.91113 | 0.069512 | 0.114536 | 173.6235 | 140.3095 | 165.7822 | 720.7836 | 714.8372 | 720.9719 | 581.2957 | 718.4661 | 657.317 | 166.9195 | 216.5579 | 194.0487 | 69.24913 | 95.90208 | 93.94495 |
| Tle3 | 21887 | 2.069753 | 6.638766 | 51.88631 | 6.00E-24 | 7.27E-22 | -0.90527 | 6.638766 | -25.4526 | 1.87E-17 | 6.08E-16 | 4700.448 | 4924.154 | 5044.639 | 21733.9 | 22148.72 | 21759.01 | 12407.61 | 12717.92 | 11397.33 | 5220.658 | 5417.76 | 5275.886 | 1089.631 | 1047.848 | 1122.48 |
| Trim34b | 434218 | 2.070279 | -0.10403 | 5.744202 | 9.96E-06 | 3.68E-05 | 0.750122 | -0.10403 | 3.100026 | 0.005352 | 0.01241 | 32.64715 | 14.20856 | 11.2836 | 93.50706 | 68.81537 | 83.8654 | 159.1812 | 143.4251 | 131.0037 | 45.38365 | 35.07628 | 41.79511 | 53.39692 | 73.89177 | 57.50079 |
| Mov10 | 17454 | 2.07028 | 6.498139 | 42.74035 | 3.62E-22 | 2.47E-20 | -1.27925 | 6.498139 | -27.8045 | 2.98E-18 | 1.17E-16 | 3883.527 | 3920.674 | 3782.612 | 17749.46 | 17216.48 | 16988.15 | 7970.431 | 7688.659 | 6732.903 | 8170.596 | 7931.052 | 7835.09 | 1131.347 | 1113.093 | 1235.862 |
| Epsti1 | 108670 | 2.071157 | 6.648199 | 44.46302 | 1.57E-22 | 1.19E-20 | -0.53508 | 6.648199 | -13.6334 | 5.24E-12 | 5.16E-11 | 4173.642 | 4380.677 | 4080.325 | 19084.53 | 19314.65 | 18254.25 | 14047.74 | 14082.47 | 12638.41 | 5996.027 | 6051.421 | 6011.032 | 1141.359 | 1139.82 | 1303.081 |
| Akt3 | 23797 | 2.072547 | 6.38358 | 49.26471 | 1.80E-23 | 1.84E-21 | -0.83944 | 6.38358 | -22.5035 | 2.36E-16 | 5.97E-15 | 3189.033 | 3269.745 | 3337.343 | 15096.2 | 14777.05 | 14085.33 | 8656.9 | 8846.784 | 8072.818 | 2985.321 | 3069.937 | 3130.155 | 2442.909 | 2485.593 | 2516.267 |
| Tmcc3 | 319880 | 2.072644 | 5.294092 | 43.88187 | 2.08E-22 | 1.50E-20 | -0.34309 | 5.294092 | -8.74398 | 1.68E-08 | 9.22E-08 | 1739.203 | 1842.673 | 1755.903 | 8089.659 | 7983.987 | 7869.821 | 6645.816 | 6565.386 | 6401.946 | 2539.946 | 2599.457 | 2361.424 | 304.5293 | 300.2836 | 311.8001 |
| Vmn2r118 | 383258 | 2.073035 | -3.0505 | 3.186259 | 0.004381 | 0.011291 | 0.682445 | -3.0505 | 1.309935 | 0.204143 | 0.285457 | 1.483961 | 1.77607 | 3.471878 | 10.38967 | 8.426372 | 13.52668 | 11.37009 | 21.44675 | 22.98311 | 2.307643 | 3.812639 | 0.746341 | 18.35519 | 13.3634 | 15.38754 |
| Kynu | 70789 | 2.073318 | 0.922061 | 8.80181 | 1.50E-08 | 7.90E-08 | -1.36262 | 0.922061 | -6.0441 | 5.00E-06 | 1.93E-05 | 37.84102 | 47.95389 | 54.68209 | 228.5728 | 212.0637 | 182.6101 | 73.90556 | 93.82952 | 82.7392 | 319.9932 | 300.436 | 247.7853 | 92.61029 | 69.17527 | 77.74755 |
| Gsap | 212167 | 2.073608 | 7.039102 | 59.36177 | 3.46E-25 | 6.36E-23 | -0.0602 | 7.039102 | -2.19676 | 0.039212 | 0.070391 | 4421.463 | 4568.052 | 4431.853 | 20293.63 | 20436.76 | 19564.99 | 20360.98 | 20338.22 | 19427.62 | 3860.687 | 3802.726 | 3897.394 | 4217.523 | 4415.426 | 4342.524 |
| Tank | 21353 | 2.073636 | 7.172116 | 55.58536 | 1.39E-24 | 2.07E-22 | -0.64334 | 7.172116 | -20.1546 | 2.24E-15 | 4.46E-14 | 4497.887 | 4544.075 | 4332.904 | 20201.42 | 20112.34 | 19805.76 | 13513.35 | 13535.58 | 12963.62 | 6932.93 | 6871.138 | 6805.886 | 5464.007 | 5989.95 | 6012.477 |
| Agrn | 11603 | 2.073711 | 6.193457 | 29.80959 | 7.00E-19 | 2.26E-17 | 0.574734 | 6.193457 | 11.17511 | 2.21E-10 | 1.63E-09 | 2092.386 | 2014.951 | 1966.819 | 9005.249 | 9297.097 | 9041.231 | 14837.96 | 14846.51 | 12876.29 | 2649.944 | 2731.375 | 2551.741 | 2193.445 | 2513.106 | 2263.587 |
| Tpst1 | 22021 | 2.074282 | 5.404383 | 44.83444 | 1.32E-22 | 1.01E-20 | -0.58165 | 5.404383 | -14.6512 | 1.30E-12 | 1.43E-11 | 1761.462 | 1866.65 | 1860.927 | 8149.4 | 8322.446 | 8172.818 | 5999.142 | 5578.835 | 5548.123 | 1839.961 | 1943.683 | 1912.873 | 651.6093 | 605.2836 | 659.2344 |
| Cpxm2 | 55987 | 2.074301 | -2.32642 | 3.054628 | 0.005944 | 0.014956 | 1.070072 | -2.32642 | 2.153983 | 0.042815 | 0.07587 | 1.483961 | 6.216245 | 4.339848 | 18.18193 | 11.23516 | 21.64268 | 35.53152 | 25.46801 | 51.712 | 7.692145 | 3.812639 | 2.239024 | 21.6925 | 33.80155 | 13.76779 |
| Ephx3 | 71932 | 2.074497 | -1.43042 | 3.510855 | 0.00204 | 0.005578 | 0.856046 | -1.43042 | 2.055984 | 0.052219 | 0.089906 | 4.451884 | 2.664105 | 19.9633 | 50.64966 | 22.47032 | 21.64268 | 54.00791 | 44.23392 | 70.09849 | 16.1535 | 11.43792 | 12.6878 | 37.54471 | 40.09021 | 27.53559 |
| Sox17 | 20671 | 2.076764 | -2.29413 | 3.112762 | 0.005197 | 0.013202 | 0.446179 | -2.29413 | 0.854289 | 0.402432 | 0.501534 | 2.225942 | 4.440175 | 4.339848 | 12.98709 | 18.25714 | 20.29002 | 12.79135 | 26.80843 | 36.77298 | 1.538429 | 7.625279 | 8.956095 | 32.53875 | 36.1598 | 24.29611 |
| Txndc2 | 213272 | 2.081874 | -2.19943 | 4.561907 | 0.000164 | 0.000522 | -2.05977 | -2.19943 | -4.15157 | 0.000439 | 0.001281 | 13.35565 | 10.65642 | 8.679696 | 50.64966 | 30.8967 | 79.8074 | 12.79135 | 13.40422 | 10.3424 | 3.076858 | 3.050111 | 5.224389 | 18.35519 | 15.72165 | 6.478962 |
| Il12rb1 | 16161 | 2.08323 | 0.723047 | 7.57699 | 1.75E-07 | 8.00E-07 | -0.80251 | 0.723047 | -3.4433 | 0.002395 | 0.006044 | 45.26083 | 45.28978 | 36.45472 | 144.1567 | 224.7032 | 223.1902 | 112.2796 | 112.5954 | 117.2139 | 140.7662 | 179.9566 | 185.839 | 75.08942 | 66.81702 | 39.68364 |
| Uba7 | 74153 | 2.08395 | 7.797064 | 36.26131 | 1.05E-16 | 2.14E-15 | -0.03614 | 7.797064 | -0.82017 | 0.424257 | 0.523204 | 7089.626 | 7431.965 | 7004.515 | 32322.27 | 32626.91 | 32409.92 | 34056.25 | 34881.8 | 30284.85 | 8862.889 | 9173.21 | 8417.237 | 6097.261 | 6092.926 | 6202.796 |
| Gm3716 | 100042198 | 2.086584 | -2.61199 | 3.4638 | 0.002281 | 0.006186 | 0.633264 | -2.61199 | 1.339354 | 0.194545 | 0.274091 | 2.225942 | 1.77607 | 4.339848 | 20.77935 | 12.63956 | 8.116006 | 22.74017 | 17.42548 | 24.13227 | 3.846072 | 6.100223 | 3.731706 | 18.35519 | 25.15464 | 17.81715 |
| Snai1 | 20613 | 2.088781 | -2.06644 | 2.782978 | 0.01104 | 0.02622 | 0.961644 | -2.06644 | 1.792163 | 0.087296 | 0.138998 | 1.483961 | 11.54445 | 6.075787 | 12.98709 | 40.72746 | 21.64268 | 42.63783 | 36.19139 | 59.75609 | 3.076858 | 1.525056 | 2.239024 | 63.40885 | 20.43815 | 51.02183 |
| Cspg4b | 408066 | 2.088782 | -0.97183 | 4.213025 | 0.000379 | 0.001157 | 2.441521 | -0.97183 | 8.172775 | 5.18E-08 | 2.65E-07 | 14.83961 | 5.32821 | 5.207818 | 37.66257 | 49.15383 | 25.70069 | 210.3466 | 187.659 | 222.9362 | 6.92293 | 12.20045 | 11.94146 | 44.21933 | 50.30929 | 31.58494 |
| Ccser1 | 232035 | 2.089673 | 1.104299 | 10.95928 | 3.16E-10 | 2.11E-09 | -1.068 | 1.104299 | -6.28848 | 2.87E-06 | 1.15E-05 | 74.19807 | 68.37869 | 66.83366 | 303.8979 | 396.0395 | 271.8862 | 143.5473 | 174.2548 | 156.2852 | 196.1497 | 240.1963 | 235.0975 | 45.05365 | 43.23454 | 42.92312 |
| Rgs16 | 19734 | 2.090909 | 0.735344 | 8.953505 | 1.13E-08 | 6.02E-08 | -1.33271 | 0.735344 | -6.02853 | 5.18E-06 | 2.00E-05 | 50.45469 | 42.62568 | 48.6063 | 185.7154 | 214.8725 | 255.6542 | 61.11422 | 104.5529 | 104.5732 | 139.2278 | 160.1308 | 132.1024 | 67.58048 | 71.53352 | 84.22651 |
| 5430405H( | 74487 | 2.09219 | 0.495031 | 11.0233 | 2.84E-10 | 1.90E-09 | -1.66238 | 0.495031 | -8.631 | 2.09E-08 | 1.13E-07 | 52.68063 | 54.17013 | 65.96569 | 277.9238 | 262.6219 | 243.4802 | 88.11817 | 89.80826 | 79.29174 | 76.92145 | 57.95212 | 55.22925 | 49.22529 | 45.59279 | 59.9304 |
| Mfsd6l | 215723 | 2.092471 | 0.424048 | 11.35457 | 1.65E-10 | 1.14E-09 | -2.98721 | 0.424048 | -11.4288 | 1.46E-10 | 1.11E-09 | 86.81175 | 122.5488 | 82.45711 | 401.3011 | 412.8922 | 495.0764 | 62.53548 | 57.63814 | 50.56285 | 58.4603 | 39.65145 | 52.24389 | 23.36115 | 18.86598 | 46.16261 |
| Stk32b | 64293 | 2.0926 | -2.61139 | 2.828445 | 0.009965 | 0.023933 | 1.21475 | -2.61139 | 2.165081 | 0.041852 | 0.074396 | 2.967923 | 0.888035 | 2.603909 | 9.090964 | 12.63956 | 9.468674 | 17.05513 | 24.12759 | 37.92213 | 9.230573 | 6.100223 | 2.985365 | 16.68654 | 16.50774 | 38.87377 |
| Ttc39c | 72747 | 2.093556 | 4.646216 | 20.51865 | 1.56E-15 | 2.49E-14 | 1.150219 | 4.646216 | 16.21532 | 1.77E-13 | 2.32E-12 | 775.3699 | 814.3281 | 725.6226 | 3236.383 | 3679.516 | 3656.261 | 8801.868 | 8452.7 | 7259.216 | 575.3724 | 486.4928 | 488.8535 | 680.8108 | 601.3532 | 698.1082 |
| Cyp7b1 | 13123 | 2.093697 | -2.68732 | 3.294129 | 0.003404 | 0.008953 | 0.366285 | -2.68732 | 0.744515 | 0.464684 | 0.563314 | 5.935846 | 3.55214 | 2.603909 | 20.77935 | 23.87472 | 13.52668 | 19.89765 | 28.14886 | 28.72889 | 3.846072 | 3.050111 | 0 | 17.52087 | 17.29382 | 18.62702 |
| Ky | 16716 | 2.095828 | -1.49585 | 4.297459 | 0.000309 | 0.000955 | -0.48044 | -1.49585 | -1.19194 | 0.246369 | 0.333973 | 6.677827 | 10.65642 | 12.15157 | 37.66257 | 54.77142 | 41.9327 | 39.7953 | 16.08506 | 50.56285 | 6.92293 | 9.150334 | 3.731706 | 44.21933 | 55.81187 | 34.82442 |
| Snord89 | 100217461 | 2.095836 | -0.12345 | 9.668983 | 2.98E-09 | 1.74E-08 | -2.27064 | -0.12345 | -8.83593 | 1.41E-08 | 7.79E-08 | 45.26083 | 65.71459 | 75.51336 | 324.6773 | 276.6659 | 228.6008 | 51.16539 | 50.93603 | 73.54596 | 23.84565 | 29.73859 | 26.12194 | 25.86413 | 18.86598 | 12.95792 |
| Foxl1 | 14241 | 2.097082 | -2.31654 | 3.190583 | 0.004337 | 0.011187 | 0.874484 | -2.31654 | 1.771226 | 0.090804 | 0.143765 | 1.483961 | 5.32821 | 3.471878 | 12.98709 | 12.63956 | 20.29002 | 25.5827 | 34.85097 | 27.57973 | 2.307643 | 0.762528 | 7.463413 | 44.21933 | 62.88661 | 53.45144 |
| Tnfrsf14 | 230979 | 2.097974 | 4.121062 | 27.35094 | 4.20E-18 | 1.16E-16 | -2.64354 | 4.121062 | -27.8607 | 2.86E-18 | 1.13E-16 | 1081.066 | 1119.812 | 1074.546 | 5080.55 | 5012.287 | 4872.309 | 976.4062 | 904.7847 | 669.9577 | 1557.659 | 1562.42 | 1389.687 | 204.4101 | 179.2268 | 150.6359 |
| Dusp28 | 67446 | 2.098661 | 2.420402 | 13.52876 | 6.07E-12 | 5.21E-11 | 0.957451 | 2.420402 | 9.567049 | 3.58E-09 | 2.18E-08 | 139.4924 | 107.4522 | 104.1564 | 516.8862 | 589.846 | 509.9557 | 1132.745 | 1159.465 | 983.6772 | 234.6104 | 257.7344 | 228.3804 | 205.2444 | 208.3119 | 198.4182 |
| Clic4 | 29876 | 2.098696 | 10.2449 | 72.92495 | 3.96E-19 | 1.36E-17 | -0.23708 | 10.2449 | -11.5724 | 1.89E-08 | 1.03E-07 | 43491.2 | 41998.73 | 43276.96 | 198948 | 200578.5 | 189960.5 | 176247.7 | 175202.5 | 168948.9 | 53408.1 | 54248.52 | 53642.53 | 25142.44 | 25057.17 | 26434.17 |
| Sertad4 | 214791 | 2.098944 | -2.13588 | 2.536815 | 0.019045 | 0.042696 | 2.12691 | -2.13588 | 3.551537 | 0.001852 | 0.004785 | 2.225942 | 4.440175 | 0 | 12.98709 | 5.617581 | 5.410671 | 31.26774 | 32.17012 | 44.81707 | 1.538429 | 1.525056 | 2.985365 | 367.9382 | 361.598 | 445.4286 |
| Irak2 | 108960 | 2.099359 | 6.671325 | 46.0223 | 7.59E-23 | 6.16E-21 | -0.83766 | 6.671325 | -20.953 | 1.02E-15 | 2.19E-14 | 4934.172 | 5153.267 | 4779.909 | 23061.18 | 23178.14 | 21752.25 | 13638.42 | 13641.47 | 12368.36 | 4516.058 | 4384.535 | 4799.721 | 1131.347 | 1338.699 | 1260.968 |
| A2ml1 | 232400 | 2.099932 | -1.81603 | 4.234879 | 0.00036 | 0.001101 | 1.063599 | -1.81603 | 3.065936 | 0.005791 | 0.013303 | 2.967923 | 7.992315 | 5.207818 | 27.27289 | 21.06593 | 22.99535 | 59.69296 | 44.23392 | 54.01031 | 7.692145 | 4.575167 | 4.478048 | 34.2074 | 33.01547 | 36.44416 |
| Rnf31 | 268749 | 2.100569 | 7.153849 | 23.69449 | 8.93E-12 | 7.45E-11 | 0.473396 | 7.153849 | 7.338089 | 7.07E-06 | 2.68E-05 | 4115.025 | 4346.931 | 4227.88 | 19624.79 | 19761.25 | 18658.7 | 30227.38 | 29317.7 | 24965.41 | 5561.42 | 5669.395 | 5233.345 | 3534.209 | 3577.462 | 3529.415 |
| Cdc20b | 238896 | 2.100956 | -3.41773 | 2.525392 | 0.019525 | 0.043642 | 0.66004 | -3.41773 | 0.963478 | 0.346098 | 0.443472 | 1.483961 | 0 | 6.943757 | 7.792255 | 4.213186 | 13.52668 | 9.948826 | 14.74464 | 14.93902 | 4.615287 | 2.287584 | 0.746341 | 10.01192 | 11.00516 | 9.718443 |
| Sim2 | 20465 | 2.102093 | -1.86116 | 3.460234 | 0.002301 | 0.006235 | 1.160223 | -1.86116 | 2.719024 | 0.01274 | 0.026666 | 2.225942 | 7.10428 | 5.207818 | 31.16902 | 14.04395 | 20.29002 | 45.48035 | 54.95729 | 48.26453 | 8.461359 | 3.812639 | 2.985365 | 33.37308 | 62.88661 | 47.78235 |
| Hivep1 | 110521 | 2.103597 | 6.984118 | 42.6968 | 3.70E-22 | 2.51E-20 | -0.42877 | 6.984118 | -10.5334 | 6.49E-10 | 4.45E-09 | 3805.619 | 3530.827 | 3589.054 | 16720.88 | 16751.63 | 16741.97 | 13244.73 | 13611.98 | 12050.05 | 7545.225 | 7611.553 | 7673.881 | 4078.19 | 3918.622 | 4411.363 |
| Usp12 | 22217 | 2.104354 | 6.637423 | 56.14588 | 1.13E-24 | 1.71E-22 | -1.13819 | 6.637423 | -32.9018 | 8.89E-20 | 5.52E-18 | 3994.082 | 4226.158 | 4052.55 | 18518.29 | 19516.88 | 18284.01 | 9097.491 | 8747.592 | 8717.494 | 4824.513 | 4733.773 | 4897.491 | 2147.557 | 2261.56 | 2436.09 |
| Ccnj | 240665 | 2.105603 | 3.723309 | 27.59454 | 3.49E-18 | 9.85E-17 | -2.68258 | 3.723309 | -27.8745 | 2.83E-18 | 1.12E-16 | 859.9557 | 927.1085 | 856.686 | 4055.869 | 4253.913 | 3832.108 | 731.9493 | 711.7639 | 552.7438 | 621.5253 | 542.1573 | 604.5364 | 198.5698 | 199.665 | 271.3065 |
| Atp1b4 | 67821 | 2.105901 | -1.09679 | 5.240169 | 3.24E-05 | 0.000113 | -0.49381 | -1.09679 | -1.5258 | 0.141743 | 0.209576 | 21.51744 | 9.768385 | 13.88751 | 54.54578 | 80.05053 | 74.39673 | 38.37404 | 62.99982 | 51.712 | 3.846072 | 6.100223 | 17.91219 | 40.88202 | 38.51805 | 43.73299 |
| Psg22 | 243862 | 2.106807 | -3.7121 | 2.68895 | 0.013623 | 0.031694 | 1.380887 | -3.7121 | 2.211154 | 0.038064 | 0.068631 | 0.741981 | 0.888035 | 0.86797 | 10.38967 | 2.808791 | 4.058003 | 8.527565 | 13.40422 | 25.28142 | 0.769214 | 1.525056 | 2.985365 | 13.34923 | 7.860826 | 7.288832 |
| Ak7 | 78801 | 2.107299 | -1.81782 | 3.617714 | 0.001582 | 0.004407 | 0.189555 | -1.81782 | 0.435067 | 0.667882 | 0.753257 | 20.03348 | 6.216245 | 1.735939 | 32.46773 | 25.27911 | 36.52203 | 45.48035 | 29.48928 | 37.92213 | 5.384501 | 3.812639 | 8.209754 | 37.54471 | 27.51289 | 24.29611 |
| Rab20 | 19332 | 2.110006 | 3.144715 | 20.72472 | 1.27E-15 | 2.07E-14 | 0.054323 | 3.144715 | 0.72825 | 0.474376 | 0.572884 | 268.597 | 226.4489 | 226.5401 | 1124.682 | 1164.244 | 1053.728 | 1245.025 | 1175.55 | 1183.63 | 572.2956 | 565.7957 | 637.3754 | 201.9071 | 234.2526 | 215.4255 |
| Kdr | 16542 | 2.112009 | 3.52927 | 24.02235 | 6.16E-17 | 1.32E-15 | -0.05568 | 3.52927 | -0.83716 | 0.411778 | 0.510809 | 320.5357 | 353.4379 | 328.9605 | 1451.957 | 1608.033 | 1566.389 | 1506.537 | 1553.549 | 1547.913 | 204.611 | 250.8717 | 201.5121 | 879.3806 | 916.5723 | 920.0126 |
| Gm19412 | 100502849 | 2.112372 | -0.0944 | 6.819508 | 8.83E-07 | 3.69E-06 | 0.341189 | -0.0944 | 1.565791 | 0.132105 | 0.197601 | 20.03348 | 33.74533 | 26.03909 | 105.1954 | 134.8219 | 121.7401 | 186.1852 | 154.1485 | 140.197 | 55.38344 | 38.88892 | 49.25852 | 27.53279 | 30.65722 | 12.95792 |
| Col6a3 | 12835 | 2.112651 | 0.020288 | 5.594402 | 1.41E-05 | 5.12E-05 | 1.169659 | 0.020288 | 4.889594 | 7.46E-05 | 0.000245 | 34.8731 | 10.65642 | 17.35939 | 96.10448 | 113.756 | 67.63339 | 228.823 | 218.4887 | 194.2073 | 27.69172 | 22.11331 | 20.89756 | 75.92375 | 112.4098 | 91.51534 |
| Pcp4 | 18546 | 2.112715 | -2.0088 | 3.945176 | 0.000722 | 0.002122 | 3.147347 | -2.0088 | 9.721114 | 2.71E-09 | 1.68E-08 | 5.935846 | 5.32821 | 3.471878 | 15.58451 | 35.10988 | 24.34802 | 207.5041 | 221.1696 | 235.5769 | 3.076858 | 6.100223 | 8.209754 | 2.502981 | 3.930413 | 6.478962 |
| Thbs4 | 21828 | 2.113607 | -2.09904 | 2.822558 | 0.010098 | 0.024224 | 1.462754 | -2.09904 | 2.781225 | 0.011084 | 0.02361 | 3.709904 | 4.440175 | 1.735939 | 10.38967 | 9.830767 | 36.52203 | 65.378 | 33.51054 | 44.81707 | 5.384501 | 15.25056 | 2.239024 | 35.04173 | 36.94588 | 24.29611 |
| Ass1 | 11898 | 2.113857 | 6.771945 | 29.84162 | 7.20E-16 | 1.25E-14 | -0.91415 | 6.771945 | -14.5324 | 7.14E-11 | 5.73E-10 | 4026.73 | 4274.112 | 3168.957 | 17401.4 | 18278.2 | 17028.73 | 10115.11 | 9955.312 | 9069.136 | 5979.104 | 5975.168 | 5777.428 | 3111.205 | 3000.477 | 3233.812 |
| 1700018A( | 71307 | 2.114089 | -4.13686 | 2.49034 | 0.021069 | 0.046676 | 1.566411 | -4.13686 | 2.325382 | 0.029981 | 0.055881 | 0.741981 | 1.77607 | 0 | 3.896127 | 4.213186 | 4.058003 | 9.948826 | 16.08506 | 16.08818 | 3.076858 | 1.525056 | 0 | 4.171635 | 6.288661 | 3.239481 |
| Csf2 | 12981 | 2.114597 | -2.84466 | 2.72046 | 0.0127 | 0.029764 | 0.785347 | -2.84466 | 1.350639 | 0.190959 | 0.269703 | 5.193865 | 0.888035 | 5.207818 | 14.2858 | 14.04395 | 18.93735 | 44.05909 | 25.46801 | 21.83396 | 3.076858 | 4.575167 | 0 | 6.674615 | 20.43815 | 13.76779 |
| Rbms3 | 207181 | 2.114877 | -1.37766 | 3.679553 | 0.001365 | 0.003844 | 0.842179 | -1.37766 | 2.085069 | 0.049251 | 0.085507 | 5.193865 | 6.216245 | 6.075787 | 29.87031 | 22.47032 | 32.46403 | 41.21656 | 61.6594 | 56.30862 | 19.99958 | 9.150334 | 9.702436 | 65.91183 | 70.74744 | 34.01455 |
| Itgb6 | 16420 | 2.115727 | -1.84898 | 3.366848 | 0.002869 | 0.00764 | 0.977508 | -1.84898 | 2.243759 | 0.035574 | 0.064845 | 5.193865 | 4.440175 | 7.811726 | 27.27289 | 25.27911 | 29.75869 | 45.48035 | 64.34024 | 60.90525 | 8.461359 | 0.762528 | 5.97073 | 34.2074 | 51.09537 | 18.62702 |
| Sema4c | 20353 | 2.116025 | 5.842854 | 54.86743 | 1.84E-24 | 2.62E-22 | -3.51763 | 5.842854 | -62.8412 | 1.03E-25 | 5.12E-23 | 5534.434 | 5915.201 | 5180.043 | 25693.66 | 26287.47 | 24878.27 | 2416.143 | 2369.866 | 2199.484 | 2489.178 | 2514.817 | 2503.975 | 604.887 | 509.3815 | 617.1211 |
| Pml | 18854 | 2.116368 | 7.989663 | 35.46041 | 6.02E-16 | 1.05E-14 | 0.27257 | 7.989663 | 6.274801 | 1.45E-05 | 5.25E-05 | 7177.18 | 7577.602 | 6781.447 | 33530.07 | 33251.87 | 32785.96 | 43356.98 | 43736.62 | 38580.6 | 12535.12 | 12659.49 | 12387.77 | 6425.152 | 6505.62 | 6541.322 |
| Stk40 | 74178 | 2.117445 | 6.446404 | 44.59077 | 1.48E-22 | 1.13E-20 | -0.30832 | 6.446404 | -7.99347 | 7.43E-08 | 3.71E-07 | 3335.945 | 3299.938 | 2992.759 | 14735.15 | 15243.31 | 14664.27 | 12906.47 | 12905.58 | 11747.82 | 4300.678 | 4331.158 | 4254.891 | 1472.587 | 1550.941 | 1481.253 |
| Fbxw22 | 382156 | 2.117461 | -3.50423 | 2.66806 | 0.01427 | 0.033033 | 0.666491 | -3.50423 | 1.035022 | 0.312244 | 0.407136 | 1.483961 | 1.77607 | 1.735939 | 5.194837 | 16.85274 | 8.116006 | 21.31891 | 8.04253 | 19.53565 | 2.307643 | 1.525056 | 0 | 18.35519 | 9.432991 | 5.669092 |
| Sez6 | 20370 | 2.118525 | -1.59119 | 3.567202 | 0.001785 | 0.004926 | 1.184637 | -1.59119 | 2.933384 | 0.007852 | 0.017451 | 8.903769 | 2.664105 | 6.075787 | 22.07806 | 28.08791 | 29.75869 | 42.63783 | 58.97856 | 91.93245 | 3.846072 | 4.575167 | 15.67317 | 45.05365 | 49.52321 | 42.11325 |
| Nebl | 74103 | 2.119407 | -2.68686 | 2.792384 | 0.010809 | 0.02572 | 0.677997 | -2.68686 | 1.159817 | 0.258934 | 0.348224 | 7.419807 | 2.664105 | 0.86797 | 20.77935 | 12.63956 | 10.82134 | 24.16143 | 14.74464 | 36.77298 | 1.538429 | 6.862751 | 0.746341 | 31.70442 | 27.51289 | 15.38754 |
| Snap91 | 20616 | 2.120335 | -2.07371 | 2.81866 | 0.010188 | 0.024405 | 1.27406 | -2.07371 | 2.417997 | 0.024622 | 0.047169 | 3.709904 | 0.888035 | 12.15157 | 27.27289 | 12.63956 | 17.58468 | 54.00791 | 42.8935 | 45.96622 | 5.384501 | 6.862751 | 3.731706 | 16.68654 | 56.59795 | 21.8665 |
| Col6a1 | 12833 | 2.12148 | -1.18658 | 5.181319 | 3.72E-05 | 0.000128 | 0.005392 | -1.18658 | 0.017696 | 0.986045 | 0.9883 | 17.06556 | 11.54445 | 6.943757 | 51.94837 | 51.96262 | 58.16471 | 79.59061 | 48.25518 | 48.26453 | 4.615287 | 8.387806 | 9.702436 | 47.55663 | 38.51805 | 48.59222 |
| Cacng5 | 140723 | 2.122678 | -2.54758 | 2.504095 | 0.02045 | 0.045458 | 0.960553 | -2.54758 | 1.483871 | 0.152462 | 0.223343 | 4.451884 | 0 | 7.811726 | 6.493546 | 18.25714 | 13.52668 | 15.63387 | 25.46801 | 33.32551 | 1.538429 | 3.050111 | 0.746341 | 81.76404 | 82.53868 | 55.88105 |
| Nlrp14 | 76858 | 2.122716 | -3.12008 | 2.754643 | 0.011765 | 0.027764 | -0.47859 | -3.12008 | -0.69525 | 0.494399 | 0.592428 | 3.709904 | 1.77607 | 2.603909 | 9.090964 | 15.44835 | 17.58468 | 2.842522 | 17.42548 | 16.08818 | 4.615287 | 0 | 2.239024 | 24.19548 | 16.50774 | 20.24676 |
| Srms | 20811 | 2.123126 | -1.354 | 3.821019 | 0.000973 | 0.002804 | 0.839798 | -1.354 | 2.206953 | 0.038396 | 0.069111 | 7.419807 | 8.88035 | 6.943757 | 20.77935 | 74.43295 | 33.81669 | 86.69691 | 49.5956 | 76.99342 | 13.84586 | 28.97606 | 12.6878 | 18.35519 | 20.43815 | 15.38754 |
| Noct | 12457 | 2.12315 | 5.458926 | 44.25048 | 1.74E-22 | 1.30E-20 | -0.08356 | 5.458926 | -2.19558 | 0.039308 | 0.070536 | 2191.811 | 2284.026 | 2266.269 | 10249.41 | 10341.97 | 10749.65 | 10892.54 | 9917.78 | 9963.179 | 474.6053 | 472.7673 | 450.7901 | 1130.513 | 1065.928 | 1163.784 |
| Cd6 | 12511 | 2.124578 | -1.23575 | 3.034553 | 0.006226 | 0.015597 | 0.152862 | -1.23575 | 0.295433 | 0.770518 | 0.843772 | 25.96933 | 16.87266 | 0.86797 | 55.84449 | 26.68351 | 41.9327 | 42.63783 | 38.87223 | 57.45778 | 9.999788 | 20.58825 | 22.39024 | 33.37308 | 54.2397 | 32.39481 |
| Efnb2 | 13642 | 2.125843 | 0.998354 | 7.567029 | 1.79E-07 | 8.15E-07 | 0.072298 | 0.998354 | 0.359035 | 0.723097 | 0.80259 | 50.45469 | 28.41712 | 46.00239 | 206.4948 | 176.9538 | 193.4315 | 206.0828 | 289.5311 | 162.0309 | 46.92208 | 62.52728 | 52.99023 | 269.4876 | 245.2578 | 285.8842 |
| Ttc34 | 242800 | 2.126643 | -2.58589 | 2.505289 | 0.020397 | 0.045356 | 1.695708 | -2.58589 | 2.704848 | 0.01315 | 0.02739 | 3.709904 | 0 | 3.471878 | 14.2858 | 4.213186 | 9.468674 | 17.05513 | 32.17012 | 44.81707 | 5.384501 | 3.812639 | 11.19512 | 28.36712 | 32.22939 | 25.10598 |
| Zpld1 | 239852 | 2.126794 | -3.82797 | 2.552586 | 0.018401 | 0.041421 | 1.541202 | -3.82797 | 2.436276 | 0.023675 | 0.045577 | 1.483961 | 0 | 5.207818 | 10.38967 | 8.426372 | 4.058003 | 18.47639 | 24.12759 | 26.43058 | 0 | 0 | 0 | 5.840288 | 21.22423 | 10.52831 |
| Saa2 | 20209 | 2.128696 | -2.87071 | 3.848905 | 0.00091 | 0.002634 | -1.7538 | -2.87071 | -3.11455 | 0.005175 | 0.012052 | 8.161788 | 12.43249 | 1.735939 | 38.96127 | 23.87472 | 24.34802 | 7.106304 | 6.702109 | 11.49156 | 5.384501 | 4.575167 | 3.731706 | 7.508942 | 3.14433 | 3.239481 |
| Nlrp9a | 233001 | 2.129551 | -3.24911 | 2.71542 | 0.012843 | 0.030065 | 0.803298 | -3.24911 | 1.296271 | 0.208724 | 0.290781 | 0.741981 | 2.664105 | 3.471878 | 9.090964 | 9.830767 | 12.17401 | 11.37009 | 24.12759 | 24.13227 | 0.769214 | 3.050111 | 0 | 20.02385 | 25.15464 | 8.908573 |
| Mymk | 66139 | 2.130548 | -3.11806 | 2.81488 | 0.010275 | 0.024583 | 0.294643 | -3.11806 | 0.472987 | 0.641024 | 0.729378 | 5.193865 | 0.888035 | 1.735939 | 18.18193 | 4.213186 | 17.58468 | 14.21261 | 14.74464 | 14.93902 | 5.384501 | 1.525056 | 2.239024 | 17.52087 | 11.79124 | 5.669092 |
| Fam102b | 329739 | 2.130667 | 7.477613 | 52.87957 | 4.01E-24 | 5.17E-22 | 1.034713 | 7.477613 | 38.12209 | 4.03E-21 | 3.73E-19 | 4928.236 | 4744.771 | 4779.041 | 22537.8 | 23777.82 | 21410.03 | 49279.38 | 47824.91 | 46987.82 | 4292.986 | 4286.169 | 4298.926 | 6108.107 | 6218.7 | 6054.59 |
| Cyp4a29 | 230639 | 2.130831 | -3.52083 | 2.845795 | 0.009582 | 0.02309 | 0.58583 | -3.52083 | 0.972958 | 0.341473 | 0.438594 | 3.709904 | 1.77607 | 0.86797 | 5.194837 | 11.23516 | 17.58468 | 21.31891 | 14.74464 | 13.78987 | 2.307643 | 0 | 0 | 16.68654 | 7.860826 | 13.76779 |
| Nrtn | 18188 | 2.132113 | -2.64101 | 2.522093 | 0.019666 | 0.043904 | 1.217522 | -2.64101 | 1.901332 | 0.070845 | 0.116406 | 6.677827 | 0 | 2.603909 | 9.090964 | 14.04395 | 6.763339 | 27.00396 | 20.10633 | 25.28142 | 1.538429 | 12.96297 | 2.985365 | 31.70442 | 33.80155 | 36.44416 |
| Cyp3a13 | 13113 | 2.132274 | -1.72646 | 2.886351 | 0.00874 | 0.021244 | 2.149633 | -1.72646 | 4.518658 | 0.000182 | 0.000565 | 0.741981 | 8.88035 | 13.88751 | 20.77935 | 30.8967 | 18.93735 | 76.74809 | 100.5316 | 151.6885 | 10.769 | 2.287584 | 5.224389 | 25.86413 | 27.51289 | 24.29611 |
| Oas1h | 246729 | 2.132616 | 2.297711 | 11.53865 | 1.22E-10 | 8.62E-10 | 0.961922 | 2.297711 | 8.123372 | 5.72E-08 | 2.91E-07 | 100.1674 | 97.68385 | 100.6845 | 507.7953 | 390.4219 | 511.3084 | 1009.095 | 989.2312 | 861.8667 | 183.8423 | 197.4947 | 148.5219 | 314.5412 | 274.3428 | 255.919 |
| Gabrd | 14403 | 2.137145 | -2.82673 | 2.74663 | 0.011978 | 0.028242 | 0.934951 | -2.82673 | 1.515201 | 0.144392 | 0.213052 | 1.483961 | 0.888035 | 2.603909 | 7.792255 | 9.830767 | 8.116006 | 8.527565 | 24.12759 | 24.13227 | 6.92293 | 19.0632 | 7.463413 | 6.674615 | 18.0799 | 15.38754 |
| Inpp5b | 16330 | 2.137348 | 6.973513 | 48.90719 | 2.10E-23 | 2.11E-21 | -0.11651 | 6.973513 | -3.4135 | 0.00257 | 0.006444 | 4412.559 | 4232.375 | 4402.342 | 20755.97 | 20858.08 | 19732.72 | 20376.62 | 20104.99 | 18489.91 | 5484.499 | 5143.25 | 5335.594 | 2567.224 | 2560.271 | 2597.254 |
| Tmem151l | 210573 | 2.137419 | -1.18728 | 4.046906 | 0.000565 | 0.001687 | 0.896833 | -1.18728 | 2.467855 | 0.022119 | 0.043022 | 6.677827 | 3.55214 | 12.15157 | 45.45482 | 25.27911 | 29.75869 | 52.58665 | 52.27645 | 88.48498 | 13.07665 | 16.01308 | 6.717071 | 72.58644 | 70.74744 | 72.88832 |
| Piwil4 | 330890 | 2.137526 | 0.677448 | 8.311518 | 3.92E-08 | 1.95E-07 | 1.89073 | 0.677448 | 12.38447 | 3.27E-11 | 2.79E-10 | 38.583 | 42.62568 | 22.56721 | 189.6115 | 132.0132 | 158.2621 | 578.4532 | 625.977 | 632.0356 | 47.6913 | 51.85189 | 50.75121 | 50.89394 | 47.95104 | 44.54286 |
| Adamts14 | 237360 | 2.138886 | -0.08607 | 4.819267 | 8.82E-05 | 0.00029 | -0.25618 | -0.08607 | -0.75611 | 0.45785 | 0.556578 | 32.64715 | 7.992315 | 16.49142 | 79.22126 | 54.77142 | 121.7401 | 71.06304 | 54.95729 | 86.18667 | 75.38302 | 70.91509 | 48.51218 | 100.9536 | 92.75775 | 81.7969 |
| Gm5082 | 328231 | 2.141728 | -1.97144 | 4.760791 | 0.000101 | 0.000332 | -0.98258 | -1.97144 | -2.45496 | 0.022742 | 0.044059 | 10.38773 | 5.32821 | 10.41564 | 38.96127 | 36.51428 | 48.69604 | 22.74017 | 13.40422 | 29.87805 | 6.153716 | 3.812639 | 5.97073 | 26.69846 | 16.50774 | 13.76779 |
| Slc34a2 | 20531 | 2.141932 | -2.6136 | 2.471143 | 0.021962 | 0.048471 | 1.475997 | -2.6136 | 2.400363 | 0.025568 | 0.04878 | 5.935846 | 0 | 8.679696 | 11.68838 | 19.66153 | 10.82134 | 56.85043 | 34.85097 | 34.47467 | 1.538429 | 4.575167 | 0 | 29.20144 | 34.58764 | 35.63429 |
| Mir665 | 751555 | 2.142349 | -5.3947 | 2.675412 | 0.014039 | 0.032541 | 0.05231 | -5.3947 | 0.072228 | 0.943093 | 0.952912 | 0.741981 | 0 | 0 | 2.597418 | 2.808791 | 1.352668 | 1.421261 | 4.021265 | 2.298311 | 0 | 0.762528 | 0 | 1.668654 | 1.572165 | 1.619741 |

| | | | | | | | | | | | | | | | | | | | | | | | | | | |
|---|---|---|---|---|---|---|---|---|---|---|---|---|---|---|---|---|---|---|---|---|---|---|---|---|---|---|
| Dspp | 666279 | 2.142582 | -2.6365 | 2.622602 | 0.015779 | 0.036151 | 0.280143 | -2.6365 | 0.437722 | 0.665985 | 0.751549 | 3.709904 | 1.77607 | 7.811726 | 20.77935 | 11.23516 | 29.75869 | 8.527565 | 32.17012 | 44.81707 | 2.307643 | 3.050111 | 0 | 35.04173 | 35.37372 | 15.38754 |
| Slc27a5 | 26459 | 2.142709 | -2.80501 | 2.568569 | 0.017769 | 0.040118 | -0.03999 | -2.80501 | -0.0567 | 0.955309 | 0.963035 | 5.935846 | 2.664105 | 0.86797 | 22.07806 | 11.23516 | 9.468674 | 9.948826 | 9.382952 | 24.13227 | 13.07665 | 9.912862 | 1.492683 | 12.5149 | 22.01031 | 6.478962 |
| Tap2 | 21355 | 2.143229 | 6.88323 | 29.12666 | 1.83E-16 | 3.60E-15 | 0.676431 | 6.88323 | 12.87654 | 1.87E-10 | 1.40E-09 | 2887.047 | 3025.535 | 3000.571 | 14262.42 | 13760.26 | 13955.47 | 24717.15 | 24489.51 | 21047.93 | 3542.233 | 3621.245 | 3610.053 | 5006.796 | 4854.846 | 5193.698 |
| Vmn2r90 | 626942 | 2.143565 | -2.80994 | 2.852755 | 0.009432 | 0.02276 | -0.153 | -2.80994 | -0.24045 | 0.81228 | 0.880544 | 2.967923 | 2.664105 | 3.471878 | 29.87031 | 9.830767 | 12.17401 | 12.79135 | 8.04253 | 26.43058 | 6.153716 | 9.150334 | 0.746341 | 10.01192 | 11.00516 | 20.24676 |
| Bnip5 | 207819 | 2.146282 | 0.355287 | 6.20202 | 3.49E-06 | 1.36E-05 | 0.656727 | 0.355287 | 2.814595 | 0.010281 | 0.022121 | 18.54952 | 53.2821 | 17.35939 | 123.3774 | 110.9472 | 128.5034 | 197.5553 | 164.8719 | 232.1294 | 53.84501 | 62.52728 | 44.78048 | 99.2849 | 68.38919 | 68.0291 |
| 9130017K | 72804 | 2.147402 | -2.47082 | 2.605228 | 0.016395 | 0.037347 | 0.823049 | -2.47082 | 1.301543 | 0.206947 | 0.288649 | 1.483961 | 0.888035 | 9.547666 | 23.37676 | 4.213186 | 20.29002 | 18.47639 | 21.44675 | 34.47467 | 3.846072 | 9.912862 | 2.985365 | 19.18952 | 29.87114 | 38.87377 |
| Zdhhc19 | 245308 | 2.149759 | -3.30234 | 2.544346 | 0.018735 | 0.042087 | 1.944935 | -3.30234 | 3.088673 | 0.005495 | 0.012699 | 2.225942 | 1.77607 | 0 | 5.194837 | 5.617581 | 6.763339 | 18.47639 | 28.14886 | 29.87805 | 3.076858 | 6.100223 | 0.746341 | 13.34923 | 17.29382 | 7.288832 |
| Olfr545 | 258837 | 2.150991 | -4.29552 | 2.634462 | 0.015372 | 0.035331 | 0.105058 | -4.29552 | 0.146158 | 0.885168 | 0.918721 | 1.483961 | 0 | 2.603909 | 5.194837 | 5.617581 | 6.763339 | 2.842522 | 13.40422 | 6.894934 | 1.538429 | 0 | 0 | 6.674615 | 5.502578 | 5.669092 |
| Dcun1d3 | 233805 | 2.153302 | 5.332485 | 40.6278 | 1.05E-21 | 6.53E-20 | -1.14171 | 5.332485 | -23.4422 | 1.02E-16 | 2.83E-15 | 1445.378 | 1514.1 | 1493.776 | 7315.629 | 7175.055 | 6648.362 | 3357.018 | 3493.139 | 3123.405 | 2053.803 | 2005.448 | 2085.277 | 1123.004 | 1154.755 | 1140.297 |
| Nr1h5 | 381463 | 2.153597 | -2.83862 | 2.833641 | 0.009849 | 0.023692 | 0.027904 | -2.83862 | 0.04473 | 0.964738 | 0.971015 | 5.193865 | 0.888035 | 5.207818 | 10.38967 | 11.23516 | 35.16936 | 21.31891 | 6.702109 | 31.0272 | 2.307643 | 1.525056 | 2.985365 | 10.01192 | 24.36856 | 14.57766 |
| Dmtn | 13829 | 2.153804 | -0.46019 | 4.957292 | 6.34E-05 | 0.000212 | -0.28081 | -0.46019 | -0.83792 | 0.411362 | 0.510342 | 22.25942 | 11.54445 | 14.75548 | 59.74062 | 67.41097 | 113.6241 | 72.4843 | 34.85097 | 103.424 | 23.07643 | 25.16342 | 25.3756 | 68.41481 | 84.11084 | 78.55742 |
| Pak5 | 241656 | 2.154623 | -2.43536 | 3.284525 | 0.003481 | 0.009142 | 1.041326 | -2.43536 | 2.205694 | 0.038496 | 0.069258 | 3.709904 | 1.77607 | 6.075787 | 27.27289 | 16.85274 | 12.17401 | 25.5827 | 36.19139 | 58.60694 | 0.769214 | 3.050111 | 2.985365 | 34.2074 | 18.0799 | 24.29611 |
| Btc | 12223 | 2.155364 | -2.96819 | 2.578506 | 0.017386 | 0.039339 | 3.195575 | -2.96819 | 5.854025 | 7.73E-06 | 2.92E-05 | 7.419807 | 3.55214 | 0 | 7.792255 | 11.23516 | 14.87935 | 142.1261 | 84.44657 | 110.3189 | 0 | 0 | 0 | 32.53875 | 21.22423 | 15.38754 |
| Gramd1a | 52857 | 2.156682 | 7.223421 | 63.06828 | 9.55E-26 | 2.17E-23 | -2.11855 | 7.223421 | -56.826 | 8.72E-25 | 2.67E-22 | 7019.88 | 6937.329 | 6767.559 | 33418.38 | 32323.56 | 32963.16 | 8320.061 | 8063.977 | 7333.911 | 7702.914 | 7776.259 | 7692.539 | 4007.272 | 4070.336 | 4234.812 |
| Tfap2e | 332937 | 2.156778 | -1.54814 | 4.166622 | 0.000424 | 0.001286 | -0.12486 | -1.54814 | -0.30558 | 0.762888 | 0.837216 | 2.967923 | 7.992315 | 9.547666 | 40.25998 | 28.08791 | 22.99535 | 27.00396 | 32.17012 | 26.43058 | 11.53822 | 12.20045 | 14.18048 | 34.2074 | 55.81187 | 55.07118 |
| Hormad1 | 67981 | 2.156928 | -2.37423 | 4.127399 | 0.000466 | 0.001405 | -1.11863 | -2.37423 | -2.32239 | 0.030171 | 0.056187 | 4.451884 | 9.768385 | 5.207818 | 25.97418 | 33.70549 | 31.11136 | 11.37009 | 17.42548 | 13.78987 | 2.307643 | 7.625279 | 5.97073 | 24.19548 | 10.21907 | 8.908573 |
| Rasd2 | 75141 | 2.158483 | -1.93861 | 2.703412 | 0.013192 | 0.030783 | 1.36207 | -1.93861 | 2.33332 | 0.029483 | 0.055116 | 1.483961 | 1.77607 | 4.339848 | 10.38967 | 21.06593 | 8.116006 | 19.89765 | 38.87223 | 44.81707 | 16.92272 | 21.35078 | 14.92683 | 59.23721 | 59.74228 | 24.29611 |
| Optn | 71648 | 2.159651 | 4.23224 | 30.80352 | 3.53E-19 | 1.23E-17 | -1.74264 | 4.23224 | -24.4539 | 4.27E-17 | 1.28E-15 | 698.9459 | 721.0844 | 685.696 | 3244.175 | 3384.593 | 3422.249 | 1034.678 | 1020.061 | 1056.074 | 1161.514 | 1362.637 | 1388.941 | 586.5318 | 536.8944 | 567.7191 |
| Gadd45g | 23882 | 2.159763 | 2.910672 | 19.74367 | 3.41E-15 | 5.10E-14 | -0.83074 | 2.910672 | -9.03545 | 9.63E-09 | 5.46E-08 | 315.3418 | 333.9012 | 307.2612 | 1601.308 | 1450.74 | 1508.225 | 967.8786 | 1002.635 | 742.3545 | 209.2263 | 202.0699 | 205.9902 | 216.0907 | 209.8841 | 187.8899 |
| Xcr1 | 23832 | 2.160305 | 1.926547 | 13.70362 | 4.75E-12 | 4.12E-11 | 0.766274 | 1.926547 | 7.369083 | 2.71E-07 | 1.25E-06 | 135.0405 | 188.2634 | 150.1587 | 810.3945 | 681.1317 | 731.7932 | 1294.769 | 1175.55 | 1446.787 | 87.69045 | 44.22662 | 69.40974 | 70.08346 | 57.38403 | 57.50079 |
| Fgf3 | 14174 | 2.160683 | -3.28118 | 2.504134 | 0.020448 | 0.045458 | 1.086288 | -3.28118 | 1.609499 | 0.1222 | 0.184934 | 7.419807 | 0 | 0.86797 | 3.896127 | 9.830767 | 10.82134 | 29.84648 | 12.0638 | 14.93902 | 0.769214 | 2.287584 | 1.492683 | 20.85817 | 24.36856 | 14.57766 |
| Cylc1 | 67407 | 2.160709 | -3.18231 | 3.283309 | 0.003491 | 0.009165 | -0.17291 | -3.18231 | -0.30924 | 0.76014 | 0.834967 | 2.225942 | 1.77607 | 4.339848 | 10.38967 | 16.85274 | 14.87935 | 19.89765 | 13.40422 | 8.044089 | 1.538429 | 1.525056 | 0.746341 | 23.36115 | 7.074744 | 12.14805 |
| Trim30e-ps | 625321 | 2.161724 | -2.83186 | 3.214929 | 0.004097 | 0.010619 | 0.79394 | -2.83186 | 1.536165 | 0.139191 | 0.206339 | 0.741981 | 4.440175 | 4.339848 | 7.792255 | 18.25714 | 16.23201 | 17.05513 | 20.10633 | 37.92213 | 3.076858 | 2.287584 | 3.731706 | 13.34923 | 13.3634 | 9.718443 |
| 9030612E | 74530 | 2.162187 | -3.10975 | 2.820082 | 0.010155 | 0.024345 | 0.561875 | -3.10975 | 0.905997 | 0.375052 | 0.47331 | 0.741981 | 2.664105 | 3.471878 | 3.896127 | 18.25714 | 16.23201 | 9.948826 | 13.40422 | 29.87805 | 1.538429 | 0.762528 | 1.492683 | 30.03577 | 21.22423 | 16.19741 |
| Acoxl | 74121 | 2.162236 | 0.025489 | 5.508985 | 1.72E-05 | 6.18E-05 | 2.616832 | 0.025489 | 11.62482 | 1.07E-10 | 8.32E-10 | 19.2915 | 18.64873 | 12.15157 | 125.9748 | 71.62416 | 55.45938 | 558.5555 | 499.9773 | 489.5403 | 13.07665 | 25.92595 | 15.67317 | 73.42077 | 69.17527 | 59.12053 |
| Endou | 19011 | 2.162257 | -2.57018 | 2.659834 | 0.014533 | 0.033575 | 0.503851 | -2.57018 | 0.801976 | 0.431406 | 0.53038 | 1.483961 | 2.664105 | 7.811726 | 9.090964 | 28.08791 | 17.58468 | 17.05513 | 21.44675 | 36.77298 | 12.30743 | 1.525056 | 7.463413 | 11.68058 | 25.15464 | 4.859222 |
| Ubqlnl | 244179 | 2.162979 | -3.85437 | 2.666844 | 0.014309 | 0.033107 | 0.530494 | -3.85437 | 0.800649 | 0.432158 | 0.531229 | 2.967923 | 0 | 5.207818 | 12.98709 | 14.04395 | 4.058003 | 14.21261 | 12.0638 | 16.08818 | 0 | 0 | 0 | 18.35519 | 9.432991 | 6.478962 |
| Mlkl | 74568 | 2.163998 | 6.160595 | 47.15683 | 4.53E-23 | 4.11E-21 | 0.035551 | 6.160595 | 0.998471 | 0.329238 | 0.425517 | 2135.421 | 2127.732 | 2135.205 | 10365 | 10443.08 | 9825.778 | 11145.53 | 10861.44 | 10618.2 | 2882.247 | 2745.1 | 2676.38 | 2233.493 | 2068.183 | 2247.39 |
| Urad | 231903 | 2.164083 | -4.18445 | 2.56993 | 0.017716 | 0.040009 | 0.661304 | -4.18445 | 0.929851 | 0.362847 | 0.460676 | 0.741981 | 0 | 2.603909 | 3.896127 | 4.213186 | 6.763339 | 11.37009 | 9.382952 | 5.745778 | 1.538429 | 0 | 0 | 14.18356 | 11.79124 | 5.669092 |
| Nhs | 195727 | 2.164982 | 2.875694 | 30.74793 | 3.66E-19 | 1.27E-17 | -3.14307 | 2.875694 | -30.9487 | 3.20E-19 | 1.67E-17 | 529.7743 | 617.1843 | 533.8013 | 2618.198 | 2822.834 | 2575.479 | 314.0986 | 312.3183 | 312.5703 | 502.297 | 526.1442 | 534.3803 | 59.23721 | 75.46393 | 64.78962 |
| Unc5c | 22253 | 2.165295 | -1.1351 | 3.392043 | 0.002703 | 0.007229 | 0.770953 | -1.1351 | 1.791404 | 0.087421 | 0.139155 | 9.64575 | 19.53677 | 9.547666 | 42.8574 | 60.389 | 79.8074 | 75.32683 | 101.8721 | 140.197 | 6.153716 | 0 | 11.19512 | 50.05962 | 110.0516 | 43.73299 |
| Olfr1380 | 404336 | 2.165529 | -4.79653 | 2.672805 | 0.014121 | 0.032714 | 1.178706 | -4.79653 | 1.747149 | 0.094987 | 0.14923 | 1.483961 | 0 | 0 | 1.298709 | 2.808791 | 5.410671 | 11.37009 | 5.361687 | 6.894934 | 0 | 0 | 0 | 5.840288 | 6.288661 | 4.859222 |
| Gm5215 | 383032 | 2.165529 | -5.18337 | 2.581326 | 0.017279 | 0.039123 | 0.125412 | -5.18337 | 0.167302 | 0.868707 | 0.918721 | 1.483961 | 0 | 0 | 1.298709 | 2.808791 | 5.410671 | 5.685043 | 2.680843 | 2.298311 | 0.769214 | 0 | 0 | 1.668654 | 2.358248 | 1.619741 |
| Fcrl6 | 677296 | 2.165586 | -2.35652 | 2.923344 | 0.008034 | 0.019672 | 0.68223 | -2.35652 | 1.23244 | 0.231189 | 0.316611 | 2.967923 | 5.32821 | 2.603909 | 11.68838 | 28.08791 | 17.58468 | 15.63387 | 41.55307 | 39.07129 | 22.30722 | 12.96297 | 10.44878 | 12.5149 | 11.00516 | 1.619741 |
| Gm9979 | 791406 | 2.16658 | -2.91224 | 3.119988 | 0.00511 | 0.013004 | 0.738585 | -2.91224 | 1.392473 | 0.178118 | 0.254298 | 0.741981 | 3.55214 | 6.943757 | 18.18193 | 16.85274 | 9.468674 | 27.00396 | 18.7659 | 31.0272 | 1.538429 | 1.525056 | 0 | 19.18952 | 22.01031 | 27.53559 |
| H2-Q8 | 15019 | 2.167047 | 2.78823 | 11.72313 | 9.12E-11 | 6.57E-10 | 2.448229 | 2.78823 | 23.11306 | 1.36E-16 | 3.66E-15 | 81.61788 | 91.4676 | 85.06102 | 425.9766 | 404.4658 | 405.8003 | 2399.088 | 2430.185 | 2209.826 | 197.6881 | 276.7976 | 261.2194 | 620.7392 | 551.83 | 600.9237 |
| Cldn1 | 12737 | 2.171217 | 0.760682 | 11.17639 | 2.21E-10 | 1.50E-09 | 0.359785 | 0.760682 | 2.714972 | 0.012856 | 0.026866 | 59.35846 | 49.72996 | 57.28599 | 287.0147 | 280.8791 | 239.4222 | 355.3152 | 352.5309 | 364.2823 | 62.30637 | 54.90201 | 55.22925 | 29.20144 | 48.73712 | 27.53559 |
| Gm15473 | 100504308 | 2.173299 | -1.70824 | 3.527926 | 0.001959 | 0.00537 | 1.494167 | -1.70824 | 3.592134 | 0.001682 | 0.004382 | 4.451884 | 7.992315 | 1.735939 | 29.87031 | 15.44835 | 18.93735 | 61.11422 | 45.57434 | 81.59005 | 17.69193 | 10.67539 | 8.209754 | 23.36115 | 36.1598 | 28.34546 |
| Nampt | 59027 | 2.173494 | 8.661119 | 69.27824 | 1.30E-26 | 4.17E-24 | 0.852107 | 8.661119 | 41.49129 | 6.77E-22 | 7.88E-20 | 9532.227 | 9950.432 | 9472.152 | 47184.7 | 48121.6 | 44126.73 | 87475.76 | 86559.07 | 87057.73 | 14818.92 | 14261.56 | 14227.5 | 14705.01 | 14504.8 | 14996.37 |
| Gm10826 | 100038717 | 2.174024 | -1.45705 | 3.731166 | 0.001207 | 0.003429 | 1.655553 | -1.45705 | 4.412417 | 0.000234 | 0.000717 | 4.451884 | 7.10428 | 5.207818 | 22.07806 | 33.70549 | 28.40602 | 92.38196 | 121.9784 | 71.24765 | 16.92272 | 6.862751 | 2.985365 | 43.385 | 45.59279 | 35.63429 |
| Ighv6-6 | 238427 | 2.174651 | -4.01516 | 2.468006 | 0.022111 | 0.048739 | 1.902752 | -4.01516 | 2.753279 | 0.011801 | 0.024964 | 1.483961 | 0 | 0 | 5.194837 | 2.808791 | 1.352668 | 19.89765 | 10.72337 | 10.3424 | 5.384501 | 1.525056 | 2.985365 | 9.177596 | 6.288661 | 3.239481 |
| Pttg1 | 30939 | 2.17473 | 4.975185 | 34.17769 | 4.00E-20 | 1.75E-18 | -1.34375 | 4.975185 | -22.3774 | 2.65E-16 | 6.64E-15 | 1386.762 | 1237.921 | 1327.126 | 6342.896 | 6437.748 | 6304.784 | 2542.636 | 2908.715 | 2400.586 | 2030.726 | 1923.095 | 1813.609 | 533.9692 | 562.0491 | 583.1066 |
| Sdr16c5 | 242285 | 2.174797 | -3.63566 | 2.664732 | 0.014376 | 0.033248 | 0.957769 | -3.63566 | 1.426575 | 0.168168 | 0.242355 | 2.225942 | 0.888035 | 0 | 5.194837 | 4.213186 | 5.410671 | 5.685043 | 17.42548 | 11.49156 | 3.846072 | 6.100223 | 1.492683 | 12.5149 | 7.860826 | 8.098703 |
| Gm20319 | 100504628 | 2.17521 | -3.67751 | 2.498178 | 0.020714 | 0.045971 | -0.39709 | -3.67751 | -0.50077 | 0.621655 | 0.712115 | 4.451884 | 0.888035 | 0 | 2.597418 | 8.426372 | 10.82134 | 5.685043 | 2.680843 | 6.894934 | 1.538429 | 2.287584 | 5.97073 | 19.18952 | 8.646909 | 12.14805 |
| Slc35f1 | 215085 | 2.175307 | -2.01193 | 3.415113 | 0.00256 | 0.006876 | 0.911279 | -2.01193 | 2.007975 | 0.05747 | 0.097586 | 4.451884 | 5.32821 | 2.603909 | 14.2858 | 23.87472 | 25.70069 | 31.26774 | 54.95729 | 39.07129 | 1.538429 | 7.625279 | 3.731706 | 56.73423 | 55.81187 | 30.77507 |
| Vnn3 | 26464 | 2.17641 | -0.53734 | 8.158302 | 5.33E-08 | 2.61E-07 | -3.35353 | -0.53734 | -8.33335 | 3.76E-08 | 1.96E-07 | 43.03488 | 79.92315 | 37.32269 | 196.1051 | 317.3933 | 231.3062 | 36.95278 | 20.10633 | 19.53565 | 35.38386 | 59.47717 | 38.0634 | 5.840288 | 9.432991 | 10.52831 |
| H1f8 | 171506 | 2.176533 | -2.49953 | 2.59827 | 0.016648 | 0.037841 | 1.133433 | -2.49953 | 1.724655 | 0.099044 | 0.154615 | 0.741981 | 1.77607 | 1.735939 | 15.58451 | 9.830767 | 2.705335 | 9.948826 | 18.7659 | 29.87805 | 6.92293 | 18.30067 | 7.463413 | 63.40885 | 41.66238 | 40.49351 |
| Robo4 | 74144 | 2.176792 | -2.15139 | 2.728104 | 0.012485 | 0.029305 | 2.008103 | -2.15139 | 3.608815 | 0.001616 | 0.00423 | 0.741981 | 4.440175 | 2.603909 | 10.38967 | 5.617581 | 24.34802 | 52.58665 | 38.87223 | 58.60694 | 11.53822 | 2.287584 | 1.492683 | 66.74615 | 65.24486 | 90.70547 |
| Optc | 269120 | 2.177291 | -3.12821 | 2.6161 | 0.016007 | 0.036597 | 1.026397 | -3.12821 | 1.556609 | 0.134268 | 0.200328 | 0 | 4.440175 | 2.603909 | 12.98709 | 2.808791 | 10.82134 | 17.05513 | 17.42548 | 16.08818 | 3.846072 | 5.337695 | 2.985365 | 15.85221 | 14.93557 | 9.718443 |
| Cysltr2 | 70086 | 2.17745 | 1.016368 | 13.262 | 8.90E-12 | 7.43E-11 | -4.02238 | 1.016368 | -13.1643 | 1.02E-11 | 9.61E-11 | 136.5245 | 85.25136 | 130.1954 | 567.5359 | 543.501 | 576.2365 | 39.7953 | 20.10633 | 51.712 | 296.1476 | 305.7737 | 260.4731 | 40.04769 | 37.73197 | 38.87377 |
| Gm12830 | 433746 | 2.179318 | -2.99874 | 2.689542 | 0.013605 | 0.031664 | 0.788282 | -2.99874 | 1.266261 | 0.219066 | 0.302704 | 1.483961 | 2.664105 | 3.471878 | 10.38967 | 25.27911 | 8.116006 | 21.31891 | 16.08506 | 35.62382 | 1.538429 | 8.387806 | 0 | 9.177596 | 17.29382 | 15.38754 |
| Slfn2 | 20556 | 2.181102 | 8.34731 | 64.30668 | 6.32E-26 | 1.51E-23 | -0.53419 | 8.34731 | -19.7839 | 3.27E-15 | 6.23E-14 | 12570.64 | 12664.27 | 12793.87 | 61784.79 | 61876.25 | 60420.96 | 45750.39 | 45145.4 | 41651.14 | 21011.86 | 20239.01 | 21302.07 | 4060.669 | 3814.073 | 3846.884 |
| Kif1a | 16560 | 2.181433 | 1.517334 | 6.54047 | 1.63E-06 | 6.59E-06 | -0.46679 | 1.517334 | -1.79603 | 0.086661 | 0.138145 | 51.93865 | 16.87266 | 45.13442 | 161.0399 | 164.3142 | 189.3735 | 85.27565 | 128.6805 | 178.1191 | 209.9955 | 228.7584 | 230.6194 | 810.9658 | 905.5672 | 926.4916 |
| Plek | 56193 | 2.181894 | 9.780334 | 97.62873 | 8.79E-30 | 1.65E-26 | -1.01396 | 9.780334 | -53.4635 | 3.18E-24 | 8.62E-22 | 31881.43 | 32701.89 | 31684.36 | 156210 | 160685.3 | 149523.9 | 80018.41 | 78101.01 | 81233.81 | 40819.13 | 40282.82 | 42204.11 | 24463.3 | 23820.66 | 25381.33 |
| Samd9l | 209086 | 2.1831 | 8.341158 | 93.70934 | 2.10E-29 | 3.25E-26 | -0.74408 | 8.341158 | -38.6413 | 3.03E-21 | 2.90E-19 | 13538.92 | 13382.69 | 13234.8 | 65310.79 | 66868.88 | 62725.91 | 39897.63 | 40432.48 | 40352.6 | 16055.04 | 16490.43 | 16073.95 | 4759.835 | 4797.462 | 4931.3 |
| Tnfsf18 | 240873 | 2.183784 | -0.38474 | 4.054169 | 0.000556 | 0.001659 | 4.948066 | -0.38474 | 16.12532 | 1.97E-13 | 2.56E-12 | 3.709904 | 11.54445 | 3.471878 | 18.18193 | 36.51428 | 29.75869 | 911.0282 | 845.8061 | 889.4464 | 19.99958 | 30.50111 | 17.91219 | 58.40288 | 70.74744 | 59.9304 |
| Pitx3 | 18742 | 2.18388 | -2.67399 | 2.492515 | 0.02097 | 0.046476 | 2.205457 | -2.67399 | 3.553489 | 0.001844 | 0.004765 | 0 | 2.664105 | 6.075787 | 15.58451 | 7.021976 | 5.410671 | 49.74413 | 38.87223 | 42.51876 | 6.92293 | 0.762528 | 0.746341 | 42.55067 | 42.44846 | 43.73299 |
| Bfar | 67118 | 2.184921 | 5.58747 | 47.64549 | 3.65E-23 | 3.39E-21 | -2.23164 | 5.58747 | -44.1773 | 1.80E-22 | 2.91E-20 | 2100.547 | 2169.469 | 2064.9 | 10150.71 | 10594.76 | 10016.5 | 2220.009 | 2454.312 | 2143.175 | 2449.948 | 2518.629 | 2518.155 | 1571.038 | 1549.369 | 1680.481 |
| Pcdha12 | 192164 | 2.184998 | -2.5514 | 2.951052 | 0.007541 | 0.01857 | 0.65445 | -2.5514 | 1.186242 | 0.248564 | 0.336377 | 1.483961 | 7.992315 | 4.339848 | 20.77935 | 14.04395 | 24.34802 | 21.31891 | 26.80843 | 51.712 | 0 | 6.862751 | 0 | 45.05365 | 41.66238 | 35.63429 |
| Susd2 | 71733 | 2.185117 | 0.431572 | 7.41276 | 2.47E-07 | 1.11E-06 | 1.598421 | 0.431572 | 9.044361 | 9.47E-09 | 5.38E-08 | 32.64715 | 21.31284 | 36.45472 | 171.4296 | 136.2263 | 129.8561 | 474.7011 | 502.6582 | 410.2485 | 28.46093 | 16.77561 | 36.57072 | 60.07154 | 71.53352 | 61.55014 |
| A530041M | 100503209 | 2.185789 | -1.1871 | 4.990061 | 5.87E-05 | 0.000197 | -0.95138 | -1.1871 | -2.48999 | 0.021085 | 0.041293 | 8.903769 | 14.20856 | 26.03909 | 55.84449 | 101.1165 | 67.63339 | 28.42522 | 41.55307 | 45.96622 | 15.38429 | 19.82572 | 17.16585 | 7.508942 | 16.50774 | 31.58494 |
| Rab38 | 72433 | 2.186037 | 1.168252 | 7.926783 | 8.51E-08 | 4.05E-07 | 1.193507 | 1.168252 | 7.003631 | 5.92E-07 | 2.60E-06 | 52.68063 | 43.51371 | 33.85081 | 257.1444 | 153.0791 | 234.0115 | 551.4492 | 466.4668 | 492.9877 | 119.2282 | 117.4293 | 158.2243 | 90.94163 | 62.88661 | 68.0291 |
| Ppp2r2b | 72930 | 2.186101 | -2.96879 | 2.888209 | 0.008703 | 0.021159 | 1.565524 | -2.96879 | 2.777448 | 0.011178 | 0.023786 | 0.741981 | 4.440175 | 0.86797 | 10.38967 | 4.213186 | 13.52668 | 22.74017 | 21.44675 | 40.22045 | 2.307643 | 3.812639 | 1.492683 | 26.69846 | 17.29382 | 14.57766 |
| Spesp1 | 66712 | 2.186225 | -2.81836 | 2.876764 | 0.008932 | 0.021666 | 1.435974 | -2.81836 | 2.526301 | 0.019487 | 0.038602 | 2.225942 | 1.77607 | 0.86797 | 11.68838 | 4.213186 | 13.52668 | 32.689 | 14.74464 | 35.62382 | 1.538429 | 2.287584 | 3.731706 | 21.6925 | 40.8763 | 35.63429 |
| Nrg3 | 18183 | 2.186642 | -2.3885 | 2.956708 | 0.007444 | 0.018348 | 0.286325 | -2.3885 | 0.514856 | 0.611945 | 0.703848 | 6.677827 | 7.992315 | 2.603909 | 28.5716 | 33.70549 | 20.29002 | 15.63387 | 29.48928 | 73.54596 | 6.153716 | 4.575167 | 0 | 16.68654 | 16.50774 | 14.57766 |
| Serpinb9d | 20726 | 2.186787 | -4.21318 | 2.475049 | 0.021778 | 0.048096 | 0.64857 | -4.21318 | 0.870711 | 0.3936 | 0.492511 | 0 | 1.77607 | 2.603909 | 11.68838 | 5.617581 | 2.705335 | 2.842522 | 14.74464 | 18.38649 | 0 | 0 | 0 | 10.84625 | 9.432991 | 9.718443 |
| Cd8a | 12525 | 2.187046 | -0.60613 | 4.978828 | 6.03E-05 | 0.000202 | 0.258361 | -0.60613 | 0.830657 | 0.415363 | 0.514577 | 16.32358 | 32.85729 | 5.207818 | 84.4161 | 66.00658 | 62.22272 | 92.38196 | 77.74446 | 93.0816 | 19.99958 | 28.97606 | 29.85365 | 38.37904 | 49.52321 | 29.9652 |
| Kctd14 | 233529 | 2.187052 | 0.553161 | 8.656638 | 1.99E-08 | 1.03E-07 | -0.13773 | 0.553161 | -0.74579 | 0.463927 | 0.562604 | 51.19667 | 26.64105 | 65.96569 | 241.5599 | 203.6373 | 220.4848 | 220.2954 | 231.893 | 183.8649 | 66.92166 | 45.75167 | 59.7073 | 44.21933 | 37.73197 | 45.35273 |
| Lix1 | 66643 | 2.187336 | -0.25415 | 7.200811 | 3.87E-07 | 1.69E-06 | -2.95425 | -0.25415 | -7.17179 | 4.12E-07 | 1.85E-06 | 33.38913 | 26.64105 | 44.26645 | 154.5464 | 157.2923 | 192.0788 | 21.31891 | 14.74464 | 31.0272 | 33.84544 | 15.25056 | 52.99023 | 75.92375 | 86.46909 | 80.17716 |
| Marveld2 | 218518 | 2.187443 | -0.83657 | 4.372441 | 0.000258 | 0.000805 | -0.13942 | -0.83657 | -0.36661 | 0.717521 | 0.797512 | 11.87169 | 8.88035 | 10.41564 | 54.54578 | 63.19779 | 40.58003 | 51.16539 | 29.48928 | 72.3968 | 24.61486 | 35.83881 | 41.04877 | 27.53279 | 56.59795 | 59.9304 |
| Fam71a | 619288 | 2.188166 | -1.80488 | 4.670765 | 0.000126 | 0.000407 | -1.64637 | -1.80488 | -3.54954 | 0.001861 | 0.004806 | 11.87169 | 16.87266 | 13.88751 | 64.93546 | 61.79339 | 82.51273 | 22.74017 | 28.14886 | 18.38649 | 9.230573 | 6.100223 | 0 | 20.02385 | 22.01031 | 21.8665 |
| Nectin4 | 71740 | 2.18873 | 5.22739 | 33.82401 | 4.98E-20 | 2.10E-18 | 1.218722 | 5.22739 | 27.60991 | 3.45E-18 | 1.33E-16 | 895.5708 | 904.0196 | 854.0821 | 4500.027 | 4470.19 | 3970.08 | 10650.93 | 10470.03 | 10147.04 | 1083.054 | 1094.99 | 1043.385 | 1292.372 | 1297.822 | 1371.11 |
| Tmem266 | 244886 | 2.190194 | -1.80635 | 3.129748 | 0.004996 | 0.01274 | 1.75999 | -1.80635 | 3.734832 | 0.001196 | 0.003224 | 13.35565 | 0.888035 | 1.735939 | 18.18193 | 18.25714 | 13.52668 | 56.85043 | 65.68067 | 57.45778 | 4.615287 | 10.67539 | 12.6878 | 54.23125 | 46.37887 | 37.25403 |
| Dmrtc1a | 70887 | 2.190241 | -3.80918 | 2.87229 | 0.009023 | 0.021865 | 3.437493 | -3.80918 | 6.463563 | 1.94E-06 | 7.93E-06 | 0.741981 | 0.888035 | 0 | 2.597418 | 4.213186 | 4.058003 | 36.95278 | 54.95729 | 49.41369 | 0 | 0 | 0.746341 | 14.18356 | 14.93557 | 14.57766 |
| Dnah17 | 69926 | 2.190601 | 3.514304 | 21.0063 | 9.66E-16 | 1.62E-14 | -0.97844 | 3.514304 | -10.8644 | 3.70E-10 | 2.63E-09 | 522.3544 | 425.3688 | 362.8113 | 2168.844 | 2074.292 | 2135.862 | 1138.43 | 1239.89 | 1011.257 | 372.2998 | 318.7366 | 292.5658 | 441.3589 | 432.3454 | 442.999 |
| Hrh2 | 15466 | 2.190682 | 2.612096 | 12.28366 | 3.81E-11 | 2.88E-10 | 2.045623 | 2.612096 | 19.44254 | 4.65E-15 | 8.54E-14 | 131.3306 | 129.6531 | 176.1978 | 711.6926 | 634.7867 | 762.9046 | 3091.242 | 3056.162 | 2899.32 | 197.6881 | 209.6952 | 126.1317 | 111.7998 | 158.0026 | 102.8535 |
| Triobp | 110253 | 2.190864 | 6.493174 | 46.15813 | 7.13E-23 | 5.91E-21 | -0.37877 | 6.493174 | -9.81573 | 2.28E-09 | 1.43E-08 | 2594.707 | 2790.206 | 2687.234 | 13218.26 | 13195.7 | 12863.87 | 11017.61 | 10428.48 | 10011.44 | 4492.982 | 4407.411 | 4319.077 | 2596.425 | 2889.64 | 2874.23 |
| Gm15987 | 100504591 | 2.191182 | 2.110804 | 27.87503 | 2.83E-18 | 8.11E-17 | -5.96279 | 2.110804 | -23.1326 | 1.34E-16 | 3.61E-15 | 667.0407 | 688.2271 | 614.5225 | 3103.915 | 3444.982 | 3073.261 | 44.05909 | 58.97856 | 55.15947 | 565.3726 | 548.2575 | 445.5657 | 13.34923 | 23.58248 | 23.48624 |
| Kalrn | 545156 | 2.191318 | 2.752026 | 18.0803 | 2.01E-14 | 2.63E-13 | 0.446665 | 2.752026 | 5.36047 | 2.44E-05 | 8.58E-05 | 236.6919 | 222.8968 | 252.5792 | 1133.773 | 1151.604 | 1190.348 | 1598.918 | 1772.038 | 1560.553 | 105.3824 | 134.9674 | 88.06827 | 256.1384 | 181.5851 | 217.0452 |
| Slc26a7 | 208890 | 2.19245 | -2.82999 | 3.441958 | 0.002402 | 0.006491 | 0.714538 | -2.82999 | 1.478114 | 0.153984 | 0.225196 | 2.225942 | 1.77607 | 5.207818 | 14.2858 | 11.23516 | 20.29002 | 28.42522 | 17.42548 | 33.32551 | 2.307643 | 4.575167 | 0.746341 | 16.68654 | 11.79124 | 10.52831 |
| Ifit1bl2 | 112419 | 2.192838 | 2.49383 | 16.00632 | 2.29E-13 | 2.48E-12 | 0.626917 | 2.49383 | 6.90563 | 7.32E-07 | 3.17E-06 | 174.3655 | 170.5027 | 141.479 | 837.6674 | 706.4108 | 835.9487 | 1236.497 | 1270.72 | 1297.397 | 142.3047 | 142.5927 | 135.0878 | 172.7057 | 236.6109 | 166.0234 |
| Tlk2 | 24086 | 2.194856 | 6.171231 | 53.84536 | 2.73E-24 | 3.67E-22 | -1.84115 | 6.171231 | -44.0516 | 1.91E-22 | 3.05E-20 | 2789.848 | 3007.774 | 2973.664 | 14262.42 | 14623.97 | 13906.78 | 4127.342 | 4245.116 | 4031.238 | 3370.698 | 3339.109 | 3424.96 | 2613.946 | 2642.81 | 2392.357 |
| Pramel6 | 347711 | 2.195193 | -4.27703 | 2.691301 | 0.013552 | 0.031549 | -0.37973 | -4.27703 | -0.5094 | 0.615696 | 0.706982 | 1.483961 | 1.77607 | 0 | 3.896127 | 5.617581 | 6.763339 | 2.842522 | 2.680843 | 8.044089 | 0 | 1.525056 | 1.492683 | 5.840288 | 10.21907 | 5.669092 |
| Olfr959 | 258501 | 2.195706 | -4.83484 | 2.46802 | 0.022111 | 0.048739 | -1.26736 | -4.83484 | -1.46482 | 0.157546 | 0.229728 | 0.741981 | 3.55214 | 0 | 6.493546 | 2.808791 | 8.116006 | 4.263783 | 0 | 4.596622 | 0 | 0.762528 | 0 | 3.337308 | 4.716496 | 2.429611 |
| Isl2 | 104360 | 2.195712 | -1.71721 | 3.93911 | 0.000733 | 0.002151 | 0.06253 | -1.71721 | 0.148131 | 0.88363 | 0.918721 | 7.419807 | 4.440175 | 6.943757 | 40.25998 | 33.70549 | 22.99535 | 18.47639 | 58.97856 | 35.62382 | 2.307643 | 0.762528 | 1.492683 | 156.8535 | 128.9175 | 172.5024 |
| Cplx3 | 235415 | 2.195828 | -2.64911 | 4.009145 | 0.000619 | 0.001838 | 1.058817 | -2.64911 | 2.692612 | 0.013513 | 0.028048 | 5.935846 | 1.77607 | 2.603909 | 16.88322 | 16.85274 | 16.23201 | 39.7953 | 30.8297 | 40.22045 | 1.538429 | 1.525056 | 0.746341 | 22.52683 | 20.43815 | 21.05663 |
| Trim21 | 20821 | 2.198464 | 6.534335 | 51.73718 | 6.37E-24 | 7.69E-22 | 0.585309 | 6.534335 | 19.21858 | 5.88E-15 | 1.05E-13 | 2615.482 | 2655.225 | 2770.559 | 13355.93 | 13229.4 | 12784.06 | 20663.71 | 20661.26 | 20051.62 | 3165.317 | 2878.543 | 2993.575 | 2462.933 | 2335.451 | 2433.66 |
| Cby2 | 67926 | 2.198734 | -3.74768 | 2.77119 | 0.011336 | 0.026849 | 0.534332 | -3.74768 | 0.81103 | 0.426301 | 0.525207 | 1.483961 | 0.888035 | 1.735939 | 5.194837 | 7.021976 | 13.52668 | 7.106304 | 8.04253 | 27.57973 | 0 | 1.525056 | 0 | 21.6925 | 10.21907 | 12.14805 |
| Cfap20dc | 74430 | 2.198907 | -2.88269 | 2.562309 | 0.018014 | 0.040616 | 1.010348 | -2.88269 | 1.534879 | 0.139506 | 0.206677 | 0 | 3.55214 | 8.679696 | 5.194837 | 16.85274 | 13.52668 | 17.05513 | 26.80843 | 25.28142 | 3.846072 | 0.762528 | 2.985365 | 21.6925 | 22.7964 | 20.24676 |
| Ace2 | 70008 | 2.200645 | -2.78901 | 3.33017 | 0.003127 | 0.008277 | 0.268949 | -2.78901 | 0.500385 | 0.621925 | 0.712392 | 5.935846 | 1.77607 | 0.86797 | 10.38967 | 12.63956 | 14.87935 | 17.05513 | 10.72337 | 20.6848 | 4.615287 | 1.525056 | 3.731706 | 27.53279 | 19.65207 | 28.34546 |
| Zscan5b | 170734 | 2.20081 | -3.81035 | 2.670304 | 0.014199 | 0.032886 | 1.516206 | -3.81035 | 2.34339 | 0.028862 | 0.05412 | 1.483961 | 0.888035 | 0 | 2.597418 | 2.808791 | 10.82134 | 22.74017 | 10.72337 | 12.64071 | 3.076858 | 1.525056 | 1.492683 | 10.01192 | 4.716496 | 8.098703 |
| Hbegf | 15200 | 2.201737 | 1.935873 | 6.048687 | 4.94E-06 | 1.89E-05 | 2.785891 | 1.935873 | 13.65409 | 5.09E-12 | 5.02E-11 | 23.0014 | 15.98463 | 22.56721 | 128.5722 | 122.1824 | 66.28072 | 790.221 | 719.8065 | 696.3883 | 456.1442 | 474.2923 | 509.7511 | 796.7822 | 805.7347 | 924.062 |
| Otop3 | 69602 | 2.202151 | -2.7797 | 2.574129 | 0.017554 | 0.039691 | 0.68634 | -2.7797 | 1.041492 | 0.309302 | 0.403993 | 5.935846 | 1.77607 | 0.86797 | 6.493546 | 22.47032 | 13.52668 | 36.95278 | 16.08506 | 17.23733 | 3.076858 | 6.100223 | 0 | 27.53279 | 41.66238 | 25.91585 |
| Rpl5-ps2 | 668936 | 2.202994 | -1.18206 | 4.489317 | 0.000195 | 0.000616 | -0.33606 | -1.18206 | -0.88325 | 0.386942 | 0.485728 | 22.25942 | 7.10428 | 7.811726 | 66.23417 | 49.15383 | 54.10671 | 24.16143 | 58.97856 | 63.20356 | 14.61507 | 24.40089 | 29.85365 | 14.18356 | 24.36856 | 20.24676 |
| Aim2 | 383619 | 2.203974 | 6.029109 | 50.06137 | 1.28E-23 | 1.35E-21 | -0.14008 | 6.029109 | -4.03373 | 0.000584 | 0.00167 | 1993.702 | 2164.141 | 2140.413 | 10428.63 | 10461.34 | 10017.86 | 9778.275 | 9848.079 | 9509.263 | 2264.567 | 2184.642 | 2219.619 | 1878.904 | 1915.683 | 2033.584 |
| Soga3 | 67412 | 2.205055 | -1.79123 | 3.629901 | 0.001537 | 0.004288 | 0.357314 | -1.79123 | 0.801589 | 0.431625 | 0.530624 | 3.709904 | 5.32821 | 9.547666 | 20.77935 | 33.70549 | 36.52203 | 34.11026 | 36.19139 | 47.11538 | 7.692145 | 3.812639 | 2.985365 | 55.06558 | 60.52836 | 23.48624 |
| 4930597O | 75373 | 2.205142 | 0.759969 | 6.259417 | 3.06E-06 | 1.20E-05 | 0.860536 | 0.759969 | 3.865222 | 0.000875 | 0.002427 | 63.81034 | 13.32052 | 56.41802 | 210.3909 | 155.8879 | 198.8422 | 335.4176 | 419.552 | 319.4653 | 46.15287 | 57.95212 | 59.7073 | 80.09538 | 79.39434 | 76.12781 |
| Mmp3 | 17392 | 2.20578 | -1.42608 | 3.077117 | 0.005644 | 0.014263 | 5.93548 | -1.42608 | 13.92762 | 3.47E-12 | 3.54E-11 | 5.935846 | 0.888035 | 3.471878 | 10.38967 | 23.87472 | 14.87935 | 1081.58 | 1020.061 | 968.7382 | 0 | 0.762528 | 4.478048 | 78.42673 | 74.67785 | 85.84625 |
| Gm20559 | 330256 | 2.206573 | 7.334863 | 63.23157 | 9.04E-26 | 2.07E-23 | -1.01112 | 7.334863 | -32.9523 | 8.60E-20 | 5.39E-18 | 6969.425 | 7042.117 | 6733.708 | 34028.78 | 34251.8 | 33977.66 | 16931.48 | 17883.91 | 17786.63 | 11301.3 | 11086.39 | 10306.23 | 1883.91 | 1759.253 | 1940.449 |
| Arel1 | 68497 | 2.207073 | 7.450048 | 62.52517 | 1.15E-25 | 2.50E-23 | -0.14063 | 7.450048 | -5.1825 | 3.71E-05 | 0.000127 | 5070.696 | 5155.043 | 5320.654 | 25963.79 | 25944.8 | 24678.07 | 24731.36 | 24205.34 | 23318.67 | 7426.766 | 7542.163 | 7529.837 | 5171.993 | 5119.756 | 5380.778 |
| Trafd1 | 231712 | 2.209758 | 8.007351 | 54.13968 | 2.44E-24 | 3.34E-22 | -0.21435 | 8.007351 | -6.91122 | 7.23E-07 | 3.13E-06 | 8930.48 | 9026.876 | 8557.312 | 43878.19 | 44086.78 | 43018.89 | 40903.89 | 39900.33 | 36935.01 | 13567.4 | 14212.76 | 13562.51 | 4120.741 | 4314.021 | 3960.266 |
| 5031434C | 100503613 | 2.211738 | -3.88624 | 2.579441 | 0.01735 | 0.039265 | 0.585343 | -3.88624 | 0.824283 | 0.418896 | 0.518245 | 2.967923 | 0 | 1.735939 | 2.597418 | 8.426372 | 13.52668 | 15.63387 | 9.382952 | 9.193245 | 1.538429 | 0 | 0 | 16.68654 | 14.14949 | 8.098703 |
| Stat3 | 20848 | 2.211864 | 7.752153 | 60.09326 | 2.66E-25 | 5.15E-23 | -0.9082 | 7.752153 | -28.7493 | 1.49E-18 | 6.41E-17 | 8076.46 | 8223.204 | 8078.193 | 40249.59 | 40990.08 | 39339.64 | 23719.42 | 22372.98 | 20929.57 | 8330.593 | 8471.684 | 8930.719 | 5640.05 | 5640.143 | 5785.713 |
| Gimd1 | 433653 | 2.212138 | -5.16477 | 2.483451 | 0.021386 | 0.047309 | -0.11862 | -5.16477 | -0.14682 | 0.884652 | 0.918721 | 1.483961 | 0 | 0 | 3.896127 | 4.213186 | 1.352668 | 1.421261 | 1.340422 | 8.044089 | 0 | 0 | 0 | 5.840288 | 1.572165 | 4.859222 |
| BE692007 | 100504727 | 2.212476 | 3.170269 | 21.20149 | 8.00E-16 | 1.36E-14 | -0.02236 | 3.170269 | -0.29383 | 0.771724 | 0.844974 | 284.1786 | 250.4259 | 276.0143 | 1283.125 | 1355.241 | 1389.19 | 1527.855 | 1262.677 | 1343.363 | 276.9172 | 269.9349 | 276.1463 | 365.4352 | 297.1392 | 327.9975 |
| H2-Q9 | 110558 | 2.213031 | 3.821444 | 11.63517 | 1.05E-10 | 7.48E-10 | 2.990866 | 3.821444 | 28.00272 | 2.57E-18 | 1.03E-16 | 142.4603 | 199.8079 | 122.3837 | 693.5107 | 832.8064 | 727.7352 | 6593.229 | 6117.685 | 5884.826 | 505.3739 | 491.8305 | 565.7267 | 1146.365 | 1029.768 | 1367.061 |
| Rilpl1 | 75695 | 2.214283 | 2.439995 | 22.78614 | 1.83E-16 | 3.59E-15 | -3.34355 | 2.439995 | -22.3544 | 2.71E-16 | 6.77E-15 | 252.2735 | 272.6267 | 289.0339 | 1372.736 | 1311.705 | 1331.025 | 112.2796 | 154.1485 | 143.6444 | 391.5302 | 402.6147 | 447.8048 | 153.5162 | 190.232 | 183.8406 |
| Etos1 | 114660 | 2.215797 | -3.40987 | 2.79732 | 0.01069 | 0.025464 | 2.912534 | -3.40987 | 5.344478 | 2.53E-05 | 8.89E-05 | 1.483961 | 3.55214 | 0 | 5.194837 | 8.426372 | 6.763339 | 71.06304 | 54.95729 | 48.26453 | 1.538429 | 0.762528 | 0 | 8.343269 | 9.432991 | 9.718443 |
| Srgap1 | 117600 | 2.216087 | 0.542646 | 6.433897 | 2.07E-06 | 8.26E-06 | 1.213289 | 0.542646 | 5.614315 | 1.35E-05 | 4.90E-05 | 12.61367 | 31.96926 | 15.62345 | 97.40319 | 102.5209 | 75.74939 | 241.6143 | 213.1271 | 210.2955 | 29.99936 | 22.11331 | 14.92683 | 599.8811 | 585.6316 | 596.0645 |
| Tnfsf4 | 22164 | 2.216474 | 2.874238 | 13.06296 | 1.19E-11 | 9.75E-11 | 4.216307 | 2.874238 | 46.32579 | 6.60E-23 | 1.21E-20 | 101.6514 | 100.348 | 131.9314 | 579.2243 | 574.3977 | 492.3711 | 10565.65 | 9919.121 | 11184.73 | 188.4575 | 201.3074 | 155.9853 | 130.155 | 145.4253 | 141.7273 |
| Cldn4 | 12740 | 2.217307 | -2.52569 | 3.436599 | 0.002433 | 0.006567 | 0.282251 | -2.52569 | 0.541584 | 0.593714 | 0.686688 | 2.967923 | 1.77607 | 2.603909 | 12.98709 | 8.426372 | 21.64268 | 11.37009 | 16.08506 | 25.28142 | 7.692145 | 3.812639 | 2.985365 | 45.88798 | 33.01547 | 29.9652 |
| Kcnh7 | 170738 | 2.218459 | -2.20694 | 3.773957 | 0.001089 | 0.003117 | 0.450206 | -2.20694 | 1.041226 | 0.309422 | 0.404117 | 5.193865 | 3.55214 | 5.207818 | 22.07806 | 32.30109 | 20.29002 | 24.16143 | 38.87223 | 42.51876 | 2.307643 | 8.387806 | 0.746341 | 22.52683 | 34.58764 | 26.72572 |
| Igfbp1 | 16006 | 2.218721 | -3.39589 | 2.817987 | 0.010203 | 0.024433 | 1.377425 | -3.39589 | 2.263363 | 0.03415 | 0.062578 | 1.483961 | 2.664105 | 0 | 6.493546 | 4.213186 | 8.116006 | 21.31891 | 26.80843 | 10.3424 | 1.538429 | 1.525056 | 1.492683 | 20.02385 | 14.14949 | 19.43689 |
| Nexmif | 245555 | 2.219116 | -0.51594 | 4.951936 | 6.42E-05 | 0.000215 | 0.522161 | -0.51594 | 1.729595 | 0.09814 | 0.153423 | 23.74338 | 21.31284 | 10.41564 | 85.7148 | 120.778 | 68.98605 | 112.2796 | 109.9146 | 180.4174 | 16.1535 | 6.100223 | 5.97073 | 43.385 | 82.53868 | 89.08573 |
| Serpina11 | 380780 | 2.219411 | -2.87509 | 2.562039 | 0.018025 | 0.040633 | 2.671324 | -2.87509 | 4.444223 | 0.000217 | 0.000668 | 0.741981 | 8.88035 | 0 | 12.98709 | 5.617581 | 5.410671 | 35.53152 | 65.68067 | 60.90525 | 0.769214 | 1.525056 | 1.492683 | 28.36712 | 31.4433 | 34.82442 |
| F930017D | 414125 | 2.220191 | -3.67301 | 2.644012 | 0.015051 | 0.034663 | 0.037628 | -3.67301 | 0.051793 | 0.959175 | 0.966451 | 2.967923 | 0 | 3.471878 | 5.194837 | 8.426372 | 13.52668 | 7.106304 | 5.361687 | 17.23733 | 3.076858 | 0.762528 | 0.746341 | 8.343269 | 14.93557 | 4.859222 |
| Gm6377 | 622976 | 2.220868 | 4.928148 | 48.2384 | 2.81E-23 | 2.73E-21 | -2.57249 | 4.928148 | -47.7299 | 3.51E-23 | 7.22E-21 | 2343.175 | 2355.957 | 2176.868 | 11261.11 | 11749.17 | 11240.67 | 1938.6 | 1832.357 | 2187.992 | 983.0561 | 1010.349 | 1095.629 | 382.9561 | 349.8068 | 390.3575 |
| Icam4 | 78369 | 2.220917 | -0.66656 | 5.935391 | 6.41E-06 | 2.42E-05 | -0.24266 | -0.66656 | -0.85068 | 0.404389 | 0.503376 | 8.161788 | 23.97694 | 24.30315 | 100.0006 | 73.02855 | 75.74939 | 85.27565 | 60.31898 | 73.54596 | 18.46115 | 18.30067 | 23.13658 | 41.71635 | 27.51289 | 38.87377 |
| Snx10 | 71982 | 2.221437 | 7.154889 | 60.28781 | 2.49E-25 | 4.92E-23 | -1.10223 | 7.154889 | -33.4732 | 6.19E-20 | 4.05E-18 | 5765.932 | 6300.608 | 5531.57 | 28654.72 | 30208.54 | 28611.63 | 14137.28 | 13866.66 | 14228.84 | 6442.94 | 6094.123 | 6561.832 | 2979.381 | 2803.957 | 3015.957 |
| Ppm1n | 232941 | 2.221715 | -0.34665 | 6.978311 | 6.25E-07 | 2.66E-06 | -1.26806 | -0.34665 | -4.43937 | 0.00022 | 0.000675 | 32.64715 | 33.74533 | 17.35939 | 119.4812 | 183.9758 | 114.9768 | 73.90556 | 71.04235 | 40.22045 | 26.15329 | 32.7887 | 24.62926 | 35.87606 | 41.66238 | 31.58494 |
| Ttll2 | 100216474 | 2.22251 | -3.03941 | 2.978654 | 0.007079 | 0.017539 | 0.745457 | -3.03941 | 1.358756 | 0.188412 | 0.26673 | 2.967923 | 1.77607 | 6.943757 | 10.38967 | 15.44835 | 37.8747 | 21.31891 | 33.51054 | 43.66791 | 2.307643 | 5.337695 | 0 | 3.337308 | 4.716496 | 5.669092 |
| Serpinb9e | 20710 | 2.22264 | -3.84812 | 2.585708 | 0.017114 | 0.038799 | 1.084203 | -3.84812 | 1.56577 | 0.13211 | 0.197601 | 0 | 1.77607 | 1.735939 | 5.194837 | 9.830767 | 2.705335 | 17.05513 | 17.42548 | 6.894934 | 0 | 0 | 1.492683 | 17.52087 | 18.0799 | 22.67637 |
| Gm11538 | 432593 | 2.223507 | -2.8136 | 2.779896 | 0.011117 | 0.026393 | 1.648688 | -2.8136 | 2.864603 | 0.009182 | 0.020041 | 2.967923 | 0 | 6.075787 | 7.792255 | 14.04395 | 9.468674 | 27.00396 | 50.93603 | 29.87805 | 3.846072 | 1.525056 | 1.492683 | 18.35519 | 29.08506 | 19.43689 |
| Adap2 | 216991 | 2.223962 | 6.544933 | 44.67487 | 1.42E-22 | 1.09E-20 | -2.34604 | 6.544933 | -42.0977 | 4.98E-22 | 6.24E-20 | 4610.668 | 4854.887 | 4345.056 | 23191.05 | 22994.16 | 22617.96 | 5112.275 | 5037.305 | 4070.309 | 6009.873 | 6174.951 | 6240.906 | 1837.188 | 1746.676 | 1735.552 |
| Gm15350 | 100504249 | 2.227063 | -2.93114 | 2.572092 | 0.017632 | 0.039844 | -0.78318 | -2.93114 | -0.99038 | 0.333083 | 0.429622 | 5.193865 | 0 | 8.679696 | 11.68838 | 12.63956 | 17.58468 | 7.106304 | 4.021265 | 16.08818 | 6.153716 | 9.912862 | 16.41951 | 15.01788 | 9.432991 | 3.239481 |
| Nfkbib | 18036 | 2.227665 | 5.600821 | 35.71043 | 1.59E-20 | 7.72E-19 | -0.11852 | 5.600821 | -2.41919 | 0.024559 | 0.047076 | 1468.38 | 1362.246 | 1625.707 | 7553.292 | 7368.862 | 7312.522 | 7356.446 | 7055.98 | 6889.188 | 2022.265 | 1907.845 | 2221.858 | 1227.295 | 1231.791 | 1409.174 |
| H2-K2 | 630499 | 2.228174 | 3.566422 | 12.80037 | 1.75E-11 | 1.40E-10 | 2.128435 | 3.566422 | 21.02416 | 9.49E-16 | 2.06E-14 | 154.332 | 137.6454 | 153.6306 | 750.6539 | 716.2416 | 769.6679 | 3699.542 | 3603.054 | 2969.418 | 916.9036 | 1042.376 | 1032.936 | 499.7618 | 460.6444 | 457.5767 |
| Car4 | 12351 | 2.228202 | -3.06369 | 2.606749 | 0.01634 | 0.037248 | -1.36419 | -3.06369 | -1.65626 | 0.112298 | 0.17231 | 6.677827 | 0 | 5.207818 | 9.090964 | 16.85274 | 13.52668 | 7.106304 | 6.702109 | 2.298311 | 2.307643 | 13.7255 | 4.478048 | 24.19548 | 11.79124 | 16.19741 |
| Capn6 | 12338 | 2.229446 | -2.63762 | 2.528122 | 0.019409 | 0.043409 | 0.458418 | -2.63762 | 0.671575 | 0.509059 | 0.606265 | 9.64575 | 0 | 6.075787 | 14.2858 | 18.25714 | 13.52668 | 24.16143 | 13.40422 | 31.0272 | 6.92293 | 0.762528 | 6.717071 | 20.02385 | 27.51289 | 12.14805 |
| Uchl4 | 93841 | 2.229843 | 2.336141 | 15.1026 | 7.18E-13 | 7.15E-12 | -0.06483 | 2.336141 | -0.60975 | 0.548463 | 0.643404 | 149.1381 | 174.9429 | 199.633 | 807.7971 | 884.769 | 902.2294 | 999.1464 | 904.7847 | 719.3714 | 122.3051 | 117.4293 | 110.4585 | 167.6997 | 150.9279 | 174.1221 |
| Serpina3n | 20716 | 2.230624 | -1.64152 | 2.793348 | 0.010786 | 0.025669 | 1.412079 | -1.64152 | 2.351611 | 0.028363 | 0.053312 | 1.483961 | 0 | 6.075787 | 9.090964 | 4.213186 | 13.52668 | 19.89765 | 30.8297 | 21.83396 | 36.92229 | 28.97606 | 32.09267 | 199.4041 | 201.2372 | 225.9538 |
| Six1 | 20471 | 2.232293 | -0.01275 | 6.469961 | 1.91E-06 | 7.65E-06 | 0.535155 | -0.01275 | 2.315535 | 0.03061 | 0.05686 | 19.2915 | 25.75301 | 13.88751 | 88.31222 | 94.09448 | 109.5661 | 167.7088 | 123.3188 | 150.5394 | 43.07601 | 44.22662 | 45.52682 | 39.21337 | 70.74744 | 71.26858 |
| Gkn1 | 66283 | 2.232999 | -4.006 | 2.50334 | 0.020484 | 0.045513 | -0.18887 | -4.006 | -0.24024 | 0.812435 | 0.880677 | 1.483961 | 0 | 7.811726 | 10.38967 | 7.021976 | 9.468674 | 5.685043 | 6.702109 | 12.64071 | 5.384501 | 0 | 0.746341 | 1.668654 | 3.930413 | 4.049351 |
| Myo3a | 667663 | 2.23372 | -2.26236 | 3.070689 | 0.005728 | 0.014455 | 1.351346 | -2.26236 | 2.697291 | 0.013373 | 0.027795 | 3.709904 | 1.77607 | 6.075787 | 24.67547 | 18.25714 | 14.87935 | 22.74017 | 60.31898 | 86.18667 | 3.076858 | 0.762528 | 1.492683 | 32.53875 | 50.30929 | 39.68364 |
| Gm19303 | 100502643 | 2.234751 | -4.02276 | 2.6618 | 0.01447 | 0.033447 | 1.463593 | -4.02276 | 2.218075 | 0.037523 | 0.06779 | 0 | 0.888035 | 2.603909 | 5.194837 | 8.426372 | 2.705335 | 9.948826 | 13.40422 | 25.28142 | 0.769214 | 0 | 0 | 14.18356 | 8.646909 | 12.95792 |
| Aspg | 104816 | 2.23484 | -2.42477 | 2.894341 | 0.008582 | 0.020893 | 1.591769 | -2.42477 | 2.88886 | 0.00869 | 0.019081 | 6.677827 | 0 | 3.471878 | 10.38967 | 12.63956 | 10.82134 | 35.53152 | 34.85097 | 39.07129 | 4.615287 | 3.812639 | 8.209754 | 21.6925 | 29.87114 | 38.87377 |
| Lin7a | 108030 | 2.236572 | -2.34341 | 2.725905 | 0.012546 | 0.029439 | 0.067417 | -2.34341 | 0.104676 | 0.91761 | 0.933475 | 1.483961 | 9.768385 | 4.339848 | 44.15611 | 15.44835 | 13.52668 | 24.16143 | 24.12759 | 21.83396 | 11.53822 | 5.337695 | 0 | 28.36712 | 29.87114 | 40.49351 |
| Tafa2 | 268354 | 2.236668 | -2.15004 | 2.731175 | 0.012399 | 0.02912 | 1.078917 | -2.15004 | 1.830445 | 0.081184 | 0.130683 | 1.483961 | 3.55214 | 5.207818 | 9.090964 | 19.66153 | 25.70069 | 14.21261 | 57.63814 | 55.15947 | 7.692145 | 3.050111 | 2.239024 | 50.89394 | 65.24486 | 33.20468 |
| Ndp | 17986 | 2.236921 | -3.7878 | 2.506108 | 0.020361 | 0.045293 | -0.25755 | -3.7878 | -0.32585 | 0.747713 | 0.824277 | 4.451884 | 2.664105 | 0 | 10.38967 | 7.021976 | 9.468674 | 4.263783 | 9.382952 | 10.3424 | 4.615287 | 0 | 0 | 6.674615 | 16.50774 | 12.95792 |
| Pamr1 | 210622 | 2.237078 | -2.20907 | 4.155431 | 0.000435 | 0.001319 | -0.76361 | -2.20907 | -1.66808 | 0.109907 | 0.169233 | 8.161788 | 2.664105 | 7.811726 | 29.87031 | 28.08791 | 32.46403 | 31.26774 | 14.74464 | 13.78987 | 3.076858 | 3.050111 | 8.956095 | 15.01788 | 22.7964 | 25.91585 |
| 9930111J2 | 667214 | 2.239039 | 6.356796 | 15.42315 | 9.93E-09 | 5.36E-08 | 0.170596 | 6.356796 | 1.563552 | 0.14657 | 0.215806 | 2421.083 | 2416.343 | 1421.734 | 10546.82 | 9958.567 | 10304.62 | 12681.91 | 12264.86 | 11235.29 | 4001.454 | 4185.515 | 4016.062 | 2663.172 | 2840.117 | 2802.151 |
| Cts8 | 56094 | 2.240565 | -3.89791 | 2.901098 | 0.008451 | 0.020605 | 1.876609 | -3.89791 | 3.26172 | 0.003672 | 0.008879 | 2.967923 | 0 | 0.86797 | 7.792255 | 2.808791 | 8.116006 | 18.47639 | 25.46801 | 27.57973 | 0 | 0 | 0 | 15.85221 | 15.72165 | 10.52831 |
| Mcoln2 | 68279 | 2.241695 | 5.720322 | 33.96609 | 4.56E-20 | 1.95E-18 | 0.463894 | 5.720322 | 9.628905 | 3.20E-09 | 1.97E-08 | 1137.456 | 1032.785 | 1077.15 | 5251.98 | 5829.645 | 5378.207 | 7890.84 | 8203.381 | 7490.196 | 6912.93 | 6755.997 | 6607.359 | 917.7596 | 1017.191 | 1028.535 |
| Pcdh17 | 219228 | 2.241929 | -0.23755 | 5.873466 | 7.39E-06 | 2.77E-05 | -0.45896 | -0.23755 | -1.5436 | 0.137383 | 0.204186 | 25.22735 | 15.09659 | 24.30315 | 100.0006 | 110.9472 | 116.3294 | 61.11422 | 54.95729 | 145.9428 | 21.538 | 28.97606 | 17.91219 | 95.9476 | 66.03094 | 76.12781 |
| Ripk2 | 192656 | 2.242762 | 6.462604 | 47.21574 | 4.42E-23 | 4.03E-21 | 0.449769 | 6.462604 | 13.03183 | 1.24E-11 | 1.15E-10 | 2090.902 | 2362.173 | 2165.584 | 11171.5 | 11329.26 | 10834.87 | 16031.82 | 15748.62 | 15511.3 | 3496.08 | 3335.297 | 3398.838 | 3021.932 | 2987.9 | 3278.355 |
| Psg25 | 114868 | 2.244402 | -3.08282 | 2.793893 | 0.010772 | 0.02564 | 1.231278 | -3.08282 | 1.991725 | 0.059351 | 0.100315 | 0 | 6.216245 | 1.735939 | 6.493546 | 11.23516 | 6.763339 | 25.5827 | 14.74464 | 21.83396 | 3.076858 | 3.812639 | 2.239024 | 15.01788 | 20.43815 | 13.76779 |
| Ak9 | 633979 | 2.244521 | -1.23683 | 4.214644 | 0.000377 | 0.001153 | 0.696954 | -1.23683 | 1.902002 | 0.070753 | 0.116291 | 5.193865 | 5.32821 | 11.2836 | 37.66257 | 49.15383 | 24.34802 | 39.7953 | 72.38277 | 72.3968 | 4.615287 | 8.387806 | 6.717071 | 120.9774 | 81.75259 | 95.56469 |
| Ptpn13 | 19249 | 2.246056 | 0.683528 | 7.586566 | 1.72E-07 | 7.85E-07 | 1.187707 | 0.683528 | 6.539099 | 1.64E-06 | 6.78E-06 | 31.90517 | 39.07354 | 26.03909 | 128.5722 | 198.0197 | 171.7888 | 322.6262 | 466.4668 | 379.2213 | 49.22973 | 37.36386 | 53.73657 | 75.08942 | 92.75775 | 88.27586 |
| Clec2g | 70809 | 2.24651 | 1.178309 | 16.56437 | 1.16E-13 | 1.33E-12 | -4.34214 | 1.178309 | -15.7467 | 3.16E-13 | 3.90E-12 | 214.4324 | 211.3523 | 164.0463 | 1050.656 | 981.6723 | 950.9254 | 36.95278 | 56.29771 | 58.60694 | 213.8416 | 249.3466 | 158.2243 | 25.86413 | 19.65207 | 21.8665 |
| Gm4876 | 232599 | 2.246856 | 2.313923 | 16.40155 | 1.41E-13 | 1.58E-12 | -0.55946 | 2.313923 | -5.26671 | 3.04E-05 | 0.000106 | 176.5914 | 166.9506 | 130.1954 | 816.8881 | 789.2701 | 791.3106 | 494.5988 | 666.1896 | 541.2523 | 160.7658 | 180.7191 | 175.3902 | 202.7414 | 176.0825 | 191.1294 |
| Tcp10c | 100041352 | 2.247841 | -1.97017 | 3.482371 | 0.002183 | 0.00594 | 0.339919 | -1.97017 | 0.708954 | 0.486026 | 0.584684 | 7.419807 | 11.54445 | 0.86797 | 27.27289 | 22.47032 | 24.34802 | 25.5827 | 26.80843 | 47.11538 | 9.230573 | 8.387806 | 2.985365 | 27.53279 | 36.1598 | 28.34546 |
| Zfp711 | 245595 | 2.249102 | -2.38057 | 3.982373 | 0.00066 | 0.001952 | -0.52142 | -2.38057 | -1.08155 | 0.291523 | 0.384617 | 4.451884 | 6.216245 | 1.735939 | 16.88322 | 18.25714 | 27.05335 | 14.21261 | 12.0638 | 17.23733 | 8.461359 | 3.050111 | 5.97073 | 30.8701 | 26.72681 | 21.8665 |
| Nr0b1 | 11614 | 2.250598 | -3.32614 | 2.891849 | 0.008631 | 0.021004 | 0.206773 | -3.32614 | 0.325113 | 0.748261 | 0.824778 | 1.483961 | 4.440175 | 1.735939 | 9.090964 | 19.66153 | 12.17401 | 8.527565 | 18.7659 | 21.83396 | 0 | 5.337695 | 0 | 9.177596 | 12.57732 | 20.24676 |
| Acsl1 | 14081 | 2.251685 | 6.370556 | 51.28144 | 7.69E-24 | 8.91E-22 | -1.91514 | 6.370556 | -42.3975 | 4.29E-22 | 5.62E-20 | 3374.528 | 3624.959 | 3872.012 | 18606.61 | 18415.84 | 18025.65 | 5252.98 | 5183.411 | 4768.996 | 4485.289 | 4284.644 | 4616.867 | 1874.733 | 2043.815 | 2003.619 |
| Il1r2 | 16178 | 2.252193 | 1.30183 | 8.839048 | 1.40E-08 | 7.39E-08 | 1.251361 | 1.30183 | 8.101249 | 5.98E-08 | 3.03E-07 | 46.00281 | 48.84192 | 59.02193 | 240.2612 | 249.9824 | 292.1762 | 703.5241 | 628.6578 | 607.9033 | 39.99915 | 41.93903 | 34.3317 | 236.1145 | 149.3557 | 246.2006 |
| Mir29b-1 | 387223 | 2.25302 | -2.40807 | 5.048807 | 5.10E-05 | 0.000173 | -1.54465 | -2.40807 | -3.56916 | 0.001776 | 0.004604 | 5.935846 | 18.64873 | 7.811726 | 48.05224 | 60.389 | 40.58003 | 14.21261 | 16.08506 | 20.6848 | 3.076858 | 7.625279 | 9.702436 | 3.337308 | 3.930413 | 1.619741 |
| Rgs9 | 19739 | 2.253794 | 1.824916 | 13.72275 | 4.62E-12 | 4.02E-11 | 0.498219 | 1.824916 | 4.63793 | 0.000136 | 0.000432 | 130.5886 | 87.02743 | 92.87275 | 540.263 | 549.1185 | 493.7237 | 787.3785 | 727.849 | 802.1106 | 109.9977 | 89.97829 | 106.7268 | 125.9834 | 96.68816 | 80.98703 |
| Prkcq | 18761 | 2.256459 | 0.786564 | 7.805563 | 1.09E-07 | 5.12E-07 | 0.694046 | 0.786564 | 3.752382 | 0.001147 | 0.003102 | 49.71271 | 39.07354 | 26.03909 | 171.4296 | 199.4241 | 211.0162 | 332.575 | 379.3394 | 278.0957 | 49.22973 | 52.61442 | 24.62926 | 122.6461 | 120.2706 | 119.8608 |

| | | | | | | | | | | | | | | | | | | | | | | | | | | |
|---|---|---|---|---|---|---|---|---|---|---|---|---|---|---|---|---|---|---|---|---|---|---|---|---|---|---|
| Lcn3 | 16820 | 2.25675 | -4.88458 | 2.579211 | 0.017359 | 0.039281 | 0.592562 | -4.88458 | 0.784444 | 0.4414 | 0.540292 | 0 | 1.77607 | 0 | 3.896127 | 1.404395 | 4.058003 | 5.685043 | 9.382952 | 2.298311 | 0 | 0 | 0 | 3.337308 | 7.074744 | 8.098703 |
| Il22 | 50929 | 2.257256 | -4.36638 | 2.522271 | 0.019658 | 0.043894 | 0.829277 | -4.36638 | 1.122251 | 0.274223 | 0.365564 | 2.225942 | 0 | 0.86797 | 9.090964 | 5.617581 | 2.705335 | 2.842522 | 21.44675 | 14.93902 | 0 | 0.762528 | 0 | 4.171635 | 6.288661 | 4.049351 |
| Kremen1 | 84035 | 2.258286 | 6.530527 | 49.42169 | 1.68E-23 | 1.73E-21 | -0.52978 | 6.530527 | -14.2227 | 2.31E-12 | 2.43E-11 | 3633.48 | 3569.013 | 3474.482 | 18735.18 | 18049.29 | 17787.58 | 13641.26 | 13333.18 | 12413.18 | 3509.926 | 3516.016 | 3413.765 | 1613.588 | 1740.387 | 1886.188 |
| Gm20597 | 100502868 | 2.259921 | -1.98196 | 3.146228 | 0.004808 | 0.012302 | 1.653344 | -1.98196 | 3.598715 | 0.001656 | 0.004323 | 15.5816 | 1.77607 | 2.603909 | 14.2858 | 46.34504 | 22.99535 | 76.74809 | 81.76573 | 86.18667 | 3.076858 | 0 | 2.985365 | 60.90587 | 40.09021 | 45.35273 |
| Itgb1bp2 | 26549 | 2.260967 | 0.867451 | 13.18906 | 9.89E-12 | 8.21E-11 | -0.28335 | 0.867451 | -2.24047 | 0.035819 | 0.065199 | 57.13252 | 67.49066 | 65.96569 | 331.1708 | 337.0549 | 304.3502 | 305.5711 | 264.0631 | 264.3058 | 63.8448 | 61.76476 | 72.3951 | 45.05365 | 40.09021 | 35.63429 |
| Gm6654 | 626175 | 2.262869 | 0.570745 | 9.081401 | 8.83E-09 | 4.80E-08 | -1.03219 | 0.570745 | -4.87113 | 7.79E-05 | 0.000256 | 55.64856 | 72.81887 | 52.07818 | 353.2489 | 273.8571 | 290.8236 | 153.4962 | 174.2548 | 140.197 | 41.53758 | 94.55345 | 58.21462 | 47.55663 | 32.22939 | 19.43689 |
| Irgm1 | 15944 | 2.264887 | 8.670471 | 87.02797 | 1.02E-28 | 9.89E-26 | 1.113101 | 8.670471 | 69.09524 | 1.37E-26 | 1.17E-23 | 9780.048 | 10375.8 | 9644.01 | 50437.97 | 51455.64 | 50852.19 | 114955.8 | 113021.7 | 114834 | 14742.76 | 14563.52 | 14188.69 | 10241.36 | 10252.09 | 10204.37 |
| Slc9a3 | 105243 | 2.266429 | -2.38201 | 4.214051 | 0.000378 | 0.001155 | 0.080327 | -2.38201 | 0.191483 | 0.849957 | 0.912581 | 3.709904 | 2.664105 | 5.207818 | 20.77935 | 22.47032 | 18.93735 | 21.31891 | 25.46801 | 21.83396 | 3.076858 | 1.525056 | 5.224389 | 25.02981 | 36.94588 | 24.29611 |
| Gm10865 | 100038602 | 2.266449 | -0.3428 | 6.11438 | 4.26E-06 | 1.64E-05 | -0.06113 | -0.3428 | -0.2265 | 0.822971 | 0.88997 | 13.35565 | 30.19319 | 26.90706 | 102.598 | 132.0132 | 105.5081 | 83.85439 | 128.6805 | 125.258 | 44.61444 | 28.97606 | 35.82438 | 10.84625 | 33.01547 | 29.9652 |
| Dusp26 | 66959 | 2.267348 | -3.65541 | 2.796881 | 0.0107 | 0.025487 | 0.859242 | -3.65541 | 1.316986 | 0.201809 | 0.282675 | 1.483961 | 0 | 3.471878 | 6.493546 | 12.63956 | 4.058003 | 9.948826 | 13.40422 | 18.38649 | 0 | 0 | 1.492683 | 23.36115 | 26.72681 | 18.62702 |
| Birc7 | 329581 | 2.267545 | -4.68975 | 2.489423 | 0.021111 | 0.046762 | 1.285762 | -4.68975 | 1.738882 | 0.096461 | 0.151193 | 0 | 0 | 3.471878 | 2.597418 | 2.808791 | 6.763339 | 4.263783 | 17.42548 | 13.78987 | 0 | 0 | 0 | 4.171635 | 2.358248 | 5.669092 |
| Ggct | 110175 | 2.26819 | 1.534403 | 13.88831 | 3.67E-12 | 3.23E-11 | -0.0222 | 1.534403 | -0.19269 | 0.849022 | 0.911881 | 70.48817 | 88.8035 | 78.98523 | 418.1844 | 398.8483 | 399.037 | 437.7483 | 406.1478 | 403.3536 | 133.0741 | 103.7038 | 117.1756 | 113.4685 | 96.68816 | 85.84625 |
| Gm11166 | 625850 | 2.268649 | -2.7193 | 3.080471 | 0.0056 | 0.014159 | 0.143344 | -2.7193 | 0.251987 | 0.803462 | 0.87264 | 7.419807 | 7.10428 | 0.86797 | 28.5716 | 11.23516 | 29.75869 | 14.21261 | 29.48928 | 33.32551 | 6.92293 | 1.525056 | 5.97073 | 10.84625 | 2.358248 | 6.478962 |
| Plpp3 | 67916 | 2.268728 | 2.796397 | 11.87594 | 7.17E-11 | 5.23E-10 | 2.430756 | 2.796397 | 22.7972 | 1.81E-16 | 4.69E-15 | 115.749 | 82.58725 | 80.72117 | 459.743 | 519.6262 | 462.6124 | 2754.404 | 2903.354 | 2452.298 | 292.3015 | 274.51 | 243.3072 | 395.471 | 392.2552 | 362.012 |
| Bhmt | 12116 | 2.268752 | -2.78624 | 2.699983 | 0.013293 | 0.031 | 0.940926 | -2.78624 | 1.503507 | 0.147362 | 0.216717 | 1.483961 | 7.992315 | 0.86797 | 6.493546 | 29.4923 | 12.17401 | 34.11026 | 20.10633 | 28.72889 | 6.153716 | 2.287584 | 2.985365 | 4.171635 | 25.15464 | 13.76779 |
| Ces3b | 13909 | 2.270995 | -3.34603 | 2.630801 | 0.015497 | 0.035574 | 1.154202 | -3.34603 | 1.731818 | 0.097736 | 0.152891 | 2.225942 | 0 | 3.471878 | 6.493546 | 5.617581 | 13.52668 | 15.63387 | 14.74464 | 28.72889 | 0 | 3.050111 | 1.492683 | 25.86413 | 13.3634 | 17.81715 |
| A730049H | 74516 | 2.271721 | -4.7518 | 2.476485 | 0.02171 | 0.047967 | 0.254754 | -4.7518 | 0.316767 | 0.754498 | 0.829986 | 0.741981 | 0 | 0.86797 | 1.298709 | 7.021976 | 5.410671 | 4.263783 | 2.680843 | 9.193245 | 0 | 3.050111 | 0 | 4.171635 | 3.14433 | 1.619741 |
| Unc45b | 217012 | 2.272074 | -1.63941 | 3.783473 | 0.001065 | 0.003053 | -0.04941 | -1.63941 | -0.11011 | 0.913353 | 0.930368 | 12.61367 | 2.664105 | 7.811726 | 23.37676 | 28.08791 | 71.69139 | 34.11026 | 38.87223 | 35.62382 | 6.153716 | 12.20045 | 7.463413 | 26.69846 | 50.30929 | 23.48624 |
| E330012B | 100039641 | 2.273762 | -3.83106 | 2.535366 | 0.019106 | 0.042795 | 1.595675 | -3.83106 | 2.363425 | 0.027661 | 0.052189 | 0 | 3.55214 | 1.735939 | 10.38967 | 7.021976 | 4.058003 | 28.42522 | 20.10633 | 20.6848 | 0 | 1.525056 | 0 | 7.508942 | 16.50774 | 3.239481 |
| Cyp2j8 | 665095 | 2.274405 | -3.60919 | 3.418237 | 0.002541 | 0.006828 | 0.747802 | -3.60919 | 1.420916 | 0.169787 | 0.244247 | 1.483961 | 2.664105 | 0.86797 | 14.2858 | 8.426372 | 6.763339 | 12.79135 | 20.10633 | 18.38649 | 0.769214 | 0.762528 | 0 | 7.508942 | 10.21907 | 12.95792 |
| Adgrg6 | 215798 | 2.275569 | 6.685446 | 61.03098 | 1.92E-25 | 3.91E-23 | -0.9441 | 6.685446 | -29.3705 | 9.53E-19 | 4.38E-17 | 5013.564 | 5377.94 | 4988.221 | 26248.21 | 27496.65 | 25691.22 | 14465.59 | 14406.85 | 13994.42 | 4199.911 | 3976.583 | 4310.867 | 1001.192 | 1131.959 | 1151.636 |
| Cdhr5 | 72040 | 2.280234 | -1.40718 | 3.60373 | 0.001636 | 0.004543 | 1.680392 | -1.40718 | 4.298362 | 0.000309 | 0.000922 | 8.903769 | 6.216245 | 4.339848 | 33.76644 | 51.96262 | 22.99535 | 162.0237 | 103.2125 | 96.52907 | 2.307643 | 19.0632 | 3.731706 | 34.2074 | 47.95104 | 25.91585 |
| Serpina9 | 71907 | 2.281064 | -3.79652 | 2.760336 | 0.011616 | 0.027442 | -0.03862 | -3.79652 | -0.05367 | 0.957701 | 0.96515 | 0.741981 | 3.55214 | 0.86797 | 11.68838 | 12.63956 | 4.058003 | 8.527565 | 2.680843 | 22.98311 | 0 | 1.525056 | 0.746341 | 9.177596 | 11.79124 | 9.718443 |
| Sfrp1 | 20377 | 2.282705 | -1.88207 | 3.860906 | 0.000884 | 0.002564 | 0.70759 | -1.88207 | 1.691985 | 0.105199 | 0.163081 | 2.225942 | 7.10428 | 6.943757 | 40.25998 | 22.47032 | 18.93735 | 29.84648 | 44.23392 | 62.0544 | 3.076858 | 9.150334 | 5.97073 | 28.36712 | 36.94588 | 26.72572 |
| Oasl2 | 23962 | 2.282887 | 8.838581 | 84.86456 | 1.74E-28 | 1.34E-25 | 0.223591 | 8.838581 | 12.1281 | 4.84E-11 | 3.98E-10 | 11761.88 | 11872.14 | 11967.56 | 62489.99 | 61402.97 | 60995.85 | 76350.13 | 74861.21 | 73297.74 | 21021.86 | 20915.38 | 20883.37 | 13525.27 | 13581.94 | 13068.88 |
| Gsdmc | 83492 | 2.283613 | -3.0632 | 3.165492 | 0.004598 | 0.011797 | 1.105406 | -3.0632 | 2.077598 | 0.049998 | 0.08661 | 2.225942 | 0.888035 | 4.339848 | 12.98709 | 12.63956 | 12.17401 | 22.74017 | 24.12759 | 41.3696 | 4.615287 | 0 | 1.492683 | 11.68058 | 11.00516 | 13.76779 |
| 2310043L1 | 75589 | 2.284677 | -3.55666 | 2.715757 | 0.012834 | 0.030051 | 1.446313 | -3.55666 | 2.311515 | 0.030871 | 0.057279 | 0 | 1.77607 | 7.811726 | 14.2858 | 7.021976 | 6.763339 | 21.31891 | 32.17012 | 26.43058 | 2.307643 | 3.812639 | 1.492683 | 0.834327 | 3.14433 | 3.239481 |
| Ddc | 13195 | 2.285402 | -2.05896 | 3.370262 | 0.002846 | 0.007584 | -1.0968 | -2.05896 | -1.81826 | 0.083087 | 0.133213 | 16.32358 | 0.888035 | 14.75548 | 40.25998 | 30.8967 | 40.58003 | 8.527565 | 16.08506 | 34.47467 | 6.92293 | 9.150334 | 2.239024 | 30.03577 | 19.65207 | 20.24676 |
| Olfr722 | 258487 | 2.286284 | -3.9464 | 2.976343 | 0.007117 | 0.017621 | 0.444648 | -3.9464 | 0.702626 | 0.489882 | 0.58806 | 4.451884 | 1.77607 | 0 | 11.68838 | 5.617581 | 8.116006 | 15.63387 | 9.382952 | 11.49156 | 0.769214 | 0 | 0 | 6.674615 | 8.646909 | 9.718443 |
| Gm20093 | 100504159 | 2.287056 | -3.27792 | 2.535692 | 0.019092 | 0.042775 | 0.563818 | -3.27792 | 0.77895 | 0.44456 | 0.543352 | 3.709904 | 0 | 3.471878 | 18.18193 | 2.808791 | 14.87935 | 11.37009 | 21.44675 | 13.78987 | 3.076858 | 0 | 1.492683 | 22.52683 | 25.15464 | 17.00728 |
| Tmc1 | 13409 | 2.287911 | -1.82046 | 4.026243 | 0.000594 | 0.001768 | 0.288606 | -1.82046 | 0.687444 | 0.499206 | 0.597071 | 3.709904 | 7.10428 | 4.339848 | 19.48064 | 18.25714 | 52.75404 | 35.53152 | 24.12759 | 43.66791 | 8.461359 | 11.43792 | 5.97073 | 28.36712 | 32.22939 | 35.63429 |
| Cep85l | 100038725 | 2.288037 | 4.166005 | 28.17149 | 2.27E-18 | 6.64E-17 | -1.27957 | 4.166005 | -17.3962 | 4.36E-14 | 6.59E-13 | 667.7827 | 680.2348 | 632.7498 | 3575.346 | 3381.784 | 3373.553 | 1701.249 | 1450.336 | 1320.38 | 1126.899 | 1017.975 | 1097.122 | 377.9501 | 433.1315 | 362.8219 |
| Gm14207 | 100043609 | 2.288889 | -3.58282 | 2.491342 | 0.021023 | 0.046579 | 0.875521 | -3.58282 | 1.189928 | 0.247143 | 0.334833 | 0 | 0.888035 | 6.943757 | 7.792255 | 4.213186 | 10.82134 | 19.89765 | 10.72337 | 13.78987 | 0 | 3.812639 | 0.746341 | 15.85221 | 16.50774 | 9.718443 |
| C030029H | 77383 | 2.289202 | -3.83791 | 2.770776 | 0.011347 | 0.026867 | 0.50971 | -3.83791 | 0.751731 | 0.460423 | 0.559153 | 4.451884 | 1.77607 | 0 | 5.194837 | 9.830767 | 10.82134 | 12.79135 | 6.702109 | 20.6848 | 0 | 1.525056 | 0 | 10.84625 | 7.860826 | 11.33818 |
| Rgl1 | 19731 | 2.290506 | 7.208464 | 66.20482 | 3.41E-26 | 9.29E-24 | -2.10565 | 7.208464 | -57.3941 | 7.07E-25 | 2.21E-22 | 6061.241 | 6198.484 | 6133.941 | 32006.69 | 32055.32 | 31974.36 | 8081.289 | 8003.658 | 7190.267 | 16189.66 | 15827.79 | 16085.15 | 2076.64 | 2285.142 | 2287.074 |
| F530104D | 676676 | 2.290928 | 0.737611 | 11.1823 | 2.18E-10 | 1.49E-09 | 0.333378 | 0.737611 | 2.44283 | 0.023344 | 0.045049 | 56.39054 | 64.82655 | 59.8899 | 288.3134 | 307.5626 | 354.3989 | 464.7523 | 373.9777 | 406.8011 | 47.6913 | 54.13948 | 36.57072 | 17.52087 | 29.87114 | 35.63429 |
| Glb1l3 | 70893 | 2.290967 | -3.37646 | 2.977826 | 0.007093 | 0.017566 | -0.0812 | -3.37646 | -0.12812 | 0.899256 | 0.9219 | 5.935846 | 0 | 8.679696 | 14.2858 | 25.27911 | 9.468674 | 11.37009 | 18.7659 | 14.93902 | 0 | 0 | 0 | 24.19548 | 24.36856 | 20.24676 |
| Obox3 | 246791 | 2.292318 | -4.12128 | 2.575013 | 0.01752 | 0.039617 | 0.289215 | -4.12128 | 0.379577 | 0.70801 | 0.78864 | 1.483961 | 1.77607 | 0 | 2.597418 | 5.617581 | 12.17401 | 7.106304 | 17.42548 | 3.447467 | 0 | 0 | 1.492683 | 15.85221 | 9.432991 | 8.908573 |
| Gm16503 | 100038537 | 2.292508 | -2.78399 | 3.25984 | 0.003689 | 0.00965 | 0.153438 | -2.78399 | 0.276404 | 0.784897 | 0.856189 | 4.451884 | 5.32821 | 0.86797 | 10.38967 | 16.85274 | 27.05335 | 28.42522 | 13.40422 | 19.53565 | 1.538429 | 3.050111 | 0 | 34.2074 | 29.08506 | 29.15533 |
| Il5ra | 16192 | 2.293614 | -1.72618 | 4.801306 | 9.21E-05 | 0.000303 | 1.192673 | -1.72618 | 3.89518 | 0.000814 | 0.002272 | 5.193865 | 6.216245 | 10.41564 | 24.67547 | 56.17581 | 39.22736 | 106.5946 | 92.4891 | 76.99342 | 2.307643 | 3.812639 | 2.985365 | 24.19548 | 22.01031 | 18.62702 |
| Kif26a | 668303 | 2.295667 | -0.75096 | 4.287339 | 0.000317 | 0.000977 | 1.18836 | -0.75096 | 3.497627 | 0.002106 | 0.005373 | 6.677827 | 7.10428 | 13.88751 | 37.66257 | 42.13186 | 64.92805 | 88.11817 | 105.8933 | 141.3461 | 18.46115 | 9.912862 | 2.985365 | 140.1669 | 152.5 | 116.6213 |
| Gpr119 | 236781 | 2.299798 | -2.70392 | 3.338367 | 0.003068 | 0.008128 | 0.31996 | -2.70392 | 0.590483 | 0.561068 | 0.655647 | 1.483961 | 3.55214 | 3.471878 | 18.18193 | 23.87472 | 8.116006 | 17.05513 | 16.08506 | 27.57973 | 3.846072 | 9.150334 | 2.239024 | 7.508942 | 19.65207 | 16.19741 |
| Lrch1 | 380916 | 2.300001 | 5.941029 | 50.19044 | 1.21E-23 | 1.29E-21 | -1.74341 | 5.941029 | -38.4909 | 3.29E-21 | 3.10E-19 | 2850.69 | 2751.132 | 2611.721 | 14627.36 | 14743.34 | 13875.67 | 4752.696 | 4426.073 | 4269.113 | 1893.806 | 1876.581 | 1982.282 | 1973.183 | 1847.294 | 2103.233 |
| Nipal2 | 223473 | 2.300758 | -2.78985 | 2.562087 | 0.018023 | 0.040632 | -0.30521 | -2.78985 | -0.40126 | 0.692214 | 0.774424 | 5.935846 | 0 | 11.2836 | 14.2858 | 19.66153 | 16.23201 | 11.37009 | 4.021265 | 44.81707 | 1.538429 | 2.287584 | 4.478048 | 18.35519 | 22.01031 | 24.29611 |
| Slc2a13 | 239606 | 2.301316 | -1.40625 | 4.382531 | 0.000252 | 0.000787 | -0.01871 | -1.40625 | -0.04867 | 0.961633 | 0.96852 | 7.419807 | 9.768385 | 11.2836 | 33.76644 | 58.9846 | 64.92805 | 52.58665 | 41.55307 | 60.90525 | 7.692145 | 1.525056 | 5.97073 | 75.92375 | 48.73712 | 30.77507 |
| Gp6 | 243816 | 2.301527 | -2.37306 | 4.21006 | 0.000382 | 0.001165 | -0.85658 | -2.37306 | -1.82855 | 0.081478 | 0.131043 | 11.12971 | 3.55214 | 5.207818 | 29.87031 | 29.4923 | 43.28537 | 19.89765 | 9.382952 | 33.32551 | 4.615287 | 3.050111 | 1.492683 | 19.18952 | 16.50774 | 10.52831 |
| Mir551b | 791072 | 2.302433 | -5.93239 | 2.627309 | 0.015616 | 0.035815 | -0.63555 | -5.93239 | -0.77763 | 0.445323 | 0.544031 | 0.741981 | 0 | 0 | 2.597418 | 4.213186 | 1.352668 | 4.263783 | 0 | 2.298311 | 0 | 0 | 0 | 0 | 0 | 0.80987 |
| Mapkbp1 | 26390 | 2.304166 | 5.383494 | 47.00504 | 4.85E-23 | 4.31E-21 | -1.07064 | 5.383494 | -24.6755 | 3.54E-17 | 1.08E-15 | 1364.503 | 1346.261 | 1340.145 | 7002.64 | 7448.912 | 6944.596 | 3638.428 | 3517.267 | 3424.484 | 1574.582 | 1524.293 | 1460.59 | 1867.224 | 1851.225 | 1904.815 |
| Dlx3 | 13393 | 2.304213 | -2.42628 | 2.856226 | 0.009358 | 0.022593 | 1.043757 | -2.42628 | 1.789976 | 0.087657 | 0.139463 | 0.741981 | 7.10428 | 3.471878 | 19.48064 | 21.06593 | 9.468674 | 29.84648 | 25.46801 | 50.56285 | 3.076858 | 6.862751 | 0 | 41.71635 | 47.95104 | 42.92312 |
| Lcn2 | 16819 | 2.30533 | 7.007447 | 56.13624 | 1.13E-24 | 1.71E-22 | -0.43441 | 7.007447 | -13.4691 | 6.61E-12 | 6.42E-11 | 5550.758 | 5680.76 | 5655.69 | 29828.75 | 28476.92 | 30784.01 | 23332.84 | 23458.72 | 21829.36 | 3233.008 | 3281.157 | 3240.614 | 2139.214 | 2215.967 | 2142.107 |
| Mir9-3hg | 101694 | 2.306097 | -2.53568 | 3.810123 | 0.000999 | 0.002874 | -0.28941 | -2.53568 | -0.59703 | 0.556772 | 0.65135 | 4.451884 | 9.768385 | 2.603909 | 18.18193 | 26.68351 | 41.9327 | 18.47639 | 14.74464 | 40.22045 | 0.769214 | 3.812639 | 0.746341 | 18.35519 | 24.36856 | 15.38754 |
| Tcim | 69068 | 2.310114 | -3.84015 | 3.317274 | 0.003224 | 0.008518 | 0.444454 | -3.84015 | 0.764885 | 0.452715 | 0.551499 | 0.741981 | 0.888035 | 1.735939 | 7.792255 | 5.617581 | 8.116006 | 9.948826 | 10.72337 | 10.3424 | 0 | 0 | 1.492683 | 11.68058 | 18.0799 | 16.19741 |
| Asz1 | 74068 | 2.310619 | -3.46977 | 3.002957 | 0.006695 | 0.016668 | 0.188943 | -3.46977 | 0.289211 | 0.775211 | 0.847838 | 0.741981 | 2.664105 | 0.86797 | 12.98709 | 5.617581 | 6.763339 | 7.106304 | 13.40422 | 9.193245 | 0 | 3.812639 | 1.492683 | 20.85817 | 22.01031 | 21.05663 |
| Parp8 | 52552 | 2.311259 | 5.883369 | 45.23847 | 1.09E-22 | 8.62E-21 | -0.22502 | 5.883369 | -5.62745 | 1.31E-05 | 4.76E-05 | 1900.213 | 1852.441 | 1928.628 | 10018.24 | 10010.53 | 10089.55 | 9262.357 | 9214.059 | 8381.941 | 1822.269 | 1863.618 | 1735.99 | 1799.643 | 1733.312 | 1806.011 |
| Gypc | 71683 | 2.312989 | 5.239236 | 38.6419 | 3.03E-21 | 1.71E-19 | 0.194227 | 5.239236 | 4.35848 | 0.000267 | 0.000807 | 1135.231 | 1141.125 | 1201.27 | 6142.894 | 6281.86 | 6018.019 | 7183.052 | 7542.553 | 7182.222 | 1185.359 | 1170.48 | 1071 | 908.582 | 1096.585 | 966.1752 |
| Slamf8 | 74748 | 2.313233 | 3.455183 | 19.2039 | 5.98E-15 | 8.56E-14 | 0.605357 | 3.455183 | 7.629992 | 1.57E-07 | 7.46E-07 | 233.7239 | 229.113 | 235.2198 | 1145.461 | 1263.956 | 1310.735 | 2082.147 | 1971.76 | 1834.052 | 1876.883 | 1873.531 | 1982.282 | 109.2968 | 85.68301 | 148.2063 |
| Snn | 20621 | 2.3142 | 2.990455 | 19.99689 | 2.63E-15 | 4.03E-14 | -2.11306 | 2.990455 | -17.0618 | 6.43E-14 | 9.28E-13 | 255.9834 | 214.0164 | 247.3713 | 1362.346 | 1308.896 | 1155.178 | 388.0042 | 261.3822 | 286.1397 | 964.5949 | 958.4975 | 991.1412 | 261.1443 | 290.0645 | 247.0104 |
| Hsh2d | 209488 | 2.314496 | 1.70705 | 10.37198 | 8.56E-10 | 5.36E-09 | 1.355476 | 1.70705 | 10.29281 | 9.83E-10 | 6.54E-09 | 79.39194 | 71.0428 | 66.83366 | 412.9895 | 367.9516 | 380.0996 | 1112.847 | 1084.401 | 921.6228 | 116.9206 | 102.1787 | 64.93169 | 102.6222 | 84.89692 | 132.0089 |
| Loxl1 | 16949 | 2.315448 | -1.2803 | 3.775188 | 0.001086 | 0.003109 | 0.685426 | -1.2803 | 1.654605 | 0.112637 | 0.172709 | 9.64575 | 7.10428 | 4.339848 | 42.8574 | 43.53625 | 28.40602 | 48.32287 | 61.6594 | 80.44089 | 14.61507 | 14.48803 | 5.97073 | 50.05962 | 77.82218 | 25.91585 |
| Ms4a4a | 666907 | 2.315473 | 5.220084 | 29.63088 | 7.93E-19 | 2.53E-17 | -1.98193 | 5.220084 | -24.5724 | 3.86E-17 | 1.17E-15 | 753.1105 | 819.6563 | 735.1703 | 4161.064 | 4265.148 | 3828.05 | 1259.237 | 1010.678 | 985.9755 | 8297.516 | 7992.054 | 8152.286 | 2040.764 | 1849.652 | 2086.226 |
| Cfd | 11537 | 2.316077 | -2.61145 | 2.755481 | 0.011743 | 0.027722 | 1.963229 | -2.61145 | 3.304304 | 0.003323 | 0.008109 | 2.967923 | 3.55214 | 0 | 15.58451 | 15.44835 | 2.705335 | 39.7953 | 36.19139 | 39.07129 | 9.230573 | 4.575167 | 4.478048 | 17.52087 | 18.86598 | 29.15533 |
| Birc3 | 11796 | 2.317027 | 6.540092 | 51.18542 | 8.00E-24 | 9.19E-22 | -0.57394 | 6.540092 | -15.5907 | 3.84E-13 | 4.65E-12 | 3285.491 | 3104.57 | 3219.299 | 17188.42 | 17720.66 | 16309.11 | 12556.84 | 12006.16 | 11270.92 | 6340.635 | 6121.574 | 6328.227 | 1340.763 | 1301.753 | 1269.067 |
| Efcab8 | 100504221 | 2.31711 | -2.46946 | 3.138367 | 0.004897 | 0.012512 | 0.01651 | -2.46946 | 0.027479 | 0.978333 | 0.981804 | 1.483961 | 5.32821 | 2.603909 | 20.77935 | 18.25714 | 10.82134 | 7.106304 | 22.78717 | 26.43058 | 19.23036 | 22.87584 | 6.717071 | 12.5149 | 7.074744 | 12.95792 |
| Dkk1 | 13380 | 2.317423 | -2.64603 | 2.658187 | 0.014586 | 0.033681 | 0.890636 | -2.64603 | 1.394837 | 0.177413 | 0.253526 | 0.741981 | 3.55214 | 6.075787 | 15.58451 | 11.23516 | 20.29002 | 31.26774 | 37.53181 | 24.13227 | 0 | 5.337695 | 7.463413 | 22.52683 | 32.22939 | 10.52831 |
| Parp12 | 243771 | 2.317587 | 8.270837 | 61.72233 | 1.51E-25 | 3.15E-23 | -0.04992 | 8.270837 | -1.84606 | 0.0788 | 0.127532 | 9472.126 | 9075.718 | 8861.102 | 48404.19 | 49660.82 | 48057.58 | 50324 | 49877.09 | 46776.38 | 12962.8 | 13171.91 | 12381.06 | 7691.66 | 7749.989 | 7576.336 |
| Gpr31b | 436440 | 2.31829 | 0.83866 | 10.72017 | 4.72E-10 | 3.07E-09 | -2.5115 | 0.83866 | -9.82224 | 2.26E-09 | 1.42E-08 | 198.1089 | 144.7497 | 74.64539 | 636.3675 | 845.4459 | 649.2805 | 96.64574 | 178.2761 | 119.5122 | 175.3809 | 154.0306 | 165.6878 | 8.343269 | 4.716496 | 8.098703 |
| Dmrt1 | 50796 | 2.318501 | -2.77091 | 3.19801 | 0.004262 | 0.011009 | 0.980535 | -2.77091 | 1.768518 | 0.091266 | 0.144332 | 2.225942 | 0.888035 | 1.735939 | 6.493546 | 9.830767 | 14.87935 | 19.89765 | 17.42548 | 25.28142 | 5.384501 | 1.525056 | 8.956095 | 26.69846 | 31.4433 | 17.81715 |
| Mip | 17339 | 2.320935 | -2.80072 | 3.766709 | 0.001108 | 0.003169 | 0.195818 | -2.80072 | 0.403689 | 0.690458 | 0.772853 | 2.225942 | 3.55214 | 2.603909 | 23.37676 | 9.830767 | 17.58468 | 18.47639 | 29.48928 | 13.78987 | 3.846072 | 5.337695 | 8.956095 | 6.674615 | 4.716496 | 9.718443 |
| Alx3 | 11694 | 2.32115 | -2.88841 | 3.408681 | 0.002599 | 0.006977 | 0.865093 | -2.88841 | 1.658057 | 0.111932 | 0.171828 | 2.967923 | 0.888035 | 1.735939 | 10.38967 | 11.23516 | 10.82134 | 15.63387 | 21.44675 | 26.43058 | 4.615287 | 1.525056 | 2.239024 | 35.04173 | 15.72165 | 19.43689 |
| Dennd6b | 69440 | 2.321158 | 3.677398 | 20.94379 | 1.03E-15 | 1.71E-14 | 0.444042 | 3.677398 | 5.82175 | 8.32E-06 | 3.12E-05 | 353.9248 | 361.4302 | 378.4347 | 2038.973 | 1815.883 | 1977.6 | 2946.274 | 2926.141 | 2440.806 | 353.8386 | 395.752 | 325.4048 | 403.8142 | 350.5928 | 367.6811 |
| Olfr1412 | 258274 | 2.322396 | -5.30027 | 2.537897 | 0.019 | 0.042606 | 1.137484 | -5.30027 | 1.460304 | 0.158771 | 0.231131 | 0 | 0 | 0 | 3.896127 | 2.808791 | 0 | 5.685043 | 6.702109 | 2.298311 | 0 | 0 | 0 | 5.840288 | 2.358248 | 4.859222 |
| Tac2 | 21334 | 2.322695 | -4.15083 | 2.506297 | 0.020353 | 0.045283 | 0.527013 | -4.15083 | 0.677849 | 0.50515 | 0.602592 | 6.677827 | 0 | 0 | 9.090964 | 4.213186 | 4.058003 | 7.106304 | 10.72337 | 8.044089 | 0.769214 | 1.525056 | 0 | 8.343269 | 10.21907 | 4.049351 |
| Rel | 19696 | 2.322835 | 6.315698 | 56.18049 | 1.11E-24 | 1.70E-22 | -0.35748 | 6.315698 | -10.9038 | 3.46E-10 | 2.47E-09 | 2349.111 | 2374.606 | 2316.611 | 12637.74 | 12593.21 | 12380.97 | 10169.12 | 10295.78 | 10021.79 | 4636.056 | 4721.572 | 4510.887 | 1822.17 | 1660.206 | 1832.736 |
| Six6 | 20476 | 2.323846 | -1.84301 | 3.064545 | 0.00581 | 0.014644 | 0.981252 | -1.84301 | 1.864018 | 0.076134 | 0.123865 | 0.741981 | 6.216245 | 13.01954 | 22.07806 | 19.66153 | 28.40602 | 29.84648 | 42.8935 | 78.14258 | 16.1535 | 3.050111 | 2.985365 | 48.39096 | 36.94588 | 42.92312 |
| Gsta1 | 14857 | 2.325287 | -5.17217 | 2.523098 | 0.019623 | 0.04383 | 0.248574 | -5.17217 | 0.300603 | 0.766626 | 0.840509 | 0 | 0 | 0 | 2.597418 | 0 | 4.058003 | 2.842522 | 1.340422 | 2.298311 | 1.538429 | 2.287584 | 1.492683 | 1.668654 | 4.716496 | 0.80987 |
| Gm19723 | 100503490 | 2.327656 | -1.98846 | 3.064199 | 0.005815 | 0.014654 | 2.097815 | -1.98846 | 4.5452 | 0.00017 | 0.000532 | 4.451884 | 9.768385 | 3.471878 | 33.76644 | 26.68351 | 29.75869 | 113.7009 | 151.4677 | 142.4953 | 10.769 | 23.63836 | 3.731706 | 12.5149 | 5.502578 | 0 |
| Gpr141 | 353346 | 2.327757 | 3.922542 | 34.65309 | 3.00E-20 | 1.34E-18 | -2.01115 | 3.922542 | -28.8238 | 1.41E-18 | 6.12E-17 | 766.4661 | 695.3314 | 729.0945 | 3896.127 | 4091.003 | 3791.528 | 935.1896 | 993.2525 | 1090.549 | 653.0631 | 703.8132 | 632.151 | 322.0502 | 355.3093 | 286.6941 |
| Olfr583 | 258752 | 2.327829 | -5.43643 | 2.552074 | 0.018422 | 0.041464 | 0.262063 | -5.43643 | 0.320893 | 0.751412 | 0.82728 | 0 | 0 | 0 | 3.896127 | 0 | 2.705335 | 5.685043 | 1.340422 | 1.149156 | 0 | 0 | 0 | 3.337308 | 5.502578 | 5.669092 |
| B3gnt2 | 53625 | 2.327971 | 7.589184 | 71.31201 | 7.02E-27 | 2.37E-24 | -1.34777 | 7.589184 | -45.2511 | 1.08E-22 | 1.84E-20 | 6713.442 | 6595.436 | 6226.814 | 35362.55 | 35911.79 | 33579.98 | 14135.86 | 14243.32 | 14336.87 | 8301.362 | 8327.567 | 7988.09 | 7668.299 | 7367.952 | 7782.853 |
| Xlr4d-ps | 667387 | 2.328557 | -4.79009 | 2.541334 | 0.018859 | 0.042321 | 1.857711 | -4.79009 | 2.579905 | 0.017333 | 0.034847 | 0 | 0 | 0.86797 | 2.597418 | 1.404395 | 4.058003 | 21.31891 | 10.72337 | 6.894934 | 0.769214 | 0 | 0 | 1.668654 | 7.860826 | 2.429611 |
| Rnd1 | 223881 | 2.330661 | 1.799855 | 12.70261 | 2.02E-11 | 1.60E-10 | 0.46213 | 1.799855 | 3.874006 | 0.000857 | 0.00238 | 83.84382 | 122.5488 | 111.1001 | 528.5746 | 514.0087 | 633.0485 | 812.9612 | 903.4443 | 699.8358 | 80.76752 | 61.00223 | 87.32193 | 114.3028 | 117.1263 | 97.9943 |
| Palmd | 114301 | 2.331163 | -3.38058 | 2.768201 | 0.011413 | 0.027018 | 0.942096 | -3.38058 | 1.472403 | 0.155506 | 0.227157 | 4.451884 | 0 | 3.471878 | 7.792255 | 18.25714 | 8.116006 | 24.16143 | 16.08506 | 25.28142 | 0 | 2.287584 | 0 | 12.5149 | 26.72681 | 13.76779 |
| Tnfrsf1b | 21938 | 2.332789 | 10.3989 | 116.0295 | 6.25E-29 | 7.15E-26 | -1.16741 | 10.3989 | -70.1685 | 1.01E-24 | 3.05E-22 | 47659.65 | 47895.28 | 48854.54 | 258979.5 | 262301.7 | 255191.6 | 122288.1 | 122202.2 | 115492.4 | 98780.21 | 98363.81 | 100508.3 | 23014.91 | 22868.72 | 23394.72 |
| Cyp4a12b | 13118 | 2.333739 | -3.03446 | 3.141406 | 0.004862 | 0.012428 | -0.02905 | -3.03446 | -0.04636 | 0.963452 | 0.969905 | 1.483961 | 4.440175 | 0.86797 | 20.77935 | 16.85274 | 4.058003 | 8.527565 | 12.0638 | 14.93902 | 5.384501 | 1.525056 | 2.985365 | 24.19548 | 16.50774 | 14.57766 |
| Plod2 | 26432 | 2.334539 | 3.470917 | 25.50043 | 1.80E-17 | 4.21E-16 | -0.68305 | 3.470917 | -9.42756 | 4.63E-09 | 2.77E-08 | 313.8579 | 276.1789 | 348.0558 | 1672.737 | 1775.156 | 1600.206 | 1094.371 | 1006.657 | 1152.603 | 118.459 | 111.3291 | 111.9512 | 2036.592 | 1916.469 | 1959.076 |
| Ndst4 | 64580 | 2.335748 | -2.67985 | 3.131158 | 0.00498 | 0.012703 | 0.587382 | -2.67985 | 1.022682 | 0.317911 | 0.413234 | 0.741981 | 4.440175 | 3.471878 | 11.68838 | 8.426372 | 27.05335 | 12.79135 | 24.12759 | 32.17636 | 2.307643 | 3.050111 | 11.94146 | 14.18356 | 18.0799 | 15.38754 |
| Fcgr4 | 246256 | 2.336906 | 7.185737 | 53.11456 | 3.65E-24 | 4.73E-22 | 0.221236 | 7.185737 | 6.971211 | 6.35E-07 | 2.77E-06 | 3678.741 | 3815.886 | 3265.302 | 19274.14 | 19908.71 | 18806.14 | 23897.08 | 23047.21 | 23230.18 | 6177.561 | 5987.369 | 6074.471 | 5195.354 | 4846.199 | 5188.029 |
| Lypd4 | 232973 | 2.337477 | -3.69244 | 2.548952 | 0.018548 | 0.041712 | 0.37021 | -3.69244 | 0.490492 | 0.628791 | 0.718864 | 2.225942 | 0 | 4.339848 | 3.896127 | 22.47032 | 8.116006 | 5.685043 | 20.10633 | 14.93902 | 0.769214 | 1.525056 | 0 | 6.674615 | 19.65207 | 7.288832 |
| Marchf5 | 69104 | 2.338693 | 6.852459 | 51.63988 | 6.63E-24 | 7.89E-22 | -0.00942 | 6.852459 | -0.2788 | 0.783079 | 0.854808 | 3141.546 | 3618.743 | 3398.101 | 18552.06 | 18554.87 | 17508.93 | 19347.62 | 18241.8 | 18749.62 | 4703.746 | 4623.206 | 4863.906 | 3042.79 | 2736.354 | 2861.272 |
| Pcdhb1 | 93872 | 2.339884 | -3.1784 | 3.122234 | 0.005084 | 0.012948 | 0.573587 | -3.1784 | 0.973469 | 0.341225 | 0.438361 | 2.225942 | 0.888035 | 2.603909 | 7.792255 | 11.23516 | 16.23201 | 11.37009 | 24.12759 | 19.53565 | 4.615287 | 1.525056 | 0 | 20.85817 | 14.93557 | 15.38754 |
| Myocos | 100038657 | 2.340406 | -4.53819 | 2.707141 | 0.013083 | 0.030555 | -0.79052 | -4.53819 | -0.98201 | 0.337098 | 0.43412 | 1.483961 | 0 | 0.86797 | 1.298709 | 8.426372 | 9.468674 | 2.842522 | 1.340422 | 4.596622 | 1.538429 | 0 | 0 | 8.343269 | 9.432991 | 8.098703 |
| Ppef1 | 237178 | 2.341869 | -2.76855 | 3.016153 | 0.006495 | 0.016213 | 0.65314 | -2.76855 | 1.099869 | 0.283641 | 0.375588 | 2.225942 | 1.77607 | 2.603909 | 14.2858 | 5.617581 | 28.40602 | 12.79135 | 29.48928 | 27.57973 | 1.538429 | 6.100223 | 0.746341 | 27.53279 | 17.29382 | 36.44416 |
| Ppil6 | 73075 | 2.343729 | -2.46511 | 3.142641 | 0.004849 | 0.012399 | 0.913412 | -2.46511 | 1.649433 | 0.113701 | 0.174065 | 1.483961 | 7.10428 | 0.86797 | 23.37676 | 18.25714 | 5.410671 | 27.00396 | 22.78717 | 31.0272 | 3.846072 | 7.625279 | 1.492683 | 33.37308 | 41.66238 | 34.82442 |
| Prdx6b | 320769 | 2.343909 | -2.34103 | 3.374326 | 0.002819 | 0.007519 | 1.882414 | -2.34103 | 3.981774 | 0.000661 | 0.001872 | 0.741981 | 5.32821 | 2.603909 | 10.38967 | 11.23516 | 21.64268 | 42.63783 | 45.57434 | 75.84427 | 4.615287 | 5.337695 | 2.985365 | 32.53875 | 21.22423 | 19.43689 |
| Stat2 | 20847 | 2.345957 | 9.027211 | 37.83107 | 3.59E-13 | 3.75E-12 | -0.47873 | 9.027211 | -10.2644 | 4.89E-07 | 2.17E-06 | 14056.08 | 14791.11 | 13914.42 | 78134.24 | 78352.62 | 75373.35 | 60096.59 | 61065.59 | 52899.08 | 29015.54 | 29269.63 | 28450.53 | 16154.24 | 16065.96 | 15799.76 |
| Rnase10 | 75019 | 2.348408 | -2.40393 | 4.964138 | 6.24E-05 | 0.000209 | -1.62451 | -2.40393 | -3.54264 | 0.001892 | 0.004876 | 6.677827 | 12.43249 | 7.811726 | 54.54578 | 32.30109 | 62.22272 | 9.948826 | 13.40422 | 26.43058 | 1.538429 | 6.100223 | 1.492683 | 7.508942 | 7.860826 | 11.33818 |
| Nfkb1 | 18033 | 2.348613 | 8.319201 | 45.68072 | 4.63E-18 | 1.26E-16 | -0.08874 | 8.319201 | -2.39998 | 0.029243 | 0.054718 | 9251.758 | 9073.053 | 9114.549 | 50487.32 | 50551.21 | 48290.24 | 50721.96 | 50717.54 | 45222.72 | 12370.51 | 12246.2 | 12854.24 | 8989.873 | 9175.156 | 9256.817 |
| Mmp14 | 17387 | 2.351746 | 7.401168 | 80.23858 | 5.72E-28 | 3.13E-25 | -2.85401 | 7.401168 | -78.3102 | 9.60E-28 | 1.85E-24 | 13126.38 | 13147.36 | 13196.61 | 72760.18 | 72382.53 | 69962.68 | 11172.53 | 10423.12 | 9497.771 | 6211.407 | 5996.519 | 6435.701 | 1752.087 | 1663.351 | 1798.722 |
| Olfr420 | 258302 | 2.351802 | -4.38072 | 2.698093 | 0.013349 | 0.031106 | 1.197451 | -4.38072 | 1.706046 | 0.102511 | 0.159356 | 2.967923 | 0 | 0 | 10.38967 | 4.213186 | 1.352668 | 8.527565 | 9.382952 | 14.93902 | 0 | 0 | 0 | 8.343269 | 13.3634 | 10.52831 |
| Zhx2 | 387609 | 2.351845 | 4.44626 | 31.16747 | 2.76E-19 | 9.85E-18 | -0.69208 | 4.44626 | -11.3078 | 1.78E-10 | 1.34E-09 | 566.8733 | 515.0603 | 632.7498 | 3125.993 | 3255.388 | 2951.521 | 2032.403 | 2108.483 | 1873.124 | 915.3652 | 875.382 | 923.2241 | 1174.732 | 1177.552 | 1156.495 |
| D16Ertd47 | 67102 | 2.352319 | 3.674378 | 26.70126 | 6.92E-18 | 1.81E-16 | -0.49118 | 3.674378 | -7.18409 | 4.01E-07 | 1.80E-06 | 447.4144 | 415.6004 | 418.3613 | 2410.404 | 2322.87 | 2271.129 | 1736.781 | 1780.08 | 1665.126 | 279.2248 | 298.1484 | 270.9219 | 577.3542 | 447.281 | 578.2474 |
| 9330154K | 319829 | 2.353563 | 1.546649 | 11.62542 | 1.07E-10 | 7.59E-10 | 0.102711 | 1.546649 | 0.75356 | 0.459347 | 0.558207 | 82.35986 | 73.7069 | 57.28599 | 354.5476 | 387.6131 | 427.443 | 491.7563 | 485.2327 | 353.9399 | 91.53652 | 81.59048 | 85.0829 | 161.0251 | 152.5 | 151.4457 |
| Cd164l2 | 69655 | 2.354823 | -3.25864 | 2.494135 | 0.020897 | 0.046333 | 0.237089 | -3.25864 | 0.306039 | 0.76254 | 0.836869 | 5.193865 | 0.888035 | 0.86797 | 14.2858 | 4.213186 | 21.64268 | 7.106304 | 9.382952 | 33.32551 | 7.692145 | 0.762528 | 0 | 15.01788 | 25.15464 | 14.57766 |
| Prr23a3 | 71979 | 2.355053 | -3.81602 | 2.777739 | 0.011171 | 0.0265 | 0.966302 | -3.81602 | 1.436788 | 0.165277 | 0.238894 | 1.483961 | 2.664105 | 0 | 2.597418 | 8.426372 | 13.52668 | 15.63387 | 9.382952 | 19.53565 | 0 | 0 | 0 | 32.53875 | 14.93557 | 26.72572 |
| Arid5b | 71371 | 2.357046 | 6.210895 | 38.41483 | 3.43E-21 | 1.92E-19 | -0.45099 | 6.210895 | -9.19769 | 7.10E-09 | 4.12E-08 | 2189.585 | 2108.195 | 1926.893 | 11280.59 | 11303.98 | 11494.97 | 8988.054 | 8944.634 | 8062.476 | 3945.301 | 4090.199 | 3563.033 | 2210.966 | 2169.588 | 2110.522 |
| Gm10099 | 791367 | 2.357112 | 2.133723 | 17.59757 | 3.47E-14 | 4.34E-13 | -1.16737 | 2.133723 | -10.2098 | 1.14E-09 | 7.49E-09 | 167.6876 | 190.0395 | 206.5768 | 1023.383 | 1068.745 | 973.9208 | 530.1303 | 485.2327 | 418.2926 | 130.7665 | 152.5056 | 94.78534 | 109.2968 | 126.5593 | 93.94495 |
| Rubcnl | 271221 | 2.357407 | 4.046456 | 27.23046 | 4.60E-18 | 1.25E-16 | -0.78044 | 4.046456 | -11.0479 | 2.73E-10 | 1.98E-09 | 513.4507 | 451.1218 | 545.0849 | 2841.576 | 2749.806 | 2664.755 | 1674.245 | 1818.952 | 1527.228 | 580.7569 | 668.7369 | 573.9364 | 696.663 | 651.6625 | 665.7134 |
| Cyp2j9 | 74519 | 2.357568 | -2.13191 | 3.300735 | 0.003351 | 0.008827 | 0.267635 | -2.13191 | 0.482512 | 0.634354 | 0.72351 | 6.677827 | 2.664105 | 0.86797 | 19.48064 | 18.25714 | 12.17401 | 12.79135 | 32.17012 | 20.6848 | 21.538 | 22.87584 | 11.19512 | 35.87606 | 15.72165 | 22.67637 |
| Gm19301 | 100502638 | 2.357619 | -3.7032 | 2.629893 | 0.015528 | 0.035627 | 0.515718 | -3.7032 | 0.706378 | 0.487593 | 0.586128 | 3.709904 | 1.77607 | 0 | 7.792255 | 8.426372 | 8.116006 | 29.84648 | 4.021265 | 14.93902 | 1.538429 | 1.525056 | 0 | 10.01192 | 12.57732 | 7.288832 |
| Olfr213 | 258020 | 2.358862 | -4.86793 | 2.514171 | 0.020008 | 0.044564 | -1.17986 | -4.86793 | -1.31279 | 0.203195 | 0.284387 | 0.741981 | 0 | 1.735939 | 2.597418 | 5.617581 | 8.116006 | 1.421261 | 0 | 10.3424 | 0 | 1.525056 | 0 | 4.171635 | 3.930413 | 2.429611 |
| Gdf10 | 14560 | 2.361553 | -2.76796 | 2.833327 | 0.009856 | 0.023705 | 0.777316 | -2.76796 | 1.221172 | 0.23534 | 0.321374 | 0 | 5.32821 | 6.075787 | 18.18193 | 9.830767 | 9.468674 | 29.84648 | 17.42548 | 20.6848 | 6.153716 | 2.287584 | 1.492683 | 17.52087 | 28.29897 | 25.10598 |
| Sos1 | 20662 | 2.362235 | 7.023543 | 74.70007 | 2.62E-27 | 9.56E-25 | -4.1515 | 7.023543 | -81.3111 | 4.31E-28 | 1.42E-24 | 7939.936 | 7878.646 | 7552.204 | 42822.34 | 43923.87 | 41679.75 | 2704.659 | 2462.355 | 2364.962 | 8796.736 | 9334.866 | 8947.139 | 3641.837 | 3707.952 | 3972.414 |
| Gm2128 | 100039269 | 2.36289 | -3.35279 | 3.447543 | 0.002371 | 0.006413 | 0.78018 | -3.35279 | 1.505573 | 0.146834 | 0.216037 | 2.225942 | 0.888035 | 3.471878 | 7.792255 | 15.44835 | 16.23201 | 25.5827 | 20.10633 | 22.98311 | 1.538429 | 0 | 0 | 9.177596 | 21.22423 | 13.76779 |
| Ralgds | 19730 | 2.362954 | 6.89967 | 43.04009 | 3.12E-22 | 2.17E-20 | 0.4191 | 6.89967 | 10.70276 | 4.86E-10 | 3.40E-09 | 2030.801 | 1908.387 | 2160.376 | 10990.98 | 11469.7 | 11037.77 | 16047.46 | 15896.06 | 14695.4 | 20690.33 | 20683.57 | 21158.03 | 2427.057 | 2474.588 | 2670.142 |
| Ankrd34c | 330998 | 2.363126 | -3.48817 | 2.494281 | 0.02089 | 0.046322 | 0.364104 | -3.48817 | 0.468766 | 0.64399 | 0.73199 | 0 | 5.32821 | 3.471878 | 18.18193 | 2.808791 | 17.58468 | 11.37009 | 8.04253 | 25.28142 | 0.769214 | 0 | 2.239024 | 14.18356 | 9.432991 | 17.00728 |
| Olfr360 | 258615 | 2.363861 | -5.06921 | 2.481841 | 0.02146 | 0.047459 | 0.801169 | -5.06921 | 0.995319 | 0.330732 | 0.427079 | 0 | 0 | 1.735939 | 1.298709 | 1.404395 | 13.52668 | 5.685043 | 4.021265 | 10.3424 | 0 | 0 | 0 | 4.171635 | 2.358248 | 0.80987 |
| Car2 | 12349 | 2.366369 | 2.52473 | 10.302 | 9.67E-10 | 6.01E-09 | 3.448224 | 2.52473 | 28.74392 | 1.49E-18 | 6.42E-17 | 92.00561 | 63.05048 | 44.26645 | 350.6515 | 410.0834 | 320.5823 | 3834.562 | 4117.776 | 4218.55 | 84.61359 | 103.7038 | 88.06827 | 497.2588 | 571.4821 | 618.7409 |
| Tead1 | 21676 | 2.36646 | 0.723918 | 8.93932 | 1.16E-08 | 6.18E-08 | 1.838626 | 0.723918 | 12.22079 | 4.19E-11 | 3.51E-10 | 24.48536 | 28.41712 | 27.77503 | 181.8193 | 134.8219 | 132.5614 | 523.024 | 522.7645 | 609.0525 | 16.1535 | 18.30067 | 12.6878 | 260.31 | 262.5516 | 267.2572 |
| Sh3bgrl2 | 212531 | 2.366768 | 5.819474 | 46.24616 | 6.85E-23 | 5.76E-21 | -1.77202 | 5.819474 | -35.2263 | 2.12E-20 | 1.57E-18 | 1864.598 | 1903.059 | 1983.311 | 10613.05 | 11209.88 | 9865.006 | 3226.262 | 3277.331 | 3121.107 | 3272.238 | 3137.04 | 3347.341 | 2181.765 | 1978.57 | 2077.317 |
| Slc27a2 | 26458 | 2.366795 | -1.75557 | 3.924412 | 0.000759 | 0.002222 | 0.274653 | -1.75557 | 0.633208 | 0.533325 | 0.628928 | 5.935846 | 2.664105 | 7.811726 | 22.07806 | 36.51428 | 32.46403 | 36.95278 | 42.8935 | 33.32551 | 23.07643 | 6.100223 | 4.478048 | 24.19548 | 43.23454 | 25.91585 |
| Gm10579 | 100038399 | 2.367911 | -2.93169 | 2.828582 | 0.009962 | 0.023932 | 0.445911 | -2.93169 | 0.674471 | 0.507252 | 0.604522 | 2.967923 | 7.10428 | 0 | 19.48064 | 5.617581 | 13.52668 | 14.21261 | 33.51054 | 10.3424 | 1.538429 | 3.050111 | 2.985365 | 20.85817 | 19.65207 | 24.29611 |
| Ier3 | 15937 | 2.36836 | 6.039609 | 30.22531 | 5.24E-19 | 1.75E-17 | 1.780736 | 6.039609 | 36.1237 | 1.25E-20 | 9.88E-19 | 871.8274 | 749.5015 | 880.9891 | 4409.118 | 4675.232 | 4732.984 | 16780.83 | 16883.95 | 15707.81 | 4508.366 | 4331.158 | 4635.526 | 3278.07 | 3169.485 | 3522.936 |
| Tmem232 | 381107 | 2.368406 | -2.80489 | 2.61574 | 0.01602 | 0.036619 | 1.54162 | -2.80489 | 2.3234 | 0.030107 | 0.056076 | 2.967923 | 2.664105 | 0 | 7.792255 | 8.426372 | 9.468674 | 29.84648 | 18.7659 | 34.47467 | 9.999788 | 9.912862 | 0.746341 | 20.85817 | 20.43815 | 24.29611 |
| Pnp | 18950 | 2.369057 | 8.447709 | 67.25457 | 2.44E-26 | 7.13E-24 | -0.45674 | 8.447709 | -17.1984 | 5.49E-14 | 8.08E-13 | 11164.58 | 11405.03 | 12024.85 | 64565.33 | 65623.18 | 60414.2 | 49631.85 | 48055.46 | 46721.22 | 12467.43 | 12853.17 | 12171.33 | 8828.013 | 9131.136 | 9036.532 |
| Rnf213 | 672511 | 2.369135 | 12.37012 | 37.50528 | 3.25E-12 | 2.90E-11 | -0.72904 | 12.37012 | -19.3522 | 2.43E-09 | 1.52E-08 | 173237.7 | 178296.1 | 167067.7 | 956611 | 970563.5 | 932423.6 | 618829.8 | 627367 | 562893.2 | 283384 | 291591.4 | 272231.7 | 113732.9 | 115880.4 | 110362.6 |
| Ighv6-4 | 628398 | 2.369831 | -4.63243 | 2.48262 | 0.021424 | 0.04739 | 0.483295 | -4.63243 | 0.591972 | 0.560089 | 0.654664 | 0.741981 | 0 | 0.86797 | 7.792255 | 1.404395 | 5.410671 | 11.37009 | 1.340422 | 12.64071 | 1.538429 | 3.050111 | 2.239024 | 0 | 2.358248 | 0.80987 |
| Spag17os | 320613 | 2.372138 | -3.58572 | 3.035397 | 0.006214 | 0.015574 | -0.67328 | -3.58572 | -0.96115 | 0.347242 | 0.444552 | 2.225942 | 3.55214 | 0.86797 | 23.37676 | 8.426372 | 9.468674 | 11.37009 | 4.021265 | 10.3424 | 3.846072 | 0 | 0 | 8.343269 | 14.93557 | 18.62702 |
| Rtn2 | 20167 | 2.374199 | 0.562948 | 10.60486 | 5.74E-10 | 3.68E-09 | -1.46679 | 0.562948 | -7.18612 | 4.00E-07 | 1.80E-06 | 36.35706 | 47.06585 | 63.36178 | 236.3651 | 272.4527 | 281.3549 | 129.3347 | 77.74446 | 95.37991 | 53.84501 | 44.98914 | 49.25852 | 92.61029 | 75.46393 | 89.8956 |
| Rab3c | 67295 | 2.374834 | -1.81642 | 3.481975 | 0.002185 | 0.005944 | 1.463958 | -1.81642 | 3.413732 | 0.002568 | 0.006442 | 7.419807 | 1.77607 | 8.679696 | 20.77935 | 35.10988 | 36.52203 | 105.1733 | 79.08488 | 79.29174 | 7.692145 | 0 | 2.239024 | 61.74019 | 49.52321 | 36.44416 |
| Hopx | 74318 | 2.37514 | 1.233694 | 9.776674 | 2.45E-09 | 1.44E-08 | 1.846777 | 1.233694 | 13.47 | 6.60E-12 | 6.42E-11 | 39.32498 | 55.05817 | 74.64539 | 306.4954 | 313.1801 | 277.2969 | 1252.131 | 1087.082 | 1038.837 | 57.69108 | 53.37695 | 32.83902 | 53.39692 | 77.82218 | 59.9304 |
| Itih4 | 16427 | 2.3764 | -2.0141 | 4.356208 | 0.000268 | 0.000835 | 0.849256 | -2.0141 | 2.244546 | 0.035516 | 0.064756 | 7.419807 | 1.77607 | 3.471878 | 22.07806 | 26.68351 | 18.93735 | 39.7953 | 37.53181 | 49.41369 | 3.076858 | 6.100223 | 5.97073 | 38.37904 | 33.01547 | 29.9652 |
| 4930579C | 319213 | 2.376771 | -3.34668 | 2.857424 | 0.009333 | 0.022542 | 0.509326 | -3.34668 | 0.783491 | 0.441947 | 0.540887 | 0.741981 | 5.32821 | 1.735939 | 14.2858 | 9.830767 | 13.52668 | 22.74017 | 14.74464 | 19.53565 | 3.846072 | 0 | 5.97073 | 2.502981 | 3.930413 | 11.33818 |
| Mettl7a2 | 393082 | 2.377211 | -3.68547 | 2.849508 | 0.009501 | 0.02292 | 0.459486 | -3.68547 | 0.672752 | 0.508324 | 0.605587 | 2.967923 | 1.77607 | 0 | 5.194837 | 12.63956 | 6.763339 | 19.89765 | 13.40422 | 5.745778 | 0.769214 | 0.762528 | 2.985365 | 5.005962 | 11.79124 | 9.718443 |
| Adgrg7 | 239853 | 2.378656 | -0.01446 | 6.624166 | 1.36E-06 | 5.53E-06 | -0.93442 | -0.01446 | -3.21464 | 0.0041 | 0.009785 | 50.45469 | 10.65642 | 29.51097 | 136.3645 | 140.4395 | 167.7308 | 62.53548 | 80.4253 | 96.52907 | 100.7671 | 77.77784 | 73.88778 | 28.36712 | 35.37372 | 22.67637 |
| Irf1 | 16362 | 2.378775 | 6.870531 | 36.5595 | 9.73E-21 | 5.00E-19 | 0.386859 | 6.870531 | 8.40315 | 3.27E-08 | 1.72E-07 | 2688.938 | 2664.993 | 2630.816 | 15075.42 | 14760.19 | 14522.24 | 21148.36 | 21233.62 | 18295.71 | 8065.983 | 8204.8 | 8594.866 | 2487.129 | 2589.356 | 2555.141 |
| Flrt3 | 71436 | 2.379802 | 2.440919 | 10.4904 | 6.98E-10 | 4.45E-09 | 1.682027 | 2.440919 | 13.10158 | 1.12E-11 | 1.05E-10 | 64.55232 | 37.29747 | 74.64539 | 362.3399 | 257.0043 | 343.5776 | 1020.465 | 1077.699 | 1084.803 | 304.6089 | 328.6495 | 341.8243 | 587.3662 | 544.7553 | 599.304 |
| Ces1h | 75704 | 2.379861 | -3.48918 | 2.82811 | 0.009972 | 0.023949 | 0.154383 | -3.48918 | 0.214594 | 0.832121 | 0.897614 | 0 | 6.216245 | 0.86797 | 12.98709 | 7.021976 | 4.058003 | 5.685043 | 13.40422 | 8.044089 | 2.307643 | 2.287584 | 0.746341 | 19.18952 | 19.65207 | 19.43689 |
| Vmn2r110 | 224582 | 2.38018 | -3.75274 | 2.8404 | 0.009699 | 0.023363 | 1.861802 | -3.75274 | 3.010093 | 0.006587 | 0.01492 | 0 | 3.55214 | 0.86797 | 6.493546 | 8.426372 | 4.058003 | 22.74017 | 22.78717 | 28.72889 | 0.769214 | 2.287584 | 0 | 5.005962 | 8.646909 | 8.908573 |
| Zeb1 | 21417 | 2.380217 | 4.016034 | 32.08604 | 1.50E-19 | 5.71E-18 | -0.93762 | 4.016034 | -15.082 | 7.37E-13 | 8.44E-12 | 625.4898 | 588.7672 | 513.838 | 3289.63 | 3119.162 | 3204.47 | 1696.985 | 1749.25 | 1758.208 | 495.3741 | 520.8065 | 450.0438 | 518.9513 | 526.6754 | 582.2967 |
| Lgals7 | 16858 | 2.382043 | -2.95564 | 3.402366 | 0.002638 | 0.007072 | -0.30821 | -2.95564 | -0.51136 | 0.614347 | 0.705864 | 2.225942 | 3.55214 | 0.86797 | 16.88322 | 7.021976 | 16.23201 | 11.37009 | 5.361687 | 17.23733 | 8.461359 | 10.67539 | 3.731706 | 13.34923 | 7.860826 | 10.52831 |
| Tcte1 | 21645 | 2.382118 | -2.71993 | 2.603717 | 0.01645 | 0.037448 | 1.716933 | -2.71993 | 2.607911 | 0.016299 | 0.033051 | 0 | 11.54445 | 0.86797 | 7.792255 | 9.830767 | 9.468674 | 35.53152 | 32.17012 | 29.87805 | 6.153716 | 4.575167 | 2.239024 | 14.18356 | 38.51805 | 25.10598 |
| Mapkapk2 | 17164 | 2.383665 | 8.758834 | 39.29056 | 3.11E-13 | 3.28E-12 | -0.28242 | 8.758834 | -6.52484 | 4.17E-05 | 0.000142 | 15447.3 | 16272.35 | 15502.81 | 88691.45 | 88840.64 | 85260 | 79306.36 | 77200.25 | 69427.38 | 15667.36 | 15005.02 | 15932.89 | 7191.064 | 7443.416 | 7065.308 |
| Cyp2b19 | 13090 | 2.384885 | -2.84215 | 2.713103 | 0.01291 | 0.030202 | 0.957845 | -2.84215 | 1.443249 | 0.163468 | 0.236667 | 6.677827 | 0 | 1.735939 | 22.07806 | 2.808791 | 14.87935 | 25.5827 | 26.80843 | 14.93902 | 3.076858 | 3.050111 | 5.97073 | 16.68654 | 22.7964 | 22.67637 |
| Mir342 | 723909 | 2.384919 | -0.17859 | 8.13148 | 5.62E-08 | 2.74E-07 | -0.37119 | -0.17859 | -1.72828 | 0.09838 | 0.153734 | 28.93725 | 19.53677 | 20.83127 | 122.0787 | 115.1604 | 156.9095 | 135.0198 | 85.78699 | 99.97654 | 37.69151 | 44.22662 | 35.07804 | 30.03577 | 32.22939 | 34.82442 |
| Epyc | 13516 | 2.387246 | -4.46989 | 2.46251 | 0.022375 | 0.049246 | 0.498078 | -4.46989 | 0.593359 | 0.559177 | 0.653757 | 0.741981 | 0 | 0 | 2.597418 | 2.808791 | 2.705335 | 2.842522 | 1.340422 | 14.93902 | 0.769214 | 3.812639 | 1.492683 | 12.5149 | 7.074744 | 3.239481 |
| Scara5 | 71145 | 2.388312 | -2.30817 | 3.07483 | 0.005674 | 0.014334 | 1.633637 | -2.30817 | 3.051583 | 0.005986 | 0.013707 | 0.741981 | 1.77607 | 6.943757 | 16.88322 | 12.63956 | 12.17401 | 41.21656 | 45.57434 | 50.56285 | 12.30743 | 0.762528 | 6.717071 | 34.2074 | 30.65722 | 27.53559 |
| Paqr9 | 75552 | 2.388654 | -1.81486 | 3.761876 | 0.001121 | 0.003202 | -0.44293 | -1.81486 | -0.90577 | 0.37517 | 0.473436 | 10.38773 | 5.32821 | 11.2836 | 62.33804 | 67.41097 | 29.75869 | 25.5827 | 36.19139 | 55.15947 | 10.769 | 0 | 4.478048 | 19.18952 | 39.30413 | 34.01455 |
| Serpinb6d | 238568 | 2.389374 | -4.19256 | 2.955952 | 0.007457 | 0.018378 | 0.267997 | -4.19256 | 0.388087 | 0.701796 | 0.783078 | 2.225942 | 0.888035 | 0 | 5.194837 | 7.021976 | 5.410671 | 5.685043 | 6.702109 | 10.3424 | 0 | 0.762528 | 2.239024 | 6.674615 | 7.860826 | 3.239481 |
| Trim30d | 209387 | 2.390017 | 6.403718 | 65.75545 | 3.94E-26 | 1.05E-23 | -1.91648 | 6.403718 | -52.5342 | 4.61E-24 | 1.19E-21 | 5354.133 | 5504.929 | 5004.713 | 29130.05 | 30545.6 | 29128.35 | 8200.675 | 8392.381 | 7852.18 | 8219.056 | 8430.508 | 7944.803 | 312.8726 | 317.5774 | 322.3284 |
| Psd | 73728 | 2.390827 | 3.712369 | 21.68157 | 5.06E-16 | 8.91E-15 | 0.876324 | 3.712369 | 12.45101 | 2.95E-11 | 2.54E-10 | 306.438 | 254.866 | 281.2222 | 1620.789 | 1490.063 | 1624.554 | 3322.908 | 2916.758 | 2837.265 | 619.9868 | 683.225 | 682.1559 | 343.7427 | 309.7166 | 349.0541 |
| Cyp2d40 | 71754 | 2.391098 | -4.14055 | 2.506082 | 0.020362 | 0.045293 | 1.229414 | -4.14055 | 1.607605 | 0.122616 | 0.185478 | 0.741981 | 0.888035 | 0 | 2.597418 | 9.830767 | 2.705335 | 7.106304 | 13.40422 | 13.78987 | 0.769214 | 0 | 2.985365 | 16.68654 | 3.14433 | 8.098703 |
| Tnfrsf9 | 21942 | 2.392789 | -0.09495 | 8.77783 | 1.58E-08 | 8.25E-08 | -0.57856 | -0.09495 | -2.79999 | 0.010626 | 0.022773 | 37.84102 | 25.75301 | 33.85081 | 171.4296 | 172.7406 | 208.3108 | 137.8623 | 92.4891 | 156.2852 | 23.84565 | 16.77561 | 32.09267 | 35.87606 | 27.51289 | 27.53559 |
| Cdx1 | 12590 | 2.394509 | -3.13795 | 2.879347 | 0.00888 | 0.021549 | 1.007801 | -3.13795 | 1.560337 | 0.133386 | 0.199216 | 0 | 2.664105 | 3.471878 | 3.896127 | 9.830767 | 14.87935 | 9.948826 | 18.7659 | 28.72889 | 3.076858 | 2.287584 | 3.731706 | 17.52087 | 13.3634 | 15.38754 |
| Prkce | 18754 | 2.39581 | 2.340354 | 14.22087 | 2.32E-12 | 2.11E-11 | -0.00263 | 2.340354 | -0.02306 | 0.981817 | 0.984759 | 108.3292 | 102.124 | 93.74072 | 537.6656 | 575.8021 | 607.3478 | 686.469 | 636.7003 | 495.2861 | 156.1505 | 160.8934 | 169.4195 | 427.1754 | 442.5645 | 469.7248 |
| Pla2r1 | 18779 | 2.396482 | 2.264083 | 21.10472 | 8.78E-16 | 1.48E-14 | -2.1212 | 2.264083 | -17.6362 | 3.32E-14 | 5.13E-13 | 169.9136 | 200.6959 | 190.0853 | 1045.461 | 1074.362 | 1014.501 | 250.1419 | 238.5951 | 256.2617 | 333.0699 | 377.4513 | 333.6145 | 110.1312 | 97.47424 | 141.7273 |
| Gm2427 | 100039794 | 2.396639 | -0.04139 | 7.052589 | 5.32E-07 | 2.28E-06 | -0.02819 | -0.04139 | -0.11935 | 0.906111 | 0.925711 | 22.25942 | 21.31284 | 74.64539 | 177.9232 | 200.8285 | 174.4941 | 171.9726 | 170.2336 | 217.1904 | 24.61486 | 25.16342 | 25.3756 | 43.385 | 12.57732 | 20.24676 |
| Ephx4 | 384214 | 2.39888 | -3.57083 | 2.595869 | 0.016736 | 0.038008 | 1.751718 | -3.57083 | 2.513011 | 0.020058 | 0.039573 | 0 | 0.888035 | 1.735939 | 2.597418 | 4.213186 | 10.82134 | 27.00396 | 13.40422 | 19.53565 | 3.846072 | 2.287584 | 0 | 15.01788 | 19.65207 | 16.19741 |
| Hoxb9 | 15417 | 2.398963 | -2.54508 | 3.146519 | 0.004805 | 0.012295 | 1.241511 | -2.54508 | 2.285994 | 0.032571 | 0.060011 | 1.483961 | 3.55214 | 1.735939 | 5.194837 | 26.68351 | 17.58468 | 32.689 | 37.53181 | 32.17636 | 8.461359 | 0.762528 | 0.746341 | 36.71038 | 45.59279 | 37.25403 |

| | | | | | | | | | | | | | | | | | | | | | | | | | | |
|---|---|---|---|---|---|---|---|---|---|---|---|---|---|---|---|---|---|---|---|---|---|---|---|---|---|---|
| Hpx | 15458 | 2.399795 | -2.1444 | 2.783435 | 0.011029 | 0.026201 | 0.581087 | -2.1444 | 0.932836 | 0.361338 | 0.459153 | 9.64575 | 0 | 9.547666 | 11.68838 | 21.06593 | 32.46403 | 27.00396 | 33.51054 | 33.32551 | 19.99958 | 5.337695 | 14.18048 | 9.177596 | 19.65207 | 21.8665 |
| Kirrel3os | 442804 | 2.400954 | -1.8933 | 2.715546 | 0.01284 | 0.03006 | -0.06101 | -1.8933 | -0.08645 | 0.931916 | 0.944266 | 0 | 15.98463 | 9.547666 | 32.46773 | 18.25714 | 13.52668 | 18.47639 | 12.0638 | 33.32551 | 11.53822 | 18.30067 | 7.463413 | 50.89394 | 55.81187 | 55.07118 |
| Olfr769 | 257667 | 2.402129 | -4.56904 | 2.682356 | 0.013824 | 0.032095 | -0.99264 | -4.56904 | -1.18203 | 0.250197 | 0.338203 | 1.483961 | 0.888035 | 0.86797 | 9.090964 | 2.808791 | 16.23201 | 0 | 8.04253 | 11.49156 | 0 | 0 | 0 | 4.171635 | 6.288661 | 4.049351 |
| Gm11725 | 100503754 | 2.402816 | -0.52771 | 7.212883 | 3.77E-07 | 1.65E-06 | -3.1709 | -0.52771 | -6.9342 | 6.88E-07 | 2.99E-06 | 28.19527 | 31.96926 | 29.51097 | 174.027 | 185.3802 | 151.4988 | 4.263783 | 45.57434 | 28.72889 | 22.30722 | 37.36386 | 32.09267 | 40.04769 | 36.94588 | 46.97248 |
| Kcnj12 | 16515 | 2.402928 | -2.41812 | 2.97806 | 0.007089 | 0.017558 | 0.504472 | -2.41812 | 0.82693 | 0.417427 | 0.516719 | 1.483961 | 4.440175 | 2.603909 | 5.194837 | 33.70549 | 24.34802 | 15.63387 | 24.12759 | 34.47467 | 3.076858 | 12.96297 | 2.239024 | 30.03577 | 36.94588 | 19.43689 |
| Lgals12 | 56072 | 2.403738 | -2.58375 | 2.864141 | 0.009192 | 0.022232 | 1.4687 | -2.58375 | 2.58264 | 0.017229 | 0.034673 | 1.483961 | 14.20856 | 0.86797 | 12.98709 | 15.44835 | 27.05335 | 38.37404 | 40.21265 | 81.59005 | 3.846072 | 0 | 2.985365 | 22.52683 | 21.22423 | 12.14805 |
| Gk | 14933 | 2.404681 | 6.732141 | 79.76225 | 6.49E-28 | 3.21E-25 | -0.99433 | 6.732141 | -38.8916 | 2.65E-21 | 2.58E-19 | 4819.907 | 5149.715 | 5135.776 | 28479.39 | 28825.21 | 27920.41 | 14948.82 | 14511.41 | 14919.49 | 3959.916 | 4000.984 | 4140.701 | 1161.383 | 1249.871 | 1233.432 |
| Gm15821 | 100502931 | 2.405817 | -2.27473 | 3.973381 | 0.000675 | 0.001992 | 0.096254 | -2.27473 | 0.220241 | 0.827777 | 0.893917 | 2.967923 | 8.88035 | 8.679696 | 31.16902 | 49.15383 | 29.75869 | 38.37404 | 72.38277 | 24.13227 | 3.846072 | 6.100223 | 5.224389 | 8.343269 | 8.646909 | 1.619741 |
| 1700016C | 69428 | 2.406158 | -3.66175 | 2.885989 | 0.008747 | 0.02126 | 0.337303 | -3.66175 | 0.491204 | 0.628295 | 0.718378 | 5.935846 | 0.888035 | 0 | 10.38967 | 5.617581 | 9.468674 | 14.21261 | 10.72337 | 9.193245 | 1.538429 | 0.762528 | 4.478048 | 3.337308 | 7.074744 | 9.718443 |
| Cbln1 | 12404 | 2.406606 | -2.84367 | 3.058371 | 0.005893 | 0.01484 | 1.677841 | -2.84367 | 2.960487 | 0.00738 | 0.016517 | 2.967923 | 2.664105 | 0 | 11.68838 | 8.426372 | 6.763339 | 25.5827 | 30.8297 | 35.62382 | 6.153716 | 3.050111 | 1.492683 | 26.69846 | 16.50774 | 29.15533 |
| Spic | 20728 | 2.407958 | 0.525399 | 13.99037 | 3.18E-12 | 2.84E-11 | -4.65863 | 0.525399 | -12.791 | 1.77E-11 | 1.59E-10 | 120.2009 | 92.35564 | 105.8923 | 594.8088 | 636.1911 | 584.3525 | 28.42522 | 9.382952 | 47.11538 | 322.3009 | 305.0111 | 284.356 | 9.177596 | 14.93557 | 6.478962 |
| Stra8 | 20899 | 2.409632 | -3.51252 | 2.744995 | 0.012022 | 0.028325 | 1.716068 | -3.51252 | 2.581346 | 0.017278 | 0.034756 | 0 | 0.888035 | 1.735939 | 1.298709 | 15.44835 | 5.410671 | 14.21261 | 17.42548 | 25.28142 | 1.538429 | 2.287584 | 3.731706 | 15.85221 | 11.00516 | 10.52831 |
| Mir501 | 751560 | 2.409939 | -2.56846 | 3.340345 | 0.003053 | 0.008095 | -0.07586 | -2.56846 | -0.12822 | 0.899176 | 0.9219 | 5.935846 | 1.77607 | 0.86797 | 10.38967 | 9.830767 | 29.75869 | 11.37009 | 13.40422 | 18.38649 | 19.99958 | 10.67539 | 5.97073 | 12.5149 | 18.0799 | 15.38754 |
| Ctxn2 | 381418 | 2.41027 | -4.5893 | 2.604482 | 0.016422 | 0.037395 | 0.337043 | -4.5893 | 0.417805 | 0.680263 | 0.763782 | 0 | 0 | 0.86797 | 2.597418 | 2.808791 | 2.705335 | 2.842522 | 8.04253 | 2.298311 | 0.769214 | 4.575167 | 1.492683 | 7.508942 | 1.572165 | 6.478962 |
| Trav18 | 547427 | 2.41027 | -5.47022 | 2.588513 | 0.017009 | 0.038571 | -0.82025 | -5.47022 | -0.93793 | 0.358775 | 0.456492 | 0 | 0 | 0.86797 | 2.597418 | 2.808791 | 2.705335 | 1.421261 | 0 | 4.596622 | 0 | 0 | 0 | 7.508942 | 2.358248 | 0.80987 |
| Mup10 | 100039008 | 2.4109 | -4.80906 | 2.777646 | 0.011173 | 0.026503 | 0.442305 | -4.80906 | 0.595126 | 0.558017 | 0.652575 | 2.225942 | 0 | 0 | 3.896127 | 2.808791 | 5.410671 | 5.685043 | 2.680843 | 11.49156 | 1.538429 | 0.762528 | 0 | 1.668654 | 1.572165 | 1.619741 |
| Carhsp1 | 52502 | 2.412075 | 6.772303 | 56.93703 | 8.37E-25 | 1.31E-22 | 0.500641 | 6.772303 | 17.02674 | 6.70E-14 | 9.64E-13 | 2668.905 | 2569.973 | 2539.679 | 14741.65 | 15143.59 | 14385.62 | 22103.45 | 22012.41 | 21004.27 | 3809.919 | 3876.692 | 3913.067 | 3625.985 | 3696.947 | 3808.82 |
| Car13 | 71934 | 2.412743 | 7.099424 | 68.74473 | 1.53E-26 | 4.79E-24 | 0.641658 | 7.099424 | 27.12445 | 4.99E-18 | 1.84E-16 | 3479.148 | 3601.87 | 3595.998 | 20487.14 | 20230.31 | 19923.44 | 31850.46 | 32419.44 | 33747.25 | 6669.859 | 6480.724 | 6925.3 | 2321.097 | 2370.825 | 2369.68 |
| Gm5095 | 328953 | 2.416308 | -4.14242 | 2.540634 | 0.018887 | 0.042382 | 1.372305 | -4.14242 | 1.817526 | 0.083204 | 0.133354 | 2.225942 | 0 | 0 | 9.090964 | 4.213186 | 1.352668 | 9.948826 | 6.702109 | 21.83396 | 1.538429 | 0 | 0.746341 | 7.508942 | 16.50774 | 8.908573 |
| Gpr161 | 240888 | 2.416414 | -2.56467 | 3.592138 | 0.001682 | 0.004657 | -0.61055 | -2.56467 | -1.06492 | 0.298813 | 0.39262 | 0.741981 | 3.55214 | 13.01954 | 22.07806 | 30.8967 | 13.52668 | 9.948826 | 13.40422 | 19.53565 | 7.692145 | 7.625279 | 8.209754 | 8.343269 | 7.860826 | 15.38754 |
| Mdfic | 16543 | 2.41831 | 7.636669 | 84.92134 | 1.71E-28 | 1.34E-25 | -2.46376 | 7.636669 | -77.3971 | 1.23E-27 | 1.91E-24 | 7952.55 | 8068.686 | 8559.916 | 46798.98 | 48270.47 | 45089.83 | 8607.156 | 8849.464 | 8874.929 | 11135.15 | 10676.91 | 10971.22 | 6409.299 | 6489.112 | 6581.006 |
| Bcl2a1c | 12046 | 2.42023 | -3.65654 | 2.76486 | 0.011499 | 0.027204 | 1.027297 | -3.65654 | 1.590035 | 0.126531 | 0.190427 | 2.967923 | 0 | 6.075787 | 10.38967 | 22.47032 | 6.763339 | 19.89765 | 36.19139 | 22.98311 | 1.538429 | 1.525056 | 0 | 3.337308 | 0.786083 | 6.478962 |
| Mhrt | 791403 | 2.421776 | -4.67047 | 2.506948 | 0.020324 | 0.045227 | 0.829547 | -4.67047 | 1.016525 | 0.320765 | 0.416327 | 0.741981 | 0 | 0 | 1.298709 | 1.404395 | 9.468674 | 2.842522 | 4.021265 | 13.78987 | 0 | 0.762528 | 1.492683 | 5.840288 | 8.646909 | 3.239481 |
| Clrn3 | 212070 | 2.422108 | -3.37731 | 2.459492 | 0.022521 | 0.049522 | 0.558953 | -3.37731 | 0.728667 | 0.474126 | 0.572648 | 9.64575 | 0 | 1.735939 | 11.68838 | 7.021976 | 20.29002 | 7.106304 | 20.10633 | 37.92213 | 4.615287 | 0 | 0 | 15.85221 | 9.432991 | 17.81715 |
| 4933409D | 71084 | 2.422933 | -3.47276 | 3.291994 | 0.003421 | 0.008994 | -0.317 | -3.47276 | -0.50603 | 0.618023 | 0.708945 | 2.225942 | 1.77607 | 2.603909 | 9.090964 | 29.4923 | 10.82134 | 5.685043 | 10.72337 | 22.98311 | 2.307643 | 1.525056 | 0 | 5.005962 | 11.00516 | 7.288832 |
| Gm19351 | 100502750 | 2.423984 | -3.50516 | 2.70401 | 0.013174 | 0.030748 | 0.61744 | -3.50516 | 0.877671 | 0.389895 | 0.488758 | 5.935846 | 1.77607 | 0 | 20.77935 | 5.617581 | 8.116006 | 17.05513 | 16.08506 | 16.08818 | 3.076858 | 0 | 0 | 11.68058 | 25.94073 | 11.33818 |
| Thbs1 | 21825 | 2.425352 | 9.380016 | 24.96944 | 1.17E-10 | 8.23E-10 | 0.821151 | 9.380016 | 13.51869 | 5.90E-08 | 2.99E-07 | 8041.587 | 8070.462 | 7591.262 | 45710.67 | 45194.84 | 45065.48 | 86594.58 | 83797.81 | 79955.95 | 26701.74 | 28362.22 | 24962.13 | 125139 | 121544.9 | 131467.1 |
| Lgr5 | 14160 | 2.426219 | -1.546 | 3.590562 | 0.001688 | 0.004673 | 2.407319 | -1.546 | 5.738144 | 1.01E-05 | 3.74E-05 | 7.419807 | 1.77607 | 1.735939 | 20.77935 | 11.23516 | 28.40602 | 90.96069 | 138.0634 | 98.82738 | 6.92293 | 12.20045 | 4.478048 | 65.91183 | 78.60826 | 55.88105 |
| Oxct2a | 64059 | 2.426574 | -2.71889 | 2.689404 | 0.013609 | 0.031671 | 1.239387 | -2.71889 | 1.913955 | 0.069131 | 0.114101 | 10.38773 | 1.77607 | 0 | 15.58451 | 21.06593 | 5.410671 | 29.84648 | 29.48928 | 33.32551 | 0.769214 | 1.525056 | 5.97073 | 40.88202 | 18.86598 | 29.9652 |
| Icam1 | 15894 | 2.427519 | 7.86909 | 60.69871 | 2.15E-25 | 4.29E-23 | -0.27995 | 7.86909 | -9.52146 | 3.90E-09 | 2.36E-08 | 7654.273 | 8019.844 | 7636.397 | 44605.46 | 45529.09 | 43662.76 | 39673.08 | 38478.15 | 36587.96 | 8500.589 | 8586.826 | 9100.139 | 4736.474 | 4240.916 | 4885.137 |
| Parp10 | 671535 | 2.427885 | 6.876258 | 34.57376 | 1.55E-17 | 3.69E-16 | 0.100826 | 6.876258 | 1.990618 | 0.062401 | 0.104658 | 3293.653 | 3426.039 | 3135.974 | 19217 | 18696.71 | 18662.76 | 21985.48 | 22535.17 | 19003.59 | 3933.763 | 4175.603 | 4083.979 | 3263.053 | 3091.663 | 3288.883 |
| Vmn2r83 | 625029 | 2.428708 | -3.98988 | 2.657044 | 0.014623 | 0.033763 | 1.250444 | -3.98988 | 1.74658 | 0.095088 | 0.14937 | 0.741981 | 0 | 1.735939 | 10.38967 | 8.426372 | 1.352668 | 18.47639 | 10.72337 | 13.78987 | 0 | 2.287584 | 0 | 13.34923 | 7.074744 | 15.38754 |
| Olfr374 | 258335 | 2.429015 | -4.75336 | 2.759176 | 0.011646 | 0.027508 | 1.408168 | -4.75336 | 1.983024 | 0.06038 | 0.101825 | 0.741981 | 0 | 0 | 5.194837 | 2.808791 | 1.352668 | 7.106304 | 10.72337 | 9.193245 | 0 | 0 | 0.746341 | 7.508942 | 7.074744 | 1.619741 |
| Sult2a8 | 76971 | 2.429015 | -4.72937 | 2.728968 | 0.012461 | 0.029251 | 0.946585 | -4.72937 | 1.27599 | 0.215671 | 0.298703 | 0.741981 | 0 | 0 | 5.194837 | 2.808791 | 1.352668 | 11.37009 | 6.702109 | 3.447467 | 0.769214 | 0 | 0 | 3.337308 | 9.432991 | 9.718443 |
| Cacng6 | 54378 | 2.432641 | -4.07566 | 2.531908 | 0.01925 | 0.0431 | 2.338779 | -4.07566 | 3.130861 | 0.004983 | 0.011648 | 0 | 0 | 0 | 1.298709 | 9.830767 | 0 | 12.79135 | 9.382952 | 13.78987 | 4.615287 | 5.337695 | 2.985365 | 6.674615 | 10.21907 | 12.14805 |
| Wfdc18 | 14038 | 2.435582 | -1.51804 | 5.773897 | 9.30E-06 | 3.45E-05 | -2.13391 | -1.51804 | -4.82682 | 8.66E-05 | 0.000283 | 16.32358 | 11.54445 | 19.9633 | 88.31222 | 115.1604 | 77.10206 | 15.63387 | 20.10633 | 28.72889 | 24.61486 | 12.96297 | 13.43414 | 1.668654 | 11.00516 | 17.00728 |
| Klre1 | 243655 | 2.436442 | -4.46061 | 2.600368 | 0.016571 | 0.037693 | 2.747682 | -4.46061 | 3.892451 | 0.00082 | 0.002285 | 0 | 0 | 0 | 1.298709 | 2.808791 | 1.352668 | 25.5827 | 9.382952 | 17.23733 | 1.538429 | 0 | 2.985365 | 5.005962 | 6.288661 | 7.288832 |
| C3 | 12266 | 2.43702 | 9.215884 | 27.43123 | 4.95E-11 | 3.68E-10 | 1.558706 | 9.215884 | 31.34044 | 1.26E-11 | 1.16E-10 | 11054.03 | 10921.94 | 10245.51 | 62104.27 | 61880.46 | 62440.5 | 197059.2 | 195347.7 | 180911.6 | 53485.79 | 53738.39 | 54614.27 | 8081.291 | 7831.741 | 8889.946 |
| Tnnt2 | 21956 | 2.439487 | -0.67097 | 6.596075 | 1.44E-06 | 5.87E-06 | -0.23727 | -0.67097 | -0.91026 | 0.372851 | 0.471006 | 34.8731 | 24.86498 | 11.2836 | 120.78 | 136.2263 | 129.8561 | 156.3387 | 92.4891 | 103.424 | 6.153716 | 16.01308 | 5.224389 | 25.02981 | 19.65207 | 38.87377 |
| Kcnj16 | 16517 | 2.439706 | -2.85877 | 3.047319 | 0.006046 | 0.015191 | 0.744839 | -2.85877 | 1.232258 | 0.231256 | 0.316686 | 5.935846 | 0 | 3.471878 | 20.77935 | 8.426372 | 10.82134 | 14.21261 | 25.46801 | 28.72889 | 4.615287 | 4.575167 | 1.492683 | 14.18356 | 21.22423 | 12.95792 |
| 4930599N | 75379 | 2.439731 | 2.828842 | 14.88781 | 9.50E-13 | 9.23E-12 | 0.788617 | 2.828842 | 7.789794 | 1.13E-07 | 5.48E-07 | 171.3976 | 150.0779 | 173.5939 | 985.7203 | 1013.973 | 880.5867 | 1733.938 | 1991.867 | 1498.499 | 221.5338 | 256.9719 | 250.0243 | 253.6354 | 183.9433 | 191.1294 |
| 4930579D | 75934 | 2.439968 | -4.11811 | 2.592582 | 0.016857 | 0.038264 | 0.393387 | -4.11811 | 0.492061 | 0.627699 | 0.717913 | 0 | 0.888035 | 1.735939 | 11.68838 | 1.404395 | 6.763339 | 7.106304 | 6.702109 | 8.044089 | 4.615287 | 0 | 1.492683 | 8.343269 | 4.716496 | 6.478962 |
| Slc25a22 | 68267 | 2.440032 | 6.736708 | 69.90437 | 1.07E-26 | 3.48E-24 | -1.71369 | 6.736708 | -51.3084 | 7.60E-24 | 1.89E-21 | 5000.208 | 5126.626 | 4972.598 | 28840.43 | 29506.34 | 29132.41 | 9620.515 | 9503.59 | 8669.23 | 4792.975 | 4681.158 | 4622.838 | 1655.305 | 1641.341 | 1650.516 |
| Gm3286 | 100041354 | 2.442011 | -5.95735 | 2.495204 | 0.020848 | 0.046237 | 0.574637 | -5.95735 | 0.67268 | 0.508369 | 0.605602 | 0 | 0 | 0 | 1.298709 | 0 | 9.468674 | 5.685043 | 1.340422 | 3.447467 | 0 | 0 | 0 | 0 | 0 | 0.80987 |
| Olfr287 | 634104 | 2.442451 | -3.99681 | 2.536384 | 0.019063 | 0.042729 | 0.924489 | -3.99681 | 1.209215 | 0.239805 | 0.326472 | 7.419807 | 0 | 0 | 14.2858 | 2.808791 | 5.410671 | 14.21261 | 12.0638 | 12.64071 | 3.076858 | 0 | 0.746341 | 5.840288 | 5.502578 | 5.669092 |
| Apoa5 | 66113 | 2.443371 | -2.91745 | 2.835057 | 0.009817 | 0.023621 | 1.005156 | -2.91745 | 1.519843 | 0.143227 | 0.211496 | 0 | 5.32821 | 1.735939 | 15.58451 | 4.213186 | 9.468674 | 21.31891 | 18.7659 | 17.23733 | 3.846072 | 10.67539 | 2.985365 | 15.01788 | 26.72681 | 15.38754 |
| Vmn2r7 | 319217 | 2.443658 | -3.81365 | 3.142259 | 0.004853 | 0.012406 | 1.185947 | -3.81365 | 1.973732 | 0.061497 | 0.10341 | 2.967923 | 0.888035 | 0 | 7.792255 | 7.021976 | 5.410671 | 14.21261 | 12.0638 | 25.28142 | 0.769214 | 0.762528 | 0 | 10.84625 | 13.3634 | 8.908573 |
| Olfr695 | 258591 | 2.4448 | -5.22682 | 2.668973 | 0.014241 | 0.032972 | -0.14635 | -5.22682 | -0.1778 | 0.860559 | 0.918721 | 0 | 0 | 0.86797 | 1.298709 | 1.404395 | 9.468674 | 1.421261 | 1.340422 | 6.894934 | 0 | 0 | 0 | 6.674615 | 2.358248 | 3.239481 |
| Traj38 | 100124351 | 2.444893 | -5.78578 | 2.546869 | 0.018632 | 0.04188 | -0.64828 | -5.78578 | -0.72055 | 0.479004 | 0.577452 | 0 | 0 | 0 | 5.194837 | 2.808791 | 0 | 2.842522 | 0 | 1.149156 | 0 | 0.762528 | 0 | 1.668654 | 0.786083 | 1.619741 |
| Olfr64 | 18366 | 2.445941 | -4.0305 | 2.605811 | 0.016374 | 0.037315 | 2.697584 | -4.0305 | 3.807575 | 0.001005 | 0.002749 | 0 | 0 | 0 | 2.597418 | 0 | 5.410671 | 14.21261 | 24.12759 | 11.49156 | 3.846072 | 4.575167 | 1.492683 | 11.68058 | 11.00516 | 11.33818 |
| Gm10688 | 100038652 | 2.445941 | -5.26869 | 2.763204 | 0.011541 | 0.027285 | 1.311108 | -5.26869 | 1.774298 | 0.090282 | 0.143068 | 0 | 0 | 0 | 2.597418 | 0 | 5.410671 | 5.685043 | 6.702109 | 4.596622 | 0 | 0 | 0 | 2.502981 | 4.716496 | 3.239481 |
| Pkdrej | 18766 | 2.447578 | -1.76098 | 3.66018 | 0.00143 | 0.00401 | -0.06922 | -1.76098 | -0.13834 | 0.891272 | 0.918721 | 5.935846 | 6.216245 | 3.471878 | 28.5716 | 19.66153 | 54.10671 | 35.53152 | 13.40422 | 56.30862 | 10.769 | 13.7255 | 2.985365 | 40.04769 | 38.51805 | 38.0639 |
| Plekhg1 | 213783 | 2.44852 | 4.670304 | 36.84772 | 8.25E-21 | 4.29E-19 | -2.73324 | 4.670304 | -34.1765 | 4.01E-20 | 2.75E-18 | 608.4242 | 675.7946 | 644.0334 | 3700.022 | 3808.72 | 3696.841 | 537.2366 | 612.5727 | 595.2626 | 3263.008 | 3409.262 | 3297.336 | 1757.927 | 1670.426 | 1858.652 |
| Etnk1 | 75320 | 2.449918 | 8.098125 | 82.98072 | 2.80E-28 | 1.84E-25 | -1.02715 | 8.098125 | -41.7663 | 5.89E-22 | 7.10E-20 | 8178.112 | 8656.565 | 8462.704 | 48866.53 | 50149.55 | 48370.05 | 25557.11 | 24962.67 | 24605.72 | 12881.27 | 13251.21 | 12617.65 | 8743.746 | 8551.007 | 9121.569 |
| Gm9753 | 791382 | 2.450603 | -3.52635 | 2.859004 | 0.009299 | 0.022467 | -0.49573 | -3.52635 | -0.65403 | 0.520078 | 0.616625 | 2.225942 | 1.77607 | 0.86797 | 9.090964 | 25.27911 | 5.410671 | 2.842522 | 9.382952 | 14.93902 | 0 | 4.575167 | 0.746341 | 16.68654 | 9.432991 | 22.67637 |
| Gm7135 | 634346 | 2.450902 | -1.27958 | 6.624748 | 1.35E-06 | 5.53E-06 | -1.38191 | -1.27958 | -4.18279 | 0.000408 | 0.001196 | 9.64575 | 14.20856 | 24.30315 | 84.4161 | 80.05053 | 98.74475 | 36.95278 | 26.80843 | 40.22045 | 12.30743 | 16.01308 | 13.43414 | 23.36115 | 7.860826 | 12.14805 |
| Pbld1 | 68371 | 2.451448 | -3.49346 | 2.648701 | 0.014895 | 0.034338 | 0.455482 | -3.49346 | 0.603802 | 0.55234 | 0.647029 | 0 | 2.664105 | 3.471878 | 3.896127 | 15.44835 | 10.82134 | 8.527565 | 12.0638 | 18.38649 | 0.769214 | 0.762528 | 2.985365 | 8.343269 | 7.074744 | 29.9652 |
| Whamm | 434204 | 2.451495 | 4.650278 | 35.62379 | 1.68E-20 | 8.06E-19 | -0.70183 | 4.650278 | -12.5643 | 2.49E-11 | 2.16E-10 | 800.5972 | 895.1393 | 914.84 | 4988.342 | 5210.306 | 4980.523 | 3261.794 | 3317.544 | 3117.659 | 785.368 | 928.7589 | 849.3363 | 558.1647 | 667.3841 | 665.7134 |
| Rbpms | 19663 | 2.451678 | 4.835045 | 35.97867 | 1.36E-20 | 6.69E-19 | 0.443645 | 4.835045 | 9.311957 | 5.74E-09 | 3.38E-08 | 715.2694 | 716.6442 | 761.2093 | 4329.896 | 4152.797 | 4316.363 | 5987.772 | 6349.578 | 5775.656 | 601.5257 | 651.9613 | 614.9852 | 1183.076 | 1213.712 | 1304.701 |
| 4933434M | 71203 | 2.455175 | -4.66544 | 2.933697 | 0.007846 | 0.019254 | 1.146211 | -4.66544 | 1.700851 | 0.103497 | 0.160737 | 0 | 0.888035 | 0.86797 | 2.597418 | 5.617581 | 5.410671 | 5.685043 | 10.72337 | 18.38649 | 0 | 0 | 0 | 1.668654 | 3.14433 | 8.098703 |
| Gabrb2 | 14401 | 2.457578 | -2.22969 | 3.919609 | 0.000768 | 0.002247 | 0.406074 | -2.22969 | 0.900109 | 0.378106 | 0.476341 | 1.483961 | 8.88035 | 4.339848 | 20.77935 | 36.51428 | 18.93735 | 55.42917 | 20.10633 | 34.47467 | 3.076858 | 1.525056 | 2.239024 | 41.71635 | 32.22939 | 29.9652 |
| Mark1 | 226778 | 2.457981 | -1.38102 | 4.170742 | 0.00042 | 0.001275 | 0.321785 | -1.38102 | 0.795541 | 0.435058 | 0.534046 | 15.5816 | 2.664105 | 5.207818 | 27.27289 | 37.91867 | 58.16471 | 35.53152 | 45.57434 | 74.69511 | 13.07665 | 14.48803 | 7.463413 | 52.5626 | 32.22939 | 42.11325 |
| Arl5c | 217151 | 2.458055 | 4.615556 | 27.70033 | 3.23E-18 | 9.16E-17 | 0.953075 | 4.615556 | 16.59692 | 1.11E-13 | 1.52E-12 | 419.2191 | 443.1295 | 466.9676 | 2579.236 | 2756.828 | 2452.387 | 5312.673 | 5068.135 | 5245.895 | 2502.255 | 2146.516 | 2281.565 | 445.5306 | 522.7449 | 502.1196 |
| Mrgprx2 | 243978 | 2.458427 | -2.79763 | 2.57368 | 0.017571 | 0.039723 | 1.453725 | -2.79763 | 2.16739 | 0.041655 | 0.074109 | 1.483961 | 0 | 13.88751 | 14.2858 | 5.617581 | 24.34802 | 31.26774 | 34.85097 | 45.96622 | 4.615287 | 3.050111 | 0 | 20.85817 | 26.72681 | 18.62702 |
| Spata13 | 219140 | 2.458453 | 6.108964 | 48.24222 | 2.80E-23 | 2.73E-21 | 0.30414 | 6.108964 | 8.518957 | 2.61E-08 | 1.39E-07 | 3042.863 | 2809.743 | 3062.197 | 17444.26 | 17814.75 | 17108.54 | 22633.58 | 23512.34 | 21199.62 | 789.214 | 727.4516 | 803.0632 | 1324.077 | 1275.026 | 1392.977 |
| Mrgprx3-ps | 100502859 | 2.45853 | -4.54722 | 2.779143 | 0.011136 | 0.026431 | 1.219374 | -4.54722 | 1.711877 | 0.101414 | 0.157902 | 1.483961 | 0 | 0 | 5.194837 | 1.404395 | 5.410671 | 15.63387 | 5.361687 | 10.3424 | 0.769214 | 0 | 0 | 4.171635 | 11.00516 | 4.859222 |
| E230013L | 102132 | 2.46296 | 2.330052 | 15.03669 | 7.82E-13 | 7.72E-12 | -0.03712 | 2.330052 | -0.33412 | 0.741551 | 0.819097 | 111.2971 | 160.7343 | 138.8751 | 846.7584 | 793.4833 | 731.7932 | 794.4848 | 864.572 | 750.3986 | 205.3803 | 185.2943 | 282.117 | 129.3207 | 147.7835 | 110.9522 |
| Sult2a1 | 20859 | 2.464032 | -4.1853 | 2.7016 | 0.013245 | 0.030902 | 0.426868 | -4.1853 | 0.553596 | 0.585609 | 0.679315 | 0 | 0 | 5.207818 | 7.792255 | 2.808791 | 5.410671 | 8.527565 | 8.04253 | 5.745778 | 0 | 1.525056 | 1.492683 | 9.177596 | 7.860826 | 5.669092 |
| Asb13 | 142688 | 2.466949 | 3.595371 | 23.67222 | 8.34E-17 | 1.73E-15 | 0.635417 | 3.595371 | 9.500704 | 4.05E-09 | 2.44E-08 | 241.1437 | 245.9857 | 287.2979 | 1448.061 | 1565.901 | 1542.041 | 2447.411 | 2542.78 | 2360.366 | 560.7573 | 545.9699 | 617.2242 | 370.4412 | 388.3248 | 316.6593 |
| Bcl2l14 | 66813 | 2.467243 | -2.14152 | 2.847574 | 0.009543 | 0.02301 | 1.18744 | -2.14152 | 1.993262 | 0.059171 | 0.100063 | 0.741981 | 10.65642 | 2.603909 | 35.06515 | 9.830767 | 17.58468 | 36.95278 | 37.53181 | 60.90525 | 24.61486 | 12.20045 | 5.97073 | 24.19548 | 4.716496 | 16.19741 |
| Gm8526 | 667230 | 2.467629 | -5.29864 | 2.544584 | 0.018725 | 0.042068 | 0.106232 | -5.29864 | 0.122322 | 0.903789 | 0.924421 | 0 | 0 | 0 | 0 | 4.213186 | 4.058003 | 2.842522 | 1.340422 | 2.298311 | 0 | 1.525056 | 0 | 4.171635 | 6.288661 | 2.429611 |
| Ly6g5c | 114652 | 2.4685 | -4.41892 | 2.558619 | 0.01816 | 0.040913 | -0.53285 | -4.41892 | -0.60293 | 0.552909 | 0.647609 | 0 | 1.77607 | 0 | 3.896127 | 7.021976 | 1.352668 | 4.263783 | 0 | 6.894934 | 2.307643 | 5.337695 | 3.731706 | 4.171635 | 4.716496 | 4.859222 |
| Olfr263 | 18341 | 2.47258 | -4.57102 | 2.772439 | 0.011305 | 0.026781 | 0.343158 | -4.57102 | 0.45061 | 0.656815 | 0.74323 | 0.741981 | 0 | 0.86797 | 5.194837 | 4.213186 | 4.058003 | 9.948826 | 4.021265 | 5.745778 | 3.076858 | 1.525056 | 0 | 1.668654 | 0.786083 | 5.669092 |
| Sult1e1 | 20860 | 2.477405 | -4.36096 | 2.894703 | 0.008575 | 0.020877 | 0.372021 | -4.36096 | 0.514698 | 0.612053 | 0.703905 | 0.741981 | 0 | 1.735939 | 1.298709 | 7.021976 | 14.87935 | 8.527565 | 5.361687 | 9.193245 | 0 | 0 | 0.746341 | 8.343269 | 6.288661 | 6.478962 |
| Klk1b16 | 16615 | 2.477553 | -3.92968 | 2.65217 | 0.014782 | 0.034097 | -0.49181 | -3.92968 | -0.58735 | 0.563132 | 0.657563 | 0.741981 | 0 | 5.207818 | 7.792255 | 14.04395 | 4.058003 | 4.263783 | 2.680843 | 12.64071 | 0 | 4.575167 | 1.492683 | 7.508942 | 11.00516 | 6.478962 |
| Poteb | 100040599 | 2.47827 | -2.3051 | 2.961327 | 0.007366 | 0.018175 | 0.473554 | -2.3051 | 0.768253 | 0.450754 | 0.549595 | 0 | 15.98463 | 7.811726 | 16.88322 | 14.04395 | 33.81669 | 24.16143 | 26.80843 | 36.77298 | 6.153716 | 3.812639 | 5.97073 | 20.02385 | 22.01031 | 18.62702 |
| Kcnj11 | 16514 | 2.478782 | -2.13358 | 3.4433 | 0.002395 | 0.006473 | 0.583369 | -2.13358 | 1.127174 | 0.272182 | 0.363126 | 4.451884 | 3.55214 | 1.735939 | 7.792255 | 28.08791 | 37.8747 | 21.31891 | 36.19139 | 39.07129 | 7.692145 | 3.050111 | 10.44878 | 40.04769 | 29.87114 | 21.8665 |
| Gcnt2 | 14538 | 2.479188 | 4.315151 | 34.89765 | 2.58E-20 | 1.19E-18 | 1.586058 | 4.315151 | 35.99415 | 1.35E-20 | 1.06E-18 | 534.9681 | 573.6706 | 645.7694 | 3528.593 | 3565.76 | 3291.041 | 10739.05 | 10683.16 | 10902.04 | 186.1499 | 200.5448 | 223.156 | 532.3006 | 576.1986 | 511.838 |
| Adam3 | 11497 | 2.479371 | -0.08338 | 12.9461 | 1.41E-11 | 1.14E-10 | -2.10214 | -0.08338 | -10.6703 | 5.14E-10 | 3.58E-09 | 53.42261 | 41.73764 | 56.41802 | 305.1967 | 296.3274 | 302.9976 | 62.53548 | 81.76573 | 73.54596 | 15.38429 | 13.7255 | 8.956095 | 37.54471 | 46.37887 | 36.44416 |
| C2 | 12263 | 2.480215 | 0.623785 | 8.826894 | 1.43E-08 | 7.54E-08 | 1.073087 | 0.623785 | 6.53908 | 1.64E-06 | 6.78E-06 | 37.84102 | 28.41712 | 27.77503 | 180.5206 | 172.7406 | 215.0742 | 449.1184 | 426.2541 | 374.6247 | 46.92208 | 40.41398 | 66.42437 | 37.54471 | 63.67269 | 60.74027 |
| Myt1l | 17933 | 2.481729 | -1.66365 | 4.397493 | 0.000243 | 0.00076 | 0.196635 | -1.66365 | 0.494329 | 0.626123 | 0.716509 | 5.193865 | 6.216245 | 5.207818 | 22.07806 | 54.77142 | 35.16936 | 42.63783 | 42.8935 | 39.07129 | 5.384501 | 7.625279 | 4.478048 | 51.72827 | 60.52836 | 26.72572 |
| Gm14636 | 100503338 | 2.484354 | 6.045374 | 54.79354 | 1.89E-24 | 2.67E-22 | -1.14908 | 6.045374 | -29.2328 | 1.05E-18 | 4.73E-17 | 2327.594 | 2315.107 | 2389.52 | 14154.63 | 13622.63 | 14231.42 | 6647.237 | 6695.407 | 6351.383 | 2469.178 | 2550.656 | 2305.448 | 1750.418 | 1881.882 | 1841.645 |
| Gm16619 | 100503470 | 2.484584 | -3.76877 | 2.601943 | 0.016514 | 0.037582 | 0.683854 | -3.76877 | 0.868562 | 0.394749 | 0.493643 | 3.709904 | 0 | 0 | 3.896127 | 11.23516 | 2.705335 | 2.842522 | 22.78717 | 9.193245 | 3.076858 | 1.525056 | 2.985365 | 9.177596 | 11.00516 | 14.57766 |
| Rasgrp1 | 19419 | 2.485149 | 5.082152 | 31.44235 | 2.30E-19 | 8.34E-18 | -2.69727 | 5.082152 | -29.0588 | 1.19E-18 | 5.25E-17 | 802.8232 | 831.2007 | 598.899 | 4542.885 | 4513.726 | 4202.739 | 697.8391 | 697.0193 | 721.6697 | 3219.932 | 3228.543 | 3330.921 | 4595.473 | 4412.282 | 4620.31 |
| Cers6 | 241447 | 2.485948 | 8.017405 | 81.13233 | 4.52E-28 | 2.64E-25 | -1.29996 | 8.017405 | -48.8683 | 2.13E-23 | 4.71E-21 | 9260.662 | 9206.259 | 9183.118 | 55512.02 | 55935.66 | 54003.91 | 23379.74 | 24374.23 | 22200.54 | 12101.28 | 12222.56 | 12371.35 | 6229.919 | 5937.282 | 6292.692 |
| Nutm2 | 328250 | 2.486177 | -3.53362 | 3.143871 | 0.004835 | 0.012367 | 0.338021 | -3.53362 | 0.533244 | 0.599374 | 0.692229 | 4.451884 | 0 | 2.603909 | 7.792255 | 9.830767 | 16.23201 | 15.63387 | 8.04253 | 21.83396 | 0.769214 | 0.762528 | 0 | 10.84625 | 19.65207 | 13.76779 |
| Mir221 | 723827 | 2.486879 | -0.20269 | 9.310696 | 5.75E-09 | 3.21E-08 | -1.3851 | -0.20269 | -5.94153 | 6.32E-06 | 2.41E-05 | 34.8731 | 36.40943 | 26.90706 | 220.7806 | 172.7406 | 197.4895 | 69.64178 | 76.40404 | 86.18667 | 23.07643 | 26.68847 | 54.48291 | 25.86413 | 21.22423 | 21.05663 |
| A230052G | 442796 | 2.487016 | -2.22628 | 2.792998 | 0.010794 | 0.025687 | 1.149657 | -2.22628 | 1.780003 | 0.089319 | 0.141773 | 0 | 7.10428 | 3.471878 | 16.88322 | 8.426372 | 12.17401 | 55.42917 | 18.7659 | 24.13227 | 13.07665 | 5.337695 | 7.463413 | 49.22529 | 59.74228 | 35.63429 |
| Gm2329 | 100039589 | 2.489693 | -4.0714 | 2.724468 | 0.012587 | 0.029523 | 0.83801 | -4.0714 | 1.130376 | 0.270861 | 0.361694 | 3.709904 | 0 | 0 | 3.896127 | 2.808791 | 10.82134 | 11.37009 | 9.382952 | 9.193245 | 0.769214 | 3.050111 | 0 | 7.508942 | 8.646909 | 9.718443 |
| Awat2 | 245532 | 2.490044 | -1.46318 | 5.725043 | 1.04E-05 | 3.84E-05 | -2.86277 | -1.46318 | -5.42557 | 2.09E-05 | 7.42E-05 | 14.83961 | 14.20856 | 14.75548 | 93.50706 | 92.69009 | 82.51273 | 9.948826 | 6.702109 | 24.13227 | 29.99936 | 44.22662 | 11.19512 | 14.18356 | 7.074744 | 12.95792 |
| Il17rd | 171463 | 2.492212 | 0.474107 | 5.717782 | 1.06E-05 | 3.90E-05 | 1.209522 | 0.474107 | 4.76696 | 1.00E-04 | 0.000324 | 23.0014 | 19.53677 | 10.41564 | 96.10448 | 106.734 | 109.5661 | 271.4608 | 257.361 | 228.682 | 59.99873 | 84.64059 | 30.59999 | 134.3266 | 193.3763 | 157.9247 |
| Ms4a18 | 76002 | 2.493503 | -3.85067 | 3.137357 | 0.004908 | 0.012537 | 0.96553 | -3.85067 | 1.583942 | 0.127913 | 0.192221 | 2.225942 | 0 | 2.603909 | 12.98709 | 7.021976 | 6.763339 | 11.37009 | 24.12759 | 20.6848 | 0.769214 | 0 | 0 | 14.18356 | 7.860826 | 4.859222 |
| Lpin3 | 64899 | 2.493881 | -1.96701 | 4.013885 | 0.000612 | 0.001818 | 0.689573 | -1.96701 | 1.610068 | 0.122075 | 0.184788 | 1.483961 | 9.768385 | 4.339848 | 37.66257 | 16.85274 | 25.70069 | 48.32287 | 37.53181 | 44.81707 | 5.384501 | 6.100223 | 3.731706 | 19.18952 | 40.8763 | 34.01455 |
| Il1f10 | 215274 | 2.495552 | -3.80158 | 3.088673 | 0.005495 | 0.01392 | -0.4891 | -3.80158 | -0.68786 | 0.498947 | 0.596844 | 0.741981 | 1.77607 | 2.603909 | 7.792255 | 21.06593 | 8.116006 | 2.842522 | 13.40422 | 11.49156 | 0.769214 | 4.575167 | 4.478048 | 5.840288 | 0.786083 | 1.619741 |
| Zswim2 | 71861 | 2.495936 | -3.11038 | 2.499209 | 0.020668 | 0.04588 | 0.877588 | -3.11038 | 1.173966 | 0.253342 | 0.341858 | 5.935846 | 2.664105 | 0 | 14.2858 | 21.06593 | 5.410671 | 8.527565 | 29.48928 | 42.51876 | 0.769214 | 8.387806 | 0.746341 | 13.34923 | 18.0799 | 6.478962 |
| Cel | 12613 | 2.497752 | -2.52851 | 2.961701 | 0.00736 | 0.018162 | 0.177192 | -2.52851 | 0.270223 | 0.789584 | 0.860232 | 4.451884 | 0 | 9.547666 | 14.2858 | 11.23516 | 27.05335 | 25.5827 | 8.04253 | 32.17636 | 11.53822 | 17.53814 | 10.44878 | 7.508942 | 13.3634 | 8.098703 |
| Mir146b | 751550 | 2.498403 | -3.3489 | 3.665484 | 0.001412 | 0.003963 | -0.01411 | -3.3489 | -0.02625 | 0.979298 | 0.982698 | 2.225942 | 0.888035 | 6.943757 | 16.88322 | 18.25714 | 16.23201 | 21.31891 | 21.44675 | 12.64071 | 1.538429 | 6.862751 | 2.985365 | 0.834327 | 0.786083 | 4.859222 |
| Gm12216 | 622459 | 2.499842 | -0.00019 | 5.829054 | 8.18E-06 | 3.06E-05 | 1.166562 | -0.00019 | 4.685293 | 0.000122 | 0.000389 | 25.96933 | 23.08891 | 12.15157 | 116.8838 | 130.6088 | 114.9768 | 329.7325 | 313.6587 | 226.3837 | 33.84544 | 44.22662 | 35.07804 | 40.04769 | 48.73712 | 12.14805 |
| Pappa2 | 23850 | 2.501707 | -0.74798 | 6.075908 | 4.65E-06 | 1.78E-05 | -0.50684 | -0.74798 | -1.65333 | 0.112899 | 0.17305 | 17.06556 | 10.65642 | 11.2836 | 85.7148 | 95.49888 | 62.22272 | 42.63783 | 56.29771 | 76.99342 | 19.99958 | 31.26364 | 11.19512 | 44.21933 | 44.02063 | 55.07118 |
| Art3 | 109979 | 2.503422 | -1.40257 | 5.551991 | 1.56E-05 | 5.62E-05 | 1.669587 | -1.40257 | 6.473661 | 1.89E-06 | 7.77E-06 | 14.09763 | 4.440175 | 11.2836 | 57.1432 | 64.60218 | 51.40137 | 179.0789 | 201.0633 | 193.0581 | 6.153716 | 4.575167 | 2.985365 | 15.01788 | 17.29382 | 5.669092 |
| Tnf | 21926 | 2.504464 | 7.123557 | 97.80553 | 8.45E-30 | 1.65E-26 | -1.50185 | 7.123557 | -65.2631 | 4.62E-26 | 2.96E-23 | 10470.09 | 10643.99 | 10059.77 | 62682.2 | 64123.28 | 62111.8 | 23436.59 | 23200.02 | 22659.05 | 5226.812 | 4941.18 | 5244.54 | 501.4305 | 485.013 | 571.7684 |
| Dlx6os1 | 320038 | 2.504493 | -3.02967 | 2.703563 | 0.013187 | 0.030775 | 1.717996 | -3.02967 | 2.601193 | 0.016541 | 0.033486 | 2.967923 | 1.77607 | 0 | 6.493546 | 8.426372 | 10.82134 | 39.7953 | 32.17012 | 22.98311 | 0 | 5.337695 | 4.478048 | 20.85817 | 30.65722 | 17.00728 |
| Gm19340 | 100502727 | 2.505418 | -4.61785 | 2.732761 | 0.012355 | 0.02903 | -1.69864 | -4.61785 | -1.89115 | 0.072256 | 0.118368 | 0 | 2.664105 | 1.735939 | 5.194837 | 14.04395 | 5.410671 | 1.421261 | 4.021265 | 1.149156 | 0 | 4.575167 | 0 | 5.005962 | 3.14433 | 1.619741 |
| Rasef | 242505 | 2.505675 | -2.48096 | 3.591714 | 0.001684 | 0.004661 | 0.829781 | -2.48096 | 1.669206 | 0.109681 | 0.168953 | 4.451884 | 3.55214 | 0.86797 | 12.98709 | 12.63956 | 31.11136 | 36.95278 | 21.44675 | 41.3696 | 6.153716 | 7.625279 | 1.492683 | 16.68654 | 20.43815 | 15.38754 |
| Insl5 | 23919 | 2.506011 | -4.17622 | 2.961273 | 0.007367 | 0.018175 | 0.517668 | -4.17622 | 0.729338 | 0.473724 | 0.572189 | 2.967923 | 0 | 0 | 6.493546 | 2.808791 | 5.410671 | 4.263783 | 9.382952 | 9.193245 | 1.538429 | 0.762528 | 3.731706 | 5.005962 | 3.930413 | 5.669092 |
| Ppfia3 | 76787 | 2.506394 | 1.614438 | 15.44393 | 4.63E-13 | 4.76E-12 | -1.00697 | 1.614438 | -7.80744 | 1.09E-07 | 5.30E-07 | 90.52165 | 97.68385 | 72.90945 | 458.4443 | 596.868 | 538.3618 | 254.4057 | 270.7652 | 287.2889 | 99.22866 | 115.9042 | 102.9951 | 152.6818 | 162.7191 | 149.826 |
| Tmem30c | 71027 | 2.507012 | -3.66197 | 2.774989 | 0.01124 | 0.02665 | 0.891724 | -3.66197 | 1.255373 | 0.222915 | 0.30723 | 2.967923 | 0 | 0.86797 | 11.68838 | 4.213186 | 6.763339 | 9.948826 | 18.7659 | 14.93902 | 0.769214 | 0 | 5.97073 | 11.68058 | 15.72165 | 8.098703 |
| 1700017N | 66605 | 2.507355 | -3.60894 | 2.543168 | 0.018783 | 0.042173 | -0.07801 | -3.60894 | -0.09386 | 0.926096 | 0.939963 | 2.225942 | 0 | 4.339848 | 3.896127 | 28.08791 | 9.468674 | 4.263783 | 18.7659 | 11.49156 | 5.384501 | 0 | 0 | 18.35519 | 16.50774 | 9.718443 |
| Fgf7 | 14178 | 2.509391 | -2.51472 | 3.372148 | 0.002833 | 0.007553 | -0.07119 | -2.51472 | -0.12344 | 0.902916 | 0.923983 | 2.225942 | 8.88035 | 1.735939 | 15.58451 | 29.4923 | 22.99535 | 14.21261 | 24.12759 | 27.57973 | 0 | 3.812639 | 7.463413 | 33.37308 | 22.7964 | 30.77507 |
| Dmrta1 | 242523 | 2.509759 | -3.24392 | 2.500204 | 0.020623 | 0.045793 | 1.105179 | -3.24392 | 1.504594 | 0.147084 | 0.216393 | 0 | 3.55214 | 6.943757 | 28.5716 | 5.617581 | 10.82134 | 22.74017 | 16.08506 | 55.15947 | 0 | 0 | 2.985365 | 26.69846 | 14.14949 | 9.718443 |
| Krt72 | 105866 | 2.510249 | -2.9815 | 2.745894 | 0.011998 | 0.028275 | 1.423133 | -2.9815 | 2.122833 | 0.045623 | 0.080065 | 0 | 2.664105 | 3.471878 | 10.38967 | 4.213186 | 16.23201 | 12.79135 | 33.51054 | 39.07129 | 3.846072 | 0.762528 | 2.239024 | 35.87606 | 22.7964 | 18.62702 |
| Mir146 | 387164 | 2.510378 | -3.51809 | 3.624721 | 0.001556 | 0.004338 | -1.37099 | -3.51809 | -2.15663 | 0.042583 | 0.075526 | 5.193865 | 7.992315 | 0.86797 | 32.46773 | 11.23516 | 33.81669 | 14.21261 | 12.0638 | 4.596622 | 5.384501 | 12.96297 | 13.43414 | 0 | 0 | 0 |
| Mep1b | 17288 | 2.510563 | -3.7723 | 3.000053 | 0.00674 | 0.01677 | 1.793192 | -3.7723 | 2.885122 | 0.008764 | 0.019225 | 0 | 0.888035 | 1.735939 | 9.090964 | 4.213186 | 4.058003 | 24.16143 | 13.40422 | 28.72889 | 0 | 0.762528 | 0.746341 | 9.177596 | 19.65207 | 17.00728 |
| Fezf1 | 73191 | 2.511137 | -3.25037 | 2.80984 | 0.010392 | 0.024828 | 2.355643 | -3.25037 | 3.651381 | 0.00146 | 0.00386 | 2.225942 | 0 | 0 | 6.493546 | 2.808791 | 4.058003 | 22.74017 | 20.10633 | 34.47467 | 3.846072 | 1.525056 | 3.731706 | 22.52683 | 47.16496 | 25.10598 |
| 8030474K | 382231 | 2.511802 | -3.17642 | 2.648041 | 0.014917 | 0.034382 | 1.182687 | -3.17642 | 1.699034 | 0.103844 | 0.16119 | 8.161788 | 0.888035 | 0 | 6.493546 | 28.08791 | 5.410671 | 32.689 | 14.74464 | 29.87805 | 2.307643 | 3.050111 | 0.746341 | 20.85817 | 13.3634 | 6.478962 |
| Olfr43 | 258706 | 2.514263 | -4.90645 | 2.496826 | 0.020775 | 0.046094 | 0.901914 | -4.90645 | 1.113629 | 0.277823 | 0.369415 | 4.451884 | 0 | 0 | 18.18193 | 2.808791 | 2.705335 | 7.106304 | 8.04253 | 21.83396 | 0 | 0 | 0 | 0 | 0.786083 | 3.239481 |
| Tcap | 21393 | 2.514866 | -1.63802 | 3.909403 | 0.000787 | 0.002298 | 0.084151 | -1.63802 | 0.184598 | 0.855287 | 0.917108 | 2.967923 | 9.768385 | 9.547666 | 36.36386 | 25.27911 | 73.04406 | 34.11026 | 49.5956 | 51.712 | 19.99958 | 15.25056 | 3.731706 | 10.84625 | 20.43815 | 25.91585 |
| Hp | 15439 | 2.514984 | 3.489389 | 32.07955 | 1.51E-19 | 5.72E-18 | -0.53674 | 3.489389 | -8.84164 | 1.39E-08 | 7.71E-08 | 639.5874 | 678.4587 | 603.2389 | 4031.193 | 3791.867 | 3887.567 | 2991.754 | 2863.141 | 2576.407 | 186.1499 | 174.6189 | 216.439 | 85.10135 | 124.2011 | 97.18443 |
| Hmgn3 | 94353 | 2.515606 | 2.770374 | 23.56474 | 9.16E-17 | 1.89E-15 | -3.23043 | 2.770374 | -21.8841 | 4.18E-16 | 9.93E-15 | 275.2749 | 412.0482 | 330.6964 | 2040.272 | 2032.16 | 1995.185 | 190.449 | 253.3397 | 227.5328 | 383.0688 | 394.2269 | 330.6292 | 205.2444 | 221.6753 | 181.4109 |
| Igkv4-63 | 628571 | 2.516413 | -4.94103 | 2.516853 | 0.019891 | 0.044347 | -1.62373 | -4.94103 | -1.66968 | 0.109586 | 0.168848 | 1.483961 | 2.664105 | 0 | 7.792255 | 19.66153 | 2.705335 | 4.263783 | 0 | 5.745778 | 0 | 0 | 0.746341 | 0 | 4.716496 | 1.619741 |
| Ccdc149 | 100503884 | 2.51682 | -1.85649 | 4.556957 | 0.000166 | 0.000528 | 0.091824 | -1.85649 | 0.227457 | 0.822233 | 0.889504 | 3.709904 | 4.440175 | 5.207818 | 29.87031 | 25.27911 | 31.11136 | 24.16143 | 29.48928 | 42.51876 | 6.153716 | 17.53814 | 10.44878 | 18.35519 | 36.1598 | 22.67637 |
| Olfr638 | 259124 | 2.51753 | -4.73967 | 2.684096 | 0.013771 | 0.032004 | 0.495047 | -4.73967 | 0.621357 | 0.540945 | 0.636089 | 1.483961 | 0 | 0 | 7.792255 | 1.404395 | 4.058003 | 4.263783 | 4.021265 | 10.3424 | 0 | 0 | 1.492683 | 1.668654 | 3.930413 | 8.908573 |
| Kctd15 | 233107 | 2.518426 | -1.97114 | 3.939014 | 0.000733 | 0.002151 | 1.547753 | -1.97114 | 3.800311 | 0.001023 | 0.002792 | 4.451884 | 3.55214 | 2.603909 | 28.5716 | 26.68351 | 16.23201 | 48.32287 | 83.10615 | 87.33582 | 1.538429 | 6.862751 | 1.492683 | 30.8701 | 55.02578 | 42.92312 |
| Cd70 | 21948 | 2.51861 | -3.42551 | 2.798546 | 0.01066 | 0.025396 | 1.270186 | -3.42551 | 1.897571 | 0.071363 | 0.117136 | 4.451884 | 0 | 0.86797 | 16.88322 | 5.617581 | 5.410671 | 29.84648 | 17.42548 | 19.53565 | 3.846072 | 1.525056 | 0 | 9.177596 | 13.3634 | 17.81715 |
| Apobec3 | 80287 | 2.519122 | 7.04081 | 58.59204 | 4.56E-25 | 8.16E-23 | -0.07025 | 7.04081 | -2.25718 | 0.034594 | 0.063289 | 3429.435 | 3297.274 | 3419.8 | 20479.34 | 20954.98 | 20728.28 | 21188.16 | 21005.75 | 19470.14 | 6486.785 | 6400.659 | 6296.135 | 3328.13 | 3272.462 | 3242.721 |
| Gm19519 | 100503033 | 2.519169 | -3.65115 | 2.926028 | 0.007985 | 0.01957 | 0.362198 | -3.65115 | 0.514423 | 0.612242 | 0.704031 | 1.483961 | 0 | 3.471878 | 5.194837 | 7.021976 | 16.23201 | 14.21261 | 5.361687 | 18.38649 | 0 | 2.287584 | 0.746341 | 10.01192 | 18.86598 | 14.57766 |
| Gm8369 | 666926 | 2.51993 | 1.528414 | 14.98379 | 8.38E-13 | 8.23E-12 | 0.308326 | 1.528414 | 2.899117 | 0.00849 | 0.018699 | 70.48817 | 80.81118 | 72.04148 | 494.8082 | 426.9362 | 445.0277 | 564.2406 | 583.0835 | 605.605 | 112.3053 | 109.0415 | 82.84388 | 84.26702 | 77.0361 | 66.40936 |
| 1700009J | 75188 | 2.521926 | -4.97287 | 2.808138 | 0.010432 | 0.024914 | -0.58144 | -4.97287 | -0.71025 | 0.485239 | 0.583925 | 0 | 0.888035 | 0.86797 | 7.792255 | 2.808791 | 4.058003 | 1.421261 | 2.680843 | 5.745778 | 0 | 0 | 0 | 0.834327 | 10.21907 | 4.859222 |
| Majin | 622554 | 2.522155 | -0.64385 | 5.139661 | 4.11E-05 | 0.000141 | 1.07901 | -0.64385 | 3.692896 | 0.001322 | 0.00353 | 11.12971 | 7.992315 | 17.35939 | 97.40319 | 60.389 | 64.92805 | 190.449 | 215.8079 | 109.1698 | 18.46115 | 9.912862 | 20.89756 | 38.37904 | 29.87114 | 36.44416 |
| Tmem121 | 69195 | 2.522806 | -2.09291 | 3.919176 | 0.000769 | 0.002249 | 0.389162 | -2.09291 | 0.827419 | 0.417155 | 0.516457 | 5.935846 | 0.888035 | 4.339848 | 20.77935 | 19.66153 | 21.64268 | 24.16143 | 22.78717 | 39.07129 | 4.615287 | 11.43792 | 3.731706 | 35.87606 | 55.81187 | 34.82442 |
| Pstpip2 | 19201 | 2.523426 | 4.360228 | 41.70356 | 6.08E-22 | 4.04E-20 | -0.31301 | 4.360228 | -6.82933 | 8.64E-07 | 3.70E-06 | 788.7255 | 747.7255 | 858.4219 | 4946.783 | 4930.832 | 4807.381 | 4178.507 | 4192.839 | 3925.516 | 795.3677 | 780.066 | 831.4242 | 205.2444 | 213.0284 | 233.2426 |
| Dusp14 | 56405 | 2.52363 | -0.45227 | 5.112272 | 4.39E-05 | 0.00015 | 2.639578 | -0.45227 | 9.786653 | 2.41E-09 | 1.50E-08 | 5.193865 | 7.10428 | 13.01954 | 35.06515 | 64.60218 | 54.10671 | 312.6774 | 282.829 | 371.1773 | 16.92272 | 17.53814 | 13.43414 | 57.56856 | 93.54383 | 72.07845 |
| Olfr788 | 258544 | 2.52405 | -4.96178 | 2.459864 | 0.022503 | 0.049486 | 1.479208 | -4.96178 | 1.754359 | 0.093717 | 0.147597 | 0 | 0 | 0 | 0 | 2.808791 | 6.763339 | 7.106304 | 2.680843 | 13.78987 | 0 | 0 | 0.746341 | 6.674615 | 3.14433 | 8.908573 |
| Unc80 | 329178 | 2.525854 | 0.487039 | 9.06233 | 9.16E-09 | 4.96E-08 | 2.285734 | 0.487039 | 15.47468 | 4.45E-13 | 5.33E-12 | 31.16319 | 33.74533 | 19.09533 | 190.9102 | 167.123 | 152.8515 | 821.4888 | 817.6573 | 940.0093 | 16.1535 | 6.100223 | 10.44878 | 98.45058 | 93.54383 | 98.80417 |
| Gm5606 | 434279 | 2.527009 | -4.04208 | 2.861328 | 0.00925 | 0.02236 | 0.709576 | -4.04208 | 0.993324 | 0.331681 | 0.428087 | 0 | 0.888035 | 2.603909 | 2.597418 | 4.213186 | 20.29002 | 19.89765 | 6.702109 | 10.3424 | 0 | 0 | 0 | 19.18952 | 20.43815 | 12.95792 |
| S100a11-p | 277089 | 2.528993 | -1.68096 | 4.081512 | 0.00052 | 0.001561 | 0.1071 | -1.68096 | 0.243956 | 0.809595 | 0.878032 | 1.483961 | 16.87266 | 10.41564 | 37.66257 | 54.77142 | 32.46403 | 62.53548 | 24.12759 | 59.75609 | 9.230573 | 10.67539 | 7.463413 | 20.02385 | 18.86598 | 14.57766 |
| Vmn2r71 | 233445 | 2.529193 | -4.26569 | 2.726585 | 0.012527 | 0.029397 | 0.731721 | -4.26569 | 0.961932 | 0.346856 | 0.444228 | 0 | 0 | 5.207818 | 5.194837 | 2.808791 | 9.468674 | 4.263783 | 13.40422 | 13.78987 | 0 | 0 | 0 | 10.01192 | 10.21907 | 20.24676 |
| Ido1 | 15930 | 2.529876 | -1.36004 | 4.573215 | 0.000159 | 0.000508 | 3.069193 | -1.36004 | 10.06151 | 1.47E-09 | 9.54E-09 | 7.419807 | 5.32821 | 3.471878 | 35.06515 | 33.70549 | 33.81669 | 378.0554 | 246.6376 | 302.2279 | 1.538429 | 4.575167 | 1.492683 | 52.5626 | 36.1598 | 43.73299 |
| Clca1 | 23844 | 2.532483 | -3.17538 | 3.071605 | 0.005716 | 0.014428 | 1.937032 | -3.17538 | 3.415894 | 0.002555 | 0.006411 | 4.451884 | 0 | 1.735939 | 9.090964 | 9.830767 | 10.82134 | 45.48035 | 38.87223 | 40.22045 | 4.615287 | 0 | 0.746341 | 10.84625 | 11.00516 | 17.81715 |
| Nbl1 | 17965 | 2.533675 | -1.97388 | 2.635879 | 0.015324 | 0.035236 | 1.471308 | -1.97388 | 2.297908 | 0.031767 | 0.05873 | 2.967923 | 18.64873 | 0 | 32.46773 | 12.63956 | 13.52668 | 41.21656 | 52.27645 | 65.50187 | 3.846072 | 0.762528 | 14.92683 | 49.22529 | 77.0361 | 74.50806 |
| Dcaf12l1 | 245404 | 2.533685 | -2.78746 | 3.184615 | 0.004397 | 0.01133 | 0.687836 | -2.78746 | 1.224737 | 0.23402 | 0.319955 | 5.935846 | 3.55214 | 0.86797 | 23.37676 | 11.23516 | 27.05335 | 39.7953 | 46.91476 | 20.6848 | 0 | 0 | 7.463413 | 22.52683 | 17.29382 | 22.67637 |
| Tat | 234724 | 2.534233 | -3.03322 | 3.438669 | 0.002421 | 0.006537 | 0.29512 | -3.03322 | 0.51932 | 0.608882 | 0.701055 | 7.419807 | 0.888035 | 0.86797 | 22.07806 | 14.04395 | 9.468674 | 15.63387 | 30.8297 | 13.78987 | 2.307643 | 0.762528 | 0.746341 | 18.35519 | 28.29897 | 13.76779 |
| Klk1b24 | 16617 | 2.536 | -3.95663 | 2.935732 | 0.00781 | 0.019174 | 0.172137 | -3.95663 | 0.23447 | 0.816855 | 0.884702 | 0 | 1.77607 | 1.735939 | 6.493546 | 5.617581 | 8.116006 | 4.263783 | 9.382952 | 11.49156 | 0.769214 | 4.575167 | 0 | 8.343269 | 9.432991 | 7.288832 |
| Aldh3a1 | 11670 | 2.536182 | -3.0504 | 3.587336 | 0.001701 | 0.004707 | 0.65624 | -3.0504 | 1.237566 | 0.22932 | 0.314525 | 1.483961 | 2.664105 | 1.735939 | 18.18193 | 16.85274 | 8.116006 | 25.5827 | 14.74464 | 29.87805 | 0 | 3.050111 | 0.746341 | 15.85221 | 25.94073 | 25.91585 |
| Cd83 | 12522 | 2.537096 | 5.509761 | 35.1065 | 2.28E-20 | 1.06E-18 | 1.66904 | 5.509761 | 37.80109 | 4.82E-21 | 4.30E-19 | 632.1676 | 661.5861 | 618.8623 | 3824.698 | 4162.628 | 3875.393 | 13452.23 | 13071.79 | 12660.25 | 3749.151 | 3742.487 | 4006.36 | 1148.034 | 1158.686 | 1260.158 |
| Dhh | 13363 | 2.538882 | -0.29591 | 5.368947 | 2.39E-05 | 8.45E-05 | 1.618437 | -0.29591 | 6.045167 | 4.98E-06 | 1.93E-05 | 10.38773 | 18.64873 | 8.679696 | 55.84449 | 103.9252 | 74.39673 | 241.6143 | 247.978 | 234.4277 | 38.46072 | 7.625279 | 8.209754 | 93.44461 | 81.75259 | 84.22651 |
| BB019430 | 103505 | 2.539067 | -4.31307 | 2.658766 | 0.014567 | 0.033643 | 0.062583 | -4.31307 | 0.074553 | 0.941265 | 0.951576 | 0 | 0 | 0.86797 | 1.298709 | 2.808791 | 6.763339 | 1.421261 | 6.702109 | 3.447467 | 0.769214 | 5.337695 | 1.492683 | 6.674615 | 12.57732 | 18.62702 |
| Phxr4 | 18689 | 2.541512 | 3.01741 | 20.08595 | 2.40E-15 | 3.70E-14 | 0.324216 | 3.01741 | 3.992256 | 0.000645 | 0.00183 | 176.5914 | 182.9352 | 210.0486 | 1241.566 | 1131.943 | 1151.12 | 1544.911 | 1666.144 | 1400.821 | 149.9968 | 190.632 | 147.0292 | 495.5902 | 490.5156 | 454.3372 |
| Gm20235 | 100504452 | 2.542017 | 3.667732 | 21.84528 | 4.34E-16 | 7.77E-15 | 0.164933 | 3.667732 | 2.091997 | 0.048566 | 0.084486 | 411.0573 | 341.0054 | 295.9776 | 2332.482 | 2091.145 | 2097.988 | 2497.155 | 2639.29 | 2460.342 | 434.6062 | 360.6757 | 288.8341 | 361.2636 | 379.6779 | 374.1601 |
| Zup1 | 72580 | 2.542788 | 5.947348 | 57.91434 | 5.83E-25 | 9.83E-23 | -0.30502 | 5.947348 | -9.16052 | 7.61E-09 | 4.39E-08 | 1859.404 | 2034.488 | 1814.056 | 11537.73 | 12229.47 | 11693.81 | 9924.665 | 9724.76 | 10091.88 | 2875.324 | 2782.464 | 2774.897 | 1105.483 | 1092.655 | 1073.078 |
| Stard5 | 170460 | 2.544741 | 5.498537 | 42.43728 | 4.21E-22 | 2.84E-20 | -0.75927 | 5.498537 | -15.6569 | 3.53E-13 | 4.32E-12 | 1729.557 | 1629.544 | 1516.343 | 10342.92 | 10323.71 | 9752.734 | 6644.394 | 6411.237 | 5713.602 | 2055.341 | 2074.076 | 2144.985 | 700.8346 | 638.2991 | 673.0022 |
| Gm6524 | 624710 | 2.545636 | -4.05539 | 2.620733 | 0.015845 | 0.036282 | 0.125924 | -4.05539 | 0.160413 | 0.874064 | 0.918721 | 4.451884 | 0 | 2.603909 | 5.194837 | 16.85274 | 16.23201 | 22.74017 | 10.72337 | 9.193245 | 0 | 1.525056 | 1.492683 | 0 | 10.21907 | 0.80987 |
| Nlgn2 | 216856 | 2.546117 | 4.423947 | 13.30776 | 4.67E-11 | 3.48E-10 | 3.198091 | 4.423947 | 31.8253 | 6.69E-18 | 2.40E-16 | 227.0461 | 193.5916 | 251.7112 | 1316.891 | 1358.05 | 1519.046 | 13926.94 | 14504.7 | 11835.15 | 726.9077 | 752.615 | 703.0535 | 1139.691 | 1104.446 | 1196.988 |
| Zfp36 | 22695 | 2.546625 | 8.330248 | 79.60426 | 6.77E-28 | 3.24E-25 | -0.80367 | 8.330248 | -32.6209 | 1.06E-19 | 6.48E-18 | 11486.6 | 11180.36 | 11287.08 | 71853.68 | 70097.58 | 69970.8 | 43347.03 | 42985.98 | 40168.73 | 15250.45 | 15599.79 | 15576.14 | 4846.605 | 5055.297 | 5269.826 |
| Gm13315 | 625342 | 2.549282 | 0.352393 | 6.924463 | 7.02E-07 | 2.97E-06 | 1.908214 | 0.352393 | 9.435931 | 4.56E-09 | 2.73E-08 | 11.87169 | 19.53677 | 14.75548 | 107.7929 | 74.43295 | 105.5081 | 375.2129 | 420.8924 | 332.106 | 23.07643 | 38.88892 | 22.39024 | 169.3684 | 161.933 | 176.5517 |
| Ehd3 | 57440 | 2.5513 | 1.129564 | 9.449995 | 4.44E-09 | 2.52E-08 | -1.50563 | 1.129564 | -6.35365 | 2.48E-06 | 1.00E-05 | 42.2929 | 25.75301 | 26.03909 | 223.378 | 210.6593 | 159.6148 | 72.4843 | 62.99982 | 78.14258 | 109.9977 | 106.7539 | 97.02436 | 896.0671 | 845.0388 | 778.2853 |
| Serpinb3b | 383548 | 2.552233 | -4.35116 | 2.531427 | 0.01927 | 0.043134 | 2.100529 | -4.35116 | 2.809088 | 0.01041 | 0.022361 | 1.483961 | 0 | 0 | 6.493546 | 5.617581 | 1.352668 | 21.31891 | 8.04253 | 37.92213 | 0.769214 | 0.762528 | 0 | 1.668654 | 3.930413 | 8.908573 |
| Mid1 | 17318 | 2.552801 | 6.083784 | 54.32879 | 2.26E-24 | 3.15E-22 | -0.87145 | 6.083784 | -22.6146 | 2.13E-16 | 5.46E-15 | 2133.937 | 2076.226 | 1968.555 | 13024.75 | 12626.92 | 13092.47 | 7518.47 | 7525.128 | 6999.507 | 1876.114 | 1744.664 | 1821.819 | 3013.589 | 3102.668 | 3163.353 |

| | | | | | | | | | | | | | | | | | | | | | | | | | | |
|---|---|---|---|---|---|---|---|---|---|---|---|---|---|---|---|---|---|---|---|---|---|---|---|---|---|---|
| Olfr1417 | 258938 | 2.5536 | -2.57859 | 3.33481 | 0.003093 | 0.008192 | 1.866066 | -2.57859 | 3.527737 | 0.00196 | 0.005033 | 0.741981 | 3.55214 | 0.86797 | 10.38967 | 5.617581 | 18.93735 | 36.95278 | 30.8297 | 58.60694 | 13.84586 | 3.812639 | 4.478048 | 16.68654 | 18.0799 | 19.43689 |
| Gpr87 | 84111 | 2.554223 | -2.28766 | 3.860239 | 0.000886 | 0.002567 | 0.978408 | -2.28766 | 2.311386 | 0.030879 | 0.05729 | 9.64575 | 1.77607 | 5.207818 | 22.07806 | 37.91867 | 37.8747 | 41.21656 | 87.12741 | 73.54596 | 1.538429 | 9.150334 | 0.746341 | 12.5149 | 6.288661 | 9.718443 |
| F930015N | 654805 | 2.554469 | -3.02193 | 4.764506 | 0.000101 | 0.000329 | -0.67113 | -3.02193 | -1.54444 | 0.137181 | 0.204003 | 4.451884 | 3.55214 | 5.207818 | 20.77935 | 37.91867 | 32.46403 | 12.79135 | 28.14886 | 18.38649 | 0 | 0 | 0 | 30.8701 | 14.93557 | 17.00728 |
| Tet2 | 214133 | 2.554614 | 7.968734 | 80.15692 | 5.84E-28 | 3.14E-25 | -1.00096 | 7.968734 | -38.8627 | 2.69E-21 | 2.60E-19 | 8526.843 | 8429.228 | 8598.975 | 53160.06 | 54303.75 | 52910.95 | 28676.78 | 28217.22 | 26534 | 9695.948 | 9350.879 | 9093.422 | 6453.519 | 6356.264 | 6221.423 |
| Ptn | 19242 | 2.554618 | -3.73125 | 2.872933 | 0.00901 | 0.021837 | 0.327064 | -3.73125 | 0.454853 | 0.653808 | 0.740464 | 0 | 0.888035 | 10.41564 | 12.98709 | 11.23516 | 6.763339 | 12.79135 | 9.382952 | 18.38649 | 0 | 1.525056 | 0 | 13.34923 | 14.14949 | 6.478962 |
| Serpina1f | 68348 | 2.555567 | -2.61358 | 3.316471 | 0.00323 | 0.008532 | 0.913332 | -2.61358 | 1.630607 | 0.117643 | 0.179111 | 0.741981 | 1.77607 | 5.207818 | 12.98709 | 8.426372 | 25.70069 | 15.63387 | 29.48928 | 44.81707 | 6.153716 | 2.287584 | 6.717071 | 15.85221 | 26.72681 | 12.14805 |
| Kcnk10 | 72258 | 2.556364 | -2.15851 | 2.882688 | 0.008813 | 0.0214 | 2.579463 | -2.15851 | 4.646592 | 0.000133 | 0.000423 | 5.193865 | 0 | 5.207818 | 7.792255 | 18.25714 | 22.99535 | 116.5434 | 112.5954 | 70.09849 | 9.999788 | 12.20045 | 6.717071 | 16.68654 | 18.86598 | 4.859222 |
| Arrdc5 | 76920 | 2.556368 | -3.77161 | 2.699994 | 0.013293 | 0.031 | 0.167696 | -3.77161 | 0.207244 | 0.837784 | 0.902612 | 4.451884 | 0 | 0 | 3.896127 | 5.617581 | 8.116006 | 1.421261 | 9.382952 | 16.08818 | 1.538429 | 3.050111 | 2.239024 | 16.68654 | 10.21907 | 15.38754 |
| Gmnc | 239789 | 2.556638 | -3.52028 | 2.576111 | 0.017478 | 0.039529 | 1.939198 | -3.52028 | 2.651864 | 0.014792 | 0.030341 | 2.967923 | 0 | 0 | 1.298709 | 5.617581 | 13.52668 | 11.37009 | 20.10633 | 37.92213 | 1.538429 | 1.525056 | 0.746341 | 27.53279 | 28.29897 | 15.38754 |
| 4930546C | 78931 | 2.557321 | -2.97921 | 2.52483 | 0.019549 | 0.043688 | 1.143378 | -2.97921 | 1.526978 | 0.141451 | 0.209217 | 2.225942 | 0 | 3.471878 | 9.090964 | 15.44835 | 6.763339 | 18.47639 | 18.7659 | 34.47467 | 0.769214 | 14.48803 | 9.702436 | 5.005962 | 23.58248 | 9.718443 |
| Rnf19b | 75234 | 2.558732 | 7.927829 | 36.1052 | 5.50E-16 | 9.64E-15 | -0.05857 | 7.927829 | -1.16475 | 0.262327 | 0.352085 | 4067.538 | 4423.302 | 4336.376 | 26598.86 | 27392.73 | 26516.35 | 29512.48 | 26860.71 | 24544.81 | 21337.24 | 21545.22 | 22244.7 | 9769.134 | 9388.185 | 10132.29 |
| Pcgf5 | 76073 | 2.559398 | 4.571671 | 39.48751 | 1.92E-21 | 1.13E-19 | 0.241265 | 4.571671 | 5.314367 | 2.72E-05 | 9.51E-05 | 634.3935 | 737.069 | 694.3757 | 4344.182 | 4355.03 | 4232.497 | 5535.811 | 5066.794 | 5279.221 | 883.8274 | 881.4822 | 853.8144 | 397.1396 | 436.2759 | 437.3299 |
| Gm8995 | 668139 | 2.559865 | 9.730662 | 69.41687 | 2.28E-17 | 5.25E-16 | -0.59958 | 9.730662 | -23.6827 | 1.18E-11 | 1.09E-10 | 30892.37 | 31535.01 | 29353 | 192632.3 | 196573.2 | 188706.6 | 135228.7 | 134554.2 | 127372.4 | 51287.37 | 51878.58 | 49669.01 | 8557.691 | 8559.654 | 8407.263 |
| Cdh9 | 12565 | 2.560959 | -3.2805 | 2.925939 | 0.007986 | 0.019572 | 0.503334 | -3.2805 | 0.733128 | 0.471457 | 0.569938 | 1.483961 | 1.77607 | 0.86797 | 12.98709 | 15.44835 | 5.410671 | 21.31891 | 12.0638 | 14.93902 | 0.769214 | 0 | 7.463413 | 23.36115 | 9.432991 | 26.72572 |
| Vmn2r25 | 545874 | 2.560962 | -3.2393 | 2.562918 | 0.01799 | 0.040565 | 0.299782 | -3.2393 | 0.372236 | 0.713388 | 0.793696 | 1.483961 | 7.10428 | 0 | 18.18193 | 9.830767 | 6.763339 | 5.685043 | 14.74464 | 27.57973 | 7.692145 | 3.812639 | 0 | 14.18356 | 13.3634 | 16.19741 |
| Parp14 | 547253 | 2.561602 | 10.19846 | 65.63138 | 1.22E-15 | 2.00E-14 | -0.69228 | 10.19846 | -25.8684 | 3.24E-11 | 2.77E-10 | 39459.28 | 40219.1 | 36823.61 | 244288.5 | 248041.5 | 242088.3 | 159738.3 | 162232.6 | 151489.7 | 66191.67 | 67676.63 | 66715.44 | 17508.35 | 17210.49 | 16471.14 |
| Tap1 | 21354 | 2.561875 | 8.337522 | 51.69243 | 2.92E-20 | 1.31E-18 | 0.223624 | 8.337522 | 6.893048 | 2.49E-06 | 1.01E-05 | 8218.921 | 8126.408 | 7927.166 | 51741.87 | 51961.22 | 49444.06 | 64270.84 | 63667.35 | 58556.37 | 11885.13 | 12449.03 | 11913.85 | 8541.005 | 8619.396 | 9016.286 |
| Psg20 | 434540 | 2.562149 | -4.21331 | 3.164487 | 0.004608 | 0.011824 | -0.06414 | -4.21331 | -0.09186 | 0.927668 | 0.941267 | 0.741981 | 0.888035 | 0.86797 | 11.68838 | 5.617581 | 5.410671 | 7.106304 | 2.680843 | 16.08818 | 0 | 0.762528 | 0 | 10.01192 | 13.3634 | 4.049351 |
| Vmn2r91 | 665210 | 2.564265 | -4.05338 | 2.659144 | 0.014555 | 0.033618 | 0.074093 | -4.05338 | 0.090897 | 0.928422 | 0.941851 | 0 | 1.77607 | 3.471878 | 7.792255 | 7.021976 | 10.82134 | 1.421261 | 24.12759 | 16.08818 | 0 | 0 | 0.746341 | 8.343269 | 13.3634 | 4.859222 |
| Dst | 13518 | 2.564994 | 8.284238 | 50.08059 | 3.79E-16 | 6.92E-15 | -1.42601 | 8.284238 | -32.178 | 1.08E-13 | 1.49E-12 | 13849.07 | 14087.79 | 12826.85 | 87199.23 | 87294.4 | 83055.15 | 35272.85 | 34711.56 | 30303.23 | 15270.45 | 15281.06 | 15640.33 | 3822.052 | 3714.24 | 3727.833 |
| Ddx58 | 230073 | 2.565045 | 8.907398 | 83.57484 | 2.40E-28 | 1.62E-25 | -0.33935 | 8.907398 | -15.8603 | 2.74E-13 | 3.42E-12 | 12782.84 | 13079.87 | 12131.61 | 80219.97 | 82116.4 | 77766.22 | 66822 | 66452.75 | 64025.2 | 23308.74 | 23621.59 | 22923.87 | 12801.08 | 12549.81 | 12332.7 |
| Cyp2j7 | 546837 | 2.565185 | -3.77965 | 2.784252 | 0.011009 | 0.026164 | 0.339661 | -3.77965 | 0.442079 | 0.662879 | 0.748546 | 0.741981 | 2.664105 | 0 | 3.896127 | 4.213186 | 16.23201 | 7.106304 | 8.04253 | 11.49156 | 0 | 4.575167 | 1.492683 | 17.52087 | 8.646909 | 13.76779 |
| Plaat3 | 225845 | 2.56704 | 3.317017 | 14.84437 | 1.01E-12 | 9.72E-12 | 1.553056 | 3.317017 | 16.05149 | 2.16E-13 | 2.78E-12 | 138.0084 | 130.5411 | 145.8189 | 1038.967 | 742.9251 | 860.2967 | 2835.415 | 2802.822 | 2423.569 | 328.4546 | 387.3641 | 332.1219 | 664.1242 | 648.5182 | 678.6713 |
| Phactr1 | 218194 | 2.567066 | 1.468709 | 8.398723 | 3.30E-08 | 1.66E-07 | 2.257609 | 1.468709 | 14.015 | 3.08E-12 | 3.17E-11 | 24.48536 | 31.08122 | 51.21021 | 237.6638 | 237.3428 | 171.7888 | 1152.643 | 1003.976 | 1041.135 | 130.7665 | 150.9805 | 94.039 | 124.3147 | 149.3557 | 113.3818 |
| Oas1b | 23961 | 2.567794 | 5.071712 | 30.8798 | 3.35E-19 | 1.17E-17 | 0.380274 | 5.071712 | 6.638149 | 1.32E-06 | 5.52E-06 | 776.8538 | 707.7639 | 753.3976 | 4857.172 | 4746.856 | 4590.954 | 6597.493 | 6893.789 | 5820.473 | 1295.357 | 1358.825 | 1213.551 | 1077.116 | 1142.964 | 1154.065 |
| Lag3 | 16768 | 2.567926 | 0.299506 | 7.924121 | 8.56E-08 | 4.07E-07 | -0.31408 | 0.299506 | -1.37888 | 0.182212 | 0.259341 | 14.09763 | 35.5214 | 42.53051 | 163.6374 | 195.2109 | 159.6148 | 177.6576 | 116.6167 | 143.6444 | 59.99873 | 51.08937 | 36.57072 | 57.56856 | 60.52836 | 70.45871 |
| Rspo4 | 228770 | 2.568045 | -2.92241 | 2.529473 | 0.019352 | 0.043296 | 0.848808 | -2.92241 | 1.093524 | 0.286353 | 0.378803 | 0.741981 | 0 | 8.679696 | 24.67547 | 4.213186 | 8.116006 | 8.527565 | 28.14886 | 25.28142 | 10.769 | 1.525056 | 1.492683 | 29.20144 | 18.86598 | 36.44416 |
| Hrc | 15464 | 2.568884 | -0.28066 | 7.136907 | 4.44E-07 | 1.92E-06 | -0.60921 | -0.28066 | -2.29687 | 0.031836 | 0.058838 | 19.2915 | 43.51371 | 24.30315 | 216.8844 | 130.6088 | 175.8468 | 142.1261 | 101.8721 | 111.4681 | 14.61507 | 22.87584 | 30.59999 | 20.02385 | 45.59279 | 12.95792 |
| Wfdc11 | 629761 | 2.570282 | -4.74445 | 3.244186 | 0.003826 | 0.009979 | 1.489992 | -4.74445 | 2.378711 | 0.026777 | 0.050724 | 0 | 0 | 0.86797 | 2.597418 | 4.213186 | 2.705335 | 14.21261 | 6.702109 | 11.49156 | 0 | 0 | 0 | 8.343269 | 3.930413 | 4.859222 |
| Htr1b | 15551 | 2.570282 | -3.79125 | 3.089132 | 0.005489 | 0.013908 | 0.454097 | -3.79125 | 0.638713 | 0.529805 | 0.625837 | 0 | 0 | 0.86797 | 2.597418 | 4.213186 | 2.705335 | 5.685043 | 4.021265 | 4.596622 | 3.846072 | 2.287584 | 1.492683 | 32.53875 | 55.81187 | 47.78235 |
| A730090N | 319798 | 2.571893 | -0.70382 | 7.17458 | 4.10E-07 | 1.78E-06 | 0.451656 | -0.70382 | 1.990173 | 0.059533 | 0.100526 | 11.12971 | 20.4248 | 12.15157 | 105.1954 | 64.60218 | 105.5081 | 142.1261 | 112.5954 | 131.0037 | 15.38429 | 17.53814 | 17.91219 | 25.86413 | 22.7964 | 19.43689 |
| Nat1 | 17960 | 2.572402 | -4.09396 | 2.460963 | 0.02245 | 0.049378 | -0.47598 | -4.09396 | -0.50599 | 0.618052 | 0.708945 | 0 | 5.32821 | 0 | 3.896127 | 14.04395 | 2.705335 | 2.842522 | 1.340422 | 11.49156 | 1.538429 | 1.525056 | 8.956095 | 6.674615 | 8.646909 | 3.239481 |
| Apol6 | 71939 | 2.574208 | -2.76446 | 2.92393 | 0.008023 | 0.019653 | 0.903183 | -2.76446 | 1.459913 | 0.158878 | 0.231222 | 3.709904 | 0.888035 | 3.471878 | 37.66257 | 7.021976 | 17.58468 | 27.00396 | 50.93603 | 28.72889 | 3.846072 | 8.387806 | 0 | 8.343269 | 17.29382 | 13.76779 |
| Igkv3-9 | 667928 | 2.574499 | -5.15101 | 3.063582 | 0.005823 | 0.014672 | -0.37994 | -5.15101 | -0.50593 | 0.618095 | 0.708962 | 1.483961 | 0 | 0 | 2.597418 | 4.213186 | 5.410671 | 4.263783 | 5.361687 | 1.149156 | 0 | 0 | 0 | 1.668654 | 5.502578 | 1.619741 |
| H2-T10 | 15024 | 2.576631 | 4.371024 | 29.49665 | 8.72E-19 | 2.74E-17 | 0.527613 | 4.371024 | 9.086588 | 8.75E-09 | 5.00E-08 | 489.7073 | 534.5971 | 467.8356 | 3210.409 | 3161.294 | 3111.136 | 5022.736 | 4617.753 | 4588.578 | 726.1384 | 735.8394 | 617.2242 | 498.0932 | 594.2785 | 599.304 |
| Gucy2g | 73707 | 2.576959 | -0.0874 | 7.124528 | 4.56E-07 | 1.97E-06 | 0.551011 | -0.0874 | 2.481905 | 0.021457 | 0.041937 | 20.77546 | 22.20087 | 13.01954 | 103.8967 | 106.734 | 150.1461 | 217.4529 | 174.2548 | 159.7326 | 29.23015 | 22.11331 | 11.94146 | 55.8999 | 75.46393 | 76.12781 |
| Vmn1r4 | 171194 | 2.578266 | -4.79054 | 2.732473 | 0.012363 | 0.029044 | -0.71312 | -4.79054 | -0.83007 | 0.41569 | 0.514885 | 0.741981 | 0 | 0.86797 | 1.298709 | 14.04395 | 5.410671 | 2.842522 | 6.702109 | 1.149156 | 0 | 1.525056 | 2.985365 | 1.668654 | 1.572165 | 1.619741 |
| Slc4a4 | 54403 | 2.579652 | -1.44148 | 4.234483 | 0.00036 | 0.001102 | -0.41958 | -1.44148 | -0.93522 | 0.360136 | 0.457913 | 13.35565 | 7.10428 | 5.207818 | 80.51997 | 36.51428 | 51.40137 | 21.31891 | 53.61687 | 57.45778 | 7.692145 | 9.150334 | 0.746341 | 54.23125 | 80.96651 | 55.88105 |
| Glipr2 | 384009 | 2.582506 | 4.775162 | 34.03539 | 4.37E-20 | 1.87E-18 | 0.670481 | 4.775162 | 13.53665 | 6.01E-12 | 5.86E-11 | 488.2233 | 486.6432 | 467.8356 | 3011.707 | 3093.883 | 3135.484 | 5054.004 | 5064.113 | 5132.129 | 1202.282 | 1269.609 | 1147.127 | 1244.816 | 1230.219 | 1409.174 |
| Gypa | 14934 | 2.582643 | -3.02239 | 3.173862 | 0.004509 | 0.011582 | 0.695299 | -3.02239 | 1.10545 | 0.28127 | 0.373044 | 2.967923 | 0 | 2.603909 | 12.98709 | 5.617581 | 13.52668 | 8.527565 | 17.42548 | 31.0272 | 2.307643 | 4.575167 | 2.239024 | 20.85817 | 27.51289 | 16.19741 |
| Hoxd3os1 | 73429 | 2.582888 | -4.38486 | 2.69429 | 0.013462 | 0.031351 | 0.999281 | -4.38486 | 1.281524 | 0.213758 | 0.296553 | 0 | 1.77607 | 0 | 11.68838 | 1.404395 | 2.705335 | 11.37009 | 5.361687 | 10.3424 | 1.538429 | 1.525056 | 0 | 5.840288 | 8.646909 | 3.239481 |
| Rin2 | 74030 | 2.584892 | 7.843966 | 82.86941 | 2.88E-28 | 1.85E-25 | -1.8465 | 7.843966 | -62.4024 | 1.20E-25 | 5.40E-23 | 10541.32 | 10565.84 | 11479.77 | 69522.5 | 71952.79 | 67039.57 | 20370.93 | 20481.64 | 19410.39 | 7432.15 | 7475.823 | 7346.983 | 4177.475 | 4343.106 | 4531.224 |
| Olfr728 | 258039 | 2.586927 | -5.60499 | 2.483957 | 0.021362 | 0.047266 | -0.22487 | -5.60499 | -0.23793 | 0.814202 | 0.882264 | 0 | 0 | 0 | 0 | 5.617581 | 4.058003 | 0 | 2.680843 | 4.596622 | 0 | 0 | 0 | 5.005962 | 3.14433 | 0.80987 |
| Usb1 | 101985 | 2.587287 | 6.667617 | 68.26907 | 1.77E-26 | 5.37E-24 | -0.98241 | 6.667617 | -31.4811 | 2.24E-19 | 1.23E-17 | 3152.676 | 3107.234 | 3258.358 | 20491.03 | 20629.16 | 19953.2 | 11145.53 | 10498.18 | 10489.49 | 4540.673 | 4479.089 | 4308.628 | 2663.172 | 2803.171 | 2628.029 |
| Dram1 | 71712 | 2.587521 | 5.8828 | 51.59953 | 6.74E-24 | 7.99E-22 | -1.81669 | 5.8828 | -38.1035 | 4.07E-21 | 3.76E-19 | 2642.193 | 2676.537 | 2516.244 | 16887.12 | 17667.29 | 15797.81 | 5116.539 | 5204.858 | 4572.49 | 3197.624 | 3338.347 | 3246.584 | 780.93 | 747.5646 | 784.7643 |
| Mtmr14 | 97287 | 2.588002 | 6.405307 | 51.67274 | 6.54E-24 | 7.82E-22 | 0.328888 | 6.405307 | 9.517 | 3.93E-09 | 2.37E-08 | 1994.444 | 2044.257 | 2061.428 | 13244.24 | 13185.87 | 12708.31 | 17335.12 | 17334.33 | 16458.21 | 4276.063 | 4434.862 | 4674.335 | 1535.996 | 1679.072 | 1760.658 |
| Olfr675 | 258147 | 2.589421 | -4.68331 | 2.724703 | 0.01258 | 0.02951 | -0.16106 | -4.68331 | -0.19297 | 0.848807 | 0.911762 | 0 | 3.55214 | 0 | 3.896127 | 2.808791 | 9.468674 | 5.685043 | 6.702109 | 2.298311 | 0 | 0 | 1.492683 | 5.005962 | 7.074744 | 1.619741 |
| Lif | 16878 | 2.590828 | 0.06511 | 5.095799 | 4.56E-05 | 0.000156 | 3.376696 | 0.06511 | 12.76374 | 1.85E-11 | 1.65E-10 | 8.161788 | 14.20856 | 5.207818 | 77.92255 | 32.30109 | 68.98605 | 608.2996 | 651.445 | 587.2185 | 33.84544 | 15.25056 | 22.39024 | 146.0072 | 121.8428 | 134.4385 |
| Hs3st4 | 628779 | 2.591926 | -4.06481 | 3.110464 | 0.005224 | 0.01327 | -0.00637 | -4.06481 | -0.00896 | 0.992933 | 0.994143 | 1.483961 | 0 | 1.735939 | 3.896127 | 7.021976 | 13.52668 | 8.527565 | 4.021265 | 11.49156 | 1.538429 | 2.287584 | 0 | 5.840288 | 4.716496 | 6.478962 |
| Tex14 | 83560 | 2.592631 | 0.458344 | 9.052081 | 9.34E-09 | 5.05E-08 | -0.15118 | 0.458344 | -0.78258 | 0.442471 | 0.541342 | 28.19527 | 24.86498 | 26.03909 | 175.3257 | 154.4835 | 188.0208 | 163.445 | 121.9784 | 201.1022 | 95.38259 | 73.9652 | 75.38047 | 51.72827 | 71.53352 | 63.97975 |
| Slc13a5 | 237831 | 2.595376 | -2.97559 | 3.778915 | 0.001076 | 0.003084 | 1.934487 | -2.97559 | 4.141137 | 0.000451 | 0.001312 | 1.483961 | 0.888035 | 1.735939 | 7.792255 | 7.021976 | 21.64268 | 49.74413 | 44.23392 | 40.22045 | 2.307643 | 0.762528 | 0.746341 | 16.68654 | 25.15464 | 14.57766 |
| Olfr391 | 258236 | 2.595657 | -4.65553 | 2.523594 | 0.019602 | 0.043787 | 0.6888 | -4.65553 | 0.803488 | 0.430551 | 0.529601 | 0 | 0 | 1.735939 | 14.2858 | 8.426372 | 0 | 5.685043 | 5.361687 | 10.3424 | 0 | 0 | 0 | 7.508942 | 7.860826 | 8.098703 |
| Klk7 | 23993 | 2.596332 | -3.43423 | 3.044211 | 0.006089 | 0.015293 | 0.718741 | -3.43423 | 1.071556 | 0.295887 | 0.389457 | 2.967923 | 0.888035 | 0 | 2.597418 | 7.021976 | 21.64268 | 15.63387 | 20.10633 | 8.044089 | 1.538429 | 2.287584 | 1.492683 | 16.68654 | 16.50774 | 15.38754 |
| Gip | 14607 | 2.596772 | -4.51285 | 2.722232 | 0.01265 | 0.029657 | 1.378497 | -4.51285 | 1.815121 | 0.083585 | 0.133875 | 0.741981 | 0 | 0 | 2.597418 | 8.426372 | 1.352668 | 2.842522 | 17.42548 | 16.08818 | 0 | 0 | 0.746341 | 12.5149 | 7.860826 | 7.288832 |
| Olfr1097 | 258840 | 2.598321 | -4.50851 | 2.45918 | 0.022536 | 0.049544 | -1.05333 | -4.50851 | -1.06707 | 0.297864 | 0.391569 | 2.967923 | 0 | 0 | 10.38967 | 2.808791 | 4.058003 | 0 | 1.340422 | 13.78987 | 0 | 1.525056 | 0 | 9.177596 | 21.22423 | 13.76779 |
| Zdhhc2 | 70546 | 2.598689 | 3.046173 | 18.74966 | 9.69E-15 | 1.34E-13 | -0.11852 | 3.046173 | -1.28661 | 0.212011 | 0.294596 | 115.749 | 154.5181 | 137.1392 | 809.0958 | 835.6152 | 964.4521 | 874.0754 | 784.1467 | 830.8395 | 128.4588 | 143.3552 | 144.7902 | 2200.12 | 2068.969 | 2234.432 |
| Nptx2 | 53324 | 2.598815 | -2.56963 | 3.605952 | 0.001627 | 0.004522 | 0.221879 | -2.56963 | 0.424102 | 0.675736 | 0.760029 | 5.193865 | 7.10428 | 0.86797 | 46.75353 | 21.06593 | 13.52668 | 32.689 | 20.10633 | 36.77298 | 1.538429 | 2.287584 | 0 | 21.6925 | 35.37372 | 34.82442 |
| Elobl | 217066 | 2.599191 | -4.27392 | 2.639933 | 0.015187 | 0.034946 | -0.492 | -4.27392 | -0.56935 | 0.575062 | 0.66882 | 2.967923 | 1.77607 | 0 | 3.896127 | 8.426372 | 21.64268 | 24.16143 | 1.340422 | 6.894934 | 0 | 1.525056 | 0 | 5.005962 | 6.288661 | 1.619741 |
| Olfr126 | 258892 | 2.599247 | -5.17217 | 2.465711 | 0.022221 | 0.048948 | -0.6054 | -5.17217 | -0.63083 | 0.534849 | 0.630217 | 1.483961 | 0 | 0 | 7.792255 | 2.808791 | 2.705335 | 0 | 2.680843 | 11.49156 | 1.538429 | 1.525056 | 0 | 0 | 0.786083 | 2.429611 |
| Gm10388 | 100038707 | 2.599548 | 1.766454 | 15.17229 | 6.56E-13 | 6.56E-12 | -0.45646 | 1.766454 | -3.83734 | 0.000936 | 0.002578 | 99.42542 | 84.36332 | 83.32508 | 602.6011 | 554.7361 | 581.6471 | 464.7523 | 508.0198 | 368.8789 | 198.4573 | 192.157 | 200.7658 | 62.57452 | 90.3995 | 78.55742 |
| Spsb1 | 74646 | 2.599606 | 3.607151 | 30.57344 | 4.13E-19 | 1.41E-17 | -0.78777 | 3.607151 | -12.0136 | 5.78E-11 | 4.71E-10 | 379.8941 | 383.6311 | 415.7574 | 2640.276 | 2380.45 | 2602.533 | 1516.485 | 1541.485 | 1522.631 | 411.5297 | 404.9023 | 485.8682 | 339.5711 | 291.6367 | 301.2717 |
| Gm10650 | 100038383 | 2.600339 | -3.74601 | 2.608889 | 0.016263 | 0.037097 | 1.436592 | -3.74601 | 1.919156 | 0.068435 | 0.113102 | 1.483961 | 0 | 0.86797 | 9.090964 | 8.426372 | 2.705335 | 24.16143 | 14.74464 | 17.23733 | 1.538429 | 12.20045 | 0 | 5.840288 | 7.860826 | 4.859222 |
| Sowahc | 268301 | 2.600632 | 5.306779 | 46.07538 | 7.40E-23 | 6.08E-21 | -0.65721 | 5.306779 | -14.771 | 1.11E-12 | 1.23E-11 | 966.0589 | 1121.588 | 986.0135 | 6650.69 | 6738.288 | 6425.172 | 4445.704 | 4401.945 | 4211.655 | 3097.627 | 2963.183 | 3022.682 | 831.8239 | 819.8842 | 846.3144 |
| Hpcal4 | 170638 | 2.601042 | -2.13772 | 3.069529 | 0.005743 | 0.01449 | 2.424232 | -2.13772 | 4.621811 | 0.000142 | 0.000448 | 6.677827 | 6.216245 | 0 | 18.18193 | 21.06593 | 13.52668 | 86.69691 | 73.7232 | 143.6444 | 5.384501 | 3.050111 | 0.746341 | 33.37308 | 40.8763 | 28.34546 |
| Gvin1 | 74558 | 2.601443 | 4.720081 | 13.04417 | 6.33E-08 | 3.07E-07 | -1.2794 | 4.720081 | -7.46218 | 1.44E-05 | 5.22E-05 | 1175.298 | 1110.932 | 512.97 | 5609.125 | 5666.735 | 5931.448 | 2444.569 | 2340.376 | 2556.871 | 2312.259 | 2389 | 2299.477 | 362.9322 | 302.6418 | 352.2936 |
| H2-T22 | 15039 | 2.60455 | 7.447185 | 48.82857 | 2.17E-23 | 2.17E-21 | 0.127943 | 7.447185 | 3.506937 | 0.00206 | 0.005263 | 4402.914 | 4362.028 | 3786.083 | 27553.41 | 27795.79 | 26147.07 | 31934.31 | 31790.78 | 29095.47 | 7182.155 | 7368.307 | 7243.988 | 4830.753 | 4799.034 | 4924.821 |
| Rgs4 | 19736 | 2.604909 | -3.2717 | 3.177843 | 0.004467 | 0.011494 | 1.296379 | -3.2717 | 2.223132 | 0.037131 | 0.067212 | 2.225942 | 5.32821 | 0 | 9.090964 | 11.23516 | 14.87935 | 42.63783 | 17.42548 | 36.77298 | 1.538429 | 1.525056 | 0 | 10.01192 | 12.57732 | 11.33818 |
| Ccdc162 | 75973 | 2.605335 | -1.23414 | 5.579061 | 1.46E-05 | 5.29E-05 | 1.373239 | -1.23414 | 5.004404 | 5.67E-05 | 0.000189 | 7.419807 | 3.55214 | 11.2836 | 45.45482 | 35.10988 | 60.87005 | 142.1261 | 115.2763 | 120.6613 | 8.461359 | 7.625279 | 9.702436 | 35.04173 | 23.58248 | 25.91585 |
| Mir194-2 | 723957 | 2.606021 | -5.2598 | 2.732879 | 0.012352 | 0.029025 | -0.91071 | -5.2598 | -1.02444 | 0.3171 | 0.412424 | 0 | 0.888035 | 0 | 1.298709 | 5.617581 | 4.058003 | 1.421261 | 0 | 5.745778 | 0 | 0 | 1.492683 | 4.171635 | 2.358248 | 1.619741 |
| Tmco5b | 75275 | 2.606066 | -4.13534 | 2.602549 | 0.016492 | 0.037538 | 1.436471 | -4.13534 | 1.859014 | 0.076869 | 0.124829 | 0 | 0 | 2.603909 | 10.38967 | 2.808791 | 2.705335 | 12.79135 | 20.10633 | 10.3424 | 2.307643 | 0 | 0 | 11.68058 | 18.86598 | 4.859222 |
| Rffl | 67338 | 2.606273 | 4.831506 | 46.21286 | 6.95E-23 | 5.82E-21 | -0.76024 | 4.831506 | -16.9135 | 7.65E-14 | 1.08E-12 | 1005.384 | 966.1821 | 931.3314 | 6384.454 | 6486.902 | 6039.661 | 3990.9 | 3922.074 | 3699.132 | 956.9028 | 976.7982 | 922.4778 | 656.6153 | 634.3687 | 615.5014 |
| Tlr11 | 239081 | 2.606667 | -1.64537 | 5.251692 | 3.15E-05 | 0.00011 | -0.879 | -1.64537 | -2.18039 | 0.040557 | 0.072441 | 2.967923 | 11.54445 | 8.679696 | 36.36386 | 44.94065 | 54.10671 | 17.05513 | 24.12759 | 34.47467 | 26.15329 | 36.60134 | 36.57072 | 13.34923 | 7.860826 | 7.288832 |
| Abcc12 | 244562 | 2.607913 | -1.88703 | 4.352853 | 0.000271 | 0.000841 | 0.506417 | -1.88703 | 1.239547 | 0.2286 | 0.313751 | 1.483961 | 7.992315 | 6.943757 | 25.97418 | 43.53625 | 24.34802 | 42.63783 | 34.85097 | 57.45778 | 6.92293 | 3.812639 | 8.209754 | 18.35519 | 36.94588 | 23.48624 |
| Plat | 18791 | 2.608215 | -0.4315 | 4.661305 | 0.000129 | 0.000416 | 5.633635 | -0.4315 | 19.41092 | 4.81E-15 | 8.80E-14 | 3.709904 | 8.88035 | 4.339848 | 25.97418 | 36.51428 | 48.69604 | 1937.179 | 1778.74 | 1911.046 | 13.07665 | 3.050111 | 2.239024 | 78.42673 | 99.04641 | 89.8956 |
| Aqp4 | 11829 | 2.608382 | -3.05344 | 3.047441 | 0.006044 | 0.015188 | 0.669664 | -3.05344 | 1.044996 | 0.307717 | 0.402189 | 0 | 7.992315 | 3.471878 | 16.88322 | 14.04395 | 10.82134 | 18.47639 | 16.08506 | 37.92213 | 1.538429 | 0 | 1.492683 | 30.8701 | 21.22423 | 17.00728 |
| Osmr | 18414 | 2.60876 | -0.45777 | 5.87067 | 7.43E-06 | 2.79E-05 | 3.795115 | -0.45777 | 16.97681 | 7.10E-14 | 1.01E-12 | 12.61367 | 6.216245 | 13.88751 | 61.03933 | 53.36702 | 105.5081 | 1083.001 | 911.4868 | 1065.267 | 6.92293 | 3.812639 | 5.224389 | 34.2074 | 32.22939 | 42.11325 |
| Gpr37 | 14763 | 2.609135 | -2.47589 | 3.778458 | 0.001078 | 0.003087 | 0.543809 | -2.47589 | 1.062483 | 0.299891 | 0.393644 | 4.451884 | 1.77607 | 0.86797 | 20.77935 | 15.44835 | 10.82134 | 34.11026 | 13.40422 | 27.57973 | 7.692145 | 6.862751 | 2.239024 | 25.86413 | 33.80155 | 27.53559 |
| Hoxc5 | 15424 | 2.609859 | -2.64197 | 3.302246 | 0.00334 | 0.0088 | 1.453892 | -2.64197 | 2.5528 | 0.018392 | 0.036688 | 2.967923 | 1.77607 | 0 | 6.493546 | 7.021976 | 16.23201 | 22.74017 | 24.12759 | 34.47467 | 3.076858 | 4.575167 | 5.224389 | 42.55067 | 59.74228 | 34.01455 |
| Gm3149 | 100041119 | 2.610092 | -5.41713 | 2.831463 | 0.009897 | 0.023796 | 0.478181 | -5.41713 | 0.598775 | 0.555625 | 0.650229 | 0 | 0 | 0 | 3.896127 | 0 | 5.410671 | 5.685043 | 2.680843 | 2.298311 | 0 | 0 | 0 | 4.171635 | 1.572165 | 3.239481 |
| Vmn2r44 | 434113 | 2.610178 | -4.65863 | 2.626844 | 0.015632 | 0.035849 | -0.02378 | -4.65863 | -0.02769 | 0.978165 | 0.981673 | 3.709904 | 0 | 0 | 1.298709 | 5.617581 | 18.93735 | 1.421261 | 9.382952 | 10.3424 | 0 | 0 | 0 | 5.840288 | 4.716496 | 4.859222 |
| Timd4 | 276891 | 2.611202 | -2.19002 | 4.815006 | 8.91E-05 | 0.000293 | 0.325849 | -2.19002 | 0.887015 | 0.384957 | 0.483629 | 8.161788 | 1.77607 | 6.943757 | 27.27289 | 36.51428 | 39.22736 | 49.74413 | 54.95729 | 33.32551 | 2.307643 | 4.575167 | 5.97073 | 9.177596 | 13.3634 | 8.908573 |
| Svs5 | 20944 | 2.612123 | -5.17113 | 2.478143 | 0.021632 | 0.047807 | 0.432752 | -5.17113 | 0.481288 | 0.635209 | 0.724295 | 0 | 0 | 0.86797 | 5.194837 | 4.213186 | 1.352668 | 2.842522 | 8.04253 | 4.596622 | 0 | 0 | 0 | 8.343269 | 4.716496 | 0 |
| Olfr1420 | 258405 | 2.613132 | -3.42455 | 2.917377 | 0.008144 | 0.019917 | 0.709119 | -3.42455 | 0.999473 | 0.328763 | 0.425113 | 1.483961 | 1.77607 | 0 | 10.38967 | 2.808791 | 12.17401 | 12.79135 | 14.74464 | 11.49156 | 3.076858 | 3.050111 | 8.956095 | 5.005962 | 4.716496 | 21.8665 |
| Olfr601 | 258311 | 2.613311 | -5.34596 | 2.514369 | 0.019999 | 0.044549 | 0.448322 | -5.34596 | 0.495516 | 0.6253 | 0.715853 | 0 | 0 | 0 | 2.597418 | 8.426372 | 0 | 1.421261 | 1.340422 | 11.49156 | 0 | 0 | 0 | 3.337308 | 2.358248 | 7.288832 |
| Vmn1r13 | 113862 | 2.613311 | -5.48176 | 2.493304 | 0.020934 | 0.046405 | -0.39116 | -5.48176 | -0.40836 | 0.687079 | 0.769825 | 0 | 0 | 0 | 2.597418 | 8.426372 | 0 | 2.842522 | 0 | 3.447467 | 0 | 1.525056 | 0 | 3.337308 | 2.358248 | 1.619741 |
| Fgf9 | 14180 | 2.614994 | -1.91165 | 3.155608 | 0.004705 | 0.012055 | 0.796093 | -1.91165 | 1.423682 | 0.168994 | 0.243386 | 8.903769 | 3.55214 | 0.86797 | 22.07806 | 14.04395 | 41.9327 | 52.58665 | 28.14886 | 52.86116 | 3.076858 | 1.525056 | 27.61463 | 43.385 | 55.02578 | 40.49351 |
| 1700093J2 | 74296 | 2.615034 | -3.42491 | 2.994519 | 0.006826 | 0.016972 | 0.92304 | -3.42491 | 1.353871 | 0.189941 | 0.268511 | 0.741981 | 2.664105 | 0 | 12.98709 | 4.213186 | 5.410671 | 14.21261 | 17.42548 | 11.49156 | 3.076858 | 1.525056 | 10.44878 | 11.68058 | 14.14949 | 5.669092 |
| Ifi204 | 15951 | 2.615389 | 8.523386 | 81.64766 | 3.95E-28 | 2.42E-25 | -0.66477 | 8.523386 | -28.0298 | 2.52E-18 | 1.01E-16 | 10867.05 | 11110.21 | 9678.729 | 69338.08 | 70514.69 | 67165.36 | 44803.83 | 44223.19 | 46276.5 | 19096.52 | 18787.16 | 18598.82 | 8230.635 | 8549.435 | 8249.339 |
| 1700022A2 | 72252 | 2.615414 | -4.45294 | 3.018 | 0.006468 | 0.016147 | -0.12773 | -4.45294 | -0.17062 | 0.866126 | 0.918721 | 0 | 1.77607 | 1.735939 | 3.896127 | 11.23516 | 8.116006 | 11.37009 | 2.680843 | 9.193245 | 0 | 0.762528 | 0 | 8.343269 | 1.572165 | 3.239481 |
| Sh2d6 | 71130 | 2.616223 | 0.665096 | 13.62007 | 5.34E-12 | 4.60E-11 | -0.95604 | 0.665096 | -6.51364 | 1.73E-06 | 7.15E-06 | 66.03629 | 70.15476 | 49.47427 | 401.3011 | 417.1054 | 393.6263 | 225.9805 | 238.5951 | 190.7598 | 46.15287 | 41.1765 | 29.85365 | 43.385 | 26.72681 | 39.68364 |
| Gm10764 | 100038674 | 2.617595 | -3.52735 | 2.515268 | 0.01996 | 0.044481 | 0.714519 | -3.52735 | 0.875847 | 0.390864 | 0.489762 | 11.12971 | 0 | 0 | 10.38967 | 4.213186 | 12.17401 | 9.948826 | 9.382952 | 28.72889 | 0 | 1.525056 | 2.985365 | 23.36115 | 16.50774 | 10.52831 |
| Eif1-ps1 | 100040439 | 2.618883 | -3.13666 | 2.928108 | 0.007947 | 0.019481 | -1.15349 | -3.13666 | -1.42452 | 0.168755 | 0.243107 | 3.709904 | 0 | 12.15157 | 15.58451 | 16.85274 | 22.99535 | 9.948826 | 2.680843 | 17.23733 | 9.230573 | 7.625279 | 1.492683 | 4.171635 | 12.57732 | 3.239481 |
| 4731417B2 | 74576 | 2.619243 | -4.26573 | 2.997545 | 0.006779 | 0.016859 | 0.16754 | -4.26573 | 0.227215 | 0.822419 | 0.889582 | 0 | 1.77607 | 1.735939 | 5.194837 | 5.617581 | 12.17401 | 5.685043 | 4.021265 | 21.83396 | 0 | 0 | 0 | 8.343269 | 16.50774 | 4.859222 |
| I730028E1 | 619328 | 2.620211 | -3.58948 | 2.73682 | 0.012244 | 0.028794 | -0.10933 | -3.58948 | -0.13342 | 0.895114 | 0.92056 | 0 | 4.440175 | 0.86797 | 5.194837 | 9.830767 | 9.468674 | 7.106304 | 10.72337 | 5.745778 | 6.153716 | 0 | 1.492683 | 23.36115 | 11.00516 | 25.91585 |
| Gm20554 | 328287 | 2.62265 | -1.82905 | 3.425016 | 0.002501 | 0.006729 | 0.894423 | -1.82905 | 1.721992 | 0.099534 | 0.155315 | 1.483961 | 5.32821 | 3.471878 | 28.5716 | 15.44835 | 22.99535 | 29.84648 | 64.34024 | 40.22045 | 3.076858 | 12.20045 | 9.702436 | 70.91779 | 67.60311 | 29.9652 |
| Ces2f | 71903 | 2.625699 | -2.95068 | 2.970588 | 0.007211 | 0.017832 | 2.58776 | -2.95068 | 4.295424 | 0.000311 | 0.000929 | 5.193865 | 0 | 0 | 6.493546 | 5.617581 | 6.763339 | 48.32287 | 60.31898 | 28.72889 | 8.461359 | 4.575167 | 7.463413 | 12.5149 | 13.3634 | 10.52831 |
| Vmn1r233 | 171236 | 2.625963 | -4.37027 | 2.89489 | 0.008572 | 0.020871 | 1.697777 | -4.37027 | 2.461414 | 0.022428 | 0.04353 | 0 | 0 | 1.735939 | 2.597418 | 2.808791 | 8.116006 | 28.42522 | 6.702109 | 17.23733 | 0 | 0.762528 | 0 | 6.674615 | 6.288661 | 7.288832 |
| Stk39 | 53416 | 2.626364 | 2.474978 | 21.26114 | 7.55E-16 | 1.30E-14 | 0.80115 | 2.474978 | 11.01235 | 2.89E-10 | 2.09E-09 | 159.5259 | 148.3018 | 131.0634 | 933.7719 | 994.3118 | 977.9788 | 1725.411 | 1869.888 | 1672.021 | 186.1499 | 179.9566 | 185.839 | 81.76404 | 81.75259 | 102.0437 |
| Sost | 74499 | 2.626524 | -2.36599 | 3.239311 | 0.00387 | 0.010084 | 0.836657 | -2.36599 | 1.445956 | 0.162715 | 0.235828 | 5.193865 | 5.32821 | 0 | 18.18193 | 21.06593 | 9.468674 | 28.42522 | 33.51054 | 26.43058 | 3.846072 | 3.812639 | 5.97073 | 39.21337 | 55.02578 | 23.48624 |
| Mir652 | 723976 | 2.627259 | -4.49823 | 3.48136 | 0.002188 | 0.005953 | -1.97939 | -4.49823 | -2.64918 | 0.01488 | 0.03051 | 0.741981 | 1.77607 | 2.603909 | 11.68838 | 14.04395 | 10.82134 | 2.842522 | 1.340422 | 4.596622 | 2.307643 | 2.287584 | 0 | 2.502981 | 0.786083 | 0 |
| Nt5el | 66763 | 2.62787 | -2.57945 | 2.958287 | 0.007417 | 0.018289 | 1.063384 | -2.57945 | 1.74797 | 0.094842 | 0.149032 | 2.967923 | 4.440175 | 0.86797 | 12.98709 | 21.06593 | 20.29002 | 31.26774 | 48.25518 | 39.07129 | 12.30743 | 5.337695 | 0 | 21.6925 | 23.58248 | 8.098703 |
| Helz2 | 229003 | 2.628942 | 10.20385 | 27.01424 | 8.75E-11 | 6.33E-10 | 0.432336 | 10.20385 | 8.437509 | 6.71E-06 | 2.55E-05 | 34362.61 | 35257.65 | 33614.73 | 227297.5 | 230097.5 | 224233.1 | 333838.5 | 333565.3 | 296662.6 | 35941.55 | 37186.96 | 34099.59 | 19282.13 | 19838.37 | 19193.93 |
| Galp | 232836 | 2.629457 | -4.19458 | 3.02953 | 0.006299 | 0.015765 | 0.757532 | -4.19458 | 1.046802 | 0.306902 | 0.401383 | 0 | 0.888035 | 0 | 2.597418 | 1.404395 | 8.116006 | 4.263783 | 6.702109 | 8.044089 | 2.307643 | 1.525056 | 1.492683 | 18.35519 | 8.646909 | 7.288832 |
| Sntn | 218739 | 2.629845 | -4.71865 | 3.214215 | 0.004104 | 0.010636 | -0.8854 | -4.71865 | -1.18667 | 0.248399 | 0.33623 | 1.483961 | 1.77607 | 0 | 10.38967 | 5.617581 | 6.763339 | 4.263783 | 4.021265 | 3.447467 | 2.307643 | 0 | 0 | 0.834327 | 3.930413 | 0.80987 |
| Csn2 | 12991 | 2.630429 | -2.8964 | 2.81067 | 0.010373 | 0.024792 | 0.43491 | -2.8964 | 0.569819 | 0.57475 | 0.668576 | 1.483961 | 0 | 1.735939 | 12.98709 | 5.617581 | 5.410671 | 15.63387 | 2.680843 | 24.13227 | 19.23036 | 34.31375 | 40.30243 | 12.5149 | 7.074744 | 10.52831 |
| Bcl2a1b | 12045 | 2.630759 | 7.026836 | 49.96878 | 1.33E-23 | 1.39E-21 | -0.12937 | 7.026836 | -3.43745 | 0.002428 | 0.006119 | 3333.719 | 3300.826 | 2816.561 | 20697.53 | 21547.64 | 20242.67 | 20837.11 | 19542.01 | 19065.64 | 6909.853 | 6688.894 | 7145.471 | 3308.941 | 3086.946 | 3247.58 |
| Bmp15 | 12155 | 2.631641 | -3.6907 | 3.069985 | 0.005737 | 0.014478 | 0.300098 | -3.6907 | 0.434213 | 0.668492 | 0.753719 | 2.967923 | 1.77607 | 0 | 15.58451 | 16.85274 | 2.705335 | 14.21261 | 12.0638 | 10.3424 | 1.538429 | 1.525056 | 0 | 11.68058 | 7.860826 | 10.52831 |
| Gpr52 | 620246 | 2.6337 | -1.70908 | 4.2117 | 0.00038 | 0.001161 | -0.04606 | -1.70908 | -0.10321 | 0.91876 | 0.93432 | 8.903769 | 7.10428 | 1.735939 | 38.96127 | 39.32307 | 28.40602 | 27.00396 | 34.85097 | 44.81707 | 15.38429 | 27.451 | 5.224389 | 21.6925 | 25.94073 | 18.62702 |
| Gm13417 | 100504544 | 2.633962 | -2.92909 | 4.058428 | 0.00055 | 0.001643 | -1.38799 | -2.92909 | -2.38016 | 0.026694 | 0.050601 | 8.903769 | 0.888035 | 4.339848 | 31.16902 | 21.06593 | 25.70069 | 12.79135 | 17.42548 | 4.596622 | 3.076858 | 10.67539 | 3.731706 | 5.005962 | 4.716496 | 4.049351 |
| Olfr1256 | 258985 | 2.634416 | -5.12282 | 2.603369 | 0.016462 | 0.037474 | 1.173208 | -5.12282 | 1.39753 | 0.176613 | 0.25252 | 0 | 0 | 0 | 0 | 1.404395 | 14.87935 | 4.263783 | 5.361687 | 8.044089 | 0 | 0 | 0 | 6.674615 | 3.14433 | 5.669092 |
| Pax3 | 18505 | 2.634783 | -2.6428 | 4.079121 | 0.000523 | 0.001569 | 0.508367 | -2.6428 | 1.051473 | 0.304801 | 0.399212 | 3.709904 | 1.77607 | 0.86797 | 12.98709 | 12.63956 | 18.93735 | 18.47639 | 18.7659 | 28.72889 | 3.846072 | 4.575167 | 1.492683 | 38.37904 | 30.65722 | 23.48624 |
| Bambi-ps1 | 81913 | 2.635723 | 3.759272 | 33.29739 | 6.92E-20 | 2.80E-18 | -0.6983 | 3.759272 | -11.5058 | 1.29E-10 | 9.92E-10 | 569.0992 | 534.5971 | 482.5911 | 3488.333 | 3659.854 | 3404.665 | 2248.435 | 2381.929 | 2135.131 | 508.4508 | 459.0418 | 421.6828 | 188.5579 | 188.6598 | 161.9741 |
| Fndc3a | 319448 | 2.63582 | 8.918797 | 106.9294 | 1.26E-30 | 4.16E-27 | -1.33884 | 8.918797 | -64.952 | 5.11E-26 | 3.13E-23 | 13278.49 | 13847.13 | 13252.16 | 89756.39 | 90448.67 | 87518.96 | 36915.83 | 37033.17 | 36012.24 | 25099.47 | 25525.62 | 24774.05 | 17892.98 | 18217.46 | 18860.26 |
| Gem | 14579 | 2.636099 | 1.095991 | 8.207792 | 4.83E-08 | 2.38E-07 | 3.399012 | 1.095991 | 21.11937 | 8.66E-16 | 1.91E-14 | 12.61367 | 28.41712 | 23.43518 | 146.7541 | 127.8 | 127.1508 | 1483.796 | 1415.485 | 1512.289 | 99.22866 | 99.89115 | 139.5658 | 68.41481 | 85.68301 | 68.83897 |
| Olfr870 | 57251 | 2.638219 | -4.98327 | 2.589604 | 0.016968 | 0.038496 | -0.84587 | -4.98327 | -0.90339 | 0.376401 | 0.474784 | 1.483961 | 0 | 0 | 7.792255 | 1.404395 | 5.410671 | 7.106304 | 0 | 3.447467 | 0 | 0 | 1.492683 | 1.668654 | 7.860826 | 3.239481 |
| Myh2 | 17882 | 2.63875 | -1.57955 | 4.415604 | 0.000233 | 0.000729 | 1.272979 | -1.57955 | 3.441436 | 0.002405 | 0.006069 | 7.419807 | 2.664105 | 3.471878 | 23.37676 | 28.08791 | 41.9327 | 52.58665 | 87.12741 | 93.0816 | 6.92293 | 7.625279 | 2.985365 | 45.05365 | 67.60311 | 43.73299 |
| Vmn2r3 | 637004 | 2.638817 | -3.62339 | 3.554951 | 0.001837 | 0.005062 | -0.40477 | -3.62339 | -0.63347 | 0.53316 | 0.628761 | 0.741981 | 3.55214 | 0.86797 | 14.2858 | 14.04395 | 6.763339 | 4.263783 | 13.40422 | 10.3424 | 0 | 0.762528 | 0.746341 | 23.36115 | 20.43815 | 10.52831 |
| Meox1 | 17285 | 2.63967 | -3.05862 | 3.011631 | 0.006563 | 0.016365 | 0.475265 | -3.05862 | 0.685608 | 0.500341 | 0.59813 | 5.935846 | 0.888035 | 0 | 9.090964 | 15.44835 | 6.763339 | 9.948826 | 6.702109 | 37.92213 | 3.076858 | 4.575167 | 2.985365 | 25.02981 | 20.43815 | 17.00728 |
| Dppa1 | 347708 | 2.640715 | -3.368 | 3.204375 | 0.004199 | 0.010862 | 0.492364 | -3.368 | 0.753233 | 0.459539 | 0.558415 | 0 | 4.440175 | 1.735939 | 6.493546 | 19.66153 | 6.763339 | 8.527565 | 16.08506 | 19.53565 | 2.307643 | 1.525056 | 1.492683 | 9.177596 | 11.00516 | 19.43689 |
| Cfap54 | 380654 | 2.640742 | 0.155357 | 8.267553 | 4.28E-08 | 2.12E-07 | -0.7357 | 0.155357 | -3.10244 | 0.005322 | 0.012349 | 24.48536 | 29.30515 | 24.30315 | 166.2348 | 158.6967 | 200.1948 | 86.69691 | 95.16994 | 144.7936 | 35.38386 | 31.26364 | 13.43414 | 124.3147 | 96.68816 | 98.80417 |
| Ccdc42 | 276920 | 2.643261 | -3.55363 | 2.801246 | 0.010595 | 0.025251 | 1.420703 | -3.55363 | 2.049582 | 0.052893 | 0.090942 | 1.483961 | 1.77607 | 0 | 9.090964 | 2.808791 | 14.87935 | 29.84648 | 20.10633 | 18.38649 | 2.307643 | 0 | 5.97073 | 10.84625 | 8.646909 | 4.859222 |
| Gpd2 | 14571 | 2.643608 | 6.177473 | 65.63396 | 4.10E-26 | 1.06E-23 | -2.8864 | 6.177473 | -62.1527 | 1.30E-25 | 5.54E-23 | 3537.764 | 3594.766 | 3357.306 | 23668.97 | 24054.48 | 22302.79 | 3452.243 | 3284.033 | 3119.957 | 4259.14 | 4232.03 | 4481.779 | 1262.337 | 1278.956 | 1388.928 |
| Ccna1 | 12427 | 2.644376 | -3.62816 | 2.517796 | 0.01985 | 0.044278 | 1.182721 | -3.62816 | 1.492206 | 0.15028 | 0.220626 | 0.741981 | 3.55214 | 0 | 5.194837 | 15.44835 | 5.410671 | 18.47639 | 9.382952 | 34.47467 | 1.538429 | 0 | 2.239024 | 13.34923 | 25.15464 | 3.239481 |
| Gm16527 | 669780 | 2.645091 | -4.43726 | 2.676516 | 0.014005 | 0.032479 | -0.26766 | -4.43726 | -0.30614 | 0.762468 | 0.836831 | 0 | 0 | 2.603909 | 10.38967 | 1.404395 | 5.410671 | 2.842522 | 2.680843 | 6.894934 | 3.076858 | 0 | 2.239024 | 3.337308 | 7.860826 | 5.669092 |
| Lck | 16818 | 2.646927 | 1.342421 | 9.21252 | 6.90E-09 | 3.81E-08 | 3.021559 | 1.342421 | 20.97917 | 9.92E-16 | 2.14E-14 | 24.48536 | 53.2821 | 24.30315 | 250.6509 | 221.8945 | 166.3781 | 1768.048 | 1837.718 | 1736.374 | 36.15308 | 47.27673 | 54.48291 | 141.0012 | 138.3505 | 156.305 |
| Hspb3 | 56534 | 2.647227 | -4.4998 | 2.597203 | 0.016687 | 0.037916 | -0.28607 | -4.4998 | -0.3189 | 0.7529 | 0.828675 | 0 | 5.32821 | 0 | 7.792255 | 8.426372 | 2.705335 | 2.842522 | 8.04253 | 4.596622 | 3.076858 | 0.762528 | 0 | 6.674615 | 0.786083 | 4.049351 |
| Src | 20779 | 2.648562 | 3.460343 | 24.93516 | 2.85E-17 | 6.48E-16 | -0.95248 | 3.460343 | -11.6704 | 9.92E-11 | 7.78E-10 | 256.7253 | 230.8891 | 240.4276 | 1592.217 | 1530.791 | 1778.758 | 797.3273 | 869.9337 | 946.9042 | 885.3658 | 886.8199 | 947.8534 | 370.4412 | 425.2707 | 388.7377 |
| Spta1 | 20739 | 2.649746 | 0.422393 | 9.274645 | 6.15E-09 | 3.41E-08 | 1.592809 | 0.422393 | 10.32451 | 9.30E-10 | 6.22E-09 | 23.74338 | 16.87266 | 23.43518 | 149.3516 | 117.9692 | 170.4361 | 478.9649 | 427.5945 | 457.3639 | 27.69172 | 26.68847 | 23.13658 | 88.43865 | 95.90208 | 81.7969 |
| Exoc3l4 | 74190 | 2.649767 | 1.465398 | 12.5527 | 2.53E-11 | 1.97E-10 | 0.305905 | 1.465398 | 2.383241 | 0.02652 | 0.050321 | 80.8759 | 58.61031 | 61.62584 | 451.9508 | 393.2307 | 516.7191 | 649.5162 | 608.5515 | 506.7776 | 87.69045 | 70.91509 | 72.3951 | 80.92971 | 110.8376 | 80.17716 |
| Gm10503 | 100038436 | 2.650458 | -4.71967 | 3.035564 | 0.006212 | 0.015571 | -1.85413 | -4.71967 | -2.17961 | 0.040622 | 0.072538 | 2.225942 | 0.888035 | 0 | 7.792255 | 5.617581 | 8.116006 | 5.685043 | 2.680843 | 0 | 0 | 0 | 1.492683 | 3.337308 | 6.288661 | 3.239481 |
| Gm9915 | 791306 | 2.651067 | -3.99383 | 3.08843 | 0.005498 | 0.013927 | 0.607015 | -3.99383 | 0.870913 | 0.393492 | 0.492422 | 0.741981 | 1.77607 | 0 | 5.194837 | 8.426372 | 5.410671 | 11.37009 | 5.361687 | 16.08818 | 0 | 3.050111 | 0 | 10.01192 | 11.00516 | 16.19741 |
| Oas1f | 243262 | 2.651184 | -3.33084 | 2.968506 | 0.007246 | 0.017908 | 1.728182 | -3.33084 | 2.765857 | 0.011473 | 0.024338 | 1.483961 | 3.55214 | 0 | 5.194837 | 11.23516 | 13.52668 | 24.16143 | 49.5956 | 31.0272 | 1.538429 | 0 | 3.731706 | 15.01788 | 11.00516 | 5.669092 |
| Pln | 18821 | 2.651295 | 1.794739 | 19.7158 | 3.51E-15 | 5.24E-14 | -1.03873 | 1.794739 | -10.0228 | 1.58E-09 | 1.02E-08 | 105.3613 | 118.1087 | 104.1564 | 676.6275 | 720.4548 | 806.19 | 373.7916 | 375.3181 | 360.8349 | 143.8431 | 138.0175 | 140.3122 | 87.60433 | 86.46909 | 99.61404 |
| Elovl2 | 54326 | 2.651973 | 0.140656 | 6.954145 | 6.59E-07 | 2.80E-06 | 0.993938 | 0.140656 | 4.548233 | 0.000169 | 0.000528 | 19.2915 | 17.7607 | 19.9633 | 151.949 | 153.0791 | 93.33407 | 264.3545 | 282.829 | 260.8583 | 23.84565 | 18.30067 | 27.61463 | 80.09538 | 51.09537 | 115.0016 |
| Tas2r108 | 57253 | 2.652102 | -3.22321 | 2.7399 | 0.01216 | 0.028624 | 0.61608 | -3.22321 | 0.838498 | 0.411044 | 0.510067 | 3.709904 | 3.55214 | 0 | 22.07806 | 19.66153 | 4.058003 | 7.106304 | 21.44675 | 43.66791 | 1.538429 | 1.525056 | 0 | 20.85817 | 14.14949 | 24.29611 |
| Mmp2 | 17390 | 2.652753 | 1.729152 | 17.18033 | 5.60E-14 | 6.75E-13 | -1.49184 | 1.729152 | -11.3576 | 1.64E-10 | 1.24E-09 | 124.6528 | 127.877 | 102.4204 | 796.1087 | 852.4679 | 739.9093 | 326.89 | 235.9142 | 321.7636 | 80.76752 | 102.1787 | 64.18535 | 149.3445 | 133.634 | 127.1496 |
| Olfr175 | 259004 | 2.654967 | -4.32385 | 2.542492 | 0.018811 | 0.042228 | -0.64356 | -4.32385 | -0.67982 | 0.503923 | 0.601538 | 1.483961 | 0 | 0 | 2.597418 | 9.830767 | 2.705335 | 5.685043 | 0 | 6.894934 | 4.615287 | 9.912862 | 2.985365 | 1.668654 | 9.432991 | 2.429611 |
| Peg10 | 170676 | 2.65592 | 2.579311 | 8.730411 | 1.73E-08 | 8.99E-08 | 4.916149 | 2.579311 | 34.03373 | 4.37E-20 | 2.96E-18 | 43.77686 | 23.08891 | 16.49142 | 200.0012 | 161.5055 | 175.8468 | 5736.209 | 5769.175 | 5407.926 | 68.46009 | 57.95212 | 64.18535 | 3523.363 | 3402.166 | 3503.499 |
| 3110009M | 73079 | 2.657336 | -4.10518 | 3.414661 | 0.002563 | 0.006882 | -0.41481 | -4.10518 | -0.62373 | 0.539412 | 0.634569 | 1.483961 | 0.888035 | 1.735939 | 5.194837 | 28.08791 | 9.468674 | 14.21261 | 8.04253 | 5.745778 | 1.538429 | 1.525056 | 0 | 0.834327 | 2.358248 | 3.239481 |
| 2900052N | 73040 | 2.657695 | -3.6011 | 2.984461 | 0.006986 | 0.017337 | -0.16995 | -3.6011 | -0.22942 | 0.820729 | 0.888166 | 3.709904 | 0.888035 | 0.86797 | 14.2858 | 5.617581 | 22.99535 | 2.842522 | 13.40422 | 29.87805 | 0 | 3.050111 | 0 | 12.5149 | 10.21907 | 14.57766 |
| 9530091C | 320440 | 2.658323 | -2.08076 | 3.033031 | 0.006248 | 0.015646 | 0.834126 | -2.08076 | 1.388342 | 0.179354 | 0.255841 | 7.419807 | 0 | 7.811726 | 24.67547 | 14.04395 | 24.34802 | 34.11026 | 24.12759 | 63.20356 | 6.92293 | 1.525056 | 10.44878 | 48.39096 | 52.66754 | 30.77507 |
| Krtap4-7 | 76444 | 2.661541 | -3.86447 | 2.505516 | 0.020387 | 0.045341 | 1.519137 | -3.86447 | 1.856532 | 0.077236 | 0.125347 | 0 | 0 | 1.735939 | 1.298709 | 11.23516 | 4.058003 | 7.106304 | 20.10633 | 16.08818 | 3.076858 | 3.050111 | 0 | 25.02981 | 12.57732 | 9.718443 |
| Pdcd1 | 18566 | 2.661642 | -1.33312 | 3.90555 | 0.000794 | 0.002318 | 5.313736 | -1.33312 | 15.07957 | 7.40E-13 | 8.47E-12 | 4.451884 | 1.77607 | 13.01954 | 40.25998 | 29.4923 | 35.16936 | 1473.848 | 1505.294 | 1420.356 | 7.692145 | 0 | 1.492683 | 26.69846 | 14.14949 | 8.908573 |
| Cd200 | 17470 | 2.662341 | 0.992966 | 13.94861 | 3.37E-12 | 3.00E-11 | -1.6181 | 0.992966 | -9.73471 | 2.64E-09 | 1.64E-08 | 88.29571 | 70.15476 | 61.62584 | 445.4572 | 557.5449 | 499.1344 | 183.3426 | 167.5527 | 156.2852 | 126.9204 | 107.5164 | 129.8634 | 19.18952 | 43.23454 | 37.25403 |
| Sppl2c | 237958 | 2.662836 | -3.03026 | 2.762625 | 0.011556 | 0.027316 | 0.58826 | -3.03026 | 0.779648 | 0.444158 | 0.542986 | 2.967923 | 0 | 1.735939 | 2.597418 | 9.830767 | 31.11136 | 8.527565 | 9.382952 | 36.77298 | 7.692145 | 3.050111 | 6.717071 | 11.68058 | 15.72165 | 19.43689 |
| Cyp2c53-p | 638988 | 2.662902 | -3.22475 | 3.743861 | 0.001171 | 0.003332 | 0.671015 | -3.22475 | 1.211137 | 0.239083 | 0.325676 | 0.741981 | 0.888035 | 1.735939 | 10.38967 | 11.23516 | 6.763339 | 7.106304 | 17.42548 | 27.57973 | 3.846072 | 5.337695 | 2.985365 | 11.68058 | 10.21907 | 8.908573 |
| Gm9271 | 668618 | 2.663596 | -2.73005 | 3.085511 | 0.005535 | 0.014013 | 0.356811 | -2.73005 | 0.549725 | 0.588215 | 0.681736 | 0 | 5.32821 | 8.679696 | 15.58451 | 28.08791 | 10.82134 | 19.89765 | 16.08506 | 33.32551 | 6.153716 | 1.525056 | 0.746341 | 17.52087 | 22.01031 | 29.9652 |
| Tnfsf8 | 21949 | 2.663778 | 0.657648 | 11.09724 | 2.51E-10 | 1.69E-09 | 1.548133 | 0.657648 | 12.0377 | 5.57E-11 | 4.54E-10 | 32.64715 | 31.96926 | 39.9266 | 206.4948 | 255.5999 | 251.5962 | 722.0005 | 760.0191 | 681.4493 | 22.30722 | 35.07628 | 20.89756 | 53.39692 | 50.30929 | 40.49351 |
| 6330407A | 70720 | 2.664714 | 3.630604 | 30.9728 | 3.15E-19 | 1.10E-17 | -1.61243 | 3.630604 | -20.9543 | 1.02E-15 | 2.19E-14 | 617.328 | 674.9066 | 680.4882 | 4438.988 | 4359.243 | 4512.5 | 1638.714 | 1525.4 | 1385.882 | 419.9911 | 411.0025 | 287.3414 | 141.8356 | 147.7835 | 121.4805 |
| Gfi1 | 14581 | 2.666102 | 1.488833 | 13.56937 | 5.73E-12 | 4.93E-11 | -1.2273 | 1.488833 | -7.82258 | 1.05E-07 | 5.15E-07 | 71.23015 | 68.37869 | 107.6282 | 518.185 | 535.0746 | 588.4105 | 306.9923 | 221.1696 | 214.8921 | 128.4588 | 99.12862 | 83.59022 | 113.4685 | 107.6933 | 137.6779 |
| Mcpt2 | 17225 | 2.666327 | -3.39219 | 4.476136 | 0.000201 | 0.000635 | -0.79598 | -3.39219 | -1.51487 | 0.144475 | 0.213139 | 1.483961 | 1.77607 | 1.735939 | 15.58451 | 14.04395 | 10.82134 | 7.106304 | 8.04253 | 8.044089 | 1.538429 | 0.762528 | 2.985365 | 12.5149 | 14.93557 | 9.718443 |
| Wfdc9 | 629754 | 2.668625 | -4.83279 | 2.676012 | 0.014021 | 0.032507 | 1.442252 | -4.83279 | 1.783193 | 0.088784 | 0.141056 | 0 | 0 | 0 | 0 | 9.830767 | 2.705335 | 7.106304 | 6.702109 | 8.044089 | 0 | 0 | 1.492683 | 5.005962 | 5.502578 | 8.908573 |
| Zfp811 | 240063 | 2.670073 | 3.167273 | 36.2069 | 1.19E-20 | 5.98E-19 | -2.21206 | 3.167273 | -29.6769 | 7.68E-19 | 3.63E-17 | 609.1662 | 690.0032 | 678.7522 | 4511.716 | 4421.036 | 4442.161 | 888.288 | 1065.635 | 1038.837 | 487.682 | 487.2553 | 445.5657 | 47.55663 | 20.43815 | 33.20468 |
| Mir466f-1 | 100124486 | 2.670135 | -5.27517 | 2.965451 | 0.007297 | 0.018018 | 0.320049 | -5.27517 | 0.417823 | 0.68025 | 0.763782 | 0 | 0 | 1.735939 | 5.194837 | 1.404395 | 8.116006 | 4.263783 | 4.021265 | 9.193245 | 2.307643 | 0 | 0.746341 | 0 | 0 | 0 |
| Duoxa2 | 66811 | 2.670514 | -2.7626 | 3.825346 | 0.000963 | 0.002777 | 1.089411 | -2.7626 | 2.194542 | 0.039392 | 0.070639 | 0.741981 | 0.888035 | 5.207818 | 7.792255 | 21.06593 | 13.52668 | 35.53152 | 21.44675 | 33.32551 | 3.076858 | 3.050111 | 1.492683 | 18.35519 | 21.22423 | 27.53559 |
| 2310005A | 100504268 | 2.670594 | -4.21103 | 2.754038 | 0.011781 | 0.027799 | 0.280473 | -4.21103 | 0.345171 | 0.733344 | 0.811703 | 0 | 5.32821 | 0 | 5.194837 | 5.617581 | 6.763339 | 7.106304 | 10.72337 | 5.745778 | 0 | 3.050111 | 0 | 15.01788 | 4.716496 | 7.288832 |
| Mir674 | 732489 | 2.670977 | -0.47655 | 10.43624 | 7.66E-10 | 4.83E-09 | -2.12124 | -0.47655 | -8.37116 | 3.49E-08 | 1.83E-07 | 24.48536 | 47.95389 | 34.71878 | 215.5857 | 231.7252 | 250.2435 | 59.69296 | 60.31898 | 47.11538 | 23.07643 | 34.31375 | 38.0634 | 5.840288 | 11.00516 | 14.57766 |
| Olfr623 | 259126 | 2.672545 | -4.75751 | 2.565618 | 0.017884 | 0.040351 | 1.118325 | -4.75751 | 1.292068 | 0.210149 | 0.292488 | 0 | 0 | 0 | 0 | 2.808791 | 9.468674 | 4.263783 | 2.680843 | 13.78987 | 1.538429 | 0 | 0.746341 | 6.674615 | 7.074744 | 8.098703 |

| | | | | | | | | | | | | | | | | | | | | | | | | | | |
|---|---|---|---|---|---|---|---|---|---|---|---|---|---|---|---|---|---|---|---|---|---|---|---|---|---|---|
| Gm19514 | 100503026 | 2.673753 | -3.27539 | 3.222985 | 0.004021 | 0.010443 | 0.901379 | -3.27539 | 1.480662 | 0.153309 | 0.224359 | 3.709904 | 0 | 2.603909 | 12.98709 | 12.63956 | 9.468674 | 29.84648 | 14.74464 | 27.57973 | 0.769214 | 3.812639 | 0 | 14.18356 | 19.65207 | 9.718443 |
| Smok4a | 272667 | 2.674998 | -3.85474 | 2.783639 | 0.011024 | 0.026191 | 0.095798 | -3.85474 | 0.116471 | 0.908368 | 0.926906 | 0.741981 | 0 | 0.86797 | 7.792255 | 1.404395 | 10.82134 | 8.527565 | 1.340422 | 13.78987 | 4.615287 | 2.287584 | 0.746341 | 12.5149 | 12.57732 | 19.43689 |
| 4930512H | 75111 | 2.67524 | 2.010759 | 14.97041 | 8.53E-13 | 8.36E-12 | 0.922272 | 2.010759 | 9.076525 | 8.91E-09 | 5.09E-08 | 62.32638 | 93.24367 | 97.2126 | 602.6011 | 501.3691 | 570.8258 | 1162.591 | 1096.465 | 1046.881 | 145.3815 | 147.1679 | 153 | 100.9536 | 128.1315 | 93.13508 |
| Nfkbia | 18035 | 2.676472 | 8.046754 | 83.90429 | 2.21E-28 | 1.53E-25 | -0.50015 | 8.046754 | -22.3064 | 2.83E-16 | 7.03E-15 | 11206.14 | 11203.45 | 11193.34 | 76806.96 | 76938.39 | 75643.89 | 58143.78 | 57069.8 | 53756.35 | 10155.17 | 10127.13 | 10063.67 | 1916.449 | 2101.985 | 2053.831 |
| Parp11 | 101187 | 2.678496 | 6.130002 | 47.64096 | 3.65E-23 | 3.39E-21 | -0.7002 | 6.130002 | -15.978 | 2.37E-13 | 3.01E-12 | 2161.39 | 2279.586 | 1961.611 | 14652.04 | 14990.52 | 14070.45 | 9907.609 | 9447.292 | 8702.555 | 4251.448 | 4535.516 | 4319.077 | 1005.364 | 1029.768 | 1010.718 |
| Vmn1r158 | 100043067 | 2.681263 | -5.07423 | 3.058563 | 0.005891 | 0.014835 | 0.577064 | -5.07423 | 0.7691 | 0.450262 | 0.549121 | 0 | 0 | 0 | 1.298709 | 2.808791 | 2.705335 | 8.527565 | 2.680843 | 2.298311 | 0 | 0 | 0 | 4.171635 | 14.14949 | 9.718443 |
| Gm5258 | 383538 | 2.681263 | -5.36155 | 2.812899 | 0.010321 | 0.024684 | -0.3543 | -5.36155 | -0.40995 | 0.685927 | 0.768829 | 0 | 0 | 0 | 1.298709 | 2.808791 | 2.705335 | 5.685043 | 0 | 2.298311 | 0 | 0 | 0.746341 | 1.668654 | 4.716496 | 7.288832 |
| Cnksr2 | 245684 | 2.68467 | -1.40188 | 5.170548 | 3.82E-05 | 0.000132 | -0.03607 | -1.40188 | -0.09961 | 0.921582 | 0.936501 | 3.709904 | 7.992315 | 7.811726 | 51.94837 | 43.53625 | 36.52203 | 29.84648 | 44.23392 | 60.90525 | 13.07665 | 8.387806 | 6.717071 | 60.07154 | 55.81187 | 34.82442 |
| Defb10 | 246085 | 2.685317 | -6.03805 | 2.896385 | 0.008543 | 0.020809 | -0.73187 | -6.03805 | -0.85177 | 0.403797 | 0.502901 | 0 | 0 | 0 | 2.597418 | 2.808791 | 1.352668 | 4.263783 | 0 | 1.149156 | 0 | 0 | 0 | 0 | 0 | 2.429611 |
| Erich6b | 75272 | 2.685868 | -4.22973 | 3.208642 | 0.004158 | 0.010761 | -0.73288 | -4.22973 | -0.9712 | 0.342328 | 0.439504 | 1.483961 | 0.888035 | 0 | 6.493546 | 4.213186 | 9.468674 | 4.263783 | 2.680843 | 4.596622 | 0.769214 | 4.575167 | 0 | 7.508942 | 7.860826 | 7.288832 |
| Tcl1b2 | 27381 | 2.686447 | -4.60095 | 2.980613 | 0.007048 | 0.017472 | 0.640143 | -4.60095 | 0.851146 | 0.404137 | 0.503157 | 0.741981 | 0 | 0 | 3.896127 | 1.404395 | 6.763339 | 5.685043 | 4.021265 | 9.193245 | 0 | 2.287584 | 1.492683 | 4.171635 | 2.358248 | 6.478962 |
| Zbp1 | 58203 | 2.686728 | 6.557136 | 62.99789 | 9.78E-26 | 2.20E-23 | -1.17305 | 6.557136 | -33.4459 | 6.30E-20 | 4.11E-18 | 5509.949 | 5503.153 | 5002.109 | 36756.07 | 36667.36 | 36692.47 | 16893.11 | 17586.33 | 16313.41 | 12579.73 | 12302.62 | 11851.9 | 103.4565 | 141.4949 | 183.0307 |
| Gm14151 | 433486 | 2.687139 | -5.78578 | 3.327022 | 0.003151 | 0.008336 | 0.957953 | -5.78578 | 1.416111 | 0.171172 | 0.24593 | 0 | 0 | 0 | 2.597418 | 1.404395 | 2.705335 | 5.685043 | 2.680843 | 8.044089 | 0 | 0 | 0 | 0 | 0.786083 | 0 |
| Fabp1 | 14080 | 2.687139 | -4.74454 | 2.980127 | 0.007056 | 0.017488 | 0.663539 | -4.74454 | 0.862586 | 0.397954 | 0.497013 | 0 | 0 | 0 | 2.597418 | 1.404395 | 2.705335 | 1.421261 | 6.702109 | 5.745778 | 3.076858 | 0.762528 | 1.492683 | 2.502981 | 10.21907 | 4.049351 |
| 1700031L1 | 67327 | 2.687139 | -5.2584 | 3.410061 | 0.002591 | 0.006955 | 0.576521 | -5.2584 | 0.854027 | 0.402574 | 0.501663 | 0 | 0 | 0 | 2.597418 | 1.404395 | 2.705335 | 2.842522 | 5.361687 | 3.447467 | 0 | 0 | 0 | 2.502981 | 4.716496 | 6.478962 |
| Olfr1036 | 258245 | 2.687139 | -5.44052 | 2.979961 | 0.007058 | 0.017492 | -0.52334 | -5.44052 | -0.6344 | 0.532562 | 0.62831 | 0 | 0 | 0 | 2.597418 | 1.404395 | 2.705335 | 7.106304 | 0 | 1.149156 | 0 | 0 | 0 | 3.337308 | 7.074744 | 4.859222 |
| Olfr930 | 258269 | 2.68726 | -4.32284 | 3.233361 | 0.003924 | 0.010215 | 0.268959 | -4.32284 | 0.385229 | 0.70388 | 0.784971 | 2.967923 | 0 | 0 | 5.194837 | 5.617581 | 5.410671 | 4.263783 | 8.04253 | 9.193245 | 0 | 0 | 1.492683 | 8.343269 | 8.646909 | 9.718443 |
| Chic1 | 12212 | 2.687744 | 2.381753 | 21.77128 | 4.65E-16 | 8.27E-15 | -2.60716 | 2.381753 | -18.8716 | 8.50E-15 | 1.47E-13 | 200.3348 | 284.1712 | 210.9166 | 1535.074 | 1721.789 | 1451.412 | 243.0356 | 273.446 | 279.2448 | 266.1482 | 208.9326 | 229.8731 | 160.1908 | 120.2706 | 116.6213 |
| Caps2 | 353025 | 2.688442 | -3.2539 | 3.359104 | 0.002921 | 0.007768 | 0.960037 | -3.2539 | 1.646458 | 0.114316 | 0.174855 | 5.193865 | 0 | 1.735939 | 11.68838 | 12.63956 | 10.82134 | 18.47639 | 22.78717 | 32.17636 | 3.076858 | 0 | 1.492683 | 12.5149 | 15.72165 | 11.33818 |
| A130040M | 319269 | 2.688661 | 8.442817 | 18.37331 | 3.99E-09 | 2.27E-08 | -0.68257 | 8.442817 | -6.31031 | 8.21E-05 | 0.000269 | 9228.757 | 9602.322 | 5246.876 | 53674.35 | 55180.09 | 52142.64 | 35355.28 | 35734.3 | 33291.04 | 22854.9 | 23143.48 | 23647.82 | 11704.77 | 11351.82 | 11748.79 |
| Cstl1 | 228756 | 2.691705 | -4.63566 | 2.663916 | 0.014402 | 0.033305 | 0.21509 | -4.63566 | 0.247942 | 0.806549 | 0.875306 | 0 | 0.888035 | 0 | 1.298709 | 2.808791 | 9.468674 | 4.263783 | 8.04253 | 2.298311 | 1.538429 | 0 | 2.239024 | 1.668654 | 7.860826 | 8.098703 |
| Rasgef1c | 74563 | 2.691706 | -1.52454 | 4.527816 | 0.000178 | 0.000564 | 2.914384 | -1.52454 | 8.672644 | 1.93E-08 | 1.05E-07 | 4.451884 | 1.77607 | 4.339848 | 16.88322 | 33.70549 | 28.40602 | 200.3978 | 226.5313 | 182.7157 | 2.307643 | 9.912862 | 4.478048 | 40.04769 | 33.01547 | 42.11325 |
| Cdkl5 | 382253 | 2.692861 | 1.567855 | 11.74576 | 8.80E-11 | 6.36E-10 | 0.777685 | 1.567855 | 5.917813 | 6.67E-06 | 2.54E-05 | 40.06696 | 52.39406 | 63.36178 | 324.6773 | 386.2087 | 344.9303 | 586.9807 | 638.0408 | 643.5271 | 74.6138 | 80.06542 | 134.3414 | 171.8713 | 154.0722 | 171.6925 |
| Olfm5 | 244180 | 2.693457 | -3.39545 | 2.639891 | 0.015188 | 0.034946 | 1.369623 | -3.39545 | 1.819381 | 0.082911 | 0.132975 | 0.741981 | 0 | 2.603909 | 1.298709 | 8.426372 | 28.40602 | 17.05513 | 28.14886 | 18.38649 | 0 | 3.050111 | 2.239024 | 18.35519 | 15.72165 | 25.91585 |
| Sp110 | 109032 | 2.693595 | 6.259036 | 16.9036 | 8.00E-09 | 4.37E-08 | -2.27677 | 6.259036 | -14.1012 | 4.78E-08 | 2.46E-07 | 3609.736 | 3976.621 | 2019.765 | 21204.02 | 21422.65 | 21335.63 | 4687.318 | 4624.455 | 4411.608 | 6065.256 | 6032.358 | 5819.223 | 1047.08 | 1069.072 | 1266.637 |
| Tchhl1 | 71325 | 2.693836 | -4.1969 | 2.869448 | 0.009082 | 0.021988 | 2.661812 | -4.1969 | 3.989877 | 0.000649 | 0.00184 | 0 | 0.888035 | 0 | 2.597418 | 5.617581 | 2.705335 | 19.89765 | 30.8297 | 28.72889 | 0 | 1.525056 | 0 | 3.337308 | 16.50774 | 9.718443 |
| Klf8 | 245671 | 2.695715 | 2.310587 | 19.04306 | 7.08E-15 | 1.00E-13 | -2.38204 | 2.310587 | -15.9734 | 2.38E-13 | 3.02E-12 | 159.5259 | 190.9275 | 188.3494 | 1135.072 | 1304.683 | 1270.155 | 233.0868 | 226.5313 | 272.3499 | 572.2956 | 489.5429 | 532.8877 | 113.4685 | 51.09537 | 86.65612 |
| Olfr689 | 258745 | 2.697792 | -5.09297 | 2.471539 | 0.021943 | 0.048437 | 1.083182 | -5.09297 | 1.197155 | 0.244374 | 0.331628 | 0 | 0 | 0 | 3.896127 | 7.021976 | 0 | 11.37009 | 2.680843 | 5.745778 | 0 | 0 | 0 | 18.35519 | 2.358248 | 3.239481 |
| Slc1a3 | 20512 | 2.697918 | -2.88157 | 3.01091 | 0.006574 | 0.016389 | 1.057407 | -2.88157 | 1.627414 | 0.118323 | 0.179946 | 0 | 7.10428 | 1.735939 | 10.38967 | 18.25714 | 8.116006 | 31.26774 | 18.7659 | 28.72889 | 1.538429 | 9.912862 | 1.492683 | 13.34923 | 29.08506 | 13.76779 |
| Pth2r | 213527 | 2.698022 | -2.74877 | 3.148403 | 0.004784 | 0.012243 | 1.19148 | -2.74877 | 1.960483 | 0.063122 | 0.105615 | 0 | 6.216245 | 2.603909 | 12.98709 | 9.830767 | 14.87935 | 48.32287 | 16.08506 | 35.62382 | 3.846072 | 0.762528 | 3.731706 | 29.20144 | 25.94073 | 21.8665 |
| Noto | 384452 | 2.698505 | -3.20926 | 3.134431 | 0.004942 | 0.012614 | 0.860526 | -3.20926 | 1.305712 | 0.20555 | 0.287059 | 2.967923 | 0 | 0.86797 | 2.597418 | 8.426372 | 22.99535 | 15.63387 | 16.08506 | 16.08818 | 1.538429 | 5.337695 | 2.985365 | 21.6925 | 13.3634 | 20.24676 |
| Tpsb2 | 17229 | 2.698902 | -4.45957 | 2.896909 | 0.008532 | 0.020786 | 1.537008 | -4.45957 | 2.125349 | 0.04539 | 0.079747 | 0 | 0 | 0.86797 | 1.298709 | 4.213186 | 6.763339 | 8.527565 | 14.74464 | 12.64071 | 2.307643 | 0 | 0 | 4.171635 | 11.79124 | 4.859222 |
| Tex24 | 541463 | 2.699562 | -3.85488 | 2.951141 | 0.00754 | 0.018568 | -0.04807 | -3.85488 | -0.06238 | 0.950843 | 0.959156 | 0 | 2.664105 | 1.735939 | 6.493546 | 14.04395 | 6.763339 | 18.47639 | 2.680843 | 11.49156 | 2.307643 | 0 | 1.492683 | 8.343269 | 12.57732 | 4.859222 |
| Itpr1 | 16438 | 2.699771 | 5.963009 | 51.20311 | 7.94E-24 | 9.16E-22 | -3.13199 | 5.963009 | -49.5414 | 1.60E-23 | 3.68E-21 | 2576.899 | 2807.967 | 3031.818 | 19807.91 | 19304.82 | 18961.7 | 2485.785 | 2380.589 | 2067.331 | 4432.214 | 4589.655 | 4154.882 | 1349.107 | 1276.598 | 1318.469 |
| Cyp2c65 | 72303 | 2.700458 | -3.70042 | 2.591355 | 0.016903 | 0.038357 | 1.529568 | -3.70042 | 1.928089 | 0.067254 | 0.111417 | 0 | 2.664105 | 0 | 5.194837 | 1.404395 | 12.17401 | 5.685043 | 22.78717 | 25.28142 | 0 | 0.762528 | 2.985365 | 30.8701 | 24.36856 | 24.29611 |
| Gm9507 | 670880 | 2.700804 | -3.97938 | 2.63011 | 0.01552 | 0.035616 | 1.127928 | -3.97938 | 1.393217 | 0.177896 | 0.254064 | 0 | 1.77607 | 0 | 2.597418 | 2.808791 | 9.468674 | 15.63387 | 22.78717 | 3.447467 | 1.538429 | 3.050111 | 1.492683 | 5.840288 | 18.0799 | 4.859222 |
| Klf14 | 619665 | 2.701267 | -2.30721 | 3.43754 | 0.002428 | 0.006553 | 0.595584 | -2.30721 | 1.08635 | 0.289442 | 0.382306 | 2.967923 | 7.10428 | 0.86797 | 28.5716 | 16.85274 | 20.29002 | 21.31891 | 45.57434 | 39.07129 | 4.615287 | 6.862751 | 0 | 37.54471 | 50.30929 | 45.35273 |
| Sis | 69983 | 2.703957 | -2.61494 | 2.856698 | 0.009348 | 0.022575 | 1.65904 | -2.61494 | 2.541569 | 0.018849 | 0.037485 | 6.677827 | 0.888035 | 0 | 7.792255 | 9.830767 | 16.23201 | 21.31891 | 33.51054 | 62.0544 | 9.999788 | 5.337695 | 0.746341 | 33.37308 | 33.80155 | 30.77507 |
| Olfr885 | 257885 | 2.705821 | -4.70604 | 3.206272 | 0.004181 | 0.010815 | -0.68327 | -4.70604 | -0.89967 | 0.378333 | 0.476591 | 0.741981 | 0 | 0.86797 | 7.792255 | 7.021976 | 2.705335 | 5.685043 | 2.680843 | 2.298311 | 0 | 1.525056 | 0 | 2.502981 | 9.432991 | 3.239481 |
| Skint4 | 320640 | 2.705931 | -3.09732 | 3.188975 | 0.004353 | 0.011226 | 1.580183 | -3.09732 | 2.594276 | 0.016795 | 0.033926 | 1.483961 | 1.77607 | 0 | 7.792255 | 21.06593 | 2.705335 | 29.84648 | 22.78717 | 25.28142 | 1.538429 | 5.337695 | 0.746341 | 25.02981 | 27.51289 | 25.10598 |
| Samsn1 | 67742 | 2.706253 | 5.251593 | 62.49425 | 1.16E-25 | 2.50E-23 | -2.95793 | 5.251593 | -59.0113 | 3.92E-25 | 1.40E-22 | 2181.423 | 2366.613 | 2232.418 | 15622.17 | 16458.11 | 15117.41 | 2045.194 | 2166.122 | 2085.717 | 1900.729 | 1865.143 | 1932.278 | 414.6605 | 408.763 | 475.3938 |
| Ano5 | 233246 | 2.706305 | -2.14138 | 3.443539 | 0.002393 | 0.00647 | 0.846703 | -2.14138 | 1.591684 | 0.126159 | 0.190009 | 11.12971 | 5.32821 | 0 | 18.18193 | 28.08791 | 18.93735 | 31.26774 | 30.8297 | 62.0544 | 6.92293 | 4.575167 | 2.985365 | 37.54471 | 33.80155 | 30.77507 |
| Cyp2c68 | 433247 | 2.707486 | -4.18695 | 2.577644 | 0.017419 | 0.039406 | 0.896831 | -4.18695 | 1.068847 | 0.297079 | 0.390752 | 2.225942 | 0 | 0 | 2.597418 | 11.23516 | 4.058003 | 2.842522 | 25.46801 | 12.64071 | 0 | 2.287584 | 0.746341 | 12.5149 | 8.646909 | 3.239481 |
| Slfn5 | 327978 | 2.707529 | 9.916834 | 84.61298 | 1.91E-21 | 1.12E-19 | -0.30333 | 9.916834 | -15.3065 | 1.55E-10 | 1.18E-09 | 26856.74 | 28321.21 | 25132.06 | 186667.4 | 190610.1 | 182221.9 | 158578.6 | 157286.4 | 155172.8 | 42417.56 | 42345.46 | 40546.48 | 19887.85 | 19648.14 | 19563.23 |
| Gm16675 | 100503498 | 2.707952 | 2.422604 | 20.06666 | 2.45E-15 | 3.76E-14 | -0.89927 | 2.422604 | -8.98127 | 1.07E-08 | 6.00E-08 | 138.0084 | 176.719 | 126.7236 | 1014.292 | 1072.958 | 959.0414 | 542.9216 | 583.0835 | 565.3846 | 291.5323 | 268.4098 | 227.6341 | 171.8713 | 156.4304 | 136.0582 |
| Pramel34 | 381654 | 2.709232 | -3.9036 | 3.14178 | 0.004858 | 0.012419 | 0.94165 | -3.9036 | 1.401842 | 0.175338 | 0.250973 | 0 | 0.888035 | 1.735939 | 10.38967 | 2.808791 | 8.116006 | 17.05513 | 16.08506 | 9.193245 | 0.769214 | 1.525056 | 0 | 5.840288 | 14.93557 | 12.14805 |
| Il18bp | 16068 | 2.711926 | 5.07075 | 41.01277 | 8.65E-22 | 5.53E-20 | 0.615987 | 5.07075 | 14.24852 | 2.23E-12 | 2.36E-11 | 679.6544 | 727.3006 | 728.2265 | 5153.278 | 5017.904 | 4737.042 | 7916.423 | 8128.317 | 7695.895 | 1173.821 | 1208.607 | 1129.214 | 1017.879 | 941.727 | 1039.873 |
| Prr23a2 | 623186 | 2.713924 | -3.95401 | 3.240144 | 0.003863 | 0.010067 | 0.861498 | -3.95401 | 1.3174 | 0.201673 | 0.282531 | 0 | 2.664105 | 0.86797 | 6.493546 | 8.426372 | 6.763339 | 8.527565 | 14.74464 | 20.6848 | 1.538429 | 0 | 0 | 7.508942 | 16.50774 | 7.288832 |
| Gm12250 | 631323 | 2.714414 | 6.76299 | 57.92052 | 5.82E-25 | 9.83E-23 | 0.463042 | 6.76299 | 15.14982 | 6.75E-13 | 7.78E-12 | 2887.047 | 2828.391 | 2616.06 | 19548.17 | 19838.49 | 19049.62 | 28246.14 | 27835.2 | 27568.24 | 5316.041 | 5036.496 | 4871.369 | 1448.392 | 1664.923 | 1422.942 |
| Gdf2 | 12165 | 2.715568 | -3.51287 | 2.839361 | 0.009722 | 0.023412 | 1.032025 | -3.51287 | 1.45817 | 0.159353 | 0.23176 | 0 | 1.77607 | 4.339848 | 15.58451 | 15.44835 | 4.058003 | 19.89765 | 14.74464 | 34.47467 | 0 | 0.762528 | 5.97073 | 7.508942 | 11.00516 | 4.049351 |
| Cylc2 | 74914 | 2.718807 | -4.28287 | 2.996711 | 0.006792 | 0.016888 | -0.02157 | -4.28287 | -0.02805 | 0.977883 | 0.981433 | 2.225942 | 0.888035 | 0 | 9.090964 | 8.426372 | 5.410671 | 1.421261 | 12.0638 | 18.38649 | 0 | 0 | 0 | 5.840288 | 18.0799 | 7.288832 |
| Dpysl3 | 22240 | 2.720504 | -1.7435 | 4.77213 | 9.88E-05 | 0.000324 | 1.183485 | -1.7435 | 3.473907 | 0.002227 | 0.005655 | 10.38773 | 4.440175 | 2.603909 | 46.75353 | 19.66153 | 59.51738 | 98.067 | 91.14868 | 86.18667 | 3.076858 | 5.337695 | 2.239024 | 28.36712 | 25.15464 | 21.05663 |
| Cxxc4 | 319478 | 2.720905 | -2.02958 | 4.1906 | 0.0004 | 0.001218 | -0.53828 | -2.02958 | -1.06697 | 0.297905 | 0.391603 | 2.967923 | 7.10428 | 3.471878 | 32.46773 | 29.4923 | 32.46403 | 21.31891 | 12.0638 | 39.07129 | 4.615287 | 4.575167 | 8.956095 | 50.05962 | 40.8763 | 17.00728 |
| Olfr629 | 258818 | 2.721679 | -4.37828 | 3.242926 | 0.003838 | 0.010005 | -0.03672 | -4.37828 | -0.05141 | 0.959473 | 0.966677 | 1.483961 | 0.888035 | 0 | 7.792255 | 2.808791 | 12.17401 | 4.263783 | 12.0638 | 5.745778 | 0 | 0 | 0 | 4.171635 | 13.3634 | 12.95792 |
| Gm11525 | 100502635 | 2.721847 | -3.4053 | 3.113673 | 0.005186 | 0.013178 | 0.535804 | -3.4053 | 0.812077 | 0.425713 | 0.524581 | 0 | 7.10428 | 2.603909 | 16.88322 | 19.66153 | 6.763339 | 12.79135 | 26.80843 | 22.98311 | 2.307643 | 0 | 0 | 10.84625 | 19.65207 | 10.52831 |
| Tnip3 | 414084 | 2.722048 | 6.501215 | 101.0576 | 4.21E-30 | 1.11E-26 | -3.01084 | 6.501215 | -94.9228 | 1.60E-29 | 1.28E-25 | 6522.753 | 6725.977 | 6690.31 | 47049.64 | 47252.28 | 46055.63 | 6064.52 | 6163.259 | 5866.439 | 9661.334 | 9709.267 | 9884.544 | 259.4757 | 282.9897 | 223.5242 |
| Olfr62 | 18363 | 2.722192 | -5.06477 | 2.774532 | 0.011252 | 0.026672 | -0.62158 | -5.06477 | -0.69771 | 0.492889 | 0.591076 | 0 | 0 | 0.86797 | 1.298709 | 5.617581 | 5.410671 | 0 | 2.680843 | 8.044089 | 0 | 0 | 1.492683 | 4.171635 | 4.716496 | 2.429611 |
| Krt2 | 16681 | 2.722902 | -2.89638 | 3.02615 | 0.006348 | 0.01588 | 1.447816 | -2.89638 | 2.275521 | 0.033293 | 0.061204 | 5.193865 | 0.888035 | 0 | 11.68838 | 15.44835 | 5.410671 | 15.63387 | 38.87223 | 39.07129 | 3.846072 | 0.762528 | 7.463413 | 22.52683 | 18.0799 | 22.67637 |
| Adamts1 | 11504 | 2.723895 | 2.285386 | 12.92552 | 1.45E-11 | 1.17E-10 | 1.261963 | 2.285386 | 11.17605 | 2.21E-10 | 1.63E-09 | 85.32779 | 43.51371 | 52.94615 | 438.9637 | 446.5977 | 381.4523 | 1075.894 | 955.7207 | 1115.83 | 153.8429 | 142.5927 | 164.9414 | 496.4245 | 564.4073 | 521.5565 |
| 1700044K( | 67321 | 2.724809 | -5.80477 | 2.686211 | 0.013706 | 0.031871 | 0.286007 | -5.80477 | 0.323481 | 0.749479 | 0.82567 | 0 | 0 | 0 | 5.194837 | 5.617581 | 0 | 1.421261 | 1.340422 | 10.3424 | 0 | 0 | 0 | 0.834327 | 0 | 0.80987 |
| Ifne | 230405 | 2.724809 | -5.19417 | 2.547004 | 0.018627 | 0.041875 | 0.182662 | -5.19417 | 0.195021 | 0.847221 | 0.910542 | 0 | 0 | 0 | 5.194837 | 5.617581 | 0 | 1.421261 | 4.021265 | 3.447467 | 0 | 3.050111 | 0 | 2.502981 | 7.074744 | 1.619741 |
| Gm8239 | 666688 | 2.726306 | -4.30543 | 2.789312 | 0.010884 | 0.025882 | 1.078179 | -4.30543 | 1.367838 | 0.185594 | 0.263356 | 0 | 0.888035 | 0 | 1.298709 | 5.617581 | 5.410671 | 2.842522 | 13.40422 | 13.78987 | 0 | 1.525056 | 0.746341 | 17.52087 | 13.3634 | 6.478962 |
| Flt1 | 14254 | 2.728364 | 4.33647 | 14.95389 | 8.71E-13 | 8.52E-12 | 1.211563 | 4.33647 | 12.20498 | 4.30E-11 | 3.58E-10 | 100.1674 | 77.25904 | 101.5524 | 609.0946 | 624.9559 | 753.4359 | 1678.509 | 1536.123 | 1551.36 | 3543.002 | 3918.631 | 3514.521 | 7206.916 | 7154.138 | 7330.136 |
| Mir19b-2 | 387195 | 2.728738 | -6.51063 | 3.500247 | 0.002092 | 0.005711 | -2.23101 | -6.51063 | -2.8633 | 0.009209 | 0.020094 | 0 | 0 | 0 | 5.194837 | 1.404395 | 1.352668 | 0 | 0 | 0 | 0 | 0 | 0 | 0 | 0 | 0 |
| D130009I1 | 320249 | 2.729168 | -3.67847 | 3.097092 | 0.005389 | 0.013673 | 1.247367 | -3.67847 | 1.873848 | 0.074708 | 0.121862 | 0.741981 | 0 | 1.735939 | 7.792255 | 2.808791 | 12.17401 | 14.21261 | 12.0638 | 27.57973 | 3.076858 | 0 | 1.492683 | 15.85221 | 11.00516 | 10.52831 |
| 1700065I1 | 78462 | 2.729221 | -4.19966 | 2.666401 | 0.014323 | 0.033137 | 1.057298 | -4.19966 | 1.309582 | 0.20426 | 0.285575 | 1.483961 | 0 | 0 | 7.792255 | 7.021976 | 1.352668 | 15.63387 | 13.40422 | 5.745778 | 0.769214 | 1.525056 | 5.97073 | 9.177596 | 3.930413 | 0.80987 |
| Gm17634 | 100504568 | 2.731002 | -3.15786 | 3.450147 | 0.002356 | 0.006378 | 1.409028 | -3.15786 | 2.399087 | 0.025638 | 0.048895 | 0.741981 | 0 | 1.735939 | 6.493546 | 15.44835 | 2.705335 | 19.89765 | 14.74464 | 24.13227 | 3.076858 | 3.050111 | 5.224389 | 21.6925 | 18.86598 | 25.91585 |
| Meikin | 74847 | 2.734 | -3.00559 | 3.766109 | 0.00111 | 0.003173 | -0.06189 | -3.00559 | -0.10794 | 0.915052 | 0.931702 | 3.709904 | 1.77607 | 0.86797 | 11.68838 | 22.47032 | 14.87935 | 12.79135 | 12.0638 | 22.98311 | 0 | 3.812639 | 2.985365 | 18.35519 | 24.36856 | 21.8665 |
| Glipr1l2 | 67537 | 2.734206 | -4.08429 | 2.908743 | 0.008306 | 0.020275 | 0.82609 | -4.08429 | 1.111031 | 0.278915 | 0.370473 | 0.741981 | 0.888035 | 0 | 2.597418 | 5.617581 | 10.82134 | 17.05513 | 22.78717 | 3.447467 | 1.538429 | 0 | 0.746341 | 11.68058 | 7.074744 | 8.098703 |
| Olfr191 | 258035 | 2.734331 | -5.49296 | 2.93462 | 0.00783 | 0.019217 | -1.27085 | -5.49296 | -1.44812 | 0.162114 | 0.2351 | 0 | 0 | 1.735939 | 5.194837 | 5.617581 | 2.705335 | 0 | 4.021265 | 2.298311 | 0 | 0 | 0 | 0 | 1.572165 | 3.239481 |
| Gpr152 | 269053 | 2.736734 | -0.36162 | 6.746371 | 1.04E-06 | 4.28E-06 | 0.996716 | -0.36162 | 4.375502 | 0.000256 | 0.000777 | 27.45329 | 10.65642 | 7.811726 | 92.20835 | 78.64613 | 136.6194 | 206.0828 | 203.7441 | 211.4446 | 26.15329 | 13.7255 | 15.67317 | 40.88202 | 45.59279 | 38.87377 |
| Scarf2 | 224024 | 2.736963 | 2.834662 | 30.38299 | 4.70E-19 | 1.58E-17 | -3.82245 | 2.834662 | -27.3958 | 4.06E-18 | 1.54E-16 | 301.2442 | 321.4687 | 360.2074 | 2298.715 | 2280.738 | 2379.343 | 137.8623 | 188.9995 | 182.7157 | 336.1467 | 354.5755 | 344.8097 | 302.8607 | 360.0258 | 294.7928 |
| Glycam1 | 14663 | 2.737535 | -4.30095 | 3.000834 | 0.006728 | 0.016741 | 0.176332 | -4.30095 | 0.231246 | 0.819326 | 0.887049 | 1.483961 | 0.888035 | 0 | 7.792255 | 7.021976 | 5.410671 | 5.685043 | 5.361687 | 14.93902 | 0 | 0.762528 | 2.985365 | 0.834327 | 3.14433 | 9.718443 |
| I830077J0 | 433638 | 2.737554 | 3.255143 | 28.83909 | 1.39E-18 | 4.24E-17 | -1.75017 | 3.255143 | -20.5703 | 1.48E-15 | 3.07E-14 | 335.3753 | 442.2414 | 408.8137 | 2961.057 | 2770.872 | 2601.18 | 857.0203 | 879.3167 | 835.4361 | 245.3794 | 301.1985 | 300.0292 | 247.7951 | 238.183 | 204.8972 |
| Glod5 | 69824 | 2.737871 | -4.04918 | 2.81888 | 0.010182 | 0.024395 | 0.3348 | -4.04918 | 0.412795 | 0.683874 | 0.767051 | 0 | 0.888035 | 0.86797 | 3.896127 | 23.87472 | 1.352668 | 4.263783 | 10.72337 | 8.044089 | 1.538429 | 1.525056 | 2.985365 | 10.84625 | 7.860826 | 2.429611 |
| Ly6c1 | 17067 | 2.737884 | -2.02299 | 3.913422 | 0.000779 | 0.002277 | 0.904597 | -2.02299 | 2.027076 | 0.055327 | 0.094521 | 2.967923 | 6.216245 | 3.471878 | 37.66257 | 30.8967 | 24.34802 | 73.90556 | 28.14886 | 96.52907 | 9.999788 | 7.625279 | 2.239024 | 19.18952 | 13.3634 | 9.718443 |
| Gm20064 | 100504102 | 2.738222 | -4.60765 | 2.559863 | 0.018111 | 0.040813 | 0.41025 | -4.60765 | 0.454752 | 0.653879 | 0.740513 | 0.741981 | 0 | 0 | 6.493546 | 0 | 14.87935 | 2.842522 | 10.72337 | 4.596622 | 1.538429 | 0 | 0 | 10.01192 | 7.074744 | 8.908573 |
| Tlcd4 | 99887 | 2.739941 | -2.04301 | 3.556636 | 0.00183 | 0.005043 | 2.316408 | -2.04301 | 5.025817 | 5.39E-05 | 0.000181 | 3.709904 | 2.664105 | 1.735939 | 31.16902 | 23.87472 | 12.17401 | 98.067 | 107.2337 | 127.5563 | 9.230573 | 1.525056 | 0 | 49.22529 | 33.80155 | 59.9304 |
| Fam71d | 70897 | 2.740196 | -3.21084 | 2.673216 | 0.014108 | 0.032689 | 0.11381 | -3.21084 | 0.138605 | 0.891062 | 0.918721 | 1.483961 | 4.440175 | 0 | 10.38967 | 14.04395 | 8.116006 | 24.16143 | 17.42548 | 4.596622 | 0 | 6.862751 | 8.209754 | 15.01788 | 18.0799 | 15.38754 |
| Jph2 | 59091 | 2.740844 | -2.19274 | 3.399607 | 0.002655 | 0.007111 | 1.724968 | -2.19274 | 3.253116 | 0.003747 | 0.009033 | 0 | 5.32821 | 5.207818 | 14.2858 | 12.63956 | 18.93735 | 49.74413 | 46.91476 | 63.20356 | 3.846072 | 5.337695 | 2.985365 | 33.37308 | 64.45877 | 42.92312 |
| Olfr1260 | 258983 | 2.740983 | -4.38948 | 2.786093 | 0.010963 | 0.026059 | 0.053053 | -4.38948 | 0.064329 | 0.949307 | 0.957866 | 0 | 0.888035 | 2.603909 | 5.194837 | 7.021976 | 10.82134 | 8.527565 | 10.72337 | 5.745778 | 0 | 0 | 0 | 11.68058 | 15.72165 | 0.80987 |
| Csrnp1 | 215418 | 2.741388 | 5.564101 | 36.07447 | 1.29E-20 | 6.39E-19 | 0.853966 | 5.564101 | 17.62626 | 3.36E-14 | 5.17E-13 | 991.2863 | 950.1974 | 1011.185 | 7128.615 | 7194.717 | 6778.218 | 13867.24 | 13748.71 | 12199.44 | 1368.433 | 1463.291 | 1491.19 | 1436.711 | 1349.704 | 1322.518 |
| 1700031F( | 73300 | 2.74526 | -3.8362 | 3.113867 | 0.005183 | 0.013173 | 0.269777 | -3.8362 | 0.380924 | 0.707025 | 0.787776 | 0.741981 | 5.32821 | 0 | 11.68838 | 8.426372 | 8.116006 | 8.527565 | 21.44675 | 9.193245 | 0 | 3.050111 | 0.746341 | 11.68058 | 5.502578 | 3.239481 |
| Tldc2 | 383766 | 2.746868 | 0.99189 | 12.75271 | 1.88E-11 | 1.49E-10 | -0.65221 | 0.99189 | -4.34591 | 0.000275 | 0.00083 | 60.10044 | 77.25904 | 72.90945 | 531.172 | 442.3845 | 522.1297 | 370.9491 | 377.9989 | 264.3058 | 82.30595 | 80.82795 | 55.97559 | 31.70442 | 36.1598 | 18.62702 |
| H2-M2 | 14990 | 2.747918 | -0.60736 | 7.825759 | 1.05E-07 | 4.93E-07 | -1.32832 | -0.60736 | -4.6002 | 0.000149 | 0.00047 | 14.83961 | 26.64105 | 25.17112 | 187.0141 | 122.1824 | 156.9095 | 59.69296 | 45.57434 | 87.33582 | 46.92208 | 24.40089 | 19.40487 | 16.68654 | 12.57732 | 16.19741 |
| Fscn1 | 14086 | 2.748491 | 4.129485 | 32.62581 | 1.06E-19 | 4.16E-18 | -0.95631 | 4.129485 | -14.5117 | 1.56E-12 | 1.70E-11 | 522.3544 | 549.6937 | 573.7279 | 3811.711 | 3911.241 | 4066.119 | 2306.706 | 2091.058 | 1946.67 | 717.6771 | 863.1815 | 796.3461 | 388.7963 | 332.5129 | 326.3777 |
| Mindy4b-ps | 100040293 | 2.748914 | -3.51746 | 2.8502 | 0.009487 | 0.022886 | -0.23802 | -3.51746 | -0.29323 | 0.772181 | 0.845368 | 0 | 1.77607 | 5.207818 | 6.493546 | 16.85274 | 12.17401 | 14.21261 | 2.680843 | 19.53565 | 3.846072 | 0 | 2.985365 | 13.34923 | 7.860826 | 12.14805 |
| Clcf1 | 56708 | 2.750569 | 0.730767 | 8.575203 | 2.33E-08 | 1.19E-07 | 0.921274 | 0.730767 | 5.206279 | 3.51E-05 | 0.000121 | 52.68063 | 31.08122 | 32.11488 | 276.6251 | 325.8197 | 235.3642 | 487.4925 | 568.3388 | 573.4286 | 44.61444 | 40.41398 | 10.44878 | 60.07154 | 52.66754 | 74.50806 |
| Chrna6 | 11440 | 2.753501 | -3.00013 | 2.604041 | 0.016438 | 0.037425 | 0.164949 | -3.00013 | 0.19845 | 0.844571 | 0.908461 | 0 | 6.216245 | 3.471878 | 18.18193 | 14.04395 | 10.82134 | 7.106304 | 13.40422 | 39.07129 | 6.153716 | 0 | 7.463413 | 8.343269 | 20.43815 | 25.91585 |
| Ppp1r36 | 210762 | 2.753809 | -3.58674 | 3.16215 | 0.004634 | 0.011885 | 0.321786 | -3.58674 | 0.472727 | 0.641207 | 0.729491 | 2.225942 | 0 | 3.471878 | 3.896127 | 19.66153 | 18.93735 | 11.37009 | 13.40422 | 20.6848 | 0 | 3.050111 | 0 | 10.84625 | 11.79124 | 13.76779 |
| Gm136 | 214568 | 2.757983 | -4.21018 | 3.009725 | 0.006592 | 0.016428 | 0.768004 | -4.21018 | 1.0464 | 0.307083 | 0.401561 | 0 | 0 | 3.471878 | 5.194837 | 11.23516 | 2.705335 | 8.527565 | 10.72337 | 11.49156 | 1.538429 | 0 | 1.492683 | 3.337308 | 9.432991 | 4.049351 |
| Gch1 | 14528 | 2.758045 | 5.680202 | 53.56735 | 3.05E-24 | 4.07E-22 | -0.57346 | 5.680202 | -14.7093 | 1.20E-12 | 1.33E-11 | 1594.517 | 1872.866 | 1613.555 | 12302.67 | 12372.72 | 11833.14 | 8472.136 | 8471.465 | 8507.199 | 2563.023 | 2503.379 | 2407.697 | 627.4138 | 573.8403 | 668.9528 |
| Eif1ad3 | 100039042 | 2.758282 | -4.44036 | 2.651381 | 0.014807 | 0.034153 | 1.110481 | -4.44036 | 1.335734 | 0.195706 | 0.275476 | 0 | 0 | 0.86797 | 10.38967 | 9.830767 | 0 | 12.79135 | 5.361687 | 10.3424 | 0.769214 | 0 | 0 | 9.177596 | 14.93557 | 10.52831 |
| Robo2 | 268902 | 2.758871 | -0.00662 | 14.83756 | 1.01E-12 | 9.80E-12 | -2.80576 | -0.00662 | -13.124 | 1.09E-11 | 1.01E-10 | 52.68063 | 46.17782 | 54.68209 | 371.4308 | 362.334 | 382.805 | 65.378 | 41.55307 | 59.75609 | 6.92293 | 10.67539 | 13.43414 | 70.91779 | 66.03094 | 65.59949 |
| Gm10240 | 100503563 | 2.759589 | -2.88742 | 5.064784 | 4.91E-05 | 0.000167 | -1.5901 | -2.88742 | -3.20889 | 0.004155 | 0.009901 | 3.709904 | 3.55214 | 4.339848 | 19.48064 | 46.34504 | 31.11136 | 8.527565 | 17.42548 | 6.894934 | 2.307643 | 4.575167 | 5.224389 | 4.171635 | 6.288661 | 5.669092 |
| Ctsm | 64139 | 2.760838 | -4.63721 | 2.52397 | 0.019586 | 0.043755 | 0.281186 | -4.63721 | 0.302055 | 0.765534 | 0.839594 | 0.741981 | 0 | 0 | 3.896127 | 8.426372 | 1.352668 | 1.421261 | 5.361687 | 11.49156 | 2.307643 | 1.525056 | 0 | 16.68654 | 2.358248 | 1.619741 |
| Gm11978 | 100041311 | 2.761335 | -3.33456 | 3.233121 | 0.003927 | 0.010217 | 1.29514 | -3.33456 | 2.13316 | 0.044674 | 0.078673 | 2.225942 | 0 | 2.603909 | 7.792255 | 18.25714 | 8.116006 | 22.74017 | 24.12759 | 36.77298 | 0 | 2.287584 | 2.239024 | 11.68058 | 18.0799 | 5.669092 |
| Eppin | 75526 | 2.762327 | -3.85539 | 3.316368 | 0.003231 | 0.008533 | 1.173721 | -3.85539 | 1.811857 | 0.084104 | 0.134567 | 0 | 2.664105 | 0 | 7.792255 | 5.617581 | 2.705335 | 12.79135 | 9.382952 | 16.08818 | 2.307643 | 1.525056 | 2.239024 | 6.674615 | 12.57732 | 6.478962 |
| AA387883 | 100043450 | 2.763134 | -4.54005 | 2.778441 | 0.011153 | 0.026468 | 0.010394 | -4.54005 | 0.012246 | 0.990343 | 0.991889 | 0 | 0 | 3.471878 | 5.194837 | 2.808791 | 10.82134 | 2.842522 | 18.7659 | 3.447467 | 0 | 0 | 0 | 19.18952 | 5.502578 | 4.859222 |
| P2ry14 | 140795 | 2.763809 | 4.080204 | 17.33128 | 4.70E-14 | 5.74E-13 | 2.565891 | 4.080204 | 32.198 | 1.40E-19 | 8.17E-18 | 94.23155 | 140.3095 | 88.5329 | 733.7707 | 745.7339 | 804.8373 | 4772.594 | 4636.519 | 4631.097 | 852.2896 | 840.3057 | 756.0437 | 3494.995 | 3318.841 | 3495.4 |
| F10 | 14058 | 2.763943 | 2.765558 | 18.87999 | 8.43E-15 | 1.18E-13 | -0.84912 | 2.765558 | -8.04766 | 6.66E-08 | 3.35E-07 | 151.3641 | 139.4215 | 158.8384 | 1038.967 | 1072.958 | 1160.589 | 743.3194 | 670.2109 | 515.9709 | 1252.281 | 1274.947 | 1206.087 | 79.26106 | 93.54383 | 83.41664 |
| 2810032G | 72669 | 2.765493 | -4.5929 | 2.908147 | 0.008317 | 0.020297 | 0.90275 | -4.5929 | 1.183533 | 0.249613 | 0.337571 | 0 | 0 | 1.735939 | 10.38967 | 2.808791 | 2.705335 | 18.47639 | 4.021265 | 10.3424 | 0 | 0 | 0.746341 | 1.668654 | 6.288661 | 7.288832 |
| Themis2 | 230787 | 2.765661 | 8.504214 | 74.12265 | 1.70E-26 | 5.21E-24 | -0.86405 | 8.504214 | -31.4103 | 6.95E-19 | 3.31E-17 | 10999.12 | 11357.97 | 11451.99 | 81560.23 | 84147.15 | 79615.32 | 48121.05 | 48056.8 | 44249.39 | 21198.78 | 22060.69 | 21042.35 | 5274.615 | 5339.073 | 5036.583 |
| Mthfr | 17769 | 2.766942 | 8.112821 | 62.1818 | 1.72E-22 | 1.29E-20 | -1.48408 | 8.112821 | -39.5085 | 5.78E-19 | 2.80E-17 | 9274.017 | 9554.368 | 8861.97 | 67130.28 | 67807.01 | 66240.14 | 25400.77 | 26654.29 | 23038.27 | 20572.64 | 20353.39 | 19792.22 | 4181.647 | 3701.663 | 4026.675 |
| 1700011B( | 100503727 | 2.770202 | -2.80496 | 4.111221 | 0.000484 | 0.001458 | 2.710954 | -2.80496 | 6.770826 | 9.82E-07 | 4.18E-06 | 1.483961 | 4.440175 | 1.735939 | 18.18193 | 16.85274 | 21.64268 | 142.1261 | 120.638 | 133.302 | 3.076858 | 0.762528 | 0.746341 | 7.508942 | 6.288661 | 0.80987 |
| Dok6 | 623279 | 2.770286 | -3.22075 | 3.432195 | 0.002459 | 0.006627 | 0.626001 | -3.22075 | 1.002967 | 0.327113 | 0.423254 | 1.483961 | 2.664105 | 0 | 11.68838 | 7.021976 | 9.468674 | 22.74017 | 6.702109 | 21.83396 | 0.769214 | 3.050111 | 2.985365 | 18.35519 | 25.94073 | 20.24676 |
| Zfp800 | 627049 | 2.77052 | 6.213219 | 66.4218 | 3.18E-26 | 8.89E-24 | -1.59987 | 6.213219 | -43.3793 | 2.65E-22 | 3.93E-20 | 2680.034 | 2794.646 | 2705.461 | 20150.77 | 20011.23 | 19387.79 | 7133.308 | 6830.789 | 6491.58 | 3121.472 | 3244.556 | 3012.98 | 1438.38 | 1488.84 | 1538.754 |
| Runx1t1 | 12395 | 2.77262 | -0.75609 | 6.109447 | 4.30E-06 | 1.66E-05 | -0.06673 | -0.75609 | -0.22726 | 0.822385 | 0.889582 | 18.54952 | 18.64873 | 5.207818 | 80.51997 | 101.1165 | 100.0974 | 78.16935 | 91.14868 | 106.8715 | 17.69193 | 9.912862 | 2.985365 | 62.57452 | 66.81702 | 48.59222 |
| Zfp872 | 619310 | 2.77285 | -2.55887 | 3.466804 | 0.002265 | 0.006146 | 1.183789 | -2.55887 | 2.166497 | 0.041731 | 0.07421 | 5.193865 | 3.55214 | 0 | 18.18193 | 9.830767 | 18.93735 | 44.05909 | 24.12759 | 44.81707 | 3.846072 | 5.337695 | 5.97073 | 9.177596 | 16.50774 | 25.91585 |
| Pramel22 | 277668 | 2.776157 | -4.13351 | 2.619188 | 0.015899 | 0.036377 | 1.445004 | -4.13351 | 1.780087 | 0.089305 | 0.141759 | 0 | 0 | 1.735939 | 5.194837 | 4.213186 | 4.058003 | 14.21261 | 6.702109 | 25.28142 | 4.615287 | 0 | 1.492683 | 7.508942 | 11.79124 | 1.619741 |
| Efhb | 211482 | 2.777078 | -1.57962 | 4.007833 | 0.000621 | 0.001844 | 0.768518 | -1.57962 | 1.775679 | 0.090048 | 0.14274 | 13.35565 | 3.55214 | 1.735939 | 41.55869 | 21.06593 | 54.10671 | 35.53152 | 91.14868 | 79.29174 | 9.999788 | 17.53814 | 8.209754 | 30.8701 | 18.0799 | 21.8665 |
| Tmprss11I | 319875 | 2.780584 | -4.00917 | 3.420609 | 0.002527 | 0.006794 | 0.558384 | -4.00917 | 0.894406 | 0.38108 | 0.479353 | 0 | 1.77607 | 4.339848 | 7.792255 | 12.63956 | 12.17401 | 22.74017 | 9.382952 | 20.6848 | 0 | 0 | 0 | 8.343269 | 8.646909 | 2.429611 |
| Gm10847 | 100038618 | 2.781042 | 2.499474 | 22.23106 | 3.03E-16 | 5.68E-15 | -1.62393 | 2.499474 | -15.2348 | 6.05E-13 | 7.05E-12 | 220.3683 | 206.9122 | 217.8604 | 1571.438 | 1700.723 | 1485.229 | 521.6027 | 557.6154 | 520.5675 | 118.459 | 195.9697 | 113.4439 | 166.8654 | 158.7887 | 200.8478 |
| Rhbdf2 | 217344 | 2.781493 | 8.162465 | 76.67417 | 1.50E-27 | 5.90E-25 | -0.78076 | 8.162465 | -29.3205 | 9.88E-19 | 4.50E-17 | 8471.936 | 8589.074 | 7996.604 | 60917.25 | 62609.35 | 60498.06 | 38233.34 | 38167.17 | 35189.44 | 15285.06 | 15339.77 | 15350 | 5104.412 | 4899.653 | 5150.775 |
| Ms4a4b | 60361 | 2.783739 | 3.143354 | 22.57547 | 2.21E-16 | 4.26E-15 | 0.93748 | 3.143354 | 13.2296 | 9.32E-12 | 8.80E-11 | 204.7867 | 165.1745 | 243.8995 | 1602.607 | 1379.116 | 1496.051 | 2950.538 | 2876.545 | 3059.052 | 206.9187 | 218.8455 | 167.9268 | 245.2921 | 223.2475 | 230.813 |
| Tiparp | 99929 | 2.78379 | 6.702011 | 77.98874 | 1.05E-27 | 4.37E-25 | -0.53688 | 6.702011 | -20.2955 | 1.95E-15 | 3.90E-14 | 2555.382 | 2719.163 | 2722.821 | 19496.22 | 19924.16 | 19282.28 | 13867.24 | 13782.22 | 14285.15 | 3163.779 | 3075.275 | 2999.545 | 4125.747 | 4170.168 | 4181.36 |
| I830134H0 | 619304 | 2.785057 | -1.9129 | 4.38028 | 0.000253 | 0.000791 | 1.165088 | -1.9129 | 2.89292 | 0.00861 | 0.018925 | 2.225942 | 1.77607 | 6.943757 | 18.18193 | 19.66153 | 43.28537 | 66.79926 | 54.95729 | 56.30862 | 6.153716 | 3.050111 | 9.702436 | 30.03577 | 41.66238 | 22.67637 |
| Gm614 | 245536 | 2.785079 | 1.165971 | 13.24524 | 9.12E-12 | 7.60E-11 | 0.44701 | 1.165971 | 3.662533 | 0.001422 | 0.003767 | 49.71271 | 55.9462 | 37.32269 | 357.145 | 356.7164 | 332.7563 | 510.2326 | 467.8072 | 503.3301 | 65.38323 | 99.12862 | 95.53168 | 60.07154 | 55.02578 | 70.45871 |
| Adgra1 | 52389 | 2.785797 | -2.29864 | 3.126943 | 0.005029 | 0.012818 | 1.5073 | -2.29864 | 2.499724 | 0.020645 | 0.040551 | 0 | 1.77607 | 9.547666 | 11.68838 | 18.25714 | 12.17401 | 25.5827 | 40.21265 | 64.35271 | 3.076858 | 6.100223 | 5.97073 | 56.73423 | 45.59279 | 28.34546 |
| Gm9639 | 675294 | 2.787381 | -4.18761 | 3.476904 | 0.002212 | 0.006011 | -0.54346 | -4.18761 | -0.77453 | 0.447114 | 0.545839 | 0.741981 | 4.440175 | 0 | 7.792255 | 7.021976 | 13.52668 | 2.842522 | 5.361687 | 13.78987 | 0 | 0 | 0 | 15.85221 | 8.646909 | 9.718443 |
| Olfr12 | 257890 | 2.789211 | -4.56909 | 3.001237 | 0.006722 | 0.016728 | 0.052353 | -4.56909 | 0.065219 | 0.948607 | 0.957342 | 0 | 0.888035 | 0 | 2.597418 | 7.021976 | 2.705335 | 1.421261 | 4.021265 | 9.193245 | 0.769214 | 2.287584 | 0 | 7.508942 | 11.79124 | 4.859222 |
| Tex28 | 385380 | 2.78956 | -4.26836 | 3.20416 | 0.004201 | 0.010867 | -0.53359 | -4.26836 | -0.70218 | 0.490157 | 0.588336 | 2.967923 | 0.888035 | 0 | 2.597418 | 19.66153 | 12.17401 | 5.685043 | 2.680843 | 12.64071 | 0 | 0 | 0 | 13.34923 | 9.432991 | 6.478962 |
| Ccdc146 | 75172 | 2.790142 | 1.416051 | 15.80166 | 2.95E-13 | 3.13E-12 | -0.29654 | 1.416051 | -2.60411 | 0.016435 | 0.033292 | 80.8759 | 61.27441 | 60.75787 | 487.0159 | 539.2878 | 484.2551 | 427.7995 | 408.8286 | 435.53 | 66.15244 | 78.54037 | 55.22925 | 97.61625 | 128.9175 | 98.80417 |
| Fam131b | 76156 | 2.790985 | -1.27558 | 3.726582 | 0.00122 | 0.003465 | 2.627327 | -1.27558 | 6.354358 | 2.47E-06 | 9.99E-06 | 9.64575 | 0.888035 | 4.339848 | 23.37676 | 26.68351 | 41.9327 | 159.1812 | 209.1058 | 212.5938 | 16.1535 | 18.30067 | 2.985365 | 38.37904 | 51.88145 | 28.34546 |
| Fmnl2 | 71409 | 2.792895 | 7.005587 | 65.65776 | 4.06E-26 | 1.06E-23 | 0.365676 | 7.005587 | 13.20447 | 9.67E-12 | 9.11E-11 | 2591.739 | 2743.14 | 2541.415 | 19249.47 | 19978.93 | 19017.16 | 26507.94 | 25933.14 | 25511.25 | 3429.158 | 3244.556 | 3418.243 | 5912.875 | 5840.594 | 6268.396 |
| A730011C | 319916 | 2.798158 | 1.72886 | 18.18298 | 1.80E-14 | 2.37E-13 | -0.59258 | 1.72886 | -5.67003 | 1.18E-05 | 4.34E-05 | 113.5231 | 112.7804 | 85.06102 | 803.901 | 775.2262 | 730.4406 | 554.2917 | 512.0411 | 525.1641 | 106.9208 | 94.55345 | 88.06827 | 102.6222 | 61.31444 | 65.59949 |
| Saxo1 | 75811 | 2.799971 | -3.62215 | 3.083056 | 0.005567 | 0.014082 | 1.244786 | -3.62215 | 1.800891 | 0.085868 | 0.137065 | 0.741981 | 0 | 0.86797 | 3.896127 | 2.808791 | 16.23201 | 9.948826 | 13.40422 | 24.13227 | 0.769214 | 3.050111 | 2.985365 | 22.52683 | 12.57732 | 7.288832 |
| Nlrp3 | 216799 | 2.800215 | 6.932631 | 60.25685 | 2.52E-25 | 4.94E-23 | 0.271758 | 6.932631 | 9.133841 | 8.00E-09 | 4.61E-08 | 4178.094 | 4336.275 | 4295.582 | 32091.1 | 32129.75 | 30965.27 | 41274.84 | 40593.33 | 37876.17 | 3529.156 | 3380.286 | 3339.877 | 1165.555 | 1048.634 | 1162.974 |
| Gm2897 | 100040671 | 2.800832 | -3.42143 | 3.335531 | 0.003088 | 0.008179 | 0.068825 | -3.42143 | 0.101329 | 0.920235 | 0.935277 | 0 | 3.55214 | 2.603909 | 11.68838 | 11.23516 | 10.82134 | 12.79135 | 5.361687 | 24.13227 | 0.769214 | 3.050111 | 2.985365 | 15.01788 | 7.860826 | 5.669092 |
| Gm15340 | 100040997 | 2.800994 | -2.87119 | 3.919094 | 0.000769 | 0.002249 | 0.499169 | -2.87119 | 0.970126 | 0.34285 | 0.44004 | 0.741981 | 1.77607 | 6.075787 | 7.792255 | 36.51428 | 20.29002 | 22.74017 | 20.10633 | 37.92213 | 10.769 | 11.43792 | 11.94146 | 1.668654 | 1.572165 | 3.239481 |
| Gm7160 | 635504 | 2.80159 | 1.013312 | 11.7374 | 8.92E-11 | 6.43E-10 | -0.25289 | 1.013312 | -1.63122 | 0.117514 | 0.178945 | 29.67923 | 74.59494 | 60.75787 | 416.8856 | 356.7164 | 346.2829 | 285.6734 | 351.1905 | 336.7026 | 103.0747 | 107.5164 | 129.117 | 37.54471 | 36.94588 | 33.20468 |
| Ovol1 | 18426 | 2.801968 | 0.050098 | 8.27959 | 4.18E-08 | 2.07E-07 | -1.02396 | 0.050098 | -4.03571 | 0.000581 | 0.001662 | 28.19527 | 39.07354 | 13.88751 | 176.6244 | 196.6153 | 192.0788 | 78.16935 | 117.9571 | 94.23076 | 24.61486 | 51.08937 | 26.86829 | 76.75808 | 44.02063 | 63.97975 |
| Sdr16c6 | 242286 | 2.805825 | -4.12398 | 3.336063 | 0.003084 | 0.00817 | -1.23889 | -4.12398 | -1.60138 | 0.123991 | 0.187262 | 0.741981 | 0 | 4.339848 | 5.194837 | 12.63956 | 12.17401 | 4.263783 | 5.361687 | 2.298311 | 0.769214 | 0.762528 | 0.746341 | 13.34923 | 8.646909 | 2.429611 |
| Gm11346 | 76024 | 2.80614 | -1.92109 | 3.651349 | 0.00146 | 0.004088 | -0.20168 | -1.92109 | -0.36846 | 0.716162 | 0.796304 | 13.35565 | 0 | 13.01954 | 35.06515 | 28.08791 | 36.52203 | 46.90161 | 20.10633 | 28.72889 | 6.153716 | 8.387806 | 5.97073 | 35.04173 | 28.29897 | 27.53559 |
| Olfr1300-p | 258199 | 2.806597 | -3.73516 | 3.434569 | 0.002445 | 0.006593 | 1.954627 | -3.73516 | 3.239108 | 0.003872 | 0.0093 | 0.741981 | 0 | 0 | 6.493546 | 5.617581 | 1.352668 | 14.21261 | 14.74464 | 24.13227 | 1.538429 | 1.525056 | 2.985365 | 16.68654 | 18.86598 | 12.95792 |
| Fam177a | 73385 | 2.807214 | -1.89834 | 3.654719 | 0.001449 | 0.004057 | 0.966436 | -1.89834 | 1.943252 | 0.065291 | 0.108595 | 8.161788 | 8.88035 | 0 | 35.06515 | 16.85274 | 22.99535 | 59.69296 | 41.55307 | 49.41369 | 16.1535 | 20.58825 | 20.15121 | 17.52087 | 8.646909 | 15.38754 |
| A730020E( | 100504262 | 2.807461 | -0.76603 | 6.134988 | 4.06E-06 | 1.57E-05 | -0.77534 | -0.76603 | -2.33724 | 0.02924 | 0.054716 | 14.09763 | 10.65642 | 11.2836 | 93.50706 | 124.9912 | 67.63339 | 51.16539 | 64.34024 | 52.86116 | 29.23015 | 48.03925 | 51.49755 | 23.36115 | 33.80155 | 8.908573 |
| Plcxd3 | 239318 | 2.808113 | -4.06196 | 2.925762 | 0.00799 | 0.019578 | 1.000758 | -4.06196 | 1.331663 | 0.197019 | 0.277146 | 0 | 0 | 2.603909 | 2.597418 | 4.213186 | 12.17401 | 14.21261 | 10.72337 | 10.3424 | 0.769214 | 7.625279 | 0.746341 | 6.674615 | 2.358248 | 4.859222 |
| Gm15354 | 100503651 | 2.808959 | 1.008502 | 20.19232 | 2.16E-15 | 3.36E-14 | -4.2221 | 1.008502 | -18.0256 | 2.14E-14 | 3.41E-13 | 152.848 | 129.6531 | 124.1197 | 1102.604 | 957.7976 | 998.2688 | 75.32683 | 54.95729 | 44.81707 | 105.3824 | 137.255 | 125.3853 | 31.70442 | 29.08506 | 17.81715 |
| Olfr715 | 258776 | 2.809486 | -5.35693 | 2.58007 | 0.017327 | 0.039217 | -0.59834 | -5.35693 | -0.60128 | 0.553985 | 0.648681 | 0 | 0 | 0 | 11.68838 | 2.808791 | 0 | 1.421261 | 0 | 5.745778 | 0 | 0 | 0 | 5.840288 | 11.79124 | 3.239481 |
| Ahr | 11622 | 2.809628 | 4.524907 | 21.78758 | 4.58E-16 | 8.16E-15 | 2.527304 | 4.524907 | 38.88152 | 2.66E-21 | 2.58E-19 | 132.8146 | 198.0318 | 138.0072 | 1164.942 | 1087.002 | 1174.116 | 6841.95 | 6648.492 | 6997.208 | 1064.593 | 1011.112 | 1110.556 | 3903.816 | 3775.555 | 3923.821 |
| Slfn8 | 276950 | 2.811025 | 8.674977 | 86.46132 | 1.17E-28 | 1.10E-25 | -0.34166 | 8.674977 | -16.0224 | 2.24E-13 | 2.86E-12 | 11900.63 | 11976.04 | 10812.3 | 86427.8 | 87816.84 | 85738.84 | 71877.43 | 71862.69 | 69519.32 | 14855.07 | 15293.26 | 14243.18 | 8270.683 | 8356.844 | 8184.549 |
| Pdcd1lg2 | 58205 | 2.812606 | 2.213491 | 11.67793 | 9.80E-11 | 7.02E-10 | -0.16513 | 2.213491 | -1.08358 | 0.29064 | 0.383684 | 45.26083 | 40.84961 | 28.643 | 271.4302 | 294.923 | 294.8816 | 220.2954 | 265.4035 | 307.9737 | 2229.953 | 2213.618 | 2083.785 | 242.7891 | 267.2681 | 276.9756 |
| Mir222 | 723828 | 2.818061 | -0.94922 | 6.748989 | 1.03E-06 | 4.26E-06 | -1.37794 | -0.94922 | -4.11203 | 0.000483 | 0.0014 | 25.96933 | 16.87266 | 9.547666 | 155.8451 | 134.8219 | 96.03941 | 68.22052 | 41.55307 | 45.96622 | 20.76879 | 29.73859 | 8.956095 | 8.343269 | 16.50774 | 20.24676 |
| Gm13077 | 100504731 | 2.818437 | -3.32484 | 2.686831 | 0.013688 | 0.03183 | -0.61 | -3.32484 | -0.67964 | 0.50404 | 0.601622 | 1.483961 | 0.888035 | 2.603909 | 20.77935 | 30.8967 | 4.058003 | 2.842522 | 8.04253 | 27.57973 | 8.461359 | 4.575167 | 0 | 21.6925 | 6.288661 | 8.098703 |
| A730036I1 | 329513 | 2.819108 | -3.9789 | 3.283031 | 0.003494 | 0.00917 | 1.072787 | -3.9789 | 1.594827 | 0.125453 | 0.189086 | 1.483961 | 0 | 0 | 2.597418 | 4.213186 | 9.468674 | 11.37009 | 6.702109 | 17.23733 | 3.076858 | 1.525056 | 0.746341 | 13.34923 | 7.074744 | 6.478962 |
| Trav8d-2 | 667604 | 2.822684 | -4.77639 | 2.752948 | 0.01181 | 0.027862 | -0.44353 | -4.77639 | -0.47897 | 0.636833 | 0.725677 | 0 | 0 | 0 | 6.493546 | 5.617581 | 0 | 4.263783 | 0 | 3.447467 | 3.076858 | 2.287584 | 2.239024 | 3.337308 | 5.502578 | 7.288832 |
| A230057D | 319893 | 2.826696 | -4.37084 | 3.199238 | 0.00425 | 0.010983 | -0.1948 | -4.37084 | -0.26054 | 0.796942 | 0.866847 | 0 | 2.664105 | 0.86797 | 7.792255 | 11.23516 | 5.410671 | 19.89765 | 8.04253 | 2.298311 | 0 | 0 | 0 | 8.343269 | 3.14433 | 9.718443 |
| 2810433D( | 77132 | 2.826703 | -3.63323 | 2.742727 | 0.012083 | 0.028451 | 0.346512 | -3.63323 | 0.42884 | 0.672337 | 0.757081 | 0 | 3.55214 | 1.735939 | 1.298709 | 19.66153 | 33.81669 | 12.79135 | 20.10633 | 10.3424 | 0 | 1.525056 | 0 | 19.18952 | 14.14949 | 16.19741 |
| Olfr479 | 257891 | 2.827956 | -4.65911 | 2.857281 | 0.009336 | 0.022547 | -1.27192 | -4.65911 | -1.37894 | 0.182195 | 0.259331 | 0 | 0 | 4.339848 | 5.194837 | 4.213186 | 10.82134 | 4.263783 | 0 | 6.894934 | 0 | 1.525056 | 0.746341 | 4.171635 | 2.358248 | 5.669092 |
| Olfr462 | 258406 | 2.828795 | -5.15366 | 2.837139 | 0.009771 | 0.023517 | -1.99996 | -5.15366 | -2.04622 | 0.053251 | 0.091461 | 1.483961 | 0 | 0.86797 | 2.597418 | 16.85274 | 8.116006 | 0 | 0 | 14.93902 | 0 | 0 | 0 | 0.834327 | 3.930413 | 1.619741 |
| Olfr1311 | 258271 | 2.829077 | -5.0436 | 2.851096 | 0.009467 | 0.022844 | 1.595531 | -5.0436 | 2.040989 | 0.053811 | 0.092291 | 0 | 0 | 0 | 5.194837 | 0 | 6.763339 | 18.47639 | 5.361687 | 8.044089 | 0 | 0.762528 | 0 | 2.502981 | 1.572165 | 3.239481 |
| Gm9839 | 408192 | 2.829713 | -4.52897 | 3.464329 | 0.002278 | 0.006179 | 0.522702 | -4.52897 | 0.776697 | 0.445861 | 0.544612 | 0 | 0 | 1.735939 | 5.194837 | 7.021976 | 2.705335 | 4.263783 | 9.382952 | 9.193245 | 0 | 0 | 0 | 13.34923 | 11.00516 | 6.478962 |
| Gpx8 | 69590 | 2.831201 | -4.49463 | 2.909298 | 0.008295 | 0.020254 | 1.018232 | -4.49463 | 1.309606 | 0.204252 | 0.285575 | 0 | 0.888035 | 0 | 5.194837 | 7.021976 | 1.352668 | 5.685043 | 14.74464 | 8.044089 | 3.846072 | 0.762528 | 0 | 1.668654 | 5.502578 | 5.669092 |
| Cyp2d11 | 545123 | 2.83147 | -3.87063 | 3.266609 | 0.003631 | 0.009508 | 1.254892 | -3.87063 | 1.896505 | 0.071511 | 0.11734 | 0 | 0 | 2.603909 | 5.194837 | 7.021976 | 4.058003 | 18.47639 | 9.382952 | 16.08818 | 0.769214 | 3.050111 | 0.746341 | 5.840288 | 9.432991 | 15.38754 |
| Tcp10a | 21461 | 2.831699 | -2.69982 | 3.679293 | 0.001366 | 0.003846 | 2.091948 | -2.69982 | 4.265849 | 0.000334 | 0.000992 | 2.225942 | 1.77607 | 0.86797 | 16.88322 | 12.63956 | 13.52668 | 41.21656 | 85.78699 | 75.84427 | 0.769214 | 0.762528 | 7.463413 | 12.5149 | 12.57732 | 23.48624 |
| Gm7732 | 665654 | 2.834199 | -5.18084 | 3.189483 | 0.004348 | 0.011214 | -2.20455 | -5.18084 | -2.49696 | 0.020769 | 0.040758 | 0.741981 | 0 | 0.86797 | 3.896127 | 5.617581 | 9.468674 | 0 | 0 | 5.745778 | 0 | 0 | 2.239024 | 1.668654 | 1.572165 | 1.619741 |
| Il2ra | 16184 | 2.834831 | 2.607862 | 34.12025 | 4.15E-20 | 1.79E-18 | -4.16573 | 2.607862 | -30.6417 | 3.94E-19 | 2.02E-17 | 594.3266 | 500.8517 | 512.97 | 4100.025 | 4100.834 | 4089.115 | 191.8702 | 302.9353 | 226.3837 | 221.5338 | 287.473 | 276.8926 | 37.54471 | 58.17011 | 47.78235 |

| | | | | | | | | | | | | | | | | | | | | | | | | | | |
|---|---|---|---|---|---|---|---|---|---|---|---|---|---|---|---|---|---|---|---|---|---|---|---|---|---|---|
| Nppb | 18158 | 2.836592 | -4.76664 | 2.873485 | 0.008999 | 0.021812 | -0.25215 | -4.76664 | -0.29262 | 0.772637 | 0.84575 | 0 | 1.77607 | 0 | 1.298709 | 8.426372 | 8.116006 | 1.421261 | 5.361687 | 6.894934 | 0 | 0 | 1.492683 | 5.005962 | 1.572165 | 8.098703 |
| Rsph4a | 212892 | 2.840434 | 0.370667 | 11.36578 | 1.62E-10 | 1.12E-09 | -0.4537 | 0.370667 | -2.72924 | 0.012453 | 0.026165 | 27.45329 | 50.61799 | 27.77503 | 284.4173 | 227.512 | 262.4175 | 207.5041 | 184.9782 | 194.2073 | 50.76815 | 30.50111 | 38.0634 | 44.21933 | 37.73197 | 38.87377 |
| Gm9731 | 732482 | 2.841382 | -4.43647 | 2.462919 | 0.022355 | 0.049211 | 1.020502 | -4.43647 | 1.074044 | 0.294796 | 0.388293 | 0 | 0 | 0 | 2.597418 | 14.04395 | 0 | 4.263783 | 4.021265 | 11.49156 | 0.769214 | 0 | 5.97073 | 15.85221 | 9.432991 | 13.76779 |
| Svs3a | 64335 | 2.841405 | -4.73509 | 2.618262 | 0.015931 | 0.036445 | 0.376367 | -4.73509 | 0.413199 | 0.683583 | 0.766789 | 0.741981 | 0 | 0 | 2.597418 | 8.426372 | 2.705335 | 1.421261 | 8.04253 | 11.49156 | 0 | 3.050111 | 0 | 6.674615 | 8.646909 | 0.80987 |
| Il7 | 16196 | 2.841419 | 0.654604 | 10.84839 | 3.80E-10 | 2.51E-09 | 0.193621 | 0.654604 | 1.243301 | 0.227242 | 0.312147 | 34.8731 | 33.74533 | 46.00239 | 283.1186 | 276.6659 | 315.1716 | 308.4136 | 414.1903 | 325.211 | 74.6138 | 95.31598 | 88.06827 | 17.52087 | 40.09021 | 14.57766 |
| Igkv8-24 | 677858 | 2.842357 | -2.73787 | 3.184392 | 0.0044 | 0.011335 | -0.24253 | -2.73787 | -0.34444 | 0.733889 | 0.81217 | 0 | 5.32821 | 7.811726 | 10.38967 | 25.27911 | 24.34802 | 17.05513 | 10.72337 | 21.83396 | 10.769 | 16.77561 | 14.92683 | 0.834327 | 8.646909 | 9.718443 |
| Ccl7 | 20306 | 2.843772 | 9.076145 | 70.11407 | 6.53E-23 | 5.57E-21 | -0.65789 | 9.076145 | -23.4831 | 1.09E-14 | 1.85E-13 | 12051.25 | 12201.6 | 11478.9 | 91081.07 | 93222.35 | 89691.34 | 60224.51 | 56849.97 | 62769.18 | 29680.91 | 28520.83 | 29123.73 | 18101.56 | 17377.93 | 19933.34 |
| Ctcfl | 664799 | 2.844798 | -2.6107 | 3.294206 | 0.003403 | 0.008952 | 0.931338 | -2.6107 | 1.57078 | 0.130941 | 0.196167 | 5.935846 | 0 | 3.471878 | 24.67547 | 15.44835 | 12.17401 | 32.689 | 21.44675 | 51.712 | 6.92293 | 0.762528 | 1.492683 | 39.21337 | 29.87114 | 17.00728 |
| Samt4 | 75185 | 2.845401 | -4.96961 | 2.811792 | 0.010347 | 0.024739 | -0.00099 | -4.96961 | -0.00112 | 0.999119 | 0.999256 | 0 | 0 | 0 | 2.597418 | 4.213186 | 1.352668 | 0 | 4.021265 | 9.193245 | 2.307643 | 0 | 0.746341 | 6.674615 | 5.502578 | 4.049351 |
| Igkv8-19 | 232065 | 2.846376 | -4.27553 | 2.924613 | 0.008011 | 0.019624 | -3.09946 | -4.27553 | -3.00541 | 0.006658 | 0.015067 | 0 | 3.55214 | 1.735939 | 5.194837 | 11.23516 | 18.93735 | 0 | 5.361687 | 0 | 9.999788 | 6.100223 | 0.746341 | 3.337308 | 3.14433 | 1.619741 |
| Gpr143 | 18241 | 2.846772 | -3.69465 | 2.958774 | 0.007409 | 0.01827 | 1.267068 | -3.69465 | 1.760033 | 0.092728 | 0.146293 | 0 | 5.32821 | 0 | 10.38967 | 9.830767 | 2.705335 | 17.05513 | 12.0638 | 25.28142 | 0.769214 | 1.525056 | 0 | 15.01788 | 26.72681 | 12.14805 |
| Pcdhb4 | 93875 | 2.848506 | -2.54448 | 3.551464 | 0.001853 | 0.005101 | 1.060747 | -2.54448 | 1.903619 | 0.070532 | 0.115992 | 5.935846 | 1.77607 | 0 | 9.090964 | 9.830767 | 28.40602 | 25.5827 | 40.21265 | 27.57973 | 6.92293 | 4.575167 | 4.478048 | 22.52683 | 33.80155 | 18.62702 |
| D730005E | 109361 | 2.849104 | 3.469253 | 22.41186 | 2.57E-16 | 4.88E-15 | -0.04924 | 3.469253 | -0.60127 | 0.553996 | 0.648681 | 293.8244 | 275.2908 | 256.051 | 2315.598 | 1983.006 | 2073.64 | 2274.017 | 2242.526 | 1922.537 | 399.2223 | 541.3948 | 461.2389 | 211.919 | 202.8093 | 244.5808 |
| Gm10466 | 100038617 | 2.849304 | -3.5611 | 2.757515 | 0.01169 | 0.027601 | 0.153714 | -3.5611 | 0.184254 | 0.855553 | 0.91732 | 2.225942 | 0 | 1.735939 | 15.58451 | 11.23516 | 5.410671 | 4.263783 | 9.382952 | 31.0272 | 0 | 10.67539 | 0 | 13.34923 | 20.43815 | 17.00728 |
| Arhgef3 | 71704 | 2.850336 | 5.566727 | 65.22731 | 4.67E-26 | 1.19E-23 | -1.74138 | 5.566727 | -44.4983 | 1.55E-22 | 2.53E-20 | 1587.839 | 1588.695 | 1628.311 | 12029.94 | 12770.17 | 12225.41 | 3749.286 | 3834.947 | 3886.444 | 2480.717 | 2478.978 | 2498.751 | 1023.719 | 903.995 | 988.8516 |
| Tma16 | 66282 | 2.850998 | 3.733082 | 32.01686 | 1.57E-19 | 5.93E-18 | -0.72013 | 3.733082 | -11.0294 | 2.81E-10 | 2.03E-09 | 476.3516 | 551.4697 | 372.359 | 3623.399 | 3561.546 | 3472.298 | 2245.592 | 2176.845 | 2285.67 | 449.2212 | 501.7433 | 521.6925 | 188.5579 | 167.4356 | 173.3122 |
| Nkx2-6 | 18092 | 2.851542 | -3.80311 | 3.363518 | 0.002891 | 0.007696 | 0.016207 | -3.80311 | 0.023262 | 0.981657 | 0.984652 | 1.483961 | 1.77607 | 0 | 6.493546 | 8.426372 | 12.17401 | 4.263783 | 30.8297 | 5.745778 | 0 | 0 | 0.746341 | 20.85817 | 27.51289 | 20.24676 |
| Asic2 | 11418 | 2.851636 | -2.00527 | 3.835534 | 0.00094 | 0.002713 | 1.383259 | -2.00527 | 2.965203 | 0.007301 | 0.016352 | 5.935846 | 8.88035 | 0 | 19.48064 | 21.06593 | 27.05335 | 52.58665 | 58.97856 | 72.3968 | 4.615287 | 6.100223 | 2.239024 | 37.54471 | 45.59279 | 36.44416 |
| Olfr461 | 258380 | 2.851887 | -4.0028 | 2.629603 | 0.015538 | 0.035641 | 1.868472 | -4.0028 | 2.31907 | 0.030383 | 0.056514 | 0 | 0 | 0.86797 | 2.597418 | 4.213186 | 5.410671 | 5.685043 | 28.14886 | 25.28142 | 0 | 6.862751 | 2.985365 | 6.674615 | 6.288661 | 8.098703 |
| Gm830 | 329892 | 2.852239 | -3.16781 | 3.643905 | 0.001487 | 0.004155 | 0.022179 | -3.16781 | 0.03549 | 0.972019 | 0.976847 | 1.483961 | 0 | 4.339848 | 11.68838 | 14.04395 | 9.468674 | 14.21261 | 14.74464 | 9.193245 | 6.153716 | 1.525056 | 3.731706 | 10.01192 | 22.7964 | 10.52831 |
| Otud6a | 408193 | 2.853199 | -3.11851 | 3.511017 | 0.00204 | 0.005576 | -0.4399 | -3.11851 | -0.6403 | 0.528791 | 0.624954 | 0 | 1.77607 | 5.207818 | 6.493546 | 12.63956 | 20.29002 | 7.106304 | 10.72337 | 9.193245 | 9.230573 | 7.625279 | 2.985365 | 14.18356 | 12.57732 | 8.908573 |
| Mndal | 100040462 | 2.853849 | 8.67617 | 78.56972 | 8.94E-28 | 4.02E-25 | 0.388912 | 8.67617 | 18.10487 | 1.96E-14 | 3.14E-13 | 8034.168 | 8390.154 | 7771.8 | 62717.26 | 62456.27 | 61425.99 | 86958.42 | 83738.83 | 83342.51 | 13893.55 | 14209.71 | 13073.66 | 15106.32 | 15154.89 | 14763.93 |
| Igtp | 16145 | 2.855106 | 7.634273 | 64.93698 | 5.14E-26 | 1.25E-23 | 1.447395 | 7.634273 | 59.7464 | 3.01E-25 | 1.15E-22 | 4057.893 | 4639.095 | 3931.034 | 32646.95 | 32813.7 | 31670.01 | 93530.33 | 89211.77 | 92218.59 | 5400.655 | 5560.353 | 5549.794 | 3606.795 | 3399.807 | 3335.046 |
| Hsd17b2 | 15486 | 2.85541 | -3.51526 | 2.83115 | 0.009904 | 0.023807 | 0.51735 | -3.51526 | 0.660137 | 0.516228 | 0.613263 | 0.741981 | 0 | 4.339848 | 7.792255 | 8.426372 | 13.52668 | 5.685043 | 18.7659 | 25.28142 | 1.538429 | 0 | 5.224389 | 18.35519 | 4.716496 | 17.81715 |
| Tbx15 | 21384 | 2.855876 | -2.03676 | 3.235166 | 0.003908 | 0.010175 | 1.069456 | -2.03676 | 1.890979 | 0.072279 | 0.118399 | 7.419807 | 0 | 6.943757 | 22.07806 | 39.32307 | 13.52668 | 54.00791 | 58.97856 | 41.3696 | 13.84586 | 8.387806 | 4.478048 | 30.8701 | 9.432991 | 33.20468 |
| Gm1604b | 381059 | 2.856025 | -4.26209 | 2.94419 | 0.00766 | 0.01884 | 0.336887 | -4.26209 | 0.418275 | 0.679925 | 0.763663 | 0 | 1.77607 | 0 | 2.597418 | 14.04395 | 2.705335 | 11.37009 | 10.72337 | 2.298311 | 0.769214 | 0.762528 | 0.746341 | 4.171635 | 11.79124 | 12.95792 |
| 1700006A | 71824 | 2.856891 | -3.56359 | 3.153759 | 0.004725 | 0.012101 | 0.756314 | -3.56359 | 1.072947 | 0.295277 | 0.38879 | 4.451884 | 0 | 0 | 3.896127 | 11.23516 | 8.116006 | 12.79135 | 6.702109 | 22.98311 | 0.769214 | 3.050111 | 3.731706 | 11.68058 | 15.72165 | 10.52831 |
| Golga7b | 71146 | 2.857117 | -2.48909 | 3.010619 | 0.006579 | 0.016397 | 1.507436 | -2.48909 | 2.373105 | 0.027098 | 0.051263 | 2.225942 | 5.32821 | 0 | 15.58451 | 11.23516 | 14.87935 | 22.74017 | 53.61687 | 56.30862 | 0 | 7.625279 | 3.731706 | 47.55663 | 46.37887 | 63.16988 |
| Gm10848 | 100038578 | 2.857223 | -0.22434 | 4.775887 | 9.79E-05 | 0.000321 | 2.361786 | -0.22434 | 7.654017 | 1.49E-07 | 7.12E-07 | 3.709904 | 13.32052 | 8.679696 | 49.35095 | 51.96262 | 82.51273 | 332.575 | 377.9989 | 270.0516 | 10.769 | 33.55123 | 5.97073 | 151.8475 | 190.232 | 161.1642 |
| Zfp936 | 668620 | 2.857945 | -2.52213 | 3.756265 | 0.001136 | 0.00324 | 0.953014 | -2.52213 | 1.828553 | 0.081477 | 0.131043 | 0.741981 | 1.77607 | 4.339848 | 19.48064 | 7.021976 | 32.46403 | 39.7953 | 34.85097 | 29.87805 | 3.076858 | 0.762528 | 7.463413 | 33.37308 | 23.58248 | 26.72572 |
| Ucn | 22226 | 2.858801 | -4.05742 | 2.705079 | 0.013143 | 0.030683 | 0.856799 | -4.05742 | 1.014722 | 0.321604 | 0.41723 | 0 | 0 | 1.735939 | 3.896127 | 2.808791 | 9.468674 | 14.21261 | 2.680843 | 19.53565 | 4.615287 | 0 | 0.746341 | 10.01192 | 18.86598 | 8.098703 |
| Gm381 | 214308 | 2.859421 | -3.79576 | 3.608148 | 0.001619 | 0.004501 | -0.54775 | -3.79576 | -0.81481 | 0.424183 | 0.523181 | 0.741981 | 1.77607 | 1.735939 | 12.98709 | 19.66153 | 8.116006 | 9.948826 | 6.702109 | 10.3424 | 4.615287 | 0 | 0 | 10.84625 | 9.432991 | 3.239481 |
| Pramel28 | 626922 | 2.860371 | -4.93949 | 2.611072 | 0.016185 | 0.036962 | 0.31205 | -4.93949 | 0.341974 | 0.735715 | 0.813813 | 1.483961 | 0 | 0 | 10.38967 | 7.021976 | 1.352668 | 1.421261 | 18.7659 | 8.044089 | 0 | 0 | 1.492683 | 4.171635 | 1.572165 | 0 |
| Lim2 | 233187 | 2.860547 | -4.25121 | 3.121239 | 0.005096 | 0.012971 | 0.92935 | -4.25121 | 1.274627 | 0.216144 | 0.299216 | 0 | 1.77607 | 0 | 2.597418 | 4.213186 | 9.468674 | 5.685043 | 6.702109 | 22.98311 | 0 | 0.762528 | 1.492683 | 10.01192 | 7.074744 | 6.478962 |
| Crygf | 12969 | 2.861185 | -4.40028 | 3.307079 | 0.003302 | 0.008705 | 1.518129 | -4.40028 | 2.301184 | 0.031549 | 0.058377 | 0 | 0.888035 | 0 | 5.194837 | 4.213186 | 2.705335 | 9.948826 | 17.42548 | 12.64071 | 0 | 0.762528 | 0 | 4.171635 | 8.646909 | 16.19741 |
| Fut9 | 14348 | 2.861954 | -1.99355 | 4.199161 | 0.000392 | 0.001195 | 0.584868 | -1.99355 | 1.307572 | 0.204929 | 0.286298 | 8.903769 | 3.55214 | 0.86797 | 27.27289 | 43.53625 | 18.93735 | 46.90161 | 28.14886 | 62.0544 | 4.615287 | 6.100223 | 2.239024 | 33.37308 | 50.30929 | 29.9652 |
| Casp4 | 12363 | 2.863313 | 5.022499 | 52.23491 | 5.20E-24 | 6.39E-22 | -0.57606 | 5.022499 | -14.1572 | 2.53E-12 | 2.64E-11 | 1357.083 | 1255.681 | 1332.333 | 10244.22 | 10216.98 | 10231.58 | 7133.308 | 7370.979 | 6899.53 | 2809.94 | 2630.721 | 2540.546 | 132.658 | 101.4047 | 93.94495 |
| Adam39 | 546055 | 2.865647 | -3.03783 | 3.25328 | 0.003746 | 0.009786 | -0.70613 | -3.03783 | -0.92984 | 0.362852 | 0.460676 | 0 | 5.32821 | 2.603909 | 19.48064 | 9.830767 | 12.17401 | 11.37009 | 5.361687 | 9.193245 | 6.153716 | 6.100223 | 5.97073 | 16.68654 | 27.51289 | 4.859222 |
| Gm11213 | 670833 | 2.866745 | -3.42811 | 3.013295 | 0.006538 | 0.016311 | 0.301108 | -3.42811 | 0.394076 | 0.697435 | 0.779151 | 0 | 10.65642 | 0 | 7.792255 | 9.830767 | 8.116006 | 11.37009 | 9.382952 | 12.64071 | 3.076858 | 15.25056 | 9.702436 | 4.171635 | 5.502578 | 4.049351 |
| Vmn1r205 | 171251 | 2.868368 | -5.25455 | 2.713368 | 0.012902 | 0.030188 | -1.85683 | -5.25455 | -1.81722 | 0.083252 | 0.133407 | 1.483961 | 0 | 0 | 2.597418 | 8.426372 | 5.410671 | 0 | 2.680843 | 1.149156 | 0 | 0 | 0.746341 | 6.674615 | 6.288661 | 0 |
| Cfap97d2 | 403185 | 2.869223 | -4.71291 | 2.763323 | 0.011538 | 0.02728 | 0.255189 | -4.71291 | 0.290652 | 0.774123 | 0.846895 | 0.741981 | 0 | 0 | 1.298709 | 2.808791 | 14.87935 | 2.842522 | 1.340422 | 24.13227 | 0 | 0 | 0 | 10.84625 | 7.074744 | 12.14805 |
| Prss23 | 76453 | 2.870968 | -1.91657 | 3.647402 | 0.001474 | 0.004124 | 3.588331 | -1.91657 | 7.908537 | 8.83E-08 | 4.37E-07 | 4.451884 | 0 | 4.339848 | 11.68838 | 19.66153 | 20.29002 | 198.9765 | 201.0633 | 243.621 | 7.692145 | 3.050111 | 1.492683 | 44.21933 | 32.22939 | 24.29611 |
| Setdb2 | 239122 | 2.876431 | 5.145682 | 50.66903 | 9.91E-24 | 1.09E-21 | -2.95613 | 5.145682 | -46.3665 | 6.48E-23 | 1.19E-20 | 1549.998 | 1587.807 | 1461.661 | 12109.16 | 12169.08 | 11783.09 | 1715.462 | 1546.847 | 1572.045 | 2368.411 | 2341.723 | 2158.419 | 705.0062 | 531.3919 | 650.3258 |
| Prr16 | 71373 | 2.876956 | -3.97283 | 2.916226 | 0.008165 | 0.01996 | -0.05911 | -3.97283 | -0.07155 | 0.943628 | 0.953379 | 0 | 0 | 7.811726 | 10.38967 | 11.23516 | 4.058003 | 12.79135 | 2.680843 | 12.64071 | 0.769214 | 0 | 1.492683 | 13.34923 | 6.288661 | 12.95792 |
| Alppl2 | 11650 | 2.880414 | -2.67425 | 3.073885 | 0.005686 | 0.014361 | 1.908768 | -2.67425 | 2.990764 | 0.006886 | 0.015526 | 1.483961 | 0 | 1.735939 | 10.38967 | 12.63956 | 5.410671 | 22.74017 | 49.5956 | 42.51876 | 13.84586 | 3.050111 | 2.985365 | 33.37308 | 47.16496 | 17.00728 |
| Spats2l | 67198 | 2.880499 | 0.327098 | 9.203219 | 7.03E-09 | 3.87E-08 | 0.933502 | 0.327098 | 5.521421 | 1.67E-05 | 6.00E-05 | 24.48536 | 12.43249 | 28.643 | 181.8193 | 162.9099 | 155.5568 | 355.3152 | 367.2756 | 286.1397 | 47.6913 | 62.52728 | 55.22925 | 35.87606 | 33.01547 | 39.68364 |
| Fmr1nb | 207854 | 2.88186 | -4.54257 | 2.491908 | 0.020998 | 0.046529 | 1.448637 | -4.54257 | 1.580512 | 0.128697 | 0.193254 | 0 | 0 | 0 | 0 | 8.426372 | 5.410671 | 9.948826 | 8.04253 | 8.044089 | 3.846072 | 0 | 4.478048 | 5.840288 | 6.288661 | 1.619741 |
| Olfr1337 | 258306 | 2.883209 | -4.81463 | 2.6092 | 0.016252 | 0.03708 | 1.1069 | -4.81463 | 1.229311 | 0.232336 | 0.317933 | 0 | 0 | 0 | 0 | 5.617581 | 8.116006 | 8.527565 | 2.680843 | 11.49156 | 0 | 0 | 2.985365 | 7.508942 | 5.502578 | 3.239481 |
| Olfr352 | 258942 | 2.883209 | -5.24143 | 3.001438 | 0.006719 | 0.016721 | 0.380451 | -5.24143 | 0.46407 | 0.647296 | 0.734735 | 0 | 0 | 0 | 0 | 5.617581 | 8.116006 | 2.842522 | 5.361687 | 3.447467 | 0 | 0 | 0 | 5.005962 | 4.716496 | 2.429611 |
| Lce1l | 73730 | 2.884708 | -4.43588 | 2.908562 | 0.008309 | 0.02028 | -0.46886 | -4.43588 | -0.53764 | 0.596388 | 0.689325 | 1.483961 | 0 | 0 | 9.090964 | 1.404395 | 8.116006 | 4.263783 | 1.340422 | 6.894934 | 3.076858 | 0.762528 | 0 | 15.85221 | 5.502578 | 5.669092 |
| Gm9881 | 791288 | 2.884851 | -2.97083 | 3.420856 | 0.002525 | 0.006791 | 1.006811 | -2.97083 | 1.73037 | 0.097999 | 0.153239 | 2.967923 | 0 | 4.339848 | 18.18193 | 18.25714 | 9.468674 | 28.42522 | 40.21265 | 27.57973 | 0 | 1.525056 | 4.478048 | 28.36712 | 14.14949 | 12.95792 |
| Amotl2 | 56332 | 2.885665 | 1.148307 | 10.95695 | 3.17E-10 | 2.11E-09 | 2.763559 | 1.148307 | 22.01303 | 3.71E-16 | 8.90E-15 | 20.03348 | 24.86498 | 32.11488 | 190.9102 | 205.0417 | 202.9002 | 1348.777 | 1465.081 | 1412.312 | 35.38386 | 41.93903 | 55.22925 | 135.161 | 132.0619 | 119.0509 |
| Itln1 | 16429 | 2.88774 | -3.22162 | 3.958349 | 0.0007 | 0.00206 | -0.19733 | -3.22162 | -0.34161 | 0.735984 | 0.814008 | 1.483961 | 1.77607 | 1.735939 | 14.2858 | 18.25714 | 14.87935 | 12.79135 | 16.08506 | 13.78987 | 7.692145 | 0.762528 | 0 | 18.35519 | 11.00516 | 12.95792 |
| A430072P | 414110 | 2.889012 | -1.67277 | 5.255196 | 3.13E-05 | 0.000109 | -0.98219 | -1.67277 | -2.30869 | 0.031055 | 0.057579 | 4.451884 | 9.768385 | 8.679696 | 70.13029 | 58.9846 | 48.69604 | 21.31891 | 20.10633 | 58.60694 | 2.307643 | 11.43792 | 2.985365 | 35.04173 | 25.15464 | 50.21196 |
| Rapgef2 | 76089 | 2.890255 | 7.797779 | 88.34858 | 7.37E-29 | 7.75E-26 | -2.17648 | 7.797779 | -70.0059 | 1.04E-26 | 9.44E-24 | 11750.75 | 12270.87 | 10938.15 | 92369.39 | 94741.91 | 89469.5 | 22156.04 | 21100.92 | 20407.85 | 9653.641 | 10266.67 | 9704.675 | 1857.212 | 1849.652 | 1851.363 |
| D930028M | 434147 | 2.890302 | -2.35498 | 4.570956 | 0.00016 | 0.000511 | -0.91472 | -2.35498 | -1.76539 | 0.091803 | 0.145016 | 2.967923 | 2.664105 | 4.339848 | 28.5716 | 28.08791 | 28.40602 | 5.685043 | 22.78717 | 24.13227 | 9.230573 | 5.337695 | 8.956095 | 8.343269 | 24.36856 | 20.24676 |
| Ctag2 | 70062 | 2.890831 | -5.53909 | 2.961035 | 0.007371 | 0.018183 | -0.83791 | -5.53909 | -0.93241 | 0.361551 | 0.459379 | 0 | 0 | 0 | 1.298709 | 8.426372 | 1.352668 | 2.842522 | 0 | 2.298311 | 0 | 0 | 0.746341 | 5.840288 | 0.786083 | 1.619741 |
| Gm7995 | 666233 | 2.890831 | -5.90521 | 3.311257 | 0.00327 | 0.00863 | -1.59196 | -5.90521 | -1.89176 | 0.07217 | 0.118261 | 0 | 0 | 0 | 1.298709 | 8.426372 | 1.352668 | 0 | 0 | 2.298311 | 0 | 0 | 0 | 1.668654 | 0.786083 | 1.619741 |
| Npr1 | 18160 | 2.891982 | 0.223148 | 9.120959 | 8.20E-09 | 4.47E-08 | -0.01575 | 0.223148 | -0.08028 | 0.936766 | 0.947793 | 17.06556 | 23.08891 | 11.2836 | 138.9619 | 148.8659 | 113.6241 | 130.756 | 143.4251 | 135.6004 | 33.07622 | 42.70156 | 28.36097 | 163.5281 | 144.6392 | 154.6852 |
| Ly6i | 57248 | 2.894854 | 0.362228 | 10.71369 | 4.77E-10 | 3.10E-09 | 1.296341 | 0.362228 | 9.270341 | 6.20E-09 | 3.63E-08 | 29.67923 | 41.73764 | 20.83127 | 224.6767 | 205.0417 | 288.1182 | 638.1461 | 625.977 | 561.9371 | 26.15329 | 24.40089 | 19.40487 | 25.02981 | 25.15464 | 27.53559 |
| Kcnk9 | 223604 | 2.896147 | -4.95983 | 2.709684 | 0.013009 | 0.03041 | 1.430009 | -4.95983 | 1.685786 | 0.106403 | 0.164618 | 0 | 0 | 0 | 2.597418 | 0 | 14.87935 | 4.263783 | 17.42548 | 8.044089 | 0 | 0 | 0 | 8.343269 | 6.288661 | 2.429611 |
| Olfr994 | 258425 | 2.896147 | -5.36957 | 2.69943 | 0.013309 | 0.031027 | -0.54731 | -5.36957 | -0.56387 | 0.578722 | 0.672423 | 0 | 0 | 0 | 2.597418 | 0 | 14.87935 | 2.842522 | 0 | 4.596622 | 0 | 0 | 0 | 5.005962 | 7.074744 | 3.239481 |
| Gm5767 | 436336 | 2.897215 | -2.52229 | 3.44208 | 0.002402 | 0.00649 | 0.239051 | -2.52229 | 0.382544 | 0.705841 | 0.78679 | 2.967923 | 7.10428 | 0 | 15.58451 | 29.4923 | 10.82134 | 17.05513 | 13.40422 | 36.77298 | 13.07665 | 5.337695 | 13.43414 | 23.36115 | 10.21907 | 11.33818 |
| Cyp2b9 | 13094 | 2.89916 | -3.29555 | 2.93904 | 0.007751 | 0.019038 | 0.938422 | -3.29555 | 1.30886 | 0.2045 | 0.285835 | 1.483961 | 0 | 3.471878 | 24.67547 | 9.830767 | 5.410671 | 12.79135 | 18.7659 | 43.66791 | 4.615287 | 2.287584 | 0 | 20.02385 | 13.3634 | 6.478962 |
| Nkx2-3 | 18089 | 2.90039 | -2.5624 | 3.439409 | 0.002417 | 0.006527 | 1.454471 | -2.5624 | 2.657273 | 0.014615 | 0.030015 | 8.903769 | 1.77607 | 0 | 18.18193 | 19.66153 | 12.17401 | 52.58665 | 45.57434 | 45.96622 | 1.538429 | 0 | 3.731706 | 51.72827 | 47.95104 | 30.77507 |
| Cxcl1 | 14825 | 2.901089 | 2.506917 | 12.64763 | 2.20E-11 | 1.73E-10 | 4.935944 | 2.506917 | 48.30162 | 2.73E-23 | 5.82E-21 | 67.52025 | 63.93852 | 69.43757 | 544.1591 | 497.1559 | 570.8258 | 16971.28 | 16995.21 | 17232.74 | 156.1505 | 137.255 | 231.3658 | 59.23721 | 55.02578 | 23.48624 |
| Gm5265 | 383613 | 2.904155 | -5.58003 | 2.71999 | 0.012713 | 0.02979 | -0.17388 | -5.58003 | -0.18463 | 0.855261 | 0.917108 | 0 | 0 | 0 | 7.792255 | 5.617581 | 0 | 4.263783 | 1.340422 | 2.298311 | 1.538429 | 0.762528 | 0 | 0 | 0 | 2.429611 |
| Cdr1 | 631990 | 2.905734 | -3.51699 | 3.199204 | 0.00425 | 0.010983 | 0.557789 | -3.51699 | 0.791253 | 0.437502 | 0.536401 | 0 | 1.77607 | 1.735939 | 6.493546 | 9.830767 | 10.82134 | 25.5827 | 5.361687 | 18.38649 | 0 | 3.050111 | 1.492683 | 13.34923 | 14.14949 | 25.10598 |
| Flt4 | 14257 | 2.905777 | 1.387691 | 10.82976 | 3.92E-10 | 2.59E-09 | 2.253893 | 1.387691 | 17.50899 | 3.84E-14 | 5.86E-13 | 43.03488 | 29.30515 | 39.9266 | 322.0799 | 334.2461 | 252.9489 | 1559.123 | 1524.06 | 1412.312 | 36.15308 | 78.54037 | 47.76584 | 114.3028 | 107.6933 | 98.80417 |
| Fam186a | 72277 | 2.908186 | -4.80304 | 3.283519 | 0.00349 | 0.009161 | 0.894638 | -4.80304 | 1.224054 | 0.234273 | 0.3202 | 0 | 0 | 0 | 7.792255 | 1.404395 | 1.352668 | 7.106304 | 2.680843 | 9.193245 | 0.769214 | 0 | 0 | 9.177596 | 7.074744 | 11.33818 |
| Dcbld2 | 73379 | 2.908472 | 5.219145 | 43.56356 | 2.42E-22 | 1.73E-20 | 0.946389 | 5.219145 | 23.38055 | 1.08E-16 | 2.96E-15 | 616.586 | 612.7441 | 698.7155 | 5392.24 | 5038.97 | 4983.228 | 10382.31 | 10340.01 | 10132.1 | 2015.342 | 1896.407 | 1992.731 | 854.3508 | 801.8043 | 804.2012 |
| Mefv | 54483 | 2.911858 | 4.912934 | 33.58307 | 5.78E-20 | 2.38E-18 | -0.22767 | 4.912934 | -3.77944 | 0.001075 | 0.002923 | 772.402 | 720.1964 | 762.9453 | 6148.089 | 6109.119 | 5896.279 | 5550.024 | 5818.771 | 4845.989 | 1346.895 | 1303.923 | 1235.195 | 698.3316 | 537.6805 | 563.6697 |
| Zfp735 | 76390 | 2.913532 | -3.40992 | 3.249363 | 0.00378 | 0.009869 | 0.893827 | -3.40992 | 1.33948 | 0.194504 | 0.274052 | 1.483961 | 0 | 1.735939 | 12.98709 | 2.808791 | 18.93735 | 9.948826 | 26.80843 | 21.83396 | 0.769214 | 1.525056 | 0.746341 | 10.01192 | 25.94073 | 23.48624 |
| Hoxd4 | 15436 | 2.915147 | -3.35904 | 2.833556 | 0.009851 | 0.023695 | 1.040571 | -3.35904 | 1.404307 | 0.174613 | 0.250153 | 0 | 0.888035 | 6.075787 | 14.2858 | 2.808791 | 28.40602 | 12.79135 | 37.53181 | 27.57973 | 0.769214 | 0.762528 | 0 | 39.21337 | 13.3634 | 15.38754 |
| Gm10309 | 100038711 | 2.915317 | -2.11582 | 3.604587 | 0.001633 | 0.004535 | 0.245495 | -2.11582 | 0.426117 | 0.674289 | 0.758727 | 0.741981 | 10.65642 | 1.735939 | 40.25998 | 9.830767 | 28.40602 | 11.37009 | 45.57434 | 39.07129 | 4.615287 | 7.625279 | 5.97073 | 39.21337 | 45.59279 | 35.63429 |
| 4933412E | 71086 | 2.915716 | 2.709351 | 20.68063 | 1.33E-15 | 2.15E-14 | 0.54189 | 2.709351 | 6.736295 | 1.06E-06 | 4.49E-06 | 130.5886 | 172.2788 | 150.1587 | 1175.332 | 1199.354 | 1232.28 | 1934.336 | 1844.42 | 1698.452 | 488.4512 | 385.0766 | 492.5852 | 62.57452 | 66.81702 | 53.45144 |
| Lrrc36 | 270091 | 2.915964 | -2.06454 | 3.940913 | 0.00073 | 0.002142 | 0.581006 | -2.06454 | 1.153881 | 0.261307 | 0.350895 | 5.193865 | 7.992315 | 0 | 23.37676 | 21.06593 | 20.29002 | 29.84648 | 44.23392 | 29.87805 | 3.846072 | 7.625279 | 9.702436 | 45.88798 | 35.37372 | 39.68364 |
| A930015D | 77810 | 2.91856 | 2.65079 | 17.66214 | 3.22E-14 | 4.04E-13 | 0.889824 | 2.65079 | 10.01701 | 1.60E-09 | 1.03E-08 | 126.8787 | 115.4445 | 83.32508 | 853.2519 | 900.2174 | 865.7074 | 1718.304 | 1847.101 | 1512.289 | 222.303 | 240.9588 | 226.1414 | 220.2623 | 198.8789 | 195.1787 |
| Nupr1 | 56312 | 2.921978 | 1.904364 | 16.8041 | 8.71E-14 | 1.02E-12 | -0.84168 | 1.904364 | -7.00426 | 5.91E-07 | 2.60E-06 | 105.3613 | 132.3172 | 113.704 | 961.0448 | 908.6437 | 950.9254 | 577.0319 | 644.7429 | 449.3198 | 256.9176 | 204.3575 | 216.439 | 35.04173 | 62.10053 | 30.77507 |
| Crp | 12944 | 2.925043 | -3.55903 | 3.389948 | 0.002717 | 0.007262 | 0.569529 | -3.55903 | 0.873673 | 0.392021 | 0.49088 | 0 | 3.55214 | 1.735939 | 7.792255 | 14.04395 | 12.17401 | 21.31891 | 10.72337 | 21.83396 | 0 | 4.575167 | 1.492683 | 6.674615 | 4.716496 | 9.718443 |
| Tnks1bp1 | 228140 | 2.925632 | 4.534012 | 30.74983 | 3.66E-19 | 1.27E-17 | 0.570242 | 4.534012 | 9.761258 | 2.52E-09 | 1.57E-08 | 391.0239 | 498.1876 | 406.2098 | 3261.059 | 3700.582 | 3460.124 | 5505.965 | 5440.772 | 5118.339 | 694.6007 | 691.6128 | 828.4388 | 849.3448 | 851.3275 | 899.7659 |
| Gm11190 | 791423 | 2.927752 | -2.76498 | 3.081353 | 0.005589 | 0.014133 | 0.801453 | -2.76498 | 1.212254 | 0.238664 | 0.325221 | 2.967923 | 0 | 5.207818 | 7.792255 | 14.04395 | 37.8747 | 17.05513 | 36.19139 | 39.07129 | 1.538429 | 7.625279 | 0 | 23.36115 | 26.72681 | 37.25403 |
| Vegfc | 22341 | 2.929611 | 4.395597 | 38.77822 | 2.82E-21 | 1.60E-19 | -1.86862 | 4.395597 | -28.0176 | 2.55E-18 | 1.02E-16 | 591.3587 | 515.9483 | 533.8013 | 4553.274 | 4485.638 | 4352.885 | 1415.576 | 1251.954 | 1169.84 | 558.4497 | 623.7478 | 606.7754 | 1659.476 | 1660.206 | 1779.285 |
| 4930447A | 74892 | 2.930214 | -4.19664 | 3.342139 | 0.003041 | 0.008063 | 1.410705 | -4.19664 | 2.031576 | 0.054832 | 0.093798 | 0 | 0 | 0 | 2.597418 | 2.808791 | 2.705335 | 5.685043 | 12.0638 | 9.193245 | 2.307643 | 6.100223 | 1.492683 | 4.171635 | 11.00516 | 12.95792 |
| Ifi203 | 15950 | 2.931213 | 8.158002 | 79.72447 | 6.56E-28 | 3.21E-25 | 0.124071 | 8.158002 | 5.474417 | 1.87E-05 | 6.66E-05 | 6371.389 | 6443.582 | 5780.678 | 49814.59 | 51789.88 | 49893.15 | 56766.58 | 57996.03 | 56592.47 | 10812.85 | 11307.53 | 10509.23 | 7512.28 | 7487.437 | 7418.412 |
| 4930594C | 77633 | 2.931616 | 2.564606 | 15.90537 | 2.59E-13 | 2.78E-12 | -0.46112 | 2.564606 | -3.84642 | 0.000916 | 0.002527 | 120.2009 | 150.0779 | 201.3689 | 1359.748 | 1162.839 | 1199.816 | 1074.473 | 1007.997 | 780.2766 | 157.689 | 144.1178 | 96.27802 | 210.2504 | 253.9047 | 242.1512 |
| Sell | 20343 | 2.932914 | 4.638337 | 38.1535 | 3.96E-21 | 2.19E-19 | -2.14613 | 4.638337 | -29.5808 | 8.22E-19 | 3.84E-17 | 1065.484 | 1143.789 | 897.4806 | 8381.869 | 8110.383 | 8743.644 | 1995.45 | 2072.292 | 1866.229 | 1073.823 | 1004.249 | 1040.4 | 423.0037 | 333.299 | 489.1616 |
| Stox1 | 216021 | 2.934558 | -2.53149 | 3.171803 | 0.004531 | 0.011633 | 0.819473 | -2.53149 | 1.255563 | 0.222847 | 0.307169 | 1.483961 | 7.992315 | 0 | 18.18193 | 18.25714 | 9.468674 | 21.31891 | 24.12759 | 37.92213 | 8.461359 | 6.100223 | 2.985365 | 29.20144 | 47.16496 | 11.33818 |
| Gm10724 | 100038688 | 2.934579 | -4.0374 | 3.59906 | 0.001654 | 0.004587 | 0.934357 | -4.0374 | 1.508504 | 0.146087 | 0.215215 | 0 | 1.77607 | 0.86797 | 10.38967 | 14.04395 | 2.705335 | 11.37009 | 12.0638 | 25.28142 | 0 | 0 | 0 | 14.18356 | 13.3634 | 9.718443 |
| BC052688 | 432812 | 2.936905 | -4.55226 | 2.921153 | 0.008074 | 0.019765 | 0.142015 | -4.55226 | 0.169342 | 0.867123 | 0.918721 | 1.483961 | 0 | 0 | 3.896127 | 2.808791 | 12.17401 | 12.79135 | 5.361687 | 3.447467 | 0 | 0 | 0.746341 | 14.18356 | 10.21907 | 1.619741 |
| Lrrc8c | 100604 | 2.937284 | 5.881222 | 62.38157 | 1.21E-25 | 2.58E-23 | -1.35641 | 5.881222 | -34.9429 | 2.51E-20 | 1.83E-18 | 1832.692 | 1800.047 | 1788.017 | 14961.13 | 15313.53 | 14120.5 | 5997.721 | 6152.536 | 5852.649 | 2131.493 | 1965.797 | 1941.234 | 1747.915 | 1638.196 | 1662.664 |
| Gm10824 | 100038582 | 2.939876 | -3.4925 | 3.638147 | 0.001507 | 0.004209 | 0.709131 | -3.4925 | 1.159319 | 0.259132 | 0.348455 | 1.483961 | 1.77607 | 0 | 10.38967 | 5.617581 | 13.52668 | 8.527565 | 21.44675 | 21.83396 | 0.769214 | 2.287584 | 0.746341 | 12.5149 | 22.01031 | 8.908573 |
| Cpvl | 71287 | 2.941501 | -2.10412 | 5.017347 | 5.50E-05 | 0.000186 | 1.261164 | -2.10412 | 3.592595 | 0.00168 | 0.004378 | 4.451884 | 1.77607 | 4.339848 | 24.67547 | 22.47032 | 48.69604 | 78.16935 | 62.99982 | 86.18667 | 1.538429 | 6.100223 | 3.731706 | 11.68058 | 17.29382 | 19.43689 |
| 9330179D | 77558 | 2.944738 | -0.67972 | 9.419195 | 4.70E-09 | 2.66E-08 | 1.124602 | -0.67972 | 6.781065 | 9.60E-07 | 4.09E-06 | 12.61367 | 13.32052 | 19.09533 | 125.9748 | 117.9692 | 125.7981 | 301.3073 | 262.7227 | 278.0957 | 16.92272 | 12.96297 | 8.956095 | 11.68058 | 10.21907 | 8.908573 |
| 2810030D | 78335 | 2.945374 | -4.38408 | 2.944518 | 0.007655 | 0.01883 | 0.864534 | -4.38408 | 1.048664 | 0.306063 | 0.400506 | 0 | 0 | 0 | 0 | 4.213186 | 12.17401 | 4.263783 | 4.021265 | 10.3424 | 0.769214 | 1.525056 | 0.746341 | 13.34923 | 16.50774 | 16.19741 |
| 5830477G | 76116 | 2.945785 | -0.42833 | 7.412068 | 2.47E-07 | 1.11E-06 | -0.59011 | -0.42833 | -2.22142 | 0.037263 | 0.0674 | 24.48536 | 22.20087 | 6.943757 | 146.7541 | 130.6088 | 124.4454 | 99.48826 | 84.44657 | 93.0816 | 59.99873 | 61.00223 | 31.34633 | 20.85817 | 12.57732 | 19.43689 |
| Gm3764 | 100042277 | 2.947196 | -3.28888 | 3.695328 | 0.001315 | 0.003711 | -0.03134 | -3.28888 | -0.04912 | 0.961281 | 0.968239 | 0 | 2.664105 | 3.471878 | 14.2858 | 11.23516 | 12.17401 | 12.79135 | 5.361687 | 26.43058 | 1.538429 | 1.525056 | 1.492683 | 15.01788 | 20.43815 | 12.14805 |
| Cflar | 12633 | 2.948289 | 8.257441 | 93.8737 | 2.03E-29 | 3.25E-26 | -1.13527 | 8.257441 | -47.8967 | 3.26E-23 | 6.84E-21 | 8598.815 | 8734.712 | 7858.597 | 70113.41 | 69485.26 | 67941.8 | 33864.38 | 32326.95 | 32061.44 | 15937.35 | 15594.46 | 15926.92 | 6768.894 | 6693.494 | 6977.842 |
| Zfp879 | 214779 | 2.949362 | -3.31422 | 2.788101 | 0.010914 | 0.025948 | 1.187986 | -3.31422 | 1.486435 | 0.151788 | 0.222554 | 2.225942 | 0 | 0 | 16.88322 | 4.213186 | 2.705335 | 4.263783 | 28.14886 | 24.13227 | 6.153716 | 7.625279 | 3.731706 | 27.53279 | 9.432991 | 15.38754 |
| Serpina1c | 20702 | 2.951835 | -4.41008 | 3.188503 | 0.004358 | 0.011238 | -2.19194 | -4.41008 | -2.4068 | 0.025219 | 0.048183 | 0.741981 | 0.888035 | 1.735939 | 9.090964 | 15.44835 | 10.82134 | 1.421261 | 0 | 12.64071 | 4.615287 | 0 | 0 | 7.508942 | 1.572165 | 4.049351 |
| Cemip2 | 83921 | 2.954293 | 6.710974 | 61.07058 | 1.89E-25 | 3.89E-23 | -0.61451 | 6.710974 | -17.7367 | 2.96E-14 | 4.64E-13 | 3318.138 | 3515.73 | 3070.876 | 27635.23 | 27594.96 | 26612.39 | 19410.16 | 18777.97 | 17524.62 | 4356.831 | 4190.091 | 4114.579 | 1417.521 | 1346.56 | 1338.716 |
| Phf11d | 219132 | 2.954414 | 6.075249 | 46.24558 | 6.85E-23 | 5.76E-21 | 0.064163 | 6.075249 | 1.50624 | 0.146664 | 0.215907 | 1626.422 | 1506.995 | 1321.918 | 12198.78 | 12649.39 | 12049.56 | 12983.22 | 13481.96 | 13514.07 | 2754.557 | 2909.806 | 2564.429 | 1529.321 | 1767.9 | 1515.267 |
| Vmn2r84 | 625068 | 2.954607 | -3.49597 | 2.745228 | 0.012016 | 0.028313 | 0.524731 | -3.49597 | 0.629306 | 0.535828 | 0.631143 | 1.483961 | 0 | 1.735939 | 9.090964 | 5.617581 | 16.23201 | 4.263783 | 12.0638 | 44.81707 | 0 | 7.625279 | 0.746341 | 12.5149 | 27.51289 | 10.52831 |
| Vmn2r86 | 625109 | 2.955972 | -4.37886 | 3.561794 | 0.001808 | 0.004988 | -0.32134 | -4.37886 | -0.45034 | 0.657004 | 0.743347 | 0.741981 | 0 | 0.86797 | 2.597418 | 8.426372 | 12.17401 | 4.263783 | 2.680843 | 11.49156 | 0 | 0 | 0.746341 | 7.508942 | 13.3634 | 8.098703 |
| Lacc1 | 210808 | 2.956165 | 8.161506 | 114.6564 | 2.86E-31 | 1.25E-27 | -1.14582 | 8.161506 | -59.0058 | 3.93E-25 | 1.40E-22 | 8943.836 | 8936.296 | 8587.691 | 74313.44 | 73442.85 | 71622.4 | 34496.84 | 34738.37 | 33777.13 | 12469.74 | 12663.3 | 12440.76 | 5305.485 | 5262.823 | 5312.749 |
| 4930433N | 114673 | 2.956483 | -3.91069 | 3.485972 | 0.002165 | 0.005894 | 1.619874 | -3.91069 | 2.549611 | 0.018521 | 0.03692 | 0 | 0.888035 | 0 | 3.896127 | 2.808791 | 6.763339 | 11.37009 | 10.72337 | 25.28142 | 3.076858 | 2.287584 | 0.746341 | 13.34923 | 9.432991 | 7.288832 |
| Crisp2 | 22024 | 2.95713 | -3.24536 | 3.471678 | 0.002239 | 0.006078 | 0.107155 | -3.24536 | 0.157079 | 0.876659 | 0.918721 | 2.967923 | 0 | 0.86797 | 3.896127 | 22.47032 | 10.82134 | 8.527565 | 8.04253 | 18.38649 | 6.92293 | 3.050111 | 2.985365 | 12.5149 | 22.7964 | 10.52831 |
| Ankrd65 | 242805 | 2.957819 | -4.23303 | 2.58439 | 0.017163 | 0.038894 | 0.966272 | -4.23303 | 1.033531 | 0.312925 | 0.407821 | 0 | 0 | 0 | 3.896127 | 12.63956 | 0 | 5.685043 | 5.361687 | 8.044089 | 1.538429 | 0 | 2.985365 | 13.34923 | 28.29897 | 35.63429 |
| Olfr1325 | 258393 | 2.958813 | -5.14626 | 2.752436 | 0.011823 | 0.027892 | 0.533996 | -5.14626 | 0.590885 | 0.560803 | 0.655396 | 0 | 0 | 0 | 1.298709 | 9.830767 | 1.352668 | 2.842522 | 5.361687 | 5.745778 | 3.076858 | 0.762528 | 0 | 1.668654 | 0 | 4.859222 |
| Vmn2r10 | 22307 | 2.959551 | -4.24237 | 2.913297 | 0.00822 | 0.020079 | 1.107825 | -4.24237 | 1.458999 | 0.159127 | 0.231508 | 2.967923 | 0 | 0 | 10.38967 | 9.830767 | 2.705335 | 15.63387 | 22.78717 | 11.49156 | 3.846072 | 0 | 0 | 6.674615 | 4.716496 | 0.80987 |
| Kcna3 | 16491 | 2.960137 | 4.27459 | 40.79822 | 9.66E-22 | 6.06E-20 | -0.49216 | 4.27459 | -9.47653 | 4.23E-09 | 2.55E-08 | 932.6698 | 813.44 | 841.9305 | 7294.849 | 6866.088 | 7400.445 | 5771.74 | 5116.39 | 5089.61 | 279.2248 | 224.1832 | 264.9511 | 222.7653 | 244.4717 | 273.7362 |
| 9330175E | 320377 | 2.960469 | 2.740423 | 21.25669 | 7.58E-16 | 1.30E-14 | 0.131065 | 2.740423 | 1.588778 | 0.126815 | 0.1908 | 149.1381 | 165.1745 | 144.083 | 1223.384 | 1436.696 | 1164.647 | 1492.324 | 1404.762 | 1431.848 | 301.5321 | 293.5732 | 220.1707 | 137.6639 | 129.7036 | 135.2483 |
| Gm4559 | 100043627 | 2.960695 | -4.99319 | 3.829861 | 0.000953 | 0.002748 | 0.984321 | -4.99319 | 1.562022 | 0.132989 | 0.198713 | 0 | 0 | 0 | 1.298709 | 5.617581 | 2.705335 | 5.685043 | 6.702109 | 6.894934 | 0 | 0 | 0 | 5.840288 | 6.288661 | 4.859222 |
| Gm19966 | 100503921 | 2.961612 | -3.30099 | 3.596134 | 0.001666 | 0.004617 | 0.926329 | -3.30099 | 1.472744 | 0.155415 | 0.227046 | 2.967923 | 0 | 0 | 6.493546 | 5.617581 | 8.116006 | 11.37009 | 12.0638 | 18.38649 | 3.846072 | 4.575167 | 2.985365 | 15.85221 | 18.86598 | 33.20468 |
| Olfr159 | 29849 | 2.964937 | -5.46139 | 3.156515 | 0.004695 | 0.012031 | -1.44328 | -5.46139 | -1.62128 | 0.119639 | 0.181623 | 0 | 0 | 0 | 2.597418 | 5.617581 | 1.352668 | 5.685043 | 0 | 0 | 0 | 0.762528 | 0.746341 | 1.668654 | 3.14433 | 3.239481 |
| Insrr | 23920 | 2.966246 | 0.457943 | 6.348922 | 2.51E-06 | 9.91E-06 | 3.671234 | 0.457943 | 16.97977 | 7.08E-14 | 1.01E-12 | 8.903769 | 16.87266 | 13.01954 | 98.7019 | 101.1165 | 117.6821 | 1475.269 | 1544.166 | 1235.342 | 12.30743 | 19.82572 | 41.79511 | 81.76404 | 94.32991 | 84.22651 |
| Lta | 16992 | 2.966258 | -1.17594 | 6.427812 | 2.10E-06 | 8.37E-06 | -0.55172 | -1.17594 | -1.75511 | 0.093587 | 0.147426 | 5.935846 | 15.98463 | 20.83127 | 137.6632 | 87.07251 | 93.33407 | 51.16539 | 81.76573 | 90.78329 | 11.53822 | 3.050111 | 8.956095 | 27.53279 | 17.29382 | 11.33818 |
| Rem1 | 19700 | 2.96645 | -2.63556 | 3.819266 | 0.000977 | 0.002815 | 0.687383 | -2.63556 | 1.234443 | 0.230457 | 0.315789 | 0.741981 | 0.888035 | 3.471878 | 11.68838 | 18.25714 | 13.52668 | 38.37404 | 9.382952 | 36.77298 | 6.153716 | 1.525056 | 4.478048 | 31.70442 | 30.65722 | 32.39481 |
| Olfr187 | 258319 | 2.968985 | -5.1774 | 2.740546 | 0.012142 | 0.028588 | 0.918061 | -5.1774 | 1.037032 | 0.311328 | 0.406203 | 0 | 0 | 0 | 2.597418 | 1.404395 | 5.410671 | 12.79135 | 4.021265 | 4.596622 | 0.769214 | 0 | 0 | 10.01192 | 2.358248 | 0 |
| Malt1 | 240354 | 2.971891 | 5.865439 | 63.90021 | 7.23E-26 | 1.68E-23 | -0.48935 | 5.865439 | -14.6854 | 1.24E-12 | 1.37E-11 | 1601.936 | 1585.142 | 1501.587 | 12911.77 | 13184.46 | 13220.97 | 9796.751 | 9853.44 | 9448.357 | 2165.339 | 2164.054 | 2329.331 | 1132.182 | 1221.572 | 1145.966 |
| Dipk1c | 240479 | 2.972484 | -3.94305 | 2.833766 | 0.009846 | 0.023688 | 1.038552 | -3.94305 | 1.313594 | 0.202929 | 0.284092 | 0.741981 | 0.888035 | 0 | 9.090964 | 1.404395 | 17.58468 | 15.63387 | 24.12759 | 9.193245 | 0 | 4.575167 | 0 | 11.68058 | 15.72165 | 4.859222 |
| Mrgprb3 | 404238 | 2.974208 | -4.49219 | 3.447552 | 0.002371 | 0.006413 | 0.39071 | -4.49219 | 0.555633 | 0.58424 | 0.677906 | 1.483961 | 0 | 0 | 7.792255 | 7.021976 | 2.705335 | 5.685043 | 13.40422 | 5.745778 | 0 | 1.525056 | 0 | 3.337308 | 5.502578 | 8.098703 |
| 1600014C | 72244 | 2.978251 | 5.685747 | 43.22094 | 2.86E-22 | 2.01E-20 | 0.746705 | 5.685747 | 17.62525 | 3.36E-14 | 5.18E-13 | 992.0283 | 917.3401 | 1046.771 | 8429.921 | 8434.798 | 8007.793 | 15304.14 | 14511.41 | 13683 | 1494.584 | 1409.151 | 1376.253 | 1588.558 | 1683.789 | 1635.128 |
| Unc93a2 | 667055 | 2.97901 | -1.32577 | 5.572183 | 1.49E-05 | 5.37E-05 | -0.04367 | -1.32577 | -0.12951 | 0.898165 | 0.921248 | 7.419807 | 7.10428 | 8.679696 | 72.72771 | 95.49888 | 44.63804 | 48.32287 | 117.9571 | 56.30862 | 10.769 | 15.25056 | 14.92683 | 16.68654 | 20.43815 | 8.098703 |
| Olfr429 | 258717 | 2.981082 | -4.55261 | 3.291079 | 0.003428 | 0.009011 | 0.826005 | -4.55261 | 1.154579 | 0.261027 | 0.350695 | 0 | 0 | 1.735939 | 10.38967 | 1.404395 | 8.116006 | 7.106304 | 8.04253 | 17.23733 | 0 | 0 | 0 | 10.01192 | 3.14433 | 6.478962 |
| Cobll1 | 319876 | 2.981239 | 3.819026 | 20.37133 | 1.80E-15 | 2.85E-14 | -1.07202 | 3.819026 | -10.2242 | 1.11E-09 | 7.32E-09 | 186.2372 | 188.2634 | 142.347 | 1302.605 | 1667.017 | 1409.48 | 730.5281 | 806.9339 | 633.1847 | 1039.978 | 977.5607 | 1000.097 | 2369.488 | 2268.634 | 2471.724 |
| AA414992 | 102402 | 2.983003 | -5.10013 | 3.628086 | 0.001544 | 0.004305 | -0.09612 | -5.10013 | -0.13635 | 0.892824 | 0.918744 | 0 | 0.888035 | 0 | 5.194837 | 2.808791 | 5.410671 | 5.685043 | 1.340422 | 8.044089 | 0 | 0 | 0 | 1.668654 | 3.930413 | 2.429611 |
| Cxcl3 | 330122 | 2.983505 | 0.859703 | 8.67803 | 1.91E-08 | 9.89E-08 | 4.835984 | 0.859703 | 31.86437 | 1.74E-19 | 9.81E-18 | 28.19527 | 18.64873 | 9.547666 | 149.3516 | 162.9099 | 142.0301 | 4552.299 | 4535.987 | 4417.354 | 52.30658 | 49.56431 | 32.09267 | 22.52683 | 31.4433 | 30.77507 |
| Med12l | 329650 | 2.983683 | 2.510772 | 13.84679 | 3.88E-12 | 3.42E-11 | 0.627314 | 2.510772 | 5.373176 | 2.37E-05 | 8.34E-05 | 78.64996 | 106.5642 | 69.43757 | 668.8352 | 630.5735 | 829.1853 | 1219.442 | 1209.06 | 999.7654 | 136.9202 | 154.7932 | 104.4878 | 479.738 | 556.5465 | 528.0354 |
| Vmn1r22 | 171196 | 2.983785 | -4.75347 | 2.682652 | 0.013815 | 0.032087 | -1.16751 | -4.75347 | -1.14422 | 0.265202 | 0.355183 | 0 | 0.888035 | 0.86797 | 2.597418 | 12.63956 | 8.116006 | 1.421261 | 0 | 21.83396 | 3.076858 | 0 | 0 | 2.502981 | 6.288661 | 1.619741 |
| Gm12159 | 100504255 | 2.98415 | -4.36601 | 3.175436 | 0.004492 | 0.011549 | 0.267563 | -4.36601 | 0.348431 | 0.73093 | 0.809746 | 0 | 0.888035 | 0.86797 | 11.68838 | 14.04395 | 1.352668 | 5.685043 | 9.382952 | 10.3424 | 3.076858 | 0 | 0 | 5.005962 | 3.930413 | 4.049351 |
| Olfr25 | 18323 | 2.985053 | -5.03896 | 2.819654 | 0.010165 | 0.024357 | 0.599897 | -5.03896 | 0.696496 | 0.493635 | 0.591737 | 0 | 0.888035 | 0 | 5.194837 | 1.404395 | 9.468674 | 14.21261 | 2.680843 | 9.193245 | 0 | 0 | 0 | 5.840288 | 3.930413 | 0 |
| Fgl2 | 14190 | 2.988307 | 7.852306 | 57.71854 | 3.03E-17 | 6.86E-16 | -4.09476 | 7.852306 | -54.4887 | 6.46E-17 | 1.87E-15 | 11913.98 | 12901.37 | 10090.15 | 97647.34 | 101425.4 | 95492.93 | 5940.87 | 5876.409 | 6048.006 | 29566.3 | 29498.39 | 28596.81 | 2494.637 | 2498.957 | 2638.557 |
| Nr4a3 | 18124 | 2.99023 | 2.721409 | 21.65791 | 5.17E-16 | 9.11E-15 | 0.579697 | 2.721409 | 7.569675 | 1.78E-07 | 8.40E-07 | 148.3961 | 142.9736 | 124.9876 | 1200.007 | 1092.62 | 1253.923 | 1894.541 | 1735.846 | 1867.378 | 216.1493 | 227.9958 | 168.6731 | 139.3326 | 159.5748 | 170.0728 |
| C4b | 12268 | 2.991981 | 2.1134 | 11.59763 | 1.11E-10 | 7.90E-10 | 2.639564 | 2.1134 | 22.16108 | 3.23E-16 | 7.92E-15 | 47.48677 | 40.84961 | 42.53051 | 335.067 | 356.7164 | 440.9697 | 2602.329 | 2553.503 | 2182.246 | 99.22866 | 88.45323 | 84.33656 | 347.08 | 300.2836 | 302.8915 |
| Dnm3os | 474332 | 2.992567 | 0.229181 | 7.44401 | 2.31E-07 | 1.04E-06 | 1.160554 | 0.229181 | 5.483177 | 1.83E-05 | 6.53E-05 | 12.61367 | 12.43249 | 10.41564 | 109.0916 | 92.69009 | 108.2134 | 214.6104 | 234.5738 | 268.9024 | 19.99958 | 22.87584 | 8.956095 | 338.7367 | 383.6083 | 302.0816 |
| Enpp5 | 83965 | 2.993749 | -0.28265 | 6.478265 | 1.88E-06 | 7.52E-06 | 1.20809 | -0.28265 | 5.005068 | 5.66E-05 | 0.000189 | 14.09763 | 5.32821 | 19.09533 | 132.4683 | 106.734 | 71.69139 | 208.9253 | 286.8503 | 237.8752 | 35.38386 | 44.22662 | 32.09267 | 28.36712 | 36.94588 | 17.00728 |
| Teddm3 | 66561 | 2.995332 | -4.7416 | 2.979969 | 0.007058 | 0.017492 | 0.836262 | -4.7416 | 1.050959 | 0.305032 | 0.399415 | 0.741981 | 0 | 0 | 5.194837 | 4.213186 | 4.058003 | 8.527565 | 18.7659 | 4.596622 | 1.538429 | 0 | 4.478048 | 0.834327 | 3.14433 | 0 |
| Gm10830 | 100038520 | 2.996729 | -3.18121 | 3.091385 | 0.00546 | 0.013838 | 2.622024 | -3.18121 | 3.934409 | 0.000741 | 0.002081 | 0 | 0 | 0.86797 | 2.597418 | 2.808791 | 10.82134 | 19.89765 | 37.53181 | 40.22045 | 6.153716 | 4.575167 | 1.492683 | 42.55067 | 20.43815 | 38.87377 |
| Myl2 | 17906 | 2.997908 | -3.38128 | 3.46734 | 0.002262 | 0.006139 | 1.639484 | -3.38128 | 2.77264 | 0.0113 | 0.024026 | 0 | 3.55214 | 0.86797 | 9.090964 | 18.25714 | 5.410671 | 45.48035 | 28.14886 | 28.72889 | 0 | 1.525056 | 2.985365 | 8.343269 | 9.432991 | 12.95792 |
| Rdh18-ps | 380674 | 3.000943 | -4.65751 | 3.153209 | 0.004731 | 0.012116 | -1.08672 | -4.65751 | -1.25046 | 0.224668 | 0.309242 | 0 | 1.77607 | 0 | 6.493546 | 4.213186 | 5.410671 | 2.842522 | 0 | 9.193245 | 0 | 0 | 1.492683 | 7.508942 | 10.21907 | 9.718443 |
| Nlrp4a | 243880 | 3.001232 | -3.62926 | 4.055722 | 0.000554 | 0.001654 | 0.586487 | -3.62926 | 1.048555 | 0.306112 | 0.40053 | 1.483961 | 1.77607 | 0 | 7.792255 | 9.830767 | 12.17401 | 24.16143 | 10.72337 | 14.93902 | 0 | 0 | 0.746341 | 28.36712 | 21.22423 | 19.43689 |
| Vmn2r28 | 665255 | 3.003367 | -3.95883 | 3.285399 | 0.003474 | 0.009124 | -0.03197 | -3.95883 | -0.04237 | 0.966601 | 0.972554 | 4.451884 | 0 | 0 | 5.194837 | 7.021976 | 13.52668 | 4.263783 | 6.702109 | 16.08818 | 0.769214 | 0.762528 | 1.492683 | 8.343269 | 16.50774 | 4.049351 |
| Ext1 | 14042 | 3.005117 | 5.507053 | 48.84236 | 2.16E-23 | 2.16E-21 | -0.37039 | 5.507053 | -8.58222 | 2.30E-08 | 1.24E-07 | 1147.102 | 1161.55 | 1169.155 | 9890.969 | 10207.14 | 9722.976 | 8223.415 | 8297.211 | 7508.583 | 853.0588 | 1033.225 | 921.7314 | 1722.885 | 1749.82 | 1698.298 |
| Gm3428 | 100041605 | 3.005356 | -4.95747 | 2.898797 | 0.008496 | 0.020705 | 0.250166 | -4.95747 | 0.282789 | 0.780063 | 0.852118 | 0 | 0 | 0 | 3.896127 | 14.04395 | 0 | 1.421261 | 5.361687 | 5.745778 | 0 | 0.762528 | 0 | 5.840288 | 6.288661 | 9.718443 |
| Pdyn | 18610 | 3.00577 | -4.27261 | 3.094993 | 0.005415 | 0.013733 | 1.508781 | -4.27261 | 2.108451 | 0.046975 | 0.082049 | 0 | 1.77607 | 0 | 2.597418 | 8.426372 | 6.763339 | 12.79135 | 24.12759 | 16.08818 | 3.076858 | 0 | 0 | 8.343269 | 1.572165 | 5.669092 |
| Epha4 | 13838 | 3.005858 | 3.386453 | 29.64737 | 7.84E-19 | 2.50E-17 | -1.8985 | 3.386453 | -21.777 | 4.63E-16 | 1.08E-14 | 362.8286 | 315.2524 | 295.1097 | 2845.472 | 2870.584 | 2662.05 | 878.3392 | 785.4871 | 695.2391 | 290.7631 | 264.5972 | 256.7414 | 564.8393 | 458.2862 | 476.2037 |
| Tnfrsf25 | 85030 | 3.006843 | -1.82034 | 5.082816 | 4.70E-05 | 0.00016 | 0.34838 | -1.82034 | 0.893938 | 0.381325 | 0.479569 | 3.709904 | 0.888035 | 7.811726 | 24.67547 | 23.87472 | 45.9907 | 41.21656 | 34.85097 | 43.66791 | 6.153716 | 6.100223 | 8.209754 | 52.5626 | 40.8763 | 38.87377 |
| G6pc2 | 14378 | 3.007723 | -3.61626 | 3.37888 | 0.002788 | 0.007443 | 0.805996 | -3.61626 | 1.17929 | 0.251262 | 0.339417 | 0 | 4.440175 | 0 | 3.896127 | 8.426372 | 10.82134 | 8.527565 | 20.10633 | 13.78987 | 6.153716 | 3.050111 | 1.492683 | 8.343269 | 5.502578 | 8.908573 |
| Cacng2 | 12300 | 3.009786 | -2.35933 | 3.350507 | 0.002981 | 0.007913 | 0.742778 | -2.35933 | 1.170258 | 0.254799 | 0.34356 | 2.967923 | 0 | 3.471878 | 27.27289 | 12.63956 | 9.468674 | 9.948826 | 30.8297 | 51.712 | 3.846072 | 6.862751 | 8.956095 | 35.87606 | 40.09021 | 60.74027 |
| Gm19382 | 100502800 | 3.00987 | -4.17258 | 3.103279 | 0.005312 | 0.013483 | 0.46595 | -4.17258 | 0.599488 | 0.555158 | 0.649755 | 0 | 0 | 3.471878 | 10.38967 | 9.830767 | 2.705335 | 7.106304 | 5.361687 | 21.83396 | 0 | 0 | 1.492683 | 5.005962 | 11.00516 | 13.76779 |
| Psg23 | 56868 | 3.010972 | -3.53288 | 3.177092 | 0.004475 | 0.011511 | 0.238885 | -3.53288 | 0.324306 | 0.748863 | 0.825265 | 0 | 1.77607 | 2.603909 | 9.090964 | 12.63956 | 10.82134 | 35.53152 | 6.702109 | 10.3424 | 0.769214 | 4.575167 | 0 | 6.674615 | 19.65207 | 14.57766 |
| Kcnh4 | 380728 | 3.012338 | 2.988209 | 19.24192 | 5.74E-15 | 8.26E-14 | -0.58162 | 2.988209 | -5.61735 | 1.34E-05 | 4.87E-05 | 223.3362 | 204.248 | 259.5229 | 2066.246 | 2047.608 | 1794.99 | 1581.863 | 1520.038 | 1095.145 | 110.7669 | 163.9435 | 136.5804 | 295.3517 | 285.348 | 273.7362 |
| Olfr1395 | 258877 | 3.015432 | -4.27736 | 3.168081 | 0.00457 | 0.011731 | 0.231681 | -4.27736 | 0.300564 | 0.766656 | 0.840509 | 2.967923 | 0 | 0 | 19.48064 | 21.06593 | 10.82134 | 5.410671 | 17.05513 | 9.193245 | 0 | 1.525056 | 1.492683 | 5.005962 | 3.14433 | 3.239481 |
| Gvin2 | 100042856 | 3.015876 | 5.106342 | 9.906143 | 1.53E-06 | 6.22E-06 | -1.55262 | 5.106342 | -6.11662 | 0.000106 | 0.000342 | 1535.158 | 1624.216 | 1454.851 | 9663.695 | 8947.402 | 8761.229 | 3311.538 | 3261.246 | 3126.852 | 3113.011 | 3182.029 | 3075.672 | 359.5949 | 406.4047 | 390.3575 |
| Spock3 | 72902 | 3.017281 | -3.18802 | 3.285399 | 0.003474 | 0.009123 | 1.467167 | -3.18802 | 2.218877 | 0.03746 | 0.067705 | 0 | 0 | 4.339848 | 14.2858 | 11.23516 | 16.23201 | 17.05513 | 14.74464 | 37.92213 | 2.307643 | 2.287584 | 2.239024 | 26.69846 | 29.87114 | 27.53559 |
| Gm4951 | 240327 | 3.019902 | 5.449869 | 47.40527 | 4.06E-23 | 3.76E-21 | -0.22351 | 5.449869 | -5.12616 | 4.24E-05 | 0.000145 | 986.0924 | 1015.912 | 867.9696 | 8050.698 | 8600.517 | 8202.577 | 7589.533 | 7468.83 | 7069.605 | 4388.368 | 4466.126 | 4254.145 | 443.0276 | 513.312 | 478.6333 |
| Cyp4b1 | 13120 | 3.021332 | -3.33391 | 3.702737 | 0.001292 | 0.003648 | 0.602672 | -3.33391 | 1.004093 | 0.326582 | 0.422709 | 4.451884 | 0.888035 | 0 | 9.090964 | 16.85274 | 10.82134 | 24.16143 | 12.0638 | 22.98311 | 0.769214 | 3.050111 | 0 | 21.6935 | 17.29382 | 16.19741 |

| | | | | | | | | | | | | | | | | | | | | | | | | | | |
|---|---|---|---|---|---|---|---|---|---|---|---|---|---|---|---|---|---|---|---|---|---|---|---|---|---|---|
| Olfr632 | 259123 | 3.021542 | -4.35894 | 3.372729 | 0.002829 | 0.007544 | -0.54934 | -4.35894 | -0.70592 | 0.487873 | 0.586317 | 2.225942 | 0 | 0 | 10.38967 | 2.808791 | 8.116006 | 2.842522 | 8.04253 | 3.447467 | 0 | 1.525056 | 1.492683 | 5.840288 | 11.00516 | 4.049351 |
| Tm4sf1 | 17112 | 3.021755 | -0.93838 | 5.07784 | 4.76E-05 | 0.000162 | 3.808112 | -0.93838 | 13.91888 | 3.51E-12 | 3.57E-11 | 17.06556 | 1.77607 | 12.15157 | 64.93546 | 74.43295 | 71.69139 | 1085.843 | 974.4866 | 1030.793 | 5.384501 | 0 | 2.985365 | 18.35519 | 26.72681 | 35.63429 |
| Olfr460 | 258381 | 3.023823 | -3.83177 | 2.865805 | 0.009157 | 0.022156 | 0.529628 | -3.83177 | 0.635735 | 0.531708 | 0.627499 | 0 | 0.888035 | 0.86797 | 5.194837 | 2.808791 | 18.93735 | 27.00396 | 5.361687 | 8.044089 | 1.538429 | 1.525056 | 3.731706 | 11.68058 | 15.72165 | 1.619741 |
| Col4a2 | 12827 | 3.024321 | 2.773376 | 15.79291 | 2.98E-13 | 3.16E-12 | 3.008043 | 2.773376 | 34.47195 | 3.34E-20 | 2.38E-18 | 82.35986 | 64.82655 | 104.1564 | 724.6797 | 790.6745 | 649.2805 | 6182.485 | 6232.961 | 5672.232 | 134.6125 | 149.4555 | 122.4 | 260.31 | 211.4562 | 234.0525 |
| Gm20100 | 100504175 | 3.027007 | 2.549602 | 18.146 | 1.87E-14 | 2.46E-13 | -0.3221 | 2.549602 | -3.13366 | 0.004951 | 0.011576 | 131.3306 | 138.5335 | 120.6478 | 1266.241 | 1115.09 | 1028.027 | 946.5597 | 1040.167 | 859.5684 | 321.5316 | 281.3728 | 226.1414 | 128.4863 | 120.2706 | 165.2135 |
| Herc6 | 67138 | 3.027064 | 7.297288 | 64.96456 | 5.09E-26 | 1.25E-23 | -0.26775 | 7.297288 | -8.68105 | 1.90E-08 | 1.03E-07 | 3701.742 | 4174.652 | 3616.829 | 33445.66 | 33413.37 | 32800.84 | 29069.05 | 28731.94 | 28206.02 | 8450.59 | 8733.994 | 7883.603 | 2467.939 | 2508.39 | 2278.165 |
| Cd38 | 12494 | 3.028114 | 6.393155 | 53.22612 | 3.49E-24 | 4.55E-22 | -0.52172 | 6.393155 | -12.9052 | 1.50E-11 | 1.36E-10 | 1803.013 | 1950.125 | 1567.553 | 15101.39 | 16278.35 | 14856.35 | 10687.88 | 10974.03 | 11643.24 | 2393.795 | 2382.137 | 2538.307 | 4356.855 | 4354.112 | 4336.045 |
| Cntnap5a | 636808 | 3.028267 | -2.82166 | 3.863518 | 0.000879 | 0.002549 | 0.62866 | -2.82166 | 1.134808 | 0.26904 | 0.359609 | 2.967923 | 0 | 3.471878 | 12.98709 | 16.85274 | 16.23201 | 28.42522 | 21.44675 | 25.28142 | 1.538429 | 4.575167 | 2.985365 | 26.69846 | 25.15464 | 8.908573 |
| 1810065E( | 69864 | 3.029431 | -3.89357 | 3.807292 | 0.001006 | 0.002892 | -0.35964 | -3.89357 | -0.53625 | 0.59733 | 0.690141 | 5.193865 | 0 | 0 | 7.792255 | 9.830767 | 8.116006 | 7.106304 | 5.361687 | 8.044089 | 1.538429 | 0.762528 | 0.746341 | 12.5149 | 9.432991 | 10.52831 |
| Fam166c | 75434 | 3.029645 | -4.4635 | 3.834443 | 0.000942 | 0.00272 | 0.818171 | -4.4635 | 1.30053 | 0.207288 | 0.289009 | 0 | 0.888035 | 0 | 2.597418 | 8.426372 | 4.058003 | 5.685043 | 8.04253 | 13.78987 | 0 | 0 | 0 | 15.01788 | 11.79124 | 12.14805 |
| Cldn34a | 635396 | 3.038055 | -5.50588 | 2.78406 | 0.011013 | 0.026173 | -1.44374 | -5.50588 | -1.39647 | 0.176929 | 0.252916 | 0 | 0 | 0 | 5.194837 | 0 | 10.82134 | 0 | 0 | 4.596622 | 0 | 0 | 1.492683 | 3.337308 | 1.572165 | 3.239481 |
| Syt7 | 54525 | 3.038902 | 1.508448 | 17.86578 | 2.56E-14 | 3.26E-13 | -1.24118 | 1.508448 | -10.071 | 1.45E-09 | 9.40E-09 | 96.4575 | 56.83424 | 73.77742 | 623.3804 | 724.668 | 645.2225 | 275.7246 | 289.5311 | 303.3771 | 140.7662 | 166.9936 | 142.5512 | 69.24913 | 66.03094 | 62.36001 |
| Krtap10-4 | 100191037 | 3.039246 | -4.14049 | 2.819755 | 0.010162 | 0.024354 | 1.907914 | -4.14049 | 2.351805 | 0.028352 | 0.0533 | 0 | 0 | 0 | 10.38967 | 1.404395 | 1.352668 | 8.527565 | 14.74464 | 18.38649 | 0 | 2.287584 | 3.731706 | 6.674615 | 13.3634 | 23.48624 |
| Olfr1502 | 258793 | 3.039246 | -4.94543 | 2.698489 | 0.013337 | 0.031084 | -0.26203 | -4.94543 | -0.26423 | 0.794135 | 0.864295 | 0 | 0 | 0 | 10.38967 | 1.404395 | 1.352668 | 2.842522 | 0 | 11.49156 | 1.538429 | 0 | 2.985365 | 4.171635 | 6.288661 | 2.429611 |
| Lemd1 | 213409 | 3.040221 | -4.30458 | 3.47183 | 0.002238 | 0.006077 | 0.928093 | -4.30458 | 1.376569 | 0.182916 | 0.26016 | 1.483961 | 0 | 0 | 7.792255 | 5.617581 | 4.058003 | 15.63387 | 9.382952 | 11.49156 | 0 | 1.525056 | 0 | 3.337308 | 5.502578 | 14.57766 |
| AI504432 | 229694 | 3.040458 | 4.981751 | 64.98007 | 5.07E-26 | 1.25E-23 | -0.73652 | 4.981751 | -21.4613 | 6.24E-16 | 1.42E-14 | 1482.478 | 1470.586 | 1474.68 | 12603.97 | 13164.8 | 13172.28 | 8321.482 | 8033.148 | 7918.831 | 410.7605 | 425.4905 | 382.1267 | 401.3112 | 350.5928 | 394.4068 |
| Ddx60 | 234311 | 3.040673 | 7.663602 | 66.72966 | 2.88E-26 | 8.33E-24 | -0.40016 | 7.663602 | -13.2397 | 9.19E-12 | 8.68E-11 | 5087.02 | 4957.011 | 4631.486 | 42692.47 | 43199.2 | 43175.8 | 34204.06 | 34199.52 | 33215.19 | 9778.254 | 10373.43 | 9015.056 | 3822.886 | 3923.338 | 3604.733 |
| Akr1c19 | 432720 | 3.042137 | -4.38749 | 3.196652 | 0.004276 | 0.01104 | -0.00091 | -4.38749 | -0.00113 | 0.99911 | 0.999256 | 0 | 0.888035 | 0 | 7.792255 | 2.808791 | 4.058003 | 1.421261 | 10.72337 | 5.745778 | 1.538429 | 0 | 3.731706 | 9.177596 | 7.074744 | 6.478962 |
| Ildr1 | 106347 | 3.042645 | -0.25653 | 6.448106 | 2.01E-06 | 8.02E-06 | 4.677082 | -0.25653 | 21.63879 | 5.27E-16 | 1.22E-14 | 7.419807 | 4.440175 | 10.41564 | 58.44191 | 80.05053 | 59.51738 | 1844.797 | 1636.655 | 1763.954 | 9.230573 | 5.337695 | 18.65853 | 38.37904 | 32.22939 | 36.44416 |
| Apobec4 | 71281 | 3.045341 | -2.03128 | 4.044379 | 0.000569 | 0.001697 | 2.379094 | -2.03128 | 5.435191 | 2.05E-05 | 7.26E-05 | 2.967923 | 2.664105 | 0.86797 | 28.5716 | 15.44835 | 18.93735 | 147.8111 | 81.76573 | 120.6613 | 12.30743 | 9.150334 | 4.478048 | 7.508942 | 29.08506 | 11.33818 |
| Gm12167 | 100503965 | 3.045671 | -5.77121 | 2.516016 | 0.019927 | 0.04442 | -2.02854 | -5.77121 | -1.71485 | 0.100859 | 0.157131 | 0 | 0 | 0 | 0 | 9.830767 | 6.763339 | 0 | 1.340422 | 0 | 6.153716 | 0 | 0 | 0 | 0.786083 | 2.429611 |
| Gm13861 | 100039184 | 3.046404 | -4.86236 | 2.88235 | 0.008819 | 0.021412 | -0.73892 | -4.86236 | -0.7833 | 0.442058 | 0.540972 | 0 | 0.888035 | 0 | 10.38967 | 5.617581 | 1.352668 | 2.842522 | 0 | 12.64071 | 1.538429 | 0.762528 | 0 | 5.840288 | 2.358248 | 3.239481 |
| Gm10860 | 100038671 | 3.047954 | -4.49173 | 3.01305 | 0.006542 | 0.016315 | 0.123008 | -4.49173 | 0.146503 | 0.8849 | 0.918721 | 1.483961 | 0 | 0 | 2.597418 | 12.63956 | 5.410671 | 2.842522 | 6.702109 | 12.64071 | 1.538429 | 1.525056 | 0 | 0.834327 | 11.00516 | 3.239481 |
| A230087F | 100503323 | 3.049204 | -3.40617 | 3.824056 | 0.000966 | 0.002785 | -0.77419 | -3.40617 | -1.13092 | 0.270638 | 0.361435 | 2.225942 | 2.664105 | 0 | 11.68838 | 16.85274 | 10.82134 | 2.842522 | 6.702109 | 18.38649 | 4.615287 | 0.762528 | 2.985365 | 13.34923 | 11.00516 | 13.76779 |
| Gm19395 | 100502820 | 3.050121 | -0.77481 | 7.574066 | 1.76E-07 | 8.04E-07 | -1.67319 | -0.77481 | -5.09218 | 4.60E-05 | 0.000156 | 17.80754 | 23.97694 | 9.547666 | 105.1954 | 137.6307 | 212.3688 | 58.27169 | 33.51054 | 51.712 | 28.46093 | 43.46409 | 28.36097 | 18.35519 | 12.57732 | 10.52831 |
| Capns2 | 69543 | 3.050692 | 0.49031 | 12.80024 | 1.75E-11 | 1.40E-10 | -1.3447 | 0.49031 | -7.56703 | 1.79E-07 | 8.44E-07 | 46.00281 | 75.48297 | 43.39848 | 420.7818 | 570.1845 | 434.2063 | 227.4017 | 183.6378 | 172.3733 | 41.53758 | 41.1765 | 22.39024 | 26.69846 | 18.0799 | 30.77507 |
| Lhx8 | 16875 | 3.052911 | -2.95234 | 3.419401 | 0.002534 | 0.006811 | 0.385937 | -2.95234 | 0.617082 | 0.543708 | 0.63871 | 0 | 2.664105 | 13.01954 | 23.37676 | 18.25714 | 20.29002 | 32.689 | 16.08506 | 40.22045 | 0.769214 | 0 | 2.239024 | 10.84625 | 33.01547 | 13.76779 |
| 4933417A1 | 66761 | 3.053273 | -4.16144 | 3.358667 | 0.002924 | 0.007774 | 0.282402 | -4.16144 | 0.385497 | 0.703684 | 0.784879 | 0 | 0.888035 | 0.86797 | 2.597418 | 9.830767 | 12.17401 | 15.63387 | 6.702109 | 6.894934 | 0 | 2.287584 | 0 | 15.85221 | 7.074744 | 5.669092 |
| Slc7a9 | 30962 | 3.053822 | -3.88782 | 3.235425 | 0.003906 | 0.010169 | 0.71243 | -3.88782 | 0.963272 | 0.346199 | 0.44358 | 1.483961 | 0 | 0 | 5.194837 | 12.63956 | 2.705335 | 11.37009 | 12.0638 | 8.044089 | 1.538429 | 4.575167 | 0 | 15.85221 | 9.432991 | 17.00728 |
| Ldoc1 | 434784 | 3.054116 | -3.78447 | 3.264589 | 0.003648 | 0.009551 | -0.63991 | -3.78447 | -0.80294 | 0.430861 | 0.529863 | 1.483961 | 2.664105 | 0 | 5.194837 | 14.04395 | 20.29002 | 2.842522 | 10.72337 | 11.49156 | 6.153716 | 2.287584 | 0 | 5.840288 | 8.646909 | 4.049351 |
| B430319G | 78575 | 3.056161 | -0.65833 | 7.829942 | 1.04E-07 | 4.90E-07 | -1.4013 | -0.65833 | -4.67388 | 0.000125 | 0.000399 | 13.35565 | 7.10428 | 39.05863 | 145.4554 | 162.9099 | 119.0348 | 54.00791 | 58.97856 | 52.86116 | 6.153716 | 3.812639 | 8.209754 | 89.27298 | 92.75775 | 117.4312 |
| Olfr668 | 259061 | 3.057678 | -5.27811 | 2.466408 | 0.022188 | 0.048883 | -1.53602 | -5.27811 | -1.30811 | 0.204751 | 0.286089 | 0 | 0 | 0 | 0 | 8.426372 | 8.116006 | 5.685043 | 0 | 0 | 1.538429 | 0 | 1.492683 | 3.337308 | 11.00516 | 0.80987 |
| Gipr | 381853 | 3.057813 | -0.88096 | 8.230427 | 4.61E-08 | 2.28E-07 | -0.50331 | -0.88096 | -2.10574 | 0.047234 | 0.082463 | 19.2915 | 7.10428 | 12.15157 | 100.0006 | 137.6307 | 101.4501 | 79.59061 | 93.82952 | 73.54596 | 6.153716 | 10.67539 | 5.97073 | 35.04173 | 44.02063 | 42.92312 |
| Gm17830 | 100415784 | 3.059011 | -3.44954 | 3.075128 | 0.00567 | 0.014326 | 1.119032 | -3.44954 | 1.53392 | 0.13974 | 0.206943 | 0 | 6.216245 | 0 | 9.090964 | 11.23516 | 5.410671 | 18.47639 | 8.04253 | 42.51876 | 7.692145 | 1.525056 | 2.239024 | 6.674615 | 9.432991 | 7.288832 |
| Olfr576 | 258248 | 3.061882 | -4.82426 | 2.883197 | 0.008803 | 0.021381 | 1.703881 | -4.82426 | 2.113685 | 0.046479 | 0.08135 | 0 | 0 | 0 | 0 | 12.63956 | 5.410671 | 9.948826 | 25.46801 | 6.894934 | 0 | 0 | 0 | 3.337308 | 5.502578 | 7.288832 |
| 4930519F1 | 75106 | 3.0638 | -4.1687 | 3.034628 | 0.006225 | 0.015596 | 0.638979 | -4.1687 | 0.815696 | 0.423684 | 0.522858 | 0 | 3.55214 | 0 | 2.597418 | 11.23516 | 10.82134 | 11.37009 | 25.46801 | 5.745778 | 0 | 0 | 1.492683 | 9.177596 | 11.79124 | 3.239481 |
| Fndc11 | 332713 | 3.064668 | -1.47228 | 5.124107 | 4.27E-05 | 0.000146 | 0.436796 | -1.47228 | 1.195121 | 0.245151 | 0.332546 | 5.935846 | 8.88035 | 1.735939 | 45.45482 | 40.72746 | 47.34337 | 44.05909 | 65.68067 | 80.44089 | 10.769 | 16.01308 | 9.702436 | 19.18952 | 19.65207 | 48.59222 |
| Neu4 | 241159 | 3.065297 | -3.3129 | 3.151705 | 0.004747 | 0.012155 | 1.841893 | -3.3129 | 2.719877 | 0.012716 | 0.026623 | 2.225942 | 0 | 0 | 1.298709 | 22.47032 | 8.116006 | 25.5827 | 26.80843 | 26.43058 | 3.076858 | 1.525056 | 0.746341 | 20.85817 | 36.94588 | 21.05663 |
| Prss32 | 69814 | 3.065316 | -3.50487 | 3.182353 | 0.004421 | 0.011387 | 0.791879 | -3.50487 | 1.110903 | 0.278969 | 0.370526 | 0 | 1.77607 | 1.735939 | 9.090964 | 19.66153 | 5.410671 | 14.21261 | 13.40422 | 29.87805 | 4.615287 | 0 | 6.717071 | 6.674615 | 5.502578 | 8.098703 |
| Flnb | 286940 | 3.065515 | 7.405654 | 43.8491 | 1.92E-18 | 5.67E-17 | 0.454118 | 7.405654 | 10.92134 | 6.09E-09 | 3.57E-08 | 3281.781 | 3235.999 | 3570.827 | 30139.14 | 30313.87 | 29600.43 | 44125.89 | 45058.28 | 39664.25 | 4953.741 | 4944.231 | 5258.72 | 5003.459 | 4869.782 | 4815.489 |
| Cdc42ep2 | 104252 | 3.06641 | 3.597702 | 20.97068 | 1.00E-15 | 1.67E-14 | 1.474872 | 3.597702 | 19.83575 | 3.10E-15 | 5.96E-14 | 157.2999 | 184.7113 | 134.5353 | 1474.035 | 1467.593 | 1294.503 | 4403.066 | 4148.605 | 3738.203 | 1329.203 | 1331.374 | 1424.765 | 149.3445 | 158.7887 | 171.6925 |
| 4930474N( | 218921 | 3.066494 | -4.73093 | 3.375764 | 0.002809 | 0.007497 | -0.19547 | -4.73093 | -0.25141 | 0.803901 | 0.872973 | 0 | 0 | 0.86797 | 2.597418 | 7.021976 | 5.410671 | 2.842522 | 5.361687 | 4.596622 | 0 | 0 | 0.746341 | 16.68654 | 6.288661 | 2.429611 |
| Nalf1 | 270028 | 3.067081 | -2.70361 | 3.879526 | 0.000846 | 0.002458 | 0.227083 | -2.70361 | 0.419462 | 0.67907 | 0.762902 | 6.677827 | 7.10428 | 0 | 32.46773 | 15.44835 | 31.11136 | 25.5827 | 34.85097 | 32.17636 | 1.538429 | 1.525056 | 0 | 24.19548 | 35.37372 | 18.62702 |
| Fgf23 | 64654 | 3.068884 | -2.67437 | 3.610664 | 0.001609 | 0.004475 | 0.89056 | -2.67437 | 1.571525 | 0.130769 | 0.195963 | 5.935846 | 0 | 2.603909 | 11.68838 | 23.87472 | 21.64268 | 27.00396 | 32.17012 | 48.26453 | 3.846072 | 4.575167 | 0 | 25.86413 | 28.29897 | 24.29611 |
| 6530411M( | 67791 | 3.069459 | -4.46855 | 2.692455 | 0.013517 | 0.031473 | -0.70539 | -4.46855 | -0.70493 | 0.488478 | 0.586794 | 0 | 3.55214 | 0 | 9.090964 | 12.63956 | 2.705335 | 14.21261 | 0 | 10.3424 | 3.076858 | 0 | 0 | 5.840288 | 5.502578 | 4.859222 |
| Rhox2g | 434766 | 3.070117 | -4.74468 | 3.569286 | 0.001776 | 0.004904 | -0.13191 | -4.74468 | -0.18095 | 0.858116 | 0.918721 | 0 | 0 | 1.735939 | 5.194837 | 8.426372 | 4.058003 | 4.263783 | 2.680843 | 11.49156 | 0.769214 | 0 | 0 | 5.005962 | 4.716496 | 1.619741 |
| Taar7f | 435207 | 3.071523 | -4.65375 | 3.091992 | 0.005453 | 0.01382 | -1.49229 | -4.65375 | -1.62171 | 0.119546 | 0.181493 | 2.967923 | 0 | 0 | 10.38967 | 12.63956 | 2.705335 | 2.842522 | 0 | 9.193245 | 0 | 1.525056 | 0 | 6.674615 | 6.288661 | 4.049351 |
| Rasl11a | 68895 | 3.072796 | -2.20007 | 4.869147 | 7.83E-05 | 0.000259 | 0.521996 | -2.20007 | 1.254509 | 0.223222 | 0.307606 | 0.741981 | 3.55214 | 5.207818 | 22.07806 | 25.27911 | 28.40602 | 48.32287 | 42.8935 | 27.57973 | 1.538429 | 5.337695 | 1.492683 | 48.39096 | 50.30929 | 38.87377 |
| Enox1 | 239188 | 3.073752 | -1.34074 | 7.309939 | 3.07E-07 | 1.36E-06 | -0.96759 | -1.34074 | -3.18728 | 0.00437 | 0.010363 | 6.677827 | 6.216245 | 13.88751 | 81.81868 | 84.26372 | 67.63339 | 46.90161 | 29.48928 | 48.26453 | 13.07665 | 9.150334 | 10.44878 | 30.03577 | 24.36856 | 16.19741 |
| Olfr586 | 259115 | 3.075489 | -4.40652 | 2.828493 | 0.009964 | 0.023933 | -0.07064 | -4.40652 | -0.07696 | 0.939372 | 0.949955 | 0 | 1.77607 | 0 | 3.896127 | 5.617581 | 8.116006 | 1.421261 | 8.04253 | 11.49156 | 1.538429 | 0 | 2.985365 | 10.84625 | 7.860826 | 0.80987 |
| Rhox2f | 434764 | 3.075955 | -5.09116 | 2.905194 | 0.008373 | 0.020423 | 0.431735 | -5.09116 | 0.486522 | 0.631556 | 0.721132 | 0 | 0 | 0 | 6.493546 | 0 | 9.468674 | 1.421261 | 12.0638 | 4.596622 | 0 | 0.762528 | 0 | 5.005962 | 2.358248 | 3.239481 |
| Crisp4 | 78081 | 3.076021 | -4.04595 | 3.600009 | 0.001651 | 0.004578 | -0.1643 | -4.04595 | -0.22741 | 0.82227 | 0.889504 | 1.483961 | 0 | 0 | 5.194837 | 7.021976 | 5.410671 | 5.685043 | 2.680843 | 9.193245 | 3.076858 | 0.762528 | 0.746341 | 8.343269 | 12.57732 | 22.67637 |
| Gvin-ps3 | 668108 | 3.076026 | -4.8654 | 2.599287 | 0.016611 | 0.037767 | -0.66838 | -4.8654 | -0.63761 | 0.530507 | 0.626383 | 0 | 0.888035 | 0 | 5.194837 | 7.021976 | 2.705335 | 0 | 6.702109 | 6.894934 | 0 | 3.812639 | 2.239024 | 2.502981 | 7.074744 | 0 |
| Il36g | 215257 | 3.076403 | 3.301592 | 30.743 | 3.68E-19 | 1.28E-17 | 0.890301 | 3.301592 | 15.92803 | 2.52E-13 | 3.18E-12 | 323.5036 | 237.9934 | 322.0167 | 2584.431 | 2574.257 | 2790.554 | 5143.543 | 5124.432 | 5022.959 | 156.1505 | 164.706 | 167.1804 | 117.6401 | 91.97167 | 123.9102 |
| Zscan4d | 545913 | 3.077505 | -4.08637 | 3.114293 | 0.005178 | 0.013161 | -1.00914 | -4.08637 | -1.15389 | 0.261305 | 0.350895 | 1.483961 | 0 | 1.735939 | 18.18193 | 4.213186 | 13.52668 | 1.421261 | 5.361687 | 12.64071 | 0 | 1.525056 | 2.985365 | 15.01788 | 1.572165 | 4.859222 |
| Tdpoz1 | 207213 | 3.080766 | -4.04725 | 2.542315 | 0.018818 | 0.042241 | 0.010846 | -4.04725 | 0.010673 | 0.991583 | 0.992942 | 0 | 1.77607 | 0 | 3.896127 | 11.23516 | 4.058003 | 17.05513 | 0 | 21.83396 | 4.615287 | 1.525056 | 1.492683 | 4.171635 | 14.93557 | 7.288832 |
| Phf11b | 236451 | 3.083312 | 5.314678 | 72.15738 | 5.47E-27 | 1.89E-24 | -1.68256 | 5.314678 | -46.4922 | 6.12E-23 | 1.13E-20 | 1875.727 | 1886.186 | 1892.174 | 17122.18 | 17224.91 | 16825.83 | 5487.488 | 5530.58 | 5520.543 | 3493.003 | 3398.587 | 3242.853 | 141.0012 | 102.1907 | 141.7273 |
| 5031426D | 68144 | 3.084044 | -2.70326 | 4.259886 | 0.000338 | 0.00104 | 0.795728 | -2.70326 | 1.565063 | 0.132275 | 0.197788 | 1.483961 | 0 | 4.339848 | 10.38967 | 15.44835 | 16.23201 | 27.00396 | 21.44675 | 27.57973 | 4.615287 | 1.525056 | 4.478048 | 35.04173 | 26.72681 | 29.9652 |
| Nfkbie | 18037 | 3.084343 | 5.176661 | 34.46749 | 3.35E-20 | 1.48E-18 | 1.270075 | 5.176661 | 25.19714 | 2.30E-17 | 7.33E-16 | 562.4214 | 531.933 | 552.0287 | 4827.302 | 5120.425 | 5000.813 | 12882.31 | 13144.18 | 11569.7 | 1250.743 | 1298.585 | 1283.707 | 1115.495 | 1021.907 | 1103.043 |
| Cdh19 | 227485 | 3.084456 | -3.98164 | 2.800938 | 0.010603 | 0.025266 | 0.939999 | -3.98164 | 1.156543 | 0.260241 | 0.349749 | 4.451884 | 0 | 0 | 15.58451 | 2.808791 | 12.17401 | 29.84648 | 5.361687 | 29.87805 | 0 | 0 | 0 | 19.18952 | 21.22423 | 4.859222 |
| Cyp2g1 | 13108 | 3.087072 | -3.48306 | 3.407952 | 0.002603 | 0.006986 | 1.201306 | -3.48306 | 1.887214 | 0.072807 | 0.119175 | 0 | 3.55214 | 0.86797 | 5.194837 | 11.23516 | 18.93735 | 22.74017 | 24.12759 | 29.87805 | 0 | 0.762528 | 2.985365 | 18.35519 | 4.716496 | 12.14805 |
| Olfr123 | 258623 | 3.089011 | -4.81463 | 2.730997 | 0.012404 | 0.029129 | -0.28761 | -4.81463 | -0.29625 | 0.7699 | 0.843258 | 0 | 0 | 1.735939 | 5.194837 | 12.63956 | 2.705335 | 0 | 8.04253 | 21.83396 | 0 | 0 | 0 | 10.84625 | 2.358248 | 3.239481 |
| Pramel7 | 347712 | 3.090693 | -4.35684 | 3.574201 | 0.001755 | 0.004852 | 0.113219 | -4.35684 | 0.157729 | 0.876153 | 0.918721 | 0 | 1.77607 | 0 | 3.896127 | 7.021976 | 6.763339 | 7.106304 | 2.680843 | 12.64071 | 0.769214 | 1.525056 | 0 | 8.343269 | 6.288661 | 5.669092 |
| Trav21-dv1 | 100101940 | 3.091495 | -5.30452 | 2.675284 | 0.014043 | 0.032548 | -1.37971 | -5.30452 | -1.2743 | 0.216257 | 0.299335 | 0 | 0 | 0 | 2.597418 | 2.808791 | 4.058003 | 0 | 8.04253 | 0 | 0 | 0 | 5.224389 | 5.005962 | 0.786083 | 6.478962 |
| Tlr3 | 142980 | 3.09278 | 5.815889 | 63.78808 | 7.51E-26 | 1.73E-23 | -1.7516 | 5.815889 | -42.2301 | 4.66E-22 | 5.99E-20 | 2098.322 | 2171.246 | 2327.894 | 19711.81 | 20497.15 | 19774.65 | 6010.512 | 5952.813 | 6476.641 | 3912.225 | 4180.178 | 3674.984 | 381.2874 | 436.2759 | 411.4141 |
| Tmc3 | 233424 | 3.093933 | 3.410467 | 42.82851 | 3.47E-22 | 2.38E-20 | -1.01699 | 3.410467 | -19.3352 | 5.21E-15 | 9.44E-14 | 449.6403 | 447.5696 | 418.3613 | 4128.596 | 3964.608 | 3914.62 | 2015.348 | 2147.356 | 2002.978 | 316.9164 | 321.0242 | 308.9853 | 105.1252 | 78.60826 | 89.8956 |
| Gm10228 | 100040214 | 3.094319 | -5.08387 | 2.989544 | 0.006905 | 0.017152 | -2.05134 | -5.08387 | -2.02822 | 0.0552 | 0.094341 | 0 | 0 | 0 | 3.896127 | 2.808791 | 2.705335 | 0 | 0 | 1.149156 | 0 | 7.625279 | 2.985365 | 6.674615 | 2.358248 | 7.288832 |
| Gm7489 | 665097 | 3.099056 | -4.04104 | 3.316637 | 0.003229 | 0.00853 | 0.166368 | -4.04104 | 0.224182 | 0.824748 | 0.891488 | 0 | 0 | 6.943757 | 10.38967 | 8.426372 | 8.116006 | 14.21261 | 10.72337 | 8.044089 | 1.538429 | 0 | 0 | 5.840288 | 4.716496 | 17.81715 |
| Fam241a | 70617 | 3.099552 | 5.027833 | 55.53385 | 1.42E-24 | 2.10E-22 | -0.94366 | 5.027833 | -22.5889 | 2.18E-16 | 5.57E-15 | 1017.256 | 1088.731 | 1118.813 | 9679.279 | 10315.28 | 9471.38 | 5429.216 | 5104.326 | 5348.17 | 1384.586 | 1474.729 | 1446.409 | 311.2039 | 378.8918 | 368.491 |
| Gm20187 | 100504347 | 3.099809 | -4.05639 | 3.246387 | 0.003807 | 0.009934 | -0.68687 | -4.05639 | -0.83809 | 0.411267 | 0.510271 | 5.935846 | 0 | 0 | 16.88322 | 7.021976 | 6.763339 | 4.263783 | 8.04253 | 5.745778 | 2.307643 | 2.287584 | 0 | 2.502981 | 7.074744 | 10.52831 |
| Smim31 | 407795 | 3.117823 | -3.87118 | 2.84737 | 0.009548 | 0.023018 | 1.076981 | -3.87118 | 1.301553 | 0.206944 | 0.288649 | 0 | 0 | 1.735939 | 2.597418 | 25.27911 | 2.705335 | 11.37009 | 13.40422 | 16.08818 | 4.615287 | 3.050111 | 0 | 4.171635 | 12.57732 | 12.95792 |
| Ptf1a | 19213 | 3.11875 | -2.43139 | 3.448706 | 0.002364 | 0.006398 | 0.597557 | -2.43139 | 0.95763 | 0.348972 | 0.446221 | 7.419807 | 1.77607 | 0 | 14.2858 | 25.27911 | 16.23201 | 15.63387 | 24.12759 | 54.01031 | 10.769 | 9.150334 | 1.492683 | 35.04173 | 29.08506 | 20.24676 |
| Sfrp4 | 20379 | 3.120055 | -2.70333 | 3.843129 | 0.000923 | 0.002667 | -0.1148 | -2.70333 | -0.19514 | 0.84713 | 0.910542 | 6.677827 | 0 | 4.339848 | 19.48064 | 23.87472 | 25.70069 | 15.63387 | 26.80843 | 24.13227 | 0 | 2.287584 | 2.985365 | 23.36115 | 49.52321 | 30.77507 |
| Fam50b | 108161 | 3.121291 | -3.50982 | 2.950948 | 0.007543 | 0.018572 | 1.860936 | -3.50982 | 2.558203 | 0.018176 | 0.036307 | 0 | 0 | 3.471878 | 2.597418 | 4.213186 | 31.11136 | 36.95278 | 22.78717 | 29.87805 | 3.846072 | 0 | 0 | 27.53279 | 18.0799 | 22.67637 |
| Klk1b22 | 13646 | 3.128997 | -4.34127 | 3.626788 | 0.001549 | 0.004317 | 1.106132 | -4.34127 | 1.616586 | 0.120654 | 0.182916 | 0 | 0 | 0 | 3.896127 | 5.617581 | 1.352668 | 15.63387 | 4.021265 | 8.044089 | 0.769214 | 3.050111 | 2.239024 | 5.840288 | 8.646909 | 6.478962 |
| Sprr2a3 | 100042514 | 3.128997 | -4.82671 | 2.988984 | 0.006914 | 0.017172 | 0.463197 | -4.82671 | 0.531132 | 0.600812 | 0.693433 | 0 | 0 | 0 | 3.896127 | 5.617581 | 1.352668 | 4.263783 | 2.680843 | 9.193245 | 7.692145 | 0 | 0 | 3.337308 | 3.14433 | 6.478962 |
| Serpina3i | 628900 | 3.129756 | -1.50292 | 4.560007 | 0.000164 | 0.000524 | 2.292361 | -1.50292 | 6.259531 | 3.06E-06 | 1.22E-05 | 2.225942 | 2.664105 | 6.075787 | 59.74062 | 22.47032 | 28.40602 | 159.1812 | 168.8931 | 197.6548 | 12.30743 | 19.82572 | 9.702436 | 5.005962 | 17.29382 | 25.10598 |
| G530011O | 654820 | 3.132158 | 5.329517 | 51.04585 | 8.47E-24 | 9.65E-22 | -0.9117 | 5.329517 | -20.1437 | 2.27E-15 | 4.50E-14 | 990.5443 | 884.4828 | 1025.072 | 9046.808 | 8948.807 | 9157.561 | 5157.756 | 4972.965 | 4872.42 | 1163.052 | 1149.892 | 1019.502 | 1635.281 | 1587.101 | 1746.89 |
| Gpr84 | 80910 | 3.132509 | 4.621261 | 84.18594 | 2.06E-28 | 1.46E-25 | -3.22533 | 4.621261 | -76.8831 | 1.42E-27 | 2.07E-24 | 2856.626 | 3005.998 | 2998.835 | 27730.04 | 27842.14 | 27281.96 | 2978.963 | 3091.013 | 3113.062 | 819.2134 | 801.4168 | 959.7949 | 38.37904 | 25.15464 | 33.20468 |
| Gzmb | 14939 | 3.135194 | -2.93049 | 4.022178 | 0.0006 | 0.001785 | -0.02562 | -2.93049 | -0.04166 | 0.967154 | 0.972942 | 2.967923 | 0.888035 | 0 | 7.792255 | 16.85274 | 10.82134 | 7.106304 | 20.10633 | 10.3424 | 1.538429 | 0.762528 | 0 | 167.6997 | 183.1573 | 175.7418 |
| Nudt10 | 102954 | 3.136492 | -3.62944 | 2.848832 | 0.009516 | 0.022949 | 0.980556 | -3.62944 | 1.162988 | 0.257673 | 0.34667 | 0 | 0 | 1.735939 | 5.194837 | 14.04395 | 2.705335 | 12.79135 | 2.680843 | 45.96622 | 2.307643 | 1.525056 | 4.478048 | 10.84625 | 13.3634 | 21.05663 |
| Olfr1496 | 258991 | 3.139026 | -5.09757 | 2.889354 | 0.00868 | 0.021115 | -0.83463 | -5.09757 | -0.84778 | 0.405966 | 0.505053 | 0 | 0 | 0 | 1.298709 | 2.808791 | 8.116006 | 0 | 1.340422 | 6.894934 | 1.538429 | 0 | 0 | 17.52087 | 6.288661 | 2.429611 |
| 1700067P | 68224 | 3.139421 | -4.65005 | 2.945341 | 0.00764 | 0.0188 | 1.034584 | -4.65005 | 1.219975 | 0.235784 | 0.32185 | 0 | 0 | 0 | 5.194837 | 0 | 13.52668 | 12.79135 | 8.04253 | 4.596622 | 0 | 0 | 0 | 10.01192 | 25.94073 | 11.33818 |
| Gm525 | 217071 | 3.140029 | -2.17043 | 5.374838 | 2.36E-05 | 8.34E-05 | 0.223391 | -2.17043 | 0.600689 | 0.554373 | 0.649036 | 2.967923 | 3.55214 | 4.339848 | 35.06515 | 26.68351 | 52.75404 | 51.16539 | 57.63814 | 32.17636 | 6.153716 | 6.100223 | 2.985365 | 5.840288 | 13.3634 | 17.00728 |
| Asap3 | 230837 | 3.14014 | 1.724048 | 11.59991 | 1.11E-10 | 7.88E-10 | 2.634589 | 1.724048 | 21.84655 | 4.33E-16 | 1.02E-14 | 44.51884 | 56.83424 | 32.98284 | 351.9502 | 463.4504 | 439.617 | 2856.734 | 2834.992 | 2397.139 | 44.61444 | 39.65145 | 38.0634 | 147.6759 | 167.4356 | 130.3891 |
| Klk5 | 68668 | 3.140669 | -4.52254 | 3.779825 | 0.001074 | 0.003078 | 0.270776 | -4.52254 | 0.40074 | 0.692595 | 0.77475 | 0 | 0 | 1.735939 | 10.38967 | 7.021976 | 2.705335 | 4.263783 | 10.72337 | 9.193245 | 0 | 0 | 0 | 10.01192 | 9.432991 | 4.859222 |
| BC051076 | 243090 | 3.141154 | -2.96327 | 3.414944 | 0.002561 | 0.006878 | 1.165403 | -2.96327 | 1.811702 | 0.084128 | 0.134598 | 0 | 0 | 10.41564 | 5.194837 | 16.85274 | 13.52668 | 31.26774 | 17.42548 | 29.87805 | 3.846072 | 3.050111 | 1.492683 | 23.36115 | 22.01031 | 27.53559 |
| C130026I2 | 620078 | 3.142962 | 1.673848 | 16.57799 | 1.14E-13 | 1.31E-12 | -0.35204 | 1.673848 | -3.11311 | 0.005192 | 0.01209 | 63.06836 | 73.7069 | 62.49381 | 598.7049 | 630.5735 | 650.6332 | 525.8665 | 481.2114 | 519.4183 | 174.6117 | 115.9042 | 166.4341 | 97.61625 | 64.45877 | 76.93768 |
| Ifna11 | 15964 | 3.145287 | -4.89186 | 3.639323 | 0.001503 | 0.004198 | 0.983946 | -4.89186 | 1.42026 | 0.169976 | 0.244492 | 0 | 0 | 0 | 2.597418 | 1.404395 | 8.116006 | 2.842522 | 12.0638 | 10.3424 | 0 | 0 | 0 | 7.508942 | 7.074744 | 4.859222 |
| Gm19763 | 100503548 | 3.145287 | -5.26303 | 2.831529 | 0.009896 | 0.023795 | -0.5223 | -5.26303 | -0.53064 | 0.601149 | 0.693731 | 0 | 0 | 0 | 2.597418 | 1.404395 | 8.116006 | 5.685043 | 0 | 4.596622 | 2.307643 | 0 | 1.492683 | 3.337308 | 2.358248 | 0 |
| A530095I0 | 210373 | 3.145422 | -1.64704 | 4.649158 | 0.000133 | 0.000427 | 0.227013 | -1.64704 | 0.545469 | 0.591086 | 0.684341 | 6.677827 | 4.440175 | 2.603909 | 41.55869 | 51.96262 | 40.58003 | 95.22448 | 42.8935 | 41.3696 | 6.153716 | 19.82572 | 10.44878 | 8.343269 | 36.94588 | 20.24676 |
| Lad1 | 16763 | 3.146513 | 3.214709 | 19.86739 | 3.00E-15 | 4.54E-14 | -0.55389 | 3.214709 | -5.53893 | 1.61E-05 | 5.77E-05 | 142.4603 | 222.8968 | 152.7627 | 1770.141 | 1585.562 | 1416.243 | 1178.225 | 1201.018 | 1013.555 | 1240.743 | 1326.798 | 1328.487 | 154.3505 | 103.7629 | 147.3964 |
| Psg28 | 114871 | 3.149053 | -4.02579 | 3.13483 | 0.004937 | 0.012604 | -0.00371 | -4.02579 | -0.00449 | 0.996456 | 0.997062 | 0 | 0.888035 | 0.86797 | 10.38967 | 7.021976 | 5.410671 | 2.842522 | 12.0638 | 11.49156 | 0.769214 | 1.525056 | 3.731706 | 12.5149 | 11.00516 | 0.80987 |
| Methig1 | 554292 | 3.150484 | -3.36274 | 3.198093 | 0.004261 | 0.011008 | -0.59608 | -3.36274 | -0.7251 | 0.476269 | 0.574656 | 0 | 3.55214 | 1.735939 | 16.88322 | 5.617581 | 21.64268 | 2.842522 | 25.46801 | 8.044089 | 4.615287 | 10.67539 | 7.463413 | 0.834327 | 8.646909 | 8.908573 |
| Olfr1168 | 258524 | 3.15065 | -4.78962 | 3.20872 | 0.004157 | 0.01076 | -0.01761 | -4.78962 | -0.0217 | 0.982887 | 0.985585 | 1.483961 | 0 | 0 | 1.298709 | 11.23516 | 13.52668 | 5.685043 | 13.40422 | 3.447467 | 0 | 0.762528 | 0 | 0.834327 | 1.572165 | 5.669092 |
| Klra8 | 16639 | 3.151945 | -4.23107 | 3.460423 | 0.0023 | 0.006233 | 0.091028 | -4.23107 | 0.121772 | 0.904219 | 0.924752 | 1.483961 | 0 | 0 | 6.493546 | 12.63956 | 2.705335 | 9.948826 | 4.021265 | 8.044089 | 0 | 3.050111 | 0 | 12.5149 | 12.57732 | 8.098703 |
| Ddx4 | 13206 | 3.155796 | 0.213413 | 9.348209 | 5.36E-09 | 3.01E-08 | -2.75633 | 0.213413 | -7.81678 | 1.07E-07 | 5.20E-07 | 14.09763 | 33.74533 | 27.77503 | 237.6638 | 207.8505 | 221.8375 | 21.31891 | 32.17012 | 50.56285 | 161.535 | 150.9805 | 139.5658 | 67.58048 | 79.39434 | 38.0639 |
| Anks4b | 72074 | 3.156459 | -4.27842 | 2.997746 | 0.006776 | 0.016853 | 0.670225 | -4.27842 | 0.802251 | 0.431251 | 0.530213 | 0 | 0 | 0.86797 | 2.597418 | 4.213186 | 10.82134 | 5.685043 | 10.72337 | 10.3424 | 0 | 3.050111 | 1.492683 | 1.668654 | 6.288661 | 19.43689 |
| Gm5321 | 384525 | 3.156459 | -4.86214 | 2.622662 | 0.015777 | 0.036149 | -1.18007 | -4.86214 | -1.07875 | 0.292741 | 0.385972 | 0 | 0 | 0.86797 | 2.597418 | 4.213186 | 10.82134 | 5.685043 | 0 | 3.447467 | 0 | 4.575167 | 2.985365 | 7.508942 | 0 | 3.239481 |
| Nav3 | 260315 | 3.156516 | -0.97549 | 7.606218 | 1.65E-07 | 7.55E-07 | -1.03422 | -0.97549 | -3.59435 | 0.001673 | 0.004362 | 19.2915 | 9.768385 | 8.679696 | 97.40319 | 101.1165 | 169.0835 | 68.22052 | 58.97856 | 54.01031 | 12.30743 | 6.100223 | 3.731706 | 40.04769 | 49.52321 | 36.44416 |
| C820005J( | 319979 | 3.15868 | -4.02301 | 2.991249 | 0.006878 | 0.017088 | -0.32383 | -4.02301 | -0.36109 | 0.721585 | 0.801256 | 0 | 1.77607 | 0 | 16.88322 | 4.213186 | 2.705335 | 12.79135 | 4.021265 | 2.298311 | 6.92293 | 1.525056 | 0 | 11.68058 | 15.72165 | 15.38754 |
| Tex37 | 74221 | 3.158738 | -4.9368 | 3.074027 | 0.005684 | 0.014358 | -1.70085 | -4.9368 | -1.75902 | 0.092904 | 0.146509 | 0 | 0 | 1.735939 | 9.090964 | 8.426372 | 2.705335 | 0 | 1.340422 | 5.745778 | 4.615287 | 0.762528 | 0.746341 | 4.171635 | 0 | 1.619741 |
| Gm5524 | 433281 | 3.159243 | -4.01607 | 3.404615 | 0.002624 | 0.007039 | 0.273332 | -4.01607 | 0.373529 | 0.71244 | 0.792936 | 0 | 0.888035 | 1.735939 | 7.792255 | 8.426372 | 10.82134 | 2.842522 | 18.7659 | 21.83396 | 3.076858 | 0 | 0 | 7.508942 | 7.860826 | 6.478962 |
| Gfpt2 | 14584 | 3.16189 | -0.05361 | 6.52912 | 1.67E-06 | 6.75E-06 | 3.638472 | -0.05361 | 16.67225 | 1.02E-13 | 1.41E-12 | 5.935846 | 6.216245 | 14.75548 | 96.10448 | 77.24174 | 70.33872 | 1060.261 | 998.6142 | 1092.847 | 6.92293 | 16.77561 | 17.91219 | 46.72231 | 66.03094 | 89.08573 |
| Inhbb | 16324 | 3.162563 | -1.42525 | 4.826977 | 8.66E-05 | 0.000286 | 1.196472 | -1.42525 | 3.098447 | 0.005372 | 0.012451 | 0.741981 | 5.32821 | 5.207818 | 19.48064 | 46.34504 | 31.11136 | 88.11817 | 57.63814 | 75.84427 | 7.692145 | 16.77561 | 14.92683 | 47.55663 | 55.81187 | 65.59949 |
| Pax4 | 18506 | 3.163318 | -3.71623 | 3.722467 | 0.001232 | 0.003495 | 0.557736 | -3.71623 | 0.891379 | 0.382665 | 0.48114 | 2.967923 | 0 | 0.86797 | 5.194837 | 19.66153 | 14.87935 | 15.63387 | 10.72337 | 31.0272 | 1.538429 | 0 | 0 | 14.18356 | 7.074744 | 11.33818 |
| Ccn2 | 14219 | 3.168249 | -1.85189 | 4.020862 | 0.000602 | 0.00179 | 2.561276 | -1.85189 | 5.880353 | 7.27E-06 | 2.75E-05 | 5.193865 | 0.888035 | 1.735939 | 20.77935 | 19.66153 | 33.81669 | 113.7009 | 166.2123 | 173.5225 | 5.384501 | 5.337695 | 0 | 62.57452 | 59.74228 | 51.02183 |
| Gm14496 | 672125 | 3.168897 | -3.50613 | 3.372568 | 0.00283 | 0.007546 | -0.38435 | -3.50613 | -0.50545 | 0.618421 | 0.709249 | 4.451884 | 0 | 0.86797 | 14.2858 | 19.66153 | 8.116006 | 2.842522 | 20.10633 | 16.08818 | 2.307643 | 0 | 3.731706 | 18.35519 | 7.860826 | 10.52831 |
| Bmp1 | 12153 | 3.169831 | 1.491595 | 19.73136 | 3.45E-15 | 5.16E-14 | -2.92992 | 1.491595 | -17.0072 | 6.86E-14 | 9.84E-13 | 108.3292 | 91.4676 | 121.5157 | 932.4732 | 1131.943 | 1034.791 | 112.2796 | 170.2336 | 140.197 | 130.7665 | 95.31598 | 128.3707 | 66.74615 | 90.3995 | 67.21923 |
| Sox8 | 20681 | 3.171482 | -2.64837 | 3.253463 | 0.003744 | 0.009784 | 1.627164 | -2.64837 | 2.484669 | 0.021329 | 0.041715 | 0 | 1.77607 | 1.735939 | 12.98709 | 2.808791 | 29.75869 | 24.16143 | 25.46801 | 66.65102 | 1.538429 | 3.050111 | 2.239024 | 55.06558 | 81.75259 | 54.26131 |
| Tacr2 | 21337 | 3.173259 | -2.2562 | 4.280855 | 0.000322 | 0.000992 | 1.379962 | -2.2562 | 3.288037 | 0.003453 | 0.008395 | 0.741981 | 6.216245 | 6.943757 | 41.55869 | 32.30109 | 31.11136 | 100.9095 | 89.80826 | 95.37991 | 0 | 3.812639 | 2.985365 | 32.53875 | 7.074744 | 9.718443 |
| Artn | 11876 | 3.173274 | -2.28434 | 3.256619 | 0.003717 | 0.009721 | 1.40906 | -2.28434 | 2.242688 | 0.035654 | 0.064944 | 6.677827 | 0 | 0.86797 | 12.98709 | 15.44835 | 17.58468 | 58.27169 | 21.44675 | 60.90525 | 13.84586 | 1.525056 | 8.209754 | 53.39692 | 46.37887 | 27.53559 |
| Krt40 | 406221 | 3.179691 | -4.16803 | 3.061237 | 0.005855 | 0.014746 | -1.02687 | -4.16803 | -1.11295 | 0.278108 | 0.369696 | 0 | 1.77607 | 0.86797 | 7.792255 | 14.04395 | 6.763339 | 0 | 6.702109 | 19.53565 | 2.307643 | 1.525056 | 0 | 6.674615 | 3.930413 | 10.52831 |
| Ifi47 | 15953 | 3.182432 | 6.815005 | 63.90612 | 7.22E-26 | 1.68E-23 | -0.61197 | 6.815005 | -18.1546 | 1.86E-14 | 2.99E-13 | 3459.114 | 3876.273 | 3356.438 | 34984.63 | 35427.27 | 32917.17 | 24064.79 | 24025.72 | 22291.32 | 6472.94 | 6684.319 | 6561.832 | 697.4973 | 773.5053 | 724.8339 |
| Gca | 227960 | 3.185978 | 3.535777 | 45.223 | 1.10E-22 | 8.63E-21 | -2.64679 | 3.535777 | -38.0335 | 4.23E-21 | 3.87E-19 | 496.3851 | 413.8243 | 453.9481 | 4571.456 | 4390.14 | 4333.947 | 763.2171 | 690.3172 | 749.2494 | 799.983 | 792.2664 | 823.2144 | 135.9953 | 140.7088 | 126.3398 |
| Kel | 23925 | 3.187098 | -2.8506 | 4.066977 | 0.000539 | 0.001612 | 1.10131 | -2.8506 | 2.056277 | 0.052188 | 0.089859 | 3.709904 | 0.888035 | 0 | 16.88322 | 16.85274 | 6.763339 | 21.31891 | 33.51054 | 32.17636 | 2.307643 | 1.525056 | 4.478048 | 22.52683 | 36.1598 | 21.05663 |
| Igkv9-124 | 243431 | 3.187175 | -5.33124 | 3.225828 | 0.003994 | 0.010381 | -3.23507 | -5.33124 | -3.16033 | 0.004653 | 0.010948 | 2.967923 | 0 | 0 | 19.48064 | 8.426372 | 2.705335 | 0 | 0 | 1.149156 | 0 | 1.525056 | 0 | 0 | 2.358248 | 3.239481 |
| Axl | 26362 | 3.19043 | 6.048205 | 52.81101 | 4.12E-24 | 5.27E-22 | 0.972356 | 6.048205 | 28.15404 | 2.30E-18 | 9.41E-17 | 1320.726 | 1215.72 | 1358.372 | 12788.39 | 12586.19 | 12598.75 | 26746.71 | 26104.71 | 24736.72 | 1488.43 | 1644.773 | 1563.585 | 1492.611 | 1444.034 | 1436.71 |
| Gm382 | 211208 | 3.191179 | -3.51274 | 3.254059 | 0.003739 | 0.009773 | 0.132375 | -3.51274 | 0.168127 | 0.868066 | 0.918721 | 0 | 5.32821 | 0 | 10.38967 | 5.617581 | 10.82134 | 4.263783 | 5.361687 | 33.32551 | 1.538429 | 5.337695 | 2.239024 | 11.68058 | 11.79124 | 18.62702 |
| Bcl2a1a | 12044 | 3.192754 | 3.226688 | 25.84267 | 1.36E-17 | 3.27E-16 | 0.319739 | 3.226688 | 4.51851 | 0.000182 | 0.000565 | 313.8579 | 296.6037 | 191.8213 | 2701.315 | 2571.448 | 2456.445 | 3366.967 | 3489.118 | 3182.012 | 636.9096 | 574.1835 | 602.2974 | 45.88798 | 44.80671 | 30.77507 |
| Ppm1k | 243382 | 3.197813 | 5.532791 | 58.18471 | 5.29E-25 | 9.09E-23 | -0.03313 | 5.532791 | -0.93772 | 0.358881 | 0.456605 | 819.8887 | 802.7836 | 851.4782 | 7853.294 | 8343.512 | 8065.958 | 8318.64 | 8228.849 | 8063.625 | 1826.884 | 1896.407 | 1820.326 | 1631.943 | 1550.941 | 1631.889 |
| Lrrc63 | 70859 | 3.199561 | 0.396208 | 9.76821 | 2.49E-09 | 1.47E-08 | 1.481959 | 0.396208 | 9.597677 | 3.39E-09 | 2.07E-08 | 25.96933 | 44.40175 | 13.88751 | 197.4038 | 276.6659 | 290.8236 | 703.5241 | 735.8915 | 738.907 | 14.61507 | 12.96297 | 26.12194 | 28.36712 | 46.37887 | 36.44416 |
| Olfr395 | 259007 | 3.201337 | -4.98593 | 2.454616 | 0.022759 | 0.049991 | 0.285798 | -4.98593 | 0.261961 | 0.795864 | 0.865889 | 0 | 0 | 0 | 1.298709 | 16.85274 | 1.352668 | 5.685043 | 2.680843 | 5.745778 | 0 | 0 | 4.478048 | 0 | 12.57732 | 4.859222 |
| Olfr350 | 258620 | 3.202119 | -4.73285 | 2.81547 | 0.010261 | 0.024557 | -0.71102 | -4.73285 | -0.71418 | 0.482853 | 0.5814 | 0 | 1.77607 | 0 | 7.792255 | 5.617581 | 5.410671 | 0 | 8.04253 | 11.49156 | 0 | 0 | 0.746341 | 11.68058 | 0.786083 | 8.908573 |
| Ceacam3 | 384557 | 3.204641 | -3.82573 | 3.664821 | 0.001414 | 0.003968 | 0.871466 | -3.82573 | 1.376673 | 0.182885 | 0.260129 | 5.193865 | 0 | 0 | 15.58451 | 7.021976 | 8.116006 | 12.79135 | 20.10633 | 25.28142 | 0 | 0.762528 | 0 | 6.674615 | 17.29382 | 12.95792 |
| Olfr1384 | 258464 | 3.210188 | -4.62416 | 2.596385 | 0.016717 | 0.037978 | 0.161583 | -4.62416 | 0.154838 | 0.878404 | 0.918721 | 0 | 0 | 0 | 10.38967 | 8.426372 | 0 | 2.842522 | 2.680843 | 8.044089 | 3.076858 | 4.575167 | 0 | 1.668654 | 7.074744 | 11.33818 |
| Olfr948 | 257912 | 3.211929 | -5.17853 | 4.167957 | 0.000422 | 0.001283 | 0.519575 | -5.17853 | 0.819243 | 0.421702 | 0.520884 | 0 | 0 | 0 | 2.597418 | 2.808791 | 5.410671 | 5.685043 | 4.021265 | 6.894934 | 0 | 0 | 0 | 1.668654 | 4.716496 | 2.429611 |
| Ddi1 | 71829 | 3.211929 | -5.44949 | 3.684732 | 0.001349 | 0.003801 | -0.52899 | -5.44949 | -0.68977 | 0.497774 | 0.595765 | 0 | 0 | 0 | 2.597418 | 2.808791 | 5.410671 | 7.106304 | 1.340422 | 1.149156 | 0 | 0 | 1.492683 | 0.834327 | 1.572165 | 0.80987 |
| Rtn1 | 104001 | 3.211987 | 1.941783 | 22.14824 | 3.27E-16 | 6.08E-15 | -3.6412 | 1.941783 | -19.2474 | 5.71E-15 | 1.03E-13 | 111.2971 | 111.0044 | 168.3861 | 1318.19 | 1296.257 | 1167.352 | 99.48826 | 83.10615 | 131.0037 | 559.9881 | 606.2096 | 638.1218 | 66.74615 | 65.24486 | 58.31066 |
| Speer4b | 73526 | 3.214293 | -3.97557 | 3.867216 | 0.000871 | 0.002527 | 0.876261 | -3.97557 | 1.361502 | 0.187556 | 0.265739 | 0 | 0 | 0.86797 | 3.896127 | 2.808791 | 12.17401 | 8.527565 | 8.04253 | 16.08818 | 2.307643 | 3.812639 | 2.239024 | 6.674615 | 3.930413 | 8.908573 |
| Serpina3h | 546546 | 3.216022 | -1.22049 | 5.579445 | 1.46E-05 | 5.29E-05 | 0.264758 | -1.22049 | 0.761011 | 0.454977 | 0.553786 | 5.193865 | 8.88035 | 1.735939 | 48.05224 | 56.17581 | 39.22736 | 75.32683 | 50.93603 | 55.15947 | 53.84501 | 32.02617 | 67.17071 | 17.52087 | 17.29382 | 12.95792 |
| Llcfc1 | 76606 | 3.216985 | -5.23032 | 2.865849 | 0.009156 | 0.022156 | -0.32142 | -5.23032 | -0.3292 | 0.745211 | 0.822208 | 0 | 0 | 0 | 5.194837 | 2.808791 | 2.705335 | 8.527565 | 0 | 5.745778 | 0 | 4.575167 | 0 | 0.834327 | 4.716496 | 0.80987 |
| Olfr345 | 258947 | 3.216985 | -5.13099 | 2.865843 | 0.009156 | 0.022156 | -0.39922 | -5.13099 | -0.40676 | 0.688234 | 0.770888 | 0 | 0 | 0 | 5.194837 | 2.808791 | 2.705335 | 7.106304 | 0 | 5.745778 | 0 | 0.762528 | 0 | 13.34923 | 0.786083 | 6.478962 |
| Vmn2r27 | 232367 | 3.217146 | -3.29304 | 2.970573 | 0.007212 | 0.017832 | 0.265496 | -3.29304 | 0.32815 | 0.745996 | 0.822891 | 10.38773 | 0 | 0 | 16.88322 | 14.04395 | 8.116006 | 21.31891 | 8.04253 | 21.83396 | 11.53822 | 0.762528 | 0 | 23.36115 | 11.00516 | 21.05663 |
| Unc93a | 381058 | 3.217585 | -1.15087 | 4.844075 | 8.31E-05 | 0.000274 | 1.342053 | -1.15087 | 3.800224 | 0.001023 | 0.002792 | 2.225942 | 2.664105 | 18.22736 | 51.94837 | 37.91867 | 67.63339 | 110.8583 | 148.7868 | 148.2411 | 15.38429 | 23.63836 | 20.15121 | 27.53279 | 7.860826 | 29.9652 |
| Peli1 | 67245 | 3.218169 | 7.026524 | 65.92104 | 3.73E-26 | 1.00E-23 | -0.20189 | 7.026524 | -6.61793 | 1.38E-06 | 5.75E-06 | 3620.124 | 4159.556 | 3719.25 | 37924.9 | 38585.76 | 37268.7 | 36020.44 | 34111.05 | 32891.13 | 3281.469 | 3138.565 | 2992.082 | 1885.579 | 1775.761 | 1861.892 |
| Irf7 | 54123 | 3.221744 | 8.717998 | 51.82489 | 1.65E-15 | 2.63E-14 | -0.16114 | 8.717998 | -4.5291 | 0.000687 | 0.001941 | 9489.192 | 9550.816 | 9229.121 | 94307.06 | 94816.34 | 92526.53 | 91296.11 | 89959.72 | 81817.58 | 20782.64 | 21014.51 | 20564.69 | 6222.41 | 6151.096 | 6448.997 |
| Nt5c3 | 107569 | 3.222177 | 5.519603 | 66.32893 | 3.27E-26 | 9.07E-24 | -1.97988 | 5.519603 | -46.8946 | 5.10E-23 | 9.72E-21 | 1607.13 | 1792.055 | 1571.025 | 16606.59 | 16859.77 | 15922.25 | 4235.357 | 4511.86 | 4251.876 | 2988.398 | 2960.896 | 3110.004 | 471.3947 | 414.2655 | 421.1325 |
| Gm19440 | 100502898 | 3.224538 | -3.30237 | 2.677124 | 0.013986 | 0.032441 | 0.078361 | -3.30237 | 0.082033 | 0.935385 | 0.946833 | 0 | 6.216245 | 0 | 12.98709 | 11.23516 | 5.410671 | 22.74017 | 4.021265 | 11.49156 | 0 | 3.812639 | 1.492683 | 76.75808 | 69.17527 | 23.48624 |
| Trim43b | 666747 | 3.226168 | -2.48365 | 4.274636 | 0.000327 | 0.001006 | 0.11342 | -2.48365 | 0.220741 | 0.827392 | 0.893649 | 0.741981 | 9.768385 | 1.735939 | 32.46773 | 25.27911 | 25.70069 | 22.74017 | 22.78717 | 52.86116 | 1.538429 | 10.67539 | 1.492683 | 10.01192 | 15.72165 | 24.29611 |
| BC051665 | 218275 | 3.226866 | -4.00486 | 3.173523 | 0.004513 | 0.01159 | 0.884329 | -4.00486 | 1.144843 | 0.264951 | 0.354973 | 0 | 1.77607 | 0 | 19.48064 | 4.213186 | 2.705335 | 21.31891 | 5.361687 | 17.23733 | 1.538429 | 0 | 2.239024 | 10.84625 | 4.716496 | 11.33818 |
| Daxx | 13163 | 3.227323 | 7.88018 | 44.88672 | 3.83E-15 | 5.66E-14 | -0.18053 | 7.88018 | -4.18774 | 0.001166 | 0.003147 | 5978.881 | 6255.318 | 5637.463 | 60279.59 | 59887.63 | 58359.5 | 57038.04 | 56268.22 | 51151.21 | 9677.487 | 10329.96 | 8911.315 | 3029.441 | 2919.511 | 2955.217 |
| Lrriq4 | 68307 | 3.229662 | -2.77215 | 3.607815 | 0.00162 | 0.004504 | 1.48509 | -2.77215 | 2.415648 | 0.024746 | 0.047386 | 0 | 0.888035 | 1.735939 | 9.090964 | 8.426372 | 10.82134 | 12.79135 | 45.57434 | 36.77298 | 3.076858 | 10.67539 | 5.224389 | 20.02385 | 39.30413 | 29.9652 |
| Tmprss2 | 50528 | 3.233986 | 0.98943 | 15.83475 | 2.83E-13 | 3.01E-12 | -1.13168 | 0.98943 | -8.19267 | 4.97E-08 | 2.55E-07 | 48.22875 | 50.61799 | 51.21021 | 481.8211 | 519.6262 | 515.3664 | 251.5632 | 247.978 | 221.787 | 71.53694 | 68.62751 | 56.72194 | 39.21337 | 64.45877 | 76.93768 |
| Dlx1as | 111970 | 3.23749 | -3.69147 | 4.487935 | 0.000195 | 0.000618 | 0.684589 | -3.69147 | 1.311529 | 0.203613 | 0.284833 | 2.225942 | 0 | 0.86797 | 12.98709 | 8.426372 | 12.17401 | 17.05513 | 20.10633 | 19.53565 | 0 | 0 | 0 | 31.70442 | 18.86598 | 15.38754 |
| Mir101a | 387143 | 3.241953 | -2.94873 | 4.076773 | 0.000526 | 0.001578 | -0.26122 | -2.94873 | -0.42615 | 0.674266 | 0.758727 | 0.741981 | 9.768385 | 0 | 24.67547 | 14.04395 | 12.17401 | 14.21261 | 17.42548 | 11.49156 | 9.230573 | 3.812639 | 4.478048 | 13.34923 | 11.79124 | 8.098703 |
| Apod | 11815 | 3.242087 | -1.66206 | 7.220866 | 3.71E-07 | 1.63E-06 | -1.39832 | -1.66206 | -4.06269 | 0.000544 | 0.001565 | 5.193865 | 7.10428 | 7.811726 | 59.74062 | 70.21976 | 81.16006 | 27.00396 | 24.12759 | 29.87805 | 13.84586 | 18.30067 | 15.67317 | 5.005962 | 17.29382 | 12.95792 |
| Tmed11 | 67366 | 3.242136 | -4.85243 | 3.627541 | 0.001546 | 0.00431 | 1.071097 | -4.85243 | 1.519875 | 0.143219 | 0.211496 | 0 | 0 | 0 | 1.298709 | 5.617581 | 5.410671 | 2.842522 | 12.0638 | 14.93902 | 0 | 0 | 0.746341 | 4.171635 | 3.930413 | 4.049351 |
| Ifi211 | 381308 | 3.242689 | 6.660065 | 39.84157 | 1.23E-18 | 3.77E-17 | -0.28316 | 6.660065 | -5.32922 | 5.01E-05 | 0.000169 | 2090.16 | 2153.485 | 1490.304 | 19809.21 | 19519.69 | 18067.58 | 16721.13 | 15508.68 | 16685.74 | 4220.68 | 4184.753 | 4069.799 | 3208.821 | 3421.032 | 3281.594 |
| Btn2a2 | 238555 | 3.242736 | -2.91083 | 3.785694 | 0.001059 | 0.003038 | 0.596445 | -2.91083 | 0.978426 | 0.338825 | 0.435742 | 0.741981 | 1.77607 | 0.86797 | 19.48064 | 5.617581 | 24.34802 | 9.948826 | 26.80843 | 37.92213 | 0.769214 | 6.862751 | 3.731706 | 25.02981 | 19.65207 | 15.38754 |
| Nlrc5 | 434341 | 3.244241 | 8.257977 | 38.2856 | 8.20E-13 | 8.06E-12 | -0.43297 | 8.257977 | -8.37104 | 5.02E-06 | 1.94E-05 | 7724.762 | 8075.79 | 7159.881 | 77369.3 | 78837.13 | 75899.54 | 61868.91 | 63617.76 | 54428.61 | 12860.5 | 13071.25 | 12419.12 | 4640.526 | 4712.565 | 4266.397 |
| Cdkl4 | 381113 | 3.244526 | 0.333243 | 15.2663 | 5.81E-13 | 5.86E-12 | -2.89252 | 0.333243 | -13.1084 | 1.11E-11 | 1.04E-10 | 37.09904 | 37.29747 | 35.58675 | 379.2231 | 397.4439 | 348.9883 | 45.48035 | 60.31898 | 50.56285 | 48.46051 | 50.32684 | 34.3317 | 65.91183 | 81.75259 | 77.74755 |
| Gpr39 | 71111 | 3.246532 | -4.18888 | 3.072242 | 0.005708 | 0.014409 | -0.42763 | -4.18888 | -0.48001 | 0.6361 | 0.725124 | 0 | 3.55214 | 0 | 19.48064 | 8.426372 | 2.705335 | 5.685043 | 5.361687 | 6.894934 | 3.846072 | 5.337695 | 0.746341 | 11.68058 | 0.786083 | 0.80987 |
| Efs | 13644 | 3.249198 | -1.67246 | 5.309669 | 2.75E-05 | 9.64E-05 | 0.88958 | -1.67246 | 2.422494 | 0.024386 | 0.046781 | 2.967923 | 1.77607 | 3.471878 | 54.54578 | 19.66153 | 25.70069 | 56.85043 | 75.06362 | 50.56285 | 8.461359 | 5.337695 | 2.985365 | 78.42673 | 65.24486 | 73.69819 |
| Marcksl1 | 17357 | 3.249891 | 8.879553 | 116.9087 | 1.89E-31 | 1.24E-27 | -0.15395 | 8.879553 | -10.3778 | 8.48E-10 | 5.69E-09 | 15591.24 | 15975.75 | 14535.02 | 158208.8 | 157167.3 | 152681 | 148081.2 | 145411.6 | 143829.5 | 15112.76 | 15127.03 | 15297.01 | 3327.296 | 3273.248 | 3410.364 |
| Olfr273 | 258821 | 3.251609 | -4.72197 | 3.67408 | 0.001383 | 0.00389 | 1.033003 | -4.72197 | 1.478231 | 0.153953 | 0.225176 | 0 | 0 | 0 | 5.194837 | 5.617581 | 1.352668 | 5.685043 | 6.702109 | 13.78987 | 0 | 1.525056 | 0 | 3.337308 | 6.288661 | 8.098703 |
| Olfr1440 | 258679 | 3.253405 | -4.83913 | 3.984783 | 0.000657 | 0.001942 | -0.43454 | -4.83913 | -0.62224 | 0.540375 | 0.635617 | 0 | 0.888035 | 0 | 6.493546 | 8.426372 | 2.705335 | 2.842522 | 9.382952 | 2.298311 | 0 | 0 | 0 | 7.508942 | 6.288661 | 4.049351 |
| Spaca3 | 75622 | 3.254344 | -4.35529 | 3.991831 | 0.000646 | 0.001911 | 2.098702 | -4.35529 | 3.570531 | 0.001771 | 0.00459 | 0 | 0 | 0 | 3.896127 | 4.213186 | 2.705335 | 11.37009 | 22.78717 | 21.83396 | 0.769214 | 0 | 0 | 13.34923 | 14.14949 | 8.098703 |

| | | | | | | | | | | | | | | | | | | | | | | | | | | |
|---|---|---|---|---|---|---|---|---|---|---|---|---|---|---|---|---|---|---|---|---|---|---|---|---|---|---|
| Vmn1r220 | 171271 | 3.255862 | -5.47352 | 3.019321 | 0.006448 | 0.016103 | -0.82475 | -5.47352 | -0.85199 | 0.403678 | 0.502814 | 0 | 0 | 0 | 5.194837 | 1.404395 | 5.410671 | 1.421261 | 0 | 9.193245 | 1.538429 | 0 | 0 | 0 | 2.358248 | 3.239481 |
| Mov10l1 | 83456 | 3.25758 | -2.22453 | 4.275559 | 0.000326 | 0.001004 | 0.852627 | -2.22453 | 1.730894 | 0.097904 | 0.153108 | 0 | 2.664105 | 8.679696 | 19.48064 | 18.25714 | 25.70069 | 45.48035 | 33.51054 | 41.3696 | 7.692145 | 6.862751 | 2.985365 | 25.02981 | 35.37372 | 31.58494 |
| Mir330 | 724063 | 3.25889 | -2.17239 | 5.755596 | 9.70E-06 | 3.59E-05 | -1.70719 | -2.17239 | -3.66172 | 0.001425 | 0.003774 | 1.483961 | 12.43249 | 10.41564 | 85.7148 | 44.94065 | 59.51738 | 27.00396 | 20.10633 | 13.78987 | 11.53822 | 12.20045 | 5.97073 | 5.005962 | 3.930413 | 5.669092 |
| Pde11a | 241489 | 3.259266 | -1.52406 | 6.077628 | 4.63E-06 | 1.78E-05 | 0.507216 | -1.52406 | 1.571389 | 0.1308 | 0.195989 | 2.225942 | 2.664105 | 6.943757 | 35.06515 | 46.34504 | 36.52203 | 65.378 | 44.23392 | 65.50187 | 9.230573 | 7.625279 | 11.94146 | 55.8999 | 38.51805 | 38.0639 |
| Fam107a | 268709 | 3.260151 | -3.39773 | 3.545178 | 0.001881 | 0.005169 | 0.287545 | -3.39773 | 0.41747 | 0.680504 | 0.763988 | 0 | 2.664105 | 1.735939 | 18.18193 | 8.426372 | 13.52668 | 15.63387 | 6.702109 | 36.77298 | 0 | 1.525056 | 3.731706 | 15.85221 | 8.646909 | 16.19741 |
| Mtmr7 | 54384 | 3.260905 | 3.341399 | 22.70391 | 1.97E-16 | 3.83E-15 | 1.163359 | 3.341399 | 16.53606 | 1.20E-13 | 1.63E-12 | 118.7169 | 126.989 | 114.572 | 1174.033 | 1268.169 | 1253.923 | 2794.199 | 3029.353 | 2774.062 | 123.8435 | 123.5295 | 182.8536 | 1375.805 | 1348.918 | 1354.103 |
| Hnf4a | 15378 | 3.26698 | -2.59104 | 3.610382 | 0.00161 | 0.004478 | 2.073778 | -2.59104 | 3.547666 | 0.00187 | 0.004826 | 4.451884 | 0 | 0 | 3.896127 | 16.85274 | 13.52668 | 36.95278 | 45.57434 | 50.56285 | 12.30743 | 3.050111 | 5.224389 | 26.69846 | 39.30413 | 35.63429 |
| Arhgap28 | 268970 | 3.267965 | 1.842463 | 11.1498 | 2.30E-10 | 1.57E-09 | 0.211213 | 1.842463 | 1.366254 | 0.186083 | 0.263922 | 44.51884 | 31.96926 | 41.66254 | 379.2231 | 448.0021 | 401.7423 | 528.709 | 411.5095 | 535.5065 | 178.4578 | 211.2202 | 144.7902 | 259.4757 | 396.1856 | 332.8567 |
| Dpp10 | 269109 | 3.26838 | -1.92473 | 3.858032 | 0.00089 | 0.002581 | 0.854237 | -1.92473 | 1.639934 | 0.115676 | 0.176555 | 6.677827 | 5.32821 | 0 | 57.1432 | 21.06593 | 14.87935 | 76.74809 | 38.87223 | 44.81707 | 13.84586 | 8.387806 | 19.40487 | 14.18356 | 28.29897 | 17.81715 |
| Inhba | 16323 | 3.273403 | 2.955255 | 11.34311 | 1.68E-10 | 1.16E-09 | 4.226452 | 2.955255 | 35.5271 | 1.78E-20 | 1.37E-18 | 26.71131 | 35.5214 | 40.79457 | 345.4566 | 344.0768 | 357.1043 | 7117.674 | 6846.874 | 6472.044 | 122.3051 | 74.72773 | 64.93169 | 3059.477 | 3036.637 | 3245.15 |
| Cd86 | 12524 | 3.276954 | 5.550128 | 50.58906 | 1.03E-23 | 1.11E-21 | 0.805489 | 5.550128 | 21.72241 | 4.87E-16 | 1.14E-14 | 864.4076 | 774.3665 | 692.6397 | 7827.32 | 8214.308 | 8132.238 | 14690.15 | 14425.62 | 14693.1 | 1740.732 | 1685.949 | 1778.531 | 1008.701 | 1114.665 | 1150.826 |
| Gcg | 14526 | 3.277373 | -3.27007 | 3.5235 | 0.00198 | 0.005423 | 0.52288 | -3.27007 | 0.773983 | 0.44743 | 0.546148 | 0.741981 | 3.55214 | 0 | 18.18193 | 7.021976 | 13.52668 | 21.31891 | 17.42548 | 17.23733 | 4.615287 | 6.100223 | 0 | 12.5149 | 8.646909 | 18.62702 |
| Vmn2r59 | 628444 | 3.288727 | -4.35098 | 2.850488 | 0.00948 | 0.022873 | 2.071615 | -4.35098 | 2.48964 | 0.021101 | 0.041321 | 0 | 0 | 0 | 10.38967 | 0 | 9.468674 | 11.37009 | 16.08506 | 29.87805 | 1.538429 | 0 | 1.492683 | 2.502981 | 4.716496 | 13.76779 |
| Gm10440 | 330086 | 3.289061 | -3.09385 | 4.358332 | 0.000267 | 0.000831 | 0.755179 | -3.09385 | 1.429717 | 0.167274 | 0.241239 | 2.967923 | 0 | 0.86797 | 11.68838 | 12.63956 | 13.52668 | 27.00396 | 24.12759 | 18.38649 | 2.307643 | 3.050111 | 1.492683 | 16.68654 | 26.72681 | 9.718443 |
| Nme8 | 73412 | 3.289865 | -2.18064 | 5.288725 | 2.89E-05 | 0.000101 | -1.65097 | -2.18064 | -3.13015 | 0.004991 | 0.011666 | 9.64575 | 3.55214 | 0.86797 | 50.64966 | 30.8967 | 37.8747 | 5.685043 | 14.74464 | 20.6848 | 10.769 | 10.67539 | 5.97073 | 18.35519 | 23.58248 | 25.91585 |
| Tsix | 22097 | 3.29021 | -2.47256 | 3.65715 | 0.00144 | 0.004037 | 0.394381 | -2.47256 | 0.609239 | 0.548797 | 0.643624 | 0.741981 | 0 | 6.075787 | 23.37676 | 14.04395 | 9.468674 | 17.05513 | 14.74464 | 31.0272 | 19.99958 | 7.625279 | 2.985365 | 27.53279 | 47.16496 | 41.30338 |
| Gldn | 235379 | 3.290352 | -2.89182 | 3.991337 | 0.000646 | 0.001913 | 1.138962 | -2.89182 | 2.116014 | 0.04626 | 0.081006 | 0.741981 | 0.888035 | 2.603909 | 19.48064 | 7.021976 | 31.11136 | 21.31891 | 33.51054 | 70.09849 | 0.769214 | 0.762528 | 2.985365 | 15.01788 | 25.94073 | 17.81715 |
| Ly6h | 23934 | 3.293589 | -3.11099 | 3.329649 | 0.003131 | 0.008286 | 0.911591 | -3.11099 | 1.301144 | 0.207081 | 0.288783 | 0 | 2.664105 | 0.86797 | 12.98709 | 8.426372 | 12.17401 | 25.5827 | 10.72337 | 36.77298 | 4.615287 | 0.762528 | 6.717071 | 32.53875 | 15.72165 | 6.478962 |
| F630111L1 | 320463 | 3.298481 | 3.165223 | 13.89687 | 3.62E-12 | 3.20E-11 | 2.840854 | 3.165223 | 28.27237 | 2.11E-18 | 8.71E-17 | 51.93865 | 37.29747 | 46.87036 | 493.5095 | 419.9142 | 531.5984 | 3637.007 | 3631.203 | 3469.301 | 323.0701 | 390.4143 | 292.5658 | 1707.867 | 1760.039 | 1651.325 |
| Rtl3 | 213436 | 3.308344 | -3.45696 | 3.222244 | 0.004028 | 0.01046 | 1.48174 | -3.45696 | 2.03986 | 0.053932 | 0.092481 | 0 | 0 | 1.735939 | 18.18193 | 5.617581 | 2.705335 | 17.05513 | 29.48928 | 18.38649 | 2.307643 | 6.100223 | 0 | 19.18952 | 18.86598 | 29.9652 |
| Gm15433 | 100503433 | 3.309702 | -1.63708 | 6.473423 | 1.90E-06 | 7.60E-06 | -0.30123 | -1.63708 | -0.93827 | 0.358605 | 0.456297 | 9.64575 | 5.32821 | 2.603909 | 54.54578 | 58.9846 | 66.28072 | 41.21656 | 48.25518 | 60.90525 | 13.07665 | 20.58825 | 19.40487 | 5.005962 | 17.29382 | 6.478962 |
| A1cf | 69865 | 3.311447 | -0.26404 | 8.376015 | 3.45E-08 | 1.73E-07 | 0.775979 | -0.26404 | 3.821317 | 0.000972 | 0.002667 | 6.677827 | 10.65642 | 12.15157 | 107.7929 | 110.9472 | 91.98141 | 184.7639 | 186.3186 | 181.5666 | 55.38344 | 45.75167 | 29.10731 | 39.21337 | 40.09021 | 33.20468 |
| Plppr5 | 75769 | 3.31376 | -2.73897 | 3.764777 | 0.001113 | 0.003182 | 0.587778 | -2.73897 | 0.995841 | 0.330484 | 0.426814 | 0 | 1.77607 | 8.679696 | 19.48064 | 15.44835 | 24.34802 | 39.7953 | 22.78717 | 32.17636 | 1.538429 | 0.762528 | 11.94146 | 14.18356 | 10.21907 | 23.48624 |
| Ceacam13 | 69785 | 3.315553 | -4.98698 | 3.528519 | 0.001957 | 0.005364 | -1.31664 | -4.98698 | -1.54417 | 0.137246 | 0.204077 | 0 | 0.888035 | 0 | 3.896127 | 5.617581 | 8.116006 | 0 | 5.361687 | 3.447467 | 2.307643 | 0.762528 | 0 | 2.502981 | 0.786083 | 3.239481 |
| Hspa1a | 193740 | 3.315617 | 4.398829 | 32.07864 | 1.51E-19 | 5.72E-18 | 1.664844 | 4.398829 | 32.48959 | 1.16E-19 | 6.95E-18 | 276.0168 | 299.2678 | 239.5596 | 2790.926 | 2935.186 | 2919.057 | 9506.814 | 9577.313 | 9364.469 | 352.3002 | 383.5515 | 328.3902 | 1108.82 | 1211.353 | 1333.856 |
| Ptgds | 19215 | 3.31643 | -4.29394 | 3.41456 | 0.002563 | 0.006882 | -0.07083 | -4.29394 | -0.08787 | 0.930796 | 0.943495 | 0 | 0.888035 | 0 | 5.194837 | 8.426372 | 4.058003 | 9.948826 | 1.340422 | 10.3424 | 0.769214 | 0 | 2.239024 | 5.005962 | 14.93557 | 14.57766 |
| Pde4b | 18578 | 3.316501 | 6.48988 | 67.41168 | 2.32E-26 | 6.94E-24 | -0.50788 | 6.48988 | -15.4828 | 4.41E-13 | 5.29E-12 | 1645.713 | 1817.808 | 1681.257 | 17920.89 | 18925.63 | 17782.17 | 13629.89 | 13440.41 | 12854.45 | 4017.607 | 4023.859 | 3927.248 | 2674.852 | 2621.586 | 2813.489 |
| Try4 | 22074 | 3.318799 | -4.82947 | 3.93282 | 0.000744 | 0.002181 | -0.97134 | -4.82947 | -1.31349 | 0.202964 | 0.284126 | 1.483961 | 0 | 0 | 10.38967 | 2.808791 | 10.82134 | 2.842522 | 4.021265 | 3.447467 | 0.769214 | 0 | 0 | 6.674615 | 1.572165 | 2.429611 |
| Rab11fip1 | 75767 | 3.32243 | 5.90989 | 62.16686 | 1.30E-25 | 2.75E-23 | -0.57453 | 5.90989 | -15.9315 | 2.51E-13 | 3.17E-12 | 1504.737 | 1271.666 | 1495.512 | 15310.48 | 15187.13 | 15143.12 | 10855.59 | 10583.97 | 10409.05 | 1731.502 | 1900.219 | 1859.136 | 1501.788 | 1363.853 | 1423.752 |
| Armcx6 | 278097 | 3.329668 | 2.037868 | 12.23745 | 4.09E-11 | 3.07E-10 | 2.4285 | 2.037868 | 20.83031 | 1.15E-15 | 2.43E-14 | 36.35706 | 33.74533 | 47.73833 | 474.0288 | 467.6636 | 332.7563 | 2401.931 | 2556.184 | 2138.579 | 82.30595 | 88.45323 | 88.06827 | 270.3219 | 250.7604 | 271.3065 |
| Gm19359 | 100502768 | 3.329913 | -3.54422 | 4.003086 | 0.000628 | 0.001863 | -0.01398 | -3.54422 | -0.02227 | 0.982441 | 0.985251 | 3.709904 | 0 | 0.86797 | 14.2858 | 14.04395 | 13.52668 | 22.74017 | 8.04253 | 16.08818 | 0 | 0 | 2.985365 | 15.85221 | 9.432991 | 21.8665 |
| Pla2g10 | 26565 | 3.331536 | -3.29702 | 3.863322 | 0.000879 | 0.00255 | -1.38478 | -3.29702 | -1.87698 | 0.07426 | 0.121213 | 3.709904 | 5.32821 | 0 | 33.76644 | 18.25714 | 18.93735 | 5.685043 | 10.72337 | 10.3424 | 4.615287 | 0 | 4.478048 | 8.343269 | 14.93557 | 4.049351 |
| Atp10a | 11982 | 3.333798 | 5.260659 | 61.86618 | 1.44E-25 | 3.02E-23 | -0.57489 | 5.260659 | -15.8013 | 2.95E-13 | 3.66E-12 | 1098.873 | 1158.886 | 1087.566 | 12089.68 | 12104.48 | 11789.85 | 8457.923 | 8743.571 | 7947.56 | 1534.583 | 1528.868 | 1559.853 | 351.2516 | 379.6779 | 366.0614 |
| Mettl21e | 403183 | 3.33517 | -3.54291 | 4.154678 | 0.000436 | 0.001321 | 0.985444 | -3.54291 | 1.668093 | 0.109904 | 0.169233 | 1.483961 | 0 | 0 | 7.792255 | 8.426372 | 5.410671 | 9.948826 | 20.10633 | 17.23733 | 5.384501 | 3.050111 | 2.239024 | 9.177596 | 17.29382 | 7.288832 |
| Nfil3 | 18030 | 3.336683 | 5.281857 | 37.88835 | 4.59E-21 | 2.50E-19 | -0.17281 | 5.281857 | -3.19317 | 0.004311 | 0.010233 | 420.7031 | 440.4654 | 401.8699 | 4636.392 | 4408.397 | 4574.722 | 4360.428 | 4325.541 | 3911.726 | 4022.222 | 3799.676 | 4026.511 | 2092.492 | 2052.462 | 2305.701 |
| Olfr1360 | 258536 | 3.338222 | -5.29587 | 3.732078 | 0.001204 | 0.003422 | 0.246769 | -5.29587 | 0.332183 | 0.742991 | 0.820344 | 0 | 0 | 0 | 1.298709 | 5.617581 | 6.763339 | 1.421261 | 5.361687 | 11.49156 | 0 | 0 | 0 | 2.502981 | 2.358248 | 0.80987 |
| Gm19684 | 100503422 | 3.341276 | 2.488077 | 15.47109 | 4.47E-13 | 4.61E-12 | 1.112184 | 2.488077 | 10.91658 | 3.39E-10 | 2.42E-09 | 89.03769 | 77.25904 | 78.11726 | 938.9667 | 790.6745 | 938.7514 | 2084.99 | 2246.547 | 1726.032 | 175.3809 | 195.2071 | 147.7756 | 157.6878 | 182.3712 | 167.6431 |
| Fcgr2b | 14130 | 3.343038 | 7.665492 | 71.4617 | 6.72E-27 | 2.29E-24 | -0.67171 | 7.665492 | -21.3205 | 7.13E-16 | 1.59E-14 | 2180.681 | 2391.478 | 2120.45 | 23301.44 | 24856.39 | 24253.33 | 16094.36 | 15812.96 | 15305.6 | 18964.21 | 18756.66 | 19113.05 | 16019.08 | 16492.8 | 16761.08 |
| Sfta3-ps | 104798 | 3.345343 | -4.03828 | 3.940844 | 0.00073 | 0.002142 | 1.512576 | -4.03828 | 2.534415 | 0.019145 | 0.037997 | 0 | 0 | 1.735939 | 3.896127 | 7.021976 | 12.17401 | 12.79135 | 26.80843 | 28.72889 | 0 | 0 | 0 | 8.343269 | 17.29382 | 17.00728 |
| Tuba8 | 53857 | 3.347463 | 2.008105 | 16.37441 | 1.46E-13 | 1.63E-12 | -0.18976 | 2.008105 | -1.69329 | 0.104948 | 0.162749 | 76.42402 | 65.71459 | 78.11726 | 776.6281 | 714.8372 | 927.9301 | 778.8509 | 768.0617 | 661.9136 | 377.6843 | 482.6801 | 384.3657 | 44.21933 | 51.09537 | 38.87377 |
| Foxl2os | 768252 | 3.347762 | -2.32582 | 3.903327 | 0.000799 | 0.002329 | 0.743619 | -2.32582 | 1.319505 | 0.200981 | 0.281709 | 8.161788 | 0 | 0.86797 | 14.2858 | 18.25714 | 24.34802 | 21.31891 | 49.5956 | 32.17636 | 6.153716 | 1.525056 | 6.717071 | 60.90587 | 57.38403 | 34.82442 |
| F830208F2 | 619308 | 3.349225 | 1.598927 | 16.51857 | 1.22E-13 | 1.39E-12 | -0.82917 | 1.598927 | -6.70255 | 1.14E-06 | 4.82E-06 | 106.1032 | 65.71459 | 78.11726 | 936.3693 | 915.6657 | 865.7074 | 594.087 | 569.6792 | 451.6181 | 70.76773 | 109.804 | 55.97559 | 68.41481 | 73.10568 | 51.8317 |
| Xkr8 | 381560 | 3.355922 | 4.699222 | 47.75622 | 3.47E-23 | 3.27E-21 | -0.62837 | 4.699222 | -13.2665 | 8.84E-12 | 8.37E-11 | 736.7869 | 768.1503 | 867.1016 | 8957.197 | 8603.325 | 8252.626 | 6020.461 | 5927.345 | 5449.296 | 851.5204 | 897.4953 | 806.7949 | 290.3458 | 262.5516 | 255.919 |
| Tmem229 | 432460 | 3.35624 | -3.27813 | 4.677605 | 0.000124 | 0.000401 | -0.52234 | -3.27813 | -0.93186 | 0.361831 | 0.459647 | 2.225942 | 3.55214 | 0 | 15.58451 | 14.04395 | 24.34802 | 14.21261 | 12.0638 | 11.49156 | 4.615287 | 2.287584 | 1.492683 | 6.674615 | 11.79124 | 4.859222 |
| D030046N | 433208 | 3.358466 | -4.17478 | 2.583835 | 0.017184 | 0.038928 | 0.903817 | -4.17478 | 0.913384 | 0.371244 | 0.469359 | 0 | 0 | 0.86797 | 15.58451 | 2.808791 | 4.058003 | 5.685043 | 8.04253 | 31.0272 | 1.538429 | 1.525056 | 0 | 16.68654 | 0.786083 | 21.8665 |
| Dusp1 | 19252 | 3.360941 | 6.577612 | 85.44541 | 1.50E-28 | 1.24E-25 | -2.50873 | 6.577612 | -68.5832 | 1.61E-26 | 1.32E-23 | 3258.779 | 3275.073 | 3208.884 | 35908.01 | 35341.61 | 35634.68 | 7076.458 | 6010.451 | 6470.895 | 2616.098 | 2511.767 | 2627.868 | 3178.786 | 3035.851 | 3136.628 |
| Gm10757 | 100038551 | 3.37085 | -4.75697 | 3.984656 | 0.000657 | 0.001942 | 0.893993 | -4.75697 | 1.347371 | 0.191992 | 0.270867 | 0 | 0 | 0 | 2.597418 | 5.617581 | 4.058003 | 18.47639 | 5.361687 | 5.745778 | 0 | 0 | 0 | 4.171635 | 11.00516 | 9.718443 |
| Reg3g | 19695 | 3.371831 | -5.02465 | 2.962277 | 0.00735 | 0.01814 | -0.12212 | -5.02465 | -0.12633 | 0.900653 | 0.922972 | 0 | 0 | 0 | 2.597418 | 4.213186 | 5.410671 | 5.685043 | 0 | 17.23733 | 0.769214 | 0.762528 | 0 | 0.834327 | 9.432991 | 2.429611 |
| Slfn1 | 20555 | 3.372544 | 4.495862 | 36.00154 | 1.34E-20 | 6.64E-19 | 0.085518 | 4.495862 | 1.517247 | 0.143878 | 0.212407 | 520.8705 | 531.933 | 590.2193 | 5983.153 | 6214.449 | 5928.743 | 6804.997 | 7114.959 | 6144.535 | 1039.209 | 1035.513 | 1016.517 | 198.5698 | 149.3557 | 252.6795 |
| Htr7 | 15566 | 3.373358 | -0.66404 | 5.019947 | 5.46E-05 | 0.000185 | 0.800777 | -0.66404 | 2.213452 | 0.037883 | 0.068357 | 8.903769 | 1.77607 | 5.207818 | 59.74062 | 64.60218 | 41.9327 | 123.6497 | 76.40404 | 103.424 | 31.53779 | 53.37695 | 43.28779 | 38.37904 | 92.75775 | 84.22651 |
| 2410017I1 | 675325 | 3.374611 | -3.94542 | 2.660411 | 0.014514 | 0.033539 | -0.35861 | -3.94542 | -0.34179 | 0.735853 | 0.813898 | 0 | 0 | 3.471878 | 14.2858 | 5.617581 | 8.116006 | 24.16143 | 0 | 24.13227 | 3.076858 | 0 | 0 | 14.18356 | 28.29897 | 14.57766 |
| Nfkb2 | 18034 | 3.378395 | 7.105479 | 54.5029 | 1.00E-23 | 1.09E-21 | 0.057568 | 7.105479 | 1.578217 | 0.129773 | 0.194589 | 2694.132 | 3033.527 | 2855.62 | 32307.99 | 31850.28 | 30794.83 | 36499.4 | 34778.58 | 31937.33 | 6995.236 | 6929.853 | 6929.779 | 1739.572 | 1760.825 | 1746.89 |
| Tcl1b1 | 27379 | 3.379841 | -4.23165 | 3.606624 | 0.001625 | 0.004515 | -0.4277 | -4.23165 | -0.54448 | 0.591756 | 0.684965 | 0 | 1.77607 | 0 | 5.194837 | 5.617581 | 12.17401 | 5.685043 | 8.04253 | 3.447467 | 4.615287 | 3.050111 | 0 | 5.840288 | 4.716496 | 4.049351 |
| Zfp957 | 105590 | 3.380858 | -2.95126 | 4.185784 | 0.000405 | 0.001232 | 0.778033 | -2.95126 | 1.414553 | 0.171623 | 0.246484 | 1.483961 | 0 | 2.603909 | 12.98709 | 19.66153 | 12.17401 | 29.84648 | 18.7659 | 32.17636 | 9.230573 | 0.762528 | 4.478048 | 10.01192 | 18.0799 | 11.33818 |
| Fas | 14102 | 3.381113 | 4.6783 | 33.22092 | 7.26E-20 | 2.93E-18 | 1.635619 | 4.6783 | 32.1914 | 1.40E-19 | 8.18E-18 | 362.0866 | 350.7738 | 328.9605 | 3674.048 | 4006.74 | 3952.495 | 12838.25 | 12892.18 | 11863.88 | 926.1342 | 949.3472 | 1088.912 | 518.117 | 476.3661 | 520.7466 |
| Erich2 | 66748 | 3.382214 | -2.92997 | 5.570821 | 1.49E-05 | 5.39E-05 | -0.52239 | -2.92997 | -1.2067 | 0.240754 | 0.327543 | 3.709904 | 4.440175 | 0.86797 | 24.67547 | 32.30109 | 43.28537 | 12.79135 | 28.14886 | 32.17636 | 0.769214 | 0 | 0 | 22.52683 | 22.01031 | 25.10598 |
| Ccer2 | 100504112 | 3.382485 | -2.14278 | 4.161413 | 0.000429 | 0.001301 | 0.413718 | -2.14278 | 0.856868 | 0.401037 | 0.500174 | 0.741981 | 6.216245 | 8.679696 | 54.54578 | 26.68351 | 54.10671 | 35.53152 | 87.12741 | 66.65102 | 5.384501 | 7.625279 | 0 | 19.18952 | 11.79124 | 18.62702 |
| Il1bos | 329514 | 3.384181 | -0.41895 | 10.25669 | 1.05E-09 | 6.47E-09 | 0.502288 | -0.41895 | 3.031174 | 0.006275 | 0.014293 | 20.77546 | 10.65642 | 13.88751 | 153.2477 | 183.9758 | 173.1415 | 237.3506 | 285.5098 | 230.9803 | 18.46115 | 12.96297 | 17.16585 | 20.85817 | 18.0799 | 12.14805 |
| Bsnd | 140475 | 3.386243 | -3.20403 | 3.39965 | 0.002655 | 0.007111 | 0.478081 | -3.20403 | 0.637707 | 0.530447 | 0.626377 | 0 | 0 | 4.339848 | 14.2858 | 2.808791 | 18.93735 | 9.948826 | 9.382952 | 26.43058 | 1.538429 | 6.100223 | 4.478048 | 20.85817 | 14.93557 | 32.39481 |
| Sct | 20287 | 3.388106 | -3.57909 | 3.655184 | 0.001447 | 0.004053 | 4.742978 | -3.57909 | 8.239366 | 4.53E-08 | 2.33E-07 | 0 | 0 | 0 | 2.597418 | 2.808791 | 8.116006 | 152.0749 | 108.5742 | 128.7054 | 0 | 3.050111 | 0 | 11.68058 | 19.65207 | 16.19741 |
| Dnase2b | 56629 | 3.38917 | -3.39392 | 3.725863 | 0.001222 | 0.00347 | 1.01413 | -3.39392 | 1.601021 | 0.124071 | 0.187346 | 4.451884 | 0 | 0 | 7.792255 | 11.23516 | 13.52668 | 17.05513 | 17.42548 | 36.77298 | 1.538429 | 0 | 2.985365 | 21.6925 | 11.79124 | 19.43689 |
| Pfkfb3 | 170768 | 3.394824 | 7.468683 | 112.6537 | 4.16E-31 | 1.56E-27 | -2.47536 | 7.468683 | -90.4717 | 4.44E-29 | 2.34E-25 | 7205.375 | 7425.749 | 7010.59 | 81771.92 | 81161.41 | 79987.3 | 15386.57 | 15685.62 | 14411.56 | 5827.569 | 6059.809 | 6099.847 | 2526.342 | 2444.717 | 2554.331 |
| Ccdc173 | 75051 | 3.396135 | 0.728162 | 11.38856 | 1.56E-10 | 1.08E-09 | -1.04947 | 0.728162 | -5.46057 | 1.93E-05 | 6.87E-05 | 21.51744 | 25.75301 | 40.79457 | 249.3522 | 367.9516 | 355.7516 | 174.8151 | 154.1485 | 151.6885 | 109.2285 | 99.12862 | 114.1902 | 83.43269 | 51.88145 | 48.59222 |
| Scgb3a2 | 117158 | 3.396579 | -5.30846 | 3.941752 | 0.000728 | 0.002138 | -1.06059 | -5.30846 | -1.36791 | 0.185571 | 0.263352 | 0 | 0 | 0 | 7.792255 | 2.808791 | 2.705335 | 1.421261 | 1.340422 | 2.298311 | 0 | 0 | 1.492683 | 0.834327 | 3.14433 | 4.049351 |
| Gm11565 | 670550 | 3.39894 | -4.54136 | 3.396929 | 0.002672 | 0.007151 | -1.48855 | -4.54136 | -1.64954 | 0.113679 | 0.174042 | 0 | 0 | 4.339848 | 6.493546 | 11.23516 | 12.17401 | 0 | 10.72337 | 5.745778 | 0 | 0 | 1.492683 | 7.508942 | 4.716496 | 2.429611 |
| Slc18a3 | 20508 | 3.401439 | -2.57948 | 3.852151 | 0.000903 | 0.002615 | 1.272775 | -2.57948 | 2.229424 | 0.03665 | 0.066473 | 1.483961 | 2.664105 | 0 | 10.38967 | 11.23516 | 27.05335 | 48.32287 | 18.7659 | 56.30862 | 3.076858 | 1.525056 | 6.717071 | 30.03577 | 50.30929 | 27.53559 |
| Ddn | 13199 | 3.403428 | -0.44576 | 10.04792 | 1.51E-09 | 9.16E-09 | -1.59638 | -0.44576 | -6.42927 | 2.09E-06 | 8.53E-06 | 17.80754 | 27.52908 | 12.15157 | 171.4296 | 240.1516 | 221.8375 | 63.95674 | 65.68067 | 82.7392 | 11.53822 | 12.96297 | 3.731706 | 69.24913 | 73.10568 | 60.74027 |
| Gm7030 | 630294 | 3.409883 | -0.69443 | 8.114408 | 5.82E-08 | 2.83E-07 | 1.188037 | -0.69443 | 5.965006 | 5.99E-06 | 2.29E-05 | 10.38773 | 5.32821 | 15.62345 | 125.9748 | 103.9252 | 110.9188 | 292.7797 | 306.9566 | 225.2345 | 10.769 | 14.48803 | 16.41951 | 15.01788 | 11.79124 | 20.24676 |
| Bpifa5 | 67135 | 3.412191 | -4.38954 | 3.74764 | 0.00116 | 0.003303 | -0.15667 | -4.38954 | -0.20319 | 0.840914 | 0.905316 | 0 | 0 | 0 | 1.298709 | 2.808791 | 14.87935 | 1.421261 | 5.361687 | 5.745778 | 2.307643 | 1.525056 | 1.492683 | 11.68058 | 7.074744 | 12.14805 |
| Orm1 | 18405 | 3.413225 | -2.02053 | 5.222725 | 3.38E-05 | 0.000117 | -0.51274 | -2.02053 | -1.17311 | 0.253678 | 0.342205 | 1.483961 | 10.65642 | 3.471878 | 54.54578 | 50.55823 | 37.8747 | 22.74017 | 40.21265 | 41.3696 | 11.53822 | 9.150334 | 11.19512 | 13.34923 | 18.0799 | 2.429611 |
| Tex16 | 83556 | 3.420341 | -3.35438 | 3.515487 | 0.002018 | 0.00552 | 0.747913 | -3.35438 | 1.072031 | 0.295679 | 0.389261 | 0 | 0 | 5.207818 | 7.792255 | 5.617581 | 24.34802 | 18.47639 | 18.7659 | 18.38649 | 0 | 1.525056 | 5.97073 | 20.85817 | 17.29382 | 20.24676 |
| Olfr513 | 258718 | 3.421636 | -4.80268 | 3.246719 | 0.003804 | 0.009927 | -1.40032 | -4.80268 | -1.4732 | 0.155292 | 0.226908 | 0 | 1.77607 | 0 | 6.493546 | 11.23516 | 5.410671 | 2.842522 | 0 | 11.49156 | 0 | 3.050111 | 0 | 1.668654 | 7.860826 | 1.619741 |
| Olfr106-ps | 257925 | 3.426257 | -4.88764 | 3.649291 | 0.001468 | 0.004107 | -1.74361 | -4.88764 | -2.00963 | 0.057281 | 0.097316 | 0.741981 | 0 | 0 | 2.597418 | 8.426372 | 10.82134 | 0 | 4.021265 | 2.298311 | 1.538429 | 1.525056 | 0 | 6.674615 | 1.572165 | 2.429611 |
| Gm13075 | 100503826 | 3.427086 | -4.26386 | 3.7875 | 0.001055 | 0.003025 | 0.201929 | -4.26386 | 0.278592 | 0.78324 | 0.854913 | 0 | 0 | 0.86797 | 1.298709 | 14.04395 | 10.82134 | 8.527565 | 4.021265 | 11.49156 | 1.538429 | 0.762528 | 0.746341 | 5.840288 | 4.716496 | 8.098703 |
| Hcrt | 15171 | 3.430358 | -0.80679 | 7.305004 | 3.10E-07 | 1.37E-06 | -0.76816 | -0.80679 | -2.62098 | 0.015836 | 0.032215 | 5.935846 | 11.54445 | 26.03909 | 151.949 | 174.145 | 104.1554 | 110.8583 | 69.70193 | 81.59005 | 16.1535 | 8.387806 | 7.463413 | 16.68654 | 32.22939 | 43.73299 |
| Mdfic2 | 330390 | 3.431162 | -4.60409 | 3.971698 | 0.000678 | 0.001999 | 0.64208 | -4.60409 | 0.929368 | 0.363091 | 0.460869 | 0 | 0 | 0 | 7.792255 | 5.617581 | 1.352668 | 4.263783 | 5.361687 | 13.78987 | 0.769214 | 0.762528 | 0 | 5.840288 | 7.860826 | 8.098703 |
| Rhox3d-ps | 546295 | 3.445412 | -4.26527 | 3.29847 | 0.003369 | 0.008869 | 0.109725 | -4.26527 | 0.133831 | 0.89479 | 0.920407 | 0 | 3.55214 | 0 | 12.98709 | 2.808791 | 18.93735 | 5.685043 | 5.361687 | 29.87805 | 0 | 3.050111 | 0 | 3.337308 | 3.14433 | 4.859222 |
| Jak2 | 16452 | 3.447689 | 7.805632 | 78.44841 | 9.24E-28 | 4.02E-25 | -0.34486 | 7.805632 | -13.0511 | 1.21E-11 | 1.12E-10 | 3916.174 | 4009.478 | 3917.147 | 46117.16 | 47259.31 | 44601.51 | 37410.43 | 38982.15 | 36528.21 | 11866.67 | 12625.17 | 12006.39 | 4969.251 | 5106.393 | 5380.778 |
| Stx11 | 74732 | 3.448441 | 5.184333 | 60.8715 | 2.03E-25 | 4.07E-23 | 0.169721 | 5.184333 | 5.053008 | 5.05E-05 | 0.00017 | 757.5623 | 744.1733 | 686.564 | 8506.545 | 8746.574 | 8293.206 | 10117.96 | 9994.185 | 9735.646 | 1306.126 | 1306.973 | 1332.219 | 580.6915 | 607.6419 | 601.7336 |
| Bcl2a1d | 12047 | 3.454501 | 4.960174 | 33.74762 | 5.22E-20 | 2.19E-18 | 0.689618 | 4.960174 | 12.20329 | 4.31E-11 | 3.59E-10 | 619.5539 | 680.2348 | 405.3418 | 6572.767 | 6555.717 | 6435.993 | 11317.5 | 10978.05 | 10536.61 | 1495.353 | 1495.317 | 1562.839 | 391.2993 | 344.3042 | 377.3995 |
| Mycl | 16918 | 3.454648 | 1.934889 | 18.47777 | 1.30E-14 | 1.76E-13 | 3.06979 | 1.934889 | 40.61141 | 1.06E-21 | 1.15E-19 | 86.06977 | 57.72227 | 60.75787 | 815.5894 | 789.2701 | 795.3686 | 7320.915 | 7038.555 | 6650.163 | 16.1535 | 22.11331 | 28.36097 | 71.75211 | 73.89177 | 77.74755 |
| Mier3 | 218613 | 3.455791 | 5.732384 | 66.18726 | 3.43E-26 | 9.29E-24 | -0.86784 | 5.732384 | -24.2106 | 5.25E-17 | 1.54E-15 | 1407.537 | 1417.304 | 1298.483 | 15996.2 | 16529.73 | 15814.04 | 9535.239 | 9090.74 | 8910.552 | 1732.271 | 1889.544 | 1771.814 | 863.5284 | 840.3223 | 963.7456 |
| 1700029E | 72250 | 3.456697 | -5.06176 | 4.17026 | 0.00042 | 0.001276 | -0.60976 | -5.06176 | -0.84745 | 0.406147 | 0.505248 | 0 | 0 | 0 | 2.597418 | 2.808791 | 9.468674 | 5.685043 | 1.340422 | 2.298311 | 0.769214 | 0 | 0 | 4.171635 | 4.716496 | 4.859222 |
| Slc28a2b | 381417 | 3.458243 | 0.735592 | 9.327907 | 5.57E-09 | 3.12E-08 | -0.02171 | 0.735592 | -0.11228 | 0.911649 | 0.929208 | 29.67923 | 13.32052 | 65.96569 | 403.8985 | 287.901 | 382.805 | 397.953 | 296.2332 | 402.2045 | 33.84544 | 44.98914 | 26.12194 | 51.72827 | 80.18043 | 91.51534 |
| Gm19304 | 100502648 | 3.458875 | -4.26732 | 3.519946 | 0.001997 | 0.005466 | 1.364933 | -4.26732 | 1.944333 | 0.065153 | 0.108414 | 0 | 0.888035 | 0 | 2.597418 | 16.85274 | 5.410671 | 27.00396 | 8.04253 | 25.28142 | 0 | 1.525056 | 0.746341 | 3.337308 | 7.860826 | 1.619741 |
| Gm6034 | 547347 | 3.45999 | 0.461452 | 12.29745 | 3.73E-11 | 2.82E-10 | 0.852568 | 0.461452 | 6.457032 | 1.97E-06 | 8.05E-06 | 20.77546 | 14.20856 | 19.9633 | 202.5986 | 241.556 | 213.7215 | 432.0633 | 395.4244 | 404.5028 | 31.53779 | 40.41398 | 44.78048 | 59.23721 | 62.10053 | 52.64157 |
| Taar7d | 435206 | 3.469153 | -4.72859 | 3.445208 | 0.002384 | 0.006446 | -0.62475 | -4.72859 | -0.71198 | 0.484186 | 0.582898 | 0 | 0 | 0 | 3.896127 | 7.021976 | 2.705335 | 0 | 4.021265 | 9.193245 | 2.307643 | 1.525056 | 0 | 10.01192 | 5.502578 | 5.669092 |
| Pim1 | 18712 | 3.472195 | 7.341149 | 66.48682 | 3.11E-26 | 8.81E-24 | -0.28254 | 7.341149 | -9.00695 | 1.02E-08 | 5.73E-08 | 3137.837 | 3496.194 | 2959.776 | 37746.98 | 38057.71 | 37489.19 | 33904.18 | 32245.19 | 30883.56 | 11918.21 | 12025.83 | 12399.71 | 1860.549 | 1675.928 | 1909.674 |
| Il10 | 16153 | 3.474459 | 4.422672 | 59.52797 | 3.26E-25 | 6.07E-23 | -3.78573 | 4.422672 | -54.2756 | 2.31E-24 | 6.46E-22 | 1643.487 | 1512.324 | 1392.223 | 18201.41 | 17991.71 | 17843.04 | 1388.572 | 1264.018 | 1410.014 | 1193.052 | 1090.415 | 1024.727 | 58.40288 | 87.25517 | 119.8608 |
| Dnase1l3 | 13421 | 3.478276 | 2.233285 | 6.965116 | 6.43E-07 | 2.74E-06 | 5.594322 | 2.233285 | 27.93131 | 2.71E-18 | 1.08E-16 | 2.967923 | 27.52908 | 23.43518 | 131.1696 | 182.5714 | 128.5034 | 7330.863 | 7392.426 | 7205.206 | 117.6898 | 134.2049 | 167.1804 | 953.6357 | 872.5517 | 894.9066 |
| Ifna5 | 15968 | 3.480465 | -3.99902 | 4.349083 | 0.000273 | 0.000848 | -4.05253 | -3.99902 | -4.64702 | 0.000133 | 0.000423 | 5.193865 | 0 | 0.86797 | 29.87031 | 14.04395 | 12.17401 | 0 | 0 | 2.298311 | 5.384501 | 5.337695 | 2.985365 | 2.502981 | 10.21907 | 4.049351 |
| Gm19426 | 100502874 | 3.484597 | -4.60034 | 3.607996 | 0.001619 | 0.004502 | 0.065716 | -4.60034 | 0.082472 | 0.93504 | 0.94663 | 0 | 0 | 0 | 1.298709 | 9.830767 | 5.410671 | 7.106304 | 12.0638 | 1.149156 | 0.769214 | 0.762528 | 0 | 9.177596 | 11.79124 | 8.098703 |
| Il20ra | 237313 | 3.48505 | -3.08032 | 4.096405 | 0.000502 | 0.001508 | 0.469875 | -3.08032 | 0.762549 | 0.454078 | 0.552898 | 4.451884 | 0 | 0 | 7.792255 | 12.63956 | 14.87935 | 17.05513 | 13.40422 | 19.53565 | 5.384501 | 1.525056 | 5.224389 | 30.8701 | 22.7964 | 14.57766 |
| Gm7337 | 654494 | 3.488418 | -3.828 | 3.254168 | 0.003738 | 0.009772 | 0.8418 | -3.828 | 1.114075 | 0.277636 | 0.36926 | 2.967923 | 0 | 0 | 20.77935 | 15.44835 | 2.705335 | 36.95278 | 16.08506 | 12.64071 | 0 | 0 | 2.985365 | 3.337308 | 16.50774 | 11.33818 |
| Nod2 | 257632 | 3.490345 | 5.259063 | 42.00386 | 5.22E-22 | 3.50E-20 | 0.423288 | 5.259063 | 8.93349 | 1.17E-08 | 6.54E-08 | 779.0798 | 714.8682 | 761.2093 | 9007.847 | 8944.593 | 9158.913 | 12939.16 | 13200.47 | 11770.8 | 963.0565 | 853.2687 | 806.0486 | 901.9074 | 751.495 | 874.6599 |
| Magea1 | 17137 | 3.491216 | -5.32919 | 3.784502 | 0.001062 | 0.003046 | 0.157875 | -5.32919 | 0.207838 | 0.837326 | 0.902303 | 0 | 0 | 0 | 2.597418 | 5.617581 | 5.410671 | 4.263783 | 2.680843 | 10.3424 | 0 | 0 | 0 | 1.668654 | 4.716496 | 0 |
| Olfr1277 | 258391 | 3.491216 | -5.3542 | 2.838818 | 0.009734 | 0.023436 | -0.37342 | -5.3542 | -0.35487 | 0.726167 | 0.80525 | 0 | 0 | 0 | 2.597418 | 5.617581 | 5.410671 | 0 | 5.361687 | 12.64071 | 2.307643 | 0 | 0 | 9.177596 | 0 | 0 |
| Cpb1 | 76703 | 3.49173 | -2.95739 | 3.894559 | 0.000816 | 0.002375 | -0.12424 | -2.95739 | -0.18634 | 0.853939 | 0.916138 | 10.38773 | 0 | 0 | 7.792255 | 15.44835 | 29.75869 | 17.05513 | 12.0638 | 14.93902 | 5.384501 | 4.575167 | 2.239024 | 26.69846 | 18.0799 | 17.00728 |
| Ccl12 | 20293 | 3.493579 | 7.573022 | 34.81613 | 1.02E-13 | 1.18E-12 | 1.161392 | 7.573022 | 22.99637 | 1.60E-11 | 1.44E-10 | 2470.054 | 2733.372 | 2148.225 | 29737.84 | 30031.59 | 28238.29 | 67714.55 | 63530.63 | 72466.9 | 7945.216 | 7599.353 | 7561.183 | 4900.836 | 4762.089 | 5732.262 |
| Vmn1r210 | 171269 | 3.496473 | -4.56487 | 3.809039 | 0.001001 | 0.002881 | -0.68774 | -4.56487 | -0.85961 | 0.399555 | 0.498705 | 0 | 0 | 0 | 5.194837 | 5.617581 | 2.705335 | 5.685043 | 0 | 6.894934 | 0 | 0 | 0 | 43.385 | 45.59279 | 34.01455 |
| Gm5796 | 545007 | 3.49757 | -4.80096 | 4.266104 | 0.000333 | 0.001026 | -0.95896 | -4.80096 | -1.35526 | 0.189505 | 0.267981 | 0 | 0 | 1.735939 | 3.896127 | 11.23516 | 10.82134 | 4.263783 | 4.021265 | 3.447467 | 1.538429 | 0 | 0 | 1.668654 | 2.358248 | 2.429611 |
| Olfr772 | 257666 | 3.500883 | -5.46626 | 3.541169 | 0.001899 | 0.005217 | -0.96964 | -5.46626 | -1.11026 | 0.27924 | 0.37086 | 0 | 0 | 0 | 5.194837 | 1.404395 | 9.468674 | 7.106304 | 4.021265 | 0 | 0 | 0 | 0 | 1.668654 | 0.786083 | 3.239481 |
| Prop1 | 19127 | 3.502578 | -4.08972 | 4.037996 | 0.000578 | 0.001722 | 0.479813 | -4.08972 | 0.714375 | 0.482736 | 0.581312 | 0 | 0 | 0.86797 | 5.194837 | 5.617581 | 9.468674 | 9.948826 | 5.361687 | 16.08818 | 1.538429 | 0 | 2.985365 | 8.343269 | 10.21907 | 7.288832 |
| F3 | 14066 | 3.50896 | -0.14259 | 6.333557 | 2.59E-06 | 1.02E-05 | 2.814468 | -0.14259 | 12.01004 | 5.81E-11 | 4.73E-10 | 5.935846 | 4.440175 | 19.09533 | 133.767 | 110.9472 | 68.98605 | 697.8391 | 839.104 | 704.4324 | 20.76879 | 25.16342 | 7.463413 | 38.37904 | 44.80671 | 51.8317 |
| Ifit3b | 667370 | 3.50933 | 6.790152 | 43.90474 | 1.22E-18 | 3.76E-17 | 0.574331 | 6.790152 | 13.2523 | 2.89E-10 | 2.09E-09 | 2179.197 | 2518.467 | 1711.636 | 25696.26 | 25550.16 | 25733.15 | 40147.78 | 38833.36 | 39965.33 | 2153.031 | 2398.15 | 1938.248 | 2634.804 | 2824.395 | 2756.798 |
| Rhox2-ps | 622311 | 3.511753 | -4.36655 | 3.777679 | 0.00108 | 0.003092 | 0.622253 | -4.36655 | 0.84556 | 0.407178 | 0.506297 | 0 | 0 | 0 | 2.597418 | 11.23516 | 2.705335 | 4.263783 | 13.40422 | 6.894934 | 1.538429 | 2.287584 | 0 | 6.674615 | 13.3634 | 8.908573 |
| Olfr376 | 258924 | 3.511871 | -4.24201 | 2.986798 | 0.006948 | 0.01725 | -0.38378 | -4.24201 | -0.39742 | 0.695002 | 0.777014 | 0 | 2.664105 | 0 | 11.68838 | 14.04395 | 4.058003 | 8.527565 | 1.340422 | 20.6848 | 0 | 2.287584 | 0.746341 | 19.18952 | 0.786083 | 6.478962 |
| Hspa1b | 15511 | 3.512941 | 4.718902 | 33.59373 | 5.74E-20 | 2.38E-18 | 0.949331 | 4.718902 | 17.42385 | 4.23E-14 | 6.41E-13 | 462.996 | 412.0482 | 356.7355 | 5002.628 | 4741.238 | 5301.105 | 10756.1 | 9967.376 | 9540.29 | 449.2212 | 481.9176 | 450.0438 | 934.4461 | 1064.356 | 1056.881 |
| Fcrlb | 435653 | 3.513312 | 1.197826 | 7.814938 | 1.07E-07 | 5.03E-07 | 3.57311 | 1.197826 | 19.97838 | 2.68E-15 | 5.25E-14 | 18.54952 | 20.4248 | 6.943757 | 219.4818 | 147.4615 | 163.6728 | 2346.502 | 2278.717 | 1946.67 | 45.38365 | 31.26364 | 23.13658 | 260.31 | 292.4227 | 268.8769 |
| Ptgs2os | 320019 | 3.515061 | -2.38361 | 4.131527 | 0.000461 | 0.001392 | 2.509547 | -2.38361 | 5.523869 | 1.66E-05 | 5.97E-05 | 2.967923 | 0 | 5.207818 | 32.46773 | 36.51428 | 12.17401 | 149.2324 | 172.9144 | 128.7054 | 4.615287 | 2.287584 | 0 | 16.68654 | 18.0799 | 10.52831 |
| Ppef2 | 19023 | 3.515093 | -0.92176 | 6.637398 | 1.32E-06 | 5.38E-06 | -0.30128 | -0.92176 | -0.98027 | 0.337935 | 0.434871 | 6.677827 | 4.440175 | 10.41564 | 62.33804 | 91.28569 | 119.0348 | 56.85043 | 83.10615 | 81.59005 | 30.76858 | 12.20045 | 12.6878 | 33.37308 | 30.65722 | 55.07118 |
| Gm15179 | 100503248 | 3.516672 | -4.69553 | 3.613476 | 0.001598 | 0.004448 | 0.329183 | -4.69553 | 0.417939 | 0.680167 | 0.763782 | 0 | 0 | 0 | 2.597418 | 2.808791 | 10.82134 | 4.263783 | 10.72337 | 4.596622 | 3.076858 | 0.762528 | 0 | 9.177596 | 3.14433 | 1.619741 |
| Oas1e | 231699 | 3.518044 | -1.42139 | 4.896234 | 7.34E-05 | 0.000244 | 2.255738 | -1.42139 | 5.980957 | 5.77E-06 | 2.22E-05 | 1.483961 | 3.55214 | 1.735939 | 42.8574 | 15.44835 | 39.22736 | 139.2836 | 144.7655 | 167.7767 | 6.153716 | 0.762528 | 10.44878 | 110.1312 | 102.9768 | 118.2411 |
| Gstp3 | 225884 | 3.520178 | -1.49787 | 4.751909 | 0.000104 | 0.000338 | 5.068679 | -1.49787 | 13.31686 | 8.22E-12 | 7.84E-11 | 0.741981 | 1.77607 | 1.735939 | 23.37676 | 21.06593 | 17.58468 | 652.3587 | 876.6358 | 707.8798 | 3.076858 | 3.812639 | 5.97073 | 30.03577 | 49.52321 | 25.91585 |
| 4932438H2 | 74387 | 3.520755 | 1.697143 | 14.58013 | 1.43E-12 | 1.35E-11 | 0.882832 | 1.697143 | 8.042519 | 6.73E-08 | 3.38E-07 | 60.10044 | 55.9462 | 44.26645 | 667.5365 | 616.5295 | 689.8605 | 1254.973 | 1470.443 | 1103.189 | 56.92187 | 75.49026 | 55.22925 | 79.26106 | 113.1959 | 105.2831 |
| Gm10610 | 100038384 | 3.524109 | -2.70433 | 3.919605 | 0.000768 | 0.002247 | 0.202983 | -2.70433 | 0.346037 | 0.732703 | 0.811232 | 2.967923 | 7.10428 | 0 | 27.27289 | 19.66153 | 35.16936 | 25.5827 | 29.48928 | 41.3696 | 8.461359 | 4.575167 | 0 | 20.85817 | 13.3634 | 7.288832 |
| Ifi209 | 236312 | 3.524141 | 7.701242 | 72.99222 | 4.28E-27 | 1.52E-24 | -0.54351 | 7.701242 | -18.6621 | 1.06E-14 | 1.81E-13 | 4804.325 | 4847.783 | 3877.22 | 54821.11 | 56483.37 | 54471.93 | 40129.3 | 38939.25 | 39028.77 | 11635.14 | 11656 | 11120.48 | 2717.403 | 2716.702 | 2554.331 |
| Olfr893 | 258333 | 3.524384 | -5.0108 | 4.09162 | 0.000508 | 0.001525 | 0.494784 | -5.0108 | 0.719171 | 0.479836 | 0.578271 | 0 | 0 | 0 | 6.493546 | 8.426372 | 1.352668 | 8.527565 | 5.361687 | 6.894934 | 0 | 0 | 0 | 2.502981 | 7.860826 | 1.619741 |
| Shroom4 | 208431 | 3.524431 | 0.053583 | 10.62121 | 5.58E-10 | 3.60E-09 | -1.10756 | 0.053583 | -5.32204 | 2.67E-05 | 9.34E-05 | 26.71131 | 19.53677 | 15.62345 | 237.6638 | 254.1955 | 275.9442 | 176.2363 | 101.8721 | 106.8715 | 13.07665 | 8.387806 | 20.89756 | 95.9476 | 89.61342 | 124.72 |
| Vgf | 381677 | 3.52475 | 0.122612 | 13.11964 | 1.09E-11 | 9.04E-11 | -1.61585 | 0.122612 | -8.60119 | 2.22E-08 | 1.20E-07 | 34.8731 | 24.86498 | 29.51097 | 359.7424 | 348.29 | 404.4477 | 112.2796 | 140.7443 | 122.9596 | 16.92272 | 19.82572 | 5.224389 | 57.56856 | 71.53352 | 72.07845 |
| F13b | 14060 | 3.526018 | -3.73631 | 3.755173 | 0.001139 | 0.003248 | 0.262855 | -3.73631 | 0.367805 | 0.716641 | 0.796702 | 0 | 3.55214 | 0 | 7.792255 | 9.830767 | 12.17401 | 4.263783 | 16.08506 | 22.98311 | 0 | 0 | 2.985365 | 19.18952 | 18.86598 | 21.8665 |
| Cp | 12870 | 3.534711 | 6.80613 | 58.18288 | 6.92E-21 | 3.68E-19 | -0.96705 | 6.80613 | -24.3735 | 1.37E-14 | 2.27E-13 | 4457.82 | 4732.338 | 3622.905 | 52731.49 | 53531.33 | 51516.35 | 28135.28 | 29257.39 | 26639.72 | 7779.066 | 7726.695 | 7530.583 | 257.807 | 295.5671 | 284.2645 |
| Ms4a8a | 64381 | 3.53755 | -1.30656 | 6.520698 | 1.71E-06 | 6.87E-06 | 2.451757 | -1.30656 | 9.14802 | 7.79E-09 | 4.50E-08 | 2.967923 | 1.77607 | 3.471878 | 29.87031 | 33.70549 | 52.75404 | 235.9293 | 193.0207 | 221.787 | 20.76879 | 18.30067 | 26.12194 | 17.52087 | 11.00516 | 15.38754 |
| Tmem67 | 329795 | 3.537893 | 4.804104 | 45.27753 | 1.07E-22 | 8.49E-21 | -1.05894 | 4.804104 | -19.8285 | 3.12E-15 | 5.99E-14 | 673.7185 | 645.6014 | 544.2169 | 7627.319 | 7899.723 | 7592.524 | 3955.369 | 4057.457 | 3557.786 | 1216.897 | 1239.87 | 1113.541 | 603.2184 | 607.6419 | 571.7684 |
| C8b | 110382 | 3.539032 | -3.72795 | 4.618774 | 0.000143 | 0.000458 | 0.773248 | -3.72795 | 1.375571 | 0.183221 | 0.260522 | 0 | 1.77607 | 0 | 6.493546 | 8.426372 | 9.468674 | 11.37009 | 14.74464 | 18.38649 | 1.538429 | 0.762528 | 1.492683 | 8.343269 | 14.93557 | 13.76779 |
| Synb | 239167 | 3.542723 | -3.12957 | 3.184853 | 0.004395 | 0.011325 | 1.299646 | -3.12957 | 1.750491 | 0.094397 | 0.148471 | 0 | 0 | 5.207818 | 23.37676 | 2.808791 | 18.93735 | 25.5827 | 17.42548 | 54.01031 | 0 | 1.525056 | 6.717071 | 25.02981 | 21.22423 | 28.34546 |
| 4921504A2 | 70832 | 3.546375 | -3.52649 | 3.375712 | 0.002809 | 0.007497 | 0.866766 | -3.52649 | 1.150941 | 0.262488 | 0.352211 | 0 | 2.664105 | 0 | 6.493546 | 7.021976 | 16.23201 | 14.21261 | 12.0638 | 29.87805 | 0 | 5.337695 | 4.478048 | 16.68654 | 17.29382 | 5.669092 |
| Olfr857 | 257963 | 3.548518 | -4.85065 | 4.066186 | 0.00054 | 0.001615 | 0.381356 | -4.85065 | 0.542585 | 0.593037 | 0.686055 | 0 | 0 | 0 | 3.896127 | 8.426372 | 2.705335 | 4.263783 | 5.361687 | 10.3424 | 1.538429 | 0 | 0 | 7.508942 | 2.358248 | 2.429611 |
| Gm10499 | 69717 | 3.551174 | -1.51436 | 4.407774 | 0.000237 | 0.000743 | 1.198214 | -1.51436 | 2.843048 | 0.009641 | 0.020898 | 8.161788 | 11.54445 | 0 | 50.64966 | 50.55823 | 33.81669 | 112.2796 | 121.9784 | 90.78329 | 14.61507 | 9.912862 | 2.985365 | 40.88202 | 30.65722 | 18.62702 |
| Gm829 | 329839 | 3.552368 | -4.8112 | 3.930548 | 0.000748 | 0.002192 | 0.884392 | -4.8112 | 1.266852 | 0.218859 | 0.302465 | 0 | 0 | 0 | 3.896127 | 2.808791 | 8.116006 | 8.527565 | 5.361687 | 16.08818 | 1.538429 | 1.525056 | 0 | 0.834327 | 3.930413 | 0.80987 |
| Plscr2 | 18828 | 3.554252 | -2.80884 | 4.543017 | 0.000171 | 0.000545 | -0.17818 | -2.80884 | -0.33878 | 0.738086 | 0.815956 | 2.225942 | 0 | 12.15157 | 28.5716 | 30.8967 | 31.11136 | 21.31891 | 22.78717 | 40.22045 | 0 | 3.050111 | 1.492683 | 18.35519 | 11.79124 | 17.81715 |
| Fap | 14089 | 3.560638 | 1.355643 | 14.96203 | 8.62E-13 | 8.45E-12 | -0.32163 | 1.355643 | -2.53394 | 0.019165 | 0.038028 | 32.64715 | 49.72996 | 41.66254 | 567.5359 | 480.3032 | 488.3131 | 362.4215 | 491.9348 | 426.3367 | 59.99873 | 77.77784 | 49.25852 | 156.0191 | 157.2165 | 142.5372 |
| Gm10512 | 100038394 | 3.562826 | -3.51772 | 3.307751 | 0.003297 | 0.008692 | 1.228354 | -3.51772 | 1.647529 | 0.114094 | 0.174556 | 1.483961 | 0 | 0 | 19.48064 | 7.021976 | 4.058003 | 21.31891 | 24.12759 | 19.53565 | 4.615287 | 3.812639 | 2.985365 | 21.6925 | 0.786083 | 10.52831 |
| Dusp16 | 70686 | 3.563018 | 6.12108 | 77.26561 | 1.28E-27 | 5.17E-25 | -1.51827 | 6.12108 | -44.2688 | 1.72E-22 | 2.80E-20 | 1867.566 | 2013.175 | 2004.142 | 24952.1 | 25236.98 | 23917.87 | 9069.065 | 9195.293 | 8633.606 | 1612.273 | 1694.337 | 1744.2 | 1742.909 | 1603.609 | 1776.045 |
| Ighv8-8 | 780938 | 3.563384 | -4.20928 | 3.485593 | 0.002166 | 0.005899 | -1.97669 | -4.20928 | -2.08398 | 0.049359 | 0.08565 | 0 | 0 | 1.735939 | 11.68838 | 11.23516 | 4.058003 | 2.842522 | 0 | 4.596622 | 6.153716 | 0 | 2.985365 | 8.343269 | 12.57732 | 14.57766 |
| Abcb1a | 18671 | 3.565611 | 2.355923 | 15.9618 | 2.42E-13 | 2.61E-12 | 0.486856 | 2.355923 | 4.608486 | 0.000146 | 0.000462 | 28.19527 | 41.73764 | 40.79457 | 441.5611 | 440.9801 | 493.7237 | 672.2564 | 596.4877 | 727.4155 | 162.3042 | 150.218 | 139.5658 | 1565.197 | 1579.24 | 1622.17 |
| Dlx1 | 13390 | 3.569106 | -2.11332 | 4.522243 | 0.00018 | 0.000571 | 0.019807 | -2.11332 | 0.038462 | 0.969677 | 0.975089 | 1.483961 | 6.216245 | 0.86797 | 32.46773 | 32.30109 | 28.40602 | 24.16143 | 20.10633 | 63.20356 | 1.538429 | 6.100223 | 20.15121 | 30.03577 | 25.94073 | 36.44416 |
| Tcp10b | 21462 | 3.571638 | -0.36243 | 11.13505 | 2.36E-10 | 1.60E-09 | -0.34221 | -0.36243 | -1.96281 | 0.062833 | 0.105227 | 20.77546 | 10.65642 | 12.15157 | 185.7154 | 190.9978 | 178.5521 | 139.2836 | 164.8719 | 150.5394 | 28.46093 | 28.21353 | 20.89756 | 19.18952 | 26.72681 | 16.19741 |
| Gm6497 | 624362 | 3.572792 | -4.56674 | 3.535965 | 0.001922 | 0.005276 | -0.44063 | -4.56674 | -0.52374 | 0.605856 | 0.698163 | 0 | 0 | 0.86797 | 15.58451 | 7.021976 | 2.705335 | 1.421261 | 8.04253 | 9.193245 | 4.615287 | 0 | 0 | 2.502981 | 5.502578 | 4.859222 |
| Fam83a | 239463 | 3.574041 | -2.57179 | 4.426933 | 0.000226 | 0.00071 | 1.276213 | -2.57179 | 2.555887 | 0.018269 | 0.036475 | 0 | 7.10428 | 0.86797 | 20.77935 | 15.44835 | 18.93735 | 28.42522 | 44.23392 | 74.69511 | 3.076858 | 4.575167 | 2.985365 | 21.6925 | 19.65207 | 17.00728 |
| Procr | 19124 | 3.584347 | 4.892865 | 44.17871 | 1.80E-22 | 1.33E-20 | 1.771605 | 4.892865 | 44.9706 | 1.24E-22 | 2.05E-20 | 506.0309 | 548.8056 | 507.7622 | 6759.781 | 6319.779 | 6904.016 | 24113.11 | 23978.8 | 22864.75 | 449.9905 | 460.5668 | 497.8096 | 488.9156 | 442.5645 | 445.4286 |
| Fgf5 | 14176 | 3.585343 | -3.50182 | 3.800693 | 0.001022 | 0.002937 | 0.395112 | -3.50182 | 0.582069 | 0.566618 | 0.66087 | 0.741981 | 1.77607 | 0 | 2.597418 | 32.30109 | 21.64268 | 12.79135 | 12.0638 | 29.87805 | 0 | 0.762528 | 0.746341 | 18.35519 | 21.22423 | 25.10598 |
| Il23a | 83430 | 3.585408 | -1.10746 | 8.568598 | 2.36E-08 | 1.20E-07 | -0.89353 | -1.10746 | -3.4501 | 0.002357 | 0.005954 | 7.419807 | 19.53677 | 7.811726 | 161.0399 | 119.3736 | 129.8561 | 75.32683 | 58.97856 | 94.23076 | 13.84586 | 9.912862 | 14.92683 | 5.840288 | 18.86598 | 11.33818 |
| Ccnd2 | 12444 | 3.586446 | 6.725792 | 47.45892 | 1.25E-19 | 4.82E-18 | 1.16258 | 6.725792 | 31.19821 | 1.53E-16 | 4.06E-15 | 2260.073 | 2323.988 | 2294.912 | 29827.45 | 29374.33 | 28972.79 | 70648.03 | 70285.01 | 64907.76 | 1690.733 | 1878.869 | 1743.453 | 1188.082 | 1147.681 | 1395.406 |
| Bco2 | 170752 | 3.588287 | 1.588008 | 22.26954 | 2.93E-16 | 5.50E-15 | -2.65883 | 1.588008 | -18.6448 | 1.09E-14 | 1.84E-13 | 92.74759 | 80.81118 | 96.34463 | 1063.643 | 1138.965 | 1291.798 | 220.2954 | 176.9357 | 179.2683 | 119.2282 | 120.4794 | 114.9366 | 90.10731 | 86.46909 | 61.55014 |
| AW01173 | 100382 | 3.594483 | 2.238275 | 20.64947 | 1.37E-15 | 2.21E-14 | -0.20584 | 2.238275 | -2.28843 | 0.032405 | 0.059775 | 83.84382 | 81.69922 | 73.77742 | 966.2396 | 1123.516 | 1023.969 | 939.4534 | 966.4441 | 894.0431 | 237.6873 | 202.8324 | 160.4634 | 120.9774 | 102.9768 | 124.72 |
| Olfr58 | 18358 | 3.595041 | -5.19478 | 4.152931 | 0.000438 | 0.001326 | -1.13156 | -5.19478 | -1.46556 | 0.157345 | 0.229485 | 0 | 0 | 0 | 5.194837 | 2.808791 | 6.763339 | 0 | 2.680843 | 5.745778 | 0 | 0 | 0 | 6.674615 | 3.14433 | 6.478962 |
| Vmn2r-ps1 | 319210 | 3.596891 | -4.50661 | 3.715072 | 0.001254 | 0.003551 | 0.98667 | -4.50661 | 1.337006 | 0.195298 | 0.275048 | 0 | 0 | 0 | 6.493546 | 2.808791 | 5.410671 | 2.842522 | 20.10633 | 16.08818 | 0 | 1.525056 | 0 | 7.508942 | 12.57732 | 4.859222 |
| Gm19261 | 100502579 | 3.603855 | -0.97069 | 6.751987 | 1.02E-06 | 4.23E-06 | -0.34531 | -0.97069 | -1.18815 | 0.247827 | 0.33557 | 18.54952 | 7.10428 | 8.679696 | 122.0787 | 209.2549 | 120.3874 | 109.4371 | 139.4039 | 109.1698 | 23.84565 | 17.53814 | 10.44878 | 20.02385 | 13.3634 | 1.619741 |
| Ccl22 | 20299 | 3.605597 | 4.857653 | 28.79378 | 1.44E-18 | 4.35E-17 | 1.473747 | 4.857653 | 24.50407 | 4.09E-17 | 1.23E-15 | 348.731 | 312.5883 | 328.9605 | 3998.726 | 4369.074 | 4578.78 | 12325.17 | 13107.98 | 11896.06 | 1372.279 | 1591.396 | 1264.302 | 635.7571 | 684.678 | 572.5783 |
| Ip6k3 | 271424 | 3.605685 | -3.13716 | 3.658061 | 0.001437 | 0.004029 | -1.47088 | -3.13716 | -1.70914 | 0.101928 | 0.158562 | 0 | 0.888035 | 3.471878 | 16.88322 | 12.63956 | 16.23201 | 12.79135 | 0 | 20.6848 | 6.153716 | 6.100223 | 8.956095 | 18.35519 | 12.57732 | 21.05663 |
| Gm20367 | 100504712 | 3.610151 | 1.079041 | 16.63298 | 1.07E-13 | 1.23E-12 | -0.07731 | 1.079041 | -0.70961 | 0.485629 | 0.584367 | 41.55092 | 36.40943 | 46.87036 | 561.0424 | 497.1559 | 578.9418 | 572.7681 | 520.0836 | 521.7166 | 109.2285 | 89.97829 | 82.09754 | 29.20144 | 36.94588 | 19.43689 |
| Rnf7l | 631906 | 3.612558 | -4.49246 | 3.26852 | 0.003614 | 0.009469 | -0.80068 | -4.49246 | -0.84831 | 0.40568 | 0.50477 | 0 | 0 | 0.86797 | 7.792255 | 8.426372 | 5.410671 | 0 | 6.702109 | 16.08818 | 0 | 0 | 2.985365 | 3.337308 | 10.21907 | 12.95792 |
| Teddm1b | 433365 | 3.614054 | -3.72506 | 4.158966 | 0.000432 | 0.001308 | -0.93452 | -3.72506 | -1.27501 | 0.216011 | 0.299064 | 0 | 0 | 3.471878 | 18.18193 | 11.23516 | 5.410671 | 4.263783 | 8.04253 | 4.596622 | 1.538429 | 1.525056 | 2.985365 | 17.52087 | 19.65207 | 7.288832 |
| Fpr3 | 14294 | 3.614085 | -2.69144 | 7.424665 | 2.41E-07 | 1.08E-06 | -2.65489 | -2.69144 | -5.93757 | 6.38E-06 | 2.43E-05 | 4.451884 | 13.32052 | 6.075787 | 89.61093 | 101.1165 | 98.74475 | 9.948826 | 17.42548 | 19.53565 | 0 | 3.812639 | 2.985365 | 3.337308 | 2.358248 | 3.239481 |
| Olfr1349 | 269862 | 3.614742 | -4.01722 | 3.074112 | 0.005683 | 0.014357 | 0.078087 | -4.01722 | 0.083868 | 0.933943 | 0.945883 | 0.741981 | 0 | 0 | 3.896127 | 4.213186 | 21.64268 | 7.106304 | 2.680843 | 21.83396 | 8.461359 | 1.525056 | 0 | 3.337308 | 14.93557 | 13.76779 |
| Olfr225 | 257886 | 3.61644 | -4.47508 | 3.272276 | 0.003583 | 0.009388 | 0.61292 | -4.47508 | 0.705037 | 0.48841 | 0.586775 | 0 | 0 | 0 | 2.597418 | 4.213186 | 9.468674 | 1.421261 | 13.40422 | 18.38649 | 0 | 0 | 5.97073 | 7.508942 | 7.860826 | 6.478962 |
| Phf11 | 628693 | 3.618444 | 1.247847 | 23.50338 | 9.66E-17 | 1.99E-15 | -3.11641 | 1.247847 | -20.3592 | 1.83E-15 | 3.68E-14 | 134.2985 | 138.5335 | 93.74072 | 1598.711 | 1513.938 | 1659.723 | 200.3978 | 197.042 | 175.8208 | 281.5325 | 300.436 | 212.7073 | 5.005962 | 4.716496 | 7.288832 |

| | | | | | | | | | | | | | | | | | | | | | | | | | | |
|---|---|---|---|---|---|---|---|---|---|---|---|---|---|---|---|---|---|---|---|---|---|---|---|---|---|---|
| Olfr204 | 258994 | 3.621176 | -5.0855 | 3.881794 | 0.000841 | 0.002446 | -0.87766 | -5.0855 | -1.07751 | 0.293283 | 0.386532 | 0 | 0 | 0 | 11.68838 | 1.404395 | 5.410671 | 2.842522 | 1.340422 | 3.447467 | 0 | 2.287584 | 0 | 3.337308 | 1.572165 | 4.859222 |
| Ifi203-ps | 100504287 | 3.624151 | 3.839588 | 37.59177 | 5.41E-21 | 2.94E-19 | 0.229103 | 3.839588 | 4.474753 | 0.000202 | 0.000624 | 245.5956 | 273.5148 | 265.5987 | 3435.086 | 3374.762 | 3489.883 | 4091.81 | 4479.689 | 3993.316 | 519.2198 | 513.1812 | 509.0047 | 219.428 | 253.9047 | 238.9117 |
| Gm20125 | 100504235 | 3.632464 | -0.9947 | 7.794999 | 1.11E-07 | 5.23E-07 | 0.047617 | -0.9947 | 0.196759 | 0.845878 | 0.909395 | 8.903769 | 6.216245 | 13.88751 | 114.2864 | 102.5209 | 173.1415 | 92.38196 | 158.1698 | 163.1801 | 4.615287 | 6.100223 | 2.239024 | 30.8701 | 36.94588 | 32.39481 |
| Ifit3 | 15959 | 3.632525 | 8.399152 | 37.32317 | 2.16E-12 | 1.98E-11 | -0.29284 | 8.399152 | -5.67129 | 0.00018 | 0.000561 | 8169.208 | 8256.949 | 5910.005 | 98985.01 | 99564.6 | 94924.81 | 86297.54 | 82779.09 | 80339.77 | 12232.05 | 12711.34 | 10968.98 | 4801.551 | 4560.065 | 4685.099 |
| Vmn2r92 | 627111 | 3.632651 | -4.23012 | 3.317396 | 0.003223 | 0.008516 | -1.56246 | -4.23012 | -1.5988 | 0.124565 | 0.187941 | 0 | 0.888035 | 0.86797 | 16.88322 | 8.426372 | 8.116006 | 0 | 2.680843 | 21.83396 | 2.307643 | 0 | 4.478048 | 1.668654 | 7.860826 | 8.098703 |
| 1110032F( | 68725 | 3.633485 | 0.371039 | 8.315164 | 3.90E-08 | 1.94E-07 | 1.950861 | 0.371039 | 10.51399 | 6.71E-10 | 4.58E-09 | 4.451884 | 10.65642 | 22.56721 | 166.2348 | 127.8 | 119.0348 | 575.6106 | 554.9346 | 527.4624 | 47.6913 | 57.18959 | 57.46828 | 80.92971 | 51.09537 | 58.31066 |
| Gnb4 | 14696 | 3.633732 | 4.68949 | 49.83558 | 1.41E-23 | 1.46E-21 | -0.79959 | 4.68949 | -17.225 | 5.32E-14 | 7.86E-13 | 490.4493 | 487.5312 | 487.7989 | 6289.648 | 6597.849 | 6571.26 | 3899.94 | 3812.159 | 3879.549 | 1464.584 | 1268.846 | 1294.902 | 542.3125 | 488.9434 | 564.4796 |
| Krt222 | 268481 | 3.639391 | 0.523999 | 13.4117 | 7.18E-12 | 6.08E-11 | -1.43188 | 0.523999 | -8.17482 | 5.16E-08 | 2.64E-07 | 51.93865 | 31.96926 | 35.58675 | 527.2759 | 516.8175 | 550.5358 | 167.7088 | 237.2546 | 210.2955 | 20.76879 | 13.7255 | 29.85365 | 88.43865 | 34.58764 | 46.97248 |
| Aldh1b1 | 72535 | 3.643339 | 4.707298 | 50.66353 | 9.94E-24 | 1.09E-21 | -0.81659 | 4.707298 | -17.5172 | 3.80E-14 | 5.81E-13 | 671.4926 | 731.7408 | 605.8428 | 9048.107 | 9158.062 | 8533.981 | 5379.472 | 5495.729 | 4936.772 | 1781.501 | 1891.832 | 1847.195 | 171.037 | 154.8583 | 156.305 |
| Dcpp2 | 630537 | 3.644065 | -4.33127 | 3.704305 | 0.001287 | 0.003635 | -0.224 | -4.33127 | -0.28094 | 0.781463 | 0.853328 | 0 | 0.888035 | 0 | 12.98709 | 2.808791 | 9.468674 | 2.842522 | 14.74464 | 5.745778 | 0 | 4.575167 | 0 | 10.01192 | 6.288661 | 6.478962 |
| Sp140 | 434484 | 3.645341 | 6.708011 | 21.33875 | 7.60E-10 | 4.80E-09 | -0.81032 | 6.708011 | -7.37994 | 2.04E-05 | 7.24E-05 | 2527.186 | 2727.155 | 1283.727 | 27681.99 | 28344.91 | 27394.23 | 16515.05 | 16752.59 | 16141.04 | 4289.909 | 4347.171 | 4148.165 | 2483.791 | 2411.701 | 2196.368 |
| Cyp2d9 | 13105 | 3.646147 | -3.7846 | 3.96063 | 0.000696 | 0.002049 | 0.380922 | -3.7846 | 0.549635 | 0.588276 | 0.681777 | 0 | 1.77607 | 0 | 3.896127 | 15.44835 | 10.82134 | 11.37009 | 6.702109 | 20.6848 | 0 | 3.050111 | 1.492683 | 14.18356 | 10.21907 | 16.19741 |
| Prr32 | 68800 | 3.658775 | -4.74141 | 3.482506 | 0.002182 | 0.005939 | -0.02601 | -4.74141 | -0.03012 | 0.976254 | 0.980271 | 0 | 0 | 0 | 5.194837 | 4.213186 | 5.410671 | 2.842522 | 2.680843 | 13.78987 | 0.769214 | 0.762528 | 0 | 13.34923 | 9.432991 | 0.80987 |
| Gm3002 | 100040852 | 3.661763 | -4.70117 | 4.59747 | 0.00015 | 0.000481 | -0.83296 | -4.70117 | -1.23369 | 0.230731 | 0.316114 | 0.741981 | 0 | 0 | 9.090964 | 5.617581 | 8.116006 | 2.842522 | 2.680843 | 8.044089 | 1.538429 | 0 | 0 | 3.337308 | 5.502578 | 3.239481 |
| Cst9 | 13013 | 3.6657 | -4.76984 | 4.185072 | 0.000405 | 0.001234 | 0.21648 | -4.76984 | 0.307141 | 0.761713 | 0.836241 | 0 | 0 | 0 | 2.597418 | 8.426372 | 5.410671 | 9.948826 | 2.680843 | 8.044089 | 0 | 0.762528 | 0 | 2.502981 | 9.432991 | 6.478962 |
| Vmn1r148 | 81011 | 3.6657 | -5.60938 | 3.868477 | 0.000868 | 0.002521 | -1.61278 | -5.60938 | -1.862 | 0.076429 | 0.124245 | 0 | 0 | 0 | 2.597418 | 8.426372 | 5.410671 | 2.842522 | 0 | 2.298311 | 0 | 0 | 0 | 1.668654 | 4.716496 | 0 |
| Tnfaip2 | 21928 | 3.667096 | 8.495798 | 80.44396 | 5.42E-28 | 3.03E-25 | -0.38746 | 8.495798 | -15.4358 | 4.68E-13 | 5.59E-12 | 5733.285 | 5695.856 | 5679.993 | 77462.81 | 77969.22 | 76713.85 | 62323.71 | 64365.71 | 58298.96 | 16171.2 | 16185.42 | 16692.67 | 10539.22 | 10505.21 | 10521.83 |
| Msantd5 | 622699 | 3.668986 | -4.59945 | 3.766978 | 0.001108 | 0.003167 | 0.985181 | -4.59945 | 1.34211 | 0.193664 | 0.272929 | 0 | 0 | 0 | 16.88322 | 4.213186 | 1.352668 | 11.37009 | 8.04253 | 13.78987 | 0 | 0 | 0 | 14.18356 | 13.3634 | 3.239481 |
| Kcna6 | 16494 | 3.670112 | -3.3271 | 3.848968 | 0.00091 | 0.002633 | 0.689814 | -3.3271 | 1.064021 | 0.29921 | 0.393044 | 0 | 0 | 6.943757 | 12.98709 | 23.87472 | 8.116006 | 24.16143 | 22.78717 | 22.98311 | 1.538429 | 3.050111 | 0 | 26.69846 | 13.3634 | 8.908573 |
| Zcwpw2 | 100039681 | 3.671208 | -2.85286 | 4.588208 | 0.000154 | 0.000491 | -0.88922 | -2.85286 | -1.52497 | 0.141948 | 0.209799 | 3.709904 | 7.992315 | 0 | 29.87031 | 44.94065 | 27.05335 | 27.00396 | 10.72337 | 20.6848 | 10.769 | 13.7255 | 11.94146 | 0 | 3.14433 | 4.859222 |
| Slc13a1 | 55961 | 3.673531 | -1.9125 | 5.761061 | 9.58E-06 | 3.55E-05 | -0.28948 | -1.9125 | -0.71766 | 0.480748 | 0.579316 | 0.741981 | 2.664105 | 9.547666 | 38.96127 | 37.91867 | 52.75404 | 38.37404 | 36.19139 | 34.47467 | 9.230573 | 17.53814 | 5.224389 | 27.53279 | 15.72165 | 16.19741 |
| Gabra6 | 14399 | 3.680536 | -3.66499 | 3.561048 | 0.001811 | 0.004995 | 0.544324 | -3.66499 | 0.736851 | 0.469236 | 0.567837 | 0 | 0 | 3.471878 | 9.090964 | 26.68351 | 5.410671 | 11.37009 | 18.7659 | 21.83396 | 1.538429 | 6.862751 | 0 | 3.337308 | 13.3634 | 8.098703 |
| Slc7a11 | 26570 | 3.682215 | 5.848545 | 69.06796 | 1.39E-26 | 4.39E-24 | -0.19161 | 5.848545 | -6.09264 | 4.47E-06 | 1.75E-05 | 1354.115 | 1402.207 | 1333.201 | 18874.14 | 18869.45 | 18290.77 | 16998.28 | 16698.97 | 17190.22 | 2553.792 | 2446.952 | 2815.946 | 370.4412 | 435.4898 | 496.4505 |
| Becn2 | 226720 | 3.682499 | -4.33903 | 2.724013 | 0.012599 | 0.02955 | -0.0341 | -4.33903 | -0.0307 | 0.975792 | 0.979891 | 0 | 0 | 0 | 2.597418 | 7.021976 | 6.763339 | 0 | 14.74464 | 14.93902 | 6.153716 | 7.625279 | 2.985365 | 4.171635 | 14.93557 | 0 |
| Gm3739 | 100042235 | 3.683213 | -5.24922 | 4.614695 | 0.000144 | 0.000463 | -2.29437 | -5.24922 | -3.07218 | 0.005708 | 0.01314 | 0.741981 | 0.888035 | 0 | 11.68838 | 9.830767 | 12.17401 | 2.842522 | 0 | 5.745778 | 0 | 0 | 0 | 0 | 0.786083 | 1.619741 |
| Gadd45b | 17873 | 3.685034 | 4.995012 | 46.95123 | 4.97E-23 | 4.40E-21 | -0.93821 | 4.995012 | -18.4019 | 1.41E-14 | 2.33E-13 | 550.5497 | 578.1108 | 601.5029 | 7748.099 | 8005.053 | 7967.213 | 4704.373 | 4225.009 | 3994.465 | 1777.655 | 1742.376 | 1952.429 | 705.0062 | 742.8481 | 666.5232 |
| Gbp2b | 14468 | 3.691851 | -0.73959 | 6.994037 | 6.04E-07 | 2.58E-06 | 1.023857 | -0.73959 | 4.244673 | 0.000351 | 0.001041 | 12.61367 | 5.32821 | 5.207818 | 127.2735 | 61.79339 | 144.7354 | 217.4529 | 211.7866 | 236.7261 | 16.1535 | 23.63836 | 10.44878 | 20.85817 | 24.36856 | 16.19741 |
| Lpcat2b | 70902 | 3.692695 | -3.2233 | 4.65513 | 0.000131 | 0.000422 | 0.464291 | -3.2233 | 0.840959 | 0.409694 | 0.508806 | 0.741981 | 1.77607 | 0 | 11.68838 | 14.04395 | 14.87935 | 19.89765 | 24.12759 | 16.08818 | 0.769214 | 3.050111 | 7.463413 | 7.508942 | 18.86598 | 9.718443 |
| Usp18 | 24110 | 3.695206 | 7.622689 | 92.97929 | 2.48E-29 | 3.35E-26 | -0.94781 | 7.622689 | -38.4925 | 3.29E-21 | 3.10E-19 | 4649.251 | 4758.091 | 4630.618 | 65245.85 | 64933.62 | 63894.61 | 35864.1 | 35088.22 | 33738.06 | 8865.966 | 8777.458 | 8026.9 | 2490.466 | 2489.524 | 2560 |
| Phf11c | 628705 | 3.697009 | 3.295071 | 35.98472 | 1.36E-20 | 6.68E-19 | -0.7566 | 3.295071 | -12.6271 | 2.26E-11 | 1.99E-10 | 241.1437 | 232.6652 | 180.5377 | 3022.096 | 2915.525 | 3111.136 | 1871.801 | 1883.293 | 1808.771 | 485.3743 | 493.3555 | 456.0145 | 110.1312 | 127.3454 | 121.4805 |
| Ifi44 | 99899 | 3.697503 | 5.512986 | 56.06364 | 1.16E-24 | 1.75E-22 | -2.32205 | 5.512986 | -41.9408 | 5.39E-22 | 6.63E-20 | 1697.652 | 1621.552 | 1518.947 | 22241.69 | 23131.79 | 21767.13 | 4604.885 | 4641.881 | 4671.317 | 6636.013 | 6924.515 | 5828.925 | 127.652 | 178.4408 | 120.6707 |
| Gja6 | 414089 | 3.707015 | -2.2644 | 5.689906 | 1.13E-05 | 4.14E-05 | 0.388495 | -2.2644 | 1.033558 | 0.312913 | 0.407821 | 2.967923 | 0.888035 | 5.207818 | 27.27289 | 42.13186 | 58.16471 | 35.53152 | 65.68067 | 67.80018 | 3.076858 | 7.625279 | 8.209754 | 10.01192 | 7.860826 | 4.049351 |
| Olfr114 | 258284 | 3.712742 | -3.33893 | 3.364394 | 0.002885 | 0.007682 | 0.506776 | -3.33893 | 0.618342 | 0.542893 | 0.637966 | 0 | 0.888035 | 0 | 7.792255 | 8.426372 | 6.763339 | 35.53152 | 10.72337 | 4.596622 | 9.999788 | 24.40089 | 11.19512 | 1.668654 | 13.3634 | 16.19741 |
| Trim30b | 244183 | 3.71897 | 3.478978 | 39.53057 | 1.88E-21 | 1.11E-19 | -0.64299 | 3.478978 | -11.6056 | 1.10E-10 | 8.57E-10 | 390.2819 | 289.4994 | 282.0901 | 4397.429 | 4732.812 | 4423.224 | 2924.955 | 3115.14 | 2961.374 | 211.534 | 220.3705 | 205.9902 | 149.3445 | 130.4897 | 150.6359 |
| Gm3285 | 100041351 | 3.725498 | -4.27046 | 3.423674 | 0.002509 | 0.006748 | 0.920856 | -4.27046 | 1.119242 | 0.275476 | 0.366887 | 0 | 0 | 0 | 2.597418 | 12.63956 | 4.058003 | 4.263783 | 8.04253 | 29.87805 | 6.153716 | 0 | 0 | 12.5149 | 9.432991 | 9.718443 |
| Hoxb6 | 15414 | 3.726033 | -1.98566 | 5.406943 | 2.19E-05 | 7.76E-05 | 2.485722 | -1.98566 | 6.220836 | 3.34E-06 | 1.33E-05 | 0.741981 | 0 | 1.735939 | 12.98709 | 16.85274 | 12.17401 | 73.90556 | 87.12741 | 89.63414 | 7.692145 | 6.100223 | 4.478048 | 85.10135 | 105.3351 | 106.9029 |
| Cnn3 | 71994 | 3.728937 | 1.610493 | 13.11357 | 1.10E-11 | 9.11E-11 | 3.038866 | 1.610493 | 28.43531 | 1.87E-18 | 7.81E-17 | 26.71131 | 39.96157 | 19.09533 | 344.1579 | 466.2592 | 374.689 | 3349.912 | 3373.842 | 3291.182 | 46.92208 | 50.32684 | 72.3951 | 100.9536 | 84.89692 | 114.1917 |
| Isg15 | 100038882 | 3.734751 | 7.343885 | 41.53368 | 8.11E-15 | 1.14E-13 | -0.39964 | 7.343885 | -7.75347 | 3.92E-06 | 1.54E-05 | 3183.097 | 3283.953 | 2639.496 | 42666.49 | 43770.79 | 42668.55 | 35818.62 | 35334.86 | 31149.01 | 7042.158 | 7379.745 | 6620.793 | 2761.622 | 2786.663 | 2801.341 |
| Olfr1322 | 257978 | 3.737081 | -4.96479 | 3.643702 | 0.001487 | 0.004157 | -0.99675 | -4.96479 | -1.1114 | 0.27876 | 0.370342 | 0 | 0 | 0 | 10.38967 | 8.426372 | 1.352668 | 0 | 2.680843 | 10.3424 | 0.769214 | 1.525056 | 0 | 3.337308 | 2.358248 | 4.859222 |
| Saa3 | 20210 | 3.73711 | 7.603851 | 41.75625 | 6.54E-13 | 6.55E-12 | -1.7262 | 7.603851 | -27.8369 | 4.26E-11 | 3.55E-10 | 8487.518 | 10074.76 | 7613.829 | 124024.1 | 124471.6 | 121011 | 41145.5 | 39372.21 | 36120.26 | 8029.83 | 7452.185 | 9660.641 | 643.2661 | 558.9047 | 647.8962 |
| Hsd3b6 | 15497 | 3.743018 | -3.61722 | 4.33898 | 0.00028 | 0.000868 | 0.186119 | -3.61722 | 0.281319 | 0.781175 | 0.853085 | 0 | 0 | 0.86797 | 7.792255 | 5.617581 | 10.82134 | 11.37009 | 13.40422 | 5.745778 | 1.538429 | 6.862751 | 1.492683 | 28.36712 | 13.3634 | 13.76779 |
| Zfhx4 | 80892 | 3.748047 | 0.458761 | 12.23486 | 4.10E-11 | 3.08E-10 | -1.0986 | 0.458761 | -6.02015 | 5.28E-06 | 2.03E-05 | 18.54952 | 30.19319 | 25.17112 | 380.5218 | 345.4812 | 316.5243 | 184.7639 | 124.6592 | 199.9531 | 13.07665 | 19.0632 | 36.57072 | 129.3207 | 117.9124 | 121.4805 |
| Mfsd7a | 243197 | 3.749996 | 3.853294 | 26.8607 | 6.11E-18 | 1.63E-16 | -0.32008 | 3.853294 | -4.20845 | 0.000383 | 0.001129 | 235.9499 | 201.5839 | 251.7112 | 3480.541 | 3343.865 | 3084.082 | 2922.112 | 2960.992 | 2430.464 | 563.8342 | 423.9655 | 532.1413 | 454.7082 | 430.7733 | 489.1616 |
| Casp12 | 12364 | 3.753278 | -0.82962 | 7.857261 | 9.81E-08 | 4.64E-07 | -0.89296 | -0.82962 | -3.1614 | 0.004642 | 0.010924 | 17.80754 | 10.65642 | 2.603909 | 129.8709 | 129.2044 | 116.3294 | 78.16935 | 45.57434 | 90.78329 | 30.76858 | 25.92595 | 32.83902 | 18.35519 | 22.7964 | 16.19741 |
| Ccl11 | 20292 | 3.757763 | -2.25318 | 3.958177 | 0.0007 | 0.002061 | 3.416265 | -2.25318 | 6.489691 | 1.83E-06 | 7.52E-06 | 0.741981 | 1.77607 | 0 | 23.37676 | 7.021976 | 16.23201 | 211.7679 | 142.0847 | 148.2411 | 6.153716 | 13.7255 | 4.478048 | 36.71038 | 14.93557 | 7.288832 |
| Olfr433 | 258712 | 3.76248 | -2.71164 | 4.843892 | 8.32E-05 | 0.000275 | -0.40874 | -2.71164 | -0.81729 | 0.422794 | 0.522013 | 5.935846 | 0 | 6.943757 | 58.44191 | 37.91867 | 28.40602 | 29.84648 | 26.80843 | 36.77298 | 7.692145 | 7.625279 | 1.492683 | 1.668654 | 2.358248 | 8.098703 |
| Fbxo2 | 230904 | 3.767457 | -3.32723 | 3.66331 | 0.001419 | 0.003981 | 1.344294 | -3.32723 | 1.973273 | 0.061553 | 0.103478 | 0 | 0 | 1.735939 | 18.18193 | 5.617581 | 8.116006 | 42.63783 | 34.85097 | 13.78987 | 0.769214 | 0 | 2.985365 | 43.385 | 40.8763 | 18.62702 |
| Vcam1 | 22329 | 3.770017 | 1.901993 | 23.89011 | 6.90E-17 | 1.46E-15 | 1.125681 | 1.901993 | 16.71947 | 9.63E-14 | 1.34E-12 | 97.94146 | 83.47529 | 80.72117 | 1201.306 | 1405.8 | 1251.218 | 2971.856 | 2895.311 | 2851.055 | 62.30637 | 45.75167 | 70.90242 | 21.6925 | 29.08506 | 36.44416 |
| Rspo2 | 239405 | 3.773336 | -2.20915 | 4.830174 | 8.59E-05 | 0.000284 | -1.71497 | -2.20915 | -2.94874 | 0.007581 | 0.016912 | 13.35565 | 3.55214 | 1.735939 | 58.44191 | 81.45493 | 83.8654 | 14.21261 | 16.08506 | 44.81707 | 1.538429 | 0 | 8.209754 | 37.54471 | 37.73197 | 8.908573 |
| Olfr341 | 258952 | 3.774852 | -5.02655 | 3.633298 | 0.001525 | 0.004254 | 0.623527 | -5.02655 | 0.778873 | 0.444605 | 0.543381 | 0 | 0 | 0 | 7.792255 | 2.808791 | 6.763339 | 2.842522 | 13.40422 | 16.08818 | 0 | 0 | 0 | 0 | 3.930413 | 6.478962 |
| Gm10319 | 381806 | 3.776208 | -4.24629 | 4.345089 | 0.000276 | 0.000856 | 0.445397 | -4.24629 | 0.706871 | 0.487293 | 0.585827 | 0 | 1.77607 | 0 | 12.98709 | 9.830767 | 6.763339 | 22.74017 | 22.78717 | 5.745778 | 0.769214 | 0 | 0 | 4.171635 | 3.930413 | 6.478962 |
| F2rl1 | 14063 | 3.778748 | -2.43209 | 4.511951 | 0.000184 | 0.000585 | 1.470434 | -2.43209 | 2.854037 | 0.009404 | 0.020454 | 1.483961 | 0.888035 | 0 | 22.07806 | 9.830767 | 13.52668 | 66.79926 | 36.19139 | 34.47467 | 4.615287 | 3.812639 | 8.209754 | 27.53279 | 62.88661 | 45.35273 |
| Cish | 12700 | 3.79209 | 3.820852 | 13.07138 | 1.17E-11 | 9.64E-11 | 4.825879 | 3.820852 | 46.97017 | 4.93E-23 | 9.47E-21 | 25.22735 | 27.52908 | 30.37894 | 483.1198 | 324.4153 | 442.3224 | 12318.07 | 12393.54 | 11824.81 | 2370.719 | 2358.499 | 2467.404 | 1254.828 | 1315.902 | 1256.109 |
| Arid5a | 214855 | 3.792178 | 5.431136 | 50.22384 | 1.20E-23 | 1.27E-21 | -1.52553 | 5.431136 | -28.5366 | 1.74E-18 | 7.32E-17 | 837.6963 | 863.17 | 808.9477 | 12980.6 | 12090.44 | 12048.21 | 4704.373 | 4538.668 | 4196.716 | 2148.416 | 2215.143 | 1883.019 | 1139.691 | 1271.096 | 1150.016 |
| Trpm6 | 225997 | 3.799731 | -0.98601 | 8.61599 | 2.16E-08 | 1.11E-07 | -1.10353 | -0.98601 | -4.10308 | 0.000494 | 0.001429 | 11.12971 | 11.54445 | 5.207818 | 135.0658 | 129.2044 | 147.4408 | 52.58665 | 65.68067 | 79.29174 | 9.999788 | 3.812639 | 5.224389 | 70.91779 | 46.37887 | 33.20468 |
| Prm1 | 19118 | 3.806241 | -2.82988 | 5.310419 | 2.74E-05 | 9.62E-05 | -0.60223 | -2.82988 | -1.27558 | 0.215814 | 0.298822 | 2.225942 | 3.55214 | 1.735939 | 59.74062 | 26.68351 | 43.28537 | 36.95278 | 20.10633 | 29.87805 | 4.615287 | 0 | 1.492683 | 3.337308 | 7.074744 | 18.62702 |
| Adam30 | 71078 | 3.806441 | -2.81891 | 4.279477 | 0.000323 | 0.000995 | 0.546876 | -2.81891 | 1.004707 | 0.326293 | 0.42246 | 0 | 5.32821 | 2.603909 | 12.98709 | 36.51428 | 39.22736 | 32.689 | 30.8297 | 59.75609 | 2.307643 | 0 | 2.239024 | 24.19548 | 17.29382 | 8.098703 |
| Tnfaip6 | 21930 | 3.815475 | -0.09679 | 10.65736 | 5.25E-10 | 3.39E-09 | 0.762494 | -0.09679 | 4.894065 | 7.38E-05 | 0.000243 | 16.32358 | 7.992315 | 13.88751 | 162.3386 | 220.4901 | 200.1948 | 335.4176 | 371.2968 | 316.0178 | 9.230573 | 9.150334 | 5.97073 | 65.91183 | 81.75259 | 88.27586 |
| Dnmt3b-ps | 668932 | 3.822288 | -1.05483 | 10.30098 | 9.69E-10 | 6.02E-09 | -2.30282 | -1.05483 | -7.7916 | 1.12E-07 | 5.46E-07 | 6.677827 | 11.54445 | 7.811726 | 131.1696 | 126.3956 | 137.9721 | 25.5827 | 22.78717 | 33.32551 | 29.23015 | 32.02617 | 37.31706 | 24.19548 | 15.72165 | 15.38754 |
| Sele | 20339 | 3.827663 | 0.060338 | 12.00228 | 5.88E-11 | 4.34E-10 | 2.457224 | 0.060338 | 20.50831 | 1.57E-15 | 3.24E-14 | 22.25942 | 10.65642 | 14.75548 | 235.0664 | 242.9604 | 244.8329 | 1422.682 | 1305.571 | 1399.672 | 3.076858 | 9.912862 | 7.463413 | 23.36115 | 28.29897 | 30.77507 |
| Il15 | 16168 | 3.829075 | 5.333139 | 59.28031 | 3.56E-25 | 6.50E-23 | 0.091534 | 5.333139 | 2.571452 | 0.017657 | 0.035401 | 678.1704 | 682.8989 | 754.2656 | 10823.44 | 10673.4 | 10534.58 | 11674.24 | 12209.9 | 11575.44 | 1569.967 | 1507.518 | 1392.673 | 612.396 | 591.9202 | 636.558 |
| Lypd11 | 210155 | 3.83226 | -4.63818 | 3.927779 | 0.000753 | 0.002206 | -0.26512 | -4.63818 | -0.33113 | 0.743778 | 0.821075 | 0 | 0 | 0 | 5.194837 | 8.426372 | 4.058003 | 9.948826 | 1.340422 | 6.894934 | 0 | 5.337695 | 0.746341 | 3.337308 | 2.358248 | 6.478962 |
| Olfr589 | 259054 | 3.838666 | -4.66736 | 4.766302 | 0.0001 | 0.000328 | 0.357724 | -4.66736 | 0.568483 | 0.57564 | 0.669404 | 0 | 0 | 0 | 7.792255 | 4.213186 | 5.410671 | 8.527565 | 4.021265 | 12.64071 | 0 | 0 | 0 | 6.674615 | 14.93557 | 5.669092 |
| Slc2a6 | 227659 | 3.842813 | 6.22211 | 40.07889 | 3.97E-19 | 1.36E-17 | -0.20369 | 6.22211 | -3.71487 | 0.001573 | 0.004129 | 1040.257 | 1046.993 | 1071.074 | 16611.79 | 16501.64 | 15289.2 | 15488.9 | 15609.21 | 12947.54 | 5737.571 | 5985.844 | 6041.632 | 1188.916 | 1157.9 | 1263.398 |
| Foxf1 | 15227 | 3.845156 | -1.0003 | 6.422122 | 2.13E-06 | 8.47E-06 | -0.14084 | -1.0003 | -0.45616 | 0.652886 | 0.739488 | 8.903769 | 7.10428 | 4.339848 | 138.9619 | 157.2923 | 55.45938 | 100.9095 | 125.9996 | 82.7392 | 19.23036 | 12.96297 | 13.43414 | 26.69846 | 24.36856 | 12.14805 |
| Nts | 67405 | 3.851097 | -3.64712 | 4.069723 | 0.000535 | 0.001602 | 0.779866 | -3.64712 | 1.097481 | 0.284659 | 0.376733 | 0 | 0 | 0 | 6.493546 | 7.021976 | 4.058003 | 5.685043 | 13.40422 | 14.93902 | 0.769214 | 4.575167 | 0 | 49.22529 | 78.60826 | 68.83897 |
| Mmp25 | 240047 | 3.856805 | 3.017939 | 30.75576 | 3.64E-19 | 1.27E-17 | 0.442834 | 3.017939 | 7.593099 | 1.69E-07 | 8.03E-07 | 113.5231 | 136.7574 | 160.5744 | 2142.87 | 2140.298 | 1973.542 | 2970.435 | 2971.715 | 2887.828 | 238.4565 | 273.7475 | 242.5609 | 129.3207 | 113.982 | 136.8681 |
| Gm12185 | 620913 | 3.857409 | 3.919991 | 40.52206 | 1.11E-21 | 6.83E-20 | 1.492412 | 3.919991 | 35.87349 | 1.45E-20 | 1.12E-18 | 229.2721 | 203.36 | 193.5572 | 3190.928 | 3376.166 | 3173.359 | 9728.531 | 9273.038 | 9439.164 | 420.7603 | 400.3271 | 397.0535 | 221.931 | 230.3222 | 244.5808 |
| 1700012B( | 69325 | 3.862512 | -3.77848 | 4.364988 | 0.000263 | 0.000819 | 1.556497 | -3.77848 | 2.672933 | 0.014117 | 0.029134 | 0 | 0.888035 | 0 | 2.597418 | 14.04395 | 16.23201 | 28.42522 | 24.12759 | 29.87805 | 0 | 1.525056 | 0.746341 | 10.01192 | 14.93557 | 7.288832 |
| Ifih1 | 71586 | 3.866639 | 8.277576 | 90.31812 | 9.04E-28 | 4.02E-25 | -1.31198 | 8.277576 | -49.8859 | 1.35E-22 | 2.22E-20 | 7247.668 | 7538.529 | 6875.187 | 111798.1 | 116375.2 | 109194.1 | 48774.83 | 47520.63 | 45127.34 | 12030.51 | 12040.31 | 11228.7 | 4889.99 | 4811.612 | 4952.357 |
| 1700048F( | 73396 | 3.868022 | -4.99349 | 4.576178 | 0.000158 | 0.000505 | -0.89104 | -4.99349 | -1.26178 | 0.220643 | 0.304546 | 0 | 0.888035 | 0 | 11.68838 | 4.213186 | 12.17401 | 2.842522 | 5.361687 | 5.745778 | 0 | 0 | 0 | 6.674615 | 0.786083 | 0.80987 |
| Misp | 78906 | 3.872257 | 0.392326 | 14.63481 | 1.33E-12 | 1.26E-11 | -1.65575 | 0.392326 | -9.61789 | 3.27E-09 | 2.00E-08 | 17.80754 | 39.07354 | 19.9633 | 384.4179 | 348.29 | 393.6263 | 116.5434 | 123.3188 | 128.7054 | 69.99852 | 75.49026 | 83.59022 | 27.53279 | 37.73197 | 37.25403 |
| Tjp1 | 21872 | 3.875218 | 4.121652 | 31.13285 | 2.82E-19 | 1.00E-17 | 0.139822 | 4.121652 | 2.264078 | 0.034099 | 0.062493 | 191.431 | 209.5763 | 197.8971 | 3202.617 | 3053.155 | 3113.841 | 3496.302 | 3617.798 | 3595.708 | 134.6125 | 165.4685 | 174.6439 | 3310.609 | 3276.392 | 3853.363 |
| 6530402F1 | 76220 | 3.877177 | 4.200875 | 40.3904 | 1.19E-21 | 7.28E-20 | 0.069534 | 4.200875 | 1.406796 | 0.173883 | 0.24928 | 268.597 | 265.5225 | 295.9776 | 4416.91 | 4197.737 | 4396.17 | 4775.436 | 4975.646 | 4469.066 | 544.6038 | 599.3469 | 592.595 | 463.0514 | 497.5903 | 502.9294 |
| Il18 | 16173 | 3.882757 | 3.592549 | 50.98458 | 8.69E-24 | 9.81E-22 | -3.01607 | 3.592549 | -43.5595 | 2.43E-22 | 3.69E-20 | 365.7965 | 456.45 | 390.5863 | 6181.856 | 6484.093 | 6275.026 | 824.3313 | 844.4657 | 767.6359 | 542.2962 | 560.458 | 591.8486 | 158.5221 | 178.4408 | 163.5938 |
| Slc28a2 | 269346 | 3.887779 | 7.894874 | 68.47362 | 9.53E-21 | 4.91E-19 | -0.32088 | 7.894874 | -11.123 | 8.12E-09 | 4.67E-08 | 6133.213 | 5954.275 | 5002.977 | 91143.41 | 90329.3 | 88333.26 | 76790.72 | 75996.55 | 72140.54 | 4415.291 | 4611.768 | 4260.116 | 3469.966 | 3340.851 | 3516.457 |
| Cpa4 | 71791 | 3.901308 | -3.19462 | 5.151513 | 4.00E-05 | 0.000137 | -0.93465 | -3.19462 | -1.58131 | 0.128514 | 0.193025 | 0 | 4.440175 | 0.86797 | 14.2858 | 28.08791 | 20.29002 | 8.527565 | 8.04253 | 16.08818 | 5.384501 | 1.525056 | 1.492683 | 15.01788 | 16.50774 | 10.52831 |
| F830016B( | 240328 | 3.917214 | 3.471318 | 24.37187 | 4.57E-17 | 1.00E-15 | 0.900185 | 3.471318 | 12.76432 | 1.84E-11 | 1.65E-10 | 132.8146 | 143.8617 | 143.215 | 2301.313 | 2269.503 | 2196.732 | 4418.7 | 4603.008 | 4133.513 | 917.6728 | 906.6456 | 874.7119 | 60.90587 | 135.2062 | 129.5792 |
| Olfr805 | 258548 | 3.923737 | -4.8262 | 4.544793 | 0.00017 | 0.000543 | -0.77895 | -4.8262 | -1.0688 | 0.297101 | 0.390762 | 0 | 0 | 0 | 7.792255 | 2.808791 | 9.468674 | 2.842522 | 4.021265 | 3.447467 | 0 | 0 | 1.492683 | 6.674615 | 2.358248 | 8.908573 |
| 5230400M( | 100502767 | 3.923798 | 1.621241 | 16.62734 | 1.07E-13 | 1.24E-12 | 1.033822 | 1.621241 | 10.89269 | 3.53E-10 | 2.52E-09 | 35.61508 | 36.40943 | 42.53051 | 594.8088 | 615.1251 | 656.0439 | 1192.438 | 1490.549 | 1293.949 | 52.30658 | 41.93903 | 52.24389 | 118.4744 | 142.281 | 147.3964 |
| Slco3a1 | 108116 | 3.925475 | 6.665524 | 65.2117 | 4.70E-26 | 1.19E-23 | -0.75881 | 6.665524 | -21.4096 | 6.55E-16 | 1.48E-14 | 2434.439 | 2639.24 | 2450.278 | 40722.32 | 41390.34 | 39745.44 | 26185.31 | 25967.99 | 23057.81 | 2690.712 | 2601.745 | 2762.955 | 1166.389 | 1139.82 | 1306.321 |
| Hcar2 | 80885 | 3.92635 | 3.908059 | 57.98995 | 5.67E-25 | 9.69E-23 | -0.95262 | 3.908059 | -22.7878 | 1.82E-16 | 4.72E-15 | 606.9403 | 560.3501 | 537.2732 | 9213.043 | 9269.009 | 9263.069 | 5147.807 | 4860.369 | 4887.359 | 442.2983 | 411.765 | 386.6048 | 72.58644 | 50.30929 | 42.11325 |
| Trp53i11 | 277414 | 3.927891 | 2.365856 | 33.80326 | 5.04E-20 | 2.12E-18 | -3.06318 | 2.365856 | -29.199 | 1.08E-18 | 4.80E-17 | 135.0405 | 146.5258 | 148.4228 | 2431.184 | 2261.076 | 2284.656 | 291.3585 | 274.7865 | 297.6313 | 326.9161 | 322.5493 | 297.0438 | 95.9476 | 73.89177 | 106.093 |
| Phf11a | 219131 | 3.945552 | 3.499555 | 50.29759 | 1.16E-23 | 1.24E-21 | -3.1292 | 3.499555 | -43.2464 | 2.82E-22 | 4.06E-20 | 483.0295 | 521.2765 | 468.7036 | 8081.867 | 8402.497 | 7731.849 | 905.3432 | 1036.146 | 933.1143 | 1200.744 | 1227.67 | 1049.356 | 30.03577 | 27.51289 | 38.87377 |
| Il36a | 54448 | 3.945696 | -0.85349 | 13.08806 | 1.15E-11 | 9.42E-11 | 0.955233 | -0.85349 | 7.837187 | 1.02E-07 | 5.01E-07 | 20.77546 | 22.20087 | 27.77503 | 403.8985 | 394.6351 | 369.2783 | 847.0715 | 682.2747 | 828.5412 | 0 | 1.525056 | 1.492683 | 1.668654 | 7.860826 | 6.478962 |
| Ccl8 | 20307 | 3.9531 | 1.858231 | 12.12307 | 4.88E-11 | 3.63E-10 | 0.237623 | 1.858231 | 1.657313 | 0.112084 | 0.172021 | 23.0014 | 19.53677 | 19.9633 | 319.4825 | 334.2461 | 408.5057 | 432.0633 | 443.6796 | 418.2926 | 2045.341 | 2189.98 | 2002.434 | 57.56856 | 58.17011 | 106.093 |
| Tent5d | 213449 | 3.953751 | -2.49212 | 6.182272 | 3.65E-06 | 1.42E-05 | -0.24244 | -2.49212 | -0.65552 | 0.519138 | 0.61576 | 3.709904 | 3.55214 | 1.735939 | 41.55869 | 68.81537 | 55.45938 | 59.69296 | 38.87223 | 45.96622 | 2.307643 | 3.812639 | 0 | 10.84625 | 4.716496 | 15.38754 |
| Meox2 | 17286 | 3.955306 | -3.64756 | 3.561164 | 0.001811 | 0.004994 | 0.409407 | -3.64756 | 0.521499 | 0.607389 | 0.699704 | 1.483961 | 0 | 0 | 18.18193 | 18.25714 | 4.058003 | 25.5827 | 6.702109 | 21.83396 | 0 | 0 | 10.44878 | 10.84625 | 25.15464 | 12.14805 |
| Ch25h | 12642 | 3.95678 | 6.462138 | 85.96491 | 1.32E-28 | 1.16E-25 | 0.412299 | 6.462138 | 18.39625 | 1.42E-14 | 2.35E-13 | 2388.436 | 2539.78 | 2269.741 | 39928.81 | 40470.46 | 38799.92 | 55964.99 | 53308.57 | 55337.59 | 995.3635 | 1026.362 | 927.7022 | 746.7226 | 739.7037 | 897.3363 |
| Olfr1196 | 258456 | 3.96346 | -5.21793 | 4.117388 | 0.000477 | 0.001438 | -2.7153 | -5.21793 | -2.9385 | 0.007761 | 0.017272 | 0 | 0 | 0 | 5.194837 | 4.213186 | 10.82134 | 2.842522 | 0 | 0 | 0 | 3.050111 | 0 | 4.171635 | 7.860826 | 1.619741 |
| Vmn1r193 | 171259 | 3.973924 | -5.21306 | 3.87374 | 0.000857 | 0.002491 | -0.82016 | -5.21306 | -0.94767 | 0.353905 | 0.451233 | 0 | 0 | 0 | 10.38967 | 1.404395 | 13.52668 | 1.421261 | 5.361687 | 4.596622 | 0 | 3.050111 | 0 | 1.668654 | 1.572165 | 0 |
| Syt17 | 110058 | 3.977002 | -2.6226 | 4.166843 | 0.000424 | 0.001286 | 2.448089 | -2.6226 | 4.555659 | 0.000166 | 0.000519 | 0 | 0 | 4.339848 | 7.792255 | 25.27911 | 16.23201 | 103.752 | 124.6592 | 54.01031 | 1.538429 | 3.050111 | 4.478048 | 22.52683 | 23.58248 | 17.00728 |
| Gbp11 | 634650 | 3.97845 | 0.122462 | 9.275909 | 6.13E-09 | 3.40E-08 | 1.790115 | 0.122462 | 10.79473 | 4.16E-10 | 2.94E-09 | 9.64575 | 8.88035 | 9.547666 | 138.9619 | 171.3362 | 186.6681 | 606.8784 | 601.8494 | 567.6829 | 78.45987 | 62.52728 | 36.57072 | 23.36115 | 23.58248 | 17.00728 |
| Mxd1 | 17119 | 3.983351 | 7.390465 | 77.01417 | 1.37E-27 | 5.45E-25 | -0.13187 | 7.390465 | -4.93214 | 6.74E-05 | 0.000223 | 2766.104 | 2616.151 | 2951.097 | 47732.76 | 46672.27 | 46192.25 | 45284.21 | 45186.96 | 43050.82 | 3569.924 | 3901.092 | 3383.165 | 4873.304 | 4711.779 | 4982.322 |
| Igkv4-70 | 385120 | 3.983856 | -5.15496 | 4.217281 | 0.000375 | 0.001147 | -2.51606 | -5.15496 | -2.80432 | 0.010522 | 0.022581 | 0 | 0 | 0 | 7.792255 | 2.808791 | 10.82134 | 0 | 0 | 3.447467 | 0 | 0 | 2.985365 | 3.337308 | 7.074744 | 3.239481 |
| Nod1 | 107607 | 3.986035 | 6.235402 | 55.08083 | 1.69E-24 | 2.47E-22 | -0.3751 | 6.235402 | -9.13472 | 7.99E-09 | 4.60E-08 | 1016.514 | 1180.198 | 1028.544 | 18280.63 | 18632.11 | 17419.66 | 14916.13 | 15217.81 | 13540.5 | 3098.396 | 3079.087 | 2859.233 | 2170.919 | 2231.689 | 2133.198 |
| Csf1 | 12977 | 3.991669 | 5.488941 | 46.85507 | 5.19E-23 | 4.54E-21 | 2.109122 | 5.488941 | 57.53067 | 6.72E-25 | 2.13E-22 | 423.671 | 516.8364 | 472.1755 | 7833.814 | 8031.737 | 7941.512 | 36307.53 | 35988.98 | 34513.74 | 1397.663 | 1488.454 | 1374.014 | 775.924 | 703.5439 | 741.0313 |
| Aanat | 11298 | 3.997754 | 1.578187 | 21.81108 | 4.48E-16 | 7.99E-15 | -1.95108 | 1.578187 | -16.037 | 2.20E-13 | 2.82E-12 | 47.48677 | 65.71459 | 78.11726 | 1055.851 | 990.0987 | 1129.478 | 265.7758 | 284.1694 | 297.6313 | 133.8433 | 164.706 | 142.5512 | 76.75808 | 64.45877 | 55.88105 |
| Slamf7 | 75345 | 3.999556 | 7.846066 | 103.9686 | 2.30E-30 | 6.72E-27 | -0.12103 | 7.846066 | -6.35965 | 2.45E-06 | 9.89E-06 | 4037.859 | 4013.918 | 3912.807 | 69070.55 | 69052.71 | 66325.36 | 66688.4 | 62566.87 | 65859.26 | 9287.495 | 9312.753 | 8867.28 | 3010.252 | 2971.392 | 3148.776 |
| Ifi213 | 623121 | 4.01688 | 5.026971 | 62.57226 | 1.13E-25 | 2.48E-23 | -1.11511 | 5.026971 | -27.5483 | 3.62E-18 | 1.39E-16 | 808.017 | 858.7298 | 720.4148 | 13518.26 | 14100.13 | 13584.84 | 6443.997 | 6482.28 | 6774.272 | 1743.809 | 1953.596 | 1715.839 | 241.1205 | 199.665 | 213.8058 |
| Cyp2j13 | 230459 | 4.018753 | -2.9067 | 4.40201 | 0.00024 | 0.000752 | 0.724216 | -2.9067 | 1.243097 | 0.227315 | 0.312232 | 0 | 0 | 6.075787 | 14.2858 | 22.47032 | 14.87935 | 22.74017 | 24.12759 | 42.51876 | 2.307643 | 0.762528 | 6.717071 | 20.85817 | 34.58764 | 13.76779 |
| Gpr15 | 71223 | 4.020565 | -2.59207 | 6.466282 | 1.93E-06 | 7.71E-06 | -2.36149 | -2.59207 | -4.52638 | 0.000178 | 0.000555 | 1.483961 | 1.77607 | 12.15157 | 50.64966 | 77.24174 | 64.92805 | 8.527565 | 9.382952 | 20.6848 | 3.846072 | 3.812639 | 5.97073 | 10.01192 | 2.358248 | 9.718443 |
| Olfr1248 | 258787 | 4.033976 | -4.76588 | 3.710328 | 0.001268 | 0.00359 | -1.21066 | -4.76588 | -1.28814 | 0.211489 | 0.293931 | 0 | 0 | 0 | 2.597418 | 7.021976 | 14.87935 | 2.842522 | 0 | 12.64071 | 0 | 3.050111 | 0 | 10.01192 | 3.930413 | 6.478962 |
| Astl | 215095 | 4.035187 | -0.00646 | 11.22369 | 2.04E-10 | 1.40E-09 | -1.77917 | -0.00646 | -7.84333 | 1.01E-07 | 4.95E-07 | 31.16319 | 10.65642 | 23.43518 | 385.7166 | 286.4966 | 431.501 | 109.4371 | 107.2337 | 111.4681 | 10.769 | 25.16342 | 27.61463 | 45.05365 | 51.09537 | 41.30338 |
| Cxcl2 | 20310 | 4.040087 | 5.552062 | 46.06973 | 7.42E-23 | 6.08E-21 | 2.954872 | 5.552062 | 83.34279 | 2.55E-28 | 9.58E-25 | 405.1215 | 487.5312 | 460.8919 | 7763.683 | 7634.293 | 8203.93 | 64253.78 | 61026.72 | 64465.33 | 1660.734 | 1734.751 | 1853.912 | 447.1992 | 411.1212 | 472.1544 |
| Pnp2 | 667034 | 4.046942 | 2.777972 | 23.05943 | 1.43E-16 | 2.86E-15 | 1.075984 | 2.777972 | 15.3376 | 5.30E-13 | 6.26E-12 | 107.5872 | 96.79581 | 55.55005 | 1601.308 | 1431.079 | 1451.412 | 3341.384 | 3304.14 | 3180.863 | 105.3824 | 105.9914 | 87.32193 | 292.0144 | 286.1341 | 272.9263 |
| Tnp2 | 21959 | 4.049038 | -2.98981 | 5.168566 | 3.84E-05 | 0.000132 | 0.73815 | -2.98981 | 1.502089 | 0.147726 | 0.217203 | 0.741981 | 0 | 1.735939 | 7.792255 | 18.25714 | 37.8747 | 25.5827 | 33.51054 | 33.32551 | 6.153716 | 3.050111 | 5.97073 | 8.343269 | 10.21907 | 4.859222 |
| Gbp9 | 236573 | 4.064849 | 7.234291 | 64.94723 | 5.12E-26 | 1.25E-23 | 0.77555 | 7.234291 | 26.42709 | 8.57E-18 | 2.99E-16 | 1570.031 | 1448.385 | 1465.133 | 26832.63 | 27375.88 | 26078.08 | 48575.85 | 47587.65 | 46663.76 | 3517.618 | 3761.55 | 3112.243 | 9065.796 | 8811.986 | 9081.075 |
| Apol11b | 328563 | 4.071079 | -4.01694 | 3.558077 | 0.001824 | 0.005028 | 0.832146 | -4.01694 | 1.006286 | 0.325551 | 0.421643 | 0 | 0 | 0 | 20.77935 | 1.404395 | 8.116006 | 11.37009 | 17.42548 | 11.49156 | 0 | 7.625279 | 0 | 22.52683 | 18.86598 | 8.098703 |
| Gm4945 | 240110 | 4.089959 | -4.78369 | 3.723164 | 0.00123 | 0.00349 | -1.25109 | -4.78369 | -1.32083 | 0.200547 | 0.281206 | 0 | 0 | 0 | 10.38967 | 5.617581 | 5.410671 | 0 | 6.702109 | 5.745778 | 1.538429 | 1.525056 | 0 | 15.01788 | 0.786083 | 4.049351 |
| Tnfrsf8 | 21941 | 4.103895 | 0.788292 | 15.87719 | 2.68E-13 | 2.87E-12 | -0.12051 | 0.788292 | -1.04542 | 0.307526 | 0.40198 | 20.77546 | 22.20087 | 19.9633 | 420.7818 | 369.356 | 384.1576 | 366.6853 | 369.9564 | 382.6688 | 83.84438 | 99.12862 | 70.15608 | 45.88798 | 44.02063 | 42.92312 |
| Ism1 | 319909 | 4.104417 | 0.009086 | 13.40963 | 7.20E-12 | 6.09E-11 | -1.63173 | 0.009086 | -8.86605 | 1.33E-08 | 7.38E-08 | 25.22735 | 16.87266 | 28.643 | 397.405 | 463.4504 | 442.3224 | 119.3859 | 152.8081 | 160.8818 | 15.38429 | 14.48803 | 30.59999 | 44.21933 | 17.29382 | 26.72572 |
| Ntng2 | 171171 | 4.111634 | 3.755036 | 30.63636 | 3.95E-19 | 1.36E-17 | -0.12474 | 3.755036 | -1.9298 | 0.06703 | 0.111101 | 218.1423 | 182.0472 | 164.9142 | 3697.425 | 3303.138 | 3465.535 | 3378.337 | 3596.352 | 3047.561 | 480.759 | 529.1943 | 467.2096 | 302.8607 | 344.3042 | 310.9902 |
| Smtnl1 | 68678 | 4.117421 | -1.68216 | 5.315263 | 2.71E-05 | 9.52E-05 | -0.57393 | -1.68216 | -1.28783 | 0.211596 | 0.294065 | 0 | 10.65642 | 17.35939 | 88.31222 | 61.79339 | 67.63339 | 52.58665 | 41.55307 | 56.30862 | 10.769 | 8.387806 | 2.985365 | 19.18952 | 24.36856 | 19.43689 |
| Cd274 | 60533 | 4.120139 | 9.226959 | 115.0346 | 2.67E-31 | 1.25E-27 | -0.55936 | 9.226959 | -32.4229 | 1.21E-19 | 7.22E-18 | 7148.243 | 7240.149 | 6538.415 | 129525.5 | 133059.4 | 126084.9 | 91723.91 | 90072.32 | 91771.57 | 33210.06 | 33409.4 | 31726.97 | 21633.26 | 21821.65 | 22111.08 |
| Lum | 17022 | 4.124455 | -0.93842 | 11.99268 | 5.97E-11 | 4.40E-10 | -4.5754 | -0.93842 | -10.5026 | 6.84E-10 | 4.67E-09 | 29.67923 | 15.09659 | 10.41564 | 348.0541 | 266.8351 | 376.0416 | 11.37009 | 10.72337 | 19.53565 | 19.99958 | 15.25056 | 20.15121 | 17.52087 | 25.15464 | 17.00728 |
| Gm7759 | 665725 | 4.133318 | -4.55283 | 4.81854 | 8.84E-05 | 0.000291 | 0.08982 | -4.55283 | 0.135985 | 0.893107 | 0.918927 | 0 | 0 | 0 | 3.896127 | 8.426372 | 10.82134 | 9.948826 | 5.361687 | 9.193245 | 0 | 0 | 2.985365 | 5.005962 | 5.502578 | 3.239481 |
| Ccl3 | 20302 | 4.147718 | 8.225407 | 151.1087 | 7.95E-34 | 2.09E-29 | -2.86833 | 8.225407 | -129 | 2.32E-32 | 6.10E-28 | 12707.9 | 12580.79 | 11931.98 | 236062.5 | 239773.8 | 229216.3 | 33320.04 | 32613.8 | 34136.82 | 16938.1 | 16574.31 | 16561.31 | 1087.962 | 1025.052 | 1290.933 |
| Batf2 | 74481 | 4.148819 | 4.839567 | 45.13844 | 1.14E-22 | 8.95E-21 | 1.376339 | 4.839567 | 34.83347 | 2.69E-20 | 1.94E-18 | 327.9555 | 321.4687 | 342.848 | 6307.83 | 6315.566 | 6183.044 | 17528.41 | 17011.29 | 16274.34 | 759.9839 | 775.4908 | 764.2534 | 488.9156 | 602.1393 | 548.2822 |
| Trim30c | 434219 | 4.151117 | 4.708271 | 66.63904 | 2.97E-26 | 8.48E-24 | -0.57825 | 4.708271 | -16.7386 | 9.41E-14 | 1.31E-12 | 552.7757 | 527.4928 | 514.706 | 10040.32 | 10207.14 | 10081.43 | 7211.478 | 6873.683 | 7004.103 | 1513.814 | 1537.256 | 1528.507 | 148.5102 | 188.6598 | 149.0161 |
| Lipk | 240633 | 4.151686 | -2.4271 | 5.185039 | 3.69E-05 | 0.000127 | 0.674583 | -2.4271 | 1.348773 | 0.191548 | 0.270404 | 0 | 6.216245 | 0 | 16.88322 | 16.85274 | 21.64268 | 22.74017 | 33.51054 | 36.77298 | 14.61507 | 18.30067 | 8.956095 | 20.85817 | 18.86598 | 20.24676 |
| A530040E: | 621875 | 4.164366 | 3.664622 | 40.23208 | 1.30E-21 | 7.84E-20 | -0.73366 | 3.664622 | -13.5338 | 6.03E-12 | 5.89E-11 | 212.9485 | 226.4489 | 191.8213 | 4218.207 | 3933.711 | 3926.794 | 2575.325 | 2600.418 | 2386.796 | 542.2962 | 597.8218 | 556.7706 | 216.0907 | 212.2423 | 193.559 |
| Klrk1 | 27007 | 4.16522 | 0.29067 | 19.37505 | 4.99E-15 | 7.28E-14 | -2.2508 | 0.29067 | -15.4401 | 4.65E-13 | 5.56E-12 | 51.93865 | 34.63336 | 41.66254 | 918.1874 | 834.2108 | 729.0879 | 170.5513 | 168.8931 | 196.5056 | 32.30701 | 39.65145 | 33.58536 | 14.18356 | 7.860826 | 6.478962 |
| Frmpd3 | 245643 | 4.171745 | -3.40696 | 4.964725 | 6.23E-05 | 0.000209 | 0.025611 | -3.40696 | 0.045788 | 0.963905 | 0.970324 | 1.483961 | 0 | 0.86797 | 14.2858 | 19.66153 | 25.70069 | 11.37009 | 42.8935 | 17.23733 | 3.076858 | 1.525056 | 0 | 8.343269 | 14.93557 | 6.478962 |
| Gm19521 | 100503038 | 4.171869 | -3.37068 | 4.558764 | 0.000165 | 0.000526 | 0.323357 | -3.37068 | 0.519462 | 0.608785 | 0.700999 | 0 | 2.664105 | 0 | 12.98709 | 28.08791 | 8.116006 | 11.37009 | 18.7659 | 28.72889 | 0 | 3.050111 | 1.492683 | 20.85817 | 28.29897 | 14.57766 |
| Ccl4 | 20303 | 4.182658 | 8.127998 | 141.0763 | 3.44E-33 | 4.52E-29 | -2.77253 | 8.127998 | -118.278 | 1.47E-31 | 1.94E-27 | 9466.932 | 9479.773 | 8557.312 | 178389.4 | 180758.3 | 174289.9 | 27286.79 | 27139.52 | 26651.22 | 13973.55 | 14003.06 | 14008.08 | 2028.249 | 2034.382 | 2317.849 |
| A530032D | 381287 | 4.192018 | 2.42226 | 24.20122 | 5.29E-17 | 1.14E-15 | -0.58224 | 2.42226 | -7.08084 | 5.01E-07 | 2.22E-06 | 83.84382 | 105.6762 | 83.32508 | 1675.335 | 1766.729 | 1853.155 | 1277.714 | 1313.613 | 1104.339 | 203.8418 | 195.9697 | 148.5219 | 85.10135 | 89.61342 | 109.3325 |
| Mrgpra3 | 233222 | 4.199569 | -4.20063 | 4.256944 | 0.000341 | 0.001047 | 0.777384 | -4.20063 | 1.103638 | 0.282039 | 0.373774 | 0 | 0 | 0 | 9.090964 | 2.808791 | 14.87935 | 12.79135 | 10.72337 | 18.38649 | 0 | 2.287584 | 2.985365 | 1.668654 | 4.716496 | 11.33818 |
| Gm16525 | 100036569 | 4.200335 | -3.80936 | 4.585501 | 0.000155 | 0.000494 | -2.42472 | -3.80936 | -2.99269 | 0.006855 | 0.015461 | 2.967923 | 0.888035 | 0 | 40.25998 | 11.23516 | 29.75869 | 5.685043 | 2.680843 | 4.596622 | 6.153716 | 0 | 1.492683 | 2.502981 | 3.930413 | 14.57766 |
| Mir691 | 751561 | 4.201794 | -5.54052 | 4.494956 | 0.000192 | 0.000608 | -2.90292 | -5.54052 | -3.23859 | 0.003877 | 0.009309 | 0 | 0 | 0 | 11.68838 | 4.213186 | 8.116006 | 0 | 0 | 2.298311 | 0 | 0 | 2.985365 | 1.668654 | 1.572165 | 0 |
| Lpar1 | 14745 | 4.2023 | 3.671782 | 34.64743 | 3.01E-20 | 1.34E-18 | -0.49547 | 3.671782 | -8.07715 | 6.27E-08 | 3.17E-07 | 253.7574 | 272.6267 | 172.726 | 4481.845 | 4744.047 | 4342.063 | 3648.377 | 3200.927 | 3178.564 | 273.0711 | 289.7606 | 273.1609 | 274.4936 | 264.9098 | 258.3486 |
| Gstt4 | 75886 | 4.210474 | -1.24497 | 7.187691 | 3.98E-07 | 1.74E-06 | 0.866473 | -1.24497 | 3.456135 | 0.002323 | 0.005877 | 5.193865 | 1.77607 | 9.547666 | 127.2735 | 127.8 | 58.16471 | 180.5001 | 215.8079 | 170.075 | 12.30743 | 12.96297 | 15.67317 | 8.343269 | 10.21907 | 4.859222 |
| Abtb2 | 99382 | 4.212993 | 4.879452 | 53.3273 | 3.36E-24 | 4.41E-22 | -0.72956 | 4.879452 | -17.0982 | 6.16E-14 | 8.95E-13 | 347.989 | 319.6926 | 336.7722 | 6684.456 | 6742.502 | 6522.564 | 4327.739 | 4188.818 | 4000.211 | 1123.822 | 1146.842 | 1171.009 | 1561.026 | 1520.284 | 1665.093 |
| Rnf133 | 386611 | 4.215014 | -3.72133 | 5.345719 | 2.53E-05 | 8.90E-05 | -0.67107 | -3.72133 | -1.14493 | 0.264914 | 0.354941 | 2.967923 | 0 | 0 | 20.77935 | 16.85274 | 13.52668 | 9.948826 | 16.08506 | 8.044089 | 3.076858 | 1.525056 | 1.492683 | 6.674615 | 7.074744 | 1.619741 |
| Clec4e | 56619 | 4.216528 | 7.407106 | 84.64183 | 1.83E-28 | 1.34E-25 | 1.236545 | 7.407106 | 61.95357 | 1.40E-25 | 5.82E-23 | 2720.101 | 2795.534 | 2734.972 | 54378.25 | 56921.54 | 52552.49 | 133990.8 | 128832 | 136988.5 | 2023.803 | 1897.932 | 1967.356 | 2633.97 | 2641.238 | 2948.738 |
| Tgtp2 | 100039796 | 4.224522 | 6.184343 | 47.43851 | 5.14E-19 | 1.72E-17 | -0.32914 | 6.184343 | -7.05804 | 2.33E-06 | 9.45E-06 | 1682.812 | 1721.9 | 1142.248 | 29671.61 | 30486.61 | 29471.92 | 25429.2 | 24689.23 | 24052.98 | 3019.167 | 3294.883 | 2879.385 | 455.5425 | 474.0078 | 451.9076 |
| Gbp7 | 229900 | 4.228348 | 7.902907 | 99.75931 | 5.55E-30 | 1.33E-26 | -0.05151 | 7.902907 | -2.58521 | 0.017132 | 0.03451 | 2973.117 | 3263.529 | 3076.952 | 61922.45 | 63499.73 | 60569.76 | 62697.5 | 62062.87 | 61572.91 | 6769.856 | 6955.779 | 6518.545 | 7463.054 | 7753.133 | 7107.421 |
| Vmn2r2 | 100125586 | 4.228572 | -4.45079 | 3.578859 | 0.001736 | 0.0048 | -0.28986 | -4.45079 | -0.30927 | 0.760114 | 0.834967 | 0 | 0 | 0 | 7.792255 | 8.426372 | 6.763339 | 0 | 13.40422 | 28.72889 | 0 | 0 | 1.492683 | 12.5149 | 5.502578 | 13.76779 |
| Gm10406 | 100038847 | 4.229469 | -4.61083 | 5.719464 | 1.05E-05 | 3.88E-05 | -0.74233 | -4.61083 | -1.22844 | 0.232657 | 0.318256 | 0 | 0 | 0 | 7.792255 | 7.021976 | 8.116006 | 5.685043 | 2.680843 | 5.745778 | 0.769214 | 0 | 0.746341 | 5.005962 | 5.502578 | 7.288832 |
| Macc1 | 238455 | 4.229789 | -1.82106 | 8.160548 | 5.30E-08 | 2.60E-07 | 0.432088 | -1.82106 | 1.867366 | 0.075646 | 0.12317 | 5.193865 | 7.992315 | 2.603909 | 87.01351 | 108.1384 | 120.3874 | 123.6497 | 182.2974 | 137.8987 | 0.769214 | 0 | 0 | 22.52683 | 43.23454 | 30.77507 |
| Bmp10 | 12154 | 4.230347 | -2.51224 | 5.449793 | 1.98E-05 | 7.06E-05 | -0.59969 | -2.51224 | -1.28506 | 0.212543 | 0.29521 | 5.935846 | 5.32821 | 0 | 48.05224 | 51.96262 | 51.40137 | 32.689 | 44.23392 | 28.72889 | 4.615287 | 3.050111 | 0 | 12.5149 | 23.58248 | 10.52831 |
| Fbn1 | 14118 | 4.231631 | 2.857112 | 23.82049 | 7.33E-17 | 1.55E-15 | -0.04504 | 2.857112 | -0.58596 | 0.564051 | 0.658418 | 97.19948 | 85.25136 | 77.24929 | 1651.958 | 1856.611 | 1743.589 | 1860.43 | 1713.059 | 1712.242 | 356.9155 | 396.5145 | 378.395 | 148.5102 | 202.8093 | 114.1917 |
| Gm906 | 380882 | 4.234096 | -4.75353 | 3.736208 | 0.001192 | 0.00339 | -1.68678 | -4.75353 | -1.6916 | 0.105273 | 0.163176 | 0 | 0 | 0 | 6.493546 | 5.617581 | 12.17401 | 0 | 1.340422 | 11.49156 | 1.538429 | 3.050111 | 1.492683 | 11.68058 | 3.14433 | 0 |
| Il36rn | 54450 | 4.242025 | -3.50582 | 4.54586 | 0.00017 | 0.000541 | 0.055569 | -3.50582 | 0.084428 | 0.933503 | 0.945656 | 0 | 1.77607 | 0 | 15.58451 | 7.021976 | 21.64268 | 9.948826 | 17.42548 | 17.23733 | 0.769214 | 5.337695 | 0.746341 | 20.02385 | 20.43815 | 5.669092 |
| Steap4 | 117167 | 4.242216 | 0.498364 | 12.10319 | 5.03E-11 | 3.74E-10 | 0.264041 | 0.498364 | 1.873401 | 0.074773 | 0.121944 | 27.45329 | 24.86498 | 15.62345 | 474.0288 | 470.4724 | 427.443 | 514.4964 | 625.977 | 571.1303 | 45.38365 | 24.40089 | 23.88292 | 29.20144 | 44.02063 | 10.52831 |
| Ms4a4c | 64380 | 4.243877 | 5.996518 | 82.33074 | 3.31E-28 | 2.07E-25 | -1.2109 | 5.996518 | -39.0774 | 2.39E-21 | 2.38E-19 | 1797.819 | 1807.151 | 1807.113 | 36624.9 | 36448.27 | 36420.58 | 15932.33 | 16081.04 | 16984.52 | 3060.704 | 3155.34 | 2655.482 | 285.3398 | 253.1186 | 229.1933 |
| Epb41l4b | 54357 | 4.244801 | 0.782697 | 16.12291 | 1.98E-13 | 2.17E-12 | -0.61679 | 0.782697 | -5.07904 | 4.75E-05 | 0.000161 | 48.22875 | 43.51371 | 50.34224 | 1036.37 | 853.8723 | 1014.501 | 632.4611 | 612.5727 | 710.1782 | 13.07665 | 9.150334 | 27.61463 | 32.53875 | 43.23454 | 10.52831 |
| Efna3 | 13638 | 4.245428 | -2.97074 | 5.964515 | 5.99E-06 | 2.27E-05 | -1.54329 | -2.97074 | -2.79064 | 0.010852 | 0.023185 | 0.741981 | 1.77607 | 0.86797 | 33.76644 | 25.27911 | 28.40602 | 22.74017 | 8.04253 | 5.745778 | 0.769214 | 1.525056 | 5.97073 | 27.53279 | 26.72681 | 14.57766 |
| Gm19551 | 100503094 | 4.248101 | 3.755586 | 34.74333 | 2.84E-20 | 1.29E-18 | -0.0704 | 3.755586 | -1.21353 | 0.238188 | 0.324724 | 224.0782 | 265.5225 | 196.1611 | 4535.092 | 4801.627 | 4478.683 | 4744.169 | 4773.242 | 4202.462 | 190.7652 | 140.3051 | 146.2829 | 442.1933 | 491.3016 | 430.851 |
| Isg20 | 57444 | 4.249125 | 4.829669 | 50.38304 | 1.12E-23 | 1.20E-21 | 0.430115 | 4.829669 | 10.76798 | 4.35E-10 | 3.06E-09 | 408.8314 | 526.6047 | 472.1755 | 9615.643 | 9283.053 | 9340.171 | 13278.84 | 13535.58 | 12755.63 | 681.524 | 629.0855 | 654.5413 | 409.6545 | 327.0104 | 387.9279 |
| Dll4 | 54485 | 4.252219 | -0.4441 | 12.99379 | 1.31E-11 | 1.07E-10 | -0.54984 | -0.4441 | -3.5983 | 0.001657 | 0.004327 | 15.5816 | 10.65642 | 13.88751 | 300.0018 | 271.0483 | 266.4755 | 207.5041 | 197.042 | 190.7598 | 2.307643 | 3.050111 | 9.702436 | 52.5626 | 49.52321 | 40.49351 |
| Ms4a4d | 66607 | 4.257471 | 1.471056 | 23.55267 | 9.26E-17 | 1.91E-15 | -1.944 | 1.471056 | -17.4897 | 3.92E-14 | 5.98E-13 | 60.10044 | 92.35564 | 100.6845 | 1593.516 | 1718.98 | 1681.366 | 466.1736 | 455.7434 | 427.4859 | 86.15202 | 89.21576 | 86.57559 | 45.88798 | 13.3634 | 18.62702 |
| Apol9b | 71898 | 4.29425 | 2.888451 | 22.29104 | 2.87E-16 | 5.40E-15 | 1.691459 | 2.888451 | 24.66039 | 3.59E-17 | 1.09E-15 | 51.93865 | 79.92315 | 67.70163 | 1305.203 | 1369.285 | 1417.596 | 4552.299 | 4747.774 | 4427.696 | 221.5338 | 250.8717 | 202.2585 | 222.7653 | 162.7191 | 170.8826 |
| Ifng | 15978 | 4.296229 | -3.81324 | 5.124985 | 4.26E-05 | 0.000146 | 0.567605 | -3.81324 | 0.946876 | 0.3543 | 0.451665 | 0 | 0 | 0 | 7.792255 | 9.830767 | 6.763339 | 9.948826 | 26.80843 | 8.044089 | 1.538429 | 3.050111 | 7.463413 | 9.177596 | 5.502578 | 7.288832 |
| Serpine1 | 18787 | 4.297346 | 4.700072 | 26.38746 | 9.70E-16 | 1.62E-14 | 3.829613 | 4.700072 | 65.87814 | 9.53E-23 | 1.66E-20 | 330.1814 | 286.8353 | 273.4104 | 6135.102 | 6262.198 | 6373.77 | 96341.59 | 93141.89 | 88609.09 | 72.30616 | 46.5142 | 52.24389 | 959.476 | 862.3326 | 986.422 |
| Selp | 20344 | 4.297422 | 0.193341 | 13.62646 | 5.29E-12 | 4.56E-11 | -1.76928 | 0.193341 | -9.58616 | 3.46E-09 | 2.11E-08 | 22.25942 | 13.32052 | 15.62345 | 353.2489 | 384.8043 | 354.3989 | 157.76 | 104.5529 | 86.18667 | 68.46009 | 57.18959 | 63.43901 | 34.2074 | 36.1598 | 29.15533 |
| Olfr1273-p | 258977 | 4.300009 | -5.00004 | 3.987368 | 0.000653 | 0.00193 | -1.05111 | -5.00004 | -1.17315 | 0.253663 | 0.342203 | 0 | 0 | 0 | 5.194837 | 23.87472 | 4.058003 | 1.421261 | 6.702109 | 4.596622 | 0 | 0.762528 | 0 | 0 | 7.860826 | 4.859222 |
| Olfr301 | 257958 | 4.315532 | -5.15658 | 4.753402 | 0.000103 | 0.000337 | -2.32357 | -5.15658 | -2.80215 | 0.010574 | 0.022679 | 0 | 0 | 0 | 9.090964 | 4.213186 | 13.52668 | 0 | 1.340422 | 3.447467 | 0 | 0 | 0 | 1.668654 | 4.716496 | 13.76779 |

| | | | | | | | | | | | | | | | | | | | | | | | | | | |
|---|---|---|---|---|---|---|---|---|---|---|---|---|---|---|---|---|---|---|---|---|---|---|---|---|---|---|
| Gm4524 | 100043571 | 4.327846 | -0.442 | 12.21152 | 4.25E-11 | 3.19E-10 | -1.6439 | -0.442 | -7.96249 | 7.92E-08 | 3.94E-07 | 11.87169 | 6.216245 | 13.88751 | 219.4818 | 237.3428 | 225.8955 | 82.43313 | 64.34024 | 79.29174 | 83.07516 | 69.39003 | 66.42437 | 10.01192 | 10.21907 | 17.00728 |
| Saa1 | 20208 | 4.329883 | -2.09609 | 7.836797 | 1.02E-07 | 4.83E-07 | -1.19517 | -2.09609 | -3.67829 | 0.001369 | 0.003643 | 2.967923 | 7.10428 | 3.471878 | 88.31222 | 77.24174 | 135.2668 | 34.11026 | 49.5956 | 48.26453 | 4.615287 | 1.525056 | 10.44878 | 5.840288 | 3.930413 | 6.478962 |
| Schip1 | 30953 | 4.33852 | 0.797153 | 12.15702 | 4.63E-11 | 3.45E-10 | 2.410631 | 0.797153 | 20.45535 | 1.66E-15 | 3.39E-14 | 11.12971 | 19.53677 | 17.35939 | 290.9109 | 356.7164 | 378.747 | 1945.706 | 1935.569 | 1765.103 | 16.92272 | 19.82572 | 7.463413 | 73.42077 | 84.89692 | 94.75482 |
| Adora2a | 11540 | 4.3405 | 2.854979 | 32.07441 | 1.51E-19 | 5.72E-18 | -2.12071 | 2.854979 | -25.0013 | 2.70E-17 | 8.48E-16 | 162.4938 | 130.5411 | 118.9118 | 2828.589 | 3206.234 | 2928.526 | 771.7446 | 690.3172 | 681.4493 | 285.3786 | 339.3249 | 274.6536 | 171.8713 | 161.1469 | 188.6998 |
| Ifit1 | 15957 | 4.340753 | 9.583864 | 106.7511 | 1.82E-24 | 2.61E-22 | 0.094146 | 9.583864 | 6.650352 | 5.28E-06 | 2.04E-05 | 11689.91 | 12281.52 | 10630.02 | 249610.6 | 254975 | 243127.1 | 276038.7 | 272346.9 | 279180.5 | 19218.82 | 19577.9 | 17616.64 | 13738.03 | 13681.77 | 13564.52 |
| Ereg | 13874 | 4.344393 | -2.52377 | 5.49129 | 1.79E-05 | 6.43E-05 | 2.680477 | -2.52377 | 6.963879 | 6.45E-07 | 2.82E-06 | 1.483961 | 1.77607 | 0 | 28.5716 | 18.25714 | 32.46403 | 156.3387 | 198.3824 | 173.5225 | 1.538429 | 0 | 5.97073 | 14.18356 | 14.14949 | 9.718443 |
| Plagl1 | 22634 | 4.352983 | 3.808213 | 29.94545 | 6.36E-19 | 2.08E-17 | 1.614984 | 3.808213 | 29.90154 | 6.56E-19 | 3.14E-17 | 115.007 | 119.8847 | 151.0267 | 2981.836 | 2651.498 | 2713.451 | 8989.475 | 8459.402 | 9066.838 | 204.611 | 171.5688 | 250.7707 | 624.0765 | 613.1444 | 660.0443 |
| Il12a | 16159 | 4.356957 | -0.70704 | 7.808291 | 1.08E-07 | 5.10E-07 | 2.72364 | -0.70704 | 14.34991 | 1.95E-12 | 2.07E-11 | 8.161788 | 3.55214 | 9.547666 | 158.4425 | 130.6088 | 174.4941 | 1092.95 | 1156.784 | 951.5008 | 1.538429 | 18.30067 | 5.224389 | 12.5149 | 10.21907 | 6.478962 |
| Il1rn | 16181 | 4.367306 | 8.262082 | 137.976 | 5.53E-33 | 4.84E-29 | -0.84154 | 8.262082 | -58.474 | 4.76E-25 | 1.63E-22 | 9034.358 | 7963.898 | 9202.214 | 192055.7 | 191458.4 | 193996.9 | 110629.5 | 110961.5 | 112654 | 6383.711 | 6388.458 | 6308.823 | 1680.334 | 1698.725 | 1876.469 |
| Wdr20rt | 70948 | 4.369779 | -2.80726 | 5.988015 | 5.68E-06 | 2.16E-05 | -0.0075 | -2.80726 | -0.01667 | 0.986857 | 0.989001 | 2.967923 | 0 | 0.86797 | 31.16902 | 32.30109 | 20.29002 | 19.89765 | 28.14886 | 37.92213 | 3.076858 | 1.525056 | 3.731706 | 15.01788 | 24.36856 | 8.908573 |
| Scimp | 327957 | 4.37348 | 2.257845 | 23.44681 | 1.02E-16 | 2.07E-15 | -0.57681 | 2.257845 | -6.96247 | 6.47E-07 | 2.82E-06 | 40.06696 | 61.27441 | 52.07818 | 1064.942 | 1119.303 | 1147.062 | 818.6463 | 721.1469 | 779.1275 | 243.841 | 250.1091 | 215.6926 | 179.3803 | 177.6547 | 198.4182 |
| Gpr141b | 319293 | 4.375308 | -1.31412 | 7.755899 | 1.21E-07 | 5.64E-07 | -0.30905 | -1.31412 | -1.10327 | 0.282195 | 0.373943 | 5.193865 | 5.32821 | 1.735939 | 100.0006 | 82.85932 | 90.62874 | 88.11817 | 101.8721 | 52.86116 | 9.230573 | 12.20045 | 11.94146 | 27.53279 | 31.4433 | 14.57766 |
| C78653 | 97640 | 4.376144 | -3.84971 | 5.177089 | 3.76E-05 | 0.00013 | 1.063961 | -3.84971 | 1.874973 | 0.074547 | 0.121628 | 0 | 0 | 0 | 7.792255 | 9.830767 | 8.116006 | 29.84648 | 13.40422 | 18.38649 | 3.076858 | 1.525056 | 0 | 9.177596 | 14.14949 | 11.33818 |
| Sprr2d | 20758 | 4.379472 | -4.05237 | 4.791904 | 9.42E-05 | 0.000309 | -2.27434 | -4.05237 | -2.87677 | 0.008932 | 0.019554 | 0 | 1.77607 | 0.86797 | 12.98709 | 49.15383 | 16.23201 | 9.948826 | 1.340422 | 4.596622 | 4.615287 | 2.287584 | 0 | 3.337308 | 3.930413 | 1.619741 |
| Gm3667 | 100042100 | 4.382902 | -4.4521 | 4.878835 | 7.65E-05 | 0.000254 | -0.92781 | -4.4521 | -1.26607 | 0.219132 | 0.302763 | 0 | 0 | 0 | 5.194837 | 11.23516 | 10.82134 | 1.421261 | 10.72337 | 4.596622 | 0 | 1.525056 | 2.239024 | 3.337308 | 9.432991 | 6.478962 |
| Gm10552 | 100503502 | 4.385419 | -2.16124 | 6.089791 | 4.50E-06 | 1.73E-05 | -0.43927 | -2.16124 | -1.13191 | 0.270228 | 0.360976 | 12.61367 | 4.440175 | 0 | 50.64966 | 78.64613 | 82.51273 | 46.90161 | 53.61687 | 56.30862 | 10.769 | 9.912862 | 6.717071 | 1.668654 | 7.074744 | 4.049351 |
| Pou3f1 | 18991 | 4.400039 | 2.355088 | 18.45904 | 1.33E-14 | 1.80E-13 | 2.334928 | 2.355088 | 30.34693 | 4.82E-19 | 2.39E-17 | 31.16319 | 44.40175 | 22.56721 | 779.2255 | 696.5801 | 674.9812 | 3837.404 | 3684.819 | 3746.247 | 164.6119 | 178.4315 | 168.6731 | 188.5579 | 159.5748 | 200.038 |
| Tnfaip3 | 21929 | 4.410253 | 7.504234 | 94.0207 | 1.96E-29 | 3.25E-26 | -0.0002 | 7.504234 | -0.00965 | 0.99239 | 0.993674 | 3066.606 | 3053.952 | 3394.629 | 71903.03 | 72101.65 | 71580.47 | 76219.38 | 75479.15 | 72564.58 | 6176.023 | 6351.857 | 6074.471 | 1412.515 | 1348.132 | 1523.366 |
| Pla1a | 85031 | 4.415271 | 1.425616 | 12.18263 | 4.45E-11 | 3.32E-10 | 1.459153 | 1.425616 | 11.78109 | 8.32E-11 | 6.60E-10 | 14.09763 | 19.53677 | 11.2836 | 346.7553 | 351.0988 | 323.2876 | 1013.359 | 1010.678 | 907.8329 | 103.844 | 86.16565 | 67.17071 | 274.4936 | 209.098 | 276.9756 |
| Gm10634 | 100039674 | 4.420472 | 1.293176 | 14.61944 | 1.35E-12 | 1.28E-11 | 0.430034 | 1.293176 | 3.839579 | 0.000931 | 0.002564 | 41.55092 | 29.30515 | 15.62345 | 674.0301 | 603.89 | 619.5218 | 932.3471 | 915.5081 | 822.7954 | 99.99788 | 110.5665 | 91.05363 | 40.88202 | 29.87114 | 67.21923 |
| Arhgef37 | 328967 | 4.427098 | 0.983168 | 16.37375 | 1.46E-13 | 1.64E-12 | -0.69605 | 0.983168 | -5.79616 | 8.83E-06 | 3.30E-05 | 31.16319 | 24.86498 | 30.37894 | 688.3159 | 662.8746 | 660.1019 | 393.6893 | 482.5518 | 417.1435 | 66.15244 | 79.3029 | 80.60486 | 23.36115 | 62.88661 | 31.58494 |
| Ptges | 64292 | 4.428717 | 3.313835 | 36.09755 | 1.27E-20 | 6.33E-19 | -1.09381 | 3.313835 | -16.6312 | 1.07E-13 | 1.47E-12 | 223.3362 | 297.4917 | 318.5448 | 6258.479 | 6246.75 | 6392.708 | 3332.857 | 3262.587 | 2693.621 | 147.6892 | 180.7191 | 164.9414 | 85.93567 | 87.25517 | 77.74755 |
| Mx1 | 17857 | 4.42891 | 8.124973 | 49.68263 | 5.03E-14 | 6.11E-13 | -1.21163 | 8.124973 | -26.4195 | 4.17E-11 | 3.49E-10 | 5925.458 | 6306.824 | 5400.507 | 134285.2 | 136121 | 134371.3 | 62717.4 | 63381.84 | 56416.64 | 10627.47 | 11858.83 | 9705.422 | 2460.43 | 2519.395 | 2567.289 |
| Ido2 | 209176 | 4.483172 | -0.19027 | 8.836972 | 1.41E-08 | 7.41E-08 | -0.59645 | -0.19027 | -2.4763 | 0.021719 | 0.042373 | 4.451884 | 7.992315 | 6.075787 | 133.767 | 209.2549 | 124.4454 | 125.071 | 91.14868 | 98.82738 | 76.15223 | 42.70156 | 50.75121 | 62.57452 | 83.32476 | 63.97975 |
| Fzd1 | 14362 | 4.487761 | 2.633306 | 27.33491 | 4.25E-18 | 1.17E-16 | -1.53384 | 2.633306 | -17.6937 | 3.11E-14 | 4.85E-13 | 77.90798 | 111.8924 | 100.6845 | 2335.079 | 2336.914 | 2161.563 | 764.6383 | 809.6147 | 865.3142 | 99.99788 | 136.4925 | 159.717 | 275.3279 | 342.732 | 293.9829 |
| Tex12 | 66654 | 4.487794 | 0.094996 | 17.28592 | 4.96E-14 | 6.02E-13 | -3.00571 | 0.094996 | -15.2299 | 6.09E-13 | 7.09E-12 | 34.13111 | 18.64873 | 30.37894 | 707.7965 | 668.4921 | 611.4058 | 66.79926 | 87.12741 | 101.1257 | 35.38386 | 42.70156 | 25.3756 | 16.68654 | 23.58248 | 20.24676 |
| Ifi214 | 545384 | 4.49737 | 1.07133 | 12.67078 | 2.12E-11 | 1.67E-10 | 0.554793 | 1.07133 | 4.34318 | 0.000277 | 0.000835 | 23.74338 | 15.09659 | 56.41802 | 602.6011 | 734.4987 | 673.6285 | 1019.044 | 1138.018 | 920.4736 | 93.84416 | 64.81487 | 54.48291 | 30.8701 | 31.4433 | 12.95792 |
| Sdc4 | 20971 | 4.504442 | 10.01965 | 60.61275 | 5.19E-15 | 7.54E-14 | -0.79872 | 10.01965 | -25.2152 | 6.20E-11 | 5.02E-10 | 8560.232 | 8650.349 | 9315.918 | 213655.8 | 221738.6 | 206764.7 | 132818.2 | 129077.3 | 122645.9 | 60894.09 | 60733.82 | 63021.8 | 60603 | 60431.67 | 64132.01 |
| Gm9895 | 100503337 | 4.51777 | -1.7176 | 6.747388 | 1.03E-06 | 4.27E-06 | 0.828068 | -1.7176 | 2.607872 | 0.0163 | 0.033051 | 2.967923 | 0.888035 | 1.735939 | 50.64966 | 60.389 | 45.9907 | 79.59061 | 93.82952 | 114.9156 | 23.84565 | 25.16342 | 38.80975 | 10.84625 | 1.572165 | 11.33818 |
| Fam3b | 52793 | 4.542369 | -0.57971 | 9.278464 | 6.10E-09 | 3.39E-08 | 0.639966 | -0.57971 | 3.298034 | 0.003373 | 0.008218 | 5.193865 | 5.32821 | 4.339848 | 137.6632 | 139.0351 | 121.7401 | 206.0828 | 209.1058 | 227.5328 | 18.46115 | 9.912862 | 11.19512 | 76.75808 | 35.37372 | 47.78235 |
| Vmn2r116 | 619697 | 4.544624 | -3.82695 | 5.828466 | 8.19E-06 | 3.06E-05 | 0.940868 | -3.82695 | 1.825959 | 0.081881 | 0.131578 | 0 | 0 | 0 | 11.68838 | 7.021976 | 10.82134 | 25.5827 | 16.08506 | 19.53565 | 0 | 1.525056 | 1.492683 | 15.01788 | 10.21907 | 14.57766 |
| Olfr56 | 18356 | 4.564765 | 0.392245 | 16.91806 | 7.61E-14 | 9.00E-13 | -1.6205 | 0.392245 | -11.5061 | 1.29E-10 | 9.91E-10 | 28.19527 | 22.20087 | 19.09533 | 609.0946 | 634.7867 | 541.0671 | 181.9214 | 215.8079 | 202.2514 | 67.69087 | 72.44015 | 51.49755 | 12.5149 | 14.14949 | 25.10598 |
| Krt16 | 16666 | 4.576761 | -2.54781 | 6.361696 | 2.43E-06 | 9.64E-06 | 2.421096 | -2.54781 | 6.657253 | 1.26E-06 | 5.30E-06 | 0 | 0.888035 | 0.86797 | 28.5716 | 19.66153 | 16.23201 | 130.756 | 105.8933 | 124.1088 | 2.307643 | 6.100223 | 1.492683 | 10.84625 | 16.50774 | 19.43689 |
| Ighv6-3 | 629845 | 4.580067 | -3.06458 | 5.91718 | 6.68E-06 | 2.52E-05 | 0.671714 | -3.06458 | 1.484806 | 0.152216 | 0.223065 | 0 | 0 | 3.471878 | 33.76644 | 18.25714 | 14.87935 | 48.32287 | 28.14886 | 33.32551 | 3.076858 | 2.287584 | 2.985365 | 7.508942 | 11.79124 | 6.478962 |
| Olr1 | 108078 | 4.584473 | 3.149345 | 25.31843 | 2.08E-17 | 4.84E-16 | 1.452998 | 3.149345 | 22.66253 | 2.04E-16 | 5.24E-15 | 103.1353 | 103.9001 | 150.1587 | 3057.161 | 2995.575 | 2933.936 | 8528.986 | 7917.871 | 9055.346 | 179.9962 | 108.279 | 170.1658 | 130.9893 | 81.75259 | 76.12781 |
| Apol9a | 223672 | 4.588922 | 1.381839 | 15.36718 | 5.10E-13 | 5.20E-12 | 1.952107 | 1.381839 | 20.89941 | 1.07E-15 | 2.30E-14 | 20.77546 | 30.19319 | 21.69924 | 619.4843 | 591.2504 | 645.2225 | 2748.718 | 2494.525 | 2274.179 | 37.69151 | 50.32684 | 44.03413 | 65.91183 | 52.66754 | 47.78235 |
| Angpt1 | 11600 | 4.690909 | 1.083385 | 16.98516 | 7.04E-14 | 8.38E-13 | -2.09437 | 1.083385 | -13.333 | 8.03E-12 | 7.66E-11 | 17.06556 | 26.64105 | 27.77503 | 658.4455 | 634.7867 | 635.7538 | 162.0237 | 162.191 | 145.9428 | 37.69151 | 41.1765 | 23.13658 | 456.3768 | 371.8171 | 396.8364 |
| Hdc | 15186 | 4.699353 | 4.062019 | 39.61976 | 1.79E-21 | 1.06E-19 | 1.268382 | 4.062019 | 28.8488 | 1.39E-18 | 6.03E-17 | 234.4659 | 237.1053 | 206.5768 | 6489.65 | 6109.119 | 6234.446 | 16250.7 | 15942.98 | 15047.04 | 236.9181 | 305.7737 | 221.6634 | 166.0311 | 178.4408 | 182.2208 |
| Gm7609 | 665378 | 4.703219 | -1.21019 | 5.728425 | 1.63E-05 | 5.86E-05 | -1.9543 | -1.21019 | -4.10432 | 0.000608 | 0.001732 | 24.48536 | 20.4248 | 0 | 158.4425 | 202.2329 | 169.0835 | 59.69296 | 67.02109 | 27.57973 | 31.53779 | 42.70156 | 26.12194 | 11.68058 | 6.288661 | 3.239481 |
| Shisa3 | 330096 | 4.721103 | 2.938459 | 39.0984 | 2.37E-21 | 1.37E-19 | -2.93813 | 2.938459 | -34.2751 | 3.77E-20 | 2.62E-18 | 281.9527 | 359.6542 | 219.5963 | 7983.165 | 8116 | 7664.215 | 1154.064 | 1036.146 | 1037.687 | 209.9955 | 173.8563 | 230.6194 | 42.55067 | 66.03094 | 24.29611 |
| Bend7 | 209645 | 4.725118 | -0.89507 | 9.495936 | 4.08E-09 | 2.32E-08 | -1.03243 | -0.89507 | -4.17946 | 0.000411 | 0.001205 | 2.225942 | 5.32821 | 10.41564 | 163.6374 | 146.0571 | 140.6774 | 78.16935 | 77.74446 | 72.3968 | 11.53822 | 18.30067 | 31.34633 | 29.20144 | 33.01547 | 22.67637 |
| Gm3636 | 100041151 | 4.730969 | -4.40956 | 6.509035 | 1.75E-06 | 7.05E-06 | -2.86227 | -4.40956 | -4.49686 | 0.000191 | 0.000594 | 4.451884 | 0 | 0 | 27.27289 | 25.27911 | 31.11136 | 5.685043 | 1.340422 | 4.596622 | 0 | 0 | 0 | 3.337308 | 3.14433 | 4.859222 |
| Mx2 | 17858 | 4.757379 | 6.344064 | 55.17298 | 2.80E-20 | 1.27E-18 | -1.42453 | 6.344064 | -29.9029 | 6.73E-16 | 1.51E-14 | 1477.284 | 1654.409 | 1378.336 | 43304.16 | 43252.57 | 43193.39 | 17620.79 | 17559.52 | 15341.23 | 2448.41 | 2855.667 | 2321.121 | 896.9014 | 979.4589 | 1064.17 |
| Gm10344 | 100038691 | 4.759102 | -3.09558 | 4.957281 | 6.34E-05 | 0.000212 | 3.27791 | -3.09558 | 6.315171 | 2.70E-06 | 1.09E-05 | 0 | 0 | 0 | 15.58451 | 8.426372 | 10.82134 | 102.3308 | 140.7443 | 119.5122 | 6.153716 | 0 | 1.492683 | 12.5149 | 30.65722 | 11.33818 |
| Olfr1396 | 258334 | 4.777611 | -0.55572 | 13.96487 | 3.30E-12 | 2.94E-11 | -1.51682 | -0.55572 | -9.0516 | 9.34E-09 | 5.31E-08 | 11.87169 | 14.20856 | 14.75548 | 446.756 | 424.1274 | 351.6936 | 147.8111 | 170.2336 | 128.7054 | 19.23036 | 27.451 | 20.15121 | 2.502981 | 7.860826 | 8.098703 |
| Dusp2 | 13537 | 4.813967 | 4.142879 | 62.71287 | 1.08E-25 | 2.38E-23 | -2.47202 | 4.142879 | -49.4877 | 1.63E-23 | 3.74E-21 | 408.0894 | 493.7474 | 476.5153 | 13339.04 | 13888.06 | 13839.14 | 2534.108 | 2717.035 | 2445.403 | 419.2219 | 481.9176 | 446.3121 | 197.7355 | 166.6495 | 181.4109 |
| Tarm1 | 245126 | 4.826096 | 4.583631 | 40.35916 | 1.21E-21 | 7.37E-20 | -2.06612 | 4.583631 | -30.5033 | 4.33E-19 | 2.18E-17 | 270.081 | 205.1361 | 237.8237 | 7423.422 | 7050.064 | 7189.429 | 1859.009 | 1959.697 | 1599.625 | 2471.486 | 2410.351 | 2291.268 | 877.7119 | 753.0671 | 841.4552 |
| Gm14023 | 100503468 | 4.84657 | 0.345114 | 12.26895 | 3.90E-11 | 2.94E-10 | 2.008574 | 0.345114 | 17.4008 | 4.34E-14 | 6.56E-13 | 12.61367 | 11.54445 | 13.01954 | 405.1973 | 356.7164 | 417.9743 | 1712.619 | 1550.868 | 1667.425 | 6.153716 | 16.77561 | 30.59999 | 21.6925 | 24.36856 | 14.57766 |
| Ifi208 | 100033459 | 4.850266 | 1.430308 | 16.69701 | 9.89E-14 | 1.15E-12 | -0.39558 | 1.430308 | -3.67267 | 0.001388 | 0.003686 | 42.2929 | 46.17782 | 12.15157 | 976.6293 | 903.0262 | 860.2967 | 706.3666 | 735.8915 | 718.2222 | 133.0741 | 118.9543 | 124.639 | 46.72231 | 50.30929 | 37.25403 |
| Ifit2 | 15958 | 4.859801 | 9.519416 | 56.15993 | 4.38E-14 | 5.38E-13 | -0.92739 | 9.519416 | -29.9901 | 2.59E-11 | 2.25E-10 | 11800.46 | 13343.61 | 11083.97 | 373373.7 | 384318.4 | 360983.7 | 204302 | 197771.2 | 207408.8 | 45759.8 | 47303.42 | 40713.66 | 3963.887 | 4317.952 | 3749.699 |
| P3h2 | 210530 | 4.939236 | -0.59804 | 9.025829 | 9.81E-09 | 5.30E-08 | -1.6432 | -0.59804 | -5.79581 | 8.84E-06 | 3.30E-05 | 20.03348 | 1.77607 | 3.471878 | 192.209 | 168.5274 | 216.4268 | 55.42917 | 52.27645 | 82.7392 | 83.84438 | 83.87806 | 61.94632 | 19.18952 | 20.43815 | 15.38754 |
| Plekha4 | 69217 | 4.971421 | 3.944828 | 20.70969 | 2.72E-15 | 4.16E-14 | 2.304895 | 3.944828 | 32.12481 | 4.12E-19 | 2.10E-17 | 92.00561 | 105.6762 | 96.34463 | 3440.281 | 3182.36 | 3212.586 | 17886.57 | 17779.35 | 15219.42 | 139.997 | 175.3814 | 167.9268 | 816.8061 | 685.464 | 766.9471 |
| Gm4841 | 225594 | 4.978209 | 1.595625 | 16.81481 | 8.60E-14 | 1.01E-12 | 1.725616 | 1.595625 | 20.43688 | 1.69E-15 | 3.44E-14 | 28.19527 | 28.41712 | 26.90706 | 1070.136 | 896.0042 | 890.0554 | 3467.876 | 3015.949 | 3322.209 | 109.9977 | 147.1679 | 85.0829 | 20.85817 | 21.22423 | 16.19741 |
| Gbp6 | 100702 | 4.983428 | 5.343222 | 52.30133 | 5.06E-24 | 6.25E-22 | 0.227511 | 5.343222 | 5.903817 | 6.89E-06 | 2.62E-05 | 531.2582 | 539.9253 | 378.4347 | 16344.25 | 16879.43 | 15396.06 | 20291.34 | 19621.09 | 19224.22 | 2441.487 | 2530.067 | 2353.214 | 254.4697 | 211.4562 | 213.8058 |
| Adamts4 | 240913 | 4.996019 | 0.708713 | 15.34593 | 5.25E-13 | 5.33E-12 | -0.22361 | 0.708713 | -1.96773 | 0.062229 | 0.104421 | 19.2915 | 13.32052 | 24.30315 | 657.1468 | 681.1317 | 597.8791 | 541.5004 | 620.6153 | 559.6388 | 13.07665 | 19.0632 | 14.18048 | 55.06558 | 91.18558 | 106.9029 |
| Enpp4 | 224794 | 4.996822 | 5.050103 | 57.36344 | 7.15E-25 | 1.16E-22 | 0.313877 | 5.050103 | 9.189533 | 7.21E-09 | 4.18E-08 | 324.9876 | 356.9901 | 300.3175 | 11037.73 | 11454.25 | 10936.32 | 14555.13 | 14456.45 | 14135.76 | 1193.821 | 1263.509 | 1106.824 | 529.7976 | 481.0826 | 460.0063 |
| Spdef | 30051 | 5.009982 | -2.60711 | 6.166611 | 3.78E-06 | 1.47E-05 | 0.3095 | -2.60711 | 0.668578 | 0.510932 | 0.60801 | 1.483961 | 0 | 0 | 27.27289 | 37.91867 | 13.52668 | 39.7953 | 33.51054 | 24.13227 | 9.230573 | 7.625279 | 2.985365 | 16.68654 | 33.01547 | 26.72572 |
| Gbp10 | 626578 | 5.011417 | 0.38954 | 13.8471 | 3.88E-12 | 3.42E-11 | 0.397805 | 0.38954 | 3.413664 | 0.002569 | 0.006443 | 11.12971 | 7.992315 | 25.17112 | 453.2495 | 477.4944 | 462.6124 | 703.5241 | 592.4664 | 619.3949 | 25.38408 | 32.7887 | 46.27316 | 15.85221 | 22.01031 | 26.72572 |
| Oasl1 | 231655 | 5.018789 | 7.911774 | 39.60641 | 6.53E-13 | 6.54E-12 | 0.978879 | 7.911774 | 23.8032 | 1.40E-10 | 1.07E-09 | 2360.241 | 2347.964 | 2283.628 | 81774.52 | 81616.43 | 78768.55 | 172643.4 | 168271.2 | 157290.7 | 4627.594 | 4726.148 | 4225.038 | 4360.192 | 4314.021 | 4461.575 |
| Sectm1a | 209588 | 5.033707 | 0.884333 | 13.53038 | 6.06E-12 | 5.20E-11 | 2.15479 | 0.884333 | 20.69734 | 1.31E-15 | 2.73E-14 | 8.903769 | 7.992315 | 9.547666 | 320.7812 | 300.5406 | 344.9303 | 1554.859 | 1506.634 | 1421.505 | 39.99915 | 39.65145 | 29.10731 | 88.43865 | 110.0516 | 106.9029 |
| Cd40 | 21939 | 5.034114 | 5.569989 | 87.55401 | 8.93E-29 | 9.03E-26 | -0.14597 | 5.569989 | -6.06816 | 4.73E-06 | 1.84E-05 | 1015.03 | 1082.515 | 951.2947 | 35537.88 | 35844.38 | 35138.25 | 34297.87 | 33584.27 | 32274.03 | 2136.878 | 2098.477 | 2070.351 | 76.75808 | 67.60311 | 73.69819 |
| Pdgfrb | 18596 | 5.036744 | 1.384398 | 15.68118 | 3.43E-13 | 3.59E-12 | 0.610717 | 1.384398 | 6.175502 | 3.71E-06 | 1.46E-05 | 30.42121 | 20.4248 | 36.45472 | 1054.552 | 1047.679 | 937.3987 | 1628.765 | 1777.399 | 1447.936 | 45.38365 | 20.58825 | 49.25852 | 75.08942 | 47.95104 | 43.73299 |
| Nt5c1a | 230718 | 5.043549 | -2.5454 | 5.56209 | 1.52E-05 | 5.49E-05 | 2.612587 | -2.5454 | 6.696458 | 1.16E-06 | 4.88E-06 | 0 | 6.216245 | 0 | 27.27289 | 36.51428 | 41.9327 | 260.0907 | 193.0207 | 222.9362 | 0 | 0 | 5.97073 | 11.68058 | 21.22423 | 10.52831 |
| Gbp3 | 55932 | 5.045996 | 7.027903 | 75.52372 | 2.07E-27 | 7.79E-25 | 0.624592 | 7.027903 | 25.51075 | 1.78E-17 | 5.83E-16 | 1233.914 | 1372.902 | 1237.725 | 45349.63 | 46467.23 | 43481.5 | 74506.76 | 70569.18 | 71610.78 | 2965.322 | 3117.976 | 2687.575 | 2359.477 | 2490.31 | 2276.545 |
| Ifitm7 | 74482 | 5.046677 | -0.4335 | 15.53532 | 4.12E-13 | 4.27E-12 | -3.19657 | -0.4335 | -14.4955 | 1.60E-12 | 1.73E-11 | 24.48536 | 15.98463 | 21.69924 | 718.1862 | 828.5932 | 688.5079 | 100.9095 | 76.40404 | 76.99342 | 4.615287 | 19.82572 | 8.956095 | 11.68058 | 24.36856 | 7.288832 |
| Olfr231 | 404222 | 5.048425 | -3.51514 | 5.782571 | 9.11E-06 | 3.38E-05 | 3.907252 | -3.51514 | 8.944255 | 1.15E-08 | 6.41E-08 | 0 | 0 | 0 | 7.792255 | 11.23516 | 31.11136 | 234.508 | 261.3822 | 197.6548 | 0 | 0 | 0 | 5.005962 | 15.72165 | 4.049351 |
| Calhm6 | 215900 | 5.055123 | 2.951582 | 30.07125 | 5.83E-19 | 1.93E-17 | -0.11643 | 2.951582 | -1.89244 | 0.072076 | 0.11814 | 100.9094 | 111.0044 | 88.5329 | 3624.697 | 3546.098 | 3487.177 | 3534.676 | 3450.246 | 3249.812 | 294.6091 | 398.8021 | 440.3413 | 48.39096 | 48.73712 | 42.11325 |
| Slfn4 | 20558 | 5.082066 | 6.757758 | 92.86267 | 2.55E-29 | 3.35E-26 | -1.85942 | 6.757758 | -64.5999 | 5.74E-26 | 3.43E-23 | 3090.35 | 3171.173 | 2850.412 | 108535.7 | 108165.1 | 112796.3 | 31399.92 | 30753.3 | 31977.55 | 2742.25 | 2834.316 | 2315.897 | 395.471 | 378.1057 | 470.5346 |
| Gm19654 | 100503358 | 5.082555 | -4.18253 | 4.368484 | 0.000261 | 0.000812 | -1.01857 | -4.18253 | -1.1542 | 0.261178 | 0.350811 | 0 | 0 | 0 | 7.792255 | 14.04395 | 27.05335 | 0 | 20.10633 | 25.28142 | 0 | 1.525056 | 0 | 13.34923 | 12.57732 | 12.14805 |
| Tpbg | 21983 | 5.085511 | 1.856372 | 22.63055 | 2.10E-16 | 4.07E-15 | 1.787596 | 1.856372 | 28.45563 | 1.84E-18 | 7.71E-17 | 23.0014 | 38.1855 | 39.9266 | 1200.007 | 1106.663 | 1234.986 | 4255.255 | 4188.818 | 4242.682 | 44.61444 | 43.46409 | 32.83902 | 69.24913 | 76.25001 | 67.21923 |
| Htra4 | 330723 | 5.086767 | 3.872316 | 26.98941 | 5.54E-18 | 1.49E-16 | 1.700256 | 3.872316 | 30.25835 | 5.12E-19 | 2.52E-17 | 76.42402 | 66.60262 | 105.0243 | 2955.862 | 3033.494 | 2881.182 | 10231.66 | 10293.1 | 9483.981 | 650.7554 | 706.8633 | 609.7608 | 317.8786 | 372.6032 | 289.9336 |
| Cfb | 14962 | 5.164688 | 5.542928 | 45.69865 | 8.56E-20 | 3.40E-18 | 0.459091 | 5.542928 | 10.81422 | 3.23E-09 | 1.98E-08 | 657.3949 | 675.7946 | 627.542 | 25287.17 | 25269.28 | 24553.62 | 37684.73 | 37198.04 | 33029.03 | 716.1387 | 804.4669 | 841.8729 | 359.5949 | 342.732 | 389.5476 |
| Socs3 | 12702 | 5.173215 | 6.046308 | 92.62302 | 2.69E-29 | 3.36E-26 | -3.10144 | 6.046308 | -79.5176 | 6.93E-28 | 1.82E-24 | 2177.713 | 2090.434 | 1749.827 | 77800.47 | 77050.74 | 76694.91 | 9701.527 | 9770.334 | 8677.274 | 2421.487 | 2464.49 | 2599.507 | 270.3219 | 271.1985 | 258.3486 |
| Draxin | 70433 | 5.219378 | -0.44043 | 13.42744 | 7.02E-12 | 5.95E-11 | -3.1303 | -0.44043 | -12.1694 | 4.54E-11 | 3.77E-10 | 7.419807 | 23.97694 | 6.943757 | 433.7689 | 471.8768 | 404.4477 | 41.21656 | 50.93603 | 60.90525 | 12.30743 | 6.100223 | 16.41951 | 64.24317 | 52.66754 | 61.55014 |
| Gm20115 | 100504215 | 5.223922 | 1.510183 | 17.94319 | 2.35E-14 | 3.02E-13 | 0.102846 | 1.510183 | 1.124649 | 0.273228 | 0.364354 | 69.00421 | 33.74533 | 29.51097 | 1840.271 | 1542.026 | 1712.477 | 2012.505 | 2028.058 | 1685.811 | 41.53758 | 64.81487 | 37.31706 | 15.85221 | 27.51289 | 25.91585 |
| Il15ra | 16169 | 5.258359 | 4.062709 | 37.51201 | 5.66E-21 | 3.05E-19 | -0.43056 | 4.062709 | -7.92419 | 8.56E-08 | 4.24E-07 | 192.915 | 172.2788 | 208.3127 | 8026.023 | 7821.077 | 7603.345 | 6507.953 | 6316.067 | 5398.733 | 502.297 | 441.5036 | 437.356 | 236.1145 | 249.9743 | 230.813 |
| Vcan | 13003 | 5.266488 | 5.534063 | 81.46285 | 4.14E-28 | 2.48E-25 | -2.60727 | 5.534063 | -65.1604 | 4.78E-26 | 2.99E-23 | 1383.052 | 1355.141 | 1372.26 | 57151 | 56428.6 | 55460.73 | 9782.538 | 10267.63 | 8900.21 | 765.3684 | 828.8678 | 699.3218 | 358.7606 | 270.4124 | 242.9611 |
| Iigp1 | 60440 | 5.321236 | 6.989527 | 85.41481 | 1.51E-28 | 1.24E-25 | 0.135456 | 6.989527 | 6.17905 | 3.68E-06 | 1.45E-05 | 1612.324 | 1697.923 | 1449.509 | 68112.1 | 69396.79 | 65423.13 | 77538.31 | 76597.06 | 77158.9 | 5949.874 | 6184.101 | 5314.696 | 579.8572 | 491.3016 | 514.2676 |
| Tnfsf10 | 22035 | 5.321782 | 4.994094 | 80.00138 | 6.09E-28 | 3.14E-25 | -1.98482 | 4.994094 | -55.7232 | 1.32E-24 | 3.82E-22 | 524.5804 | 632.2809 | 560.7084 | 23954.69 | 25199.06 | 23739.32 | 6313.241 | 6505.067 | 6287.03 | 1244.589 | 1218.52 | 1159.068 | 238.6175 | 245.2578 | 221.0946 |
| Cmpk2 | 22169 | 5.328875 | 9.990639 | 37.42014 | 3.31E-12 | 2.94E-11 | 0.536362 | 9.990639 | 18.92987 | 3.02E-09 | 1.86E-08 | 10102.07 | 11101.33 | 9935.648 | 444859.8 | 456723.4 | 431732.3 | 680681.6 | 679910.2 | 654461.3 | 28281.71 | 29886.52 | 24250.12 | 9631.47 | 9946.303 | 9894.995 |
| Ms4a20 | 69369 | 5.354426 | -2.10381 | 6.456301 | 1.97E-06 | 7.88E-06 | 3.380307 | -2.10381 | 10.50459 | 6.81E-10 | 4.66E-09 | 0 | 0 | 5.207818 | 33.76644 | 56.17581 | 37.8747 | 447.6972 | 491.9348 | 428.635 | 6.92293 | 1.525056 | 0.746341 | 17.52087 | 7.074744 | 9.718443 |
| Traf1 | 22029 | 5.387536 | 5.095723 | 61.16356 | 1.83E-25 | 3.79E-23 | -0.86963 | 5.095723 | -23.3429 | 1.11E-16 | 3.05E-15 | 435.5427 | 358.7661 | 363.6793 | 17068.93 | 17434.16 | 17462.94 | 10096.64 | 10189.89 | 9325.398 | 1291.511 | 1383.226 | 1417.302 | 380.4531 | 428.415 | 385.4982 |
| Hap1 | 15114 | 5.393969 | 2.445797 | 17.83282 | 2.66E-14 | 3.38E-13 | 0.722664 | 2.445797 | 8.429111 | 3.11E-08 | 1.64E-07 | 43.03488 | 33.74533 | 45.13442 | 1757.153 | 1768.134 | 1996.538 | 3281.691 | 3321.565 | 2892.425 | 96.15181 | 123.5295 | 51.49755 | 206.0787 | 154.0722 | 187.08 |
| Gm15998 | 100038411 | 5.403154 | -2.4575 | 7.09683 | 4.84E-07 | 2.09E-06 | -4.04052 | -2.4575 | -5.92391 | 6.58E-06 | 2.51E-05 | 1.483961 | 5.32821 | 1.735939 | 140.2606 | 112.3516 | 128.5034 | 1.421261 | 10.72337 | 18.38649 | 6.153716 | 8.387806 | 0 | 15.85221 | 20.43815 | 21.8665 |
| Gm5544 | 433632 | 5.414554 | -1.78021 | 7.382461 | 2.63E-07 | 1.18E-06 | -0.17503 | -1.78021 | -0.51214 | 0.613809 | 0.705431 | 0.741981 | 6.216245 | 0 | 66.23417 | 74.43295 | 58.16471 | 88.11817 | 49.5956 | 52.86116 | 16.92272 | 17.53814 | 8.956095 | 20.85817 | 11.79124 | 23.48624 |
| Dll1 | 13388 | 5.41733 | 1.721722 | 19.05223 | 7.01E-15 | 9.93E-14 | 1.29064 | 1.721722 | 17.34462 | 4.63E-14 | 6.95E-13 | 16.32358 | 30.19319 | 20.83127 | 1018.188 | 1005.547 | 960.3941 | 2610.856 | 2544.12 | 2440.806 | 50.76815 | 41.1765 | 29.85365 | 146.8415 | 143.8531 | 110.9522 |
| Tmem132 | 270893 | 5.425935 | 0.186742 | 9.563444 | 3.61E-09 | 2.07E-08 | 2.858888 | 0.186742 | 20.39513 | 1.76E-15 | 3.57E-14 | 9.64575 | 15.98463 | 1.735939 | 296.1057 | 381.9955 | 312.4662 | 2333.71 | 2633.929 | 2445.403 | 9.230573 | 6.100223 | 2.239024 | 53.39692 | 29.87114 | 60.74027 |
| Gbp2 | 14469 | 5.50386 | 7.98086 | 80.09696 | 3.55E-23 | 3.33E-21 | 0.179091 | 7.98086 | 8.63582 | 1.37E-07 | 6.58E-07 | 2601.384 | 2775.997 | 2354.802 | 125702.1 | 126279 | 121861.8 | 146866 | 149290.8 | 143857 | 7174.463 | 7743.47 | 6676.769 | 2232.659 | 2353.531 | 2390.737 |
| Styk1 | 243659 | 5.506972 | -1.14073 | 11.6871 | 9.66E-11 | 6.93E-10 | -1.39797 | -1.14073 | -7.21586 | 3.75E-07 | 1.70E-06 | 7.419807 | 5.32821 | 5.207818 | 272.7289 | 328.6285 | 331.4036 | 112.2796 | 119.2975 | 131.0037 | 9.999788 | 1.525056 | 0.746341 | 25.02981 | 18.0799 | 21.8665 |
| H2al1o | 333452 | 5.639412 | -4.51341 | 5.559594 | 1.53E-05 | 5.52E-05 | -1.14826 | -4.51341 | -1.63511 | 0.11669 | 0.177896 | 0 | 0 | 0 | 40.25998 | 8.426372 | 27.05335 | 11.37009 | 14.74464 | 5.745778 | 3.076858 | 0 | 0 | 5.005962 | 1.572165 | 0 |
| Socs1 | 12703 | 5.645323 | 6.27472 | 59.81056 | 2.95E-25 | 5.61E-23 | -0.59269 | 6.27472 | -16.3454 | 1.51E-13 | 2.01E-12 | 460.77 | 507.068 | 538.1412 | 27113.15 | 27245.27 | 25720.98 | 19691.57 | 18413.37 | 17290.2 | 7619.069 | 7394.995 | 7561.93 | 1150.537 | 1066.714 | 1217.235 |
| Tgtp1 | 21822 | 5.675446 | 3.249649 | 27.48362 | 3.80E-18 | 1.06E-16 | 3.294375 | 3.249649 | 70.68818 | 8.47E-27 | 8.46E-24 | 38.583 | 65.71459 | 37.32269 | 2564.951 | 2467.522 | 2411.807 | 26043.18 | 24590.04 | 25244.65 | 119.9975 | 127.3422 | 113.4439 | 181.0489 | 172.1521 | 158.7346 |
| BC023105 | 667597 | 5.683796 | 0.870549 | 12.3826 | 3.27E-11 | 2.50E-10 | 1.513294 | 0.870549 | 13.97909 | 3.23E-12 | 3.32E-11 | 15.5816 | 8.88035 | 17.35939 | 725.9784 | 848.2547 | 731.7932 | 2529.844 | 2336.355 | 2016.768 | 71.53694 | 48.03925 | 66.42437 | 3.337308 | 14.93557 | 19.43689 |
| Mir155 | 387173 | 5.71874 | -2.31111 | 9.55458 | 3.67E-09 | 2.11E-08 | -0.59928 | -2.31111 | -2.62919 | 0.015552 | 0.031688 | 1.483961 | 7.10428 | 1.735939 | 153.2477 | 224.7032 | 150.1461 | 135.0198 | 127.3401 | 98.82738 | 0.769214 | 0.762528 | 3.731706 | 0.834327 | 3.930413 | 1.619741 |
| Rsad2 | 58185 | 5.725121 | 10.85311 | 48.30104 | 2.52E-13 | 2.71E-12 | 0.287001 | 10.85311 | 19.66739 | 2.07E-09 | 1.31E-08 | 17103.4 | 18304.18 | 16941.9 | 984642.3 | 1001800 | 965633 | 1264354 | 1226785 | 1246532 | 58275.69 | 63035.89 | 50503.42 | 13476.88 | 13169.24 | 14026.95 |
| Ifit1bl1 | 667373 | 5.730314 | 4.615736 | 78.6613 | 8.72E-28 | 4.02E-25 | -4.63266 | 4.615736 | -73.9105 | 3.28E-27 | 3.92E-24 | 914.8623 | 849.8495 | 769.0211 | 48454.84 | 48371.59 | 47001.15 | 1995.45 | 1983.824 | 2031.707 | 1682.272 | 1785.84 | 1372.522 | 41.71635 | 55.81187 | 59.12053 |
| A630012P( | 100504594 | 5.739035 | -1.30996 | 11.12425 | 2.40E-10 | 1.63E-09 | -3.68248 | -1.30996 | -10.1284 | 1.31E-09 | 8.56E-09 | 2.967923 | 5.32821 | 9.547666 | 342.8592 | 297.7318 | 328.6983 | 9.948826 | 37.53181 | 39.07129 | 20.76879 | 17.53814 | 11.94146 | 10.01192 | 13.3634 | 7.288832 |
| Timp1 | 21857 | 5.750176 | 0.861865 | 19.89549 | 2.92E-15 | 4.43E-14 | -2.6847 | 0.861865 | -19.2927 | 5.44E-15 | 9.83E-14 | 17.80754 | 23.08891 | 32.11488 | 1336.372 | 1551.857 | 1228.222 | 230.2443 | 207.7654 | 221.787 | 66.15244 | 78.54037 | 84.33656 | 33.37308 | 31.4433 | 18.62702 |
| Mmp13 | 17386 | 5.761507 | 9.752719 | 41.77965 | 8.67E-13 | 8.49E-12 | 1.179259 | 9.752719 | 39.31851 | 1.61E-12 | 1.74E-11 | 3315.912 | 3170.285 | 3570.827 | 195090.8 | 193956.8 | 192799.8 | 456192 | 442081.8 | 466964 | 6604.475 | 6325.169 | 6040.886 | 192852.2 | 188257.4 | 205089.1 |
| Gm7582 | 665317 | 5.773318 | -1.17136 | 9.127753 | 8.09E-09 | 4.42E-08 | -1.31027 | -1.17136 | -5.04974 | 5.09E-05 | 0.000171 | 4.451884 | 1.77607 | 4.339848 | 190.9102 | 242.9604 | 224.5428 | 90.96069 | 109.9146 | 76.99342 | 13.07665 | 26.68847 | 11.19512 | 13.34923 | 25.15464 | 3.239481 |
| Tnfaip8l3 | 244882 | 5.911347 | 0.047428 | 14.79696 | 1.07E-12 | 1.03E-11 | -3.34766 | 0.047428 | -14.8154 | 1.04E-12 | 1.17E-11 | 14.83961 | 7.10428 | 6.943757 | 581.8217 | 707.8152 | 581.6471 | 52.58665 | 76.40404 | 60.90525 | 77.69066 | 83.11554 | 59.7073 | 29.20144 | 40.8763 | 29.15533 |
| Ccrl2 | 54199 | 5.921887 | 5.210638 | 57.39643 | 7.06E-25 | 1.15E-22 | 0.629948 | 5.210638 | 20.97788 | 9.93E-16 | 2.14E-14 | 267.855 | 201.5839 | 257.787 | 15602.69 | 16278.35 | 15267.56 | 25903.9 | 25296.44 | 24617.21 | 1296.896 | 1218.52 | 1234.448 | 444.6962 | 433.1315 | 454.3372 |
| Itgb8 | 320910 | 5.978444 | 1.121788 | 13.7336 | 4.55E-12 | 3.96E-11 | 1.27416 | 1.121788 | 13.1654 | 1.02E-11 | 9.60E-11 | 16.32358 | 5.32821 | 8.679696 | 587.0165 | 744.3295 | 665.5125 | 1533.54 | 1750.591 | 1692.706 | 51.53737 | 41.1765 | 29.85365 | 99.2849 | 89.61342 | 75.31793 |
| Cd69 | 12515 | 6.065676 | 6.37325 | 92.44264 | 2.81E-29 | 3.36E-26 | 0.029414 | 6.37325 | 1.570927 | 0.130907 | 0.196137 | 1282.143 | 1388.887 | 1335.805 | 95881.1 | 98373.68 | 91870.49 | 102232.7 | 97337.41 | 103106.8 | 1745.348 | 1768.302 | 1535.224 | 126.8177 | 122.6289 | 169.2629 |
| Ifnb1 | 15977 | 6.066922 | -0.32046 | 8.627643 | 2.11E-08 | 1.08E-07 | 2.809782 | -0.32046 | 17.49687 | 3.89E-14 | 5.94E-13 | 7.419807 | 15.09659 | 0 | 267.5341 | 328.6285 | 225.8955 | 2035.246 | 1946.292 | 1942.073 | 21.538 | 16.77561 | 31.34633 | 5.840288 | 3.930413 | 5.669092 |
| Acod1 | 16365 | 6.126652 | 9.027207 | 64.28183 | 1.11E-14 | 1.52E-13 | -0.7712 | 9.027207 | -41.8973 | 8.74E-13 | 9.88E-12 | 9740.723 | 9644.948 | 8475.723 | 692719.8 | 710440 | 676274.4 | 424280.5 | 427443.1 | 415275 | 39313.78 | 38987.29 | 36474.44 | 254.4697 | 260.9794 | 345.0047 |
| Gm3696 | 100042149 | 6.139464 | -4.39151 | 6.526139 | 1.69E-06 | 6.80E-06 | -4.13123 | -4.39151 | -4.95313 | 6.41E-05 | 0.000213 | 0 | 0 | 0 | 24.67547 | 22.47032 | 51.40137 | 2.842522 | 2.680843 | 0 | 0 | 5.337695 | 1.492683 | 1.668654 | 6.288661 | 7.288832 |
| AA467197 | 433470 | 6.19006 | 3.328246 | 44.56811 | 1.50E-22 | 1.14E-20 | -1.27352 | 3.328246 | -27.799 | 3.00E-18 | 1.18E-16 | 102.3933 | 77.25904 | 128.4595 | 7888.359 | 7941.855 | 7856.294 | 3408.184 | 3391.267 | 3369.324 | 313.0703 | 270.6974 | 305.2536 | 85.93567 | 113.1959 | 93.94495 |
| Fst | 14313 | 6.252578 | 0.315498 | 15.70399 | 3.33E-13 | 3.50E-12 | -0.94096 | 0.315498 | -8.43184 | 3.09E-08 | 1.63E-07 | 9.64575 | 7.992315 | 14.75548 | 887.0184 | 907.2393 | 854.886 | 478.9649 | 415.5307 | 534.3574 | 4.615287 | 10.67539 | 2.239024 | 121.8117 | 80.18043 | 87.46599 |
| Il1b | 16176 | 6.284428 | 5.306162 | 77.57245 | 1.17E-27 | 4.82E-25 | -1.08569 | 5.306162 | -38.286 | 3.68E-21 | 3.46E-19 | 546.8398 | 594.0954 | 505.1583 | 45214.56 | 45944.79 | 45574.08 | 23354.16 | 22519.09 | 21237.54 | 1513.045 | 1450.328 | 1424.765 | 97.61625 | 73.10568 | 102.8535 |
| Lhx2 | 16870 | 6.288018 | 2.805764 | 19.75357 | 3.37E-15 | 5.06E-14 | -2.26559 | 2.805764 | -19.8026 | 3.21E-15 | 6.14E-14 | 45.26083 | 17.7607 | 28.643 | 2423.391 | 2526.507 | 2472.677 | 589.8233 | 526.7857 | 496.4352 | 206.1495 | 180.7191 | 134.3414 | 1945.65 | 1760.039 | 2121.05 |
| AW11201( | 107350 | 6.303618 | 5.789762 | 70.9008 | 7.94E-27 | 2.64E-24 | 1.512006 | 5.789762 | 70.81015 | 8.16E-27 | 8.46E-24 | 301.2442 | 301.0439 | 262.9948 | 24602.75 | 24652.75 | 23823.18 | 72767.13 | 70381.52 | 73051.82 | 971.5179 | 843.3558 | 926.9558 | 842.6702 | 916.5723 | 811.49 |
| Il33 | 77125 | 6.470987 | -0.42961 | 12.29861 | 3.72E-11 | 2.82E-10 | 1.515547 | -0.42961 | 14.62478 | 1.35E-12 | 1.47E-11 | 11.12971 | 0.888035 | 14.75548 | 592.2114 | 636.1911 | 570.8258 | 1841.954 | 1687.591 | 1806.473 | 0.769214 | 3.050111 | 1.492683 | 12.5149 | 11.00516 | 16.19741 |
| Gbp4 | 17472 | 6.481838 | 5.632987 | 41.33545 | 7.33E-22 | 4.76E-20 | 2.213726 | 5.632987 | 66.16857 | 3.45E-26 | 2.39E-23 | 275.2749 | 205.1361 | 183.1416 | 21200.13 | 21283.61 | 20781.03 | 103037.1 | 104077 | 98348.18 | 689.2162 | 740.4145 | 650.0632 | 734.2077 | 697.2553 | 690.8193 |
| Tnn | 329278 | 6.566554 | -0.28339 | 13.75541 | 4.41E-12 | 3.85E-11 | -4.5935 | -0.28339 | -14.0806 | 2.81E-12 | 2.91E-11 | 2.967923 | 25.75301 | 24.30315 | 1253.254 | 1228.846 | 1199.816 | 27.00396 | 61.6594 | 76.99342 | 9.230573 | 12.96297 | 10.44878 | 38.37904 | 34.58764 | 22.67637 |
| Ifi205 | 226695 | 6.573269 | 4.737448 | 64.18659 | 6.58E-26 | 1.56E-23 | -0.72975 | 4.737448 | -23.8444 | 7.18E-17 | 2.06E-15 | 270.081 | 284.1712 | 322.0167 | 29293.68 | 30260.5 | 29144.58 | 18990.89 | 17886.59 | 18623.22 | 719.2155 | 821.2425 | 605.2828 | 88.43865 | 93.54383 | 89.8956 |
| Tnfsf15 | 326623 | 6.582091 | 3.853292 | 47.95703 | 3.18E-23 | 3.03E-21 | -0.22393 | 3.853292 | -6.39465 | 2.26E-06 | 9.18E-06 | 89.77967 | 84.36332 | 98.94853 | 9535.123 | 9541.461 | 8937.076 | 8487.77 | 8002.318 | 8393.432 | 684.6009 | 677.1247 | 699.3218 | 104.2909 | 108.4794 | 85.03638 |
| Upp1 | 22271 | 6.602606 | 4.216659 | 35.17796 | 2.18E-20 | 1.02E-18 | 0.871846 | 4.216659 | 20.97418 | 9.96E-16 | 2.15E-14 | 86.06977 | 89.69153 | 78.11726 | 8875.378 | 8594.899 | 8966.834 | 17451.66 | 16909.42 | 16021.53 | 423.8372 | 382.0265 | 342.5706 | 330.3935 | 323.08 | 413.0338 |
| Heatr9 | 629303 | 6.604312 | 0.428658 | 21.89747 | 4.13E-16 | 7.45E-15 | -4.0723 | 0.428658 | -24.7926 | 3.21E-17 | 9.89E-16 | 22.25942 | 17.7607 | 31.24691 | 2213 | 2579.874 | 2574.127 | 159.1812 | 162.191 | 134.4512 | 29.99936 | 13.7255 | 12.6878 | 27.53279 | 16.50774 | 21.05663 |
| Tnc | 21923 | 6.737767 | 2.7424 | 35.13707 | 2.24E-20 | 1.04E-18 | 0.194659 | 2.7424 | 4.430657 | 0.000224 | 0.000688 | 74.94006 | 87.91546 | 65.09772 | 8416.934 | 9131.378 | 8368.955 | 10586.97 | 10471.37 | 9779.314 | 10.769 | 19.82572 | 26.86829 | 105.1252 | 73.10568 | 73.69819 |
| Clic5 | 224796 | 6.755971 | 2.713253 | 17.8975 | 2.47E-14 | 3.16E-13 | 3.003323 | 2.713253 | 48.65619 | 2.34E-23 | 5.04E-21 | 11.87169 | 15.09659 | 19.09533 | 1832.479 | 1815.883 | 1659.723 | 14891.97 | 14404.17 | 14849.39 | 69.99852 | 109.0415 | 52.99023 | 367.1038 | 275.1289 | 290.7434 |
| Csf3 | 12985 | 6.960673 | 1.085591 | 19.3356 | 5.20E-15 | 7.55E-14 | 0.113968 | 1.085591 | 1.619303 | 0.120065 | 0.182165 | 12.61367 | 16.87266 | 20.83127 | 2335.079 | 2203.496 | 2103.398 | 2603.75 | 2289.44 | 2569.512 | 28.46093 | 22.87584 | 8.209754 | 19.18952 | 27.51289 | 13.76779 |
| Serpina3g | 20715 | 7.023718 | 6.801158 | 49.57688 | 1.57E-23 | 1.63E-21 | -0.37143 | 6.801158 | -10.7446 | 4.53E-10 | 3.18E-09 | 209.2386 | 253.978 | 153.6306 | 27905.36 | 28034.54 | 28163.9 | 22511.35 | 23729.49 | 21451.29 | 13888.94 | 13839.88 | 13510.27 | 7508.108 | 7634.434 | 7381.967 |
| Cxcl10 | 15945 | 7.06015 | 7.766443 | 59.95302 | 9.53E-15 | 1.32E-13 | -0.06584 | 7.766443 | -3.62014 | 0.00428 | 0.010162 | 2973.859 | 2761.789 | 2639.496 | 402436.2 | 404314.2 | 388433.4 | 404162.5 | 388289.4 | 394350 | 5631.419 | 6031.595 | 4501.93 | 170.2027 | 143.067 | 163.5938 |
| Slc7a2 | 11988 | 7.064614 | 8.283696 | 35.17396 | 1.70E-14 | 2.25E-13 | 0.341581 | 8.283696 | 7.815495 | 2.52E-06 | 1.02E-05 | 310.148 | 278.843 | 280.3542 | 42050.9 | 42025.12 | 40677.42 | 55478.92 | 56199.86 | 52832.43 | 18086.54 | 18486.73 | 17639.03 | 186200.9 | 183908.7 | 193674 |
| Gbp5 | 229898 | 7.099983 | 6.943702 | 98.21404 | 7.74E-30 | 1.65E-26 | -0.72992 | 6.943702 | -42.4161 | 4.25E-22 | 5.59E-20 | 1008.352 | 1081.627 | 956.5025 | 148443.8 | 151964 | 145303.6 | 93831.64 | 94565.41 | 90947.62 | 2788.402 | 2799.24 | 2486.809 | 553.1587 | 601.3532 | 561.2401 |
| Il23r | 209590 | 7.206169 | -0.68955 | 16.50297 | 1.25E-13 | 1.42E-12 | -4.87823 | -0.68955 | -18.3411 | 1.51E-14 | 2.48E-13 | 9.64575 | 4.440175 | 6.943757 | 1094.812 | 1238.677 | 1090.25 | 49.74413 | 32.17012 | 39.07129 | 10.769 | 18.30067 | 9.702436 | 17.52087 | 25.15464 | 9.718443 |
| Il27 | 246779 | 7.284006 | 3.160583 | 36.87105 | 8.14E-21 | 4.25E-19 | 0.414926 | 3.160583 | 10.96268 | 3.14E-10 | 2.26E-09 | 86.06977 | 79.03511 | 48.6063 | 11700.07 | 11596.09 | 11770.91 | 16428.35 | 16757.95 | 15479.13 | 86.92123 | 89.21576 | 103.7414 | 45.05365 | 34.58764 | 27.53559 |
| 8030451A( | 100504061 | 7.324759 | -3.35488 | 8.012237 | 7.15E-08 | 3.45E-07 | 1.625576 | -3.35488 | 5.690133 | 1.13E-05 | 4.15E-05 | 0 | 0 | 0 | 49.35095 | 85.66811 | 83.8654 | 247.2994 | 243.9568 | 198.8039 | 0 | 0 | 0 | 4.171635 | 9.432991 | 0 |
| Cxcl9 | 17329 | 7.34235 | 2.937575 | 28.93703 | 1.30E-18 | 3.98E-17 | 1.972144 | 2.937575 | 48.29056 | 2.74E-23 | 5.82E-21 | 62.32638 | 87.91546 | 32.11488 | 9759.799 | 10353.2 | 9329.349 | 39586.38 | 38025.08 | 41877.53 | 33.84544 | 29.73859 | 35.82438 | 35.04173 | 29.87114 | 18.62702 |
| Serpina3f | 238393 | 7.40735 | 3.871446 | 30.39185 | 4.67E-19 | 1.57E-17 | 0.28867 | 3.871446 | 6.482518 | 1.86E-06 | 7.63E-06 | 61.5844 | 38.1855 | 40.79457 | 8676.676 | 8549.958 | 8297.264 | 11114.26 | 10912.37 | 10409.05 | 1674.58 | 1673.749 | 1398.644 | 80.09538 | 69.96135 | 78.55742 |
| Ptgs2 | 19225 | 7.639313 | 6.444138 | 29.20677 | 2.35E-11 | 1.84E-10 | 1.62053 | 6.444138 | 41.293 | 6.49E-13 | 7.50E-12 | 432.5748 | 468.8825 | 368.8871 | 90105.74 | 92503.3 | 86992.77 | 285384.9 | 272088.2 | 299813.5 | 916.9036 | 829.6303 | 862.0241 | 434.6843 | 286.9202 | 535.3242 |
| Serpinb2 | 18788 | 8.091379 | 1.791217 | 21.10497 | 8.78E-16 | 1.48E-14 | 2.801341 | 1.791217 | 62.39697 | 1.20E-25 | 5.40E-23 | 5.193865 | 11.54445 | 19.09533 | 3093.525 | 3155.676 | 3023.212 | 22600.89 | 22032.51 | 22441.86 | 16.92272 | 23.63836 | 16.41951 | 20.85817 | 19.65207 | 42.11325 |
| Cxcl11 | 56066 | 8.124414 | 0.959975 | 19.84815 | 3.06E-15 | 4.62E-14 | 0.904593 | 0.959975 | 16.65295 | 1.04E-13 | 1.43E-12 | 7.419807 | 15.98463 | 9.547666 | 3046.772 | 3272.241 | 3228.818 | 6142.689 | 6088.196 | 6283.583 | 10.769 | 16.77561 | 4.478048 | 5.005962 | 22.7964 | 8.098703 |
| Ccl5 | 20304 | 8.214494 | 3.230278 | 56.25297 | 1.08E-24 | 1.66E-22 | -4.09352 | 3.230278 | -78.2179 | 9.84E-28 | 1.85E-24 | 128.3627 | 153.6301 | 149.2908 | 45288.59 | 44846.55 | 45783.75 | 3020.179 | 2840.354 | 2476.43 | 143.8431 | 138.0175 | 137.3268 | 31.70442 | 25.94073 | 12.14805 |
| Edn1 | 13614 | 8.236639 | 2.126905 | 33.05062 | 8.08E-20 | 3.22E-18 | -4.75391 | 2.126905 | -47.5941 | 3.73E-23 | 7.60E-21 | 25.96933 | 41.73764 | 35.58675 | 10987.08 | 10367.25 | 11235.26 | 412.1656 | 406.1478 | 430.9333 | 360.7616 | 408.7149 | 338.8389 | 29.20144 | 21.22423 | 14.57766 |
| Il12b | 16160 | 8.387023 | 1.802385 | 30.49061 | 4.37E-19 | 1.48E-17 | -1.95046 | 1.802385 | -35.0386 | 2.37E-20 | 1.73E-18 | 25.22735 | 74.59494 | 46.87036 | 15602.69 | 16632.25 | 14558.76 | 4191.298 | 3883.202 | 4449.53 | 8.461359 | 13.7255 | 11.94146 | 15.01788 | 11.79124 | 8.098703 |
| Il6 | 16193 | 8.557099 | 3.672188 | 41.43377 | 6.97E-22 | 4.55E-20 | -1.05541 | 3.672188 | -29.9822 | 6.20E-19 | 2.98E-17 | 68.26223 | 117.2206 | 69.43757 | 32520.98 | 33462.53 | 32553.3 | 16074.46 | 15704.38 | 17300.54 | 283.0709 | 224.1832 | 244.0536 | 30.03577 | 9.432991 | 41.30338 |
| Ptx3 | 19288 | 8.561358 | 0.408743 | 10.48396 | 7.06E-10 | 4.49E-09 | 5.029829 | 0.408743 | 45.45788 | 9.85E-23 | 1.70E-20 | 1.483961 | 0 | 1.735939 | 427.2753 | 547.7142 | 519.4244 | 16723.98 | 17615.82 | 16101.97 | 24.61486 | 13.7255 | 8.209754 | 23.36115 | 27.51289 | 14.57766 |
| Lipg | 16891 | 8.639674 | 3.516947 | 40.94744 | 8.94E-22 | 5.68E-20 | -1.0336 | 3.516947 | -30.4622 | 4.45E-19 | 2.23E-17 | 46.74479 | 44.40175 | 56.41802 | 21309.22 | 20459.23 | 21165.19 | 11023.3 | 10890.93 | 10190.12 | 127.6896 | 131.1548 | 123.1463 | 97.61625 | 110.8376 | 110.1424 |
| Il1a | 16175 | 8.792424 | 3.158634 | 33.43915 | 6.53E-20 | 2.58E-18 | 1.860887 | 3.158634 | 66.76882 | 2.85E-26 | 2.02E-23 | 29.67923 | 35.5214 | 28.643 | 15011.78 | 14875.35 | 14776.54 | 55839.92 | 54229.44 | 58033.51 | 69.99852 | 93.79093 | 75.38047 | 23.36115 | 11.00516 | 32.39481 |
| Il19 | 329244 | 9.012558 | -1.64398 | 15.83423 | 2.83E-13 | 3.01E-12 | -5.67766 | -1.64398 | -19.5176 | 4.30E-15 | 7.97E-14 | 7.419807 | 1.77607 | 1.735939 | 1894.817 | 1956.323 | 1800.401 | 29.84648 | 49.5956 | 35.62382 | 0 | 3.050111 | 0.746341 | 5.005962 | 15.72165 | 8.908573 |
| U90926 | 57425 | 9.119503 | -1.66679 | 17.44827 | 4.11E-14 | 5.09E-13 | -2.56248 | -1.66679 | -22.2003 | 3.12E-16 | 7.66E-15 | 2.225942 | 0.888035 | 3.471878 | 1467.541 | 1453.549 | 1339.141 | 279.9884 | 238.5951 | 234.4277 | 0 | 0.762528 | 0 | 5.005962 | 6.288661 | 3.239481 |
| Nos2 | 18126 | 11.0084 | 5.017067 | 32.13612 | 9.41E-15 | 1.30E-13 | -1.04119 | 5.017067 | -39.8597 | 4.45E-16 | 1.05E-14 | 59.35846 | 101.236 | 72.90945 | 165601 | 165654 | 165423.1 | 86787.87 | 85733.37 | 79244.62 | 616.1408 | 696.1879 | 617.9706 | 43.385 | 30.65722 | 46.16261 |

| Supplementary Table S4 | | |
|---|---|---|
| ENTREZID | SYMBOL | GENENAME |
| 20290 | Ccl1 | chemokine (C-C motif) ligand 1 |
| 20292 | Ccl11 | chemokine (C-C motif) ligand 11 |
| 20295 | Ccl17 | chemokine (C-C motif) ligand 17 |
| 24047 | Ccl19 | chemokine (C-C motif) ligand 19 |
| 20296 | Ccl2 | chemokine (C-C motif) ligand 2 |
| 20297 | Ccl20 | chemokine (C-C motif) ligand 20 |
| 20299 | Ccl22 | chemokine (C-C motif) ligand 22 |
| 56221 | Ccl24 | chemokine (C-C motif) ligand 24 |
| 20300 | Ccl25 | chemokine (C-C motif) ligand 25 |
| 541307 | Ccl26 | chemokine (C-C motif) ligand 26 |
| 20301 | Ccl27a | chemokine (C-C motif) ligand 27A |
| 56838 | Ccl28 | chemokine (C-C motif) ligand 28 |
| 20302 | Ccl3 | chemokine (C-C motif) ligand 3 |
| 20303 | Ccl4 | chemokine (C-C motif) ligand 4 |
| 20304 | Ccl5 | chemokine (C-C motif) ligand 5 |
| 20306 | Ccl7 | chemokine (C-C motif) ligand 7 |
| 20307 | Ccl8 | chemokine (C-C motif) ligand 8 |
| 12977 | Csf1 | colony stimulating factor 1 (macrophage) |
| 12981 | Csf2 | colony stimulating factor 2 (granulocyte-macrophage) |
| 12985 | Csf3 | colony stimulating factor 3 (granulocyte) |
| 20312 | Cx3cl1 | chemokine (C-X3-C motif) ligand 1 |
| 14825 | Cxcl1 | chemokine (C-X-C motif) ligand 1 |
| 15945 | Cxcl10 | chemokine (C-X-C motif) ligand 10 |
| 56066 | Cxcl11 | chemokine (C-X-C motif) ligand 11 |
| 20315 | Cxcl12 | chemokine (C-X-C motif) ligand 12 |
| 55985 | Cxcl13 | chemokine (C-X-C motif) ligand 13 |
| 57266 | Cxcl14 | chemokine (C-X-C motif) ligand 14 |
| 66102 | Cxcl16 | chemokine (C-X-C motif) ligand 16 |
| 232983 | Cxcl17 | chemokine (C-X-C motif) ligand 17 |
| 20310 | Cxcl2 | chemokine (C-X-C motif) ligand 2 |
| 330122 | Cxcl3 | chemokine (C-X-C motif) ligand 3 |
| 20311 | Cxcl5 | chemokine (C-X-C motif) ligand 5 |
| 20311 | Cxcl5 | chemokine (C-X-C motif) ligand 5 |
| 17329 | Cxcl9 | chemokine (C-X-C motif) ligand 9 |
| 50498 | Ebi3 | Epstein-Barr virus induced gene 3 |
| 15962 | Ifna1 | interferon alpha 1 |
| 110296 | Ifna10 | interferon alpha 10 |
| 230396 | Ifna13 | interferon alpha 13 |
| 404549 | Ifna14 | interferon alpha 14 |
| 230398 | Ifna16 | interferon alpha 16 |
| 15965 | Ifna2 | interferon alpha 2 |
| 15967 | Ifna4 | interferon alpha 4 |
| 15968 | Ifna5 | interferon alpha 5 |
| 15969 | Ifna6 | interferon alpha 6 |
| 15969 | Ifna6 | interferon alpha 6 |
| 15970 | Ifna7 | interferon alpha 7 |
| 15977 | Ifnb1 | interferon beta 1, fibroblast |

| | | |
|---|---|---|
| 230405 | Ifne | interferon epsilon |
| 15978 | Ifng | interferon gamma |
| 387510 | Ifnk | interferon kappa |
| 330496 | Ifnl2 | interferon lambda 2 |
| 338374 | Ifnl3 | interferon lambda 3 |
| 16153 | Il10 | interleukin 10 |
| 16156 | Il11 | interleukin 11 |
| 16159 | Il12a | interleukin 12a |
| 16160 | Il12b | interleukin 12b |
| 16163 | Il13 | interleukin 13 |
| 16168 | Il15 | interleukin 15 |
| 16170 | Il16 | interleukin 16 |
| 16171 | Il17a | interleukin 17A |
| 56069 | Il17b | interleukin 17B |
| 234836 | Il17c | interleukin 17C |
| 239114 | Il17d | interleukin 17D |
| 257630 | Il17f | interleukin 17F |
| 16173 | Il18 | interleukin 18 |
| 329244 | Il19 | interleukin 19 |
| 16175 | Il1a | interleukin 1 alpha |
| 16176 | Il1b | interleukin 1 beta |
| 16181 | Il1rn | interleukin 1 receptor antagonist |
| 16183 | Il2 | interleukin 2 |
| 58181 | Il20 | interleukin 20 |
| 60505 | Il21 | interleukin 21 |
| 50929 | Il22 | interleukin 22 |
| 83430 | Il23a | interleukin 23, alpha subunit p19 |
| 93672 | Il24 | interleukin 24 |
| 140806 | Il25 | interleukin 25 |
| 246779 | Il27 | interleukin 27 |
| 16187 | Il3 | interleukin 3 |
| 76399 | Il31 | interleukin 31 |
| 77125 | Il33 | interleukin 33 |
| 76527 | Il34 | interleukin 34 |
| NA | Il36a | NA |
| NA | Il36b | NA |
| NA | Il36g | NA |
| 16189 | Il4 | interleukin 4 |
| 16191 | Il5 | interleukin 5 |
| 16193 | Il6 | interleukin 6 |
| 16196 | Il7 | interleukin 7 |
| 16198 | Il9 | interleukin 9 |
| 17311 | Kitl | kit ligand |
| 16878 | Lif | leukemia inhibitory factor |
| 16992 | Lta | lymphotoxin A |
| 16994 | Ltb | lymphotoxin B |
| 28106 | Mydgf | myeloid derived growth factor |
| 18413 | Osm | oncostatin M |
| 57349 | Ppbp | pro-platelet basic protein |

| | | |
|---|---|---|
| 21803 | Tgfb1 | transforming growth factor, beta 1 |
| 21808 | Tgfb2 | transforming growth factor, beta 2 |
| 21809 | Tgfb3 | transforming growth factor, beta 3 |
| 21926 | Tnf | tumor necrosis factor |
| 22035 | Tnfsf10 | tumor necrosis factor (ligand) superfamily, member 10 |
| 21943 | Tnfsf11 | tumor necrosis factor (ligand) superfamily, member 11 |
| 21944 | Tnfsf12 | tumor necrosis factor (ligand) superfamily, member 12 |
| 69583 | Tnfsf13 | tumor necrosis factor (ligand) superfamily, member 13 |
| 24099 | Tnfsf13b | tumor necrosis factor (ligand) superfamily, member 13b |
| 50930 | Tnfsf14 | tumor necrosis factor (ligand) superfamily, member 14 |
| 326623 | Tnfsf15 | tumor necrosis factor (ligand) superfamily, member 15 |
| 240873 | Tnfsf18 | tumor necrosis factor (ligand) superfamily, member 18 |
| 22164 | Tnfsf4 | tumor necrosis factor (ligand) superfamily, member 4 |
| 21949 | Tnfsf8 | tumor necrosis factor (ligand) superfamily, member 8 |
| 21950 | Tnfsf9 | tumor necrosis factor (ligand) superfamily, member 9 |
| 16963 | Xcl1 | chemokine (C motif) ligand 1 |

**Supplementary Table S5**

| | LPS vs CNT | | | | | DB+LPS vs LPS | | | | | Read counts | | | | | | | | |
|---|---|---|---|---|---|---|---|---|---|---|---|---|---|---|---|---|---|---|---|
| SYMBOL | logFC | AveExpr | t | P.Value | adj.P.Val | logFC | AveExpr | t | P.Value | adj.P.Val | CNT1 | CNT2 | CNT3 | LPS1 | LPS2 | LPS3 | DB+LPS1 | DB+LPS2 | DB+LPS3 |
| Cxcl10 | 7.06015 | 7.766443 | 59.95302 | 9.53E-15 | 1.32E-13 | -0.06584 | 7.766443 | -3.62014 | 0.00428 | 0.010162 | 2973.859 | 2761.789 | 2639.496 | 402436.2 | 404314.2 | 388433.4 | 404162.5 | 388289.4 | 394350 |
| Ccl3 | 4.147718 | 8.225407 | 151.1087 | 7.95E-34 | 2.09E-29 | -2.86833 | 8.225407 | -129 | 2.32E-32 | 6.10E-28 | 12707.9 | 12580.79 | 11931.98 | 236062.5 | 239773.8 | 229216.3 | 33320.04 | 32613.8 | 34136.82 |
| Il1rn | 4.367306 | 8.262082 | 137.976 | 5.53E-33 | 4.84E-29 | -0.84154 | 8.262082 | -58.474 | 4.76E-25 | 1.63E-22 | 9034.358 | 7963.898 | 9202.214 | 192055.7 | 191458.4 | 193996.9 | 110629.5 | 110961.5 | 112654 |
| Ccl4 | 4.182658 | 8.127998 | 141.0763 | 3.44E-33 | 4.52E-29 | -2.77253 | 8.127998 | -118.278 | 1.47E-31 | 1.94E-27 | 9466.932 | 9479.773 | 8557.312 | 178389.4 | 180758.3 | 174289.9 | 27286.79 | 27139.52 | 26651.22 |
| Ccl7 | 2.843772 | 9.076145 | 70.11407 | 6.53E-23 | 5.57E-21 | -0.65789 | 9.076145 | -23.4831 | 1.09E-14 | 1.85E-13 | 12051.25 | 12201.6 | 11478.9 | 91081.07 | 93222.35 | 89691.34 | 60224.51 | 56849.97 | 62769.18 |
| Ccl2 | 2.042757 | 8.976411 | 37.75698 | 4.63E-14 | 5.66E-13 | -0.42548 | 8.976411 | -9.82095 | 3.64E-07 | 1.65E-06 | 15253.64 | 15069.07 | 15373.48 | 66995.21 | 67417.99 | 66667.58 | 54131.56 | 50383.77 | 51169.6 |
| Tnf | 2.504464 | 7.123557 | 97.80553 | 8.45E-30 | 1.65E-26 | -1.50185 | 7.123557 | -65.2631 | 4.62E-26 | 2.96E-23 | 10470.09 | 10643.99 | 10059.77 | 62682.2 | 64123.28 | 62111.8 | 23436.59 | 23200.02 | 22659.05 |
| Ccl5 | 8.214494 | 3.230278 | 56.25297 | 1.08E-24 | 1.66E-22 | -4.09352 | 3.230278 | -78.2179 | 9.84E-28 | 1.85E-24 | 128.3627 | 153.6301 | 149.2908 | 45288.59 | 44846.55 | 45783.75 | 3020.179 | 2840.354 | 2476.43 |
| Il1b | 6.284428 | 5.306162 | 77.57245 | 1.17E-27 | 4.82E-25 | -1.08569 | 5.306162 | -38.286 | 3.68E-21 | 3.46E-19 | 546.8398 | 594.0954 | 505.1583 | 45214.56 | 45944.79 | 45574.08 | 23354.16 | 22519.09 | 21237.54 |
| Il6 | 8.557099 | 3.672188 | 41.43377 | 6.97E-22 | 4.55E-20 | -1.05541 | 3.672188 | -29.9822 | 6.20E-19 | 2.98E-17 | 68.26223 | 117.2206 | 69.43757 | 32520.98 | 33462.53 | 32553.3 | 16074.46 | 15704.38 | 17300.54 |
| Tnfsf10 | 5.321782 | 4.994094 | 80.00138 | 6.09E-28 | 3.14E-25 | -1.98482 | 4.994094 | -55.7232 | 1.32E-24 | 3.82E-22 | 524.5804 | 632.2809 | 560.7084 | 23954.69 | 25199.06 | 23739.32 | 6313.241 | 6505.067 | 6287.03 |
| Il10 | 3.474459 | 4.422672 | 59.52797 | 3.26E-25 | 6.07E-23 | -3.78573 | 4.422672 | -54.2756 | 2.31E-24 | 6.46E-22 | 1643.487 | 1512.324 | 1392.223 | 18201.41 | 17991.71 | 17843.04 | 1388.572 | 1264.018 | 1410.014 |
| Il12b | 8.387023 | 1.802385 | 30.49061 | 4.37E-19 | 1.48E-17 | -1.95046 | 1.802385 | -35.0386 | 2.37E-20 | 1.73E-18 | 25.22735 | 74.59494 | 46.87036 | 15602.69 | 16632.25 | 14558.76 | 4191.298 | 3883.202 | 4449.53 |
| Il1a | 8.792424 | 3.158634 | 33.43915 | 6.33E-20 | 2.58E-18 | 1.860887 | 3.158634 | 66.76882 | 2.85E-26 | 2.02E-23 | 29.67923 | 35.5214 | 28.643 | 15011.78 | 14875.35 | 14776.54 | 55839.92 | 54229.44 | 58033.51 |
| Il27 | 7.284006 | 3.160583 | 36.87105 | 8.14E-21 | 4.25E-19 | 0.414926 | 3.160583 | 10.96268 | 3.14E-10 | 2.26E-09 | 86.06977 | 79.03511 | 48.6063 | 11700.07 | 11596.09 | 11770.91 | 16428.35 | 16757.95 | 15479.13 |
| Il15 | 3.829075 | 5.333139 | 59.28031 | 3.56E-25 | 6.50E-23 | 0.091534 | 5.333139 | 2.571452 | 0.017657 | 0.035401 | 678.1704 | 682.8989 | 754.2656 | 10823.44 | 10673.4 | 10534.58 | 11674.24 | 12209.9 | 11575.44 |
| Cxcl9 | 7.34235 | 2.937575 | 28.93703 | 1.30E-18 | 3.98E-17 | 1.972144 | 2.937575 | 48.29056 | 2.74E-23 | 5.82E-21 | 62.32638 | 87.91546 | 32.11488 | 9759.799 | 10353.2 | 9329.349 | 39586.38 | 38025.08 | 41877.53 |
| Tnfsf15 | 6.582091 | 3.853292 | 47.95703 | 3.18E-23 | 3.03E-21 | -0.22393 | 3.853292 | -6.39465 | 2.26E-06 | 9.18E-06 | 89.77967 | 84.36332 | 98.94853 | 9535.123 | 9541.461 | 8937.076 | 8487.77 | 8002.318 | 8393.432 |
| Csf1 | 3.991669 | 5.488941 | 46.85507 | 5.19E-23 | 4.54E-21 | 2.109122 | 5.488941 | 57.53067 | 6.72E-25 | 2.13E-22 | 423.671 | 516.8364 | 472.1755 | 7833.814 | 8031.737 | 7941.512 | 36307.53 | 35988.98 | 34513.74 |
| Cxcl2 | 4.040087 | 5.552062 | 46.06973 | 7.42E-23 | 6.08E-21 | 2.954872 | 5.552062 | 83.34279 | 2.55E-28 | 9.58E-25 | 405.1215 | 487.5312 | 460.8919 | 7763.683 | 7634.293 | 8203.93 | 64253.78 | 61026.72 | 64465.33 |
| Il18 | 3.882757 | 3.592549 | 50.98458 | 8.69E-24 | 9.81E-22 | -3.01607 | 3.592549 | -43.5595 | 2.43E-22 | 3.69E-20 | 365.7965 | 456.45 | 390.5863 | 6181.856 | 6484.093 | 6275.026 | 824.3313 | 844.4657 | 767.6359 |
| Ccl22 | 3.605597 | 4.857653 | 28.79378 | 1.44E-18 | 4.35E-17 | 1.473747 | 4.857653 | 24.50407 | 4.09E-17 | 1.23E-15 | 348.731 | 312.5883 | 328.9605 | 3998.726 | 4369.074 | 4578.78 | 12325.17 | 13107.98 | 11896.06 |
| Cxcl11 | 8.124414 | 0.959975 | 19.84815 | 3.06E-15 | 4.62E-14 | 0.904593 | 0.959975 | 16.65295 | 1.04E-13 | 1.43E-12 | 7.419807 | 15.98463 | 9.547666 | 3046.772 | 3272.241 | 3228.818 | 6142.689 | 6088.196 | 6283.583 |
| Il36g | 3.076403 | 3.301592 | 30.743 | 3.68E-19 | 1.28E-17 | 0.890301 | 3.301592 | 15.92803 | 2.52E-13 | 3.18E-12 | 323.5036 | 237.9934 | 322.0167 | 2584.431 | 2574.257 | 2790.554 | 5143.543 | 5124.432 | 5022.959 |
| Csf3 | 6.960673 | 1.085591 | 19.3356 | 5.20E-15 | 7.55E-14 | 0.113968 | 1.085591 | 1.619303 | 0.120065 | 0.182165 | 12.61367 | 16.87266 | 20.83127 | 2335.079 | 2203.496 | 2103.398 | 2603.75 | 2289.44 | 2569.512 |
| Tnfsf9 | 1.636028 | 3.887836 | 23.17624 | 1.29E-16 | 2.59E-15 | 1.021555 | 3.887836 | 19.19007 | 6.06E-15 | 1.08E-13 | 667.7827 | 613.6322 | 637.0897 | 2132.48 | 2089.74 | 2157.505 | 4603.464 | 4556.094 | 4315.079 |
| Il19 | 9.012558 | -1.64398 | 15.83423 | 2.83E-13 | 3.01E-12 | -5.67766 | -1.64398 | -19.5176 | 4.30E-15 | 7.97E-14 | 7.419807 | 1.77607 | 1.735939 | 1894.817 | 1956.323 | 1800.401 | 29.84648 | 49.5956 | 35.62382 |
| Osm | 1.64244 | 4.66686 | 17.78033 | 2.82E-14 | 3.57E-13 | 3.494029 | 4.66686 | 59.37689 | 3.44E-25 | 1.27E-22 | 348.731 | 413.8243 | 414.0215 | 1207.8 | 1379.116 | 1312.088 | 15397.94 | 15075.72 | 15041.3 |
| Il33 | 6.470987 | -0.42961 | 12.29861 | 3.72E-11 | 2.82E-10 | 1.515547 | -0.42961 | 14.62478 | 1.35E-12 | 1.47E-11 | 11.12971 | 0.888035 | 14.75548 | 592.2114 | 636.1911 | 570.8258 | 1841.954 | 1687.591 | 1806.473 |
| Tnfsf4 | 2.216474 | 2.874238 | 13.06296 | 1.19E-11 | 9.75E-11 | 4.216307 | 2.874238 | 46.32579 | 6.60E-23 | 1.21E-20 | 101.6514 | 100.348 | 131.9314 | 579.2243 | 574.3977 | 492.3711 | 10565.65 | 9919.121 | 11184.73 |
| Cxcl1 | 2.901089 | 2.506917 | 12.64763 | 2.20E-11 | 1.73E-10 | 4.935944 | 2.506917 | 48.30162 | 2.73E-23 | 5.82E-21 | 67.52025 | 63.93852 | 69.43757 | 544.1591 | 497.1559 | 570.8258 | 16971.28 | 16995.21 | 17232.74 |
| Il36a | 3.945696 | -0.85349 | 13.08806 | 1.15E-11 | 9.42E-11 | 0.955233 | -0.85349 | 7.837187 | 1.02E-07 | 5.01E-07 | 20.77546 | 22.20087 | 27.77503 | 403.8985 | 394.6351 | 369.2783 | 847.0715 | 682.2747 | 828.5412 |
| Ccl8 | 3.9531 | 1.858231 | 12.12307 | 4.88E-11 | 3.63E-10 | 0.237623 | 1.858231 | 1.657313 | 0.112084 | 0.172021 | 23.0014 | 19.53677 | 19.9633 | 319.4825 | 334.2461 | 408.5057 | 432.0633 | 443.6796 | 418.2926 |
| Il7 | 2.841419 | 0.654604 | 10.84839 | 3.80E-10 | 2.51E-09 | 0.193621 | 0.654604 | 1.243301 | 0.227242 | 0.312147 | 34.8731 | 33.74533 | 46.00239 | 283.1186 | 276.6659 | 315.1716 | 308.4136 | 414.1903 | 325.211 |
| Ifnb1 | 6.066922 | -0.32046 | 8.627643 | 2.11E-08 | 1.08E-07 | 2.809782 | -0.32046 | 17.49687 | 3.89E-14 | 5.94E-13 | 7.419807 | 15.09659 | 0 | 267.5341 | 328.6285 | 225.8955 | 2035.246 | 1946.292 | 1942.073 |
| Tnfsf8 | 2.663778 | 0.657648 | 11.09724 | 2.51E-10 | 1.69E-09 | 1.548133 | 0.657648 | 12.0377 | 5.57E-11 | 4.54E-10 | 32.64715 | 31.96926 | 39.9266 | 206.4948 | 255.5999 | 251.5962 | 722.0005 | 760.0191 | 681.4493 |
| Il23a | 3.585408 | -1.10746 | 8.568598 | 2.36E-08 | 1.20E-07 | -0.89353 | -1.10746 | -3.4501 | 0.002357 | 0.005954 | 7.419807 | 19.53677 | 7.811726 | 161.0399 | 119.3736 | 129.8561 | 75.32683 | 58.97856 | 94.23076 |
| Il12a | 4.356957 | -0.70704 | 7.808291 | 1.08E-07 | 5.10E-07 | 2.72364 | -0.70704 | 14.34991 | 1.95E-12 | 2.07E-11 | 8.161788 | 3.55214 | 9.547666 | 158.4425 | 130.6088 | 174.4941 | 1092.95 | 1156.784 | 951.5008 |
| Cxcl3 | 2.983505 | 0.859703 | 8.67803 | 1.91E-08 | 9.89E-08 | 4.835984 | 0.859703 | 31.86437 | 1.74E-19 | 9.81E-18 | 28.19527 | 18.64873 | 9.547666 | 149.3516 | 162.9099 | 142.0301 | 4552.299 | 4535.987 | 4417.354 |
| Lta | 2.966258 | -1.17594 | 6.427812 | 2.10E-06 | 8.37E-06 | -0.55172 | -1.17594 | -1.75511 | 0.093587 | 0.147426 | 5.935846 | 15.98463 | 20.83127 | 137.6632 | 87.07251 | 93.33407 | 51.16539 | 81.76573 | 90.78329 |
| Lif | 2.590828 | 0.06511 | 5.095799 | 4.56E-05 | 0.000156 | 3.376696 | 0.06511 | 12.76374 | 1.85E-11 | 1.65E-10 | 8.161788 | 14.20856 | 5.207818 | 77.92255 | 32.30109 | 68.98605 | 608.2996 | 651.445 | 587.2185 |
| Ifna5 | 3.480465 | -3.99902 | 4.349083 | 0.000273 | 0.000848 | -4.05253 | -3.99902 | -4.64702 | 0.000133 | 0.000423 | 5.193865 | 0 | 0.86797 | 29.87031 | 14.04395 | 12.17401 | 0 | 0 | 2.298311 |
| Ccl11 | 3.757763 | -2.25318 | 3.958177 | 0.0007 | 0.002061 | 3.416265 | -2.25318 | 6.489691 | 1.83E-06 | 7.52E-06 | 0.741981 | 1.77607 | 0 | 23.37676 | 7.021976 | 16.23201 | 211.7679 | 142.0847 | 148.2411 |
| Tnfsf18 | 2.183784 | -0.38474 | 4.054169 | 0.000556 | 0.001659 | 4.948066 | -0.38474 | 16.12532 | 1.97E-13 | 2.56E-12 | 3.709904 | 11.54445 | 3.471878 | 18.18193 | 36.51428 | 29.75869 | 911.0282 | 845.8061 | 889.4464 |
| Csf2 | 2.114597 | -2.84466 | 2.72046 | 0.0127 | 0.029764 | 0.785347 | -2.84466 | 1.350639 | 0.190959 | 0.269703 | 5.193865 | 0.888035 | 5.207818 | 14.2858 | 14.04395 | 18.93735 | 44.05909 | 25.46801 | 21.83396 |
| Il13 | 1.965449 | -3.02529 | 2.911599 | 0.008252 | 0.020151 | 0.03998 | -3.02529 | 0.069352 | 0.945355 | 0.954831 | 5.193865 | 0.888035 | 3.471878 | 11.68838 | 9.830767 | 16.23201 | 8.527565 | 17.42548 | 14.93902 |
| Il22 | 2.257256 | -4.36638 | 2.522271 | 0.019658 | 0.043894 | 0.829277 | -4.36638 | 1.122251 | 0.274223 | 0.365564 | 2.225942 | 0 | 0.86797 | 9.090964 | 5.617581 | 2.705335 | 2.842522 | 21.44675 | 14.93902 |
| Ifng | 4.296229 | -3.81324 | 5.124985 | 4.26E-05 | 0.000146 | 0.567605 | -3.81324 | 0.946876 | 0.3543 | 0.451665 | 0 | 0 | 0 | 7.792255 | 9.830767 | 6.763339 | 9.948826 | 26.80843 | 8.044089 |
| Ifne | 2.724809 | -5.19417 | 2.547004 | 0.018627 | 0.041875 | 0.182662 | -5.19417 | 0.195021 | 0.847221 | 0.910542 | 0 | 0 | 0 | 5.194837 | 5.617581 | 0 | 1.421261 | 4.021265 | 3.447467 |

**Supplementary Table S6**

| SYMBOL | Gene ID | FC in Log2 | Adj P |
|---|---|---|---|
| Dmbt1 | 12945 | -5.15140344 | 0.03765 |
| Krt8 | 16691 | -4.91459097 | 0.001062 |
| Krt18 | 16668 | -4.15383813 | 0.006624 |
| Krt19 | 16669 | -2.46681119 | 0.018023 |
| Nr1i2 | 18171 | -1.07823524 | 0.00361 |
| Hspa1b | 15511 | 1.029913253 | 0.000159 |
| Rgs16 | 19734 | 1.062747523 | 0.044919 |
| Hspa1a | 193740 | 1.068970143 | 0.012221 |
| Cxcl9 | 17329 | 1.074382752 | 0.01098 |
| Hbegf | 15200 | 1.081050545 | 0.049708 |
| Hspb1 | 15507 | 1.088775634 | 0.012221 |
| Ccn1 | 16007 | 1.166112611 | 0.027244 |
| Sdcbp2 | 228765 | 1.178966234 | 0.023988 |
| A130040M | 319269 | 1.184151945 | 0.023988 |
| Stk32c | 57740 | 1.224519902 | 0.049124 |
| Ear2 | 13587 | 1.23420969 | 0.006964 |
| Prdm1 | 12142 | 1.236140532 | 0.041507 |
| Nedd9 | 18003 | 1.276436742 | 0.027093 |
| Gimap8 | 243374 | 1.286247891 | 0.034398 |
| Akap12 | 83397 | 1.314042939 | 0.025467 |
| Gem | 14579 | 1.316258751 | 0.006562 |
| 2900026A( | 243219 | 1.334555941 | 0.049124 |
| AA467197 | 433470 | 1.341398734 | 0.031555 |
| Ms4a4d | 66607 | 1.36833793 | 0.041259 |
| Thy1 | 21838 | 1.418840209 | 0.033629 |
| Cdc42ep3 | 260409 | 1.441630669 | 0.040397 |
| Angptl4 | 57875 | 1.47146538 | 0.042044 |
| Ccl11 | 20292 | 1.520432792 | 0.037246 |
| Cfap69 | 207686 | 1.521606141 | 0.023988 |
| Spata13 | 219140 | 1.524043769 | 0.0105 |
| Ecscr | 68545 | 1.541232523 | 0.031335 |
| Kank3 | 80880 | 1.551560229 | 0.043539 |
| 4930523C( | 67647 | 1.552384727 | 0.023988 |
| Spry1 | 24063 | 1.553222325 | 0.015398 |
| Rnf122 | 68867 | 1.575502369 | 0.022946 |
| Gimap1 | 16205 | 1.576677016 | 0.006624 |
| Cav1 | 12389 | 1.595116747 | 0.022804 |
| Id1 | 15901 | 1.618530249 | 0.02843 |
| Ece1 | 230857 | 1.61968325 | 0.040397 |
| Ramp2 | 54409 | 1.639924875 | 0.037345 |
| Gimap6 | 231931 | 1.645401558 | 0.019611 |
| Zeb1 | 21417 | 1.677131629 | 0.014427 |
| Fabp4 | 11770 | 1.700044325 | 0.025467 |
| Kitl | 17311 | 1.704188314 | 0.033075 |
| F2r | 14062 | 1.715130437 | 0.044451 |
| Prr5 | 109270 | 1.71882282 | 0.040081 |
| Id3 | 15903 | 1.724542004 | 0.023079 |

| | | | |
|---|---|---|---|
| Kcnj2 | 16518 | 1.725837959 | 0.037664 |
| Trp53i11 | 277414 | 1.728618898 | 0.013179 |
| Cd200 | 17470 | 1.74017338 | 0.026506 |
| Pcdh17 | 219228 | 1.757542287 | 0.025467 |
| Cyb561 | 13056 | 1.757906729 | 0.038744 |
| Gimap5 | 317757 | 1.763417466 | 0.013712 |
| 6430562O | 320893 | 1.765221988 | 0.049124 |
| Kcnj15 | 16516 | 1.775307786 | 0.023988 |
| Adamts7 | 108153 | 1.776164697 | 0.031888 |
| Msrb3 | 320183 | 1.78221032 | 0.030809 |
| Dll1 | 13388 | 1.78550618 | 0.023201 |
| Angpt2 | 11601 | 1.787200556 | 0.021727 |
| Ttc9 | 69480 | 1.793236428 | 0.021814 |
| Inhbb | 16324 | 1.795667159 | 0.049124 |
| Adamts9 | 101401 | 1.797277776 | 0.016143 |
| Gimap4 | 107526 | 1.798442499 | 0.006562 |
| Epas1 | 13819 | 1.819465985 | 0.020349 |
| Nid1 | 18073 | 1.821496752 | 0.028683 |
| Itga2 | 16398 | 1.822484183 | 0.049489 |
| Arhgef5 | 54324 | 1.827154465 | 0.028683 |
| Lamb1 | 16777 | 1.827249018 | 0.042013 |
| Celf4 | 108013 | 1.829809831 | 0.044177 |
| Tll1 | 21892 | 1.838497568 | 0.049708 |
| Igfbp7 | 29817 | 1.8444599 | 0.043872 |
| Epha2 | 13836 | 1.856672305 | 0.023923 |
| Clca3a1 | 12722 | 1.857596672 | 0.028683 |
| Piezo2 | 667742 | 1.858316961 | 0.036319 |
| Adgrg3 | 54672 | 1.858626418 | 0.014208 |
| Pxdc1 | 66895 | 1.861012492 | 0.026971 |
| Pxdn | 69675 | 1.871972915 | 0.049124 |
| Pdgfb | 18591 | 1.882992151 | 0.027093 |
| Ddah2 | 51793 | 1.887762859 | 0.044441 |
| Trim72 | 434246 | 1.897104706 | 0.026983 |
| Chst1 | 76969 | 1.899972456 | 0.028427 |
| Adgrf5 | 224792 | 1.901344228 | 0.016946 |
| Edn1 | 13614 | 1.905370993 | 0.002152 |
| Mpp3 | 13384 | 1.905718832 | 0.023431 |
| Pde2a | 207728 | 1.906250739 | 0.022946 |
| Rhbdl2 | 230726 | 1.912017757 | 0.023988 |
| Flt1 | 14254 | 1.918196661 | 0.019802 |
| Plcb1 | 18795 | 1.919614016 | 0.023431 |
| Vash1 | 238328 | 1.922584993 | 0.017869 |
| Tshz2 | 228911 | 1.927135701 | 0.023988 |
| Kctd12b | 207474 | 1.928605506 | 0.031034 |
| Nrarp | 67122 | 1.934914029 | 0.033629 |
| Bean1 | 65115 | 1.939336155 | 0.016946 |
| Notch4 | 18132 | 1.948018912 | 0.018933 |
| Tafa3 | 329731 | 1.948504965 | 0.041507 |
| Arhgef28 | 110596 | 1.949407619 | 0.021398 |

| | | | |
|---|---|---|---|
| NA | 1.01E+08 | 1.957321315 | 0.000906 |
| Itga3 | 16400 | 1.957824646 | 0.006562 |
| Afap1l1 | 106877 | 1.958414151 | 0.030523 |
| Nt5e | 23959 | 1.961424879 | 0.013695 |
| Rab3b | 69908 | 1.962831485 | 0.015969 |
| Csgalnact1 | 234356 | 1.964393252 | 0.049708 |
| Lysmd2 | 70082 | 1.966862951 | 0.012221 |
| Fam131a | 78408 | 1.968206668 | 0.049124 |
| Erg | 13876 | 1.975675689 | 0.021814 |
| Prex2 | 109294 | 1.975938617 | 0.008868 |
| F11r | 16456 | 1.983814932 | 0.013179 |
| Ptk7 | 71461 | 1.986166583 | 0.037592 |
| Meox1 | 17285 | 1.993136631 | 0.047994 |
| Dchs1 | 233651 | 1.993178479 | 0.027519 |
| Atp8b1 | 54670 | 1.993574248 | 0.048446 |
| Dysf | 26903 | 1.993966962 | 0.014024 |
| Sncaip | 67847 | 1.996377307 | 0.039008 |
| Ubd | 24108 | 1.996526216 | 0.006624 |
| Trim16 | 94092 | 1.999001242 | 0.027093 |
| S1pr1 | 13609 | 1.999473198 | 0.00995 |
| Egfl7 | 353156 | 2.000655982 | 0.013141 |
| Lrrc3b | 218763 | 2.004213092 | 0.042561 |
| Shank3 | 58234 | 2.008019185 | 0.013088 |
| Septin4 | 18952 | 2.010371938 | 0.023988 |
| Nos3 | 18127 | 2.010661878 | 0.028683 |
| Tmem204 | 407831 | 2.010876456 | 0.023988 |
| Pcdh12 | 53601 | 2.01206906 | 0.044441 |
| Hoxd8 | 15437 | 2.012457518 | 0.025722 |
| Apln | 30878 | 2.013710977 | 0.01098 |
| Alox12 | 11684 | 2.01462351 | 0.012617 |
| Fgd5 | 232237 | 2.01772702 | 0.023923 |
| Hoxb7 | 15415 | 2.017743242 | 0.037592 |
| Ror2 | 26564 | 2.018718109 | 0.033629 |
| Ackr3 | 12778 | 2.019831506 | 0.030057 |
| Kcnq1 | 16535 | 2.025731118 | 0.043028 |
| Emp2 | 13731 | 2.026373918 | 0.023431 |
| Kcne3 | 57442 | 2.032077128 | 0.027093 |
| Vsig2 | 57276 | 2.033058264 | 0.016411 |
| Lama3 | 16774 | 2.03522978 | 0.016946 |
| Spns2 | 216892 | 2.035258401 | 0.016527 |
| Chst7 | 60322 | 2.037257529 | 0.046753 |
| Ppm1j | 71887 | 2.044525005 | 0.006562 |
| Ushbp1 | 234395 | 2.053189262 | 0.0169 |
| Col13a1 | 12817 | 2.058585604 | 0.025914 |
| Ccn2 | 14219 | 2.065833515 | 0.006562 |
| Fam110d | 72690 | 2.069803231 | 0.023708 |
| Ly6c1 | 17067 | 2.071345079 | 0.006964 |
| Cavin3 | 109042 | 2.076870943 | 0.023079 |
| Ct55 | 75013 | 2.079975117 | 0.041507 |

| | | | |
|---|---|---|---|
| Hecw2 | 329152 | 2.080503974 | 0.014427 |
| Disp2 | 214240 | 2.081758181 | 0.049708 |
| Sox18 | 20672 | 2.082367318 | 0.026983 |
| Tcim | 69068 | 2.08356601 | 0.023988 |
| Plxna2 | 18845 | 2.090028475 | 0.006101 |
| Plvap | 84094 | 2.090413153 | 0.013695 |
| C1qtnf1 | 56745 | 2.095800046 | 0.048446 |
| Bok | 51800 | 2.096431447 | 0.046327 |
| Cdh5 | 12562 | 2.100065119 | 0.010057 |
| Rasip1 | 69903 | 2.101290731 | 0.014427 |
| Cxcl12 | 20315 | 2.103862183 | 0.001062 |
| Fam171a2 | 217219 | 2.103947285 | 0.030523 |
| Pecam1 | 18613 | 2.105021137 | 0.015424 |
| Sox13 | 20668 | 2.106698818 | 0.027093 |
| Kdr | 16542 | 2.112120739 | 0.014427 |
| Gpr4 | 319197 | 2.114697028 | 0.006964 |
| Clca2 | 229933 | 2.117446204 | 0.01098 |
| Gm14207 | 1E+08 | 2.120121423 | 0.042591 |
| Clec1a | 243653 | 2.1209357 | 0.023923 |
| Bcl6b | 12029 | 2.121422429 | 0.009416 |
| Cmtm8 | 70031 | 2.125223395 | 0.049708 |
| Gprc5a | 232431 | 2.12730262 | 0.01098 |
| Sema3f | 20350 | 2.127474857 | 0.014014 |
| Wscd1 | 216881 | 2.129014972 | 0.030005 |
| Cdkn2b | 12579 | 2.131305048 | 0.025467 |
| Tbxa2r | 21390 | 2.13646579 | 0.045124 |
| Prnd | 26434 | 2.14124199 | 0.014452 |
| Gpihbp1 | 68453 | 2.147667402 | 0.015175 |
| Ctla2a | 13024 | 2.148528191 | 0.007667 |
| Sema3a | 20346 | 2.149242276 | 0.006254 |
| Lpar4 | 78134 | 2.153355128 | 0.033387 |
| Fam167b | 230766 | 2.170567778 | 0.011432 |
| Mmrn2 | 105450 | 2.171582268 | 0.013972 |
| Tspan18 | 241556 | 2.17235272 | 0.016106 |
| Vwa1 | 246228 | 2.174633095 | 0.015462 |
| Dll4 | 54485 | 2.175284155 | 0.014191 |
| Ebf3 | 13593 | 2.179744454 | 0.025467 |
| Cd300lg | 52685 | 2.192302078 | 0.016143 |
| Esam | 69524 | 2.19281575 | 0.013141 |
| Sox17 | 20671 | 2.193353115 | 0.023708 |
| Robo4 | 74144 | 2.193936585 | 0.006624 |
| C1qtnf9 | 239126 | 2.194202778 | 0.012287 |
| Adgrl4 | 170757 | 2.196834184 | 0.012617 |
| Palmd | 114301 | 2.199178814 | 0.007667 |
| Grb10 | 14783 | 2.204507385 | 0.014452 |
| Pcdh19 | 279653 | 2.208024692 | 0.006101 |
| Cpne8 | 66871 | 2.211816565 | 0.049124 |
| Emcn | 59308 | 2.21251932 | 0.013088 |
| Smoc2 | 64074 | 2.213998581 | 0.033326 |

| | | | |
|---|---|---|---|
| Lmcd1 | 30937 | 2.219813801 | 0.02741 |
| Arhgef15 | 442801 | 2.219889708 | 0.00858 |
| Khdrbs3 | 13992 | 2.22094729 | 0.019169 |
| Ldb2 | 16826 | 2.225152596 | 0.01076 |
| Esm1 | 71690 | 2.226436072 | 0.018813 |
| Myct1 | 68632 | 2.229229917 | 0.0105 |
| Prrg3 | 208748 | 2.233343346 | 0.006964 |
| Hey1 | 15213 | 2.233700243 | 0.023708 |
| Adcy4 | 104110 | 2.234012144 | 0.006624 |
| Fat4 | 329628 | 2.240037779 | 0.008329 |
| Efnb2 | 13642 | 2.240476681 | 0.0105 |
| Efna1 | 13636 | 2.243676818 | 0.014427 |
| Tie1 | 21846 | 2.247249947 | 0.013141 |
| Sox7 | 20680 | 2.259909238 | 0.012617 |
| Ccdc3 | 74186 | 2.262091897 | 0.049124 |
| Chst2 | 54371 | 2.268951538 | 0.010057 |
| Cyyr1 | 224405 | 2.275643365 | 0.046753 |
| She | 214547 | 2.278782988 | 0.012617 |
| Tmem47 | 192216 | 2.285007899 | 0.012415 |
| Unc45b | 217012 | 2.287300867 | 0.006562 |
| Rnd1 | 223881 | 2.303232012 | 0.008385 |
| Tmem88 | 67020 | 2.303911317 | 0.006562 |
| Igfbp3 | 16009 | 2.304388803 | 0.010057 |
| Podxl | 27205 | 2.309314007 | 0.006624 |
| NA | 1.01E+08 | 2.326714879 | 0.037664 |
| Exoc3l2 | 74463 | 2.343345074 | 0.012382 |
| Mall | 228576 | 2.363022132 | 0.006102 |
| Ptprb | 19263 | 2.366416999 | 0.006102 |
| Meox2 | 17286 | 2.370850242 | 0.006562 |
| Ccm2l | 228788 | 2.378367299 | 0.011065 |
| Gm19590 | 1.01E+08 | 2.383476328 | 0.033387 |
| C130074G | 226777 | 2.394788001 | 0.006964 |
| Tacr1 | 21336 | 2.463040837 | 0.026009 |
| Nkd2 | 72293 | 2.475574902 | 0.043028 |
| Fzd8 | 14370 | 2.475975929 | 0.023708 |
| Ablim3 | 319713 | 2.4953019 | 0.016143 |
| Hoxd9 | 15438 | 2.50269715 | 0.049708 |
| Cpne7 | 102278 | 2.507331542 | 0.013003 |
| 2200002D( | 72275 | 2.521631138 | 0.003898 |
| Pcsk6 | 18553 | 2.528633356 | 0.013141 |
| Tmem252 | 226040 | 2.545798477 | 0.000132 |
| Slco2a1 | 24059 | 2.54761947 | 0.006101 |
| Bik | 12124 | 2.559192872 | 0.021814 |
| Lurap1l | 52829 | 2.559382318 | 0.022946 |
| Gja5 | 14613 | 2.56312002 | 0.006562 |
| Lhx6 | 16874 | 2.567215596 | 0.003495 |
| Gnai1 | 14677 | 2.627304138 | 0.025467 |
| Tnfsf18 | 240873 | 2.633475167 | 0.003439 |
| Rbp7 | 63954 | 2.636829607 | 0.006885 |

| | | | |
|---|---|---|---|
| Rbfox3 | 52897 | 2.638586075 | 0.023431 |
| Selp | 20344 | 2.708465338 | 0.006624 |
| Klhl4 | 237010 | 2.731264723 | 0.006562 |
| Depp1 | 213393 | 2.738136446 | 0.000376 |
| Add2 | 11519 | 2.897728121 | 0.043028 |
| Dkk2 | 56811 | 2.914374507 | 0.013141 |
| Tek | 21687 | 2.922081953 | 0.006562 |
| Akr1c14 | 105387 | 2.989983667 | 0.017176 |
| Clec14a | 66864 | 3.094740137 | 0.001378 |
| Car2 | 12349 | 3.337550961 | 7.59E-05 |

**Supplementary Table S7**

Combo vs DB node

| EnrichmentMap::Ge | EnrichmentMap::Name | EnrichmentM | EnrichmentMap::fdr_ |
|---|---|---|---|
| KAZN\|NUMA1\|CNFN | GOBP_KERATINOCYTE_DIFFERENTIATION | 1.7682 | 0.0263 |
| SHH\|NSUN2\|IRF6\|W | GOBP_SKIN_DEVELOPMENT | 1.7636 | 0.0469 |
| NUMA1\|SHH\|ACVR1 | GOBP_SKIN_EPIDERMIS_DEVELOPMENT | 1.753 | 0.0211 |
| EREG\|NOTCH2\|IRF | GOBP_KERATINOCYTE_PROLIFERATION | 1.7458 | 0.0388 |
| FA2H\|KLF4\|ACER1 | GOBP_REGULATION_OF_WATER_LOSS_VIA_SKIN | 1.7324 | 0.0612 |
| SPRY1\|SHH\|SAPCD | GOBP_REGULATION_OF_ANIMAL_ORGAN_MORPHOGENESIS | 1.7304 | 0.0035 |
| SHH\|KRT80\|EPPK1 | GOBP_INTERMEDIATE_FILAMENT_ORGANIZATION | 1.7298 | 0.0337 |
| NOTCH2\|IRF6\|OVO | GOBP_REGULATION_OF_KERATINOCYTE_PROLIFERATION | 1.7297 | 0.0207 |
| KAZN\|CNFN\|KRT80 | GOBP_KERATINIZATION | 1.7201 | 0.0425 |
| FOXF1\|WNT5A\|SAF | GOBP_MORPHOGENESIS_OF_A_POLARIZED_EPITHELIUM | 1.7167 | 0.0409 |
| AIDA\|SHH\|2210016 | GOBP_NEGATIVE_REGULATION_OF_WNT_SIGNALING_PATHW | 1.7119 | 0.0468 |
| SHH\|SVIL\|MYOZ1\|C | GOBP_SKELETAL_MUSCLE_ORGAN_DEVELOPMENT | 1.7053 | 0.0305 |
| SPRY1\|GDF6\|SHH\| | GOBP_KIDNEY_EPITHELIUM_DEVELOPMENT | 1.6992 | 0.0388 |
| SHH\|KRT80\|EPPK1 | GOBP_INTERMEDIATE_FILAMENT_BASED_PROCESS | 1.6955 | 0.0471 |
| SPRY1\|FOXA1\|DAE | GOBP_REGULATION_OF_EPITHELIAL_TO_MESENCHYMAL_TR | 1.695 | 0.0344 |
| WNT5A\|SAPCD2\|EX | GOBP_ESTABLISHMENT_OF_TISSUE_POLARITY | 1.6942 | 0.0522 |
| EMX2\|FOXF1\|LHX1 | GOBP_URETER_DEVELOPMENT | 1.6917 | 0.0764 |
| CCN1\|ACVR2B\|ACV | GOBP_POSITIVE_REGULATION_OF_OSSIFICATION | 1.6891 | 0.0914 |
| CHST1\|ALDH1A7\|B | GOBP_AMINO_SUGAR_METABOLIC_PROCESS | 1.6788 | 0.0615 |
| FOXF1\|MSX1\|SHH\|V | GOBP_MESENCHYMAL_CELL_PROLIFERATION | 1.6768 | 0.0102 |
| SPRY1\|SHH\|HS3ST | GOBP_MESONEPHROS_DEVELOPMENT | 1.6731 | 0.0941 |
| SHH\|NSUN2\|IRF6\|S | GOBP_EPIDERMIS_DEVELOPMENT | 1.6703 | 0.0283 |
| WNT5A\|ZNRF3\|RSF | GOBP_NON_CANONICAL_WNT_SIGNALING_PATHWAY | 1.6688 | 0.0239 |
| SPRY1\|SHH\|LGR4\|V | GOBP_BRANCHING_MORPHOGENESIS_OF_AN_EPITHELIAL_T | 1.6676 | 0.0712 |
| FGFR2\|CTNNB1\|SH | GOBP_EPIDERMIS_MORPHOGENESIS | 1.6669 | 0.0051 |
| SHH\|TNN\|GLI1\|ZNR | GOBP_NEGATIVE_REGULATION_OF_CANONICAL_WNT_SIGNA | 1.6604 | 0.0472 |
| SHH\|TNN\|LGR4\|MIR | GOBP_OSTEOBLAST_DIFFERENTIATION | 1.657 | 0.0039 |
| CPLANE2\|MSX1\|MS | GOBP_ENDOCARDIAL_CUSHION_MORPHOGENESIS | 1.6549 | 0.0285 |
| SHH\|GRHL2\|ACVR2 | GOBP_EMBRYONIC_SKELETAL_SYSTEM_DEVELOPMENT | 1.6542 | 0.0246 |
| SEC24B\|SFRP1\|WN | GOBP_REGULATION_OF_ESTABLISHMENT_OF_PLANAR_POL | 1.6538 | 0.0332 |
| GRHL2\|NIPBL\|PAX9 | GOBP_EMBRYONIC_SKELETAL_SYSTEM_MORPHOGENESIS | 1.6502 | 0.093 |
| NUMA1\|SHH\|ACVR1 | GOBP_MOLTING_CYCLE | 1.6496 | 0.0264 |
| NOTCH2\|LGR4\|CFI | GOBP_GLOMERULUS_DEVELOPMENT | 1.6485 | 0.0976 |
| SPRY1\|GDF6\|AKR1 | GOBP_METANEPHROS_DEVELOPMENT | 1.6477 | 0.0949 |
| SHH\|HS3ST3A1\|LG | GOBP_NEPHRON_DEVELOPMENT | 1.6442 | 0.0541 |
| SPRY1\|WNT16\|FOX | GOBP_EPITHELIAL_TO_MESENCHYMAL_TRANSITION | 1.6434 | 0.0543 |
| CCDC3\|LMO3\|HNR | GOBP_POSITIVE_REGULATION_OF_FAT_CELL_DIFFERENTIA | 1.6427 | 0.034 |
| SHH\|FOXO4\|NEUR | GOBP_MUSCLE_ORGAN_DEVELOPMENT | 1.6411 | 0.0349 |
| ACVR2A\|TNN\|GLI1\| | GOBP_REGULATION_OF_OSTEOBLAST_DIFFERENTIATION | 1.6376 | 0.075 |
| ALOX5\|AKR1C18\|A | GOBP_ICOSANOID_METABOLIC_PROCESS | 1.6339 | 0.0485 |
| SPRY1\|SFRP1\|IL17 | GOBP_NEGATIVE_REGULATION_OF_EPITHELIAL_TO_MESEN | 1.6309 | 0.0443 |
| WNT5A\|GLI1\|SOX9 | GOBP_NOTOCHORD_DEVELOPMENT | 1.6295 | 0.0271 |
| CLIC1\|CCN1\|LRP3\| | GOBP_POSITIVE_REGULATION_OF_OSTEOBLAST_DIFFEREN | 1.6267 | 0.061 |
| CCN1\|ACVR2B\|ACV | GOBP_POSITIVE_REGULATION_OF_BONE_MINERALIZATION | 1.6265 | 0.0414 |
| STFA1\|STFA2\|TGM | GOBP_PEPTIDE_CROSS_LINKING | 1.626 | 0.0657 |
| SEC24B\|SFRP1\|PT | GOBP_ESTABLISHMENT_OF_PLANAR_POLARITY_OF_EMBRY | 1.6254 | 0.0425 |
| MFN2\|CFL2\|FLII\|NE | GOBP_STRIATED_MUSCLE_CELL_DEVELOPMENT | 1.6247 | 0.0057 |
| FLII\|PBRM1\|HNRNP | GOBP_MUSCLE_CELL_DEVELOPMENT | 1.6243 | 0.0542 |
| IRF6\|SULT2B1\|WN | GOBP_EPIDERMAL_CELL_DIFFERENTIATION | 1.6227 | 0.0345 |
| CLEC3B\|TNN\|NIPBL | GOBP_OSSIFICATION | 1.6206 | 0.0346 |
| RPS7\|SHH\|GRHL2\| | GOBP_MORPHOGENESIS_OF_EMBRYONIC_EPITHELIUM | 1.6159 | 0.0471 |
| LOX\|SHH\|NACA\|TN | GOBP_REGULATION_OF_MUSCLE_TISSUE_DEVELOPMENT | 1.6143 | 0.0468 |
| LGR4\|HESX1\|AREG | GOBP_EPITHELIAL_TUBE_MORPHOGENESIS | 1.614 | 0.0446 |
| GDF6\|MTSS1\|LGR4 | GOBP_METANEPHRIC_EPITHELIUM_DEVELOPMENT | 1.6139 | 0.0468 |
| EPHA1\|RAPGEF3\|I | GOBP_POSITIVE_REGULATION_OF_ACTIN_FILAMENT_BUNDL | 1.6136 | 0.0421 |
| SHH\|CDKN1C\|CRK | GOBP_DIGESTIVE_SYSTEM_DEVELOPMENT | 1.6124 | 0.0786 |
| FOXF1\|MSX1\|BTRC | GOBP_MAMMARY_GLAND_EPITHELIUM_DEVELOPMENT | 1.6109 | 0.0442 |
| SHH\|EPPK1\|CFLAR | GOBP_POSITIVE_REGULATION_OF_EPITHELIAL_CELL_PROL | 1.6103 | 0.0272 |
| FGF18\|TMEM204\|N | GOBP_VASCULAR_ENDOTHELIAL_GROWTH_FACTOR_RECE | 1.6096 | 0.0399 |
| TNN\|TNNC1\|MYOZ1 | GOBP_MUSCLE_TISSUE_DEVELOPMENT | 1.609 | 0.0245 |
| BHLHA15\|BHLHE41 | GOBP_REGULATION_OF_MYOTUBE_DIFFERENTIATION | 1.607 | 0.05 |
| SHH\|HS3ST3A1\|LG | GOBP_NEPHRON_EPITHELIUM_DEVELOPMENT | 1.6056 | 0.0087 |
| FOXF1\|WNT5A\|NUI | GOBP_POLARIZED_EPITHELIAL_CELL_DIFFERENTIATION | 1.6047 | 0.0616 |
| SPRY1\|SHH\|LGR4\|V | GOBP_MORPHOGENESIS_OF_A_BRANCHING_STRUCTURE | 1.6039 | 0.0367 |
| SEC24B\|SFRP1\|PT | GOBP_ESTABLISHMENT_OF_PLANAR_POLARITY_INVOLVED_ | 1.6039 | 0.0731 |
| FGF18\|TMEM204\|F | GOBP_REGULATION_OF_VASCULAR_ENDOTHELIAL_GROWT | 1.6031 | 0.0522 |
| RHOA\|FOXF1\|CDC | GOBP_ESTABLISHMENT_OF_EPITHELIAL_CELL_APICAL_BAS | 1.6026 | 0.0539 |
| NTRK1\|SHH\|EPHA1 | GOBP_POSITIVE_REGULATION_OF_VASCULATURE_DEVELO | 1.6024 | 0.0038 |
| SHH\|FLII\|WNT1\|HNF | GOBP_STRIATED_MUSCLE_CELL_DIFFERENTIATION | 1.6021 | 0.045 |
| ARNTL\|SFRP1\|SIRT | GOBP_NEGATIVE_REGULATION_OF_INTRACELLULAR_STER | 1.6017 | 0.0341 |
| SHH\|ADAMTS16\|GA | GOBP_RENAL_TUBULE_MORPHOGENESIS | 1.6004 | 0.0617 |
| FGFR2\|BTRC\|MSX | GOBP_MAMMARY_GLAND_DUCT_MORPHOGENESIS | 1.5998 | 0.0443 |
| TEAD1\|TEAD2\|SP1 | GOBP_RESPONSE_TO_RETINOIC_ACID | 1.599 | 0.0319 |
| ARNTL\|SHH\|CITED | GOBP_SKELETAL_MUSCLE_CELL_DIFFERENTIATION | 1.5986 | 0.0406 |
| SHH\|ADAMTS16\|GA | GOBP_MESONEPHRIC_TUBULE_MORPHOGENESIS | 1.5974 | 0.0205 |
| CTNNB1\|IGF1\|KLHL | GOBP_REGULATION_OF_MYOBLAST_PROLIFERATION | 1.5971 | 0.0521 |
| CTNNB1\|MSX1\|SHH | GOBP_FORELIMB_MORPHOGENESIS | 1.5969 | 0.0507 |
| SHH\|HS3ST3A1\|LG | GOBP_RENAL_TUBULE_DEVELOPMENT | 1.5968 | 0.0357 |
| SERPINB5\|EXOC8\| | GOBP_EXTERNAL_ENCAPSULATING_STRUCTURE_ORGANIZA | 1.5948 | 0.054 |
| KITL\|PYCARD\|IL12 | GOBP_POSITIVE_REGULATION_OF_LEUKOCYTE_MIGRATION | 1.5944 | 0.0865 |
| EDNRA\|RET\|GATA | GOBP_REGULATION_OF_KIDNEY_DEVELOPMENT | 1.5942 | 0.0469 |
| TBX5\|TGFB2\|ABI3E | GOBP_REGULATION_OF_CARDIOCYTE_DIFFERENTIATION | 1.5932 | 0.0542 |
| SMAD2\|DLX2\|GRHL | GOBP_EMBRYONIC_CRANIAL_SKELETON_MORPHOGENESIS | 1.5927 | 0.0445 |
| SHH\|SIRT1\|MESP1\| | GOBP_POSITIVE_REGULATION_OF_STRIATED_MUSCLE_CEL | 1.5926 | 0.0346 |
| MYOZ1\|MYPN\|CFL | GOBP_MUSCLE_CELL_DIFFERENTIATION | 1.5924 | 0.0947 |
| RPS7\|SPRY1\|SHH\|V | GOBP_MESENCHYMAL_CELL_DIFFERENTIATION | 1.5908 | 0.047 |
| NEUROG1\|CHRM2\| | GOBP_SMOOTH_MUSCLE_CONTRACTION | 1.5895 | 0.0599 |
| SHH\|LGMN\|CCBE1\| | GOBP_REGULATION_OF_ENDOTHELIAL_CELL_MIGRATION | 1.5893 | 0.015 |
| FOXF1\|SHH\|WNT5A | GOBP_ESTABLISHMENT_OF_EPITHELIAL_CELL_POLARITY | 1.5887 | 0.0195 |
| FERMT1\|EPPK1\|MT | GOBP_KERATINOCYTE_MIGRATION | 1.5881 | 0.0386 |
| WNT5A\|NEUROG1\| | GOBP_COCHLEA_MORPHOGENESIS | 1.5877 | 0.0806 |
| CTNNB1\|RET\|EGR1 | GOBP_METANEPHRIC_NEPHRON_DEVELOPMENT | 1.5873 | 0.0425 |
| FGFR2\|CCN1\|SFRI | GOBP_REGULATION_OF_OSTEOBLAST_PROLIFERATION | 1.587 | 0.0755 |
| CYP2C65\|CYP2S1\| | GOBP_ARACHIDONIC_ACID_METABOLIC_PROCESS | 1.5863 | 0.0286 |
| SHH\|ZFP418\|BMP4 | GOBP_REGULATION_OF_STRIATED_MUSCLE_CELL_DIFFERE | 1.5857 | 0.0286 |
| SHH\|FOXO4\|CCBE1 | GOBP_REGULATION_OF_VASCULATURE_DEVELOPMENT | 1.5855 | 0.0269 |
| FOXF1\|CPLANE2\|S | GOBP_MESENCHYME_MORPHOGENESIS | 1.5853 | 0.0104 |
| SPRY1\|CTNNB1\|RU | GOBP_REGULATION_OF_FIBROBLAST_GROWTH_FACTOR_R | 1.5828 | 0.015 |
| SHH\|FOXA1\|ISL1\|H | GOBP_NEURON_FATE_SPECIFICATION | 1.5823 | 0.0089 |
| IRF6\|EPPK1\|SGPP2 | GOBP_REGULATION_OF_EPITHELIAL_CELL_PROLIFERATION | 1.5817 | 0.0487 |
| SHH\|NOTCH2\|GATA | GOBP_CELL_DIFFERENTIATION_INVOLVED_IN_KIDNEY_DEVE | 1.5817 | 0.0366 |
| FGFR2\|LRRC17\|CC | GOBP_OSTEOBLAST_PROLIFERATION | 1.5815 | 0.003 |
| CPLANE2\|SMAD2\|D | GOBP_CRANIAL_SKELETAL_SYSTEM_DEVELOPMENT | 1.5809 | 0.071 |
| EDNRA\|RET\|ARX\|K | GOBP_ENTERIC_NERVOUS_SYSTEM_DEVELOPMENT | 1.5801 | 0.0688 |
| ECSCR\|FOXO4\|ALC | GOBP_NEGATIVE_REGULATION_OF_VASCULATURE_DEVELO | 1.5796 | 0.0098 |
| RET\|TEAD1\|OSR1\| | GOBP_CELLULAR_RESPONSE_TO_RETINOIC_ACID | 1.5788 | 0.0669 |
| MSX1\|MSX2\|SNAI1\| | GOBP_ENDOCARDIAL_CUSHION_FORMATION | 1.5782 | 0.0042 |
| SPRY1\|KIF16B\|FGF | GOBP_RESPONSE_TO_FIBROBLAST_GROWTH_FACTOR | 1.5769 | 0.0481 |
| SORBS2\|PDCD4\|PI | GOBP_CARDIOCYTE_DIFFERENTIATION | 1.5747 | 0.0265 |
| SPRY1\|KIF16B\|RUN | GOBP_FIBROBLAST_GROWTH_FACTOR_RECEPTOR_SIGNAL | 1.574 | 0.0417 |
| SFRP1\|SIRT1\|NOD | GOBP_REGULATION_OF_ANDROGEN_RECEPTOR_SIGNALING | 1.5732 | 0.0345 |
| SHH\|HS3ST3A1\|LG | GOBP_KIDNEY_MORPHOGENESIS | 1.5731 | 0.0986 |
| FGFR2\|SERPINB5\| | GOBP_PROSTATE_GLAND_MORPHOGENESIS | 1.5722 | 0.0305 |
| SERPINC1\|SHH\|SY | GOBP_WOUND_HEALING | 1.5716 | 0.0258 |
| CTNNB1\|MEIS2\|AT | GOBP_MYOBLAST_PROLIFERATION | 1.5715 | 0.0349 |
| CTNNB1\|MSX1\|SHH | GOBP_EMBRYONIC_HINDLIMB_MORPHOGENESIS | 1.5712 | 0.0919 |
| NUMA1\|SHH\|GRHL2 | GOBP_LUNG_EPITHELIUM_DEVELOPMENT | 1.5703 | 0.0616 |
| CAPN1\|PDCD4\|IL18 | GOBP_REGULATION_OF_NIK_NF_KAPPAB_SIGNALING | 1.5701 | 0.0616 |
| GDF6\|GLG1\|TRIP1 | GOBP_CARTILAGE_DEVELOPMENT | 1.5682 | 0.0391 |
| CCN1\|TMEFF2\|MSX | GOBP_MORPHOGENESIS_OF_AN_EPITHELIAL_SHEET | 1.568 | 0.0394 |
| TIMP1\|CD44\|ALOX | GOBP_INFLAMMATORY_RESPONSE_TO_WOUNDING | 1.568 | 0.0561 |
| FOXF1\|SPRY1\|SHH | GOBP_LUNG_MORPHOGENESIS | 1.5679 | 0.0303 |
| FOXF1\|WNT5A\|MY | GOBP_ESTABLISHMENT_OR_MAINTENANCE_OF_BIPOLAR_C | 1.5675 | 0.0183 |
| LYVE1\|CAPN1\|GCN | GOBP_REGULATION_OF_CELLULAR_EXTRAVASATION | 1.5673 | 0.0246 |
| EDNRA\|CTNNB1\|NO | GOBP_EPITHELIAL_CELL_DIFFERENTIATION_INVOLVED_IN_K | 1.5673 | 0.0388 |
| RETREG3\|ADAM15 | GOBP_COLLAGEN_CATABOLIC_PROCESS | 1.5664 | 0.0262 |
| DPP4\|IL12A\|LGMN\| | GOBP_REGULATION_OF_LEUKOCYTE_CHEMOTAXIS | 1.5663 | 0.0784 |
| SPRY1\|STUB1\|CDK | GOBP_REGULATION_OF_CELLULAR_RESPONSE_TO_GROW | 1.5655 | 0.0855 |
| RETREG3\|ADAM15 | GOBP_COLLAGEN_METABOLIC_PROCESS | 1.5652 | 0.0246 |
| SERPINC1\|ALOX5\|E | GOBP_REGULATION_OF_RESPONSE_TO_WOUNDING | 1.565 | 0.0424 |
| FGFR2\|FOXF1\|SPF | GOBP_EPITHELIAL_TUBE_BRANCHING_INVOLVED_IN_LUNG_ | 1.5645 | 0.0995 |
| CTNNB1\|MSX1\|SHH | GOBP_HINDLIMB_MORPHOGENESIS | 1.5643 | 0.0443 |
| NACA\|P2RX2\|WNT1 | GOBP_MYOTUBE_CELL_DEVELOPMENT | 1.5633 | 0.0749 |
| TNNC1\|GRCC10\|PA | GOBP_MULTICELLULAR_ORGANISMAL_MOVEMENT | 1.5625 | 0.098 |
| LOX\|CLEC3B\|ACVR | GOBP_BONE_MINERALIZATION | 1.5614 | 0.0119 |
| OVGP1\|MGAT1\|AL | GOBP_AMINO_SUGAR_CATABOLIC_PROCESS | 1.5602 | 0.0204 |
| UBASH3B\|ALOX5\|S | GOBP_NEGATIVE_REGULATION_OF_RESPONSE_TO_WOUND | 1.5602 | 0.0888 |
| SHH\|PLD3\|WNT1\|C | GOBP_MYOTUBE_DIFFERENTIATION | 1.56 | 0.0808 |
| REG3G\|NOTCH2\|L | GOBP_POSITIVE_REGULATION_OF_KERATINOCYTE_PROLIF | 1.5598 | 0.0348 |
| KITL\|SART3\|SHH\|IR | GOBP_STEM_CELL_PROLIFERATION | 1.5596 | 0.0302 |
| JAGN1\|IRAK4\|CXCL | GOBP_MYELOID_LEUKOCYTE_MIGRATION | 1.5595 | 0.0371 |
| SLAMF9\|CXCL16\|C | GOBP_LEUKOCYTE_CHEMOTAXIS | 1.5589 | 0.0472 |
| SPRY1\|SPRY4\|GAT | GOBP_NEGATIVE_REGULATION_OF_FIBROBLAST_GROWTH_ | 1.5574 | 0.0485 |
| CAPN1\|GCNT1\|KLF | GOBP_REGULATION_OF_LEUKOCYTE_ADHESION_TO_VASC | 1.557 | 0.0482 |
| SHH\|MSX2\|RUNX2\|T | GOBP_OSTEOBLAST_DEVELOPMENT | 1.5561 | 0.0434 |
| NEUROG1\|TNNC1\| | GOBP_MUSCLE_CONTRACTION | 1.5551 | 0.0347 |
| CTNNB1\|MSX1\|SHH | GOBP_EMBRYONIC_FORELIMB_MORPHOGENESIS | 1.5546 | 0.0454 |
| EID2B\|CMTM5\|BMF | GOBP_NEGATIVE_REGULATION_OF_MYOBLAST_DIFFERENT | 1.5541 | 0.0047 |
| SHH\|CCBE1\|DAB2I | GOBP_ENDOTHELIAL_CELL_MIGRATION | 1.5541 | 0.0442 |
| KITL\|SHH\|IRF6\|CDK | GOBP_REGULATION_OF_STEM_CELL_PROLIFERATION | 1.5538 | 0.0569 |
| CHMP7\|SYT7\|ARL8 | GOBP_PLASMA_MEMBRANE_REPAIR | 1.5531 | 0.0469 |
| CHST1\|HS3ST1\|HS | GOBP_AMINOGLYCAN_BIOSYNTHETIC_PROCESS | 1.5531 | 0.0978 |
| RPS7\|SPRY1\|SHH\|V | GOBP_MESENCHYME_DEVELOPMENT | 1.5529 | 0.0334 |
| EDNRA\|SPRY1\|CTN | GOBP_POSITIVE_REGULATION_OF_ANIMAL_ORGAN_MORPH | 1.5525 | 0.0288 |
| GDF6\|NUMA1\|LEFT | GOBP_RESPONSE_TO_BMP | 1.5512 | 0.0266 |
| CTNNB1\|SHH\|NACA | GOBP_REGULATION_OF_SKELETAL_MUSCLE_TISSUE_DEVE | 1.5511 | 0.0472 |
| FOXO4\|DRD4\|NF1\| | GOBP_SMOOTH_MUSCLE_CELL_MIGRATION | 1.5511 | 0.0695 |
| IL12A\|LGMN\|PTK2E | GOBP_POSITIVE_REGULATION_OF_LEUKOCYTE_CHEMOTAX | 1.5507 | 0.0643 |
| SHH\|DLX2\|EPOP\|F | GOBP_NEURON_FATE_COMMITMENT | 1.5501 | 0.0404 |
| SIRT1\|ETS1\|NF1\|A | GOBP_REGULATION_OF_BLOOD_VESSEL_ENDOTHELIAL_CE | 1.5501 | 0.0272 |
| SHH\|SIRT1\|MESP1\| | GOBP_POSITIVE_REGULATION_OF_MUSCLE_CELL_DIFFERE | 1.5493 | 0.0301 |
| SMAD2\|SNAI1\|ISL1 | GOBP_POSITIVE_REGULATION_OF_EPITHELIAL_TO_MESENC | 1.5483 | 0.0335 |
| PLCG2\|APP\|CAMP\| | GOBP_RESPONSE_TO_YEAST | 1.5483 | 0.0241 |
| GDF6\|AKR1B3\|LGF | GOBP_UROGENITAL_SYSTEM_DEVELOPMENT | 1.5477 | 0.0348 |
| CD5L\|SERPING1\|C | GOBP_REGULATION_OF_COMPLEMENT_ACTIVATION | 1.5473 | 0.0169 |
| SHH\|WNT16\|PBRM | GOBP_HEART_MORPHOGENESIS | 1.5454 | 0.0531 |
| MSX1\|SHH\|ACVR2E | GOBP_ODONTOGENESIS_OF_DENTIN_CONTAINING_TOOTH | 1.5449 | 0.013 |
| SHH\|GATA3\|WNT5A | GOBP_REGULATION_OF_MORPHOGENESIS_OF_AN_EPITHEL | 1.5441 | 0.0338 |
| CAPN1\|GCNT1\|SEL | GOBP_LEUKOCYTE_ADHESION_TO_VASCULAR_ENDOTHELIA | 1.5439 | 0.0541 |
| PTGIS\|CYP2S1\|SIR | GOBP_PROSTANOID_METABOLIC_PROCESS | 1.5439 | 0.0642 |
| TNNC1\|GRCC10\|M | GOBP_MUSCLE_SYSTEM_PROCESS | 1.5438 | 0.0302 |
| FGFR2\|FOXF1\|CTN | GOBP_REGULATION_OF_MESENCHYMAL_CELL_PROLIFERAT | 1.5435 | 0.0422 |
| IRF6\|CDKN1C\|IL12 | GOBP_NEGATIVE_REGULATION_OF_EPITHELIAL_CELL_PRO | 1.5434 | 0.0603 |
| ALOX12\|ZFP703\|F | GOBP_NEGATIVE_REGULATION_OF_HOMOTYPIC_CELL_CEL | 1.5432 | 0.0128 |
| FOXF1\|SMAD2\|SHH | GOBP_DIGESTIVE_TRACT_MORPHOGENESIS | 1.5431 | 0.0428 |
| FOXF1\|LFNG\|SHH\| | GOBP_SOMITE_DEVELOPMENT | 1.543 | 0.0302 |
| SHH\|CCBE1\|EPPK1 | GOBP_REGULATION_OF_EPITHELIAL_CELL_MIGRATION | 1.543 | 0.05 |
| SERPING1\|C1QBP | GOBP_REGULATION_OF_HUMORAL_IMMUNE_RESPONSE | 1.5427 | 0.095 |
| SHH\|ACVR2A\|ITGB | GOBP_ODONTOGENESIS | 1.5425 | 0.0397 |
| HTR2B\|ACVR2A\|C | GOBP_REGULATION_OF_MONOOXYGENASE_ACTIVITY | 1.5417 | 0.0424 |
| MSX2\|NACA\|TGFB2 | GOBP_TRABECULA_MORPHOGENESIS | 1.5416 | 0.0612 |
| CAPN1\|NLRP12\|RE | GOBP_NEGATIVE_REGULATION_OF_NIK_NF_KAPPAB_SIGNA | 1.5411 | 0.0444 |
| SHH\|NEUROG1\|RB | GOBP_SEGMENTATION | 1.5401 | 0.027 |
| ETS1\|NF1\|PTK2B\|F | GOBP_BLOOD_VESSEL_ENDOTHELIAL_CELL_MIGRATION | 1.5398 | 0.0266 |
| FOXF1\|SLC7A11\|F | GOBP_LUNG_ALVEOLUS_DEVELOPMENT | 1.5397 | 0.0369 |
| EDNRA\|EGR1\|NOT | GOBP_RENAL_SYSTEM_VASCULATURE_DEVELOPMENT | 1.5393 | 0.0394 |
| FOXF1\|LFNG\|DLL3 | GOBP_SOMITOGENESIS | 1.5391 | 0.0288 |
| SERPINC1\|ALOX5\|E | GOBP_REGULATION_OF_WOUND_HEALING | 1.5391 | 0.0939 |
| RPS7\|SHH\|GRHL2\| | GOBP_TUBE_FORMATION | 1.5387 | 0.0788 |
| SLAMF9\|JAGN1\|CX | GOBP_LEUKOCYTE_MIGRATION | 1.5384 | 0.015 |
| SLAMF9\|SHH\|CXCL | GOBP_CELL_CHEMOTAXIS | 1.5376 | 0.0346 |
| CTNNB1\|GATA3\|AC | GOBP_POSITIVE_REGULATION_OF_MORPHOGENESIS_OF_A | 1.5366 | 0.0678 |
| GDF6\|GPS2\|NIPBL | GOBP_FAT_CELL_DIFFERENTIATION | 1.5364 | 0.0947 |
| AKR1B3\|ABCA12\|A | GOBP_WATER_HOMEOSTASIS | 1.5357 | 0.0609 |
| SHH\|LGMN\|CCBE1\| | GOBP_POSITIVE_REGULATION_OF_ENDOTHELIAL_CELL_MIG | 1.5354 | 0.0208 |
| MYSM1\|NUMA1\|DK | GOBP_REGULATION_OF_HAIR_FOLLICLE_DEVELOPMENT | 1.5348 | 0.0344 |
| SHH\|SLC7A11\|IRF6 | GOBP_APPENDAGE_DEVELOPMENT | 1.5347 | 0.0139 |
| GDF6\|GLG1\|TRIP1 | GOBP_CHONDROCYTE_DIFFERENTIATION | 1.5345 | 0.0257 |
| RET\|GATA3\|XYLT1 | GOBP_ANIMAL_ORGAN_MATURATION | 1.534 | 0.054 |
| CAPN1\|BTRC\|MSX2 | GOBP_MAMMARY_GLAND_MORPHOGENESIS | 1.5337 | 0.0286 |
| CHRM2\|TNNC1\|GR | GOBP_REGULATION_OF_MUSCLE_CONTRACTION | 1.5335 | 0.0902 |
| IL12A\|IL12B\|ALOX5 | GOBP_ENDOTHELIAL_CELL_PROLIFERATION | 1.5335 | 0.0486 |
| SHH\|IL12A\|LGMN\|P | GOBP_POSITIVE_REGULATION_OF_CHEMOTAXIS | 1.533 | 0.0032 |
| GRHL2\|SNAI1\|MYO | GOBP_APICAL_JUNCTION_ASSEMBLY | 1.5326 | 0.0938 |
| PTK2B\|CCL27A\|TM | GOBP_REGULATION_OF_LEUKOCYTE_MIGRATION | 1.5314 | 0.0199 |
| SHH\|CCBE1\|EPPK1 | GOBP_TISSUE_MIGRATION | 1.5307 | 0.054 |
| NTRK1\|WNT16\|FYN | GOBP_INOSITOL_LIPID_MEDIATED_SIGNALING | 1.5304 | 0.0473 |
| FOXF1\|CPLANE2\|M | GOBP_ENDOCARDIAL_CUSHION_DEVELOPMENT | 1.5298 | 0.0457 |
| CEND1\|KIFAP3\|WN | GOBP_NEGATIVE_REGULATION_OF_NEURAL_PRECURSOR_ | 1.5284 | 0.0353 |
| PTGIS\|SIRT1\|ALOX | GOBP_ICOSANOID_BIOSYNTHETIC_PROCESS | 1.5281 | 0.048 |
| CHST1\|HS3ST1\|HS | GOBP_AMINOGLYCAN_METABOLIC_PROCESS | 1.528 | 0.0756 |
| SHH\|TNN\|JRK\|CCAF | GOBP_REGULATION_OF_CANONICAL_WNT_SIGNALING_PATH | 1.5275 | 0.0864 |
| CTNNB1\|DLX2\|GLI1 | GOBP_PROXIMAL_DISTAL_PATTERN_FORMATION | 1.5272 | 0.0416 |
| GDF6\|CCN1\|RUNX2 | GOBP_REGULATION_OF_CARTILAGE_DEVELOPMENT | 1.527 | 0.026 |
| PYCARD\|PLCG2\|SI | GOBP_MACROPHAGE_CYTOKINE_PRODUCTION | 1.5269 | 0.053 |
| FOXO4\|DRD4\|NF1\| | GOBP_MUSCLE_CELL_MIGRATION | 1.5258 | 0.0286 |
| FOS\|RAF1\|JUN\|REL | GOBP_RESPONSE_TO_MUSCLE_STRETCH | 1.5257 | 0.0979 |
| CCL26\|CCL24\|LGA | GOBP_EOSINOPHIL_MIGRATION | 1.5255 | 0.0276 |
| CAPN1\|GCNT1\|SEL | GOBP_CELLULAR_EXTRAVASATION | 1.5252 | 0.0215 |
| RET\|GATA1\|TGFB2 | GOBP_REGULATION_OF_EXTRINSIC_APOPTOTIC_SIGNALING | 1.5251 | 0.0338 |
| SHH\|NEUROG1\|FEZ | GOBP_ANTERIOR_POSTERIOR_PATTERN_SPECIFICATION | 1.525 | 0.0749 |
| FLII\|LURAP1\|MYOZ | GOBP_ACTOMYOSIN_STRUCTURE_ORGANIZATION | 1.525 | 0.0243 |
| CAPN1\|SERPINB5\| | GOBP_GLAND_MORPHOGENESIS | 1.5243 | 0.035 |
| MTPN\|NEUROG1\|T | GOBP_REGULATION_OF_MUSCLE_SYSTEM_PROCESS | 1.523 | 0.0646 |
| ALOX5\|AKR1C18\|A | GOBP_UNSATURATED_FATTY_ACID_METABOLIC_PROCESS | 1.5228 | 0.0938 |
| EREG\|FOXF1\|SHH\| | GOBP_SMOOTH_MUSCLE_CELL_DIFFERENTIATION | 1.5224 | 0.0059 |
| SHH\|IRF6\|GABRB3 | GOBP_ROOF_OF_MOUTH_DEVELOPMENT | 1.5214 | 0.0422 |
| SMARCC1\|XKR8\|A | GOBP_REGULATION_OF_MYOBLAST_DIFFERENTIATION | 1.5213 | 0.0265 |
| ACVR2A\|GATA1\|NI | GOBP_REGULATION_OF_OSSIFICATION | 1.5211 | 0.0308 |
| C1QTNF12\|LIN28A | GOBP_POSITIVE_REGULATION_OF_PROTEIN_KINASE_B_SIG | 1.5207 | 0.0363 |
| SORBS2\|HNRNPU\| | GOBP_CARDIAC_MUSCLE_CELL_DIFFERENTIATION | 1.5203 | 0.0417 |
| EPHA1\|RHPN1\|GPF | GOBP_REGULATION_OF_ACTIN_FILAMENT_BUNDLE_ASSEM | 1.5186 | 0.0429 |
| CAPN1\|PDCD4\|IL18 | GOBP_NIK_NF_KAPPAB_SIGNALING | 1.5184 | 0.0542 |
| GLG1\|NIPBL\|NEUR | GOBP_SKELETAL_SYSTEM_MORPHOGENESIS | 1.5177 | 0.0102 |
| TNN\|MIR137\|MIR20 | GOBP_NEGATIVE_REGULATION_OF_OSTEOBLAST_DIFFERE | 1.5172 | 0.0366 |
| RPS7\|SHH\|MIR125 | GOBP_STEM_CELL_DIFFERENTIATION | 1.5171 | 0.042 |
| NUMA1\|FSTL3\|WN | GOBP_REGULATION_OF_BMP_SIGNALING_PATHWAY | 1.515 | 0.0395 |
| DPP4\|JAGN1\|CKLF | GOBP_GRANULOCYTE_MIGRATION | 1.5149 | 0.0388 |
| SHH\|SMARCC1\|AC | GOBP_MYOBLAST_DIFFERENTIATION | 1.5148 | 0.0414 |
| CHMP7\|2610002M | GOBP_MULTIVESICULAR_BODY_ORGANIZATION | 1.5133 | 0.0919 |
| CCN1\|ACVR2B\|ACV | GOBP_POSITIVE_REGULATION_OF_BIOMINERALIZATION | 1.5133 | 0.0387 |
| SHH\|WNT16\|TNN\|J | GOBP_CANONICAL_WNT_SIGNALING_PATHWAY | 1.5125 | 0.0389 |
| GRHL2\|SMARCD3\| | GOBP_CARDIAC_CHAMBER_MORPHOGENESIS | 1.5123 | 0.006 |
| CCBE1\|EPPK1\|ITG | GOBP_AMEBOIDAL_TYPE_CELL_MIGRATION | 1.5109 | 0.047 |
| AIDA\|SHH\|FOXA1\|G | GOBP_DORSAL_VENTRAL_PATTERN_FORMATION | 1.5108 | 0.0506 |
| DPP4\|JAGN1\|CKLF | GOBP_NEUTROPHIL_MIGRATION | 1.5105 | 0.099 |
| SHH\|FEZF2\|PTK2B | GOBP_REGULATION_OF_CHEMOTAXIS | 1.5102 | 0.0279 |
| SHH\|FOXO4\|PDCD | GOBP_REGULATION_OF_MUSCLE_CELL_DIFFERENTIATION | 1.5096 | 0.0541 |
| TNN\|FLRT2\|NIPBL\| | GOBP_REGULATION_OF_NEURON_MIGRATION | 1.5093 | 0.0127 |
| SHH\|EYA2\|TNNC1\|H | GOBP_STRIATED_MUSCLE_TISSUE_DEVELOPMENT | 1.509 | 0.0155 |
| MSX1\|GRHL2\|WNT | GOBP_FACE_DEVELOPMENT | 1.5087 | 0.0439 |
| UBASH3B\|ALOX5\|S | GOBP_NEGATIVE_REGULATION_OF_WOUND_HEALING | 1.5085 | 0.0435 |
| SHH\|TNN\|ATP1B2\|E | GOBP_NEGATIVE_REGULATION_OF_LOCOMOTION | 1.5083 | 0.0529 |
| CDKN1C\|GATA3\|W | GOBP_UTERUS_DEVELOPMENT | 1.5082 | 0.0488 |
| MACIR\|EPPK1\|SP1 | GOBP_NEGATIVE_REGULATION_OF_EPITHELIAL_CELL_MIGR | 1.508 | 0.0542 |
| SND1\|LIN28A\|ETS1 | GOBP_MIRNA_METABOLIC_PROCESS | 1.5078 | 0.0286 |
| PYCARD\|TNFSF10 | GOBP_POSITIVE_REGULATION_OF_EXTRINSIC_APOPTOTIC_ | 1.5077 | 0.0266 |
| MFN2\|CFL2\|NEB\|M | GOBP_SARCOMERE_ORGANIZATION | 1.5075 | 0.0569 |
| SHH\|EIF2B4\|NF1\|E | GOBP_OLIGODENDROCYTE_DIFFERENTIATION | 1.5074 | 0.0244 |
| DPP4\|CKLF\|CXCL1 | GOBP_GRANULOCYTE_CHEMOTAXIS | 1.5069 | 0.0832 |
| SHH\|LGMN\|CCBE1\| | GOBP_POSITIVE_REGULATION_OF_EPITHELIAL_CELL_MIGRA | 1.5066 | 0.0897 |
| MSX1\|TGFB2\|NIPB | GOBP_BODY_MORPHOGENESIS | 1.5065 | 0.0274 |
| APOE\|DPP4\|CRHR | GOBP_LOCOMOTORY_EXPLORATION_BEHAVIOR | 1.5042 | 0.0388 |
| SHH\|IL12A\|TNN\|IL1 | GOBP_SPROUTING_ANGIOGENESIS | 1.5039 | 0.0941 |
| APOA5\|LONP2\|THF | GOBP_REGULATION_OF_LIPID_CATABOLIC_PROCESS | 1.5031 | 0.0903 |
| SPRY1\|NUMA1\|GR | GOBP_REGULATION_OF_EPITHELIAL_CELL_DIFFERENTIATIO | 1.5029 | 0.0425 |
| NTRK1\|GATA3\|NF1 | GOBP_AUTONOMIC_NERVOUS_SYSTEM_DEVELOPMENT | 1.4997 | 0.006 |
| GJA5\|CCN1\|NOTCH | GOBP_HEART_VALVE_MORPHOGENESIS | 1.4994 | 0.0443 |

Combo vs Veh nodes

| EnrichmentM | EnrichmentMap::Name | EnrichmentMap::NES (D | EnrichmentMap::fdr_qvalue (Da |
|---|---|---|---|
| FUT7\|CCL21 | GOBP_REGULATION_OF_LEUKOCYTE_TETH | 1.753 | 0.0273 |
| CCL21B\|SPN | GOBP_LEUKOCYTE_TETHERING_OR_ROLLIN | 1.753 | 0.0547 |
| SOX17\|SIX1\| | GOBP_URETER_DEVELOPMENT | 1.7135 | 0.0737 |
| PIN1\|PIN1RT | GOBP_REGULATION_OF_MUSCLE_HYPERTR | 1.7048 | 0.0741 |
| AKAP5\|ADR | GOBP_REGULATION_OF_CALCIUM_ION_IMPO | 1.6966 | 0.0771 |
| IL18R1\|ABH | GOBP_NEGATIVE_REGULATION_OF_COLD_I | 1.6908 | 0.0766 |
| CASP6\|PPA | GOBP_HEPATOCYTE_APOPTOTIC_PROCESS | 1.6891 | 0.0696 |
| CCL21B\|ITG | GOBP_LEUKOCYTE_ADHESION_TO_VASCUL | 1.6869 | 0.0652 |
| ADORA2B\|N | GOBP_SMOOTH_MUSCLE_CONTRACTION | 1.6817 | 0.0667 |
| CLEC4B1\|C | GOBP_ANTIFUNGAL_INNATE_IMMUNE_RESPO | 1.6806 | 0.0623 |
| AKAP5\|ADR | GOBP_POSITIVE_REGULATION_OF_CALCIUM | 1.6743 | 0.0682 |
| SLC9A3R2\| | GOBP_INOSITOL_LIPID_MEDIATED_SIGNALIN | 1.6732 | 0.0644 |
| PIN1\|PIN1RT | GOBP_REGULATION_OF_MUSCLE_ADAPTAT | 1.6728 | 0.0601 |
| FUT7\|ST3G | GOBP_POSITIVE_REGULATION_OF_LEUKOC | 1.6667 | 0.0663 |
| SORL1\|S10 | GOBP_SMOOTH_MUSCLE_CELL_MIGRATION | 1.6632 | 0.0676 |
| RIPOR2\|CCL | GOBP_REGULATION_OF_CELLULAR_EXTRAV | 1.6613 | 0.0664 |
| CTBP2\|CCD | GOBP_FAT_CELL_DIFFERENTIATION | 1.6549 | 0.0732 |
| TRIM27\|TRI | GOBP_NEGATIVE_REGULATION_OF_TUMOR_ | 1.6495 | 0.0805 |
| CCL21B\|ITG | GOBP_CELLULAR_EXTRAVASATION | 1.6488 | 0.0772 |
| FOXP1\|PAK | GOBP_NEGATIVE_REGULATION_OF_MUSCLE | 1.647 | 0.077 |
| ADORA2B\|N | GOBP_REGULATION_OF_SMOOTH_MUSCLE_ | 1.6461 | 0.0753 |
| FGF7\|SIX1\|F | GOBP_MESENCHYMAL_CELL_PROLIFERATIO | 1.6392 | 0.0861 |
| IL18R1\|SOR | GOBP_ADAPTIVE_THERMOGENESIS | 1.6359 | 0.0896 |
| MEF2C\|PAK | GOBP_VASCULAR_ASSOCIATED_SMOOTH_M | 1.6325 | 0.0891 |
| GPR27\|OPR | GOBP_PHOSPHOLIPASE_C_ACTIVATING_G_F | 1.627 | 0.0884 |
| TWIST1\|PPA | GOBP_POSITIVE_REGULATION_OF_FATTY_A | 1.6247 | 0.0901 |
| FZD5\|HOXA | GOBP_ANTERIOR_POSTERIOR_PATTERN_SP | 1.6243 | 0.0876 |
| AVPR2\|MLL | GOBP_RENAL_SODIUM_EXCRETION | 1.6224 | 0.0888 |
| IL18R1\|PTG | GOBP_TEMPERATURE_HOMEOSTASIS | 1.6215 | 0.0852 |
| FGF10\|IL4\|P | GOBP_FAT_CELL_PROLIFERATION | 1.6177 | 0.0907 |
| TLE5\|DACT | GOBP_NEGATIVE_REGULATION_OF_WNT_SI | 1.6165 | 0.0905 |
| PIN1\|PIN1RT | GOBP_MUSCLE_HYPERTROPHY | 1.6159 | 0.0889 |
| SHISA7\|GAB | GOBP_GAMMA_AMINOBUTYRIC_ACID_SIGNA | 1.6147 | 0.0891 |
| ENTPD1\|BL | GOBP_PLATELET_ACTIVATION | 1.6134 | 0.0891 |
| MYF6\|SFRP | GOBP_SOMITOGENESIS | 1.6121 | 0.0894 |
| PRDM16\|SIX | GOBP_BROWN_FAT_CELL_DIFFERENTIATIO | 1.6074 | 0.0863 |
| TLE5\|DACT | GOBP_NEGATIVE_REGULATION_OF_CANONI | 1.6065 | 0.0845 |
| MBL1\|HCK\|P | GOBP_REGULATION_OF_PHAGOCYTOSIS | 1.6065 | 0.0862 |
| CNR2\|CNR1 | GOBP_SYNAPTIC_TRANSMISSION_GABAERG | 1.5998 | 0.0905 |
| PIN1\|PIN1RT | GOBP_MUSCLE_ADAPTATION | 1.5958 | 0.0933 |
| ADCY4\|ADC | GOBP_ACTIVATION_OF_ADENYLATE_CYCLA | 1.5903 | 0.0943 |
| SIX1\|SMO\|SI | GOBP_REGULATION_OF_KIDNEY_DEVELOPM | 1.5881 | 0.0969 |
| FZD9\|MEF2 | GOBP_POSITIVE_REGULATION_OF_BONE_M | 1.5868 | 0.0966 |
| GPC3\|SMAD | GOBP_MESONEPHROS_DEVELOPMENT | 1.5848 | 0.0998 |
| MEF2C\|EDN | GOBP_GLOMERULUS_DEVELOPMENT | 1.5844 | 0.0988 |
| SCG2\|NKX2- | GOBP_NEGATIVE_REGULATION_OF_EPITHEL | 1.5828 | 0.0993 |
| ITGB1BP1\|C | GOBP_SPROUTING_ANGIOGENESIS | 1.5712 | 0.098 |
| NKX3-2\|HOX | GOBP_EMBRYONIC_SKELETAL_SYSTEM_DEV | 1.5695 | 0.0989 |
| ITGB1BP1\|V | GOBP_REGULATION_OF_BLOOD_VESSEL_E | 1.5691 | 0.0976 |
| SLC9A3R2\| | GOBP_PHOSPHATIDYLINOSITOL_3_KINASE_S | 1.5653 | 0.0985 |
| FZD5\|COBL | GOBP_SEGMENTATION | 1.5648 | 0.0983 |
| ADCYAP1R1 | GOBP_REGULATION_OF_LIPASE_ACTIVITY | 1.5612 | 0.1 |
| KRT83\|KRT | GOBP_KERATINOCYTE_DIFFERENTIATION | -1.8933 | 0.0411 |
| KRT83\|KRT | GOBP_KERATINIZATION | -2.0527 | 0.0035 |

aCXCR3 vs Veh nodes

| EnrichmentMap::Genes | EnrichmentMap::Name | EnrichmentMap::NES (Data | EnrichmentMap::fdr_qvalue (Dat |
|---|---|---|---|
| CAV3\|RGS2\|TOMM70A\|DDX39B\|AKAF | GOBP_PHYSIOLOGICAL_CARDIAC_MUSCLE | 1.9927 | 0.0313 |
| CAV3\|RGS2\|TOMM70A\|DDX39B\|AKAF | GOBP_REGULATION_OF_CELL_GROWTH_I | 1.9857 | 0.0029 |
| PPARD\|FGF9\|PTK2\|RPS6KB1\|ITGB1E | GOBP_MUSCLE_CELL_MIGRATION | 1.9313 | 0.0148 |
| PPARD\|FGF9\|PTK2\|RPS6KB1\|SRC\|MI | GOBP_SMOOTH_MUSCLE_CELL_MIGRATIO | 1.9202 | 0.032 |
| RGS2\|ANG2\|PI16\|AKAP1\|YY1\|COL14A | GOBP_MUSCLE_HYPERTROPHY | 1.911 | 0.0502 |
| ATP7A\|IQGAP1\|DOCK7\|FGF9\|RHOA\| | GOBP_VASCULAR_ASSOCIATED_SMOOTH_ | 1.9015 | 0.0723 |
| ZPR1\|SALL4\|DICER1\|BRCA2\|PELO\|DN | GOBP_BLASTOCYST_GROWTH | 1.8948 | 0.0307 |
| RGS2\|DDX39B\|SLC9A1\|PI16\|AKAP1\|Y | GOBP_REGULATION_OF_MUSCLE_HYPERT | 1.8835 | 0.0271 |
| RGS2\|PI16\|AKAP1\|YY1\|PTK2\|SELENO | GOBP_REGULATION_OF_MUSCLE_ADAPTA | 1.8756 | 2.00E-04 |
| LRP6\|ABL1\|ABL2\|NKD1\|IFT80\|SFRP1 | GOBP_REGULATION_OF_NON_CANONICAL_ | 1.846 | 0.0762 |
| IFT172\|FOXA2\|GORAB\|FGF9\|GAS8\|G | GOBP_POSITIVE_REGULATION_OF_SMOOT | 1.8397 | 0.0606 |
| TRPV1\|ATP6V0A4\|ABCG2\|ACE\|KCNM | GOBP_EXCRETION | 1.837 | 0.017 |
| PPARD\|PPARG\|S1PR2\|RHOA\|NFE2L2 | GOBP_NEGATIVE_REGULATION_OF_SMOO | 1.8363 | 1.00E-04 |
| COMT\|AVPR1A\|AGTR2\|OXT\|GAS6\|AV | GOBP_REGULATION_OF_RENAL_SYSTEM_ | 1.8281 | 0.0825 |
| NR0B1\|EP300\|TCF21\|IGF1\|FOXH1\|SR | GOBP_REGULATION_OF_INTRACELLULAR_ | 1.8219 | 0.02 |
| PARP2\|DDX39B\|AKAP6\|CCN4\|IGF1\|FD | GOBP_POSITIVE_REGULATION_OF_CELL_C | 1.8174 | 0.0095 |
| CASP6\|STK4\|GSN\|RB1\|PRKAA1\|ATF2 | GOBP_HEPATOCYTE_APOPTOTIC_PROCES | 1.8018 | 0.0433 |
| HSPG2\|MEGF8\|FGF9\|MYF5\|SHH\|MEF | GOBP_EMBRYONIC_SKELETAL_SYSTEM_D | 1.7958 | 0.0292 |
| HJV\|NUMA1\|ELAPOR2\|MIR675\|KCP\|H | GOBP_REGULATION_OF_BMP_SIGNALING_ | 1.7895 | 1.00E-04 |
| LRP6\|GPC3\|ANKRD6\|MKS1\|ABL1\|ABL | GOBP_REGULATION_OF_WNT_SIGNALING_ | 1.7814 | 0.0667 |
| RGS2\|RAP1GDS1\|CHGA\|CAV1\|CDC42 | GOBP_G_PROTEIN_COUPLED_RECEPTOR_ | 1.7807 | 0.073 |
| PHB\|EP300\|TCF21\|DDX5\|SMARCA4\|I | GOBP_REGULATION_OF_ANDROGEN_RECE | 1.78 | 0.0959 |
| IDH2\|NAMPT\|HAAO\|KYNU\|NMNAT1\|G6 | GOBP_NICOTINAMIDE_NUCLEOTIDE_BIOSY | 1.7798 | 0.0099 |
| ZPR1\|COPS2\|SALL4\|BRCA2\|PELO\|DN | GOBP_INNER_CELL_MASS_CELL_PROLIFER | 1.7782 | 0.0128 |
| NR0B1\|CALR\|PHB\|TCF21\|CNOT2\|CYF | GOBP_NEGATIVE_REGULATION_OF_INTRA | 1.7775 | 0.0583 |
| CAV3\|RGS2\|TOMM70A\|PPARG\|FBXO | GOBP_NEGATIVE_REGULATION_OF_MUSCL | 1.7766 | 0.0025 |
| FGF9\|FGFR1\|SHH\|IRS1\|SIX1\|TBX1\|SO | GOBP_MESENCHYMAL_CELL_PROLIFERATI | 1.7663 | 0.0203 |
| NEUROG1\|TACR2\|TBX3\|AGT\|HTR2B\| | GOBP_PHASIC_SMOOTH_MUSCLE_CONTRA | 1.7656 | 0.083 |
| VANGL2\|CTNNB1\|VEGFA\|STK4\|PLXNI | GOBP_BRANCHING_INVOLVED_IN_BLOOD_ | 1.7651 | 0.0511 |
| PGR\|CDH2\|HNF4A\|SLC9A4\|BHLHA15\| | GOBP_GLANDULAR_EPITHELIAL_CELL_DEV | 1.763 | 0.0317 |
| COMT\|AVPR1A\|AGTR2\|OXT\|AVPR2\|A | GOBP_RENAL_SODIUM_EXCRETION | 1.7619 | 0.0383 |
| FOXP1\|PHB\|TCF21\|NODAL\|NCOR1\|SI | GOBP_NEGATIVE_REGULATION_OF_ANDRO | 1.7602 | 0.0489 |
| PXDN\|COL14A1\|LMX1B\|ADAMTS12\|M | GOBP_COLLAGEN_FIBRIL_ORGANIZATION | 1.7597 | 0.0843 |
| PPARD\|RGS2\|KLK1B1\|CHGA\|PTK2\|O | GOBP_REGULATION_OF_TUBE_SIZE | 1.7559 | 0.014 |
| IQGAP1\|FGF9\|PDE4D\|PTK2\|RPS6KB1 | GOBP_POSITIVE_REGULATION_OF_SMOOT | 1.7555 | 0.0016 |
| NPPC\|NPPB\|ADCY3\|NPPA\|ADCY10\|A | GOBP_CYCLIC_NUCLEOTIDE_BIOSYNTHETI | 1.7537 | 0.008 |
| FGF2\|CLEC14A\|VEGFA\|NR2F2\|PKD1\| | GOBP_LYMPH_VESSEL_DEVELOPMENT | 1.7518 | 0.0647 |
| BMP6\|JAK2\|PPARG\|BMP2\|D130043K2 | GOBP_POSITIVE_REGULATION_OF_SMAD_ | 1.7507 | 0.0294 |
| SLC12A6\|PRKG2\|LRRC8A\|SHANK3\|G | GOBP_CELL_VOLUME_HOMEOSTASIS | 1.7436 | 0.035 |
| EP300\|NUMA1\|FGF9\|ELAPOR2\|ACVR | GOBP_POSITIVE_REGULATION_OF_TRANS | 1.7414 | 0.0114 |
| CDKN1A\|NQO2\|DDX39B\|FGF2\|FGF9\| | GOBP_VASCULAR_ASSOCIATED_SMOOTH_ | 1.7408 | 0.0521 |
| CDH2\|HES1\|HNF4A\|BHLHA15\|GDF11\| | GOBP_ENTEROENDOCRINE_CELL_DIFFERE | 1.7361 | 0.0227 |
| RGS2\|HOMER1\|ANG2\|COL14A1\|TMT | GOBP_MUSCLE_CELL_DEVELOPMENT | 1.734 | 0.0376 |
| NMU\|PRAP1\|GPR39\|TRPV1\|OXTR\|AB | GOBP_REGULATION_OF_DIGESTIVE_SYSTE | 1.7339 | 1.00E-04 |
| IGFBP4\|BMP2\|ZFAND2B\|PIK3R1\|IGF1 | GOBP_INSULIN_LIKE_GROWTH_FACTOR_R | 1.733 | 0.0584 |
| AVPR1A\|BRINP1\|NPAS1\|MBD2\|PTEN\| | GOBP_PARENTAL_BEHAVIOR | 1.7327 | 0.0313 |
| PRKG2\|GOLGB1\|SIX2\|MUSTN1\|COMP | GOBP_CHONDROCYTE_PROLIFERATION | 1.7319 | 0.0387 |
| PPARD\|NR0B1\|EP300\|TCF21\|NR1D1\| | GOBP_HORMONE_MEDIATED_SIGNALING_P | 1.7313 | 0.052 |
| HJV\|NUMA1\|OGT\|FGF9\|NREP\|MIR675 | GOBP_REGULATION_OF_TRANSMEMBRANE | 1.7255 | 0.0127 |
| FOXA2\|LRP6\|MEGF8\|CTNNB1\|GDF3\| | GOBP_GASTRULATION_WITH_MOUTH_FOR | 1.722 | 0.0064 |
| RGS2\|NMU\|TRPV1\|OXTR\|LCK\|GRIP2\| | GOBP_SMOOTH_MUSCLE_CONTRACTION | 1.721 | 0.0668 |
| LRP6\|CTNNB1\|FGF9\|FGFR1\|SHH\|IRS | GOBP_REGULATION_OF_MESENCHYMAL_C | 1.7208 | 0.0977 |
| IDH2\|NAMPT\|HAAO\|KYNU\|NMNAT1\|NM | GOBP_PYRIDINE_NUCLEOTIDE_BIOSYNTHE | 1.7207 | 0.0477 |
| CNOT2\|CYP7B1\|UBA5\|LBH\|FOXA1\|FC | GOBP_REGULATION_OF_INTRACELLULAR_ | 1.7195 | 0 |
| VEGFB\|CADM4\|MAPK14\|CD59A\|CLEC | GOBP_VASCULAR_ENDOTHELIAL_GROWTH | 1.7186 | 0.064 |
| NR0B1\|EP300\|TCF21\|FOXH1\|SRC\|GH | GOBP_STEROID_HORMONE_MEDIATED_SIG | 1.718 | 0.0263 |
| HSPG2\|FGF9\|SLC39A14\|LTBP3\|RELA | GOBP_CHONDROCYTE_DIFFERENTIATION | 1.7152 | 0.0516 |
| LRP6\|CTNNB1\|FGF9\|FGFR1\|SHH\|IRS | GOBP_POSITIVE_REGULATION_OF_MESEN | 1.7128 | 0.047 |
| GNG7\|SHANK3\|ATP1A2\|EPHB2\|RPS6 | GOBP_FEAR_RESPONSE | 1.712 | 0.0309 |
| OLFR734\|CHGA\|DEFB1\|PDE3B\|SOX9 | GOBP_CYCLIC_NUCLEOTIDE_MEDIATED_SI | 1.7104 | 0.0645 |
| ATP7A\|CLASP2\|FKBP10\|SMPD3\|RGC | GOBP_EXTRACELLULAR_MATRIX_ASSEMBL | 1.7078 | 0.0768 |
| RGS2\|PI16\|AKAP1\|YY1\|PTK2\|SELENO | GOBP_MUSCLE_ADAPTATION | 1.7067 | 0.0662 |
| AVPR1A\|RNF165\|NR5A1\|WRN\|COMP\| | GOBP_MULTICELLULAR_ORGANISM_AGING | 1.7061 | 1.00E-04 |
| FKBP4\|PHB\|EP300\|TCF21\|DDX5\|RHO | GOBP_ANDROGEN_RECEPTOR_SIGNALING | 1.701 | 0.012 |
| PPARD\|RGS2\|KLK1B1\|PTK2\|KCNMA1 | GOBP_VASODILATION | 1.6987 | 3.00E-04 |
| DDX39B\|SLC9A1\|AKAP6\|PRKCA\|ROC | GOBP_POSITIVE_REGULATION_OF_MUSCL | 1.6984 | 0.0169 |
| KCNE1\|HCN1\|HCN2\|AKAP6\|GPD1\|PDE | GOBP_CELLULAR_RESPONSE_TO_CAMP | 1.696 | 0.0149 |
| SLC12A6\|ATP12A\|ATP1A2\|ATP1B1\|S | GOBP_POTASSIUM_ION_HOMEOSTASIS | 1.6955 | 0.0079 |
| KCNE1\|SIX3\|PGR\|VEGFA\|TFCP2L1\|EI | GOBP_EPITHELIAL_CELL_MATURATION | 1.6931 | 0.0351 |
| DCC\|EPHB2\|PTK2\|LMX1B\|SCN1B\|NIN\| | GOBP_CENTRAL_NERVOUS_SYSTEM_NEUR | 1.6927 | 0.0039 |
| ZFPM1\|ZBTB14\|HEY1\|HEY2\|JAG1\|ME | GOBP_HEART_VALVE_DEVELOPMENT | 1.6921 | 0.0441 |
| COMT\|AVPR1A\|AGTR2\|OXT\|AVPR2\|A | GOBP_REGULATION_OF_EXCRETION | 1.6855 | 0.0446 |
| STRA6L\|SCARB1\|SLC25A19\|SLC46A | GOBP_VITAMIN_TRANSMEMBRANE_TRANSF | 1.6821 | 0.0536 |
| IGFBP4\|BMP2\|ZFAND2B\|IGF1\|IGFBP2 | GOBP_REGULATION_OF_INSULIN_LIKE_GRO | 1.68 | 0.0593 |
| USP15\|DEFB34\|CX3CR1\|PLCG2\|CLE | GOBP_ANTIFUNGAL_INNATE_IMMUNE_RESF | 1.6791 | 0.0204 |
| CHGA\|KISS1\|PLCB2\|PLCD1\|P2RY10B | GOBP_PHOSPHOLIPASE_C_ACTIVATING_G_ | 1.6762 | 0.0826 |
| CHGA\|OXTR\|HMGCR\|GRIP2\|SNTA1\|M | GOBP_VASOCONSTRICTION | 1.6743 | 0.0158 |
| CAV3\|RGS2\|TOMM70A\|BMP2\|PI16\|YY | GOBP_NEGATIVE_REGULATION_OF_STRIAT | 1.6665 | 0 |
| KLRE1\|KLRK1\|CLNK\|PARP9\|EREG\|CD | GOBP_POSITIVE_REGULATION_OF_DEFENS | -1.5592 | 0.075 |
| RBMX\|UPF1\|SAA3\|MYLK3\|TANK\|NR1D | GOBP_RESPONSE_TO_INTERLEUKIN_1 | -1.56 | 0.0745 |
| CD226\|KLRB1F\|SERPINB9H\|CLEC12E | GOBP_REGULATION_OF_NATURAL_KILLER | -1.5603 | 9.00E-04 |
| YY1\|SUPT6\|THOC1\|PARP3\|IL27RA\|AP | GOBP_SOMATIC_DIVERSIFICATION_OF_IMM | -1.5619 | 0.032 |
| IFNA13\|IFNA16\|IFNE\|IFNA15\|IFNA12\|S | GOBP_LEUKOCYTE_PROLIFERATION | -1.5626 | 0.0662 |
| SERPINI1\|SERPINA3K\|SERPINA3G\|SE | GOBP_NEGATIVE_REGULATION_OF_ENDOP | -1.574 | 0.0385 |
| CTLA4\|TBC1D10C\|PGLYRP3\|BANK1\| | GOBP_NEGATIVE_REGULATION_OF_LYMPH | -1.5796 | 0.0755 |
| PGLYRP3\|KLRE1\|PTPN6\|BCR\|SLAMF | GOBP_NEGATIVE_REGULATION_OF_IMMUN | -1.581 | 0.0951 |
| C8B\|C9\|FGL2\|GZMB\|HC\|PRF1\|C8A\|C8 | GOBP_ADAPTIVE_IMMUNE_RESPONSE_BAS | -1.582 | 0.0342 |
| SFN\|BCL11B\|OVOL2\|IFT74\|SRSF6\|MD | GOBP_REGULATION_OF_KERATINOCYTE_F | -1.5831 | 0.0139 |
| ASAH1\|HEXA\|HEXB\|ST3GAL1\|SPHK2\| | GOBP_CERAMIDE_METABOLIC_PROCESS | -1.5853 | 0.0473 |
| BST2\|BCR\|TWIST2\|FGL2\|EVPL\|LBP\|IV | GOBP_REGULATION_OF_IMMUNE_EFFECTO | -1.5911 | 0.0231 |
| CFHR2\|CD59A\|SPNS2\|PTPN6\|NOD2\|C | GOBP_REGULATION_OF_HUMORAL_IMMUN | -1.5934 | 0.0608 |
| TBC1D10C\|BANK1\|SUPT6\|THOC1\|ZFI | GOBP_REGULATION_OF_B_CELL_ACTIVAT | -1.5984 | 0.0168 |
| FOXP3\|IL2\|BCL6\|CARMIL2\|LGALS9\|T | GOBP_POSITIVE_REGULATION_OF_REGULA | -1.5986 | 0.0642 |
| C8B\|C9\|HC\|C8A\|C8G\|KRT1\|PRKDC\|PF | GOBP_ACTIVATION_OF_IMMUNE_RESPONS | -1.6009 | 0.0653 |
| PGLYRP3\|KLRE1\|KLRK1\|C9\|PGLYRP | GOBP_CELL_KILLING | -1.601 | 0.0693 |
| EPHB2\|MIR324\|IL13\|IL5\|SLAMF1\|CLNK | GOBP_POSITIVE_REGULATION_OF_PRODU | -1.6058 | 0.0617 |
| IFNA13\|IFNA16\|IFNE\|PGLYRP3\|IFNA1 | GOBP_LYMPHOCYTE_ACTIVATION_INVOLVE | -1.6106 | 0.0349 |
| ZBTB1\|SLAMF1\|CD160\|IL1A\|TYK2\|ICA | GOBP_POSITIVE_REGULATION_OF_CELL_C | -1.6106 | 0.0021 |
| BST2\|AXL\|SLAMF1\|CLNK\|SUPT6\|THO | GOBP_REGULATION_OF_PRODUCTION_OF | -1.6126 | 0.0831 |
| BST2\|KLRE1\|BCR\|ZBTB1\|KLRK1\|SLA | GOBP_REGULATION_OF_LEUKOCYTE_MED | -1.6143 | 0.0351 |
| CD8A\|ZBTB1\|SLAMF1\|GZMB\|NECTIN2 | GOBP_T_CELL_MEDIATED_IMMUNITY | -1.6161 | 0 |
| IL21\|IL10\|IL2\|MLH1\|MSH2\|PMS2\|ATAD | GOBP_POSITIVE_REGULATION_OF_B_CELL | -1.6202 | 0.0451 |
| AXL\|LYN\|MEF2C\|LGALS3\|MIF\|ST6GAL | GOBP_REGULATION_OF_LEUKOCYTE_APO | -1.6225 | 7.00E-04 |
| XRCC6\|BCL11B\|YY1\|SUPT6\|THOC1\|P | GOBP_SOMATIC_DIVERSIFICATION_OF_IMM | -1.6382 | 0.0527 |
| FGL2\|CASP3\|LRG1\|IL1A\|CCL28\|ADAM | GOBP_LEUKOCYTE_CELL_CELL_ADHESION | -1.6404 | 0.0306 |
| KLRE1\|ZBTB1\|KLRK1\|SLAMF1\|CLNK\| | GOBP_POSITIVE_REGULATION_OF_IMMUNE | -1.6418 | 0.0128 |
| RBMX\|UPF1\|SAA3\|MYLK3\|ZC3H12A\|T | GOBP_CELLULAR_RESPONSE_TO_INTERLE | -1.6459 | 0.0733 |
| EXOG\|REXO4\|DICER1\|AIFM1\|FOXL2\|D | GOBP_DNA_CATABOLIC_PROCESS_ENDON | -1.6468 | 0.0517 |
| PTGDR\|UOX\|XDH\|NT5C1A\|URAD\|NT5 | GOBP_ADENOSINE_METABOLIC_PROCESS | -1.6492 | 0.0733 |
| SOCS5\|NOD2\|NDFIP1\|CCR2\|IL27RA\|C | GOBP_REGULATION_OF_TYPE_2_IMMUNE_ | -1.6513 | 0.0775 |
| SLAMF1\|CASP3\|FCGR2B\|IL1A\|TYK2\|I | GOBP_REGULATION_OF_LEUKOCYTE_PRO | -1.6531 | 0.0669 |
| KLRE1\|KLRK1\|CLNK\|PARP9\|MIF\|CGA | GOBP_POSITIVE_REGULATION_OF_RESPO | -1.6554 | 2.00E-04 |
| ZFP36L1\|TWIST2\|FGL2\|FOXN1\|HOXA | GOBP_REGULATION_OF_HEMOPOIESIS | -1.6645 | 0.0604 |
| BCL11B\|GPAM\|LYN\|IL3\|LGALS3\|MIF\|R | GOBP_REGULATION_OF_LYMPHOCYTE_AP | -1.6647 | 0.0453 |
| CRKL\|AXL\|LYN\|MEF2C\|LGALS3\|MIF\|S | GOBP_LEUKOCYTE_APOPTOTIC_PROCESS | -1.666 | 0.0514 |
| IL1R1\|IL18\|NLRP10\|SLC11A1\|SLAMF1 | GOBP_POSITIVE_REGULATION_OF_T_HELF | -1.6667 | 0.0022 |
| MALT1\|CCL19\|NFKBID\|IL18\|SOCS5\|M | GOBP_POSITIVE_REGULATION_OF_T_HELF | -1.6676 | 0.0168 |
| PHPT1\|LAPTM5\|ELF1\|SLA2\|LILRB4A\| | GOBP_NEGATIVE_REGULATION_OF_T_CEL | -1.6678 | 0.0249 |
| CD226\|KLRE1\|SH2D1B1\|KLRK1\|LAG3 | GOBP_POSITIVE_REGULATION_OF_CELL_K | -1.6696 | 0.0138 |
| ZFPM1\|ZP3\|RBPJ\|VTCN1\|NDFIP1\|CD1 | GOBP_INTERLEUKIN_4_PRODUCTION | -1.6747 | 0.0234 |
| IKZF1\|AXL\|ZBTB1\|ZFP36L1\|NINJ1\|IL21 | GOBP_POSITIVE_REGULATION_OF_HEMOP | -1.675 | 0.0719 |
| PEMT\|PTGDR\|UOX\|XDH\|ENPP4\|NT5C | GOBP_PURINE_RIBONUCLEOSIDE_METABO | -1.6766 | 0.0354 |
| CRKL\|LYN\|LGALS3\|MIF\|TNFRSF4\|WN | GOBP_LYMPHOCYTE_APOPTOTIC_PROCES | -1.6768 | 0.0146 |
| ZP3\|NLRP6\|TREM2\|NOD2\|CD28\|CCR7 | GOBP_REGULATION_OF_INFLAMMATORY_ | -1.6774 | 3.00E-04 |
| CCL21A\|TNFRSF14\|CCL19\|CD86\|CA | GOBP_LYMPHOCYTE_COSTIMULATION | -1.6786 | 0.0206 |
| IL17RA\|ZC3H12A\|EPHB2\|IL27RA\|TYK2 | GOBP_T_HELPER_17_TYPE_IMMUNE_RESP | -1.6789 | 0.0045 |
| CD226\|KLRE1\|KLRK1\|PVR\|CRTAM\|NE | GOBP_IMMUNE_RESPONSE_TO_TUMOR_CE | -1.6796 | 0.0037 |
| SLAMF1\|VSIR\|SASH3\|IL25\|TRAF6\|GB | GOBP_T_CELL_CYTOKINE_PRODUCTION | -1.682 | 0.0135 |
| SLAMF1\|GATA3\|IFNB1\|IL12A\|IL12B\|IL | GOBP_CD4_POSITIVE_ALPHA_BETA_T_CEL | -1.6878 | 0.0298 |
| TRAF2\|MALT1\|MAP3K7\|SLAMF1\|GAT | GOBP_POSITIVE_REGULATION_OF_T_CELL | -1.6882 | 0.0642 |
| TRAF2\|MALT1\|DEFB25\|CARMIL2\|PNF | GOBP_POSITIVE_REGULATION_OF_INTERL | -1.6884 | 0.0661 |
| RGMA\|LRIG2\|SH3D19\|APOE\|PTPN3\|S | GOBP_REGULATION_OF_MEMBRANE_PROT | -1.6886 | 5.00E-04 |
| PPARD\|SFN\|BCL11B\|OVOL2\|IFT74\|SF | GOBP_KERATINOCYTE_PROLIFERATION | -1.6895 | 0.0622 |
| SUPT6\|THOC1\|PARP3\|IL27RA\|APLF\|M | GOBP_SOMATIC_DIVERSIFICATION_OF_IMM | -1.6905 | 0.0856 |
| CD226\|PTPN6\|NOD2\|GIMAP5\|SUPT6\|T | GOBP_REGULATION_OF_B_CELL_MEDIATE | -1.6906 | 2.00E-04 |
| H2-OA\|H2-EB2\|H2-DMB1\|H2-DMB2\|IG | GOBP_IMMUNOGLOBULIN_PRODUCTION_IN | -1.6923 | 0.0147 |
| RGN\|EXOG\|REXO4\|TREX2\|DICER1\|XF | GOBP_DNA_CATABOLIC_PROCESS | -1.6943 | 0.0808 |
| KLRE1\|KLRK1\|CD160\|KLRC3\|PRKCH\| | GOBP_IMMUNE_RESPONSE_REGULATING_C | -1.6949 | 1.00E-04 |
| TNFRSF14\|ZC3H12A\|TARM1\|RUNX1\|F | GOBP_NEGATIVE_REGULATION_OF_CD4_F | -1.6951 | 0.0855 |
| KLRE1\|PTPN6\|KLRK1\|CTSG\|GZMB\|N | GOBP_LEUKOCYTE_MEDIATED_CYTOTOXI | -1.6987 | 0.0688 |
| KLRE1\|ZBTB1\|KLRK1\|SLAMF1\|CLNK\| | GOBP_REGULATION_OF_LYMPHOCYTE_ME | -1.6997 | 2.00E-04 |
| GADD45G\|IL27\|SLAMF1\|IL27RA\|TRAF | GOBP_T_HELPER_1_TYPE_IMMUNE_RESPO | -1.7047 | 0.0131 |
| C8B\|C9\|FGL2\|GZMB\|HC\|PRF1\|C8A\|C8 | GOBP_LYMPHOCYTE_MEDIATED_IMMUNITY | -1.7108 | 0.0102 |
| TRAF2\|MALT1\|LILRB4A\|MAP3K7\|SLA | GOBP_REGULATION_OF_T_CELL_CYTOKIN | -1.7153 | 3.00E-04 |
| BCL6\|IL18\|SOCS5\|BCL3\|HLX\|BATF\|AN | GOBP_T_HELPER_2_CELL_DIFFERENTIATIO | -1.7159 | 0.0221 |
| BCL2\|CRKL\|TNFRSF21\|LYN\|IL3\|BAK1 | GOBP_B_CELL_APOPTOTIC_PROCESS | -1.7171 | 0.0653 |
| H2-EA\|FCER1G\|H2-OA\|H2-EB2\|H2-DM | GOBP_ANTIGEN_PROCESSING_AND_PRESE | -1.7191 | 0.0495 |
| SLAMF1\|CASP3\|IL1A\|TYK2\|IL21\|CDKN | GOBP_REGULATION_OF_T_CELL_PROLIFE | -1.7193 | 0.0757 |
| CD226\|KLRB1F\|SERPINB9H\|H60B\|H60 | GOBP_NATURAL_KILLER_CELL_MEDIATED_ | -1.7215 | 0.0234 |
| IKZF1\|PGLYRP3\|AXL\|ZBTB1\|ZFP36L1 | GOBP_REGULATION_OF_LEUKOCYTE_DIFF | -1.7222 | 0.0637 |
| CD226\|NOD2\|GIMAP5\|NECTIN2\|TNF\|T | GOBP_POSITIVE_REGULATION_OF_B_CELL | -1.7269 | 0.0111 |
| CD226\|ZFPM1\|ZP3\|CARMIL2\|PDE4D\|H | GOBP_POSITIVE_REGULATION_OF_INTERF | -1.7286 | 3.00E-04 |
| ZBTB1\|SLAMF1\|SUPT6\|THOC1\|PARP | GOBP_REGULATION_OF_ADAPTIVE_IMMUN | -1.7291 | 0.09 |
| ZBTB1\|SLAMF1\|FGL2\|CASP3\|CD160\| | GOBP_REGULATION_OF_T_CELL_ACTIVAT | -1.7305 | 0.0318 |
| ERO1A\|DNAJB5\|DNAJC7\|DNAJB12\|SD | GOBP_CHAPERONE_COFACTOR_DEPENDE | -1.7378 | 0.0954 |
| TICAM1\|IL17RA\|KPNA6\|IL17A\|IL17RC\| | GOBP_POSITIVE_REGULATION_OF_CYTOK | -1.7401 | 0.0687 |
| AP1G1\|IFNA13\|RAB27A\|IFNA16\|IFNE\| | GOBP_NATURAL_KILLER_CELL_ACTIVATIO | -1.7422 | 0.028 |
| PHB\|CCL1\|RFTN1\|NOD2\|MIR301\|CAR | GOBP_INTERLEUKIN_17_PRODUCTION | -1.7425 | 3.00E-04 |
| SH3GLB1\|CAPN1\|PCSK9\|SNX25\|ABCA | GOBP_RECEPTOR_CATABOLIC_PROCESS | -1.7439 | 0.0135 |
| PSAP\|SFN\|BCL11B\|KRT2\|DNASE1L2\| | GOBP_KERATINOCYTE_DEVELOPMENT | -1.7442 | 0.0739 |
| ZBTB1\|SLAMF1\|CD160\|IL1A\|TYK2\|ICA | GOBP_POSITIVE_REGULATION_OF_LEUKO | -1.7483 | 0.0532 |
| GRIN1\|GRIN2B\|KRT1\|H2-KE6\|PKD1\|P | GOBP_PROTEIN_HETEROTETRAMERIZATIO | -1.7522 | 0.0177 |
| KLRE1\|ZBTB1\|KLRK1\|SLAMF1\|CLNK\| | GOBP_POSITIVE_REGULATION_OF_LEUKO | -1.7539 | 0.0884 |
| BCL2\|LYN\|IL3\|MIF\|BAX\|CD74\|SLC39A1 | GOBP_REGULATION_OF_B_CELL_APOPTO | -1.7542 | 0.0812 |
| CD226\|AP1G1\|KLRE1\|SH2D1B1\|KLRK | GOBP_POSITIVE_REGULATION_OF_NATUR | -1.7555 | 0.0431 |
| CCL19\|SOCS5\|IL27\|NLRP10\|SLAMF1\| | GOBP_REGULATION_OF_T_HELPER_1_TYP | -1.7564 | 0.0225 |
| MALT1\|CCL19\|NFKBID\|SOCS5\|MIR32 | GOBP_POSITIVE_REGULATION_OF_CD4_P | -1.7573 | 0.038 |
| MALT1\|ZC3H12A\|NFKBID\|RC3H2\|RC3 | GOBP_REGULATION_OF_T_HELPER_17_TY | -1.7605 | 0.0026 |
| IL21\|PRDM1\|PGLYRP3\|LGALS9\|FLT3 | GOBP_REGULATION_OF_NATURAL_KILLER | -1.7617 | 0.0622 |
| LPCAT4\|PLA2G10\|ABHD12B\|PLA2G2 | GOBP_PHOSPHATIDYLSERINE_METABOLIC_ | -1.7622 | 0.0146 |
| KCNE1\|KCNK4\|RAB11B\|SLC9A1\|NOX | GOBP_CELLULAR_RESPONSE_TO_PH | -1.763 | 0.0637 |
| SLAMF1\|CASP3\|IL1A\|TYK2\|ELF4\|IL21 | GOBP_T_CELL_PROLIFERATION | -1.7652 | 0.0356 |
| IFT172\|SFN\|CASK\|KLF9\|NFATC1\|PTP | GOBP_NEGATIVE_REGULATION_OF_KERAT | -1.7661 | 0.0024 |
| TNFRSF14\|TARM1\|CARD11\|LILRB4A | GOBP_CD4_POSITIVE_ALPHA_BETA_T_CEL | -1.7667 | 0.0354 |
| CD247\|CARMIL2\|PDE4D\|GPAM\|VTCN | GOBP_INTERLEUKIN_2_PRODUCTION | -1.7702 | 0.0516 |
| ZP3\|CD6\|NOD2\|MIR21A\|MIR155\|IL36R | GOBP_INFLAMMATORY_RESPONSE_TO_AN | -1.7713 | 0.0866 |
| KMT5B\|HMCES\|KMT5C\|NDFIP1\|CD28 | GOBP_REGULATION_OF_ISOTYPE_SWITCH | -1.772 | 0.0146 |
| TICAM1\|AP1G1\|BLOC1S3\|FLT3L\|GAS | GOBP_POSITIVE_REGULATION_OF_NATUR | -1.7726 | 0.0092 |
| STK11\|ZAP70\|CD1D1\|SHH\|TOX\|CD3D | GOBP_POSITIVE_THYMIC_T_CELL_SELECT | -1.777 | 0.0135 |
| ERO1A\|SH3GLB1\|DNAJB5\|DNAJC7\|DN | GOBP_DE_NOVO_PROTEIN_FOLDING | -1.7801 | 0.0505 |
| MALT1\|CCL19\|ZC3H12A\|NFKBID\|RC3 | GOBP_REGULATION_OF_T_HELPER_CELL_ | -1.788 | 0.0221 |
| AXL\|SHH\|ZBTB1\|LCK\|WNT10B\|IL21\|AD | GOBP_POSITIVE_REGULATION_OF_LYMPHO | -1.7912 | 0.0959 |
| SHH\|GIMAP5\|SASH3\|GIMAP3\|NCKAP1 | GOBP_POSITIVE_REGULATION_OF_ALPHA_ | -1.7943 | 0.0315 |
| IL1A\|ADAM8\|B2M\|EGR3\|ANXA1\|IL7R\|F | GOBP_POSITIVE_REGULATION_OF_LYMPHO | -1.795 | 0.01 |
| ZBTB1\|SLAMF1\|MIF\|NECTIN2\|TNF\|TR | GOBP_POSITIVE_REGULATION_OF_ADAPTI | -1.7999 | 0.0354 |
| CDKN1A\|CSF1\|CSF1R\|CSF2\|CSF2RA | GOBP_POSITIVE_REGULATION_OF_LEUKO | -1.804 | 0.0056 |
| IKZF1\|ZBTB1\|ZFP36L1\|FGL2\|FOXN1\| | GOBP_T_CELL_DIFFERENTIATION | -1.8056 | 0.0457 |
| VTCN1\|BCL3\|GBP8\|IL12A\|TRBV13-3\| | GOBP_RESPONSE_TO_PROTOZOAN | -1.812 | 0.0524 |
| KRT83\|KRT71\|KRT72\|KRT81\|KRT75\| | GOBP_INTERMEDIATE_FILAMENT_BASED_F | -1.8215 | 0.0514 |
| CTLA2A\|DROSHA\|CTLA4\|FANCA\|DIC | GOBP_REGULATORY_T_CELL_DIFFERENTI | -1.8226 | 0.0148 |
| C6\|FCER2A\|IL17A\|PHB\|KLK7\|IL17F\|KL | GOBP_POSITIVE_REGULATION_OF_HUMOR | -1.8369 | 0.0656 |
| ZC3H12A\|GADD45G\|IL27\|GIMAP5\|SA | GOBP_CD4_POSITIVE_ALPHA_BETA_T_CEL | -1.8378 | 0.0092 |
| PGLYRP3\|AXL\|ZBTB1\|ZFP36L1\|PGLY | GOBP_REGULATION_OF_LYMPHOCYTE_DIF | -1.8379 | 3.00E-04 |
| ACTR2\|VPS26B\|DAPK1\|RAB43\|RPS6 | GOBP_CELLULAR_RESPONSE_TO_INTERFE | -1.8416 | 0.0039 |
| CD160\|TYK2\|ELF4\|IL21\|NKG7\|ANXA1 | GOBP_ALPHA_BETA_T_CELL_ACTIVATION | -1.8464 | 0.0317 |
| FOXP3\|CD40\|IL2\|NDFIP1\|CD28\|SLC15 | GOBP_ISOTYPE_SWITCHING_TO_IGG_ISOT | -1.8511 | 0.0319 |
| SFN\|IGF1\|HRNR\|KDF1\|TFAP2B\|TMEM | GOBP_WATER_HOMEOSTASIS | -1.8561 | 0.0503 |
| CARD11\|CD28\|CD3E\|BLM\|CCR2\|CD8 | GOBP_POSITIVE_REGULATION_OF_ALPHA_ | -1.858 | 0.0093 |
| ZC3H12A\|IL27\|SHH\|GIMAP5\|SASH3\|G | GOBP_REGULATION_OF_ALPHA_BETA_T_C | -1.8607 | 0.0763 |
| SHH\|IL21\|ANXA1\|SATB1\|PSAP\|PRR7\|E | GOBP_ALPHA_BETA_T_CELL_DIFFERENTIA | -1.8652 | 0.0382 |
| CD226\|MALT1\|PHPT1\|CARD11\|RC3H | GOBP_REGULATION_OF_T_CELL_RECEPT | -1.8657 | 0.0482 |
| ZC3H12A\|IL27\|GIMAP5\|SASH3\|GIMAP | GOBP_REGULATION_OF_CD4_POSITIVE_AL | -1.8742 | 0.0665 |
| SHH\|ZBTB1\|SOS1\|ZFP36L1\|FOXN1\|IL | GOBP_T_CELL_DIFFERENTIATION_IN_THYM | -1.8802 | 0.0955 |
| CCL28\|RRAGA\|C8G\|LYZL6\|MMD2\|TG | GOBP_CYTOLYSIS | -1.8835 | 1.00E-04 |
| IFITM3\|ACTR2\|IFITM1\|VPS26B\|BST2\| | GOBP_RESPONSE_TO_INTERFERON_GAMM | -1.8995 | 0.0281 |
| IKZF1\|PRDM1\|PGLYRP3\|FLT3L\|GAS6 | GOBP_NATURAL_KILLER_CELL_DIFFERENT | -1.9082 | 0.0464 |
| SHH\|SLAMF1\|IL1A\|TRAF6\|TYK2\|IL21\| | GOBP_POSITIVE_REGULATION_OF_T_CELL | -1.9148 | 2.00E-04 |
| CD274\|VSIR\|TYK2\|ELF4\|ARG2\|IL18\|C | GOBP_ALPHA_BETA_T_CELL_PROLIFERAT | -1.9276 | 0.0597 |
| ZC3H12A\|NLRP6\|CD276\|PGLYRP3\|G | GOBP_NEGATIVE_REGULATION_OF_INTER | -1.9337 | 0.0383 |
| MALT1\|CCL19\|NFKBID\|CARD11\|SOC | GOBP_POSITIVE_REGULATION_OF_CD4_P | -1.9376 | 0.0015 |
| CARD11\|AIRE\|ATG5\|CD1D1\|SHH\|CD2 | GOBP_THYMIC_T_CELL_SELECTION | -1.9425 | 0.0038 |
| BLOC1S3\|PGLYRP3\|AXL\|ZBTB1\|CLNK | GOBP_REGULATION_OF_NATURAL_KILLER | -1.9557 | 0.0092 |
| KRT83\|KRT71\|KRT72\|KRT81\|KRT75\| | GOBP_INTERMEDIATE_FILAMENT_ORGANIZ | -1.9562 | 0.0171 |
| SHH\|GIMAP5\|CD160\|SASH3\|TYK2\|GIM | GOBP_POSITIVE_REGULATION_OF_ALPHA_ | -1.9654 | 0.0139 |
| SHH\|CD160\|CD300A\|TYK2\|ANXA1\|CA | GOBP_REGULATION_OF_ALPHA_BETA_T_C | -1.9717 | 0.0293 |
| BCL2\|BCL11B\|BRAF\|CD1D1\|SHH\|CD3 | GOBP_POSITIVE_T_CELL_SELECTION | -1.9752 | 0.0689 |
| CD160\|IL21\|NKG7\|ANXA1\|SATB1\|CAR | GOBP_CD4_POSITIVE_ALPHA_BETA_T_CEL | -1.9954 | 0.0085 |
| CACNB3\|CACNB4\|PSEN1\|CTLA4\|PSE | GOBP_T_CELL_RECEPTOR_SIGNALING_PA | -1.9954 | 0.0055 |
| CSTDC5\|CSTDC6\|COL3A1\|DSP\|SPRR | GOBP_PEPTIDE_CROSS_LINKING | -2.0121 | 1.00E-04 |
| ZC3H12A\|IL27\|GIMAP5\|CD160\|CD274 | GOBP_REGULATION_OF_CD4_POSITIVE_AL | -2.033 | 0.0018 |
| PGLYRP3\|KLRE1\|AXL\|KLRK1\|SLAMF | GOBP_INTERFERON_GAMMA_PRODUCTION | -2.0336 | 0.0526 |
| CSTDC5\|CSTDC6\|KRT83\|NUMA1\|KRT | GOBP_EPIDERMIS_DEVELOPMENT | -2.0356 | 0.0635 |
| IKZF1\|IFNA13\|IFNA16\|IFNE\|BLOC1S3\| | GOBP_NATURAL_KILLER_CELL_ACTIVATIO | -2.0633 | 0.0069 |
| CSTDC5\|CSTDC6\|KRT83\|NUMA1\|KRT | GOBP_EPIDERMAL_CELL_DIFFERENTIATIO | -2.0703 | 0.0599 |
| BCL11B\|SHH\|FOXN1\|SRF\|BCL2\|BTNL | GOBP_T_CELL_SELECTION | -2.0717 | 0.0127 |
| CSTDC5\|CSTDC6\|KRT83\|NUMA1\|KRT | GOBP_SKIN_DEVELOPMENT | -2.1118 | 0.0319 |
| CERS3\|SFN\|KRT83\|KRT71\|KRT72\|SH | GOBP_KERATINIZATION | -2.1822 | 0.0033 |
| SFN\|GRHL1\|HRNR\|KDF1\|TMEM79\|AB | GOBP_REGULATION_OF_WATER_LOSS_VIA | -2.1964 | 2.00E-04 |
| CSTDC5\|CSTDC6\|KRT83\|NUMA1\|KRT | GOBP_KERATINOCYTE_DIFFERENTIATION | -2.2978 | 0.0477 |

DB vs Veh nodes

| EnrichmentMap::Genes | EnrichmentMap::Name | EnrichmentMap::NES (Dataset 1) | EnrichmentMap::fdr_qvalue (Dataset 1) |
|---|---|---|---|
| TFRC\|CFH\|PTPN6\|MIR181B-1\|MIR181B-2\|CD74\|C1RL\| | GOBP_B_CELL_MEDIATED_IMMUNITY | 1.8693 | 0.0181 |
| TFRC\|ZFP36L1\|CASP3\|PTPN6\|IL4I1\|MIR181B-1\|MIR181 | GOBP_REGULATION_OF_B_CELL_ACTIVATION | 1.847 | 0.0176 |
| STX11\|STAT5A\|KLRC1\|KLRC2\|KLRC3\|CEBPG\|PTPN6\| | GOBP_NATURAL_KILLER_CELL_MEDIATED_IMMUNITY | 1.8391 | 0.0114 |
| STAT5A\|KLRC1\|KLRC2\|KLRC3\|CD160\|LAMP1\|SERPINE | GOBP_REGULATION_OF_NATURAL_KILLER_CELL_MEDIATED_IMMUNITY | 1.839 | 0.0092 |
| CD81\|NCKAP1L\|LCK\|CD79A\|CD79B\|LPXN\|PTPN6\|PRKC | GOBP_B_CELL_RECEPTOR_SIGNALING_PATHWAY | 1.8376 | 0.0081 |
| SH2D1B1\|SH2D1A\|SH2D1B2\|RAET1D\|STAT5A\|LAG3\|U | GOBP_POSITIVE_REGULATION_OF_NATURAL_KILLER_CELL_MEDIATED_IMMUNI | 1.8348 | 0.0076 |
| CD47\|EIF2B5\|RAB29\|NR4A3\|KLRC1\|KLRC2\|KLRC3\|PTF | GOBP_IMMUNE_RESPONSE_REGULATING_CELL_SURFACE_RECEPTOR_SIGNAL | 1.8268 | 0.0092 |
| SREBF1\|PARP9\|TLR8\|RPS19\|KLRC1\|TASL\|KLRC2\|KLR | GOBP_POSITIVE_REGULATION_OF_INNATE_IMMUNE_RESPONSE | 1.814 | 0.0115 |
| TYROBP\|ZNFX1\|KLRC1\|KLRC2\|KLRC3\|NLRC4\|CASP6\| | GOBP_ACTIVATION_OF_INNATE_IMMUNE_RESPONSE | 1.7998 | 0.0153 |
| KLRC1\|KLRC2\|KLRC3\|PRKCB\|FOXP1\|KRT1\|C1RL\|C1R | GOBP_ACTIVATION_OF_IMMUNE_RESPONSE | 1.794 | 0.015 |
| WDFY4\|MARCHF1\|PSMB9\|PSMB8\|UNC93B1\|RAB32\|PS | GOBP_ANTIGEN_PROCESSING_AND_PRESENTATION | 1.7843 | 0.0176 |
| GM8909\|MARCHF1\|UNC93B1\|H2-T10\|H2-T23\|CTSS\|CL | GOBP_ANTIGEN_PROCESSING_AND_PRESENTATION_OF_PEPTIDE_ANTIGEN | 1.7821 | 0.0176 |
| TFRC\|ZBTB1\|IFNA13\|IFNA16\|IFNE\|IFNA15\|IFNA12\|ZFP3 | GOBP_B_CELL_ACTIVATION | 1.7744 | 0.0195 |
| TFRC\|BAD\|CD74\|TLR4\|CD40LG\|NSD2\|ATAD5\|TNFRSF | GOBP_POSITIVE_REGULATION_OF_B_CELL_ACTIVATION | 1.7711 | 0.0196 |
| RAC2\|RHOA\|ALOX15\|CDC42\|ALKBH4\|FCER1G\|FCGR3 | GOBP_MEMBRANE_INVAGINATION | 1.77 | 0.0189 |
| TFRC\|MCM3AP\|EZH2\|MIR181B-1\|MIR181B-2\|SIGLECG | GOBP_IMMUNOGLOBULIN_PRODUCTION | 1.747 | 0.0278 |
| PLD4\|MESD\|CD47\|RAC2\|BCR\|ALOX15\|CLN3\|PIP5K1C\|F | GOBP_PHAGOCYTOSIS | 1.7373 | 0.0338 |
| FCER1G\|FCGR3\|IFI30\|UNC93B1\|FCGR1\|CTSS\|CLEC4 | GOBP_ANTIGEN_PROCESSING_AND_PRESENTATION_OF_EXOGENOUS_PEPTID | 1.7239 | 0.0409 |
| CD81\|BCL3\|TNF\|PTPN6\|IGHA\|TRDV4\|TRDC\|TRBC1\|TR | GOBP_HUMORAL_IMMUNE_RESPONSE_MEDIATED_BY_CIRCULATING_IMMUNOG | 1.7201 | 0.0427 |
| TFRC\|ZBTB1\|KLRC1\|KLRC2\|SART1\|KLRC3\|ARID1A\|IC | GOBP_POSITIVE_REGULATION_OF_LYMPHOCYTE_ACTIVATION | 1.7178 | 0.0436 |
| TFRC\|ZBTB1\|KLRC1\|KLRC2\|KLRC3\|WAS\|IL4I1\|MIR181 | GOBP_LYMPHOCYTE_MEDIATED_IMMUNITY | 1.709 | 0.0489 |
| PYCARD\|FCER1G\|MARCHF1\|IFI30\|UNC93B1\|CTSS\|TR | GOBP_ANTIGEN_PROCESSING_AND_PRESENTATION_OF_PEPTIDE_OR_POLYS | 1.7048 | 0.0499 |
| CD47\|TULP1\|IGHA\|TRDV4\|TRDC\|TRBC1\|TRBC2\|IGLC1 | GOBP_PHAGOCYTOSIS_RECOGNITION | 1.7033 | 0.0499 |
| FCER1G\|FCGR3\|IFI30\|UNC93B1\|FCGR1\|PSME1\|PSME | GOBP_ANTIGEN_PROCESSING_AND_PRESENTATION_OF_EXOGENOUS_ANTIGE | 1.6988 | 0.0532 |
| STX11\|HPS1\|ZBTB1\|RHBDD3\|GATA3\|KLRC1\|IFNA13\|KL | GOBP_NATURAL_KILLER_CELL_ACTIVATION | 1.6758 | 0.0731 |
| LILRB4B\|CD81\|RAB29\|LAPTM5\|EZR\|LCK\|LPXN\|PTPN6\| | GOBP_REGULATION_OF_ANTIGEN_RECEPTOR_MEDIATED_SIGNALING_PATHW | 1.6729 | 0.0747 |
| PMS2\|TFRC\|MSH2\|TNF\|RIF1\|PAGR1A\|TGFB1\|STAT6\|E | GOBP_POSITIVE_REGULATION_OF_B_CELL_MEDIATED_IMMUNITY | 1.6715 | 0.0749 |
| TFRC\|CD81\|NCKAP1L\|CDKN1A\|IFNA13\|IFNA16\|CD24A | GOBP_B_CELL_PROLIFERATION | 1.667 | 0.0801 |
| PMS2\|TFRC\|EXO1\|MSH2\|FOXP3\|POU2F2\|MSH6\|NBN\|R | GOBP_IMMUNOGLOBULIN_PRODUCTION_INVOLVED_IN_IMMUNOGLOBULIN_MED | 1.6436 | 0.0998 |
| SOS1\|SOX18\|SOX9\|NUMA1\|TNF\|SOX21\|WNT10A\|WNT1 | GOBP_SKIN_EPIDERMIS_DEVELOPMENT | -1.8064 | 0.0105 |
| SFN\|SRF\|TFAP2B\|TMPRSS13\|HRNR\|KDF1\|FLG2\|TMEM | GOBP_WATER_HOMEOSTASIS | -1.813 | 0.0097 |
| SOS1\|SOX18\|SOX9\|NUMA1\|TNF\|SOX21\|WNT10A\|WNT1 | GOBP_MOLTING_CYCLE | -1.8653 | 0.0023 |
| CSTDC5\|CSTDC6\|DSP\|SPRR1A\|SPRR3\|STFA1\|STFA2 | GOBP_PEPTIDE_CROSS_LINKING | -1.938 | 1.00E-04 |
| SFN\|STARD7\|GRHL1\|SRF\|FLG\|TMPRSS13\|GBA\|HRNR | GOBP_REGULATION_OF_WATER_LOSS_VIA_SKIN | -2.0286 | 0 |
| CSTDC5\|CSTDC6\|KRT83\|NUMA1\|KRT72\|SHARPIN\|OVC | GOBP_EPIDERMAL_CELL_DIFFERENTIATION | -2.2004 | 0 |
| CSTDC5\|CSTDC6\|KRT83\|NUMA1\|KRT72\|SHARPIN\|OVC | GOBP_EPIDERMIS_DEVELOPMENT | -2.208 | 0 |
| CSTDC5\|CSTDC6\|KRT83\|NUMA1\|KRT72\|SHARPIN\|OVC | GOBP_SKIN_DEVELOPMENT | -2.2244 | 0 |
| CERS3\|SFN\|KRT83\|KRT71\|KRT72\|SHARPIN\|KRT81\|KR | GOBP_KERATINIZATION | -2.272 | 0 |
| CSTDC5\|CSTDC6\|KRT83\|NUMA1\|KRT72\|SHARPIN\|OVC | GOBP_KERATINOCYTE_DIFFERENTIATION | -2.387 | 0 |
| LILRB4B\|MAPK8IP1\|WDFY4\|NCKAP1L\|HFE\|IFNG\|CD27 | GOBP_CD8_POSITIVE_ALPHA_BETA_T_CELL_ACTIVATION | 1.8432 | 0.0137 |
| ARMC5\|RHOA\|CLEC4A2\|CLEC4A1\|CLEC4A4\|CLEC4F\| | GOBP_ALPHA_BETA_T_CELL_ACTIVATION | 1.7542 | 0.0248 |
| TNFSF18\|SRF\|FOXP3\|ZAP70\|BCL11B\|STK11\|BRAF\|CD | GOBP_POSITIVE_T_CELL_SELECTION | 1.6978 | 0.0528 |
| TNFSF18\|SRF\|FOXP3\|BCL11B\|STK11\|GATA3\|TOX\|IL6\| | GOBP_T_CELL_SELECTION | 1.6833 | 0.0657 |
| ARMC5\|NCKAP1L\|BCL3\|GATA3\|EOMES\|TOX\|RHOA\|TG | GOBP_ALPHA_BETA_T_CELL_DIFFERENTIATION | 1.6659 | 0.0796 |
| KRT83\|KRT71\|KRT72\|KRT81\|KRT75\|DSP\|KRT88\|KRT5 | GOBP_INTERMEDIATE_FILAMENT_BASED_PROCESS | -2.0984 | 0 |
| KRT83\|KRT71\|KRT72\|KRT81\|KRT75\|DSP\|KRT88\|KRT5 | GOBP_INTERMEDIATE_FILAMENT_ORGANIZATION | -2.1053 | 0 |
| ENO1B\|PPARGC1A\|VCP\|BCL2L1\|MYC\|MAP2K1\|STAT3\| | GOBP_POSITIVE_REGULATION_OF_ATP_BIOSYNTHETIC_PROCESS | 1.7062 | 0.0505 |
| NDUFA3\|DNAJC30\|NDUFC1\|NDUFS1\|NDUFA12\|NDUFB | GOBP_ATP_BIOSYNTHETIC_PROCESS | 1.6451 | 0.0993 |
| HAS2\|KLF9\|SFN\|MDK\|NOTCH2\|PRKD1\|BCL11B\|PTPRK | GOBP_REGULATION_OF_KERATINOCYTE_PROLIFERATION | -1.7193 | 0.0637 |
| HAS2\|KLF9\|SFN\|MDK\|NOTCH2\|PRKD1\|BCL11B\|PTPRK | GOBP_KERATINOCYTE_PROLIFERATION | -1.8316 | 0.006 |
| SLAMF1\|ISG15\|TLR8\|BCL3\|TNF\|CEBPG\|IL1B\|WNT5A\|C | GOBP_POSITIVE_REGULATION_OF_INTERFERON_GAMMA_PRODUCTION | 1.7167 | 0.0427 |
| ISG15\|CD47\|TLR8\|LAPTM5\|BCL3\|GATA3\|TNF\|CEBPG\| | GOBP_INTERFERON_GAMMA_PRODUCTION | 1.654 | 0.0954 |
| TLR8\|DDX58\|IFIH1\|NMB\|FLT3\|ZC3HAV1\|MMP12\|MAVS\| | GOBP_POSITIVE_REGULATION_OF_INTERFERON_ALPHA_PRODUCTION | 1.8124 | 0.0112 |
| TLR8\|DDX58\|IFIH1\|NMB\|FLT3\|ZC3HAV1\|MMP12\|PTPRS | GOBP_INTERFERON_ALPHA_PRODUCTION | 1.7265 | 0.0406 |
| MYC\|PRKN\|NDUFC2\|AKT2\|STOML2\|GRIA1\|MFN2\|NNT\| | GOBP_POSITIVE_REGULATION_OF_MEMBRANE_POTENTIAL | 1.7958 | 0.0154 |
| WDFY1\|PIK3AP1\|TASL\|NINJ1\|CAV1\|TREML4\|LBP\|CD30 | GOBP_POSITIVE_REGULATION_OF_TOLL_LIKE_RECEPTOR_SIGNALING_PATHW | 1.7688 | 0.0183 |
| ATF7\|PIK3CG\|STK4\|BCL2L1\|ARF6\|CASP6\|ATF2\|KRT8\| | GOBP_HEPATOCYTE_APOPTOTIC_PROCESS | 1.6882 | 0.0615 |
| PARP9\|PYCARD\|CCL5\|DDX58\|EIF2AK4\|IL12B\|TREML4\| | GOBP_POSITIVE_REGULATION_OF_DEFENSE_RESPONSE_TO_VIRUS_BY_HOST | 1.6523 | 0.0962 |
| KLF2\|SH2B3\|ZBTB7A\|LYAR\|SRF\|CITED2\|SLC11A2\|RHA | GOBP_ERYTHROCYTE_DEVELOPMENT | -1.7292 | 0.0532 |
| WNT5A\|SEMA3D\|NRP1\|PLXNA3\|RYK\|SEMA3A\|SEMA3B | GOBP_NEGATIVE_REGULATION_OF_AXON_EXTENSION_INVOLVED_IN_AXON_GU | -1.7339 | 0.0515 |
| SFN\|FGF10\|RBPJ\|SNAI2\|OVOL2\|CDKN2C\|CEBPA\|FBLN | GOBP_NEGATIVE_REGULATION_OF_STEM_CELL_PROLIFERATION | -1.7349 | 0.0539 |
| SPDEF\|TIGAR\|SOX9\|GSDMC2\|MIR203\|MIR7-2\|MIR7-1\|S | GOBP_INTESTINAL_EPITHELIAL_CELL_DIFFERENTIATION | -1.7758 | 0.0208 |

| | | | |
|---|---|---|---|
| GPS2\|LMO3\|ALOX5 | GOBP_REGULATION_OF_FAT_CELL_DIFFERENTIATION | 1.4991 | 0.0384 |
| CLDN4\|NEDD4L\|RC | GOBP_CELL_CELL_JUNCTION_ORGANIZATION | 1.4981 | 0.0424 |
| IL12A\|CCBE1\|IL12B | GOBP_VASCULAR_ENDOTHELIAL_GROWTH_FACTOR_SIGNA | 1.498 | 0.0453 |
| SHH\|GRHL2\|NIPBL\| | GOBP_EMBRYONIC_APPENDAGE_MORPHOGENESIS | 1.4976 | 0.0536 |
| PYCARD\|WNT5A\|C | GOBP_POSITIVE_REGULATION_OF_INTERLEUKIN_8_PRODUC | 1.4974 | 0.0543 |
| APP\|SREBF1\|GATA | GOBP_MIRNA_TRANSCRIPTION | 1.4968 | 0.0426 |
| SHH\|STUB1\|LCN5\|F | GOBP_HORMONE_METABOLIC_PROCESS | 1.4968 | 0.0623 |
| SORBS1\|C1QTNF1 | GOBP_REGULATION_OF_CARBOHYDRATE_BIOSYNTHETIC_F | 1.4966 | 0.0537 |
| RPS7\|FBH1\|PYCAR | GOBP_POSITIVE_REGULATION_OF_APOPTOTIC_SIGNALING_ | 1.4963 | 0.0337 |
| FGFR2\|FOXF1\|ID2\| | GOBP_EMBRYONIC_DIGESTIVE_TRACT_DEVELOPMENT | 1.4959 | 0.0378 |
| REG3A\|ORM1\|TRP | GOBP_ACUTE_PHASE_RESPONSE | 1.4956 | 0.014 |
| SIRT1\|ETS1\|ATP5A | GOBP_POSITIVE_REGULATION_OF_BLOOD_VESSEL_ENDOT | 1.4951 | 0.0749 |
| GDF6\|SHH\|FOXA1\| | GOBP_POSITIVE_REGULATION_OF_NEURON_DIFFERENTIATI | 1.494 | 0.0396 |
| NTRK1\|SIRT1\|WNT | GOBP_POSITIVE_REGULATION_OF_PHOSPHATIDYLINOSITOL | 1.4918 | 0.0414 |
| KITL\|SHH\|ISL1\|SMO | GOBP_MESENCHYMAL_CELL_MIGRATION | 1.4917 | 0.0613 |
| PDPN\|ZFP703\|FGG | GOBP_REGULATION_OF_HOMOTYPIC_CELL_CELL_ADHESIO | 1.4914 | 0.0423 |
| GJA5\|MSX2\|ELN\|CI | GOBP_OUTFLOW_TRACT_MORPHOGENESIS | 1.4907 | 0.0429 |
| SORBS2\|PDCD4\|PE | GOBP_CARDIAC_CELL_DEVELOPMENT | 1.4905 | 0.0364 |
| ALOX5\|TERF2\|PTK | GOBP_POSITIVE_REGULATION_OF_OXIDOREDUCTASE_ACTI | 1.4893 | 0.0468 |
| SPRY1\|FSTL3\|IL12 | GOBP_NEGATIVE_REGULATION_OF_CELLULAR_RESPONSE_ | 1.488 | 0.0612 |
| SHH\|WNT5A\|LGR4\| | GOBP_REGULATION_OF_MORPHOGENESIS_OF_A_BRANCHI | 1.4875 | 0.0137 |
| CHRM2\|CALCA\|GP | GOBP_REGULATION_OF_SMOOTH_MUSCLE_CONTRACTION | 1.4865 | 0.06 |
| CITED2\|ISL1\|OSM\|N | GOBP_PERIPHERAL_NERVOUS_SYSTEM_DEVELOPMENT | 1.4864 | 0.0339 |
| RPL30\|CXCL15\|CX | GOBP_ANTIMICROBIAL_HUMORAL_RESPONSE | 1.485 | 0.0032 |
| DPP4\|CKLF\|CXCL1 | GOBP_NEUTROPHIL_CHEMOTAXIS | 1.484 | 0.0753 |
| SPRY1\|SHH\|SLC7A | GOBP_RESPIRATORY_SYSTEM_DEVELOPMENT | 1.4822 | 0.0284 |
| HINT1\|CD3E\|PLCG | GOBP_REGULATION_OF_CALCIUM_MEDIATED_SIGNALING | 1.4814 | 0.0058 |
| SERPINC1\|AKR1B3 | GOBP_REGULATION_OF_BODY_FLUID_LEVELS | 1.479 | 0.0423 |
| FEZF2\|GATA5\|PAX | GOBP_REGIONALIZATION | 1.4774 | 0.0258 |
| AKR1B3\|IRAK4\|IL12 | GOBP_SMOOTH_MUSCLE_CELL_PROLIFERATION | 1.4768 | 0.0542 |
| NTRK1\|WNT16\|FYN | GOBP_PHOSPHATIDYLINOSITOL_3_KINASE_SIGNALING | 1.4765 | 0.0507 |
| SMARCD3\|TNNC1\| | GOBP_CARDIAC_CHAMBER_DEVELOPMENT | 1.4765 | 0.0423 |
| AKR1B3\|SHH\|IRAK4 | GOBP_MUSCLE_CELL_PROLIFERATION | 1.4765 | 0.0289 |
| CAPN1\|IRF6\|PINC\|A | GOBP_MAMMARY_GLAND_DEVELOPMENT | 1.4759 | 0.0237 |
| AKR1B3\|SLC7A11\| | GOBP_PHENOL_CONTAINING_COMPOUND_METABOLIC_PRO | 1.4759 | 0.0965 |
| CKLF\|MAPK1\|STAF | GOBP_MACROPHAGE_MIGRATION | 1.4739 | 0.0099 |
| GRHL2\|SNAI1\|MYO | GOBP_TIGHT_JUNCTION_ORGANIZATION | 1.4731 | 0.0138 |
| ALOX5\|PAM\|CD36\| | GOBP_LONG_CHAIN_FATTY_ACID_METABOLIC_PROCESS | 1.4708 | 0.0397 |
| GDF6\|GLG1\|TRIP1 | GOBP_CONNECTIVE_TISSUE_DEVELOPMENT | 1.4704 | 0.0273 |
| NF1\|BMP4\|NRAS\|P | GOBP_POSITIVE_REGULATION_OF_ENDOTHELIAL_CELL_PR | 1.4704 | 0.0132 |
| GATA3\|CFLAR\|SEF | GOBP_NEGATIVE_REGULATION_OF_EPITHELIAL_CELL_APOI | 1.4684 | 0.0542 |
| RPS7\|GRHL2\|BMP4 | GOBP_PRIMARY_NEURAL_TUBE_FORMATION | 1.4678 | 0.0648 |
| ATP1B2\|UTS2R\|SN | GOBP_SODIUM_ION_HOMEOSTASIS | 1.4647 | 0.054 |
| SLC7A11\|UBASH3E | GOBP_HOMOTYPIC_CELL_CELL_ADHESION | 1.4645 | 0.0263 |
| KITL\|SHH\|BMP4\|BM | GOBP_STEM_CELL_DEVELOPMENT | 1.464 | 0.0305 |
| C1QTNF12\|GPER1 | GOBP_PROTEIN_KINASE_B_SIGNALING | 1.4622 | 0.0421 |
| SHH\|2210016L21R | GOBP_REGULATION_OF_WNT_SIGNALING_PATHWAY | 1.4609 | 0.0046 |
| GATA1\|GLI1\|AURK | GOBP_REGENERATION | 1.4602 | 0.044 |
| PYCARD\|SLAMF9\| | GOBP_CHEMOKINE_PRODUCTION | 1.458 | 0.0542 |
| CACNB4\|FGF14\|AL | GOBP_POSITIVE_REGULATION_OF_TRANSPORTER_ACTIVIT | 1.457 | 0.0642 |
| SHH\|PBRM1\|PROK | GOBP_VASCULOGENESIS | 1.4568 | 0.056 |
| IFITM5\|CCN1\|SGMS | GOBP_REGULATION_OF_BONE_MINERALIZATION | 1.4561 | 0.0064 |
| MTPN\|TNNC1\|ZFP4 | GOBP_REGULATION_OF_MUSCLE_ADAPTATION | 1.4559 | 0.0416 |
| NUMA1\|MESP1\|SUL | GOBP_POSITIVE_REGULATION_OF_EPITHELIAL_CELL_DIFFE | 1.4549 | 0.0368 |
| AKR1B3\|LCN5\|PLP | GOBP_ISOPRENOID_METABOLIC_PROCESS | 1.454 | 0.0206 |
| SORBS1\|EPHA1\|PT | GOBP_STRESS_FIBER_ASSEMBLY | 1.4538 | 0.0242 |
| SSTR5\|SHH\|NUP62 | GOBP_POSITIVE_REGULATION_OF_CELL_DIVISION | 1.4527 | 0.0377 |
| CFLAR\|HCRTR1\|G | GOBP_POSITIVE_REGULATION_OF_ERK1_AND_ERK2_CASC | 1.4519 | 0.0267 |
| EPHA1\|RHPN1\|GPF | GOBP_REGULATION_OF_ACTOMYOSIN_STRUCTURE_ORGAI | 1.4514 | 0.0895 |
| SPRY1\|PYCARD\|AI | GOBP_NEGATIVE_REGULATION_OF_PROTEIN_SERINE_THRE | 1.4512 | 0.0407 |
| SORBS1\|PTK2B\|C | GOBP_POSITIVE_REGULATION_OF_LIPID_METABOLIC_PROC | 1.4512 | 0.0311 |
| FLII\|MYOZ1\|MYPN\| | GOBP_CELLULAR_COMPONENT_ASSEMBLY_INVOLVED_IN_M | 1.4507 | 0.0272 |
| KITL\|RPS7\|SHH\|BM | GOBP_NEURAL_CREST_CELL_DIFFERENTIATION | 1.4499 | 0.0339 |
| ALOX5\|AKR1C18\|A | GOBP_OLEFINIC_COMPOUND_METABOLIC_PROCESS | 1.4498 | 0.0274 |
| DRD5\|ACVR2A\|GZ | GOBP_REGULATION_OF_OXIDOREDUCTASE_ACTIVITY | 1.4497 | 0.0838 |
| GRHL2\|CLDN4\|NED | GOBP_CELL_CELL_JUNCTION_ASSEMBLY | 1.4493 | 0.0201 |
| LOX\|RIPOR1\|CXCL | GOBP_RESPONSE_TO_CHEMOKINE | 1.4468 | 0.0771 |
| 2210016L21RIK\|TN | GOBP_CELL_CELL_SIGNALING_BY_WNT | 1.4453 | 0.0026 |
| ALOX5\|UTS2R\|ADD | GOBP_REGULATION_OF_TUBE_SIZE | 1.4453 | 0.0833 |
| SHH\|WNT1\|NF1\|VIT | GOBP_SPINAL_CORD_DEVELOPMENT | 1.4443 | 0.0679 |
| SERPINC1\|SHH\|SL | GOBP_COAGULATION | 1.4438 | 0.0043 |
| SHH\|WNT16\|NEUR | GOBP_CELL_FATE_COMMITMENT | 1.443 | 0.0532 |
| SORBS1\|AOAH\|ND | GOBP_POLYSACCHARIDE_METABOLIC_PROCESS | 1.4418 | 0.0666 |
| PTPRS\|NEDD4L\|C | GOBP_REGULATION_OF_CELL_JUNCTION_ASSEMBLY | 1.4411 | 0.0267 |
